# The chicken retrovirus-like gene *ENS-1/ERNI* and its LTR *Soprano* are involved in primordial germ cell development

**DOI:** 10.1101/2025.07.30.667597

**Authors:** Yuya Okuzaki, Akane Kawaguchi, Yumi Ozaki, Takeo Uemura, Daisuke Saito, Ken-ichi Nishijima

**Affiliations:** Avian Bioscience Research Center, Graduate School of Bioagricultural Sciences, Nagoya University, Furo-cho, Chikusa-ku, Nagoya, 464-8601, Japan; Department of Animal Sciences, Graduate School of Bioagricultural Sciences, Nagoya University, Furo-cho, Chikusa-ku, Nagoya, 464-8601, Japan; Molecular Life History Laboratory, National Institute of Genetics, Mishima, Shizuoka, 411-8540, Japan; Department of Genetics, Sokendai (Graduate University for Advanced Studies), Mishima, Shizuoka, 411-8540, Japan; Graduate School of Science, Kyushu University, Fukuoka-shi, Fukuoka, 819-0395, Japan

## Abstract

Cultured chicken primordial germ cells (PGCs) are attractive tools for reproductive biology and for preserving and modifying genetic resources. However, it remains difficult to establish cell lines from some breeds or females. In the present study, we performed transcriptome analyses of cultured PGCs from different breeds and found that the expression of the *Galliformes*-specific endogenous retrovirus-like gene *ENS-1/ERNI* correlated with the establishment efficiency of PGC lines. CRISPRi targeting the *ENS-1/ERNI* long terminal repeat (LTR) sequence (*Soprano*) broadly reduced homeostasis-related genes including ribosome, endoplasmic reticulum, and oxidative phosphorylation. These PGCs exhibited abnormal cell cycle arrest under DNA-damaged conditions, suggesting involvement in the quality control mechanisms of chicken PGCs. Furthermore, when transplanted into blastoderms, these PGCs showed a reduction in colonization of the genital ridges. These results indicate the involvement of *Soprano* LTR and *ENS-1/ERNI* in the development of chicken PGCs and their adaptation to the in vitro environment.

## Introduction

Primordial germ cells (PGCs) are intrinsic stem cells in the embryonic development of animals, and only PGCs form gametes and transmit their genetic information to subsequent generations. Although the mechanisms underlying developmental processes are an important issue in developmental and reproductive biology, the knowledge on species other than rodents and human is limited. Chickens are the most widely raised livestock in the world and there are many regionally specific breeds. The preservation of these breeds is important for species diversity and agricultural purposes; however, the cryopreservation of fertilized eggs, which is commonly performed for mammals, is challenging for avian species due to the presence of huge yolk and eggshells. Therefore, avian PGCs are an attractive alternative for preserving genetic resources. Extensive research on chickens has made it possible to culture PGCs in vitro for long periods without the loss of their capacity to differentiate into sperm and oocytes when transplanted into recipient embryos [1–3]. This opens the possibility to cryopreserve genetic resources and to create genetically modified chickens [4–7]. However, PGCs obtained from some breeds and females remain difficult to expand in vitro [1,3,8–10].

Transposable elements (TEs), also known as jumping genes, are genetic elements that have the ability to move their sequences from one locus to another within a genome. Recent studies demonstrated that TEs are not merely selfish parasitic sequences but they are also involved in various physiological processes, including germ cell development, embryonic development, and cancer [11]. Among various TEs, long terminal repeat (LTR) retrotransposons are considered to have originated from a common ancestor with retroviruses early in the evolution of vertebrates [12]. Functional LTR retrotransposons encode the viral *Gag*, *Pol*, and *Env* genes, and some of these have co-evolved with the host genes and become involved in mammalian embryonic development, such as Syncytin-1 and PEG10 [11,13,14]. Furthermore, these LTR sequences spread throughout the genome and function as cis-regulatory elements, such as promoters and enhancers, widely affecting the regulation of gene expression in various cell types [11,15,16]. Birds have the unique feature of having a markedly lower percentage of TEs in their genome than mammals and reptiles; however, they possess several families of LTR transposons [17]. Four main types of LTR retrotransposons (*Birddawg*, *Hitchcock*, *Kronos*, and *Soprano*) have been identified in chickens [18]. Of these, *Soprano* exists in more than 1,000 copies in the chicken genome; most of these are solo LTRs or truncated sequences that lack functional coding sequences. While, some *Soprano* contain the functional proteins other than *Gag*, *Pol*, and *Env*, ENS-1 and ERNI, as internal domains. *ENS-1/ERNI* is a *Galliformes*-specific endogenous retrovirus-like gene that was originally reported to be expressed in prospective neural plates and the area pellucida of blastoderms [19]. Subsequent studies showed that it was also expressed in the hypoblast, the pluripotent epiblast, PGCs, and in embryonic stem cells established from the pluripotent epiblast [20–24]. The ENS-1/ERNI protein plays a key role in neural plate development by regulating the timing of *SOX2* expression together with the epigenetic regulator CBX3 (also known as HP1γ) [21]. However, the role of ENS-1/ERNI in other types of cells and those of solo *Soprano* LTRs remain unclear.

We herein performed a transcriptome analysis of cultured PGCs from chicken breeds with different cultured PGC establishment efficiencies (EE) and found that *ENS-1/ERNI* expression correlated with EE. Furthermore, we demonstrated that ENS-1/ERNI and *Soprano* broadly regulated gene expression in cultured PGCs and were involved in the development of PGCs in ovo. Collectively, this study sheds new light on the role of LTR retrotransposons in the development of germ cells in birds.

## Results

### Establishment of in vitro cultured PGCs from various chicken strains

During the course of efforts to cryopreserve various chicken breeds that are maintained at the Avian Bioscience Research Center at Nagoya University, [25] we attempted to establish in vitro cultured PGCs from various inbred or closed chicken strains. Blood collected from a single embryo at HH St.14-16[26] was seeded into microwell plates and expanded using the medium previously reported by Whyte et al. [1]. PGCs that proliferated to more than 1×10^5^ cells were considered to be successfully established as a cell line. These cells highly expressed PGC marker genes (Supplementary Fig. S1A) [1,27–29], and retained the ability to produce functional gametes, as confirmed in transplantation experiments on several strains (GSP, GSN/1, and M/O) of cultured PGCs (Supplementary Fig. S1B, Supplementary Table S1). We obtained several transgenic chicken lines of the M/O strain (reference 25 and manuscript in preparation). On the other hand, EE markedly varied depending on the chicken strain. The EE of cultured PGCs in each chicken strain and their sex are shown in Table 1. The establishment of female PGCs was more difficult in strains with low EE (Fig. 1A), which was consistent with previous findings [1,9].

**Figure 1.**
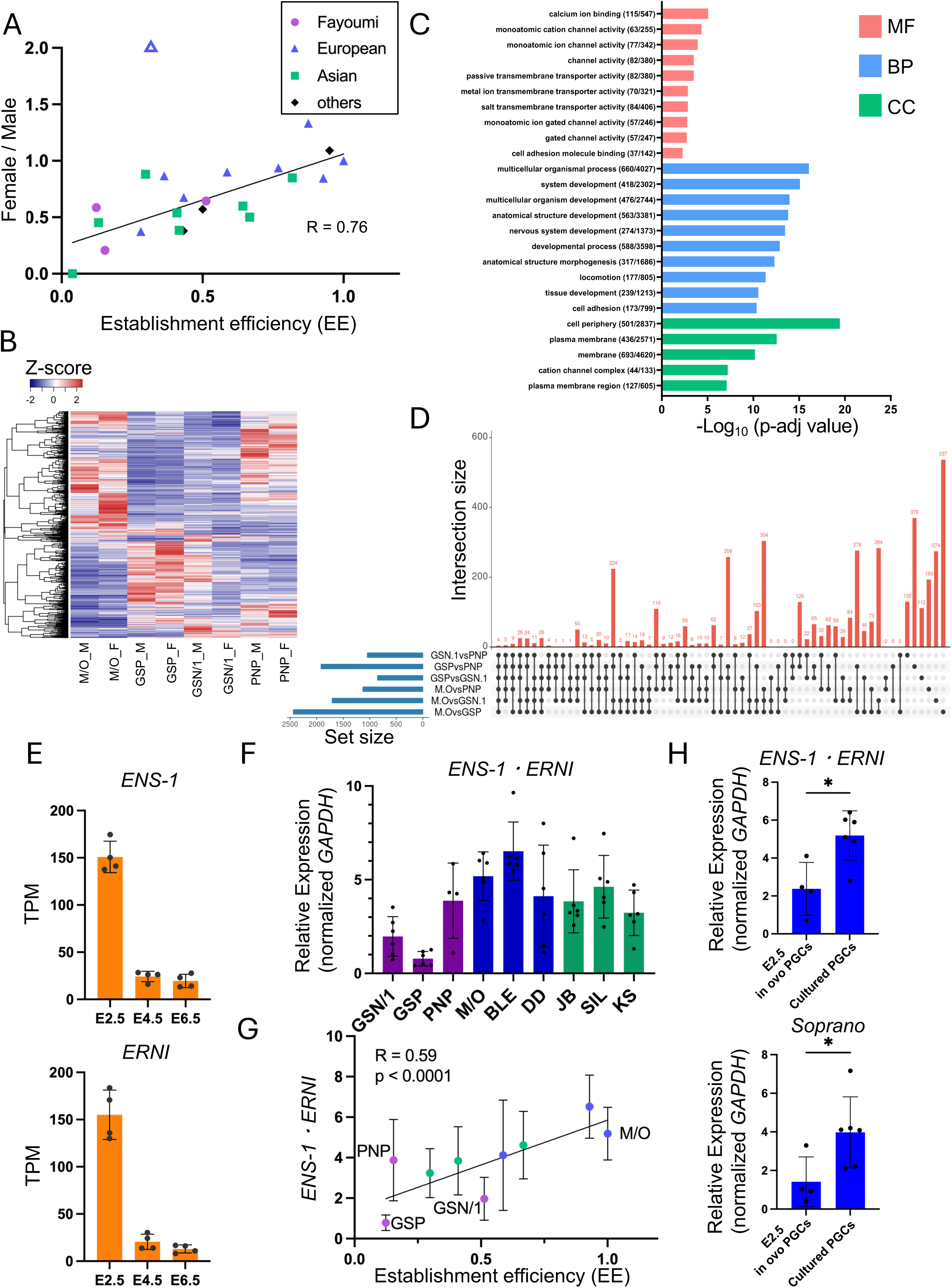
The retrovirus-like gene *ENS-1/ERNI* shows high levels of expression in cultured PGCs and correlates with their establishment efficiency. A) The correlation between the establishment efficiency (EE) of in vitro cultured PGCs and their sex ratio. The outlier (BMC strain) is shown as an open triangle. B) A heat map showing differentially expressed genes (p-adj < 0.01) based on an RNA-seq variance analysis comparing male (M) and female (F) of four chicken strains of in vitro cultured PGCs: GSP, GSN/1, PNP, and M/O. C) A GO analysis of differentially expressed genes (p-adj < 0.01) based on an RNA-seq variance analysis comparing the four chicken strains of in vitro cultured PGCs: GSP, GSN/1, PNP, and M/O. D) An upset plot showing the inclusive relation of DEGs (p-adj < 0.01) based on an RNA-seq expression analysis comparing the four chicken strains of in vitro cultured PGCs: GSP, GSN/1, PNP, and M/O. E) *ENS-1* and *ERNI* expression levels in each embryonic stage of in ovo PGCs based on an RNA-seq expression re-analysis [33]. F) The RNA from in vitro cultured PGCs established from each chicken strain was subjected to qRT-PCR. The expression levels of *ENS-1/ERNI* were normalized by those of *GAPDH*. Data show the mean ± SD of individual clones. Purple bars show the Fayoumi breeds, blue bars show the European breeds, and green bars show the Asian breeds. Note: The primers used in qRT-PCR do not distinguish between *ENS-1* and *ERNI*. G) The correlation between the establishment efficiency (EE) of in vitro cultured PGCs and the expression level of *ENS-1/ERNI*. Expression was evaluated for individual clones by qRT-PCR and is shown as the mean ± SD after normalization by the expression of *GAPDH*. Purple dots show the Fayoumi breeds, blue dots show the European breeds, and green dots show the Asian breeds. H) The RNA from in vitro cultured PGCs established from the M/O strain and in ovo PGCs collected from the E2.5 embryonic blood of the M/O strain was subjected to qRT-PCR. The expression levels of *ENS-1/ERNI* and UTR of *Soprano* were normalized by those of *GAPDH*. Data show the mean ± SD of biologically different samples.

**Table 1.** The establishment efficiency of in vitro cultured PGC lines from various chicken strains.

| Chicken strains | Group | Inoculated number |  | Established number |  | Establishment efficiency (EE) |  |  |
| --- | --- | --- | --- | --- | --- | --- | --- | --- |
|  |  | Male | Female | Male | Female | total | Male | Female |
| GSN/1 (Fayoumi breed) | Fayoumi | 17 | 24 | 11 | 10 | 0.51 | 0.65 | 0.42 |
| PNP (Fayoumi breed) |  | 22 | 25 | 3 | 1 | 0.09 | 0.14 | 0.04 |
| GSP (Fayoumi breed) |  | 25 | 32 | 4 | 3 | 0.12 | 0.16 | 0.09 |
| WLG (White Leghorn breed) | European | 5 | 8 | 4 | 6 | 0.77 | 0.8 | 0.75 |
| M/O (White Leghorn breed) |  | 5 | 8 | 5 | 8 | 1 | 1 | 1 |
| BLE (Brown Leghorn breed) |  | 15 | 13 | 15 | 11 | 0.93 | 1 | 0.85 |
| GB (Game Bantam breed) |  | 11 | 19 | 6 | 7 | 0.43 | 0.55 | 0.37 |
| OS |  | 24 | 8 | 8 | 1 | 0.28 | 0.33 | 0.13 |
| DD (Dandarawi breed) |  | 29 | 34 | 18 | 19 | 0.59 | 0.62 | 0.56 |
| BMC (Black Minorca breed) |  | 16 | 8 | 4 | 4 | 0.33 | 0.25 | 0.5 |
| POL (Polish breed) |  | 4 | 4 | 3 | 4 | 0.88 | 0.75 | 1 |
| BRB (Brahma Bantam breed) |  | 13 | 9 | 5 | 3 | 0.36 | 0.38 | 0.33 |
| PC (Petit-Cocco breed) | Asian | 17 | 16 | 15 | 12 | 0.82 | 0.88 | 0.75 |
| JB (Japanese Bantam breed) |  | 29 | 37 | 16 | 11 | 0.41 | 0.55 | 0.3 |
| EJ (Ehime-Jidori breed) |  | 85 | 75 | 15 | 6 | 0.13 | 0.18 | 0.08 |
| SIL (White Silkie breed) |  | 5 | 10 | 5 | 5 | 0.67 | 1 | 0.5 |
| CHN (Okinawa Crower breed) |  | 6 | 8 | 5 | 4 | 0.64 | 0.83 | 0.5 |
| CB (Cochin Bantam breed) |  | 27 | 25 | 2 | 0 | 0.04 | 0.07 | 0 |
| NGY (Nagoya breed) |  | 20 | 28 | 13 | 7 | 0.42 | 0.65 | 0.25 |
| KS (Ko-Shamo breed) |  | 22 | 25 | 7 | 7 | 0.3 | 0.32 | 0.28 |
| CAL (White Wyandotte breed) | European x Asian | 12 | 6 | 7 | 2 | 0.5 | 0.58 | 0.33 |
| TK | Fayoumi x Asian | 12 | 8 | 11 | 8 | 0.95 | 0.92 | 1 |
| GRN | N/A | 23 | 14 | 13 | 3 | 0.43 | 0.57 | 0.21 |

Note: The grouping of each chicken strain is based on a phylogenetic tree analysis using microsatellite markers by Nunome et al. [70].

### The expression level of *ENS-1/ERNI* correlated with the EE of in vitro cultured PGCs

To elucidate the mechanisms underlying differences in the EE of cultured PGCs between sexes and strains, we performed an RNA-seq analysis of the cultured PGCs of males and females derived from several strains. These included three strains of the Fayoumi breed (GSP, GSN/1, and PNP), which exhibited low EE with a variable magnitude, and an M/O strain of the White Leghorn breed, which had the highest EE of those tested. We then investigated differentially expressed genes (DEGs) between the sexes and strains.

Although EE markedly differed between male and female PGCs (Table 1, Fig. 1A), relatively small differences in gene expression were observed in cultured PGCs (431 genes, p < 0.01; and 226 genes, p < 0.01, and |Fold Change (FC)| > 2) (Supplementary Fig. S2A). The majority of DEGs between males and females were located on the sex chromosome (Chr Z/W) (Supplementary Fig. S2B, Supplementary Table S2). Consistent with previous reports showing no chromosome-wide dosage compensation of sex chromosomes in birds [30,31], the present results demonstrated that the average expression levels of genes on the Z chromosome were two-fold higher in males than in females (Supplementary Fig. S2C). On the other hand, the expression of some genes on the Z chromosome showed no sex difference, suggesting that the dosage compensation of sex chromosomes exists at the level of individual genes in chicken PGCs, as reported in previous studies that examined various types of cells [10,30–32] (Supplementary Fig. S2D).

Regarding transcriptome differences among strains, we performed a variance analysis to extract DEGs in either strain and identified 2868 DEGs (p < 0.01) (Fig. 1B, Supplementary Table S3). Subsequent GO analyses showed that enriched GO terms were classified as molecular function (MF), biological process (BP), or cellular component (CC) (Fig. 1C and Supplementary Table S4). In terms of MF, ion channels and transmembrane transporters, particularly those related to calcium ions, accumulated as the major term. Cultured PGCs are sensitive to calcium ion concentrations and require a medium with reduced calcium ions to prevent cell-cell aggregation [1]. Strain differences in calcium dynamics are consistent with our observation showing that the magnitude of PGC aggregation varied among various breeds. Consistent with MF terms, locomotion and cell adhesion terms were enriched in the BP category. In addition, BP terms related to development were enriched, including many neurogenesis-related terms. In CC, plasma membrane and cell adhesion terms were mainly enriched, which is consistent with MF results. Furthermore, terms related to neurons were enriched as well as BP. To clarify EE differences among chicken strains, we analyzed transcriptome data and extracted DEGs between the M/O strain and each strain of the Fayoumi breed (Fig. 1D). A total of 409 genes were identified as DEGs between the M/O strain and Fayoumi breeds (Supplementary Table S5). We found that *ENS-1/ERNI* was the top hit gene, which was the highest in the M/O strain and lower in Fayoumi breeds, among the genes that were expressed more highly in PGCs than in chicken embryonic fibroblasts (CEFs), the counterpart for non-germ and non-pluripotent stem cells (Supplementary Fig. S2E, Supplementary Table S6). A reanalysis of RNA-seq data from endogenous and non-cultured in ovo PGCs by Ichikawa et al. [33] revealed that *ENS-1/ERNI* was highly expressed in blood circulating PGCs at E2.5, and also that its expression decreased after PGCs had settled into the genital ridge (Fig. 1E); although, a specific level of *ENS-1/ERNI* expression was maintained in gonadal PGCs at E6.5 [34]. The qRT-PCR analysis confirmed that *ENS-1/ERNI* was expressed in cultured PGCs established from various chicken strains, but that expression levels varied between strains (Fig. 1F), which is consistent with the results of the RNA-seq analysis. Furthermore, a moderate correlation was observed between *ENS-1/ERNI* expression levels and EE (Fig. 1G). In addition, the expression of *ENS-1/ERNI* and the untranslated region in *Soprano* LTR was higher in cultured PGCs than in E2.5 blood circulating in ovo PGCs (Fig. 1H). These results imply the involvement of *ENS-1/ERNI* in the in vivo development and in vitro expansion capacity of chicken PGCs.

### Regulation of gene expression in cultured PGCs by *ENS-1/ERNI* and *Soprano* LTR

*ENS-1/ERNI* is an endogenous retrovirus-like gene, and its LTR, *Soprano*, is present in more than 1000 copies in the chicken genome [18]. Furthermore, dozens of truncated pseudogenes have been identified in *ENS-1/ERNI* [21] . Therefore, instead of examining each of them individually, we initially investigated the comprehensive functions of these factors in PGCs by suppressing the *Soprano* LTR using CRISPRi. We established a PGC line stably expressing dCas9-KRAB-MeCP2 (CRISPRi PGCs) [35]. We induced CRISPRi by transfecting this cell line with in vitro synthesized gRNA against *Soprano* LTR and analyzed transcriptome changes (Supplementary Fig. S3A). Based on RNA-seq results, a moderate decrease was observed in the expression level of *ENS-1/ERNI*, which may have been due to the low stability of gRNA; however, the results obtained showed significant changes in the expression of multiple genes (256 genes in sgSoprano#1 and 248 genes in sgSoprano#2, p < 0.05, and |FC| > 1.5) (Supplementary Table S7,8, Supplementary Fig. S3B). To increase the efficiencies of CRISPRi against *Soprano* LTR, we examined gRNA expression plasmid vector transfection. To achieve efficient CRISPRi, we constructed a plasmid expressing three different gRNAs against the *Soprano* LTR (3× sgSoprano) (Supplementary Fig. S3A). This vector successfully reduced the expression levels of *ENS-1/ERNI* (less than 9% and 5% for *ENS-1* and *ERNI*, respectively) and many *ENS-1/ERNI*-like pseudogenes (Fig. 2A, Supplementary Table S9). In addition, 1422 (p < 0.01) and 307 (p < 0.01, |FC| > 2) of genes showed changes in expression levels. The GO analysis revealed that *Soprano* LTR CRISPRi reduced the expression of genes encoding ribosomal proteins, endoplasmic reticulum-related proteins, and oxidative phosphorylation-related proteins, suggesting the involvement of *Soprano* LTR in the regulation of broad metabolic processes in cultured PGCs (Fig. 2B, Supplementary Table S10, 11). On the other hand, the GO analysis of genes with increased expression showed cell migration and development, particularly neuronal development-related terms. These results appear to be consistent with those of the GO analysis of cultured PGCs among chicken strains (Fig. 1C) and with previous studies showing the involvement of ENS-1/ERNI in the transcriptional regulation of neuronal development [19–21]. There were 199 genes that were significant (p<0.05) DEGs shared under three different *Soprano* LTR CRISPRi conditions (Fig. 2C). Among 25 genes with p<0.05 and |FC| > 2 from 3× sgSoprano plasmid-transfected PGC samples (Supplementary Table S12), there were 10 protein-coding genes (*AGK*, *CDH7*, *DENND11*, *ENOX1*, *GUCA1C*, *SLC16A7*, *TMPRSS15*, *LOC418414*, *ENS-1*, and *ERNI*). Regarding *CDH7*, *SLC16A7*, and *TMPRSS15*, *Soprano*-like sequences were present in their flanking regions (within 500 kb) (Supplementary Fig. S3C), suggesting that these genes might be regulated by the enhancer activity of *Soprano* LTR. Notably, several genes were located in clusters in the chicken genome: *AGK*/*DENND11*/*WEE2* and *GUCA1C*/*LOC418414* were located as clusters within a very short region (70 kb in Chr 1 and 110 kb in Chr 1, respectively), and we did not detect any *Soprano*-like sequences in the neighboring region by a BLAST analysis (within 500 kb) (Supplementary Fig. S3C). The qRT-PCR analysis confirmed the expression reduction of these genes in cultured PGCs under *Soprano* LTR CRISPRi conditions, which is consistent with the results from RNA-seq, except for *DENND11* (Fig. 2D). Regarding *DENND11*, we examined in detail the mapped reads and found that unspliced RNA was transcribed in control (3× sgOVA) samples (Supplementary Fig. S3D). A strand-specific read count analysis also suggested that this long non-coding RNA was expressed from the reverse strand of *DENND11* (Supplementary Fig. S3E). Taken together, these results indicate that transcription from the positive strand of the *AGK*/*DENND11*/*WEE2* locus was activated in cultured PGCs via *Soprano* activities, and that the repression of *Soprano* decreased this transcription and increased that of reverse strand-encoded *DENND11*. A reanalysis of RNA-seq data in embryonic PGCs [33] showed that these genes were highly expressed in E2.5 embryonic PGCs, and their expression tended to decreased in subsequent embryonic stage samples, except for *ENOX1*, *LOC418414*, and *TMPRSS15* (Supplementary Fig. S3F), which is consistent with the reduction in *ENS-1/ERNI* expression levels during development.

**Figure 2.**
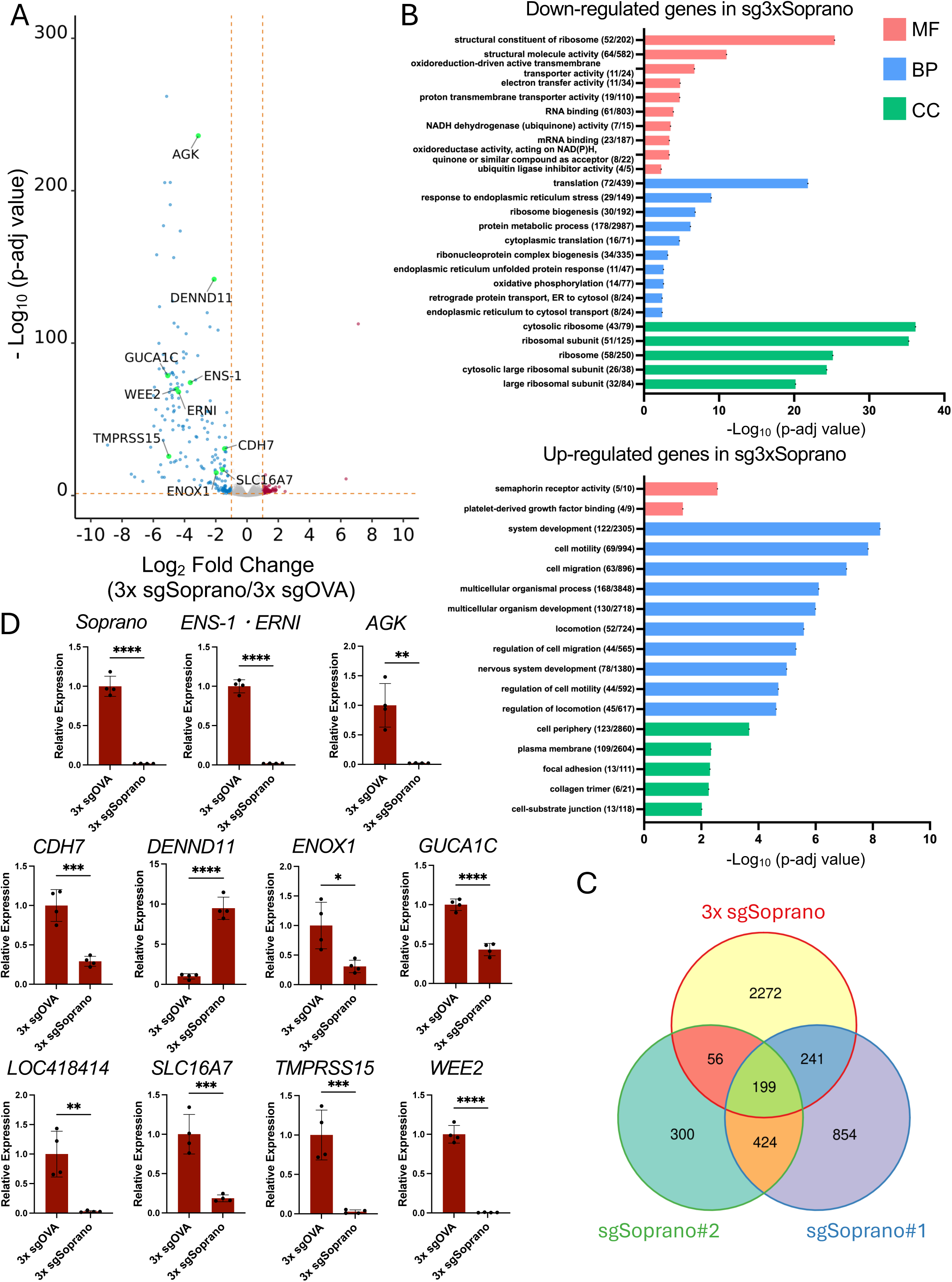
Analysis of gene expression variations in cultured PGCs under *Soprano* LTR CRISPRi. A) A volcano plot showing an RNA-seq expression analysis comparing 3× sgOVA and 3× sgSoprano expression plasmid-transfected CRISPRi PGCs. Dashed lines indicate -Log_10_ p-adj > 2 or |Log2 FC| > 1. B) GO analyses of differentially expressed genes (p< 0.01) from the RNA-seq expression analysis comparing 3× sgOVA and 3× sgSoprano expression plasmids transfected into CRISPRi PGCs. The upper panel shows the results of the GO analysis of genes down-regulated under 3×sgSoprano conditions, and the lower panel shows the results of the GO analysis of genes up-regulated under 3×sgSoprano conditions. C) A Venn plot showing the inclusive relation of DEGs (p< 0.05, sgOVA vs sgSoprano) based on an RNA-seq expression analysis comparing the 3× sgSoprano expression plasmid or synthesized sgRNA (sgSoprano#1 and #2)-transfected CRISPRi PGCs. D) The 3× sgOVA or 3× sgSoprano expression plasmid was transfected into CRISPRi PGCs, and RNA purified from them was subjected to qRT-PCR. The expression levels of each gene were normalized by *GAPDH* and that of 3× sgOVA was set as 1. Data show the mean ± SD of four different biological replicates. ** p < 0.01, *** p < 0.001, **** p < 0.0001; the Student’s *t*-test.

To clarify whether these results were due to the ENS-1/ERNI protein and/or *Soprano*, we performed knockout experiments using CRISPR-Cas9. We designed three different sgRNAs targeting the *ENS-1/ERNI* coding region, and conducted knockout experiments using Cas9 and sgRNA expression plasmids in cultured PGCs (Supplementary Fig. 4A). Following the transient selection of transfected PGCs, we observed a gradual decrease in cell numbers (Supplementary Fig. 4B). However, since the chicken genome had several dozen *ENS-1/ERNI* like pseudogene sequences [20,22], we were unable to rule out the possibility that this was caused by the DNA damage response triggered due to multiple breakage in genomic DNA by Cas9. Therefore, instead of knockout *ENS-1/ERNI*, we conducted siRNA knockdown experiments on cultured PGCs. We designed siRNAs on the *ENS-1/ERNI*-coding region (#1 was the central region and #2 was the 3’ terminal region (Supplementary Fig. 4A)). Both siRNAs reduced the expression of *ENS-1/ERNI* by approximately 50-70%. The expression of the UTR region inside the *Soprano* LTR was also reduced. Genes with expression decreased by *Soprano* LTR CRISPRi also showed modest but significant reductions with the knockdown of *ENS-1/ERNI* (Fig. 3A). Similar decreases in expression were observed in experiments using cultured PGCs derived from the GSP strain, which expresses lower levels of *ENS-1/ERNI* (Supplementary Fig. S5). We then performed *ENS-1/ERN*I rescue experiments using CRISPRi PGCs. Since *ENS-1/ERNI* was highly expressed in cultured PGCs, the transient transfection of an *ERNI* expression vector did not induce *ERNI* expression compatible with the endogenous level (Fig. 3B). However, the qRT-PCR analysis showed that *WEE2* expression was partially restored by a similar magnitude to *ERNI* expression, and *DENND11* expression was further increased by *ERNI* under *Soprano* LTR CRISPRi conditions. On the other hand, the expression of other genes was not rescued, suggesting that the regulation of their expression required *Soprano* activity and/or higher levels of *ERNI* expression. Regarding the *AGK*/*DENND11/WEE2* locus, *Soprano* up-regulated *WEE2* expression and down-regulated *DENND11* expression, while ERNI slightly up-regulated the expression of both genes. These results suggest the partial involvement of *ENS-1/ERNI* in regulating the expression of these genes, and also that *Soprano* and *ENS-1/ERNI* regulated gene expression in independent manners.

**Figure 3.**
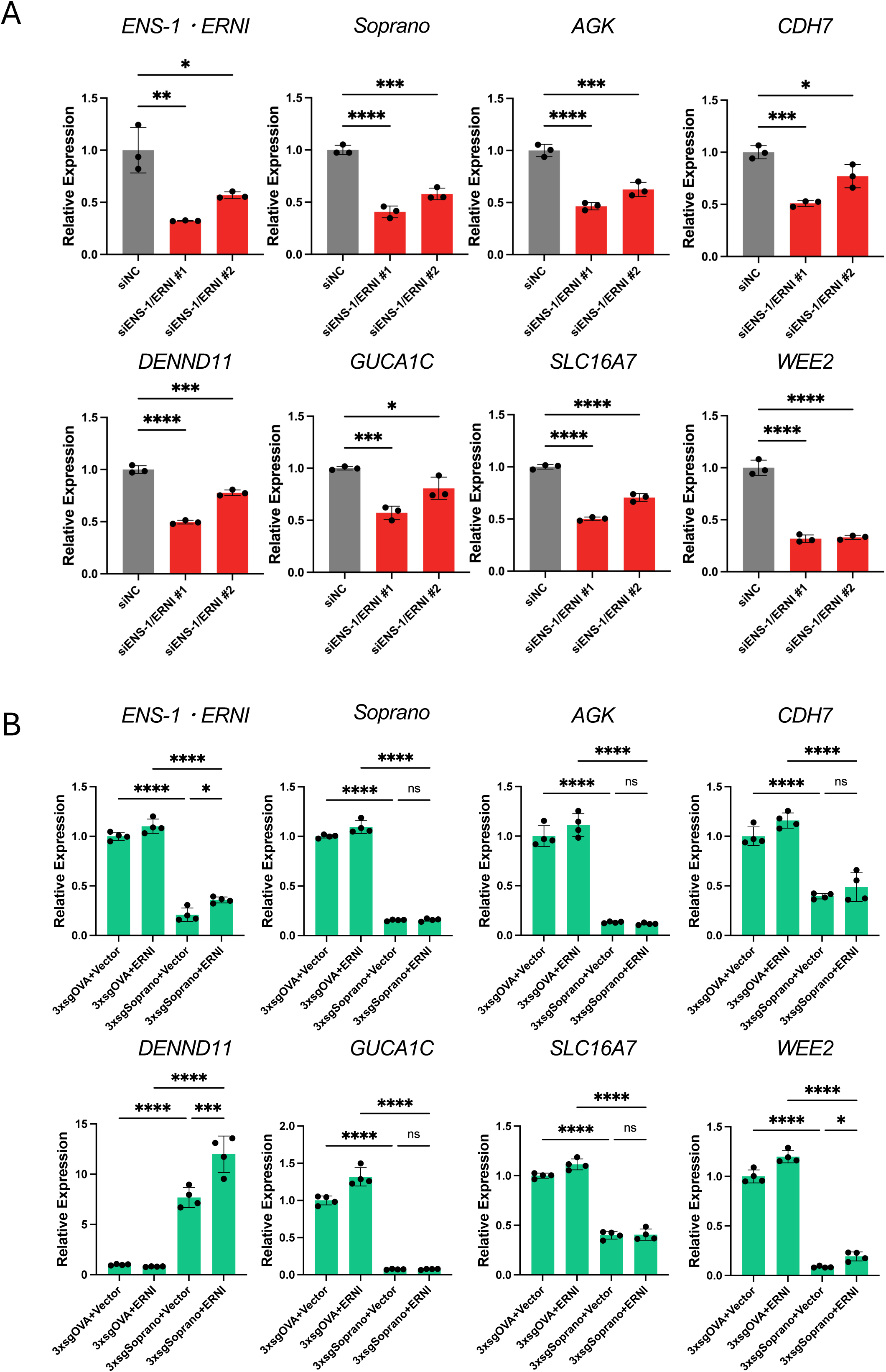
ENS-1/ERNI partially regulates the expression of gene sets that exhibited significant variations in *Soprano* LTR CRISPRi. A) *ENS-1/ERNI* knockdown experiments. The RNA from DsiRNA-transfected cultured PGCs was subjected to qRT-PCR. The expression levels of each gene were normalized by *GAPDH*, and that of control DsiRNA was set as 1. Data show the mean ± SD of three different biological replicates. * p < 0.05, ** p < 0.01, *** p < 0.001, **** p < 0.0001; ANOVA with Dunnett’s post hoc test. B) *ENS-1/ERNI* rescue experiments. The *ENRI* expression plasmid and 3× sgOVA or 3× sgSoprano expression plasmid were co-transfected into CRISPRi PGCs, and RNA purified from them was subjected to qRT-PCR. The expression levels of each gene were normalized by *GAPDH*, and that of 3× sgOVA + Vector was set as 1. Data show the mean ± SD of four different biological replicates. * p < 0.05, *** p < 0.001, **** p < 0.0001, ns: not significant; ANOVA with Tukey’s multiple comparisons post hoc test.

### Regulation of *Soprano* LTR in cultured PGCs

The results of the CRISPRi and siRNA experiments suggest that *Soprano* regulated gene expression more broadly than *ENS-1/ERNI* in cultured PGCs. Therefore, we investigated the regulatory mechanism of *Soprano* as a promoter and enhancer in PGCs. Since there are more than 1000 copies of *Soprano* in the chicken genome [18], we cloned several *Soprano* LTRs with different sequences and examined their promoter activity using a luciferase assay. As expected, some of the cloned *Soprano* LTRs exhibited high promoter activity, while others lacked promoter activity in cultured PGCs (Supplementary Fig. S6A, Supplementary Table S13). A consensus motif analysis performed on clones exhibiting higher promoter activity using MEME suite [36] indicated the presence of consensus motifs in the proximal and distal regions of the *Soprano* LTR, while the central parts were not conserved (Supplementary Fig. S6B). The luciferase assay using proximal and distal region sequences showed that only the distal region sequence exhibited promoter activity in cultured PGCs (Fig. 4A, Supplementary Fig. S6C). These results are consistent with the findings of previous studies that investigated *Soprano* LTR promoter activity using chicken embryonic stem cells [37,38]. Furthermore, *Soprano* LTR did not exhibit any promoter activity in chicken fibroblast cell line DF-1 cells, indicating that its promoter activity was specific to the cell type (Supplementary Fig. S6D). We then examined the consensus motifs in the distal region that were essential for promoter activity, and revealed that 35 bp of the motif 7 (motif 7 core) sequence was critical for activity (Fig. 4A). A transcription factor-binding site prediction analysis using JASPAR[39] suggested that the motif 7 sequence contained the binding motifs for GATA/PRDM1, ELK/ETS, and homeobox/NANOG (Fig. 4B, Supplementary Table S14). Therefore, we performed a luciferase assay with point mutants of motif 7 and found that promoter activity was significantly reduced when a mutation was introduced into the predicted homeobox/NANOG-binding site (Fig. 4C). We also demonstrated that the forced expression of NANOG in DF-1 cells increased the promoter activity of *Soprano* LTR (Fig. 4D). Since NANOG is the central transcription factor regulating the pluripotency of stem cells in chickens [40,41] and is highly expressed specifically in PGCs, epiblasts, and ESCs [24,27,42], these results indicate that NANOG was also an important transcription factor for the regulation of *Soprano* in cultured PGCs. PRDM1 (also called BLIMP1) is a key regulator of PGC specification in mice, non-human primates, and humans [43–45], and is also involved in the development of chicken PGCs [27]. Therefore, we conducted a luciferase assay utilizing mutant vectors with a predicted GATA/PRDM1-binding site. However, the promoter activities of mutant vectors were similar to those of the wild-type vector (Fig. 4C). Furthermore, the forced expression of PRDM1 in DF-1 cells did not affect the promoter activity of *Soprano* LTR (Fig. 4D). These results suggest that PRDM1 did not enhance *Soprano* LTR activity. A previous study suggested that GATA4 bound to this site, and also that the mutation of this site reduced promoter activity in chicken ESCs [37]. However, our experiment showed that its effects on promoter activity in cultured PGCs were negligible (Fig. 4C). This may be due to the low expression of GATA family genes in cultured PGCs (Supplementary Fig. S6E). We also investigated the enhancer activity of the motif 7 core sequence of *Soprano* LTR. Since enhancers function independently of their relative position [46], we evaluated the activity of the 3× motif 7 core sequence when placed upstream of the promoter sequence and downstream of the luciferase gene, respectively. The vector containing the 3× motif 7 core sequence upstream of the minimal TK promoter and downstream of the luciferase gene both exhibited significant luciferase activity in cultured PGCs; however, latter vector activity was weaker than former vector activity (Fig. 4E). In DF-1, the former vector showed a significant increase in luciferase activity upon the forced expression of NANOG, while the latter vector showed no luciferase activity even with the forced expression of NANOG (Supplementary Fig. S6F). These results suggest that the *Soprano* LTR might also possess enhancer activity, but that its activity was dependent on the cell type and also required other undifferentiated cell-specific factors in addition of NANOG.

**Figure 4.**
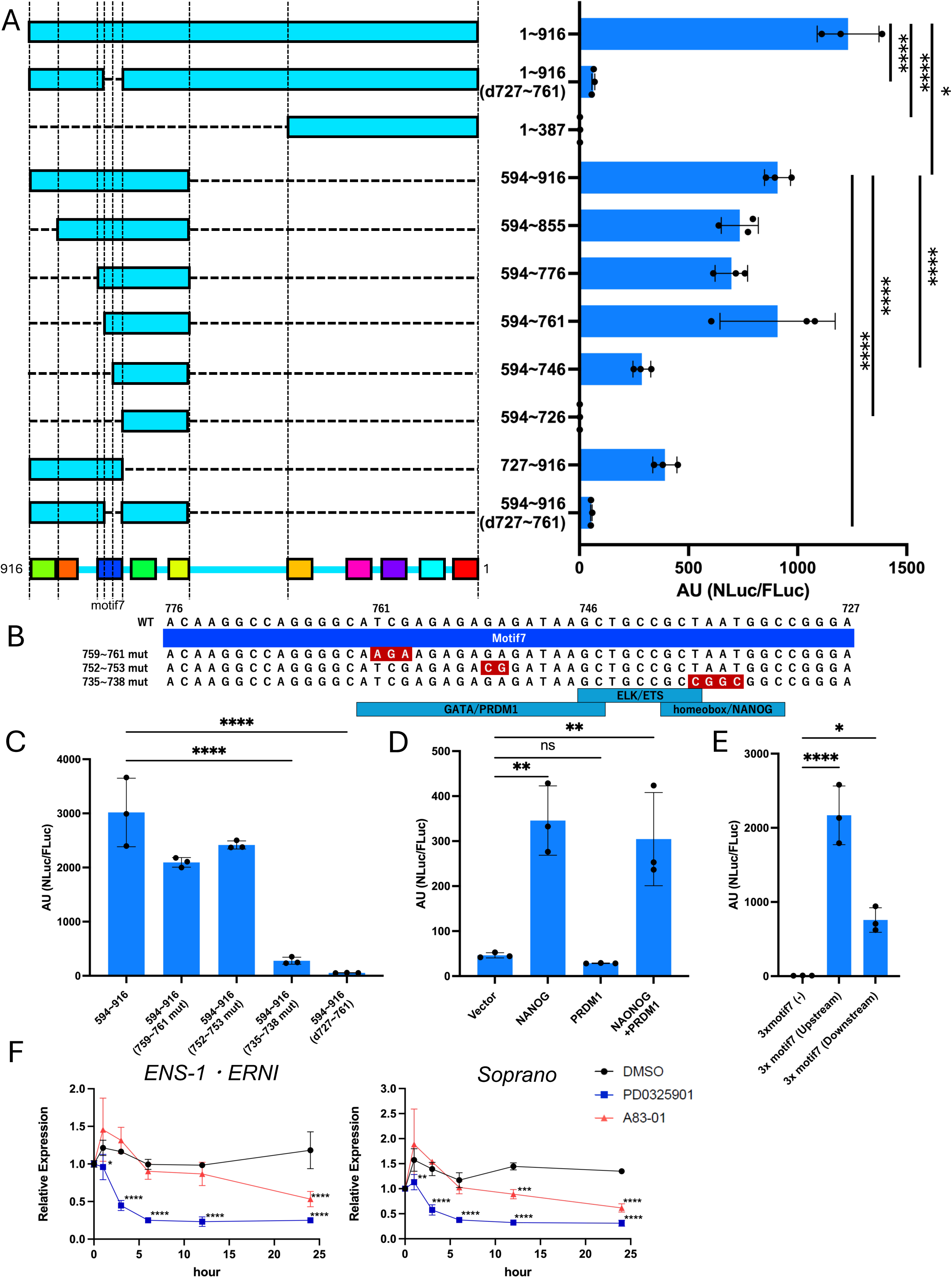
FGF signaling and NANOG regulate *Soprano* LTR expression in cultured PGCs. A-E) Dual luciferase assay investigating *Soprano* LTR activities. (A) Promoter activities of serial deletion mutants of *Soprano* LTR in cultured PGCs. (B) Sequence analysis of motif 7 in *Soprano* LTR. The nucleotides substituted in the mutants used in (C) are shown in red. The predicted transcription factor-binding sites based on the JASPAR analysis are shown in light blue. (C) Promoter activities of the motif 7 point mutants of *Soprano* LTR in cultured PGCs. (D) The effects of transcription factors against *Soprano* LTR in DF-1. (E) The enhancer activity of the motif 7 core sequence of *Soprano* LTR in cultured PGCs. The expression levels of NanoLuc (NLuc) were normalized by those of firefly luciferase (FLuc). Data show the mean ± SD of three different biological replicates. * p < 0.05, ** p < 0.01, *** p < 0.001, **** p < 0.0001, ns: not significant; ANOVA with Tukey’s multiple comparisons post hoc test (A). ANOVA with Dunnett’s post hoc test (C-E). F) Inhibitor treatment experiment on cultured PGCs. PGCs were treated with PD0325901 or A83-01 for the indicated period. RNAs were then subjected to qRT-PCR. The expression levels of *ENS-1/ERNI* and *Soprano* LTR were normalized by *GAPDH* and expression levels at 0 hr were set as 1. Data show the mean ± SD of three different biological replicates. * p < 0.05, ** p < 0.01, *** p < 0.001, **** p < 0.0001; ANOVA with Dunnett’s post hoc test.

The pluripotency and proliferation of chicken cultured PGCs are fundamentally controlled by FGF2/MAPK and activin A/SMAD signaling [1]. To identify the signaling pathway regulating the expression of *Soprano* LTR, we performed inhibitor treatment experiments on cultured PGCs. The culture medium of PGCs was changed to fresh medium, and PD0325902 (MEK inhibitor) or A83-01 (ALK4/5/7 inhibitor) was added at 0 hr. The expression levels of UTR in *Soprano* LTR and *ENS-1/ERNI* were measured by qRT-PCR at each time point (Fig. 4F). In vehicle or A83-01-treated samples, transient increases in the expression of *Soprano* LTR and *ENS-1/ERNI* were observed 1 hr after medium replacement, possibly due to the stimulation induced by the addition of new growth factors. The expression of these genes gradually decreased from 12 to 24 hr in A83-01-treated samples. On the other hand, in samples treated with PD0325901, their expression levels remained unchanged for 1 hr and then rapidly decreased between 3 to 6 hr. These results suggest that the expression of *Soprano* LTR and *ENS-1/ERNI* was basically regulated by FGF/MAPK signaling in cultured PGCs, which is consistent with previous findings showing that ectopic FGF8 administration induced *ENS-1/ERNI* expression in chick embryos [19]. Activin A/SMAD signaling also contributed to their expression; however, its role appeared to be indirect.

Collectively, these results suggest that *Soprano* LTR exhibited strong promoter activity in chicken PGCs and also acted as enhancer, and its expression was regulated by FGF signaling and NANOG, which bound to motif 7.

### *Soprano* LTR activity is involved in cell cycle progression during the induction of DNA damage in cultured PGCs

We investigated whether *Soprano* LTR CRISPRi affected the cell cycle in cultured PGCs. However, *Soprano* LTR CRISPRi alone did not affect the cell cycle of cultured PGCs (Fig. 5A). On the other hand, inducing DNA damages by a treatment with camptothecin (CPT), which inhibits topoisomerase I by which accumulates subsequent DNA damages leading to apoptosis, changed the rate of each phase of the cell cycle in cultured PGCs. CPT decreased the percentage of G0/G1 phase cells and increased that of G2/M phase cells, and at higher CPT concentrations also in the S phase. *Soprano* LTR CRISPRi significantly increased the percentage of G2/M phase cells in a CPT dosage-dependent manner (Fig. 5B), suggesting that *Soprano* regulated the cell cycle progression under DNA-damaged conditions. On the other hand, the percentage of apoptotic cells induced by the CPT treatment was not changed by *Soprano* LTR CRISPRi, indicating a modest role for the DNA damage response by *Soprano* (Fig. 5C).

**Figure 5.**
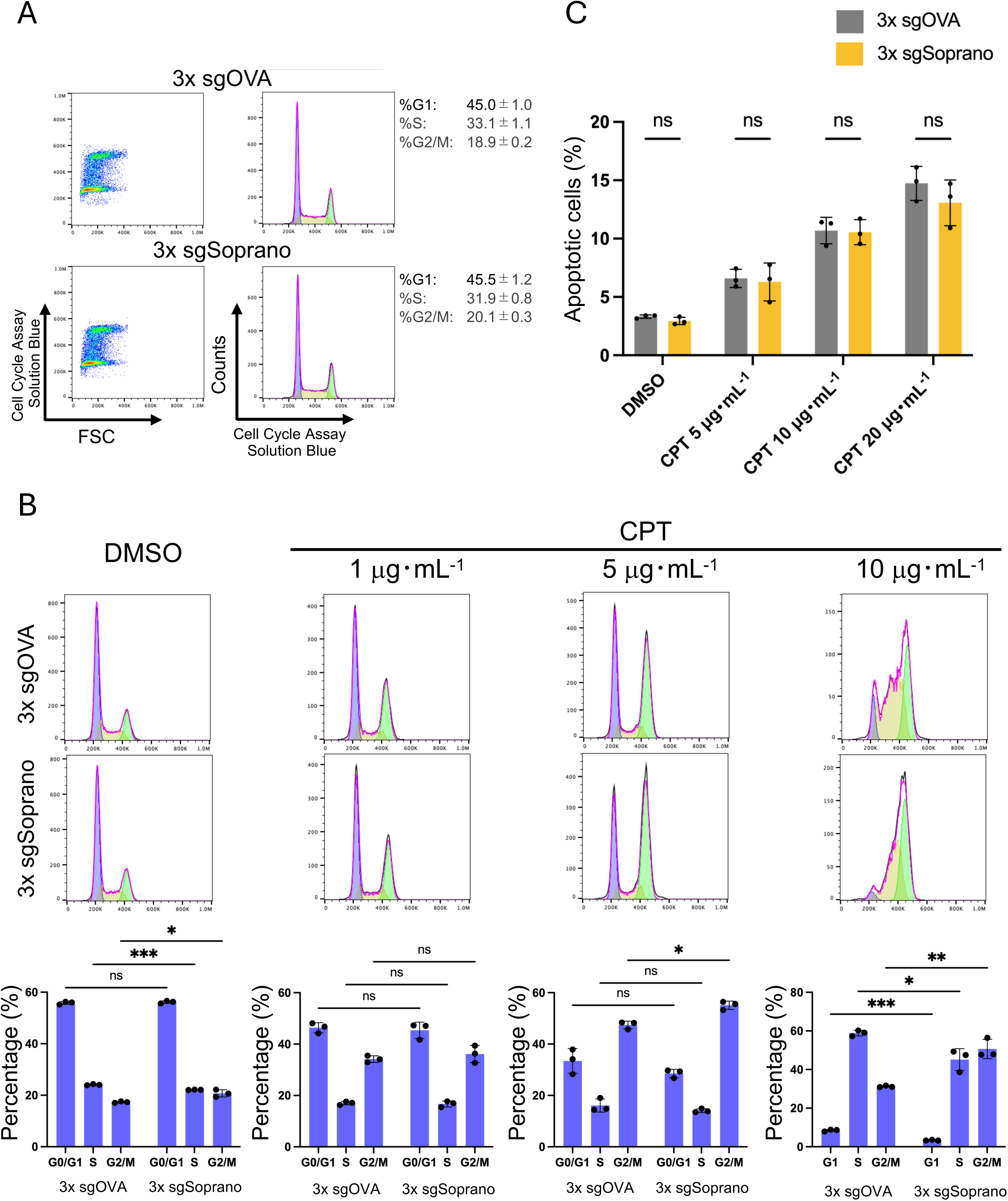
*Soprano* LTR CRISPRi affects the cell cycle of cultured PGCs in the presence of a DNA damage inducer. A) Flow cytometry diagram showing representative results of cell cycle assays on *Soprano* LTR CRISPRi PGCs. The values on the right side of the figure show the mean ± SD of three different biological replicates. B) Flow cytometry diagram showing representative results of cell cycle assays in cultured PGCs treated with camptothecin (CPT) at the indicated concentration under *Soprano* LTR CRISPRi conditions. The graphs on the bottom side of the figures show the mean ± SD of three different biological replicates. * p < 0.05, ** p < 0.01, *** p < 0.001, ns: not significant; multiple *t*-test. C) The percentage of apoptotic cells by the JC-1 assay. Data show the mean ± SD of three different biological replicates. ns: not significant; the Student’s *t*-test.

### *Soprano* LTR activity is critical for early chicken PGC development in ovo

To elucidate the role of *Soprano* in PGC development in ovo, we performed transplantation experiments on *Soprano* LTR CRISPRi PGCs (Fig. 6A). The EGFP-expressing 3× sgSoprano plasmid or mCherry-expressing 3× sgOVA control plasmid was transfected into CRISPRi PGCs and sorted by a cell sorter. One day after transfection, an equal number of EGFP-or mCherry-positive sorted PGCs were mixed and transplanted into EGK St.X [47] blastoderms to examine their effects on early PGC development, or into HH St.14-16 embryo blood to investigate their effects on the settlement of PGCs into the gonads from the blood and their further development. After a further incubation for 3.5 days, embryos were dissected and the genital ridges or gonads were harvested and stained with an anti-SSEA-1 antibody as a PGC marker. The percentage of EGFP-or mCherry-labeled cells among SSEA-1-positive cells was analyzed by flow cytometry, and the ratio of *Soprano* LTR CRISPRi (EGFP-positive) PGCs in all transplanted (sum of EGFP-positive and mCherry-positive) PGCs was calculated (Fig. 6B, C). The percentage of *Soprano* LTR CRISPRi PGCs colonizing the E3.5 genital ridges was lower than that of control (3× sgOVA transfected) cells when transplanted into blastoderms. On the other hand, they did not affect the colonization of E6 gonads when transplanted into HH St.14–16 embryos. These results suggest the importance of *Soprano* for the development of early-stage PGCs, whereas it was dispensable for their colonization into the genital ridges from the circulation.

**Figure 6.**
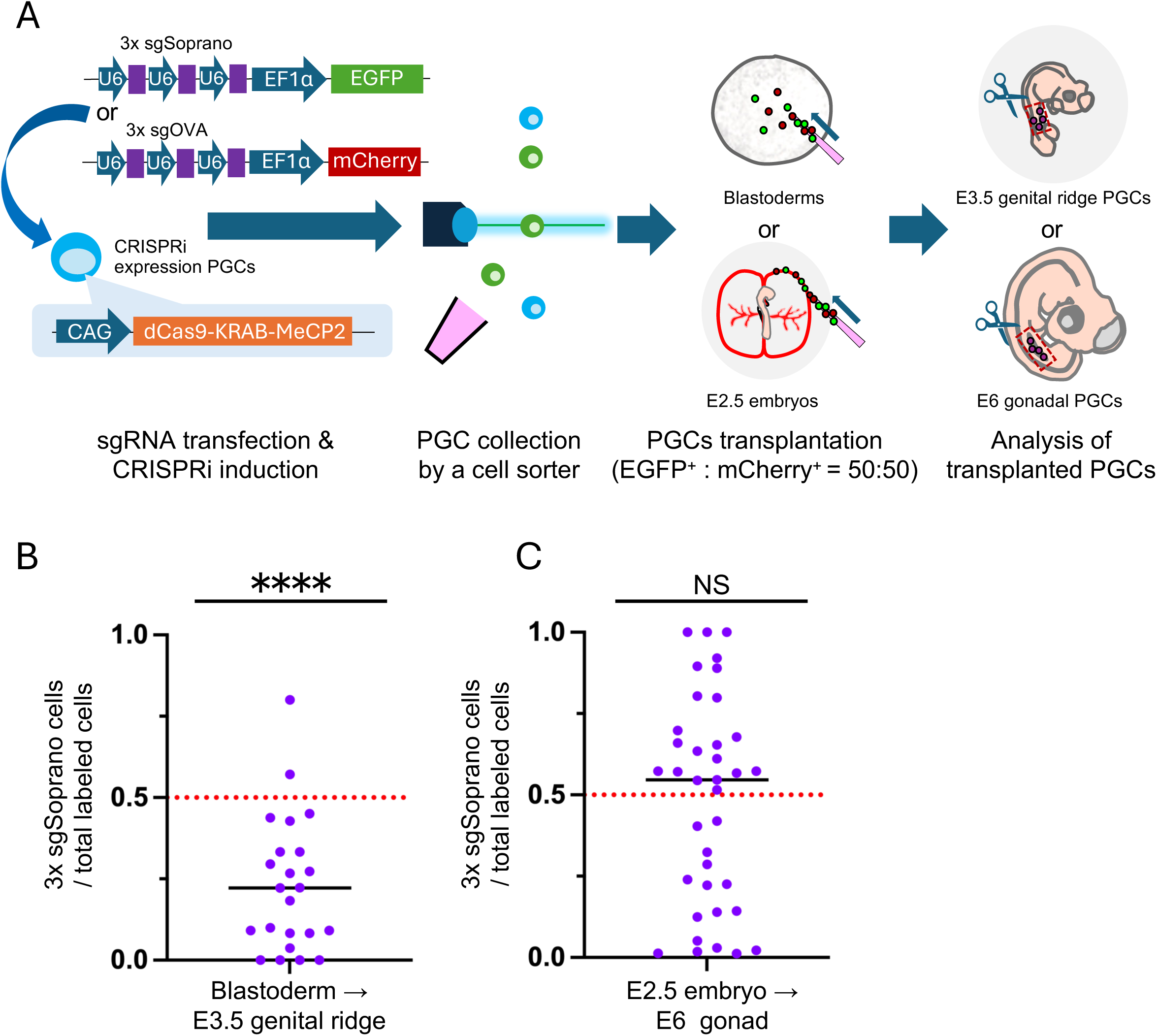
*Soprano* LTR CRISPRi inhibits PGC development in ovo. A) Experimental scheme of *Soprano* LTR CRISPRi PGC transplantation. B, C) Ratio of *Soprano* LTR CRISPRi PGCs among all transplanted PGCs settled in gonadal regions. (B) PGCs were transplanted into the EGK St. X blastoderm. (C) PGCs were transplanted into HH St.14-16 blood. After an incubation for 3.5 days, the genital ridges (B) or gonads (C) were harvested and stained with SSEA-1. SSEA-1-positive cells and EGFP-or mCherry-positive cells were analyzed by flow cytometry, and the ratio of *Soprano* LTR CRISPRi (EGFP-positive) cells in SSEA-1-positive and EGFP-or mCherry-positive PGCs was calculated. Data show individual embryo values and medians. The red dashed line shows the expected ratio of transplanted PGCs (set as 0.5). **** p < 0.0001, ns: not significant; one-sample Wilcoxon signed-rank test.

## Discussion

In this study, we performed transcriptome analyses of cultured PGCs derived from closed or inbred strains of chickens to elucidate why EE differs among chicken sexes and strains. Most of the DEGs associated with sex differences in cultured PGCs were encoded in sex chromosomes (Supplementary Fig. S2A-C), which is consistent with a previous study that compared transcriptomes between male and female cultured PGCs from broiler chickens [10]. The present results indicate that an inhibitory SMAD protein coding *SMAD7B*, which is encoded onto the W chromosome, was only expressed in female cultured PGCs. This difference in SMAD7B expression may be one of the factors affecting cell proliferation and cell-cell aggregation in female cultured PGCs through the modulation of the TGFβ/Activin signaling pathway based on sex differences in PGCs collected in ovo [10]. Although growth speeds were similar between our PGC lines after their establishment, sex differences in the transcriptome may have been maintained after cultivation. Differences in gene expression between chicken strains were greater than those between sexes (Fig. 1B). The GO analysis of DEGs in different strains of cultured PGCs accumulated several terms related to ion channels, transporters, and cell migration/cell adhesion (Fig. 1C). A previous study showed that reducing the calcium ion concentration in the PGC culture medium inhibited cell aggregation and increased culture efficiency, particularly in female PGCs [1]. We also observed that PGCs in some chicken strains were more likely to aggregate and attach to the plate, which induced cell differentiation and cell death during cell line establishment. It might be possible that the differences in cell-cell aggregation and adhesion by PGCs may have been due to differences in the expression of ion channels and transporters, which affected the early adaptation processes and EE of cultured PGCs.

Among DEGs between the strains, we selected *ENS-1/ERNI* for further analyses for several reasons. *ENS-1/ERNI* was specifically expressed in undifferentiated cells and was highly expressed in cultured PGCs established from various strains (Fig. 1F). Furthermore, the expression level of *ENS-1/ERNI* correlated with EE in cultured PGCs (Fig. 1G). Moreover, *ENS-1/ERNI* was highly expressed in earlier stage PGCs in ovo, suggesting its involvement in the early development of PGCs (Fig. 1E). *ENS-1/ERNI* and the UTR of *Soprano* LTR were also more highly expressed in cultured PGCs than in E2.5 blood circulating PGCs, the source of cultured PGCs, suggesting functions in the adaptation of endogenous PGCs to the in vitro environment (Fig. 1H). CRISPRi targeting the *Soprano* LTR was performed to comprehensively examine the function of *ENS-1/ERNI* genes and *Soprano* as cis-regulatory elements, and resulted in a broad reduction in gene expression involved in PGC homeostasis, including ribosomal proteins-, ER-associated proteins-, and mitochondria-related genes, and a induction of neural development-related genes (Fig. 2B). In addition, significant decreases in the expression levels of some protein-coding genes (*AGK*, *CDH7*, *ENOX1*, *GUCA1C*, *SLC16A7*, *TMPRSS15*, and *LOC418414*) were noted (Fig. 2D). Some of these genes had solo *Soprano* LTRs in their vicinity. Furthermore, *Soprano* LTR exhibited enhancer activity in a luciferase assay with cultured PGCs (Fig. 4E), suggesting that *Soprano* LTR functioned not only as a promoter of *ENS-1/ERNI*, but also acted as a distal enhancer of other genes. In addition to its function as a cis-element of *Soprano*, knockdown and rescue experiments on *ENS-1/ERNI* indicated that the ENS-1/ERNI protein independently up-regulated the expression of most of these genes (Fig. 3A, B). ENS-1/ERNI proteins have been shown to localize to both the cytoplasm and nucleus in undifferentiated cells, and act as transcriptional repressors in neural plate cells through a coiled-coil domain belonging to the SMC superfamily on the N terminus and a CBX3-binding domain on the C terminus [20,21]. Therefore, although ENS-1/ERNI acts as a transcriptional regulator in cultured PGCs, further investigations are required to clarify whether ENS-1/ERNI proteins regulate these genes directly or through other transcriptional and/or epigenetic regulators.

While *Soprano* LTR CRISPRi did not affect the cell cycle of healthy PGCs, it increased the percentage of G2/M phase-arrested cells in CPT-treated PGCs; however, there was no obvious change in the percentage of apoptotic cells (Fig. 5A-C). CPT has been reported to increase S phase-arrested cells and delay progression to the G2/M phase, if it does not induce apoptosis, through its activity to induce replication-mediated DNA damage and subsequently activate intra-S phase checkpoints [48]. These findings suggest that intra-S phase checkpoints were repressed by *Soprano* LTR CRISPRi, leading to premature entry into mitosis and a subsequent increase in G2/M phase-arrested cells. Notably, *Soprano* LTR CRISPRi significantly down-regulated *WEE2* expression in cultured PGCs. WEE2 is one of three Wee family kinases (WEE1, WEE2, and PKMYT1) in vertebrates that regulate cell cycle progression in G2/M phase checkpoints by inhibiting CDK1 activity through phosphorylation [49–52]. In addition, WEE1, which is widely expressed in somatic cells and regulates mitosis, mediates intra-S phase checkpoints by regulating CDK2 activity through phosphorylation independent of ATR/CHK1 activities. On the other hand, the role of WEE2, which is specifically expressed in the ovary and testis, is not well known, except for its induction of meiotic arrest in oocytes [50,52,53]: Mutations in WEE2 have been shown to cause infertility in humans due to oocyte maturation failure, although *Wee2* knockout only mildly affected the fecundity in mice [51,54]. Although the functions of WEE2 in PGCs were not previously reported, and in addition to *WEE2*, *WEE1* is also highly expressed in cultured chicken PGCs (Supplementary Fig. S7), we cannot exclude the possibility that *Soprano* activity partially regulates cell cycle progression via WEE2 and is involved in the quality control of genome stability in chicken PGCs.

The transplantation of cultured PGCs with *Soprano* LTR CRISPRi into the blood of HH St.14– 16 embryos did not affect subsequent gonadal settlement, whereas gonadal settlement was reduced when they were transplanted in blastoderms (Fig. 6B, C). These results suggest that *Soprano* and/or *ENS-1/ERNI* play important roles in the early-stage development of PGCs up to the blood circulation stage. Chicken PGCs are located in the central region of the area pellucida in EGK St.X blastoderms. They then migrate through the prospective neural plate region to the germinal crescent, located in the anterior part of the embryo outside the neural plate by HH St.4 [55], and begin intravasation around HH St.8 and circulate in the blood [56]. Previous studies indicated that ENS-1/ERNI controlled the timing of neurogenesis by regulating the expression of *SOX2* in the nascent neural plate [20,21]. In addition, our *Soprano* LTR CRISPRi experiments on cultured PGCs showed the up-regulated expression of genes related to neurogenesis. Therefore, *Soprano* and ENS-1/ERNI possibly suppress misdirected neurogenic signaling in in ovo PGCs during the gastrulation stage. On the other hand, the present results are based on the inhibition of *Soprano* LTR activity by CRISPRi and, thus, the comprehensive functions of *Soprano* as a cis-regulatory element plus ENS-1/ERNI proteins appear to have been included. Therefore, future studies need to address the individual contributions and detailed functions of these factors to in ovo PGC development and also in vitro cultured PGCs. Since PGCs are the fundamental cells for animal gametogenesis, they have multi-layered defense mechanisms to silence TEs, which cause genomic instability [57]. In chicken PGCs, the piRNA pathway suppresses TEs, and its down-regulation increases genomic DNA double-strand breakage by reactivating TEs [58]. In contrast, specific TEs escape these silencing mechanisms and organisms sometimes use TEs for their own purposes. For example, LTR retrotransposons also act as enhancers and are involved in human PGC specification [15]. Therefore, as with somatic cells and early-stage embryo development [11,13,14], TEs that are integrated into the host gene expression circuitry and are precisely regulated may play significant roles in PGC development. In the present study, we revealed that the *Galliformes*-specific LTR retrotransposon *Soprano* and its encoded protein ENS-1/ERNI regulated gene expression in cultured chicken PGCs and was involved in PGC development in ovo. Additionally, we demonstrated that *Soprano* acted as a cis-regulatory element regulated by NANOG and FGF signaling, and potentially functions as an enhancer in cultured chicken PGCs. In comparisons with other amniote animals, birds have evolved in a direction that has reduced their genome size, and the percentage of TEs in birds is very small [17]. Despite this evolutionary selection pressure, the small number of remaining TEs involved in the development of PGCs, as in mammals, is an important result when considering the relationship between TEs and animal evolution. On the other hand, since *Soprano* and *ENS-1/ERNI* are *Galliformes*-specific TEs, other TEs or different genes may contribute to the development of PGCs in other bird species, which is also interesting in terms of the diversity of germline development mechanisms involving TEs. In addition, we showed that variations in *Soprano* sequences affected its activity as a cis-regulatory element. Therefore, even within the same chicken species, polymorphisms in the *Soprano* sequence and possibly differences in insertion sites into the genome due to breed differences may affect the properties of PGCs. Collectively, these results provide valuable insights into establishing in vitro culture methods for avian PGCs and elucidating the mechanisms of avian PGC development and the evolution of PGCs in animals.

## Materials and Methods

### Chickens and eggs

Fertilized chicken eggs for the in vitro PGC culture were obtained from the Avian Bioscience Research Center, Nagoya University. Host eggs (white Leghorn) used for PGC transplantation experiments were purchased from the Takeuchi hatchery (Nara, Japan). All animal experiments were performed according to the ethical guidelines and approval for animal experimentation of Nagoya University.

### PGC culture medium

The medium for PGC cultivation was based on a previously reported method with some modifications [1]. Avian KO-DMEM basal medium (250 mOsm·kg^-1^, 12.0 mM glucose, and CaCl_2_ free) was custom ordered and purchased from Thermo Fisher Scientific. 1× B-27 minus vitamin A (Thermo Fisher Scientific), 1× NEAA (Thermo Fisher Scientific), 1× nucleosides (Merck), 1× monothioglycerol (Fujifilm), 2.0 mM GlutaMax (Thermo Fisher Scientific), 1.2 mM pyruvate (Thermo Fisher Scientific), 0.2% chicken serum (Thermo Fisher Scientific), 1× Penicillin-Streptomycin (Thermo Fisher Scientific), 0.2% ovalbumin (Merck), 0.2% sodium heparin (Merck), 10 μg·mL^-1^ ovotransferrin (Merck), 4 ng·mL^-1^ FGF2 (R&D Biosystems), and 25 ng·mL^-1^ Activin A (PeproTech) were added to avian KO-DMEM.

### Establishment of cultured PGCs

To establish various strains of chicken cultured PGCs, 1-2 μL of blood was collected from HH St.14-16 embryos and individually inoculated onto 96-well plates (WATSON) with 200 μL of the PGC culture medium. Half of the medium was changed every 2 or 3 days. When the total cell number reached approximately 1×10^4^ cells, cells were transferred to 24-well plates (IWAKI) and expanded to 2-3×10^5^ cells. Cells were frozen using CELLBANKER® 2 (Takara), stored in liquid nitrogen, and used in further experiments.

### Sex determination of PGCs and embryos

The sex of PGCs and embryos was identified as previously reported [59]. Briefly, genomic DNA was purified using easy DNA extraction kit version 2 (Kaneka). PCR was performed by KOD Fx Neo (Toyobo) under the following conditions: pre-denaturation at 95°C for 60 s, followed by 10 cycles at 98°C for 10 s, 60°C for 30 s (touchdown -1°C for each cycle), and 68°C for 30 s, and then 30 cycles at 98°C for 10 s, 50°C for 30 s, and 68°C for 30 s. The primers used are listed in Supplementary Table S15.

### Embryo transplantation of cultured PGCs and confirmation of germline transmission

The PGC lines established from GSP and GSN/1 chickens were used to confirm germline transmission. Regarding the transplantation of cultured PGCs, a 0.8-cm window was opened in the eggshell of a host embryo incubated for 52 hr and blood was collected using a glass capillary. After the sex of the host embryos was confirmed using blood samples, 1×10^4^ PGCs were transplanted into the blood of a sexually matched host embryo using glass capillaries. The window was sealed by scotch Ultra Transparent Tape (3M) and embryos were incubated until hatching. To confirm germline transmission, hatched germline chimeric chicks were raised to sexual maturity. Chimeric hens, into which PGC lines established from GSP were transplanted, were mated with GSP strain rooster. The fertilized eggs obtained from these hens were incubated and germline transmission was confirmed by the plumage color of hatched chicks. Chimeric hens, into which PGC lines established from GSN/1were transplanted, were mated with M/O strain rooster. The fertilized eggs obtained from these hens were incubated for 2 days, dissected, and tissues were collected. Genomic DNA was extracted using cell lysis buffer (50 mM KCl (Fujifilm), 10 mM Tris-HCl (pH 8.3) (Nacalai), 2.5 mM MgCl_2_ (Fujifilm), 0.45% NP-40 (Nacalai), 0.45% Tween-20 (Fujifilm), and 200 μg·mL^-1^ proteinase K (Nippongene)) and incubated at 56°C overnight. The Fayoumi breed-specific deletion on *SOX10* loci [60] was detected using PCR amplification. PCR was performed with KOD Fx NEO, and cycle conditions and the primers used were the same as those previously reported[60].

### Purification of blood circulating PGCs

To isolate PGCs circulating in the blood, blood was collected from HH St.14-16 M/O strain embryos and incubated with 10 μL of a BV421-labeled anti-SSEA-1 antibody (BD Biosciences, 562705) on ice for 1 hr in 1% FBS containing PBS(-). After washing with PBS(-), labeled cells were collected using a cell sorter (BD Biosciences, FACSJazz or Sony, sh800s) and subjected to a qRT-PCR analysis.

### CEF preparation

M/O strain chicken embryos incubated for 9 days were dissected, and the limbs, head, and organs were removed. Tissue was minced and digested in 0.1% trypsin-EDTA solution (Thermo Fisher Scientific). Isolated cells were filtered with a 100-μm cell strainer and inoculated onto 10-cm dishes (IWAKI) with DMEM high glucose (Fujifilm) supplemented with 10% FBS (Merck) and 1× Penicillin-Streptomycin. After 2-3 passages, cells were used in RNA-seq experiments.

### RT-qPCR

Total RNA was purified by ReliaPrep RNA Miniprep Systems (Promega) and cDNA was synthesized by the ReverTra Ace qPCR RT Master Mix (Toyobo). PCR was performed using the LightCycler 96 System (Roche) and THUNDERBIRD Next SYBR qPCR Mix reagent (Toyobo) under the following conditions: pre-denaturation at 95°C for 60 s, followed by 40 cycles at 95°C for 3 s, 60°C for 10 s, and 72°C for 30 s. The primers used are listed in Supplementary Table S15.

### RNA-seq data acquisition and mapping to the genome

Four strains (M/O, GSN/1, GSP, and PNP) of in vitro cultured male and female PGC lines established from a single embryo, CRISPRi-induced cultured PGCs established from the M/O strain, and CEFs were subjected to RNA-seq analyses. Total RNA was purified using the ReliaPrep RNA Cell Miniprep System. The NEBNext Poly(A) mRNA Magnetic Isolation Module and NEBNext UltraTMII Directional RNA Library Prep Kit were used for library preparation. Sequencing was performed by Illumina NovaSeq 6000 with 150-bp paired-end reads. The qualities of the acquired raw read data were assessed using fastp [61] (Version 0.23.2) and low-quality bases and adapter sequences were trimmed with default parameters. Trimmed reads were mapped to a chicken reference genome (GRCg7b, release 109) using HISAT2 [62] (Ver.2.2.1) with default parameters. The SAM files acquired were converted to BAM files using Samtools [63] (Ver.1.11). Publicly accessible embryonic PGC RNA-seq data [33] were downloaded from the NCBI SRA database and analyzed with the same pipeline as described above.

### Bioinformatical analyses

The number of mapped reads on genes were counted using featureCounts [64] (Ver.2.0.1). Raw read counts were filtered to exclude genes with low counts and normalized with default parameters, and a DEG analysis was performed using edgeR [65] (Version 3.40.2). A GO analysis was conducted using the g:Profiler online tool [66]. The transcripts per million (TPM) of each transcript was analyzed using TPMCalculator [67] (Version 0.0.3). A consensus motif analysis of *Soprano* LTR was performed using the MEME suite online tool [36]. A transcription factor-binding site prediction analysis was conducted using the JASPAR online tool [39] with human and mouse CORE transcription factor collection and default parameters.

### *ENS-1/ERNI* knockout and cell growth analysis

The Cas9 and gRNA expression vectors were constructed from pX330_CBH::spCas9-T2A-Puro^R^-pA_mU6::sgRNA (a vector in which the mCherry gene of Addgene #64324 [68] was replaced with a puromycin-resistant gene). This plasmid was digested by BbsI-HF (NEB) and ligated with a pre-annealed sgRNA spacer sequence containing oligo DNA to make pX330_CBH::spCas9-T2A-PuroR-pA_mU6::sgRNA (ENS-1/ERNI #1∼#3). In the cell growth analysis, 10 μg of these vectors was transfected using Lipofectamine 2000 (Thermo Fisher Scientific) into 1×10^6^ of cultured PGCs. PGCs were then cultured for 24 hr, and selected with 1 μg·mL^-1^ of puromycin (Fujifilm) for another 24 hr. After removing dead cells using the Viahance Dead Cell Removal Kit (BioPal), PGCs were counted and 2×10^4^ cells were re-inoculated into 48-well plates (IWAKI). After the indicated culture period, the number of cells was counted in each culture period using the Countess 3 Cell Counter (Thermo Fisher Scientific).

### CRISPRi

The dCas9-KRAB-MeCP2 gene sequence was amplified by KOD One (Toyobo) from the Addgene #110821 plasmid [35], and cloned into a blasticidin S-resistant gene expressing a piggyBac vector by Gibson assembly cloning using the NEBuilder HiFi DNA Assembly Master Mix to make pPV_CAG::dCas9-KRAB-MeCP2-pA_EF1α::Bsd^R^-pA. This plasmid and piggyBac transposase expression plasmid were transfected into the M/O strain of cultured PGCs using Lipofectamine 2000, and stably transfected PGCs were selected by 10 μg·mL^-1^ of blasticidin S (Fujifilm) for two weeks. Regarding in vitro-synthesized sgRNA transfection, 20 pmol of Alt-R CRISPR-Cas9 sgRNA purchased from IDT was transfected using Lipofectamine RNAiMAX (Thermo scientific) into 1×10^5^ of CRISPRi-expressing cultured PGCs. PGCs were cultured for 24 hr and then subjected to further analyses. To construct 3× sgRNA expression plasmids, each sgRNA spacer sequence was cloned into the pPV_ mU6::sgRNA_ EF1α::EGFP (or mCherry)-IRES-Puro^R^-pA plasmid using the same method as that described above. Each mU6::sgRNA cassette was amplified using KOD One (Toyobo) and cloned by Gibson assembly cloning using the NEBuilder HiFi DNA Assembly Master Mix to make pPV_3×(mU6::sgSoprano)_EF1α::EGFP-IRES-Puro^R^-pA or pPV_3×(mU6::sgOVA)_EF1α::mCherry-IRES-Puro^R^-pA. In the transfection of sgRNA expression plasmids, 10 μg of these plasmids was introduced into 1×10^6^ of CRISPRi-expressing cultured PGCs using Lipofectamine 2000. PGCs were then cultured for 48 hr, and EGFP-or mCherry-positive cells were collected using a cell sorter (Sony, sh800s) and subjected to further analyses.

### ERNI rescue experiments

*ERNI* cDNA was amplified by KOD One from a PGC line of the M/O strain and cloned by replacing the EGFP gene in the pcDNA4_CMV::EGFP-pA plasmid by Gibson assembly cloning using the NEBuilder HiFi DNA Assembly Master Mix to make pcDNA4_CMV::ERNI-pA. Regarding sgRNA expression plasmids and *ERNI* expression plasmid transfection, 5 μg of the pPV_3×(mU6::sgSoprano)_EF1α::EGFP-IRES-Puro^R^-pA or pPV_3×(mU6::sgOVA)_EF1α::mCherry-IRES-Puro^R^-pA plasmid and 5 μg of the pcDNA4_CMV::EGFP-pA or pcDNA4_CMV::ERNI-pA plasmid were introduced into 1×10^6^ of CRISPRi-expressing cultured PGCs using Lipofectamine 2000. PGCs were then cultured for 48 hr, and EGFP-or mCherry-positive cells were collected using a cell sorter and subjected to further analyses.

### siRNA treatment

DsiRNAs targeted to *ENS-1/ERNI* (Supplementary Table S15) and the negative control DsiRNA were purchased from IDT. A total of 20 pmol of DsiRNAs was transfected using Lipofectamine RNAiMAX into 1×10^5^ of cultured PGCs established from the M/O or GSP strain. PGCs were then cultured for 48 hr and subjected to further analyses.

### Luciferase assay

*NANOG* and *PRDM1* cDNAs were amplified by KOD Fx Neo from the blastodermal cells of M/O strain embryos and cloned into a pcDNA4_CMV::EGFP-pA vector by restriction enzyme digestion and ligation to make pcDNA4_CMV::EGFP-NANOG-pA or pcDNA4_CMV::EGFP-PRDM1-pA. *Soprano* sequences were amplified by KOD One (Toyobo) from the genomic DNA of cultured PGCs established from the M/O strain and cloned into a pNL1.1[Nluc] vector (Promega) by Gibson assembly cloning using the NEBuilder HiFi DNA Assembly Master Mix to make pNL1.1_Soprano::Nluc-pA. Soprano deletion and point mutants were made from these vectors by inverse PCR using KOD One. These plasmid vectors (100 ng) and the control firefly luciferase expression pGL4.54[luc2/TK] vector (10 ng) (Promega) were transfected into 1.5×10^4^ cells of cultured PGCs from the M/O strain using Lipofectamine 2000 and inoculated onto 96-well plates. In DF-1 cells, 1.5×10^4^ cells were seeded onto 96-well plates the day before transfection, and the same amount of vectors was transfected using Lipofectamine 3000 (Thermo Fisher Scientific). To investigate the effects of transcription factors, 100 ng of transcription factor expression vectors was transfected together with luciferase-expressing vectors. After 24 hr, luciferase activity was measured using the Nano-Glo Dual-Luciferase reporter assay system (Promega) and TECAN Infinite200PRO (Tecan) microplate reader.

### Inhibition of activin A and FGF2 signaling

The M/O strain of cultured PGCs was seeded onto the fresh PGC culture medium described above in the presence of each inhibitor (1 mM A83-01 (Fujifilm) or 1 mM PD0325901(Fujifilm)) or vehicle (DMSO (Fujifilm), 0.15%) for the indicated period and then subjected to further analyses.

### Cell cycle assay

In the cell cycle assay, 10 μg of the pPV_3×(mU6::sgSoprano)_EF1α::EGFP-IRES-Puro^R^-pA or pPV_3×(mU6::sgOVA)_EF1α::mCherry-IRES-Puro^R^-pA plasmid was introduced into 1×10^6^ of CRISPRi-expressing PGCs using Lipofectamine 2000. PGCs were cultured for 48 hr and collected using a cell sorter based on fluorescence. Sorted cells were stained by Cell Cycle Assay Solution Blue (Dojindo) at 37°C for 15 min. Stained cells were re-analyzed using a cell sorter. To investigate the effects of DNA damage inducers on the cell cycle, one day after sgRNA expression plasmid transfection, CPT (Fujifilm) was applied at a predetermined dose for 1 hr. After washing with PBS(-), cells were cultured for a further 24 hr for the cell cycle analysis.

### JC-1 assay

The pPV_3×(mU6::sgSoprano)_EF1α::BFP-WPRE-pA and pPV_3×(mU6::sgOVA)_EF1α::BFP-WPRE-pA plasmids were constructed from the pPV_3×(mU6::sgSoprano)_EF1α::EGFP-IRES-Puro^R^-pA or pPV_3×(mU6::sgOVA)_EF1α::mCherry-IRES-Puro^R^-pA plasmid, respectively, by replacing the EGFP (or mCherry)-IRES-Puro^R^-pA cassette to BFP-WPRE-pA by PCR amplification using KOD One and Gibson assembly cloning using the NEBuilder HiFi DNA Assembly Master Mix. In the JC-1 assay, 10 μg of these plasmids was introduced into 1×10^6^ of CRISPRi-expressing PGCs using Lipofectamine 2000. PGCs were cultured for 24 hr and then treated with CPT at a predetermined dose for 5 hr. Cells were stained with the MitoPT JC-1 Assay kit (ImmunoChemistry Technologies) and BFP-positive apoptotic cells were analyzed using a cell sorter.

### *Soprano* LTR CRISPRi PGC transplantation

Regarding sgRNA expression plasmid transfection, 8 μg of pPV_3×(mU6::sgSoprano)_EF1α::EGFP-IRES-Puro^R^-pA or pPV_3×(mU6::sgOVA)_EF1α::mCherry-IRES-Puro^R^-pA and 2 μg of the piggyBac transposase expression plasmid were introduced into 1×10^6^ of CRISPRi-expressing PGCs using Lipofectamine 2000. After PGCs were cultured for 24 hr, EGFP- or mCherry-positive cells were collected using a cell sorter. A total of 2 to 3×10^3^ sorted EGFP- or mCherry-positive PGCs were injected into the subgerminal cavity of EGK St.X blastoderms or the blood vessels of HH St.14-16 embryos. The eggshell was sealed with a PTFE membrane (Kokugo) and plastic wrap (Asahi KASEI) (blastoderms) or scotch tape (3M) (HH St.14-16 embryos). Injected embryos were incubated sealed side down (blastoderms) or up (HH St.14-16 embryos) at 38°C under 60% humidity with a rocking angle of 90° (blastoderms) or 30° (HH St.14-16 embryos) every hr. After 3.5 days, embryos were sacrificed and embryonic genital ridges or gonads were isolated. The tissues were dissociated by TrypLE Express (Thermo Fisher Scientific), filtrated through 100-μm cell strainers, and stained with 3 μL of a BV421-labeled anti-SSEA-1 antibody (BD Biosciences, 562705) on ice for 1 hr in 1% FBS containing PBS(-). The ratio of SSEA-1- and EGFP- or mCherry-positive cells was analyzed using a flow cytometer.

### Data availability

Raw sequencing data from the present study are publicly available in the DDBJ BioProject (PRJDB35733). Other datasets and plasmids used in this study are available from the corresponding author upon reasonable request.

## Acknowledgments

This work was supported a “Strategic Research Projects” grant from ROIS (Research Organization of Information and Systems) (A.K), Grant-in-Aid for Scientific Research KAKENHI (C) 22K15027 (Y. Okuzaki), and 23K05798 (K.N.), and MEXT National BioResource Project JPNBRP202205 (K.N.). The piggyBac plasmid and piggyBac transposase expression plasmid were kindly gifted from Dr. Akitsu Hotta [69].

## Author Contributions

Conceptualization: Y. Okuzaki and K.N.; methodology: Y. Okuzaki and A.K; investigation: Y. Okuzaki, A.K., Y. Ozaki, and T.U.; supervision: S.D and K.N.; writing -original draft: Y. Okuzaki and K.N.; writing - review & editing: Y. Okuzaki, A.K., D.S., and K.N.

## Competing Interests

The authors declare no competing interests.

## Notes

### Competing Interest Statement

The authors have declared no competing interest.

