## Supplementary Figures for "The chicken retrovirus-like gene *ENS-1/ERNI* and its LTR *Soprano* are involved in primordial germ cell development"

**Supplementary Figure S1. Confirmation of PGC marker gene expression and the germline transmission of cultured PGC lines.**

A) The RNA from in vitro cultured PGCs established from each chicken strain was subjected to qRT-PCR. The expression levels of each PGC marker gene were normalized by those of *GAPDH*. Data show the mean ± SD of individual clones.

B) Representative photo of transplanted PGC-derived offspring identified by the plumage color. Germline chimeric white Leghorn hens transplanted with cultured PGCs established from the GSP strain were crossed with roosters of the GSP strain to produce offspring. Chicks derived from transplanted PGCs have a brown plumage, while chicks derived from white Leghorn with dominant white genes have a black-flecked yellow plumage. The chick derived from transplanted PGCs is indicated by the arrow.

**Supplementary Figure S2. RNA-seq analysis of cultured PGCs of males and females derived from several strains.**

A) A volcano plot showing an RNA-seq expression analysis comparing male and female cultured PGCs. Dashed lines indicate $-\text{Log}_{10}$ p-adj > 2 or |Log2 FC| > 1.

B) A pie chart showing the percentage of chromosomal locations of DEGs (p-adj < 0.01, |FC| > 2) in male and female cultured PGCs.

C) Violin plots showing the ratio of gene expression levels in male and female cultured PGCs for genes located on chromosome Z or autosomes. Genes that were identified as low expression genes in the edgeR analysis were excluded from the graph.

D) A figure showing the ratio of gene expression levels in male and female cultured PGCs and the chromosomal location of genes on chromosome Z. Genes that were identified as low expression genes in the edgeR analysis were excluded from the graph.

E) A Benn plot showing the inclusive relation of DEGs comparing cultured PGCs from the M/O strain and Fayoumi strains versus DEGs that were more highly expressed in cultured PGCs from the M/O strain than CEFs (p-adj < 0.01, FC > 1.5) from the M/O strain.

**Supplementary Figure S3. Transcriptome analyses of *Soprano* LTR CRISPRi PGCs**

A) Schematic diagram showing the *ENS-1/ERNI* gene structure and sgRNA location for CRISPRi (red arrows).

B) A volcano plot showing an RNA-seq expression analysis comparing synthesized sgRNA (sgOVA vs sgSoprano#1 and #2)-transfected CRISPRi expression PGCs. Dashed lines indicate $-\mathrm{Log}_{10}$ p-adj > 1.3 (p-adj < 0.05) or |Log2 FC| > 0.58 (|FC| > 1.5).

C) Schematic diagram showing the chromosomal location of gene sets that showed significant expression variations in *Soprano* LTR CRISPRi PGCs. DEGs in *Soprano* LTR CRISPRi are shown in blue arrows. *Soprano*-like sequences present in the surrounding region are shown in red. Red arrows indicate the direction of *Soprano*-like sequences (5'-3').

D) An overview of RNA-seq reads mapping on the *AGK/DENND11/WEE2* locus.

E) A strand-specific count analysis of the *DENND11* gene. Data show the mean ± SD of three biologically different samples.

F) Expression levels of each gene in each embryonic stage of in ovo PGCs based on an RNA-seq expression re-analysis [1].

**Supplementary Figure S4. Cell proliferation assay of cultured PGCs under *ENS-1/ERNI* knockout conditions**

A) Schematic diagram showing the *ENS-1/ERNI* gene structure and sgRNA locations for gene knockout (orange arrows) and DsiRNAs for knockdown (red arrows).

B) PGC number after the knockout of *ENS-1/ERNI* by CRISPR-Cas9. Data show the mean ± SD of three different biological replicates. *** $p < 0.001$, **** $p < 0.0001$; ANOVA with Dunnett's post hoc test.

**Supplementary Figure S5. *ENS-1/ERNI* knockdown experiments on a cultured PGC line established from the GSP strain.**

The RNA from DsiRNA-transfected cultured PGCs established from the GSP strain was subjected to qRT-PCR. The expression levels of each gene were normalized by *GAPDH*, and that of control DsiRNA was set as 1. Data show the mean ± SD of eight biologically different samples. ** $p < 0.01$, **** $p < 0.0001$; the Student's *t*-test.

**Supplementary Figure S6. Investigation of *Soprano* LTR activities.**

A, C, D, F) Dual luciferase assay investigating *Soprano* LTR activities. (A) Promoter activity of various clones of *Soprano* LTR in cultured PGCs. (C) Activity of deletion mutants of *Soprano* LTR in cultured PGCs. (D) Cell specificity of *Soprano* LTR in cultured PGCs and DF-1. (F) The effects of NANOG and the enhancer activity of the motif 7 core sequence of *Soprano* LTR in DF-1. The expression level of NanoLuc (NLuc) was normalized by that of firefly luciferase (FLuc). Data show the mean ± SD of three different biological replicates (C,D,F). In (A), the three different experimental results were combined after adjusting the lot differences by the expression levels of NLuc activity under the TK promoter measured in each experiment and are showed as the mean ± SD. * $p < 0.05$, *** $p < 0.001$; **** $p < 0.0001$, ns: not significant; ANOVA with Dunnett's post hoc test was used to compare each clone sample individually (C). The Student's *t*-test (D). ANOVA with Tukey's multiple comparisons post hoc test (F).

B) Consensus motif analysis using MEME suite.

E) Expression levels of each GATA family gene in cultured PGCs based on an RNA-seq expression analysis. Data show the mean ± SD of eight biologically different samples (male and female GSP, GSN/1, PNP, and M/O). The *GATA1* gene was not mapped in the current chicken genome database.

**Supplementary Figure S7. *WEE1* expression in cultured PGCs.**

A) Expression levels of *WEE1* and *WEE2* in cultured PGCs based on an RNA-seq expression analysis. Data show the mean ± SD of eight biologically different samples (male and female GSP, GSN/1, PNP, and M/O).

B) The 3× sgOVA or 3× sgSoprano expression plasmid was transfected into CRISPRi PGCs, and RNA purified from them was subjected to qRT-PCR. The expression levels of *WEE1* were normalized by *GAPDH* and that of 3× sgOVA was set as 1. Data show the mean ± SD of four different biological replicates. ns: not significant; the Student's *t*-test.

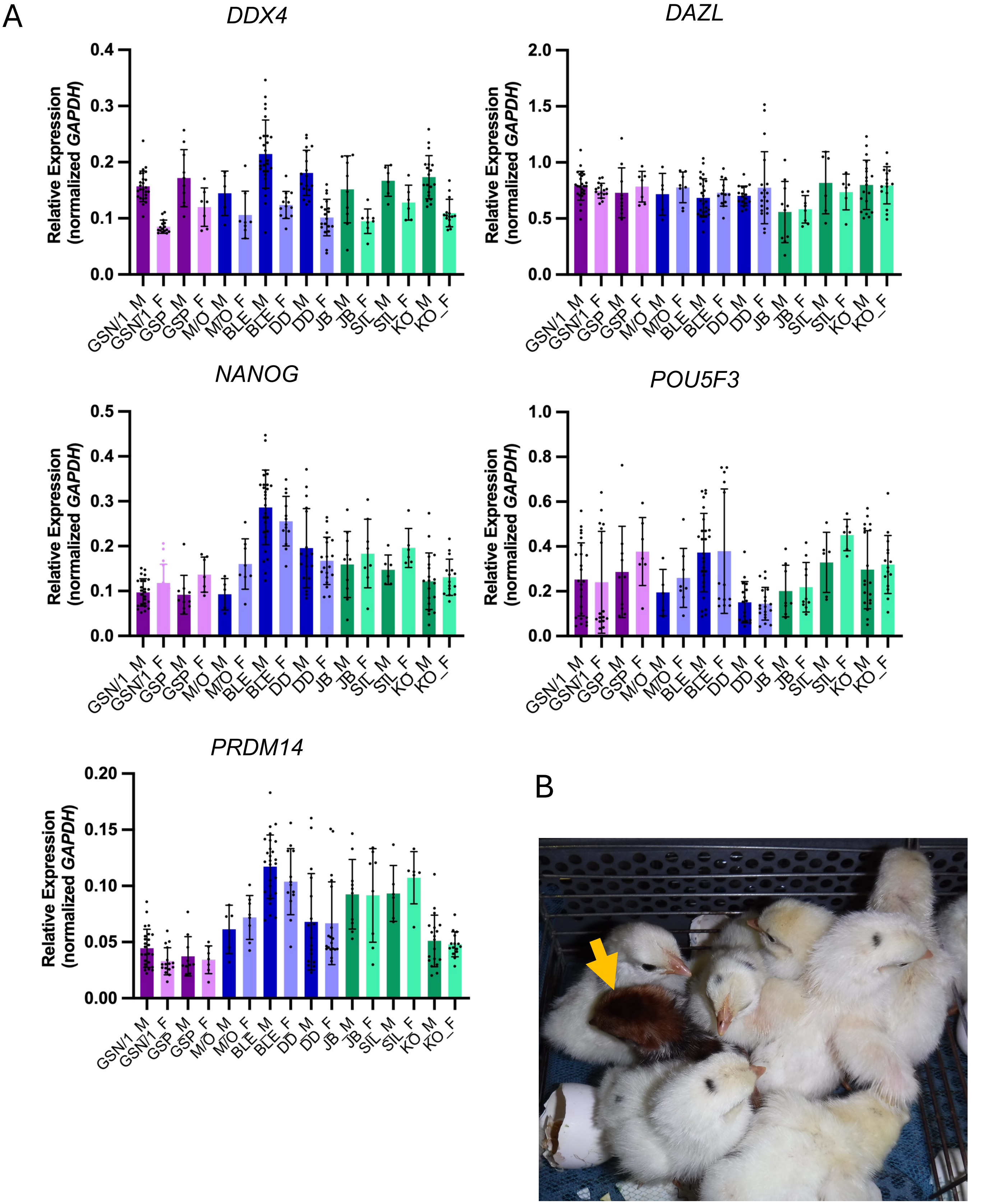
A
DDX4
DAZL
NANOG
POU5F3
PRDM14
Relative Expression
(normalized GAPDH)
0.0
0.1
0.2
0.3
0.4
0.5
0.5
1.0
1.5
2.0
0.6
0.8
0.00
0.05
0.10
0.15
0.20
GSN/1_M
GSN/1_F
GSP_M
GSP_F
M/O_M
M/O_F
BLE_M
BLE_F
DD_M
DD_F
JB_M
JB_F
SIL_M
SIL_F
KO_M
KO_F
B

Fig. S1

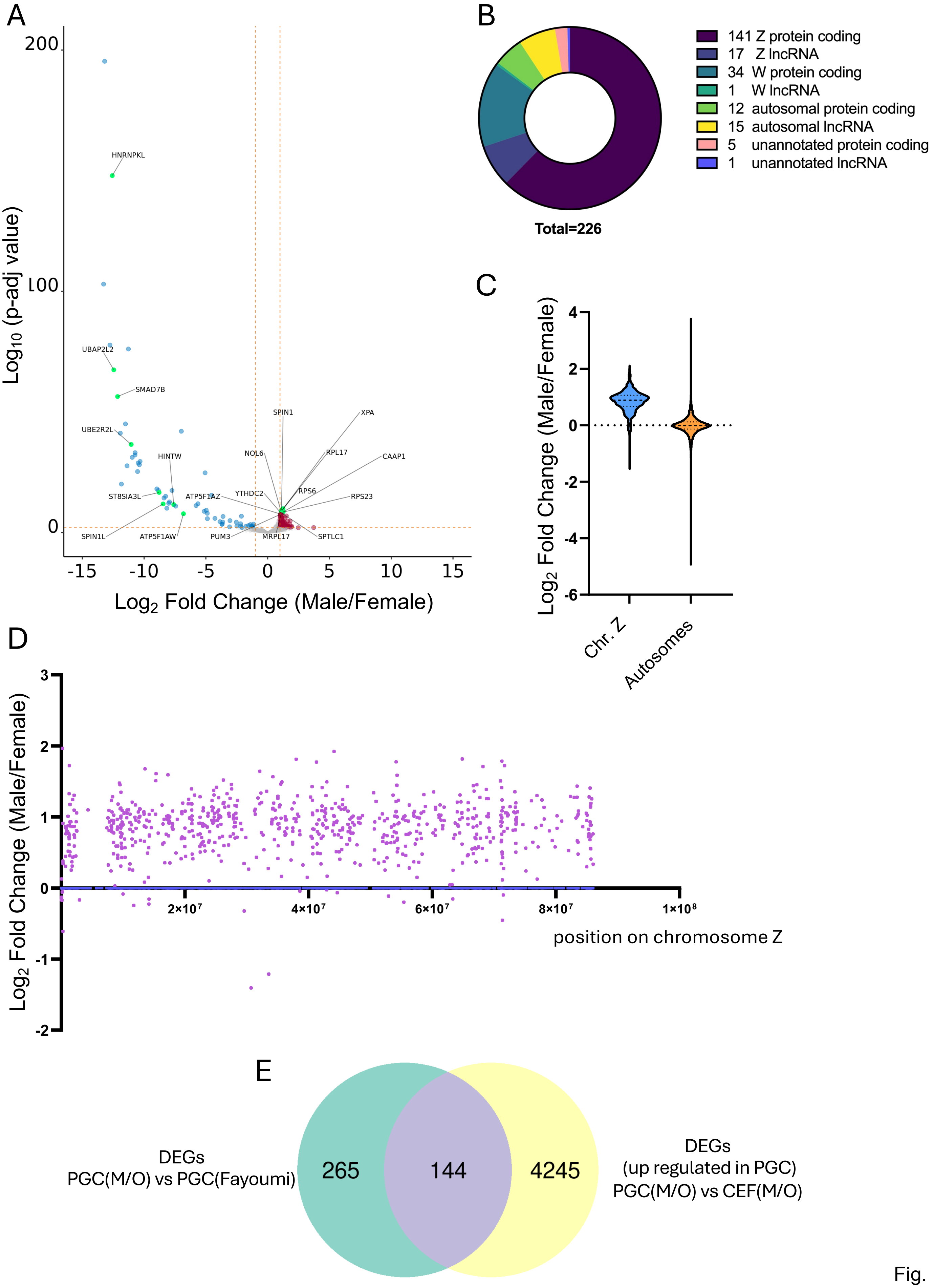
A
B
C
D
E
Log10 (p-adj value)
Log2 Fold Change (Male/Female)
HNRNPKL
UBAP2L2
SMAD7B
UBE2R2L
HINTW
ST8SIA3L
ATP5F1AZ
SPIN1L
ATP5F1AW
PUM3
MRPL17
SPTLC1
SPIN1
NOL6
YTHDC2
RPS6
RPL17
XPA
CAAP1
RPS23
141 Z protein coding
17 Z lncRNA
34 W protein coding
1 W lncRNA
12 autosomal protein coding
15 autosomal lncRNA
5 unannotated protein coding
1 unannotated lncRNA
Total=226
Chr. Z
Autosomes
position on chromosome Z
2×10^7
4×10^7
6×10^7
8×10^7
1×10^8
DEGs
PGC(M/O) vs PGC(Fayoumi)
265
144
4245
DEGs
(up regulated in PGC)
PGC(M/O) vs CEF(M/O)

Fig. S2

A

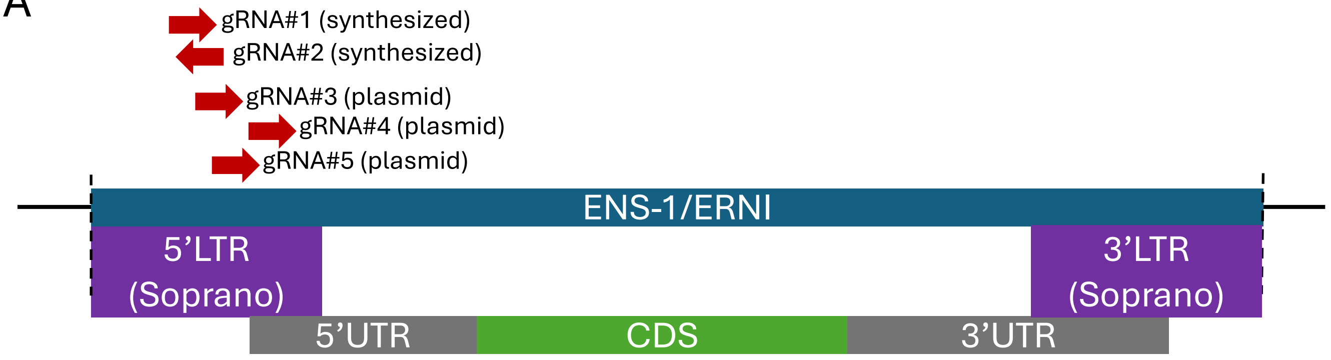

gRNA#1 (synthesized)
gRNA#2 (synthesized)
gRNA#3 (plasmid)
gRNA#4 (plasmid)
gRNA#5 (plasmid)
ENS-1/ERNI
5'LTR (Soprano)
3'LTR (Soprano)
5'UTR
CDS
3'UTR


B

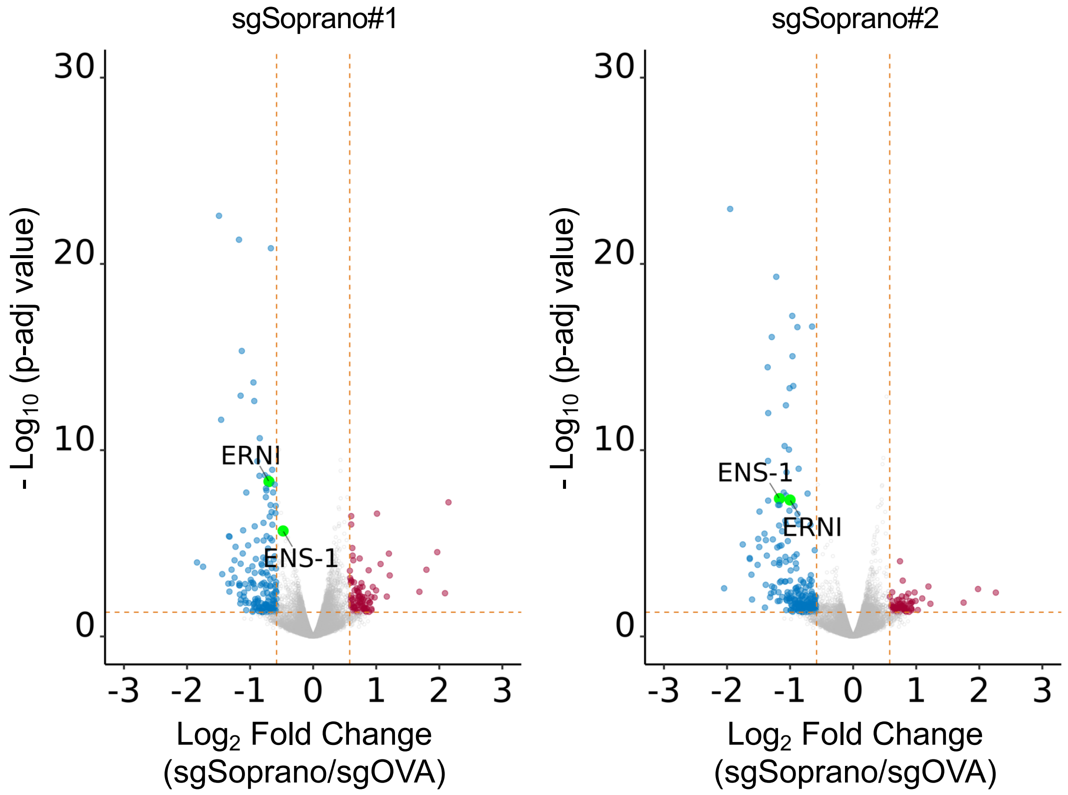

sgSoprano#1
sgSoprano#2
- Log10 (p-adj value)
30
20
10
0
ERNI
ENS-1
-3 -2 -1 0 1 2 3
Log2 Fold Change (sgSoprano/sgOVA)


Fig. S3

C

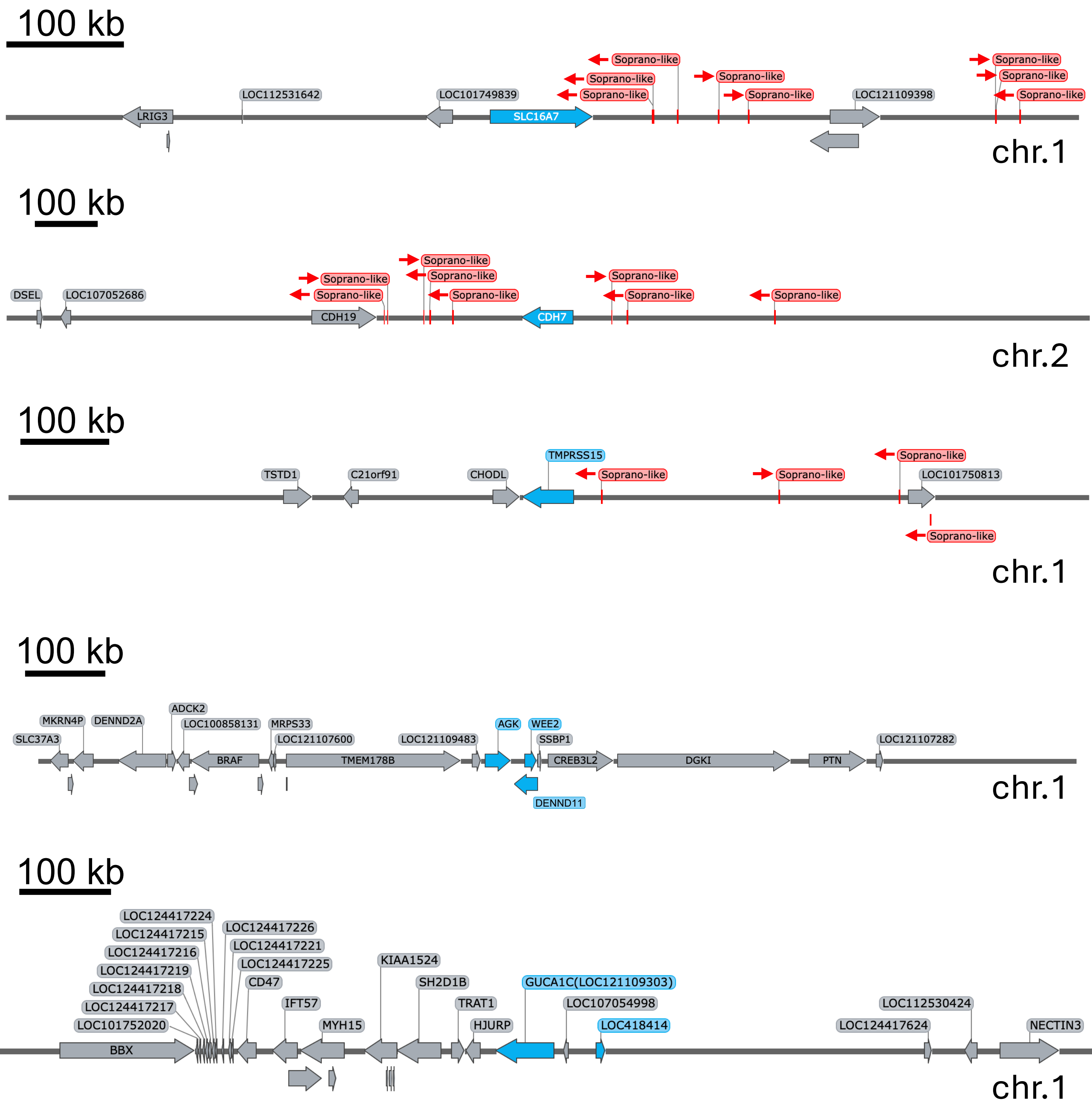

100 kb
Soprano-like
LRIG3
LOC112531642
LOC101749839
SLC16A7
LOC121109398
chr.1
100 kb
DSEL
LOC107052686
CDH19
CDH7
Soprano-like
chr.2
100 kb
TSTD1
C21orf91
CHODL
TMPRSS15
LOC101750813
Soprano-like
chr.1
100 kb
SLC37A3
MKRN4P
DENND2A
ADCK2
LOC100858131
BRAF
MRPS33
LOC121107600
TMEM178B
LOC121109483
AGK
WEE2
SSBP1
DENND11
CREB3L2
DGKI
PTN
LOC121107282
chr.1
100 kb
LOC124417224
LOC124417215
LOC124417216
LOC124417219
LOC124417218
LOC124417217
LOC101752020
BBX
LOC124417226
LOC124417221
LOC124417225
CD47
IFT57
MYH15
KIAA1524
SH2D1B
TRAT1
HJURP
GUCA1C(LOC121109303)
LOC107054998
LOC418414
LOC124417624
LOC112530424
NECTIN3
chr.1


Fig. S3

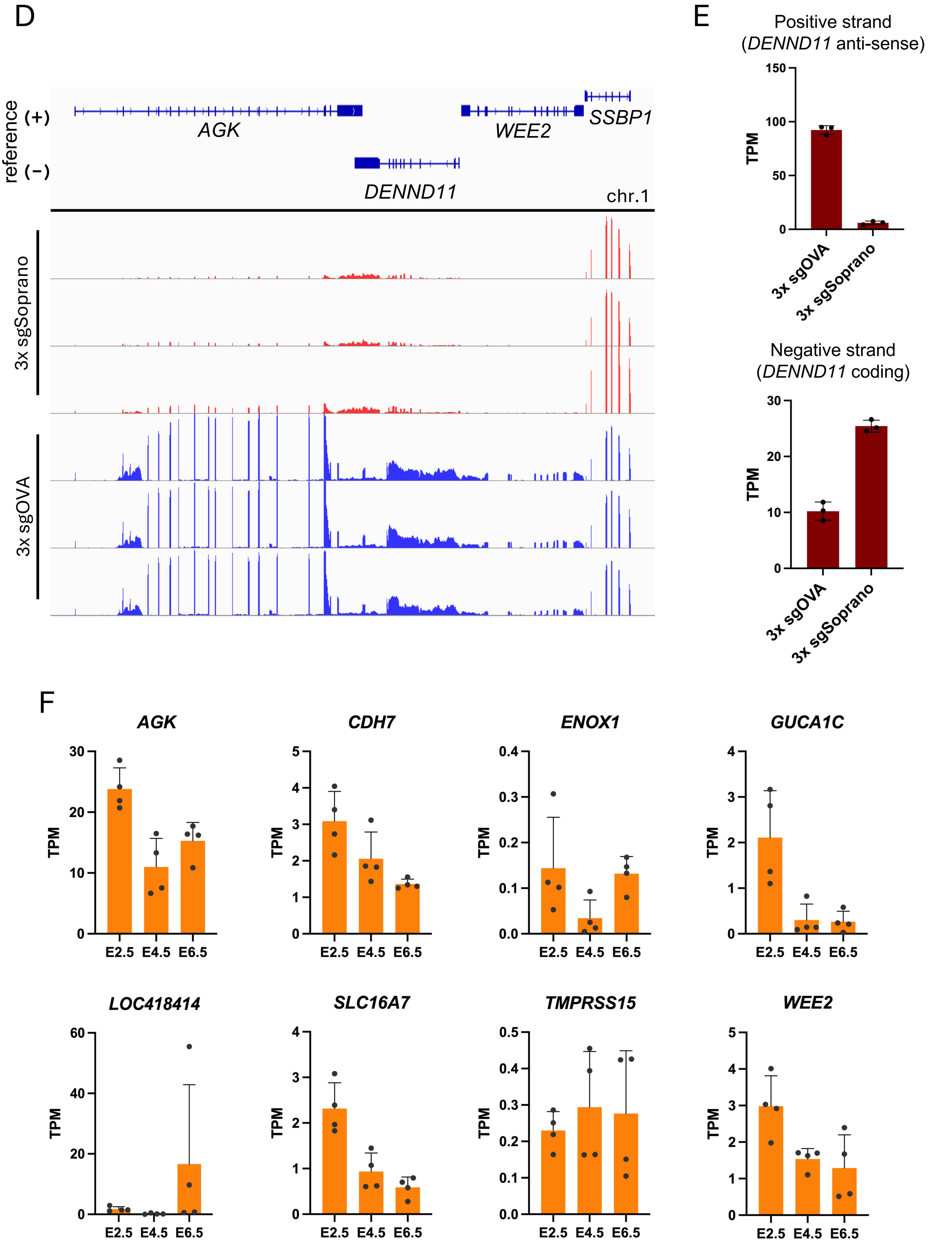
D
reference
(+)
(−)
AGK
WEE2
SSBP1
DENND11
chr.1
3x sgSoprano
3x sgOVA
E
Positive strand
(DENND11 anti-sense)
150
100
50
0
TPM
3x sgOVA
3x sgSoprano
Negative strand
(DENND11 coding)
30
20
10
0
TPM
3x sgOVA
3x sgSoprano
F
AGK
CDH7
ENOX1
GUCA1C
LOC418414
SLC16A7
TMPRSS15
WEE2
TPM
E2.5
E4.5
E6.5

Fig. S3

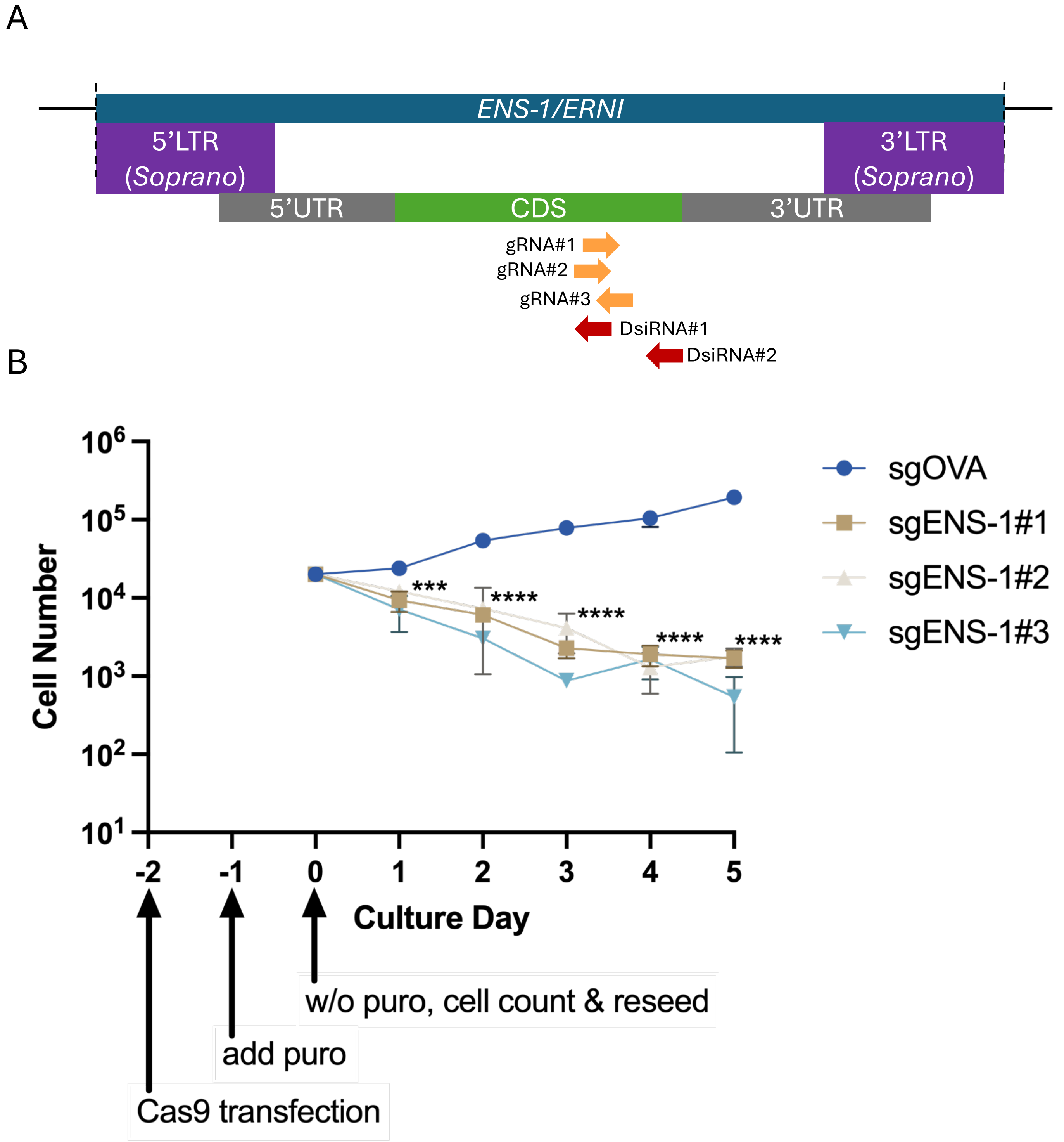

A
ENS-1/ERNI
5'LTR (Soprano)
3'LTR (Soprano)
5'UTR
CDS
3'UTR
gRNA#1
gRNA#2
gRNA#3
DsiRNA#1
DsiRNA#2
B
Cell Number
Culture Day
sgOVA
sgENS-1#1
sgENS-1#2
sgENS-1#3
***
****
w/o puro, cell count & reseed
add puro
Cas9 transfection


Fig. S4

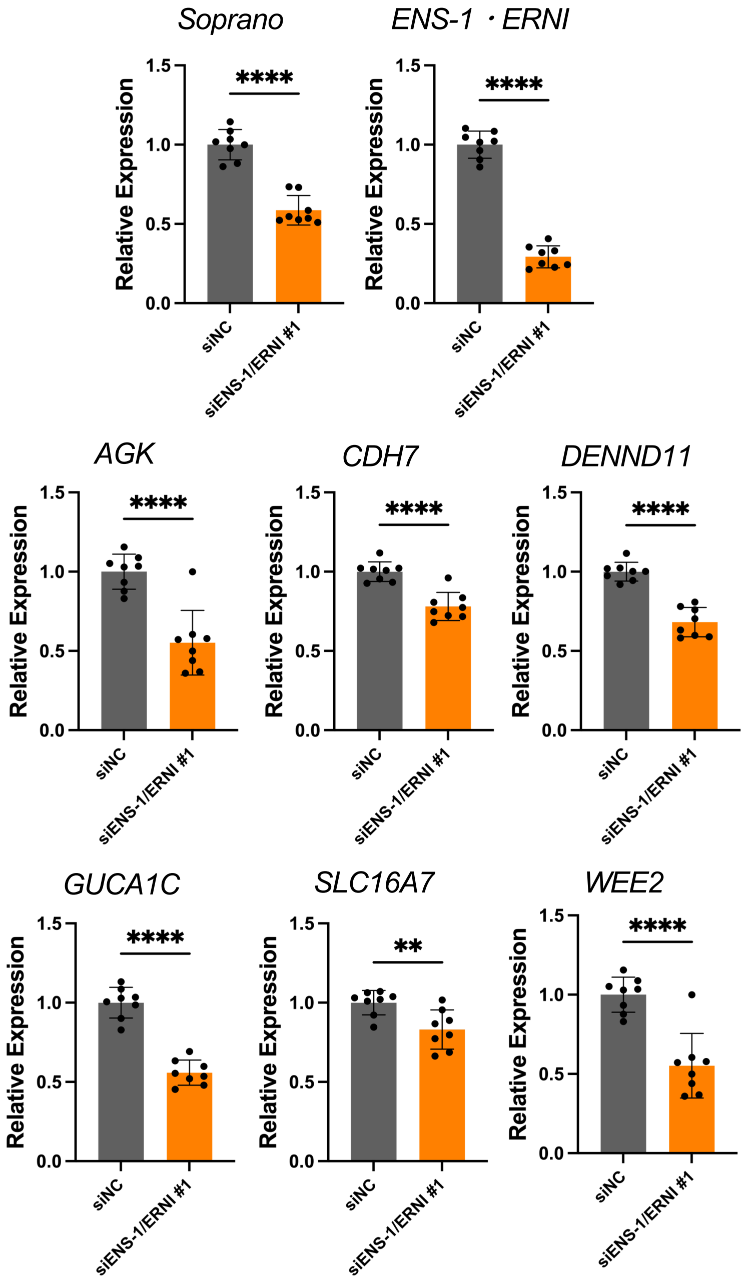
Soprano
ENS-1・ERNI
AGK
CDH7
DENND11
GUCA1C
SLC16A7
WEE2
Relative Expression
1.5
1.0
0.5
0.0
****
**
siNC
siENS-1/ERNI #1

Fig. S5

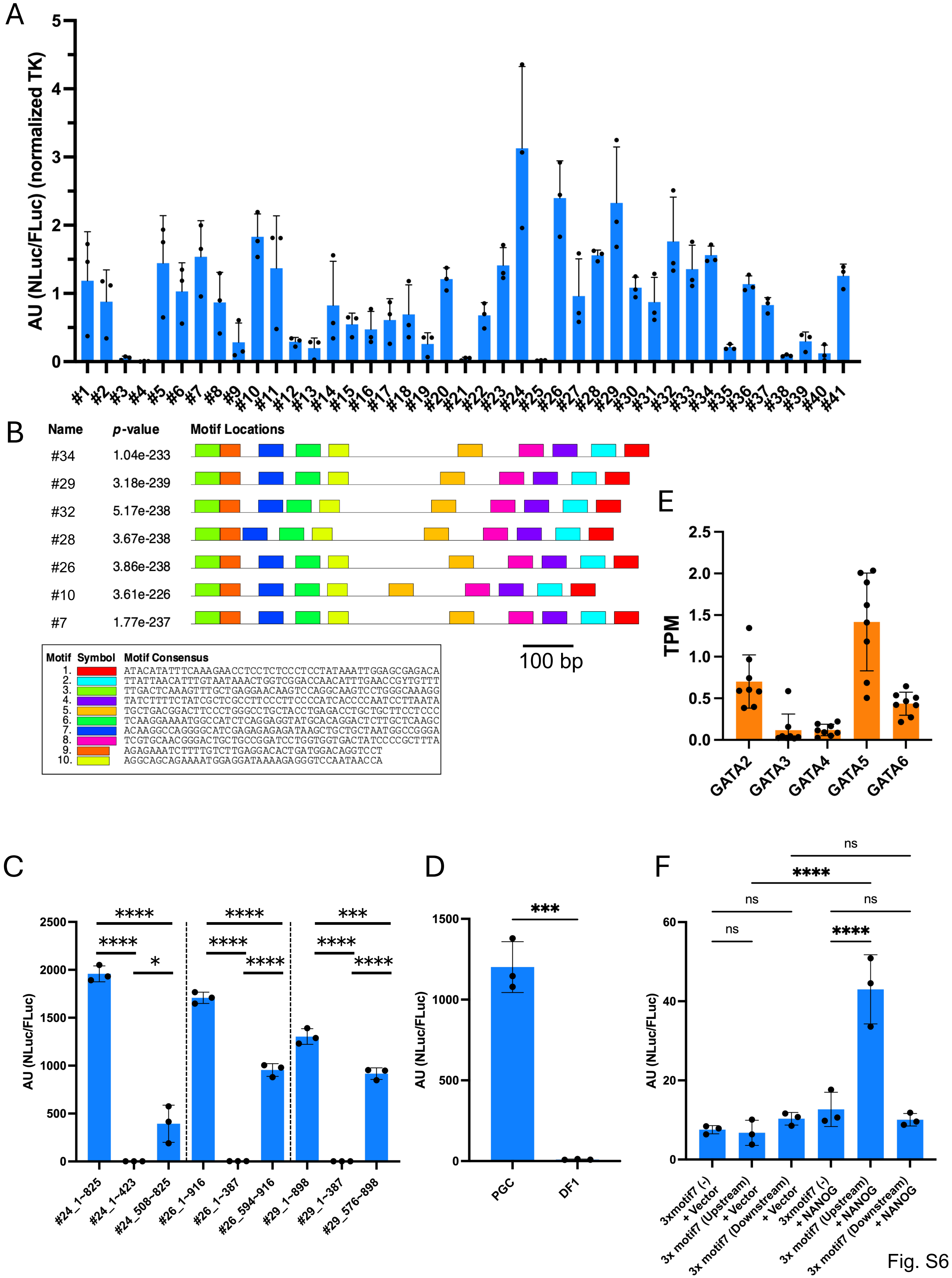
A
AU (NLuc/FLuc) (normalized TK)
#1 #2 #3 #4 #5 #6 #7 #8 #9 #10 #11 #12 #13 #14 #15 #16 #17 #18 #19 #20 #21 #22 #23 #24 #25 #26 #27 #28 #29 #30 #31 #32 #33 #34 #35 #36 #37 #38 #39 #40 #41
B
Name p-value Motif Locations
#34 1.04e-233
#29 3.18e-239
#32 5.17e-238
#28 3.67e-238
#26 3.86e-238
#10 3.61e-226
#7 1.77e-237
100 bp
Motif Symbol Motif Consensus
1. ATACATATTTCAAAGAACCTCCTCTCCCTCCTATAAATTGGAGCGAGACA
2. TTATTAACATTTGTAATAAACTGGTCGGACCAACATTTGAACCGYTGTTT
3. TTGACTCAAAGTTTGCTGAGGAACAAGTCCAGGCAAGTCCTGGGCAAAGG
4. TATCTTTTCTATCGCTCGCCTTCCCTTCCCCATCACCCCAATCCTTAATA
5. TGCTGACGGACTTCCCTGGGCCTGCTACCTGAGACCTGCTGCTTCCTCCC
6. TCAAGGAAAATGGCCATCTCAGGAGGTATGCACAGGACTCTTGCTCAAGC
7. ACAAGGCCAGGGGCATCGAGAGAGAGATAAGCTGCTGCTAATGGCCGGGA
8. TCGTGCAACGGGACTGCTGCCGGATCCTGGTGGTGACTATCCCCGCTTTA
9. AGAGAAATCTTTTGTCTTGAGGACACTGATGGACAGGTCCT
10. AGGCAGCAGAAAATGGAGGATAAAAGAGGGTCCAATAACCA
E
TPM
GATA2 GATA3 GATA4 GATA5 GATA6
C
AU (NLuc/FLuc)
#24_1-825 #24_1-423 #24_508-825 #26_1-916 #26_1-387 #26_594-916 #29_1-898 #29_1-387 #29_576-898
D
AU (NLuc/FLuc)
PGC DF1
F
AU (NLuc/FLuc)
3xmotif7 (-) + Vector
3x motif7 (Upstream) + Vector
3x motif7 (Downstream) + Vector
3xmotif7 (-) + NANOG
3x motif7 (Upstream) + NANOG
3x motif7 (Downstream) + NANOG
ns
****

Fig. S6

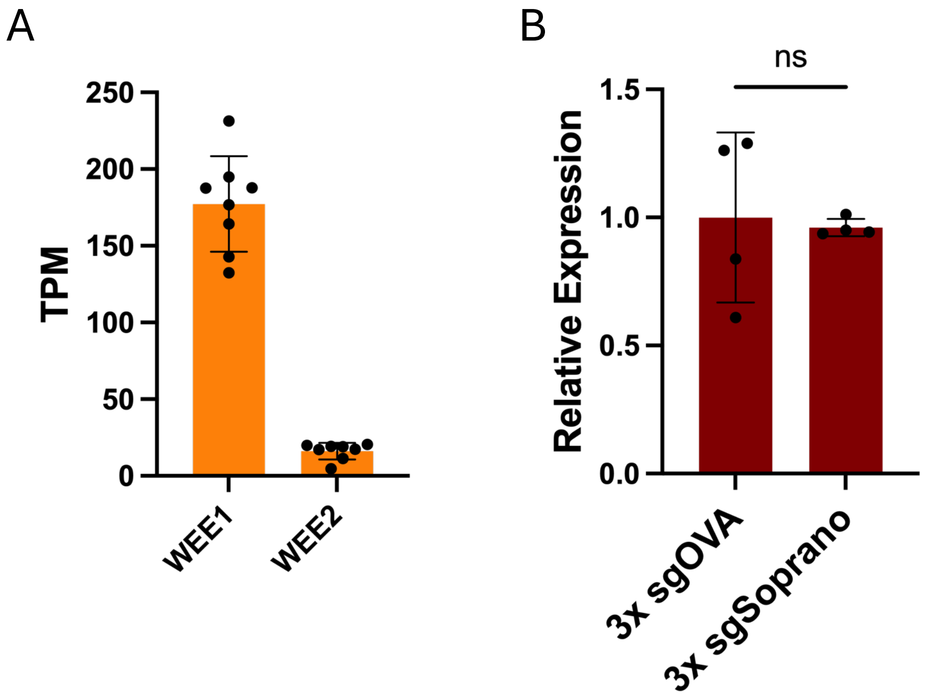
A
B
250
200
150
100
50
0
TPM
WEE1
WEE2
ns
1.5
1.0
0.5
0.0
Relative Expression
3x sgOVA
3x sgSoprano

Fig. S7
