## Supplementary Tables for "The chicken retrovirus-like gene *ENS-1/ERNI* and its LTR *Soprano* are involved in primordial germ cell development"

### Supplementally tableS1. Germ-line transmission of cultured PGCs

Generation of germ line chimeras

GSP

| | No. of transplantation | No. of hatched |
|---|---|---|
| | 45 | 15 |

GSN/1

| | No. of transplantation | No. of hatched |
|---|---|---|
| PGC lot#1 | 26 | 4 |
| PGC lot#2 | 41 | 10 |

Mating results of germ-line chimeras and non-transgenic chickens

GSP

| Hen ID | No. of offspring from transplanted PGCs | No. of analyzed chicks | transmission ratio |
|---|---|---|---|
| #3213 | 0 | 30 | 0.000 |
| #3709 | 3 | 313 | 0.010 |
| #5238 | 26 | 194 | 0.134 |

GSN/1

| Hen ID | PGCs lot | No. of offspring from transplanted PGCs | No. of analyzed embryos | transmission ratio |
|---|---|---|---|---|
| #2520 | #1 | 4 | 10 | 0.400 |
| #2722 | #2 | 2 | 18 | 0.111 |
| #2723 | | 8 | 24 | 0.333 |
| #2725 | | 13 | 32 | 0.406 |

### Supplementally table S2. DEGs comparing male and female cultured PGCs (p-adj < 0.01)

| Emsembl_ID | Symbol | entrez_ID | logFC | p-adj_value | FDR | genetype | Chr |
|---|---|---|---|---|---|---|---|
| ENSGALG00010013512 | LOC430910 | 430910 | -13.27887576 | 1.05E-103 | 5.47E-100 | protein-coding | W |
| ENSGALG00010013319 | LOC430766 | 430766 | -13.19428637 | 4.50E-196 | 7.05E-192 | protein-coding | W |
| ENSGALG00010012927 | LOC769000 | 769000 | -12.76276746 | 1.86E-78 | 7.27E-75 | protein-coding | W |
| ENSGALG00010000262 | HNRNPKL | 426516 | -12.57571255 | 1.14E-148 | 8.94E-145 | protein-coding | JAENSK010000547.1 |
| ENSGALG00010011436 | UBAP2L2 | 407091 | -12.45242663 | 3.66E-68 | 9.56E-65 | protein-coding | W |
| ENSGALG00010012323 | SMAD7B | 395544 | -12.14679124 | 4.19E-57 | 9.38E-54 | protein-coding | W |
| ENSGALG00010012336 | LOC426615 | 426615 | -11.92581939 | 6.43E-42 | 1.01E-38 | protein-coding | W |
| ENSGALG00010012286 | LOC107049046 | 107049046 | -11.83602494 | 8.05E-21 | 6.01E-18 | protein-coding | W |
| ENSGALG00010000326 | NA | NA | -11.50887128 | 9.03E-46 | 1.77E-42 | protein-coding | JAENSK010000532.1 |
| ENSGALG00010012437 | LOC374195 | 374195 | -11.38110469 | 2.10E-28 | 1.83E-25 | protein-coding | W |
| ENSGALG00010012393 | LOC427025 | 427025 | -11.2713594 | 8.08E-77 | 2.53E-73 | protein-coding | W |
| ENSGALG00010011442 | UBE2R2L | 427134 | -11.04341274 | 3.04E-37 | 4.33E-34 | protein-coding | W |
| ENSGALG00010013248 | LOC107049459 | 107049459 | -10.95913064 | 6.71E-32 | 7.50E-29 | protein-coding | W |
| ENSGALG00010011555 | LOC100859602 | 100859602 | -10.73519106 | 7.05E-33 | 8.49E-30 | protein-coding | W |
| ENSGALG00010012888 | LOC431003 | 431003 | -10.72866566 | 7.91E-34 | 1.03E-30 | protein-coding | W |
| ENSGALG00010000309 | LOC121108809 | 121108809 | -10.52896838 | 5.07E-26 | 4.18E-23 | protein-coding | JAENSK010000532.1 |
| ENSGALG00010013420 | LOC107049174 | 107049174 | -10.47736291 | 1.51E-29 | 1.48E-26 | protein-coding | W |
| ENSGALG00010012211 | LOC425347 | 425347 | -10.38323382 | 7.54E-29 | 6.94E-26 | protein-coding | W |
| ENSGALG00010011467 | LOC100859467 | 100859467 | -10.32347641 | 3.29E-30 | 3.44E-27 | protein-coding | W |
| ENSGALG00010011964 | NA | NA | -8.969711969 | 8.29E-19 | 5.90E-16 | protein-coding | W |
| ENSGALG00010000369 | LOC121108844 | 121108844 | -8.837912469 | 3.91E-18 | 2.55E-15 | protein-coding | JAENSK010000532.1 |
| ENSGALG00010012906 | ST8SIA3L | 776262 | -8.785695068 | 2.38E-17 | 1.49E-14 | protein-coding | W |
| ENSGALG00010012781 | SPIN1L | 374014 | -8.459987498 | 1.35E-12 | 6.59E-10 | protein-coding | W |
| ENSGALG00010000276 | NA | NA | -8.387099975 | 4.81E-15 | 2.69E-12 | ncRNA | JAENSK010000533.1 |
| ENSGALG00010013134 | NEDD4L | 374119 | -8.256700113 | 9.38E-16 | 5.44E-13 | protein-coding | W |
| ENSGALG00010000318 | NA | NA | -8.159317853 | 8.29E-11 | 3.61E-08 | protein-coding | JAENSK010000532.1 |
| ENSGALG00010011978 | NA | NA | -8.007931191 | 7.60E-13 | 3.97E-10 | protein-coding | W |
| ENSGALG00010012897 | NA | NA | -7.94813718 | 1.91E-13 | 1.03E-10 | protein-coding | W |
| ENSGALG00010013058 | LOC121108214 | 121108214 | -7.741334169 | 3.67E-18 | 2.50E-15 | protein-coding | W |
| ENSGALG00010011988 | HINTW | 395423 | -7.603994404 | 2.56E-12 | 1.21E-09 | protein-coding | W |
| ENSGALG00010013712 | NA | NA | -7.440245514 | 1.14E-11 | 5.11E-09 | ncRNA | W |
| ENSGALG00010013483 | NA | NA | -6.986614655 | 1.04E-42 | 1.80E-39 | protein-coding | W |
| ENSGALG00010011931 | ATP5F1AW | 431564 | -6.81462674 | 1.60E-08 | 5.11E-06 | protein-coding | W |
| ENSGALG00010013185 | NA | NA | -5.790680994 | 6.60E-12 | 3.04E-09 | protein-coding | W |
| ENSGALG00010013154 | NA | NA | -5.645575538 | 1.12E-12 | 5.67E-10 | protein-coding | W |
| ENSGALG00010013079 | NA | NA | -5.13754388 | 1.62E-09 | 6.04E-07 | protein-coding | W |
| ENSGALG00010012519 | LOC427010 | 427010 | -5.063147774 | 1.75E-25 | 1.37E-22 | protein-coding | W |
| ENSGALG00010011577 | NA | NA | -4.971968512 | 6.12E-10 | 2.39E-07 | protein-coding | W |
| ENSGALG00010011493 | NA | NA | -4.885459879 | 1.04E-08 | 3.41E-06 | protein-coding | W |
| ENSGALG00010028331 | NA | NA | -4.879928344 | 1.77E-06 | 0.00027982 | ncRNA | 12 |
| ENSGALG00010013347 | NA | NA | -4.537910809 | 3.39E-16 | 2.04E-13 | protein-coding | W |
| ENSGALG00010011541 | NA | NA | -4.267639731 | 1.31E-06 | 0.000213995 | ncRNA | 3 |
| ENSGALG00010005791 | NA | NA | -3.854803361 | 3.06E-05 | 0.003110606 | ncRNA | 2 |
| ENSGALG00010011537 | NA | NA | -3.693685476 | 0.000384931 | 0.02777887 | ncRNA | 3 |
| ENSGALG00010000826 | NA | NA | -3.683413465 | 0.000320673 | 0.023913038 | ncRNA | 2 |
| ENSGALG00010012884 | NA | NA | -3.632641782 | 4.41E-05 | 0.004319582 | protein-coding | W |
| ENSGALG00010007107 | NA | NA | -3.61163161 | 2.16E-07 | 4.46E-05 | ncRNA | 5 |
| ENSGALG00010011767 | NA | NA | -3.081183409 | 0.004223525 | 0.183723324 | ncRNA | 3 |
| ENSGALG00010001092 | NA | NA | -3.052943719 | 1.13E-05 | 0.001342028 | ncRNA | 1 |
| ENSGALG00010004360 | NA | NA | -2.537694935 | 4.92E-05 | 0.004755317 | ncRNA | 1 |
| ENSGALG00010007901 | FSTL5 | 428726 | -2.493248551 | 0.002072822 | 0.104710945 | protein-coding | 4 |
| ENSGALG00010010189 | NA | NA | -2.194149123 | 0.000221255 | 0.01776848 | ncRNA | 2 |
| ENSGALG00010028330 | NA | NA | -2.182900191 | 0.002307791 | 0.112585687 | ncRNA | 12 |
| ENSGALG00010004357 | NA | NA | -2.124744493 | 1.69E-07 | 3.67E-05 | ncRNA | 1 |
| ENSGALG00010003227 | PLAC8L1 | 770777 | -2.081624053 | 0.008940278 | 0.334140222 | protein-coding | 4 |
| ENSGALG00010006138 | NA | NA | -1.752353431 | 0.002465583 | 0.118438759 | ncRNA | 1 |
| ENSGALG00010019838 | ALX4 | 373976 | -1.54238928 | 0.001247247 | 0.068532963 | protein-coding | 5 |
| ENSGALG00010002843 | NA | NA | -1.392470044 | 0.000497472 | 0.034021471 | ncRNA | 6 |
| ENSGALG00010016880 | AR | 422165 | -1.331478019 | 0.002344766 | 0.113501303 | protein-coding | 4 |
| ENSGALG00010028386 | NCAN | 395493 | -1.273108751 | 0.004039635 | 0.176213614 | protein-coding | 28 |
| ENSGALG00010001884 | NA | NA | -1.195773275 | 0.001814334 | 0.094445015 | protein-coding | 6 |
| ENSGALG00010016872 | EDA2R | 422166 | -1.181402753 | 0.006925319 | 0.270450121 | protein-coding | 4 |
| ENSGALG00010018627 | MYLK | 396445 | -1.146236567 | 0.00031809 | 0.02383391 | protein-coding | 7 |
| ENSGALG00010026062 | ITGA4 | 424121 | -0.941191551 | 0.000237047 | 0.01865404 | protein-coding | 7 |
| ENSGALG00010021703 | CD82 | 423172 | -0.802445034 | 0.002998658 | 0.138872872 | protein-coding | 5 |
| ENSGALG00010001877 | OAT | 426430 | -0.67389874 | 0.000879955 | 0.052000377 | protein-coding | 6 |
| ENSGALG00010024655 | IGF2BP2 | 424865 | -0.643751946 | 0.007800147 | 0.298656 | protein-coding | 9 |
| ENSGALG00010008029 | LY6E | 395550 | -0.570037306 | 0.008746199 | 0.328220842 | protein-coding | 2 |
| ENSGALG00010027927 | LOC426626 | 426626 | -0.545030257 | 0.004906025 | 0.207644197 | protein-coding | 28 |
| ENSGALG00010009593 | NCBP1 | 427379 | 0.507516428 | 0.006468638 | 0.257757932 | protein-coding | Z |
| ENSGALG00010011029 | HMGCR | 395145 | 0.521703226 | 0.006852626 | 0.268952673 | protein-coding | Z |
| ENSGALG00010009492 | CDC42SE2 | 768825 | 0.541603068 | 0.007628215 | 0.294957656 | protein-coding | Z |
| ENSGALG00010001169 | SLC25A51 | 100859749 | 0.55875236 | 0.005106641 | 0.212686165 | protein-coding | Z |
| ENSGALG00010014907 | ZFR2 | 427424 | 0.568511196 | 0.001816596 | 0.094445015 | protein-coding | Z |
| ENSGALG00010014677 | HNRNPK | 427458 | 0.574940777 | 0.001759592 | 0.09215791 | protein-coding | Z |
| ENSGALG00010010751 | SLC30A5 | 427173 | 0.584766869 | 0.003796442 | 0.167821789 | protein-coding | Z |

| | | | | | | | |
|---|---|---|---|---|---|---|---|
| ENSGALG00010014736 | KIF2A | 427156 | 0.585375134 | 0.002740534 | 0.129267338 | protein-coding | Z |
| ENSGALG00010011544 | STOML2 | 427412 | 0.590939559 | 0.001215987 | 0.06752608 | protein-coding | Z |
| ENSGALG00010015616 | NADK2 | 427438 | 0.598501828 | 0.001642374 | 0.086890484 | protein-coding | Z |
| ENSGALG00010009724 | ISOC1 | 415601 | 0.61398276 | 0.00850432 | 0.32102589 | protein-coding | Z |
| ENSGALG00010016798 | NA | NA | 0.619192744 | 0.002136213 | 0.106878885 | ncRNA | Z |
| ENSGALG00010012640 | PRRC1 | 427127 | 0.62011729 | 0.002305059 | 0.112585687 | protein-coding | Z |
| ENSGALG00010009388 | TMED7 | 769360 | 0.626982808 | 0.002082266 | 0.104849811 | protein-coding | Z |
| ENSGALG00010012551 | PCGF3 | 427287 | 0.637857215 | 0.002458439 | 0.118438759 | protein-coding | Z |
| ENSGALG00010010905 | ENC1 | 427203 | 0.640230312 | 0.003070668 | 0.141016611 | protein-coding | Z |
| ENSGALG00010010457 | AGGF1 | 416364 | 0.642402796 | 0.003976099 | 0.173926553 | protein-coding | Z |
| ENSGALG00010015413 | CARNMT1 | 427256 | 0.642678819 | 0.006384838 | 0.255720116 | protein-coding | Z |
| ENSGALG00010001152 | FEM1C | 431262 | 0.64348383 | 0.00938357 | 0.345756949 | protein-coding | Z |
| ENSGALG00010008149 | DNAJA1 | 427376 | 0.651327528 | 0.000725064 | 0.04502089 | protein-coding | Z |
| ENSGALG00010013438 | RPL37 | 427186 | 0.656389523 | 0.000590492 | 0.038819462 | protein-coding | Z |
| ENSGALG00010009486 | B4GALT1 | 396121 | 0.670566949 | 0.009020853 | 0.336348959 | protein-coding | Z |
| ENSGALG00010013967 | MLLT3 | 427234 | 0.681933439 | 0.005171954 | 0.214835028 | protein-coding | Z |
| ENSGALG00010012969 | SLC12A2 | 427123 | 0.687473486 | 0.009294529 | 0.34328379 | protein-coding | Z |
| ENSGALG00010010445 | LMAN1L | 426849 | 0.695679604 | 0.000499127 | 0.034021471 | protein-coding | Z |
| ENSGALG00010013132 | TTC37 | 427114 | 0.696123768 | 0.002565242 | 0.122102391 | protein-coding | Z |
| ENSGALG00010012011 | HOOK3 | 427297 | 0.69889477 | 0.001878701 | 0.09614529 | protein-coding | Z |
| ENSGALG00010015628 | UBQLN1 | 427454 | 0.70074232 | 0.000655892 | 0.041763286 | protein-coding | Z |
| ENSGALG00010012889 | RIC1 | 427226 | 0.703729386 | 0.001089548 | 0.062044824 | protein-coding | Z |
| ENSGALG00010010186 | TNPO1 | 427218 | 0.708474443 | 0.00115026 | 0.064795202 | protein-coding | Z |
| ENSGALG00010012903 | RGMB | NA | 0.7095072 | 0.005449408 | 0.223983558 | protein-coding | Z |
| ENSGALG00010012439 | PAPD4 | 427178 | 0.709569048 | 0.003006252 | 0.138872872 | protein-coding | Z |
| ENSGALG00010012973 | GLDC | 374222 | 0.713786149 | 0.001455316 | 0.078316987 | protein-coding | Z |
| ENSGALG00010009722 | GBA2 | 100859407 | 0.716189418 | 0.000432288 | 0.030631795 | protein-coding | Z |
| ENSGALG00010001204 | SNX2 | 426797 | 0.724674825 | 0.002503964 | 0.119914614 | protein-coding | Z |
| ENSGALG00010011793 | TLN1 | 395194 | 0.729814049 | 0.000122111 | 0.010564949 | protein-coding | Z |
| ENSGALG00010012121 | SREK1 | 427168 | 0.730738125 | 0.000157548 | 0.013193627 | protein-coding | Z |
| ENSGALG00010014539 | PAIP1 | 427195 | 0.733161246 | 2.98E-05 | 0.003074648 | protein-coding | Z |
| ENSGALG00010004350 | MOXD1 | 395333 | 0.736694955 | 0.009669526 | 0.354168992 | protein-coding | 3 |
| ENSGALG00010013183 | CLTA | 427284 | 0.737029503 | 0.000635381 | 0.04094676 | protein-coding | Z |
| ENSGALG00010013085 | LYSMD3 | 427106 | 0.739077812 | 0.005096688 | 0.212686165 | protein-coding | Z |
| ENSGALG00010014176 | ABHD17B | 427252 | 0.741767119 | 0.008580219 | 0.322995747 | protein-coding | Z |
| ENSGALG00010014579 | NNT | 427196 | 0.745511461 | 0.003056304 | 0.140769755 | protein-coding | Z |
| ENSGALG00010014031 | RPL22L1 | 100858296 | 0.746918441 | 0.006787669 | 0.267745346 | protein-coding | 9 |
| ENSGALG00010009199 | SEMA4D | 396331 | 0.748267488 | 6.60E-05 | 0.00618657 | protein-coding | Z |
| ENSGALG00010011367 | IER3IP1 | 101749130 | 0.748761125 | 0.00078629 | 0.047541685 | protein-coding | Z |
| ENSGALG00010011881 | RIOK2 | 427278 | 0.749579282 | 0.000898783 | 0.052913309 | protein-coding | Z |
| ENSGALG00010010463 | ACAA2 | 426847 | 0.749906182 | 0.00069267 | 0.043215997 | protein-coding | Z |
| ENSGALG00010006851 | TMEM167A | 770790 | 0.750356848 | 0.001850514 | 0.095013295 | protein-coding | Z |
| ENSGALG00010006355 | PTGR1 | 427337 | 0.755288097 | 0.000123508 | 0.01062712 | protein-coding | Z |
| ENSGALG00010014265 | ZNF462 | 431622 | 0.756570369 | 0.000266089 | 0.0206745 | protein-coding | Z |
| ENSGALG00010010809 | CENPH | 395316 | 0.760574019 | 0.000238348 | 0.018662659 | protein-coding | Z |
| ENSGALG00010010122 | THBS4 | 396306 | 0.762178459 | 0.007773004 | 0.298656 | protein-coding | Z |
| ENSGALG00010015852 | ZFAND5 | 427254 | 0.762178725 | 9.34E-05 | 0.008266026 | protein-coding | Z |
| ENSGALG00010001237 | MELK | 428391 | 0.762552374 | 0.000154875 | 0.013039494 | protein-coding | Z |
| ENSGALG00010012322 | MYO3AL | 427419 | 0.763461286 | 0.000735375 | 0.045179724 | protein-coding | Z |
| ENSGALG00010012711 | ELL2 | 427119 | 0.768594612 | 0.001821903 | 0.094445015 | protein-coding | Z |
| ENSGALG00010010368 | ZNF532 | 100857356 | 0.770513692 | 0.000361934 | 0.026485433 | protein-coding | Z |
| ENSGALG00010012830 | ARRDC3 | 427107 | 0.774486094 | 0.00438367 | 0.189603727 | protein-coding | Z |
| ENSGALG00010010671 | SIGMAR1 | 100859748 | 0.774674387 | 0.000313157 | 0.023577104 | protein-coding | Z |
| ENSGALG00010014035 | PSIP1 | 431605 | 0.779803088 | 6.41E-06 | 0.000833926 | protein-coding | Z |
| ENSGALG00010000453 | ACHE | NA | 0.782723132 | 0.001827384 | 0.094445015 | protein-coding | JAENSK010000584.1 |
| ENSGALG00010012638 | NA | NA | 0.790249143 | 0.005039074 | 0.212108793 | ncRNA | Z |
| ENSGALG00010014759 | NA | NA | 0.792643162 | 0.003368672 | 0.151590227 | ncRNA | Z |
| ENSGALG00010014170 | CEMIP2 | 427250 | 0.792689741 | 2.89E-05 | 0.002992494 | protein-coding | Z |
| ENSGALG00010008231 | ZCCHC6 | 100857368 | 0.793455128 | 0.000735685 | 0.045179724 | protein-coding | Z |
| ENSGALG00010008259 | MCC | 415603 | 0.794652772 | 0.002983807 | 0.138872872 | protein-coding | Z |
| ENSGALG00010010062 | RGS7BP | 770874 | 0.795179357 | 0.006828951 | 0.268696901 | protein-coding | Z |
| ENSGALG00010015493 | OSTF1 | 427258 | 0.797525222 | 0.004525503 | 0.193632168 | protein-coding | Z |
| ENSGALG00010010322 | NEDD4L | 374119 | 0.798862869 | 0.000650029 | 0.041719051 | protein-coding | Z |
| ENSGALG00010008292 | REEP5 | 770021 | 0.802145596 | 0.003579702 | 0.160625016 | protein-coding | Z |
| ENSGALG00010012119 | EPG5 | 416379 | 0.802624304 | 0.000106548 | 0.009373822 | protein-coding | Z |
| ENSGALG00010015894 | CEP78 | 431612 | 0.804582533 | 0.000951998 | 0.05542113 | protein-coding | Z |
| ENSGALG00010012326 | SGTB | 427166 | 0.806499668 | 0.009130483 | 0.338022139 | protein-coding | Z |
| ENSGALG00010011406 | UBE2R2 | 427021 | 0.80997969 | 1.42E-05 | 0.001630075 | protein-coding | Z |
| ENSGALG00010009451 | ROR2 | 769516 | 0.810063142 | 0.004508394 | 0.193632168 | protein-coding | Z |
| ENSGALG00010012082 | HAUS1 | 416380 | 0.810338705 | 0.002563965 | 0.122102391 | protein-coding | Z |
| ENSGALG00010009120 | ATG12 | 427390 | 0.819166227 | 0.001187635 | 0.066343745 | protein-coding | Z |
| ENSGALG00010009595 | LYRM7 | 768932 | 0.821002763 | 0.004957208 | 0.209244961 | protein-coding | Z |
| ENSGALG00010013256 | BRIX1 | 427433 | 0.825267545 | 5.75E-06 | 0.000762654 | protein-coding | Z |
| ENSGALG00010001400 | SNX24 | 426798 | 0.825892473 | 0.005858815 | 0.238929789 | protein-coding | Z |
| ENSGALG00010015148 | GPBP1 | 427148 | 0.826979431 | 0.003837799 | 0.16882005 | protein-coding | Z |
| ENSGALG00010010252 | WDR7 | 426855 | 0.831467549 | 0.007768709 | 0.298656 | protein-coding | Z |
| ENSGALG00010008424 | PPIP5K2 | 427276 | 0.832643134 | 0.000129469 | 0.011018968 | protein-coding | Z |

| | | | | | | | |
|---|---|---|---|---|---|---|---|
| ENSGALG00010014737 | RMI1 | 431669 | 0.832753149 | 7.59E-05 | 0.007044799 | protein-coding | Z |
| ENSGALG00010005887 | CCNH | 427328 | 0.83316661 | 0.000684202 | 0.04303052 | protein-coding | Z |
| ENSGALG00010014758 | FXN | 427244 | 0.835050991 | 0.000592455 | 0.038819462 | protein-coding | Z |
| ENSGALG00010012861 | WDR70 | 427444 | 0.838450183 | 0.0002813 | 0.021635693 | protein-coding | Z |
| ENSGALG00010009360 | TNFAIP8 | 427384 | 0.838561981 | 0.00093925 | 0.05488304 | protein-coding | Z |
| ENSGALG00010005871 | EGLN3 | 423316 | 0.841186164 | 0.004395029 | 0.189603727 | protein-coding | 5 |
| ENSGALG00010014754 | DIMT1 | 427157 | 0.842398241 | 0.000488727 | 0.033910118 | protein-coding | Z |
| ENSGALG00010009574 | TAF1C | 101750393 | 0.843898171 | 0.000630694 | 0.040812691 | protein-coding | Z |
| ENSGALG00010009603 | HINT2 | 395424 | 0.845106639 | 1.66E-05 | 0.001875755 | protein-coding | Z |
| ENSGALG00010011456 | KIAA1328 | 426890 | 0.847648069 | 0.006888231 | 0.269674254 | protein-coding | Z |
| ENSGALG00010008333 | APC | 415607 | 0.850167317 | 0.000112967 | 0.009883061 | protein-coding | Z |
| ENSGALG00010008928 | CFC1 | 395437 | 0.851464062 | 0.000478243 | 0.033434286 | protein-coding | Z |
| ENSGALG00010013033 | MAP3K1 | 427144 | 0.855106761 | 0.00234004 | 0.113501303 | protein-coding | Z |
| ENSGALG00010007066 | INIP | 427331 | 0.856082147 | 0.001401336 | 0.076197655 | protein-coding | Z |
| ENSGALG00010011464 | FANCG | 378893 | 0.857215235 | 0.000121661 | 0.010564949 | protein-coding | Z |
| ENSGALG00010012795 | TBCA | 416367 | 0.858298118 | 6.44E-06 | 0.000833926 | protein-coding | Z |
| ENSGALG00010012833 | JAK2 | 374199 | 0.85985876 | 0.001407404 | 0.076262789 | protein-coding | Z |
| ENSGALG00010001195 | ZNF608 | 426810 | 0.865163128 | 0.004708118 | 0.200350904 | protein-coding | Z |
| ENSGALG00010012904 | ERMP1 | 426644 | 0.86582105 | 3.01E-05 | 0.003083649 | protein-coding | Z |
| ENSGALG00010011759 | CREB3L4 | 427417 | 0.868288997 | 7.60E-05 | 0.007044799 | protein-coding | Z |
| ENSGALG00010010288 | TXNL1 | 426854 | 0.868710169 | 7.03E-06 | 0.000888393 | protein-coding | Z |
| ENSGALG00010006575 | SSBP2 | 770958 | 0.870872803 | 0.000626761 | 0.040726489 | protein-coding | Z |
| ENSGALG00010010447 | BTF3 | 426154 | 0.871061071 | 3.97E-06 | 0.000555043 | protein-coding | Z |
| ENSGALG00010014397 | PRKAA1 | 427185 | 0.871097011 | 0.000396808 | 0.028504669 | protein-coding | Z |
| ENSGALG00010008141 | APTX | 395173 | 0.87281834 | 0.000564482 | 0.037456719 | protein-coding | Z |
| ENSGALG00010005495 | TXN | 396437 | 0.873682411 | 3.23E-06 | 0.000455441 | protein-coding | Z |
| ENSGALG00010011846 | UBAP1 | 431652 | 0.874058426 | 0.008760939 | 0.328220842 | protein-coding | Z |
| ENSGALG00010012751 | AP3B1 | 427646 | 0.876830361 | 1.54E-06 | 0.00024543 | protein-coding | Z |
| ENSGALG00010010382 | SEC11C | 426850 | 0.880315035 | 0.005067233 | 0.212173433 | protein-coding | Z |
| ENSGALG00010005899 | RASA1 | 427327 | 0.880704073 | 4.60E-06 | 0.000637614 | protein-coding | Z |
| ENSGALG00010001182 | ZCCHC7 | 426807 | 0.884022065 | 4.47E-05 | 0.004345364 | protein-coding | Z |
| ENSGALG00010001171 | POLR1E | 426805 | 0.884394491 | 5.71E-06 | 0.000762654 | protein-coding | Z |
| ENSGALG00010009640 | NPR2 | 100859339 | 0.885670268 | 0.002225597 | 0.109673918 | protein-coding | Z |
| ENSGALG00010014359 | LOC100857745 | 100857745 | 0.887506339 | 0.007359776 | 0.285282403 | protein-coding | Z |
| ENSGALG00010006223 | ATG10 | 427322 | 0.88806663 | 0.001034715 | 0.059353993 | protein-coding | Z |
| ENSGALG00010010302 | WDR36 | 415609 | 0.888916978 | 2.58E-06 | 0.000384886 | protein-coding | Z |
| ENSGALG00010009116 | S1PR3 | 431264 | 0.889777706 | 0.000796695 | 0.047985565 | protein-coding | Z |
| ENSGALG00010013306 | ARSB | 771459 | 0.893440107 | 0.001903533 | 0.097098768 | protein-coding | Z |
| ENSGALG00010011649 | MRPS27 | 427216 | 0.895802824 | 1.83E-05 | 0.002022076 | protein-coding | Z |
| ENSGALG00010009431 | SMN | 374025 | 0.896601456 | 2.74E-06 | 0.0004043 | protein-coding | Z |
| ENSGALG00010008699 | NAA35 | 427461 | 0.90006585 | 8.00E-06 | 0.00099384 | protein-coding | Z |
| ENSGALG00010012796 | MOCS2 | 427199 | 0.901052632 | 0.000663601 | 0.04207281 | protein-coding | Z |
| ENSGALG00010010393 | FANCC | 427468 | 0.90130674 | 4.88E-06 | 0.000663869 | protein-coding | Z |
| ENSGALG00010013106 | SETD9 | 770222 | 0.901944846 | 0.002095694 | 0.105187735 | protein-coding | Z |
| ENSGALG00010015674 | RFKL | 431449 | 0.902288111 | 3.50E-05 | 0.003489246 | protein-coding | Z |
| ENSGALG00010012060 | KCMF1 | 770239 | 0.903978221 | 5.41E-07 | 0.00010335 | protein-coding | Z |
| ENSGALG00010013002 | NA | NA | 0.904083424 | 0.008507391 | 0.32102589 | ncRNA | Z |
| ENSGALG00010016168 | PLIN2 | 427237 | 0.904532077 | 7.70E-05 | 0.007095312 | protein-coding | Z |
| ENSGALG00010016887 | TJP2 | 395751 | 0.906010818 | 6.60E-06 | 0.000847416 | protein-coding | Z |
| ENSGALG00010012247 | TRIM23 | 427164 | 0.906353211 | 0.000225606 | 0.018025423 | protein-coding | Z |
| ENSGALG00010014167 | SLC38A9 | 427139 | 0.911045718 | 0.001232853 | 0.067980551 | protein-coding | Z |
| ENSGALG00010010834 | MRPS36 | 771273 | 0.913222537 | 0.001099856 | 0.062404886 | protein-coding | Z |
| ENSGALG00010010990 | MED18 | 100857707 | 0.913363166 | 0.000739861 | 0.045258676 | protein-coding | Z |
| ENSGALG00010005637 | TOPORS | 427342 | 0.915371933 | 4.02E-05 | 0.00398303 | protein-coding | Z |
| ENSGALG00010009167 | DCTN3 | 427404 | 0.915633649 | 2.32E-05 | 0.002525228 | protein-coding | Z |
| ENSGALG00010008177 | NREP | 396353 | 0.920682734 | 0.000334795 | 0.024730602 | protein-coding | Z |
| ENSGALG00010009663 | FER | 415614 | 0.921727455 | 0.005282457 | 0.21826722 | protein-coding | Z |
| ENSGALG00010009153 | SECISBP2 | 426815 | 0.922013932 | 2.80E-06 | 0.000410294 | protein-coding | Z |
| ENSGALG00010013238 | RAD1 | 427432 | 0.924118156 | 0.001109892 | 0.062746948 | protein-coding | Z |
| ENSGALG00010011203 | C18orf32 | 426844 | 0.926245377 | 0.000656052 | 0.041763286 | protein-coding | Z |
| ENSGALG00010012334 | ARHGEF39 | 431661 | 0.928319941 | 8.25E-05 | 0.007509197 | protein-coding | Z |
| ENSGALG00010008497 | PAM | 427274 | 0.930253224 | 8.86E-06 | 0.001092979 | protein-coding | Z |
| ENSGALG00010011429 | UBAP2 | 407092 | 0.930415485 | 4.00E-07 | 7.82E-05 | protein-coding | Z |
| ENSGALG00010015239 | NA | NA | 0.931378582 | 0.003330702 | 0.150662531 | ncRNA | Z |
| ENSGALG00010001191 | TOMM5 | 426803 | 0.932182601 | 6.32E-07 | 0.000116407 | protein-coding | Z |
| ENSGALG00010016114 | HAUS6 | 427238 | 0.932200134 | 5.88E-07 | 0.00010961 | protein-coding | Z |
| ENSGALG00010009457 | BDP1 | 427394 | 0.933856163 | 3.42E-05 | 0.003452825 | protein-coding | Z |
| ENSGALG00010012067 | CZH18ORF25 | 416381 | 0.934680267 | 0.001441241 | 0.077827001 | protein-coding | Z |
| ENSGALG00010008136 | IKBKAP | 427375 | 0.937646227 | 8.75E-07 | 0.000150557 | protein-coding | Z |
| ENSGALG00010014427 | TMEM267 | 427193 | 0.938872879 | 0.00051308 | 0.034632868 | protein-coding | Z |
| ENSGALG00010014189 | DDX4 | 395447 | 0.942263626 | 1.44E-07 | 3.22E-05 | protein-coding | Z |
| ENSGALG00010010733 | GFM2 | 427205 | 0.947261953 | 0.002348303 | 0.113501303 | protein-coding | Z |
| ENSGALG00010013246 | LNPEP | 427279 | 0.947767371 | 0.000438741 | 0.030810272 | protein-coding | Z |
| ENSGALG00010009152 | RPP25L | 427403 | 0.949438265 | 1.34E-06 | 0.00021581 | protein-coding | Z |
| ENSGALG00010006252 | COX7C | 431629 | 0.949736469 | 8.02E-07 | 0.000142677 | protein-coding | Z |
| ENSGALG00010013065 | FKTN | 427303 | 0.949919218 | 2.15E-05 | 0.002350983 | protein-coding | Z |
| ENSGALG00010011747 | UTP15 | 426152 | 0.951905343 | 2.40E-06 | 0.000365265 | protein-coding | Z |

| | | | | | | | |
|---|---|---|---|---|---|---|---|
| ENSGALG00010012803 | NUP155 | 427443 | 0.952791237 | 1.18E-06 | 0.00019475 | protein-coding | Z |
| ENSGALG00010011911 | NUDT2 | 427399 | 0.953809331 | 0.009071532 | 0.337435142 | protein-coding | Z |
| ENSGALG00010009446 | LOC107052389 | 107052389 | 0.954139751 | 3.05E-06 | 0.000433932 | protein-coding | Z |
| ENSGALG00010005477 | TMEM161B | 426919 | 0.95420126 | 0.000382151 | 0.027705929 | protein-coding | Z |
| ENSGALG00010008879 | HABP4 | 395444 | 0.95492788 | 0.000301381 | 0.022910831 | protein-coding | Z |
| ENSGALG00010012536 | C5orf51 | 427191 | 0.955437802 | 0.001021593 | 0.058941666 | protein-coding | Z |
| ENSGALG00010012309 | TRAPPC13 | 427165 | 0.958081192 | 0.001561823 | 0.083760797 | protein-coding | Z |
| ENSGALG00010013953 | DHX29 | 427136 | 0.958584378 | 0.00612853 | 0.247352519 | protein-coding | Z |
| ENSGALG00010012578 | NDUFS4 | 374122 | 0.965091455 | 1.82E-06 | 0.000285121 | protein-coding | Z |
| ENSGALG00010011335 | VCP | 427410 | 0.965633432 | 1.22E-07 | 2.85E-05 | protein-coding | Z |
| ENSGALG00010007097 | ZCCHC9 | 427320 | 0.966669852 | 0.000228831 | 0.018190309 | protein-coding | Z |
| ENSGALG00010008311 | SRP19 | 770061 | 0.968907767 | 0.00015237 | 0.012897922 | protein-coding | Z |
| ENSGALG00010009408 | GTF2H2 | 100858972 | 0.97104328 | 9.04E-06 | 0.00110554 | protein-coding | Z |
| ENSGALG00010011133 | CDC37L1 | 426535 | 0.973850061 | 0.000160668 | 0.013383306 | protein-coding | Z |
| ENSGALG00010013225 | GNE | 427285 | 0.975687552 | 2.94E-07 | 5.98E-05 | protein-coding | Z |
| ENSGALG00010006029 | HSDL2 | 100858057 | 0.976425886 | 5.86E-08 | 1.53E-05 | protein-coding | Z |
| ENSGALG00010015038 | PCSK5 | 395456 | 0.977064864 | 0.001023763 | 0.058941666 | protein-coding | Z |
| ENSGALG00010005549 | RNF20 | 427310 | 0.977233908 | 1.43E-07 | 3.22E-05 | protein-coding | Z |
| ENSGALG00010012965 | UHRF2 | 431601 | 0.98254367 | 1.53E-05 | 0.001733105 | protein-coding | Z |
| ENSGALG00010012141 | SREK1IP1 | 427158 | 0.982920958 | 0.000216949 | 0.017512458 | protein-coding | Z |
| ENSGALG00010012532 | FAM174A | 427273 | 0.983148177 | 9.29E-06 | 0.001118756 | protein-coding | Z |
| ENSGALG00010014971 | SMC5 | 431608 | 0.983177774 | 3.34E-08 | 9.69E-06 | protein-coding | Z |
| ENSGALG00010012110 | FCHO2 | NA | 0.983648251 | 0.001044853 | 0.05971678 | protein-coding | Z |
| ENSGALG00010009811 | PJA2 | 415612 | 0.984741986 | 0.000161675 | 0.013395909 | protein-coding | Z |
| ENSGALG00010010257 | NA | NA | 0.985098938 | 0.002937754 | 0.137740187 | protein-coding | Z |
| ENSGALG00010001141 | MTAP | 431261 | 0.985542607 | 0.001326228 | 0.0724805 | protein-coding | Z |
| ENSGALG00010011458 | TPGS2 | 426891 | 0.986469229 | 0.000501849 | 0.034021471 | protein-coding | Z |
| ENSGALG00010012154 | CWC27 | 427159 | 0.986550615 | 2.93E-06 | 0.000423318 | protein-coding | Z |
| ENSGALG00010009594 | HINT1 | 100859112 | 0.987549383 | 0.005563081 | 0.228057202 | protein-coding | Z |
| ENSGALG00010007001 | SNX30 | 427330 | 0.98779487 | 1.14E-05 | 0.001347385 | protein-coding | Z |
| ENSGALG00010007322 | AK6 | 425215 | 0.989901079 | 0.000401571 | 0.028715092 | protein-coding | Z |
| ENSGALG00010010003 | NA | NA | 0.989985429 | 0.005637205 | 0.23049249 | ncRNA | Z |
| ENSGALG00010010621 | LOC107052340 | 107052340 | 0.99150235 | 0.006268059 | 0.252333706 | ncRNA | Z |
| ENSGALG00010009143 | MCCC2 | 427395 | 0.993562051 | 0.000377646 | 0.02750665 | protein-coding | Z |
| ENSGALG00010010937 | CDK7 | 771294 | 0.994725961 | 0.003975974 | 0.173926553 | protein-coding | Z |
| ENSGALG00010013746 | LOC427439 | 427439 | 0.995656976 | 1.80E-07 | 3.86E-05 | protein-coding | Z |
| ENSGALG00010010713 | NSA2 | 427206 | 0.997327725 | 7.87E-08 | 1.96E-05 | protein-coding | Z |
| ENSGALG00010011572 | PTCD2 | 427217 | 1.000949736 | 0.000576168 | 0.038070857 | protein-coding | Z |
| ENSGALG00010006465 | PLAA | 427365 | 1.00259 | 2.17E-07 | 4.46E-05 | protein-coding | Z |
| ENSGALG00010014083 | NIPSNAP3A | 427300 | 1.00290794 | 0.000266683 | 0.0206745 | protein-coding | Z |
| ENSGALG00010010681 | NA | NA | 1.004450536 | 0.007933173 | 0.302271259 | ncRNA | Z |
| ENSGALG00010015212 | GNAQ | 427262 | 1.006456409 | 0.000500833 | 0.034021471 | protein-coding | Z |
| ENSGALG00010014220 | C9orf85 | 770483 | 1.006512009 | 6.88E-06 | 0.000875647 | protein-coding | Z |
| ENSGALG00010009139 | CKS2 | 768830 | 1.007615154 | 0.000405355 | 0.028853897 | protein-coding | Z |
| ENSGALG00010013313 | GAK | 427291 | 1.009146418 | 0.00365671 | 0.163145503 | protein-coding | Z |
| ENSGALG00010015245 | NA | NA | 1.009158363 | 0.000438147 | 0.030810272 | ncRNA | Z |
| ENSGALG00010015054 | CSPG4B | 425372 | 1.009351844 | 0.005271241 | 0.21826722 | protein-coding | Z |
| ENSGALG00010007703 | LOC107052414 | 107052414 | 1.010081937 | 0.001176967 | 0.066062005 | ncRNA | Z |
| ENSGALG00010012745 | 3-Mar | 768965 | 1.010560949 | 0.001570861 | 0.083957989 | protein-coding | Z |
| ENSGALG00010012353 | PIGG | 427290 | 1.011650338 | 2.34E-05 | 0.002532572 | protein-coding | Z |
| ENSGALG00010009704 | RGP1 | 431658 | 1.012396747 | 0.000236405 | 0.01865404 | protein-coding | Z |
| ENSGALG00010010254 | WDR41 | 416366 | 1.01255227 | 5.05E-07 | 9.77E-05 | protein-coding | Z |
| ENSGALG00010014184 | SLC44A1 | 427301 | 1.014070472 | 2.47E-05 | 0.002645652 | protein-coding | Z |
| ENSGALG00010006345 | SMC2 | 396156 | 1.014386747 | 8.37E-08 | 2.05E-05 | protein-coding | Z |
| ENSGALG00010009367 | DMXL1 | 427385 | 1.01627486 | 0.004236373 | 0.18377174 | protein-coding | Z |
| ENSGALG00010001151 | CCDC112 | 770644 | 1.017353268 | 0.003801153 | 0.167821789 | protein-coding | Z |
| ENSGALG00010014041 | SKIV2L2 | 427137 | 1.020246775 | 3.56E-08 | 1.01E-05 | protein-coding | Z |
| ENSGALG00010008127 | TBC1D2 | 427374 | 1.024393167 | 0.002216401 | 0.109673918 | protein-coding | Z |
| ENSGALG00010001252 | SNCAIP | 426796 | 1.024818957 | 0.000340795 | 0.0250556 | protein-coding | Z |
| ENSGALG00010010810 | GALT | 431509 | 1.026149006 | 0.000689824 | 0.043210598 | protein-coding | Z |
| ENSGALG00010011222 | RCL1 | 427222 | 1.027525638 | 3.16E-08 | 9.33E-06 | protein-coding | Z |
| ENSGALG00010009236 | DTWD2 | 427386 | 1.031136708 | 8.79E-05 | 0.007958761 | protein-coding | Z |
| ENSGALG00010014592 | C9orf64 | 427457 | 1.03273963 | 9.74E-07 | 0.000163971 | protein-coding | Z |
| ENSGALG00010014476 | C5orf34 | 427194 | 1.036456309 | 0.005052144 | 0.212108793 | protein-coding | Z |
| ENSGALG00010001156 | ALDH7A1 | 426812 | 1.037251482 | 9.07E-07 | 0.000154353 | protein-coding | Z |
| ENSGALG00010014371 | TTC33 | 427184 | 1.038032343 | 4.18E-05 | 0.004119844 | protein-coding | Z |
| ENSGALG00010015181 | VPS13A | 427050 | 1.038699195 | 0.000127489 | 0.010909743 | protein-coding | Z |
| ENSGALG00010012470 | CHD1 | 395783 | 1.04311616 | 8.59E-08 | 2.07E-05 | protein-coding | Z |
| ENSGALG00010006480 | IFT74 | 427367 | 1.045207571 | 3.10E-07 | 6.22E-05 | protein-coding | Z |
| ENSGALG00010012688 | NDUFAF2 | 769549 | 1.045784891 | 0.003259697 | 0.149259791 | protein-coding | Z |
| ENSGALG00010007224 | DHFR | 427317 | 1.050670758 | 2.57E-06 | 0.000384886 | protein-coding | Z |
| ENSGALG00010011384 | PUM3 | 427350 | 1.050923028 | 2.51E-08 | 7.56E-06 | protein-coding | Z |
| ENSGALG00010012574 | PHAX | 427124 | 1.051241463 | 1.66E-07 | 3.65E-05 | protein-coding | Z |
| ENSGALG00010012952 | KIAA2026 | 427227 | 1.051557411 | 6.31E-06 | 0.000830738 | protein-coding | Z |
| ENSGALG00010010769 | HEXB | 427204 | 1.051664994 | 2.50E-05 | 0.002659416 | protein-coding | Z |
| ENSGALG00010013116 | TARS | 427427 | 1.052350885 | 2.34E-06 | 0.000358636 | protein-coding | Z |
| ENSGALG00010001286 | SRFBP1 | 426795 | 1.052563442 | 1.82E-05 | 0.002018998 | protein-coding | Z |

| | | | | | | | |
|---|---|---|---|---|---|---|---|
| ENSGALG00010014537 | GKAP1 | 427455 | 1.053537664 | 0.002280743 | 0.11196375 | protein-coding | Z |
| ENSGALG00010009280 | HSD17B4 | 395785 | 1.053683701 | 8.88E-05 | 0.007994898 | protein-coding | Z |
| ENSGALG00010013117 | ARSK | 427116 | 1.054819043 | 0.006415951 | 0.256310686 | protein-coding | Z |
| ENSGALG00010012088 | ATP5F1AZ | 374159 | 1.055929112 | 2.15E-08 | 6.59E-06 | protein-coding | Z |
| ENSGALG00010009103 | AP3S1 | 427389 | 1.059529424 | 5.50E-08 | 1.49E-05 | protein-coding | Z |
| ENSGALG00010014877 | GOLPH3 | 427422 | 1.06274707 | 7.79E-08 | 1.96E-05 | protein-coding | Z |
| ENSGALG00010007259 | ANKRD34B | 431626 | 1.062853387 | 5.02E-06 | 0.000678275 | protein-coding | Z |
| ENSGALG00010008234 | COMMD10 | 427388 | 1.069639805 | 4.75E-06 | 0.00065197 | protein-coding | Z |
| ENSGALG00010011774 | ANKRA2 | 426153 | 1.070393454 | 1.90E-07 | 4.01E-05 | protein-coding | Z |
| ENSGALG00010012913 | EGFLAM | 427445 | 1.071893143 | 0.002218506 | 0.109673918 | protein-coding | Z |
| ENSGALG00010013431 | TMEM175 | 427293 | 1.072183276 | 2.95E-06 | 0.000423318 | protein-coding | Z |
| ENSGALG00010012669 | ERCC8 | 427153 | 1.072700706 | 0.000166132 | 0.013692752 | protein-coding | Z |
| ENSGALG00010014350 | PTGER4 | 427183 | 1.073274789 | 0.006368071 | 0.255702541 | protein-coding | Z |
| ENSGALG00010013327 | SLF1 | 427111 | 1.075321307 | 1.37E-05 | 0.00159119 | protein-coding | Z |
| ENSGALG00010012893 | C5orf63 | 427125 | 1.077833004 | 0.008059927 | 0.306355482 | protein-coding | Z |
| ENSGALG00010013446 | ATP5I | 769146 | 1.078871302 | 3.77E-08 | 1.05E-05 | protein-coding | Z |
| ENSGALG00010011972 | IPO11 | 769679 | 1.080638868 | 1.17E-07 | 2.77E-05 | protein-coding | Z |
| ENSGALG00010008243 | YTHDC2 | 415602 | 1.081495804 | 9.34E-09 | 3.11E-06 | protein-coding | Z |
| ENSGALG00010010064 | ERCC6L2 | 427469 | 1.082413924 | 0.000727349 | 0.04502089 | protein-coding | Z |
| ENSGALG00010001221 | GRAMD3 | 426813 | 1.08450163 | 5.47E-05 | 0.005158308 | protein-coding | Z |
| ENSGALG00010013487 | MIER3 | 427146 | 1.084680738 | 0.000862497 | 0.051356285 | protein-coding | Z |
| ENSGALG00010016139 | LURAP1L | 768654 | 1.087302922 | 0.000749563 | 0.045673738 | protein-coding | Z |
| ENSGALG00010010133 | MIR2954 | 100498678 | 1.087566169 | 0.001665514 | 0.087818023 | ncRNA | Z |
| ENSGALG00010010985 | COL4A3BP | 427209 | 1.088809292 | 5.86E-08 | 1.53E-05 | protein-coding | Z |
| ENSGALG00010010870 | NOL6 | 100858366 | 1.091166678 | 2.30E-09 | 8.36E-07 | protein-coding | Z |
| ENSGALG00010010496 | MRPL17 | 769700 | 1.091607357 | 1.95E-08 | 6.12E-06 | protein-coding | Z |
| ENSGALG00010012458 | MRPS30 | 395781 | 1.092356899 | 7.38E-08 | 1.89E-05 | protein-coding | Z |
| ENSGALG00010012845 | HDHD2 | 416385 | 1.09369726 | 0.003289625 | 0.149373556 | protein-coding | Z |
| ENSGALG00010014153 | FOCAD | 427233 | 1.094108934 | 8.11E-07 | 0.000142736 | protein-coding | Z |
| ENSGALG00010012087 | TMEM171 | 427220 | 1.096187485 | 0.000872111 | 0.05173205 | protein-coding | Z |
| ENSGALG00010012196 | CENPK | 427162 | 1.097550848 | 1.09E-05 | 0.001304929 | protein-coding | Z |
| ENSGALG00010013203 | AMACR | 427429 | 1.099256497 | 2.74E-05 | 0.002865238 | protein-coding | Z |
| ENSGALG00010009548 | PIK3C3 | 100857877 | 1.100151568 | 0.000531091 | 0.035542219 | protein-coding | Z |
| ENSGALG00010016251 | MPDZ | 427360 | 1.107234109 | 3.16E-07 | 6.26E-05 | protein-coding | Z |
| ENSGALG00010013696 | CETN3 | 427104 | 1.108481241 | 2.59E-05 | 0.002740082 | protein-coding | Z |
| ENSGALG00010009464 | SPTLC1 | 426145 | 1.109519292 | 6.76E-09 | 2.30E-06 | protein-coding | Z |
| ENSGALG00010013357 | RICTOR | 427447 | 1.110965667 | 0.000681284 | 0.043019782 | protein-coding | Z |
| ENSGALG00010016308 | RPS6 | 396148 | 1.114123091 | 1.35E-09 | 5.14E-07 | protein-coding | Z |
| ENSGALG00010008862 | ZNF367 | 100858851 | 1.116285838 | 8.14E-05 | 0.007458405 | protein-coding | Z |
| ENSGALG00010005531 | SUSD1 | 427334 | 1.119682663 | 0.000324075 | 0.024052181 | protein-coding | Z |
| ENSGALG00010009031 | SPIN1 | 395344 | 1.12001292 | 5.63E-10 | 2.32E-07 | protein-coding | Z |
| ENSGALG00010002957 | ADGRG2 | 418611 | 1.124355833 | 0.003804389 | 0.167821789 | protein-coding | 1 |
| ENSGALG00010009683 | ADAMTS19 | 415600 | 1.126001496 | 0.007249198 | 0.282394113 | protein-coding | Z |
| ENSGALG00010011059 | ANKRD31 | 427208 | 1.129586702 | 5.54E-07 | 0.000104458 | protein-coding | Z |
| ENSGALG00010012587 | MBLAC2 | 431565 | 1.135102927 | 8.24E-07 | 0.000143316 | protein-coding | Z |
| ENSGALG00010009376 | CDO1 | 427391 | 1.136771879 | 0.00981709 | 0.357524728 | protein-coding | Z |
| ENSGALG00010011437 | DMRT1 | 769693 | 1.138234622 | 2.15E-06 | 0.000333597 | protein-coding | Z |
| ENSGALG00010001160 | TRMT10B | 768672 | 1.139358931 | 3.45E-05 | 0.003465334 | protein-coding | Z |
| ENSGALG00010010951 | POC5 | 427210 | 1.143102746 | 0.000195366 | 0.015851953 | protein-coding | Z |
| ENSGALG00010014810 | FAM189A2 | 427245 | 1.14362691 | 0.000768878 | 0.04666913 | protein-coding | Z |
| ENSGALG00010013497 | FAM172A | 427109 | 1.149520786 | 7.96E-06 | 0.00099384 | protein-coding | Z |
| ENSGALG00010011837 | VLDLR | 396154 | 1.152030173 | 0.005916174 | 0.240018867 | protein-coding | Z |
| ENSGALG00010013102 | NA | NA | 1.153072339 | 0.006096028 | 0.246676465 | ncRNA | Z |
| ENSGALG00010001165 | LOC768709 | 768709 | 1.15332378 | 9.26E-06 | 0.001118756 | protein-coding | Z |
| ENSGALG00010014914 | PTAR1 | 431607 | 1.167607665 | 0.001586388 | 0.084499451 | protein-coding | Z |
| ENSGALG00010013771 | PELO | 430689 | 1.168159759 | 6.64E-07 | 0.000120952 | protein-coding | Z |
| ENSGALG00010013777 | CPLANE1 | 427440 | 1.169230685 | 7.24E-07 | 0.000130372 | protein-coding | Z |
| ENSGALG00010011192 | RPL17 | 426845 | 1.177576517 | 5.80E-10 | 2.33E-07 | protein-coding | Z |
| ENSGALG00010009313 | UNC13B | 100858799 | 1.181764808 | 0.002636306 | 0.125104706 | protein-coding | Z |
| ENSGALG00010012210 | PPWD1 | 427163 | 1.183473797 | 4.72E-08 | 1.30E-05 | protein-coding | Z |
| ENSGALG00010016445 | ZDHHC21 | 427356 | 1.185039273 | 0.003290797 | 0.149373556 | protein-coding | Z |
| ENSGALG00010012188 | ADAMTS6 | 427160 | 1.186747789 | 0.003632457 | 0.162526504 | protein-coding | Z |
| ENSGALG00010011287 | CBWD1 | 427347 | 1.187984488 | 5.33E-05 | 0.005057493 | protein-coding | Z |
| ENSGALG00010012722 | ZSWIM6 | 770670 | 1.193918983 | 0.000176936 | 0.014506891 | protein-coding | Z |
| ENSGALG00010009393 | MARVELD2 | 427393 | 1.200452687 | 9.34E-05 | 0.008266026 | protein-coding | Z |
| ENSGALG00010005594 | MRPL50 | 427307 | 1.201095065 | 0.000181841 | 0.014831408 | protein-coding | Z |
| ENSGALG00010014372 | CCDC171 | 427242 | 1.204742742 | 0.000614101 | 0.040070119 | protein-coding | Z |
| ENSGALG00010013514 | LMBRD2 | 429640 | 1.207560933 | 0.003000871 | 0.138872872 | protein-coding | Z |
| ENSGALG00010006178 | RPS23 | 427323 | 1.208522083 | 3.79E-09 | 1.32E-06 | protein-coding | Z |
| ENSGALG00010007217 | MSH3 | 427318 | 1.212021171 | 5.08E-05 | 0.00488373 | protein-coding | Z |
| ENSGALG00010011253 | KDM4C | 427231 | 1.231137896 | 0.000534101 | 0.035591612 | protein-coding | Z |
| ENSGALG00010010980 | POLK | 374003 | 1.232173066 | 0.002227095 | 0.109673918 | protein-coding | Z |
| ENSGALG00010011904 | KIF24 | 427398 | 1.232198862 | 2.71E-05 | 0.002845444 | protein-coding | Z |
| ENSGALG00010014570 | KIF27 | 427456 | 1.233670884 | 0.001741738 | 0.091528928 | protein-coding | Z |
| ENSGALG00010009534 | XPA | 395659 | 1.238499646 | 1.27E-10 | 5.37E-08 | protein-coding | Z |
| ENSGALG00010010623 | SMAD2 | 395247 | 1.245989235 | 1.29E-05 | 0.001509956 | protein-coding | Z |
| ENSGALG00010011419 | RNF180 | 101747752 | 1.246906458 | 0.002675708 | 0.126590902 | protein-coding | Z |

| | | | | | | | |
|---|---|---|---|---|---|---|---|
| ENSGALG00010001352 | NA | NA | 1.25552014 | 0.006749156 | 0.266898456 | ncRNA | Z |
| ENSGALG00010007770 | HRC8L | 427362 | 1.258405733 | 0.002032509 | 0.103006788 | protein-coding | Z |
| ENSGALG00010006378 | CAAP1 | 427364 | 1.258848043 | 2.63E-09 | 9.35E-07 | protein-coding | Z |
| ENSGALG00010016711 | LOC107052314 | 107052314 | 1.26170493 | 0.007319994 | 0.284444432 | ncRNA | Z |
| ENSGALG00010013219 | LOC769359 | 769359 | 1.261848266 | 0.001838896 | 0.094727358 | ncRNA | Z |
| ENSGALG00010016418 | NFIB | 396209 | 1.266007518 | 0.009605537 | 0.35310494 | protein-coding | Z |
| ENSGALG00010015541 | RASEF | 427451 | 1.279594322 | 0.00049544 | 0.034021471 | protein-coding | Z |
| ENSGALG00010012990 | NA | NA | 1.285473311 | 0.001328346 | 0.0724805 | ncRNA | Z |
| ENSGALG00010010754 | DNAI1 | 431654 | 1.285997619 | 0.000521361 | 0.035040807 | protein-coding | Z |
| ENSGALG00010010844 | CDC14B | 427473 | 1.300615208 | 8.96E-05 | 0.008014159 | protein-coding | Z |
| ENSGALG00010011422 | KATNAL2 | 416384 | 1.30453586 | 0.003276932 | 0.149373556 | protein-coding | Z |
| ENSGALG00010010381 | MAST4 | 427169 | 1.306021626 | 0.000295253 | 0.02255445 | protein-coding | Z |
| ENSGALG00010010625 | FAM169A | 431595 | 1.308746254 | 0.000803206 | 0.048192363 | protein-coding | Z |
| ENSGALG00010013410 | KIAA0825 | 427110 | 1.315990516 | 0.001221238 | 0.067578056 | protein-coding | Z |
| ENSGALG00010012707 | NA | NA | 1.327509222 | 0.004624848 | 0.197343656 | ncRNA | Z |
| ENSGALG00010016468 | TTC39B | 427354 | 1.336153929 | 0.000905329 | 0.05309909 | protein-coding | Z |
| ENSGALG00010016566 | CNTLN | 769413 | 1.361865796 | 5.13E-05 | 0.00489671 | protein-coding | Z |
| ENSGALG00010005519 | PTBP3 | 427333 | 1.364198368 | 0.001190459 | 0.066343745 | protein-coding | Z |
| ENSGALG00010007437 | TUSC1 | 101751372 | 1.388738166 | 1.05E-06 | 0.000175229 | protein-coding | Z |
| ENSGALG00010011937 | SLC1A1 | 427352 | 1.395004321 | 0.001619813 | 0.085987335 | protein-coding | Z |
| ENSGALG00010013136 | NA | NA | 1.3955073 | 0.000860464 | 0.051356285 | ncRNA | Z |
| ENSGALG00010009520 | SPINK4 | 768734 | 1.413560541 | 0.009096102 | 0.337547284 | protein-coding | Z |
| ENSGALG00010011724 | ARHGEF28 | 427201 | 1.418082286 | 0.007920578 | 0.302271259 | protein-coding | Z |
| ENSGALG00010015621 | IDNK | 427453 | 1.428952015 | 0.007798672 | 0.298656 | protein-coding | Z |
| ENSGALG00010001327 | LOC107052436 | 107052436 | 1.432430071 | 0.004521271 | 0.193632168 | ncRNA | Z |
| ENSGALG00010005997 | KIAA1958 | 431637 | 1.44135561 | 0.00970429 | 0.354240522 | protein-coding | Z |
| ENSGALG00010005772 | CR1 | 427338 | 1.453808688 | 0.000303162 | 0.022934889 | protein-coding | Z |
| ENSGALG00010013222 | LOC107052279 | 107052279 | 1.457747352 | 0.000970956 | 0.056315428 | protein-coding | Z |
| ENSGALG00010014934 | MAMDC2 | 427247 | 1.462013909 | 0.002172207 | 0.108333638 | protein-coding | Z |
| ENSGALG00010005881 | CORO2A | 427373 | 1.508722757 | 0.006733702 | 0.266898456 | protein-coding | Z |
| ENSGALG00010013898 | EMB | 395242 | 1.511941313 | 1.28E-07 | 2.94E-05 | protein-coding | Z |
| ENSGALG00010014527 | NA | NA | 1.584977349 | 0.00048938 | 0.033910118 | ncRNA | Z |
| ENSGALG00010014014 | PARP8 | 427198 | 1.610979825 | 1.47E-05 | 0.001679345 | protein-coding | Z |
| ENSGALG00010016670 | NA | NA | 1.643470914 | 0.002958767 | 0.138311328 | ncRNA | Z |
| ENSGALG00010012497 | PLCXD3 | 100857889 | 1.677230463 | 0.002027939 | 0.103006788 | protein-coding | Z |
| ENSGALG00010007686 | NA | NA | 1.710812546 | 0.00472892 | 0.200690748 | ncRNA | Z |
| ENSGALG00010012713 | NA | NA | 1.722506234 | 0.003765397 | 0.167517386 | ncRNA | Z |
| ENSGALG00010009726 | PRR16 | 769337 | 1.723638816 | 0.003338435 | 0.150662531 | protein-coding | Z |
| ENSGALG00010008636 | AGTPBP1 | 427460 | 1.771040212 | 0.000281844 | 0.021635693 | protein-coding | Z |
| ENSGALG00010013455 | PDE6B | 395092 | 1.777250725 | 0.00587442 | 0.238943923 | protein-coding | Z |
| ENSGALG00010007300 | CCDC125 | 427313 | 1.816038251 | 1.74E-05 | 0.001946134 | protein-coding | Z |
| ENSGALG00010018885 | EFEMP1 | 428543 | 1.851365546 | 0.00660698 | 0.262602302 | protein-coding | 3 |
| ENSGALG00010010686 | NA | NA | 1.920974492 | 0.005395886 | 0.222367301 | ncRNA | Z |
| ENSGALG00010013321 | PDCL2 | 422749 | 1.951339972 | 0.002752098 | 0.129422998 | protein-coding | 4 |
| ENSGALG00010029560 | TRIM58 | 770718 | 2.474282988 | 0.009679714 | 0.354168992 | protein-coding | 19 |
| ENSGALG00010001989 | NA | NA | 3.72411401 | 0.008349253 | 0.316584273 | ncRNA | 1 |

### Supplementally table S3. DEGs comparing four strains of cultured PGCs (M/O, GSP, GSN/1, and PNP) (p-adj <0.01)

| Emsembl_ID | Symbol | entrez_ID | MO_Male_Counts | MO_Female_Counts | GSP_Male_Counts | GSP_FemaleCounts | GSN_Male_Counts | GSN_Female_Counts | PNP_Male_Counts | PNP_Female_Counts | p-adj_value | FDR |
|---|---|---|---|---|---|---|---|---|---|---|---|---|
| ENSGALG00010000003 | NA | NA | 11310 | 15661 | 9954 | 11642 | 15916 | 15575 | 24668 | 15171 | 0.000407688 | 0.004491284 |
| ENSGALG00010000005 | NA | NA | 21790 | 24639 | 24297 | 18016 | 26577 | 23319 | 48651 | 36392 | 6.73E-05 | 0.000984442 |
| ENSGALG00010000012 | NA | NA | 124 | 124 | 66 | 56 | 40 | 25 | 22 | 65 | 0.005019096 | 0.033801838 |
| ENSGALG00010000020 | COX2 | COX2 | 114545 | 118479 | 83982 | 80364 | 61399 | 43613 | 92202 | 114639 | 0.000420168 | 0.004598461 |
| ENSGALG00010000022 | ATP8 | ATP8 | 1609 | 2206 | 2385 | 1769 | 1143 | 1123 | 2211 | 3536 | 0.003892987 | 0.027673159 |
| ENSGALG00010000023 | ATP6 | ATP6 | 97578 | 108617 | 82230 | 82297 | 59258 | 42591 | 81385 | 129946 | 0.005633108 | 0.037046543 |
| ENSGALG00010000028 | ND4L | ND4L | 5260 | 5704 | 4422 | 4084 | 3248 | 2874 | 6432 | 6242 | 9.22E-05 | 0.001295146 |
| ENSGALG00010000029 | ND4 | ND4 | 68474 | 71667 | 56125 | 53240 | 41050 | 30242 | 64067 | 81624 | 0.001513151 | 0.012954803 |
| ENSGALG00010000033 | ND5 | ND5 | 90897 | 84813 | 65776 | 65150 | 51523 | 37331 | 84536 | 100731 | 0.000649875 | 0.006606272 |
| ENSGALG00010000034 | CYTB | CYTB | 111542 | 111890 | 84565 | 82075 | 63083 | 43097 | 93213 | 126796 | 0.001131102 | 0.010290136 |
| ENSGALG00010000037 | ND6 | ND6 | 4410 | 3659 | 3118 | 3280 | 2883 | 2548 | 5238 | 4368 | 0.007326878 | 0.045451666 |
| ENSGALG00010000040 | NA | NA | 6 | 10 | 51 | 41 | 36 | 12 | 21 | 21 | 0.002566524 | 0.019738497 |
| ENSGALG00010000042 | CKMT1B | CKMT1 | 70 | 105 | 230 | 161 | 72 | 32 | 95 | 86 | 0.004051174 | 0.02859089 |
| ENSGALG00010000044 | MAP1A | MAP1A | 485 | 541 | 1107 | 694 | 553 | 351 | 1496 | 704 | 0.007413028 | 0.045934935 |
| ENSGALG00010000049 | FES | c-fps | 3 | 3 | 0 | 3 | 6 | 1 | 13 | 21 | 0.001421006 | 0.01233655 |
| ENSGALG00010000050 | CIB1 | CIB4 | 1002 | 1105 | 611 | 603 | 762 | 569 | 1273 | 1737 | 2.27E-06 | 5.40E-05 |
| ENSGALG00010000058 | FURIN | FURIN | 581 | 599 | 942 | 985 | 759 | 487 | 392 | 371 | 8.19E-05 | 0.001171973 |
| ENSGALG00010000070 | NA | NA | 40 | 53 | 6 | 10 | 10 | 23 | 30 | 11 | 0.004232318 | 0.029631544 |
| ENSGALG00010000071 | LOC112532885 | LOC112532885 | 30 | 84 | 15 | 11 | 16 | 2 | 6 | 19 | 0.00124957 | 0.011155353 |
| ENSGALG00010000086 | NA | NA | 277 | 347 | 20 | 18 | 370 | 228 | 299 | 446 | 1.06E-19 | 2.09E-17 |
| ENSGALG00010000087 | NA | NA | 14 | 3 | 1 | 0 | 12 | 10 | 5 | 6 | 0.003240018 | 0.023916601 |
| ENSGALG00010000090 | NA | NA | 299 | 538 | 777 | 699 | 724 | 520 | 860 | 863 | 0.009147475 | 0.054403064 |
| ENSGALG00010000091 | LOC121108173 | LOC121108173 | 13 | 22 | 9 | 2 | 55 | 28 | 20 | 1 | 0.002799118 | 0.021205285 |
| ENSGALG00010000095 | NA | NA | 34 | 98 | 9 | 13 | 16 | 19 | 15 | 28 | 0.000457566 | 0.004917587 |
| ENSGALG00010000096 | NA | NA | 22 | 43 | 6 | 19 | 87 | 41 | 16 | 5 | 2.77E-05 | 0.000467039 |
| ENSGALG00010000103 | NA | NA | 269 | 395 | 378 | 362 | 190 | 139 | 519 | 660 | 0.000206448 | 0.002567079 |
| ENSGALG00010000105 | LOC101752139 | LOC101752139 | 1288 | 772 | 452 | 385 | 614 | 417 | 729 | 798 | 0.000304422 | 0.003537905 |
| ENSGALG00010000106 | NA | NA | 370 | 439 | 262 | 206 | 230 | 273 | 409 | 697 | 0.009453522 | 0.055727427 |
| ENSGALG00010000111 | NA | NA | 145 | 149 | 89 | 53 | 88 | 107 | 229 | 286 | 0.000831744 | 0.00805803 |
| ENSGALG00010000112 | NA | NA | 95 | 107 | 111 | 141 | 22 | 52 | 237 | 158 | 0.002447113 | 0.019048048 |
| ENSGALG00010000113 | LOC121108785 | LOC121108785 | 44 | 222 | 2 | 2 | 103 | 79 | 4 | 2 | 1.78E-17 | 2.70E-15 |
| ENSGALG00010000117 | NA | NA | 253 | 222 | 150 | 152 | 351 | 260 | 196 | 196 | 0.003471475 | 0.025245995 |
| ENSGALG00010000119 | LOC124416997 | LOC124416997 | 189 | 67 | 6 | 1 | 130 | 72 | 16 | 29 | 6.44E-12 | 5.03E-10 |
| ENSGALG00010000122 | LOC121112494 | LOC121112494 | 110 | 67 | 41 | 52 | 113 | 95 | 37 | 37 | 0.000739406 | 0.007329158 |
| ENSGALG00010000123 | NA | NA | 304 | 33 | 447 | 1083 | 588 | 695 | 1029 | 830 | 0.00485046 | 0.033038995 |
| ENSGALG00010000129 | NA | NA | 100 | 129 | 157 | 142 | 22 | 41 | 51 | 67 | 0.001070959 | 0.009871958 |
| ENSGALG00010000130 | NA | NA | 1076 | 823 | 354 | 371 | 660 | 605 | 916 | 732 | 1.06E-05 | 0.000205231 |
| ENSGALG00010000138 | NA | NA | 286 | 273 | 278 | 264 | 76 | 43 | 44 | 57 | 8.88E-11 | 5.52E-09 |
| ENSGALG00010000142 | LOC121108808 | LOC121108808 | 234 | 147 | 155 | 38 | 15 | 15 | 13 | 11 | 5.30E-10 | 2.86E-08 |
| ENSGALG00010000146 | LOC121108947 | LOC121108947 | 333 | 430 | 932 | 792 | 517 | 366 | 475 | 629 | 0.004977866 | 0.033652952 |
| ENSGALG00010000156 | NA | NA | 0 | 809 | 0 | 2 | 760 | 654 | 851 | 914 | 2.57E-06 | 6.01E-05 |
| ENSGALG00010000160 | LOC121106527 | LOC121106527 | 292 | 428 | 520 | 465 | 614 | 640 | 604 | 795 | 0.003844723 | 0.027411697 |
| ENSGALG00010000162 | NA | NA | 1 | 0 | 15 | 14 | 5 | 0 | 6 | 13 | 0.000890397 | 0.008483643 |
| ENSGALG00010000163 | NA | NA | 22 | 20 | 47 | 49 | 94 | 73 | 59 | 132 | 0.000193605 | 0.002425436 |
| ENSGALG00010000164 | NA | NA | 12 | 10 | 8 | 12 | 2 | 10 | 52 | 54 | 2.06E-05 | 0.000360925 |
| ENSGALG00010000165 | LOC107050249 | LOC107050249 | 284 | 916 | 163 | 244 | 541 | 710 | 975 | 1265 | 1.15E-06 | 2.95E-05 |
| ENSGALG00010000166 | LOC107050614 | PFDN6 | 3959 | 3807 | 2343 | 2505 | 2865 | 2195 | 2061 | 2514 | 0.000139383 | 0.001829821 |
| ENSGALG00010000168 | NA | NA | 5 | 8 | 17 | 23 | 20 | 16 | 3 | 5 | 0.001782352 | 0.014782465 |
| ENSGALG00010000169 | NA | NA | 4 | 37 | 32 | 28 | 120 | 62 | 65 | 61 | 0.001526404 | 0.013041547 |
| ENSGALG00010000170 | LOC107051372 | LOC107051372 | 980 | 414 | 803 | 995 | 2291 | 777 | 1853 | 2168 | 0.000340542 | 0.00387726 |
| ENSGALG00010000175 | LOC107049971 | LOC107049971 | 61 | 99 | 272 | 226 | 187 | 105 | 128 | 179 | 0.007454377 | 0.046139831 |
| ENSGALG00010000176 | NA | NA | 251 | 621 | 152 | 50 | 93 | 100 | 94 | 111 | 2.43E-05 | 0.000417744 |
| ENSGALG00010000189 | LOC121108943 | LOC121108943 | 787 | 618 | 14 | 14 | 52 | 36 | 18 | 8 | 2.57E-50 | 2.52E-47 |
| ENSGALG00010000193 | NA | NA | 159 | 341 | 180 | 98 | 59 | 46 | 135 | 165 | 0.001841545 | 0.015153079 |
| ENSGALG00010000196 | KSR2 | KSR2 | 279 | 353 | 102 | 158 | 128 | 122 | 271 | 156 | 0.006440879 | 0.041004267 |
| ENSGALG00010000200 | GAL3ST1 | GAL3ST1 | 328 | 313 | 10 | 49 | 72 | 43 | 191 | 120 | 4.91E-09 | 2.28E-07 |
| ENSGALG00010000204 | LOC107049147 | LOC107049147 | 2004 | 1809 | 700 | 723 | 1395 | 1312 | 1664 | 1196 | 1.64E-07 | 5.27E-06 |
| ENSGALG00010000206 | NA | NA | 569 | 819 | 551 | 649 | 562 | 541 | 289 | 244 | 5.16E-05 | 0.000790936 |
| ENSGALG00010000220 | LOC107049648 | LOC107049648 | 196 | 307 | 234 | 262 | 368 | 336 | 580 | 476 | 0.001389612 | 0.012139676 |
| ENSGALG00010000222 | NA | NA | 8 | 13 | 23 | 18 | 31 | 26 | 6 | 9 | 0.0034368 | 0.02504832 |
| ENSGALG00010000224 | MYBPC2 | MYBPC2 | 1036 | 1326 | 1013 | 448 | 1608 | 1214 | 1102 | 1249 | 0.005037943 | 0.033894565 |
| ENSGALG00010000228 | NA | NA | 19 | 16 | 0 | 1 | 2 | 2 | 5 | 2 | 1.26E-05 | 0.000239999 |
| ENSGALG00010000247 | LOC417013 | LOC417013 | 10 | 2 | 78 | 79 | 20 | 9 | 12 | 7 | 1.61E-08 | 6.53E-07 |
| ENSGALG00010000272 | LOC112531323 | LOC112531323 | 423 | 386 | 148 | 212 | 232 | 191 | 343 | 298 | 0.005955682 | 0.038563099 |
| ENSGALG00010000285 | MYLPF | DTNB | 18 | 22 | 3 | 1 | 9 | 9 | 21 | 5 | 0.005588745 | 0.036829301 |
| ENSGALG00010000289 | QPRT | QPRT | 1036 | 857 | 976 | 1104 | 538 | 371 | 442 | 475 | 2.75E-05 | 0.000464799 |
| ENSGALG00010000291 | NA | NA | 15 | 4 | 17 | 18 | 32 | 56 | 32 | 33 | 0.001712936 | 0.014313294 |
| ENSGALG00010000312 | CABP7 | CABP7 | 332 | 622 | 184 | 222 | 145 | 153 | 341 | 412 | 0.000328066 | 0.003762961 |
| ENSGALG00010000316 | NA | NA | 201 | 1411 | 110 | 208 | 17 | 25 | 199 | 254 | 9.14E-11 | 5.63E-09 |
| ENSGALG00010000320 | NA | NA | 82 | 227 | 33 | 60 | 45 | 14 | 182 | 182 | 8.97E-06 | 0.000178637 |
| ENSGALG00010000330 | NA | NA | 16 | 36 | 0 | 5 | 7 | 2 | 5 | 6 | 0.000478982 | 0.005092076 |
| ENSGALG00010000333 | NA | NA | 5 | 29 | 0 | 0 | 0 | 9 | 4 | 6 | 0.001507914 | 0.01292365 |
| ENSGALG00010000335 | NA | NA | 183 | 341 | 65 | 89 | 54 | 50 | 125 | 134 | 6.57E-06 | 0.000136324 |
| ENSGALG00010000337 | LOC121108736 | LOC121108736 | 339 | 471 | 175 | 168 | 392 | 385 | 134 | 228 | 2.57E-05 | 0.000438157 |
| ENSGALG00010000344 | LOC112531461 | LOC112531461 | 103 | 395 | 25 | 72 | 14 | 25 | 51 | 39 | 3.06E-07 | 9.17E-06 |
| ENSGALG00010000351 | NA | NA | 28 | 7 | 2 | 0 | 11 | 6 | 2 | 3 | 0.000658147 | 0.006682231 |
| ENSGALG00010000363 | NA | NA | 62 | 38 | 3 | 0 | 35 | 8 | 29 | 1 | 0.000389006 | 0.004331163 |
| ENSGALG00010000365 | NA | NA | 45 | 283 | 21 | 29 | 26 | 26 | 65 | 67 | 9.24E-05 | 0.001295146 |
| ENSGALG00010000374 | NA | NA | 24 | 51 | 33 | 48 | 8 | 8 | 69 | 34 | 0.004175756 | 0.029358531 |
| ENSGALG00010000381 | LOC107051255 | LOC107051255 | 486 | 198 | 90 | 96 | 123 | 93 | 461 | 247 | 2.91E-05 | 0.000485472 |
| ENSGALG00010000383 | NA | NA | 170 | 263 | 93 | 74 | 103 | 83 | 152 | 146 | 0.008259226 | 0.050046475 |
| ENSGALG00010000386 | NA | NA | 836 | 849 | 946 | 949 | 363 | 459 | 1573 | 935 | 0.001474842 | 0.012685312 |
| ENSGALG00010000390 | NA | NA | 8 | 11 | 3 | 1 | 25 | 13 | 4 | 9 | 0.00082252 | 0.007987188 |
| ENSGALG00010000402 | PFKFB1 | 6PF-2-K/Fru-2 | 101 | 121 | 63 | 77 | 52 | 45 | 135 | 278 | 0.000345941 | 0.003924898 |
| ENSGALG00010000423 | CARD8 | CARD8 | 4 | 10 | 88 | 105 | 113 | 44 | 74 | 64 | 3.25E-08 | 1.22E-06 |
| ENSGALG00010000429 | NA | NA | 9 | 8 | 4 | 3 | 4 | 0 | 19 | 19 | 0.00134275 | 0.011822698 |
| ENSGALG00010000430 | NA | NA | 13 | 33 | 3 | 1 | 4 | 6 | 31 | 14 | 0.000235628 | 0.002857628 |
| ENSGALG00010000441 | SLC7A4 | SLC7A4 | 578 | 446 | 205 | 299 | 263 | 188 | 375 | 445 | 0.004022461 | 0.028448312 |
| ENSGALG00010000454 | NA | NA | 16 | 20 | 44 | 83 | 79 | 99 | 375 | 161 | 4.89E-10 | 2.67E-08 |
| ENSGALG00010000465 | CUNH17orf49 | C31H17orf49 | 6115 | 5542 | 3632 | 4133 | 3777 | 2643 | 3446 | 4389 | 0.001110365 | 0.01017348 |
| ENSGALG00010000467 | RNASEK | RNASEK | 7690 | 7866 | 4646 | 6474 | 4454 | 4261 | 5290 | 5835 | 0.008373661 | 0.050648074 |
| ENSGALG00010000468 | TESC | TSC | 163 | 214 | 48 | 30 | 76 | 63 | 143 | 186 | 4.55E-06 | 9.95E-05 |
| ENSGALG00010000480 | NA | NA | 25 | 125 | 0 | 0 | 0 | 0 | 0 | 1 | 1.37E-19 | 2.67E-17 |
| ENSGALG00010000509 | LOC107055124 | EDMTF2 | 4 | 18 | 58 | 73 | 48 | 30 | 21 | 29 | 0.000564388 | 0.005862059 |
| ENSGALG00010000543 | NA | NA | 112 | 203 | 10 | 5 | 20 | 13 | 11 | 9 | 4.82E-18 | 7.98E-16 |
| ENSGALG00010000544 | NA | NA | 46 | 64 | 23 | 29 | 71 | 49 | 22 | 21 | 0.001026824 | 0.009549409 |
| ENSGALG00010000546 | FLRT1 | FLRT1 | 508 | 359 | 107 | 125 | 155 | 129 | 292 | 209 | 5.84E-06 | 0.000123228 |
| ENSGALG00010000548 | NA | NA | 252 | 675 | 84 | 110 | 478 | 215 | 82 | 107 | 1.09E-08 | 4.55E-07 |
| ENSGALG00010000553 | NA | NA | 81 | 69 | 177 | 201 | 121 | 182 | 259 | 261 | 0.00206109 | 0.016653818 |
| ENSGALG00010000559 | NA | NA | 98 | 286 | 81 | 133 | 29 | 44 | 51 | 115 | 0.006777675 | 0.042646272 |
| ENSGALG00010000560 | LOC121108768 | LOC121108768 | 1256 | 942 | 276 | 275 | 802 | 549 | 364 | 416 | 2.21E-12 | 1.82E-10 |
| ENSGALG00010000569 | SIPA1 | SIPA1L3L | 912 | 940 | 2245 | 2042 | 1995 | 1630 | 1912 | 1876 | 5.56E-07 | 1.55E-05 |
| ENSGALG00010000579 | NA | NA | 94 | 98 | 39 | 64 | 150 | 152 | 120 | 72 | 0.001216682 | 0.010925945 |
| ENSGALG00010000585 | SPTBN2 | SPTBN2 | 4581 | 3992 | 2581 | 2755 | 2693 | 1805 | 1879 | 2521 | 2.17E-05 | 0.000377586 |
| ENSGALG00010000590 | DNAH5 | DNAH5 | 55 | 69 | 40 | 68 | 175 | 130 | 53 | 27 | 1.19E-05 | 0.000227352 |
| ENSGALG00010000598 | NA | NA | 8 | 0 | 0 | 0 | 40 | 10 | 62 | 29 | 7.79E-09 | 3.42E-07 |
| ENSGALG00010000599 | FAM134B | FAM134B | 418 | 598 | 1382 | 1249 | 875 | 661 | 754 | 675 | 1.72E-05 | 0.000309666 |
| ENSGALG00010000601 | LOC121107771 | LOC121107771 | 7 | 0 | 0 | 0 | 69 | 43 | 103 | 33 | 3.47E-14 | 3.56E-12 |
| ENSGALG00010000602 | NA | NA | 3 | 10 | 0 | 0 | 7 | 7 | 22 | 12 | 3.18E-05 | 0.000524816 |
| ENSGALG00010000607 | NA | NA | 28 | 7 | 2 | 0 | 56 | 9 | 30 | 10 | 0.00038304 | 0.004280651 |
| ENSGALG00010000610 | MED10 | MED10 | 2658 | 2794 | 1666 | 2090 | 1946 | 1437 | 1507 | 1680 | 0.001337649 | 0.011796722 |
| ENSGALG00010000612 | NA | NA | 10 | 1 | 1 | 0 | 24 | 5 | 2 | 0 | 0.001009278 | 0.009417674 |
| ENSGALG00010000613 | LIN37 | LIN37 | 1346 | 1099 | 1804 | 2065 | 973 | 624 | 889 | 1138 | 0.000495897 | 0.005248533 |
| ENSGALG00010000628 | NA | NA | 26 | 0 | 2 | 0 | 85 | 11 | 0 | 3 | 6.98E-05 | 0.001016492 |
| ENSGALG00010000641 | LOC121108149 | LOC121108149 | 65 | 67 | 221 | 113 | 48 | 32 | 54 | 54 | 0.000688442 | 0.006910054 |
| ENSGALG00010000642 | ZPBP1 | ZPBP | 11 | 12 | 80 | 52 | 33 | 16 | 38 | 31 | 0.000752999 | 0.007441824 |
| ENSGALG00010000645 | LOC121112235 | LOC121112235 | 26 | 1 | 2 | 2 | 139 | 68 | 1 | 7 | 2.24E-10 | 1.29E-08 |
| ENSGALG00010000646 | NA | NA | 983 | 1212 | 1502 | 1594 | 1991 | 1381 | 2279 | 2576 | 5.83E-06 | 0.000123124 |
| ENSGALG00010000651 | MYO7L2 | SM1 | 306 | 298 | 107 | 146 | 286 | 151 | 410 | 357 | 0.000317992 | 0.003667548 |
| ENSGALG00010000658 | NA | NA | 1435 | 325 | 1593 | 1530 | 163 | 142 | 239 | 355 | 2.31E-09 | 1.12E-07 |
| ENSGALG00010000659 | LOC121109782 | LOC121109782 | 204 | 214 | 385 | 1107 | 492 | 329 | 774 | 751 | 4.56E-05 | 0.000713917 |
| ENSGALG00010000662 | NA | NA | 382 | 12 | 493 | 392 | 8 | 9 | 11 | 29 | 5.22E-08 | 1.87E-06 |
| ENSGALG00010000685 | LOC112530906 | LOC112530906 | 43 | 72 | 1874 | 1969 | 2645 | 1598 | 2129 | 1732 | 4.81E-64 | 6.70E-61 |
| ENSGALG00010000688 | LOC121108155 | LOC121108155 | 24 | 13 | 404 | 415 | 499 | 342 | 487 | 482 | 1.13E-26 | 3.58E-24 |
| ENSGALG00010000697 | LOC107050463 | LOC107050463 | 325 | 295 | 262 | 186 | 507 | 442 | 903 | 634 | 8.24E-07 | 2.18E-05 |
| ENSGALG00010000699 | NA | NA | 0 | 1 | 10 | 4 | 13 | 8 | 4 | 3 | 0.002192805 | 0.017492198 |
| ENSGALG00010000700 | NA | NA | 9 | 16 | 282 | 317 | 161 | 108 | 267 | 324 | 3.39E-19 | 6.44E-17 |
| ENSGALG00010000702 | DDC | DDC | 4 | 2 | 27 | 75 | 291 | 213 | 26 | 50 | 5.85E-21 | 1.32E-18 |
| ENSGALG00010000705 | LOC769743 | LOC769743 | 68 | 27 | 36 | 68 | 177 | 111 | 87 | 68 | 0.001214875 | 0.010915583 |

| | | | | | | | | | | | | |
|---|---|---|---|---|---|---|---|---|---|---|---|---|
| ENSGALG00010000714 | FBXL7 | FBXL7 | 237 | 550 | 652 | 1004 | 363 | 227 | 348 | 335 | 0.002354785 | 0.018466999 |
| ENSGALG00010000715 | PCDH20 | protocadherin-20 | 10 | 15 | 9 | 12 | 29 | 69 | 168 | 123 | 5.02E-11 | 3.29E-09 |
| ENSGALG00010000716 | NA | NA | 15 | 0 | 2 | 1 | 162 | 44 | 0 | 7 | 1.34E-09 | 6.80E-08 |
| ENSGALG00010000717 | NDFIP2 | NDFIP2 | 2850 | 2622 | 1118 | 1831 | 2199 | 2265 | 3599 | 2963 | 2.38E-05 | 0.000410854 |
| ENSGALG00010000719 | NA | NA | 13 | 6 | 3 | 0 | 30 | 29 | 52 | 42 | 5.20E-09 | 2.40E-07 |
| ENSGALG00010000721 | LOC107051303 | LOC107051303 | 38 | 23 | 34 | 11 | 150 | 47 | 89 | 61 | 0.000884756 | 0.008449171 |
| ENSGALG00010000726 | POU4F1 | BRN-3A | 436 | 517 | 64 | 100 | 38 | 59 | 209 | 104 | 4.61E-11 | 3.07E-09 |
| ENSGALG00010000728 | FAM105B | FAM105B | 76 | 47 | 157 | 140 | 121 | 89 | 62 | 56 | 0.008001059 | 0.048907718 |
| ENSGALG00010000729 | EDNRB | EDNRB | 2937 | 2779 | 960 | 688 | 726 | 731 | 2407 | 2312 | 4.67E-16 | 6.24E-14 |
| ENSGALG00010000730 | INTS5 | INTS5 | 2281 | 1620 | 1357 | 1399 | 2500 | 1384 | 2242 | 2921 | 0.003839515 | 0.027386243 |
| ENSGALG00010000731 | NA | NA | 25 | 1 | 0 | 11 | 134 | 48 | 68 | 23 | 0.000296378 | 0.003456432 |
| ENSGALG00010000732 | NA | NA | 1966 | 2379 | 1570 | 2116 | 1235 | 981 | 1249 | 1460 | 0.004006292 | 0.02836998 |
| ENSGALG00010000736 | NA | NA | 17 | 2 | 62 | 110 | 101 | 116 | 71 | 94 | 2.12E-06 | 5.08E-05 |
| ENSGALG00010000737 | LOC121107768 | LOC121107768 | 5 | 0 | 0 | 0 | 11 | 1 | 23 | 11 | 0.00026558 | 0.00317027 |
| ENSGALG00010000738 | NA | NA | 36 | 7 | 68 | 88 | 126 | 163 | 145 | 127 | 2.65E-05 | 0.000450138 |
| ENSGALG00010000743 | NA | NA | 5 | 0 | 24 | 12 | 55 | 45 | 64 | 120 | 2.04E-10 | 1.19E-08 |
| ENSGALG00010000760 | CDH10 | cadherin-10 | 214 | 386 | 278 | 486 | 364 | 112 | 72 | 120 | 0.001964669 | 0.015930885 |
| ENSGALG00010000767 | NA | NA | 39 | 2 | 90 | 114 | 175 | 163 | 141 | 127 | 0.000323986 | 0.003723834 |
| ENSGALG00010000768 | NA | NA | 1534 | 1708 | 718 | 1622 | 1861 | 1390 | 1907 | 3495 | 0.002091615 | 0.016837706 |
| ENSGALG00010000769 | NA | NA | 29 | 3 | 63 | 59 | 124 | 87 | 133 | 123 | 6.44E-05 | 0.000945746 |
| ENSGALG00010000770 | CDH12 | CDH12 | 328 | 199 | 5 | 39 | 136 | 118 | 52 | 91 | 3.49E-07 | 1.03E-05 |
| ENSGALG00010000771 | NA | NA | 1317 | 221 | 2863 | 2793 | 5257 | 4042 | 4598 | 4380 | 2.36E-06 | 5.60E-05 |
| ENSGALG00010000773 | NA | NA | 23 | 3 | 126 | 109 | 223 | 173 | 344 | 279 | 5.86E-12 | 4.59E-10 |
| ENSGALG00010000774 | NA | NA | 71 | 99 | 7 | 6 | 5 | 3 | 50 | 59 | 7.95E-14 | 7.90E-12 |
| ENSGALG00010000780 | NA | NA | 12 | 19 | 1 | 0 | 7 | 11 | 28 | 11 | 0.000105949 | 0.001447767 |
| ENSGALG00010000785 | CLN5 | CLN5 | 595 | 598 | 222 | 262 | 471 | 415 | 692 | 692 | 4.47E-06 | 9.78E-05 |
| ENSGALG00010000786 | ANKRD33B | ANKRD33B | 27 | 66 | 375 | 291 | 55 | 14 | 22 | 32 | 1.90E-12 | 1.60E-10 |
| ENSGALG00010000788 | GATD3AL1 | ES1ML1 | 496 | 319 | 955 | 832 | 1040 | 646 | 1019 | 1203 | 5.71E-05 | 0.000856688 |
| ENSGALG00010000791 | ACOD1 | IRG1 | 4 | 10 | 123 | 91 | 41 | 19 | 12 | 16 | 2.77E-10 | 1.57E-08 |
| ENSGALG00010000802 | SEMA5A | SEMA5A | 585 | 570 | 761 | 991 | 437 | 575 | 1998 | 1491 | 8.85E-08 | 3.07E-06 |
| ENSGALG00010000811 | LOC770405 | LOC770405 | 28 | 30 | 5 | 11 | 84 | 45 | 18 | 25 | 1.00E-06 | 2.60E-05 |
| ENSGALG00010000813 | LOC101750376 | LOC101750376 | 637 | 458 | 172 | 139 | 284 | 223 | 297 | 388 | 1.63E-05 | 0.000296365 |
| ENSGALG00010000822 | BASP1 | CAP-23 | 2510 | 3993 | 3056 | 3817 | 2140 | 1302 | 2294 | 2754 | 0.009461961 | 0.055737852 |
| ENSGALG00010000825 | NA | NA | 482 | 818 | 165 | 253 | 271 | 237 | 340 | 322 | 3.30E-05 | 0.000542108 |
| ENSGALG00010000826 | NA | NA | 4 | 759 | 11 | 35 | 29 | 48 | 20 | 10 | 5.11E-05 | 0.000783629 |
| ENSGALG00010000827 | NA | NA | 8 | 14 | 55 | 93 | 53 | 25 | 3 | 14 | 2.24E-06 | 5.35E-05 |
| ENSGALG00010000830 | NA | NA | 948 | 1325 | 1 | 27 | 45 | 41 | 16 | 29 | 8.49E-36 | 4.44E-33 |
| ENSGALG00010000831 | NA | NA | 3760 | 3298 | 517 | 1285 | 393 | 1527 | 3485 | 3100 | 6.43E-05 | 0.000945333 |
| ENSGALG00010000835 | NA | NA | 903 | 850 | 5 | 56 | 110 | 235 | 10 | 94 | 1.06E-08 | 4.44E-07 |
| ENSGALG00010000847 | NA | NA | 11 | 2 | 3 | 638 | 685 | 534 | 0 | 2 | 2.23E-10 | 1.29E-08 |
| ENSGALG00010000861 | NA | NA | 2 | 4 | 30 | 24 | 24 | 16 | 11 | 12 | 0.000221035 | 0.002712138 |
| ENSGALG00010000888 | NA | NA | 4 | 4 | 6 | 140 | 33 | 34 | 45 | 33 | 0.001715826 | 0.014330279 |
| ENSGALG00010000896 | NA | NA | 28 | 17 | 0 | 3 | 3 | 33 | 3 | 7 | 0.001678297 | 0.014087238 |
| ENSGALG00010000901 | PCDH9 | protocadherin-9 | 31 | 48 | 234 | 475 | 24 | 1 | 5 | 14 | 7.70E-16 | 1.01E-13 |
| ENSGALG00010000909 | NA | NA | 0 | 10 | 1 | 14 | 0 | 34 | 0 | 0 | 0.00890442 | 0.053241741 |
| ENSGALG00010000925 | NA | NA | 399 | 399 | 37 | 84 | 59 | 64 | 139 | 105 | 1.64E-10 | 9.66E-09 |
| ENSGALG00010000935 | NA | NA | 22 | 0 | 0 | 0 | 23 | 14 | 35 | 34 | 1.40E-05 | 0.000259715 |
| ENSGALG00010000962 | NA | NA | 1 | 3 | 18 | 34 | 10 | 4 | 4 | 2 | 4.62E-05 | 0.000721245 |
| ENSGALG00010001024 | NA | NA | 538 | 3680 | 1964 | 3545 | 0 | 0 | 0 | 0 | 3.70E-59 | 4.76E-56 |
| ENSGALG00010001025 | NA | NA | 341 | 3173 | 1196 | 2349 | 0 | 0 | 0 | 0 | 4.94E-46 | 4.13E-43 |
| ENSGALG00010001026 | NA | NA | 1103 | 5542 | 1445 | 2924 | 9 | 38 | 287 | 268 | 4.93E-21 | 1.13E-18 |
| ENSGALG00010001031 | NA | NA | 120 | 171 | 0 | 12 | 0 | 48 | 80 | 126 | 0.001791839 | 0.014828291 |
| ENSGALG00010001096 | NA | NA | 0 | 171 | 0 | 7 | 0 | 2 | 2 | 13 | 0.000414584 | 0.004551093 |
| ENSGALG00010001103 | NA | NA | 1479 | 2131 | 40 | 298 | 27 | 795 | 958 | 1186 | 0.004833525 | 0.032953928 |
| ENSGALG00010001114 | MED12L | MED12L | 303 | 374 | 972 | 814 | 698 | 587 | 785 | 570 | 0.000117793 | 0.001583711 |
| ENSGALG00010001134 | TUBA3E | TUBA3D | 6439 | 8036 | 9639 | 11017 | 9108 | 7104 | 6194 | 7652 | 0.009160875 | 0.054436829 |
| ENSGALG00010001137 | NA | NA | 2 | 0 | 31 | 46 | 26 | 26 | 15 | 35 | 6.21E-07 | 1.71E-05 |
| ENSGALG00010001138 | SPTSSB | C9H3orf57 | 614 | 241 | 790 | 817 | 411 | 351 | 1741 | 743 | 0.001481378 | 0.01273497 |
| ENSGALG00010001158 | FRMPD1 | FRMPD1 | 161 | 102 | 47 | 30 | 60 | 50 | 271 | 99 | 0.000274052 | 0.003257432 |
| ENSGALG00010001176 | IL12A | IL12A | 89 | 236 | 19 | 27 | 48 | 24 | 32 | 14 | 1.10E-07 | 3.73E-06 |
| ENSGALG00010001181 | MFSD1 | MFSD1 | 3895 | 3668 | 4326 | 7074 | 4185 | 3476 | 2997 | 3042 | 0.0011024 | 0.010122693 |
| ENSGALG00010001196 | LSMEM1 | C1H7orf53 | 4 | 10 | 12 | 9 | 15 | 16 | 1 | 2 | 0.00169961 | 0.01422328 |
| ENSGALG00010001201 | TUBB6 | TUBB6 | 308 | 356 | 913 | 730 | 866 | 413 | 765 | 863 | 0.000278504 | 0.003299506 |
| ENSGALG00010001202 | CFTR | CFTR | 20 | 19 | 43 | 37 | 128 | 95 | 49 | 39 | 2.64E-07 | 8.01E-06 |
| ENSGALG00010001203 | TMX3 | TXNDC10 | 1749 | 1940 | 1194 | 1309 | 1020 | 770 | 1375 | 1575 | 0.003058051 | 0.022805065 |
| ENSGALG00010001207 | ASZ1 | GASZ | 3534 | 2328 | 5144 | 7269 | 4597 | 3328 | 3082 | 2992 | 1.94E-05 | 0.000343393 |
| ENSGALG00010001208 | MOV10L1 | MOV10L1 | 2048 | 2034 | 3810 | 3895 | 3407 | 2337 | 4296 | 3352 | 0.000107673 | 0.001467728 |
| ENSGALG00010001209 | CDH7 | cadherin-7 | 256 | 607 | 116 | 148 | 140 | 83 | 123 | 87 | 1.78E-06 | 4.37E-05 |
| ENSGALG00010001216 | SHOX2 | SHOX2 | 6 | 22 | 7 | 12 | 3 | 1 | 2 | 2 | 0.008957446 | 0.053501286 |
| ENSGALG00010001220 | PLXNB2 | PLXNB2 | 1490 | 2020 | 1445 | 1415 | 1722 | 1791 | 3683 | 2455 | 0.000120907 | 0.001620947 |
| ENSGALG00010001253 | ZBBX | ZBBX | 475 | 91 | 12 | 13 | 61 | 87 | 434 | 48 | 1.30E-05 | 0.00024649 |
| ENSGALG00010001272 | NA | NA | 143 | 85 | 24 | 16 | 29 | 17 | 44 | 18 | 1.02E-05 | 0.000199897 |
| ENSGALG00010001276 | LOC121108470 | LOC121108470 | 43 | 22 | 8 | 3 | 9 | 2 | 4 | 4 | 0.000121587 | 0.001626875 |
| ENSGALG00010001292 | LOX | LOX | 549 | 616 | 99 | 51 | 336 | 162 | 914 | 127 | 9.52E-05 | 0.00133067 |
| ENSGALG00010001313 | EMILIN2 | EMILIN2 | 1367 | 1636 | 2580 | 3686 | 668 | 389 | 1028 | 918 | 3.60E-19 | 6.76E-17 |
| ENSGALG00010001318 | NA | NA | 15 | 13 | 81 | 100 | 60 | 15 | 7 | 7 | 7.69E-08 | 2.71E-06 |
| ENSGALG00010001334 | EBF3 | EBF3 | 9 | 10 | 53 | 62 | 61 | 32 | 60 | 51 | 5.36E-05 | 0.000815865 |
| ENSGALG00010001335 | NA | NA | 71 | 82 | 20 | 20 | 19 | 3 | 34 | 18 | 0.000106856 | 0.001458969 |
| ENSGALG00010001350 | LOC107049524 | SPDYC | 405 | 130 | 1071 | 1011 | 858 | 714 | 1015 | 994 | 3.39E-05 | 0.000554318 |
| ENSGALG00010001351 | CAPN1 | calpain | 875 | 681 | 2903 | 2751 | 2334 | 1615 | 1890 | 2073 | 1.36E-11 | 9.99E-10 |
| ENSGALG00010001380 | SCRN1 | SCRN1 | 2329 | 1978 | 1160 | 1728 | 1082 | 736 | 1063 | 1515 | 0.00094534 | 0.008915642 |
| ENSGALG00010001386 | FKBP14 | FKBP14 | 6208 | 4871 | 1770 | 1926 | 2780 | 2374 | 2983 | 2649 | 2.92E-13 | 2.64E-11 |
| ENSGALG00010001407 | FOXP2 | FOXP2 | 136 | 99 | 319 | 283 | 220 | 166 | 106 | 119 | 0.000918013 | 0.008721909 |
| ENSGALG00010001429 | LOC112533497 | LOC112533497 | 91 | 136 | 27 | 17 | 24 | 15 | 36 | 29 | 7.64E-07 | 2.05E-05 |
| ENSGALG00010001441 | HMGCLL1 | HMGCLL1 | 12 | 11 | 74 | 39 | 65 | 34 | 40 | 23 | 0.001465247 | 0.012615772 |
| ENSGALG00010001448 | GPR37 | SCGPR1 | 54 | 44 | 51 | 51 | 63 | 43 | 569 | 693 | 2.83E-30 | 1.18E-27 |
| ENSGALG00010001463 | DSEL | DSEL | 753 | 1130 | 664 | 1147 | 448 | 353 | 420 | 315 | 2.43E-05 | 0.000417744 |
| ENSGALG00010001465 | TACC2 | TACC2 | 895 | 1064 | 331 | 578 | 465 | 463 | 772 | 587 | 0.001690837 | 0.014163335 |
| ENSGALG00010001480 | JAKMIP3 | C10orf39 | 68 | 49 | 27 | 35 | 56 | 28 | 135 | 72 | 0.009652775 | 0.056662161 |
| ENSGALG00010001483 | LOC121110182 | LOC121110182 | 1218 | 2857 | 34 | 597 | 52 | 616 | 12 | 259 | 0.003834142 | 0.027371284 |
| ENSGALG00010001497 | LEKR1 | LEKR1 | 65 | 71 | 255 | 241 | 130 | 118 | 113 | 107 | 0.000306813 | 0.003559777 |
| ENSGALG00010001499 | TIPARP | TIPARP | 714 | 665 | 1305 | 1289 | 852 | 438 | 581 | 503 | 0.000183032 | 0.002319048 |
| ENSGALG00010001510 | AP1S2 | AP1S2 | 1684 | 1694 | 3350 | 3977 | 3177 | 1581 | 1738 | 1991 | 1.55E-05 | 0.000284231 |
| ENSGALG00010001520 | KCNAB1 | KCNAB3 | 61 | 60 | 125 | 260 | 200 | 125 | 88 | 103 | 0.00111555 | 0.010204195 |
| ENSGALG00010001537 | NA | NA | 5 | 10 | 76 | 36 | 15 | 10 | 27 | 14 | 0.000218946 | 0.002692435 |
| ENSGALG00010001550 | DDX43 | DDX43 | 1359 | 1573 | 2604 | 3126 | 2190 | 1505 | 1730 | 1534 | 0.000177058 | 0.002258773 |
| ENSGALG00010001554 | ARHGAP33L4 | ARHGAP33L4 | 162 | 179 | 73 | 60 | 253 | 209 | 334 | 323 | 3.22E-08 | 1.21E-06 |
| ENSGALG00010001556 | EPB41L3 | EPB41L3 | 840 | 546 | 257 | 266 | 611 | 549 | 680 | 643 | 1.61E-05 | 0.000294659 |
| ENSGALG00010001558 | CCKBR | CCK-CHR | 171 | 209 | 110 | 97 | 52 | 27 | 146 | 167 | 7.47E-05 | 0.001080268 |
| ENSGALG00010001559 | HDAC10 | HDAC10 | 562 | 931 | 462 | 525 | 617 | 568 | 348 | 415 | 0.004566216 | 0.031520281 |
| ENSGALG00010001568 | NA | NA | 17 | 19 | 31 | 65 | 55 | 54 | 65 | 52 | 0.008583209 | 0.051653792 |
| ENSGALG00010001570 | SMPD1 | SMPD1 | 733 | 691 | 249 | 332 | 257 | 241 | 279 | 422 | 8.68E-05 | 0.001227521 |
| ENSGALG00010001576 | NA | NA | 1 | 3 | 11 | 15 | 18 | 1 | 0 | 2 | 0.007986252 | 0.048835067 |
| ENSGALG00010001588 | ARHGAP33L1 | ARHGAP33L1 | 15 | 11 | 5 | 6 | 16 | 12 | 34 | 34 | 0.000939507 | 0.008870646 |
| ENSGALG00010001611 | MAPK12 | MAPK12 | 416 | 350 | 668 | 745 | 705 | 530 | 787 | 683 | 0.006382519 | 0.040727243 |
| ENSGALG00010001612 | HPX | HPX | 86 | 72 | 15 | 32 | 32 | 25 | 51 | 97 | 0.002526812 | 0.019513899 |
| ENSGALG00010001629 | NA | NA | 83 | 76 | 116 | 163 | 147 | 154 | 255 | 184 | 0.009695896 | 0.056875328 |
| ENSGALG00010001633 | NA | NA | 347 | 492 | 134 | 165 | 166 | 200 | 351 | 392 | 0.000225742 | 0.00276584 |
| ENSGALG00010001648 | NA | NA | 17 | 65 | 157 | 144 | 98 | 104 | 51 | 57 | 0.001768351 | 0.014695512 |
| ENSGALG00010001652 | NA | NA | 2 | 11 | 48 | 29 | 33 | 13 | 12 | 11 | 0.002709403 | 0.020694486 |
| ENSGALG00010001666 | PIEZO2 | FAM38B | 125 | 133 | 181 | 192 | 184 | 140 | 582 | 302 | 9.24E-05 | 0.001295146 |
| ENSGALG00010001672 | NA | NA | 30 | 25 | 1 | 6 | 7 | 4 | 6 | 3 | 0.000105626 | 0.001447763 |
| ENSGALG00010001691 | CTPS2 | CTPS2 | 711 | 511 | 1041 | 1480 | 1101 | 813 | 834 | 890 | 0.004118293 | 0.02901556 |
| ENSGALG00010001692 | ALDH18A1 | ALDH18A1 | 1789 | 1561 | 1949 | 1339 | 1801 | 2550 | 4922 | 5541 | 8.98E-10 | 4.69E-08 |
| ENSGALG00010001699 | ASB15 | ASB15 | 9 | 13 | 1 | 0 | 11 | 10 | 16 | 8 | 0.000452624 | 0.004870732 |
| ENSGALG00010001703 | NA | NA | 364 | 223 | 18 | 31 | 36 | 21 | 108 | 122 | 7.22E-15 | 8.29E-13 |
| ENSGALG00010001705 | HTRA1 | HTRA1 | 19245 | 15694 | 3900 | 5023 | 7644 | 5580 | 10912 | 5720 | 3.19E-13 | 2.87E-11 |
| ENSGALG00010001707 | NA | NA | 212 | 281 | 16 | 12 | 60 | 82 | 58 | 34 | 1.44E-13 | 1.36E-11 |
| ENSGALG00010001708 | AADAC | AADACL2 | 20 | 46 | 46 | 68 | 95 | 132 | 85 | 105 | 0.00076268 | 0.007502011 |
| ENSGALG00010001713 | PNPLA4 | PNPLA4 | 26 | 50 | 190 | 193 | 114 | 65 | 70 | 44 | 1.53E-05 | 0.000281704 |
| ENSGALG00010001716 | TRIL | TRIL | 1918 | 2456 | 1295 | 1354 | 1832 | 1800 | 1312 | 1166 | 3.39E-05 | 0.000554318 |
| ENSGALG00010001726 | NA | NA | 97 | 89 | 8 | 5 | 27 | 33 | 68 | 26 | 4.27E-07 | 1.23E-05 |
| ENSGALG00010001743 | LOC107049019 | LOC107049019 | 291 | 390 | 52 | 35 | 77 | 68 | 264 | 88 | 1.78E-07 | 5.67E-06 |
| ENSGALG00010001746 | NA | NA | 8 | 11 | 27 | 20 | 0 | 0 | 6 | 13 | 5.13E-06 | 0.00011022 |
| ENSGALG00010001755 | CUZD1 | CUZD1 | 42 | 82 | 6 | 3 | 26 | 17 | 22 | 18 | 4.96E-07 | 1.41E-05 |
| ENSGALG00010001756 | ARHGAP33L6 | ARHGAP33L6 | 37 | 50 | 144 | 121 | 211 | 179 | 228 | 207 | 3.46E-07 | 1.02E-05 |
| ENSGALG00010001763 | LOC107049808 | LOC107049808 | 37 | 58 | 56 | 32 | 96 | 91 | 181 | 161 | 3.47E-05 | 0.000565971 |
| ENSGALG00010001771 | WDR73 | WDR73 | 1590 | 1796 | 1490 | 1584 | 994 | 667 | 1413 | 1821 | 0.007159784 | 0.044530822 |
| ENSGALG00010001773 | NRCAM | NRCAM | 6 | 4 | 22 | 18 | 14 | 16 | 51 | 31 | 0.00035407 | 0.004006243 |
| ENSGALG00010001785 | CNTN1 | F11 | 34 | 42 | 182 | 123 | 117 | 59 | 110 | 112 | 0.001320802 | 0.011660453 |
| ENSGALG00010001811 | ACADSB | ACADSB | 738 | 777 | 1055 | 1480 | 585 | 622 | 1222 | 1242 | 0.005419717 | 0.035942187 |

| | | | | | | | | | | | | |
|---|---|---|---|---|---|---|---|---|---|---|---|---|
| ENSGALG00010001818 | ADGRB3 | BAI3 | 20 | 58 | 151 | 158 | 43 | 29 | 46 | 44 | 0.000103432 | 0.001419173 |
| ENSGALG00010001828 | LOC100858869 | LOC100858869 | 2366 | 1901 | 1038 | 2149 | 73 | 174 | 1591 | 1392 | 1.72E-14 | 1.87E-12 |
| ENSGALG00010001856 | CHST15 | GALNAC4S-6ST | 633 | 1161 | 377 | 353 | 299 | 268 | 637 | 766 | 7.58E-06 | 0.000153938 |
| ENSGALG00010001871 | LOC107049095 | LOC107049095 | 1850 | 1683 | 830 | 1105 | 955 | 905 | 1410 | 1143 | 0.001644962 | 0.013870136 |
| ENSGALG00010001874 | NA | NA | 1 | 1 | 13 | 9 | 3 | 9 | 19 | 11 | 0.00504572 | 0.033933228 |
| ENSGALG00010001919 | LOC429348 | LOC429348 | 93 | 152 | 28 | 47 | 154 | 128 | 170 | 94 | 0.000168738 | 0.002165856 |
| ENSGALG00010001922 | NA | NA | 69 | 30 | 127 | 146 | 179 | 102 | 32 | 46 | 6.79E-05 | 0.000991623 |
| ENSGALG00010001924 | NA | NA | 19 | 26 | 85 | 58 | 22 | 25 | 52 | 68 | 0.007474496 | 0.046230118 |
| ENSGALG00010001933 | NA | NA | 27 | 24 | 69 | 63 | 130 | 123 | 40 | 34 | 2.35E-07 | 7.22E-06 |
| ENSGALG00010001967 | LAMA1 | LAMA1 | 1608 | 3119 | 1175 | 1850 | 3066 | 2370 | 2226 | 1912 | 0.000446774 | 0.004816495 |
| ENSGALG00010001976 | THRSP | THRSPA | 1 | 0 | 7 | 5 | 16 | 10 | 11 | 3 | 0.001355886 | 0.011907289 |
| ENSGALG00010001989 | NA | NA | 3800 | 203 | 4 | 4 | 20 | 17 | 88 | 88 | 1.75E-25 | 5.23E-23 |
| ENSGALG00010002022 | ERNI | ERNI | 21221 | 38187 | 3478 | 6343 | 3855 | 6621 | 16380 | 14682 | 2.33E-15 | 2.86E-13 |
| ENSGALG00010002024 | KLHL31 | KLHL31 | 5 | 4 | 19 | 20 | 23 | 17 | 12 | 9 | 0.005670071 | 0.037233787 |
| ENSGALG00010002035 | NA | NA | 14 | 19 | 0 | 2 | 1 | 0 | 0 | 1 | 1.03E-06 | 2.66E-05 |
| ENSGALG00010002046 | FAM126A | DRCTNNB1A | 5106 | 4829 | 8060 | 9482 | 8230 | 6544 | 6295 | 5834 | 1.38E-05 | 0.000256961 |
| ENSGALG00010002061 | NA | NA | 633 | 1221 | 1 | 13 | 1 | 1 | 0 | 3 | 9.87E-65 | 1.50E-61 |
| ENSGALG00010002063 | NA | NA | 634 | 544 | 46 | 77 | 102 | 71 | 354 | 324 | 1.76E-18 | 3.13E-16 |
| ENSGALG00010002088 | OZFL | RJF.Contig1.LOC05.OZF1 | 7 | 2 | 17 | 20 | 2 | 0 | 13 | 20 | 0.00032257 | 0.003710107 |
| ENSGALG00010002096 | NA | NA | 0 | 2 | 9 | 14 | 13 | 18 | 5 | 4 | 0.000234063 | 0.002846913 |
| ENSGALG00010002113 | GDPD4 | GDPD4 | 6 | 3 | 25 | 7 | 7 | 12 | 107 | 125 | 1.24E-11 | 9.18E-10 |
| ENSGALG00010002126 | MGAM | MGAM | 130 | 162 | 28 | 4 | 21 | 27 | 40 | 46 | 3.00E-06 | 6.86E-05 |
| ENSGALG00010002138 | MYO7A | myosin-VIIa | 21 | 16 | 7 | 4 | 13 | 23 | 50 | 57 | 2.88E-05 | 0.000481633 |
| ENSGALG00010002160 | CAPN5 | calpain-5 | 170 | 161 | 73 | 100 | 82 | 67 | 231 | 198 | 0.001916683 | 0.015617559 |
| ENSGALG00010002185 | NA | NA | 1601 | 1181 | 190 | 212 | 154 | 110 | 226 | 251 | 1.18E-30 | 5.06E-28 |
| ENSGALG00010002186 | ICK | ICK | 760 | 1074 | 2134 | 1811 | 1888 | 1201 | 1205 | 829 | 3.26E-05 | 0.000536126 |
| ENSGALG00010002208 | NA | NA | 24 | 25 | 0 | 0 | 1 | 0 | 1 | 1 | 1.54E-11 | 1.11E-09 |
| ENSGALG00010002220 | NA | NA | 23 | 51 | 111 | 127 | 98 | 57 | 38 | 37 | 0.000890388 | 0.008483643 |
| ENSGALG00010002239 | AGPAT5 | AGPAT5 | 2670 | 2487 | 1477 | 1419 | 1766 | 1238 | 2413 | 2633 | 6.25E-05 | 0.000923401 |
| ENSGALG00010002271 | LOC418658 | ARSD | 10 | 6 | 5 | 3 | 21 | 18 | 19 | 11 | 0.005752862 | 0.037614173 |
| ENSGALG00010002274 | LOC417067 | LOC417067 | 21 | 11 | 3 | 6 | 31 | 31 | 6 | 14 | 0.000327217 | 0.003755805 |
| ENSGALG00010002282 | TSKU | C-TSK | 524 | 640 | 186 | 150 | 178 | 116 | 423 | 436 | 4.98E-09 | 2.31E-07 |
| ENSGALG00010002285 | CAV1 | caveolin-1 | 63 | 150 | 641 | 652 | 329 | 196 | 140 | 381 | 9.22E-06 | 0.000183024 |
| ENSGALG00010002297 | NA | NA | 31 | 21 | 105 | 129 | 100 | 67 | 88 | 95 | 0.000267189 | 0.00318492 |
| ENSGALG00010002325 | NA | NA | 17 | 15 | 3 | 3 | 1 | 4 | 3 | 4 | 0.003180984 | 0.023564095 |
| ENSGALG00010002333 | MHCYL | MR1 | 37 | 64 | 5 | 1 | 45 | 45 | 39 | 49 | 8.18E-08 | 2.87E-06 |
| ENSGALG00010002335 | TMEM47 | TMEM47 | 460 | 289 | 884 | 1137 | 1485 | 1095 | 392 | 382 | 2.47E-14 | 2.63E-12 |
| ENSGALG00010002338 | CD99 | CD99 | 763 | 746 | 2055 | 1985 | 1266 | 882 | 1257 | 1360 | 1.31E-06 | 3.32E-05 |
| ENSGALG00010002341 | FAM49A | CYRIA | 5227 | 5494 | 3170 | 4189 | 3520 | 3021 | 4834 | 4446 | 0.007290898 | 0.045245263 |
| ENSGALG00010002346 | FBXO15 | FBXO15 | 163 | 148 | 378 | 475 | 369 | 191 | 203 | 166 | 0.000700618 | 0.007007015 |
| ENSGALG00010002356 | WNT11 | WNT-11 | 125 | 242 | 56 | 30 | 77 | 75 | 105 | 175 | 0.000747746 | 0.007398656 |
| ENSGALG00010002358 | FREM2 | FREM2 | 74 | 67 | 168 | 257 | 63 | 84 | 236 | 309 | 3.88E-05 | 0.000622796 |
| ENSGALG00010002359 | CMTM6 | CKLFSF6 | 328 | 582 | 143 | 249 | 287 | 304 | 638 | 585 | 0.000216057 | 0.00266279 |
| ENSGALG00010002362 | TRPC4 | TRPC4 | 1 | 0 | 5 | 5 | 0 | 0 | 11 | 12 | 1.89E-05 | 0.000337113 |
| ENSGALG00010002367 | STK31 | STK31 | 3516 | 3909 | 6843 | 8252 | 6224 | 5204 | 5726 | 5236 | 2.00E-06 | 4.87E-05 |
| ENSGALG00010002392 | DFNA5 | GSDME | 676 | 858 | 939 | 1041 | 1781 | 984 | 364 | 386 | 8.91E-13 | 7.68E-11 |
| ENSGALG00010002397 | NA | NA | 90 | 106 | 172 | 162 | 41 | 37 | 0 | 1 | 2.19E-21 | 5.15E-19 |
| ENSGALG00010002399 | CSMD1 | CSMD1 | 238 | 875 | 925 | 1035 | 550 | 244 | 266 | 301 | 0.002200467 | 0.017536581 |
| ENSGALG00010002403 | ZNFY3 | RJF.Contig3.LOC04.ZNF3 | 677 | 52 | 46 | 85 | 20 | 23 | 3 | 9 | 2.24E-10 | 1.29E-08 |
| ENSGALG00010002414 | MHCY4 | RJF.Contig1.LOC17.YF4 | 42 | 5 | 15 | 10 | 97 | 124 | 15 | 22 | 1.23E-06 | 3.15E-05 |
| ENSGALG00010002423 | LOC121106436 | LOC121106436 | 19 | 33 | 1 | 1 | 4 | 0 | 12 | 3 | 1.39E-05 | 0.000257591 |
| ENSGALG00010002430 | NUMA1 | NUMA | 5253 | 4357 | 8366 | 7802 | 5619 | 3180 | 5283 | 5821 | 0.008237286 | 0.049967886 |
| ENSGALG00010002436 | MHCY4 | RJF.Contig1.LOC17.YF4 | 92 | 40 | 13 | 2 | 0 | 2 | 3 | 14 | 8.63E-08 | 3.01E-06 |
| ENSGALG00010002445 | EDRF1 | C10orf137 | 1343 | 1005 | 1811 | 2092 | 1813 | 1254 | 1284 | 1153 | 0.008483938 | 0.051185402 |
| ENSGALG00010002457 | LOC417058 | LOC417058 | 40 | 108 | 31 | 35 | 26 | 36 | 2 | 16 | 0.001156349 | 0.010474203 |
| ENSGALG00010002466 | UROS | UROS | 177 | 153 | 457 | 488 | 355 | 180 | 134 | 140 | 1.06E-05 | 0.00020569 |
| ENSGALG00010002473 | LOC107049124 | LOC107049124 | 234 | 97 | 13 | 22 | 32 | 42 | 2 | 5 | 1.82E-14 | 1.96E-12 |
| ENSGALG00010002474 | HPCAL1 | REM-1 | 47 | 128 | 398 | 208 | 223 | 70 | 99 | 102 | 0.005885913 | 0.038274465 |
| ENSGALG00010002475 | NA | NA | 171 | 217 | 396 | 376 | 296 | 185 | 128 | 149 | 0.0010515 | 0.009730157 |
| ENSGALG00010002487 | BCCIP | BCCIP | 1445 | 1936 | 2754 | 3538 | 2346 | 1545 | 1714 | 2091 | 0.001864228 | 0.015267483 |
| ENSGALG00010002493 | LOC121110175 | LOC121110175 | 5 | 7 | 17 | 12 | 36 | 32 | 38 | 45 | 1.35E-05 | 0.000253467 |
| ENSGALG00010002501 | TCEANC | TCEANC | 4 | 10 | 38 | 48 | 24 | 13 | 8 | 11 | 0.000194216 | 0.002431269 |
| ENSGALG00010002503 | NA | NA | 7159 | 10215 | 5137 | 9498 | 8 | 10 | 2 | 5 | 4.08E-239 | 6.81E-235 |
| ENSGALG00010002533 | LPAR2 | EDG4 | 974 | 847 | 598 | 744 | 409 | 330 | 469 | 641 | 0.003293637 | 0.024248134 |
| ENSGALG00010002554 | EGFL6 | EGFL6 | 7 | 16 | 82 | 63 | 63 | 30 | 38 | 21 | 0.00023284 | 0.002838242 |
| ENSGALG00010002557 | NA | NA | 92 | 122 | 29 | 17 | 19 | 23 | 35 | 25 | 8.67E-06 | 0.000173596 |
| ENSGALG00010002586 | GPR63 | GPR63 | 1519 | 1587 | 1004 | 1153 | 1237 | 1174 | 2185 | 1734 | 0.002731054 | 0.02079334 |
| ENSGALG00010002599 | KIFC1 | HSET | 7371 | 9567 | 5211 | 5384 | 5000 | 2941 | 5977 | 7094 | 4.95E-05 | 0.000763155 |
| ENSGALG00010002600 | TRPA1 | TRPA1 | 79 | 36 | 9 | 32 | 21 | 5 | 70 | 80 | 0.001034132 | 0.009585365 |
| ENSGALG00010002604 | BLEC3 | BLEC3 | 110 | 36 | 12 | 16 | 9 | 17 | 67 | 79 | 4.07E-05 | 0.000643892 |
| ENSGALG00010002605 | EOMES | EOMES | 4430 | 4065 | 1454 | 2419 | 3811 | 3230 | 3204 | 2344 | 5.39E-07 | 1.50E-05 |
| ENSGALG00010002609 | PDIA6 | TXNDC7 | 14966 | 12102 | 7592 | 9533 | 7741 | 5438 | 9641 | 10941 | 0.000146882 | 0.00191473 |
| ENSGALG00010002614 | GPAT3 | AGPAT 10 | 5522 | 4579 | 4621 | 5735 | 5113 | 3823 | 8907 | 8106 | 0.000123661 | 0.001651972 |
| ENSGALG00010002632 | HPGD | HPGD | 72 | 49 | 13 | 19 | 36 | 34 | 43 | 41 | 0.004915853 | 0.033382261 |
| ENSGALG00010002636 | NA | NA | 12 | 29 | 2 | 7 | 5 | 2 | 31 | 35 | 3.98E-05 | 0.000632875 |
| ENSGALG00010002637 | LOC121110113 | LOC121110113 | 15 | 4 | 31 | 29 | 16 | 24 | 47 | 51 | 0.008350792 | 0.050546335 |
| ENSGALG00010002640 | NA | NA | 5 | 2 | 18 | 25 | 5 | 1 | 7 | 3 | 0.002273218 | 0.017996223 |
| ENSGALG00010002647 | FAM168A | FAM168A | 1257 | 1645 | 3071 | 3064 | 2199 | 1296 | 1804 | 2222 | 0.000383077 | 0.004280651 |
| ENSGALG00010002661 | ACBD7 | ACBD7 | 90 | 88 | 120 | 129 | 35 | 34 | 187 | 105 | 0.002877813 | 0.021673729 |
| ENSGALG00010002678 | CCNI | CCNI | 2342 | 2546 | 4228 | 4112 | 3354 | 2212 | 2443 | 2912 | 0.002794443 | 0.02118908 |
| ENSGALG00010002701 | TMEM196 | TMEM196 | 55 | 75 | 13 | 47 | 19 | 8 | 9 | 20 | 0.00211711 | 0.017010165 |
| ENSGALG00010002713 | OLFM2 | OLFM2 | 1215 | 847 | 464 | 543 | 689 | 468 | 846 | 926 | 0.001128919 | 0.010281468 |
| ENSGALG00010002716 | NHLRC3 | NHLRC3 | 237 | 214 | 184 | 213 | 159 | 176 | 427 | 426 | 0.005677731 | 0.037268752 |
| ENSGALG00010002731 | NA | NA | 5 | 25 | 41 | 83 | 32 | 11 | 11 | 6 | 0.000923479 | 0.008753927 |
| ENSGALG00010002733 | RDH8 | RDH8 | 442 | 180 | 83 | 93 | 438 | 281 | 319 | 236 | 2.55E-05 | 0.000435252 |
| ENSGALG00010002754 | MIR12263 | gga-mir-12263 | 1 | 2 | 12 | 13 | 2 | 2 | 4 | 5 | 0.005911113 | 0.038373386 |
| ENSGALG00010002758 | GREB1 | GREB1 | 13 | 89 | 4 | 0 | 28 | 36 | 14 | 13 | 1.77E-05 | 0.000318129 |
| ENSGALG00010002763 | SEPTIN11 | 11-Sep | 1098 | 1565 | 628 | 1200 | 539 | 552 | 1025 | 1246 | 0.008611399 | 0.05177804 |
| ENSGALG00010002764 | NA | NA | 11 | 10 | 69 | 67 | 107 | 15 | 11 | 8 | 9.91E-06 | 0.000194462 |
| ENSGALG00010002824 | NXPH1 | NXPH1 | 25 | 35 | 5 | 6 | 11 | 7 | 15 | 17 | 0.001744717 | 0.014528008 |
| ENSGALG00010002848 | TRIB2 | TRB2 | 2510 | 3836 | 2039 | 2077 | 2443 | 1560 | 1710 | 2074 | 0.003702473 | 0.02659034 |
| ENSGALG00010002863 | NA | NA | 2 | 6 | 29 | 16 | 11 | 17 | 43 | 21 | 0.003431875 | 0.025023338 |
| ENSGALG00010002868 | PPP1R9A | PPP1R9A | 103 | 168 | 1165 | 772 | 487 | 256 | 662 | 317 | 1.24E-08 | 5.14E-07 |
| ENSGALG00010002870 | FABP5 | FABP5 | 18 | 43 | 40 | 21 | 39 | 16 | 79 | 158 | 0.000483768 | 0.005139684 |
| ENSGALG00010002871 | LOC423967 | C10orf143 | 660 | 1186 | 1338 | 2228 | 1273 | 639 | 596 | 778 | 0.00083007 | 0.008051149 |
| ENSGALG00010002873 | MRAP2 | C6orf117 | 229 | 92 | 13 | 10 | 14 | 22 | 30 | 23 | 8.07E-11 | 5.09E-09 |
| ENSGALG00010002880 | RASGEF1B | RASGEF1B | 2580 | 1481 | 771 | 1193 | 1513 | 1027 | 1632 | 936 | 0.006679931 | 0.042126418 |
| ENSGALG00010002904 | PON1 | PON2 | 440 | 508 | 1019 | 882 | 468 | 369 | 546 | 581 | 0.003155886 | 0.023398918 |
| ENSGALG00010002907 | GABRG3 | GABRG3 | 70 | 21 | 4 | 6 | 5 | 11 | 90 | 31 | 1.39E-05 | 0.000257591 |
| ENSGALG00010002913 | NXNL1 | NXNL1 | 100 | 82 | 27 | 39 | 41 | 29 | 62 | 65 | 0.009354632 | 0.055300536 |
| ENSGALG00010002949 | LOC107052718 | pol | 0 | 0 | 1703 | 1534 | 377 | 343 | 1 | 1 | 1.24E-120 | 6.92E-117 |
| ENSGALG00010002956 | SCRG1 | SCRG1 | 53 | 68 | 16 | 14 | 19 | 34 | 35 | 32 | 0.006599686 | 0.041793846 |
| ENSGALG00010002965 | PRSS35 | PRSS35 | 687 | 586 | 116 | 119 | 245 | 154 | 629 | 352 | 3.74E-10 | 2.06E-08 |
| ENSGALG00010002972 | ME1 | NADP-ME | 513 | 238 | 68 | 102 | 103 | 107 | 255 | 216 | 8.83E-06 | 0.000176242 |
| ENSGALG00010002982 | HAND2 | dHAND | 13 | 30 | 40 | 199 | 49 | 58 | 12 | 24 | 0.000280826 | 0.003316725 |
| ENSGALG00010002991 | BLB1 | B-LB | 120 | 139 | 88 | 82 | 105 | 72 | 34 | 27 | 0.000136464 | 0.001795742 |
| ENSGALG00010003001 | MAP6 | NAU | 1138 | 2181 | 245 | 445 | 592 | 488 | 806 | 854 | 3.87E-11 | 2.61E-09 |
| ENSGALG00010003018 | MOGAT2 | MOGAT2 | 24 | 168 | 1 | 2 | 5 | 8 | 9 | 27 | 5.69E-09 | 2.59E-07 |
| ENSGALG00010003019 | TAPBP | TPN | 331 | 119 | 26 | 69 | 93 | 62 | 93 | 91 | 0.000477776 | 0.00508897 |
| ENSGALG00010003027 | SCIN | SCIN | 65 | 215 | 87 | 160 | 402 | 105 | 64 | 74 | 0.006598481 | 0.041793846 |
| ENSGALG00010003035 | DGAT2 | DGAT2 | 209 | 380 | 50 | 66 | 78 | 73 | 168 | 264 | 4.41E-07 | 1.27E-05 |
| ENSGALG00010003053 | NA | NA | 24 | 84 | 4 | 6 | 22 | 32 | 40 | 18 | 0.000282765 | 0.003330738 |
| ENSGALG00010003058 | NA | NA | 16 | 86 | 5 | 5 | 23 | 25 | 18 | 8 | 0.000309469 | 0.003579138 |
| ENSGALG00010003061 | ABCB5 | XXbac-BPG246D15.9 | 431 | 145 | 8 | 14 | 166 | 145 | 293 | 92 | 2.07E-09 | 1.03E-07 |
| ENSGALG00010003073 | NA | NA | 0 | 0 | 12 | 6 | 18 | 14 | 4 | 5 | 4.12E-06 | 9.12E-05 |
| ENSGALG00010003079 | SP8 | SP8 | 6 | 12 | 38 | 96 | 9 | 17 | 15 | 12 | 4.53E-05 | 0.000710784 |
| ENSGALG00010003094 | LOC107053759 | LOC107053759 | 120 | 179 | 385 | 553 | 471 | 314 | 348 | 483 | 6.21E-05 | 0.000921037 |
| ENSGALG00010003098 | NA | NA | 157 | 188 | 63 | 159 | 2 | 8 | 4 | 12 | 2.48E-15 | 3.02E-13 |
| ENSGALG00010003099 | NA | NA | 36 | 14 | 123 | 88 | 61 | 22 | 50 | 45 | 0.009397836 | 0.055477868 |
| ENSGALG00010003108 | PPP1R3B | PPP1R3B | 2813 | 4447 | 814 | 872 | 1094 | 831 | 1689 | 2193 | 2.00E-18 | 3.52E-16 |
| ENSGALG00010003110 | BF1 | B-F | 936 | 1400 | 457 | 603 | 462 | 460 | 613 | 968 | 0.000383445 | 0.004280651 |
| ENSGALG00010003131 | TAP1 | TAP1 | 797 | 855 | 290 | 446 | 411 | 391 | 589 | 745 | 0.000537273 | 0.005636478 |
| ENSGALG00010003157 | NA | NA | 1 | 4 | 21 | 23 | 19 | 17 | 16 | 23 | 0.00057269 | 0.005937217 |
| ENSGALG00010003168 | NA | NA | 143 | 21 | 12 | 10 | 21 | 33 | 91 | 60 | 0.001173686 | 0.010596781 |
| ENSGALG00010003176 | PRAG1 | PRAGMIN | 401 | 694 | 145 | 242 | 162 | 263 | 645 | 586 | 6.24E-05 | 0.000923401 |
| ENSGALG00010003187 | RHOG | RHOG | 602 | 511 | 196 | 207 | 224 | 219 | 358 | 303 | 8.90E-05 | 0.001256847 |
| ENSGALG00010003192 | RGCC | C13orf15 | 78 | 156 | 485 | 460 | 301 | 136 | 117 | 186 | 3.93E-05 | 0.000627842 |
| ENSGALG00010003193 | ACOT9 | ACOT9 | 279 | 354 | 784 | 1072 | 876 | 529 | 723 | 696 | 3.79E-06 | 8.50E-05 |
| ENSGALG00010003197 | NA | NA | 7 | 1 | 28 | 8 | 49 | 23 | 18 | 21 | 0.0015549 | 0.013247688 |
| ENSGALG00010003198 | PRKG2 | PRKG2 | 898 | 1074 | 330 | 520 | 684 | 526 | 894 | 463 | 0.002303174 | 0.018147406 |
| ENSGALG00010003199 | NA | NA | 71 | 109 | 15 | 7 | 27 | 31 | 64 | 30 | 3.12E-05 | 0.000516559 |

| | | | | | | | | | | | | |
|---|---|---|---|---|---|---|---|---|---|---|---|---|
| ENSGALG00010003205 | TNX | TN | 69 | 51 | 223 | 265 | 267 | 135 | 271 | 187 | 4.61E-05 | 0.000720269 |
| ENSGALG00010003219 | NA | NA | 11 | 242 | 0 | 0 | 1 | 20 | 0 | 0 | 1.95E-11 | 1.38E-09 |
| ENSGALG00010003227 | PLAC8L1 | PLAC8 | 12 | 245 | 17 | 23 | 32 | 61 | 20 | 8 | 0.001566461 | 0.013329273 |
| ENSGALG00010003240 | PLACL2 | PLACL2 | 725 | 2613 | 2877 | 4162 | 2586 | 2781 | 791 | 1155 | 1.99E-05 | 0.000350367 |
| ENSGALG00010003252 | PHEX | PHEX | 182 | 89 | 32 | 52 | 34 | 40 | 90 | 50 | 0.003417445 | 0.024929524 |
| ENSGALG00010003260 | ACP5 | ACP5 | 1349 | 444 | 515 | 527 | 368 | 281 | 327 | 276 | 0.001111581 | 0.010179041 |
| ENSGALG00010003268 | SMS | SMS | 22623 | 16201 | 8336 | 10766 | 9882 | 10176 | 12168 | 13336 | 4.43E-05 | 0.000696412 |
| ENSGALG00010003295 | IAH1 | IAH1 | 954 | 573 | 1654 | 1599 | 1280 | 931 | 1274 | 1740 | 0.005569423 | 0.03673094 |
| ENSGALG00010003305 | CALR | CALR | 37727 | 33414 | 20709 | 25170 | 22088 | 13955 | 21308 | 27185 | 0.001130577 | 0.010290136 |
| ENSGALG00010003323 | STMN2 | SCG10 | 832 | 1424 | 413 | 862 | 835 | 181 | 1147 | 2009 | 0.004245427 | 0.029698441 |
| ENSGALG00010003334 | TAC1 | TAC1 | 31 | 49 | 28 | 2 | 2 | 7 | 47 | 80 | 0.000844904 | 0.00816659 |
| ENSGALG00010003338 | NA | NA | 0 | 0 | 0 | 1087 | 14 | 14 | 75 | 13 | 1.36E-07 | 4.50E-06 |
| ENSGALG00010003358 | DNAJC15 | DNAJD1 | 735 | 1664 | 269 | 297 | 635 | 516 | 689 | 849 | 8.67E-08 | 3.01E-06 |
| ENSGALG00010003360 | NA | NA | 0 | 1 | 22 | 27 | 28 | 33 | 16 | 20 | 4.19E-08 | 1.54E-06 |
| ENSGALG00010003364 | ANKRD6 | ANKRD6 | 258 | 399 | 545 | 887 | 976 | 584 | 315 | 246 | 1.52E-07 | 4.92E-06 |
| ENSGALG00010003367 | NA | NA | 16 | 0 | 1 | 4 | 25 | 37 | 38 | 38 | 0.00013063 | 0.001729378 |
| ENSGALG00010003372 | ENOX1 | ENOX1 | 79 | 21 | 45 | 15 | 24 | 9 | 192 | 261 | 1.47E-07 | 4.80E-06 |
| ENSGALG00010003374 | SYCE2 | SYCE2 | 74 | 64 | 225 | 282 | 159 | 121 | 75 | 93 | 4.33E-05 | 0.000683657 |
| ENSGALG00010003382 | ASNS | ASNS | 1266 | 1534 | 1734 | 1130 | 1465 | 1907 | 3499 | 3634 | 1.70E-06 | 4.22E-05 |
| ENSGALG00010003400 | PKIA | PKI | 632 | 650 | 1588 | 2101 | 1330 | 1016 | 761 | 808 | 9.32E-10 | 4.85E-08 |
| ENSGALG00010003431 | PM20D2 | ACY1L2 | 760 | 928 | 1668 | 2271 | 1678 | 1427 | 1553 | 1681 | 6.16E-06 | 0.000128771 |
| ENSGALG00010003434 | NA | NA | 4 | 2 | 6 | 3 | 39 | 23 | 32 | 46 | 8.71E-09 | 3.77E-07 |
| ENSGALG00010003438 | KCNS3 | KCNS3 | 79 | 50 | 12 | 23 | 21 | 22 | 77 | 72 | 0.000211046 | 0.002616468 |
| ENSGALG00010003439 | GMIP | GMIP | 40 | 22 | 13 | 15 | 55 | 46 | 54 | 39 | 0.002587571 | 0.019863795 |
| ENSGALG00010003452 | NA | NA | 2 | 9 | 1 | 0 | 23 | 8 | 10 | 12 | 0.000307729 | 0.003561468 |
| ENSGALG00010003453 | NA | NA | 8 | 100 | 746 | 988 | 218 | 258 | 57 | 175 | 2.42E-07 | 7.42E-06 |
| ENSGALG00010003459 | NA | NA | 10 | 38 | 44 | 56 | 62 | 35 | 1 | 2 | 2.89E-08 | 1.10E-06 |
| ENSGALG00010003466 | TSC22D1 | TSC22D1 | 4326 | 3560 | 2429 | 2083 | 2603 | 2191 | 4604 | 3561 | 0.00016182 | 0.00208507 |
| ENSGALG00010003476 | NA | NA | 45 | 29 | 43 | 53 | 19 | 11 | 7 | 19 | 0.009840978 | 0.057584881 |
| ENSGALG00010003479 | SYDE1 | SYDE1 | 218 | 26 | 12 | 7 | 13 | 14 | 33 | 62 | 1.29E-05 | 0.000243558 |
| ENSGALG00010003480 | NA | NA | 564 | 341 | 51 | 100 | 69 | 89 | 240 | 256 | 2.16E-09 | 1.06E-07 |
| ENSGALG00010003505 | NA | NA | 2 | 4 | 17 | 19 | 16 | 27 | 36 | 27 | 0.000191367 | 0.002404602 |
| ENSGALG00010003535 | OR52L1 | OR52M1 | 3 | 7 | 4 | 4 | 0 | 2 | 16 | 11 | 0.00873774 | 0.052432717 |
| ENSGALG00010003536 | NA | NA | 794 | 790 | 297 | 565 | 1092 | 738 | 1038 | 577 | 0.000928223 | 0.008788929 |
| ENSGALG00010003543 | SOX11 | SOX11 | 3095 | 5134 | 1308 | 1751 | 2436 | 1553 | 2042 | 2363 | 1.94E-08 | 7.62E-07 |
| ENSGALG00010003557 | LOC121107015 | LOC121107015 | 463 | 595 | 648 | 757 | 636 | 2034 | 2481 | 1002 | 0.000478486 | 0.005091106 |
| ENSGALG00010003586 | NA | NA | 0 | 0 | 16 | 12 | 3 | 0 | 0 | 0 | 9.29E-07 | 2.44E-05 |
| ENSGALG00010003603 | NA | NA | 5 | 3 | 1 | 0 | 8 | 10 | 22 | 12 | 9.67E-05 | 0.001345158 |
| ENSGALG00010003632 | TECRL | SRD5A2L2 | 3 | 0 | 20 | 29 | 14 | 21 | 7 | 16 | 0.000344403 | 0.003910095 |
| ENSGALG00010003633 | LOC121106978 | LOC121106978 | 18 | 21 | 2 | 4 | 6 | 132 | 26 | 3 | 0.000191674 | 0.002405672 |
| ENSGALG00010003639 | NA | NA | 11994 | 13268 | 4882 | 9187 | 8884 | 7953 | 4007 | 4496 | 1.56E-10 | 9.26E-09 |
| ENSGALG00010003668 | NA | NA | 89 | 91 | 204 | 132 | 53 | 29 | 67 | 106 | 0.005600336 | 0.036876602 |
| ENSGALG00010003692 | ZNFY2 | RJF.Contig1.LOC11.ZNF2 | 10 | 39 | 72 | 85 | 25 | 8 | 6 | 3 | 1.57E-06 | 3.94E-05 |
| ENSGALG00010003700 | REST | REST | 9692 | 7063 | 4323 | 6579 | 3961 | 3957 | 5058 | 3006 | 0.000413163 | 0.00454262 |
| ENSGALG00010003701 | SARAF | TMEM66 | 1689 | 1325 | 639 | 1067 | 652 | 594 | 849 | 912 | 0.000682687 | 0.006864656 |
| ENSGALG00010003719 | RPGR | RPGR | 895 | 1055 | 1972 | 1962 | 1501 | 1012 | 1506 | 1306 | 0.001602001 | 0.013583276 |
| ENSGALG00010003722 | MHCY38 | Y-FA | 3 | 44 | 5 | 7 | 4 | 8 | 51 | 94 | 3.91E-05 | 0.000625831 |
| ENSGALG00010003723 | NA | NA | 271 | 379 | 34 | 22 | 120 | 123 | 146 | 123 | 1.66E-11 | 1.18E-09 |
| ENSGALG00010003724 | NA | NA | 736 | 780 | 299 | 345 | 1030 | 785 | 994 | 1080 | 2.72E-11 | 1.88E-09 |
| ENSGALG00010003725 | NA | NA | 33 | 26 | 151 | 148 | 60 | 57 | 65 | 48 | 5.53E-05 | 0.000834142 |
| ENSGALG00010003728 | NA | NA | 1 | 0 | 20 | 15 | 31 | 22 | 18 | 15 | 3.44E-07 | 1.02E-05 |
| ENSGALG00010003732 | NA | NA | 57 | 99 | 7 | 11 | 15 | 11 | 16 | 13 | 1.58E-08 | 6.42E-07 |
| ENSGALG00010003740 | PITX2 | Ptx2 | 178 | 208 | 691 | 567 | 613 | 324 | 138 | 133 | 7.90E-11 | 5.01E-09 |
| ENSGALG00010003743 | ARL9 | ARL9 | 805 | 799 | 619 | 768 | 690 | 440 | 395 | 435 | 0.007729711 | 0.047509718 |
| ENSGALG00010003749 | MHCY15 | RJF.Contig1.LOC44.YF15 | 4 | 30 | 7 | 2 | 38 | 19 | 2 | 2 | 0.000133327 | 0.001760004 |
| ENSGALG00010003756 | MHCY7 | RJF.Contig1.LOC27.YF7 | 79 | 456 | 8 | 9 | 65 | 113 | 21 | 16 | 4.82E-12 | 3.82E-10 |
| ENSGALG00010003759 | PRDM14 | PRDM14 | 5160 | 5163 | 3066 | 3109 | 3369 | 2375 | 5212 | 4437 | 1.88E-05 | 0.000334393 |
| ENSGALG00010003765 | SPINK2 | SPIK2 | 474 | 629 | 150 | 114 | 118 | 164 | 417 | 428 | 3.17E-08 | 1.20E-06 |
| ENSGALG00010003767 | LOC121106918 | LOC121106918 | 51 | 1 | 0 | 1 | 12 | 5 | 0 | 0 | 9.63E-06 | 0.000190015 |
| ENSGALG00010003772 | MHCY6 | MHCIA9 | 84 | 289 | 4 | 6 | 22 | 30 | 150 | 189 | 3.03E-14 | 3.15E-12 |
| ENSGALG00010003774 | NA | NA | 39 | 68 | 27 | 17 | 51 | 43 | 94 | 66 | 0.008122061 | 0.049425912 |
| ENSGALG00010003779 | NA | NA | 6 | 3 | 43 | 33 | 9 | 7 | 22 | 29 | 5.96E-05 | 0.000886504 |
| ENSGALG00010003786 | LOC121106502 | LOC121106502 | 12 | 16 | 20 | 22 | 10 | 5 | 3 | 0 | 0.000517688 | 0.005454979 |
| ENSGALG00010003793 | TRAM1L1 | TRAM1 | 4689 | 3765 | 2822 | 3102 | 2384 | 1911 | 3246 | 3816 | 0.002497138 | 0.019338356 |
| ENSGALG00010003796 | NA | NA | 207 | 398 | 24 | 31 | 159 | 99 | 1 | 0 | 1.18E-30 | 5.06E-28 |
| ENSGALG00010003804 | PSME1 | PSME1 | 2971 | 2218 | 456 | 566 | 897 | 1140 | 2365 | 1674 | 9.25E-14 | 9.04E-12 |
| ENSGALG00010003812 | BG8 | 15iTBGa-1 | 82 | 64 | 97 | 92 | 98 | 87 | 280 | 266 | 2.06E-05 | 0.000360843 |
| ENSGALG00010003814 | TPP1 | TPP1 | 3876 | 3990 | 2358 | 2545 | 2489 | 1879 | 2828 | 3379 | 0.000673211 | 0.006802114 |
| ENSGALG00010003817 | BF2 | B-F | 1935 | 1672 | 1172 | 1564 | 1300 | 1100 | 2234 | 3113 | 0.000178934 | 0.002279186 |
| ENSGALG00010003820 | SH3YL1 | SH3YL1 | 1011 | 544 | 429 | 363 | 855 | 700 | 1068 | 813 | 0.000792906 | 0.007749149 |
| ENSGALG00010003827 | RRH | RRH | 16 | 45 | 7 | 6 | 6 | 6 | 5 | 10 | 0.002575646 | 0.019790433 |
| ENSGALG00010003835 | DCAF11 | DCAF11 | 954 | 946 | 665 | 699 | 432 | 358 | 706 | 775 | 0.002717493 | 0.020736678 |
| ENSGALG00010003845 | CFI | CFI | 137 | 76 | 228 | 227 | 257 | 231 | 236 | 295 | 0.006265333 | 0.040071277 |
| ENSGALG00010003854 | NA | NA | 22 | 16 | 2 | 3 | 14 | 13 | 45 | 23 | 6.04E-05 | 0.000896838 |
| ENSGALG00010003918 | HTR1E | HTR1E | 64 | 66 | 4 | 8 | 22 | 18 | 43 | 27 | 9.91E-07 | 2.58E-05 |
| ENSGALG00010003933 | LOC107050163 | TECR | 685 | 700 | 1085 | 965 | 973 | 631 | 1276 | 1451 | 0.003923286 | 0.027864834 |
| ENSGALG00010003937 | LOC121108129 | LOC121108129 | 1368 | 1262 | 960 | 1082 | 1476 | 778 | 788 | 635 | 0.002438699 | 0.019000254 |
| ENSGALG00010003965 | SNTG1 | SNTG1 | 283 | 188 | 26 | 84 | 183 | 138 | 290 | 144 | 0.000610082 | 0.006247355 |
| ENSGALG00010003972 | PENK | PENK | 68 | 33 | 149 | 73 | 190 | 102 | 58 | 79 | 0.009567956 | 0.056263082 |
| ENSGALG00010003983 | NA | NA | 7 | 18 | 34 | 36 | 10 | 8 | 48 | 35 | 0.002210474 | 0.017597056 |
| ENSGALG00010003990 | ICA1 | ICA1 | 785 | 1085 | 2768 | 3675 | 1358 | 868 | 869 | 841 | 7.20E-17 | 1.04E-14 |
| ENSGALG00010004006 | NKAIN3 | FAM77D | 261 | 380 | 610 | 802 | 557 | 305 | 327 | 236 | 0.000814518 | 0.007918687 |
| ENSGALG00010004036 | ADCYAP1 | PACAP | 353 | 146 | 21 | 21 | 238 | 164 | 186 | 122 | 1.11E-09 | 5.71E-08 |
| ENSGALG00010004044 | RNF138 | RNF138 | 2581 | 2219 | 1477 | 1660 | 1296 | 1255 | 1735 | 1743 | 0.008800271 | 0.052717402 |
| ENSGALG00010004053 | NA | NA | 53 | 34 | 129 | 150 | 121 | 71 | 156 | 111 | 0.004997044 | 0.03372803 |
| ENSGALG00010004063 | NA | NA | 66 | 144 | 187 | 181 | 213 | 163 | 45 | 27 | 2.15E-06 | 5.16E-05 |
| ENSGALG00010004079 | AP1AR | C4orf16 | 475 | 583 | 588 | 635 | 1068 | 698 | 764 | 1023 | 0.001025103 | 0.009538706 |
| ENSGALG00010004096 | COL12A1 | COL12A1 | 38 | 47 | 50 | 43 | 31 | 17 | 100 | 128 | 0.000329736 | 0.003779527 |
| ENSGALG00010004147 | SPRYD7 | C13orf1 | 1664 | 2411 | 1440 | 1651 | 1468 | 1110 | 1195 | 1095 | 0.005055251 | 0.033956333 |
| ENSGALG00010004159 | NA | NA | 124 | 490 | 1 | 1 | 204 | 190 | 2 | 97 | 4.53E-09 | 2.12E-07 |
| ENSGALG00010004176 | NA | NA | 0 | 6 | 2 | 17 | 19 | 3 | 0 | 0 | 0.003453507 | 0.02514815 |
| ENSGALG00010004188 | BMI1 | PCGF4 | 2691 | 1758 | 1216 | 1221 | 1247 | 1079 | 1784 | 1634 | 0.002333761 | 0.018336539 |
| ENSGALG00010004208 | NA | NA | 135 | 0 | 1 | 1 | 0 | 0 | 246 | 353 | 1.01E-11 | 7.60E-10 |
| ENSGALG00010004229 | PTF1A | PTF1A | 4 | 0 | 97 | 1 | 28 | 1 | 0 | 2 | 0.001547971 | 0.013205559 |
| ENSGALG00010004233 | NHSL1 | NHSL1 | 5164 | 4784 | 2755 | 1641 | 3192 | 2377 | 5510 | 3083 | 0.000244283 | 0.002945492 |
| ENSGALG00010004235 | LOC107051715 | LOC107051715 | 2853 | 3545 | 6767 | 7093 | 4991 | 3531 | 4063 | 4105 | 1.97E-07 | 6.19E-06 |
| ENSGALG00010004240 | LOC421792 | TSTD3 | 304 | 238 | 540 | 480 | 492 | 356 | 271 | 245 | 0.003752864 | 0.026882923 |
| ENSGALG00010004256 | DSC1 | DSC2 | 166 | 45 | 36 | 17 | 40 | 50 | 174 | 76 | 0.006029112 | 0.038902902 |
| ENSGALG00010004281 | VIM | VIM | 37321 | 51125 | 16600 | 23561 | 16288 | 7950 | 20597 | 33340 | 1.95E-08 | 7.63E-07 |
| ENSGALG00010004312 | PDLIM5 | PDLIM5 | 1337 | 2092 | 522 | 988 | 512 | 471 | 1442 | 1406 | 1.46E-07 | 4.80E-06 |
| ENSGALG00010004325 | LTBP2 | LTBP3 | 152 | 301 | 350 | 299 | 168 | 211 | 447 | 786 | 0.001278628 | 0.011354107 |
| ENSGALG00010004328 | LOC121107840 | LOC121107840 | 4 | 4 | 1 | 1 | 53 | 37 | 0 | 6 | 3.39E-12 | 2.73E-10 |
| ENSGALG00010004329 | COQ3 | COQ3 | 156 | 97 | 397 | 280 | 372 | 176 | 186 | 154 | 0.004531325 | 0.031331199 |
| ENSGALG00010004332 | LOC107049387 | LOC107049387 | 115 | 106 | 80 | 142 | 630 | 603 | 0 | 94 | 4.73E-05 | 0.000734391 |
| ENSGALG00010004333 | NA | NA | 2 | 8 | 59 | 95 | 22 | 8 | 12 | 13 | 1.73E-08 | 6.90E-07 |
| ENSGALG00010004340 | SLC4A4 | SLC4A4 | 130 | 112 | 117 | 86 | 188 | 197 | 272 | 221 | 0.003288793 | 0.024233823 |
| ENSGALG00010004341 | LOC101748192 | ZNF629L | 712 | 1267 | 487 | 631 | 179 | 202 | 150 | 252 | 1.54E-10 | 9.16E-09 |
| ENSGALG00010004342 | ZNF438 | ZNF438 | 491 | 629 | 980 | 1079 | 860 | 576 | 541 | 544 | 0.002387718 | 0.018663955 |
| ENSGALG00010004347 | OPN5L1 | COPN5L1 | 21 | 6 | 3 | 3 | 19 | 24 | 5 | 3 | 0.000462057 | 0.004962663 |
| ENSGALG00010004358 | GC | VTDB | 101 | 20 | 23 | 12 | 96 | 145 | 231 | 98 | 9.67E-05 | 0.001345158 |
| ENSGALG00010004366 | UNC5C | UNC5C | 536 | 1224 | 286 | 480 | 325 | 407 | 820 | 577 | 0.00693768 | 0.043420804 |
| ENSGALG00010004367 | TSPAN13 | TM4SF13 | 4916 | 4610 | 2501 | 2825 | 3453 | 2806 | 4156 | 4353 | 1.67E-05 | 0.000302028 |
| ENSGALG00010004375 | NA | NA | 13 | 88 | 3 | 4 | 35 | 45 | 95 | 63 | 6.42E-06 | 0.00013357 |
| ENSGALG00010004401 | NA | NA | 183 | 242 | 180 | 217 | 152 | 115 | 87 | 64 | 0.003417517 | 0.024929524 |
| ENSGALG00010004405 | NA | NA | 4 | 0 | 0 | 2 | 0 | 0 | 34 | 18 | 3.14E-08 | 1.19E-06 |
| ENSGALG00010004429 | LOC112531814 | LOC112531814 | 124 | 45 | 6 | 10 | 4 | 6 | 3 | 10 | 2.77E-10 | 1.57E-08 |
| ENSGALG00010004457 | AGA | AGA | 2373 | 2870 | 2074 | 3613 | 1044 | 1176 | 1661 | 2094 | 0.001005218 | 0.009385029 |
| ENSGALG00010004464 | NA | NA | 0 | 0 | 6 | 2 | 4 | 8 | 13 | 5 | 0.002308721 | 0.018182534 |
| ENSGALG00010004473 | JDP2 | JUNDM2 | 73 | 44 | 99 | 77 | 137 | 115 | 172 | 174 | 0.003976528 | 0.028195054 |
| ENSGALG00010004481 | ZNF704 | ZNF704 | 886 | 824 | 1321 | 1130 | 798 | 687 | 1521 | 1452 | 0.008063249 | 0.049197889 |
| ENSGALG00010004493 | NA | NA | 5 | 12 | 52 | 49 | 32 | 16 | 30 | 36 | 0.001154009 | 0.010458676 |
| ENSGALG00010004514 | NA | NA | 1 | 0 | 19 | 9 | 24 | 10 | 8 | 9 | 0.000125423 | 0.001671513 |
| ENSGALG00010004566 | NA | NA | 5 | 0 | 85 | 79 | 53 | 51 | 65 | 33 | 3.56E-09 | 1.67E-07 |
| ENSGALG00010004570 | LOC107051826 | LOC107051826 | 29259 | 40078 | 1322 | 2419 | 908 | 2077 | 9983 | 8906 | 7.17E-37 | 3.99E-34 |
| ENSGALG00010004631 | NA | NA | 16 | 200 | 81 | 150 | 9 | 22 | 6 | 12 | 6.43E-05 | 0.000945333 |
| ENSGALG00010004648 | LOC421806 | LOC421806 | 3501 | 1998 | 1384 | 2716 | 5527 | 4035 | 3200 | 3657 | 5.18E-06 | 0.000111039 |
| ENSGALG00010004669 | ELMOD1 | ELMOD1 | 409 | 302 | 167 | 128 | 218 | 186 | 456 | 315 | 0.001702069 | 0.014229603 |
| ENSGALG00010004699 | NA | NA | 13417 | 9328 | 5949 | 9195 | 6291 | 4651 | 3696 | 3981 | 1.10E-09 | 5.69E-08 |
| ENSGALG00010004702 | NA | NA | 0 | 1 | 12 | 15 | 5 | 4 | 1 | 1 | 5.87E-05 | 0.000875878 |
| ENSGALG00010004721 | LAMA2 | LAMA2 | 71 | 163 | 216 | 355 | 122 | 85 | 109 | 48 | 0.003694033 | 0.026541132 |
| ENSGALG00010004734 | NA | NA | 67 | 63 | 93 | 136 | 104 | 117 | 13 | 8 | 1.63E-09 | 8.20E-08 |

| | | | | | | | | | | | | |
|---|---|---|---|---|---|---|---|---|---|---|---|---|
| ENSGALG00010004747 | NA | NA | 34 | 48 | 8 | 13 | 11 | 12 | 13 | 22 | 0.004299239 | 0.029999532 |
| ENSGALG00010004749 | NA | NA | 182 | 309 | 53 | 86 | 61 | 69 | 30 | 146 | 0.004084982 | 0.028793008 |
| ENSGALG00010004751 | PTER | PTER | 789 | 369 | 1490 | 1512 | 1101 | 979 | 511 | 527 | 4.20E-06 | 9.28E-05 |
| ENSGALG00010004758 | FLT3 | FLT3 | 0 | 1 | 15 | 7 | 14 | 8 | 7 | 6 | 0.001285025 | 0.011404852 |
| ENSGALG00010004760 | THBS2 | thrombospondin-2 | 38 | 27 | 25 | 17 | 22 | 24 | 79 | 69 | 0.003692052 | 0.026538313 |
| ENSGALG00010004762 | IFT57 | IFT57 | 617 | 263 | 487 | 476 | 1064 | 1092 | 1104 | 1213 | 2.02E-06 | 4.89E-05 |
| ENSGALG00010004778 | NA | NA | 22 | 28 | 3 | 4 | 18 | 6 | 13 | 1 | 0.009205549 | 0.054631794 |
| ENSGALG00010004785 | NKX2-8 | NKX2-8 | 19 | 20 | 80 | 26 | 102 | 73 | 36 | 245 | 0.002469125 | 0.019169112 |
| ENSGALG00010004801 | NA | NA | 51 | 91 | 10 | 19 | 25 | 42 | 51 | 40 | 0.002290254 | 0.018097411 |
| ENSGALG00010004805 | NA | NA | 1 | 0 | 0 | 3 | 92 | 116 | 65 | 95 | 2.47E-25 | 7.25E-23 |
| ENSGALG00010004809 | NA | NA | 7 | 11 | 63 | 31 | 31 | 38 | 15 | 27 | 0.002474195 | 0.019187352 |
| ENSGALG00010004830 | NA | NA | 0 | 1 | 10 | 22 | 13 | 11 | 3 | 4 | 4.72E-05 | 0.000733843 |
| ENSGALG00010004835 | HHLA2 | HHLA2 | 297 | 113 | 155 | 208 | 571 | 661 | 584 | 767 | 1.61E-07 | 5.16E-06 |
| ENSGALG00010004838 | NA | NA | 26 | 0 | 75 | 243 | 165 | 165 | 38 | 72 | 0.00013818 | 0.001816889 |
| ENSGALG00010004855 | ZEB1 | TCF8 | 172 | 364 | 722 | 815 | 458 | 257 | 354 | 329 | 0.000519634 | 0.005472034 |
| ENSGALG00010004866 | SLC25A21 | SLC25A21 | 149 | 196 | 286 | 476 | 222 | 153 | 120 | 180 | 0.007180651 | 0.044610792 |
| ENSGALG00010004871 | C11orf87 | C1H11ORF87 | 1320 | 662 | 1996 | 2165 | 1415 | 1165 | 1031 | 988 | 0.002534745 | 0.019559225 |
| ENSGALG00010004880 | STAP1 | STAP1 | 5 | 18 | 39 | 37 | 36 | 32 | 11 | 13 | 0.002219953 | 0.017649788 |
| ENSGALG00010004881 | MIPOL1 | MIPOL1 | 524 | 574 | 1177 | 1199 | 849 | 835 | 785 | 590 | 0.0003788 | 0.004248667 |
| ENSGALG00010004887 | MYH15 | MYH6 | 121 | 12 | 49 | 74 | 263 | 252 | 204 | 138 | 0.001447736 | 0.012523064 |
| ENSGALG00010004888 | FOXA1 | FOXA1 | 96 | 187 | 252 | 372 | 148 | 141 | 48 | 117 | 0.00320275 | 0.023683344 |
| ENSGALG00010004890 | ACAT1 | ACAT1 | 3027 | 3173 | 4088 | 4799 | 2317 | 1845 | 3117 | 3795 | 0.002276524 | 0.018013857 |
| ENSGALG00010004892 | HEY2 | HEY2 | 393 | 374 | 190 | 114 | 484 | 235 | 497 | 336 | 0.000991241 | 0.009275261 |
| ENSGALG00010004925 | NA | NA | 8 | 4 | 2 | 1 | 5 | 9 | 23 | 10 | 0.007621362 | 0.04693007 |
| ENSGALG00010004929 | ACCSL | ACCSL | 137 | 188 | 74 | 63 | 60 | 31 | 114 | 79 | 0.002602413 | 0.019968562 |
| ENSGALG00010004935 | SVIL | SVIL | 551 | 720 | 1388 | 1372 | 736 | 729 | 1260 | 1082 | 0.000602023 | 0.006178863 |
| ENSGALG00010004952 | PEX1 | PEX1 | 911 | 866 | 737 | 652 | 1104 | 928 | 847 | 767 | 0.007840265 | 0.048065482 |
| ENSGALG00010004965 | KIAA1462 | JCAD | 21 | 34 | 54 | 42 | 73 | 46 | 114 | 79 | 0.004812615 | 0.032868173 |
| ENSGALG00010005001 | GUCA1C | GUCA1C | 804 | 3971 | 4 | 10 | 851 | 1461 | 1544 | 2234 | 9.40E-27 | 3.02E-24 |
| ENSGALG00010005016 | STX11 | syntaxin-11 | 141 | 107 | 487 | 459 | 177 | 149 | 172 | 162 | 3.75E-06 | 8.42E-05 |
| ENSGALG00010005023 | KDELC2 | KDELC2 | 548 | 552 | 1065 | 1002 | 850 | 694 | 697 | 611 | 0.004817529 | 0.032888293 |
| ENSGALG00010005024 | NA | NA | 34 | 323 | 2 | 1 | 0 | 0 | 3 | 5 | 1.56E-20 | 3.26E-18 |
| ENSGALG00010005037 | THNSL1 | THNSL1 | 469 | 694 | 1595 | 1846 | 1100 | 814 | 1058 | 1031 | 4.45E-07 | 1.27E-05 |
| ENSGALG00010005063 | NA | NA | 0 | 0 | 12 | 10 | 7 | 14 | 9 | 21 | 5.86E-05 | 0.000875061 |
| ENSGALG00010005065 | ENKUR | C10orf63 | 119 | 122 | 50 | 51 | 112 | 98 | 76 | 84 | 0.008142463 | 0.049466476 |
| ENSGALG00010005090 | NA | NA | 3355 | 3254 | 26 | 558 | 396 | 549 | 2486 | 2679 | 2.03E-06 | 4.91E-05 |
| ENSGALG00010005092 | ESCO1 | ESCO1 | 746 | 827 | 1175 | 1320 | 929 | 603 | 727 | 559 | 0.007014925 | 0.04379284 |
| ENSGALG00010005114 | PHLDB2 | PHLDB2 | 1229 | 801 | 484 | 537 | 1117 | 886 | 1670 | 1104 | 1.30E-05 | 0.000245319 |
| ENSGALG00010005146 | TGFB3 | TGF-beta-3 | 622 | 672 | 1330 | 1302 | 751 | 540 | 763 | 867 | 0.001551139 | 0.013225836 |
| ENSGALG00010005171 | NA | NA | 19 | 17 | 0 | 1 | 6 | 10 | 19 | 7 | 0.000207623 | 0.002579774 |
| ENSGALG00010005220 | ANGEL1 | ANGEL1 | 1969 | 2277 | 4308 | 5402 | 4627 | 4029 | 4503 | 2839 | 2.27E-07 | 7.01E-06 |
| ENSGALG00010005232 | GATA3 | cGATA-3 | 5 | 6 | 16 | 81 | 5 | 1 | 2 | 3 | 5.28E-07 | 1.48E-05 |
| ENSGALG00010005233 | CD200R1B | CD200R-S1 | 9 | 3 | 22 | 20 | 21 | 17 | 4 | 4 | 0.001371392 | 0.012011896 |
| ENSGALG00010005239 | SFMBT2 | SFMBT2 | 99 | 123 | 128 | 86 | 11 | 8 | 45 | 36 | 1.34E-08 | 5.48E-07 |
| ENSGALG00010005240 | VIP | VIP | 131 | 40 | 12 | 18 | 24 | 11 | 24 | 16 | 6.29E-05 | 0.000928213 |
| ENSGALG00010005241 | C4H4orf54 | C4H4orf54 | 2296 | 3040 | 433 | 507 | 1959 | 1422 | 3232 | 1617 | 3.76E-16 | 5.11E-14 |
| ENSGALG00010005245 | KIAA1217 | KIAA1217 | 665 | 1186 | 398 | 480 | 712 | 514 | 792 | 746 | 0.002188453 | 0.017474163 |
| ENSGALG00010005248 | TMEM200A | TMEM200A | 2805 | 3427 | 1664 | 2121 | 2297 | 1384 | 2431 | 1874 | 0.003536336 | 0.025628471 |
| ENSGALG00010005249 | PDGFD | PDGFD | 95 | 85 | 783 | 690 | 538 | 199 | 189 | 147 | 7.89E-13 | 6.83E-11 |
| ENSGALG00010005255 | BZW2 | 5MP1 | 19955 | 16680 | 10099 | 13017 | 15136 | 12657 | 10341 | 10552 | 1.81E-05 | 0.000323685 |
| ENSGALG00010005257 | SAMD3 | SAMD3 | 1255 | 1471 | 267 | 778 | 1120 | 845 | 524 | 353 | 1.92E-06 | 4.68E-05 |
| ENSGALG00010005260 | ANXA5 | ANX5 | 363 | 314 | 742 | 879 | 471 | 289 | 648 | 761 | 0.000401653 | 0.004436497 |
| ENSGALG00010005281 | TMED8 | TMED8 | 908 | 1284 | 2050 | 2292 | 1882 | 1651 | 1728 | 1629 | 0.000264264 | 0.00315681 |
| ENSGALG00010005282 | FLT1 | FLT1 | 70 | 40 | 18 | 3 | 6 | 13 | 123 | 46 | 5.37E-05 | 0.000815865 |
| ENSGALG00010005300 | QRFPR | GPR103 | 24 | 28 | 7 | 7 | 7 | 15 | 66 | 37 | 0.000108307 | 0.001475168 |
| ENSGALG00010005339 | LOC121106475 | LOC121106475 | 21 | 29 | 7 | 6 | 5 | 6 | 10 | 11 | 0.007103743 | 0.044248139 |
| ENSGALG00010005340 | LOC101747944 | LOC101747944 | 23 | 98 | 13 | 5 | 16 | 13 | 51 | 76 | 0.000128969 | 0.001710582 |
| ENSGALG00010005345 | NOX3 | NOX3 | 1 | 2 | 15 | 11 | 12 | 13 | 5 | 2 | 0.001037389 | 0.009610223 |
| ENSGALG00010005354 | M1AP | C2orf65 | 64 | 98 | 274 | 263 | 396 | 210 | 94 | 88 | 1.90E-08 | 7.48E-07 |
| ENSGALG00010005355 | FDX1 | FDX1 | 1016 | 1037 | 1589 | 1873 | 1382 | 872 | 915 | 946 | 0.002619481 | 0.020062676 |
| ENSGALG00010005359 | MEIS2 | Meis2a.1 | 32 | 89 | 1034 | 949 | 363 | 356 | 107 | 468 | 2.42E-09 | 1.17E-07 |
| ENSGALG00010005360 | CAMK1D | CAMK1D | 182 | 459 | 374 | 340 | 27 | 24 | 41 | 35 | 4.81E-17 | 7.05E-15 |
| ENSGALG00010005363 | RDX | RDX | 2752 | 3014 | 5120 | 4839 | 4365 | 3515 | 4014 | 4561 | 0.000885601 | 0.008452402 |
| ENSGALG00010005368 | CCDC3 | CCDC3 | 114 | 391 | 37 | 32 | 22 | 5 | 197 | 207 | 1.40E-11 | 1.02E-09 |
| ENSGALG00010005381 | FGF9 | FGF9 | 5 | 19 | 45 | 54 | 48 | 20 | 13 | 12 | 0.001323567 | 0.011678695 |
| ENSGALG00010005402 | ABCC4 | ABCC4 | 3041 | 4091 | 494 | 591 | 2461 | 4614 | 6555 | 3957 | 6.25E-18 | 1.02E-15 |
| ENSGALG00010005413 | GAD2 | GAD65 | 52 | 78 | 89 | 119 | 27 | 38 | 294 | 426 | 1.33E-11 | 9.79E-10 |
| ENSGALG00010005437 | PDE7B | PDE7B | 134 | 259 | 447 | 554 | 404 | 298 | 352 | 272 | 0.004915724 | 0.033382261 |
| ENSGALG00010005457 | SEMA3D | COLL2 | 138 | 202 | 32 | 57 | 100 | 69 | 190 | 125 | 0.000218826 | 0.002692435 |
| ENSGALG00010005472 | PFKP | PFKP | 5289 | 5355 | 6729 | 9159 | 4731 | 3500 | 4875 | 5552 | 0.00480608 | 0.032836965 |
| ENSGALG00010005493 | KLF6 | KLF6 | 786 | 759 | 494 | 583 | 470 | 222 | 194 | 270 | 9.67E-06 | 0.000190581 |
| ENSGALG00010005494 | C8orf22 | C2H8ORF22 | 8028 | 5071 | 1984 | 2535 | 5325 | 3347 | 4066 | 1840 | 7.31E-07 | 1.97E-05 |
| ENSGALG00010005509 | COCH | COCH | 210 | 1725 | 1382 | 2474 | 488 | 500 | 140 | 329 | 7.29E-05 | 0.00105769 |
| ENSGALG00010005526 | STRN3 | STRN3 | 2579 | 4100 | 4270 | 5573 | 3875 | 2622 | 2853 | 2748 | 0.005274654 | 0.035189627 |
| ENSGALG00010005530 | NA | NA | 252 | 331 | 2415 | 2385 | 2057 | 1070 | 851 | 566 | 5.16E-25 | 1.46E-22 |
| ENSGALG00010005544 | AQP4 | aquaporin-4 | 25 | 33 | 3 | 3 | 7 | 13 | 17 | 18 | 0.000348275 | 0.003948656 |
| ENSGALG00010005547 | LRP11 | LRP11 | 709 | 471 | 473 | 401 | 962 | 724 | 896 | 589 | 0.001869774 | 0.01530248 |
| ENSGALG00010005553 | PPP1R14C | PPP1R14C | 2712 | 4321 | 1074 | 1169 | 2362 | 1952 | 2234 | 2570 | 2.69E-11 | 1.87E-09 |
| ENSGALG00010005555 | PRSS12 | PRSS12 | 736 | 328 | 179 | 518 | 1422 | 800 | 684 | 233 | 0.0007594 | 0.007487367 |
| ENSGALG00010005569 | NEUROG2 | NGN2/ATH4A | 18 | 32 | 27 | 49 | 56 | 58 | 18 | 9 | 0.000931228 | 0.008807401 |
| ENSGALG00010005583 | ALDOB | ALDB | 0 | 1 | 19 | 2 | 11 | 2 | 21 | 18 | 0.002481626 | 0.019227135 |
| ENSGALG00010005585 | LOC431250 | LOC431250 | 9 | 21 | 22 | 23 | 27 | 10 | 67 | 56 | 0.002180262 | 0.017433748 |
| ENSGALG00010005610 | MTHFD1L | MTHFD1L | 2541 | 2889 | 2522 | 3145 | 2236 | 2085 | 4327 | 3905 | 0.003337421 | 0.024451984 |
| ENSGALG00010005635 | MOS | c-mos | 31 | 48 | 15 | 8 | 63 | 51 | 14 | 8 | 1.93E-06 | 4.72E-05 |
| ENSGALG00010005652 | NA | NA | 22 | 3 | 1 | 4 | 22 | 17 | 7 | 3 | 0.005619833 | 0.036975844 |
| ENSGALG00010005702 | NA | NA | 15 | 28 | 19 | 43 | 18 | 9 | 4 | 3 | 0.001419411 | 0.012330698 |
| ENSGALG00010005705 | ECHDC3 | ECHDC3 | 813 | 784 | 1200 | 1305 | 776 | 583 | 694 | 654 | 0.008491509 | 0.051194119 |
| ENSGALG00010005718 | NA | NA | 5 | 15 | 35 | 46 | 44 | 46 | 23 | 23 | 0.000767528 | 0.007536389 |
| ENSGALG00010005720 | COL25A1 | COL25A1 | 57 | 40 | 3 | 5 | 18 | 64 | 102 | 35 | 3.65E-05 | 0.000589749 |
| ENSGALG00010005738 | ELAVL2 | ELAVL2 | 2778 | 2237 | 1803 | 1249 | 2980 | 1483 | 1632 | 1045 | 0.001148017 | 0.010410018 |
| ENSGALG00010005754 | NA | NA | 99 | 45 | 109 | 54 | 30 | 25 | 161 | 163 | 0.001114959 | 0.010204195 |
| ENSGALG00010005791 | NA | NA | 78 | 130 | 26 | 41 | 64 | 1880 | 40 | 294 | 6.77E-07 | 1.84E-05 |
| ENSGALG00010005792 | ADGRG6 | GPR126 | 109 | 47 | 22 | 31 | 109 | 62 | 56 | 43 | 0.003808007 | 0.027207957 |
| ENSGALG00010005799 | UFSP2 | C4orf20 | 198 | 320 | 542 | 651 | 195 | 104 | 215 | 212 | 8.02E-06 | 0.000161956 |
| ENSGALG00010005803 | NA | NA | 1058 | 1165 | 0 | 31 | 5 | 5 | 1256 | 1074 | 4.78E-39 | 2.85E-36 |
| ENSGALG00010005819 | NA | NA | 1092 | 1839 | 481 | 848 | 1066 | 1304 | 916 | 730 | 0.000199415 | 0.002488889 |
| ENSGALG00010005837 | LOC107052005 | LOC107052005 | 310 | 331 | 64 | 75 | 64 | 66 | 70 | 68 | 3.18E-10 | 1.78E-08 |
| ENSGALG00010005865 | NA | NA | 370 | 550 | 177 | 238 | 107 | 128 | 351 | 359 | 4.60E-05 | 0.00071861 |
| ENSGALG00010005903 | PRR18 | PRR18 | 148 | 92 | 49 | 64 | 96 | 101 | 54 | 27 | 0.002871244 | 0.021634008 |
| ENSGALG00010005905 | CITED2 | CITED2 | 1565 | 1726 | 763 | 1268 | 824 | 678 | 620 | 601 | 5.98E-07 | 1.66E-05 |
| ENSGALG00010005911 | TBXT | CH-T | 297 | 166 | 91 | 82 | 142 | 93 | 137 | 85 | 0.009869416 | 0.057711848 |
| ENSGALG00010005949 | SLITRK1 | SLITRK1 | 23 | 33 | 8 | 16 | 7 | 7 | 4 | 7 | 0.004434014 | 0.030709174 |
| ENSGALG00010005970 | SHISA2 | SHISA | 4982 | 11599 | 12737 | 14384 | 9525 | 4619 | 3281 | 5441 | 5.53E-05 | 0.000834499 |
| ENSGALG00010005982 | LOC121107804 | LOC121107804 | 21 | 9 | 6 | 11 | 1216 | 1337 | 4 | 10 | 5.72E-93 | 2.39E-89 |
| ENSGALG00010005987 | LOC121107845 | LOC121107845 | 14 | 7 | 4 | 34 | 589 | 458 | 2 | 1 | 2.72E-36 | 1.46E-33 |
| ENSGALG00010005995 | NA | NA | 25 | 23 | 12 | 4 | 0 | 1 | 4 | 2 | 9.28E-06 | 0.000183849 |
| ENSGALG00010005999 | PDE10A | PDE10A | 566 | 821 | 268 | 380 | 281 | 233 | 601 | 436 | 0.000552191 | 0.005774859 |
| ENSGALG00010006009 | MALRD1 | C10ORF112 | 2 | 17 | 32 | 245 | 8 | 9 | 5 | 8 | 2.85E-08 | 1.09E-06 |
| ENSGALG00010006016 | MTMR6 | MTMR6 | 1314 | 2228 | 2387 | 3073 | 2797 | 2038 | 1621 | 1562 | 0.000563883 | 0.005860458 |
| ENSGALG00010006023 | NA | NA | 13 | 18 | 0 | 1 | 0 | 0 | 0 | 0 | 1.01E-08 | 4.27E-07 |
| ENSGALG00010006055 | LOC107056277 | LOC107056277 | 15 | 39 | 49 | 25 | 10 | 5 | 39 | 51 | 0.008800954 | 0.052717402 |
| ENSGALG00010006072 | LOC107056274 | LOC107056274 | 2476 | 3717 | 1946 | 2092 | 1962 | 1425 | 2143 | 2505 | 0.007866105 | 0.048170883 |
| ENSGALG00010006075 | SOX21 | CH31 | 6802 | 6048 | 4206 | 4999 | 5474 | 3256 | 3004 | 4058 | 0.000446419 | 0.004816495 |
| ENSGALG00010006076 | PODXL | MEP21 | 167 | 833 | 339 | 415 | 683 | 632 | 1517 | 1305 | 0.00019051 | 0.002397444 |
| ENSGALG00010006079 | LOC107055200 | LOC107055200 | 3858 | 2067 | 1150 | 2089 | 781 | 1951 | 4454 | 4936 | 0.000566721 | 0.005878988 |
| ENSGALG00010006080 | LOC107056280 | LOC107056280 | 774 | 884 | 255 | 243 | 383 | 248 | 799 | 979 | 3.08E-11 | 2.11E-09 |
| ENSGALG00010006086 | LOC100857117 | LOC100857117 | 13031 | 8839 | 1474 | 2910 | 1034 | 1695 | 5878 | 6718 | 1.31E-17 | 2.06E-15 |
| ENSGALG00010006087 | LOC107051465 | LOC107051465 | 18543 | 31674 | 1982 | 3862 | 2825 | 4786 | 8592 | 9466 | 1.02E-20 | 2.26E-18 |
| ENSGALG00010006088 | NA | NA | 756 | 363 | 62 | 130 | 33 | 111 | 465 | 415 | 3.18E-07 | 9.48E-06 |
| ENSGALG00010006131 | LOC101749299 | LOC101749299 | 547 | 428 | 298 | 162 | 257 | 166 | 483 | 529 | 0.00216436 | 0.017323169 |
| ENSGALG00010006146 | CCDC142 | CCDC142 | 970 | 1100 | 1288 | 1478 | 805 | 547 | 735 | 843 | 0.006631379 | 0.041899282 |
| ENSGALG00010006179 | NA | NA | 1328 | 372 | 215 | 336 | 108 | 255 | 518 | 564 | 0.001272416 | 0.011323015 |
| ENSGALG00010006207 | LOC107056270 | LOC107056270 | 1055 | 897 | 729 | 770 | 1269 | 676 | 668 | 553 | 0.00501082 | 0.033780085 |
| ENSGALG00010006213 | NA | NA | 50 | 118 | 39 | 57 | 142 | 111 | 164 | 113 | 0.003520847 | 0.025549455 |
| ENSGALG00010006214 | NA | NA | 3316 | 3710 | 651 | 888 | 192 | 736 | 2488 | 1721 | 3.76E-08 | 1.39E-06 |
| ENSGALG00010006253 | NA | NA | 5 | 37 | 29 | 51 | 5 | 1 | 27 | 23 | 0.002617462 | 0.020056408 |
| ENSGALG00010006271 | PAX4 | PAX4 | 1138 | 860 | 132 | 208 | 382 | 252 | 1125 | 748 | 5.49E-15 | 6.37E-13 |
| ENSGALG00010006276 | NA | NA | 5 | 4 | 20 | 28 | 30 | 4 | 5 | 0 | 0.002547184 | 0.019634936 |
| ENSGALG00010006277 | GNG10 | GNG10 | 12711 | 8251 | 4155 | 2373 | 8227 | 4352 | 8231 | 4919 | 5.99E-06 | 0.000125631 |
| ENSGALG00010006280 | VRTN | C14orf115 | 3988 | 3371 | 1354 | 1659 | 1752 | 2024 | 3671 | 3293 | 3.29E-08 | 1.23E-06 |
| ENSGALG00010006281 | TRMT11 | MDS024 | 2010 | 2225 | 2858 | 3103 | 1634 | 1166 | 2690 | 2273 | 0.002944532 | 0.022096554 |

| | | | | | | | | | | | | |
|---|---|---|---|---|---|---|---|---|---|---|---|---|
| ENSGALG00010006285 | FZD8 | FZD8 | 2966 | 3171 | 1493 | 1660 | 1643 | 1507 | 2657 | 2911 | 1.53E-06 | 3.86E-05 |
| ENSGALG00010006293 | SLC22A3 | SLC22A3 | 69 | 34 | 2 | 14 | 18 | 16 | 36 | 37 | 0.000959585 | 0.009027376 |
| ENSGALG00010006294 | NA | NA | 0 | 2 | 12 | 13 | 0 | 3 | 7 | 14 | 0.002065859 | 0.016678566 |
| ENSGALG00010006300 | GCC1 | GCC1 | 1099 | 896 | 625 | 576 | 538 | 457 | 894 | 791 | 0.009427602 | 0.055613867 |
| ENSGALG00010006304 | NA | NA | 3 | 4 | 16 | 27 | 27 | 9 | 17 | 23 | 0.006194592 | 0.039740507 |
| ENSGALG00010006359 | SGK1 | SGK | 940 | 770 | 174 | 130 | 537 | 268 | 591 | 608 | 8.39E-11 | 5.25E-09 |
| ENSGALG00010006361 | SERTM1 | SERTM1 | 385 | 370 | 176 | 150 | 275 | 188 | 419 | 426 | 0.000446385 | 0.004816495 |
| ENSGALG00010006366 | CCNA1 | CCNA1 | 18 | 22 | 105 | 56 | 35 | 13 | 23 | 40 | 0.004856082 | 0.033043501 |
| ENSGALG00010006368 | NA | NA | 1 | 0 | 6 | 6 | 10 | 11 | 9 | 6 | 0.001644673 | 0.013870136 |
| ENSGALG00010006374 | OLIG3 | OLIG3 | 49 | 81 | 27 | 11 | 45 | 28 | 69 | 105 | 0.001805204 | 0.014912788 |
| ENSGALG00010006404 | SMAD1 | SMAD1 | 709 | 441 | 1242 | 1228 | 1020 | 613 | 615 | 542 | 0.0006439 | 0.006553508 |
| ENSGALG00010006410 | ADM2 | ADM2 | 70 | 47 | 54 | 22 | 25 | 143 | 223 | 394 | 0.00013372 | 0.0017638 |
| ENSGALG00010006492 | GAP43 | GAP-43 | 401 | 379 | 74 | 79 | 216 | 128 | 305 | 118 | 9.45E-06 | 0.000186953 |
| ENSGALG00010006504 | NA | NA | 5 | 12 | 2 | 0 | 9 | 10 | 20 | 21 | 0.000253348 | 0.003041628 |
| ENSGALG00010006516 | NBEA | NBEA | 83 | 81 | 225 | 304 | 164 | 115 | 231 | 195 | 0.001241719 | 0.011108711 |
| ENSGALG00010006522 | NA | NA | 175 | 159 | 634 | 522 | 147 | 81 | 192 | 266 | 1.51E-07 | 4.91E-06 |
| ENSGALG00010006523 | NA | NA | 3 | 5 | 13 | 5 | 26 | 11 | 18 | 24 | 0.008138692 | 0.049466476 |
| ENSGALG00010006527 | NA | NA | 4 | 1 | 14 | 23 | 22 | 17 | 4 | 8 | 0.000427286 | 0.004655675 |
| ENSGALG00010006530 | LOC101749214 | LOC101749214 | 18 | 7 | 6 | 8 | 26 | 27 | 24 | 22 | 0.008378625 | 0.050659761 |
| ENSGALG00010006546 | SMOC2 | SMOC2 | 293 | 132 | 13 | 13 | 98 | 68 | 229 | 102 | 1.68E-09 | 8.39E-08 |
| ENSGALG00010006580 | ALDH6A1 | ALDH6A1 | 324 | 659 | 594 | 701 | 198 | 185 | 341 | 459 | 0.002605594 | 0.01998379 |
| ENSGALG00010006585 | TTC29 | TTC29 | 22 | 25 | 118 | 114 | 137 | 60 | 51 | 47 | 1.70E-05 | 0.000306084 |
| ENSGALG00010006602 | EDNRA | EDNRA | 204 | 248 | 572 | 581 | 596 | 341 | 272 | 404 | 0.000631229 | 0.006440233 |
| ENSGALG00010006621 | DACT2 | DACT2 | 268 | 234 | 145 | 139 | 120 | 124 | 377 | 352 | 0.000363965 | 0.004104309 |
| ENSGALG00010006623 | LOC107052071 | LOC107052071 | 45 | 100 | 1 | 0 | 0 | 4 | 19 | 8 | 2.09E-13 | 1.91E-11 |
| ENSGALG00010006641 | PLA2G10L | PLA2G10L | 11 | 37 | 0 | 1 | 5 | 8 | 59 | 71 | 3.00E-11 | 2.07E-09 |
| ENSGALG00010006653 | CFL2 | cofilin-2 | 2854 | 3007 | 5010 | 5596 | 4240 | 2827 | 2905 | 3482 | 0.000131756 | 0.001742019 |
| ENSGALG00010006673 | BAZ1A | BAZ1A | 220 | 263 | 928 | 767 | 619 | 346 | 308 | 223 | 5.11E-08 | 1.85E-06 |
| ENSGALG00010006704 | NA | NA | 65 | 68 | 12 | 13 | 62 | 29 | 40 | 33 | 0.00015304 | 0.001985721 |
| ENSGALG00010006719 | NA | NA | 5 | 26 | 47 | 30 | 66 | 52 | 66 | 78 | 0.002142821 | 0.017175453 |
| ENSGALG00010006728 | N4BP2L1 | N4BP2L1 | 584 | 804 | 233 | 444 | 317 | 314 | 636 | 621 | 0.002761064 | 0.020974047 |
| ENSGALG00010006730 | HAPLN1 | CRTL1 | 1 | 28 | 204 | 199 | 11 | 8 | 17 | 6 | 7.81E-11 | 4.98E-09 |
| ENSGALG00010006738 | NA | NA | 6 | 4 | 50 | 16 | 9 | 1 | 6 | 8 | 0.0016916 | 0.014163335 |
| ENSGALG00010006752 | ZAR1L | ZAR1 | 2406 | 1272 | 308 | 423 | 352 | 398 | 1219 | 1155 | 6.34E-14 | 6.38E-12 |
| ENSGALG00010006785 | NA | NA | 4 | 3 | 14 | 32 | 22 | 4 | 1 | 0 | 3.14E-05 | 0.000519495 |
| ENSGALG00010006815 | NA | NA | 3 | 3 | 17 | 28 | 34 | 16 | 12 | 9 | 0.000213351 | 0.002637218 |
| ENSGALG00010006835 | FRY | FRY | 40 | 36 | 165 | 133 | 129 | 75 | 77 | 66 | 0.000400349 | 0.004427765 |
| ENSGALG00010006840 | GINS2 | GINS2 | 677 | 710 | 1270 | 1262 | 1227 | 843 | 1049 | 1162 | 0.003605894 | 0.026076025 |
| ENSGALG00010006854 | FOXG1 | CBF-1 | 16 | 13 | 86 | 55 | 36 | 63 | 23 | 49 | 0.003971209 | 0.028169289 |
| ENSGALG00010006865 | EFCAB2 | EFCAB2 | 6 | 11 | 54 | 48 | 67 | 33 | 4 | 10 | 7.33E-08 | 2.59E-06 |
| ENSGALG00010006884 | NA | NA | 3 | 8 | 18 | 25 | 3 | 5 | 1 | 3 | 0.000849008 | 0.008196779 |
| ENSGALG00010006885 | RHOU | RHOU | 3843 | 1894 | 706 | 1102 | 1091 | 1067 | 1959 | 1155 | 7.96E-06 | 0.00016088 |
| ENSGALG00010006886 | NA | NA | 14 | 20 | 37 | 62 | 16 | 19 | 9 | 15 | 0.005834762 | 0.038000992 |
| ENSGALG00010006902 | NA | NA | 162 | 680 | 2 | 0 | 148 | 281 | 250 | 459 | 1.75E-17 | 2.68E-15 |
| ENSGALG00010006909 | NA | NA | 13 | 38 | 0 | 0 | 0 | 2 | 3 | 0 | 4.86E-08 | 1.78E-06 |
| ENSGALG00010006910 | NA | NA | 4 | 23 | 0 | 0 | 14 | 14 | 55 | 22 | 3.09E-07 | 9.24E-06 |
| ENSGALG00010006916 | NA | NA | 53 | 167 | 0 | 0 | 51 | 105 | 218 | 169 | 2.34E-16 | 3.24E-14 |
| ENSGALG00010006926 | ID4 | ID4 | 1340 | 1391 | 690 | 638 | 978 | 513 | 758 | 1060 | 0.001354627 | 0.011902482 |
| ENSGALG00010006933 | NA | NA | 332 | 841 | 130 | 211 | 166 | 142 | 154 | 170 | 2.58E-06 | 6.03E-05 |
| ENSGALG00010006944 | MAB21L2 | MAB21L2 | 103 | 147 | 112 | 202 | 584 | 147 | 61 | 75 | 6.18E-06 | 0.000129198 |
| ENSGALG00010006957 | PALM2AKAP2 | AKAP2 | 1225 | 815 | 524 | 603 | 598 | 245 | 769 | 508 | 0.008490611 | 0.051194119 |
| ENSGALG00010006958 | KIF26B | KIF26B | 567 | 942 | 593 | 1095 | 577 | 858 | 278 | 102 | 1.58E-05 | 0.00028924 |
| ENSGALG00010006966 | PGBD5 | PGBD5 | 14 | 10 | 44 | 75 | 8 | 4 | 30 | 25 | 6.77E-06 | 0.000140117 |
| ENSGALG00010006983 | GTPBP2 | GTPBP2 | 513 | 586 | 518 | 586 | 720 | 508 | 312 | 247 | 0.000159839 | 0.002064315 |
| ENSGALG00010006999 | NA | NA | 14 | 13 | 85 | 93 | 17 | 24 | 12 | 21 | 1.63E-06 | 4.06E-05 |
| ENSGALG00010007014 | LOC101750035 | LOC101750035 | 24 | 23 | 3 | 5 | 29 | 20 | 34 | 24 | 0.000170506 | 0.002186873 |
| ENSGALG00010007018 | NA | NA | 1 | 2 | 7 | 24 | 36 | 16 | 19 | 18 | 0.00010057 | 0.001384447 |
| ENSGALG00010007027 | NA | NA | 11 | 28 | 37 | 74 | 64 | 78 | 53 | 58 | 0.004737554 | 0.032488308 |
| ENSGALG00010007049 | NA | NA | 9102 | 8892 | 1775 | 3194 | 1248 | 2725 | 7883 | 7776 | 3.55E-10 | 1.98E-08 |
| ENSGALG00010007054 | NA | NA | 2992 | 3003 | 1682 | 1907 | 1960 | 1532 | 2143 | 1960 | 0.000594178 | 0.006114471 |
| ENSGALG00010007069 | NA | NA | 13 | 27 | 369 | 683 | 189 | 57 | 42 | 82 | 4.99E-15 | 5.83E-13 |
| ENSGALG00010007088 | NA | NA | 18 | 11 | 61 | 81 | 90 | 68 | 25 | 4 | 4.92E-06 | 0.000106409 |
| ENSGALG00010007092 | DYNC1LI2 | DNCLI2 | 378 | 432 | 894 | 806 | 671 | 442 | 550 | 505 | 0.005742154 | 0.03757356 |
| ENSGALG00010007103 | NA | NA | 17 | 43 | 5 | 3 | 2 | 2 | 6 | 2 | 1.01E-05 | 0.000198573 |
| ENSGALG00010007106 | SLC7A1 | CAT-1 | 4457 | 4093 | 5184 | 4742 | 5250 | 4752 | 8245 | 7787 | 3.76E-05 | 0.000605568 |
| ENSGALG00010007115 | CD69L | CD69 | 865 | 421 | 131 | 170 | 193 | 142 | 134 | 182 | 1.35E-08 | 5.51E-07 |
| ENSGALG00010007130 | IRF2 | IRF-2 | 16 | 40 | 115 | 129 | 94 | 45 | 24 | 33 | 6.37E-05 | 0.00093818 |
| ENSGALG00010007135 | TK2 | TK2 | 4561 | 3819 | 7050 | 8790 | 6289 | 5426 | 5401 | 5786 | 0.000222338 | 0.002726127 |
| ENSGALG00010007137 | LOC121113333 | LOC121113333 | 367 | 300 | 83 | 78 | 198 | 129 | 120 | 200 | 1.98E-05 | 0.000349158 |
| ENSGALG00010007160 | LOC121113327 | LOC121113327 | 428 | 356 | 134 | 178 | 99 | 102 | 101 | 236 | 0.000358527 | 0.004051191 |
| ENSGALG00010007161 | NA | NA | 177 | 661 | 1 | 10 | 103 | 293 | 447 | 367 | 3.82E-12 | 3.06E-10 |
| ENSGALG00010007165 | LOC101747821 | LOC101747821 | 440 | 1720 | 3 | 8 | 36 | 12 | 23 | 18 | 2.42E-41 | 1.69E-38 |
| ENSGALG00010007167 | NA | NA | 250 | 591 | 3 | 0 | 94 | 212 | 292 | 315 | 1.09E-20 | 2.40E-18 |
| ENSGALG00010007171 | NA | NA | 701 | 610 | 0 | 0 | 345 | 636 | 832 | 1038 | 3.96E-45 | 3.15E-42 |
| ENSGALG00010007175 | FOXC2 | FOXC2 | 14 | 27 | 20 | 54 | 8 | 5 | 10 | 2 | 0.002995106 | 0.022405648 |
| ENSGALG00010007181 | GJB2 | GJB2 | 90 | 88 | 19 | 19 | 64 | 57 | 63 | 27 | 0.00040476 | 0.004464918 |
| ENSGALG00010007188 | MAB21L1 | MAB21L1 | 26 | 26 | 181 | 540 | 71 | 25 | 14 | 25 | 9.68E-15 | 1.09E-12 |
| ENSGALG00010007190 | NA | NA | 9 | 10 | 60 | 91 | 44 | 47 | 52 | 15 | 0.000204881 | 0.002549491 |
| ENSGALG00010007204 | LANCL2 | LANCL2 | 2937 | 3148 | 4272 | 5554 | 3241 | 2694 | 2604 | 3079 | 0.001886243 | 0.015414619 |
| ENSGALG00010007222 | NA | NA | 0 | 5 | 23 | 24 | 11 | 18 | 14 | 31 | 0.002713758 | 0.020718288 |
| ENSGALG00010007230 | NA | NA | 47 | 21 | 2 | 1 | 20 | 10 | 32 | 21 | 7.21E-06 | 0.000148059 |
| ENSGALG00010007267 | NA | NA | 9 | 5 | 33 | 13 | 2 | 9 | 46 | 107 | 2.28E-06 | 5.41E-05 |
| ENSGALG00010007271 | CADM2 | IGSF4D | 38 | 63 | 100 | 147 | 100 | 75 | 54 | 46 | 0.009956592 | 0.058118165 |
| ENSGALG00010007272 | NA | NA | 18 | 61 | 14 | 8 | 13 | 0 | 3 | 3 | 0.000680868 | 0.006858747 |
| ENSGALG00010007275 | DPY30 | DPY30 | 3307 | 2705 | 1451 | 1875 | 2057 | 1779 | 2973 | 1935 | 0.003118741 | 0.0231827 |
| ENSGALG00010007287 | LOC124418405 | LOC124418405 | 16551 | 1690 | 376 | 766 | 940 | 1321 | 1853 | 1781 | 8.12E-10 | 4.30E-08 |
| ENSGALG00010007302 | CDH11 | cadherin-11 | 613 | 1409 | 452 | 392 | 394 | 372 | 1504 | 1957 | 1.25E-10 | 7.54E-09 |
| ENSGALG00010007369 | NA | NA | 310 | 119 | 49 | 39 | 161 | 313 | 114 | 77 | 1.26E-05 | 0.000239999 |
| ENSGALG00010007381 | EGFR | c-erbB | 409 | 370 | 496 | 619 | 1173 | 791 | 1158 | 756 | 2.00E-07 | 6.28E-06 |
| ENSGALG00010007386 | NA | NA | 1 | 3 | 19 | 61 | 33 | 25 | 26 | 34 | 7.03E-06 | 0.000144787 |
| ENSGALG00010007403 | SDR42E1 | HSPC105 | 108 | 93 | 262 | 233 | 258 | 197 | 142 | 111 | 0.000819134 | 0.007958937 |
| ENSGALG00010007404 | FOLH1 | FOLH1 | 116 | 92 | 237 | 249 | 247 | 178 | 236 | 214 | 0.009121558 | 0.054287565 |
| ENSGALG00010007430 | MEF2C | MEF2C | 2 | 16 | 46 | 17 | 53 | 15 | 8 | 4 | 0.004046523 | 0.028570131 |
| ENSGALG00010007455 | TRPC6 | TRPC6 | 9 | 5 | 16 | 24 | 3 | 1 | 11 | 15 | 0.005062454 | 0.033991054 |
| ENSGALG00010007467 | CHODL | CHODL | 269 | 292 | 771 | 905 | 702 | 166 | 192 | 176 | 1.34E-05 | 0.000251578 |
| ENSGALG00010007472 | NA | NA | 6 | 18 | 90 | 109 | 13 | 5 | 7 | 6 | 1.07E-11 | 8.01E-10 |
| ENSGALG00010007485 | CTSC | CTSC | 4046 | 2718 | 3214 | 4697 | 2150 | 1402 | 1941 | 2316 | 0.00043182 | 0.004689128 |
| ENSGALG00010007514 | VIT | AKH | 10 | 24 | 54 | 59 | 105 | 34 | 22 | 8 | 0.000172255 | 0.002204968 |
| ENSGALG00010007529 | CDH13 | cadherin-13 | 1374 | 1291 | 1679 | 4502 | 1642 | 1461 | 889 | 773 | 1.29E-06 | 3.28E-05 |
| ENSGALG00010007536 | GABBR2 | GABBR2 | 167 | 135 | 70 | 72 | 50 | 32 | 166 | 106 | 0.000977491 | 0.009167129 |
| ENSGALG00010007547 | LOC100858777 | LOC100858777 | 63 | 64 | 121 | 78 | 213 | 124 | 76 | 95 | 0.002357485 | 0.018470835 |
| ENSGALG00010007570 | NA | NA | 8 | 17 | 62 | 24 | 65 | 25 | 21 | 5 | 0.006394321 | 0.040786984 |
| ENSGALG00010007576 | LOC100859691 | LOC100859691 | 2 | 7 | 63 | 42 | 69 | 9 | 7 | 6 | 7.28E-06 | 0.000149158 |
| ENSGALG00010007583 | NA | NA | 2 | 0 | 17 | 8 | 16 | 9 | 9 | 2 | 0.005190573 | 0.034711827 |
| ENSGALG00010007595 | ATP2C2 | ATP2C2 | 14 | 8 | 42 | 44 | 45 | 34 | 36 | 48 | 0.003211726 | 0.023739212 |
| ENSGALG00010007596 | LOC121109412 | LOC121109412 | 110 | 7 | 13 | 1 | 5 | 4 | 8 | 8 | 0.000600928 | 0.006176327 |
| ENSGALG00010007601 | TMEM154 | TMEM154 | 7 | 12 | 8 | 4 | 10 | 14 | 71 | 52 | 1.31E-07 | 4.37E-06 |
| ENSGALG00010007655 | SLC9A3 | SLC9A3 | 3 | 0 | 2 | 6 | 21 | 17 | 7 | 1 | 0.000435583 | 0.00472079 |
| ENSGALG00010007672 | NA | NA | 183 | 311 | 349 | 186 | 22 | 25 | 65 | 45 | 7.12E-11 | 4.58E-09 |
| ENSGALG00010007695 | RASGRP3 | RASGRP3 | 19 | 35 | 31 | 36 | 18 | 30 | 114 | 89 | 0.000434074 | 0.00470749 |
| ENSGALG00010007698 | ADAMTS18 | ADAMTS18 | 156 | 207 | 267 | 633 | 280 | 287 | 216 | 171 | 0.007676033 | 0.047214524 |
| ENSGALG00010007724 | RBM11 | RBM11 | 7 | 1 | 52 | 38 | 36 | 3 | 13 | 9 | 0.002187246 | 0.017472873 |
| ENSGALG00010007767 | VAT1L | VAT1L | 1572 | 909 | 871 | 750 | 1234 | 1041 | 2602 | 1874 | 3.32E-05 | 0.000545076 |
| ENSGALG00010007781 | MBP | MBP | 585 | 579 | 1381 | 2968 | 1741 | 873 | 452 | 491 | 1.92E-12 | 1.61E-10 |
| ENSGALG00010007783 | NOL7 | NOL7 | 1413 | 1523 | 1492 | 1752 | 911 | 586 | 1434 | 1768 | 0.002219945 | 0.017649788 |
| ENSGALG00010007800 | MAF | MAF | 153 | 211 | 190 | 294 | 131 | 71 | 58 | 95 | 0.003612549 | 0.026090281 |
| ENSGALG00010007802 | LOC770839 | LOC770839 | 98 | 52 | 1 | 28 | 239 | 106 | 179 | 182 | 3.27E-06 | 7.45E-05 |
| ENSGALG00010007803 | ME3 | ME3 | 159 | 50 | 400 | 396 | 375 | 325 | 523 | 484 | 4.73E-05 | 0.000734391 |
| ENSGALG00010007811 | PRCP | PRCP | 260 | 331 | 402 | 489 | 540 | 520 | 533 | 647 | 0.003119788 | 0.0231827 |
| ENSGALG00010007826 | LOC107052660 | LOC107052660 | 0 | 0 | 2 | 0 | 72 | 112 | 61 | 60 | 2.73E-25 | 7.87E-23 |
| ENSGALG00010007850 | NA | NA | 47 | 57 | 54 | 39 | 10 | 3 | 23 | 14 | 5.58E-05 | 0.000840791 |
| ENSGALG00010007873 | NA | NA | 0 | 0 | 258 | 240 | 30 | 38 | 0 | 0 | 1.67E-41 | 1.27E-38 |
| ENSGALG00010007882 | CPE | CPE | 1655 | 1774 | 747 | 782 | 770 | 599 | 1475 | 1537 | 7.93E-09 | 3.47E-07 |
| ENSGALG00010007888 | MIOX | MIOX | 38 | 22 | 29 | 5 | 13 | 90 | 152 | 217 | 0.000255861 | 0.003067397 |
| ENSGALG00010007889 | DCAF6 | IQWD1 | 701 | 865 | 1466 | 1374 | 1231 | 701 | 764 | 748 | 0.001799998 | 0.014884498 |
| ENSGALG00010007892 | LOC121109261 | LOC121109261 | 78 | 106 | 10 | 12 | 14 | 13 | 12 | 6 | 3.64E-11 | 2.46E-09 |
| ENSGALG00010007894 | CD88 | CD80 | 7 | 8 | 31 | 32 | 22 | 49 | 41 | 25 | 0.003610011 | 0.026083225 |
| ENSGALG00010007901 | FSTL5 | SPIG2-A | 34 | 423 | 72 | 676 | 68 | 23 | 14 | 33 | 0.000240656 | 0.002904032 |
| ENSGALG00010007907 | IRX2 | IRX2 | 52 | 128 | 39 | 26 | 18 | 16 | 40 | 79 | 0.002508374 | 0.019407385 |
| ENSGALG00010007913 | PABPN1L | PABPN1L | 178 | 207 | 464 | 506 | 270 | 233 | 229 | 316 | 0.005381056 | 0.035724447 |
| ENSGALG00010007917 | POGLUT1 | KTELC1 | 2832 | 4125 | 3530 | 5726 | 3599 | 2457 | 2397 | 2221 | 0.001342204 | 0.011822698 |

| | | | | | | | | | | | | |
|---|---|---|---|---|---|---|---|---|---|---|---|---|
| ENSGALG00010007936 | COL15A1 | COL15A1 | 3 | 0 | 29 | 24 | 25 | 12 | 14 | 3 | 0.000233992 | 0.002846913 |
| ENSGALG00010007950 | ARC | ARG3.1 | 105 | 293 | 81 | 121 | 4 | 22 | 47 | 170 | 0.000277472 | 0.003291064 |
| ENSGALG00010007955 | TMA16 | C4H4ORF43 | 716 | 679 | 1151 | 1412 | 869 | 637 | 676 | 819 | 0.009270151 | 0.054976139 |
| ENSGALG00010007962 | ADCY8 | ADCY8 | 106 | 58 | 18 | 15 | 10 | 24 | 72 | 78 | 5.22E-05 | 0.000796177 |
| ENSGALG00010007964 | LOC101748706 | LOC101748706 | 586 | 781 | 386 | 425 | 425 | 424 | 1034 | 696 | 0.002297961 | 0.018131972 |
| ENSGALG00010007968 | SLC22A31 | SLC22A31 | 162 | 103 | 19 | 31 | 51 | 28 | 82 | 64 | 2.44E-05 | 0.000417936 |
| ENSGALG00010007969 | NA | NA | 0 | 0 | 7 | 0 | 5 | 2 | 22 | 23 | 5.94E-06 | 0.000124792 |
| ENSGALG00010007976 | NA | NA | 0 | 0 | 15 | 23 | 35 | 13 | 15 | 27 | 1.96E-07 | 6.18E-06 |
| ENSGALG00010007981 | TDRD15 | TDRD15 | 527 | 403 | 1446 | 1275 | 1099 | 724 | 456 | 281 | 3.07E-10 | 1.73E-08 |
| ENSGALG00010007984 | EIF3H | EIF3S3 | 3931 | 3714 | 2909 | 2895 | 2557 | 2059 | 3720 | 4207 | 0.007559963 | 0.046620702 |
| ENSGALG00010007987 | NA | NA | 271 | 248 | 77 | 92 | 81 | 97 | 380 | 384 | 1.55E-08 | 6.32E-07 |
| ENSGALG00010007992 | LAPTM4A | LAPTM4A | 5502 | 5885 | 2870 | 3723 | 3614 | 3223 | 4322 | 4138 | 0.000130697 | 0.001729378 |
| ENSGALG00010008003 | NA | NA | 8 | 11 | 95 | 116 | 30 | 32 | 18 | 64 | 1.20E-05 | 0.000229541 |
| ENSGALG00010008008 | BTC | probetacellulin | 180 | 171 | 87 | 75 | 115 | 84 | 259 | 224 | 0.00092458 | 0.008759398 |
| ENSGALG00010008022 | SLC35F3 | SLC35F3 | 607 | 1217 | 964 | 1507 | 1051 | 466 | 376 | 420 | 0.000211249 | 0.002617047 |
| ENSGALG00010008028 | KCNK7 | KCNK1 | 2796 | 3625 | 2349 | 1964 | 2726 | 1842 | 3383 | 3176 | 0.009959934 | 0.058118165 |
| ENSGALG00010008034 | DPEP1 | DPEP1 | 18 | 2 | 1 | 0 | 8 | 1 | 9 | 24 | 0.005690459 | 0.037300012 |
| ENSGALG00010008045 | DCDC2 | DCDC2 | 630 | 495 | 287 | 602 | 293 | 127 | 269 | 195 | 0.003545559 | 0.025684174 |
| ENSGALG00010008055 | MLC1 | MLC1 | 31 | 49 | 58 | 104 | 23 | 23 | 16 | 22 | 0.001812083 | 0.014962217 |
| ENSGALG00010008066 | PCNX2 | PCNXL2 | 1032 | 924 | 428 | 570 | 677 | 729 | 1290 | 867 | 0.000797063 | 0.007762536 |
| ENSGALG00010008069 | DLG2 | DLG2 | 86 | 79 | 67 | 60 | 28 | 12 | 45 | 38 | 0.00471726 | 0.032402319 |
| ENSGALG00010008079 | CCDC127 | CCDC127 | 719 | 927 | 1275 | 1080 | 1214 | 917 | 711 | 508 | 0.000581429 | 0.006010631 |
| ENSGALG00010008092 | SPIRE2 | Spir-2 | 15 | 38 | 14 | 8 | 13 | 5 | 49 | 56 | 0.000384393 | 0.004288364 |
| ENSGALG00010008094 | PIM3 | PIM3 | 2430 | 2880 | 1447 | 1778 | 1935 | 1321 | 1786 | 1172 | 0.00130859 | 0.011570982 |
| ENSGALG00010008097 | LOC420808 | LOC420808 | 1 | 6 | 80 | 25 | 54 | 7 | 2 | 3 | 2.25E-06 | 5.35E-05 |
| ENSGALG00010008100 | LOC121106514 | LOC121106514 | 0 | 4 | 44 | 40 | 2 | 10 | 12 | 8 | 1.38E-05 | 0.000256777 |
| ENSGALG00010008107 | TUBB3 | TUBB3 | 7926 | 7109 | 4915 | 5447 | 4050 | 2208 | 4543 | 6850 | 0.000142299 | 0.001863716 |
| ENSGALG00010008118 | NA | NA | 0 | 0 | 582 | 705 | 1119 | 722 | 579 | 831 | 1.47E-65 | 2.46E-62 |
| ENSGALG00010008126 | CDH3 | K-CAM | 3741 | 3436 | 2567 | 2235 | 2055 | 1671 | 3522 | 5629 | 1.54E-05 | 0.000283154 |
| ENSGALG00010008129 | TMCO7 | TANGO6 | 666 | 775 | 1612 | 1595 | 1040 | 662 | 785 | 1166 | 0.000689429 | 0.006912811 |
| ENSGALG00010008150 | TERT | chTERT | 257 | 278 | 223 | 192 | 213 | 148 | 526 | 416 | 0.001674176 | 0.014066785 |
| ENSGALG00010008152 | FRAS1 | FRAS1 | 1793 | 2784 | 1828 | 1801 | 1869 | 1312 | 3934 | 2433 | 0.004800211 | 0.032810274 |
| ENSGALG00010008159 | SUPT3H | SUPT3H | 436 | 543 | 949 | 980 | 806 | 613 | 746 | 678 | 0.007750614 | 0.047585695 |
| ENSGALG00010008165 | MRPL1 | MRPL1 | 655 | 643 | 1271 | 1333 | 1199 | 829 | 726 | 879 | 0.000418473 | 0.004585916 |
| ENSGALG00010008169 | HTR4 | HTR4 | 81 | 133 | 203 | 358 | 142 | 196 | 733 | 429 | 1.87E-06 | 4.58E-05 |
| ENSGALG00010008173 | LOC100859549 | LOC100859549 | 1 | 1 | 3 | 4 | 6 | 3 | 13 | 11 | 0.00475008 | 0.032520827 |
| ENSGALG00010008178 | PGR | NR3C3 | 728 | 528 | 1550 | 1302 | 1003 | 706 | 1127 | 823 | 0.003249467 | 0.023975756 |
| ENSGALG00010008179 | SEMA6A | SEMA6A | 2623 | 1384 | 534 | 453 | 1405 | 689 | 1579 | 886 | 1.11E-06 | 2.86E-05 |
| ENSGALG00010008191 | NA | NA | 3 | 0 | 28 | 54 | 5 | 4 | 6 | 0 | 3.98E-07 | 1.15E-05 |
| ENSGALG00010008199 | TPRKB | TPRKB | 546 | 621 | 1200 | 1108 | 830 | 572 | 666 | 769 | 0.005310362 | 0.035356486 |
| ENSGALG00010008208 | ILDR2 | C1orf32 | 2725 | 3404 | 2791 | 2782 | 3352 | 3748 | 4414 | 4181 | 0.002765849 | 0.021000851 |
| ENSGALG00010008214 | CCNG2 | CCNG2 | 2379 | 1315 | 779 | 732 | 1216 | 952 | 2870 | 2216 | 5.42E-08 | 1.94E-06 |
| ENSGALG00010008218 | ARHGAP42 | ARHGAP42 | 56 | 55 | 167 | 190 | 172 | 119 | 136 | 111 | 0.001012598 | 0.009443376 |
| ENSGALG00010008225 | NA | NA | 44 | 32 | 15 | 24 | 48 | 48 | 79 | 56 | 0.006172388 | 0.039643714 |
| ENSGALG00010008261 | OTX5 | CRX | 345 | 299 | 635 | 492 | 120 | 73 | 311 | 396 | 2.32E-07 | 7.14E-06 |
| ENSGALG00010008264 | CNTNAP2 | CNTNAP2 | 302 | 352 | 501 | 576 | 564 | 404 | 327 | 243 | 0.004375839 | 0.03039444 |
| ENSGALG00010008271 | NA | NA | 2 | 12 | 22 | 16 | 4 | 1 | 2 | 3 | 0.005670176 | 0.037233787 |
| ENSGALG00010008284 | BCKDHA | BCKDHA | 1056 | 1576 | 2198 | 2877 | 1183 | 1070 | 1279 | 2024 | 0.004739788 | 0.032490292 |
| ENSGALG00010008285 | ADSS2 | ADSS | 11553 | 11208 | 7248 | 9164 | 9621 | 6400 | 7582 | 8060 | 0.006754313 | 0.042531302 |
| ENSGALG00010008290 | FAM13A | FAM13A1 | 675 | 662 | 1071 | 919 | 1261 | 982 | 1489 | 1414 | 7.17E-05 | 0.001040809 |
| ENSGALG00010008296 | PALLD | PALLD | 283 | 445 | 477 | 529 | 356 | 254 | 595 | 840 | 0.009007606 | 0.053705001 |
| ENSGALG00010008302 | NA | NA | 291 | 527 | 133 | 109 | 123 | 172 | 348 | 290 | 0.000190068 | 0.002393676 |
| ENSGALG00010008306 | NA | NA | 4 | 18 | 17 | 11 | 6 | 1 | 1 | 1 | 0.006732494 | 0.04240989 |
| ENSGALG00010008318 | ADPRH | ADPRH | 90 | 303 | 43 | 48 | 35 | 54 | 38 | 36 | 1.93E-05 | 0.00034131 |
| ENSGALG00010008327 | TGFB1 | TGF-beta4 | 2961 | 4171 | 3168 | 2544 | 1935 | 1598 | 1398 | 2487 | 0.004661863 | 0.032061341 |
| ENSGALG00010008340 | PDIA4 | PDIA4 | 17199 | 13919 | 11617 | 14704 | 8521 | 6314 | 8440 | 9053 | 1.61E-05 | 0.000294659 |
| ENSGALG00010008352 | LOC101748353 | C19orf47 | 4573 | 5484 | 2913 | 3419 | 2770 | 2356 | 3687 | 4450 | 0.00016256 | 0.002092985 |
| ENSGALG00010008355 | MAFA | L-MAF | 1012 | 1878 | 729 | 853 | 542 | 563 | 1086 | 1354 | 0.000393022 | 0.004364246 |
| ENSGALG00010008364 | NA | NA | 59 | 15 | 15 | 12 | 4 | 2 | 2 | 11 | 0.000680264 | 0.006856797 |
| ENSGALG00010008365 | NA | NA | 0 | 10 | 44 | 15 | 17 | 5 | 2 | 0 | 0.000674433 | 0.006810343 |
| ENSGALG00010008368 | SNCA | SNCA | 236 | 376 | 68 | 78 | 137 | 150 | 165 | 154 | 2.57E-05 | 0.000437987 |
| ENSGALG00010008390 | NA | NA | 8 | 4 | 38 | 24 | 17 | 7 | 2 | 0 | 6.52E-06 | 0.000135525 |
| ENSGALG00010008393 | NA | NA | 3 | 3 | 0 | 2 | 1 | 4 | 14 | 13 | 0.002233988 | 0.017736059 |
| ENSGALG00010008407 | GNAI1 | GNAI1 | 5078 | 6426 | 4423 | 4945 | 5272 | 2710 | 3064 | 3029 | 0.000270757 | 0.00322056 |
| ENSGALG00010008414 | NA | NA | 14 | 2 | 4 | 5 | 1 | 1 | 23 | 23 | 0.000557479 | 0.005808351 |
| ENSGALG00010008419 | NA | NA | 4 | 5 | 0 | 0 | 14 | 15 | 5 | 9 | 6.28E-06 | 0.000131089 |
| ENSGALG00010008428 | NA | NA | 68 | 98 | 34 | 21 | 85 | 47 | 114 | 144 | 0.000277122 | 0.003289249 |
| ENSGALG00010008430 | NA | NA | 6 | 16 | 74 | 34 | 46 | 4 | 5 | 8 | 0.002149995 | 0.017224697 |
| ENSGALG00010008433 | EBAG9 | EBAG9 | 1691 | 1391 | 926 | 1058 | 855 | 714 | 1099 | 1203 | 0.008433286 | 0.050934976 |
| ENSGALG00010008439 | NA | NA | 3 | 0 | 59 | 32 | 13 | 0 | 3 | 1 | 1.37E-06 | 3.46E-05 |
| ENSGALG00010008444 | SLC7A11 | SLC7A11 | 20 | 6 | 22 | 22 | 24 | 18 | 60 | 67 | 0.001488543 | 0.012783419 |
| ENSGALG00010008454 | NA | NA | 7 | 2 | 3 | 4 | 0 | 1 | 11 | 28 | 0.000278578 | 0.003299506 |
| ENSGALG00010008458 | NA | NA | 3 | 12 | 65 | 31 | 14 | 6 | 6 | 2 | 2.24E-05 | 0.000389596 |
| ENSGALG00010008465 | NA | NA | 4 | 10 | 82 | 22 | 23 | 3 | 5 | 5 | 0.000237603 | 0.002877693 |
| ENSGALG00010008469 | NOCT | CCRN4L | 631 | 876 | 1238 | 1470 | 890 | 763 | 746 | 699 | 0.003501199 | 0.02542896 |
| ENSGALG00010008474 | NA | NA | 1814 | 1711 | 1489 | 1178 | 1278 | 723 | 1008 | 893 | 0.00496946 | 0.033623323 |
| ENSGALG00010008498 | LOC107050955 | LOC107050955 | 8170 | 8719 | 2841 | 1828 | 3246 | 1754 | 2414 | 2524 | 6.21E-19 | 1.13E-16 |
| ENSGALG00010008509 | MGARP | MGARP | 60 | 51 | 22 | 29 | 45 | 37 | 91 | 80 | 0.005760368 | 0.037633801 |
| ENSGALG00010008522 | NUDT12 | NUDT12 | 124 | 61 | 160 | 109 | 176 | 21 | 12 | 4 | 2.59E-05 | 0.000439666 |
| ENSGALG00010008541 | NA | NA | 108 | 68 | 86 | 74 | 164 | 133 | 204 | 155 | 0.009274471 | 0.05498225 |
| ENSGALG00010008546 | NA | NA | 69 | 40 | 4 | 6 | 14 | 12 | 32 | 22 | 2.62E-06 | 6.11E-05 |
| ENSGALG00010008602 | NA | NA | 638 | 1019 | 3 | 5 | 8 | 3 | 6 | 10 | 7.56E-77 | 1.80E-73 |
| ENSGALG00010008603 | NA | NA | 44 | 78 | 8 | 10 | 25 | 10 | 3 | 3 | 2.95E-09 | 1.41E-07 |
| ENSGALG00010008639 | NA | NA | 1 | 5 | 11 | 26 | 8 | 1 | 0 | 0 | 5.29E-05 | 0.000806178 |
| ENSGALG00010008656 | CCDC6 | CCDC6 | 1133 | 1515 | 2912 | 2825 | 2199 | 1481 | 2344 | 2367 | 4.38E-05 | 0.000690078 |
| ENSGALG00010008666 | SLC16A9 | MCT9 | 638 | 864 | 967 | 1174 | 655 | 416 | 450 | 255 | 4.18E-05 | 0.000660226 |
| ENSGALG00010008700 | LOC107051508 | LOC107051508 | 1074 | 361 | 210 | 160 | 375 | 307 | 800 | 632 | 4.02E-05 | 0.000639159 |
| ENSGALG00010008713 | NA | NA | 10 | 5 | 16 | 6 | 0 | 0 | 12 | 5 | 0.0017387 | 0.014490832 |
| ENSGALG00010008793 | C11orf54 | C1H11ORF54 | 137 | 199 | 394 | 371 | 375 | 265 | 288 | 341 | 0.00706961 | 0.044101275 |
| ENSGALG00010008832 | MTSS1 | MTSS1 | 336 | 221 | 752 | 617 | 491 | 368 | 839 | 585 | 0.001368637 | 0.012000345 |
| ENSGALG00010008840 | NA | NA | 14 | 21 | 130 | 54 | 87 | 20 | 21 | 7 | 0.000233705 | 0.002846706 |
| ENSGALG00010008844 | NA | NA | 311 | 395 | 145 | 128 | 258 | 168 | 306 | 200 | 0.004194375 | 0.029439896 |
| ENSGALG00010008852 | KHDRBS3 | KHDRBS3 | 1670 | 1406 | 1546 | 2084 | 1380 | 1076 | 858 | 1067 | 0.006915135 | 0.043364235 |
| ENSGALG00010008858 | NA | NA | 5 | 3 | 7 | 30 | 18 | 25 | 3 | 0 | 0.000127084 | 0.001688255 |
| ENSGALG00010008880 | FMN2 | FMN2 | 1 | 8 | 10 | 20 | 7 | 1 | 29 | 55 | 5.82E-05 | 0.00086964 |
| ENSGALG00010008883 | NA | NA | 2 | 9 | 0 | 0 | 10 | 9 | 2 | 0 | 0.000179252 | 0.002279801 |
| ENSGALG00010008897 | NSMCE2 | NSMCE2 | 505 | 548 | 729 | 725 | 570 | 322 | 269 | 240 | 0.000105862 | 0.001447763 |
| ENSGALG00010008906 | DBN1 | DBN1 | 3996 | 6216 | 2761 | 3488 | 4050 | 2823 | 4381 | 4230 | 0.003293246 | 0.024248134 |
| ENSGALG00010008914 | NA | NA | 453 | 368 | 108 | 211 | 244 | 254 | 546 | 406 | 0.000375079 | 0.004218254 |
| ENSGALG00010008933 | NA | NA | 840 | 1076 | 41 | 100 | 95 | 53 | 150 | 85 | 6.06E-25 | 1.69E-22 |
| ENSGALG00010008946 | COL22A1 | COL22A1 | 50 | 99 | 284 | 536 | 150 | 85 | 116 | 115 | 6.14E-07 | 1.69E-05 |
| ENSGALG00010008956 | LVRN | LVRN | 77 | 20 | 5 | 7 | 19 | 10 | 61 | 25 | 0.000476524 | 0.005085361 |
| ENSGALG00010008968 | NA | NA | 4414 | 843 | 479 | 695 | 80 | 252 | 1418 | 1303 | 1.34E-07 | 4.44E-06 |
| ENSGALG00010008982 | NA | NA | 30 | 30 | 1 | 4 | 2 | 0 | 1 | 0 | 4.71E-10 | 2.58E-08 |
| ENSGALG00010009024 | PMAIP1 | PMAIP1 | 12 | 19 | 95 | 89 | 49 | 26 | 33 | 67 | 0.000414751 | 0.004551093 |
| ENSGALG00010009038 | RAD21 | RAD21 | 13975 | 14618 | 9477 | 11027 | 9532 | 7469 | 11140 | 10134 | 0.007205023 | 0.044745579 |
| ENSGALG00010009048 | NA | NA | 14 | 13 | 65 | 28 | 41 | 27 | 5 | 11 | 0.000331766 | 0.003794981 |
| ENSGALG00010009050 | PIK3CG | PIK3CG | 39 | 52 | 154 | 138 | 125 | 79 | 54 | 53 | 0.000339876 | 0.003874497 |
| ENSGALG00010009054 | RYR2 | RYR2 | 373 | 367 | 1017 | 1263 | 681 | 331 | 846 | 693 | 8.79E-06 | 0.000175716 |
| ENSGALG00010009067 | NA | NA | 0 | 0 | 89 | 98 | 33 | 54 | 0 | 0 | 6.31E-25 | 1.73E-22 |
| ENSGALG00010009092 | EED | EED | 881 | 863 | 1327 | 1746 | 1119 | 823 | 939 | 874 | 0.009461264 | 0.055737852 |
| ENSGALG00010009099 | CXCR4 | CXCR4 | 828 | 649 | 217 | 348 | 543 | 264 | 369 | 318 | 0.000240297 | 0.002903717 |
| ENSGALG00010009104 | MTR | MTR | 725 | 732 | 1351 | 1359 | 934 | 751 | 760 | 647 | 0.001028419 | 0.009550751 |
| ENSGALG00010009110 | NA | NA | 0 | 0 | 0 | 7 | 23 | 61 | 0 | 18 | 2.47E-06 | 5.81E-05 |
| ENSGALG00010009111 | LRFN1 | LRFN1 | 2097 | 1649 | 1086 | 1235 | 1269 | 869 | 1172 | 797 | 0.001643777 | 0.013870136 |
| ENSGALG00010009114 | SETD7 | SETD7 | 2539 | 3042 | 3272 | 3920 | 4288 | 3363 | 3980 | 4084 | 0.001846488 | 0.015178803 |
| ENSGALG00010009132 | SHC3 | SHC3 | 7 | 4 | 17 | 13 | 37 | 10 | 5 | 4 | 0.003080433 | 0.022949825 |
| ENSGALG00010009142 | LGALS8 | galectin-8 | 307 | 369 | 834 | 833 | 419 | 305 | 405 | 518 | 0.000516128 | 0.005441978 |
| ENSGALG00010009157 | ABHD1 | ABHD1 | 1476 | 1168 | 560 | 756 | 783 | 582 | 1103 | 1226 | 0.000185658 | 0.002343444 |
| ENSGALG00010009168 | MRPS12 | MRPS12 | 1371 | 1463 | 993 | 1150 | 969 | 687 | 1495 | 1969 | 0.001833145 | 0.015106273 |
| ENSGALG00010009172 | SPOPL | SPOPL | 2344 | 1360 | 554 | 710 | 1132 | 1033 | 2653 | 1368 | 4.06E-06 | 9.00E-05 |
| ENSGALG00010009191 | NA | NA | 12 | 25 | 49 | 71 | 53 | 62 | 106 | 61 | 0.002127124 | 0.017074206 |
| ENSGALG00010009204 | NA | NA | 57 | 45 | 202 | 186 | 152 | 74 | 82 | 97 | 0.000835175 | 0.008081902 |
| ENSGALG00010009213 | DUS4L | DUS4L | 513 | 441 | 763 | 828 | 562 | 389 | 353 | 315 | 0.001363042 | 0.011957562 |
| ENSGALG00010009231 | NA | NA | 9 | 4 | 7 | 11 | 18 | 40 | 38 | 26 | 0.000738977 | 0.007329158 |
| ENSGALG00010009257 | LOC424300 | LOC424300 | 6 | 4 | 22 | 16 | 43 | 40 | 26 | 17 | 2.26E-05 | 0.00039065 |
| ENSGALG00010009258 | DNAJC12 | DNAJC12 | 335 | 497 | 951 | 868 | 930 | 666 | 520 | 425 | 1.76E-05 | 0.000316866 |
| ENSGALG00010009279 | ELMOD2 | ELMOD2 | 469 | 668 | 1192 | 860 | 902 | 488 | 520 | 504 | 0.007052235 | 0.044009315 |
| ENSGALG00010009297 | CTNNA3 | CTNNA3 | 2062 | 2342 | 2299 | 3132 | 1742 | 1678 | 36 | 37 | 2.86E-82 | 7.97E-79 |
| ENSGALG00010009299 | LOC107055388 | MAP4K1 | 65 | 69 | 31 | 35 | 27 | 24 | 71 | 99 | 0.004143581 | 0.029158987 |
| ENSGALG00010009307 | DIRAS2 | DIRAS2 | 677 | 308 | 60 | 47 | 154 | 95 | 215 | 123 | 7.11E-10 | 3.78E-08 |

| | | | | | | | | | | | | | |
|---|---|---|---|---|---|---|---|---|---|---|---|---|---|
| ENSGALG00010009313 | UNC13B | UNC13B | | 5432 | 1496 | 1200 | 1003 | 1550 | 1247 | 4489 | 1612 | 0.001840708 | 0.015153079 |
| ENSGALG00010009328 | SLC22A15L | SLC22A15L | | 457 | 395 | 122 | 167 | 246 | 198 | 517 | 581 | 1.58E-07 | 5.08E-06 |
| ENSGALG00010009338 | KYNU | KYNU | | 50 | 46 | 8 | 31 | 92 | 69 | 139 | 40 | 0.002751184 | 0.020908499 |
| ENSGALG00010009340 | SYK | SYK | | 93 | 94 | 4 | 4 | 65 | 64 | 66 | 62 | 9.57E-12 | 7.27E-10 |
| ENSGALG00010009344 | ENPP2 | ENPP2 | | 17 | 9 | 19 | 7 | 35 | 24 | 36 | 59 | 0.0045255 | 0.03130387 |
| ENSGALG00010009346 | STC2 | STC2 | | 1652 | 1508 | 1810 | 1611 | 1363 | 1693 | 3757 | 3957 | 1.53E-07 | 4.97E-06 |
| ENSGALG00010009359 | REEP3 | REEP3 | | 1363 | 1279 | 408 | 488 | 696 | 690 | 1360 | 1243 | 1.07E-09 | 5.53E-08 |
| ENSGALG00010009365 | CA5A | CA5 | | 1500 | 1252 | 677 | 962 | 608 | 733 | 1122 | 1393 | 0.007611207 | 0.04688481 |
| ENSGALG00010009366 | NA | NA | | 1 | 3 | 23 | 16 | 22 | 14 | 9 | 9 | 0.000400295 | 0.004427765 |
| ENSGALG00010009373 | ARHGAP15 | ARHGAP15 | | 109 | 180 | 26 | 73 | 110 | 453 | 143 | 83 | 0.000177406 | 0.002261482 |
| ENSGALG00010009382 | NA | NA | | 99 | 70 | 119 | 86 | 7 | 2 | 3 | 3 | 1.66E-17 | 2.57E-15 |
| ENSGALG00010009384 | NA | NA | | 235 | 263 | 410 | 368 | 594 | 486 | 465 | 544 | 0.000297605 | 0.00346832 |
| ENSGALG00010009385 | GPI | GPI | | 14057 | 16281 | 19787 | 25714 | 17735 | 13606 | 13127 | 14765 | 0.006242911 | 0.039973769 |
| ENSGALG00010009398 | WTIP | WTIP | | 401 | 478 | 1377 | 1530 | 956 | 648 | 703 | 851 | 1.84E-08 | 7.25E-07 |
| ENSGALG00010009404 | GNG4 | GNG4 | | 74 | 66 | 231 | 217 | 220 | 119 | 108 | 119 | 0.000796807 | 0.007762536 |
| ENSGALG00010009416 | COL14A1 | COL14 | | 203 | 124 | 43 | 34 | 51 | 23 | 346 | 289 | 6.69E-12 | 5.18E-10 |
| ENSGALG00010009458 | NA | NA | | 9 | 9 | 48 | 43 | 42 | 34 | 27 | 43 | 0.000920828 | 0.00874368 |
| ENSGALG00010009467 | LOC101748577 | LOC101748577 | | 208 | 223 | 59 | 73 | 93 | 101 | 287 | 227 | 1.37E-05 | 0.000255967 |
| ENSGALG00010009468 | NA | NA | | 113 | 155 | 234 | 196 | 209 | 146 | 68 | 68 | 0.000602031 | 0.006178863 |
| ENSGALG00010009470 | ZNF469 | ZNF469 | | 109 | 158 | 66 | 80 | 59 | 63 | 484 | 174 | 9.48E-06 | 0.000187282 |
| ENSGALG00010009480 | NA | NA | | 29 | 26 | 47 | 92 | 62 | 67 | 27 | 29 | 0.004955027 | 0.033580055 |
| ENSGALG00010009499 | LOC101751218 | LOC101751218 | | 909 | 792 | 469 | 600 | 596 | 522 | 1024 | 930 | 0.006061355 | 0.039065701 |
| ENSGALG00010009502 | ARID5B | ARID5B | | 218 | 214 | 759 | 547 | 536 | 439 | 458 | 357 | 6.80E-05 | 0.000992385 |
| ENSGALG00010009519 | NA | NA | | 0 | 0 | 0 | 36 | 48 | 33 | 0 | 0 | 5.50E-09 | 2.52E-07 |
| ENSGALG00010009542 | WDR1 | WDR1 | | 17352 | 16259 | 8784 | 12375 | 12545 | 10281 | 12180 | 10696 | 0.001245405 | 0.011130059 |
| ENSGALG00010009546 | GAREM2 | FAM59B | | 385 | 473 | 122 | 148 | 147 | 89 | 254 | 202 | 2.54E-06 | 5.95E-05 |
| ENSGALG00010009554 | ABCB10 | ABCB10 | | 802 | 873 | 2104 | 2037 | 1470 | 1031 | 1078 | 1033 | 7.72E-07 | 2.07E-05 |
| ENSGALG00010009567 | NSD1 | NSD1 | | 13841 | 13790 | 8515 | 9685 | 8600 | 8796 | 18831 | 13302 | 0.000447007 | 0.004816495 |
| ENSGALG00010009653 | ABLIM2 | ABLIM2 | | 50 | 57 | 11 | 17 | 34 | 34 | 51 | 31 | 0.005593165 | 0.036843898 |
| ENSGALG00010009656 | NA | NA | | 164 | 161 | 290 | 736 | 440 | 476 | 240 | 222 | 3.08E-05 | 0.000511666 |
| ENSGALG00010009668 | TFAP2A | TFAP2 | | 35 | 42 | 143 | 98 | 110 | 39 | 27 | 30 | 0.000429825 | 0.0046705 |
| ENSGALG00010009678 | JARID2 | JMJ | | 16963 | 11136 | 6371 | 6731 | 10483 | 8225 | 12907 | 10172 | 1.59E-05 | 0.000290495 |
| ENSGALG00010009679 | ANK3 | CHANK3 | | 1053 | 1039 | 356 | 417 | 639 | 490 | 1445 | 960 | 9.97E-09 | 4.22E-07 |
| ENSGALG00010009686 | MAT2B | MAT2B | | 1910 | 1931 | 3034 | 3821 | 2936 | 2237 | 2148 | 2138 | 0.000149057 | 0.001940064 |
| ENSGALG00010009688 | NA | NA | | 1 | 9 | 11 | 17 | 10 | 12 | 37 | 27 | 0.005867557 | 0.038169952 |
| ENSGALG00010009695 | ATAD2B | ATAD2B | | 78 | 121 | 380 | 268 | 273 | 144 | 274 | 224 | 0.001414453 | 0.01230523 |
| ENSGALG00010009702 | NA | NA | | 292 | 327 | 159 | 131 | 121 | 68 | 140 | 126 | 0.000553061 | 0.00578034 |
| ENSGALG00010009707 | NA | NA | | 26 | 25 | 1 | 2 | 4 | 2 | 8 | 13 | 4.87E-06 | 0.000105558 |
| ENSGALG00010009708 | LOC100859314 | LOC100859314 | | 22 | 49 | 6 | 8 | 12 | 18 | 24 | 21 | 0.007429291 | 0.046002103 |
| ENSGALG00010009720 | RELN | RELN | | 234 | 353 | 72 | 89 | 321 | 242 | 254 | 143 | 5.12E-06 | 0.00011022 |
| ENSGALG00010009727 | CDH20 | CDH20 | | 1590 | 1175 | 210 | 355 | 846 | 616 | 1512 | 734 | 2.58E-09 | 1.24E-07 |
| ENSGALG00010009743 | CCNG1 | CCNG1 | | 2414 | 2762 | 5013 | 8797 | 3891 | 3660 | 3121 | 4897 | 1.58E-05 | 0.00028924 |
| ENSGALG00010009749 | NDRG1 | NDRG1 | | 132 | 147 | 413 | 285 | 520 | 682 | 859 | 1166 | 2.48E-13 | 2.25E-11 |
| ENSGALG00010009762 | MAN2A1 | MAN2A1 | | 4907 | 2477 | 1918 | 1857 | 2351 | 1843 | 4979 | 2817 | 0.00175966 | 0.014630566 |
| ENSGALG00010009777 | ST3GAL1 | SIAT4A | | 206 | 249 | 599 | 460 | 516 | 235 | 248 | 286 | 0.004549601 | 0.031431554 |
| ENSGALG00010009795 | NDUFA6 | NDUFA6 | | 966 | 1027 | 2048 | 2469 | 2057 | 1560 | 2094 | 2205 | 6.62E-07 | 1.80E-05 |
| ENSGALG00010009801 | LOC101750251 | LOC101750251 | | 11100 | 14801 | 2276 | 6212 | 1799 | 7071 | 11589 | 11331 | 0.003187998 | 0.02359514 |
| ENSGALG00010009809 | SMDT1 | C1H22orf32 | | 546 | 707 | 2614 | 2868 | 2457 | 1792 | 2199 | 2489 | 6.08E-20 | 1.22E-17 |
| ENSGALG00010009822 | NA | NA | | 5 | 3 | 19 | 40 | 21 | 12 | 14 | 80 | 0.001161982 | 0.010513832 |
| ENSGALG00010009844 | NA | NA | | 2 | 6 | 31 | 28 | 3 | 0 | 127 | 29 | 5.51E-10 | 2.96E-08 |
| ENSGALG00010009858 | ST8SIA5 | SIAT8E | | 6 | 6 | 47 | 34 | 31 | 17 | 70 | 34 | 0.000110093 | 0.001493403 |
| ENSGALG00010009859 | NA | NA | | 24 | 105 | 15 | 7 | 5 | 9 | 22 | 18 | 0.000172576 | 0.002204968 |
| ENSGALG00010009866 | NAGA | NAGA | | 640 | 525 | 229 | 332 | 385 | 318 | 558 | 473 | 0.005854701 | 0.038115999 |
| ENSGALG00010009878 | NA | NA | | 49 | 134 | 13 | 14 | 5 | 24 | 36 | 15 | 0.000400597 | 0.004427765 |
| ENSGALG00010009884 | GFOD1 | GFOD1 | | 30 | 30 | 41 | 40 | 58 | 61 | 119 | 64 | 0.008779846 | 0.052647576 |
| ENSGALG00010009891 | NA | NA | | 546 | 901 | 71 | 120 | 41 | 47 | 58 | 111 | 1.12E-20 | 2.42E-18 |
| ENSGALG00010009906 | NA | NA | | 0 | 3 | 26 | 24 | 7 | 2 | 0 | 0 | 3.72E-08 | 1.38E-06 |
| ENSGALG00010009925 | NA | NA | | 35 | 27 | 128 | 47 | 48 | 7 | 16 | 14 | 0.004725706 | 0.032433674 |
| ENSGALG00010009928 | NA | NA | | 1389 | 668 | 70 | 142 | 76 | 84 | 298 | 363 | 6.31E-18 | 1.02E-15 |
| ENSGALG00010009941 | CACNA1E | CACNA1E | | 360 | 851 | 254 | 266 | 208 | 211 | 517 | 737 | 0.000214647 | 0.002649319 |
| ENSGALG00010009954 | NA | NA | | 231 | 192 | 264 | 217 | 105 | 49 | 118 | 69 | 0.001792314 | 0.014828291 |
| ENSGALG00010009978 | NA | NA | | 3 | 1 | 35 | 12 | 35 | 6 | 31 | 10 | 0.006249701 | 0.040001919 |
| ENSGALG00010009979 | NA | NA | | 20 | 29 | 54 | 108 | 30 | 20 | 13 | 24 | 0.001121587 | 0.010232309 |
| ENSGALG00010009991 | LHFPL3 | LHFPL3 | | 685 | 1002 | 219 | 172 | 484 | 342 | 596 | 402 | 4.59E-08 | 1.68E-06 |
| ENSGALG00010009995 | SEPTIN3 | | 3-Sep | 29 | 44 | 2 | 0 | 6 | 4 | 29 | 30 | 2.14E-09 | 1.05E-07 |
| ENSGALG00010010007 | CTIF | CTIF | | 256 | 541 | 1138 | 984 | 863 | 415 | 572 | 415 | 0.000858531 | 0.008269604 |
| ENSGALG00010010019 | NA | NA | | 67 | 42 | 85 | 261 | 137 | 89 | 30 | 52 | 0.000945981 | 0.008916662 |
| ENSGALG00010010035 | ATP10B | ATP10B | | 436 | 142 | 217 | 322 | 209 | 88 | 83 | 80 | 0.00431975 | 0.030093509 |
| ENSGALG00010010054 | NA | NA | | 55 | 127 | 23 | 19 | 42 | 58 | 87 | 130 | 0.000405687 | 0.004472187 |
| ENSGALG00010010075 | NA | NA | | 105 | 56 | 145 | 65 | 14 | 5 | 7 | 4 | 1.59E-10 | 9.37E-09 |
| ENSGALG00010010076 | NA | NA | | 14 | 26 | 3 | 3 | 1 | 1 | 3 | 4 | 3.07E-05 | 0.000511218 |
| ENSGALG00010010084 | HK3 | HK3 | | 266 | 532 | 151 | 131 | 193 | 163 | 306 | 446 | 0.0005465 | 0.005726088 |
| ENSGALG00010010091 | NA | NA | | 4 | 11 | 4 | 2 | 5 | 5 | 29 | 32 | 4.39E-05 | 0.000690802 |
| ENSGALG00010010092 | NA | NA | | 1 | 3 | 17 | 27 | 20 | 14 | 17 | 38 | 0.000192221 | 0.0024099 |
| ENSGALG00010010094 | SLC2A9 | SLC2A9 | | 1620 | 2010 | 915 | 981 | 1331 | 946 | 1900 | 1463 | 0.00024067 | 0.002904032 |
| ENSGALG00010010099 | TNR | tenascin-R | | 15 | 7 | 4 | 6 | 42 | 36 | 21 | 14 | 1.63E-05 | 0.000296282 |
| ENSGALG00010010115 | NEB | NEB | | 1810 | 1577 | 1668 | 2001 | 5497 | 4073 | 5804 | 3143 | 1.08E-14 | 1.21E-12 |
| ENSGALG00010010128 | NA | NA | | 9 | 28 | 160 | 68 | 77 | 9 | 4 | 6 | 2.84E-06 | 6.58E-05 |
| ENSGALG00010010156 | IFNAR2 | IFNAR2 | | 299 | 298 | 480 | 598 | 423 | 203 | 209 | 217 | 0.004241106 | 0.02968064 |
| ENSGALG00010010161 | SEPSECS | SEPSECS | | 725 | 678 | 883 | 763 | 823 | 647 | 1286 | 1339 | 0.004253569 | 0.02974295 |
| ENSGALG00010010165 | NA | NA | | 1 | 8 | 11 | 10 | 9 | 11 | 1 | 0 | 0.00261409 | 0.020039761 |
| ENSGALG00010010166 | NA | NA | | 120 | 184 | 6 | 11 | 1 | 4 | 142 | 125 | 6.38E-23 | 1.58E-20 |
| ENSGALG00010010169 | NA | NA | | 256 | 1027 | 0 | 0 | 687 | 795 | 349 | 1348 | 1.54E-24 | 4.09E-22 |
| ENSGALG00010010172 | NA | NA | | 25 | 137 | 0 | 0 | 71 | 96 | 38 | 127 | 1.34E-10 | 8.03E-09 |
| ENSGALG00010010174 | NA | NA | | 44 | 239 | 0 | 0 | 150 | 209 | 130 | 288 | 2.95E-15 | 3.50E-13 |
| ENSGALG00010010178 | LGI2 | LGI2 | | 286 | 295 | 471 | 541 | 421 | 282 | 223 | 182 | 0.001269225 | 0.011303036 |
| ENSGALG00010010206 | SCARA5 | SCARA5 | | 66 | 97 | 13 | 21 | 13 | 19 | 45 | 35 | 5.07E-05 | 0.000779218 |
| ENSGALG00010010220 | LHX9 | LH-2 | | 8 | 16 | 34 | 97 | 48 | 37 | 27 | 24 | 0.001234772 | 0.011070552 |
| ENSGALG00010010228 | NA | NA | | 6 | 69 | 7 | 4 | 2 | 10 | 42 | 41 | 0.002564523 | 0.019732188 |
| ENSGALG00010010245 | GALR3 | GALR3 | | 23 | 1 | 12 | 14 | 2 | 5 | 55 | 53 | 0.000513438 | 0.005417026 |
| ENSGALG00010010273 | NA | NA | | 3 | 2 | 186 | 150 | 17 | 3 | 184 | 135 | 1.17E-21 | 2.83E-19 |
| ENSGALG00010010280 | HSP90B1 | hsp108 | | 55988 | 46789 | 28846 | 36780 | 28763 | 19042 | 27663 | 29758 | 3.81E-05 | 0.000613424 |
| ENSGALG00010010301 | CDHR3 | CDHR3 | | 94 | 51 | 373 | 221 | 227 | 152 | 98 | 103 | 7.03E-05 | 0.001022563 |
| ENSGALG00010010321 | NUP133 | NUP133 | | 3204 | 3298 | 2134 | 2226 | 2820 | 2170 | 3133 | 3251 | 0.004154099 | 0.029218565 |
| ENSGALG00010010327 | FBP1 | FBP1 | | 265 | 103 | 13 | 32 | 98 | 61 | 153 | 122 | 4.04E-06 | 8.96E-05 |
| ENSGALG00010010349 | RREB1 | RREB-1 | | 244 | 409 | 1032 | 953 | 805 | 398 | 488 | 593 | 6.46E-05 | 0.000948109 |
| ENSGALG00010010376 | NA | NA | | 1 | 4 | 6 | 26 | 13 | 14 | 63 | 23 | 0.000194676 | 0.002433375 |
| ENSGALG00010010378 | NA | NA | | 2 | 4 | 18 | 27 | 3 | 1 | 1 | 2 | 7.49E-06 | 0.000153092 |
| ENSGALG00010010383 | SLIT2 | Slit | | 685 | 756 | 822 | 1157 | 1158 | 671 | 669 | 388 | 0.006268421 | 0.040075691 |
| ENSGALG00010010396 | NA | NA | | 543 | 876 | 694 | 1363 | 635 | 630 | 142 | 103 | 1.28E-13 | 1.22E-11 |
| ENSGALG00010010399 | FGF18 | FGF-18 | | 65 | 216 | 34 | 19 | 35 | 55 | 117 | 127 | 0.000421799 | 0.004612265 |
| ENSGALG00010010415 | CPLX4 | CPLX4 | | 41 | 3 | 2 | 0 | 3 | 1 | 11 | 4 | 0.001923742 | 0.015659803 |
| ENSGALG00010010418 | BRINP2 | FAM5B | | 11 | 57 | 24 | 13 | 19 | 6 | 46 | 90 | 0.009676263 | 0.056780091 |
| ENSGALG00010010422 | CDH1 | cadherin-1 | | 796 | 634 | 418 | 387 | 466 | 605 | 1339 | 1421 | 2.09E-07 | 6.51E-06 |
| ENSGALG00010010430 | ASTN1 | ASTN1 | | 314 | 608 | 135 | 205 | 200 | 151 | 536 | 354 | 0.000125285 | 0.001670998 |
| ENSGALG00010010431 | SOCS2 | SOCS2 | | 183 | 218 | 120 | 253 | 40 | 30 | 37 | 83 | 7.51E-06 | 0.00015315 |
| ENSGALG00010010461 | SLC7A5 | SLC7A5 | | 498 | 399 | 268 | 191 | 375 | 656 | 2217 | 2048 | 2.92E-18 | 5.09E-16 |
| ENSGALG00010010468 | NA | NA | | 1 | 18 | 2 | 0 | 5 | 8 | 30 | 23 | 0.000982673 | 0.009210559 |
| ENSGALG00010010475 | CCDC71L | CCDC71L | | 158 | 227 | 501 | 475 | 332 | 185 | 192 | 225 | 0.001457114 | 0.012565301 |
| ENSGALG00010010486 | LOC769726 | LOC769726 | | 51 | 143 | 28 | 56 | 27 | 24 | 103 | 83 | 0.006460765 | 0.041085354 |
| ENSGALG00010010499 | NR4A2 | NR4A2 | | 41 | 92 | 204 | 154 | 134 | 67 | 73 | 43 | 0.00663085 | 0.041899282 |
| ENSGALG00010010511 | GABRP | GABRP | | 339 | 759 | 49 | 151 | 35 | 65 | 237 | 306 | 2.45E-08 | 9.49E-07 |
| ENSGALG00010010515 | RXRG | RXRG | | 117 | 70 | 17 | 23 | 7 | 14 | 50 | 54 | 2.16E-06 | 5.18E-05 |
| ENSGALG00010010517 | STMN4 | STMN4 | | 7 | 2 | 3 | 11 | 32 | 17 | 3 | 14 | 0.009195224 | 0.054589905 |
| ENSGALG00010010544 | NA | NA | | 88 | 119 | 295 | 336 | 259 | 163 | 186 | 161 | 0.000776282 | 0.007613399 |
| ENSGALG00010010556 | POMC | ACTH | | 1 | 7 | 1 | 2 | 1 | 2 | 16 | 66 | 1.49E-07 | 4.86E-06 |
| ENSGALG00010010567 | ERMN | ERMN | | 2 | 7 | 101 | 79 | 41 | 35 | 6 | 17 | 4.91E-10 | 2.67E-08 |
| ENSGALG00010010569 | NA | NA | | 51 | 23 | 8 | 5 | 17 | 23 | 17 | 9 | 0.002970323 | 0.022260106 |
| ENSGALG00010010578 | GLUD2 | GDH 1 | | 801 | 998 | 1510 | 1831 | 1165 | 852 | 966 | 1019 | 0.003398637 | 0.024813464 |
| ENSGALG00010010582 | NA | NA | | 25 | 33 | 54 | 50 | 9 | 12 | 11 | 2 | 8.38E-05 | 0.00119255 |
| ENSGALG00010010615 | NA | NA | | 7 | 9 | 67 | 40 | 46 | 15 | 27 | 27 | 0.001142099 | 0.010367602 |
| ENSGALG00010010626 | ADTRP | AIG1L | | 40 | 7 | 6 | 3 | 18 | 16 | 23 | 46 | 0.00931398 | 0.055088806 |
| ENSGALG00010010631 | NA | NA | | 14 | 5 | 67 | 104 | 16 | 10 | 14 | 12 | 4.99E-08 | 1.82E-06 |
| ENSGALG00010010639 | NA | NA | | 0 | 0 | 387 | 606 | 0 | 0 | 0 | 0 | 1.25E-72 | 2.32E-69 |
| ENSGALG00010010678 | KCNIP1 | KCNIP1 | | 732 | 1768 | 70 | 406 | 53 | 119 | 1257 | 1641 | 8.51E-10 | 4.46E-08 |
| ENSGALG00010010688 | IQGAP2 | IQGAP2 | | 899 | 752 | 2701 | 2141 | 2292 | 629 | 1430 | 749 | 0.00173137 | 0.014438455 |
| ENSGALG00010010690 | MTPN | MTPN | | 7708 | 8086 | 5443 | 5940 | 5320 | 4516 | 7128 | 8046 | 0.006049538 | 0.039004584 |
| ENSGALG00010010703 | LOC121108570 | LOC121108570 | | 164 | 65 | 97 | 43 | 71 | 43 | 21 | 30 | 0.006793438 | 0.042729371 |
| ENSGALG00010010705 | NA | NA | | 40 | 114 | 89 | 122 | 145 | 80 | 27 | 43 | 0.006949612 | 0.043445443 |
| ENSGALG00010010734 | AMN1 | AMN1 | | 33 | 71 | 151 | 192 | 172 | 83 | 53 | 58 | 0.000249932 | 0.00300494 |
| ENSGALG00010010753 | LOC107051636 | LOC107051636 | | 215 | 108 | 19 | 24 | 22 | 12 | 12 | 48 | 5.29E-08 | 1.90E-06 |

| | | | | | | | | | | | | |
|---|---|---|---|---|---|---|---|---|---|---|---|---|
| ENSGALG00010010761 | MEI1 | MEI1 | 161 | 260 | 725 | 630 | 547 | 447 | 525 | 509 | 1.03E-05 | 0.000200182 |
| ENSGALG00010010778 | UPP2 | UPP2 | 31 | 30 | 81 | 137 | 58 | 33 | 40 | 51 | 0.005124006 | 0.03433536 |
| ENSGALG00010010779 | DOCK2 | DOCK2 | 143 | 225 | 953 | 632 | 788 | 312 | 148 | 161 | 2.12E-10 | 1.23E-08 |
| ENSGALG00010010789 | ZNF541 | ZNF541 | 3683 | 4610 | 6979 | 7039 | 5685 | 3894 | 4216 | 4941 | 0.001594073 | 0.013522915 |
| ENSGALG00010010796 | PRKCDBP | CAVIN2 | 142 | 71 | 57 | 82 | 74 | 89 | 300 | 420 | 1.12E-06 | 2.88E-05 |
| ENSGALG00010010836 | DIP2A | DIP2A | 537 | 650 | 1148 | 1258 | 954 | 603 | 928 | 709 | 0.006801524 | 0.042762669 |
| ENSGALG00010010845 | LOC121108177 | LOC121108177 | 1481 | 1415 | 676 | 738 | 1114 | 854 | 859 | 1336 | 0.000862706 | 0.008290711 |
| ENSGALG00010010856 | CSDC2 | CSDC2 | 242 | 571 | 127 | 131 | 379 | 344 | 356 | 286 | 0.000101857 | 0.001401011 |
| ENSGALG00010010858 | LHX4 | LHX4 | 101 | 181 | 59 | 35 | 69 | 17 | 206 | 180 | 9.78E-05 | 0.001356661 |
| ENSGALG00010010867 | QSOX1 | QSCN6 | 3234 | 6554 | 3128 | 3034 | 2162 | 1525 | 2781 | 3649 | 0.000232149 | 0.002831878 |
| ENSGALG00010010871 | GCNT2 | GCNT2 | 300 | 151 | 37 | 108 | 275 | 197 | 312 | 248 | 0.00042793 | 0.004655675 |
| ENSGALG00010010873 | NA | NA | 1 | 12 | 41 | 35 | 29 | 5 | 4 | 3 | 0.000732579 | 0.007287418 |
| ENSGALG00010010885 | SNRPD2 | SNRPD2 | 5037 | 4703 | 2689 | 3258 | 2035 | 1778 | 3951 | 4942 | 3.86E-08 | 1.42E-06 |
| ENSGALG00010010907 | NA | NA | 26 | 57 | 1 | 0 | 0 | 0 | 0 | 0 | 7.84E-17 | 1.12E-14 |
| ENSGALG00010010912 | BAALC | BAALC | 26 | 61 | 34 | 38 | 150 | 123 | 171 | 171 | 8.63E-08 | 3.01E-06 |
| ENSGALG00010010915 | APP | APP | 2822 | 2594 | 2414 | 2224 | 2382 | 1387 | 3416 | 3930 | 0.002557813 | 0.019698693 |
| ENSGALG00010010919 | CDHR1 | PCDH21 | 28 | 20 | 1 | 5 | 5 | 2 | 35 | 19 | 1.97E-05 | 0.000347959 |
| ENSGALG00010010930 | FAM163A | C1orf76 | 1319 | 1927 | 126 | 227 | 155 | 165 | 391 | 304 | 4.86E-28 | 1.77E-25 |
| ENSGALG00010010931 | CYYR1 | CYYR1 | 62 | 119 | 106 | 73 | 120 | 73 | 214 | 374 | 0.000246773 | 0.00297337 |
| ENSGALG00010010938 | FZD6 | FZ-6 | 435 | 396 | 553 | 424 | 3002 | 2622 | 3836 | 3019 | 1.84E-48 | 1.71E-45 |
| ENSGALG00010010945 | PI4K2B | PI4K2B | 896 | 905 | 1432 | 1569 | 1322 | 1089 | 1549 | 1641 | 0.005806048 | 0.037858245 |
| ENSGALG00010010966 | LOC768760 | LOC768760 | 12 | 41 | 30 | 22 | 25 | 15 | 48 | 160 | 0.000975551 | 0.009154079 |
| ENSGALG00010010967 | PRXL2A | C10ORF58 | 42 | 66 | 92 | 79 | 99 | 49 | 154 | 176 | 0.00640092 | 0.040797931 |
| ENSGALG00010010998 | SOAT1 | SOAT1 | 2741 | 4249 | 4318 | 5523 | 4186 | 4377 | 6993 | 5479 | 0.006664226 | 0.042075007 |
| ENSGALG00010011003 | FECH | FECH | 1160 | 963 | 2424 | 2025 | 1901 | 1058 | 1799 | 1191 | 0.006309655 | 0.040308467 |
| ENSGALG00010011010 | NA | NA | 15 | 8 | 57 | 52 | 29 | 18 | 44 | 20 | 0.009498336 | 0.055893026 |
| ENSGALG00010011019 | NECAB1 | EFCBP1 | 43 | 41 | 29 | 34 | 102 | 64 | 67 | 68 | 0.003133789 | 0.023266055 |
| ENSGALG00010011036 | NA | NA | 6 | 19 | 92 | 114 | 77 | 40 | 28 | 35 | 2.96E-06 | 6.78E-05 |
| ENSGALG00010011037 | C8orf88 | C2H8orf88 | 11 | 7 | 65 | 37 | 80 | 27 | 24 | 17 | 0.000179067 | 0.002279186 |
| ENSGALG00010011056 | NA | NA | 2 | 19 | 42 | 33 | 54 | 56 | 19 | 11 | 0.000375873 | 0.004224338 |
| ENSGALG00010011057 | ASCL1 | CASH1 | 28 | 58 | 355 | 396 | 280 | 40 | 19 | 27 | 1.29E-09 | 6.61E-08 |
| ENSGALG00010011080 | EMP1 | EMP1 | 6 | 6 | 4 | 0 | 11 | 8 | 27 | 27 | 6.08E-05 | 0.000902385 |
| ENSGALG00010011088 | NA | NA | 4 | 5 | 32 | 44 | 22 | 20 | 12 | 22 | 0.000302089 | 0.003515672 |
| ENSGALG00010011090 | GRIN2B | GRIN2B | 106 | 109 | 27 | 41 | 57 | 85 | 183 | 78 | 0.004080703 | 0.028774981 |
| ENSGALG00010011102 | NA | NA | 10 | 6 | 1 | 0 | 16 | 9 | 6 | 3 | 0.000487688 | 0.005174755 |
| ENSGALG00010011109 | NA | NA | 0 | 0 | 26 | 17 | 22 | 6 | 10 | 1 | 1.75E-05 | 0.000315911 |
| ENSGALG00010011116 | TAPT1 | TAPT1 | 822 | 675 | 514 | 532 | 405 | 261 | 381 | 442 | 0.006731157 | 0.04240989 |
| ENSGALG00010011118 | GMPR2 | GMPR | 327 | 700 | 1154 | 1185 | 859 | 599 | 799 | 712 | 0.005012964 | 0.033780907 |
| ENSGALG00010011126 | LOC121106455 | LOC121106455 | 12 | 25 | 120 | 145 | 53 | 41 | 67 | 32 | 2.06E-05 | 0.000360843 |
| ENSGALG00010011131 | RET | RET | 1152 | 956 | 949 | 1700 | 689 | 648 | 547 | 248 | 8.71E-05 | 0.001230495 |
| ENSGALG00010011136 | NA | NA | 5 | 18 | 94 | 85 | 65 | 47 | 35 | 69 | 7.13E-05 | 0.001035556 |
| ENSGALG00010011142 | DUSP19 | DUSP19 | 1462 | 1288 | 1440 | 2048 | 1046 | 693 | 918 | 758 | 0.000709228 | 0.007080413 |
| ENSGALG00010011161 | RABGAP1L | RABGAP1L | 537 | 583 | 1349 | 1209 | 849 | 651 | 757 | 761 | 0.000230892 | 0.002822421 |
| ENSGALG00010011166 | SOX7 | CTD-2135J3.4 | 174 | 391 | 42 | 65 | 132 | 51 | 89 | 82 | 7.07E-06 | 0.000145445 |
| ENSGALG00010011172 | NA | NA | 0 | 0 | 10 | 6 | 17 | 1 | 2 | 5 | 0.003358941 | 0.024545091 |
| ENSGALG00010011181 | NA | NA | 12 | 54 | 151 | 118 | 97 | 61 | 63 | 48 | 0.005094125 | 0.034176238 |
| ENSGALG00010011187 | STK32B | STK32B | 49 | 42 | 3 | 5 | 7 | 7 | 27 | 30 | 1.38E-07 | 4.56E-06 |
| ENSGALG00010011211 | SLC37A1 | SLC37A1 | 447 | 632 | 1365 | 2005 | 1156 | 604 | 832 | 669 | 7.73E-07 | 2.07E-05 |
| ENSGALG00010011216 | NA | NA | 8 | 6 | 8 | 21 | 24 | 16 | 2 | 3 | 0.001079423 | 0.009939017 |
| ENSGALG00010011238 | NA | NA | 38 | 9 | 100 | 113 | 143 | 83 | 10 | 17 | 1.07E-07 | 3.64E-06 |
| ENSGALG00010011240 | NA | NA | 2 | 1 | 1 | 0 | 0 | 1 | 118 | 145 | 1.47E-33 | 7.01E-31 |
| ENSGALG00010011247 | CDYL | CDYL | 1039 | 685 | 414 | 511 | 510 | 597 | 1004 | 884 | 0.006160046 | 0.039579655 |
| ENSGALG00010011250 | RESF1 | C12orf35 | 609 | 1250 | 2189 | 2369 | 1931 | 1098 | 1366 | 1260 | 0.000248194 | 0.00298619 |
| ENSGALG00010011252 | NA | NA | 5 | 8 | 3 | 9 | 13 | 35 | 16 | 17 | 0.005985395 | 0.038710496 |
| ENSGALG00010011260 | NA | NA | 12 | 12 | 54 | 37 | 24 | 53 | 75 | 45 | 0.005699842 | 0.037325923 |
| ENSGALG00010011274 | NA | NA | 5 | 0 | 28 | 32 | 10 | 7 | 9 | 5 | 0.000288791 | 0.003379743 |
| ENSGALG00010011278 | TDH | TDH | 224 | 148 | 1787 | 1671 | 912 | 600 | 351 | 593 | 5.10E-19 | 9.46E-17 |
| ENSGALG00010011283 | LONRF3 | RNF127 | 235 | 374 | 793 | 837 | 528 | 429 | 528 | 465 | 0.000382895 | 0.004280651 |
| ENSGALG00010011295 | FAM167A | C8orf13 | 0 | 0 | 0 | 0 | 1 | 3 | 20 | 14 | 2.05E-08 | 8.04E-07 |
| ENSGALG00010011297 | OSR2 | OSR2 | 9 | 20 | 66 | 64 | 30 | 13 | 8 | 7 | 4.81E-06 | 0.00010439 |
| ENSGALG00010011307 | NA | NA | 144 | 46 | 9 | 14 | 32 | 20 | 28 | 39 | 2.86E-05 | 0.000478849 |
| ENSGALG00010011314 | NA | NA | 138 | 313 | 421 | 523 | 217 | 163 | 190 | 170 | 0.005304654 | 0.035356486 |
| ENSGALG00010011321 | MAP2 | MAP2 | 839 | 936 | 672 | 573 | 734 | 775 | 1716 | 1005 | 0.002868708 | 0.021624648 |
| ENSGALG00010011351 | KANK1 | ANKRD15 | 5052 | 3279 | 1970 | 2074 | 3641 | 2494 | 5211 | 2178 | 0.005990482 | 0.03872841 |
| ENSGALG00010011353 | LRP1B | LRP1B | 1106 | 1493 | 2569 | 2592 | 1838 | 1853 | 2518 | 1593 | 0.006091832 | 0.03923187 |
| ENSGALG00010011358 | POU4F1L | BRN-3 | 80 | 199 | 31 | 45 | 31 | 21 | 94 | 141 | 1.96E-05 | 0.000346178 |
| ENSGALG00010011362 | NA | NA | 59 | 74 | 5 | 6 | 14 | 19 | 72 | 23 | 3.79E-06 | 8.50E-05 |
| ENSGALG00010011366 | NA | NA | 73 | 104 | 7 | 10 | 43 | 45 | 140 | 60 | 1.83E-07 | 5.79E-06 |
| ENSGALG00010011376 | NA | NA | 3 | 4 | 5 | 14 | 9 | 12 | 333 | 723 | 2.38E-41 | 1.69E-38 |
| ENSGALG00010011386 | CTSB | CTSB | 4073 | 4410 | 1625 | 1751 | 1596 | 1144 | 2231 | 2746 | 9.80E-14 | 9.47E-12 |
| ENSGALG00010011390 | LOC100857497 | FAM13AL | 68 | 104 | 133 | 147 | 333 | 415 | 278 | 335 | 5.40E-09 | 2.49E-07 |
| ENSGALG00010011424 | ACADL | ACADL | 953 | 1191 | 2004 | 2492 | 1823 | 1157 | 1363 | 1715 | 0.000753978 | 0.007447092 |
| ENSGALG00010011451 | CRHBP | CRHBP | 5 | 36 | 110 | 51 | 81 | 33 | 203 | 247 | 8.89E-06 | 0.000177348 |
| ENSGALG00010011457 | SYCP3 | SYCP3 | 38 | 71 | 190 | 134 | 146 | 85 | 81 | 96 | 0.009766134 | 0.05720702 |
| ENSGALG00010011460 | LAMC2 | LAMC2 | 7 | 8 | 21 | 12 | 32 | 51 | 18 | 27 | 0.000442403 | 0.004785393 |
| ENSGALG00010011461 | IRF4 | IRF4 | 59 | 204 | 49 | 50 | 63 | 61 | 31 | 42 | 0.006437375 | 0.040999011 |
| ENSGALG00010011469 | CPS1 | CPS1 | 152 | 232 | 348 | 495 | 322 | 309 | 546 | 408 | 0.004605792 | 0.031727944 |
| ENSGALG00010011535 | NA | NA | 0 | 0 | 4 | 11 | 4 | 20 | 0 | 19 | 0.005356156 | 0.035619609 |
| ENSGALG00010011538 | PINLYP | PINLYP | 6 | 8 | 50 | 40 | 33 | 14 | 8 | 15 | 0.000234291 | 0.002847611 |
| ENSGALG00010011539 | LANCL1 | LANCL1 | 3780 | 3438 | 5292 | 6000 | 3806 | 2787 | 3599 | 3585 | 0.004794908 | 0.032787439 |
| ENSGALG00010011548 | LOC416197 | PAR2L | 98 | 4 | 1 | 18 | 27 | 11 | 3 | 4 | 0.005501706 | 0.036370454 |
| ENSGALG00010011570 | RGL1 | RGL1 | 351 | 616 | 956 | 1107 | 717 | 639 | 649 | 609 | 0.005639594 | 0.037062088 |
| ENSGALG00010011573 | CMAS | CMAS | 374 | 399 | 612 | 777 | 579 | 383 | 298 | 386 | 0.007580366 | 0.046726837 |
| ENSGALG00010011575 | STARD10 | STARD10 | 28 | 23 | 4 | 0 | 11 | 10 | 58 | 38 | 1.47E-07 | 4.80E-06 |
| ENSGALG00010011588 | COLGALT1 | COLGALT2 | 712 | 832 | 2161 | 2138 | 1858 | 1305 | 1781 | 2208 | 5.43E-09 | 2.49E-07 |
| ENSGALG00010011619 | PCDH1 | protocadherin-1 | 60 | 86 | 69 | 64 | 39 | 39 | 298 | 250 | 2.33E-09 | 1.13E-07 |
| ENSGALG00010011625 | ERBB4 | ERBB4 | 140 | 131 | 512 | 456 | 359 | 257 | 548 | 229 | 0.000499937 | 0.00528794 |
| ENSGALG00010011659 | CD14 | CD14 | 12 | 17 | 19 | 15 | 16 | 19 | 53 | 55 | 0.001507965 | 0.01292365 |
| ENSGALG00010011677 | EDEM3 | EDEM3 | 1911 | 1947 | 1098 | 1134 | 1101 | 914 | 2222 | 1507 | 0.000510196 | 0.005389632 |
| ENSGALG00010011694 | NA | NA | 28 | 29 | 3 | 7 | 1 | 4 | 0 | 3 | 3.54E-06 | 7.98E-05 |
| ENSGALG00010011713 | AVPR1A | VT4 | 143 | 72 | 162 | 285 | 51 | 61 | 103 | 57 | 0.008142788 | 0.049466476 |
| ENSGALG00010011718 | NA | NA | 50 | 18 | 44 | 53 | 13 | 7 | 58 | 52 | 0.00712527 | 0.044365687 |
| ENSGALG00010011723 | SPRY3 | SPRY3 | 467 | 611 | 152 | 249 | 154 | 257 | 834 | 900 | 5.20E-08 | 1.87E-06 |
| ENSGALG00010011760 | NA | NA | 54 | 39 | 175 | 155 | 88 | 60 | 73 | 40 | 0.001187571 | 0.010710567 |
| ENSGALG00010011767 | NA | NA | 12 | 269 | 0 | 6 | 3 | 87 | 33 | 24 | 0.001418335 | 0.012330698 |
| ENSGALG00010011772 | GRHL2 | GRHL2 | 101 | 55 | 60 | 50 | 78 | 125 | 264 | 118 | 0.008242013 | 0.049978421 |
| ENSGALG00010011784 | GRPEL2 | GRPEL2 | 1846 | 2319 | 2812 | 3621 | 2977 | 2187 | 2207 | 2334 | 0.009289102 | 0.055034964 |
| ENSGALG00010011796 | METAP2 | METAP2 | 12156 | 13738 | 8046 | 10043 | 8659 | 7067 | 11387 | 12675 | 0.004564962 | 0.031520281 |
| ENSGALG00010011808 | SMARCA2 | BRM | 180 | 140 | 1973 | 737 | 1332 | 387 | 458 | 246 | 1.54E-08 | 6.29E-07 |
| ENSGALG00010011819 | USP44 | USP44 | 20248 | 21482 | 8127 | 13590 | 11552 | 12254 | 21672 | 17732 | 0.000121245 | 0.001623597 |
| ENSGALG00010011822 | NA | NA | 63 | 91 | 6 | 15 | 4 | 5 | 4 | 8 | 1.01E-11 | 7.60E-10 |
| ENSGALG00010011841 | KCNMB4 | KCNMB4 | 142 | 286 | 181 | 124 | 117 | 120 | 353 | 687 | 4.76E-05 | 0.000738171 |
| ENSGALG00010011849 | NA | NA | 1769 | 2285 | 494 | 1051 | 1066 | 923 | 863 | 1515 | 0.000202446 | 0.002522954 |
| ENSGALG00010011860 | CECR1 | ADA2 | 2 | 3 | 16 | 20 | 30 | 15 | 23 | 15 | 0.000397986 | 0.004410568 |
| ENSGALG00010011893 | LIX1 | LIX1 | 4 | 5 | 8 | 12 | 28 | 19 | 23 | 8 | 0.005912606 | 0.038373386 |
| ENSGALG00010011920 | NA | NA | 2195 | 1655 | 910 | 837 | 1102 | 1231 | 1949 | 1938 | 2.76E-05 | 0.000466704 |
| ENSGALG00010011927 | LOC770996 | LOC770996 | 141 | 87 | 29 | 28 | 34 | 44 | 57 | 39 | 0.000856935 | 0.008263757 |
| ENSGALG00010011929 | ELK3 | ELK3 | 538 | 516 | 506 | 575 | 1060 | 574 | 626 | 581 | 0.005832851 | 0.038000992 |
| ENSGALG00010011931 | ATP5F1AW | ATP5A1W | 17 | 980 | 2 | 3985 | 18 | 292 | 5 | 6 | 0.000555703 | 0.005798494 |
| ENSGALG00010011935 | ST8SIA4 | SIAT8D | 276 | 320 | 620 | 537 | 353 | 163 | 237 | 160 | 0.00100082 | 0.009354419 |
| ENSGALG00010011949 | ABCA12 | ABCA12 | 73 | 119 | 288 | 522 | 214 | 178 | 113 | 97 | 8.37E-07 | 2.21E-05 |
| ENSGALG00010011977 | GATD3A | C1H21ORF33 | 915 | 642 | 1636 | 1847 | 1419 | 988 | 1188 | 1183 | 0.000671566 | 0.006789602 |
| ENSGALG00010011984 | FN1 | FN | 983 | 581 | 1594 | 1954 | 1069 | 1562 | 1744 | 3931 | 0.000194504 | 0.002433052 |
| ENSGALG00010011992 | YEATS4 | GAS41 | 10186 | 13572 | 7983 | 9295 | 7885 | 5861 | 6564 | 7832 | 0.00082725 | 0.008028458 |
| ENSGALG00010011998 | MPPED1 | MPPED1 | 337 | 449 | 783 | 777 | 617 | 386 | 410 | 420 | 0.005963934 | 0.038601573 |
| ENSGALG00010012014 | LOC121110687 | LOC121110687 | 3 | 8 | 18 | 12 | 13 | 8 | 1 | 0 | 0.000326302 | 0.003747878 |
| ENSGALG00010012039 | ODF1 | ODF1 | 4 | 7 | 12 | 16 | 16 | 9 | 30 | 36 | 0.00207792 | 0.016751668 |
| ENSGALG00010012048 | PDE6H | PDE6H | 0 | 5 | 6 | 1 | 5 | 1 | 30 | 47 | 1.56E-06 | 3.93E-05 |
| ENSGALG00010012055 | VEGFRKDRL | VEGFRKDRL | 30 | 75 | 44 | 36 | 45 | 25 | 145 | 206 | 9.89E-06 | 0.000194236 |
| ENSGALG00010012061 | AZIN1 | OAZIN | 2424 | 2528 | 3847 | 4104 | 4088 | 2012 | 2416 | 2521 | 0.005893998 | 0.03831213 |
| ENSGALG00010012066 | STXBP3 | STXBP3 | 935 | 731 | 459 | 492 | 497 | 456 | 961 | 888 | 0.001342887 | 0.011822698 |
| ENSGALG00010012084 | TMEM50B | TMEM50B | 3510 | 3516 | 2174 | 2541 | 2300 | 2044 | 2345 | 2741 | 0.00875439 | 0.052513769 |
| ENSGALG00010012091 | LOC121108229 | LOC121108229 | 795 | 98 | 142 | 131 | 112 | 61 | 112 | 130 | 0.001108957 | 0.010166153 |
| ENSGALG00010012096 | LOC422320 | LOC422320 | 3298 | 2395 | 1361 | 1138 | 1999 | 1704 | 2234 | 1481 | 7.62E-05 | 0.001097257 |
| ENSGALG00010012099 | UPK3A | UPK3A | 1 | 7 | 24 | 68 | 21 | 40 | 91 | 51 | 1.83E-05 | 0.000326085 |
| ENSGALG00010012101 | GPSM2 | GPSM2 | 1981 | 2472 | 1371 | 1555 | 1624 | 1315 | 2499 | 2087 | 0.006031762 | 0.038904984 |
| ENSGALG00010012104 | SMC1B | SMC1L2 | 48 | 75 | 565 | 373 | 281 | 215 | 260 | 144 | 1.69E-08 | 6.79E-07 |
| ENSGALG00010012118 | LOC121108231 | LOC121108231 | 32 | 4 | 3 | 3 | 31 | 24 | 3 | 4 | 0.000146399 | 0.001911423 |
| ENSGALG00010012120 | RBM44 | RBM44 | 214 | 287 | 613 | 588 | 489 | 371 | 437 | 414 | 0.002452629 | 0.01907321 |
| ENSGALG00010012123 | RASL11AL | RASL11AL | 1 | 5 | 37 | 23 | 16 | 13 | 22 | 54 | 0.000120387 | 0.001615996 |
| ENSGALG00010012136 | NA | NA | 6 | 8 | 58 | 78 | 33 | 8 | 14 | 20 | 1.42E-05 | 0.000264002 |

| | | | | | | | | | | | | |
|---|---|---|---|---|---|---|---|---|---|---|---|---|
| ENSGALG00010012156 | NA | NA | 3 | 515 | 0 | 0 | 1 | 62 | 1 | 17 | 4.43E-06 | 9.72E-05 |
| ENSGALG00010012178 | NA | NA | 5 | 6 | 57 | 34 | 51 | 13 | 9 | 9 | 3.09E-05 | 0.000512247 |
| ENSGALG00010012179 | NA | NA | 12 | 12 | 9 | 7 | 3 | 0 | 18 | 20 | 0.00224781 | 0.0178204 |
| ENSGALG00010012236 | PRLH | PrRP | 171 | 38 | 3 | 5 | 6 | 15 | 134 | 104 | 3.70E-10 | 2.06E-08 |
| ENSGALG00010012259 | IL1RAPL2 | IL1RAPL2 | 58 | 200 | 550 | 633 | 254 | 289 | 400 | 411 | 0.000105843 | 0.001447763 |
| ENSGALG00010012260 | LOC121108245 | LOC121108245 | 70 | 93 | 338 | 288 | 195 | 137 | 134 | 235 | 0.000473048 | 0.005054721 |
| ENSGALG00010012274 | B4GALNT3 | B4GALNT3 | 189 | 179 | 79 | 49 | 92 | 132 | 341 | 132 | 0.00558381 | 0.036811297 |
| ENSGALG00010012277 | HIST2H3A | HIST2H3 | 90 | 246 | 69 | 41 | 31 | 33 | 131 | 201 | 7.54E-05 | 0.001088263 |
| ENSGALG00010012283 | LOC101751171 | LOC101751171 | 258 | 26 | 2 | 4 | 11 | 25 | 15 | 13 | 1.16E-08 | 4.80E-07 |
| ENSGALG00010012293 | SRGAP1 | SRGAP1 | 1800 | 1849 | 3559 | 3023 | 2641 | 2025 | 2434 | 1908 | 0.002316614 | 0.018218946 |
| ENSGALG00010012301 | NA | NA | 343 | 371 | 55 | 102 | 96 | 139 | 154 | 260 | 2.89E-05 | 0.000483794 |
| ENSGALG00010012304 | ESRP1 | RBM35A | 867 | 499 | 133 | 111 | 251 | 367 | 770 | 370 | 1.29E-06 | 3.28E-05 |
| ENSGALG00010012306 | LOC121108207 | LOC121108207 | 270 | 160 | 9 | 50 | 11 | 47 | 224 | 101 | 8.57E-05 | 0.001215169 |
| ENSGALG00010012325 | MLPH | MLPH | 45 | 13 | 17 | 1 | 4 | 2 | 75 | 56 | 2.78E-05 | 0.000469087 |
| ENSGALG00010012337 | ACSS3 | ACSS3 | 159 | 184 | 495 | 505 | 372 | 252 | 240 | 190 | 6.24E-05 | 0.000923401 |
| ENSGALG00010012344 | NA | NA | 8 | 6 | 35 | 60 | 77 | 60 | 47 | 69 | 5.07E-07 | 1.42E-05 |
| ENSGALG00010012373 | WNT7B | WNT7B | 8 | 48 | 10 | 13 | 0 | 2 | 0 | 3 | 5.43E-05 | 0.00082322 |
| ENSGALG00010012376 | VGLL2 | VGLL2 | 46 | 9 | 2 | 5 | 123 | 52 | 27 | 13 | 1.54E-07 | 4.98E-06 |
| ENSGALG00010012384 | NRK | MAP4K4 | 136 | 262 | 771 | 490 | 481 | 479 | 654 | 584 | 8.00E-05 | 0.001150001 |
| ENSGALG00010012401 | RIPK2 | RIPK2 | 370 | 231 | 571 | 868 | 570 | 377 | 455 | 391 | 0.00931098 | 0.055088806 |
| ENSGALG00010012410 | CACNA1I | CACNA1I | 3 | 8 | 8 | 2 | 8 | 8 | 28 | 28 | 0.001775427 | 0.01473966 |
| ENSGALG00010012412 | PURA | PURB | 69 | 110 | 321 | 231 | 218 | 116 | 157 | 161 | 0.005487151 | 0.036302958 |
| ENSGALG00010012425 | CHRNB3 | NACHR | 28 | 22 | 0 | 5 | 53 | 11 | 50 | 8 | 0.002311639 | 0.018188376 |
| ENSGALG00010012429 | CXCR7 | ACKR3 | 38 | 93 | 337 | 268 | 323 | 130 | 110 | 282 | 0.001116645 | 0.010208624 |
| ENSGALG00010012431 | LAMP2 | LAMP2 | 1028 | 1511 | 2570 | 2328 | 2198 | 1681 | 1874 | 2060 | 0.001075762 | 0.00991077 |
| ENSGALG00010012454 | IQCA1 | IQCA | 2 | 6 | 28 | 29 | 33 | 6 | 4 | 10 | 0.003013986 | 0.022536792 |
| ENSGALG00010012456 | LOC100859468 | SLAINL | 900 | 887 | 1572 | 1736 | 1016 | 800 | 761 | 746 | 8.05E-05 | 0.001154494 |
| ENSGALG00010012473 | NA | NA | 13 | 27 | 207 | 136 | 112 | 43 | 185 | 316 | 9.04E-09 | 3.86E-07 |
| ENSGALG00010012476 | NA | NA | 6 | 18 | 133 | 95 | 91 | 47 | 99 | 93 | 8.33E-07 | 2.20E-05 |
| ENSGALG00010012477 | LOC430496 | PCDHB15 | 20 | 30 | 188 | 212 | 84 | 64 | 38 | 29 | 1.47E-10 | 8.79E-09 |
| ENSGALG00010012478 | NA | NA | 34 | 35 | 0 | 0 | 2 | 0 | 218 | 161 | 2.38E-34 | 1.17E-31 |
| ENSGALG00010012479 | NA | NA | 13 | 1 | 0 | 1 | 11 | 15 | 9 | 16 | 0.002502974 | 0.019374572 |
| ENSGALG00010012480 | NA | NA | 9 | 208 | 625 | 306 | 443 | 229 | 75 | 113 | 0.005174167 | 0.034615966 |
| ENSGALG00010012483 | SLC7A7 | SLC7AL | 42 | 40 | 121 | 177 | 117 | 76 | 76 | 75 | 0.001159282 | 0.010495085 |
| ENSGALG00010012489 | GBX2 | GBX1 | 228 | 475 | 192 | 130 | 141 | 62 | 146 | 300 | 0.009397908 | 0.055477868 |
| ENSGALG00010012508 | MDM1 | MDM1 | 977 | 1132 | 2226 | 2153 | 1554 | 1050 | 1306 | 1197 | 0.000156302 | 0.002022923 |
| ENSGALG00010012530 | NA | NA | 6 | 3 | 24 | 29 | 18 | 13 | 11 | 5 | 0.003056269 | 0.022801951 |
| ENSGALG00010012537 | NA | NA | 0 | 1 | 4 | 34 | 13 | 4 | 1 | 6 | 0.002723158 | 0.020761597 |
| ENSGALG00010012555 | BHMT2 | BHMT | 1016 | 543 | 235 | 275 | 237 | 181 | 370 | 189 | 6.88E-06 | 0.000142112 |
| ENSGALG00010012582 | CD24 | CD24 | 16928 | 15786 | 6237 | 10188 | 7665 | 6489 | 10036 | 12047 | 1.87E-06 | 4.58E-05 |
| ENSGALG00010012633 | USP41 | USP18 | 144 | 264 | 388 | 412 | 319 | 245 | 437 | 772 | 0.00134515 | 0.011831654 |
| ENSGALG00010012650 | DSE | DSE | 949 | 805 | 387 | 542 | 316 | 383 | 658 | 488 | 0.003958977 | 0.028094445 |
| ENSGALG00010012654 | FAM46B | FAM46B | 644 | 792 | 304 | 277 | 206 | 145 | 377 | 349 | 1.71E-07 | 5.46E-06 |
| ENSGALG00010012655 | NA | NA | 33 | 0 | 0 | 0 | 14 | 4 | 88 | 0 | 0.002740815 | 0.020848654 |
| ENSGALG00010012662 | SLC35E3 | SLC35E3 | 5707 | 3017 | 2200 | 2925 | 3947 | 3731 | 5545 | 5758 | 0.000310347 | 0.003584327 |
| ENSGALG00010012670 | NA | NA | 830 | 480 | 288 | 290 | 436 | 446 | 862 | 470 | 0.005862462 | 0.038151661 |
| ENSGALG00010012677 | NA | NA | 78 | 411 | 83 | 56 | 61 | 30 | 306 | 320 | 1.46E-05 | 0.00027079 |
| ENSGALG00010012683 | MDM2 | MDM2 | 2783 | 2283 | 2265 | 2457 | 2104 | 1816 | 3942 | 3269 | 0.009138839 | 0.054371051 |
| ENSGALG00010012702 | PTPRO | PTPRO | 274 | 295 | 17 | 9 | 57 | 27 | 118 | 100 | 6.98E-17 | 1.01E-14 |
| ENSGALG00010012720 | ACO2 | ACO2 | 3112 | 4150 | 5759 | 6458 | 5082 | 3458 | 4545 | 5326 | 0.008048049 | 0.049141026 |
| ENSGALG00010012733 | NA | NA | 2 | 23 | 0 | 0 | 1 | 11 | 0 | 1 | 0.000715341 | 0.007135592 |
| ENSGALG00010012744 | ARL4C | ARL4C | 2212 | 2523 | 1329 | 1620 | 2546 | 1970 | 3544 | 3085 | 1.31E-07 | 4.37E-06 |
| ENSGALG00010012749 | RGS4 | RGS4 | 118 | 308 | 408 | 497 | 381 | 150 | 137 | 151 | 0.005687913 | 0.037300012 |
| ENSGALG00010012754 | NA | NA | 16 | 33 | 325 | 199 | 195 | 165 | 92 | 80 | 1.82E-10 | 1.07E-08 |
| ENSGALG00010012755 | CLVS2 | RLBP1L2 | 197 | 135 | 401 | 368 | 150 | 85 | 236 | 182 | 0.002290332 | 0.018097411 |
| ENSGALG00010012759 | ATP1B4 | ATP1B4 | 28 | 32 | 18 | 17 | 54 | 58 | 44 | 54 | 0.001621748 | 0.013722865 |
| ENSGALG00010012760 | PLEKHF2 | PLEKHF2 | 547 | 543 | 870 | 987 | 832 | 554 | 558 | 513 | 0.009714598 | 0.056945057 |
| ENSGALG00010012761 | SMPDL3A | SMPDL3A | 344 | 421 | 308 | 367 | 339 | 184 | 624 | 664 | 0.003956453 | 0.02808846 |
| ENSGALG00010012769 | NEMP2 | TMEM194B | 1768 | 2294 | 3293 | 3729 | 2755 | 1965 | 2724 | 2057 | 0.007643928 | 0.04703436 |
| ENSGALG00010012780 | FABP7 | FABP7 | 2120 | 3698 | 2594 | 3099 | 390 | 256 | 2910 | 2923 | 2.31E-29 | 9.18E-27 |
| ENSGALG00010012801 | DMC1 | DMC1 | 81 | 104 | 65 | 85 | 85 | 127 | 282 | 264 | 0.000109258 | 0.001483286 |
| ENSGALG00010012812 | DECR1 | DECR1 | 450 | 826 | 1368 | 1220 | 666 | 447 | 843 | 821 | 0.00282941 | 0.021385966 |
| ENSGALG00010012821 | ZBTB33 | ZBTB33 | 460 | 584 | 1228 | 974 | 907 | 582 | 696 | 573 | 0.001993041 | 0.016145277 |
| ENSGALG00010012832 | CHADL | CHADL | 255 | 20 | 242 | 262 | 22 | 18 | 262 | 63 | 0.003887184 | 0.027643671 |
| ENSGALG00010012839 | TGFBI | RGD-CAP | 5932 | 7422 | 2503 | 3763 | 2003 | 2158 | 5728 | 6528 | 6.69E-12 | 5.18E-10 |
| ENSGALG00010012842 | DDX17 | Cp68 | 2577 | 3743 | 5107 | 4865 | 4473 | 3482 | 5547 | 4807 | 0.004853443 | 0.033038995 |
| ENSGALG00010012867 | A4GALT | A4GALT | 1198 | 1374 | 2175 | 2802 | 1642 | 1324 | 1114 | 1309 | 4.69E-05 | 0.000729982 |
| ENSGALG00010012874 | CETN1L | CETN1L | 52 | 27 | 109 | 147 | 106 | 97 | 73 | 90 | 0.006921304 | 0.043364235 |
| ENSGALG00010012876 | KDELR3 | KDELR3 | 739 | 1053 | 133 | 155 | 207 | 424 | 541 | 843 | 1.73E-08 | 6.90E-07 |
| ENSGALG00010012877 | SLC25A32 | SLC25A32 | 930 | 700 | 1597 | 1704 | 1310 | 819 | 1018 | 924 | 0.002476975 | 0.019200003 |
| ENSGALG00010012883 | DCAF13 | WDSOF1 | 6763 | 7637 | 7159 | 9274 | 5179 | 4248 | 4977 | 5105 | 0.001561321 | 0.013292307 |
| ENSGALG00010012891 | CAPS2 | CAPS2 | 69 | 33 | 21 | 14 | 43 | 51 | 22 | 5 | 0.002669423 | 0.020435823 |
| ENSGALG00010012894 | CLIC6 | CLIC6 | 657 | 758 | 188 | 240 | 178 | 144 | 498 | 358 | 3.22E-09 | 1.53E-07 |
| ENSGALG00010012905 | NKAIN1 | FAM77C | 1691 | 3902 | 1616 | 1074 | 1596 | 824 | 3243 | 4569 | 1.94E-06 | 4.73E-05 |
| ENSGALG00010012907 | GLIPR1L1 | GLIPR1L2 | 13 | 37 | 82 | 76 | 88 | 62 | 48 | 54 | 0.006261333 | 0.040061027 |
| ENSGALG00010012910 | XPNPEP3 | XPNPEP3 | 978 | 1231 | 1691 | 1781 | 2396 | 1583 | 1312 | 1155 | 5.91E-06 | 0.000124325 |
| ENSGALG00010012912 | NKAPL | NKAP | 2449 | 2489 | 2563 | 3084 | 2355 | 1550 | 1578 | 1770 | 0.008983752 | 0.053601024 |
| ENSGALG00010012915 | SELP | SELP | 1 | 2 | 10 | 31 | 8 | 5 | 13 | 10 | 0.002371652 | 0.018555737 |
| ENSGALG00010012939 | GJA1 | CX43 | 3909 | 2427 | 1596 | 1205 | 1422 | 1008 | 3107 | 2123 | 7.54E-06 | 0.000153338 |
| ENSGALG00010012956 | FABP3 | FABP | 459 | 573 | 371 | 679 | 242 | 232 | 207 | 249 | 0.001859132 | 0.01524525 |
| ENSGALG00010012957 | DERA | DERA | 728 | 651 | 887 | 871 | 630 | 509 | 387 | 465 | 0.006879185 | 0.043169771 |
| ENSGALG00010012958 | OLFML2B | OLFML2B | 269 | 242 | 83 | 86 | 76 | 80 | 257 | 235 | 9.73E-06 | 0.000191556 |
| ENSGALG00010012967 | C2orf40 | C1H2ORF40 | 93 | 74 | 24 | 28 | 43 | 43 | 54 | 66 | 0.008124281 | 0.049425912 |
| ENSGALG00010012970 | UNC5A | UNC5A | 614 | 978 | 386 | 308 | 463 | 276 | 1086 | 840 | 2.90E-06 | 6.67E-05 |
| ENSGALG00010012979 | HTR1A | HTR1A | 400 | 170 | 133 | 85 | 114 | 33 | 146 | 71 | 0.005369706 | 0.035681325 |
| ENSGALG00010012985 | ARNTL2 | ARNTL2 | 152 | 139 | 383 | 369 | 349 | 239 | 312 | 328 | 0.001876493 | 0.015349952 |
| ENSGALG00010012998 | MAFF | MAFF | 114 | 159 | 244 | 125 | 183 | 229 | 358 | 499 | 0.001957947 | 0.015907252 |
| ENSGALG00010013001 | ADGRV1 | GPR98 | 1 | 10 | 87 | 51 | 147 | 11 | 67 | 49 | 0.000622569 | 0.006359644 |
| ENSGALG00010013018 | ID1 | ID1 | 532 | 632 | 127 | 157 | 262 | 175 | 465 | 784 | 1.27E-09 | 6.49E-08 |
| ENSGALG00010013040 | SLC25A43 | SLC25A43 | 380 | 426 | 645 | 699 | 754 | 625 | 814 | 643 | 0.00383671 | 0.027377926 |
| ENSGALG00010013044 | NOS1AP | NOS1AP | 3284 | 5663 | 394 | 487 | 933 | 1051 | 2937 | 4575 | 8.78E-31 | 3.97E-28 |
| ENSGALG00010013056 | SPATA46 | C1orf111 | 261 | 526 | 36 | 49 | 111 | 95 | 282 | 487 | 2.13E-12 | 1.77E-10 |
| ENSGALG00010013078 | COL5A2 | COL5A2 | 419 | 835 | 1243 | 1915 | 1127 | 892 | 900 | 1128 | 0.001057325 | 0.009767832 |
| ENSGALG00010013086 | CXCL14 | JSC | 34 | 114 | 25 | 14 | 184 | 433 | 44 | 69 | 6.91E-11 | 4.46E-09 |
| ENSGALG00010013113 | NA | NA | 256 | 46 | 11 | 14 | 11 | 49 | 6 | 33 | 9.58E-05 | 0.001337101 |
| ENSGALG00010013126 | NA | NA | 2106 | 1127 | 528 | 1227 | 83 | 110 | 1367 | 1130 | 1.28E-15 | 1.64E-13 |
| ENSGALG00010013128 | NA | NA | 54 | 199 | 7 | 29 | 9 | 34 | 43 | 40 | 0.000833324 | 0.008068661 |
| ENSGALG00010013129 | RASSF3 | RASSF3 | 2976 | 2955 | 5285 | 5938 | 4520 | 3048 | 3741 | 3215 | 5.14E-05 | 0.000787565 |
| ENSGALG00010013137 | TMEM121 | TMEM121 | 836 | 2743 | 748 | 684 | 438 | 549 | 1446 | 1926 | 3.96E-05 | 0.000632582 |
| ENSGALG00010013156 | ABCC9 | ABCC9 | 2 | 1 | 25 | 85 | 60 | 12 | 4 | 2 | 6.52E-09 | 2.91E-07 |
| ENSGALG00010013158 | CRIP1 | CRIP1 | 52 | 90 | 3 | 10 | 14 | 20 | 85 | 87 | 7.01E-09 | 3.10E-07 |
| ENSGALG00010013164 | FAM83F | FAM83F | 905 | 118 | 130 | 102 | 331 | 239 | 1162 | 484 | 0.0005656 | 0.005870994 |
| ENSGALG00010013170 | COL3A1 | collagen | 2192 | 4435 | 8053 | 8106 | 7382 | 6258 | 10061 | 7363 | 4.74E-07 | 1.35E-05 |
| ENSGALG00010013172 | TNFSF13B | TNFSF13B | 9 | 4 | 3 | 0 | 18 | 20 | 10 | 10 | 0.000415493 | 0.00455625 |
| ENSGALG00010013173 | KCNJ8 | Kir6.1 | 89 | 98 | 138 | 157 | 177 | 143 | 57 | 51 | 0.000258655 | 0.003096451 |
| ENSGALG00010013177 | OXR1 | OXR1 | 802 | 904 | 1439 | 2054 | 1388 | 929 | 875 | 844 | 5.25E-05 | 0.000800104 |
| ENSGALG00010013187 | KDF1 | C1orf172 | 1180 | 875 | 387 | 441 | 598 | 513 | 919 | 805 | 3.41E-05 | 0.000557544 |
| ENSGALG00010013213 | SH2D1A | SH2D1A | 91 | 205 | 124 | 57 | 144 | 155 | 248 | 369 | 0.006459838 | 0.041085354 |
| ENSGALG00010013214 | RAI14 | RAI14 | 482 | 510 | 1217 | 974 | 1095 | 755 | 942 | 723 | 0.000322115 | 0.003707433 |
| ENSGALG00010013226 | DUSP6 | DUSP6 | 4100 | 3720 | 2805 | 2655 | 2687 | 1762 | 3660 | 4186 | 0.001583785 | 0.013449299 |
| ENSGALG00010013227 | CITED4 | CITED3 | 49 | 84 | 164 | 235 | 167 | 105 | 188 | 421 | 0.000266338 | 0.003177042 |
| ENSGALG00010013243 | NA | NA | 75 | 63 | 81 | 105 | 30 | 62 | 306 | 145 | 0.000831071 | 0.008056181 |
| ENSGALG00010013245 | NA | NA | 64 | 79 | 4 | 10 | 4 | 4 | 40 | 53 | 3.39E-11 | 2.30E-09 |
| ENSGALG00010013252 | PPM1H | PPM1H | 1803 | 1075 | 331 | 462 | 360 | 492 | 1433 | 1077 | 9.12E-09 | 3.89E-07 |
| ENSGALG00010013285 | ATP2B1 | ATP2B1 | 11408 | 9196 | 5811 | 6116 | 6505 | 5988 | 11057 | 9154 | 9.22E-05 | 0.001295146 |
| ENSGALG00010013288 | CA3A | CA3 | 1 | 5 | 37 | 10 | 37 | 5 | 6 | 5 | 0.005798321 | 0.037822615 |
| ENSGALG00010013290 | AKR1D1 | AKR1D1 | 781 | 629 | 286 | 175 | 535 | 356 | 882 | 964 | 9.53E-08 | 3.27E-06 |
| ENSGALG00010013295 | CA3B | CA3 | 88 | 71 | 344 | 318 | 281 | 166 | 143 | 106 | 2.90E-06 | 6.67E-05 |
| ENSGALG00010013310 | MYO16 | MYR8 | 1897 | 1954 | 1270 | 988 | 1309 | 905 | 2222 | 1650 | 0.001609279 | 0.013631154 |
| ENSGALG00010013336 | MTNR1A | CKA | 94 | 100 | 206 | 380 | 33 | 38 | 90 | 70 | 9.11E-08 | 3.15E-06 |
| ENSGALG00010013344 | OSMR | OSMR | 14 | 11 | 104 | 45 | 42 | 8 | 95 | 55 | 0.000857627 | 0.008265661 |
| ENSGALG00010013355 | FAM171B | FAM171B | 2236 | 1906 | 1224 | 1235 | 1229 | 938 | 1908 | 1452 | 0.002473054 | 0.019187352 |
| ENSGALG00010013359 | BTG1 | BTG1 | 3047 | 3287 | 4069 | 5053 | 3205 | 2330 | 2058 | 2993 | 0.004980297 | 0.033655773 |
| ENSGALG00010013369 | MTMR8 | MTMR8 | 1426 | 1448 | 1468 | 1665 | 2192 | 1592 | 1355 | 959 | 0.000682516 | 0.006864656 |
| ENSGALG00010013372 | SPRY4 | SPRY4 | 202 | 327 | 138 | 105 | 91 | 64 | 203 | 300 | 0.000476425 | 0.005085361 |
| ENSGALG00010013373 | NA | NA | 10 | 37 | 0 | 6 | 3 | 1 | 9 | 6 | 0.0018524 | 0.015212433 |
| ENSGALG00010013380 | PHGDH | PHGDH | 1447 | 936 | 1261 | 599 | 858 | 1941 | 4460 | 6758 | 2.62E-09 | 1.26E-07 |
| ENSGALG00010013382 | ASB12 | ASB12 | 75 | 218 | 454 | 523 | 225 | 128 | 339 | 419 | 0.000587688 | 0.006055145 |
| ENSGALG00010013387 | NDST1 | NDST1 | 2932 | 3172 | 1777 | 2264 | 2090 | 1396 | 2570 | 2219 | 0.006025205 | 0.038902902 |
| ENSGALG00010013388 | NA | NA | 217 | 213 | 470 | 480 | 423 | 279 | 449 | 407 | 0.009386262 | 0.055467898 |
| ENSGALG00010013390 | SLC16A2 | SLC16A2 | 683 | 1078 | 838 | 1051 | 365 | 304 | 591 | 726 | 0.000382392 | 0.004280651 |

| | | | | | | | | | | | | |
|---|---|---|---|---|---|---|---|---|---|---|---|---|
| ENSGALG00010013396 | FYB1 | FYB | 2 | 11 | 44 | 47 | 68 | 12 | 19 | 15 | 0.001371185 | 0.012011896 |
| ENSGALG00010013399 | FGF1 | ECGF | 12 | 11 | 2 | 3 | 2 | 5 | 8 | 29 | 0.008386905 | 0.05067316 |
| ENSGALG00010013403 | CPNE3 | CPNE3 | 1110 | 656 | 674 | 1048 | 1198 | 1130 | 1634 | 1500 | 0.005171846 | 0.034614293 |
| ENSGALG00010013419 | NA | NA | 95 | 55 | 25 | 13 | 54 | 73 | 21 | 12 | 3.11E-05 | 0.000515082 |
| ENSGALG00010013435 | SDC3 | syndecan-3 | 1159 | 1182 | 136 | 171 | 162 | 150 | 618 | 677 | 7.70E-27 | 2.53E-24 |
| ENSGALG00010013452 | LSS | LSS | 2505 | 2774 | 2084 | 2340 | 2951 | 2496 | 2911 | 3601 | 0.00847178 | 0.05113051 |
| ENSGALG00010013458 | LAMA4 | LAMA4 | 2 | 4 | 5 | 73 | 12 | 13 | 8 | 11 | 0.004715808 | 0.032402319 |
| ENSGALG00010013492 | B3GALT5 | B3GALT5 | 275 | 340 | 21 | 26 | 141 | 159 | 262 | 170 | 7.72E-13 | 6.72E-11 |
| ENSGALG00010013510 | MRPS33 | MRPS33 | 1237 | 1478 | 2233 | 2870 | 2095 | 1502 | 1308 | 1395 | 5.07E-05 | 0.000779218 |
| ENSGALG00010013515 | RBP4 | RBP4B | 16 | 11 | 195 | 71 | 24 | 9 | 28 | 25 | 7.88E-08 | 2.77E-06 |
| ENSGALG00010013525 | RFX8 | RFX8 | 7 | 6 | 5 | 14 | 18 | 40 | 17 | 5 | 0.006587374 | 0.041763353 |
| ENSGALG00010013549 | PCP4 | PCP4 | 1021 | 933 | 67 | 77 | 300 | 248 | 1072 | 410 | 3.55E-17 | 5.25E-15 |
| ENSGALG00010013572 | DENND11 | KIAA1147 | 1988 | 1634 | 677 | 1040 | 1045 | 1182 | 2134 | 1668 | 0.000111871 | 0.001516286 |
| ENSGALG00010013574 | NR2F1 | COUP-TFI | 11 | 8 | 57 | 30 | 30 | 12 | 16 | 10 | 0.005898298 | 0.03832518 |
| ENSGALG00010013580 | RANBP3L | RANBP3L | 75 | 89 | 818 | 584 | 332 | 64 | 118 | 78 | 1.97E-09 | 9.79E-08 |
| ENSGALG00010013589 | DSCAM | DSCAM | 141 | 62 | 30 | 8 | 29 | 22 | 224 | 66 | 7.35E-05 | 0.001065018 |
| ENSGALG00010013592 | HIST2H3A | HIST2H3 | 2 | 0 | 9 | 9 | 0 | 1 | 24 | 29 | 3.09E-08 | 1.17E-06 |
| ENSGALG00010013605 | NA | NA | 0 | 0 | 0 | 0 | 0 | 0 | 25 | 27 | 2.20E-14 | 2.36E-12 |
| ENSGALG00010013606 | NA | NA | 0 | 0 | 0 | 0 | 0 | 0 | 1097 | 1151 | 3.93E-124 | 3.28E-120 |
| ENSGALG00010013624 | SST1 | PSS1 | 7 | 38 | 38 | 28 | 13 | 13 | 79 | 171 | 1.10E-05 | 0.000212047 |
| ENSGALG00010013632 | SLC1A3 | SLC1A3 | 0 | 1 | 142 | 151 | 18 | 12 | 55 | 100 | 6.00E-20 | 1.22E-17 |
| ENSGALG00010013640 | SNPH | SNPH | 124 | 107 | 19 | 34 | 55 | 55 | 100 | 69 | 0.000375023 | 0.004218254 |
| ENSGALG00010013661 | GABRA4 | GABRA4 | 29 | 78 | 19 | 20 | 13 | 13 | 17 | 18 | 0.005951669 | 0.038552054 |
| ENSGALG00010013667 | SOWAHA | ANKRD43 | 246 | 344 | 93 | 90 | 150 | 92 | 276 | 224 | 5.81E-05 | 0.00086854 |
| ENSGALG00010013672 | SHROOM1 | SHROOM1 | 421 | 290 | 50 | 66 | 83 | 93 | 280 | 176 | 1.81E-08 | 7.20E-07 |
| ENSGALG00010013687 | NA | NA | 0 | 0 | 4 | 1 | 704 | 522 | 644 | 806 | 1.44E-91 | 4.82E-88 |
| ENSGALG00010013715 | IL1R1 | IL1R1 | 138 | 130 | 993 | 425 | 282 | 91 | 152 | 129 | 1.78E-07 | 5.66E-06 |
| ENSGALG00010013729 | NA | NA | 40 | 34 | 11 | 3 | 1 | 17 | 59 | 28 | 0.00803609 | 0.049085939 |
| ENSGALG00010013744 | IL1RL2 | IL1RL2 | 54 | 71 | 434 | 334 | 238 | 148 | 151 | 118 | 1.58E-08 | 6.42E-07 |
| ENSGALG00010013749 | NELL2 | nel | 2025 | 3034 | 607 | 1126 | 1266 | 1385 | 1820 | 1509 | 3.21E-06 | 7.32E-05 |
| ENSGALG00010013751 | FUT10 | FUT10 | 257 | 90 | 37 | 22 | 82 | 25 | 160 | 92 | 0.000302003 | 0.003515672 |
| ENSGALG00010013753 | PLEKHA5 | PLEKHA5 | 671 | 1035 | 960 | 1494 | 876 | 667 | 630 | 560 | 0.009567885 | 0.056263082 |
| ENSGALG00010013774 | NA | NA | 11 | 7 | 7 | 5 | 36 | 39 | 11 | 9 | 6.92E-06 | 0.000142776 |
| ENSGALG00010013828 | PTN | HB-GAM | 79 | 227 | 349 | 248 | 248 | 119 | 363 | 688 | 0.002457441 | 0.019101745 |
| ENSGALG00010013833 | TMPRSS2 | TMPRSS2 | 96 | 54 | 10 | 12 | 33 | 18 | 56 | 30 | 8.93E-05 | 0.001259324 |
| ENSGALG00010013837 | IFI27L2 | IFI27L1 | 49 | 55 | 26 | 21 | 17 | 6 | 39 | 29 | 0.007159665 | 0.044530822 |
| ENSGALG00010013859 | MYOC | MYOC | 238 | 252 | 460 | 510 | 369 | 255 | 244 | 237 | 0.009574771 | 0.056283355 |
| ENSGALG00010013860 | NA | NA | 15 | 14 | 0 | 2 | 3 | 10 | 25 | 10 | 0.001790937 | 0.014828291 |
| ENSGALG00010013866 | KCTD8 | KCTD8 | 1060 | 1584 | 295 | 586 | 305 | 246 | 426 | 317 | 6.68E-11 | 4.32E-09 |
| ENSGALG00010013869 | NA | NA | 20 | 20 | 1 | 5 | 17 | 23 | 34 | 23 | 0.000286732 | 0.003365073 |
| ENSGALG00010013874 | NA | NA | 59 | 61 | 8 | 13 | 27 | 39 | 56 | 24 | 0.001205849 | 0.010853528 |
| ENSGALG00010013944 | METTL24 | C3H6ORF186 | 862 | 1101 | 416 | 577 | 719 | 444 | 593 | 769 | 0.004259768 | 0.029773839 |
| ENSGALG00010013959 | STK32A | STK32A | 89 | 250 | 68 | 71 | 43 | 70 | 212 | 345 | 0.000165378 | 0.002125996 |
| ENSGALG00010013964 | SLC38A1 | SLC38A1 | 2290 | 2353 | 1758 | 1551 | 1801 | 1776 | 3588 | 2424 | 0.004588657 | 0.03162294 |
| ENSGALG00010014003 | NA | NA | 0 | 0 | 0 | 0 | 4 | 1 | 74 | 35 | 4.36E-18 | 7.36E-16 |
| ENSGALG00010014019 | IRAG2 | LRMP | 1 | 1 | 71 | 6 | 31 | 1 | 6 | 4 | 0.000873483 | 0.008366362 |
| ENSGALG00010014020 | PDXP | PDXP | 1948 | 2334 | 2371 | 2962 | 2647 | 1586 | 1522 | 1574 | 0.004975333 | 0.033649441 |
| ENSGALG00010014029 | NA | NA | 132 | 141 | 355 | 368 | 347 | 231 | 388 | 207 | 0.004769631 | 0.032641306 |
| ENSGALG00010014038 | NA | NA | 466 | 389 | 433 | 435 | 764 | 581 | 552 | 462 | 0.005098275 | 0.034190358 |
| ENSGALG00010014051 | NA | NA | 119 | 391 | 318 | 316 | 93 | 95 | 103 | 74 | 0.000761474 | 0.007497111 |
| ENSGALG00010014054 | LOC419276 | BPIL | 23 | 27 | 2 | 0 | 18 | 21 | 35 | 40 | 9.14E-08 | 3.15E-06 |
| ENSGALG00010014058 | NA | NA | 81 | 76 | 240 | 255 | 229 | 209 | 198 | 154 | 0.000219214 | 0.002693759 |
| ENSGALG00010014096 | JAKMIP2 | JAKMIP2 | 887 | 909 | 445 | 428 | 421 | 376 | 979 | 761 | 3.87E-05 | 0.000621034 |
| ENSGALG00010014102 | TRIM2 | TRIM2 | 26 | 53 | 607 | 660 | 749 | 123 | 138 | 62 | 1.23E-10 | 7.44E-09 |
| ENSGALG00010014108 | LOC100859898 | LOC100859898 | 9 | 3 | 95 | 67 | 68 | 37 | 18 | 20 | 2.31E-08 | 8.96E-07 |
| ENSGALG00010014122 | ABCA1 | ABCA1 | 634 | 1281 | 129 | 238 | 687 | 567 | 752 | 337 | 3.52E-07 | 1.03E-05 |
| ENSGALG00010014124 | AK9 | AKD1 | 197 | 76 | 41 | 41 | 67 | 72 | 61 | 22 | 0.00438222 | 0.030426113 |
| ENSGALG00010014125 | PLPP1 | PPAP2A | 6867 | 3486 | 3085 | 1870 | 2265 | 1271 | 4738 | 3590 | 9.60E-05 | 0.001337859 |
| ENSGALG00010014128 | COL4A1 | COL4A1 | 1564 | 2075 | 626 | 747 | 779 | 764 | 2379 | 1414 | 2.64E-08 | 1.01E-06 |
| ENSGALG00010014145 | METTL11B | C1orf184 | 76 | 114 | 155 | 167 | 188 | 233 | 266 | 215 | 0.003983203 | 0.028230402 |
| ENSGALG00010014147 | HTR1D | 5-HT-1D | 215 | 213 | 532 | 511 | 599 | 559 | 490 | 356 | 1.76E-05 | 0.000316866 |
| ENSGALG00010014148 | KCNG1 | KCNG1 | 1027 | 1619 | 962 | 1302 | 2846 | 2151 | 1871 | 2024 | 5.59E-09 | 2.54E-07 |
| ENSGALG00010014157 | NA | NA | 204 | 33 | 9 | 25 | 6 | 11 | 30 | 68 | 3.64E-05 | 0.00058842 |
| ENSGALG00010014161 | GFRA3 | GFRA3 | 677 | 633 | 155 | 233 | 643 | 1117 | 1202 | 582 | 5.19E-08 | 1.87E-06 |
| ENSGALG00010014162 | COL18A1 | COL18A1 | 3684 | 4057 | 2832 | 2756 | 2558 | 2136 | 4224 | 4140 | 0.001108775 | 0.010166153 |
| ENSGALG00010014193 | CARD10 | CARD10 | 57 | 20 | 3 | 8 | 22 | 13 | 59 | 32 | 0.000358085 | 0.004048934 |
| ENSGALG00010014197 | SH3GL2 | SH3P4 | 242 | 116 | 676 | 912 | 764 | 353 | 402 | 323 | 4.70E-06 | 0.000102405 |
| ENSGALG00010014211 | NA | NA | 13 | 14 | 2 | 2 | 4 | 3 | 14 | 10 | 0.003636222 | 0.026215935 |
| ENSGALG00010014226 | SLC16A7 | SLC16A7 | 134 | 129 | 22 | 24 | 30 | 25 | 57 | 35 | 4.98E-07 | 1.41E-05 |
| ENSGALG00010014238 | PHLDA1 | PHLDA1 | 232 | 304 | 451 | 429 | 224 | 191 | 472 | 647 | 0.00226469 | 0.017945706 |
| ENSGALG00010014268 | NA | NA | 2 | 6 | 1 | 4 | 14 | 11 | 14 | 14 | 0.003494702 | 0.025403854 |
| ENSGALG00010014270 | NA | NA | 1699 | 2050 | 347 | 475 | 1310 | 1202 | 965 | 585 | 7.84E-15 | 8.86E-13 |
| ENSGALG00010014274 | C1orf112 | C8H1ORF112 | 633 | 988 | 1116 | 1228 | 1067 | 859 | 519 | 532 | 8.65E-05 | 0.00122432 |
| ENSGALG00010014282 | NA | NA | 17 | 16 | 0 | 0 | 0 | 0 | 0 | 0 | 2.70E-10 | 1.54E-08 |
| ENSGALG00010014283 | NA | NA | 123 | 153 | 0 | 0 | 0 | 0 | 0 | 1 | 9.41E-40 | 6.05E-37 |
| ENSGALG00010014295 | CACNA1C | CHCACHA1C | 19 | 43 | 87 | 34 | 23 | 20 | 102 | 143 | 0.000437338 | 0.004736739 |
| ENSGALG00010014298 | METTL18 | MGC9084L | 1114 | 914 | 828 | 982 | 839 | 722 | 522 | 422 | 0.000321701 | 0.003705211 |
| ENSGALG00010014303 | GABRA2 | GABRA2 | 111 | 159 | 39 | 55 | 28 | 38 | 132 | 67 | 0.001385329 | 0.01211492 |
| ENSGALG00010014324 | COL4A2 | COL4A2 | 892 | 1609 | 511 | 539 | 501 | 521 | 1322 | 840 | 7.34E-05 | 0.001064378 |
| ENSGALG00010014334 | HMGCS1 | HMGCS1 | 3835 | 4225 | 2828 | 2803 | 3100 | 3430 | 4226 | 4479 | 0.00773774 | 0.047524115 |
| ENSGALG00010014338 | BHLHE41 | BHLHE41 | 17 | 31 | 55 | 47 | 101 | 70 | 22 | 23 | 2.19E-05 | 0.000380648 |
| ENSGALG00010014340 | EFNB1 | EFNB1 | 7806 | 6900 | 3369 | 3053 | 4209 | 3874 | 5767 | 7297 | 2.49E-08 | 9.60E-07 |
| ENSGALG00010014390 | SPARC | SPARC | 9831 | 4954 | 4699 | 6023 | 2708 | 2898 | 6744 | 11024 | 0.000144619 | 0.001891131 |
| ENSGALG00010014393 | ATP8A1 | ATP8A1 | 1334 | 2698 | 856 | 1526 | 1117 | 937 | 710 | 331 | 1.16E-05 | 0.000223044 |
| ENSGALG00010014399 | GAS6 | GAS6 | 449 | 278 | 165 | 261 | 185 | 158 | 504 | 405 | 0.00430636 | 0.030024152 |
| ENSGALG00010014425 | NA | NA | 13 | 31 | 80 | 148 | 85 | 82 | 36 | 28 | 1.32E-05 | 0.000248656 |
| ENSGALG00010014426 | NA | NA | 1 | 10 | 9 | 5 | 64 | 22 | 4 | 2 | 7.99E-07 | 2.12E-05 |
| ENSGALG00010014428 | NA | NA | 13 | 19 | 19 | 14 | 53 | 27 | 1 | 2 | 8.64E-08 | 3.01E-06 |
| ENSGALG00010014434 | NA | NA | 16 | 29 | 0 | 1 | 17 | 13 | 7 | 15 | 1.05E-05 | 0.000204279 |
| ENSGALG00010014492 | CPLX2 | CPLX2 | 627 | 836 | 339 | 334 | 398 | 251 | 1092 | 1493 | 2.12E-12 | 1.76E-10 |
| ENSGALG00010014493 | NA | NA | 20 | 20 | 104 | 76 | 98 | 50 | 51 | 37 | 0.000307637 | 0.003561468 |
| ENSGALG00010014507 | LOC101749281 | LOC101749281 | 18 | 32 | 36 | 37 | 57 | 58 | 94 | 59 | 0.007511991 | 0.0464105 |
| ENSGALG00010014536 | NA | NA | 49 | 52 | 18 | 14 | 17 | 7 | 32 | 20 | 0.003226399 | 0.023826596 |
| ENSGALG00010014547 | DHDDS | DHDDS | 1127 | 1444 | 821 | 993 | 678 | 552 | 807 | 847 | 0.006443107 | 0.041004267 |
| ENSGALG00010014601 | NA | NA | 95 | 129 | 5 | 67 | 18 | 35 | 482 | 263 | 6.79E-07 | 1.84E-05 |
| ENSGALG00010014603 | TSC22D4 | GILZ | 190 | 129 | 75 | 40 | 156 | 164 | 281 | 279 | 4.56E-05 | 0.000713917 |
| ENSGALG00010014609 | NA | NA | 7 | 4 | 9 | 39 | 24 | 22 | 10 | 3 | 0.007517919 | 0.04641989 |
| ENSGALG00010014617 | GRTP1 | GRTP1 | 1012 | 802 | 449 | 466 | 678 | 577 | 1093 | 936 | 0.000180207 | 0.00229021 |
| ENSGALG00010014632 | NWD2 | NWD2 | 54 | 59 | 850 | 1090 | 105 | 134 | 478 | 423 | 3.32E-26 | 1.03E-23 |
| ENSGALG00010014636 | NA | NA | 77 | 102 | 109 | 110 | 166 | 167 | 311 | 191 | 0.001514551 | 0.012960153 |
| ENSGALG00010014643 | CLDN16 | claudin-16 | 6 | 13 | 18 | 12 | 36 | 83 | 13 | 24 | 2.16E-05 | 0.00037716 |
| ENSGALG00010014656 | NA | NA | 78 | 78 | 22 | 10 | 25 | 16 | 38 | 28 | 0.000216689 | 0.002668611 |
| ENSGALG00010014657 | NA | NA | 3 | 7 | 28 | 28 | 16 | 6 | 4 | 4 | 0.00048689 | 0.005169575 |
| ENSGALG00010014666 | NA | NA | 46 | 55 | 79 | 102 | 175 | 150 | 238 | 109 | 0.000235553 | 0.002857628 |
| ENSGALG00010014680 | LOC107053920 | LOC107053920 | 1308 | 2155 | 848 | 1198 | 641 | 814 | 437 | 788 | 0.00013495 | 0.001778616 |
| ENSGALG00010014682 | IL13RA1 | IL13RA1 | 1379 | 1924 | 765 | 1220 | 1182 | 783 | 2503 | 2035 | 8.64E-06 | 0.000173277 |
| ENSGALG00010014688 | NA | NA | 8 | 8 | 51 | 83 | 18 | 3 | 2 | 2 | 8.42E-10 | 4.43E-08 |
| ENSGALG00010014690 | CSF3R | CSF3R | 7 | 8 | 25 | 19 | 9 | 14 | 48 | 33 | 0.00230176 | 0.018144818 |
| ENSGALG00010014698 | NA | NA | 1 | 5 | 33 | 35 | 34 | 12 | 4 | 10 | 3.91E-05 | 0.000625796 |
| ENSGALG00010014710 | NPEPL1 | NPEPL1 | 1947 | 2236 | 2494 | 3353 | 2506 | 1718 | 1704 | 1615 | 0.003881552 | 0.027626259 |
| ENSGALG00010014731 | NA | NA | 6 | 9 | 18 | 18 | 31 | 29 | 14 | 15 | 0.003875933 | 0.027598889 |
| ENSGALG00010014755 | LOC121108473 | LOC121108473 | 51 | 23 | 6 | 8 | 26 | 22 | 93 | 12 | 0.007799418 | 0.047867745 |
| ENSGALG00010014760 | NA | NA | 17 | 53 | 42 | 30 | 79 | 65 | 127 | 111 | 0.001059714 | 0.009784502 |
| ENSGALG00010014766 | F10 | FX | 266 | 122 | 34 | 66 | 56 | 72 | 100 | 102 | 0.001212465 | 0.010904237 |
| ENSGALG00010014772 | LOC101751179 | LOC101751179 | 0 | 4 | 4 | 0 | 5 | 2 | 12 | 24 | 0.003186166 | 0.023592031 |
| ENSGALG00010014780 | NA | NA | 65 | 28 | 21 | 29 | 37 | 22 | 105 | 74 | 0.0066154 | 0.041845784 |
| ENSGALG00010014782 | NA | NA | 142 | 180 | 88 | 397 | 429 | 421 | 36 | 185 | 0.004573579 | 0.031558076 |
| ENSGALG00010014785 | NA | NA | 270 | 472 | 43 | 693 | 619 | 662 | 9 | 126 | 0.006216711 | 0.039859472 |
| ENSGALG00010014797 | ZDHHC9 | ZDHHC9 | 5208 | 5711 | 3771 | 3573 | 3919 | 2784 | 4398 | 4645 | 0.004361791 | 0.03031132 |
| ENSGALG00010014824 | ANXA6 | ANXA6 | 36 | 32 | 40 | 19 | 11 | 6 | 99 | 171 | 6.99E-09 | 3.10E-07 |
| ENSGALG00010014826 | NA | NA | 16 | 10 | 72 | 66 | 63 | 24 | 74 | 57 | 0.000612472 | 0.006267995 |
| ENSGALG00010014845 | NA | NA | 60 | 39 | 13 | 18 | 17 | 16 | 33 | 38 | 0.008861233 | 0.053021454 |
| ENSGALG00010014847 | PAICS | AIRC | 2655 | 2388 | 1352 | 1410 | 1362 | 1062 | 2843 | 3005 | 1.09E-08 | 4.56E-07 |
| ENSGALG00010014860 | NA | NA | 5 | 8 | 36 | 17 | 70 | 24 | 46 | 31 | 0.000387648 | 0.00431892 |
| ENSGALG00010014867 | TP63 | TAp63alpha | 181 | 187 | 675 | 632 | 530 | 475 | 396 | 353 | 2.27E-07 | 7.02E-06 |
| ENSGALG00010014873 | FAM26F | CALHM6 | 12 | 20 | 3 | 4 | 0 | 2 | 4 | 4 | 0.000878444 | 0.008403292 |
| ENSGALG00010014902 | AASDH | AASDH | 645 | 622 | 389 | 355 | 369 | 329 | 957 | 642 | 0.000908324 | 0.008644595 |
| ENSGALG00010014903 | MCF2L | MCF2L | 96 | 111 | 26 | 39 | 60 | 72 | 106 | 74 | 0.004967532 | 0.033623323 |
| ENSGALG00010014905 | TEKT2 | tektin-2 | 48 | 260 | 49 | 39 | 25 | 33 | 82 | 119 | 0.003318652 | 0.024389316 |
| ENSGALG00010014935 | PCYT1A | PCYT1A | 790 | 1447 | 1654 | 1838 | 812 | 593 | 1212 | 1399 | 0.004616059 | 0.031785569 |
| ENSGALG00010014954 | NA | NA | 29 | 60 | 72 | 67 | 167 | 109 | 81 | 56 | 0.000764728 | 0.007513308 |

| | | | | | | | | | | | | | |
|---|---|---|---|---|---|---|---|---|---|---|---|---|---|
| ENSGALG00010014957 | NA | NA | | 34 | 54 | 121 | 157 | 92 | 51 | 60 | 43 | 0.003322658 | 0.024407172 |
| ENSGALG00010014980 | LOC121107687 | LOC121107687 | | 1559 | 2985 | 393 | 530 | 203 | 256 | 558 | 868 | 6.01E-19 | 1.10E-16 |
| ENSGALG00010014982 | NA | NA | | 49 | 35 | 1 | 5 | 4 | 9 | 37 | 19 | 3.82E-06 | 8.55E-05 |
| ENSGALG00010014984 | LPP | LPP | | 112 | 145 | 86 | 99 | 79 | 169 | 342 | 232 | 0.007377323 | 0.045734855 |
| ENSGALG00010014989 | NA | NA | | 21 | 40 | 78 | 54 | 116 | 62 | 30 | 32 | 0.001100729 | 0.010112906 |
| ENSGALG00010015002 | ADAM19 | ADAM19 | | 71 | 63 | 26 | 24 | 38 | 43 | 101 | 67 | 0.006607338 | 0.04182645 |
| ENSGALG00010015021 | RBM46 | RBM46 | | 139 | 90 | 544 | 474 | 675 | 431 | 261 | 108 | 8.79E-09 | 3.78E-07 |
| ENSGALG00010015028 | MCM9 | MCM9 | | 489 | 457 | 536 | 576 | 505 | 487 | 1126 | 1013 | 0.00012697 | 0.001688083 |
| ENSGALG00010015032 | C1orf216 | C23H1orf216 | | 47 | 69 | 27 | 13 | 25 | 19 | 66 | 89 | 0.000968419 | 0.009102484 |
| ENSGALG00010015039 | FAM114A1 | FAM114A1 | | 246 | 227 | 508 | 631 | 310 | 223 | 336 | 390 | 0.004703515 | 0.032334491 |
| ENSGALG00010015042 | EFNB2 | EFNB2 | | 3297 | 3907 | 1798 | 2294 | 1763 | 1461 | 3655 | 3086 | 3.60E-07 | 1.05E-05 |
| ENSGALG00010015050 | HIVEP3 | HIVEP3 | | 789 | 1058 | 267 | 270 | 525 | 704 | 1746 | 1078 | 1.03E-10 | 6.30E-09 |
| ENSGALG00010015060 | NA | NA | | 6 | 6 | 25 | 59 | 9 | 23 | 33 | 49 | 0.001319117 | 0.011651736 |
| ENSGALG00010015061 | MMP9 | MMP9 | | 8 | 7 | 37 | 21 | 4 | 10 | 30 | 42 | 0.001668334 | 0.014024751 |
| ENSGALG00010015071 | SNAI1 | SNAI1 | | 1143 | 915 | 442 | 549 | 751 | 677 | 857 | 990 | 0.00045331 | 0.004874979 |
| ENSGALG00010015084 | FOXO6 | FOXO6 | | 1079 | 1011 | 663 | 430 | 681 | 374 | 1155 | 925 | 0.001276238 | 0.011338912 |
| ENSGALG00010015087 | GABRA3 | GABRA3 | | 277 | 527 | 167 | 161 | 138 | 102 | 374 | 291 | 0.000160181 | 0.002067138 |
| ENSGALG00010015088 | LOC107050476 | gag-env | | 205 | 56 | 30 | 26 | 33 | 19 | 20 | 25 | 1.33E-05 | 0.000250467 |
| ENSGALG00010015094 | CDK5RAP1 | CDK5RAP1 | | 1074 | 876 | 477 | 476 | 485 | 470 | 1062 | 1150 | 7.22E-06 | 0.000148135 |
| ENSGALG00010015096 | GABRE | GABRG4 | | 9 | 10 | 1 | 6 | 2 | 1 | 29 | 35 | 1.16E-06 | 2.97E-05 |
| ENSGALG00010015108 | FAAH2 | FAAH2 | | 1584 | 1420 | 941 | 890 | 584 | 485 | 1205 | 1563 | 4.33E-06 | 9.54E-05 |
| ENSGALG00010015117 | RASD2 | RASD2 | | 0 | 0 | 24 | 8 | 9 | 1 | 18 | 50 | 8.74E-07 | 2.30E-05 |
| ENSGALG00010015121 | UNC79 | UNC79 | | 184 | 103 | 196 | 170 | 93 | 119 | 487 | 298 | 0.001293418 | 0.011455006 |
| ENSGALG00010015126 | MCM5 | MCM5 | | 5744 | 6709 | 9556 | 9740 | 6954 | 4826 | 4504 | 6311 | 0.001289867 | 0.011435681 |
| ENSGALG00010015132 | GCNT1 | GCNT1 | | 47 | 140 | 420 | 285 | 576 | 100 | 75 | 65 | 3.57E-05 | 0.00058009 |
| ENSGALG00010015138 | PRUNE2 | PRUNE2 | | 10 | 2 | 138 | 39 | 72 | 15 | 25 | 12 | 0.000102555 | 0.001409453 |
| ENSGALG00010015149 | VSIG10L | VSIG10L | | 3 | 8 | 4 | 0 | 0 | 1 | 13 | 27 | 0.000152813 | 0.001984313 |
| ENSGALG00010015154 | CYP24A1 | CP450CC24 | | 33 | 37 | 10 | 17 | 59 | 41 | 84 | 102 | 2.92E-06 | 6.71E-05 |
| ENSGALG00010015173 | ENS-1 | ENS-1 | | 19353 | 4399 | 1339 | 2617 | 1806 | 2443 | 4739 | 4955 | 3.56E-08 | 1.32E-06 |
| ENSGALG00010015177 | TC2N | MTAC2D1 | | 37 | 30 | 206 | 198 | 170 | 131 | 91 | 86 | 1.10E-07 | 3.72E-06 |
| ENSGALG00010015190 | FBLN5 | FBLN5 | | 14 | 10 | 21 | 41 | 11 | 12 | 31 | 69 | 0.004413538 | 0.030618116 |
| ENSGALG00010015194 | TMLHE | TMLHE | | 891 | 1097 | 1573 | 1679 | 1390 | 985 | 2410 | 2336 | 1.57E-06 | 3.94E-05 |
| ENSGALG00010015198 | NME5 | NME5 | | 272 | 199 | 209 | 303 | 359 | 374 | 183 | 161 | 0.003677289 | 0.026443566 |
| ENSGALG00010015205 | BPIFB3 | BPIFB8 | | 9 | 9 | 5 | 7 | 7 | 7 | 43 | 42 | 3.84E-06 | 8.57E-05 |
| ENSGALG00010015218 | PLCL1 | PLCL1 | | 1783 | 2653 | 1856 | 2833 | 1593 | 1159 | 1467 | 897 | 0.003382506 | 0.024706485 |
| ENSGALG00010015222 | NA | NA | | 33 | 46 | 128 | 233 | 131 | 124 | 202 | 248 | 4.98E-06 | 0.000107704 |
| ENSGALG00010015233 | BPIFB2 | BPIFB7 | | 5 | 3 | 0 | 2 | 1 | 1 | 37 | 21 | 2.59E-08 | 9.96E-07 |
| ENSGALG00010015243 | NA | NA | | 1 | 0 | 19 | 19 | 60 | 52 | 107 | 89 | 1.36E-17 | 2.13E-15 |
| ENSGALG00010015285 | ALG13 | GLT28D1 | | 1644 | 596 | 907 | 1082 | 2619 | 2420 | 1850 | 942 | 9.83E-05 | 0.001360314 |
| ENSGALG00010015306 | BCAT1 | BCAT1 | | 314 | 716 | 702 | 803 | 567 | 720 | 1527 | 1025 | 0.004581424 | 0.031586121 |
| ENSGALG00010015320 | TRPM6 | TRPM6 | | 141 | 61 | 142 | 88 | 370 | 162 | 467 | 161 | 0.003642282 | 0.026228113 |
| ENSGALG00010015362 | ATP9A | ATP9A | | 1516 | 2402 | 967 | 1260 | 1250 | 1279 | 2360 | 1580 | 0.006669177 | 0.042090363 |
| ENSGALG00010015367 | NA | NA | | 15 | 7 | 3 | 1 | 7 | 3 | 22 | 12 | 0.007893369 | 0.048302448 |
| ENSGALG00010015374 | LIMS1 | PINCH1 | | 2405 | 3693 | 3042 | 3183 | 1558 | 1049 | 2579 | 2953 | 5.19E-05 | 0.000793112 |
| ENSGALG00010015376 | NA | NA | | 175 | 86 | 123 | 165 | 143 | 137 | 44 | 36 | 0.000539031 | 0.005651368 |
| ENSGALG00010015393 | DCX | 18C15.5 | | 22 | 54 | 20 | 14 | 34 | 90 | 126 | 46 | 0.004834605 | 0.032953928 |
| ENSGALG00010015401 | NA | NA | | 9 | 13 | 25 | 32 | 81 | 88 | 88 | 38 | 8.58E-07 | 2.26E-05 |
| ENSGALG00010015472 | LGMN | LGMN | | 1872 | 1699 | 1481 | 1702 | 3079 | 1959 | 1366 | 1096 | 2.51E-07 | 7.68E-06 |
| ENSGALG00010015478 | FBXO7 | FBXO7 | | 1988 | 2087 | 3502 | 3815 | 2843 | 1902 | 2197 | 2340 | 0.000621745 | 0.006355113 |
| ENSGALG00010015483 | SGCD | SGCD | | 286 | 345 | 112 | 143 | 80 | 199 | 491 | 304 | 0.003468781 | 0.02523738 |
| ENSGALG00010015485 | BCAS1 | BCAS1 | | 4 | 8 | 76 | 11 | 8 | 0 | 3 | 7 | 0.000600441 | 0.006175121 |
| ENSGALG00010015499 | NA | NA | | 148 | 341 | 571 | 576 | 586 | 600 | 1125 | 559 | 0.000201443 | 0.002512323 |
| ENSGALG00010015502 | LOC107049084 | BPIFB6 | | 4 | 7 | 0 | 1 | 4 | 5 | 26 | 28 | 2.93E-07 | 8.82E-06 |
| ENSGALG00010015505 | CHGA | CHGA | | 43 | 20 | 11 | 2 | 11 | 3 | 31 | 77 | 0.000139771 | 0.001833475 |
| ENSGALG00010015523 | MRPL22 | MRPL22 | | 1107 | 1196 | 2094 | 2481 | 1758 | 1426 | 1583 | 2005 | 0.001347882 | 0.011849446 |
| ENSGALG00010015533 | PSAT1 | PSAT1 | | 292 | 119 | 197 | 64 | 96 | 237 | 1006 | 1145 | 2.02E-07 | 6.32E-06 |
| ENSGALG00010015549 | LOC101750367 | LOC101750367 | | 10 | 6 | 2 | 0 | 2 | 9 | 31 | 26 | 4.32E-05 | 0.000682504 |
| ENSGALG00010015553 | ASCL4 | ASCL4 | | 1 | 0 | 11 | 10 | 17 | 5 | 4 | 6 | 0.001632503 | 0.013799894 |
| ENSGALG00010015560 | NA | NA | | 1 | 0 | 0 | 1 | 4 | 10 | 16 | 5 | 0.000576799 | 0.005972402 |
| ENSGALG00010015561 | NA | NA | | 11 | 12 | 63 | 45 | 43 | 21 | 12 | 11 | 8.40E-05 | 0.001194322 |
| ENSGALG00010015563 | B3GNT7 | B3GNT7 | | 383 | 275 | 165 | 173 | 120 | 118 | 162 | 185 | 0.006399296 | 0.040797931 |
| ENSGALG00010015570 | LIMCH1 | LIMCH1 | | 515 | 509 | 606 | 532 | 227 | 259 | 870 | 527 | 0.009062414 | 0.054006302 |
| ENSGALG00010015608 | NA | NA | | 16 | 6 | 5 | 1 | 23 | 14 | 4 | 6 | 0.00461821 | 0.031787285 |
| ENSGALG00010015610 | MYOT | TTID | | 4 | 6 | 18 | 18 | 36 | 21 | 4 | 7 | 5.45E-05 | 0.000825068 |
| ENSGALG00010015632 | COL4A5 | COL4A5 | | 1334 | 1526 | 1536 | 1609 | 1429 | 1135 | 2733 | 2612 | 4.34E-05 | 0.000684449 |
| ENSGALG00010015661 | NA | NA | | 328 | 351 | 2 | 1 | 5 | 8 | 1 | 2 | 3.12E-51 | 3.26E-48 |
| ENSGALG00010015668 | LOC107048990 | ZNF217 | | 3147 | 3497 | 700 | 921 | 1113 | 1393 | 3287 | 2593 | 1.22E-15 | 1.56E-13 |
| ENSGALG00010015669 | INPP4A | INPP4A | | 776 | 1123 | 1275 | 1797 | 1354 | 841 | 769 | 639 | 0.000604518 | 0.006197975 |
| ENSGALG00010015670 | PTMA | PTMA | | 55941 | 42612 | 25515 | 28155 | 28720 | 19632 | 33579 | 39759 | 0.000102876 | 0.00141271 |
| ENSGALG00010015681 | LOC112532815 | LOC112532815 | | 4 | 7 | 3 | 5 | 11 | 21 | 5 | 3 | 0.006922916 | 0.043364235 |
| ENSGALG00010015694 | NA | NA | | 5 | 5 | 47 | 53 | 48 | 36 | 15 | 29 | 2.71E-06 | 6.31E-05 |
| ENSGALG00010015695 | ASAH1 | ASAH1 | | 1375 | 1126 | 1947 | 2289 | 2174 | 1418 | 1401 | 899 | 0.000685524 | 0.006889043 |
| ENSGALG00010015727 | NA | NA | | 0 | 1 | 11 | 13 | 12 | 1 | 2 | 2 | 0.003331211 | 0.024427909 |
| ENSGALG00010015733 | COL4A6 | COL4A6 | | 2063 | 2356 | 2821 | 2685 | 3789 | 3022 | 5529 | 4369 | 1.82E-08 | 7.23E-07 |
| ENSGALG00010015738 | CRACDL | KIAA1211L | | 90 | 59 | 28 | 21 | 94 | 44 | 181 | 69 | 0.00142929 | 0.012376319 |
| ENSGALG00010015758 | PCDH19 | protocadherin-19 | | 884 | 1181 | 505 | 629 | 692 | 550 | 1060 | 1166 | 0.000284999 | 0.003349445 |
| ENSGALG00010015790 | NA | NA | | 11 | 3 | 92 | 46 | 38 | 16 | 12 | 13 | 1.27E-05 | 0.00024035 |
| ENSGALG00010015801 | VSTM2L | VSTM2L | | 379 | 783 | 209 | 260 | 125 | 157 | 402 | 474 | 3.00E-05 | 0.000500866 |
| ENSGALG00010015804 | CKB | CPK-B | | 3678 | 6735 | 7245 | 6631 | 2724 | 2054 | 4754 | 6603 | 2.34E-05 | 0.000403592 |
| ENSGALG00010015806 | NA | NA | | 35 | 287 | 0 | 0 | 0 | 0 | 0 | 1 | 8.25E-25 | 2.22E-22 |
| ENSGALG00010015809 | PCOLCE2 | PCOLCE2 | | 391 | 611 | 839 | 879 | 842 | 600 | 575 | 403 | 0.006358275 | 0.040603552 |
| ENSGALG00010015817 | LOC107053589 | LOC107053589 | | 205 | 350 | 436 | 592 | 434 | 251 | 228 | 137 | 0.002173089 | 0.017384712 |
| ENSGALG00010015843 | RXFP2 | RXFP2L | | 8 | 4 | 0 | 4 | 5 | 3 | 48 | 74 | 1.21E-11 | 9.02E-10 |
| ENSGALG00010015874 | CLIC4 | CLIC4 | | 5665 | 12734 | 6674 | 9064 | 7062 | 5705 | 4780 | 5031 | 0.009300377 | 0.055057702 |
| ENSGALG00010015876 | SKP1 | CTD-2410N18.5 | | 5325 | 5497 | 7067 | 8029 | 6310 | 4407 | 4317 | 5067 | 0.006983506 | 0.043612985 |
| ENSGALG00010015951 | NA | NA | | 47 | 44 | 15 | 21 | 10 | 6 | 28 | 42 | 0.000794719 | 0.007757797 |
| ENSGALG00010015956 | MESP1 | CMESO-1 | | 15 | 14 | 1 | 1 | 17 | 3 | 16 | 11 | 0.002104755 | 0.01692717 |
| ENSGALG00010015960 | CHRNA9 | NACHR alpha-9 | | 132 | 515 | 66 | 93 | 191 | 211 | 199 | 154 | 0.001739385 | 0.014490832 |
| ENSGALG00010015969 | NA | NA | | 2309 | 2367 | 1222 | 1132 | 1202 | 1245 | 1320 | 1915 | 0.000666217 | 0.006743682 |
| ENSGALG00010015972 | FSTL4 | FSTL4 | | 908 | 1638 | 407 | 377 | 557 | 684 | 1266 | 1080 | 3.71E-07 | 1.08E-05 |
| ENSGALG00010015973 | NA | NA | | 13 | 23 | 7 | 6 | 16 | 16 | 41 | 24 | 0.008354898 | 0.050552881 |
| ENSGALG00010016002 | NA | NA | | 195 | 277 | 631 | 705 | 502 | 334 | 190 | 207 | 1.69E-07 | 5.40E-06 |
| ENSGALG00010016014 | NA | NA | | 20 | 23 | 89 | 83 | 51 | 42 | 57 | 27 | 0.005555722 | 0.036684009 |
| ENSGALG00010016022 | EPHB3 | CEK10 | | 1444 | 5445 | 1010 | 936 | 2788 | 1869 | 1991 | 1616 | 3.05E-05 | 0.000508149 |
| ENSGALG00010016039 | LOC107054849 | LOC107054849 | | 40 | 24 | 7 | 4 | 2 | 4 | 18 | 10 | 0.000125682 | 0.001673619 |
| ENSGALG00010016053 | DRP2 | DRP2 | | 23 | 30 | 7 | 5 | 3 | 11 | 35 | 35 | 0.000592436 | 0.006100307 |
| ENSGALG00010016062 | NA | NA | | 0 | 3 | 18 | 20 | 16 | 7 | 2 | 11 | 0.004334804 | 0.030166298 |
| ENSGALG00010016079 | NA | NA | | 275 | 461 | 1366 | 1057 | 763 | 541 | 538 | 688 | 5.45E-06 | 0.000115808 |
| ENSGALG00010016084 | NA | NA | | 3 | 6 | 23 | 11 | 3 | 0 | 3 | 7 | 0.009596355 | 0.056370575 |
| ENSGALG00010016086 | TDRD9 | TDRD9 | | 3110 | 3834 | 5649 | 6121 | 4945 | 3909 | 4545 | 3977 | 0.001269495 | 0.011303036 |
| ENSGALG00010016099 | NA | NA | | 40 | 60 | 11 | 41 | 9 | 9 | 49 | 37 | 0.007178341 | 0.044610792 |
| ENSGALG00010016117 | MEIKIN | MEIKIN | | 160 | 201 | 498 | 524 | 400 | 339 | 145 | 110 | 7.93E-09 | 3.47E-07 |
| ENSGALG00010016127 | ACSL6 | ACSL6 | | 2292 | 3121 | 1201 | 1292 | 1681 | 1420 | 2562 | 2493 | 3.50E-07 | 1.03E-05 |
| ENSGALG00010016159 | ASPG | ASPG | | 2736 | 2518 | 1084 | 1729 | 1169 | 1932 | 3108 | 2862 | 0.001629571 | 0.013782082 |
| ENSGALG00010016173 | GLRA4 | GLRA4 | | 690 | 558 | 183 | 145 | 433 | 421 | 1016 | 869 | 4.90E-12 | 3.86E-10 |
| ENSGALG00010016190 | THNSL2 | THNSL2 | | 92 | 57 | 722 | 621 | 416 | 196 | 280 | 234 | 2.13E-11 | 1.50E-09 |
| ENSGALG00010016209 | ADAMTS7 | ADAMTS7 | | 575 | 644 | 1193 | 1246 | 1047 | 853 | 840 | 601 | 0.000464382 | 0.004984427 |
| ENSGALG00010016227 | NA | NA | | 5 | 3 | 13 | 11 | 4 | 5 | 28 | 45 | 1.64E-05 | 0.000297899 |
| ENSGALG00010016243 | NA | NA | | 1 | 1 | 18 | 14 | 8 | 2 | 7 | 7 | 0.002285587 | 0.018077012 |
| ENSGALG00010016253 | EDN3 | EDN3 | | 179 | 145 | 70 | 70 | 212 | 138 | 192 | 162 | 0.001787359 | 0.0148101 |
| ENSGALG00010016307 | RAD50 | RAD50 | | 2398 | 2570 | 3501 | 3816 | 3282 | 2105 | 2011 | 1764 | 5.88E-05 | 0.00087607 |
| ENSGALG00010016320 | NA | NA | | 164 | 121 | 23 | 35 | 31 | 37 | 119 | 99 | 2.56E-06 | 6.00E-05 |
| ENSGALG00010016322 | LOC107050199 | LOC107050199 | | 2588 | 3201 | 1976 | 2031 | 1744 | 1375 | 2628 | 2772 | 0.001805034 | 0.014912788 |
| ENSGALG00010016323 | NA | NA | | 4 | 1 | 30 | 18 | 27 | 6 | 7 | 11 | 0.004032834 | 0.028497559 |
| ENSGALG00010016328 | USH2A | USH2A | | 59 | 61 | 130 | 181 | 354 | 189 | 127 | 64 | 2.06E-06 | 4.96E-05 |
| ENSGALG00010016374 | FRMD5 | FRMD5 | | 133 | 147 | 104 | 128 | 86 | 102 | 38 | 53 | 0.005309393 | 0.035356486 |
| ENSGALG00010016377 | FRMPD3 | FRMPD3 | | 54 | 84 | 71 | 48 | 30 | 19 | 99 | 116 | 0.004749768 | 0.032520827 |
| ENSGALG00010016387 | CCDC85C | CCDC85C | | 837 | 824 | 816 | 900 | 576 | 398 | 1214 | 1184 | 0.000923077 | 0.008753927 |
| ENSGALG00010016395 | HHIPL1 | HHIPL1 | | 23 | 43 | 170 | 107 | 76 | 19 | 54 | 64 | 0.004355998 | 0.030294398 |
| ENSGALG00010016399 | ACSM5 | ACSM5 | | 0 | 0 | 15 | 24 | 14 | 17 | 2 | 6 | 3.22E-07 | 9.57E-06 |
| ENSGALG00010016402 | SEPTIN8 | | 8-Sep | 956 | 1063 | 678 | 774 | 618 | 524 | 1069 | 1490 | 0.001222072 | 0.010968454 |
| ENSGALG00010016404 | ESRRG | ERRgamma | | 229 | 332 | 510 | 646 | 516 | 386 | 273 | 258 | 0.000583766 | 0.006022154 |
| ENSGALG00010016412 | TMSB15B | TMSB15B | | 29032 | 25749 | 9481 | 10743 | 12586 | 9965 | 18801 | 23412 | 6.50E-13 | 5.68E-11 |
| ENSGALG00010016413 | NA | NA | | 32 | 19 | 3 | 0 | 4 | 8 | 14 | 4 | 0.000687033 | 0.006900061 |
| ENSGALG00010016416 | CYP46A1 | CYP46A1 | | 117 | 149 | 375 | 334 | 302 | 194 | 274 | 308 | 0.003669889 | 0.026401718 |
| ENSGALG00010016425 | TRIM63 | TRIM63 | | 299 | 648 | 135 | 165 | 290 | 192 | 241 | 248 | 0.000384916 | 0.004291343 |
| ENSGALG00010016428 | ADRA1D | ADRA1D | | 327 | 216 | 2 | 5 | 38 | 20 | 122 | 41 | 1.15E-17 | 1.82E-15 |
| ENSGALG00010016441 | LOC772096 | LOC772096 | | 256 | 504 | 100 | 155 | 186 | 108 | 433 | 293 | 0.000181229 | 0.002297955 |
| ENSGALG00010016447 | ID3 | ID3 | | 811 | 330 | 164 | 74 | 183 | 207 | 743 | 775 | 2.13E-07 | 6.61E-06 |
| ENSGALG00010016457 | SMOX | SMOX | | 1555 | 2020 | 308 | 370 | 452 | 311 | 1050 | 1118 | 9.32E-24 | 2.36E-21 |
| ENSGALG00010016474 | MAPK8IP3 | MAPK8IP3 | | 961 | 1372 | 2251 | 2414 | 2009 | 1545 | 1541 | 1200 | 9.62E-05 | 0.001340295 |

| | | | | | | | | | | | | |
|---|---|---|---|---|---|---|---|---|---|---|---|---|
| ENSGALG00010016485 | NA | NA | 19 | 30 | 17 | 21 | 75 | 51 | 66 | 55 | 0.000156392 | 0.002022923 |
| ENSGALG00010016490 | NA | NA | 577 | 1097 | 669 | 924 | 1758 | 1092 | 641 | 948 | 0.0002306 | 0.00282122 |
| ENSGALG00010016496 | PLP1 | PLP | 176 | 207 | 38 | 27 | 78 | 87 | 207 | 178 | 1.17E-07 | 3.94E-06 |
| ENSGALG00010016519 | MEX3B | RKHD3 | 3135 | 4863 | 2530 | 3591 | 2774 | 1708 | 1804 | 1868 | 3.70E-05 | 0.000597069 |
| ENSGALG00010016523 | LOC121107350 | LOC121107350 | 3 | 4 | 3 | 6 | 6 | 11 | 40 | 16 | 0.000563296 | 0.005857993 |
| ENSGALG00010016535 | BNC2 | BNC2 | 3 | 17 | 54 | 55 | 45 | 8 | 17 | 5 | 0.005909315 | 0.038373386 |
| ENSGALG00010016541 | UBE2J2 | UBE2J2 | 7079 | 7892 | 7980 | 10143 | 8417 | 5698 | 5532 | 5189 | 0.000922431 | 0.008753927 |
| ENSGALG00010016543 | TCEA3 | TCEA3 | 25 | 0 | 3 | 0 | 0 | 2 | 38 | 27 | 0.000736416 | 0.007316877 |
| ENSGALG00010016544 | ADCY9 | AC9 | 606 | 827 | 1413 | 1402 | 1101 | 688 | 1184 | 1073 | 0.005048034 | 0.033935131 |
| ENSGALG00010016562 | GLIS2 | GLIS2 | 1939 | 2608 | 1528 | 1939 | 1531 | 983 | 1070 | 1160 | 0.000172386 | 0.002204968 |
| ENSGALG00010016574 | NA | NA | 38 | 68 | 101 | 85 | 55 | 44 | 159 | 141 | 0.005020115 | 0.033801838 |
| ENSGALG00010016578 | GRIK3 | GRIK3 | 144 | 349 | 249 | 191 | 589 | 286 | 610 | 389 | 0.004143884 | 0.029158987 |
| ENSGALG00010016581 | H2AFY | MACROH2A1 | 942 | 645 | 259 | 425 | 396 | 333 | 511 | 498 | 0.001262816 | 0.011255566 |
| ENSGALG00010016583 | VPS41 | VPS41 | 838 | 615 | 1528 | 1711 | 1006 | 712 | 703 | 588 | 8.62E-06 | 0.000173131 |
| ENSGALG00010016588 | ATP11B | ATP11B | 2807 | 3036 | 5432 | 5811 | 4560 | 4131 | 5778 | 4606 | 3.54E-05 | 0.000575409 |
| ENSGALG00010016595 | CLCNKB | CLCNKA | 70 | 149 | 126 | 91 | 123 | 105 | 265 | 270 | 0.007693177 | 0.047302567 |
| ENSGALG00010016596 | LOC419322 | LOC419322 | 50 | 28 | 11 | 10 | 29 | 25 | 137 | 53 | 1.92E-05 | 0.000340157 |
| ENSGALG00010016599 | LOC416263 | LOC416263 | 391 | 2375 | 268 | 381 | 396 | 749 | 963 | 827 | 0.003710741 | 0.026626836 |
| ENSGALG00010016601 | NA | NA | 6 | 10 | 1 | 0 | 7 | 2 | 31 | 13 | 5.06E-05 | 0.000779218 |
| ENSGALG00010016612 | GRIA1 | GluR1 | 412 | 1075 | 65 | 103 | 182 | 67 | 116 | 111 | 4.87E-13 | 4.31E-11 |
| ENSGALG00010016614 | DNAJC19 | DNAJC19 | 1121 | 1300 | 839 | 939 | 870 | 686 | 1557 | 1718 | 0.000289816 | 0.003389369 |
| ENSGALG00010016622 | KIAA1143 | KIAA1143 | 3167 | 3403 | 1912 | 2358 | 2406 | 1772 | 3980 | 4023 | 3.43E-06 | 7.77E-05 |
| ENSGALG00010016645 | NA | NA | 2984 | 3397 | 1359 | 1210 | 13 | 21 | 5162 | 2717 | 1.51E-73 | 3.15E-70 |
| ENSGALG00010016671 | CASS4 | C20orf32 | 22 | 10 | 1 | 2 | 1 | 7 | 20 | 27 | 0.000305956 | 0.003553258 |
| ENSGALG00010016680 | BPIL3 | BPIL3 | 42 | 19 | 1 | 0 | 16 | 8 | 71 | 55 | 5.24E-11 | 3.42E-09 |
| ENSGALG00010016688 | NA | NA | 7 | 8 | 28 | 48 | 34 | 40 | 65 | 49 | 8.33E-05 | 0.001186831 |
| ENSGALG00010016706 | RTN4 | NOGO | 6718 | 7365 | 9840 | 12856 | 8124 | 5187 | 6913 | 6918 | 0.0018201 | 0.015020995 |
| ENSGALG00010016718 | LIN28A | LIN28 | 14253 | 19020 | 7947 | 9632 | 5259 | 6078 | 8738 | 16456 | 2.31E-05 | 0.000400331 |
| ENSGALG00010016722 | BMP15 | GDF-9B | 4 | 4 | 20 | 18 | 21 | 19 | 10 | 11 | 0.003341758 | 0.02446231 |
| ENSGALG00010016729 | ELMO1 | ELMO1 | 619 | 879 | 1543 | 1754 | 672 | 462 | 510 | 559 | 3.30E-09 | 1.56E-07 |
| ENSGALG00010016731 | C4orf48 | C4H4ORF48 | 21 | 190 | 8 | 17 | 5 | 31 | 43 | 39 | 0.004268822 | 0.02981218 |
| ENSGALG00010016745 | BEGAIN | BEGAIN | 27 | 88 | 3 | 2 | 0 | 2 | 9 | 9 | 8.37E-11 | 5.25E-09 |
| ENSGALG00010016754 | NA | NA | 104 | 144 | 60 | 55 | 110 | 150 | 222 | 121 | 0.007538564 | 0.04652307 |
| ENSGALG00010016760 | DLK1 | DLK1 | 133 | 59 | 2 | 6 | 15 | 17 | 35 | 29 | 1.21E-09 | 6.22E-08 |
| ENSGALG00010016761 | GPR160 | GPR160 | 2708 | 2453 | 922 | 1716 | 701 | 691 | 1133 | 1191 | 2.79E-08 | 1.07E-06 |
| ENSGALG00010016762 | EML6 | EML6 | 44 | 20 | 182 | 253 | 94 | 54 | 79 | 67 | 3.62E-06 | 8.15E-05 |
| ENSGALG00010016797 | SEC62 | TLOC1 | 5041 | 6354 | 4380 | 5611 | 3483 | 2202 | 3609 | 4182 | 0.000703015 | 0.007025303 |
| ENSGALG00010016812 | LRRC31 | LRRC31 | 1266 | 912 | 542 | 581 | 523 | 494 | 1078 | 908 | 0.000391441 | 0.004355364 |
| ENSGALG00010016832 | DYNC1H1 | DYNC1H1 | 2202 | 2941 | 4101 | 4065 | 2818 | 2776 | 7075 | 3380 | 0.00538036 | 0.035724447 |
| ENSGALG00010016847 | EPHB2 | EPHB2 | 462 | 650 | 488 | 518 | 362 | 244 | 802 | 746 | 0.003875129 | 0.027598889 |
| ENSGALG00010016859 | RPIA | RPIA | 1314 | 1258 | 1796 | 2247 | 2716 | 1789 | 1878 | 1647 | 0.000163308 | 0.002101 |
| ENSGALG00010016868 | NIPAL1 | NPAL1 | 1272 | 1592 | 1020 | 1954 | 1018 | 677 | 529 | 379 | 2.61E-07 | 7.93E-06 |
| ENSGALG00010016874 | ADAMTSL3 | ADAMTSL3 | 133 | 321 | 757 | 699 | 1245 | 336 | 231 | 132 | 6.80E-07 | 1.84E-05 |
| ENSGALG00010016875 | SESN2 | SESN2 | 182 | 174 | 131 | 69 | 125 | 246 | 380 | 670 | 3.70E-05 | 0.000597069 |
| ENSGALG00010016891 | LOC107052288 | LOC107052288 | 402 | 186 | 90 | 116 | 22 | 18 | 29 | 33 | 5.86E-14 | 5.94E-12 |
| ENSGALG00010016893 | TBC1D9B | TBC1D9B | 618 | 1007 | 1812 | 1588 | 1255 | 731 | 1078 | 1036 | 0.003582969 | 0.025921464 |
| ENSGALG00010016919 | ARHGDIG | ARHGDIG | 537 | 649 | 383 | 607 | 300 | 189 | 545 | 734 | 0.004865532 | 0.033080866 |
| ENSGALG00010016928 | RGS11 | RGS11 | 8 | 13 | 23 | 15 | 5 | 3 | 31 | 72 | 1.53E-05 | 0.000282309 |
| ENSGALG00010016931 | SLC30A4 | SLC30A4 | 4096 | 3730 | 7107 | 6984 | 6497 | 4758 | 5302 | 4433 | 0.000100145 | 0.001379744 |
| ENSGALG00010016935 | PRIMA1 | PRIMA1 | 105 | 281 | 70 | 56 | 107 | 69 | 249 | 302 | 9.03E-05 | 0.001270702 |
| ENSGALG00010016941 | SLC8A1 | NCX1 | 997 | 1195 | 152 | 128 | 355 | 485 | 1322 | 916 | 9.75E-18 | 1.57E-15 |
| ENSGALG00010016945 | BDH2 | DHRS6 | 5 | 24 | 75 | 87 | 135 | 29 | 32 | 20 | 0.000383218 | 0.004280651 |
| ENSGALG00010016955 | LOC424998 | MEGF6L | 1682 | 1345 | 586 | 998 | 816 | 940 | 2315 | 2342 | 4.10E-07 | 1.18E-05 |
| ENSGALG00010016979 | PLSCR5 | PLSCR5 | 8 | 5 | 5 | 6 | 19 | 11 | 29 | 22 | 0.002396643 | 0.01870747 |
| ENSGALG00010017006 | DNAH3 | DNAH3 | 408 | 624 | 1055 | 1889 | 1124 | 1113 | 791 | 1265 | 9.99E-05 | 0.001377046 |
| ENSGALG00010017016 | ZIC1 | ZIC1 | 6 | 20 | 171 | 51 | 59 | 79 | 16 | 34 | 4.77E-05 | 0.000739234 |
| ENSGALG00010017020 | GJB5 | GJB5 | 2 | 0 | 22 | 8 | 5 | 2 | 18 | 26 | 0.000113572 | 0.001538094 |
| ENSGALG00010017022 | CCR4 | CCR4 | 30 | 13 | 67 | 26 | 7 | 7 | 4 | 4 | 4.89E-05 | 0.000756425 |
| ENSGALG00010017031 | NA | NA | 2 | 0 | 24 | 5 | 10 | 5 | 1 | 4 | 0.007478992 | 0.046240811 |
| ENSGALG00010017033 | TMEM159 | TMEM159 | 2433 | 2825 | 4492 | 7773 | 4606 | 4316 | 2667 | 4195 | 2.06E-05 | 0.000360843 |
| ENSGALG00010017037 | NA | NA | 8 | 1 | 19 | 33 | 27 | 12 | 6 | 9 | 0.005312359 | 0.035356486 |
| ENSGALG00010017043 | NA | NA | 4 | 11 | 3 | 3 | 17 | 27 | 6 | 2 | 0.000286485 | 0.003364537 |
| ENSGALG00010017045 | GLB1 | GLB1 | 2232 | 2460 | 2240 | 2526 | 1478 | 835 | 1388 | 1661 | 0.000554059 | 0.00578715 |
| ENSGALG00010017068 | ZP2 | ZPA | 0 | 0 | 12 | 11 | 5 | 2 | 7 | 7 | 0.00019174 | 0.002405672 |
| ENSGALG00010017070 | BLOC1S6 | PLDN | 320 | 332 | 937 | 729 | 635 | 460 | 484 | 483 | 0.000303767 | 0.00353274 |
| ENSGALG00010017090 | NA | NA | 13 | 12 | 1 | 4 | 2 | 3 | 13 | 15 | 0.003200963 | 0.023680608 |
| ENSGALG00010017094 | SORBS2 | 9430041O17RIK | 124 | 145 | 44 | 102 | 129 | 135 | 249 | 197 | 0.006157948 | 0.039579655 |
| ENSGALG00010017104 | FANCL | FANCL | 728 | 873 | 1313 | 1521 | 1530 | 953 | 1212 | 1131 | 0.003577399 | 0.025892378 |
| ENSGALG00010017122 | NCBP2 | NCBP2 | 5251 | 6162 | 3271 | 4149 | 3201 | 2760 | 3402 | 4154 | 0.000393009 | 0.004364246 |
| ENSGALG00010017131 | TLR3 | TLR3 | 136 | 99 | 7 | 15 | 47 | 66 | 77 | 62 | 3.05E-07 | 9.16E-06 |
| ENSGALG00010017132 | NA | NA | 35 | 61 | 119 | 152 | 114 | 67 | 30 | 59 | 0.002987765 | 0.022370754 |
| ENSGALG00010017135 | LOC100859848 | LOC100859848 | 109 | 204 | 766 | 635 | 587 | 426 | 627 | 557 | 2.09E-08 | 8.14E-07 |
| ENSGALG00010017139 | COL9A2 | COL9A2 | 1662 | 2064 | 994 | 1130 | 1130 | 1282 | 2601 | 2859 | 3.04E-07 | 9.14E-06 |
| ENSGALG00010017143 | FAM149A | FAM149A | 531 | 533 | 95 | 150 | 122 | 106 | 239 | 168 | 2.11E-09 | 1.04E-07 |
| ENSGALG00010017154 | PIGZ | PIGZ | 38 | 23 | 77 | 93 | 130 | 69 | 66 | 55 | 0.002968048 | 0.022253036 |
| ENSGALG00010017155 | BFSP2 | CP49INS | 395 | 9 | 35 | 10 | 1 | 11 | 37 | 49 | 0.000252518 | 0.003033849 |
| ENSGALG00010017158 | NA | NA | 40 | 50 | 14 | 6 | 10 | 4 | 16 | 24 | 0.000307654 | 0.003561468 |
| ENSGALG00010017166 | NA | NA | 3 | 7 | 32 | 23 | 14 | 10 | 10 | 8 | 0.005823282 | 0.037955809 |
| ENSGALG00010017180 | NA | NA | 132 | 123 | 38 | 39 | 132 | 51 | 79 | 66 | 0.002836271 | 0.021409107 |
| ENSGALG00010017188 | ITGAD | cd11b | 712 | 771 | 1368 | 1563 | 1042 | 830 | 882 | 1031 | 0.002126238 | 0.017074206 |
| ENSGALG00010017190 | STK26 | MST4 | 12092 | 11449 | 6361 | 7115 | 8638 | 7781 | 12906 | 11087 | 1.38E-05 | 0.000257128 |
| ENSGALG00010017195 | NA | NA | 88 | 89 | 1 | 3 | 13 | 16 | 84 | 33 | 1.42E-11 | 1.03E-09 |
| ENSGALG00010017200 | KCNQ4 | KCNQ4 | 693 | 930 | 455 | 564 | 403 | 281 | 450 | 519 | 0.003804743 | 0.027196265 |
| ENSGALG00010017215 | FRMD7 | FRMD7 | 26 | 36 | 4 | 1 | 7 | 15 | 30 | 23 | 8.61E-05 | 0.001219391 |
| ENSGALG00010017228 | NA | NA | 10529 | 11800 | 6773 | 7915 | 6667 | 5435 | 9160 | 10706 | 0.000268578 | 0.003199197 |
| ENSGALG00010017253 | NA | NA | 678 | 758 | 1406 | 1487 | 1371 | 898 | 882 | 792 | 9.95E-05 | 0.001373304 |
| ENSGALG00010017255 | NA | NA | 2 | 3 | 25 | 33 | 8 | 8 | 5 | 9 | 9.59E-05 | 0.001337583 |
| ENSGALG00010017270 | NA | NA | 21 | 23 | 3 | 15 | 6 | 5 | 54 | 31 | 0.000396591 | 0.004398028 |
| ENSGALG00010017272 | NRXN1 | NRXN1 | 1277 | 1503 | 1713 | 2775 | 2622 | 1729 | 1735 | 1344 | 0.001755787 | 0.014605634 |
| ENSGALG00010017318 | ADSS1 | ADSSL1 | 193 | 177 | 498 | 609 | 354 | 295 | 296 | 301 | 0.000168228 | 0.00216097 |
| ENSGALG00010017327 | CLN6 | CLN6 | 752 | 827 | 373 | 615 | 450 | 450 | 1080 | 861 | 0.002384204 | 0.018645211 |
| ENSGALG00010017343 | DLG1 | DLG1 | 1891 | 1648 | 1022 | 1340 | 1004 | 1056 | 2187 | 1619 | 0.009959736 | 0.058118165 |
| ENSGALG00010017346 | GM2A | GM2A | 466 | 2155 | 873 | 978 | 214 | 93 | 680 | 1249 | 2.41E-06 | 5.68E-05 |
| ENSGALG00010017374 | NA | NA | 9 | 10 | 22 | 46 | 41 | 36 | 42 | 19 | 0.008134726 | 0.049466476 |
| ENSGALG00010017375 | BDH1A | BDH | 2056 | 2258 | 2590 | 2914 | 1752 | 1303 | 1462 | 1747 | 0.00576646 | 0.037644173 |
| ENSGALG00010017376 | GALNT14 | GALNT14 | 14 | 45 | 29 | 15 | 10 | 17 | 85 | 164 | 6.08E-06 | 0.000127411 |
| ENSGALG00010017378 | LOC422926 | LOC422926 | 43 | 27 | 5 | 7 | 9 | 6 | 65 | 38 | 1.05E-06 | 2.73E-05 |
| ENSGALG00010017380 | MARCKSL1 | MARCKSL1 | 18886 | 20578 | 9232 | 13578 | 11220 | 8097 | 12595 | 16739 | 0.000126045 | 0.001677118 |
| ENSGALG00010017398 | SLC22A5 | OCTN2 | 784 | 1244 | 1737 | 2176 | 914 | 718 | 709 | 929 | 2.83E-05 | 0.000475173 |
| ENSGALG00010017399 | CPXM1 | CPXM1 | 240 | 390 | 90 | 189 | 128 | 92 | 268 | 342 | 0.001015874 | 0.009468646 |
| ENSGALG00010017401 | NA | NA | 0 | 0 | 13 | 55 | 0 | 1 | 0 | 2 | 3.30E-10 | 1.85E-08 |
| ENSGALG00010017402 | SLC2A5 | SLC2A5 | 4 | 9 | 17 | 17 | 31 | 21 | 46 | 34 | 0.000392928 | 0.004364246 |
| ENSGALG00010017403 | ITGA11 | ITGA11 | 128 | 137 | 14 | 22 | 115 | 60 | 130 | 52 | 7.70E-06 | 0.00015596 |
| ENSGALG00010017409 | GPC3 | GPC3 | 47 | 165 | 20 | 17 | 38 | 18 | 88 | 55 | 0.000294398 | 0.003435738 |
| ENSGALG00010017413 | ZIC3 | ZIC3 | 4569 | 6263 | 2758 | 2328 | 2940 | 2045 | 3494 | 4601 | 5.51E-07 | 1.54E-05 |
| ENSGALG00010017422 | NA | NA | 56 | 27 | 21 | 18 | 100 | 48 | 29 | 46 | 0.002851231 | 0.021502607 |
| ENSGALG00010017430 | CACNA1H | CACNA1H | 845 | 1016 | 399 | 441 | 351 | 264 | 800 | 714 | 4.55E-07 | 1.30E-05 |
| ENSGALG00010017442 | LCK | P56LCK | 8 | 6 | 23 | 47 | 35 | 21 | 23 | 18 | 0.004389052 | 0.030460898 |
| ENSGALG00010017443 | NA | NA | 474 | 517 | 121 | 235 | 355 | 224 | 287 | 216 | 0.000478586 | 0.005091106 |
| ENSGALG00010017446 | UNCX | UNCX | 18 | 20 | 6 | 9 | 67 | 10 | 37 | 70 | 0.001053417 | 0.009742506 |
| ENSGALG00010017447 | CAPN13 | calpain-13 | 1 | 12 | 8 | 2 | 7 | 9 | 20 | 45 | 0.006549348 | 0.041569582 |
| ENSGALG00010017450 | LOC416755 | LOC416755 | 223 | 150 | 85 | 129 | 56 | 51 | 236 | 264 | 8.37E-05 | 0.001190804 |
| ENSGALG00010017452 | ARMH4 | C14orf37 | 1173 | 1688 | 483 | 741 | 689 | 868 | 1140 | 1007 | 0.000477709 | 0.00508897 |
| ENSGALG00010017457 | ZMAT4 | ZMAT4 | 2 | 6 | 4 | 3 | 5 | 3 | 40 | 29 | 6.63E-07 | 1.80E-05 |
| ENSGALG00010017464 | SH3BP2 | SH3BP2 | 215 | 133 | 58 | 49 | 73 | 112 | 169 | 108 | 0.007864276 | 0.048170883 |
| ENSGALG00010017467 | SFRP1 | CSFRP1 | 2039 | 3144 | 1993 | 2507 | 1215 | 987 | 1580 | 2743 | 0.006495847 | 0.041261342 |
| ENSGALG00010017478 | NA | NA | 1 | 0 | 14 | 15 | 4 | 1 | 19 | 10 | 6.35E-05 | 0.000935806 |
| ENSGALG00010017479 | SSTR5 | SS5R | 12 | 15 | 0 | 2 | 1 | 0 | 7 | 8 | 5.53E-05 | 0.000834142 |
| ENSGALG00010017490 | SCNN1B | SCNN1B | 204 | 250 | 111 | 225 | 84 | 117 | 432 | 379 | 0.000490919 | 0.005202435 |
| ENSGALG00010017497 | NA | NA | 50 | 28 | 7 | 10 | 2 | 3 | 9 | 2 | 5.59E-06 | 0.000118458 |
| ENSGALG00010017498 | NA | NA | 52 | 44 | 104 | 139 | 104 | 108 | 67 | 55 | 0.005531889 | 0.036543323 |
| ENSGALG00010017509 | RASD1 | RASD1 | 214 | 394 | 192 | 196 | 103 | 94 | 386 | 693 | 1.97E-06 | 4.78E-05 |
| ENSGALG00010017518 | PEMT | PEMT | 549 | 582 | 1032 | 985 | 815 | 539 | 544 | 565 | 0.005922319 | 0.038391696 |
| ENSGALG00010017526 | PLEKHA2 | TAPP2 | 170 | 162 | 43 | 92 | 56 | 43 | 87 | 94 | 0.003446867 | 0.025110742 |
| ENSGALG00010017532 | MIXL1 | CMIX | 285 | 49 | 7 | 4 | 153 | 64 | 137 | 26 | 5.12E-06 | 0.00011022 |
| ENSGALG00010017554 | FAM131A | FAM131A | 33 | 12 | 15 | 15 | 1 | 2 | 31 | 40 | 5.19E-05 | 0.000793548 |
| ENSGALG00010017563 | ALK | ALK | 156 | 84 | 38 | 40 | 52 | 40 | 42 | 25 | 0.000738952 | 0.007329158 |
| ENSGALG00010017568 | CDCP1 | CDCP1 | 3657 | 2109 | 400 | 617 | 806 | 1082 | 3261 | 2298 | 8.09E-14 | 7.95E-12 |
| ENSGALG00010017607 | NA | NA | 11 | 40 | 1 | 9 | 38 | 59 | 122 | 89 | 1.67E-07 | 5.33E-06 |
| ENSGALG00010017609 | TRIM71 | LIN-41 | 4454 | 4754 | 2754 | 3155 | 2631 | 2322 | 4715 | 3575 | 0.000419905 | 0.004598461 |
| ENSGALG00010017612 | CLDN11 | CLDN11 | 1435 | 1431 | 171 | 307 | 253 | 330 | 819 | 573 | 2.58E-14 | 2.73E-12 |

| | | | | | | | | | | | | |
|---|---|---|---|---|---|---|---|---|---|---|---|---|
| ENSGALG00010017619 | POLN | POLN | 47 | 57 | 200 | 185 | 264 | 137 | 143 | 137 | 2.57E-05 | 0.000438157 |
| ENSGALG00010017623 | NA | NA | 0 | 0 | 0 | 0 | 5 | 11 | 7 | 23 | 3.95E-07 | 1.15E-05 |
| ENSGALG00010017632 | AWAT1 | AWAT1 | 3 | 7 | 6 | 1 | 0 | 0 | 13 | 23 | 4.95E-05 | 0.000763289 |
| ENSGALG00010017635 | NPPC | CNPP | 278 | 920 | 136 | 251 | 665 | 523 | 326 | 402 | 0.000212725 | 0.002632715 |
| ENSGALG00010017637 | NA | NA | 56 | 93 | 9 | 22 | 3 | 19 | 13 | 17 | 0.000287854 | 0.003373892 |
| ENSGALG00010017638 | GFRA2 | NTNRALPHA | 373 | 484 | 111 | 63 | 90 | 82 | 307 | 447 | 4.99E-10 | 2.71E-08 |
| ENSGALG00010017642 | 1-Mar | 2-Mar | 2910 | 2976 | 1992 | 2352 | 2039 | 1603 | 2977 | 3211 | 0.005738106 | 0.03756178 |
| ENSGALG00010017645 | SYNC | SYNC1 | 219 | 361 | 43 | 66 | 89 | 123 | 200 | 235 | 2.34E-06 | 5.54E-05 |
| ENSGALG00010017670 | FAM179A | TOGARAM2 | 117 | 63 | 115 | 137 | 136 | 72 | 36 | 17 | 0.000433074 | 0.004699696 |
| ENSGALG00010017690 | ARR3 | ARR3 | 23 | 56 | 82 | 142 | 116 | 86 | 35 | 55 | 0.002675264 | 0.02046179 |
| ENSGALG00010017691 | LZTS1 | LZTS1 | 790 | 1407 | 187 | 139 | 208 | 136 | 1291 | 1354 | 1.77E-27 | 6.29E-25 |
| ENSGALG00010017692 | EPCAM | TACSTD1 | 27739 | 9957 | 2970 | 4231 | 6027 | 7253 | 11569 | 8435 | 8.73E-09 | 3.77E-07 |
| ENSGALG00010017695 | FNDC5 | FNDC5 | 27 | 61 | 161 | 128 | 52 | 30 | 87 | 112 | 0.001903194 | 0.015537951 |
| ENSGALG00010017700 | C16orf62 | C14H16orf62 | 3428 | 3990 | 5879 | 7287 | 4541 | 3294 | 3936 | 3741 | 0.000108637 | 0.001478452 |
| ENSGALG00010017702 | ECEL1 | ECEL1 | 324 | 944 | 73 | 51 | 79 | 65 | 354 | 370 | 3.20E-14 | 3.30E-12 |
| ENSGALG00010017720 | ADAM32L1 | ADAM9L | 38 | 85 | 21 | 33 | 18 | 14 | 48 | 60 | 0.008846871 | 0.052954479 |
| ENSGALG00010017723 | KCNK12 | KCNK12 | 501 | 653 | 219 | 280 | 553 | 552 | 663 | 525 | 3.82E-05 | 0.000614 |
| ENSGALG00010017764 | IGDCC3 | PUNC | 1262 | 1568 | 571 | 635 | 581 | 659 | 1164 | 1143 | 5.17E-06 | 0.000110931 |
| ENSGALG00010017814 | LOC771069 | NADB-LER2 | 576 | 384 | 173 | 286 | 201 | 253 | 584 | 422 | 0.006531214 | 0.041470232 |
| ENSGALG00010017817 | REEP1 | REEP1 | 1294 | 1056 | 336 | 575 | 991 | 912 | 969 | 686 | 1.08E-05 | 0.000208984 |
| ENSGALG00010017820 | FLRT2 | FLRT2 | 2214 | 2287 | 196 | 295 | 305 | 284 | 271 | 446 | 8.48E-35 | 4.29E-32 |
| ENSGALG00010017828 | SLC12A9 | SLC12A9 | 1266 | 1860 | 927 | 977 | 779 | 679 | 1137 | 1328 | 0.002515626 | 0.019454486 |
| ENSGALG00010017845 | NA | NA | 17 | 21 | 1 | 5 | 10 | 16 | 24 | 33 | 0.000734436 | 0.00730154 |
| ENSGALG00010017858 | GPR65 | GPR65 | 0 | 7 | 29 | 42 | 38 | 18 | 10 | 3 | 4.06E-05 | 0.000643053 |
| ENSGALG00010017860 | CYSLTR1 | CYSLTR1 | 219 | 417 | 75 | 118 | 223 | 246 | 191 | 309 | 0.000782337 | 0.007664254 |
| ENSGALG00010017861 | PRSS56 | PRSS56 | 6 | 12 | 3 | 2 | 0 | 1 | 14 | 10 | 0.001030301 | 0.009559278 |
| ENSGALG00010017872 | LOC121107474 | LOC121107474 | 6 | 2 | 19 | 10 | 26 | 19 | 36 | 15 | 0.005162832 | 0.034567806 |
| ENSGALG00010017886 | DDRGK1 | C20orf116 | 1620 | 1702 | 1391 | 1608 | 2403 | 1947 | 1848 | 1569 | 0.000602284 | 0.006178863 |
| ENSGALG00010017908 | NA | NA | 35 | 38 | 4 | 10 | 19 | 6 | 17 | 25 | 0.003850439 | 0.027440738 |
| ENSGALG00010017913 | NA | NA | 78 | 151 | 25 | 29 | 65 | 62 | 93 | 127 | 0.000257619 | 0.003086256 |
| ENSGALG00010017922 | MERTK | C-EYK | 205 | 162 | 68 | 88 | 133 | 150 | 226 | 169 | 0.009968581 | 0.058148316 |
| ENSGALG00010017933 | SIX1 | SIX1 | 115 | 227 | 520 | 501 | 384 | 262 | 132 | 224 | 9.86E-05 | 0.001362108 |
| ENSGALG00010017973 | PKD1 | PKD1 | 352 | 632 | 245 | 358 | 250 | 243 | 606 | 564 | 0.008544475 | 0.051457752 |
| ENSGALG00010017976 | GPR137C | GPR137C | 731 | 1515 | 359 | 410 | 500 | 555 | 1497 | 1161 | 1.45E-07 | 4.77E-06 |
| ENSGALG00010017986 | NA | NA | 0 | 0 | 0 | 1 | 4 | 1 | 21 | 19 | 9.20E-09 | 3.90E-07 |
| ENSGALG00010017988 | NA | NA | 0 | 0 | 2 | 0 | 0 | 2 | 12 | 22 | 1.63E-06 | 4.06E-05 |
| ENSGALG00010017994 | NA | NA | 4 | 3 | 2 | 2 | 14 | 11 | 10 | 10 | 0.002310372 | 0.018186971 |
| ENSGALG00010018003 | NA | NA | 23 | 33 | 7 | 9 | 42 | 15 | 60 | 21 | 0.008250455 | 0.050011462 |
| ENSGALG00010018007 | DNAAF9 | C20orf194 | 336 | 649 | 1524 | 1270 | 1414 | 681 | 809 | 795 | 4.67E-05 | 0.0007276 |
| ENSGALG00010018012 | DVL1 | DVL1 | 90 | 107 | 264 | 272 | 261 | 122 | 161 | 101 | 0.006113465 | 0.039340865 |
| ENSGALG00010018029 | LOC121107471 | LOC121107471 | 46 | 31 | 10 | 10 | 28 | 31 | 54 | 44 | 0.001400999 | 0.012213615 |
| ENSGALG00010018048 | GFRA4 | GFRA4 | 153 | 163 | 52 | 69 | 82 | 62 | 170 | 143 | 0.00168424 | 0.014130031 |
| ENSGALG00010018060 | CASKIN1 | CASKIN1 | 255 | 565 | 187 | 154 | 156 | 153 | 704 | 821 | 9.17E-09 | 3.90E-07 |
| ENSGALG00010018064 | TTC8 | TTC8 | 338 | 261 | 893 | 827 | 665 | 453 | 397 | 349 | 5.66E-06 | 0.000119834 |
| ENSGALG00010018066 | NA | NA | 15 | 14 | 0 | 5 | 7 | 11 | 61 | 20 | 0.00012095 | 0.001620947 |
| ENSGALG00010018081 | DIO3 | DIO3 | 68 | 55 | 14 | 15 | 40 | 16 | 62 | 62 | 0.00046473 | 0.004984956 |
| ENSGALG00010018108 | NA | NA | 81 | 66 | 153 | 177 | 201 | 121 | 76 | 88 | 0.002902245 | 0.02183884 |
| ENSGALG00010018110 | CFAP65 | CCDC108 | 9 | 11 | 5 | 0 | 24 | 15 | 27 | 17 | 0.000758329 | 0.007481226 |
| ENSGALG00010018119 | GRID2IP | GRID2IP | 1 | 0 | 12 | 23 | 19 | 8 | 19 | 16 | 5.41E-05 | 0.000821204 |
| ENSGALG00010018122 | NA | NA | 15 | 42 | 50 | 123 | 57 | 29 | 18 | 23 | 0.006437153 | 0.040999011 |
| ENSGALG00010018123 | USP13 | USP13 | 285 | 414 | 751 | 783 | 867 | 527 | 644 | 333 | 0.002061797 | 0.016653818 |
| ENSGALG00010018124 | FBXO41 | FBXO41 | 67 | 73 | 5 | 5 | 16 | 23 | 89 | 53 | 6.24E-09 | 2.80E-07 |
| ENSGALG00010018142 | DDHD1 | DDHD1 | 1597 | 1969 | 2427 | 2853 | 2106 | 1620 | 1667 | 1308 | 0.004965698 | 0.033623323 |
| ENSGALG00010018176 | NA | NA | 3919 | 4999 | 1444 | 3033 | 1900 | 2454 | 2859 | 2437 | 0.001963558 | 0.0159296 |
| ENSGALG00010018193 | ANK1 | CHANK1 | 45 | 43 | 25 | 27 | 31 | 29 | 68 | 90 | 0.009579934 | 0.056293903 |
| ENSGALG00010018211 | NA | NA | 0 | 0 | 0 | 0 | 11 | 30 | 0 | 0 | 1.49E-11 | 1.08E-09 |
| ENSGALG00010018212 | LOC112530238 | LOC112530238 | 23 | 32 | 27 | 13 | 10 | 1 | 6 | 4 | 0.003706312 | 0.026606479 |
| ENSGALG00010018240 | SOUL | SOUL | 34 | 54 | 3 | 10 | 41 | 45 | 111 | 90 | 2.20E-08 | 8.56E-07 |
| ENSGALG00010018247 | NOTO | CNOT | 5521 | 5273 | 1122 | 1450 | 3830 | 3535 | 4121 | 3779 | 6.86E-24 | 1.76E-21 |
| ENSGALG00010018265 | PDCD4 | PDCD4 | 7802 | 10623 | 11664 | 14018 | 10400 | 6298 | 6607 | 6384 | 0.000116237 | 0.001567894 |
| ENSGALG00010018266 | GNOT2 | GNOT2 | 2696 | 2223 | 739 | 724 | 1314 | 1021 | 2125 | 2183 | 1.15E-13 | 1.10E-11 |
| ENSGALG00010018270 | NA | NA | 21 | 22 | 221 | 106 | 96 | 52 | 42 | 66 | 7.18E-06 | 0.000147588 |
| ENSGALG00010018279 | NA | NA | 6 | 2 | 95 | 21 | 125 | 19 | 18 | 40 | 0.000184609 | 0.002331963 |
| ENSGALG00010018283 | POF1B | POF1B | 1061 | 742 | 572 | 314 | 603 | 509 | 1268 | 1115 | 0.000237627 | 0.002877693 |
| ENSGALG00010018303 | RHBDL1 | RHBDL1 | 57 | 90 | 12 | 20 | 30 | 20 | 58 | 51 | 0.000264174 | 0.00315681 |
| ENSGALG00010018329 | LOC100857883 | LOC100857883 | 5 | 12 | 1 | 3 | 1 | 1 | 11 | 16 | 0.001399079 | 0.012209614 |
| ENSGALG00010018358 | FBXL16 | FBXL16 | 263 | 286 | 15 | 10 | 47 | 50 | 177 | 165 | 4.56E-18 | 7.62E-16 |
| ENSGALG00010018359 | CACNG3 | CACNG3 | 229 | 547 | 81 | 147 | 176 | 144 | 373 | 265 | 0.000438687 | 0.00474828 |
| ENSGALG00010018365 | HPSE2 | HPSE2 | 71 | 27 | 11 | 13 | 22 | 13 | 71 | 60 | 0.00028281 | 0.003330738 |
| ENSGALG00010018369 | MNR2 | MNR2 | 500 | 1295 | 94 | 102 | 435 | 325 | 851 | 654 | 9.65E-14 | 9.37E-12 |
| ENSGALG00010018383 | METRN | METRN | 1830 | 1690 | 200 | 281 | 975 | 960 | 1526 | 1367 | 2.69E-24 | 7.02E-22 |
| ENSGALG00010018386 | NA | NA | 21 | 17 | 0 | 0 | 4 | 11 | 0 | 0 | 8.18E-09 | 3.56E-07 |
| ENSGALG00010018390 | SLC12A1 | SLC12A1 | 3 | 2 | 20 | 13 | 19 | 16 | 17 | 24 | 0.001256466 | 0.011210922 |
| ENSGALG00010018397 | FAM173A | C16orf24 | 49 | 56 | 6 | 2 | 18 | 31 | 50 | 36 | 3.93E-06 | 8.75E-05 |
| ENSGALG00010018398 | LRFN5 | LRFN5 | 464 | 564 | 1435 | 1753 | 1035 | 871 | 629 | 560 | 9.49E-11 | 5.83E-09 |
| ENSGALG00010018403 | CCDC78 | CCDC78 | 43 | 36 | 2 | 4 | 23 | 23 | 56 | 27 | 2.73E-06 | 6.35E-05 |
| ENSGALG00010018413 | SLC15A2 | PEPT2 | 10 | 10 | 10 | 7 | 20 | 22 | 26 | 30 | 0.009450401 | 0.055727427 |
| ENSGALG00010018414 | EPAS1 | HIF2a | 932 | 641 | 236 | 257 | 389 | 380 | 887 | 384 | 0.00019079 | 0.002399163 |
| ENSGALG00010018420 | RHOQ | CHTC10 | 286 | 291 | 273 | 218 | 380 | 274 | 816 | 746 | 4.28E-07 | 1.23E-05 |
| ENSGALG00010018423 | IGFALS | IGFALS | 91 | 98 | 1 | 1 | 34 | 41 | 99 | 50 | 3.21E-13 | 2.87E-11 |
| ENSGALG00010018425 | CYP26B1 | CYP26B1 | 217 | 120 | 490 | 874 | 765 | 326 | 303 | 379 | 2.82E-05 | 0.00047482 |
| ENSGALG00010018458 | SEMA5B | SEMA5B | 825 | 692 | 78 | 109 | 324 | 358 | 920 | 486 | 2.03E-13 | 1.88E-11 |
| ENSGALG00010018466 | DYSF | DYSF | 2281 | 4007 | 2403 | 2705 | 2659 | 1420 | 1663 | 1262 | 0.000730574 | 0.007271804 |
| ENSGALG00010018471 | HHEX | PROBOX | 1782 | 1771 | 741 | 1018 | 1126 | 940 | 784 | 649 | 1.56E-07 | 5.04E-06 |
| ENSGALG00010018484 | NLGN1 | neuroligin-1 | 268 | 659 | 308 | 263 | 218 | 125 | 1239 | 650 | 7.98E-07 | 2.12E-05 |
| ENSGALG00010018510 | MRPL43 | MRPL43 | 1188 | 1102 | 1407 | 1614 | 630 | 496 | 1025 | 1388 | 0.000447465 | 0.004818317 |
| ENSGALG00010018512 | FERMT1 | C20orf42 | 441 | 521 | 962 | 1042 | 843 | 606 | 580 | 555 | 0.000426744 | 0.004655187 |
| ENSGALG00010018514 | EAF2 | EAF2 | 3198 | 3252 | 888 | 1536 | 1132 | 1555 | 1946 | 1527 | 7.38E-06 | 0.000151043 |
| ENSGALG00010018520 | LOC421255 | CUL7 | 547 | 532 | 1289 | 1241 | 1094 | 718 | 1179 | 1233 | 2.06E-05 | 0.000360843 |
| ENSGALG00010018528 | NA | NA | 170 | 190 | 183 | 203 | 222 | 186 | 38 | 39 | 2.63E-08 | 1.01E-06 |
| ENSGALG00010018535 | FAM174B | FAM174B | 2377 | 3023 | 4564 | 4800 | 4446 | 3042 | 3545 | 2898 | 0.000279334 | 0.003303766 |
| ENSGALG00010018537 | LOC112530649 | LOC112530649 | 9397 | 7573 | 1652 | 2895 | 1998 | 3541 | 11342 | 11770 | 5.16E-14 | 5.25E-12 |
| ENSGALG00010018540 | TNFSF10 | TRAIL | 5567 | 6210 | 4371 | 5514 | 3513 | 3198 | 6681 | 6943 | 0.000555839 | 0.005798494 |
| ENSGALG00010018541 | PRPH2 | CRDS1 | 3 | 0 | 5 | 10 | 10 | 13 | 21 | 12 | 0.004211211 | 0.029520872 |
| ENSGALG00010018544 | HACD2 | PTPLB | 4082 | 3483 | 4682 | 6977 | 4524 | 3786 | 10229 | 7563 | 1.65E-06 | 4.11E-05 |
| ENSGALG00010018549 | GHSR | GHSR | 49 | 42 | 11 | 14 | 7 | 9 | 36 | 40 | 8.56E-05 | 0.001215169 |
| ENSGALG00010018558 | MSLN | MSLN | 16 | 20 | 128 | 166 | 99 | 104 | 51 | 47 | 1.08E-08 | 4.54E-07 |
| ENSGALG00010018563 | NA | NA | 2 | 4 | 6 | 4 | 10 | 7 | 19 | 17 | 0.005428273 | 0.035956124 |
| ENSGALG00010018566 | LOC107052850 | LOC107052850 | 84 | 78 | 190 | 183 | 87 | 53 | 56 | 84 | 0.006917931 | 0.043364235 |
| ENSGALG00010018571 | NA | NA | 0 | 0 | 11 | 11 | 1 | 1 | 11 | 0 | 0.002239028 | 0.017759204 |
| ENSGALG00010018573 | SEC23B | SEC23A | 2106 | 2222 | 1426 | 1685 | 1387 | 939 | 1307 | 1534 | 0.007378002 | 0.045734855 |
| ENSGALG00010018582 | RCC2 | RCC2 | 3685 | 3429 | 5200 | 5893 | 4686 | 2922 | 2970 | 3203 | 0.000393303 | 0.004364459 |
| ENSGALG00010018590 | TGFA | TGFA | 1079 | 1395 | 484 | 543 | 527 | 314 | 726 | 1085 | 9.68E-07 | 2.53E-05 |
| ENSGALG00010018594 | GLTSCR1L | BICRAL | 167 | 277 | 638 | 499 | 504 | 233 | 272 | 229 | 0.001399937 | 0.012210725 |
| ENSGALG00010018597 | NA | NA | 16 | 19 | 42 | 38 | 17 | 14 | 7 | 11 | 0.006233426 | 0.03992833 |
| ENSGALG00010018598 | CUL9 | cullin-9 | 1783 | 2193 | 1057 | 1232 | 1212 | 921 | 1310 | 1140 | 0.000625815 | 0.006388897 |
| ENSGALG00010018616 | LOC770705 | LOC770705 | 0 | 0 | 94 | 88 | 16 | 9 | 0 | 0 | 3.46E-26 | 1.05E-23 |
| ENSGALG00010018624 | NA | NA | 13 | 40 | 9 | 6 | 58 | 33 | 66 | 67 | 1.51E-05 | 0.00027894 |
| ENSGALG00010018649 | HAGH | HAGH | 149 | 255 | 452 | 414 | 260 | 207 | 120 | 151 | 0.000428182 | 0.004655675 |
| ENSGALG00010018668 | OVOL3 | OVOL2 | 247 | 212 | 82 | 104 | 119 | 84 | 260 | 189 | 0.001659015 | 0.013954892 |
| ENSGALG00010018672 | PKDCCA | PKDCC | 3123 | 2941 | 1893 | 1890 | 1785 | 1274 | 2081 | 2089 | 0.000363164 | 0.004098034 |
| ENSGALG00010018696 | NA | NA | 89 | 90 | 30 | 64 | 76 | 85 | 164 | 144 | 0.004927527 | 0.033447944 |
| ENSGALG00010018721 | LRCH2 | LRCH2 | 286 | 273 | 129 | 142 | 154 | 146 | 384 | 227 | 0.008188032 | 0.04970519 |
| ENSGALG00010018724 | RRBP1 | RRBP1 | 2647 | 3784 | 1901 | 2023 | 1821 | 1113 | 2467 | 3440 | 0.000117392 | 0.001579598 |
| ENSGALG00010018726 | EBF2 | EBF2 | 36 | 137 | 161 | 196 | 279 | 80 | 49 | 58 | 0.005748968 | 0.037603426 |
| ENSGALG00010018753 | SYNM | DMN | 5 | 7 | 16 | 13 | 21 | 13 | 41 | 32 | 0.001050603 | 0.009727246 |
| ENSGALG00010018763 | NA | NA | 0 | 4 | 5 | 2 | 1 | 1 | 12 | 24 | 0.000469391 | 0.00502528 |
| ENSGALG00010018796 | PER3 | gPER3 | 980 | 873 | 663 | 730 | 1288 | 830 | 913 | 762 | 0.005995074 | 0.038736922 |
| ENSGALG00010018808 | KIAA0895 | KIAA0895 | 227 | 180 | 661 | 620 | 445 | 256 | 210 | 193 | 9.61E-07 | 2.51E-05 |
| ENSGALG00010018823 | RAB40C | RAB40C | 1553 | 2233 | 1623 | 1558 | 1275 | 852 | 1919 | 2449 | 0.009869582 | 0.057711848 |
| ENSGALG00010018838 | WFIKKN1 | WFIKKN1 | 11 | 98 | 47 | 19 | 14 | 7 | 93 | 129 | 0.00120602 | 0.010853528 |
| ENSGALG00010018839 | RPL3L | RPL3L | 71 | 71 | 27 | 36 | 48 | 36 | 115 | 100 | 0.001687297 | 0.014141477 |
| ENSGALG00010018847 | CGRRF1 | CGRRF1 | 419 | 554 | 406 | 478 | 637 | 480 | 314 | 331 | 0.007809887 | 0.047914405 |
| ENSGALG00010018851 | NA | NA | 1237 | 2330 | 855 | 840 | 982 | 1042 | 1567 | 1123 | 0.003529228 | 0.025593546 |
| ENSGALG00010018854 | LYSMD4 | LYSMD4 | 1801 | 1695 | 1091 | 1352 | 866 | 752 | 1163 | 1494 | 0.005007362 | 0.033780085 |
| ENSGALG00010018855 | SYNDIG1 | C20orf39 | 871 | 1197 | 361 | 231 | 423 | 312 | 845 | 1193 | 1.65E-09 | 8.28E-08 |
| ENSGALG00010018858 | EPB41L5 | EPB41L5 | 2756 | 2254 | 1567 | 1382 | 2447 | 2935 | 3499 | 2181 | 0.000292727 | 0.003418622 |
| ENSGALG00010018884 | NA | NA | 18 | 14 | 6 | 1 | 20 | 24 | 14 | 4 | 0.005273216 | 0.035189627 |
| ENSGALG00010018886 | NA | NA | 237 | 297 | 252 | 222 | 265 | 350 | 474 | 625 | 0.003564849 | 0.025812721 |
| ENSGALG00010018901 | TCF7L2 | TCF4 | 1519 | 1830 | 1436 | 1963 | 833 | 677 | 684 | 1123 | 0.000227038 | 0.00277968 |
| ENSGALG00010018908 | NA | NA | 1354 | 561 | 242 | 422 | 75 | 117 | 754 | 653 | 1.65E-09 | 8.27E-08 |

| | | | | | | | | | | | |
|---|---|---|---|---|---|---|---|---|---|---|---|
| ENSGALG00010018939 | NA | NA | 237 | 210 | 106 | 93 | 123 | 133 | 290 | 220 | 0.005338086 | 0.035513569 |
| ENSGALG00010018943 | ACKR4 | CCRL1 | 4 | 6 | 2 | 4 | 0 | 1 | 14 | 11 | 0.001911861 | 0.015593474 |
| ENSGALG00010018944 | NEFL | NEFL | 97 | 195 | 294 | 99 | 191 | 97 | 422 | 805 | 8.97E-05 | 0.001264121 |
| ENSGALG00010018946 | NA | NA | 43 | 181 | 61 | 49 | 59 | 26 | 239 | 175 | 0.000772887 | 0.007584553 |
| ENSGALG00010018958 | PELI3 | PELI1 | 553 | 715 | 1062 | 1483 | 999 | 1195 | 802 | 851 | 0.000561531 | 0.005843275 |
| ENSGALG00010018959 | UBA5 | UBAL | 973 | 877 | 720 | 737 | 581 | 482 | 1117 | 1252 | 0.002438501 | 0.019000254 |
| ENSGALG00010018973 | NEFM | NEF3 | 425 | 378 | 459 | 312 | 321 | 392 | 979 | 1462 | 2.59E-07 | 7.90E-06 |
| ENSGALG00010018983 | MDGA2 | MDGA2 | 1571 | 1346 | 1619 | 2061 | 742 | 747 | 2270 | 1900 | 5.88E-05 | 0.00087607 |
| ENSGALG00010018986 | CTDSP1 | CTDSP1 | 588 | 654 | 417 | 560 | 216 | 243 | 444 | 747 | 0.008096014 | 0.049307795 |
| ENSGALG00010019001 | ALDH1A3 | ALDH6 | 53 | 51 | 79 | 80 | 132 | 82 | 178 | 226 | 7.87E-05 | 0.001131734 |
| ENSGALG00010019011 | VIL1 | VIL | 101 | 95 | 14 | 45 | 26 | 23 | 51 | 46 | 0.00189155 | 0.015450432 |
| ENSGALG00010019013 | LRRK1 | LRRK1 | 860 | 1545 | 1246 | 2479 | 405 | 336 | 1164 | 2702 | 1.29E-06 | 3.28E-05 |
| ENSGALG00010019019 | TBX20 | TBX20 | 28 | 70 | 45 | 51 | 367 | 170 | 140 | 336 | 1.30E-08 | 5.35E-07 |
| ENSGALG00010019020 | KCNK2 | KCNK2 | 76 | 85 | 12 | 20 | 34 | 32 | 93 | 43 | 0.000312781 | 0.003609944 |
| ENSGALG00010019041 | KATNIP | KIAA0556 | 193 | 179 | 403 | 540 | 393 | 322 | 347 | 288 | 0.001719252 | 0.014351714 |
| ENSGALG00010019046 | SELENOS | SELS | 1769 | 1553 | 951 | 1098 | 885 | 836 | 1237 | 1366 | 0.00576471 | 0.037644173 |
| ENSGALG00010019059 | BRWD3 | BRWD3 | 307 | 608 | 1120 | 1377 | 783 | 879 | 1320 | 1065 | 8.19E-05 | 0.001171973 |
| ENSGALG00010019068 | SNRPA1 | SNRPA1 | 4814 | 3493 | 3981 | 5163 | 2986 | 2287 | 2155 | 2202 | 2.43E-05 | 0.000417042 |
| ENSGALG00010019072 | AEBP1 | AEBP1 | 1917 | 1764 | 525 | 686 | 1004 | 847 | 1269 | 1223 | 7.99E-10 | 4.24E-08 |
| ENSGALG00010019076 | ESPN | ESPN | 36 | 74 | 17 | 7 | 1 | 21 | 52 | 95 | 0.002940316 | 0.022074823 |
| ENSGALG00010019079 | CAPG | CAPG | 89 | 360 | 54 | 32 | 77 | 37 | 43 | 58 | 5.75E-05 | 0.000861649 |
| ENSGALG00010019081 | PCSK6 | PACE4 | 1251 | 1370 | 952 | 1178 | 1014 | 1147 | 2409 | 2140 | 5.41E-05 | 0.00082121 |
| ENSGALG00010019082 | POLR3K | POLR3K | 650 | 419 | 519 | 676 | 255 | 198 | 638 | 798 | 0.000951965 | 0.008962956 |
| ENSGALG00010019085 | NA | NA | 173 | 259 | 443 | 502 | 470 | 251 | 250 | 214 | 0.002955344 | 0.022167735 |
| ENSGALG00010019101 | POLE2 | POLE2 | 1081 | 1109 | 1264 | 1359 | 650 | 474 | 1185 | 1196 | 0.000882385 | 0.008431338 |
| ENSGALG00010019114 | FAM46D | TENT5D | 1 | 3 | 31 | 16 | 17 | 6 | 8 | 16 | 0.003036505 | 0.022684881 |
| ENSGALG00010019141 | ACOT7 | ACOT7 | 1755 | 2244 | 1224 | 1344 | 1036 | 893 | 1693 | 2126 | 0.000535019 | 0.00561989 |
| ENSGALG00010019145 | RAC1 | CRAC1A | 6149 | 6464 | 8411 | 9771 | 7168 | 4902 | 5001 | 5807 | 0.00234503 | 0.018410463 |
| ENSGALG00010019159 | NA | NA | 4 | 5 | 1 | 0 | 17 | 8 | 2 | 2 | 0.000115969 | 0.00156675 |
| ENSGALG00010019169 | GPR153 | GPR153 | 192 | 224 | 95 | 79 | 127 | 96 | 103 | 106 | 0.00931552 | 0.055088806 |
| ENSGALG00010019174 | KCNIP3 | KCNIP3 | 295 | 301 | 705 | 605 | 539 | 385 | 474 | 542 | 0.008341819 | 0.050510321 |
| ENSGALG00010019257 | SLC1A4 | SLC1A4 | 3318 | 2660 | 3550 | 4329 | 2742 | 3528 | 6868 | 5985 | 0.000157493 | 0.002035598 |
| ENSGALG00010019264 | HK2 | hexokinase-2 | 1026 | 1502 | 424 | 579 | 641 | 423 | 775 | 749 | 3.89E-06 | 8.66E-05 |
| ENSGALG00010019270 | NA | NA | 17 | 51 | 8 | 2 | 0 | 4 | 4 | 2 | 3.97E-05 | 0.000632582 |
| ENSGALG00010019277 | IGFBP5 | IGFBP5 | 591 | 697 | 1240 | 1952 | 1036 | 451 | 444 | 913 | 0.001573962 | 0.01337948 |
| ENSGALG00010019278 | HN1L | JPT2 | 6316 | 6541 | 4469 | 5297 | 4432 | 3937 | 6344 | 7423 | 0.00882708 | 0.052854949 |
| ENSGALG00010019281 | LOC421195 | LOC421195 | 451 | 379 | 35 | 200 | 254 | 341 | 390 | 402 | 0.003995464 | 0.028305298 |
| ENSGALG00010019282 | IGFBP2 | IGFBP-2 | 2139 | 957 | 326 | 239 | 1229 | 687 | 2216 | 1457 | 1.05E-10 | 6.39E-09 |
| ENSGALG00010019315 | PRRT3 | PRRT3 | 144 | 349 | 41 | 63 | 34 | 62 | 132 | 255 | 2.69E-05 | 0.000455947 |
| ENSGALG00010019327 | NA | NA | 7 | 4 | 32 | 17 | 23 | 33 | 25 | 14 | 0.006613409 | 0.041845784 |
| ENSGALG00010019341 | NA | NA | 3 | 4 | 39 | 31 | 2 | 4 | 12 | 7 | 5.43E-06 | 0.000115651 |
| ENSGALG00010019346 | POU3F4 | POU3F3 | 536 | 972 | 181 | 209 | 220 | 88 | 193 | 226 | 2.17E-09 | 1.06E-07 |
| ENSGALG00010019347 | NA | NA | 659 | 715 | 457 | 636 | 944 | 1076 | 1367 | 1347 | 6.48E-07 | 1.77E-05 |
| ENSGALG00010019376 | CIITA | CIITA | 11 | 11 | 54 | 26 | 51 | 31 | 23 | 11 | 0.003090751 | 0.022997611 |
| ENSGALG00010019394 | NA | NA | 16 | 8 | 30 | 34 | 43 | 69 | 75 | 19 | 0.006226489 | 0.039899189 |
| ENSGALG00010019399 | CD248 | C1QR1 | 41 | 38 | 168 | 157 | 90 | 80 | 73 | 35 | 0.000247799 | 0.00298358 |
| ENSGALG00010019404 | NA | NA | 2 | 12 | 50 | 65 | 17 | 9 | 10 | 10 | 9.09E-06 | 0.000180772 |
| ENSGALG00010019426 | GRM7 | GRM7 | 2316 | 2204 | 403 | 669 | 979 | 831 | 1893 | 2108 | 4.36E-16 | 5.88E-14 |
| ENSGALG00010019431 | NA | NA | 124 | 220 | 458 | 387 | 372 | 205 | 212 | 253 | 0.009750666 | 0.057136443 |
| ENSGALG00010019440 | NA | NA | 8 | 4 | 33 | 38 | 36 | 2 | 1 | 2 | 0.000153285 | 0.001987355 |
| ENSGALG00010019441 | LMCD1 | LMCD1 | 73 | 265 | 145 | 183 | 63 | 63 | 205 | 439 | 0.004185972 | 0.029405619 |
| ENSGALG00010019442 | NA | NA | 0 | 3 | 11 | 11 | 11 | 2 | 1 | 1 | 0.006979178 | 0.043602251 |
| ENSGALG00010019515 | NENF | NENF | 1020 | 955 | 1319 | 1431 | 1099 | 804 | 732 | 767 | 0.007232532 | 0.044899732 |
| ENSGALG00010019532 | NA | NA | 9 | 8 | 22 | 22 | 46 | 28 | 40 | 20 | 0.003046601 | 0.022739975 |
| ENSGALG00010019534 | PPP2R5A | PPP2R5A | 4542 | 4467 | 4574 | 5382 | 5984 | 3206 | 3235 | 3264 | 0.004852641 | 0.033038995 |
| ENSGALG00010019541 | NA | NA | 0 | 0 | 10 | 10 | 11 | 6 | 6 | 3 | 0.000180543 | 0.002292734 |
| ENSGALG00010019546 | HSD17B12 | HSD17B12 | 6575 | 7770 | 4615 | 5475 | 4075 | 3668 | 5550 | 6555 | 0.002841553 | 0.021439294 |
| ENSGALG00010019566 | NA | NA | 316 | 607 | 1339 | 1529 | 729 | 742 | 691 | 1130 | 0.000107564 | 0.001467431 |
| ENSGALG00010019585 | PDE1C | PDE1C | 23 | 20 | 95 | 66 | 30 | 20 | 43 | 54 | 0.003330545 | 0.024427909 |
| ENSGALG00010019594 | TMX1 | TXNDC1 | 40 | 63 | 146 | 175 | 166 | 113 | 164 | 143 | 0.000910578 | 0.008656184 |
| ENSGALG00010019610 | CORO7 | coronin-7 | 309 | 471 | 849 | 932 | 660 | 468 | 532 | 551 | 0.001963278 | 0.0159296 |
| ENSGALG00010019611 | NA | NA | 423 | 838 | 311 | 301 | 665 | 646 | 661 | 601 | 0.000427798 | 0.004655675 |
| ENSGALG00010019617 | ANO3 | TMEM16C | 419 | 271 | 137 | 152 | 294 | 196 | 494 | 292 | 0.002535022 | 0.019559225 |
| ENSGALG00010019621 | LOC421441 | SPTBB4L | 11172 | 13672 | 8154 | 7814 | 4095 | 2479 | 7343 | 8752 | 1.48E-14 | 1.62E-12 |
| ENSGALG00010019634 | GNG2 | GNG2 | 1077 | 1383 | 688 | 894 | 703 | 527 | 904 | 1257 | 0.009926917 | 0.058006516 |
| ENSGALG00010019641 | CAPN11 | CANP | 6219 | 6606 | 5576 | 5990 | 2720 | 2581 | 5596 | 8424 | 4.49E-06 | 9.83E-05 |
| ENSGALG00010019657 | PPP1R17 | C2H7orf16 | 900 | 3669 | 155 | 266 | 274 | 383 | 797 | 1467 | 2.25E-11 | 1.58E-09 |
| ENSGALG00010019671 | NA | NA | 13 | 7 | 36 | 49 | 27 | 13 | 11 | 12 | 0.003090601 | 0.022997611 |
| ENSGALG00010019673 | NID2 | NID2 | 3162 | 4267 | 1734 | 2690 | 2359 | 2303 | 3194 | 3227 | 0.004294401 | 0.029978289 |
| ENSGALG00010019683 | TMEM184A | TMEM184A | 96 | 189 | 54 | 94 | 49 | 31 | 49 | 46 | 0.00227113 | 0.01798821 |
| ENSGALG00010019685 | NA | NA | 1428 | 1813 | 838 | 706 | 989 | 741 | 1353 | 1692 | 2.86E-05 | 0.000478849 |
| ENSGALG00010019726 | PEBP4 | PEBP4 | 37 | 13 | 6 | 4 | 4 | 2 | 15 | 21 | 0.001004082 | 0.009379665 |
| ENSGALG00010019729 | NA | NA | 6 | 20 | 11 | 8 | 5 | 3 | 2 | 0 | 0.008941923 | 0.053427752 |
| ENSGALG00010019738 | ADRA2A | ADRA2A | 34 | 39 | 172 | 119 | 93 | 61 | 96 | 123 | 0.001119277 | 0.010225099 |
| ENSGALG00010019755 | NA | NA | 45 | 19 | 3 | 12 | 80 | 38 | 42 | 25 | 0.000189698 | 0.002390827 |
| ENSGALG00010019774 | NA | NA | 90 | 74 | 203 | 337 | 248 | 171 | 219 | 132 | 0.001515343 | 0.012960289 |
| ENSGALG00010019801 | FIBIN | FIBIN | 4180 | 2491 | 962 | 1058 | 4455 | 2085 | 3944 | 1507 | 5.40E-06 | 0.000115095 |
| ENSGALG00010019820 | BBOX1 | BBOX1 | 854 | 515 | 385 | 473 | 1267 | 638 | 839 | 412 | 0.002726869 | 0.020780409 |
| ENSGALG00010019830 | JAG2 | JAG2 | 202 | 241 | 495 | 691 | 429 | 366 | 229 | 179 | 5.55E-06 | 0.000117645 |
| ENSGALG00010019836 | MINAR1 | DD1 | 7 | 38 | 3 | 4 | 0 | 2 | 20 | 14 | 0.000255193 | 0.00306158 |
| ENSGALG00010019842 | NA | NA | 8 | 9 | 34 | 39 | 15 | 9 | 11 | 5 | 0.001659189 | 0.013954892 |
| ENSGALG00010019860 | SIX6 | OPTX2 | 5 | 166 | 874 | 533 | 571 | 102 | 69 | 203 | 0.004939827 | 0.033490625 |
| ENSGALG00010019896 | DDB1 | DDB1 | 8353 | 9739 | 5581 | 6092 | 6669 | 4381 | 7042 | 9010 | 0.001675785 | 0.01407323 |
| ENSGALG00010019906 | ZBTB42 | ZBTB42 | 475 | 719 | 961 | 1245 | 939 | 474 | 549 | 520 | 0.00556398 | 0.036709525 |
| ENSGALG00010019919 | LIN7C | LIN7B | 1556 | 3112 | 3394 | 4688 | 2043 | 1408 | 2255 | 2283 | 0.00172317 | 0.014377242 |
| ENSGALG00010019938 | FAM19A4 | FAM19A4 | 223 | 299 | 41 | 57 | 42 | 59 | 101 | 111 | 8.99E-08 | 3.11E-06 |
| ENSGALG00010019947 | LGALSL2 | LEGLA | 64 | 56 | 232 | 232 | 255 | 183 | 179 | 132 | 1.22E-05 | 0.000232178 |
| ENSGALG00010019982 | PLCE1 | PLCE1 | 781 | 1283 | 606 | 566 | 565 | 520 | 1319 | 803 | 0.006091446 | 0.03923187 |
| ENSGALG00010019985 | UMPS | UMPS | 9231 | 10902 | 6265 | 6296 | 5049 | 4372 | 7019 | 8397 | 1.84E-05 | 0.000329012 |
| ENSGALG00010019996 | ITGB5 | ITGB5 | 4449 | 5316 | 2137 | 2275 | 2851 | 2156 | 5660 | 4504 | 7.54E-12 | 5.80E-10 |
| ENSGALG00010019997 | FSHR | FSHR | 27 | 27 | 28 | 39 | 10 | 2 | 3 | 3 | 8.24E-06 | 0.000165837 |
| ENSGALG00010020005 | EOGT | AER61 | 2114 | 2792 | 655 | 888 | 1392 | 1054 | 952 | 899 | 2.20E-13 | 2.01E-11 |
| ENSGALG00010020007 | NA | NA | 1189 | 1064 | 1134 | 1347 | 1569 | 1513 | 2500 | 1628 | 0.001392022 | 0.012154373 |
| ENSGALG00010020009 | MACROD2 | MACROD2 | 222 | 254 | 447 | 361 | 514 | 436 | 591 | 391 | 0.00431991 | 0.030093509 |
| ENSGALG00010020015 | ATP6V1D | ATP6V1D | 2758 | 3169 | 3669 | 4573 | 3306 | 2369 | 2014 | 2560 | 0.001834864 | 0.015112986 |
| ENSGALG00010020021 | LOC112532694 | LOC112532694 | 363 | 336 | 62 | 70 | 210 | 245 | 419 | 216 | 4.07E-07 | 1.18E-05 |
| ENSGALG00010020023 | FOSL2 | fra-2 | 259 | 265 | 947 | 858 | 534 | 371 | 409 | 540 | 2.08E-06 | 5.01E-05 |
| ENSGALG00010020041 | CDH23 | CDH23 | 156 | 363 | 80 | 105 | 96 | 97 | 332 | 252 | 0.000239573 | 0.002897061 |
| ENSGALG00010020051 | NA | NA | 4 | 5 | 23 | 8 | 30 | 24 | 17 | 27 | 0.003350583 | 0.024505448 |
| ENSGALG00010020057 | RBKS | RBKS | 51 | 47 | 164 | 178 | 62 | 53 | 53 | 50 | 9.81E-05 | 0.001359043 |
| ENSGALG00010020089 | CASP14 | CASP17 | 10 | 20 | 0 | 7 | 25 | 23 | 11 | 21 | 0.004420947 | 0.030644075 |
| ENSGALG00010020094 | CLEC3B | CLEC3B | 40 | 39 | 7 | 48 | 128 | 72 | 94 | 103 | 0.001389317 | 0.012139676 |
| ENSGALG00010020095 | LRFN2 | LRFN2 | 154 | 99 | 290 | 515 | 76 | 92 | 183 | 74 | 0.00030982 | 0.003580709 |
| ENSGALG00010020100 | MOCS1 | MOCS1B | 477 | 512 | 1021 | 996 | 923 | 933 | 1563 | 1421 | 9.35E-08 | 3.22E-06 |
| ENSGALG00010020109 | WDFY4 | WDFY4 | 44 | 44 | 114 | 158 | 107 | 66 | 60 | 60 | 0.003642619 | 0.026228113 |
| ENSGALG00010020112 | AMZ2 | AMZ1 | 49 | 43 | 7 | 1 | 14 | 12 | 50 | 30 | 3.45E-06 | 7.79E-05 |
| ENSGALG00010020125 | NAV2 | NAV2 | 721 | 801 | 537 | 591 | 405 | 379 | 1430 | 762 | 0.000691455 | 0.006927815 |
| ENSGALG00010020136 | KCNK17 | KCNK17 | 102 | 55 | 13 | 11 | 39 | 34 | 84 | 42 | 0.000118051 | 0.001585903 |
| ENSGALG00010020143 | KCNK5 | KCNK5 | 366 | 607 | 182 | 137 | 527 | 266 | 440 | 204 | 0.000873582 | 0.008366362 |
| ENSGALG00010020153 | AXL | REK | 1222 | 1678 | 582 | 583 | 1016 | 746 | 1134 | 1300 | 1.82E-06 | 4.46E-05 |
| ENSGALG00010020155 | MFGE8 | MFGE8 | 1082 | 2643 | 1099 | 1167 | 864 | 922 | 2416 | 4163 | 2.42E-06 | 5.70E-05 |
| ENSGALG00010020170 | SORCS1 | SORCS1 | 4804 | 4174 | 4067 | 4520 | 4005 | 4165 | 7287 | 6135 | 0.007582766 | 0.046726837 |
| ENSGALG00010020176 | LOC112533469 | LOC112533469 | 570 | 186 | 86 | 94 | 184 | 179 | 547 | 274 | 0.000184569 | 0.002331963 |
| ENSGALG00010020183 | XPNPEP1 | XPNPEP1 | 3128 | 5179 | 6379 | 7047 | 5178 | 4282 | 4001 | 4398 | 0.00511079 | 0.034260538 |
| ENSGALG00010020194 | PRICKLE2 | PRICKLE2 | 912 | 1474 | 486 | 615 | 320 | 368 | 958 | 1086 | 5.05E-07 | 1.42E-05 |
| ENSGALG00010020207 | DRGX | Drg11 | 494 | 148 | 42 | 41 | 337 | 404 | 670 | 277 | 5.57E-08 | 1.99E-06 |
| ENSGALG00010020211 | ADD3 | gamma-adducin | 731 | 1127 | 692 | 666 | 648 | 468 | 1020 | 1244 | 0.009108521 | 0.054229284 |
| ENSGALG00010020246 | ADAMTS9 | ADAMTS9 | 3216 | 4770 | 2656 | 2980 | 1908 | 1435 | 2814 | 2911 | 4.52E-05 | 0.00070886 |
| ENSGALG00010020252 | DMTN | EPB49 | 1305 | 987 | 409 | 580 | 647 | 360 | 747 | 611 | 0.000114391 | 0.001547933 |
| ENSGALG00010020254 | LOC419409 | LOC419409 | 423 | 572 | 447 | 414 | 405 | 300 | 793 | 1141 | 8.23E-05 | 0.001174914 |
| ENSGALG00010020269 | UGT8L | UGT8L | 193 | 178 | 20 | 65 | 59 | 117 | 191 | 138 | 0.001505363 | 0.012916144 |
| ENSGALG00010020282 | MAGI1 | MAGI1 | 3242 | 2659 | 1340 | 1568 | 1476 | 994 | 3036 | 1673 | 5.58E-05 | 0.000840588 |
| ENSGALG00010020287 | GALNT16 | GALNTL1 | 158 | 147 | 335 | 467 | 329 | 237 | 330 | 242 | 0.004966243 | 0.033623323 |
| ENSGALG00010020306 | NA | NA | 6 | 2 | 112 | 112 | 144 | 48 | 37 | 24 | 1.93E-11 | 1.37E-09 |
| ENSGALG00010020307 | SHQ1 | SHQ1 | 1279 | 1739 | 1458 | 2122 | 824 | 705 | 974 | 992 | 0.000399813 | 0.004427765 |
| ENSGALG00010020326 | LOC101751780 | LOC101751780 | 16 | 5 | 22 | 22 | 77 | 72 | 18 | 52 | 3.58E-05 | 0.00058114 |
| ENSGALG00010020337 | NA | NA | 7 | 2 | 0 | 0 | 0 | 16 | 22 | 21 | 0.000402305 | 0.004440764 |
| ENSGALG00010020339 | NA | NA | 6 | 3 | 2 | 0 | 1 | 3 | 18 | 14 | 0.00058157 | 0.006010631 |
| ENSGALG00010020345 | AGBL1 | AGBL1 | 118 | 310 | 56 | 92 | 101 | 75 | 301 | 324 | 3.50E-05 | 0.000569289 |
| ENSGALG00010020372 | NA | NA | 3 | 4 | 17 | 16 | 15 | 3 | 0 | 0 | 1.99E-05 | 0.000350993 |
| ENSGALG00010020379 | COL13A1 | COL13A1 | 92 | 105 | 23 | 40 | 26 | 6 | 57 | 63 | 0.000116241 | 0.001567894 |

| | | | | | | | | | | | | |
|---|---|---|---|---|---|---|---|---|---|---|---|---|
| ENSGALG00010020382 | MFAP2 | MFAP2 | 96 | 966 | 114 | 29 | 22 | 17 | 217 | 621 | 4.08E-07 | 1.18E-05 |
| ENSGALG00010020389 | ACTN1 | ACTN1 | 3604 | 2050 | 5338 | 6218 | 4196 | 3919 | 6513 | 4922 | 0.001787456 | 0.0148101 |
| ENSGALG00010020436 | CNTN4 | FAR-1 | 6970 | 6004 | 2029 | 3643 | 4329 | 5051 | 10245 | 5312 | 2.33E-05 | 0.000402451 |
| ENSGALG00010020440 | PAX7 | PAX-7 | 2 | 13 | 23 | 25 | 38 | 29 | 11 | 16 | 0.003776836 | 0.027031467 |
| ENSGALG00010020451 | NA | NA | 555 | 628 | 867 | 1250 | 597 | 491 | 343 | 612 | 0.006951479 | 0.043445443 |
| ENSGALG00010020471 | NA | NA | 18 | 10 | 63 | 67 | 45 | 54 | 30 | 68 | 0.004073056 | 0.028733181 |
| ENSGALG00010020476 | TMEM132A | TMEM132A | 3945 | 3940 | 2058 | 2649 | 2837 | 1690 | 2736 | 3186 | 0.000909341 | 0.008649347 |
| ENSGALG00010020532 | SIX3 | CSIX3 | 28 | 149 | 676 | 478 | 382 | 360 | 121 | 1080 | 0.003715496 | 0.026649517 |
| ENSGALG00010020545 | STYX | STYX | 760 | 1113 | 1259 | 1445 | 1370 | 899 | 631 | 784 | 0.001652509 | 0.013919725 |
| ENSGALG00010020557 | NA | NA | 5 | 3 | 9 | 16 | 33 | 18 | 25 | 30 | 0.000290367 | 0.003393434 |
| ENSGALG00010020562 | LRRN1 | LRCH4 | 5523 | 5761 | 3010 | 3659 | 3214 | 2600 | 5328 | 5571 | 7.34E-07 | 1.98E-05 |
| ENSGALG00010020570 | NA | NA | 81 | 237 | 29 | 100 | 6 | 6 | 63 | 251 | 7.24E-07 | 1.95E-05 |
| ENSGALG00010020572 | PPP1R3C | PPP1R3C | 44 | 68 | 20 | 16 | 20 | 13 | 31 | 33 | 0.00610415 | 0.039296054 |
| ENSGALG00010020588 | PALD1 | PALD | 8305 | 8612 | 2058 | 3636 | 3482 | 3331 | 5028 | 6995 | 2.72E-10 | 1.55E-08 |
| ENSGALG00010020618 | NA | NA | 27 | 16 | 171 | 177 | 183 | 149 | 136 | 118 | 2.43E-09 | 1.17E-07 |
| ENSGALG00010020619 | QARS | QARS | 6111 | 7359 | 4813 | 5127 | 3887 | 3508 | 6354 | 8686 | 0.000338444 | 0.003860799 |
| ENSGALG00010020622 | ABCG2 | ABCG2 | 441 | 492 | 205 | 229 | 253 | 231 | 227 | 231 | 0.001939635 | 0.015766136 |
| ENSGALG00010020623 | NEK2 | NEK2 | 1648 | 1767 | 1006 | 904 | 1122 | 779 | 1185 | 955 | 0.001121684 | 0.010232309 |
| ENSGALG00010020629 | TRIM66 | TRIM66 | 119 | 161 | 46 | 38 | 46 | 29 | 70 | 28 | 0.000583674 | 0.006022154 |
| ENSGALG00010020640 | PTPN14 | PTPN14 | 186 | 222 | 157 | 184 | 173 | 237 | 701 | 298 | 0.002033618 | 0.0164421 |
| ENSGALG00010020649 | MYH11 | MYH11 | 258 | 289 | 524 | 485 | 499 | 461 | 350 | 221 | 0.002830646 | 0.021385966 |
| ENSGALG00010020651 | LMO1 | LMO-1 | 92 | 184 | 380 | 350 | 274 | 101 | 68 | 89 | 6.47E-05 | 0.000948109 |
| ENSGALG00010020658 | NA | NA | 2 | 0 | 16 | 15 | 12 | 15 | 10 | 7 | 0.000606837 | 0.006217935 |
| ENSGALG00010020664 | ESYT3 | FAM62C | 172 | 78 | 54 | 90 | 34 | 27 | 233 | 78 | 0.003826605 | 0.027329157 |
| ENSGALG00010020695 | LOC395991 | LOC395991 | 15 | 4 | 16 | 22 | 13 | 9 | 24 | 67 | 0.00962043 | 0.056492138 |
| ENSGALG00010020700 | MTHFD2 | MTHFD2 | 1618 | 1198 | 970 | 948 | 887 | 1038 | 1610 | 2116 | 0.005984845 | 0.038710496 |
| ENSGALG00010020716 | RRAS | RRAS2 | 877 | 1020 | 1568 | 2047 | 1355 | 1055 | 997 | 1100 | 0.000983389 | 0.009212109 |
| ENSGALG00010020719 | RPH3A | RPH3A | 203 | 161 | 70 | 82 | 111 | 72 | 245 | 385 | 7.54E-06 | 0.000153338 |
| ENSGALG00010020744 | CHST3 | CHST3 | 541 | 636 | 133 | 205 | 137 | 298 | 644 | 731 | 5.39E-06 | 0.000114941 |
| ENSGALG00010020753 | SYDE2 | SYDE2 | 242 | 201 | 386 | 374 | 297 | 284 | 493 | 560 | 0.007469401 | 0.046215709 |
| ENSGALG00010020756 | H2AFY2 | H2AFY2 | 3826 | 4196 | 2155 | 3553 | 1627 | 1856 | 2739 | 3622 | 0.004224254 | 0.029587484 |
| ENSGALG00010020765 | GLT1D1 | GLT1D1 | 2243 | 1370 | 1058 | 1401 | 1375 | 1615 | 2596 | 2334 | 0.006881538 | 0.043169771 |
| ENSGALG00010020770 | B4GALNT4 | B4GALNT4 | 5343 | 5998 | 5322 | 4824 | 4240 | 3136 | 6500 | 7357 | 0.00457637 | 0.031564298 |
| ENSGALG00010020778 | SLIT1 | SLIT1 | 783 | 2708 | 577 | 490 | 631 | 770 | 1693 | 2086 | 1.65E-05 | 0.000299362 |
| ENSGALG00010020781 | SNN | SNN | 375 | 285 | 124 | 173 | 153 | 133 | 354 | 211 | 0.008160583 | 0.049556562 |
| ENSGALG00010020797 | DGCR8 | DGCR8 | 2541 | 2903 | 2108 | 2181 | 1969 | 1876 | 4303 | 3157 | 0.000869 | 0.008341597 |
| ENSGALG00010020807 | NA | NA | 8 | 35 | 6 | 1 | 7 | 6 | 27 | 33 | 0.001555287 | 0.013247688 |
| ENSGALG00010020810 | IFITM5 | Dspa1 | 497 | 399 | 273 | 299 | 280 | 190 | 492 | 646 | 0.00333502 | 0.024445114 |
| ENSGALG00010020828 | CAMK2N1 | CAMK2N1 | 81 | 89 | 77 | 59 | 18 | 9 | 188 | 333 | 2.33E-12 | 1.91E-10 |
| ENSGALG00010020838 | NA | NA | 7 | 14 | 9 | 12 | 16 | 6 | 42 | 36 | 0.002708931 | 0.020694486 |
| ENSGALG00010020901 | NA | NA | 3 | 29 | 64 | 61 | 53 | 43 | 18 | 4 | 0.00174657 | 0.014536194 |
| ENSGALG00010020915 | EN1 | En-1 | 9 | 9 | 161 | 61 | 22 | 14 | 7 | 12 | 6.99E-10 | 3.73E-08 |
| ENSGALG00010020921 | LDB1 | CLIM2 | 2805 | 4185 | 2111 | 2441 | 2336 | 1506 | 2013 | 2538 | 0.004935997 | 0.033478241 |
| ENSGALG00010020945 | NUMB | NUMB | 978 | 960 | 1753 | 2107 | 1336 | 1083 | 1255 | 1121 | 0.001144981 | 0.010388127 |
| ENSGALG00010020946 | CTBS | CTBS | 160 | 221 | 71 | 72 | 71 | 65 | 136 | 102 | 0.003252444 | 0.023976549 |
| ENSGALG00010020960 | CLEC19A | CLEC19A | 85 | 406 | 52 | 36 | 20 | 14 | 41 | 47 | 1.23E-08 | 5.09E-07 |
| ENSGALG00010020962 | MYO5C | MYO5C | 52 | 32 | 29 | 18 | 11 | 3 | 59 | 35 | 0.00148588 | 0.012767106 |
| ENSGALG00010020963 | CCDC93 | CCDC93 | 638 | 730 | 1103 | 1299 | 984 | 682 | 680 | 621 | 0.002542712 | 0.019609506 |
| ENSGALG00010020965 | DHX37 | DHX37 | 5490 | 5826 | 7271 | 8601 | 9202 | 7620 | 9339 | 9047 | 6.73E-05 | 0.000984442 |
| ENSGALG00010020984 | PSEN1 | PS-1 | 849 | 1245 | 1912 | 2250 | 1751 | 1252 | 1361 | 1334 | 0.000642451 | 0.006542739 |
| ENSGALG00010021012 | KCNIP2 | KCNIP2 | 50 | 102 | 28 | 26 | 26 | 28 | 86 | 96 | 0.001132672 | 0.01029321 |
| ENSGALG00010021016 | FHIT | FHIT | 107 | 105 | 152 | 217 | 859 | 446 | 116 | 172 | 9.47E-16 | 1.24E-13 |
| ENSGALG00010021018 | F3 | F3 | 80 | 70 | 15 | 16 | 28 | 27 | 116 | 129 | 3.18E-08 | 1.20E-06 |
| ENSGALG00010021040 | FGF8 | FGF-8 | 232 | 87 | 18 | 11 | 43 | 29 | 153 | 44 | 4.72E-06 | 0.000102787 |
| ENSGALG00010021043 | PTPRG | PTPRG | 472 | 718 | 950 | 1117 | 1109 | 956 | 1125 | 1025 | 0.001087059 | 0.009998312 |
| ENSGALG00010021054 | FEZF2 | FEZF2 | 159 | 259 | 1005 | 706 | 574 | 485 | 409 | 840 | 1.66E-05 | 0.000301086 |
| ENSGALG00010021058 | CCP110 | CCP110 | 345 | 449 | 858 | 797 | 748 | 531 | 818 | 519 | 0.008616248 | 0.05177804 |
| ENSGALG00010021083 | SCD | SCD | 6811 | 6739 | 3715 | 4509 | 5785 | 5109 | 6975 | 7902 | 1.34E-05 | 0.000251534 |
| ENSGALG00010021108 | PCGF6 | PCGF6 | 2191 | 2036 | 2632 | 3705 | 2723 | 2756 | 1911 | 2130 | 0.002391857 | 0.018678841 |
| ENSGALG00010021124 | CALHM2 | FAM26B | 145 | 336 | 618 | 715 | 233 | 145 | 126 | 118 | 3.19E-08 | 1.20E-06 |
| ENSGALG00010021125 | LY75 | DEC205 | 133 | 97 | 484 | 441 | 316 | 195 | 209 | 198 | 1.12E-05 | 0.00021581 |
| ENSGALG00010021137 | PLAU | PLAU | 9 | 35 | 77 | 75 | 23 | 8 | 21 | 28 | 0.002936514 | 0.022066109 |
| ENSGALG00010021142 | WNT6 | WNT6 | 17 | 31 | 2 | 3 | 3 | 0 | 15 | 28 | 3.30E-06 | 7.50E-05 |
| ENSGALG00010021143 | ABCA4 | ABCA4 | 248 | 62 | 8 | 6 | 116 | 59 | 412 | 55 | 1.72E-06 | 4.25E-05 |
| ENSGALG00010021151 | WNT10A | WNT10A | 7 | 12 | 39 | 18 | 1 | 4 | 5 | 11 | 0.002913376 | 0.021902084 |
| ENSGALG00010021156 | DDOST | OST48 | 11621 | 10234 | 7716 | 10250 | 5796 | 4330 | 7037 | 7600 | 8.20E-05 | 0.001172725 |
| ENSGALG00010021157 | NA | NA | 3 | 0 | 10 | 19 | 2 | 0 | 2 | 5 | 0.002816098 | 0.0213146 |
| ENSGALG00010021163 | INHA | INHA | 5 | 2 | 35 | 8 | 3 | 1 | 7 | 12 | 0.005690801 | 0.037300012 |
| ENSGALG00010021172 | SLC29A4 | SLC29A4 | 3085 | 1166 | 502 | 739 | 269 | 255 | 639 | 850 | 1.17E-11 | 8.71E-10 |
| ENSGALG00010021195 | GCLM | GCLM | 3268 | 3194 | 4832 | 5230 | 4720 | 2833 | 2979 | 3414 | 0.003308176 | 0.024341537 |
| ENSGALG00010021205 | CDK5R2 | CDK5R2 | 135 | 228 | 32 | 21 | 22 | 30 | 71 | 140 | 1.07E-07 | 3.65E-06 |
| ENSGALG00010021209 | LOC396365 | LOC396365 | 49 | 134 | 1 | 7 | 34 | 35 | 41 | 48 | 6.74E-08 | 2.39E-06 |
| ENSGALG00010021215 | SH3PXD2A | SH3MD1 | 188 | 230 | 102 | 117 | 75 | 73 | 255 | 189 | 0.002224439 | 0.017677045 |
| ENSGALG00010021217 | FBXO2 | FBXO2 | 86 | 166 | 36 | 28 | 74 | 54 | 79 | 78 | 0.001296695 | 0.011477948 |
| ENSGALG00010021219 | STRA6 | STRA6 | 2 | 7 | 20 | 0 | 1 | 3 | 132 | 129 | 6.30E-11 | 4.09E-09 |
| ENSGALG00010021221 | HAP1 | HAP1 | 136 | 476 | 77 | 104 | 90 | 77 | 150 | 165 | 0.00145713 | 0.012565301 |
| ENSGALG00010021228 | ESRP2 | RBM35B | 5899 | 2036 | 702 | 378 | 1993 | 3031 | 4144 | 3298 | 6.62E-09 | 2.95E-07 |
| ENSGALG00010021234 | ISLR2 | SST273 | 229 | 857 | 91 | 28 | 114 | 26 | 616 | 295 | 5.99E-07 | 1.66E-05 |
| ENSGALG00010021237 | FSCN1 | fascin | 10037 | 10882 | 3010 | 2807 | 3769 | 3687 | 6751 | 11650 | 1.30E-15 | 1.64E-13 |
| ENSGALG00010021240 | SLK | SLK | 2469 | 2940 | 2420 | 2806 | 2056 | 2280 | 6047 | 3100 | 0.005496643 | 0.036351365 |
| ENSGALG00010021245 | ZFAND3 | ZFAND3 | 3951 | 4847 | 4912 | 5520 | 5986 | 3711 | 3825 | 3485 | 0.008073448 | 0.049216103 |
| ENSGALG00010021246 | PECR | PECR | 817 | 640 | 1191 | 1598 | 1164 | 844 | 1032 | 917 | 0.009064605 | 0.054006302 |
| ENSGALG00010021261 | JUP | JUP | 3976 | 5421 | 2030 | 1587 | 2928 | 2071 | 3976 | 4813 | 8.25E-10 | 4.35E-08 |
| ENSGALG00010021264 | CBFB | CBFB | 3211 | 2915 | 3305 | 4394 | 4956 | 3115 | 2104 | 2416 | 1.03E-05 | 0.000201281 |
| ENSGALG00010021267 | LRRC47 | LRRC47 | 895 | 1221 | 2110 | 2134 | 1663 | 1140 | 1330 | 1472 | 0.001065081 | 0.009828621 |
| ENSGALG00010021313 | KPNA7 | KPNA7 | 110 | 47 | 1 | 9 | 5 | 18 | 85 | 78 | 2.21E-07 | 6.85E-06 |
| ENSGALG00010021316 | PLA2G12B | PLA2G12B | 23 | 37 | 9 | 9 | 7 | 4 | 36 | 22 | 0.000794119 | 0.007756465 |
| ENSGALG00010021354 | ARHGAP39L | ARHGAP39L | 1259 | 1295 | 705 | 726 | 543 | 509 | 1027 | 868 | 0.000208993 | 0.002594866 |
| ENSGALG00010021358 | MMP15 | MMP15 | 10860 | 13366 | 5256 | 5744 | 6409 | 4453 | 7322 | 10322 | 1.11E-08 | 4.61E-07 |
| ENSGALG00010021373 | CEP250 | CEP250 | 748 | 869 | 1877 | 1605 | 1281 | 805 | 1144 | 1111 | 0.000666644 | 0.006743923 |
| ENSGALG00010021381 | DNAJB2 | DNAJB2 | 904 | 895 | 2374 | 2316 | 1368 | 1162 | 1175 | 1471 | 1.77E-06 | 4.37E-05 |
| ENSGALG00010021385 | GDF5 | GDF-5 | 20 | 20 | 177 | 85 | 55 | 24 | 18 | 32 | 6.36E-06 | 0.000132524 |
| ENSGALG00010021387 | SLC2A14 | CEF-GT3 | 23210 | 24560 | 7253 | 7352 | 14271 | 10435 | 20687 | 14561 | 1.12E-14 | 1.24E-12 |
| ENSGALG00010021390 | GSTO2 | GSTO1 | 890 | 1043 | 382 | 541 | 494 | 726 | 760 | 1022 | 0.005608781 | 0.036917666 |
| ENSGALG00010021408 | TTC9 | TTC9 | 0 | 6 | 27 | 19 | 15 | 2 | 24 | 35 | 0.002232537 | 0.017732961 |
| ENSGALG00010021411 | MCU | CCDC109A | 1740 | 1676 | 546 | 739 | 610 | 871 | 1632 | 1822 | 2.74E-07 | 8.28E-06 |
| ENSGALG00010021416 | PLEKHH3 | PLEKHH3 | 237 | 615 | 1109 | 1197 | 582 | 505 | 494 | 688 | 0.001919368 | 0.015631811 |
| ENSGALG00010021419 | MANSC1 | MANSC1 | 281 | 541 | 181 | 195 | 575 | 532 | 929 | 556 | 6.00E-07 | 1.66E-05 |
| ENSGALG00010021425 | PTPN9 | PTPN9 | 1475 | 2233 | 875 | 946 | 757 | 525 | 975 | 1547 | 1.13E-05 | 0.000218209 |
| ENSGALG00010021447 | OCSTAMP | C20H20ORF123 | 7 | 12 | 1 | 5 | 6 | 2 | 22 | 15 | 0.00807299 | 0.049216103 |
| ENSGALG00010021456 | NA | NA | 4 | 14 | 0 | 2 | 1 | 2 | 31 | 6 | 0.001167541 | 0.0105527 |
| ENSGALG00010021471 | NANOG | NANOG | 32139 | 23305 | 13728 | 19001 | 15561 | 14728 | 23592 | 16872 | 0.005921108 | 0.038391696 |
| ENSGALG00010021479 | SKAP1 | SKAP1 | 35 | 80 | 14 | 14 | 59 | 55 | 58 | 52 | 0.000575139 | 0.005958908 |
| ENSGALG00010021486 | LOC107052926 | LOC107052926 | 2 | 0 | 37 | 57 | 18 | 22 | 5 | 22 | 1.22E-06 | 3.12E-05 |
| ENSGALG00010021526 | KCNJ12 | Kir2.2 | 5 | 7 | 0 | 7 | 4 | 7 | 30 | 38 | 9.74E-05 | 0.001351389 |
| ENSGALG00010021531 | RAB15 | RAB15 | 837 | 936 | 342 | 336 | 387 | 305 | 717 | 764 | 3.15E-07 | 9.39E-06 |
| ENSGALG00010021532 | DPP10 | DPP10 | 1029 | 1911 | 2590 | 3537 | 1217 | 1225 | 2296 | 1625 | 0.001426154 | 0.012361976 |
| ENSGALG00010021536 | LOC112532074 | LOC112532074 | 9 | 12 | 48 | 43 | 44 | 10 | 9 | 6 | 0.000465471 | 0.004989709 |
| ENSGALG00010021540 | STK4 | STK4 | 3532 | 3731 | 4980 | 5963 | 5482 | 3967 | 4907 | 5362 | 0.007429373 | 0.046002103 |
| ENSGALG00010021551 | NA | NA | 1 | 3 | 1 | 4 | 18 | 15 | 19 | 9 | 0.000130631 | 0.001729378 |
| ENSGALG00010021555 | NA | NA | 6 | 10 | 3 | 6 | 23 | 19 | 26 | 19 | 0.000840676 | 0.008130427 |
| ENSGALG00010021573 | NATD1 | C14H17ORF103 | 1016 | 1243 | 1027 | 1017 | 1531 | 1329 | 1124 | 1116 | 0.003649832 | 0.026268728 |
| ENSGALG00010021583 | NFE2L1 | NRF1 | 3033 | 3095 | 2694 | 2529 | 2527 | 2320 | 4153 | 4839 | 0.000528118 | 0.005554376 |
| ENSGALG00010021616 | NA | NA | 26 | 22 | 2 | 3 | 6 | 19 | 29 | 19 | 0.001125878 | 0.010264956 |
| ENSGALG00010021622 | PTHLH | PTHrP | 205 | 56 | 21 | 67 | 24 | 24 | 17 | 59 | 0.006189109 | 0.039723422 |
| ENSGALG00010021628 | REC114 | C10H15ORF60 | 40 | 29 | 129 | 115 | 82 | 60 | 67 | 71 | 0.0047247 | 0.032433674 |
| ENSGALG00010021630 | VTG1 | LVII | 45 | 76 | 48 | 195 | 35 | 23 | 22 | 31 | 0.003118931 | 0.0231827 |
| ENSGALG00010021638 | SRC | PP60C-SCR | 1011 | 1219 | 564 | 587 | 462 | 327 | 840 | 515 | 9.32E-05 | 0.001305753 |
| ENSGALG00010021642 | NPTN | SDFR1 | 4383 | 4074 | 2711 | 2912 | 2714 | 2015 | 3313 | 3195 | 0.001999847 | 0.01619256 |
| ENSGALG00010021647 | ARHGAP11B | ARHGAP11A | 4502 | 4578 | 2635 | 3517 | 3247 | 2561 | 4012 | 4232 | 0.005204139 | 0.034774716 |
| ENSGALG00010021669 | SAR1A | SAR1A | 7148 | 7749 | 4757 | 5654 | 4623 | 2997 | 3853 | 4146 | 2.82E-06 | 6.52E-05 |
| ENSGALG00010021679 | MASP2 | MAp19 | 110 | 51 | 78 | 95 | 334 | 377 | 425 | 410 | 4.61E-11 | 3.07E-09 |
| ENSGALG00010021682 | MYL4 | L23 | 65 | 147 | 42 | 70 | 109 | 153 | 301 | 430 | 6.37E-07 | 1.75E-05 |
| ENSGALG00010021710 | KEL | KEL | 162 | 184 | 1 | 0 | 6 | 1 | 0 | 0 | 6.57E-39 | 3.79E-36 |
| ENSGALG00010021721 | TP53I11 | TP53I11 | 7435 | 8644 | 3308 | 5727 | 4447 | 4772 | 5643 | 5570 | 0.001535146 | 0.013109533 |
| ENSGALG00010021729 | RBPJL | RBPJL | 17 | 36 | 0 | 4 | 2 | 1 | 7 | 15 | 1.91E-05 | 0.000340075 |
| ENSGALG00010021742 | MATN4 | MATN4 | 82 | 108 | 10 | 9 | 16 | 5 | 36 | 72 | 6.20E-09 | 2.79E-07 |
| ENSGALG00010021744 | LITAF | TNF-alpha | 2239 | 2267 | 388 | 523 | 889 | 773 | 2445 | 809 | 1.68E-08 | 6.74E-07 |
| ENSGALG00010021748 | CDCA7 | CDCA7 | 1135 | 1040 | 2179 | 2160 | 1610 | 1309 | 1497 | 1521 | 0.00102743 | 0.009549731 |
| ENSGALG00010021775 | CCDC36 | CCDC36 | 75 | 95 | 29 | 25 | 22 | 19 | 69 | 59 | 0.000283418 | 0.003335547 |
| ENSGALG00010021792 | KLHDC8B | KLHDC8B | 504 | 954 | 256 | 124 | 225 | 131 | 505 | 868 | 3.53E-07 | 1.03E-05 |

| | | | | | | | | | | | | |
|---|---|---|---|---|---|---|---|---|---|---|---|---|
| ENSGALG00010021796 | NA | NA | 87 | 38 | 7 | 1 | 49 | 13 | 32 | 13 | 7.60E-05 | 0.001095904 |
| ENSGALG00010021799 | LOC416055 | LOC416055 | 818 | 1262 | 468 | 313 | 427 | 337 | 772 | 1119 | 2.94E-06 | 6.75E-05 |
| ENSGALG00010021803 | TLL2 | TLL2 | 155 | 189 | 51 | 47 | 62 | 65 | 144 | 106 | 0.000320056 | 0.003688807 |
| ENSGALG00010021837 | NA | NA | 19 | 16 | 0 | 1 | 0 | 1 | 0 | 0 | 8.92E-09 | 3.82E-07 |
| ENSGALG00010021846 | NA | NA | 31 | 49 | 10 | 5 | 18 | 9 | 20 | 11 | 0.002361024 | 0.018489895 |
| ENSGALG00010021848 | LARGE2 | GYLTL1B | 2046 | 4514 | 1576 | 1542 | 1404 | 1083 | 2452 | 3930 | 1.35E-05 | 0.000253467 |
| ENSGALG00010021854 | TAPBPL | TAPBPL | 2 | 1 | 27 | 16 | 12 | 6 | 15 | 14 | 0.000677412 | 0.006836296 |
| ENSGALG00010021864 | GFI1 | GFI1 | 33 | 32 | 203 | 181 | 88 | 58 | 85 | 53 | 3.96E-06 | 8.79E-05 |
| ENSGALG00010021866 | SCNN1A | SCNN1A | 294 | 431 | 693 | 743 | 1075 | 642 | 554 | 434 | 9.11E-06 | 0.000180953 |
| ENSGALG00010021867 | CXCL12 | SDF1 | 1044 | 2695 | 831 | 408 | 682 | 715 | 2154 | 2408 | 2.25E-06 | 5.35E-05 |
| ENSGALG00010021870 | LOC101747373 | LOC101747373 | 1266 | 1378 | 318 | 400 | 308 | 235 | 370 | 279 | 1.92E-17 | 2.89E-15 |
| ENSGALG00010021886 | LIPML5 | LIPML5 | 2 | 7 | 0 | 2 | 6 | 9 | 17 | 10 | 0.005200991 | 0.034767584 |
| ENSGALG00010021902 | LMTK2 | LMTK2 | 1180 | 1060 | 548 | 452 | 489 | 537 | 1237 | 758 | 0.000280821 | 0.003316725 |
| ENSGALG00010021919 | PARVA | PARVA | 73 | 75 | 250 | 175 | 187 | 191 | 321 | 187 | 0.000970814 | 0.009114747 |
| ENSGALG00010021945 | C1QTNF4 | C1QTNF4 | 79 | 288 | 432 | 828 | 411 | 224 | 87 | 145 | 2.55E-05 | 0.000435252 |
| ENSGALG00010021956 | NA | NA | 5 | 0 | 2 | 2 | 19 | 14 | 6 | 4 | 0.000658891 | 0.006685723 |
| ENSGALG00010021985 | NA | NA | 3 | 4 | 9 | 11 | 10 | 12 | 30 | 23 | 0.001419594 | 0.012330698 |
| ENSGALG00010022001 | NA | NA | 10 | 10 | 60 | 25 | 19 | 10 | 16 | 7 | 0.008615883 | 0.05177804 |
| ENSGALG00010022010 | NA | NA | 19 | 6 | 8 | 8 | 58 | 73 | 86 | 68 | 6.19E-09 | 2.79E-07 |
| ENSGALG00010022029 | LOC107049117 | LOC107049117 | 2022 | 1360 | 1103 | 1219 | 698 | 392 | 502 | 924 | 1.62E-05 | 0.000295579 |
| ENSGALG00010022071 | ACTA2 | asma | 1 | 2 | 29 | 12 | 3 | 13 | 20 | 27 | 0.001238315 | 0.011096366 |
| ENSGALG00010022085 | MLLT6 | MLLT6 | 254 | 539 | 789 | 1072 | 540 | 446 | 376 | 517 | 0.004329934 | 0.030150774 |
| ENSGALG00010022100 | ENTPD1 | ENTPD1 | 378 | 1052 | 197 | 339 | 177 | 230 | 470 | 506 | 0.001054704 | 0.009749015 |
| ENSGALG00010022121 | NA | NA | 35 | 36 | 62 | 66 | 73 | 56 | 235 | 239 | 1.60E-09 | 8.08E-08 |
| ENSGALG00010022129 | ELMO3 | ELMO3 | 511 | 436 | 235 | 218 | 317 | 231 | 610 | 609 | 8.11E-05 | 0.001162663 |
| ENSGALG00010022144 | SNTA1 | SNTA1 | 564 | 628 | 308 | 417 | 274 | 210 | 383 | 510 | 0.009480913 | 0.055810153 |
| ENSGALG00010022164 | PDGFA | PDGF-A | 36 | 26 | 141 | 67 | 86 | 21 | 186 | 88 | 0.00856125 | 0.051540203 |
| ENSGALG00010022184 | PRKAR1B | PRKAR1B | 132 | 299 | 1426 | 1292 | 1197 | 737 | 428 | 308 | 3.08E-15 | 3.62E-13 |
| ENSGALG00010022214 | NA | NA | 75 | 82 | 172 | 265 | 106 | 85 | 87 | 93 | 0.004471651 | 0.030944193 |
| ENSGALG00010022223 | DOK4 | DOK4 | 521 | 681 | 131 | 245 | 192 | 168 | 256 | 318 | 4.42E-06 | 9.71E-05 |
| ENSGALG00010022240 | SPINT1 | SPINT1 | 730 | 1426 | 249 | 260 | 317 | 300 | 1021 | 1231 | 1.18E-13 | 1.13E-11 |
| ENSGALG00010022246 | RHOV | RHOV | 631 | 646 | 97 | 119 | 461 | 418 | 916 | 575 | 2.82E-14 | 2.95E-12 |
| ENSGALG00010022248 | FIGNL2 | FIGNL2 | 1062 | 1510 | 362 | 417 | 535 | 453 | 992 | 1360 | 4.98E-11 | 3.28E-09 |
| ENSGALG00010022249 | DRD4 | D4 | 160 | 110 | 12 | 18 | 46 | 38 | 67 | 61 | 6.33E-08 | 2.25E-06 |
| ENSGALG00010022254 | LOC776720 | DYNC1LI2L | 377 | 271 | 915 | 818 | 691 | 366 | 437 | 502 | 0.000795675 | 0.007762536 |
| ENSGALG00010022264 | SCN8A | SCN8A | 51 | 115 | 56 | 32 | 27 | 11 | 108 | 91 | 0.001317911 | 0.011647238 |
| ENSGALG00010022271 | SLC18A2 | SLC18A2 | 202 | 195 | 39 | 38 | 180 | 159 | 122 | 85 | 1.94E-07 | 6.14E-06 |
| ENSGALG00010022299 | MRGBP | C20H20orf20 | 990 | 1521 | 1490 | 1688 | 1484 | 938 | 812 | 900 | 0.007846012 | 0.048083077 |
| ENSGALG00010022306 | NA | NA | 1666 | 1071 | 597 | 961 | 743 | 487 | 831 | 709 | 0.00279894 | 0.021205285 |
| ENSGALG00010022309 | WSCD2 | WSCD2 | 33 | 82 | 12 | 8 | 12 | 17 | 91 | 83 | 7.80E-07 | 2.08E-05 |
| ENSGALG00010022311 | DLL4 | DLL4 | 32 | 84 | 1396 | 307 | 885 | 254 | 272 | 115 | 2.28E-07 | 7.04E-06 |
| ENSGALG00010022340 | GFOD2 | GFOD2 | 4553 | 6678 | 2791 | 3727 | 2890 | 1963 | 2315 | 2741 | 1.07E-07 | 3.65E-06 |
| ENSGALG00010022353 | SLC39A13 | ZIP13 | 3496 | 4111 | 4223 | 5828 | 4975 | 3267 | 3328 | 2907 | 0.003797306 | 0.027154718 |
| ENSGALG00010022361 | NA | NA | 588 | 1048 | 743 | 762 | 361 | 252 | 315 | 511 | 0.000859974 | 0.008278263 |
| ENSGALG00010022373 | AK5 | AK5 | 525 | 876 | 999 | 1225 | 591 | 420 | 418 | 529 | 0.000930572 | 0.008806187 |
| ENSGALG00010022382 | CALB2 | CALB2 | 80 | 69 | 28 | 51 | 23 | 12 | 37 | 51 | 0.0087957 | 0.052717402 |
| ENSGALG00010022399 | NA | NA | 0 | 0 | 44 | 41 | 0 | 0 | 0 | 0 | 2.35E-19 | 4.52E-17 |
| ENSGALG00010022411 | ADRB1 | ADRB1 | 104 | 108 | 39 | 35 | 143 | 60 | 94 | 78 | 0.004265153 | 0.02979901 |
| ENSGALG00010022412 | CHAC1 | CHAC1 | 206 | 92 | 129 | 64 | 88 | 384 | 541 | 949 | 3.25E-05 | 0.00053499 |
| ENSGALG00010022416 | CCDC186 | C10orf118 | 2906 | 2510 | 1334 | 1724 | 1824 | 1477 | 2959 | 1872 | 0.002903969 | 0.021841195 |
| ENSGALG00010022421 | CMKLR1 | ChemR23 | 15 | 26 | 14 | 3 | 5 | 11 | 49 | 37 | 0.003883083 | 0.027626259 |
| ENSGALG00010022425 | VWA2 | VWA2 | 193 | 151 | 39 | 40 | 84 | 98 | 193 | 129 | 4.94E-05 | 0.000762739 |
| ENSGALG00010022436 | AFAP1L2 | AFAP1L2 | 1243 | 2168 | 682 | 771 | 843 | 756 | 1823 | 1853 | 1.18E-07 | 3.97E-06 |
| ENSGALG00010022453 | LOC121106446 | LOC121106446 | 785 | 780 | 302 | 401 | 539 | 624 | 913 | 761 | 0.000122468 | 0.001637345 |
| ENSGALG00010022454 | ABLIM1 | ABLIM1 | 178 | 206 | 68 | 101 | 122 | 194 | 382 | 174 | 0.00680385 | 0.042762669 |
| ENSGALG00010022476 | OIP5 | OIP5 | 1038 | 1262 | 1689 | 2225 | 1844 | 1688 | 1613 | 1539 | 0.001927993 | 0.015686769 |
| ENSGALG00010022484 | NA | NA | 36 | 24 | 35 | 28 | 251 | 126 | 25 | 30 | 5.08E-13 | 4.47E-11 |
| ENSGALG00010022491 | LOC107053365 | LOC107053365 | 339 | 342 | 1343 | 1232 | 562 | 377 | 600 | 1021 | 1.08E-07 | 3.67E-06 |
| ENSGALG00010022508 | USF3 | KIAA2018 | 204 | 302 | 159 | 196 | 184 | 350 | 1001 | 280 | 0.003529986 | 0.025593546 |
| ENSGALG00010022512 | PRAP1 | PRAP1 | 13 | 4 | 6 | 6 | 4 | 4 | 29 | 32 | 0.000613387 | 0.006273518 |
| ENSGALG00010022515 | CHRNA1 | NACHRA1 | 20 | 27 | 34 | 47 | 28 | 20 | 9 | 6 | 0.002006146 | 0.016235697 |
| ENSGALG00010022516 | NA | NA | 1236 | 1271 | 669 | 955 | 1094 | 882 | 843 | 845 | 0.007914275 | 0.048412649 |
| ENSGALG00010022520 | TMOD2 | TMOD2 | 716 | 764 | 1076 | 1457 | 1646 | 885 | 1018 | 616 | 0.001649788 | 0.013903809 |
| ENSGALG00010022522 | TNS4 | TNS4 | 79 | 232 | 100 | 155 | 56 | 44 | 207 | 267 | 0.002577301 | 0.019794047 |
| ENSGALG00010022525 | ISCU | ISCU | 3865 | 4581 | 5775 | 7296 | 5248 | 3961 | 3918 | 3956 | 0.0008115 | 0.007893942 |
| ENSGALG00010022548 | NA | NA | 136 | 440 | 625 | 1078 | 228 | 252 | 236 | 294 | 0.000330879 | 0.003788754 |
| ENSGALG00010022562 | GHRH | GHRH | 76 | 69 | 31 | 54 | 7 | 10 | 16 | 16 | 3.30E-06 | 7.50E-05 |
| ENSGALG00010022568 | SCRN3 | SCRN3 | 433 | 438 | 893 | 987 | 718 | 475 | 747 | 679 | 0.003622737 | 0.026152554 |
| ENSGALG00010022569 | KIAA1549L | C11ORF41 | 667 | 810 | 454 | 402 | 806 | 793 | 523 | 304 | 6.23E-05 | 0.000922464 |
| ENSGALG00010022572 | LTK | LTK | 162 | 204 | 127 | 126 | 241 | 468 | 488 | 462 | 4.69E-06 | 0.000102327 |
| ENSGALG00010022577 | LOC770870 | LOC770870 | 0 | 3 | 35 | 20 | 5 | 14 | 12 | 11 | 0.000680249 | 0.006856797 |
| ENSGALG00010022583 | FKBP11 | FKBP11 | 1012 | 359 | 167 | 336 | 250 | 198 | 340 | 323 | 0.001935557 | 0.015740645 |
| ENSGALG00010022594 | NA | NA | 3 | 4 | 21 | 24 | 15 | 15 | 5 | 9 | 0.001827605 | 0.015075484 |
| ENSGALG00010022597 | SLCO4A1 | SLCO4A1 | 227 | 848 | 268 | 191 | 151 | 167 | 309 | 833 | 0.006189552 | 0.039723422 |
| ENSGALG00010022598 | NA | NA | 1 | 3 | 0 | 0 | 4 | 9 | 11 | 10 | 0.000155023 | 0.002008327 |
| ENSGALG00010022622 | MPPED2 | MPPED2 | 2805 | 2992 | 1579 | 1589 | 2063 | 1648 | 1906 | 1812 | 5.18E-05 | 0.000792893 |
| ENSGALG00010022650 | NR1H3 | LXR | 1018 | 1333 | 2335 | 2058 | 1613 | 1223 | 1588 | 1964 | 0.008501617 | 0.051236576 |
| ENSGALG00010022655 | NRIP2 | NRIP3 | 20 | 127 | 55 | 107 | 249 | 133 | 43 | 83 | 0.009977663 | 0.05816069 |
| ENSGALG00010022660 | METAP1D | METAP1D | 189 | 245 | 462 | 493 | 432 | 281 | 245 | 304 | 0.005635011 | 0.037046543 |
| ENSGALG00010022671 | DBR1 | DBR1 | 1825 | 1862 | 1242 | 1346 | 1062 | 761 | 1468 | 1488 | 0.004461089 | 0.030883894 |
| ENSGALG00010022675 | DMBX1 | ATX | 353 | 246 | 5 | 55 | 86 | 61 | 144 | 72 | 1.68E-05 | 0.000303777 |
| ENSGALG00010022691 | ASIC2 | ACCN1 | 27 | 36 | 100 | 60 | 105 | 105 | 96 | 67 | 0.002347622 | 0.018419462 |
| ENSGALG00010022695 | PIK3IP1 | PIK3IP1 | 906 | 1025 | 879 | 1211 | 2130 | 1532 | 1265 | 1300 | 5.16E-07 | 1.45E-05 |
| ENSGALG00010022697 | NA | NA | 16 | 41 | 49 | 83 | 71 | 31 | 16 | 18 | 0.00542757 | 0.035956124 |
| ENSGALG00010022700 | NA | NA | 36 | 68 | 8 | 22 | 7 | 10 | 39 | 52 | 0.000344194 | 0.003910095 |
| ENSGALG00010022705 | CRHR1 | CRHR1 | 232 | 533 | 163 | 132 | 103 | 103 | 466 | 580 | 3.29E-07 | 9.75E-06 |
| ENSGALG00010022707 | SALL1 | SAL1 | 2005 | 4258 | 1048 | 1372 | 1889 | 1590 | 3021 | 2324 | 1.06E-05 | 0.000205231 |
| ENSGALG00010022742 | IFT46 | C11orf60 | 274 | 268 | 117 | 112 | 141 | 143 | 277 | 238 | 0.003737854 | 0.026791092 |
| ENSGALG00010022760 | ELAVL4 | HUD | 4285 | 7155 | 13361 | 15045 | 10946 | 7706 | 6031 | 5938 | 7.39E-11 | 4.73E-09 |
| ENSGALG00010022764 | MFAP5 | MFAP5 | 73 | 81 | 129 | 134 | 190 | 136 | 99 | 46 | 0.006637109 | 0.041919642 |
| ENSGALG00010022769 | FXYD6 | FXYD6 | 295 | 342 | 145 | 171 | 107 | 77 | 237 | 200 | 0.000370376 | 0.004170978 |
| ENSGALG00010022770 | HAT1 | HAT1 | 1877 | 2091 | 2661 | 3262 | 2282 | 1669 | 1565 | 1790 | 0.003164386 | 0.023451535 |
| ENSGALG00010022772 | NA | NA | 92 | 209 | 39 | 41 | 249 | 173 | 99 | 100 | 7.08E-07 | 1.91E-05 |
| ENSGALG00010022778 | GDF11 | GDF11 | 46 | 141 | 34 | 25 | 22 | 8 | 43 | 36 | 0.00075624 | 0.007465021 |
| ENSGALG00010022789 | LOC428824 | LOC428824 | 20 | 75 | 12 | 6 | 15 | 14 | 176 | 169 | 1.56E-11 | 1.12E-09 |
| ENSGALG00010022792 | DDB2 | DDB2 | 1187 | 1490 | 3537 | 3848 | 3078 | 2224 | 1960 | 1707 | 3.74E-12 | 3.01E-10 |
| ENSGALG00010022801 | LSM14B | LSM14B | 5562 | 6509 | 3414 | 4268 | 3310 | 3498 | 5436 | 5473 | 0.001085955 | 0.009993656 |
| ENSGALG00010022803 | LOC770492 | LOC770492 | 269 | 266 | 69 | 110 | 80 | 82 | 206 | 157 | 0.000231036 | 0.002822421 |
| ENSGALG00010022819 | PACSIN3 | PACSIN3 | 539 | 665 | 1302 | 1095 | 947 | 636 | 742 | 935 | 0.009976974 | 0.05816069 |
| ENSGALG00010022821 | A2ML2 | A2ML2 | 9 | 17 | 81 | 17 | 47 | 9 | 6 | 5 | 0.002390964 | 0.018678841 |
| ENSGALG00010022824 | PUS1 | PUS1 | 2929 | 3876 | 3265 | 3933 | 2371 | 1700 | 2245 | 2124 | 0.000878997 | 0.008403778 |
| ENSGALG00010022839 | SYT12 | synaptotagmin-12 | 262 | 384 | 427 | 191 | 92 | 72 | 234 | 501 | 0.001582898 | 0.013448601 |
| ENSGALG00010022841 | DKK4 | DKK1 | 225 | 184 | 34 | 118 | 156 | 191 | 230 | 309 | 0.003641412 | 0.026228113 |
| ENSGALG00010022843 | NA | NA | 33 | 109 | 11 | 5 | 7 | 9 | 5 | 11 | 3.66E-07 | 1.07E-05 |
| ENSGALG00010022848 | SGIP1 | SGIP1 | 22 | 69 | 17 | 8 | 13 | 12 | 44 | 54 | 0.002730324 | 0.02079334 |
| ENSGALG00010022849 | NA | NA | 375 | 392 | 0 | 2 | 2 | 3 | 3 | 6 | 6.52E-56 | 7.78E-53 |
| ENSGALG00010022879 | LOC415616 | LOC415616 | 23 | 53 | 147 | 122 | 114 | 59 | 58 | 64 | 0.003789687 | 0.027111834 |
| ENSGALG00010022891 | NTM | CEPU | 606 | 921 | 1573 | 1719 | 1272 | 743 | 945 | 1051 | 0.001223369 | 0.010974205 |
| ENSGALG00010022892 | NA | NA | 44 | 55 | 9 | 14 | 15 | 19 | 49 | 40 | 0.000661152 | 0.0067046 |
| ENSGALG00010022921 | INPP1 | INPP1 | 70 | 113 | 25 | 24 | 39 | 39 | 53 | 57 | 0.002526615 | 0.019513899 |
| ENSGALG00010022932 | OPCML | OBCAM | 1000 | 593 | 142 | 281 | 1408 | 1391 | 1459 | 1146 | 2.93E-15 | 3.50E-13 |
| ENSGALG00010022933 | UNC119B | UNC119B | 3666 | 2961 | 2094 | 2731 | 2815 | 2753 | 4440 | 3908 | 0.003345599 | 0.024479707 |
| ENSGALG00010022948 | PDZK1IP1 | PDZK1IP1 | 74 | 10 | 2 | 0 | 9 | 14 | 40 | 20 | 3.93E-05 | 0.000628189 |
| ENSGALG00010022949 | INSC | INSC | 14 | 26 | 46 | 94 | 52 | 25 | 15 | 12 | 0.00047151 | 0.005041504 |
| ENSGALG00010022977 | NA | NA | 216 | 363 | 89 | 98 | 77 | 140 | 295 | 195 | 0.002104148 | 0.01692717 |
| ENSGALG00010022979 | LAMA5 | LAMA5 | 3333 | 5217 | 2135 | 1980 | 1272 | 1548 | 3659 | 3544 | 7.52E-08 | 2.65E-06 |
| ENSGALG00010022980 | NA | NA | 118 | 140 | 40 | 51 | 24 | 53 | 100 | 88 | 0.006629473 | 0.041899282 |
| ENSGALG00010022984 | NA | NA | 201 | 227 | 19 | 24 | 13 | 49 | 72 | 55 | 7.10E-08 | 2.51E-06 |
| ENSGALG00010022997 | ARHGAP1 | ARHGAP1 | 5764 | 7034 | 912 | 1436 | 3539 | 5283 | 9086 | 7643 | 1.13E-20 | 2.42E-18 |
| ENSGALG00010023013 | VWA5A2 | VWA5A | 28 | 44 | 74 | 215 | 98 | 77 | 81 | 83 | 0.006157371 | 0.039579655 |
| ENSGALG00010023028 | TMEM61 | TMEM61 | 61 | 125 | 28 | 31 | 66 | 47 | 45 | 41 | 0.006470116 | 0.041129167 |
| ENSGALG00010023078 | TAGLN | SM22 | 5 | 4 | 0 | 6 | 4 | 1 | 12 | 28 | 0.004749007 | 0.032520827 |
| ENSGALG00010023099 | DNAJC22 | DNAJC22 | 29 | 25 | 91 | 190 | 4 | 19 | 10 | 28 | 2.19E-06 | 5.24E-05 |
| ENSGALG00010023101 | CLCA1 | CLCA1 | 3 | 2 | 0 | 2 | 12 | 2 | 24 | 9 | 0.001864584 | 0.015267483 |
| ENSGALG00010023104 | EFCAB4B | CRACR2B | 1503 | 1709 | 818 | 821 | 827 | 633 | 978 | 945 | 1.37E-05 | 0.000255702 |
| ENSGALG00010023119 | NDRG4 | NDRG4 | 454 | 383 | 116 | 172 | 221 | 324 | 325 | 247 | 0.001403758 | 0.012231287 |
| ENSGALG00010023127 | CERS5 | LASS5 | 1851 | 2124 | 2361 | 3190 | 2088 | 1509 | 1414 | 1674 | 0.00693906 | 0.043420804 |
| ENSGALG00010023136 | MDK | RI-HB | 15378 | 14312 | 6829 | 9949 | 7294 | 6502 | 10250 | 12772 | 2.59E-05 | 0.000439666 |
| ENSGALG00010023141 | LOC121108102 | LOC121108102 | 2 | 28 | 32 | 16 | 1 | 2 | 52 | 17 | 0.002938005 | 0.022067388 |
| ENSGALG00010023158 | NA | NA | 17 | 20 | 6 | 10 | 17 | 9 | 0 | 1 | 2.55E-05 | 0.000435252 |
| ENSGALG00010023176 | NA | NA | 435 | 305 | 327 | 314 | 65 | 81 | 457 | 223 | 8.04E-05 | 0.001154494 |

| | | | | | | | | | | | | | |
|---|---|---|---|---|---|---|---|---|---|---|---|---|---|
| ENSGALG00010023178 | CFAP20 | C11H16ORF80 | | 4175 | 5934 | 3004 | 3471 | 3800 | 2928 | 3462 | 3576 | 0.001770541 | 0.014706406 |
| ENSGALG00010023180 | EEF1A2 | EEF1A2 | | 208 | 370 | 109 | 125 | 97 | 55 | 140 | 192 | 0.000556473 | 0.005801486 |
| ENSGALG00010023181 | PLEKHA7 | PLEKHA7 | | 382 | 424 | 406 | 456 | 242 | 254 | 761 | 562 | 0.008875925 | 0.053090359 |
| ENSGALG00010023182 | EPHB6 | CEK9 | | 739 | 576 | 511 | 491 | 680 | 498 | 62 | 128 | 1.42E-12 | 1.21E-10 |
| ENSGALG00010023194 | MRC2 | MRC2 | | 614 | 1199 | 1472 | 1272 | 677 | 456 | 1825 | 1750 | 1.27E-05 | 0.000241073 |
| ENSGALG00010023195 | COL24A1 | COL24A1 | | 97 | 53 | 141 | 112 | 70 | 158 | 391 | 276 | 0.000866307 | 0.008320529 |
| ENSGALG00010023197 | LOC107049478 | LOC107049478 | | 532 | 430 | 234 | 195 | 405 | 381 | 307 | 207 | 0.000582894 | 0.006020594 |
| ENSGALG00010023212 | RIMBP2 | RBP2 | | 283 | 348 | 633 | 662 | 575 | 506 | 671 | 490 | 0.005444532 | 0.036049532 |
| ENSGALG00010023230 | PXDNL | PXDNL | | 311 | 566 | 891 | 1000 | 841 | 734 | 626 | 654 | 0.001239898 | 0.011098647 |
| ENSGALG00010023231 | ASIC1 | ACCN2 | | 310 | 743 | 10 | 15 | 11 | 10 | 574 | 496 | 8.61E-40 | 5.76E-37 |
| ENSGALG00010023238 | NUCB2 | nucleobindin-2 | | 3814 | 3002 | 2011 | 2475 | 1798 | 1459 | 3150 | 2498 | 0.000889295 | 0.008482817 |
| ENSGALG00010023242 | MYT1 | MYT1 | | 2 | 10 | 66 | 34 | 77 | 135 | 31 | 64 | 2.02E-07 | 6.32E-06 |
| ENSGALG00010023243 | HK1 | hexokinase-1 | | 2966 | 3437 | 2022 | 2358 | 2122 | 1701 | 3162 | 3215 | 0.001511358 | 0.012946082 |
| ENSGALG00010023244 | PPIB | PPIB | | 15685 | 12111 | 8598 | 10229 | 8360 | 5531 | 9491 | 11727 | 0.001425246 | 0.012360513 |
| ENSGALG00010023247 | LOC121106503 | LOC121106503 | | 515 | 419 | 806 | 879 | 1064 | 791 | 1221 | 850 | 0.00013244 | 0.001749673 |
| ENSGALG00010023254 | MYO3B | MYO3B | | 11 | 7 | 57 | 27 | 51 | 19 | 28 | 25 | 0.009472691 | 0.055781397 |
| ENSGALG00010023259 | TSPAN15 | tetraspanin-15 | | 339 | 288 | 135 | 124 | 150 | 180 | 497 | 437 | 1.31E-05 | 0.000247349 |
| ENSGALG00010023262 | POLD4 | POLD4 | | 41 | 73 | 24 | 21 | 59 | 37 | 79 | 93 | 0.003311003 | 0.024343812 |
| ENSGALG00010023269 | OPRL1 | OPRL1 | | 34 | 259 | 35 | 14 | 38 | 33 | 96 | 134 | 0.001701505 | 0.014229603 |
| ENSGALG00010023274 | ZBTB38 | ZBTB38 | | 203 | 144 | 638 | 410 | 491 | 301 | 383 | 349 | 0.000874185 | 0.008367345 |
| ENSGALG00010023281 | B2M | b2m-W01 | | 2578 | 2797 | 840 | 961 | 961 | 1256 | 2474 | 3183 | 9.02E-11 | 5.58E-09 |
| ENSGALG00010023305 | PLCB2 | PLCB2 | | 50 | 12 | 180 | 160 | 201 | 156 | 232 | 163 | 2.78E-05 | 0.000469087 |
| ENSGALG00010023306 | NA | NA | | 6 | 7 | 0 | 0 | 0 | 0 | 31 | 26 | 1.96E-12 | 1.64E-10 |
| ENSGALG00010023311 | ETS1 | ETS-1 | | 589 | 910 | 112 | 202 | 499 | 439 | 529 | 611 | 3.02E-09 | 1.44E-07 |
| ENSGALG00010023314 | CYB561 | CYB561 | | 20 | 3 | 2 | 5 | 1 | 1 | 17 | 11 | 0.009413586 | 0.055550794 |
| ENSGALG00010023317 | RAPGEF3 | RAPGEF3 | | 462 | 1215 | 311 | 243 | 358 | 298 | 693 | 907 | 4.06E-05 | 0.000643053 |
| ENSGALG00010023343 | TMEM233 | TMEM233 | | 265 | 1381 | 1186 | 1832 | 510 | 304 | 327 | 434 | 0.00428118 | 0.004655675 |
| ENSGALG00010023346 | NA | NA | | 128 | 154 | 13 | 29 | 55 | 119 | 152 | 207 | 2.86E-06 | 6.59E-05 |
| ENSGALG00010023369 | ACE | DCP1 | | 335 | 424 | 147 | 132 | 194 | 182 | 287 | 314 | 0.000806257 | 0.007847509 |
| ENSGALG00010023375 | ST14 | ST14 | | 1045 | 615 | 788 | 611 | 703 | 469 | 1395 | 1466 | 0.001343424 | 0.011822698 |
| ENSGALG00010023384 | AP3B2 | AP3B2 | | 1665 | 1728 | 803 | 1144 | 741 | 822 | 1574 | 1656 | 0.000269012 | 0.003202088 |
| ENSGALG00010023422 | WLS | GPR177 | | 3402 | 6060 | 2779 | 3625 | 3294 | 3023 | 2667 | 3031 | 0.007733542 | 0.04751579 |
| ENSGALG00010023471 | NA | NA | | 688 | 606 | 1289 | 1219 | 1185 | 956 | 700 | 591 | 1.78E-05 | 0.000319464 |
| ENSGALG00010023489 | ETV5 | ERM | | 5129 | 3332 | 2278 | 2560 | 2455 | 1740 | 2727 | 3510 | 0.000969765 | 0.009110013 |
| ENSGALG00010023496 | PLEKHA3 | PLEKHA3 | | 1217 | 1092 | 2095 | 1909 | 1933 | 1163 | 1426 | 1238 | 0.009157692 | 0.054436829 |
| ENSGALG00010023498 | DISP2 | DISP2 | | 112 | 83 | 282 | 251 | 235 | 248 | 346 | 203 | 0.002007696 | 0.016240378 |
| ENSGALG00010023499 | PEBP1 | PBP | | 6958 | 6223 | 10003 | 11647 | 6834 | 5720 | 5245 | 7824 | 0.007074017 | 0.044112301 |
| ENSGALG00010023500 | NA | NA | | 2 | 21 | 11 | 4 | 2 | 9 | 51 | 49 | 0.001176336 | 0.01061497 |
| ENSGALG00010023508 | NA | NA | | 24 | 80 | 1 | 11 | 7 | 15 | 31 | 61 | 0.000446681 | 0.004816495 |
| ENSGALG00010023516 | NA | NA | | 0 | 1 | 4 | 5 | 11 | 13 | 2 | 4 | 0.000348508 | 0.003948656 |
| ENSGALG00010023519 | DGKG | DGKG | | 50 | 173 | 43 | 51 | 44 | 42 | 149 | 180 | 0.001454207 | 0.01255954 |
| ENSGALG00010023527 | TMEM98 | TMEM98 | | 183 | 14 | 21 | 10 | 30 | 13 | 271 | 658 | 4.16E-09 | 1.95E-07 |
| ENSGALG00010023546 | SLC11A2 | NRAMP2 | | 5271 | 3892 | 2661 | 2213 | 1799 | 1365 | 2851 | 3996 | 6.15E-07 | 1.69E-05 |
| ENSGALG00010023549 | NA | NA | | 965 | 706 | 230 | 308 | 241 | 524 | 2119 | 936 | 3.54E-07 | 1.03E-05 |
| ENSGALG00010023553 | NA | NA | | 126 | 63 | 15 | 31 | 11 | 12 | 99 | 49 | 1.05E-05 | 0.000204891 |
| ENSGALG00010023557 | NA | NA | | 102 | 49 | 0 | 1 | 3 | 2 | 166 | 215 | 3.28E-27 | 1.14E-24 |
| ENSGALG00010023562 | NA | NA | | 14 | 6 | 2 | 0 | 4 | 15 | 18 | 16 | 0.004835031 | 0.032953928 |
| ENSGALG00010023580 | RDM1 | RAD52B | | 487 | 0 | 2 | 0 | 3 | 26 | 13 | 14 | 1.77E-05 | 0.000318129 |
| ENSGALG00010023591 | NA | NA | | 55 | 680 | 45 | 91 | 254 | 227 | 28 | 78 | 0.00028257 | 0.003330738 |
| ENSGALG00010023607 | NA | NA | | 38 | 9 | 0 | 0 | 0 | 0 | 44 | 22 | 3.09E-11 | 2.11E-09 |
| ENSGALG00010023620 | LOC107051846 | LOC107051846 | | 1003 | 1178 | 568 | 634 | 753 | 551 | 861 | 746 | 0.004138925 | 0.029148638 |
| ENSGALG00010023625 | NA | NA | | 180 | 82 | 32 | 32 | 49 | 130 | 309 | 176 | 0.00011676 | 0.001572917 |
| ENSGALG00010023635 | SCN4A | SCN4A | | 5 | 0 | 0 | 0 | 4 | 1 | 18 | 16 | 2.41E-05 | 0.000414728 |
| ENSGALG00010023636 | PDE3B | PDE3B | | 1173 | 1193 | 2085 | 2699 | 1381 | 1215 | 1292 | 1081 | 7.48E-05 | 0.00108085 |
| ENSGALG00010023641 | LRRC40 | LRRC40 | | 1097 | 1619 | 1477 | 1942 | 550 | 383 | 1251 | 1383 | 5.77E-08 | 2.06E-06 |
| ENSGALG00010023644 | NA | NA | | 3 | 5 | 5 | 19 | 17 | 12 | 1 | 1 | 0.00072376 | 0.007212573 |
| ENSGALG00010023655 | C1R | C1R | | 663 | 675 | 272 | 205 | 394 | 379 | 777 | 865 | 1.96E-07 | 6.18E-06 |
| ENSGALG00010023657 | NA | NA | | 13 | 13 | 73 | 41 | 89 | 38 | 89 | 38 | 0.001452904 | 0.01255954 |
| ENSGALG00010023659 | NA | NA | | 2 | 1 | 16 | 4 | 13 | 16 | 8 | 3 | 0.008706409 | 0.052263471 |
| ENSGALG00010023664 | SLC6A9 | SLC6A9 | | 470 | 636 | 817 | 647 | 544 | 783 | 1103 | 1354 | 0.005691421 | 0.037300012 |
| ENSGALG00010023665 | C16orf87 | C11H16ORF87 | | 99 | 112 | 322 | 394 | 255 | 105 | 127 | 128 | 0.00016112 | 0.002077649 |
| ENSGALG00010023666 | MUC1 | MUC1 | | 370 | 668 | 212 | 252 | 205 | 148 | 425 | 412 | 0.00023558 | 0.002857628 |
| ENSGALG00010023668 | RCN1 | RCN1L | | 1362 | 1595 | 3115 | 3930 | 2549 | 1722 | 1815 | 2088 | 4.69E-07 | 1.33E-05 |
| ENSGALG00010023680 | MPZL3 | MPZL3 | | 2428 | 977 | 360 | 376 | 704 | 1014 | 2567 | 1719 | 1.62E-08 | 6.54E-07 |
| ENSGALG00010023686 | MYLK3 | MYLK3 | | 73 | 83 | 185 | 287 | 143 | 70 | 99 | 52 | 0.001904609 | 0.015541907 |
| ENSGALG00010023707 | MPZL2 | EVA1 | | 454 | 227 | 47 | 22 | 188 | 153 | 87 | 114 | 4.25E-09 | 1.99E-07 |
| ENSGALG00010023724 | CYP2R1 | CYP2R1 | | 137 | 168 | 651 | 584 | 367 | 250 | 292 | 220 | 4.66E-07 | 1.33E-05 |
| ENSGALG00010023736 | LOC107050879 | KIF5A | | 885 | 759 | 156 | 155 | 186 | 126 | 203 | 258 | 1.88E-15 | 2.34E-13 |
| ENSGALG00010023742 | NA | NA | | 11 | 20 | 7 | 2 | 1 | 1 | 9 | 20 | 0.001860346 | 0.015247721 |
| ENSGALG00010023745 | NA | NA | | 51 | 67 | 205 | 196 | 211 | 141 | 149 | 103 | 0.00028425 | 0.003342984 |
| ENSGALG00010023747 | NA | NA | | 25 | 25 | 160 | 242 | 129 | 122 | 97 | 113 | 5.01E-08 | 1.82E-06 |
| ENSGALG00010023749 | RPP25 | RPP25 | | 118 | 79 | 28 | 28 | 68 | 161 | 35 | 21 | 1.19E-05 | 0.000227852 |
| ENSGALG00010023766 | NA | NA | | 0 | 1 | 8 | 11 | 14 | 12 | 17 | 24 | 1.57E-05 | 0.000287706 |
| ENSGALG00010023805 | GALNT6 | GALNT6 | | 425 | 354 | 211 | 117 | 298 | 156 | 360 | 374 | 0.009092588 | 0.054153716 |
| ENSGALG00010023806 | PATJ | INADL | | 625 | 343 | 183 | 253 | 288 | 311 | 865 | 433 | 0.0022966 | 0.018131972 |
| ENSGALG00010023831 | SATB2 | SATB2 | | 108 | 127 | 563 | 453 | 314 | 198 | 186 | 160 | 2.62E-07 | 7.96E-06 |
| ENSGALG00010023863 | EYA2 | EYA2 | | 33 | 74 | 168 | 177 | 129 | 104 | 152 | 227 | 0.001850773 | 0.015206551 |
| ENSGALG00010023864 | EPHA1 | EPHA9 | | 730 | 466 | 131 | 139 | 186 | 198 | 467 | 508 | 8.80E-09 | 3.78E-07 |
| ENSGALG00010023875 | NA | NA | | 2 | 0 | 4 | 13 | 0 | 2 | 18 | 5 | 0.009942419 | 0.058076795 |
| ENSGALG00010023883 | TBX21 | TBX21 | | 100 | 114 | 16 | 19 | 38 | 11 | 80 | 51 | 8.52E-06 | 0.000171435 |
| ENSGALG00010023888 | CTH | CTH | | 1387 | 1248 | 838 | 784 | 628 | 804 | 1837 | 2431 | 9.11E-07 | 2.39E-05 |
| ENSGALG00010023902 | PTGER3 | PTGER3 | | 355 | 731 | 1240 | 1670 | 445 | 502 | 679 | 685 | 3.61E-05 | 0.000585314 |
| ENSGALG00010023937 | XBP1 | XBP1 | | 2324 | 1784 | 1186 | 1293 | 1341 | 1163 | 1718 | 1810 | 0.007834307 | 0.048046584 |
| ENSGALG00010023959 | NA | NA | | 1 | 1 | 3 | 1 | 7 | 6 | 11 | 21 | 0.000212829 | 0.002632715 |
| ENSGALG00010023965 | NA | NA | | 14 | 7 | 2 | 2 | 21 | 20 | 27 | 8 | 0.001169402 | 0.010563806 |
| ENSGALG00010023967 | PRPH | PRPH | | 478 | 335 | 134 | 179 | 121 | 98 | 250 | 274 | 4.04E-05 | 0.000641202 |
| ENSGALG00010023978 | TFDP2 | TFDP2 | | 4581 | 6429 | 8031 | 9636 | 7450 | 6002 | 5729 | 4415 | 0.000184155 | 0.002331515 |
| ENSGALG00010023992 | FAM131B | FAM131B | | 194 | 149 | 128 | 77 | 110 | 94 | 449 | 216 | 0.001274964 | 0.011333613 |
| ENSGALG00010024001 | LOC424473 | LOC424473 | | 37 | 29 | 114 | 93 | 64 | 56 | 118 | 81 | 0.00892152 | 0.053324905 |
| ENSGALG00010024023 | LHX8 | LHX7 | | 0 | 5 | 27 | 19 | 8 | 1 | 4 | 2 | 0.001505544 | 0.012916144 |
| ENSGALG00010024038 | NKD1 | NKD1 | | 122 | 256 | 211 | 344 | 517 | 520 | 207 | 294 | 6.34E-05 | 0.000934853 |
| ENSGALG00010024049 | RCN2 | RCN2 | | 4827 | 5305 | 10631 | 12847 | 9418 | 5767 | 6375 | 6451 | 6.74E-09 | 3.00E-07 |
| ENSGALG00010024063 | BRD7 | BRD7 | | 1206 | 1862 | 3373 | 2598 | 2626 | 1513 | 2469 | 2602 | 0.005052483 | 0.033951384 |
| ENSGALG00010024073 | RABGGTB | RABGGTB | | 2086 | 2238 | 3328 | 4003 | 2655 | 2114 | 2230 | 2356 | 0.001913897 | 0.015602459 |
| ENSGALG00010024096 | MYO1D | MYO1D | | 998 | 1049 | 624 | 602 | 632 | 400 | 432 | 558 | 0.000782385 | 0.007664254 |
| ENSGALG00010024128 | PHLDA2 | PHLDA2 | | 179 | 213 | 162 | 211 | 67 | 71 | 216 | 289 | 0.003251969 | 0.023976549 |
| ENSGALG00010024162 | SRCIN1 | SRCIN1 | | 1133 | 1505 | 639 | 627 | 890 | 488 | 1368 | 1142 | 5.10E-05 | 0.000783629 |
| ENSGALG00010024163 | HS6ST1 | CHS6ST-1 | | 58 | 105 | 221 | 208 | 145 | 97 | 69 | 70 | 0.001140192 | 0.010355921 |
| ENSGALG00010024172 | ABCG4 | ABCG4 | | 231 | 253 | 35 | 26 | 48 | 42 | 135 | 135 | 3.94E-11 | 2.65E-09 |
| ENSGALG00010024173 | CBLN1 | CBLN1 | | 592 | 990 | 2224 | 2058 | 1878 | 938 | 892 | 897 | 6.26E-07 | 1.72E-05 |
| ENSGALG00010024193 | GSTK1 | GSTK1 | | 315 | 241 | 707 | 797 | 533 | 444 | 380 | 504 | 0.000500468 | 0.00529021 |
| ENSGALG00010024194 | NKAIN4 | C20orf58 | | 598 | 984 | 107 | 127 | 105 | 87 | 408 | 490 | 7.59E-19 | 1.36E-16 |
| ENSGALG00010024210 | NXPH3 | NXPH3 | | 551 | 297 | 43 | 27 | 40 | 36 | 150 | 146 | 1.11E-15 | 1.44E-13 |
| ENSGALG00010024214 | CRABP1 | CRABP-I | | 401 | 189 | 42 | 89 | 92 | 133 | 234 | 176 | 0.000376263 | 0.004225884 |
| ENSGALG00010024218 | NGFR | NGFR | | 789 | 1408 | 527 | 358 | 672 | 380 | 1371 | 913 | 5.65E-05 | 0.000850425 |
| ENSGALG00010024222 | KCNK15 | KCNK15 | | 93 | 101 | 88 | 45 | 96 | 162 | 307 | 201 | 0.001359217 | 0.011930272 |
| ENSGALG00010024228 | LOC418303 | CD163 | | 24 | 17 | 81 | 61 | 100 | 54 | 4 | 7 | 4.36E-10 | 2.40E-08 |
| ENSGALG00010024233 | FAIM2 | FAIM2 | | 4 | 5 | 1 | 0 | 4 | 0 | 17 | 22 | 1.33E-05 | 0.000250262 |
| ENSGALG00010024234 | ARHGEF4 | ARHGEF4 | | 16 | 19 | 92 | 84 | 72 | 14 | 9 | 20 | 0.000176664 | 0.002255469 |
| ENSGALG00010024266 | WISP2 | WISP2 | | 0 | 1 | 3 | 7 | 10 | 8 | 14 | 8 | 0.001604832 | 0.01360038 |
| ENSGALG00010024280 | DUSP3 | DUSP3 | | 398 | 482 | 923 | 897 | 694 | 466 | 499 | 453 | 0.001517343 | 0.012970763 |
| ENSGALG00010024286 | ADA | ADA | | 383 | 736 | 976 | 1330 | 1033 | 1127 | 844 | 1012 | 0.001382719 | 0.012104764 |
| ENSGALG00010024310 | NA | NA | | 2 | 2 | 31 | 11 | 6 | 4 | 8 | 3 | 0.003326024 | 0.02441129 |
| ENSGALG00010024325 | PCBP2 | PCBP2 | | 24953 | 21501 | 15248 | 16329 | 15039 | 10148 | 15336 | 19712 | 0.005138398 | 0.034417999 |
| ENSGALG00010024327 | PPP1R1B | PPP1R1B | | 186 | 239 | 119 | 102 | 55 | 57 | 126 | 253 | 0.002368456 | 0.018539406 |
| ENSGALG00010024335 | AMHR2 | AMHR2 | | 1134 | 1005 | 761 | 1134 | 764 | 596 | 461 | 579 | 0.002105818 | 0.016927574 |
| ENSGALG00010024347 | OTOG | OTOGL | | 630 | 185 | 82 | 140 | 501 | 853 | 1672 | 490 | 2.38E-06 | 5.63E-05 |
| ENSGALG00010024370 | HERPUD1 | HERPUD1 | | 1813 | 1533 | 990 | 1060 | 992 | 805 | 1199 | 1125 | 0.005363524 | 0.035654419 |
| ENSGALG00010024386 | TPPP3 | TPPP3 | | 299 | 441 | 164 | 128 | 101 | 35 | 270 | 135 | 5.93E-05 | 0.000881949 |
| ENSGALG00010024421 | DAPK2 | DAPK2 | | 465 | 590 | 996 | 1123 | 1062 | 681 | 613 | 541 | 9.49E-05 | 0.00132824 |
| ENSGALG00010024424 | SAMHD1 | SAMHD1 | | 1470 | 2056 | 3753 | 4234 | 2460 | 1843 | 1631 | 1441 | 5.25E-10 | 2.84E-08 |
| ENSGALG00010024431 | FEZ1 | FEZ1 | | 801 | 1252 | 491 | 672 | 596 | 598 | 339 | 338 | 3.51E-06 | 7.93E-05 |
| ENSGALG00010024437 | HERC1 | HERC1 | | 3618 | 3422 | 4144 | 4001 | 5699 | 5402 | 8569 | 3931 | 0.001567397 | 0.013330452 |
| ENSGALG00010024452 | NA | NA | | 7029 | 7147 | 3347 | 4404 | 3662 | 3160 | 5189 | 4610 | 1.59E-06 | 3.97E-05 |
| ENSGALG00010024475 | ADGRG1 | GPR56 | | 1241 | 1444 | 807 | 669 | 890 | 636 | 1277 | 1481 | 0.000343934 | 0.003910088 |
| ENSGALG00010024476 | KCNB1 | KCNB1 | | 245 | 136 | 48 | 31 | 47 | 45 | 218 | 188 | 2.06E-07 | 6.41E-06 |
| ENSGALG00010024478 | NA | NA | | 3 | 10 | 16 | 10 | 17 | 18 | 42 | 32 | 0.002526653 | 0.019513899 |
| ENSGALG00010024502 | VPS8 | VPS8 | | 2010 | 2338 | 2181 | 2283 | 1414 | 848 | 1160 | 1290 | 7.53E-05 | 0.001088263 |
| ENSGALG00010024509 | SEPTIN5 | | 5-Sep | 2867 | 2577 | 2426 | 3997 | 2584 | 3263 | 5701 | 5191 | 0.000848454 | 0.008196166 |
| ENSGALG00010024516 | EHHADH | EHHADH | | 264 | 309 | 545 | 724 | 404 | 306 | 246 | 289 | 0.000872505 | 0.008365637 |
| ENSGALG00010024520 | MAP3K13 | MAP3K13 | | 270 | 220 | 162 | 154 | 240 | 233 | 456 | 336 | 0.005996435 | 0.038736922 |

| | | | | | | | | | | | | |
|---|---|---|---|---|---|---|---|---|---|---|---|---|
| ENSGALG00010024545 | PHACTR3 | PHACTR3 | 494 | 479 | 217 | 228 | 241 | 241 | 512 | 309 | 0.006569317 | 0.041680494 |
| ENSGALG00010024550 | KCNA10 | cKv1.2 | 15 | 23 | 1 | 0 | 1 | 0 | 1 | 2 | 1.13E-07 | 3.79E-06 |
| ENSGALG00010024555 | EPHB1 | CEK6 | 801 | 1027 | 697 | 696 | 487 | 454 | 1023 | 1121 | 0.002832921 | 0.021393485 |
| ENSGALG00010024556 | LOC107049457 | LOC107049457 | 429 | 584 | 490 | 663 | 424 | 263 | 231 | 285 | 0.008534391 | 0.051415555 |
| ENSGALG00010024561 | ROBO3 | ROBO3 | 122 | 421 | 72 | 53 | 79 | 57 | 240 | 182 | 8.27E-05 | 0.001179786 |
| ENSGALG00010024569 | ASCL2 | ASCL2 | 15 | 39 | 121 | 136 | 79 | 57 | 42 | 73 | 0.000948563 | 0.008935954 |
| ENSGALG00010024571 | KCNA3 | cKv1.1 | 700 | 777 | 341 | 298 | 461 | 234 | 691 | 612 | 0.000221034 | 0.002712138 |
| ENSGALG00010024577 | CRYGS | CRYGS | 34 | 18 | 18 | 5 | 3 | 2 | 10 | 17 | 0.00751152 | 0.0464105 |
| ENSGALG00010024588 | TTC39A | C1orf34 | 20657 | 14970 | 11123 | 17270 | 19514 | 18578 | 13503 | 11977 | 0.001291137 | 0.011440872 |
| ENSGALG00010024597 | DNAJB11 | DNAJB11 | 6452 | 6432 | 4616 | 6100 | 4265 | 3203 | 4304 | 4216 | 0.003608224 | 0.026081591 |
| ENSGALG00010024605 | LSP1P1 | LSP1 | 374 | 705 | 166 | 166 | 196 | 188 | 397 | 453 | 9.84E-06 | 0.000193421 |
| ENSGALG00010024606 | NOBOX | NOBOX | 38 | 15 | 2 | 0 | 5 | 13 | 56 | 24 | 6.80E-06 | 0.000140592 |
| ENSGALG00010024608 | DENND2D | DENND2D | 785 | 414 | 249 | 282 | 221 | 369 | 756 | 653 | 0.002403547 | 0.018743855 |
| ENSGALG00010024618 | VPS13C | VPS13C | 310 | 380 | 735 | 559 | 538 | 565 | 1040 | 613 | 0.005229565 | 0.034930651 |
| ENSGALG00010024622 | PRMT1 | PRMT8 | 12 | 11 | 63 | 65 | 34 | 21 | 12 | 6 | 1.77E-06 | 4.37E-05 |
| ENSGALG00010024629 | ESAM | ESAM | 385 | 370 | 40 | 39 | 92 | 53 | 134 | 109 | 7.99E-14 | 7.90E-12 |
| ENSGALG00010024643 | ETV7 | ETV7 | 113 | 68 | 226 | 255 | 246 | 180 | 207 | 202 | 0.004545131 | 0.031413662 |
| ENSGALG00010024659 | BEND5 | C1orf165 | 2490 | 2839 | 1671 | 2042 | 2817 | 1954 | 1559 | 1586 | 1.82E-05 | 0.000324927 |
| ENSGALG00010024662 | MOB2 | MOB2 | 3104 | 4594 | 2128 | 2535 | 1956 | 1630 | 2803 | 3374 | 0.000172484 | 0.002204968 |
| ENSGALG00010024674 | DHH | DHH | 2 | 4 | 6 | 2 | 5 | 0 | 36 | 20 | 5.47E-05 | 0.000827234 |
| ENSGALG00010024678 | AMOTL2 | AMOTL2 | 986 | 1664 | 480 | 575 | 480 | 601 | 865 | 962 | 5.74E-05 | 0.000860277 |
| ENSGALG00010024682 | LOC100859737 | TUBA1A1 | 42938 | 41655 | 10269 | 11482 | 10581 | 7940 | 31212 | 38640 | 4.48E-32 | 2.08E-29 |
| ENSGALG00010024684 | CADNL | CADNL | 326 | 243 | 925 | 692 | 438 | 401 | 457 | 218 | 0.00109954 | 0.010107539 |
| ENSGALG00010024686 | CMPK1 | CK | 4626 | 4919 | 3292 | 4193 | 3396 | 2078 | 2899 | 3405 | 0.005307455 | 0.035356486 |
| ENSGALG00010024688 | CADM1 | CRTAM | 4905 | 6464 | 4090 | 4097 | 3529 | 2667 | 5600 | 6612 | 0.000105861 | 0.001447763 |
| ENSGALG00010024690 | LOC100857858 | TUBA1C | 2373 | 2287 | 8829 | 9833 | 8382 | 5734 | 1223 | 1862 | 1.31E-39 | 8.10E-37 |
| ENSGALG00010024693 | ANXA2 | ANX2 | 3188 | 5038 | 4159 | 7510 | 4052 | 2731 | 2511 | 2470 | 0.000125051 | 0.001669214 |
| ENSGALG00010024695 | TRABD2B | TRABD2B | 689 | 597 | 284 | 382 | 505 | 497 | 391 | 387 | 0.002469549 | 0.019169112 |
| ENSGALG00010024700 | TOLLIP | TOLLIP | 10089 | 12523 | 14166 | 20134 | 15446 | 11415 | 11301 | 11121 | 0.004932514 | 0.033468197 |
| ENSGALG00010024704 | FOXB1 | FOXB1 | 340 | 496 | 139 | 136 | 263 | 193 | 399 | 396 | 2.15E-05 | 0.000375507 |
| ENSGALG00010024707 | LOC107057295 | MANBAL1 | 6499 | 4171 | 6301 | 7336 | 1869 | 1385 | 4228 | 3931 | 4.44E-12 | 3.54E-10 |
| ENSGALG00010024723 | ITCH | ITCH | 1989 | 1367 | 1043 | 1043 | 1339 | 1378 | 1020 | 854 | 0.001068273 | 0.009852635 |
| ENSGALG00010024725 | NA | NA | 20 | 6 | 79 | 57 | 48 | 49 | 50 | 46 | 0.00213458 | 0.017117608 |
| ENSGALG00010024745 | PMEL | MMP115 | 3476 | 3433 | 1088 | 1840 | 1265 | 1129 | 1936 | 2550 | 4.95E-08 | 1.81E-06 |
| ENSGALG00010024752 | MPP2 | MPP2 | 647 | 616 | 347 | 270 | 404 | 358 | 882 | 779 | 3.24E-05 | 0.000534529 |
| ENSGALG00010024772 | ELAPOR1 | KIAA1324 | 595 | 662 | 376 | 409 | 505 | 393 | 697 | 788 | 0.008384351 | 0.05067316 |
| ENSGALG00010024796 | ACTBL2 | ACTBL2L | 85 | 54 | 12 | 20 | 35 | 30 | 118 | 40 | 0.001410252 | 0.012281466 |
| ENSGALG00010024802 | CD59 | CD59 | 405 | 1534 | 274 | 365 | 739 | 621 | 600 | 1008 | 0.002708266 | 0.020694486 |
| ENSGALG00010024820 | NUDT19 | NUDT19 | 731 | 590 | 1167 | 1220 | 666 | 434 | 672 | 507 | 0.002451549 | 0.01907321 |
| ENSGALG00010024829 | ERBB3 | ERBB3 | 1418 | 799 | 311 | 307 | 470 | 456 | 962 | 535 | 5.89E-06 | 0.0001242 |
| ENSGALG00010024833 | MAPT | MAPT | 253 | 761 | 1944 | 2081 | 1129 | 478 | 396 | 587 | 3.28E-07 | 9.75E-06 |
| ENSGALG00010024863 | LMOD1 | LMOD1 | 12 | 18 | 50 | 51 | 20 | 24 | 46 | 78 | 0.003044614 | 0.022735291 |
| ENSGALG00010024865 | SNAP47 | C1orf142 | 472 | 524 | 310 | 257 | 221 | 124 | 270 | 337 | 0.002398218 | 0.018711025 |
| ENSGALG00010024872 | RUNDC3A | RUNDC3A | 1408 | 1259 | 118 | 288 | 153 | 85 | 554 | 292 | 9.97E-17 | 1.40E-14 |
| ENSGALG00010024883 | ING5 | ING5 | 2973 | 2766 | 1177 | 1216 | 806 | 590 | 1572 | 1487 | 2.50E-15 | 3.02E-13 |
| ENSGALG00010024909 | SLC39A5 | SLC39A5 | 183 | 217 | 86 | 109 | 72 | 60 | 66 | 80 | 0.000893531 | 0.008508656 |
| ENSGALG00010024930 | TMEM237 | ALS2CR4 | 1533 | 1677 | 2483 | 2114 | 2222 | 1282 | 1424 | 1048 | 0.004989166 | 0.033702079 |
| ENSGALG00010024950 | NCAM1 | N-CAM | 3346 | 5562 | 7876 | 9301 | 8123 | 4708 | 5045 | 4248 | 3.44E-05 | 0.000561177 |
| ENSGALG00010024951 | LOC124417104 | LOC124417104 | 50 | 45 | 263 | 248 | 133 | 88 | 53 | 63 | 3.40E-08 | 1.26E-06 |
| ENSGALG00010024970 | CAB39 | CAB39 | 3403 | 4306 | 2495 | 2623 | 4120 | 3146 | 2715 | 2843 | 2.84E-05 | 0.000477452 |
| ENSGALG00010024972 | LOC107049570 | PYY | 50 | 19 | 3 | 8 | 15 | 9 | 60 | 64 | 3.83E-06 | 8.56E-05 |
| ENSGALG00010024974 | CD44 | CD44 | 201 | 120 | 46 | 61 | 158 | 116 | 147 | 87 | 0.003144586 | 0.023330745 |
| ENSGALG00010024984 | FAM159A | FAM159A | 236 | 139 | 27 | 28 | 48 | 114 | 193 | 128 | 2.88E-05 | 0.000481709 |
| ENSGALG00010025003 | IGFN1 | IGFN1 | 86 | 168 | 235 | 178 | 283 | 263 | 346 | 256 | 0.007101913 | 0.044248139 |
| ENSGALG00010025006 | LOC101751434 | RETSATL | 69 | 23 | 3 | 1 | 53 | 53 | 111 | 40 | 1.08E-07 | 3.67E-06 |
| ENSGALG00010025010 | SLC7A10 | SLC7A10 | 43 | 46 | 100 | 176 | 48 | 42 | 68 | 39 | 0.005382609 | 0.035724447 |
| ENSGALG00010025028 | LOC107051857 | LOC107051857 | 270 | 950 | 1845 | 1760 | 1354 | 791 | 331 | 838 | 0.000749817 | 0.007414759 |
| ENSGALG00010025031 | ECHDC2 | ECHDC2 | 763 | 674 | 1432 | 1509 | 1254 | 994 | 1339 | 1545 | 0.000306943 | 0.003559777 |
| ENSGALG00010025032 | FSTL1 | FLIK | 4852 | 6581 | 9349 | 11097 | 7298 | 6042 | 5630 | 7352 | 0.001030745 | 0.009559278 |
| ENSGALG00010025035 | NA | NA | 28 | 42 | 8 | 2 | 12 | 8 | 7 | 11 | 0.000410087 | 0.004511764 |
| ENSGALG00010025063 | TEX12 | TEX12 | 23 | 13 | 133 | 74 | 92 | 56 | 47 | 57 | 0.000180786 | 0.002294079 |
| ENSGALG00010025076 | NA | NA | 71 | 116 | 28 | 5 | 30 | 11 | 38 | 35 | 0.000649099 | 0.006602398 |
| ENSGALG00010025092 | COL1A1 | COL1A1 | 3116 | 2647 | 497 | 171 | 1036 | 751 | 2330 | 2457 | 7.24E-15 | 8.29E-13 |
| ENSGALG00010025111 | NA | NA | 3 | 3 | 42 | 40 | 24 | 30 | 53 | 35 | 1.59E-06 | 3.97E-05 |
| ENSGALG00010025117 | NA | NA | 65 | 189 | 28 | 36 | 75 | 53 | 70 | 71 | 0.003624301 | 0.026152554 |
| ENSGALG00010025127 | PAMR1 | PAMR1 | 1 | 0 | 15 | 12 | 18 | 6 | 3 | 9 | 0.000932994 | 0.008819119 |
| ENSGALG00010025130 | ATP2A2 | SERCA2 | 9413 | 8790 | 6790 | 7569 | 5222 | 4198 | 6408 | 6156 | 0.000640161 | 0.006523399 |
| ENSGALG00010025142 | NA | NA | 1 | 1 | 21 | 12 | 8 | 10 | 15 | 15 | 0.00073038 | 0.007271804 |
| ENSGALG00010025164 | C11orf52 | C24H11orf52 | 8 | 21 | 92 | 66 | 51 | 20 | 21 | 34 | 0.001198061 | 0.010799353 |
| ENSGALG00010025225 | DMRTB1 | DMRTB1 | 3715 | 6838 | 7773 | 13577 | 11903 | 6634 | 4422 | 4639 | 5.02E-07 | 1.42E-05 |
| ENSGALG00010025233 | WDR12 | WDR12 | 2226 | 2189 | 2350 | 3204 | 2321 | 1671 | 1565 | 1663 | 0.00924794 | 0.054863887 |
| ENSGALG00010025238 | IGSF9B | IGSF9B | 152 | 248 | 113 | 95 | 87 | 144 | 674 | 187 | 0.001132403 | 0.01029321 |
| ENSGALG00010025252 | PPP1R19B | NUR2L | 810 | 1108 | 482 | 265 | 603 | 430 | 661 | 814 | 0.00066401 | 0.006725414 |
| ENSGALG00010025253 | NA | NA | 7 | 12 | 42 | 36 | 27 | 14 | 52 | 62 | 0.000261434 | 0.003127472 |
| ENSGALG00010025263 | SAMD14 | SAMD14 | 1221 | 889 | 98 | 79 | 215 | 148 | 309 | 361 | 6.44E-23 | 1.58E-20 |
| ENSGALG00010025286 | ITGA3 | ITGA3 | 1707 | 1593 | 663 | 378 | 804 | 501 | 1219 | 1305 | 5.11E-08 | 1.85E-06 |
| ENSGALG00010025298 | NACAD | NACAD | 1191 | 1241 | 394 | 325 | 758 | 670 | 1577 | 1801 | 7.64E-16 | 1.01E-13 |
| ENSGALG00010025320 | JAM3 | JAM3 | 1129 | 1402 | 691 | 971 | 501 | 577 | 1000 | 1212 | 0.004015147 | 0.028408606 |
| ENSGALG00010025323 | KLHDC8A | KLHDC8A | 56 | 156 | 412 | 361 | 219 | 164 | 169 | 188 | 0.000760041 | 0.007489269 |
| ENSGALG00010025325 | DPEP2 | DPEP3 | 2 | 7 | 111 | 262 | 49 | 31 | 5 | 5 | 7.42E-20 | 1.48E-17 |
| ENSGALG00010025329 | MYO1A | MYO1A | 722 | 806 | 189 | 283 | 296 | 280 | 552 | 743 | 1.91E-07 | 6.03E-06 |
| ENSGALG00010025330 | USH1C | USH1C | 90 | 139 | 106 | 103 | 265 | 160 | 200 | 143 | 0.009292004 | 0.055034964 |
| ENSGALG00010025331 | NA | NA | 36 | 55 | 12 | 20 | 94 | 68 | 81 | 69 | 1.16E-05 | 0.000222595 |
| ENSGALG00010025357 | OLFML3 | ONT1 | 54 | 40 | 85 | 67 | 45 | 68 | 122 | 193 | 0.006880996 | 0.043169771 |
| ENSGALG00010025368 | LRP1 | A2MR | 9164 | 12232 | 7472 | 7056 | 7491 | 4761 | 11077 | 9828 | 0.000959888 | 0.009027376 |
| ENSGALG00010025370 | HPD | HPD | 321 | 297 | 141 | 129 | 127 | 141 | 210 | 335 | 0.005696845 | 0.037320925 |
| ENSGALG00010025375 | NA | NA | 0 | 1 | 53 | 36 | 45 | 12 | 0 | 0 | 2.83E-14 | 2.95E-12 |
| ENSGALG00010025378 | PAX3 | PAX3 | 131 | 180 | 49 | 61 | 78 | 36 | 108 | 104 | 0.002978408 | 0.022310694 |
| ENSGALG00010025392 | SLC6A2 | SLC6A2 | 130 | 197 | 304 | 445 | 432 | 171 | 150 | 160 | 0.002737548 | 0.020833292 |
| ENSGALG00010025406 | GLB1L2 | GLB1L3 | 2067 | 1925 | 3620 | 3871 | 2435 | 1962 | 2075 | 1995 | 5.71E-05 | 0.000856688 |
| ENSGALG00010025414 | MMP2 | MMP2 | 11 | 72 | 108 | 385 | 124 | 53 | 21 | 98 | 0.00847071 | 0.05113051 |
| ENSGALG00010025428 | VANGL1 | VANGL1 | 1469 | 1501 | 941 | 1159 | 1588 | 1689 | 2778 | 1251 | 0.008053793 | 0.049158139 |
| ENSGALG00010025431 | NA | NA | 2 | 9 | 0 | 0 | 3 | 8 | 17 | 5 | 0.002073582 | 0.016732807 |
| ENSGALG00010025432 | NA | NA | 29 | 66 | 821 | 823 | 215 | 143 | 156 | 197 | 4.45E-21 | 1.03E-18 |
| ENSGALG00010025438 | EXOSC6 | EXOSC6 | 848 | 589 | 501 | 451 | 836 | 595 | 452 | 225 | 0.000936329 | 0.008845639 |
| ENSGALG00010025445 | B3GNT4 | B3GNT4 | 5395 | 4257 | 1236 | 1355 | 2526 | 2837 | 4277 | 4039 | 7.85E-15 | 8.86E-13 |
| ENSGALG00010025451 | LRRC43 | LRRC43 | 26 | 13 | 68 | 59 | 71 | 50 | 59 | 44 | 0.009392465 | 0.055477868 |
| ENSGALG00010025454 | NA | NA | 1 | 1 | 1 | 0 | 13 | 22 | 34 | 11 | 1.12E-07 | 3.76E-06 |
| ENSGALG00010025464 | VPS33A | VPS33A | 3782 | 3091 | 1302 | 1336 | 1992 | 1920 | 3399 | 2654 | 1.03E-08 | 4.34E-07 |
| ENSGALG00010025488 | SLC4A8 | SLC4A8 | 635 | 961 | 388 | 563 | 300 | 254 | 585 | 596 | 0.000689544 | 0.006912811 |
| ENSGALG00010025491 | ZMYND11 | ZMYND11 | 61 | 78 | 319 | 284 | 231 | 102 | 149 | 101 | 7.68E-05 | 0.001105088 |
| ENSGALG00010025499 | LOC107050760 | STAC3 | 43 | 30 | 11 | 6 | 7 | 12 | 33 | 28 | 0.002082341 | 0.016779207 |
| ENSGALG00010025521 | NA | NA | 0 | 0 | 0 | 0 | 20 | 14 | 1 | 0 | 8.44E-11 | 5.26E-09 |
| ENSGALG00010025546 | INHBE | INHBE | 43 | 27 | 4 | 1 | 0 | 5 | 12 | 15 | 8.09E-06 | 0.000163083 |
| ENSGALG00010025559 | SLC6A17 | SLC6A17 | 595 | 755 | 137 | 96 | 211 | 99 | 537 | 356 | 3.19E-11 | 2.18E-09 |
| ENSGALG00010025568 | MYL3 | MLC | 25 | 15 | 212 | 260 | 54 | 28 | 18 | 26 | 1.18E-14 | 1.29E-12 |
| ENSGALG00010025573 | CLMP | ASAM | 27 | 26 | 90 | 133 | 60 | 30 | 54 | 79 | 0.00235651 | 0.018470835 |
| ENSGALG00010025576 | NA | NA | 10 | 25 | 47 | 50 | 73 | 60 | 31 | 29 | 0.000693437 | 0.006943509 |
| ENSGALG00010025580 | RASSF7 | RASSF7 | 224 | 346 | 147 | 188 | 174 | 106 | 412 | 392 | 0.000742555 | 0.007356005 |
| ENSGALG00010025583 | NA | NA | 46 | 58 | 171 | 160 | 103 | 93 | 75 | 90 | 0.003340248 | 0.024461973 |
| ENSGALG00010025588 | LOC101747602 | LOC101747602 | 112 | 60 | 34 | 67 | 148 | 160 | 223 | 155 | 0.000287887 | 0.003373892 |
| ENSGALG00010025589 | LOC121111680 | LOC121111680 | 106 | 179 | 276 | 278 | 316 | 330 | 204 | 175 | 0.001458373 | 0.012568269 |
| ENSGALG00010025590 | HRAS | H-RAS | 5032 | 6266 | 3548 | 4071 | 3776 | 2549 | 4092 | 4202 | 0.001429174 | 0.012376319 |
| ENSGALG00010025597 | SCN2A | SCN2A | 3 | 3 | 23 | 11 | 32 | 38 | 22 | 11 | 4.91E-05 | 0.000758849 |
| ENSGALG00010025613 | CDKN1A | CIP1 | 8 | 48 | 143 | 162 | 63 | 67 | 64 | 326 | 0.00207458 | 0.016732807 |
| ENSGALG00010025623 | RAB44 | RAB44 | 6 | 3 | 9 | 9 | 3 | 0 | 13 | 28 | 0.001686093 | 0.014138477 |
| ENSGALG00010025641 | LOC107054281 | LOC107054281 | 1 | 8 | 14 | 35 | 20 | 29 | 17 | 6 | 0.009820605 | 0.057505939 |
| ENSGALG00010025644 | PI16 | PI16 | 21 | 35 | 47 | 30 | 83 | 55 | 99 | 100 | 0.000872078 | 0.008365637 |
| ENSGALG00010025646 | NA | NA | 20 | 49 | 162 | 133 | 202 | 158 | 69 | 73 | 6.59E-07 | 1.80E-05 |
| ENSGALG00010025649 | NA | NA | 1 | 0 | 8 | 5 | 18 | 15 | 15 | 3 | 0.000413443 | 0.004542704 |
| ENSGALG00010025651 | NA | NA | 0 | 0 | 0 | 0 | 72 | 60 | 3 | 1 | 7.72E-27 | 2.53E-24 |
| ENSGALG00010025658 | PDE1B | PDE1B | 676 | 450 | 147 | 101 | 301 | 205 | 603 | 516 | 1.63E-08 | 6.57E-07 |
| ENSGALG00010025665 | NA | NA | 30 | 7 | 2 | 3 | 11 | 31 | 55 | 26 | 0.000579849 | 0.006000275 |
| ENSGALG00010025666 | CUX2 | CUTL2 | 207 | 227 | 84 | 86 | 93 | 64 | 245 | 175 | 0.000362246 | 0.004090445 |
| ENSGALG00010025668 | NA | NA | 4 | 3 | 18 | 12 | 16 | 21 | 7 | 0 | 0.002820971 | 0.021336134 |
| ENSGALG00010025671 | CHDSD | CHDSD | 6 | 14 | 0 | 0 | 1 | 3 | 15 | 23 | 1.68E-06 | 4.17E-05 |
| ENSGALG00010025672 | NA | NA | 0 | 0 | 17 | 15 | 6 | 0 | 0 | 1 | 7.53E-06 | 0.000153338 |
| ENSGALG00010025674 | GRB14 | GRB14 | 54 | 54 | 145 | 136 | 79 | 50 | 57 | 48 | 0.007089866 | 0.044194641 |
| ENSGALG00010025687 | NA | NA | 13 | 32 | 5 | 10 | 38 | 38 | 41 | 25 | 0.001023511 | 0.009529197 |
| ENSGALG00010025691 | NA | NA | 1068 | 789 | 421 | 477 | 533 | 337 | 687 | 642 | 0.00086042 | 0.008278263 |
| ENSGALG00010025713 | LOC107054297 | LOC107054297 | 778 | 1333 | 280 | 1038 | 154 | 141 | 153 | 827 | 0.001285767 | 0.011405384 |

| | | | | | | | | | | | | |
|---|---|---|---|---|---|---|---|---|---|---|---|---|
| ENSGALG00010025723 | SETD2 | SETD2 | 15193 | 15752 | 10576 | 10239 | 9264 | 7574 | 16779 | 13399 | 0.000340815 | 0.00387726 |
| ENSGALG00010025733 | NA | NA | 2 | 3 | 11 | 35 | 29 | 6 | 7 | 13 | 0.008211634 | 0.049830368 |
| ENSGALG00010025742 | LOC107054293 | LOC107054293 | 618 | 720 | 1531 | 1567 | 840 | 760 | 835 | 756 | 3.34E-05 | 0.000547774 |
| ENSGALG00010025750 | NA | NA | 2 | 6 | 17 | 21 | 4 | 9 | 1 | 1 | 0.000534363 | 0.00561652 |
| ENSGALG00010025753 | SORL1 | LR11 | 701 | 606 | 532 | 522 | 443 | 470 | 1139 | 992 | 0.001028683 | 0.009550751 |
| ENSGALG00010025765 | NA | NA | 171 | 114 | 222 | 239 | 17 | 6 | 275 | 199 | 1.01E-12 | 8.62E-11 |
| ENSGALG00010025767 | NA | NA | 143 | 108 | 20 | 15 | 9 | 19 | 18 | 14 | 3.73E-10 | 2.06E-08 |
| ENSGALG00010025778 | NA | NA | 1 | 24 | 21 | 40 | 1 | 0 | 28 | 19 | 0.000281891 | 0.00332695 |
| ENSGALG00010025780 | NA | NA | 0 | 12 | 65 | 68 | 0 | 0 | 61 | 60 | 1.97E-13 | 1.83E-11 |
| ENSGALG00010025783 | NA | NA | 38 | 69 | 186 | 191 | 17 | 17 | 182 | 183 | 4.83E-11 | 3.20E-09 |
| ENSGALG00010025801 | PTPN7 | PTPN7 | 25 | 15 | 3 | 0 | 9 | 16 | 29 | 14 | 0.000739241 | 0.007329158 |
| ENSGALG00010025832 | ENKD1 | C11H16ORF48 | 1549 | 1944 | 879 | 1125 | 899 | 781 | 1037 | 1106 | 0.000445785 | 0.004816495 |
| ENSGALG00010025833 | NA | NA | 66 | 71 | 128 | 133 | 157 | 94 | 65 | 26 | 0.002994896 | 0.022405648 |
| ENSGALG00010025839 | NA | NA | 8 | 5 | 77 | 40 | 49 | 15 | 10 | 15 | 3.25E-05 | 0.00053499 |
| ENSGALG00010025843 | NA | NA | 21 | 20 | 137 | 102 | 80 | 38 | 53 | 13 | 0.000378683 | 0.004248667 |
| ENSGALG00010025850 | NA | NA | 292 | 228 | 65 | 105 | 118 | 137 | 78 | 108 | 0.000424256 | 0.00463107 |
| ENSGALG00010025860 | NA | NA | 273 | 577 | 243 | 298 | 148 | 133 | 187 | 206 | 0.005273734 | 0.035189627 |
| ENSGALG00010025868 | NA | NA | 92 | 156 | 148 | 152 | 42 | 52 | 326 | 137 | 0.003016104 | 0.022542542 |
| ENSGALG00010025870 | LOC415664 | NADB-LER5 | 70 | 47 | 6 | 4 | 16 | 57 | 36 | 32 | 8.61E-05 | 0.001219391 |
| ENSGALG00010025943 | CATH1 | CATH-1 | 3 | 0 | 10 | 9 | 2 | 0 | 16 | 11 | 0.000941639 | 0.008885758 |
| ENSGALG00010025952 | NA | NA | 170 | 88 | 57 | 108 | 133 | 153 | 319 | 211 | 0.006766304 | 0.04259076 |
| ENSGALG00010025954 | TUBA1A | TUBA1 | 14622 | 18491 | 13587 | 14947 | 14821 | 8458 | 8235 | 10433 | 0.001458978 | 0.012568269 |
| ENSGALG00010025956 | LOC100859726 | SDR9C7 | 14 | 27 | 5 | 7 | 9 | 6 | 21 | 31 | 0.006595088 | 0.041793846 |
| ENSGALG00010025961 | NA | NA | 69 | 9 | 55 | 93 | 188 | 252 | 76 | 102 | 0.000288249 | 0.003375761 |
| ENSGALG00010025975 | NA | NA | 52 | 16 | 43 | 46 | 252 | 127 | 55 | 61 | 4.19E-07 | 1.21E-05 |
| ENSGALG00010026005 | DERL3 | DERL3 | 2193 | 1332 | 784 | 1118 | 771 | 640 | 1211 | 1093 | 0.000512564 | 0.005411221 |
| ENSGALG00010026006 | ITGA2B | ITGA2B | 988 | 664 | 218 | 160 | 255 | 180 | 446 | 614 | 2.90E-09 | 1.39E-07 |
| ENSGALG00010026009 | GUCY1B4 | GUCY1B4 | 269 | 368 | 2385 | 2796 | 1031 | 491 | 392 | 435 | 7.33E-30 | 2.99E-27 |
| ENSGALG00010026011 | GJC2 | GJA12 | 84 | 90 | 212 | 225 | 23 | 35 | 289 | 212 | 1.75E-08 | 6.95E-07 |
| ENSGALG00010026023 | THY1 | THY1 | 495 | 840 | 301 | 337 | 221 | 145 | 512 | 553 | 8.66E-06 | 0.000173578 |
| ENSGALG00010026045 | NA | NA | 10 | 4 | 41 | 23 | 13 | 1 | 6 | 4 | 0.004823818 | 0.032917779 |
| ENSGALG00010026056 | NA | NA | 65 | 33 | 51 | 74 | 145 | 104 | 130 | 91 | 0.005425067 | 0.035956124 |
| ENSGALG00010026066 | SELENOP2 | SELPB | 72 | 118 | 38 | 47 | 24 | 22 | 89 | 129 | 0.000187849 | 0.002369309 |
| ENSGALG00010026111 | LOC771811 | LOC771811 | 37727 | 37706 | 19163 | 23664 | 17128 | 15290 | 30644 | 27074 | 2.80E-06 | 6.48E-05 |
| ENSGALG00010026121 | GSTT1L | GSTT1L | 397 | 386 | 124 | 158 | 78 | 75 | 314 | 197 | 5.06E-07 | 1.42E-05 |
| ENSGALG00010026124 | PACSIN1 | PACSIN1 | 16 | 26 | 6 | 5 | 8 | 12 | 117 | 134 | 2.53E-15 | 3.05E-13 |
| ENSGALG00010026196 | NA | NA | 177 | 123 | 32 | 32 | 269 | 199 | 269 | 219 | 1.09E-10 | 6.58E-09 |
| ENSGALG00010026199 | TBX6 | CH-TBX6L | 10101 | 6998 | 335 | 281 | 2610 | 1955 | 7013 | 3769 | 1.03E-47 | 9.08E-45 |
| ENSGALG00010026208 | NA | NA | 39 | 16 | 4 | 0 | 2 | 7 | 15 | 9 | 0.000855701 | 0.008256623 |
| ENSGALG00010026235 | NA | NA | 14 | 7 | 6 | 3 | 9 | 9 | 35 | 18 | 0.009830494 | 0.057543683 |
| ENSGALG00010026242 | CCDC141 | CCDC141 | 159 | 317 | 330 | 403 | 917 | 336 | 363 | 403 | 0.001246643 | 0.011135164 |
| ENSGALG00010026246 | NA | NA | 21 | 45 | 24 | 7 | 52 | 71 | 66 | 40 | 0.005310976 | 0.035356486 |
| ENSGALG00010026252 | ENO2 | ENO2 | 217 | 179 | 64 | 62 | 115 | 109 | 186 | 182 | 0.000586187 | 0.006043399 |
| ENSGALG00010026264 | FZD7 | Fz-7 | 906 | 1503 | 829 | 732 | 854 | 443 | 1154 | 1660 | 0.003916113 | 0.027825718 |
| ENSGALG00010026268 | GARS | GARS | 3790 | 3497 | 4298 | 4334 | 4442 | 4004 | 5561 | 6569 | 0.002130322 | 0.017091668 |
| ENSGALG00010026269 | NA | NA | 1 | 5 | 2 | 1 | 19 | 8 | 9 | 7 | 0.002209721 | 0.017597056 |
| ENSGALG00010026284 | LOC124417382 | LOC124417382 | 42 | 43 | 36 | 43 | 43 | 39 | 133 | 125 | 0.000214343 | 0.002647524 |
| ENSGALG00010026300 | HMGA1 | HMGA1 | 15844 | 15280 | 9941 | 12275 | 10225 | 7619 | 8656 | 12272 | 0.00788325 | 0.048258197 |
| ENSGALG00010026309 | NA | NA | 22 | 25 | 109 | 100 | 35 | 17 | 22 | 25 | 9.24E-06 | 0.000183082 |
| ENSGALG00010026315 | SYT2 | SYT2 | 38 | 108 | 67 | 56 | 38 | 29 | 275 | 202 | 1.11E-06 | 2.86E-05 |
| ENSGALG00010026333 | NA | NA | 11 | 32 | 69 | 112 | 94 | 57 | 46 | 53 | 0.001431133 | 0.01238586 |
| ENSGALG00010026343 | NA | NA | 4 | 5 | 62 | 28 | 58 | 4 | 20 | 33 | 0.006027294 | 0.038902902 |
| ENSGALG00010026350 | LOC101749943 | LOC101749943 | 189 | 317 | 624 | 967 | 412 | 638 | 621 | 654 | 0.000471093 | 0.005040268 |
| ENSGALG00010026366 | NA | NA | 259 | 205 | 782 | 719 | 501 | 297 | 289 | 241 | 2.57E-06 | 6.01E-05 |
| ENSGALG00010026377 | LRRC75B | C15H22orf36 | 769 | 810 | 345 | 474 | 370 | 354 | 585 | 459 | 0.004040029 | 0.028536333 |
| ENSGALG00010026396 | NA | NA | 4 | 4 | 0 | 2 | 12 | 5 | 19 | 10 | 0.001634999 | 0.013814009 |
| ENSGALG00010026403 | GUCD1 | C15H22ORF13 | 278 | 207 | 439 | 580 | 326 | 240 | 207 | 188 | 0.001960004 | 0.015916227 |
| ENSGALG00010026409 | VILL | AVIL | 33 | 44 | 44 | 61 | 115 | 78 | 69 | 56 | 0.006939742 | 0.043420804 |
| ENSGALG00010026410 | NA | NA | 76 | 60 | 13 | 15 | 17 | 12 | 36 | 91 | 3.41E-05 | 0.000557544 |
| ENSGALG00010026421 | NA | NA | 52 | 30 | 5 | 10 | 6 | 5 | 31 | 37 | 9.97E-06 | 0.000195248 |
| ENSGALG00010026422 | P2RX3 | P2RX3 | 276 | 316 | 113 | 125 | 139 | 116 | 213 | 243 | 0.00158887 | 0.013485624 |
| ENSGALG00010026424 | NA | NA | 3 | 1 | 6 | 6 | 4 | 1 | 26 | 21 | 3.49E-05 | 0.000567954 |
| ENSGALG00010026436 | PLCD1 | PLCD1 | 467 | 253 | 113 | 158 | 274 | 300 | 401 | 315 | 0.001619134 | 0.01370768 |
| ENSGALG00010026484 | TNKS1BP1 | TNKS1BP1 | 4675 | 4941 | 3539 | 3995 | 3148 | 2207 | 3828 | 3946 | 0.007519067 | 0.04641989 |
| ENSGALG00010026546 | NA | NA | 20 | 14 | 0 | 3 | 4 | 8 | 13 | 6 | 0.005785354 | 0.037752764 |
| ENSGALG00010026550 | MYD88 | MYD88 | 219 | 184 | 338 | 331 | 461 | 317 | 270 | 277 | 0.008646088 | 0.051938687 |
| ENSGALG00010026553 | PLPP3 | PPAP2B | 446 | 618 | 203 | 272 | 181 | 148 | 348 | 446 | 6.86E-05 | 0.001000429 |
| ENSGALG00010026591 | NA | NA | 12 | 15 | 3 | 8 | 1 | 3 | 33 | 10 | 0.004861219 | 0.033064998 |
| ENSGALG00010026597 | NA | NA | 129 | 129 | 36 | 66 | 55 | 46 | 120 | 110 | 0.004894525 | 0.03326446 |
| ENSGALG00010026618 | NA | NA | 4 | 18 | 31 | 23 | 22 | 12 | 3 | 5 | 0.006951413 | 0.043445443 |
| ENSGALG00010026626 | LOC101751123 | SPX2 | 15 | 13 | 1 | 1 | 1 | 6 | 8 | 2 | 0.009192446 | 0.054589905 |
| ENSGALG00010026681 | PLXDC1 | PLXDC1 | 217 | 277 | 168 | 209 | 179 | 209 | 431 | 478 | 0.003081582 | 0.022949825 |
| ENSGALG00010026689 | NA | NA | 277 | 191 | 46 | 63 | 102 | 145 | 322 | 161 | 0.000108938 | 0.001481315 |
| ENSGALG00010026697 | CHN1 | N-chimaerin | 1965 | 2421 | 3131 | 4627 | 2267 | 1395 | 1572 | 1652 | 1.71E-06 | 4.23E-05 |
| ENSGALG00010026704 | WDFY1 | WDFY1 | 1218 | 1279 | 1804 | 2432 | 1553 | 1278 | 1352 | 1159 | 0.00637791 | 0.04071338 |
| ENSGALG00010026706 | KRT24 | GK-19 | 5 | 9 | 67 | 83 | 9 | 36 | 74 | 188 | 5.91E-07 | 1.64E-05 |
| ENSGALG00010026717 | NA | NA | 48 | 27 | 10 | 10 | 43 | 53 | 99 | 45 | 0.000138713 | 0.001822465 |
| ENSGALG00010026731 | LOC420039 | KRTC42L | 1 | 5 | 4 | 5 | 9 | 19 | 33 | 37 | 1.08E-05 | 0.000208427 |
| ENSGALG00010026737 | SCN11A | SCN11A | 1 | 0 | 26 | 6 | 22 | 9 | 8 | 3 | 0.000525955 | 0.005535118 |
| ENSGALG00010026769 | NA | NA | 4 | 5 | 1 | 1 | 8 | 17 | 9 | 13 | 0.00130506 | 0.011545876 |
| ENSGALG00010026770 | SCN1A | SCN1A | 14 | 31 | 62 | 59 | 87 | 60 | 61 | 33 | 0.006850341 | 0.04302251 |
| ENSGALG00010026786 | NA | NA | 13 | 8 | 1 | 2 | 2 | 5 | 9 | 25 | 0.003633028 | 0.026204217 |
| ENSGALG00010026788 | NA | NA | 8 | 15 | 30 | 29 | 32 | 33 | 14 | 5 | 0.003065652 | 0.022851548 |
| ENSGALG00010026810 | NA | NA | 47 | 25 | 3 | 3 | 11 | 15 | 28 | 13 | 0.000170887 | 0.002190078 |
| ENSGALG00010026821 | NA | NA | 2 | 4 | 25 | 22 | 13 | 14 | 8 | 20 | 0.005757327 | 0.037628645 |
| ENSGALG00010026825 | ADAM23 | ADAM23 | 1243 | 1406 | 963 | 915 | 1224 | 955 | 1913 | 1407 | 0.007175547 | 0.044610792 |
| ENSGALG00010026844 | NA | NA | 0 | 3 | 2 | 0 | 5 | 8 | 20 | 7 | 0.002718649 | 0.020736678 |
| ENSGALG00010026909 | CYP2J22 | CYP2J2L2 | 115 | 153 | 21 | 52 | 19 | 35 | 52 | 55 | 0.000340665 | 0.00387726 |
| ENSGALG00010026949 | NA | NA | 848 | 1091 | 1609 | 1902 | 1136 | 742 | 774 | 664 | 1.67E-05 | 0.000302259 |
| ENSGALG00010026960 | SEC61A1 | SEC61A1 | 17755 | 13636 | 7232 | 9966 | 10550 | 7534 | 12722 | 13999 | 0.000116802 | 0.001572917 |
| ENSGALG00010026987 | NA | NA | 108 | 112 | 333 | 298 | 288 | 220 | 345 | 338 | 0.000136041 | 0.001791578 |
| ENSGALG00010026999 | MLEC | MLEC | 7124 | 5638 | 4372 | 5295 | 4359 | 3924 | 7817 | 7958 | 0.000745732 | 0.007383105 |
| ENSGALG00010027010 | NA | NA | 13 | 18 | 56 | 72 | 45 | 42 | 38 | 59 | 0.004012755 | 0.028403712 |
| ENSGALG00010027036 | NUCKS1 | JC7 | 4252 | 4705 | 6685 | 8125 | 5629 | 4616 | 4725 | 4461 | 0.000491815 | 0.005208629 |
| ENSGALG00010027044 | NA | NA | 82 | 92 | 146 | 186 | 232 | 163 | 133 | 80 | 0.005400321 | 0.035827776 |
| ENSGALG00010027046 | HESX1 | GANF | 265 | 467 | 397 | 491 | 188 | 232 | 95 | 157 | 0.00037973 | 0.004256235 |
| ENSGALG00010027050 | GSTT1 | GSTT1 | 428 | 200 | 92 | 129 | 165 | 163 | 311 | 207 | 0.005922028 | 0.038391696 |
| ENSGALG00010027079 | PM20D1 | PM20D1 | 177 | 291 | 648 | 601 | 544 | 358 | 463 | 456 | 0.000763644 | 0.007507074 |
| ENSGALG00010027085 | PIM1 | PIM1 | 1733 | 2260 | 905 | 1110 | 971 | 980 | 1624 | 1599 | 9.86E-05 | 0.001362108 |
| ENSGALG00010027103 | ACVR2B | ACTR-IIB | 659 | 643 | 307 | 353 | 363 | 247 | 508 | 510 | 0.002809371 | 0.021273316 |
| ENSGALG00010027104 | NA | NA | 83 | 129 | 4 | 7 | 76 | 47 | 42 | 33 | 7.92E-11 | 5.01E-09 |
| ENSGALG00010027110 | IKBKE | IKBKE | 380 | 237 | 100 | 84 | 75 | 67 | 329 | 159 | 2.34E-05 | 0.000403592 |
| ENSGALG00010027117 | RASSF5 | RASSF5 | 743 | 940 | 21 | 19 | 47 | 74 | 1337 | 473 | 2.90E-29 | 1.13E-26 |
| ENSGALG00010027121 | ELF3 | ELF3 | 8 | 7 | 1 | 2 | 6 | 3 | 19 | 22 | 0.000184403 | 0.002331963 |
| ENSGALG00010027125 | CRYGN | CRYGN | 47 | 59 | 56 | 34 | 16 | 9 | 22 | 20 | 0.004219465 | 0.029566328 |
| ENSGALG00010027153 | ACTR3B | ACTR3B | 3004 | 3151 | 5189 | 6019 | 4679 | 3044 | 2956 | 3228 | 7.77E-06 | 0.000157275 |
| ENSGALG00010027154 | PLPP2 | PPAP2C | 992 | 678 | 430 | 325 | 380 | 381 | 726 | 856 | 0.000477405 | 0.00508897 |
| ENSGALG00010027157 | CD55 | CD55 | 48 | 46 | 19 | 20 | 36 | 24 | 71 | 64 | 0.004791561 | 0.032777967 |
| ENSGALG00010027161 | C4BPG | CR1L | 222 | 188 | 93 | 64 | 91 | 56 | 189 | 341 | 9.50E-05 | 0.001329195 |
| ENSGALG00010027167 | ASIC1L | ASIC3 | 186 | 503 | 104 | 49 | 31 | 20 | 127 | 89 | 8.18E-09 | 3.56E-07 |
| ENSGALG00010027169 | ABCB8 | ABCB8 | 1549 | 2021 | 1180 | 1250 | 905 | 603 | 1066 | 1359 | 0.000661823 | 0.006707325 |
| ENSGALG00010027171 | LOC107055991 | ATG9B | 490 | 597 | 157 | 283 | 149 | 151 | 363 | 417 | 2.47E-05 | 0.000423518 |
| ENSGALG00010027183 | CD34 | CD34 | 27 | 7 | 0 | 0 | 1 | 3 | 28 | 33 | 2.38E-08 | 9.22E-07 |
| ENSGALG00010027186 | PLXNA2 | plexin-A2 | 138 | 78 | 105 | 41 | 78 | 37 | 384 | 306 | 4.23E-06 | 9.32E-05 |
| ENSGALG00010027188 | KCNH2 | KCNH2 | 321 | 712 | 62 | 51 | 90 | 38 | 116 | 138 | 1.60E-13 | 1.49E-11 |
| ENSGALG00010027193 | CAMK1G | CAMK1G | 14 | 11 | 30 | 12 | 9 | 9 | 65 | 51 | 0.000210994 | 0.002616468 |
| ENSGALG00010027194 | NA | NA | 157 | 323 | 122 | 70 | 64 | 49 | 157 | 161 | 0.00098593 | 0.009230731 |
| ENSGALG00010027196 | LOC112529948 | LOC112529948 | 31 | 95 | 54 | 30 | 5 | 1 | 5 | 34 | 0.000144745 | 0.001891303 |
| ENSGALG00010027210 | GIMAP8 | GIMAP8 | 238 | 586 | 144 | 164 | 85 | 61 | 274 | 361 | 2.01E-06 | 4.87E-05 |
| ENSGALG00010027212 | IRF6 | IRF6 | 542 | 226 | 119 | 78 | 169 | 166 | 585 | 506 | 1.01E-06 | 2.62E-05 |
| ENSGALG00010027213 | KCNC4 | KCNC4 | 664 | 958 | 234 | 243 | 171 | 143 | 501 | 485 | 2.67E-11 | 1.87E-09 |
| ENSGALG00010027214 | PLIN4 | PLIN4 | 5681 | 7580 | 2078 | 2356 | 3035 | 2167 | 2401 | 4158 | 4.91E-11 | 3.24E-09 |
| ENSGALG00010027215 | NA | NA | 947 | 937 | 432 | 627 | 446 | 374 | 651 | 751 | 0.001854215 | 0.015215724 |
| ENSGALG00010027216 | LRG1 | LRG1 | 83 | 105 | 15 | 5 | 22 | 9 | 37 | 52 | 2.75E-07 | 8.32E-06 |
| ENSGALG00010027217 | SEMA6B | semaphorin-6B | 3217 | 3917 | 927 | 900 | 1819 | 1036 | 1705 | 1685 | 2.54E-16 | 3.48E-14 |
| ENSGALG00010027228 | SSPO | SSPO | 148 | 242 | 73 | 93 | 62 | 56 | 90 | 99 | 0.002345373 | 0.018410463 |
| ENSGALG00010027233 | NA | NA | 521 | 895 | 987 | 1307 | 525 | 402 | 372 | 459 | 4.83E-05 | 0.000748011 |
| ENSGALG00010027235 | NA | NA | 34 | 78 | 205 | 239 | 54 | 109 | 54 | 65 | 0.000551367 | 0.005769846 |
| ENSGALG00010027275 | CPAMD8 | CPAMD8 | 662 | 340 | 93 | 75 | 141 | 217 | 715 | 296 | 1.20E-06 | 3.08E-05 |
| ENSGALG00010027277 | KCND3 | KCND1 | 21 | 137 | 472 | 782 | 18 | 11 | 13 | 5 | 1.24E-20 | 2.63E-18 |
| ENSGALG00010027278 | H1FX | H1FX | 24322 | 16690 | 11047 | 11826 | 11437 | 7354 | 14201 | 15486 | 5.47E-05 | 0.000827234 |

| | | | | | | | | | | | | |
|---|---|---|---|---|---|---|---|---|---|---|---|---|
| ENSGALG00010027279 | F2RL3 | F2RL3 | 71 | 55 | 15 | 7 | 24 | 21 | 49 | 41 | 0.000204637 | 0.002548362 |
| ENSGALG00010027290 | CSPG5 | CALEB | 1007 | 1726 | 598 | 825 | 496 | 349 | 666 | 706 | 2.78E-06 | 6.46E-05 |
| ENSGALG00010027297 | LOC121107553 | LOC121107553 | 128 | 252 | 35 | 75 | 1 | 1 | 65 | 62 | 8.10E-17 | 1.15E-14 |
| ENSGALG00010027311 | NFASC | NFASC | 510 | 458 | 132 | 90 | 399 | 426 | 1228 | 1039 | 3.62E-18 | 6.18E-16 |
| ENSGALG00010027322 | NA | NA | 317 | 1220 | 450 | 266 | 508 | 351 | 882 | 1829 | 0.000682574 | 0.006864656 |
| ENSGALG00010027326 | MRPL24 | MRPL24 | 6 | 1 | 22 | 36 | 1 | 5 | 3 | 5 | 0.000215068 | 0.002652553 |
| ENSGALG00010027340 | IL12RB1 | IL12RB1 | 38 | 122 | 16 | 24 | 31 | 10 | 41 | 30 | 0.004423538 | 0.030649326 |
| ENSGALG00010027344 | NA | NA | 18 | 19 | 2 | 1 | 9 | 13 | 12 | 14 | 0.000656144 | 0.006665942 |
| ENSGALG00010027356 | NA | NA | 1209 | 1225 | 1612 | 1269 | 864 | 528 | 955 | 761 | 0.009293243 | 0.055034964 |
| ENSGALG00010027357 | GHRHR-LR | GHRH-LR | 407 | 175 | 13 | 19 | 93 | 219 | 448 | 218 | 1.34E-09 | 6.80E-08 |
| ENSGALG00010027363 | VIPR1 | CVIPR | 55 | 71 | 33 | 31 | 38 | 44 | 120 | 84 | 0.008075069 | 0.049216103 |
| ENSGALG00010027372 | PIK3C2B | PIK3C2B | 1439 | 1602 | 776 | 1035 | 946 | 808 | 1176 | 1162 | 0.006685235 | 0.042143966 |
| ENSGALG00010027377 | Pou5f3 | POUV | 110158 | 102696 | 58010 | 66815 | 63713 | 50589 | 81354 | 71696 | 0.000792537 | 0.007749149 |
| ENSGALG00010027383 | ASS1 | ASS1 | 366 | 674 | 133 | 106 | 74 | 82 | 428 | 539 | 3.12E-12 | 2.53E-10 |
| ENSGALG00010027403 | ACKR2 | CCBP2 | 29 | 15 | 61 | 80 | 61 | 39 | 24 | 33 | 0.009162925 | 0.054436829 |
| ENSGALG00010027412 | LOC101750583 | LOC101750583 | 145 | 207 | 246 | 321 | 258 | 243 | 67 | 80 | 4.38E-06 | 9.63E-05 |
| ENSGALG00010027421 | PLPPR3 | LPPR3 | 990 | 1415 | 442 | 529 | 424 | 333 | 641 | 849 | 2.57E-07 | 7.86E-06 |
| ENSGALG00010027425 | CFD | CFD | 654 | 418 | 88 | 73 | 76 | 148 | 393 | 379 | 3.41E-09 | 1.61E-07 |
| ENSGALG00010027440 | TWF2 | PTK9L | 2838 | 2866 | 1772 | 1989 | 2229 | 1656 | 2517 | 2914 | 0.006217906 | 0.039859472 |
| ENSGALG00010027457 | CAPZA1 | CAPZA1 | 2613 | 3729 | 4688 | 6530 | 4521 | 3204 | 3754 | 3989 | 0.003356026 | 0.024534517 |
| ENSGALG00010027468 | BSG | Bsg1 | 12945 | 16196 | 6460 | 10270 | 7717 | 7880 | 10267 | 12417 | 0.001260488 | 0.011240813 |
| ENSGALG00010027479 | HCN2 | HCN2 | 126 | 197 | 43 | 67 | 27 | 41 | 164 | 141 | 3.05E-05 | 0.000507972 |
| ENSGALG00010027492 | PCP4L1 | PCP4L1 | 39 | 37 | 5 | 4 | 31 | 23 | 32 | 49 | 1.04E-05 | 0.000202914 |
| ENSGALG00010027510 | RHOC | ARHC | 111 | 163 | 108 | 62 | 96 | 60 | 220 | 270 | 0.002152936 | 0.017239991 |
| ENSGALG00010027513 | SEMA3G | SEMA3G | 147 | 280 | 105 | 95 | 758 | 562 | 153 | 300 | 2.63E-12 | 2.14E-10 |
| ENSGALG00010027520 | FANCE | FANCE | 428 | 470 | 949 | 948 | 568 | 431 | 548 | 645 | 0.004302878 | 0.030012394 |
| ENSGALG00010027525 | CUTAL | CUTA | 25 | 39 | 224 | 132 | 111 | 38 | 46 | 103 | 0.000761734 | 0.007497111 |
| ENSGALG00010027530 | PHF19 | PHF19 | 296 | 418 | 1039 | 984 | 623 | 416 | 686 | 698 | 2.67E-05 | 0.000452589 |
| ENSGALG00010027533 | WDR18 | WDR18 | 4187 | 4304 | 2594 | 3197 | 2844 | 2284 | 3366 | 3590 | 0.007631635 | 0.046976014 |
| ENSGALG00010027538 | NA | NA | 425 | 394 | 690 | 811 | 606 | 465 | 332 | 361 | 0.001127216 | 0.010271553 |
| ENSGALG00010027557 | C6orf132 | C26H6ORF132 | 352 | 138 | 115 | 139 | 98 | 164 | 414 | 406 | 0.003765979 | 0.026965315 |
| ENSGALG00010027563 | WDR77 | WDR77 | 10944 | 11335 | 14069 | 17636 | 10994 | 8667 | 9123 | 9876 | 0.004995411 | 0.03372803 |
| ENSGALG00010027583 | GLE1 | GLE1L | 2134 | 2565 | 3184 | 3965 | 2099 | 1443 | 2774 | 2989 | 0.004190312 | 0.029423736 |
| ENSGALG00010027590 | SEMA3F | semaphorin-3F | 4200 | 4328 | 2042 | 1592 | 2189 | 1321 | 3079 | 3515 | 6.42E-09 | 2.88E-07 |
| ENSGALG00010027622 | FCN3 | FIBCD1 | 376 | 682 | 69 | 84 | 98 | 147 | 346 | 219 | 8.58E-09 | 3.72E-07 |
| ENSGALG00010027627 | OARD1 | C26H6ORF130 | 4717 | 5676 | 3633 | 5183 | 5719 | 4851 | 3250 | 3378 | 1.47E-05 | 0.000272475 |
| ENSGALG00010027635 | CD48 | CD48 | 15 | 13 | 0 | 0 | 1 | 7 | 8 | 1 | 0.0007922 | 0.007749149 |
| ENSGALG00010027649 | FBLN2 | fibulin-2 | 3417 | 1977 | 814 | 645 | 1305 | 1332 | 3584 | 1953 | 3.31E-08 | 1.23E-06 |
| ENSGALG00010027699 | VANGL2 | VANGL2 | 4027 | 5628 | 3334 | 3039 | 2969 | 2061 | 4795 | 5105 | 3.97E-05 | 0.000632875 |
| ENSGALG00010027701 | NHLH1 | NHLH1 | 248 | 201 | 86 | 85 | 71 | 63 | 115 | 144 | 0.000715609 | 0.007135592 |
| ENSGALG00010027707 | HIGD1C | HIGD1A | 994 | 1638 | 1222 | 1565 | 785 | 512 | 653 | 833 | 0.000700227 | 0.007007015 |
| ENSGALG00010027708 | LOC101750889 | LOC101750889 | 25 | 41 | 15 | 17 | 63 | 34 | 76 | 44 | 0.003309237 | 0.024341537 |
| ENSGALG00010027711 | WNK2 | WNK2 | 4950 | 5057 | 4884 | 5605 | 8188 | 4894 | 7678 | 5854 | 0.004182305 | 0.02939221 |
| ENSGALG00010027715 | NINJ1 | NINJ1 | 45 | 50 | 156 | 168 | 259 | 153 | 171 | 194 | 3.34E-06 | 7.58E-05 |
| ENSGALG00010027720 | DAB2IP | DAB2IP | 306 | 421 | 949 | 699 | 692 | 446 | 570 | 502 | 0.004209337 | 0.029520115 |
| ENSGALG00010027727 | GSTM2 | GSTM2 | 1063 | 1059 | 161 | 164 | 475 | 310 | 500 | 579 | 1.33E-16 | 1.85E-14 |
| ENSGALG00010027729 | LOC107050775 | LOC107050775 | 125 | 48 | 14 | 11 | 23 | 19 | 44 | 37 | 3.99E-05 | 0.000635082 |
| ENSGALG00010027739 | FGD3 | FGD3 | 554 | 731 | 1484 | 1267 | 1405 | 862 | 1103 | 864 | 0.000467321 | 0.005006331 |
| ENSGALG00010027743 | C1orf74 | C26H1orf74 | 106 | 97 | 184 | 199 | 223 | 148 | 80 | 82 | 0.001453581 | 0.01255954 |
| ENSGALG00010027751 | ANGPTL4 | ANGPTL4 | 1360 | 1917 | 430 | 496 | 587 | 410 | 968 | 1155 | 4.45E-13 | 3.95E-11 |
| ENSGALG00010027811 | MEX3D | RKHD1 | 2201 | 2191 | 1414 | 1577 | 1451 | 950 | 1498 | 1478 | 0.005927187 | 0.038408358 |
| ENSGALG00010027813 | LRRC71 | LRRC71 | 420 | 518 | 215 | 214 | 650 | 389 | 434 | 533 | 5.69E-05 | 0.000855258 |
| ENSGALG00010027820 | NA | NA | 60 | 66 | 297 | 248 | 259 | 94 | 148 | 107 | 0.000231238 | 0.002822832 |
| ENSGALG00010027822 | DPH7 | WDR85 | 15 | 12 | 1 | 1 | 9 | 11 | 31 | 13 | 0.0001991 | 0.00248681 |
| ENSGALG00010027829 | NA | NA | 8 | 5 | 33 | 13 | 29 | 26 | 52 | 21 | 0.008336057 | 0.050493723 |
| ENSGALG00010027830 | CACNA1D | CACNA1D | 165 | 295 | 195 | 162 | 542 | 406 | 307 | 144 | 0.0001432 | 0.001874047 |
| ENSGALG00010027831 | ATCAY | ATCAY | 144 | 189 | 145 | 126 | 413 | 382 | 65 | 80 | 8.36E-12 | 6.41E-10 |
| ENSGALG00010027833 | NA | NA | 1 | 2 | 3 | 11 | 9 | 13 | 19 | 17 | 0.002196346 | 0.017512088 |
| ENSGALG00010027835 | NA | NA | 2 | 0 | 0 | 1 | 3 | 8 | 16 | 6 | 0.002236549 | 0.017747965 |
| ENSGALG00010027860 | LOC101750908 | LOC101750908 | 87 | 111 | 154 | 164 | 229 | 212 | 195 | 228 | 0.004025012 | 0.028454312 |
| ENSGALG00010027862 | NA | NA | 692 | 742 | 311 | 397 | 347 | 304 | 523 | 463 | 0.001882448 | 0.015391132 |
| ENSGALG00010027866 | KCNA2 | KCNA2 | 27 | 70 | 25 | 10 | 28 | 11 | 67 | 81 | 0.003324002 | 0.024407172 |
| ENSGALG00010027875 | WNT5A | WNT5A | 91 | 105 | 368 | 325 | 215 | 83 | 114 | 187 | 0.001212959 | 0.010904237 |
| ENSGALG00010027908 | PBXIP1 | PBXIP1 | 636 | 724 | 408 | 348 | 368 | 242 | 481 | 649 | 0.006151959 | 0.039573341 |
| ENSGALG00010027939 | BSPRY | BSPRY | 304 | 231 | 48 | 118 | 132 | 137 | 133 | 104 | 0.001845408 | 0.015177395 |
| ENSGALG00010027945 | DCST2 | DCST2 | 151 | 197 | 734 | 1098 | 43 | 49 | 1735 | 2165 | 1.41E-55 | 1.57E-52 |
| ENSGALG00010027957 | LOC107049672 | LOC107049672 | 432 | 518 | 156 | 185 | 205 | 180 | 483 | 633 | 1.48E-07 | 4.83E-06 |
| ENSGALG00010027969 | HSPA5 | GRP78 | 32600 | 24885 | 14097 | 17562 | 13440 | 9058 | 16944 | 17122 | 2.89E-07 | 8.73E-06 |
| ENSGALG00010027976 | LRSAM1 | LRSAM1 | 612 | 553 | 157 | 207 | 431 | 289 | 434 | 574 | 5.26E-06 | 0.000112461 |
| ENSGALG00010027992 | NA | NA | 34 | 29 | 3 | 7 | 24 | 42 | 55 | 20 | 0.000489077 | 0.005186207 |
| ENSGALG00010028003 | NA | NA | 2360 | 1990 | 328 | 375 | 1873 | 1675 | 2698 | 2053 | 6.23E-27 | 2.12E-24 |
| ENSGALG00010028016 | PRF1 | P1 | 15 | 14 | 2 | 4 | 20 | 12 | 11 | 17 | 0.006849767 | 0.04302251 |
| ENSGALG00010028025 | TNNC1 | PCTNC1 | 639 | 757 | 232 | 206 | 572 | 561 | 1183 | 1513 | 2.00E-15 | 2.48E-13 |
| ENSGALG00010028030 | LOC121107545 | LOC121107545 | 22 | 14 | 2 | 1 | 8 | 6 | 8 | 27 | 0.001472378 | 0.012670643 |
| ENSGALG00010028034 | MUM1L1 | MUM1 | 1619 | 1552 | 1259 | 1238 | 1298 | 932 | 2460 | 1915 | 0.001542685 | 0.01316719 |
| ENSGALG00010028056 | SLC25A25 | SLC25A25 | 3312 | 6122 | 2776 | 3581 | 2360 | 2001 | 2691 | 3193 | 0.002574056 | 0.019787313 |
| ENSGALG00010028057 | APC2 | APC2 | 844 | 2306 | 311 | 203 | 637 | 308 | 996 | 734 | 4.78E-09 | 2.23E-07 |
| ENSGALG00010028059 | GZMM | GZMM | 51 | 109 | 22 | 21 | 40 | 23 | 51 | 38 | 0.005483792 | 0.036295102 |
| ENSGALG00010028061 | NAIF1 | C9orf90 | 668 | 721 | 1151 | 1252 | 1117 | 807 | 712 | 723 | 0.001654519 | 0.013929632 |
| ENSGALG00010028082 | AK1 | AK1 | 225 | 822 | 786 | 958 | 688 | 419 | 86 | 287 | 0.001274336 | 0.011333613 |
| ENSGALG00010028101 | NA | NA | 1731 | 1758 | 1420 | 1702 | 1007 | 700 | 956 | 1236 | 0.002768222 | 0.02100932 |
| ENSGALG00010028105 | PRKCD | PRKCD | 1269 | 1182 | 706 | 697 | 1128 | 960 | 1078 | 828 | 0.001239091 | 0.011097372 |
| ENSGALG00010028112 | NA | NA | 7632 | 8587 | 4364 | 5256 | 4640 | 4381 | 6271 | 7510 | 0.000148718 | 0.001937158 |
| ENSGALG00010028132 | CACNA2D2 | CACNA2D2 | 620 | 1008 | 1090 | 1098 | 1038 | 824 | 1514 | 1708 | 0.003127566 | 0.023230168 |
| ENSGALG00010028138 | LOC101747452 | LOC101747452 | 27 | 27 | 13 | 14 | 99 | 43 | 13 | 7 | 1.39E-07 | 4.59E-06 |
| ENSGALG00010028141 | CISH | CIS | 47 | 68 | 170 | 243 | 125 | 123 | 166 | 214 | 0.000631974 | 0.0064439 |
| ENSGALG00010028146 | MAPKAPK3 | MAPKAPK3 | 2312 | 1921 | 2360 | 2777 | 1185 | 808 | 1404 | 1333 | 4.80E-06 | 0.00010439 |
| ENSGALG00010028163 | GBP | GBP | 18 | 10 | 15 | 10 | 19 | 21 | 54 | 44 | 0.002551976 | 0.019662807 |
| ENSGALG00010028179 | MCM2 | MCM2 | 3731 | 4713 | 6421 | 7258 | 4460 | 3604 | 3886 | 4611 | 0.002211215 | 0.017597056 |
| ENSGALG00010028181 | BGLAP | BGP | 491 | 378 | 84 | 160 | 448 | 300 | 235 | 410 | 2.18E-05 | 0.000379436 |
| ENSGALG00010028193 | BRINP1 | BRINP | 316 | 633 | 545 | 1140 | 317 | 167 | 400 | 296 | 0.001424066 | 0.012356691 |
| ENSGALG00010028194 | LOC107055115 | LOC107055115 | 455 | 915 | 437 | 322 | 289 | 221 | 274 | 286 | 0.001384388 | 0.012113036 |
| ENSGALG00010028197 | INTS3 | INTS3 | 6405 | 7169 | 4624 | 5125 | 4527 | 3419 | 4981 | 5330 | 0.005073362 | 0.034050611 |
| ENSGALG00010028200 | NA | NA | 5 | 0 | 0 | 0 | 11 | 10 | 0 | 0 | 2.17E-05 | 0.000377816 |
| ENSGALG00010028203 | LHX3 | LIM3 | 63 | 24 | 1 | 2 | 1 | 4 | 23 | 17 | 3.18E-08 | 1.20E-06 |
| ENSGALG00010028212 | NA | NA | 4 | 0 | 0 | 0 | 12 | 8 | 6 | 0 | 0.003456737 | 0.025160709 |
| ENSGALG00010028213 | NA | NA | 146 | 111 | 19 | 25 | 83 | 85 | 174 | 81 | 1.14E-05 | 0.000219758 |
| ENSGALG00010028217 | NA | NA | 6 | 11 | 14 | 20 | 10 | 11 | 2 | 1 | 0.002444693 | 0.019038077 |
| ENSGALG00010028228 | LOC107054353 | LOC107054353 | 118 | 186 | 630 | 640 | 168 | 169 | 189 | 448 | 1.62E-05 | 0.000295648 |
| ENSGALG00010028230 | NA | NA | 7 | 11 | 72 | 101 | 12 | 12 | 11 | 9 | 2.55E-10 | 1.46E-08 |
| ENSGALG00010028240 | NA | NA | 10 | 83 | 16 | 19 | 161 | 183 | 52 | 61 | 1.37E-06 | 3.46E-05 |
| ENSGALG00010028278 | NA | NA | 9 | 10 | 17 | 12 | 3 | 5 | 63 | 36 | 5.13E-06 | 0.00011022 |
| ENSGALG00010028283 | CERS4 | LASS4 | 907 | 966 | 1798 | 1780 | 1438 | 933 | 1345 | 1287 | 0.005563636 | 0.036709525 |
| ENSGALG00010028287 | NA | NA | 2 | 0 | 0 | 0 | 1 | 0 | 19 | 11 | 1.48E-06 | 3.73E-05 |
| ENSGALG00010028301 | FBN3 | fibrillin-3 | 1393 | 2170 | 2730 | 3235 | 3488 | 2524 | 2885 | 3003 | 0.000128664 | 0.001707891 |
| ENSGALG00010028310 | NA | NA | 71 | 60 | 796 | 699 | 849 | 148 | 103 | 200 | 1.90E-09 | 9.46E-08 |
| ENSGALG00010028311 | CTXN1 | CTXN1 | 155 | 168 | 54 | 65 | 93 | 74 | 91 | 64 | 0.004640423 | 0.031927029 |
| ENSGALG00010028319 | NA | NA | 20 | 19 | 62 | 56 | 98 | 31 | 42 | 25 | 0.008960633 | 0.053501286 |
| ENSGALG00010028325 | NA | NA | 0 | 3 | 6 | 12 | 0 | 0 | 1 | 12 | 0.007142788 | 0.044458204 |
| ENSGALG00010028331 | NA | NA | 153 | 11421 | 99 | 103 | 138 | 91 | 152 | 5104 | 1.16E-06 | 2.97E-05 |
| ENSGALG00010028335 | NA | NA | 0 | 0 | 22 | 11 | 17 | 13 | 6 | 4 | 1.80E-06 | 4.43E-05 |
| ENSGALG00010028347 | NA | NA | 15 | 18 | 18 | 19 | 13 | 9 | 36 | 68 | 0.001781245 | 0.01478062 |
| ENSGALG00010028379 | HAPLN4 | HAPLN4 | 11 | 52 | 1 | 3 | 8 | 3 | 10 | 13 | 0.000239484 | 0.002897061 |
| ENSGALG00010028386 | NCAN | CSPG3 | 590 | 3797 | 597 | 1013 | 666 | 414 | 582 | 842 | 0.00111967 | 0.010225099 |
| ENSGALG00010028431 | S100A14 | S100A14 | 156 | 127 | 12 | 56 | 19 | 25 | 75 | 95 | 9.83E-05 | 0.001360314 |
| ENSGALG00010028441 | NA | NA | 24 | 43 | 8 | 14 | 1 | 1 | 5 | 6 | 7.92E-07 | 2.11E-05 |
| ENSGALG00010028445 | SYT11 | SYT11 | 5450 | 5702 | 3520 | 4199 | 3317 | 2815 | 5451 | 4464 | 0.001639826 | 0.013847788 |
| ENSGALG00010028447 | OLFM1 | NOEL | 230 | 893 | 466 | 444 | 260 | 154 | 713 | 1010 | 0.001854621 | 0.015215724 |
| ENSGALG00010028460 | CRLF1 | CRLF1 | 1440 | 1444 | 242 | 385 | 539 | 471 | 611 | 941 | 9.43E-12 | 7.20E-10 |
| ENSGALG00010028473 | CDC34 | CDC34 | 2360 | 2961 | 1398 | 1882 | 1579 | 1297 | 1787 | 2055 | 0.003738441 | 0.026791092 |
| ENSGALG00010028480 | NA | NA | 1880 | 1250 | 428 | 441 | 592 | 677 | 1307 | 1090 | 5.58E-09 | 2.54E-07 |
| ENSGALG00010028483 | EXD3 | EXD3 | 249 | 471 | 767 | 715 | 883 | 525 | 304 | 319 | 1.26E-05 | 0.000239999 |
| ENSGALG00010028487 | ENTPD8 | ENTPD1 | 18 | 18 | 79 | 54 | 79 | 47 | 31 | 30 | 0.000549176 | 0.00575052 |
| ENSGALG00010028497 | ADMP | ADMP | 3628 | 2953 | 690 | 516 | 1574 | 1040 | 2818 | 2679 | 1.56E-21 | 3.72E-19 |
| ENSGALG00010028501 | USP20 | USP20 | 639 | 714 | 1538 | 1385 | 1157 | 751 | 945 | 761 | 0.00421981 | 0.004612265 |
| ENSGALG00010028510 | CILP2 | CILP2 | 68 | 88 | 12 | 22 | 36 | 32 | 75 | 68 | 0.000140894 | 0.001846756 |
| ENSGALG00010028517 | RNF208 | RNF208 | 1407 | 1606 | 1118 | 1172 | 993 | 846 | 1722 | 2179 | 0.000992616 | 0.009282933 |
| ENSGALG00010028525 | TM6SF2 | TM6SF2 | 8 | 113 | 5 | 10 | 3 | 2 | 15 | 16 | 4.50E-05 | 0.00070661 |
| ENSGALG00010028540 | LOC427775 | LOC427775 | 11 | 10 | 2 | 1 | 6 | 1 | 13 | 27 | 0.000535724 | 0.00562376 |
| ENSGALG00010028552 | LOC769139 | LOC769139 | 210 | 76 | 853 | 549 | 1078 | 306 | 211 | 72 | 2.52E-06 | 5.91E-05 |
| ENSGALG00010028554 | LOC100858797 | LOC100858797 | 15 | 15 | 82 | 80 | 111 | 33 | 23 | 11 | 5.91E-06 | 0.000124325 |

| | | | | | | | | | | | | | |
|---|---|---|---|---|---|---|---|---|---|---|---|---|---|
| ENSGALG00010028574 | ENDOG | ENDOG | | 249 | 558 | 20 | 21 | 464 | 343 | 38 | 28 | 2.71E-28 | 1.01E-25 |
| ENSGALG00010028582 | KYAT1 | CCBL1 | | 45 | 73 | 19 | 23 | 71 | 63 | 39 | 30 | 0.001021225 | 0.009513221 |
| ENSGALG00010028612 | DIRAS1 | DIRAS1 | | 603 | 651 | 152 | 198 | 271 | 167 | 362 | 421 | 9.78E-08 | 3.35E-06 |
| ENSGALG00010028649 | LOC101751615 | PRARP1 | | 347 | 112 | 34 | 57 | 107 | 76 | 188 | 218 | 0.000109025 | 0.001481315 |
| ENSGALG00010028676 | LOC112530443 | LOC112530443 | | 122 | 74 | 268 | 268 | 228 | 217 | 271 | 480 | 0.000423696 | 0.004627982 |
| ENSGALG00010028677 | LOC107055358 | LOC107055358 | | 566 | 457 | 240 | 257 | 281 | 199 | 292 | 311 | 0.004727945 | 0.032435721 |
| ENSGALG00010028678 | NA | NA | | 44 | 112 | 219 | 166 | 152 | 93 | 31 | 53 | 0.000330995 | 0.003788754 |
| ENSGALG00010028692 | RALGPS1 | RALGPS1 | | 314 | 339 | 862 | 663 | 643 | 401 | 676 | 426 | 0.008008384 | 0.048934592 |
| ENSGALG00010028698 | LOC777082 | LOC777082 | | 349 | 190 | 40 | 82 | 88 | 44 | 147 | 144 | 2.00E-05 | 0.000351789 |
| ENSGALG00010028711 | NA | NA | | 139 | 217 | 58 | 31 | 109 | 100 | 248 | 102 | 0.001242352 | 0.011108711 |
| ENSGALG00010028732 | NA | NA | | 119 | 80 | 77 | 104 | 122 | 86 | 46 | 18 | 0.003500869 | 0.02542896 |
| ENSGALG00010028736 | NTRK1 | CTRKA | | 74 | 113 | 52 | 50 | 28 | 20 | 53 | 41 | 0.007558856 | 0.046620702 |
| ENSGALG00010028756 | LOC107055126 | EDMTF3 | | 0 | 0 | 85 | 102 | 96 | 64 | 51 | 50 | 3.45E-18 | 5.94E-16 |
| ENSGALG00010028757 | MAT2A | EDMTF1 | | 0 | 0 | 12 | 16 | 13 | 5 | 10 | 1 | 0.000336484 | 0.00384106 |
| ENSGALG00010028760 | SSR2 | SSR2 | | 6052 | 4009 | 2842 | 3822 | 3064 | 2269 | 3395 | 4259 | 0.009712116 | 0.056945057 |
| ENSGALG00010028763 | CTSK | CTSK | | 78 | 82 | 37 | 23 | 37 | 29 | 78 | 112 | 0.002300404 | 0.018142688 |
| ENSGALG00010028782 | MLLT11 | MLLT11 | | 43 | 86 | 130 | 143 | 104 | 58 | 25 | 34 | 0.000408839 | 0.004500999 |
| ENSGALG00010028809 | NA | NA | | 10 | 13 | 60 | 83 | 74 | 35 | 38 | 49 | 9.19E-05 | 0.001292769 |
| ENSGALG00010028815 | NA | NA | | 464 | 798 | 80 | 104 | 49 | 38 | 159 | 187 | 4.49E-20 | 9.26E-18 |
| ENSGALG00010028849 | CCDC180 | BDAG1 | | 7 | 14 | 3 | 1 | 6 | 3 | 27 | 15 | 0.001106408 | 0.010153925 |
| ENSGALG00010028866 | HORMAD1 | HORMAD1 | | 339 | 326 | 853 | 872 | 807 | 468 | 513 | 517 | 6.91E-05 | 0.00100686 |
| ENSGALG00010028870 | CACFD1 | C17H9ORF7 | | 704 | 1127 | 228 | 516 | 600 | 498 | 611 | 805 | 0.002318269 | 0.018223379 |
| ENSGALG00010028877 | FAM163B | FAM163B | | 42 | 111 | 209 | 344 | 138 | 109 | 99 | 106 | 0.001456346 | 0.012565301 |
| ENSGALG00010028880 | LOC426218 | LOC426218 | | 1 | 0 | 18 | 17 | 7 | 5 | 7 | 5 | 0.000150956 | 0.001963246 |
| ENSGALG00010028890 | LOC101747382 | COL2A1L | | 248 | 434 | 36 | 58 | 66 | 195 | 974 | 815 | 6.45E-14 | 6.45E-12 |
| ENSGALG00010028905 | LOC425607 | LOC425607 | | 1578 | 1111 | 1767 | 2102 | 1409 | 897 | 759 | 1000 | 0.002189815 | 0.017476693 |
| ENSGALG00010028906 | NA | NA | | 2 | 2 | 11 | 26 | 22 | 15 | 4 | 4 | 6.63E-05 | 0.000970466 |
| ENSGALG00010028917 | NR6A1 | NR6A1 | | 5793 | 8312 | 3148 | 3949 | 2980 | 3106 | 6165 | 4951 | 1.51E-06 | 3.80E-05 |
| ENSGALG00010028944 | NA | NA | | 1 | 0 | 104 | 0 | 81 | 0 | 0 | 0 | 8.33E-05 | 0.001186831 |
| ENSGALG00010028966 | NA | NA | | 10 | 3 | 1 | 0 | 2 | 4 | 15 | 11 | 0.002562037 | 0.019722135 |
| ENSGALG00010028993 | NA | NA | | 1 | 0 | 8 | 2 | 4 | 11 | 9 | 16 | 0.006583555 | 0.041754977 |
| ENSGALG00010029047 | NA | NA | | 1 | 0 | 10 | 16 | 12 | 6 | 10 | 4 | 0.001829698 | 0.015085308 |
| ENSGALG00010029106 | NA | NA | | 26 | 12 | 63 | 71 | 6 | 4 | 23 | 26 | 4.23E-06 | 9.32E-05 |
| ENSGALG00010029162 | ELN | ELN | | 5 | 12 | 25 | 20 | 23 | 15 | 52 | 82 | 1.91E-05 | 0.000340075 |
| ENSGALG00010029186 | ZNF302 | ZNF184 | | 181 | 218 | 61 | 39 | 41 | 28 | 255 | 226 | 1.93E-10 | 1.13E-08 |
| ENSGALG00010029190 | DHRS7C | DHRS7C | | 7 | 0 | 34 | 27 | 17 | 9 | 5 | 3 | 0.000333253 | 0.003809389 |
| ENSGALG00010029191 | GSG1L2 | GSG1-like protein | | 11 | 7 | 109 | 59 | 79 | 45 | 30 | 36 | 2.08E-06 | 5.01E-05 |
| ENSGALG00010029200 | GAS7 | GAS7 | | 326 | 398 | 479 | 415 | 434 | 557 | 1364 | 535 | 0.004199224 | 0.029461558 |
| ENSGALG00010029208 | CACNG5 | CACNG7 | | 1039 | 1526 | 525 | 886 | 639 | 482 | 1083 | 1042 | 0.000862023 | 0.008288916 |
| ENSGALG00010029214 | PROCA1 | PROCA1 | | 36 | 122 | 136 | 126 | 143 | 85 | 315 | 774 | 1.46E-07 | 4.80E-06 |
| ENSGALG00010029229 | CBX4 | CBX4 | | 59 | 104 | 433 | 226 | 225 | 88 | 178 | 290 | 0.002088874 | 0.016823738 |
| ENSGALG00010029233 | CBX2 | CBX2 | | 4335 | 5290 | 8036 | 10670 | 9126 | 7092 | 7200 | 5639 | 1.23E-06 | 3.14E-05 |
| ENSGALG00010029256 | RABEP1 | RABEP1 | | 2282 | 2986 | 3889 | 4393 | 3555 | 2378 | 2526 | 2535 | 0.002182552 | 0.017443719 |
| ENSGALG00010029268 | PCTP | PCTP | | 532 | 548 | 1184 | 1278 | 1023 | 563 | 593 | 584 | 3.82E-05 | 0.000614 |
| ENSGALG00010029275 | PCYT2 | SIRT7 | | 2901 | 3305 | 4118 | 5122 | 3311 | 2117 | 2033 | 2340 | 2.25E-05 | 0.000390055 |
| ENSGALG00010029288 | PYCR1 | PYCR2 | | 2289 | 1630 | 2323 | 2286 | 2065 | 3283 | 3867 | 4755 | 0.000791612 | 0.007749149 |
| ENSGALG00010029301 | RNF213 | RNF213 | | 167 | 281 | 98 | 113 | 192 | 333 | 234 | 286 | 0.002902349 | 0.02183884 |
| ENSGALG00010029318 | C17orf58 | C18H17orf58 | | 32 | 45 | 100 | 106 | 157 | 66 | 86 | 67 | 0.00649099 | 0.041246168 |
| ENSGALG00010029324 | BCAS3 | BCAS3 | | 2970 | 3694 | 3525 | 3992 | 3683 | 2255 | 2240 | 2315 | 0.004362059 | 0.03031132 |
| ENSGALG00010029325 | LRRC75A | C17orf76 | | 365 | 756 | 324 | 304 | 370 | 201 | 919 | 985 | 3.41E-06 | 7.74E-05 |
| ENSGALG00010029343 | SCARF1 | SCARF1 | | 56 | 26 | 1 | 12 | 4 | 4 | 20 | 27 | 0.000146764 | 0.001914693 |
| ENSGALG00010029353 | GCGR | GIPR | | 130 | 362 | 51 | 36 | 80 | 64 | 334 | 398 | 1.42E-09 | 7.20E-08 |
| ENSGALG00010029356 | LOC124417312 | LOC124417312 | | 97 | 36 | 4 | 0 | 12 | 4 | 29 | 11 | 1.32E-07 | 4.38E-06 |
| ENSGALG00010029365 | PIK3R5 | PIK3R5 | | 319 | 368 | 119 | 151 | 98 | 125 | 386 | 326 | 3.08E-05 | 0.000511938 |
| ENSGALG00010029366 | MGAT5B | MGAT5B | | 315 | 275 | 34 | 27 | 51 | 48 | 244 | 150 | 1.11E-12 | 9.47E-11 |
| ENSGALG00010029367 | DUSP14 | DUSP14 | | 1672 | 2202 | 3294 | 3965 | 2475 | 1642 | 2376 | 2602 | 0.001413972 | 0.01230523 |
| ENSGALG00010029380 | RGS9 | RGS9 | | 384 | 528 | 218 | 246 | 282 | 238 | 624 | 498 | 0.000559849 | 0.005829403 |
| ENSGALG00010029385 | CLIP2 | CLIP2 | | 1127 | 1650 | 463 | 648 | 745 | 612 | 1311 | 1236 | 4.42E-07 | 1.27E-05 |
| ENSGALG00010029392 | RTN4RL1 | RTN4RL1 | | 426 | 550 | 35 | 28 | 98 | 51 | 275 | 152 | 2.94E-17 | 4.39E-15 |
| ENSGALG00010029394 | CYGB | CYGB | | 387 | 663 | 196 | 86 | 130 | 80 | 460 | 930 | 7.62E-09 | 3.36E-07 |
| ENSGALG00010029396 | WIPI1 | WIPI1 | | 620 | 728 | 760 | 728 | 746 | 835 | 1355 | 1434 | 0.000368083 | 0.004147942 |
| ENSGALG00010029400 | AANAT | SNAT | | 6 | 11 | 9 | 9 | 19 | 10 | 33 | 37 | 0.002297429 | 0.018131972 |
| ENSGALG00010029410 | VTN | VTN | | 1832 | 2125 | 705 | 1017 | 1012 | 931 | 1982 | 2810 | 5.83E-09 | 2.64E-07 |
| ENSGALG00010029411 | SARM1 | SARM1 | | 475 | 590 | 231 | 242 | 232 | 311 | 737 | 829 | 2.91E-06 | 6.69E-05 |
| ENSGALG00010029428 | FN3K | FN3K | | 72 | 118 | 159 | 132 | 55 | 37 | 45 | 34 | 0.000703287 | 0.007025303 |
| ENSGALG00010029442 | LOC107049986 | ODF3L6 | | 8 | 11 | 64 | 78 | 33 | 20 | 51 | 76 | 7.68E-06 | 0.000155713 |
| ENSGALG00010029470 | SSTR2 | SS2R | | 177 | 262 | 73 | 53 | 63 | 51 | 221 | 175 | 5.12E-06 | 0.00011022 |
| ENSGALG00010029491 | NAA38 | LSMD1 | | 147 | 156 | 55 | 38 | 28 | 24 | 56 | 66 | 6.57E-06 | 0.000136324 |
| ENSGALG00010029492 | SDK2 | SDK2 | | 497 | 1957 | 689 | 886 | 940 | 561 | 1767 | 2389 | 0.00246883 | 0.019169112 |
| ENSGALG00010029494 | MYH7 | AMHC1 | | 165 | 102 | 51 | 32 | 96 | 211 | 365 | 368 | 6.15E-07 | 1.69E-05 |
| ENSGALG00010029502 | BTBD17 | BTBD17 | | 926 | 1682 | 44 | 128 | 424 | 590 | 819 | 755 | 1.54E-15 | 1.93E-13 |
| ENSGALG00010029503 | LOC430700 | LOC430700 | | 26 | 45 | 115 | 122 | 61 | 24 | 67 | 69 | 0.008086164 | 0.049265755 |
| ENSGALG00010029511 | CUEDC1 | CUEDC1 | | 458 | 654 | 1288 | 1358 | 1144 | 784 | 656 | 632 | 2.02E-06 | 4.89E-05 |
| ENSGALG00010029516 | LOC123575613 | LOC123575613 | | 40 | 48 | 27 | 152 | 75 | 66 | 18 | 19 | 0.005532229 | 0.036543323 |
| ENSGALG00010029524 | BAHCC1 | BAHCC1 | | 593 | 2419 | 4241 | 3674 | 3835 | 1426 | 1524 | 1574 | 0.002777371 | 0.021069186 |
| ENSGALG00010029537 | MYH1B | MYH3 | | 1 | 0 | 21 | 13 | 28 | 22 | 10 | 5 | 7.97E-07 | 2.12E-05 |
| ENSGALG00010029560 | TRIM58 | TRIM58 | | 215 | 7 | 9 | 2 | 12 | 10 | 34 | 28 | 0.000115671 | 0.001563987 |
| ENSGALG00010029574 | SGSM2 | RUTBC1 | | 435 | 466 | 1040 | 835 | 669 | 546 | 681 | 554 | 0.008106059 | 0.049350985 |
| ENSGALG00010029575 | ANKFN1 | ANKFN1 | | 26 | 20 | 129 | 141 | 115 | 65 | 116 | 68 | 3.62E-05 | 0.000586889 |
| ENSGALG00010029577 | NOG | NOG | | 28 | 46 | 146 | 116 | 63 | 47 | 67 | 95 | 0.005026537 | 0.033831449 |
| ENSGALG00010029584 | TRIM25 | TRIM56 | | 72 | 72 | 63 | 81 | 47 | 37 | 148 | 132 | 0.008607526 | 0.05177804 |
| ENSGALG00010029588 | RAB11FIP4 | RAB11FIP4 | | 542 | 934 | 468 | 506 | 537 | 554 | 1401 | 845 | 0.002821497 | 0.021336134 |
| ENSGALG00010029599 | RNF43 | RNF43 | | 66 | 66 | 34 | 33 | 40 | 55 | 127 | 91 | 0.008973892 | 0.053561317 |
| ENSGALG00010029603 | RHBDL3 | RHBDL3 | | 138 | 161 | 11 | 48 | 14 | 32 | 111 | 59 | 0.000151679 | 0.001971114 |
| ENSGALG00010029617 | POR | CPR | | 1476 | 1630 | 1044 | 1082 | 957 | 620 | 1059 | 1173 | 0.0086929 | 0.052201133 |
| ENSGALG00010029618 | TEX2 | TEX2 | | 1296 | 1477 | 755 | 907 | 1182 | 823 | 1060 | 874 | 0.006675911 | 0.042116962 |
| ENSGALG00010029638 | EPN3 | EPN3 | | 157 | 511 | 871 | 774 | 606 | 322 | 786 | 1110 | 0.003145295 | 0.023330745 |
| ENSGALG00010029640 | LOC112530082 | LOC112530082 | | 1 | 2 | 3 | 5 | 17 | 16 | 1 | 1 | 3.45E-06 | 7.79E-05 |
| ENSGALG00010029649 | PSPH | PSPH | | 134 | 68 | 63 | 43 | 64 | 247 | 463 | 548 | 9.39E-07 | 2.46E-05 |
| ENSGALG00010029665 | ASL1 | ASL | | 106 | 154 | 85 | 117 | 131 | 87 | 209 | 341 | 0.003513404 | 0.025506522 |
| ENSGALG00010029675 | DHRS13 | DHRS13 | | 15 | 6 | 28 | 25 | 49 | 82 | 132 | 236 | 1.00E-10 | 6.13E-09 |
| ENSGALG00010029676 | FADS6 | FADS6 | | 116 | 136 | 33 | 52 | 40 | 19 | 119 | 168 | 5.23E-06 | 0.000111993 |
| ENSGALG00010029678 | USH1G | USH1G | | 3 | 1 | 1 | 1 | 0 | 4 | 14 | 13 | 0.002671398 | 0.020441574 |
| ENSGALG00010029693 | HN1 | ARM2 | | 16708 | 12994 | 5510 | 6819 | 7937 | 5851 | 9351 | 8419 | 5.87E-10 | 3.14E-08 |
| ENSGALG00010029718 | ABR | ABR | | 1194 | 2575 | 481 | 983 | 1848 | 1917 | 1406 | 1554 | 1.44E-05 | 0.000266384 |
| ENSGALG00010029721 | EXOC7 | EXOC7 | | 1215 | 1287 | 1974 | 2344 | 1738 | 1322 | 1869 | 2237 | 0.00899609 | 0.05365548 |
| ENSGALG00010029724 | CACNA1G | CACNA1G | | 831 | 1703 | 210 | 199 | 389 | 541 | 1432 | 1183 | 1.36E-13 | 1.29E-11 |
| ENSGALG00010029726 | SEPTIN9 | | 9-Sep | 3887 | 5069 | 2235 | 2486 | 2509 | 2060 | 3398 | 5287 | 1.14E-05 | 0.000219758 |
| ENSGALG00010029743 | MYOCD | MYOCD | | 77 | 6 | 10 | 23 | 75 | 82 | 82 | 90 | 0.009892037 | 0.057822919 |
| ENSGALG00010029745 | SHPK | CARKL | | 182 | 195 | 283 | 311 | 454 | 366 | 267 | 322 | 0.001199407 | 0.010805653 |
| ENSGALG00010029758 | DNAH9 | DNAH9 | | 116 | 137 | 572 | 278 | 359 | 226 | 335 | 177 | 0.00274541 | 0.02087411 |
| ENSGALG00010029780 | TEX14 | TEX14 | | 899 | 1203 | 2246 | 2074 | 1514 | 1045 | 1447 | 1173 | 0.000796391 | 0.007762536 |
| ENSGALG00010029788 | LOC417372 | LOC417372 | | 12 | 14 | 2 | 2 | 3 | 15 | 28 | 26 | 0.001167512 | 0.0105527 |
| ENSGALG00010029817 | NA | NA | | 627 | 952 | 887 | 841 | 1270 | 1482 | 2949 | 1065 | 0.000350867 | 0.003972687 |
| ENSGALG00010029837 | NA | NA | | 1040 | 1622 | 1081 | 1585 | 876 | 695 | 453 | 850 | 0.005009625 | 0.033780085 |
| ENSGALG00010029841 | SMIM5 | SMIM5 | | 1 | 3 | 6 | 0 | 22 | 16 | 33 | 22 | 5.46E-06 | 0.00011603 |
| ENSGALG00010029864 | SLC16A3 | MCT4 | | 7296 | 9433 | 4050 | 5867 | 6677 | 4445 | 5754 | 3896 | 0.000279243 | 0.003303766 |
| ENSGALG00010029907 | NTN1 | netrin-1 | | 152 | 428 | 210 | 161 | 123 | 126 | 481 | 1103 | 7.79E-07 | 2.08E-05 |
| ENSGALG00010029927 | CFAP52 | WDR16 | | 70 | 131 | 438 | 358 | 330 | 241 | 232 | 224 | 2.08E-05 | 0.000362919 |
| ENSGALG00010029937 | PMP22 | PMP22 | | 7 | 23 | 68 | 39 | 16 | 8 | 30 | 81 | 0.003213877 | 0.02374461 |
| ENSGALG00010029948 | MYADML2 | MYADML2 | | 24 | 48 | 9 | 3 | 28 | 43 | 50 | 57 | 9.72E-05 | 0.001350552 |
| ENSGALG00010029988 | NA | NA | | 4 | 18 | 61 | 44 | 43 | 29 | 28 | 19 | 0.004416841 | 0.03062832 |
| ENSGALG00010030008 | NA | NA | | 2108 | 2065 | 1231 | 1502 | 1413 | 995 | 1099 | 1332 | 0.001988044 | 0.016112608 |
| ENSGALG00010030013 | NA | NA | | 0 | 2 | 22 | 9 | 8 | 8 | 16 | 6 | 0.00629105 | 0.040204981 |
| ENSGALG00010030063 | NA | NA | | 5 | 1 | 11 | 3 | 20 | 14 | 43 | 21 | 0.000334051 | 0.003815897 |
| ENSGALG00010030065 | NA | NA | | 13 | 0 | 0 | 2 | 0 | 1 | 9 | 23 | 0.004335774 | 0.030166298 |
| ENSGALG00010030104 | P4HB | PDI | | 27871 | 21054 | 13448 | 16753 | 12428 | 11115 | 20039 | 21600 | 0.000275743 | 0.003275212 |
| ENSGALG00010030109 | FBF1 | FBF1 | | 1192 | 730 | 58 | 64 | 1186 | 695 | 523 | 480 | 1.06E-28 | 4.01E-26 |

### Supplementally table S4. GO analysis of DEGs comparing four strains of cultured PGCs

| source | term_name | term_id | adjusted_p_value | negative_log10_of_adjusted_p_value | term_size | query_size | intersection_size |
|---|---|---|---|---|---|---|---|
| GO:MF | calcium ion binding | GO:0005509 | 4.00385E-05 | 4.39752272 | 542 | 1620 | 112 |
| GO:MF | monoatomic cation channel activity | GO:0005261 | 9.28535E-05 | 4.03220168 | 259 | 1620 | 63 |
| GO:MF | monoatomic ion channel activity | GO:0005216 | 0.000171533 | 3.76565149 | 344 | 1620 | 77 |
| GO:MF | passive transmembrane transporter activity | GO:0022803 | 0.000321605 | 3.49267743 | 385 | 1620 | 83 |
| GO:MF | channel activity | GO:0015267 | 0.000321605 | 3.49267743 | 385 | 1620 | 83 |
| GO:MF | transmembrane transporter activity | GO:0022857 | 0.004663723 | 2.33126728 | 866 | 1620 | 153 |
| GO:MF | extracellular matrix structural constituent | GO:0005201 | 0.005777592 | 2.23825316 | 28 | 1620 | 13 |
| GO:MF | metal ion transmembrane transporter activity | GO:0046873 | 0.006485458 | 2.18805932 | 334 | 1620 | 70 |
| GO:MF | monoatomic cation transmembrane transporter activity | GO:0008324 | 0.00871574 | 2.05969574 | 468 | 1620 | 91 |
| GO:MF | gated channel activity | GO:0022836 | 0.009014262 | 2.04506981 | 247 | 1620 | 55 |
| GO:MF | transporter activity | GO:0005215 | 0.009907697 | 2.00402726 | 939 | 1620 | 162 |
| GO:MF | monoatomic ion transmembrane transporter activity | GO:0015075 | 0.009938636 | 2.00267323 | 566 | 1620 | 106 |
| GO:MF | inorganic molecular entity transmembrane transporter activity | GO:0015318 | 0.026845564 | 1.57112746 | 514 | 1620 | 96 |
| GO:MF | inorganic cation transmembrane transporter activity | GO:0022890 | 0.039642916 | 1.40183441 | 435 | 1620 | 83 |
| GO:BP | multicellular organismal process | GO:0032501 | 9.47E-18 | 17.0234877 | 3848 | 1548 | 642 |
| GO:BP | system development | GO:0048731 | 6.28E-16 | 15.2018023 | 2305 | 1548 | 420 |
| GO:BP | anatomical structure development | GO:0048856 | 1.44E-15 | 14.8408294 | 3372 | 1548 | 568 |
| GO:BP | multicellular organism development | GO:0007275 | 5.28E-15 | 14.2772321 | 2718 | 1548 | 475 |
| GO:BP | developmental process | GO:0032502 | 9.15E-15 | 14.0384582 | 3583 | 1548 | 593 |
| GO:BP | nervous system development | GO:0007399 | 5.81E-14 | 13.2355507 | 1380 | 1548 | 275 |
| GO:BP | anatomical structure morphogenesis | GO:0009653 | 2.08E-13 | 12.6829719 | 1657 | 1548 | 315 |
| GO:BP | cell adhesion | GO:0007155 | 1.39E-11 | 10.855508 | 793 | 1548 | 174 |
| GO:BP | tissue development | GO:0009888 | 1.00E-10 | 9.99826474 | 1209 | 1548 | 237 |
| GO:BP | locomotion | GO:0040011 | 2.13E-10 | 9.67140439 | 724 | 1548 | 159 |
| GO:BP | animal organ development | GO:0048513 | 2.23E-10 | 9.65080376 | 1740 | 1548 | 315 |
| GO:BP | regulation of multicellular organismal process | GO:0051239 | 4.44E-09 | 8.3530686 | 1635 | 1548 | 294 |
| GO:BP | anatomical structure formation involved in morphogenesis | GO:0048646 | 5.14E-09 | 8.28890338 | 714 | 1548 | 153 |
| GO:BP | cell migration | GO:0016477 | 5.59E-09 | 8.25266558 | 896 | 1548 | 182 |
| GO:BP | generation of neurons | GO:0048699 | 1.50E-08 | 7.82462866 | 893 | 1548 | 180 |
| GO:BP | neurogenesis | GO:0022008 | 3.11E-08 | 7.50741447 | 1036 | 1548 | 201 |
| GO:BP | neuron differentiation | GO:0030182 | 3.94E-08 | 7.40461158 | 839 | 1548 | 170 |
| GO:BP | cell junction assembly | GO:0034329 | 6.22E-08 | 7.20596088 | 247 | 1548 | 69 |
| GO:BP | system process | GO:0003008 | 8.96E-08 | 7.04782814 | 963 | 1548 | 188 |
| GO:BP | neuron projection guidance | GO:0097485 | 1.23E-07 | 6.90977291 | 136 | 1548 | 46 |
| GO:BP | cell motility | GO:0048870 | 1.41E-07 | 6.85064758 | 994 | 1548 | 192 |
| GO:BP | cell morphogenesis | GO:0000902 | 1.53E-07 | 6.81538722 | 584 | 1548 | 127 |
| GO:BP | cellular developmental process | GO:0048869 | 1.63E-07 | 6.78793728 | 2365 | 1548 | 390 |
| GO:BP | cell differentiation | GO:0030154 | 1.63E-07 | 6.78793728 | 2365 | 1548 | 390 |
| GO:BP | axon guidance | GO:0007411 | 3.43E-07 | 6.46439973 | 135 | 1548 | 45 |
| GO:BP | regulation of locomotion | GO:0040012 | 4.11E-07 | 6.3856652 | 617 | 1548 | 131 |
| GO:BP | regulation of cell adhesion | GO:0030155 | 4.25E-07 | 6.37130091 | 431 | 1548 | 100 |
| GO:BP | regulation of biological quality | GO:0065008 | 6.88E-07 | 6.16239249 | 1567 | 1548 | 274 |
| GO:BP | circulatory system development | GO:0072359 | 6.91E-07 | 6.16077168 | 709 | 1548 | 145 |
| GO:BP | cell morphogenesis involved in neuron differentiation | GO:0048667 | 8.39E-07 | 6.07603228 | 338 | 1548 | 83 |
| GO:BP | plasma membrane bounded cell projection morphogenesis | GO:0120039 | 1.10267E-06 | 5.95755439 | 374 | 1548 | 89 |
| GO:BP | axonogenesis | GO:0007409 | 1.27003E-06 | 5.89618475 | 263 | 1548 | 69 |
| GO:BP | cell projection morphogenesis | GO:0048858 | 1.27425E-06 | 5.89474624 | 375 | 1548 | 89 |
| GO:BP | tube development | GO:0035295 | 2.24155E-06 | 5.64945091 | 689 | 1548 | 140 |
| GO:BP | regulation of cell migration | GO:0030334 | 2.40901E-06 | 5.61816074 | 565 | 1548 | 120 |
| GO:BP | cell junction organization | GO:0034330 | 2.57594E-06 | 5.58906482 | 415 | 1548 | 95 |
| GO:BP | regulation of cell motility | GO:2000145 | 3.10899E-06 | 5.50738107 | 592 | 1548 | 124 |
| GO:BP | axon development | GO:0061564 | 4.85187E-06 | 5.31409087 | 293 | 1548 | 73 |
| GO:BP | central nervous system development | GO:0007417 | 5.23498E-06 | 5.28108482 | 572 | 1548 | 120 |
| GO:BP | positive regulation of multicellular organismal process | GO:0051240 | 8.68674E-06 | 5.06114304 | 884 | 1548 | 168 |
| GO:BP | neuron projection development | GO:0031175 | 1.32803E-05 | 4.87679319 | 562 | 1548 | 117 |
| GO:BP | nervous system process | GO:0050877 | 0.00001443 | 4.84073455 | 569 | 1548 | 118 |
| GO:BP | vasculature development | GO:0001944 | 1.50071E-05 | 4.82370224 | 477 | 1548 | 103 |
| GO:BP | neuron projection morphogenesis | GO:0048812 | 1.58219E-05 | 4.80074263 | 364 | 1548 | 84 |
| GO:BP | positive regulation of developmental process | GO:0051094 | 1.98245E-05 | 4.702798 | 776 | 1548 | 150 |
| GO:BP | blood vessel development | GO:0001568 | 2.22104E-05 | 4.65344401 | 450 | 1548 | 98 |
| GO:BP | tube morphogenesis | GO:0035239 | 3.33296E-05 | 4.47716936 | 552 | 1548 | 114 |
| GO:BP | regulation of developmental process | GO:0050793 | 3.35397E-05 | 4.47444045 | 1445 | 1548 | 248 |
| GO:BP | cell-cell signaling | GO:0007267 | 4.97245E-05 | 4.30342929 | 638 | 1548 | 127 |
| GO:BP | monoatomic ion transmembrane transport | GO:0034220 | 0.000060078 | 4.22128487 | 514 | 1548 | 107 |
| GO:BP | regulation of neuron projection development | GO:0010975 | 6.22218E-05 | 4.20605775 | 220 | 1548 | 57 |
| GO:BP | epithelium development | GO:0060429 | 0.000077572 | 4.11029511 | 733 | 1548 | 141 |
| GO:BP | neuron development | GO:0048666 | 7.81385E-05 | 4.10713519 | 662 | 1548 | 130 |
| GO:BP | head development | GO:0060322 | 7.92411E-05 | 4.10104929 | 461 | 1548 | 98 |
| GO:BP | metal ion transport | GO:0030001 | 0.000111935 | 3.95103393 | 495 | 1548 | 103 |
| GO:BP | monoatomic cation transmembrane transport | GO:0098655 | 0.000154437 | 3.81124745 | 418 | 1548 | 90 |
| GO:BP | regulation of plasma membrane bounded cell projection organization | GO:0120035 | 0.000176664 | 3.75285133 | 347 | 1548 | 78 |
| GO:BP | regulation of cell projection organization | GO:0031344 | 0.000178055 | 3.74944643 | 353 | 1548 | 79 |
| GO:BP | cell-cell adhesion | GO:0098609 | 0.000179065 | 3.74698896 | 487 | 1548 | 101 |
| GO:BP | monoatomic cation transport | GO:0006812 | 0.000247677 | 3.60611504 | 572 | 1548 | 114 |
| GO:BP | positive regulation of locomotion | GO:0040017 | 0.000256618 | 3.59071363 | 344 | 1548 | 77 |
| GO:BP | regulation of multicellular organismal development | GO:2000026 | 0.000283421 | 3.5475673 | 801 | 1548 | 149 |
| GO:BP | transmembrane transport | GO:0055085 | 0.000309667 | 3.50910559 | 956 | 1548 | 172 |
| GO:BP | embryonic morphogenesis | GO:0048598 | 0.000375382 | 3.42552629 | 432 | 1548 | 91 |
| GO:BP | chemotaxis | GO:0006935 | 0.000379257 | 3.42106598 | 231 | 1548 | 57 |
| GO:BP | positive regulation of cell motility | GO:2000147 | 0.000419407 | 3.37736392 | 336 | 1548 | 75 |
| GO:BP | synapse assembly | GO:0007416 | 0.000446774 | 3.34991238 | 110 | 1548 | 34 |
| GO:BP | blood vessel morphogenesis | GO:0048514 | 0.000466631 | 3.3310263 | 385 | 1548 | 83 |
| GO:BP | mesenchyme development | GO:0060485 | 0.000469096 | 3.32873846 | 199 | 1548 | 51 |
| GO:BP | taxis | GO:0042330 | 0.000516966 | 3.28653834 | 233 | 1548 | 57 |
| GO:BP | brain development | GO:0007420 | 0.000539946 | 3.26764953 | 423 | 1548 | 89 |
| GO:BP | monoatomic ion transport | GO:0006811 | 0.000604729 | 3.21843894 | 718 | 1548 | 135 |
| GO:BP | negative regulation of neuron projection development | GO:0010977 | 0.000786437 | 3.10433614 | 61 | 1548 | 23 |
| GO:BP | positive regulation of cell migration | GO:0030335 | 0.000874466 | 3.05825728 | 324 | 1548 | 72 |
| GO:BP | inorganic ion transmembrane transport | GO:0098660 | 0.00091287 | 3.03959124 | 459 | 1548 | 94 |
| GO:BP | cell development | GO:0048468 | 0.001072824 | 2.96947158 | 1610 | 1548 | 263 |
| GO:BP | inorganic cation transmembrane transport | GO:0098662 | 0.001094938 | 2.96061034 | 411 | 1548 | 86 |
| GO:BP | mesenchymal cell differentiation | GO:0048762 | 0.00173811 | 2.7599227 | 163 | 1548 | 43 |
| GO:BP | plasma membrane bounded cell projection organization | GO:0120036 | 0.001994476 | 2.70017108 | 935 | 1548 | 165 |
| GO:BP | regulation of membrane potential | GO:0042391 | 0.002224459 | 2.65277554 | 220 | 1548 | 53 |
| GO:BP | developmental growth | GO:0048589 | 0.002350388 | 2.62886049 | 412 | 1548 | 85 |
| GO:BP | behavior | GO:0007610 | 0.002561014 | 2.59158807 | 388 | 1548 | 81 |
| GO:BP | regulation of nervous system development | GO:0051960 | 0.002715146 | 2.56620688 | 256 | 1548 | 59 |
| GO:BP | cell projection organization | GO:0030030 | 0.00282663 | 2.54873107 | 947 | 1548 | 166 |
| GO:BP | cell surface receptor protein tyrosine kinase signaling pathway | GO:0007169 | 0.003169833 | 2.49896355 | 390 | 1548 | 81 |
| GO:BP | biological regulation | GO:0065007 | 0.003333853 | 2.47705359 | 7514 | 1548 | 1019 |
| GO:BP | autonomic nervous system development | GO:0048483 | 0.003494709 | 2.45658901 | 36 | 1548 | 16 |
| GO:BP | tissue morphogenesis | GO:0048729 | 0.003641065 | 2.43877154 | 410 | 1548 | 84 |
| GO:BP | synapse organization | GO:0050808 | 0.003978842 | 2.40024334 | 253 | 1548 | 58 |
| GO:BP | forebrain development | GO:0030900 | 0.003978842 | 2.40024334 | 253 | 1548 | 58 |
| GO:BP | angiogenesis | GO:0001525 | 0.004209609 | 2.37575826 | 319 | 1548 | 69 |
| GO:BP | response to chemical | GO:0042221 | 0.004739956 | 2.32422566 | 1776 | 1548 | 282 |
| GO:BP | morphogenesis of an epithelium | GO:0002009 | 0.005136101 | 2.28936644 | 339 | 1548 | 72 |
| GO:BP | regulation of heart rate | GO:0002027 | 0.005336431 | 2.27274908 | 58 | 1548 | 21 |
| GO:BP | renal system development | GO:0072001 | 0.005900876 | 2.22908353 | 221 | 1548 | 52 |
| GO:BP | animal organ morphogenesis | GO:0009887 | 0.006691867 | 2.17445271 | 669 | 1548 | 123 |
| GO:BP | learning or memory | GO:0007611 | 0.006743114 | 2.1711395 | 149 | 1548 | 39 |
| GO:BP | embryo development | GO:0009790 | 0.008544027 | 2.06833737 | 719 | 1548 | 130 |
| GO:BP | neuromuscular process | GO:0050905 | 0.008764586 | 2.05726857 | 103 | 1548 | 30 |

| | | | | | | | |
|---|---|---|---|---|---|---|---|
| GO:BP | sensory organ development | GO:0007423 | 0.010464978 | 1.98026167 | 408 | 1548 | 82 |
| GO:BP | cell-substrate adhesion | GO:0031589 | 0.011183953 | 1.95140468 | 214 | 1548 | 50 |
| GO:BP | regulation of cell junction assembly | GO:1901888 | 0.011252423 | 1.94875396 | 125 | 1548 | 34 |
| GO:BP | kidney development | GO:0001822 | 0.016476166 | 1.78314384 | 211 | 1548 | 49 |
| GO:BP | adenylate cyclase-modulating G protein-coupled receptor signaling pathway | GO:0007188 | 0.0196805 | 1.70596386 | 128 | 1548 | 34 |
| GO:BP | regulation of anatomical structure morphogenesis | GO:0022603 | 0.01998651 | 1.69926303 | 479 | 1548 | 92 |
| GO:BP | muscle tissue development | GO:0060537 | 0.020523482 | 1.68774896 | 266 | 1548 | 58 |
| GO:BP | negative regulation of multicellular organismal process | GO:0051241 | 0.023816947 | 1.6231139 | 586 | 1548 | 108 |
| GO:BP | regulation of developmental growth | GO:0048638 | 0.024632522 | 1.60849112 | 185 | 1548 | 44 |
| GO:BP | negative regulation of developmental process | GO:0051093 | 0.025316859 | 1.59659018 | 534 | 1548 | 100 |
| GO:BP | positive regulation of synaptic transmission | GO:0050806 | 0.025590686 | 1.59191806 | 68 | 1548 | 22 |
| GO:BP | cell surface receptor signaling pathway | GO:0007166 | 0.025621076 | 1.59140263 | 1558 | 1548 | 247 |
| GO:BP | chemical synaptic transmission | GO:0007268 | 0.026365848 | 1.57895825 | 367 | 1548 | 74 |
| GO:BP | anterograde trans-synaptic signaling | GO:0098916 | 0.026365848 | 1.57895825 | 367 | 1548 | 74 |
| GO:BP | neuromuscular process controlling balance | GO:0050885 | 0.026760599 | 1.57250416 | 41 | 1548 | 16 |
| GO:BP | negative regulation of nervous system development | GO:0051961 | 0.026871596 | 1.57070653 | 73 | 1548 | 23 |
| GO:BP | morphogenesis of embryonic epithelium | GO:0016331 | 0.030183259 | 1.52023387 | 109 | 1548 | 30 |
| GO:BP | heart development | GO:0007507 | 0.031080093 | 1.50751769 | 375 | 1548 | 75 |
| GO:BP | gastrulation | GO:0007369 | 0.033571024 | 1.47403541 | 131 | 1548 | 34 |
| GO:BP | regulation of biological process | GO:0050789 | 0.034736935 | 1.4592085 | 7293 | 1548 | 983 |
| GO:BP | trans-synaptic signaling | GO:0099537 | 0.035311125 | 1.45208845 | 370 | 1548 | 74 |
| GO:BP | central nervous system neuron differentiation | GO:0021953 | 0.036376598 | 1.43917792 | 148 | 1548 | 37 |
| GO:BP | equilibrioception | GO:0050957 | 0.04134056 | 1.38362365 | 7 | 1548 | 6 |
| GO:BP | positive regulation of cell adhesion | GO:0045785 | 0.042469377 | 1.37192411 | 236 | 1548 | 52 |
| GO:BP | regulation of axonogenesis | GO:0050770 | 0.043902525 | 1.35751051 | 75 | 1548 | 23 |
| GO:BP | negative regulation of growth | GO:0045926 | 0.04998225 | 1.3011842 | 117 | 1548 | 31 |
| GO:CC | cell periphery | GO:0071944 | 9.12E-23 | 22.039784 | 2860 | 1473 | 509 |
| GO:CC | plasma membrane | GO:0005886 | 4.46E-16 | 15.3504978 | 2604 | 1473 | 448 |
| GO:CC | plasma membrane region | GO:0098590 | 1.29E-08 | 7.89024454 | 597 | 1473 | 127 |
| GO:CC | membrane | GO:0016020 | 1.98E-08 | 7.7040844 | 5127 | 1473 | 734 |
| GO:CC | cation channel complex | GO:0034703 | 3.49E-08 | 7.4576766 | 147 | 1473 | 47 |
| GO:CC | cell junction | GO:0030054 | 6.93E-08 | 7.15951632 | 1121 | 1473 | 204 |
| GO:CC | monoatomic ion channel complex | GO:0034702 | 6.99E-08 | 7.15541495 | 211 | 1473 | 59 |
| GO:CC | extracellular region | GO:0005576 | 4.55E-07 | 6.34215752 | 960 | 1473 | 177 |
| GO:CC | external encapsulating structure | GO:0030312 | 7.83E-07 | 6.10607705 | 218 | 1473 | 58 |
| GO:CC | extracellular matrix | GO:0031012 | 7.83E-07 | 6.10607705 | 218 | 1473 | 58 |
| GO:CC | neuron projection | GO:0043005 | 8.71E-07 | 6.06007593 | 597 | 1473 | 121 |
| GO:CC | plasma membrane protein complex | GO:0098797 | 9.24E-07 | 6.03430002 | 362 | 1473 | 83 |
| GO:CC | transporter complex | GO:1990351 | 2.72459E-06 | 5.56469826 | 270 | 1473 | 66 |
| GO:CC | transmembrane transporter complex | GO:1902495 | 2.7615E-06 | 5.55885483 | 253 | 1473 | 63 |
| GO:CC | collagen-containing extracellular matrix | GO:0062023 | 3.62678E-06 | 5.44047859 | 131 | 1473 | 40 |
| GO:CC | extracellular space | GO:0005615 | 6.07364E-06 | 5.21655094 | 649 | 1473 | 126 |
| GO:CC | cell surface | GO:0009986 | 1.64286E-05 | 4.78440052 | 372 | 1473 | 81 |
| GO:CC | synaptic membrane | GO:0097060 | 9.29474E-05 | 4.03176292 | 173 | 1473 | 45 |
| GO:CC | synapse | GO:0045202 | 0.000420609 | 3.37612115 | 733 | 1473 | 131 |
| GO:CC | neuronal cell body | GO:0043025 | 0.000463863 | 3.33361018 | 160 | 1473 | 41 |
| GO:CC | somatodendritic compartment | GO:0036477 | 0.000464618 | 3.33290428 | 326 | 1473 | 69 |
| GO:CC | cell projection | GO:0042995 | 0.000671597 | 3.17289106 | 1141 | 1473 | 188 |
| GO:CC | plasma membrane bounded cell projection | GO:0120025 | 0.000939014 | 3.02732791 | 1118 | 1473 | 184 |
| GO:CC | basal part of cell | GO:0045178 | 0.00104391 | 2.98133715 | 148 | 1473 | 38 |
| GO:CC | cellular anatomical structure | GO:0110165 | 0.001483128 | 2.82882146 | 11921 | 1473 | 1461 |
| GO:CC | axon | GO:0030424 | 0.001751431 | 2.75660702 | 257 | 1473 | 56 |
| GO:CC | postsynapse | GO:0098794 | 0.002254051 | 2.64703617 | 290 | 1473 | 61 |
| GO:CC | neuronal cell body membrane | GO:0032809 | 0.002848781 | 2.54534092 | 9 | 1473 | 7 |
| GO:CC | glutamatergic synapse | GO:0098978 | 0.003376272 | 2.47156257 | 244 | 1473 | 53 |
| GO:CC | basolateral plasma membrane | GO:0016323 | 0.004033984 | 2.39426585 | 117 | 1473 | 31 |
| GO:CC | potassium channel complex | GO:0034705 | 0.006378402 | 2.19528811 | 67 | 1473 | 21 |
| GO:CC | calcium channel complex | GO:0034704 | 0.007628025 | 2.11758791 | 44 | 1473 | 16 |
| GO:CC | cell body membrane | GO:0044298 | 0.00850047 | 2.07055706 | 10 | 1473 | 7 |
| GO:CC | postsynaptic membrane | GO:0045211 | 0.008877655 | 2.05170172 | 127 | 1473 | 32 |
| GO:CC | basement membrane | GO:0005604 | 0.016567018 | 1.78075566 | 56 | 1473 | 18 |
| GO:CC | cell projection membrane | GO:0031253 | 0.017065027 | 1.76789303 | 166 | 1473 | 38 |
| GO:CC | membrane protein complex | GO:0098796 | 0.024822999 | 1.60514575 | 752 | 1473 | 125 |
| GO:CC | basal plasma membrane | GO:0009925 | 0.027112632 | 1.56682832 | 134 | 1473 | 32 |
| GO:CC | postsynaptic specialization membrane | GO:0099634 | 0.027136461 | 1.56644679 | 53 | 1473 | 17 |
| GO:CC | presynaptic membrane | GO:0042734 | 0.027714169 | 1.55729814 | 58 | 1473 | 18 |
| GO:CC | anchoring junction | GO:0070161 | 0.032177013 | 1.49245427 | 382 | 1473 | 71 |
| GO:CC | sarcolemma | GO:0042383 | 0.034924303 | 1.45687225 | 64 | 1473 | 19 |
| GO:CC | cell body | GO:0044297 | 0.041151132 | 1.38561821 | 191 | 1473 | 41 |
| GO:CC | voltage-gated potassium channel complex | GO:0008076 | 0.04498441 | 1.34693797 | 60 | 1473 | 18 |
| KEGG | ECM-receptor interaction | KEGG:04512 | 0.001278762 | 2.89321025 | 81 | 679 | 25 |
| KEGG | Biosynthesis of amino acids | KEGG:01230 | 0.003874814 | 2.41174918 | 57 | 679 | 19 |
| KEGG | Calcium signaling pathway | KEGG:04020 | 0.020807751 | 1.68177485 | 226 | 679 | 48 |

### Supplementally table S5. DEGs comparing M/O PGCs and Fayoumi (GSP, GSN/1 and PNP) PGCs

| Emsembl_Id | Symbol | entrez_ID | MO_Male_TPM | MO_Female_TPM | GSP_Male_TPM | GSP_Female_TPM | GSN_Male_TPM | GSN_Female_TPM | PNP_Female_TPM | PNP_Male_TPM |
|---|---|---|---|---|---|---|---|---|---|---|
| ENSGALG00010015173 | ENS-1 | 769415 | 4908.89 | 2147.68 | 780.488 | 1365.18 | 828.345 | 1266.53 | 1942.95 | 1854.11 |
| ENSGALG00010006087 | LOC107051465 | 107051465 | 4498.56 | 4942.55 | 812.534 | 1354.39 | 817.725 | 1424.1 | 2215.66 | 1903.72 |
| ENSGALG00010004570 | LOC107051826 | 107051826 | 3568.04 | 3913.24 | 449.668 | 748.079 | 321.21 | 663.504 | 1526.97 | 1539.11 |
| ENSGALG00010002022 | ERNI | 769673 | 2470.02 | 3300.43 | 508.579 | 843.743 | 534.697 | 991.468 | 1798.01 | 1991.1 |
| ENSGALG00010003305 | CALR | 100859104 | 2310.24 | 1947.66 | 1345.27 | 1490.57 | 1658.41 | 1095.45 | 1579.51 | 1147.91 |
| ENSGALG00010022849 | NA | NA | 1992.4 | 2760.85 | 167.359 | 183.13 | 134.601 | 240.361 | 288.964 | 356.146 |
| ENSGALG00010027969 | HSPA5 | 396487 | 1629.1 | 1183.86 | 773.343 | 891.108 | 841.608 | 573.956 | 801.575 | 741.628 |
| ENSGALG00010004699 | NA | NA | 1077.42 | 720.459 | 541.999 | 765.195 | 633.861 | 480.861 | 305.939 | 259.896 |
| ENSGALG00010004281 | VIM | 420519 | 944.37 | 1262.56 | 483.274 | 618.071 | 520.383 | 261.116 | 810.233 | 457.868 |
| ENSGALG00010010280 | HSP90B1 | 374163 | 933.585 | 763.463 | 547.027 | 629.8 | 608.077 | 407.145 | 478.336 | 402.941 |
| ENSGALG00010011992 | YEATS4 | 771226 | 721.508 | 929.455 | 639.532 | 679.585 | 701.108 | 525.85 | 531.058 | 403.691 |
| ENSGALG00010027214 | PLIN4 | 100857433 | 609.998 | 774.585 | 265.123 | 260.413 | 413.265 | 260.739 | 443.961 | 284.228 |
| ENSGALG00010012582 | CD24 | 100859742 | 561.179 | 510.303 | 235.977 | 348.877 | 333.078 | 289.575 | 390.637 | 304.987 |
| ENSGALG00010026300 | HMGA1 | 378795 | 445.939 | 429.073 | 317.348 | 348.643 | 374.909 | 281.346 | 325.602 | 207.663 |
| ENSGALG00010024452 | NA | NA | 399.796 | 396.481 | 216.759 | 261.341 | 263.684 | 232.447 | 254.516 | 259.752 |
| ENSGALG00010024690 | LOC100857858 | 100857858 | 355.891 | 340.128 | 511.904 | 531.206 | 559.832 | 378.172 | 307.931 | 236.301 |
| ENSGALG00010021669 | SAR1A | 423711 | 334.644 | 352.202 | 252.211 | 276.838 | 274.009 | 183.277 | 187.511 | 159.196 |
| ENSGALG00010014980 | LOC121107687 | 121107687 | 315.323 | 477.574 | 102.892 | 124.401 | 70.9123 | 81.2255 | 205.718 | 132.711 |
| ENSGALG00010007287 | LOC124418405 | 124418405 | 270.722 | 77.2361 | 28.3746 | 46.1016 | 36.4591 | 60.8022 | 76.7835 | 76.4577 |
| ENSGALG00010009928 | NA | NA | 208.534 | 204.309 | 42.146 | 69.0822 | 37.5266 | 58.7747 | 115.061 | 96.8406 |
| ENSGALG00010007054 | NA | NA | 194.492 | 188.53 | 115.161 | 118.707 | 149.459 | 120.231 | 112.263 | 114.493 |
| ENSGALG00010027297 | LOC121107553 | 121107553 | 180.359 | 123.943 | 265.71 | 293.337 | 59.5395 | 49.8493 | 362.667 | 295.262 |
| ENSGALG00010001386 | FKBP14 | 420636 | 161.286 | 123.909 | 56.103 | 52.9252 | 94.648 | 83.1791 | 69.9671 | 72.7787 |
| ENSGALG00010000465 | CUNH17orf49 | 112531156 | 161.062 | 141.209 | 109.543 | 109.152 | 133.821 | 90.5472 | 109.4 | 77.5342 |
| ENSGALG00010028815 | NA | NA | 148.775 | 205.569 | 133.186 | 130.05 | 43.0863 | 35.2628 | 173.548 | 129.8 |
| ENSGALG00010029693 | HN1 | 422119 | 148.747 | 112.529 | 57.4234 | 62.7092 | 89.1761 | 69.2423 | 75.0255 | 78.4965 |
| ENSGALG00010003700 | REST | 422623 | 148.286 | 105.343 | 75.4373 | 104.258 | 76.891 | 79.6219 | 44.0224 | 69.4097 |
| ENSGALG00010019621 | LOC421441 | 421441 | 146.2 | 175.004 | 122.929 | 105.379 | 67.6482 | 41.9766 | 110.221 | 84.697 |
| ENSGALG00010022306 | NA | NA | 145.086 | 92.8646 | 60.1356 | 89.2263 | 87.0695 | 60.2258 | 59.7482 | 69.3523 |
| ENSGALG00010027727 | GSTM2 | 395976 | 141.23 | 107.961 | 17.9044 | 17.4902 | 57.417 | 57.4763 | 52.4985 | 42.0888 |
| ENSGALG00010001705 | HTRA1 | 100857572 | 137.422 | 107.867 | 31.8769 | 37.5029 | 68.9331 | 51.9575 | 39.7011 | 68.3162 |
| ENSGALG00010025783 | NA | NA | 137.313 | 105.538 | 290.922 | 295.999 | 126.21 | 108.278 | 253.413 | 254.129 |
| ENSGALG00010022340 | GFOD2 | 415642 | 123.191 | 176.316 | 87.8199 | 107.522 | 99.4511 | 69.8276 | 71.9306 | 55.471 |
| ENSGALG00010006885 | RHOU | 428589 | 113.163 | 53.9499 | 23.8339 | 33.6915 | 40.5878 | 40.066 | 33.5192 | 52.4195 |
| ENSGALG00010022984 | NA | NA | 103.55 | 153.172 | 23.3223 | 29.0818 | 23.6365 | 46.3633 | 46.1757 | 62.6039 |
| ENSGALG00010001989 | NA | NA | 102.972 | 18.2628 | 3.31675 | 5.77199 | 3.33092 | 4.79363 | 8.43459 | 7.326 |
| ENSGALG00010003268 | SMS | 418602 | 99.0386 | 68.4314 | 41.234 | 48.1178 | 54.5316 | 57.6635 | 54.561 | 47.2027 |
| ENSGALG00010021172 | SLC29A4 | 425593 | 96.0327 | 35.5829 | 18.6649 | 23.7836 | 11.1051 | 10.4214 | 25.5293 | 17.6515 |
| ENSGALG00010021870 | LOC101747373 | 101747373 | 95.2043 | 104.86 | 33.0788 | 35.2919 | 35.9317 | 27.6611 | 23.8055 | 27.8305 |
| ENSGALG00010027217 | SEMA6B | 428330 | 81.5564 | 97.7483 | 27.16 | 23.5981 | 59.5006 | 34.2849 | 41.3541 | 38.5813 |
| ENSGALG00010024883 | ING5 | 424852 | 80.7132 | 72.3974 | 35.6531 | 34.2978 | 27.895 | 20.5842 | 39.5329 | 38.4325 |
| ENSGALG00010009795 | NDUFA6 | 427897 | 78.4365 | 81.4837 | 200.274 | 212.998 | 229.955 | 167.974 | 181.392 | 151.623 |
| ENSGALG00010011386 | CTSB | 396329 | 73.2217 | 77.9289 | 33.2809 | 33.1838 | 35.8683 | 26.4943 | 47.3904 | 35.0075 |
| ENSGALG00010029491 | NAA38 | 426258 | 73.129 | 52.1275 | 30.3053 | 37.8611 | 49.3163 | 44.2851 | 50.4218 | 51.5062 |
| ENSGALG00010002185 | NA | NA | 70.3007 | 51.191 | 7.16425 | 7.94206 | 11.0204 | 4.49645 | 8.97508 | 7.28989 |
| ENSGALG00010001703 | NA | NA | 67.1982 | 57.1758 | 2.59042 | 6.22446 | 4.17186 | 4.2966 | 13.4786 | 14.9425 |
| ENSGALG00010023247 | LOC121106503 | 121106503 | 65.432 | 67.8065 | 104.797 | 111.423 | 109.998 | 104.044 | 97.1778 | 126.119 |
| ENSGALG00010007165 | LOC101747821 | 101747821 | 64.7641 | 302.987 | 5.62937 | 3.24595 | 11.5552 | 4.72766 | 10.6878 | 8.9632 |
| ENSGALG00010000925 | NA | NA | 57.3106 | 55.3505 | 6.3037 | 11.5716 | 11.1814 | 13.0224 | 15.9351 | 19.821 |
| ENSGALG00010009891 | NA | NA | 57.273 | 56.4047 | 14.4416 | 27.4946 | 12.8764 | 26.4902 | 24.6619 | 29.3009 |
| ENSGALG00010009809 | SMDT1 | 770755 | 56.5521 | 70.2591 | 298.932 | 306.438 | 312.313 | 238.678 | 243.664 | 195.641 |
| ENSGALG00010003543 | SOX11 | 396104 | 56.3927 | 90.8963 | 27.0515 | 33.0387 | 55.6925 | 36.496 | 41.2335 | 32.7764 |
| ENSGALG00010028460 | CRLF1 | 100858730 | 53.297 | 50.6266 | 13.5216 | 18.6716 | 26.3016 | 23.8082 | 34.5688 | 21.1421 |
| ENSGALG00010001570 | SMPD1 | 419077 | 53.1862 | 47.8514 | 20.7167 | 25.8105 | 23.4281 | 21.9742 | 30.4241 | 19.4541 |
| ENSGALG00010012842 | DDX17 | 418024 | 51.7646 | 70.6233 | 105.1 | 93.0095 | 109.331 | 81.3904 | 83.3466 | 78.1949 |
| ENSGALG00010016688 | NA | NA | 50.065 | 62.4003 | 98.9317 | 113.276 | 117.126 | 119.812 | 118.262 | 104.61 |
| ENSGALG00010027811 | MEX3D | 420082 | 49.8514 | 48.9164 | 36.1833 | 36.8349 | 41.693 | 27.6534 | 32.7611 | 30.9368 |
| ENSGALG00010024872 | RUNDC3A | 100858314 | 48.4614 | 42.8551 | 4.46124 | 10.1101 | 6.66865 | 3.59349 | 9.75818 | 17.1417 |
| ENSGALG00010000726 | POU4F1 | 100857623 | 48.3195 | 54.579 | 8.54258 | 12.0277 | 4.62084 | 8.82001 | 12.1996 | 20.6328 |
| ENSGALG00010018514 | EAF2 | 424243 | 47.9617 | 46.8335 | 14.9027 | 24.0623 | 21.8058 | 30.7404 | 22.5079 | 26.2756 |
| ENSGALG00010007115 | CD69L | 417816 | 47.7368 | 15.2704 | 5.56187 | 8.27224 | 7.83536 | 11.5512 | 15.0152 | 14.1275 |
| ENSGALG00010015523 | MRPL22 | 416253 | 47.3798 | 44.7721 | 72.3937 | 85.1886 | 65.1813 | 53.0564 | 48.2068 | 42.5019 |
| ENSGALG00010008498 | LOC107050955 | 107050955 | 46.2506 | 46.2851 | 17.4109 | 10.2249 | 23.2617 | 13.6264 | 13.2924 | 13.2038 |
| ENSGALG00010000316 | NA | NA | 46.0825 | 127.179 | 66.8064 | 68.9898 | 34.1461 | 38.7943 | 86.9962 | 69.9068 |
| ENSGALG00010000344 | LOC112531461 | 112531461 | 44.723 | 112.657 | 47.1532 | 49.9036 | 54.0657 | 51.4365 | 63.367 | 56.9174 |
| ENSGALG00010003701 | SARAF | 422624 | 43.1136 | 31.826 | 18.9341 | 28.3046 | 20.8244 | 19.2665 | 21.8879 | 18.6181 |
| ENSGALG00010025832 | ENKD1 | 415643 | 39.1753 | 49.1598 | 25.4228 | 29.0923 | 29.1215 | 25.4517 | 27.6831 | 23.7944 |
| ENSGALG00010008602 | NA | NA | 38.9502 | 61.2086 | 0.218648 | 0.331997 | 0.647599 | 0.208426 | 0.460599 | 0.281215 |
| ENSGALG00010000330 | NA | NA | 38.8728 | 104.817 | 53.4995 | 55.1002 | 46.3828 | 44.3501 | 74.999 | 63.8499 |
| ENSGALG00010001672 | NA | NA | 36.7091 | 30.2989 | 34.8947 | 40.4623 | 34.6547 | 22.9784 | 33.7896 | 20.1407 |
| ENSGALG00010000771 | NA | NA | 36.5502 | 4.5007 | 77.9919 | 84.7498 | 164.676 | 197.405 | 159.578 | 170.013 |
| ENSGALG00010024210 | NXPH3 | 429523 | 34.0025 | 17.681 | 3.03819 | 1.70581 | 3.06057 | 2.90961 | 8.78164 | 8.17863 |
| ENSGALG00010000176 | NA | NA | 33.8201 | 72.1251 | 20.7223 | 7.16095 | 15.5678 | 16.0333 | 13.617 | 12.2161 |
| ENSGALG00010010930 | FAM163A | 771080 | 33.5846 | 47.3221 | 3.72132 | 6.11039 | 5.19055 | 5.56023 | 7.54755 | 8.89338 |
| ENSGALG00010016761 | GPR160 | 424992 | 32.694 | 28.9849 | 13.4919 | 22.1153 | 11.4984 | 11.5731 | 14.7278 | 13.1144 |
| ENSGALG00010017612 | CLDN11 | 424990 | 32.4547 | 31.1074 | 4.5751 | 7.15036 | 7.27155 | 9.37594 | 12.673 | 16.4892 |
| ENSGALG00010007160 | LOC121113327 | 121113327 | 31.1515 | 32.5582 | 9.24737 | 8.53165 | 8.62861 | 8.34964 | 11.965 | 5.56036 |
| ENSGALG00010002403 | ZNFY3 | 100858774 | 30.981 | 15.6437 | 15.1113 | 17.0505 | 16.828 | 14.8666 | 8.73423 | 10.6882 |
| ENSGALG00010027290 | CSPG5 | 395466 | 30.6357 | 51.0336 | 20.7165 | 26.018 | 18.6644 | 13.5065 | 20.4831 | 17.4001 |
| ENSGALG00010025263 | SAMD14 | 100857537 | 30.0209 | 23.5526 | 3.27243 | 2.1487 | 7.36752 | 5.24401 | 8.85613 | 7.23465 |
| ENSGALG00010000585 | SPTBN2 | 101750255 | 29.5834 | 25.6189 | 19.2743 | 18.6626 | 22.239 | 15.5692 | 16.093 | 10.9129 |
| ENSGALG00010019346 | POU3F4 | 100857422 | 29.5372 | 51.2123 | 10.9178 | 11.8058 | 15.3751 | 6.26388 | 11.6358 | 9.5538 |
| ENSGALG00010006840 | GINS2 | 415824 | 29.0406 | 30.8213 | 63.4817 | 57.8791 | 68.8489 | 48.1587 | 49.488 | 41.3715 |
| ENSGALG00010020921 | LDB1 | 395952 | 28.3121 | 42.5592 | 25.254 | 26.8288 | 30.2659 | 20.0948 | 25.5182 | 19.0853 |
| ENSGALG00010021837 | NA | NA | 27.4698 | 23.9843 | 23.7375 | 21.0641 | 21.1861 | 15.472 | 17.3393 | 15.8795 |
| ENSGALG00010017637 | NA | NA | 27.0255 | 46.526 | 33.3281 | 35.2813 | 34.4102 | 32.3515 | 50.0356 | 27.5301 |
| ENSGALG00010006933 | NA | NA | 26.6572 | 60.8917 | 14.4629 | 22.2925 | 24.2825 | 21.6259 | 16.1429 | 15.4976 |
| ENSGALG00010005220 | ANGEL1 | 423370 | 26.3078 | 29.2495 | 64.7604 | 74.1046 | 77.0926 | 70.0288 | 35.9018 | 52.8613 |
| ENSGALG00010008933 | NA | NA | 25.4556 | 29.1244 | 3.04055 | 7.34057 | 12.5965 | 8.41076 | 8.33158 | 18.0687 |
| ENSGALG00010000543 | NA | NA | 25.2425 | 43.0612 | 7.39006 | 6.31404 | 4.12528 | 4.29242 | 7.71126 | 9.07134 |
| ENSGALG00010003108 | PPP1R3B | 422630 | 25.2288 | 38.6995 | 8.33588 | 8.05587 | 12.4129 | 9.69925 | 18.9378 | 13.3235 |
| ENSGALG00010013388 | NA | NA | 24.4279 | 22.087 | 55.8041 | 51.8001 | 54.6491 | 39.8487 | 40.7121 | 45.4606 |
| ENSGALG00010022223 | DOK4 | 430610 | 24.3129 | 30.6833 | 7.03966 | 12.7133 | 12.0166 | 10.2618 | 14.4184 | 10.6085 |
| ENSGALG00010018520 | LOC421255 | 421255 | 24.2947 | 24.7719 | 48.1509 | 43.2433 | 46.8031 | 30.934 | 40.6293 | 33.3625 |
| ENSGALG00010017122 | NCBP2 | 424889 | 24.2585 | 27.4615 | 17.165 | 19.6533 | 18.4674 | 16.4132 | 17.9829 | 13.6513 |
| ENSGALG00010000736 | NA | NA | 23.9784 | 3.27059 | 49.0381 | 52.9852 | 103.487 | 122.768 | 99.1482 | 105.556 |
| ENSGALG00010000738 | NA | NA | 23.3317 | 3.16276 | 46.0892 | 49.837 | 98.5818 | 117.499 | 95.5028 | 101.279 |
| ENSGALG00010000569 | SIPA1 | 101749913 | 23.2819 | 23.1311 | 63.8056 | 53.924 | 64.1479 | 54.3257 | 45.1164 | 42.0559 |
| ENSGALG00010025031 | ECHDC2 | 424646 | 22.8118 | 19.8908 | 48.4902 | 46.6796 | 51.0645 | 38.9729 | 44.2075 | 35.431 |
| ENSGALG00010015222 | NA | NA | 22.1185 | 53.6886 | 227.651 | 381.551 | 259.661 | 249.344 | 290.353 | 249.727 |
| ENSGALG00010029442 | LOC107049986 | 107049986 | 21.652 | 37.2218 | 103.308 | 92.0184 | 81.9242 | 53.083 | 75.678 | 63.2205 |
| ENSGALG00010019566 | NA | NA | 21.203 | 38.5125 | 95.1587 | 101.533 | 63.3688 | 61.4424 | 68.3515 | 38.0966 |
| ENSGALG00010016891 | LOC107052288 | 107052288 | 20.7654 | 7.81302 | 5.35512 | 5.81221 | 2.03988 | 2.01241 | 1.87337 | 1.9096 |
| ENSGALG00010018598 | CUL9 | 421258 | 20.3768 | 23.7533 | 16.8719 | 17.4196 | 20.0906 | 14.8863 | 15.4533 | 15.0364 |
| ENSGALG00010000700 | NA | NA | 20.0367 | 28.5765 | 87.8506 | 97.5789 | 72.6028 | 50.484 | 85.9538 | 70.6047 |
| ENSGALG00010010888 | GNPAT | 421548 | 20.0162 | 19.8387 | 37.4499 | 30.293 | 39.3017 | 25.9693 | 29.0882 | 26.9934 |
| ENSGALG00010002367 | STK31 | 420621 | 18.8933 | 20.3611 | 41.8534 | 45.9127 | 42.1837 | 36.825 | 26.8958 | 27.1091 |
| ENSGALG00010018672 | PKDCCA | 421390 | 18.6591 | 18.2067 | 13.8236 | 12.4763 | 14.0249 | 10.6991 | 12.4891 | 11.7316 |
| ENSGALG00010002325 | NA | NA | 18.1582 | 21.2773 | 22.6415 | 20.8898 | 23.0793 | 16.4577 | 15.3113 | 13.656 |
| ENSGALG00010000743 | NA | NA | 18.0301 | 20.7206 | 33.4486 | 30.9921 | 41.8054 | 43.6861 | 56.0412 | 41.4199 |
| ENSGALG00010010907 | NA | NA | 17.7175 | 22.3873 | 22.4757 | 18.6237 | 8.7174 | 6.93757 | 8.02641 | 12.7742 |
| ENSGALG00010001350 | LOC107049524 | 107049524 | 17.124 | 6.9259 | 47.4225 | 43.9694 | 46.1847 | 39.9557 | 36.7142 | 35.8713 |
| ENSGALG00010004429 | LOC112531814 | 112531814 | 16.7427 | 7.63336 | 1.55306 | 2.16396 | 2.58744 | 1.09727 | 2.00147 | 0.704987 |

| | | | | | | | | | | |
|---|---|---|---|---|---|---|---|---|---|---|
| ENSGALG00010005905 | CITED2 | 395399 | 16.4571 | 17.367 | 8.60817 | 13.0781 | 11.1129 | 9.38819 | 5.82021 | 5.81776 |
| ENSGALG00010019264 | HK2 | 374044 | 15.4694 | 20.5145 | 7.76757 | 9.23159 | 11.9596 | 7.92414 | 10.7393 | 9.79643 |
| ENSGALG00010012476 | NA | NA | 14.8964 | 21.3677 | 256.778 | 98.8142 | 133.701 | 59.2152 | 47.2549 | 41.295 |
| ENSGALG00010020005 | EOGT | 426961 | 14.8184 | 19.0454 | 5.31977 | 6.51275 | 12.8202 | 9.94822 | 6.24111 | 6.23981 |
| ENSGALG00010002473 | LOC107049124 | 107049124 | 14.7914 | 18.1778 | 2.53132 | 3.60537 | 6.77135 | 10.5219 | 0.931293 | 1.18304 |
| ENSGALG00010025780 | NA | NA | 13.6454 | 126.646 | 34.7336 | 34.6963 | 9.17937 | 10.7314 | 33.126 | 30.4234 |
| ENSGALG00010001208 | MOV10L1 | 417738 | 12.9076 | 12.5063 | 27.7209 | 25.7207 | 26.0975 | 18.4583 | 20.205 | 24.2723 |
| ENSGALG00010007976 | NA | NA | 12.8051 | 15.3559 | 26.3271 | 23.4081 | 27.1613 | 19.846 | 24.2957 | 18.9988 |
| ENSGALG00010012478 | NA | NA | 12.5881 | 8.65466 | 7.86881 | 5.16155 | 8.04059 | 0.900108 | 87.9204 | 72.5637 |
| ENSGALG00010005363 | RDX | 395511 | 12.3941 | 13.3457 | 26.9772 | 23.0464 | 24.5576 | 20.4142 | 19.8803 | 16.2189 |
| ENSGALG00010028676 | LOC112530443 | 112530443 | 12.2938 | 5.93796 | 30.3903 | 28.4209 | 28.448 | 28.5085 | 46.3171 | 23.9404 |
| ENSGALG00010007596 | LOC121109412 | 121109412 | 12.238 | 2.15159 | 1.9318 | 1.23672 | 1.50774 | 0.995395 | 1.02654 | 0.819242 |
| ENSGALG00010001272 | NA | NA | 11.8955 | 7.15929 | 2.45654 | 1.3835 | 3.62139 | 1.78819 | 1.58069 | 3.6535 |
| ENSGALG00010012555 | BHMT2 | 416371 | 11.783 | 6.05316 | 3.13671 | 3.29867 | 3.51257 | 2.67997 | 2.12643 | 3.86494 |
| ENSGALG00010001351 | CAPN1 | 693249 | 11.2742 | 7.91161 | 42.6763 | 36.3664 | 39.0904 | 27.2955 | 25.7758 | 21.845 |
| ENSGALG00010010945 | PI4K2B | 422807 | 10.9839 | 10.5671 | 19.2907 | 19.1178 | 20.0344 | 16.9118 | 18.7217 | 16.3317 |
| ENSGALG00010012654 | FAM46B | 430080 | 10.9784 | 13.4377 | 5.71347 | 4.66513 | 4.40464 | 3.18011 | 5.61529 | 5.6079 |
| ENSGALG00010020618 | NA | NA | 10.9305 | 11.0055 | 12.983 | 12.7129 | 15.9116 | 10.8357 | 9.44349 | 10.0826 |
| ENSGALG00010001176 | IL12A | 407090 | 10.8483 | 25.0237 | 2.80952 | 4.61641 | 8.46992 | 3.78769 | 3.67791 | 5.03304 |
| ENSGALG00010000189 | LOC121108943 | 121108943 | 10.8047 | 12.9445 | 0.265938 | 0.231513 | 3.24255 | 2.22408 | 0.124493 | 0.232585 |
| ENSGALG00010011694 | NA | NA | 10.6271 | 10.8958 | 3.36276 | 9.4849 | 5.18343 | 9.8037 | 4.76304 | 0.0605839 |
| ENSGALG00010025767 | NA | NA | 10.5577 | 15.0892 | 10.3565 | 10.7556 | 6.06288 | 7.1398 | 7.72067 | 9.62897 |
| ENSGALG00010000773 | NA | NA | 10.3923 | 2.1707 | 23.787 | 27.0302 | 42.0069 | 59.4154 | 59.1402 | 56.0045 |
| ENSGALG00010009458 | NA | NA | 10.2932 | 11.7823 | 17.8123 | 14.7153 | 12.6078 | 10.7266 | 13.875 | 9.34835 |
| ENSGALG00010021054 | FEZF2 | 416074 | 10.2189 | 16.0302 | 69.8582 | 45.6663 | 47.0524 | 39.5408 | 49.5044 | 22.655 |
| ENSGALG00010014492 | CPLX2 | 100857281 | 10.1764 | 13.7818 | 6.72145 | 5.82567 | 8.61888 | 5.75272 | 24.5589 | 16.5995 |
| ENSGALG00010024629 | ESAM | 419707 | 9.93899 | 9.32779 | 1.20416 | 1.11042 | 2.91955 | 1.84778 | 2.68514 | 2.96903 |
| ENSGALG00010003431 | PM20D2 | 421816 | 9.90584 | 11.7516 | 23.9523 | 29.5884 | 26.6487 | 23.1095 | 20.2688 | 17.4982 |
| ENSGALG00010013073 | NA | NA | 9.87801 | 4.74687 | 29.8477 | 11.6942 | 25.021 | 8.72729 | 6.95007 | 14.52 |
| ENSGALG00010013435 | SDC3 | 396343 | 9.67586 | 9.57736 | 1.31618 | 1.49995 | 1.73828 | 1.64791 | 5.40006 | 4.72014 |
| ENSGALG00010024909 | SLC39A5 | 100858866 | 9.46373 | 10.7809 | 7.26125 | 7.98398 | 7.23065 | 5.06003 | 6.22403 | 4.17037 |
| ENSGALG00010023736 | LOC107050879 | 107050879 | 9.143 | 6.48861 | 1.95277 | 1.61678 | 2.81275 | 1.91518 | 2.51867 | 1.96925 |
| ENSGALG00010020389 | ACTN1 | 373918 | 8.9952 | 4.9141 | 15.1788 | 16.0484 | 13.14 | 12.6104 | 11.6946 | 14.2242 |
| ENSGALG00010012091 | LOC121108229 | 121108229 | 8.92639 | 7.62802 | 7.26578 | 6.27039 | 7.49763 | 5.59229 | 5.00838 | 3.58318 |
| ENSGALG00010011822 | NA | NA | 8.88208 | 11.6926 | 0.721782 | 2.19191 | 0.534449 | 0.91738 | 0.878504 | 0.495104 |
| ENSGALG00010001313 | EMILIN2 | 421058 | 8.85726 | 10.1721 | 18.8838 | 24.6062 | 5.73005 | 3.4408 | 5.70582 | 5.85434 |
| ENSGALG00010027167 | ASIC1L | 107055878 | 8.84454 | 22.3648 | 5.70321 | 2.69415 | 2.26079 | 1.05436 | 4.59535 | 5.39395 |
| ENSGALG00010013170 | COL3A1 | 396340 | 8.79022 | 17.337 | 37.1579 | 33.9633 | 37.8299 | 32.9702 | 28.4139 | 36.3586 |
| ENSGALG00010000767 | NA | NA | 8.75869 | 2.15568 | 39.5112 | 33.7382 | 56.4938 | 41.7374 | 32.2908 | 35.9611 |
| ENSGALG00010000830 | NA | NA | 8.72521 | 9.81881 | 0.0403789 | 0.496623 | 1.20503 | 3.10842 | 1.70527 | 0.719647 |
| ENSGALG00010011166 | SOX7 | 771337 | 8.64078 | 18.6788 | 2.3593 | 3.32649 | 8.42285 | 3.2771 | 3.97604 | 3.94474 |
| ENSGALG00010028386 | NCAN | 395493 | 8.38599 | 53.7883 | 9.81291 | 15.0904 | 12.0449 | 7.71301 | 11.5094 | 7.27319 |
| ENSGALG00010027277 | KCND3 | 374228 | 8.30983 | 18.0036 | 16.1036 | 27.9245 | 1.4663 | 1.87571 | 4.02567 | 4.13605 |
| ENSGALG00010010753 | LOC107051636 | 107051636 | 8.14358 | 3.38414 | 1.50128 | 1.55509 | 1.85019 | 0.929229 | 1.53363 | 1.11091 |
| ENSGALG00010001756 | ARHGAP33L6 | 107050670 | 7.84906 | 10.6751 | 16.6546 | 12.4613 | 32.5093 | 29.9683 | 25.5393 | 22.4959 |
| ENSGALG00010001707 | NA | NA | 7.84507 | 7.69789 | 0.384196 | 0.228858 | 2.7901 | 3.74402 | 0.759532 | 1.74467 |
| ENSGALG00010023104 | EFCAB4B | 423106 | 7.65882 | 8.45076 | 4.7907 | 4.21897 | 5.2766 | 4.22808 | 4.77728 | 4.45112 |
| ENSGALG00010027530 | PHF19 | 772213 | 7.60153 | 10.2382 | 30.5377 | 26.5295 | 20.7065 | 13.9306 | 17.6206 | 15.5573 |
| ENSGALG00010000123 | NA | NA | 7.40008 | 6.12692 | 36.2968 | 19.6303 | 17.5666 | 17.2533 | 18.3153 | 17.339 |
| ENSGALG00010000659 | LOC121109782 | 121109782 | 7.06182 | 5.45741 | 12.6231 | 32.2535 | 25.2277 | 16.7616 | 29.2966 | 28.5651 |
| ENSGALG00010005037 | THNSL1 | 100859434 | 6.94251 | 10.0178 | 26.9839 | 28.5423 | 20.7343 | 15.7988 | 14.7897 | 13.8536 |
| ENSGALG00010002557 | NA | NA | 6.62304 | 9.57756 | 1.34798 | 0.598291 | 1.15168 | 1.34426 | 2.41733 | 2.14713 |
| ENSGALG00010014283 | NA | NA | 6.55513 | 7.88129 | 0 | 0.114188 | 0.0348029 | 0 | 0.0528069 | 0 |
| ENSGALG00010000788 | GATD3AL1 | 418811 | 6.30959 | 4.20879 | 15.1724 | 11.8864 | 16.8442 | 10.9796 | 14.4312 | 11.4121 |
| ENSGALG00010012473 | NA | NA | 6.30503 | 14.2001 | 149.11 | 56.4864 | 83.242 | 30.6518 | 72.0449 | 50.6139 |
| ENSGALG00010001429 | LOC112533497 | 112533497 | 6.19733 | 8.20604 | 1.09116 | 0.662728 | 1.39839 | 0.896122 | 1.13162 | 1.48976 |
| ENSGALG00010002338 | CD99 | 418662 | 6.18414 | 5.63761 | 17.8772 | 15.7493 | 12.4518 | 8.77441 | 10.0881 | 8.38721 |
| ENSGALG00010003001 | MAP6 | 395753 | 6.0978 | 11.4561 | 1.48804 | 2.42467 | 3.96202 | 3.50188 | 4.41084 | 3.75129 |
| ENSGALG00010008603 | NA | NA | 6.07041 | 10.4996 | 1.33131 | 1.5161 | 4.34359 | 1.9036 | 0.420676 | 0.25684 |
| ENSGALG00010007892 | LOC121109261 | 121109261 | 5.96844 | 8.6076 | 1.30679 | 1.37729 | 1.10992 | 1.4362 | 0.842046 | 0.889794 |
| ENSGALG00010016588 | ATP11B | 424964 | 5.94965 | 6.31134 | 12.9594 | 12.8045 | 12.1769 | 11.2359 | 9.32957 | 10.8742 |
| ENSGALG00010010076 | NA | NA | 5.76693 | 4.53705 | 153.793 | 135.937 | 49.1741 | 32.4691 | 112.858 | 97.7911 |
| ENSGALG00010005530 | NA | NA | 5.59705 | 6.22595 | 43.3111 | 39.5472 | 41.2572 | 22.3304 | 9.99064 | 12.6909 |
| ENSGALG00010005837 | LOC107052005 | 107052005 | 5.54195 | 6.11438 | 1.6127 | 1.3989 | 1.53395 | 1.55038 | 1.29385 | 1.23788 |
| ENSGALG00010006304 | NA | NA | 5.52581 | 10.8638 | 7.14295 | 11.4228 | 13.9148 | 9.65446 | 8.94038 | 9.21567 |
| ENSGALG00010028335 | NA | NA | 5.51856 | 11.1572 | 102.486 | 87.6275 | 29.9503 | 21.0087 | 13.8035 | 11.1099 |
| ENSGALG00010023766 | NA | NA | 5.45386 | 21.9375 | 6.9827 | 9.23114 | 18.7356 | 25.7856 | 19.9059 | 18.9036 |
| ENSGALG00010017820 | FLRT2 | 428901 | 5.44718 | 5.48702 | 0.581541 | 0.816056 | 1.07141 | 1.02936 | 1.10546 | 0.657202 |
| ENSGALG00010000888 | NA | NA | 5.17276 | 2.6818 | 14.4587 | 25.4205 | 35.8933 | 24.3155 | 33.1728 | 30.3017 |
| ENSGALG00010011278 | TDH | 422034 | 5.10114 | 3.25301 | 46.7379 | 39.9105 | 26.6512 | 17.878 | 13.1115 | 6.77534 |
| ENSGALG00010000719 | NA | NA | 5.07581 | 0.223797 | 0.787451 | 0.478268 | 14.5768 | 13.5114 | 15.04 | 18.4326 |
| ENSGALG00010016457 | SMOX | 422934 | 4.9371 | 6.29149 | 1.08172 | 1.16902 | 1.78432 | 1.33807 | 3.5989 | 3.15874 |
| ENSGALG00010003723 | NA | NA | 4.8868 | 6.77219 | 1.76784 | 2.20137 | 7.39069 | 6.82735 | 4.71601 | 5.65769 |
| ENSGALG00010014058 | NA | NA | 4.75256 | 3.98526 | 14.831 | 13.4311 | 13.6103 | 12.7744 | 7.28109 | 9.93391 |
| ENSGALG00010011469 | CPS1 | 428994 | 4.6917 | 4.49444 | 7.66997 | 8.11958 | 6.2052 | 4.89214 | 4.70239 | 4.53128 |
| ENSGALG00010001201 | TUBB6 | 421037 | 4.62786 | 5.18877 | 15.6479 | 11.4132 | 15.8492 | 8.14824 | 12.3699 | 10.1574 |
| ENSGALG00010003728 | NA | NA | 4.62271 | 6.2735 | 2.06489 | 2.39787 | 7.49406 | 6.87534 | 4.81954 | 5.83143 |
| ENSGALG00010028301 | FBN3 | 395079 | 4.49255 | 6.76914 | 10.06 | 10.8957 | 14.2779 | 10.6898 | 9.46272 | 8.3362 |
| ENSGALG00010011136 | NA | NA | 4.38572 | 16.6851 | 84.0446 | 54.8729 | 63.4789 | 54.7031 | 28.8843 | 20.877 |
| ENSGALG00010010273 | NA | NA | 4.30882 | 3.3899 | 114.964 | 101.618 | 36.7617 | 24.2596 | 84.3701 | 73.0655 |
| ENSGALG00010002436 | MHCY4 | 427737 | 4.22189 | 3.56584 | 2.343 | 1.59025 | 2.32909 | 2.59974 | 0.473827 | 0.171766 |
| ENSGALG00010001611 | MAPK12 | 769763 | 4.1383 | 3.18916 | 5.5047 | 5.7003 | 6.94565 | 5.628 | 5.6583 | 5.84683 |
| ENSGALG00010002061 | NA | NA | 4.09455 | 8.46188 | 0.00956133 | 0.0627175 | 0.00637178 | 0.0196869 | 0.0370606 | 0.0206595 |
| ENSGALG00010002873 | MRAP2 | 421836 | 4.0563 | 1.58664 | 0.210103 | 0.252938 | 0.21669 | 0.309003 | 0.512149 | 0.486403 |
| ENSGALG00010011588 | COLGALT1 | 424447 | 4.05092 | 4.66189 | 13.694 | 12.35 | 13.2638 | 9.45906 | 11.6657 | 8.91952 |
| ENSGALG00010017143 | FAM149A | 422721 | 4.03189 | 4.10143 | 0.845548 | 1.26925 | 1.18244 | 1.10253 | 1.40651 | 1.65661 |
| ENSGALG00010008656 | CCDC6 | 423666 | 3.94536 | 5.16121 | 12.3431 | 10.8551 | 10.0694 | 7.10123 | 7.99392 | 7.24827 |
| ENSGALG00010008118 | NA | NA | 3.88841 | 4.39298 | 54.9066 | 66.5225 | 152.836 | 120.852 | 71.8767 | 57.9136 |
| ENSGALG00010019079 | CAPG | 431244 | 3.87571 | 14.492 | 2.56025 | 1.70618 | 4.02848 | 1.95243 | 2.41702 | 1.82937 |
| ENSGALG00010003205 | TNX | 396106 | 3.82515 | 2.63965 | 11.0539 | 11.4292 | 14.4762 | 8.27503 | 8.41047 | 9.35459 |
| ENSGALG00010003732 | NA | NA | 3.76843 | 5.35335 | 5.18621 | 6.262 | 7.48064 | 6.16977 | 4.07872 | 4.94131 |
| ENSGALG00010024096 | MYO1D | 419963 | 3.76237 | 3.87361 | 2.58463 | 2.20797 | 2.97366 | 2.14658 | 1.86804 | 1.34962 |
| ENSGALG00010024643 | ETV7 | 419813 | 3.74223 | 2.62555 | 10.2303 | 9.75327 | 11.7299 | 9.45645 | 7.78443 | 7.03337 |
| ENSGALG00010009307 | DIRAS2 | 425241 | 3.62074 | 1.80526 | 0.326216 | 0.221495 | 2.48328 | 1.42046 | 1.34457 | 1.68379 |
| ENSGALG00010025111 | NA | NA | 3.62026 | 6.21006 | 5.46268 | 4.36451 | 5.22464 | 5.25686 | 3.28787 | 5.52029 |
| ENSGALG00010020023 | FOSL2 | 421416 | 3.30413 | 3.35483 | 13.829 | 11.7844 | 8.88484 | 6.31881 | 6.90793 | 5.05873 |
| ENSGALG00010017006 | DNAH3 | 427004 | 3.16438 | 3.70035 | 4.7657 | 8.96377 | 6.19448 | 8.0117 | 8.31448 | 3.94583 |
| ENSGALG00010025049 | NA | NA | 3.15623 | 1.4502 | 9.44942 | 7.2314 | 12.1746 | 7.24212 | 4.29968 | 8.3129 |
| ENSGALG00010000769 | NA | NA | 3.13172 | 0.261876 | 13.8216 | 11.137 | 29.6112 | 21.7832 | 23.5 | 23.1333 |
| ENSGALG00010017135 | LOC100859848 | 100859848 | 3.11883 | 5.51123 | 24.8067 | 18.5803 | 20.7329 | 15.5999 | 15.1442 | 15.8335 |
| ENSGALG00010020100 | MOCS1 | 395270 | 3.09191 | 3.27261 | 7.47929 | 6.72725 | 7.59074 | 7.86608 | 8.79408 | 9.24504 |
| ENSGALG00010013866 | KCTD8 | 770600 | 3.07337 | 4.85743 | 1.08055 | 2.05244 | 0.993525 | 0.87164 | 0.790251 | 0.979383 |
| ENSGALG00010005240 | VIP | 396323 | 3.0719 | 1.01938 | 0.367877 | 0.442879 | 0.715043 | 0.330638 | 0.309984 | 0.486663 |
| ENSGALG00010001276 | LOC121108470 | 121108470 | 3.04758 | 0.986914 | 0.386438 | 0.325862 | 0.379303 | 0.236442 | 0.249898 | 0.11651 |
| ENSGALG00010020960 | CLEC19A | 771758 | 3.02281 | 13.1945 | 1.89861 | 1.2728 | 0.795756 | 0.573684 | 1.41871 | 1.21634 |
| ENSGALG00010029392 | RTN4RL1 | 427828 | 2.9769 | 3.78653 | 0.318575 | 0.234952 | 0.95199 | 0.555301 | 1.15046 | 1.94624 |
| ENSGALG00010014029 | NA | NA | 2.92921 | 2.45099 | 8.73052 | 8.08481 | 8.66213 | 7.52662 | 4.30181 | 6.70849 |
| ENSGALG00010025432 | NA | NA | 2.83292 | 6.68836 | 96.4454 | 87.5888 | 27.0841 | 18.6423 | 19.2665 | 14.125 |
| ENSGALG00010027945 | DCST2 | 121107585 | 2.82647 | 3.29001 | 15.4642 | 21.4923 | 0.898296 | 1.20381 | 36.5464 | 28.5518 |
| ENSGALG00010015243 | NA | NA | 2.81991 | 12.7788 | 4.78126 | 6.17277 | 15.9314 | 21.4944 | 16.8877 | 16.8806 |
| ENSGALG00010000480 | NA | NA | 2.75465 | 6.37942 | 0 | 0.107067 | 0.217549 | 0.0448107 | 0.132036 | 0.12092 |
| ENSGALG00010008318 | ADPRH | 418452 | 2.71739 | 9.0448 | 1.73704 | 2.52449 | 1.94079 | 3.25251 | 1.66407 | 1.42024 |
| ENSGALG00010026987 | NA | NA | 2.6173 | 2.5695 | 6.92381 | 6.1191 | 6.94468 | 5.2148 | 6.22898 | 4.84245 |
| ENSGALG00010024725 | NA | NA | 2.55268 | 0.691002 | 10.806 | 7.13745 | 7.12626 | 7.33932 | 5.23566 | 5.1597 |

| | | | | | | | | | | |
|---|---|---|---|---|---|---|---|---|---|---|
| ENSGALG00010027079 | PM20D1 | 419838 | 2.54901 | 4.07534 | 10.5797 | 9.01461 | 9.92989 | 6.50947 | 6.50806 | 5.64527 |
| ENSGALG00010016760 | DLK1 | 423459 | 2.47931 | 1.16683 | 0.0626812 | 0.133245 | 0.348095 | 0.430204 | 0.492959 | 0.580445 |
| ENSGALG00010003827 | RRH | 771805 | 2.4157 | 3.63118 | 1.28211 | 1.82726 | 2.03734 | 1.79331 | 2.09116 | 1.2 |
| ENSGALG00010002035 | NA | NA | 2.35265 | 2.92104 | 1.07062 | 4.29169 | 2.37825 | 0.61234 | 0.721712 | 1.15666 |
| ENSGALG00010011109 | NA | NA | 2.35112 | 14.233 | 78.9231 | 60.3932 | 56.2677 | 51.1302 | 22.8375 | 22.1831 |
| ENSGALG00010019947 | LGALSL2 | 423277 | 2.34332 | 1.78405 | 7.85834 | 7.4417 | 9.39954 | 7.092 | 3.70918 | 4.67673 |
| ENSGALG00010000598 | NA | NA | 2.33333 | 0.341138 | 0.192361 | 0.231328 | 12.5243 | 5.68583 | 4.1365 | 10.3643 |
| ENSGALG00010016190 | THNSL2 | 422927 | 2.30644 | 1.24769 | 21.2568 | 16.8603 | 12.2041 | 6.04641 | 5.52762 | 6.59067 |
| ENSGALG00010003094 | LOC107053759 | 107053759 | 2.19809 | 2.82805 | 6.47755 | 8.74123 | 8.23609 | 5.89989 | 7.22879 | 4.55642 |
| ENSGALG00010003193 | ACOT9 | 418600 | 2.18326 | 2.82246 | 6.75349 | 8.51704 | 9.34178 | 5.82458 | 5.51027 | 5.48259 |
| ENSGALG00010002208 | NA | NA | 2.09318 | 2.55023 | 0.943341 | 0.859423 | 0.945535 | 0.381627 | 0.814275 | 0.648068 |
| ENSGALG00010009702 | NA | NA | 2.07149 | 1.44916 | 2.60355 | 2.46142 | 2.1918 | 1.5778 | 2.08103 | 1.61517 |
| ENSGALG00010016428 | ADRA1D | 422933 | 2.05783 | 1.48375 | 0.0267729 | 0.0379419 | 0.307275 | 0.173544 | 0.308315 | 0.702453 |
| ENSGALG00010027188 | KCNH2 | 100858122 | 1.96705 | 4.43286 | 0.456362 | 0.311054 | 0.709625 | 0.301617 | 0.783331 | 0.594776 |
| ENSGALG00010012283 | LOC101751171 | 101751171 | 1.85867 | 0.692753 | 0.138408 | 0.0723882 | 0.216358 | 0.240418 | 0.190058 | 0.140433 |
| ENSGALG00010000825 | NA | NA | 1.83815 | 11.1837 | 1.35011 | 2.03114 | 2.45583 | 2.4682 | 1.47827 | 1.77583 |
| ENSGALG00010000553 | NA | NA | 1.80981 | 1.53878 | 4.48654 | 4.66638 | 3.43032 | 5.27751 | 5.59042 | 5.21784 |
| ENSGALG00010011288 | NA | NA | 1.80273 | 4.50409 | 17.0911 | 13.5607 | 9.49142 | 6.5405 | 6.99126 | 6.21083 |
| ENSGALG00010024289 | NA | NA | 1.79324 | 4.01574 | 9.48282 | 9.83245 | 9.25958 | 9.81014 | 6.56856 | 5.41303 |
| ENSGALG00010019738 | ADRA2A | 428980 | 1.78463 | 1.93778 | 9.84863 | 6.26696 | 5.99106 | 4.19434 | 6.07724 | 4.34959 |
| ENSGALG00010014147 | HTR1D | 769107 | 1.72076 | 1.49515 | 4.45146 | 3.81599 | 5.44359 | 5.15531 | 2.64926 | 3.3097 |
| ENSGALG00010023306 | NA | NA | 1.7186 | 2.83077 | 1.79684 | 1.70258 | 1.83261 | 2.95462 | 5.6171 | 3.99314 |
| ENSGALG00010007103 | NA | NA | 1.66769 | 2.4219 | 0.128873 | 0.415821 | 0.489051 | 0.485246 | 0.514304 | 0.283155 |
| ENSGALG00010020532 | SIX3 | 378786 | 1.6371 | 9.15258 | 45.9529 | 29.5554 | 28.585 | 27.3479 | 59.4088 | 5.93201 |
| ENSGALG00010008003 | NA | NA | 1.52584 | 3.84816 | 28.7005 | 29.3103 | 10.4115 | 11.1199 | 15.6025 | 3.84015 |
| ENSGALG00010000601 | LOC121107771 | 121107771 | 1.48584 | 0.244834 | 0.430737 | 0.303471 | 9.64482 | 3.98992 | 2.08093 | 4.62103 |
| ENSGALG00010016416 | CYP46A1 | 423450 | 1.40014 | 1.74928 | 5.1674 | 4.41167 | 4.7769 | 3.12676 | 3.52739 | 2.88132 |
| ENSGALG00010021710 | KEL | 418306 | 1.38574 | 1.66368 | 0.0344596 | 0 | 0.086116 | 0.00985452 | 0 | 0 |
| ENSGALG00010009398 | WTIP | 769595 | 1.3194 | 1.47198 | 4.29252 | 4.33072 | 3.52747 | 2.61012 | 2.24915 | 1.86996 |
| ENSGALG00010022843 | NA | NA | 1.30521 | 3.93243 | 0.494969 | 0.184475 | 0.274878 | 0.463248 | 0.37916 | 0.156257 |
| ENSGALG00010023498 | DISP2 | 428836 | 1.2889 | 1.03467 | 3.13181 | 2.75532 | 3.34156 | 2.95706 | 2.2475 | 3.15816 |
| ENSGALG00010020471 | NA | NA | 1.25911 | 0.67852 | 5.0932 | 5.07513 | 3.80076 | 5.00679 | 4.69403 | 1.84236 |
| ENSGALG00010019594 | TMX1 | 423580 | 1.23784 | 2.03413 | 5.28513 | 5.96212 | 6.36696 | 4.37787 | 4.23004 | 4.07848 |
| ENSGALG00010028141 | CISH | 395335 | 1.2367 | 1.7657 | 5.37492 | 6.95631 | 4.31936 | 4.43042 | 5.67466 | 4.06238 |
| ENSGALG00010019059 | BRWD3 | 422281 | 1.22546 | 2.39401 | 5.03767 | 5.71571 | 4.03235 | 4.62838 | 4.10808 | 4.66269 |
| ENSGALG00010010761 | MEI1 | 417985 | 1.21712 | 1.7481 | 5.6071 | 4.50191 | 4.82508 | 4.14339 | 3.31005 | 3.32947 |
| ENSGALG00010001712 | ATP6AP1 | 427854 | 1.21406 | 1.46692 | 3.66383 | 1.95519 | 3.59888 | 2.0737 | 2.56623 | 2.43229 |
| ENSGALG00010001209 | CDH7 | 374007 | 1.17532 | 2.4591 | 0.845084 | 1.29394 | 0.918942 | 0.768672 | 0.585754 | 0.557477 |
| ENSGALG00010014282 | NA | NA | 1.11224 | 1.04617 | 0 | 0 | 0 | 0 | 0 | 0 |
| ENSGALG00010009749 | NDRG1 | 420321 | 0.859277 | 0.948569 | 3.12607 | 1.87761 | 4.44442 | 5.99181 | 7.36754 | 5.09775 |
| ENSGALG00010012985 | ARNTL2 | 373925 | 0.853229 | 0.852787 | 2.51379 | 2.15039 | 2.61507 | 1.91701 | 1.94168 | 1.72584 |
| ENSGALG00010007404 | FOLH1 | 419011 | 0.82141 | 0.673122 | 1.92239 | 1.87365 | 2.34123 | 1.80617 | 1.54669 | 1.54746 |
| ENSGALG00010012260 | LOC121108245 | 121108245 | 0.816062 | 1.31753 | 4.28691 | 3.7381 | 2.30804 | 1.72844 | 2.04241 | 1.50502 |
| ENSGALG00010016612 | GRIA1 | 414893 | 0.806731 | 2.19833 | 0.14934 | 0.232465 | 0.456967 | 0.185537 | 0.259177 | 0.238121 |
| ENSGALG00010001114 | MED12L | 425036 | 0.796144 | 1.02877 | 2.57858 | 2.02359 | 2.14947 | 1.91109 | 1.50666 | 1.74535 |
| ENSGALG00010019270 | NA | NA | 0.792604 | 2.08437 | 0.320615 | 0.21907 | 0 | 0.183374 | 0.20262 | 0.216488 |
| ENSGALG00010002285 | CAV1 | 373996 | 0.787703 | 1.80498 | 8.4615 | 7.72062 | 4.76755 | 3.02331 | 4.33376 | 1.46328 |
| ENSGALG00010000454 | NA | NA | 0.783482 | 1.45481 | 4.99156 | 6.61247 | 10.6921 | 12.1479 | 13.219 | 47.8151 |
| ENSGALG00010015088 | LOC107050476 | 107050476 | 0.742127 | 0.183336 | 0.162325 | 0.144044 | 0.155707 | 0.103693 | 0.0888182 | 0.0715798 |
| ENSGALG00010015661 | NA | NA | 0.72597 | 0.806749 | 0 | 0.00617394 | 10.8632 | 16.9593 | 0.00285516 | 0.00261479 |
| ENSGALG00010006623 | LOC107052071 | 107052071 | 0.709923 | 1.41728 | 0.0187829 | 0 | 0 | 0.0859424 | 0.126616 | 0.304385 |
| ENSGALG00010023212 | RIMBP2 | 374105 | 0.683982 | 0.767895 | 1.61935 | 1.57047 | 1.50849 | 1.40524 | 1.08068 | 1.27257 |
| ENSGALG00010007069 | NA | NA | 0.668675 | 1.32217 | 20.6569 | 34.0134 | 11.6342 | 3.58121 | 3.72284 | 1.716 |
| ENSGALG00010004493 | NA | NA | 0.6587 | 0.7967 | 1.86885 | 1.93017 | 2.46618 | 1.46574 | 1.10895 | 0.922774 |
| ENSGALG00010002126 | MGAM | 425545 | 0.638114 | 0.753111 | 0.13935 | 0.0245717 | 0.132307 | 0.15426 | 0.210221 | 0.154365 |
| ENSGALG00010020629 | TRIM66 | 423056 | 0.62437 | 0.813224 | 0.313751 | 0.292701 | 0.376357 | 0.31583 | 0.251686 | 0.360272 |
| ENSGALG00010019938 | FAM19A4 | 771511 | 0.609429 | 0.806463 | 0.136218 | 0.179876 | 0.141096 | 0.208342 | 0.305654 | 0.282283 |
| ENSGALG00010006909 | NA | NA | 0.59758 | 1.1761 | 0.0344854 | 0.0448823 | 0.0766048 | 0.140884 | 0.0456632 | 0.053224 |
| ENSGALG00010018279 | NA | NA | 0.595125 | 0.384847 | 8.85822 | 1.79909 | 14.3507 | 1.87167 | 3.13782 | 1.43682 |
| ENSGALG00010026011 | GJC2 | 420397 | 0.59211 | 0.601958 | 1.62254 | 1.62868 | 0.190645 | 0.281551 | 1.4327 | 1.8869 |
| ENSGALG00010023274 | ZBTB38 | 424783 | 0.586904 | 0.471759 | 1.88522 | 1.35913 | 1.89163 | 1.28589 | 0.84493 | 1.01341 |
| ENSGALG00010029927 | CFAP52 | 417322 | 0.558543 | 0.977856 | 3.58017 | 2.74278 | 2.96942 | 2.22939 | 1.61286 | 1.57873 |
| ENSGALG00010028809 | NA | NA | 0.554414 | 2.72731 | 5.22617 | 6.32098 | 5.05403 | 3.45292 | 4.06206 | 2.29463 |
| ENSGALG00010012384 | NRK | 422380 | 0.537192 | 0.792175 | 3.26636 | 2.48108 | 2.14417 | 1.97919 | 1.57363 | 1.65306 |
| ENSGALG00010005024 | NA | NA | 0.514998 | 5.4915 | 0.0165043 | 0.00751805 | 0 | 0 | 0.0347675 | 0.0191043 |
| ENSGALG00010003505 | NA | NA | 0.510273 | 0.775889 | 6.0563 | 5.93208 | 7.75482 | 7.95078 | 4.30998 | 8.68125 |
| ENSGALG00010026343 | NA | NA | 0.496473 | 1.34086 | 9.25873 | 3.38009 | 6.8885 | 1.33021 | 2.91163 | 2.10244 |
| ENSGALG00010027715 | NINJ1 | 415986 | 0.492635 | 0.56096 | 1.91287 | 1.81486 | 3.62671 | 2.28568 | 2.12517 | 1.59837 |
| ENSGALG00010009191 | NA | NA | 0.489834 | 0.98987 | 2.11299 | 2.94043 | 2.75951 | 3.24043 | 2.32831 | 3.76287 |
| ENSGALG00010000228 | NA | NA | 0.460184 | 0.480714 | 0.0268483 | 0.0366898 | 0.20874 | 0.138202 | 0.101804 | 0.176107 |
| ENSGALG00010004566 | NA | NA | 0.449072 | 0 | 7.6124 | 6.79558 | 5.84472 | 5.72726 | 2.71214 | 5.12532 |
| ENSGALG00010006610 | NA | NA | 0.435491 | 0.426692 | 1.11601 | 0.661103 | 1.03387 | 0.646885 | 0.809657 | 0.892108 |
| ENSGALG00010002297 | NA | NA | 0.405945 | 0.251097 | 0.947095 | 1.19518 | 1.07794 | 0.777121 | 0.775491 | 0.624979 |
| ENSGALG00010015499 | NA | NA | 0.392669 | 0.895363 | 1.66909 | 1.60083 | 1.9315 | 2.0085 | 1.33441 | 2.51708 |
| ENSGALG00010007803 | ME3 | 419019 | 0.388034 | 0.234756 | 0.866395 | 0.873773 | 0.952098 | 0.86928 | 0.992992 | 0.978093 |
| ENSGALG00010014226 | SLC16A7 | 417815 | 0.380481 | 0.401665 | 0.0791993 | 0.0870823 | 0.110715 | 0.0937196 | 0.089748 | 0.168599 |
| ENSGALG00010026821 | NA | NA | 0.379415 | 0.736063 | 4.85608 | 4.22747 | 2.87658 | 3.45634 | 3.36443 | 1.24913 |
| ENSGALG00010000688 | LOC121108155 | 121108155 | 0.376464 | 0.1787 | 4.37407 | 4.53704 | 5.71346 | 3.9246 | 4.10037 | 4.33531 |
| ENSGALG00010003018 | MOGAT2 | 426437 | 0.353531 | 2.01598 | 0.03047 | 0.0277595 | 0.108296 | 0.160331 | 0.374854 | 0.0987563 |
| ENSGALG00010003360 | NA | NA | 0.351864 | 0.183781 | 1.38568 | 1.82349 | 2.6335 | 2.88836 | 1.40114 | 1.33071 |
| ENSGALG00010019138 | NA | NA | 0.340939 | 0.515908 | 2.07903 | 1.58312 | 0.792694 | 1.72153 | 1.45116 | 1.48464 |
| ENSGALG00010014867 | TP63 | 374269 | 0.331941 | 0.35073 | 1.42001 | 1.20504 | 1.30005 | 1.20889 | 0.625059 | 0.707739 |
| ENSGALG00010009978 | NA | NA | 0.32489 | 0.131309 | 3.7578 | 1.12246 | 3.25004 | 0.951314 | 0.622906 | 2.30562 |
| ENSGALG00010012702 | PTPRO | 373911 | 0.311151 | 0.314052 | 0.0215264 | 0.00980573 | 0.0843459 | 0.0451439 | 0.114375 | 0.122742 |
| ENSGALG00010003073 | NA | NA | 0.294555 | 0.158732 | 1.09841 | 0.966777 | 2.46066 | 2.00183 | 0.658869 | 0.948199 |
| ENSGALG00010007222 | NA | NA | 0.284998 | 0.594962 | 1.55774 | 2.00995 | 1.9572 | 2.22535 | 1.47296 | 1.23473 |
| ENSGALG00010012104 | SMC1B | 418243 | 0.278822 | 0.420994 | 3.63894 | 2.26831 | 2.08733 | 1.65338 | 0.791785 | 1.32324 |
| ENSGALG00010020658 | NA | NA | 0.271357 | 0.0658039 | 0.81389 | 0.575292 | 0.732532 | 0.44945 | 0.236485 | 0.292377 |
| ENSGALG00010011537 | NA | NA | 0.270871 | 0.349476 | 1.29249 | 6.94244 | 0.897222 | 19.2818 | 12.7389 | 0.0784994 |
| ENSGALG00010000071 | LOC112532885 | 112532885 | 0.266344 | 0.730655 | 0.222526 | 0.0992084 | 0.189311 | 0.0487428 | 0.23937 | 0.0949945 |
| ENSGALG00010021043 | PTPRG | 374208 | 0.263132 | 0.389899 | 0.593504 | 0.638393 | 0.763483 | 0.696475 | 0.539419 | 0.544099 |
| ENSGALG00010014632 | NWD2 | 428789 | 0.26279 | 0.301422 | 4.0612 | 4.82124 | 0.530816 | 0.766359 | 1.84436 | 1.96527 |
| ENSGALG00010024550 | KCNA10 | 395116 | 0.26278 | 0.294547 | 0.0332177 | 0.0181576 | 0.0811677 | 0.0227985 | 0.0559804 | 0.0358872 |
| ENSGALG00010023305 | PLCB2 | 423014 | 0.256469 | 0.0574096 | 1.02684 | 0.815363 | 1.32435 | 1.08153 | 0.832149 | 1.03922 |
| ENSGALG00010027010 | NA | NA | 0.254159 | 0.403419 | 1.56405 | 1.97572 | 1.31382 | 1.54855 | 1.53941 | 0.821542 |
| ENSGALG00010015177 | TC2N | 423412 | 0.248833 | 0.193094 | 1.69855 | 1.28632 | 1.54462 | 1.09292 | 0.647044 | 0.589839 |
| ENSGALG00010017497 | NA | NA | 0.24739 | 0.237287 | 0.179248 | 0.259278 | 0.0558844 | 0.0557567 | 0.166939 | 0.177152 |
| ENSGALG00010021919 | PARVA | 423030 | 0.242275 | 0.249106 | 0.931632 | 0.637822 | 0.88168 | 0.842881 | 0.602314 | 0.98977 |
| ENSGALG00010024228 | LOC418303 | 418303 | 0.236589 | 0.143918 | 0.784675 | 0.590185 | 1.16035 | 0.720155 | 0.0653502 | 0.0352049 |
| ENSGALG00010009695 | ATAD2B | 421972 | 0.232336 | 0.335397 | 1.27808 | 0.835754 | 1.03099 | 0.574481 | 0.639359 | 0.723515 |
| ENSGALG00010018558 | MSLN | 416534 | 0.226648 | 0.267815 | 1.58462 | 1.72555 | 1.56735 | 1.39434 | 0.63602 | 0.752513 |
| ENSGALG00010029191 | GSG1L2 | 768455 | 0.222474 | 0.113579 | 2.30458 | 1.33499 | 1.79768 | 0.906668 | 0.634207 | 0.483155 |
| ENSGALG00010028880 | LOC426218 | 426218 | 0.221362 | 0 | 4.02942 | 3.55625 | 2.23771 | 1.29634 | 0.742723 | 1.26322 |
| ENSGALG00010002868 | PPP1R9A | 420568 | 0.217201 | 0.271178 | 2.04616 | 1.30096 | 1.03623 | 0.639954 | 0.487904 | 0.947148 |
| ENSGALG00010007595 | ATP2C2 | 100859316 | 0.217111 | 0.147788 | 0.745342 | 0.730361 | 0.750832 | 0.669186 | 0.514853 | 0.424692 |
| ENSGALG00010004053 | NA | NA | 0.212692 | 0.136932 | 0.593161 | 0.628184 | 0.618382 | 0.401719 | 0.442074 | 0.543664 |
| ENSGALG00010009106 | THSD7B | 424297 | 0.206857 | 0.172804 | 0.519441 | 0.414129 | 0.482519 | 0.299907 | 0.337415 | 0.354928 |
| ENSGALG00010015806 | NA | NA | 0.205633 | 2.44294 | 0.0115814 | 0.0168818 | 0.0205813 | 0 | 0.00390353 | 0.0035749 |
| ENSGALG00010026706 | KRT24 | 395861 | 0.205326 | 0.318665 | 3.34041 | 3.61785 | 0.415121 | 2.0575 | 7.63716 | 2.68591 |
| ENSGALG00010010911 | ADAP1L | 100858460 | 0.203383 | 0 | 0.892485 | 0.331259 | 0.532349 | 0.869662 | 0.557063 | 0.854524 |
| ENSGALG00010012754 | NA | NA | 0.188938 | 0.28417 | 2.22679 | 1.362 | 1.63096 | 1.47252 | 0.496564 | 0.538628 |
| ENSGALG00010001137 | NA | NA | 0.173826 | 0 | 5.53724 | 7.83723 | 5.05189 | 5.20294 | 5.8323 | 2.21282 |
| ENSGALG00010025649 | NA | NA | 0.172956 | 0.484776 | 1.75523 | 1.44512 | 3.07577 | 2.63057 | 0.730348 | 0.701465 |
| ENSGALG00010026350 | LOC101749943 | 101749943 | 0.16369 | 0.253084 | 0.56658 | 0.789691 | 0.444182 | 0.647677 | 0.498974 | 0.463063 |

| | | | | | | | | | | |
|---|---|---|---|---|---|---|---|---|---|---|
| ENSGALG00010000685 | LOC112530906 | 112530906 | 0.154693 | 0.217883 | 9.11295 | 8.55706 | 14.2854 | 8.86612 | 7.01253 | 7.95327 |
| ENSGALG00010015694 | NA | NA | 0.148859 | 0.160436 | 1.56181 | 1.56002 | 1.92278 | 1.46368 | 0.808645 | 0.421106 |
| ENSGALG00010008982 | NA | NA | 0.148619 | 0.112515 | 0.0154647 | 0.0244209 | 0.024047 | 0.00353799 | 0.0026062 | 0.00159119 |
| ENSGALG00010006023 | NA | NA | 0.137866 | 0.101353 | 0.00495305 | 0.010529 | 0.0275064 | 0.0132201 | 0.0222591 | 0.017837 |
| ENSGALG00010017619 | POLN | 422892 | 0.136488 | 0.245352 | 0.797764 | 0.59182 | 1.01897 | 0.585014 | 0.39733 | 0.419945 |
| ENSGALG00010007894 | CD88 | 768950 | 0.12875 | 0.176312 | 0.822853 | 0.596583 | 0.703392 | 0.818055 | 0.301303 | 0.575145 |
| ENSGALG00010023863 | EYA2 | 395745 | 0.117494 | 0.178386 | 0.45448 | 0.417227 | 0.391175 | 0.349687 | 0.539667 | 0.336366 |
| ENSGALG00010001334 | EBF3 | 395175 | 0.115256 | 0.158059 | 0.624873 | 0.523974 | 0.530321 | 0.47894 | 0.499867 | 0.533849 |
| ENSGALG00010004464 | NA | NA | 0.110649 | 0 | 0.566472 | 0.114684 | 0.559265 | 0.503988 | 0.238663 | 0.60714 |
| ENSGALG00010014860 | NA | NA | 0.10914 | 0.163611 | 0.84659 | 0.339363 | 1.74268 | 0.67143 | 0.580201 | 0.818808 |
| ENSGALG00010023747 | NA | NA | 0.108507 | 0.109828 | 0.73352 | 0.955133 | 0.584207 | 0.615999 | 0.399495 | 0.328585 |
| ENSGALG00010007190 | NA | NA | 0.103953 | 0.0682114 | 0.413928 | 0.652805 | 0.367022 | 0.405848 | 0.0967228 | 0.300634 |
| ENSGALG00010010376 | NA | NA | 0.102213 | 0.396584 | 0.69771 | 2.17178 | 1.61445 | 1.72923 | 1.51877 | 5.15981 |
| ENSGALG00010000423 | CARD8 | 416998 | 0.100196 | 0.179427 | 1.54326 | 1.75747 | 2.19129 | 1.18357 | 1.13785 | 1.13678 |
| ENSGALG00010011260 | NA | NA | 0.0969763 | 0.117583 | 0.510266 | 0.318292 | 0.255291 | 0.678343 | 0.375736 | 0.567592 |
| ENSGALG00010014826 | NA | NA | 0.0960066 | 0.0771202 | 0.324261 | 0.303189 | 0.360152 | 0.175699 | 0.238718 | 0.267349 |
| ENSGALG00010024353 | UMOD | 404754 | 0.0924438 | 0.112088 | 0.788785 | 0.814432 | 0.992905 | 0.390991 | 0.708964 | 0.385508 |
| ENSGALG00010030013 | NA | NA | 0.0907962 | 0.176144 | 2.32418 | 1.03519 | 1.03257 | 1.06345 | 0.696328 | 1.35512 |
| ENSGALG00010016745 | BEGAIN | 428909 | 0.090438 | 0.305129 | 0.0126748 | 0.00339625 | 0 | 0.0102343 | 0.0150779 | 0.0270416 |
| ENSGALG00010020306 | NA | NA | 0.0820804 | 0.0364915 | 1.5369 | 1.16622 | 2.09595 | 1.04593 | 0.295071 | 0.339288 |
| ENSGALG00010022577 | LOC770870 | 770870 | 0.0788316 | 0.216655 | 1.73391 | 0.953246 | 0.697283 | 0.854917 | 0.466023 | 0.715159 |
| ENSGALG00010006368 | NA | NA | 0.0786153 | 0 | 0.536634 | 0.488896 | 0.993385 | 1.07424 | 0.452183 | 0.586662 |
| ENSGALG00010009858 | ST8SIA5 | 414797 | 0.0745345 | 0.0492943 | 0.279442 | 0.15275 | 0.21405 | 0.121247 | 0.159143 | 0.22754 |
| ENSGALG00010007386 | NA | NA | 0.0731695 | 0.0642148 | 0.594596 | 0.935338 | 0.761671 | 0.49878 | 0.494344 | 0.315072 |
| ENSGALG00010029575 | ANKFN1 | 427807 | 0.0703197 | 0.0591153 | 0.417784 | 0.350655 | 0.401827 | 0.223698 | 0.189501 | 0.243515 |
| ENSGALG00010023657 | NA | NA | 0.0689968 | 0.137145 | 0.549473 | 0.246193 | 0.587424 | 0.398909 | 0.184334 | 0.381323 |
| ENSGALG00010009257 | LOC424300 | 424300 | 0.066435 | 0.0439117 | 0.154508 | 0.194996 | 0.55123 | 0.403212 | 0.0822863 | 0.210591 |
| ENSGALG00010021854 | TAPBPL | 426340 | 0.0663317 | 0.0321708 | 0.767218 | 0.481257 | 0.502901 | 0.273356 | 0.286147 | 0.339703 |
| ENSGALG00010011625 | ERBB4 | 395671 | 0.0639518 | 0.0556028 | 0.269288 | 0.242908 | 0.223705 | 0.205527 | 0.100184 | 0.226807 |
| ENSGALG00010012099 | UPK3A | 418242 | 0.0617251 | 0.0957972 | 0.194285 | 0.667487 | 0.19499 | 0.366831 | 0.536495 | 0.659319 |
| ENSGALG00010012259 | IL1RAPL2 | 422379 | 0.0599184 | 0.206315 | 0.563112 | 0.613385 | 0.293164 | 0.37621 | 0.357709 | 0.308397 |
| ENSGALG00010018390 | SLC12A1 | 415433 | 0.0581774 | 0.0507888 | 0.249305 | 0.128638 | 0.173981 | 0.222086 | 0.141287 | 0.166848 |
| ENSGALG00010018119 | GRID2IP | 427671 | 0.0563478 | 0.0273286 | 0.219791 | 0.321216 | 0.259376 | 0.199039 | 0.181344 | 0.166077 |
| ENSGALG00010004514 | NA | NA | 0.0554529 | 0 | 0.977857 | 0.517279 | 1.57658 | 0.721654 | 0.478435 | 0.267762 |
| ENSGALG00010025142 | NA | NA | 0.0543331 | 0.105406 | 2.10166 | 1.29524 | 1.02983 | 1.20204 | 1.77092 | 1.43103 |
| ENSGALG00010001785 | CNTN1 | 417786 | 0.0505382 | 0.0560251 | 0.20078 | 0.134279 | 0.149007 | 0.0762092 | 0.0992031 | 0.10881 |
| ENSGALG00010012429 | CXCR7 | 428997 | 0.0496382 | 0.137478 | 0.647945 | 0.465745 | 0.667669 | 0.260093 | 0.396335 | 0.15138 |
| ENSGALG00010012123 | RASL11AL | 422321 | 0.0496228 | 0.192536 | 0.606888 | 0.295738 | 0.26649 | 0.274457 | 0.784912 | 0.343078 |
| ENSGALG00010029047 | NA | NA | 0.0488581 | 0 | 0.555848 | 0.810241 | 0.709978 | 0.381499 | 0.327862 | 0.557625 |
| ENSGALG00010005063 | NA | NA | 0.0469374 | 0 | 0.436097 | 0.462169 | 0.444827 | 0.956975 | 0.599949 | 0.336532 |
| ENSGALG00010003157 | NA | NA | 0.0459809 | 0.0892027 | 0.889294 | 1.00082 | 0.929625 | 0.79785 | 0.866891 | 0.44405 |
| ENSGALG00010012344 | NA | NA | 0.0434364 | 0.0300951 | 0.150603 | 0.211168 | 0.254829 | 0.234185 | 0.183414 | 0.151629 |
| ENSGALG00010007018 | NA | NA | 0.0378288 | 0.0917345 | 0.55948 | 1.19586 | 2.62903 | 1.30459 | 0.652756 | 0.846885 |
| ENSGALG00010005359 | MEIS2 | 395588 | 0.0367071 | 0.102983 | 1.38646 | 1.14548 | 0.536618 | 0.548988 | 0.50997 | 0.112049 |
| ENSGALG00010019114 | FAM46D | 422282 | 0.032471 | 0.0354339 | 0.327857 | 0.189311 | 0.17438 | 0.110925 | 0.147858 | 0.0855223 |
| ENSGALG00010003632 | TECRL | 422618 | 0.0312524 | 0.033683 | 0.207405 | 0.169159 | 0.107502 | 0.212394 | 0.0965372 | 0.0533507 |
| ENSGALG00010011535 | NA | NA | 0.0269808 | 0.0994512 | 0.417459 | 2.58396 | 0.368205 | 4.2486 | 2.30198 | 0.00947497 |
| ENSGALG00010029537 | MYH1B | 374069 | 0.0236757 | 0.0145742 | 0.076144 | 0.020292 | 0.121968 | 0.0817679 | 0.0196412 | 0.0371744 |
| ENSGALG00010021486 | LOC107052926 | 107052926 | 0.0221042 | 0.085764 | 0.892733 | 1.22284 | 0.809995 | 0.956469 | 0.874086 | 0.138268 |
| ENSGALG00010011860 | CECR1 | 418160 | 0.0203251 | 0.0295729 | 0.231234 | 0.210664 | 0.410925 | 0.211605 | 0.199716 | 0.258739 |
| ENSGALG00010013001 | ADGRV1 | 431566 | 0.0192057 | 0.0203544 | 0.0942781 | 0.0593498 | 0.190551 | 0.0175883 | 0.0511426 | 0.0568289 |
| ENSGALG00010000702 | DDC | 420947 | 0.0164774 | 0.00710357 | 0.120807 | 0.313103 | 1.43664 | 1.09123 | 0.187796 | 0.0980478 |
| ENSGALG00010019541 | NA | NA | 0.0150611 | 0.160702 | 0.58258 | 0.686859 | 4.54847 | 0.607609 | 1.84809 | 0.568575 |
| ENSGALG00010025127 | PAMR1 | 423158 | 0.0143463 | 0.00869742 | 0.0979288 | 0.096652 | 0.117832 | 0.0606776 | 0.0807986 | 0.0551036 |
| ENSGALG00010010092 | NA | NA | 0.0120012 | 0.0349234 | 1.65207 | 1.39316 | 2.6235 | 0.827763 | 1.90981 | 1.04309 |
| ENSGALG00010025597 | SCN2A | 395945 | 0.00780606 | 0.0126198 | 0.0947284 | 0.0768623 | 0.156176 | 0.17947 | 0.0424049 | 0.0879496 |
| ENSGALG00010023242 | MYT1 | 419249 | 0.00708325 | 0.0584013 | 0.308236 | 0.156009 | 0.425145 | 0.686742 | 0.276705 | 0.102607 |
| ENSGALG00010004758 | FLT3 | 100858422 | 0.0056106 | 0.0081634 | 0.0861712 | 0.0523371 | 0.0992539 | 0.062063 | 0.0457177 | 0.0394059 |
| ENSGALG00010010854 | LRIT1 | 428935 | 0.00557572 | 0.024338 | 0.193473 | 0.0693489 | 0.151478 | 0.0907018 | 0.0427609 | 0.0807694 |
| ENSGALG00010015117 | RASD2 | 418057 | 0 | 0 | 0.660086 | 0.212247 | 0.287509 | 0.0296105 | 1.06879 | 0.359563 |
| ENSGALG00010003338 | NA | NA | 0 | 0.0424981 | 0.0498446 | 21.038 | 0.695975 | 0.89598 | 0.198003 | 1.33527 |
| ENSGALG00010007969 | NA | NA | 0 | 0 | 0.783666 | 0 | 0.870406 | 0.199207 | 4.18215 | 2.75496 |
| ENSGALG00010013632 | SLC1A3 | 395443 | 0 | 0.00863806 | 0.579173 | 0.636873 | 0.0956478 | 0.0656717 | 0.399813 | 0.204577 |
| ENSGALG00010005583 | ALDOB | 427308 | 0 | 0.0212993 | 0.574568 | 0.0455179 | 0.48556 | 0.114303 | 0.399948 | 0.491582 |
| ENSGALG00010016399 | ACSM5 | 426992 | 0 | 0 | 0.183903 | 0.268069 | 0.170215 | 0.231402 | 0.0619849 | 0.0189221 |
| ENSGALG00010017068 | ZP2 | 427006 | 0 | 0 | 0.334321 | 0.317271 | 0.0928314 | 0.063738 | 0.187806 | 0.193494 |
| ENSGALG00010028756 | LOC107055126 | 107055126 | 0 | 0 | 10.5366 | 11.3663 | 11.9939 | 8.87464 | 4.01959 | 4.20707 |
| ENSGALG00010028757 | MAT2A | 110231148 | 0 | 0 | 1.71629 | 1.56361 | 2.70889 | 1.85992 | 0.0761157 | 1.39415 |

### Supplementally table S6. Gene list of Intersection in supplementally figure S2E

| Emsembl_Id | Symbol | entrez_ID | CEF_Male_TPM | CEF_F_TPM | MO_Male_TPM | MO_Female_TPM | GSP_Male_TPM | GSP_Female_TPM | GSN_Male_TPM | GSN_Female_TPM | PNP_Female_TPM | PNP_Male_TPM |
|---|---|---|---|---|---|---|---|---|---|---|---|---|
| ENSGALG00010015173 | ENS-1 | 769415 | 146.961 | 82.2245 | 2147.68 | 4908.89 | 1266.53 | 828.345 | 1365.18 | 780.488 | 1942.95 | 1854.11 |
| ENSGALG00010006087 | LOC107051465 | 107051465 | 369.942 | 146.716 | 4942.55 | 4498.56 | 1424.1 | 817.725 | 1354.39 | 812.534 | 2215.66 | 1903.72 |
| ENSGALG00010004570 | LOC107051826 | 107051826 | 113.862 | 86.917 | 3913.24 | 3568.04 | 663.504 | 321.21 | 748.079 | 449.668 | 1526.97 | 1539.11 |
| ENSGALG00010002022 | ERNI | 769673 | 65.8803 | 47.9637 | 3300.43 | 2470.02 | 991.468 | 534.697 | 843.743 | 508.579 | 1798.01 | 1991.1 |
| ENSGALG00010022849 | NA | NA | 0.748309 | 0.731964 | 2760.85 | 1992.4 | 240.361 | 134.601 | 183.13 | 167.359 | 288.964 | 356.146 |
| ENSGALG00010004699 | NA | NA | 96.9509 | 99.6954 | 720.459 | 1077.42 | 480.861 | 633.861 | 765.195 | 541.999 | 305.939 | 259.896 |
| ENSGALG00010011992 | YEATS4 | 771226 | 35.9622 | 37.6458 | 929.455 | 721.508 | 525.85 | 701.108 | 679.585 | 639.532 | 531.058 | 403.691 |
| ENSGALG00010027214 | PLIN4 | 100857433 | 162.182 | 103.335 | 774.585 | 609.998 | 260.739 | 413.265 | 260.413 | 265.123 | 443.961 | 284.228 |
| ENSGALG00010012582 | CD24 | 100859742 | 30.7028 | 47.9549 | 510.303 | 561.179 | 289.575 | 333.078 | 348.877 | 235.977 | 390.637 | 304.987 |
| ENSGALG00010026300 | HMGA1 | 378795 | 103.992 | 87.9607 | 429.073 | 445.939 | 281.346 | 374.909 | 348.643 | 317.348 | 325.602 | 207.663 |
| ENSGALG00010024452 | NA | NA | 97.2907 | 85.3967 | 396.481 | 399.796 | 232.447 | 263.684 | 261.341 | 216.759 | 254.516 | 259.752 |
| ENSGALG00010009928 | NA | NA | 12.7987 | 7.09204 | 204.309 | 208.534 | 58.7747 | 37.5266 | 69.0822 | 42.146 | 115.061 | 96.8406 |
| ENSGALG00010007054 | NA | NA | 24.74 | 29.0463 | 188.53 | 194.492 | 120.231 | 149.459 | 118.707 | 115.161 | 112.263 | 114.493 |
| ENSGALG00010027297 | LOC121107553 | 121107553 | 0.254304 | 0.183858 | 123.943 | 180.359 | 49.8493 | 59.5395 | 293.337 | 265.71 | 362.667 | 295.262 |
| ENSGALG00010001386 | FKBP14 | 420636 | 55.5081 | 53.8916 | 123.909 | 161.286 | 83.1791 | 94.648 | 52.9252 | 56.103 | 69.9671 | 72.7787 |
| ENSGALG00010000465 | CUNH17orf49 | 112531156 | 35.416 | 62.1348 | 141.209 | 161.062 | 90.5472 | 133.821 | 109.152 | 109.543 | 109.4 | 77.5342 |
| ENSGALG00010028815 | NA | NA | 0.193348 | 0 | 205.569 | 148.775 | 35.2628 | 43.0863 | 130.05 | 133.186 | 173.548 | 129.8 |
| ENSGALG00010003700 | REST | 422623 | 11.7696 | 14.2723 | 105.343 | 148.286 | 79.6219 | 76.891 | 104.258 | 75.4373 | 44.0224 | 69.4097 |
| ENSGALG00010001705 | HTRA1 | 100857572 | 37.1018 | 36.9449 | 107.867 | 137.422 | 51.9575 | 68.9331 | 37.5029 | 31.8769 | 39.7011 | 68.3162 |
| ENSGALG00010025783 | NA | NA | 0 | 1.00784 | 105.538 | 137.313 | 108.278 | 126.21 | 295.999 | 290.922 | 253.413 | 254.129 |
| ENSGALG00010022340 | GFOD2 | 415642 | 45.5446 | 39.6776 | 176.316 | 123.191 | 69.8276 | 99.4511 | 107.522 | 87.8199 | 71.9306 | 55.471 |
| ENSGALG00010006885 | RHOU | 428589 | 5.06896 | 4.51169 | 53.9499 | 113.163 | 40.066 | 40.5878 | 33.6915 | 23.8339 | 33.5192 | 52.4195 |
| ENSGALG00010022984 | NA | NA | 0.2272 | 0.0676376 | 153.172 | 103.55 | 46.3633 | 23.6365 | 29.0818 | 23.3223 | 46.1757 | 62.6039 |
| ENSGALG00010001989 | NA | NA | 3.14401 | 2.86669 | 18.2628 | 102.972 | 4.79363 | 3.33092 | 5.77199 | 3.31675 | 8.43459 | 7.326 |
| ENSGALG00010003268 | SMS | 418602 | 6.49249 | 7.7496 | 68.4314 | 99.0386 | 57.6635 | 54.5316 | 48.1178 | 41.234 | 54.561 | 47.2027 |
| ENSGALG00010021172 | SLC29A4 | 425593 | 7.74564 | 6.19744 | 35.5829 | 96.0327 | 10.4214 | 11.1051 | 23.7836 | 18.6649 | 25.5293 | 17.6515 |
| ENSGALG00010021870 | LOC101747373 | 101747373 | 33.127 | 39.1473 | 104.86 | 95.2043 | 27.6611 | 35.9317 | 35.2919 | 33.0788 | 23.8055 | 27.8305 |
| ENSGALG00010027217 | SEMA6B | 428330 | 12.4585 | 8.28778 | 97.7483 | 81.5564 | 34.2849 | 59.5006 | 23.5981 | 27.16 | 41.3541 | 38.5813 |
| ENSGALG00010024883 | ING5 | 424852 | 6.89171 | 8.58612 | 72.3974 | 80.7132 | 20.5842 | 27.895 | 34.2978 | 35.6531 | 39.5329 | 38.4325 |
| ENSGALG00010002185 | NA | NA | 0.384878 | 8.02174 | 51.191 | 70.3007 | 4.49645 | 11.0204 | 7.94206 | 7.16425 | 8.97508 | 7.28989 |
| ENSGALG00010001703 | NA | NA | 0.0604824 | 0 | 57.1758 | 67.1982 | 4.2966 | 4.17186 | 6.22446 | 2.59042 | 13.4786 | 14.9425 |
| ENSGALG00010023247 | LOC121106503 | 121106503 | 0.0698723 | 1.48653 | 67.8065 | 65.432 | 104.044 | 109.998 | 111.423 | 104.797 | 97.1778 | 126.119 |
| ENSGALG00010000925 | NA | NA | 0 | 0 | 55.3505 | 57.3106 | 13.0224 | 11.1814 | 11.5716 | 6.3037 | 15.9351 | 19.821 |
| ENSGALG00010009891 | NA | NA | 14.7645 | 20.453 | 56.4047 | 57.273 | 26.4902 | 12.8764 | 27.4946 | 14.4416 | 24.6619 | 29.3009 |
| ENSGALG00010024872 | RUNDC3A | 100858314 | 8.85385 | 2.86984 | 42.8551 | 48.4614 | 3.59349 | 6.66865 | 10.1101 | 4.46124 | 9.75818 | 17.1417 |
| ENSGALG00010000726 | POU4F1 | 100857623 | 0 | 0.202701 | 54.579 | 48.3195 | 8.82001 | 4.62084 | 12.0277 | 8.54258 | 12.1996 | 20.6328 |
| ENSGALG00010018514 | EAF2 | 424243 | 9.44017 | 8.13892 | 46.8335 | 47.9617 | 30.7404 | 21.8058 | 24.0623 | 14.9027 | 22.5079 | 26.2756 |
| ENSGALG00010000316 | NA | NA | 1.98369 | 0.154864 | 127.179 | 46.0825 | 38.7943 | 34.1461 | 68.9898 | 66.8064 | 86.9962 | 69.9068 |
| ENSGALG00010000344 | LOC112531461 | 112531461 | 1.21636 | 0.243644 | 112.657 | 44.723 | 51.4365 | 54.0657 | 49.9036 | 47.1532 | 63.367 | 56.9174 |
| ENSGALG00010025832 | ENKD1 | 415643 | 16.4061 | 15.3451 | 49.1598 | 39.1753 | 25.4517 | 29.1215 | 29.0923 | 25.4228 | 27.6831 | 23.7944 |
| ENSGABL00010008602 | NA | NA | 0.272272 | 1.19846 | 61.2086 | 38.9502 | 0.208426 | 0.647599 | 0.331997 | 0.218648 | 0.460599 | 0.281215 |
| ENSGALG00010000330 | NA | NA | 1.36281 | 0.251566 | 104.817 | 38.8728 | 44.3501 | 46.3828 | 55.1002 | 53.4995 | 74.999 | 63.8499 |
| ENSGALG00010000771 | NA | NA | 3.69169 | 4.56641 | 4.5007 | 36.5502 | 197.405 | 164.676 | 84.7498 | 77.9919 | 159.578 | 170.013 |
| ENSGALG00010024210 | NXPH3 | 429523 | 0.230958 | 0.0645466 | 17.681 | 34.0025 | 2.90961 | 3.06057 | 1.70581 | 3.03819 | 8.78164 | 8.17863 |
| ENSGALG00010000176 | NA | NA | 12.1151 | 12.1015 | 72.1251 | 33.8201 | 16.0333 | 15.5678 | 7.16095 | 20.7223 | 13.617 | 12.2161 |
| ENSGALG00010010930 | FAM163A | 771080 | 0.0134709 | 0.11859 | 47.3221 | 33.5846 | 5.56023 | 5.19055 | 6.11039 | 3.72132 | 7.54755 | 8.89338 |
| ENSGALG00010016761 | GPR160 | 424992 | 1.7391 | 2.76968 | 28.9849 | 32.694 | 11.5731 | 11.4984 | 22.1153 | 13.4919 | 14.7278 | 13.1144 |
| ENSGALG00010017612 | CLDN11 | 424990 | 0.505521 | 0.164826 | 31.1074 | 32.4547 | 9.37594 | 7.27155 | 7.15036 | 4.5751 | 12.673 | 16.4892 |
| ENSGALG00010027290 | CSPG5 | 395466 | 1.17914 | 2.14433 | 51.0336 | 30.6357 | 13.5065 | 18.6644 | 26.018 | 20.7165 | 20.4831 | 17.4001 |
| ENSGALG00010025263 | SAMD14 | 100857537 | 0.991215 | 0.861835 | 23.5526 | 30.0209 | 5.24401 | 7.36752 | 2.1487 | 3.27243 | 8.85613 | 7.23465 |
| ENSGALG00010006840 | GINS2 | 415824 | 10.5675 | 5.48158 | 30.8213 | 29.0406 | 48.1587 | 68.8489 | 57.8791 | 63.4817 | 49.488 | 41.3715 |
| ENSGALG00010021837 | NA | NA | 11.4805 | 11.2298 | 23.9843 | 27.4698 | 15.472 | 21.1861 | 21.0641 | 23.7375 | 17.3393 | 15.8795 |
| ENSGALG00010017637 | NA | NA | 0.2192 | 0.214412 | 46.526 | 27.0255 | 32.3515 | 34.4102 | 35.2813 | 33.3281 | 50.0356 | 27.5301 |
| ENSGALG00010006933 | NA | NA | 1.62299 | 1.48512 | 60.8917 | 26.6572 | 21.6259 | 24.2825 | 22.2925 | 14.4629 | 16.1429 | 15.4976 |
| ENSGALG00010008933 | NA | NA | 1.42873 | 3.0618 | 29.1244 | 25.4556 | 8.41076 | 12.5965 | 7.34057 | 3.04055 | 8.33158 | 18.0687 |
| ENSGALG00010000543 | NA | NA | 0.464892 | 0 | 43.0612 | 25.2425 | 4.29242 | 4.12528 | 6.31404 | 7.39006 | 7.71126 | 9.07134 |
| ENSGALG00010003108 | PPP1R3B | 422630 | 4.16262 | 4.59994 | 38.6995 | 25.2288 | 9.69925 | 12.4129 | 8.05587 | 8.33588 | 18.9378 | 13.3235 |
| ENSGALG00010017122 | NCBP2 | 424889 | 6.76786 | 7.87171 | 27.4615 | 24.2585 | 16.4132 | 18.4674 | 19.6533 | 17.165 | 17.9829 | 13.6513 |
| ENSGALG00010000569 | SIPA1 | 101749913 | 0.362761 | 0.39578 | 23.1311 | 23.2819 | 54.3257 | 64.1479 | 53.924 | 63.8056 | 45.1164 | 42.0559 |
| ENSGALG00010025031 | ECHDC2 | 424646 | 6.76983 | 8.25263 | 19.8908 | 22.8118 | 38.9729 | 51.0645 | 46.6796 | 48.4902 | 44.2075 | 35.431 |
| ENSGALG00010015222 | NA | NA | 0.437293 | 0.0611059 | 53.6886 | 22.1185 | 249.344 | 259.661 | 381.551 | 227.651 | 290.353 | 249.727 |
| ENSGALG00010016891 | LOC107052288 | 107052288 | 0.451385 | 0.883051 | 7.81302 | 20.7654 | 2.01241 | 2.03988 | 5.81221 | 5.35512 | 1.87337 | 1.9096 |
| ENSGALG00010010888 | GNPAT | 421548 | 6.0494 | 7.71584 | 19.8387 | 20.0162 | 25.9693 | 39.3017 | 30.293 | 37.4499 | 29.0882 | 26.9934 |
| ENSGALG00010002367 | STK31 | 420621 | 0.024225 | 0.0315945 | 20.3611 | 18.8933 | 36.825 | 42.1837 | 45.9127 | 41.8534 | 26.8958 | 27.1091 |
| ENSGALG00010001350 | LOC107049524 | 107049524 | 3.23081 | 3.99189 | 6.9259 | 17.124 | 39.9557 | 46.1847 | 43.9694 | 47.4225 | 36.7142 | 35.8713 |
| ENSGALG00010004429 | LOC112531814 | 112531814 | 0 | 0.500745 | 7.63336 | 16.7427 | 1.09727 | 2.58744 | 2.16396 | 1.55306 | 2.00147 | 0.704987 |
| ENSGALG00010001208 | MOV10L1 | 417738 | 0 | 0.0179625 | 12.5063 | 12.9076 | 18.4583 | 26.0975 | 25.7207 | 27.7209 | 20.205 | 24.2723 |
| ENSGALG00010001272 | NA | NA | 0 | 0.16321 | 7.15929 | 11.8955 | 1.78819 | 3.62139 | 1.3835 | 2.45654 | 1.58069 | 3.6535 |
| ENSGALG00010001176 | IL12A | 407090 | 0.0937114 | 0.274993 | 25.0237 | 10.8483 | 3.78769 | 8.46992 | 4.61641 | 2.80952 | 3.67791 | 5.03304 |
| ENSGALG00010011694 | NA | NA | 0.199598 | 0.219145 | 10.8958 | 10.6271 | 9.8037 | 5.18343 | 9.4849 | 3.36276 | 4.76304 | 0.0605839 |
| ENSGALG00010025767 | NA | NA | 0 | 0.0304377 | 15.0892 | 10.5577 | 7.1398 | 6.06288 | 10.7556 | 10.3565 | 7.72067 | 9.62897 |
| ENSGALG00010021054 | FEZF2 | 416074 | 0 | 0 | 16.0302 | 10.2189 | 39.5408 | 47.0524 | 45.6663 | 69.8582 | 49.5044 | 22.655 |
| ENSGALG00010014492 | CPLX2 | 100857281 | 1.35607 | 1.52849 | 13.7818 | 10.1764 | 5.75272 | 8.61888 | 5.82567 | 6.72145 | 24.5589 | 16.5995 |
| ENSGALG00010024629 | ESAM | 419707 | 3.41498 | 1.12688 | 9.32779 | 9.93899 | 1.84778 | 2.91955 | 1.11042 | 1.20416 | 2.68514 | 2.96903 |
| ENSGALG00010013073 | NA | NA | 0.19258 | 0.706399 | 4.74687 | 9.87801 | 8.72729 | 25.021 | 11.6942 | 29.8477 | 6.95007 | 14.52 |
| ENSGALG00010013435 | SDC3 | 396343 | 0.666017 | 0.445973 | 9.57736 | 9.67586 | 1.64791 | 1.73828 | 1.49995 | 1.31618 | 5.40006 | 4.72014 |
| ENSGALG00010012091 | LOC121108229 | 121108229 | 2.62058 | 4.76682 | 7.62802 | 8.92639 | 5.59229 | 7.49763 | 6.27039 | 7.26578 | 5.00838 | 3.58318 |
| ENSGALG00010011822 | NA | NA | 0.2996 | 0 | 11.6926 | 8.88208 | 0.91738 | 0.534449 | 2.19191 | 0.721782 | 0.878504 | 0.495104 |
| ENSGALG00010027167 | ASIC1L | 107055878 | 2.11623 | 1.19289 | 22.3648 | 8.84454 | 1.05436 | 2.26079 | 2.69415 | 5.70321 | 4.59535 | 5.39395 |
| ENSGALG00010000830 | NA | NA | 0.150397 | 0.165116 | 9.81881 | 8.72521 | 3.10842 | 1.20503 | 0.496623 | 0.0403789 | 1.70527 | 0.719647 |
| ENSGALG00010011166 | SOX7 | 771337 | 1.25911 | 1.10331 | 18.6788 | 8.64078 | 3.2771 | 8.42285 | 3.32649 | 2.3593 | 3.97604 | 3.94474 |
| ENSGALG00010028386 | NCAN | 395493 | 0.385393 | 0.045237 | 53.7883 | 8.38599 | 7.71301 | 12.0449 | 15.0904 | 9.81291 | 11.5094 | 7.27319 |
| ENSGALG00010027277 | KCND3 | 374228 | 1.13528 | 0.263289 | 18.0036 | 8.30983 | 1.87571 | 1.4663 | 27.9245 | 16.1036 | 4.02567 | 4.13605 |
| ENSGALG00010010753 | LOC107051636 | 107051636 | 0.297707 | 0.216055 | 3.38414 | 8.14358 | 0.929229 | 1.85019 | 1.55509 | 1.50128 | 1.53363 | 1.11091 |
| ENSGALG00010001707 | NA | NA | 1.42836 | 1.12043 | 7.69789 | 7.84507 | 3.74402 | 2.7901 | 0.228858 | 0.384196 | 0.759532 | 1.74467 |
| ENSGALG00010023104 | EFCAB4B | 423106 | 2.13141 | 1.72724 | 8.45076 | 7.65882 | 4.22808 | 5.2766 | 4.21897 | 4.7907 | 4.77728 | 4.45112 |
| ENSGALG00010000659 | LOC121109782 | 121109782 | 0.407588 | 0.295994 | 5.45741 | 7.06182 | 16.7616 | 25.2277 | 32.2535 | 12.6231 | 29.2966 | 28.5651 |
| ENSGALG00010014283 | NA | NA | 0.585291 | 0.830134 | 7.88129 | 6.55513 | 0 | 0.0348029 | 0.114188 | 0 | 0.0528069 | 0 |
| ENSGALG00010003001 | MAP6 | 395753 | 0.112607 | 0.190744 | 11.4561 | 6.0978 | 3.50188 | 3.96202 | 2.42467 | 1.48804 | 4.41084 | 3.75129 |
| ENSGALG00010008603 | NA | NA | 1.39878 | 1.14019 | 10.4996 | 6.07041 | 1.9036 | 4.34359 | 1.5161 | 1.33131 | 0.420676 | 0.25684 |
| ENSGALG00010010076 | NA | NA | 0.362504 | 0.240203 | 4.53705 | 5.76693 | 32.4691 | 49.1741 | 135.937 | 153.793 | 112.858 | 97.7911 |
| ENSGALG00010005530 | NA | NA | 0.983564 | 1.56736 | 6.22595 | 5.59705 | 22.3304 | 41.2572 | 39.5472 | 43.3111 | 9.99064 | 12.6909 |
| ENSGALG00010005837 | LOC107052005 | 107052005 | 0.16023 | 0.352642 | 6.11438 | 5.54195 | 1.55038 | 1.53395 | 1.3989 | 1.6127 | 1.29385 | 1.23788 |
| ENSGALG00010016457 | SMOX | 422934 | 1.84345 | 1.3377 | 6.29149 | 4.9371 | 1.33807 | 1.78432 | 1.16902 | 1.08172 | 3.5989 | 3.15874 |
| ENSGALG00010003723 | NA | NA | 1.05698 | 0.773533 | 6.77219 | 4.8868 | 6.82735 | 7.39069 | 2.20137 | 1.76784 | 4.71601 | 5.65769 |
| ENSGALG00010014058 | NA | NA | 0.0943868 | 0.0230813 | 3.98526 | 4.75256 | 12.7744 | 13.6103 | 13.4311 | 14.831 | 7.28109 | 9.93391 |
| ENSGALG00010011469 | CPS1 | 428994 | 2.39743 | 6.57803 | 4.49444 | 4.6917 | 4.89214 | 6.2052 | 8.11958 | 7.66997 | 4.70239 | 4.53128 |
| ENSGALG00010002061 | NA | NA | 1.41625 | 2.46142 | 8.46188 | 4.09455 | 0.0196869 | 0.00637178 | 0.0627175 | 0.00956133 | 0.0370606 | 0.0206595 |
| ENSGALG00010002873 | MRAP2 | 421836 | 0.0280319 | 0 | 1.58664 | 4.0563 | 0.309003 | 0.21669 | 0.252938 | 0.210103 | 0.512149 | 0.486403 |
| ENSGALG00010011588 | COLGALT1 | 424447 | 0.980048 | 0.672731 | 4.66189 | 4.05092 | 9.45906 | 13.2638 | 12.35 | 13.694 | 11.6657 | 8.91952 |
| ENSGALG00010017143 | FAM149A | 422721 | 1.4198 | 1.41681 | 4.10143 | 4.03189 | 1.10253 | 1.18244 | 1.26925 | 0.845548 | 1.40651 | 1.65661 |
| ENSGALG00010003205 | TNX | 396106 | 0.203622 | 0.167307 | 2.63965 | 3.82515 | 8.27503 | 14.4762 | 11.4292 | 11.0539 | 8.41047 | 9.35459 |
| ENSGALG00010009307 | DIRAS2 | 425241 | 0.130776 | 0.0983997 | 1.80526 | 3.62074 | 1.42046 | 2.48328 | 0.221495 | 0.326216 | 1.34457 | 1.68379 |
| ENSGALG00010017006 | DNAH3 | 427004 | 0.462791 | 0.542307 | 3.70035 | 3.16438 | 8.0117 | 6.19448 | 8.96377 | 4.7657 | 8.31448 | 3.94583 |
| ENSGALG00010013866 | KCTD8 | 770600 | 0.0166608 | 0.0279376 | 4.85743 | 3.07337 | 0.87164 | 0.993525 | 2.05244 | 1.08055 | 0.790251 | 0.979383 |
| ENSGALG00010001276 | LOC121108470 | 121108470 | 0.0889684 | 0.133001 | 0.986914 | 3.04758 | 0.236442 | 0.379303 | 0.325862 | 0.386438 | 0.249898 | 0.11651 |
| ENSGALG00010020960 | CLEC19A | 771758 | 0.167281 | 0.130902 | 13.1945 | 3.02281 | 0.573684 | 0.795756 | 1.2728 | 1.89861 | 1.41871 | 1.21634 |
| ENSGALG00010029392 | RTN4RL1 | 427828 | 0.212521 | 0.180162 | 3.78653 | 2.9769 | 0.555301 | 0.95199 | 0.234952 | 0.318575 | 1.15046 | 1.94624 |
| ENSGALG00010014029 | NA | NA | 0.0397744 | 0.0194528 | 2.45099 | 2.92921 | 7.52662 | 8.66213 | 8.08481 | 8.73052 | 4.30181 | 6.70849 |
| ENSGALG00010027945 | DCST2 | 121107585 | 0.0728045 | 0 | 3.29001 | 2.82647 | 1.20381 | 0.898296 | 21.4923 | 15.4642 | 36.5464 | 28.5518 |
| ENSGALG00010016760 | DLK1 | 423459 | 0.117081 | 0.143154 | 1.16683 | 2.47931 | 0.430204 | 0.348095 | 0.133245 | 0.0626812 | 0.492959 | 0.580445 |
| ENSGALG00010003094 | LOC107053759 | 107053759 | 0.0810942 | 0.0793229 | 2.82805 | 2.19809 | 5.89989 | 8.23609 | 8.74123 | 6.47755 | 7.22879 | 4.55642 |
| ENSGALG00010002208 | NA | NA | 1.11739 | 0.977382 | 2.55023 | 2.09318 | 0.381627 | 0.945535 | 0.859423 | 0.943341 | 0.814275 | 0.648068 |
| ENSGALG00010016428 | ADRA1D | 422933 | 0 | 0 | 1.48375 | 2.05783 | 0.173544 | 0.307275 | 0.0379419 | 0.0267729 | 0.308315 | 0.702453 |
| ENSGALG00010000825 | NA | NA | 0.161803 | 0.246195 | 11.1837 | 1.83815 | 2.4682 | 2.45583 | 2.03114 | 1.35011 | 1.47827 | 1.77583 |
| ENSGALG00010020532 | SIX3 | 378786 | 0.316265 | 0.494971 | 9.15258 | 1.6371 | 27.3479 | 28.585 | 29.5554 | 45.9529 | 59.4088 | 5.93201 |

| | | | | | | | | | | | | |
|---|---|---|---|---|---|---|---|---|---|---|---|---|
| ENSGALG00010021710 | KEL | 418306 | 0.00804578 | 0.0314801 | 1.66368 | 1.38574 | 0.00985452 | 0.086116 | 0 | 0.0344596 | 0 | 0 |
| ENSGALG00010022843 | NA | NA | 0.0840492 | 0.0411067 | 3.93243 | 1.30521 | 0.463248 | 0.274878 | 0.184475 | 0.494969 | 0.37916 | 0.156257 |
| ENSGALG00010023498 | DISP2 | 428836 | 0.0295148 | 0.0433051 | 1.03467 | 1.2889 | 2.95706 | 3.34156 | 2.75532 | 3.13181 | 2.2475 | 3.15816 |
| ENSGALG00010010761 | MEI1 | 417985 | 0.40517 | 0.380032 | 1.7481 | 1.21712 | 4.14339 | 4.82508 | 4.50191 | 5.6071 | 3.31005 | 3.32947 |
| ENSGALG00010014282 | NA | NA | 0 | 0.245204 | 1.04617 | 1.11224 | 0 | 0 | 0 | 0 | 0 | 0 |
| ENSGALG00010016612 | GRIA1 | 414893 | 0.0720465 | 0.334294 | 2.19833 | 0.806731 | 0.185537 | 0.456967 | 0.232465 | 0.14934 | 0.259177 | 0.238121 |
| ENSGALG00010015088 | LOC107050476 | 107050476 | 0.0511902 | 0.087626 | 0.183336 | 0.742127 | 0.103693 | 0.155707 | 0.144044 | 0.162325 | 0.0888182 | 0.0715798 |
| ENSGALG00010015661 | NA | NA | 0.0379746 | 0.0448837 | 0.806749 | 0.72597 | 16.9593 | 10.8632 | 0.00617394 | 0 | 0.00285516 | 0.00261479 |
| ENSGALG00010002126 | MGAM | 425545 | 0.0167928 | 0.145781 | 0.753111 | 0.638114 | 0.15426 | 0.132307 | 0.0245717 | 0.13935 | 0.210221 | 0.154365 |
| ENSGALG00010029927 | CFAP52 | 417322 | 0.0506615 | 0.0566342 | 0.977856 | 0.558543 | 2.22939 | 2.96942 | 2.74278 | 3.58017 | 1.61286 | 1.57873 |
| ENSGALG00010028809 | NA | NA | 0 | 0.0823156 | 2.72731 | 0.554414 | 3.45292 | 5.05403 | 6.32098 | 5.22617 | 4.06206 | 2.29463 |
| ENSGALG00010005024 | NA | NA | 0.192675 | 0.173389 | 5.4915 | 0.514998 | 0 | 0 | 0.00751805 | 0.0165043 | 0.0347675 | 0.0191043 |
| ENSGALG00010000228 | NA | NA | 0.18806 | 0.0490538 | 0.480714 | 0.460184 | 0.138202 | 0.20874 | 0.0366898 | 0.0268483 | 0.101804 | 0.176107 |
| ENSGALG00010006610 | NA | NA | 0.0280435 | 0.0822929 | 0.426692 | 0.435491 | 0.646885 | 1.03387 | 0.661103 | 1.11601 | 0.809657 | 0.892108 |
| ENSGALG00010015499 | NA | NA | 0.128189 | 0.120657 | 0.895363 | 0.392669 | 2.0085 | 1.9315 | 1.60083 | 1.66909 | 1.33441 | 2.51708 |
| ENSGALG00010003018 | MOGAT2 | 426437 | 0.00569142 | 0.0445368 | 2.01598 | 0.353531 | 0.160331 | 0.108296 | 0.0277595 | 0.03047 | 0.374854 | 0.0987563 |
| ENSGALG00010012104 | SMC1B | 418243 | 0.00279485 | 0.0328056 | 0.420994 | 0.278822 | 1.65338 | 2.08733 | 2.26831 | 3.63894 | 0.791785 | 1.32324 |
| ENSGALG00010014632 | NWD2 | 428789 | 0.0366859 | 0.0318974 | 0.301422 | 0.26279 | 0.766359 | 0.530816 | 4.82124 | 4.0612 | 1.84436 | 1.96527 |
| ENSGALG00010024550 | KCNA10 | 395116 | 0.0124093 | 0.00606912 | 0.294547 | 0.26278 | 0.0227985 | 0.0811677 | 0.0181576 | 0.0332177 | 0.0559804 | 0.0358872 |
| ENSGALG00010017497 | NA | NA | 0.00734239 | 0 | 0.237287 | 0.24739 | 0.0557567 | 0.0558844 | 0.259278 | 0.179248 | 0.166939 | 0.177152 |
| ENSGALG00010015806 | NA | NA | 0.00432652 | 0.00423202 | 2.44294 | 0.205633 | 0 | 0.0205813 | 0.0168818 | 0.0115814 | 0.00390353 | 0.0035749 |
| ENSGALG00010026350 | LOC101749943 | 101749943 | 0.00913541 | 0.0113417 | 0.253084 | 0.16369 | 0.647677 | 0.444182 | 0.789691 | 0.56658 | 0.498974 | 0.463063 |
| ENSGALG00010000685 | LOC112530906 | 112530906 | 0.0180125 | 0.028631 | 0.217883 | 0.154693 | 8.86612 | 14.2854 | 8.55706 | 9.11295 | 7.01253 | 7.95327 |
| ENSGALG00010008982 | NA | NA | 0.00577722 | 0.00753471 | 0.112515 | 0.148619 | 0.00353799 | 0.024047 | 0.0244209 | 0.0154647 | 0.0026062 | 0.00159119 |
| ENSGALG00010006023 | NA | NA | 0 | 0 | 0.101353 | 0.137866 | 0.0132201 | 0.0275064 | 0.010529 | 0.00495305 | 0.0222591 | 0.017837 |
| ENSGALG00010016745 | BEGAIN | 428909 | 0.00905217 | 0.00817333 | 0.305129 | 0.090438 | 0.0102343 | 0 | 0.00339625 | 0.0126748 | 0.0150779 | 0.0270416 |
| ENSGALG00010011625 | ERBB4 | 395671 | 0.0126371 | 0.0172244 | 0.0556028 | 0.0639518 | 0.205527 | 0.223705 | 0.242908 | 0.269288 | 0.100184 | 0.226807 |
| ENSGALG00010012259 | IL1RAPL2 | 422379 | 0.0253451 | 0.0354165 | 0.206315 | 0.0599184 | 0.37621 | 0.293164 | 0.613385 | 0.563112 | 0.357709 | 0.308397 |

Supplementally table S7. DEGs comparing sgSoprano#1 and sgOVA in CRISPRi PGCs (p-adj < 0.05)

| Ensembl_ID | Symbol | entrez_ID | logFC | logCPM | p-adj_value | FDR |
|---|---|---|---|---|---|---|
| ENSGALG00010018908 | NA | NA | -1.493179561 | 7.02467312 | 2.61E-23 | 4.00E-19 |
| ENSGALG00010007105 | NA | NA | -1.176074809 | 5.22453252 | 5.02E-22 | 3.85E-18 |
| ENSGALG00010007287 | LOC124418405 | 124418405 | -0.670374171 | 9.50645294 | 1.44E-21 | 7.34E-18 |
| ENSGALG00010006797 | NA | NA | -1.132040684 | 5.60951112 | 4.71E-16 | 1.80E-12 |
| ENSGALG00010013595 | WEE2 | 427918 | -0.946989705 | 6.02073927 | 2.31E-14 | 7.07E-11 |
| ENSGALG00010016645 | NA | NA | -1.150409575 | 7.59278481 | 1.19E-13 | 3.04E-10 |
| ENSGALG00010013729 | NA | NA | -0.934121163 | 3.51234535 | 2.30E-13 | 5.04E-10 |
| ENSGALG00010004208 | NA | NA | -1.459519689 | 1.91622363 | 2.33E-12 | 4.45E-09 |
| ENSGALG00010009553 | NA | NA | -0.846753298 | 7.22060104 | 2.27E-11 | 3.86E-08 |
| ENSGALG00010001103 | NA | NA | -0.540336707 | 6.68537542 | 9.38E-11 | 1.44E-07 |
| ENSGALG00010017060 | CLSTN1 | 395372 | 0.43338252 | 8.24311508 | 3.33E-10 | 4.64E-07 |
| ENSGALG00010004527 | NA | NA | -0.885533819 | 4.63156822 | 3.93E-10 | 5.01E-07 |
| ENSGALG00010006086 | LOC100857117 | 100857117 | -0.649171627 | 9.33668308 | 1.13E-09 | 1.34E-06 |
| ENSGALG00010010905 | ENC1 | 427203 | 0.434857581 | 7.24363622 | 1.29E-09 | 1.41E-06 |
| ENSGALG00010006859 | NA | NA | -0.765343884 | 3.57568613 | 2.21E-09 | 2.26E-06 |
| ENSGALG00010006798 | NA | NA | -0.851828133 | 4.29104721 | 2.38E-09 | 2.28E-06 |
| ENSGALG00010006087 | LOC107051465 | 107051465 | -0.711279589 | 9.85439916 | 3.00E-09 | 2.71E-06 |
| ENSGALG00010002022 | ERNI | 769673 | -0.700979307 | 10.1600576 | 4.83E-09 | 4.11E-06 |
| ENSGALG00010000831 | NA | NA | -0.608800042 | 7.52734459 | 7.03E-09 | 5.67E-06 |
| ENSGALG00010006214 | NA | NA | -0.747958514 | 7.83627045 | 1.13E-08 | 8.64E-06 |
| ENSGALG00010009928 | NA | NA | -0.740831142 | 5.78052686 | 1.46E-08 | 1.06E-05 |
| ENSGALG00010002503 | NA | NA | -0.634496686 | 8.72872606 | 1.85E-08 | 1.25E-05 |
| ENSGALG00010014283 | NA | NA | -1.060298304 | 3.01304444 | 1.88E-08 | 1.25E-05 |
| ENSGALG00010006179 | NA | NA | -0.749954334 | 6.20234634 | 3.32E-08 | 2.06E-05 |
| ENSGALG00010024824 | DNAJC5 | 419256 | 0.540996121 | 4.52241936 | 3.35E-08 | 2.06E-05 |
| ENSGALG00010018244 | CHM | 422266 | 0.436655678 | 5.78014024 | 3.85E-08 | 2.27E-05 |
| ENSGALG00010000091 | LOC121108173 | 121108173 | 2.145770562 | 0.12218626 | 6.30E-08 | 3.57E-05 |
| ENSGALG00010001026 | NA | NA | -0.549846073 | 5.20704775 | 6.52E-08 | 3.57E-05 |
| ENSGALG00010004570 | LOC107051826 | 107051826 | -0.674931446 | 10.7155499 | 8.84E-08 | 4.67E-05 |
| ENSGALG00010018537 | LOC112530649 | 112530649 | -0.593870081 | 9.03669587 | 9.83E-08 | 5.02E-05 |
| ENSGALG00010019204 | ARF6 | 428927 | 0.403049784 | 7.190115 | 1.17E-07 | 5.79E-05 |
| ENSGALG00010026334 | CALCOCO2 | 419993 | 0.412399749 | 5.29303928 | 1.45E-07 | 6.95E-05 |
| ENSGALG00010007049 | NA | NA | -0.659242519 | 9.36655066 | 2.10E-07 | 9.73E-05 |
| ENSGALG00010011693 | NA | NA | -0.508683547 | 5.49687356 | 2.33E-07 | 0.0001051 |
| ENSGALG00010009801 | LOC101750251 | 101750251 | -0.598120281 | 9.21883 | 2.41E-07 | 0.00010554 |
| ENSGALG00010025449 | LOC121108098 | 121108098 | 1.012218034 | 2.06947963 | 2.55E-07 | 0.00010834 |
| ENSGALG00010025965 | NA | NA | 0.603340169 | 4.46678148 | 3.46E-07 | 0.00014325 |
| ENSGALG00010001828 | LOC100858869 | 100858869 | -0.689586063 | 7.85185665 | 3.64E-07 | 0.0001468 |
| ENSGALG00010024154 | VPS29L | 416931 | 0.494286042 | 4.4128635 | 7.23E-07 | 0.0002842 |
| ENSGALG00010015617 | SRSF10 | 419689 | -0.787308759 | 2.984672 | 7.98E-07 | 0.00030548 |
| ENSGALG00010028355 | RBM18 | 417111 | 0.411420873 | 5.79279257 | 8.75E-07 | 0.00032694 |
| ENSGALG00010012112 | MBD2 | 497269 | 0.603119007 | 3.41336779 | 9.71E-07 | 0.00035421 |
| ENSGALG00010005001 | GUCA1C | 427960 | -0.646637217 | 5.61632872 | 9.98E-07 | 0.00035572 |
| ENSGALG00010008262 | CYB5B | 415865 | 0.412834746 | 6.10259983 | 1.13E-06 | 0.00039228 |
| ENSGALG00010023095 | NA | NA | -0.92681823 | 2.92534208 | 1.22E-06 | 0.00040885 |
| ENSGALG00010025365 | SIKE1 | 419887 | 0.401239573 | 6.66681068 | 1.23E-06 | 0.00040885 |
| ENSGALG00010003982 | GLCCI1 | 420580 | 0.335718594 | 7.65153446 | 1.42E-06 | 0.000463 |
| ENSGALG00010009327 | NID1 | 395531 | 0.346928591 | 7.32543965 | 1.68E-06 | 0.00053716 |
| ENSGALG00010002947 | LOC121107416 | 121107416 | -0.538282098 | 10.7106449 | 1.85E-06 | 0.00057795 |
| ENSGALG00010006902 | NA | NA | -1.110781471 | 2.4697438 | 2.01E-06 | 0.00061512 |
| ENSGALG00010015173 | ENS-1 | 769415 | -0.476616655 | 10.9764083 | 2.17E-06 | 0.00065229 |
| ENSGALG00010001989 | NA | NA | -0.534291657 | 7.79824485 | 2.72E-06 | 0.00080098 |

| | | | | | | |
|---|---|---|---|---|---|---|
| ENSGALG00010003307 | COL1A2 | 396243 | 0.493954682 | 7.59986776 | 3.05E-06 | 0.0008813 |
| ENSGALG00010002060 | NA | NA | -0.738279349 | 2.88401753 | 4.13E-06 | 0.00115558 |
| ENSGALG00010014282 | NA | NA | -1.335623676 | 1.52159873 | 4.15E-06 | 0.00115558 |
| ENSGALG00010018386 | NA | NA | -1.32589352 | 0.59206744 | 4.41E-06 | 0.00120701 |
| ENSGALG00010003817 | BF2 | 425389 | 0.359266063 | 5.95757858 | 4.69E-06 | 0.00126044 |
| ENSGALG00010017976 | GPR137C | 428932 | -0.398766848 | 4.907209 | 4.97E-06 | 0.00131196 |
| ENSGALG00010001031 | NA | NA | -0.680697511 | 3.60503555 | 5.33E-06 | 0.00138388 |
| ENSGALG00010021964 | NA | NA | -0.412944774 | 5.57560037 | 5.64E-06 | 0.00143958 |
| ENSGALG00010027908 | PBXIP1 | 100857958 | 0.530795326 | 3.68534888 | 6.76E-06 | 0.00169188 |
| ENSGALG00010023092 | NA | NA | -0.893287424 | 3.28940292 | 6.94E-06 | 0.00169188 |
| ENSGALG00010002195 | NA | NA | -0.651182969 | 4.6362102 | 6.96E-06 | 0.00169188 |
| ENSGALG00010003639 | NA | NA | -0.786442681 | 9.23607216 | 7.12E-06 | 0.00170546 |
| ENSGALG00010011997 | DENND5B | 418139 | 0.41418522 | 4.89532035 | 7.56E-06 | 0.0017584 |
| ENSGALG00010000830 | NA | NA | -0.436301991 | 5.71167695 | 7.57E-06 | 0.0017584 |
| ENSGALG00010019767 | SUCLG2 | 416087 | 0.329784963 | 6.95262866 | 9.13E-06 | 0.00208706 |
| ENSGALG00010020476 | TMEM132A | 107053338 | 0.364918318 | 5.9898151 | 1.13E-05 | 0.00255311 |
| ENSGALG00010006088 | NA | NA | -0.574712665 | 4.99927621 | 1.25E-05 | 0.00276918 |
| ENSGALG00010013681 | NA | NA | -1.031564892 | 1.91027311 | 1.29E-05 | 0.0028264 |
| ENSGALG00010009859 | NA | NA | -1.231031662 | 2.5182047 | 1.54E-05 | 0.00331303 |
| ENSGALG00010027169 | ABCB8 | 107055879 | 0.490629636 | 5.01272194 | 1.56E-05 | 0.00331367 |
| ENSGALG00010012578 | NDUFS4 | 374122 | 0.393149969 | 6.49181353 | 1.72E-05 | 0.00360928 |
| ENSGALG00010013151 | LOC107055505 | 107055505 | 0.621104415 | 3.27272696 | 1.79E-05 | 0.00370692 |
| ENSGALG00010006916 | NA | NA | -0.715968342 | 3.66793643 | 1.90E-05 | 0.00388351 |
| ENSGALG00010005396 | MICU2 | 418946 | 0.374269584 | 5.03133958 | 2.03E-05 | 0.00409766 |
| ENSGALG00010018568 | NA | NA | 0.422750015 | 5.67007301 | 2.06E-05 | 0.00410512 |
| ENSGALG00010013385 | PGRMC1 | 772196 | 0.373981796 | 5.89979652 | 2.22E-05 | 0.00436847 |
| ENSGALG00010017139 | COL9A2 | 396524 | 0.413129146 | 5.12366212 | 2.48E-05 | 0.00480787 |
| ENSGALG00010001423 | FAM210A | 421045 | -0.344649013 | 5.91348183 | 2.67E-05 | 0.00511925 |
| ENSGALG00010006575 | SSBP2 | 770958 | -0.287771793 | 7.42531695 | 2.86E-05 | 0.00541156 |
| ENSGALG00010000093 | LOC121108174 | 121108174 | 1.967386807 | 1.10875835 | 2.98E-05 | 0.00556982 |
| ENSGALG00010023487 | GRN | 101748909 | 0.335668002 | 5.74145992 | 3.17E-05 | 0.0057835 |
| ENSGALG00010002837 | DNMT1 | 396011 | 0.33373991 | 7.69447396 | 3.17E-05 | 0.0057835 |
| ENSGALG00010013572 | DENND11 | 427917 | -0.448886342 | 6.79019328 | 3.32E-05 | 0.00598519 |
| ENSGALG00010007504 | TMPRSS15 | 427967 | -1.116661401 | 1.23418181 | 3.51E-05 | 0.00625234 |
| ENSGALG00010000135 | NA | NA | 1.200844867 | 0.629242 | 3.57E-05 | 0.00628709 |
| ENSGALG00010027156 | ISYNA1 | 107055367 | 0.343647875 | 6.09271664 | 3.78E-05 | 0.00658788 |
| ENSGALG00010013772 | KIAA0319L | 419634 | 0.309196474 | 5.98074554 | 3.90E-05 | 0.00671364 |
| ENSGALG00010016809 | SYTL1 | 771987 | 0.625696321 | 2.89247946 | 4.56E-05 | 0.00776095 |
| ENSGALG00010003814 | TPP1 | 425576 | 0.331746435 | 6.78098166 | 4.68E-05 | 0.00787654 |
| ENSGALG00010024848 | ESYT1 | 107055399 | 0.384352688 | 7.68249223 | 4.79E-05 | 0.00797953 |
| ENSGALG00010007167 | NA | NA | -0.610947157 | 4.70982123 | 4.95E-05 | 0.00815728 |
| ENSGALG00010028586 | BABAM1 | 776677 | 0.297870758 | 6.18647363 | 5.19E-05 | 0.00845839 |
| ENSGALG00010009232 | ECH1 | 100858352 | 0.345775066 | 5.63644196 | 5.24E-05 | 0.00845839 |
| ENSGALG00010006306 | NA | NA | -0.792688594 | 2.76873996 | 5.79E-05 | 0.00924224 |
| ENSGALG00010022409 | CARNS1 | 100359387 | 0.737970053 | 1.94383178 | 6.51E-05 | 0.0102884 |
| ENSGALG00010022807 | LOC107050028 | 107050028 | 0.37016176 | 4.79202547 | 6.68E-05 | 0.01044822 |
| ENSGALG00010005791 | NA | NA | -0.909623347 | 1.79983312 | 6.88E-05 | 0.01062425 |
| ENSGALG00010007117 | NA | NA | -0.750356571 | 1.89436281 | 7.05E-05 | 0.01062425 |
| ENSGALG00010014214 | REEP2 | 771891 | -0.339755967 | 4.82165538 | 7.11E-05 | 0.01062425 |
| ENSGALG00010007963 | LDAH | 421967 | 0.291754342 | 5.79647378 | 7.14E-05 | 0.01062425 |
| ENSGALG00010021802 | NA | NA | -0.458705457 | 4.96324449 | 7.14E-05 | 0.01062425 |
| ENSGALG00010009771 | NA | NA | -0.625972795 | 3.30953172 | 7.23E-05 | 0.01065578 |
| ENSGALG00010004281 | VIM | 420519 | 0.36984993 | 10.1578059 | 7.63E-05 | 0.01106667 |

| | | | | | | |
|---|---|---|---|---|---|---|
| ENSGALG00010019115 | DBNL | 425994 | 0.304490793 | 6.27977283 | 7.66E-05 | 0.01106667 |
| ENSGALG00010010296 | NA | NA | -0.445517394 | 4.05882499 | 7.75E-05 | 0.01110017 |
| ENSGALG00010029932 | RASA4B | 417513 | 0.636364893 | 3.26662856 | 8.13E-05 | 0.01144782 |
| ENSGALG00010013245 | NA | NA | -1.246409542 | 0.72500194 | 8.14E-05 | 0.01144782 |
| ENSGALG00010026917 | AMT | 395566 | 0.360928634 | 4.7805794 | 8.47E-05 | 0.01180299 |
| ENSGALG00010012306 | LOC121108207 | 121108207 | -0.539250871 | 4.22130539 | 8.56E-05 | 0.01180952 |
| ENSGALG00010001416 | AP4M1 | 396046 | 0.402866603 | 4.30688266 | 8.76E-05 | 0.0119879 |
| ENSGALG00010001138 | SPTSSB | 425008 | 0.325749675 | 5.29225966 | 9.13E-05 | 0.01237702 |
| ENSGALG00010023764 | COX5A | 769735 | 0.326908349 | 6.94587636 | 9.52E-05 | 0.01279327 |
| ENSGALG00010012294 | LOC100857299 | 100857299 | -0.535459516 | 7.93300724 | 9.64E-05 | 0.01284076 |
| ENSGALG00010022478 | ATP6V1A | 395821 | 0.26498837 | 8.59745214 | 9.79E-05 | 0.01287683 |
| ENSGALG00010028088 | APOA1BP | 426569 | 0.329762605 | 5.87057098 | 9.83E-05 | 0.01287683 |
| ENSGALG00010004371 | NA | NA | -0.334302557 | 9.24734869 | 9.95E-05 | 0.01292503 |
| ENSGALG00010017667 | SLC2A1 | 396130 | 0.304029049 | 5.20443481 | 0.00010374 | 0.01335567 |
| ENSGALG00010023412 | GSS | 428135 | 0.320820486 | 5.55574273 | 0.00010644 | 0.01348156 |
| ENSGALG00010008642 | NA | NA | -1.841758494 | -0.7074821 | 0.00010647 | 0.01348156 |
| ENSGALG00010003536 | NA | NA | -0.545793889 | 5.52686151 | 0.00011389 | 0.01430213 |
| ENSGALG00010028057 | APC2 | 429363 | 0.339253463 | 5.01107576 | 0.00012108 | 0.0150248 |
| ENSGALG00010027359 | JTB | 770053 | 0.37121969 | 5.59920394 | 0.00012171 | 0.0150248 |
| ENSGALG00010020571 | MRTO4 | 428192 | 0.441661988 | 4.55414474 | 0.00012262 | 0.0150248 |
| ENSGALG00010010193 | NA | NA | -0.681530032 | 5.26506443 | 0.00012454 | 0.0150248 |
| ENSGALG00010027034 | NA | NA | -1.149827479 | 0.44515179 | 0.00012455 | 0.0150248 |
| ENSGALG00010008968 | NA | NA | -0.819995051 | 7.64584258 | 0.00012561 | 0.01503536 |
| ENSGALG00010028027 | NIT1 | 429566 | 1.069114361 | 1.40797641 | 0.00012877 | 0.0152566 |
| ENSGALG00010009186 | PREB | 100858986 | 0.387158234 | 4.91081568 | 0.00012945 | 0.0152566 |
| ENSGALG00010028686 | TAGLN2 | 121106445 | 0.337278781 | 6.06058432 | 0.00013359 | 0.01559211 |
| ENSGALG00010009180 | NFKBIB | 100858419 | 0.392336747 | 4.83040968 | 0.00013434 | 0.01559211 |
| ENSGALG00010018542 | LOC107054982 | 107054982 | 0.301719817 | 6.27228408 | 0.00014328 | 0.01650485 |
| ENSGALG00010028310 | NA | NA | -0.809574572 | 1.8127822 | 0.00015032 | 0.0171364 |
| ENSGALG00010000768 | NA | NA | -0.531048774 | 6.09707193 | 0.00015156 | 0.0171364 |
| ENSGALG00010006079 | LOC107055200 | 107055200 | -0.446431181 | 7.79937 | 0.00015254 | 0.0171364 |
| ENSGALG00010027356 | NA | NA | 0.367533585 | 5.31911594 | 0.00015323 | 0.0171364 |
| ENSGALG00010006266 | CENPW | 421716 | 0.299410846 | 6.67592945 | 0.00015836 | 0.01745873 |
| ENSGALG00010024611 | NA | NA | 0.461736744 | 4.47010849 | 0.0001584 | 0.01745873 |
| ENSGALG00010020277 | IDH2 | 431056 | 0.317810895 | 6.39856702 | 0.00016518 | 0.01807704 |
| ENSGALG00010015392 | TMEM165 | 428780 | 0.286344474 | 6.19231917 | 0.00016828 | 0.01828521 |
| ENSGALG00010008982 | NA | NA | -0.963945707 | 1.64640457 | 0.00017194 | 0.01855078 |
| ENSGALG00010012230 | PTPRQ | 772163 | -1.745824419 | -0.5755629 | 0.00018024 | 0.0193112 |
| ENSGALG00010021233 | NA | NA | 0.274713523 | 5.95803613 | 0.00018477 | 0.01964946 |
| ENSGALG00010019132 | H2AZ2 | 426617 | 0.316452584 | 6.90130152 | 0.00018597 | 0.01964946 |
| ENSGALG00010021804 | NA | NA | -0.475001987 | 4.82489313 | 0.00018925 | 0.01981939 |
| ENSGALG00010005477 | TMEM161B | 426919 | -0.305867582 | 5.87951812 | 0.00019016 | 0.01981939 |
| ENSGALG00010011371 | LOC424430 | 424430 | 0.426748751 | 3.80714932 | 0.00019484 | 0.02016966 |
| ENSGALG00010009891 | NA | NA | -0.656126701 | 3.41904324 | 0.00021236 | 0.02183631 |
| ENSGALG00010007161 | NA | NA | -0.590451905 | 4.36532609 | 0.00022453 | 0.02293367 |
| ENSGALG00010016320 | NA | NA | -0.553160453 | 2.68082771 | 0.00023776 | 0.02412408 |
| ENSGALG00010023499 | PEBP1 | 416990 | -0.286938902 | 7.75653441 | 0.0002566 | 0.02560395 |
| ENSGALG00010019246 | IL17RC | 101750586 | 0.353459398 | 4.51982648 | 0.00025829 | 0.02560395 |
| ENSGALG00010005730 | NA | NA | -1.293742399 | 0.20896922 | 0.00026037 | 0.02560395 |
| ENSGALG00010023706 | CDK5RAP3 | 430350 | 0.299402376 | 5.46176833 | 0.00026207 | 0.02560395 |
| ENSGALG00010008297 | NA | NA | 1.79600785 | -0.8268632 | 0.0002621 | 0.02560395 |
| ENSGALG00010003191 | NA | NA | 0.395615779 | 5.03019893 | 0.00026283 | 0.02560395 |
| ENSGALG00010006111 | WBP1 | 107049494 | 0.376517709 | 4.59474321 | 0.00026404 | 0.02560395 |

| | | | | | | |
|---|---|---|---|---|---|---|
| ENSGALG00010005008 | LOC418414 | 418414 | -1.031199203 | 0.93652199 | 0.00027808 | 0.02679298 |
| ENSGALG00010026806 | LOC101750445 | 101750445 | 0.879862915 | 1.122238 | 0.0002798 | 0.02679298 |
| ENSGALG00010028188 | SMAD4 | 107055136 | 0.300120074 | 5.80556154 | 0.00028329 | 0.02695813 |
| ENSGALG00010001697 | TAF10 | 107052460 | 0.385479011 | 4.289108 | 0.0002896 | 0.02738903 |
| ENSGALG00010027883 | ATP8B2 | 770035 | 0.28357259 | 6.31887501 | 0.00029281 | 0.02752248 |
| ENSGALG00010029216 | RAB34 | 100858849 | 0.403402872 | 4.21900476 | 0.00029494 | 0.02752487 |
| ENSGALG00010000176 | NA | NA | 0.582466962 | 2.59020353 | 0.00029734 | 0.02752487 |
| ENSGALG00010011353 | LRP1B | 424301 | -0.264044251 | 7.08236085 | 0.00029823 | 0.02752487 |
| ENSGALG00010003913 | PSMB5 | 396003 | 0.294818864 | 6.2990411 | 0.00031487 | 0.02888684 |
| ENSGALG00010008498 | LOC107050955 | 107050955 | 0.307749316 | 8.02158001 | 0.00032715 | 0.0298345 |
| ENSGALG00010015483 | SGCD | 416251 | -0.283227238 | 5.41303725 | 0.00033355 | 0.03023856 |
| ENSGALG00010001483 | LOC121110182 | 121110182 | -0.824668418 | 5.32885152 | 0.00034708 | 0.03127985 |
| ENSGALG00010017006 | DNAH3 | 427004 | -0.309057298 | 4.754251 | 0.00035553 | 0.03185429 |
| ENSGALG00010008650 | NA | NA | -0.74096968 | 4.27491763 | 0.00036338 | 0.03236805 |
| ENSGALG00010019752 | GPAM | 423895 | 0.276184692 | 7.21385286 | 0.00036882 | 0.03250063 |
| ENSGALG00010027923 | AMPD2 | 112530327 | 0.31371097 | 5.48925328 | 0.00036911 | 0.03250063 |
| ENSGALG00010018834 | SAMD4A | 423559 | 0.379072368 | 5.05000918 | 0.00037246 | 0.0326081 |
| ENSGALG00010010169 | NA | NA | -0.926542222 | 3.32497644 | 0.00038622 | 0.03362069 |
| ENSGALG00010000825 | NA | NA | -0.484497212 | 4.01228481 | 0.00039365 | 0.03407377 |
| ENSGALG00010014782 | NA | NA | -0.80018151 | 2.23013947 | 0.00040012 | 0.03443943 |
| ENSGALG00010028386 | NCAN | 395493 | 0.417701966 | 4.05217551 | 0.00041168 | 0.03518152 |
| ENSGALG00010018899 | ACSS1A | 416714 | 0.377653885 | 4.36602915 | 0.00041353 | 0.03518152 |
| ENSGALG00010015218 | PLCL1 | 424060 | 0.253132103 | 6.81535559 | 0.00041563 | 0.03518152 |
| ENSGALG00010017685 | PHC2 | 100857390 | 0.271216836 | 6.59198294 | 0.00042027 | 0.0353792 |
| ENSGALG00010015709 | RBMXL3 | 416302 | 0.285295535 | 8.73399131 | 0.00042551 | 0.03550524 |
| ENSGALG00010029492 | SDK2 | 395215 | 0.379642255 | 4.42269615 | 0.00042641 | 0.03550524 |
| ENSGALG00010022216 | ROMO1 | 768833 | 0.333771851 | 5.28641715 | 0.00043081 | 0.03567806 |
| ENSGALG00010002034 | WWC3 | 418649 | 0.252534175 | 6.5584497 | 0.00045187 | 0.03705492 |
| ENSGALG00010008427 | NA | NA | -1.442281393 | -0.3803962 | 0.00045227 | 0.03705492 |
| ENSGALG00010016852 | AHDC1 | 112530201 | 0.410762843 | 4.36116452 | 0.00047807 | 0.03896043 |
| ENSGALG00010018590 | TGFA | 414743 | 0.360810354 | 5.05375024 | 0.00048818 | 0.03957348 |
| ENSGALG00010018502 | ARHGEF10L | 419360 | 0.419232238 | 4.05434038 | 0.00049209 | 0.03968067 |
| ENSGALG00010009113 | RPLP1 | 396262 | 0.29592147 | 8.61047877 | 0.00051303 | 0.04069842 |
| ENSGALG00010014157 | NA | NA | -0.721379689 | 3.29896392 | 0.00051445 | 0.04069842 |
| ENSGALG00010020097 | RETSAT | 426549 | 0.461815656 | 3.92883394 | 0.00051448 | 0.04069842 |
| ENSGALG00010019281 | LOC421195 | 421195 | -0.547391506 | 4.63839328 | 0.00051623 | 0.04069842 |
| ENSGALG00010021779 | UNC13C | 415413 | -0.474557031 | 3.45421074 | 0.00051799 | 0.04069842 |
| ENSGALG00010021155 | RADIL | 416479 | 1.214152878 | 0.25846061 | 0.00052871 | 0.0413286 |
| ENSGALG00010022815 | LRRC8D | 424513 | -0.277672699 | 5.8378708 | 0.00053965 | 0.04196922 |
| ENSGALG00010028078 | LAMTOR2 | 100859842 | 0.317587254 | 4.56336951 | 0.00054797 | 0.04226298 |
| ENSGALG00010004648 | LOC421806 | 421806 | -0.32500623 | 8.15619404 | 0.00054894 | 0.04226298 |
| ENSGALG00010001479 | TMEM168 | 417777 | -0.277588827 | 5.3584508 | 0.000569 | 0.04358821 |
| ENSGALG00010018374 | HPS1 | 429879 | 0.417641579 | 3.65737545 | 0.00059127 | 0.04506883 |
| ENSGALG00010014122 | ABCA1 | 373945 | 0.275164977 | 6.12900238 | 0.00060348 | 0.04577204 |
| ENSGALG00010001418 | RNMT | 421046 | 0.292980161 | 7.38438608 | 0.00061174 | 0.04616976 |
| ENSGALG00010001271 | NA | NA | 0.405783308 | 3.85861232 | 0.00064013 | 0.04807562 |
| ENSGALG00010014226 | SLC16A7 | 417815 | -0.455135995 | 3.27706282 | 0.00067415 | 0.05038388 |
| ENSGALG00010006178 | RPS23 | 427323 | 0.312957613 | 8.12213161 | 0.0006823 | 0.05074487 |
| ENSGALG00010024205 | ANKRD13D | 107049270 | 0.560318559 | 2.77674926 | 0.00068676 | 0.05081153 |
| ENSGALG00010029277 | WFIKKN2 | 422097 | -0.624534977 | 2.26491664 | 0.00069135 | 0.05081153 |
| ENSGALG00010002078 | NA | NA | -1.246228064 | -0.0987862 | 0.00069314 | 0.05081153 |
| ENSGALG00010025355 | STAT6 | 100859196 | 0.474057852 | 3.33342302 | 0.0007074 | 0.05161002 |
| ENSGALG00010012753 | GTPBP1 | 418022 | -0.250285465 | 6.7161904 | 0.00071828 | 0.05215492 |

| | | | | | | |
|---|---|---|---|---|---|---|
| ENSGALG00010005419 | NA | NA | 0.585931552 | 2.50519716 | 0.00072764 | 0.05258573 |
| ENSGALG00010008209 | PPARGC1B | 416148 | 0.485950493 | 3.31027984 | 0.00073479 | 0.05285338 |
| ENSGALG00010007115 | CD69L | 417816 | -0.265405576 | 5.28492682 | 0.00075239 | 0.05386629 |
| ENSGALG00010027362 | RPS27 | 107055140 | 0.273745663 | 7.66713106 | 0.00076244 | 0.05420179 |
| ENSGALG00010009286 | MPV17 | 100857465 | 0.379663193 | 4.07819347 | 0.00076415 | 0.05420179 |
| ENSGALG00010013808 | HRASLS | 424903 | 0.531154499 | 2.94686788 | 0.00077719 | 0.05451427 |
| ENSGALG00010024149 | TCIRG1 | 395472 | 0.348878282 | 4.66894773 | 0.00077785 | 0.05451427 |
| ENSGALG00010019072 | AEBP1 | 100859431 | 0.412811347 | 5.52728913 | 0.00078186 | 0.05451427 |
| ENSGALG00010013164 | FAM83F | 770747 | 0.235926077 | 6.02096806 | 0.00078374 | 0.05451427 |
| ENSGALG00010006059 | BIVM | 374099 | 0.245166087 | 6.79879458 | 0.00078635 | 0.05451427 |
| ENSGALG00010021509 | DDIT4 | 107053650 | 0.441291961 | 5.94985075 | 0.00080256 | 0.05538775 |
| ENSGALG00010020617 | DGUOK | 107054975 | 0.382239305 | 3.98468941 | 0.00081304 | 0.05585929 |
| ENSGALG00010026003 | NECTIN1 | 100857875 | 0.237711884 | 6.70093298 | 0.00083738 | 0.05727457 |
| ENSGALG00010014243 | HIGD2A | 416227 | 0.409420337 | 3.37254942 | 0.00084775 | 0.05772619 |
| ENSGALG00010016647 | EGR1 | 373931 | -0.67928439 | 2.80651386 | 0.000855 | 0.05796188 |
| ENSGALG00010008906 | DBN1 | 396496 | 0.252666334 | 6.58012708 | 0.00086433 | 0.05833049 |
| ENSGALG00010002795 | NA | NA | -0.459772479 | 3.19831298 | 0.00086805 | 0.05833049 |
| ENSGALG00010028101 | NA | NA | 0.353640797 | 5.66529975 | 0.00087888 | 0.05880082 |
| ENSGALG00010028442 | CHTOP | 426357 | 0.24668252 | 7.48353955 | 0.00088824 | 0.05905302 |
| ENSGALG00010023930 | SRPRA | 419713 | 0.294254955 | 6.7444634 | 0.00089036 | 0.05905302 |
| ENSGALG00010004159 | NA | NA | -0.989737459 | 2.61634411 | 0.00090769 | 0.05985611 |
| ENSGALG00010000022 | ATP8 | 63549490 | -0.40484233 | 7.00130559 | 0.00091029 | 0.05985611 |
| ENSGALG00010027219 | LOC101749138 | 101749138 | 0.272845933 | 5.93771324 | 0.00092268 | 0.060412 |
| ENSGALG00010016896 | SPTBN1 | 421216 | 0.226549145 | 8.19986796 | 0.00092944 | 0.06052886 |
| ENSGALG00010027706 | GSN | 395774 | 0.244030005 | 5.76206949 | 0.00093237 | 0.06052886 |
| ENSGALG00010028912 | EXFABP | 396393 | 0.33826403 | 4.31191818 | 0.00094191 | 0.06063723 |
| ENSGALG00010013018 | ID1 | 395282 | 0.598987759 | 2.52682167 | 0.00094195 | 0.06063723 |
| ENSGALG00010024856 | SORT1 | 100858592 | 0.345444342 | 4.9634835 | 0.0010042 | 0.06437355 |
| ENSGALG00010028858 | LOC776719 | 776719 | 0.278796443 | 5.18790978 | 0.0010141 | 0.06473748 |
| ENSGALG00010025745 | COX14 | 426880 | 0.379717117 | 4.1044969 | 0.00102835 | 0.06527494 |
| ENSGALG00010023546 | SLC11A2 | 751817 | 0.339974241 | 7.04564835 | 0.00103194 | 0.06527494 |
| ENSGALG00010013056 | SPATA46 | 424370 | 0.434758077 | 3.55316316 | 0.0010367 | 0.06527494 |
| ENSGALG00010014931 | RNF145 | 416239 | -0.229094493 | 6.09853473 | 0.00103956 | 0.06527494 |
| ENSGALG00010000255 | NA | NA | 0.356188043 | 7.33370072 | 0.00108886 | 0.06809141 |
| ENSGALG00010001810 | OPN2SW | 396525 | 0.298996851 | 5.48423122 | 0.00110591 | 0.06865108 |
| ENSGALG00010022101 | CISD3 | 100857847 | 0.60900194 | 2.39083295 | 0.00111029 | 0.06865108 |
| ENSGALG00010029556 | NPTX1 | 422078 | 0.47439528 | 3.56667067 | 0.00111125 | 0.06865108 |
| ENSGALG00010011206 | LOC112532760 | 112532760 | -0.445066526 | 3.25348306 | 0.00114854 | 0.07066986 |
| ENSGALG00010020540 | PCGF5 | 423796 | -0.344119203 | 4.23196181 | 0.0011568 | 0.07089343 |
| ENSGALG00010016599 | LOC416263 | 416263 | 0.433492798 | 3.48479808 | 0.00118752 | 0.07232315 |
| ENSGALG00010029524 | BAHCC1 | 776014 | 0.395659422 | 4.35758818 | 0.00118957 | 0.07232315 |
| ENSGALG00010000636 | ZNF646 | 121108148 | 0.285559764 | 5.74628642 | 0.00120114 | 0.07273748 |
| ENSGALG00010015674 | RFKL | 431449 | -0.280795758 | 6.83845057 | 0.00120677 | 0.07279102 |
| ENSGALG00010009878 | NA | NA | -0.587982866 | 2.71308739 | 0.00121153 | 0.07279147 |
| ENSGALG00010025044 | TMEM9 | 421157 | 0.648007304 | 1.63719334 | 0.00124087 | 0.07426321 |
| ENSGALG00010023967 | PRPH | 101748212 | 0.420581945 | 3.41745358 | 0.00127656 | 0.07600475 |
| ENSGALG00010000060 | HYPK | 770307 | 0.241467289 | 6.22274914 | 0.00128584 | 0.07600475 |
| ENSGALG00010007762 | SH2D4A | 422487 | -0.957509214 | 0.42407824 | 0.00129422 | 0.07600475 |
| ENSGALG00010024965 | STAT2 | 100858732 | 0.393785341 | 3.76530814 | 0.00129862 | 0.07600475 |
| ENSGALG00010010189 | NA | NA | -0.777538398 | 3.7712161 | 0.00129907 | 0.07600475 |
| ENSGALG00010009863 | NA | NA | -0.984199816 | 0.58329526 | 0.00129974 | 0.07600475 |
| ENSGALG00010028301 | FBN3 | 395079 | 0.349879711 | 6.01553389 | 0.00130476 | 0.07600844 |
| ENSGALG00010023287 | RGS19 | 374070 | 0.596064785 | 1.81069214 | 0.00132522 | 0.0766311 |

| | | | | | | |
|---|---|---|---|---|---|---|
| ENSGALG00010029424 | MAP2K6 | 417445 | 0.494177432 | 3.46861137 | 0.00133088 | 0.0766311 |
| ENSGALG00010026881 | LOC121111826 | 121111826 | -0.244450006 | 6.08553562 | 0.00133115 | 0.0766311 |
| ENSGALG00010008272 | DCP2 | 769972 | -0.243673899 | 6.1792425 | 0.00133591 | 0.0766311 |
| ENSGALG00010021413 | NA | NA | 0.389176835 | 4.08932808 | 0.00134046 | 0.0766311 |
| ENSGALG00010020293 | SEMA4B | 415582 | 0.422831897 | 3.70666559 | 0.00134573 | 0.07664669 |
| ENSGALG00010023317 | RAPGEF3 | 426888 | 0.765899046 | 3.53686685 | 0.00136601 | 0.07746166 |
| ENSGALG00010029547 | HIP1 | 770683 | 0.287393442 | 6.1614237 | 0.00137015 | 0.07746166 |
| ENSGALG00010022923 | CD63 | 107049249 | 0.278185994 | 5.81349628 | 0.00139047 | 0.07815187 |
| ENSGALG00010021477 | LOC100858626 | 100858626 | 0.4904828 | 3.46624774 | 0.00139256 | 0.07815187 |
| ENSGALG00010029591 | SUZ12 | 417406 | -0.24086331 | 6.29084825 | 0.00140057 | 0.07823388 |
| ENSGALG00010015503 | NA | NA | -1.290413294 | -0.2253607 | 0.0014099 | 0.07823388 |
| ENSGALG00010027213 | KCNC4 | 419806 | 0.391412445 | 4.01799012 | 0.00141065 | 0.07823388 |
| ENSGALG00010005803 | NA | NA | -0.310176445 | 5.6456744 | 0.00141445 | 0.07823388 |
| ENSGALG00010011450 | PTDSS1 | 428374 | -0.275348972 | 5.29745104 | 0.00142193 | 0.07836471 |
| ENSGALG00010008126 | CDH3 | 414845 | 0.249403615 | 6.68920926 | 0.00143523 | 0.07881415 |
| ENSGALG00010010174 | NA | NA | -1.170185063 | 2.57935295 | 0.00144333 | 0.07897584 |
| ENSGALG00010013727 | NA | NA | -1.345956376 | -0.2575697 | 0.00145292 | 0.07921776 |
| ENSGALG00010006910 | NA | NA | -0.808810326 | 1.07753689 | 0.00146898 | 0.07980961 |
| ENSGALG00010027473 | DNAH1 | 415943 | 0.361530169 | 4.26511373 | 0.00148225 | 0.08021251 |
| ENSGALG00010004343 | NA | NA | -0.314795984 | 7.56056137 | 0.00149009 | 0.08021251 |
| ENSGALG00010013168 | DSCR3 | 418519 | 0.260082225 | 5.09015963 | 0.00149211 | 0.08021251 |
| ENSGALG00010003155 | NA | NA | -0.526412136 | 2.63175522 | 0.00152392 | 0.08162995 |
| ENSGALG00010014608 | NA | NA | -1.079245673 | 0.0714221 | 0.00152913 | 0.08162995 |
| ENSGALG00010019724 | HES5 | 419392 | -0.945863809 | 0.50526944 | 0.00154454 | 0.08216608 |
| ENSGALG00010025933 | SKIV2L | 421171 | 0.290333306 | 6.14261972 | 0.00156817 | 0.08313452 |
| ENSGALG00010027382 | PALM | 101750199 | 0.478342212 | 2.6554254 | 0.00163205 | 0.08620115 |
| ENSGALG00010009168 | MRPS12 | 770472 | 0.379968876 | 4.0343661 | 0.00163727 | 0.08620115 |
| ENSGALG00010011866 | KLF8 | 101749773 | 0.335331807 | 4.13943353 | 0.00165774 | 0.08652749 |
| ENSGALG00010008284 | BCKDHA | 374210 | 0.299887735 | 4.88276092 | 0.00165991 | 0.08652749 |
| ENSGALG00010015407 | TIMP3 | 396483 | -0.392157249 | 3.45904956 | 0.00166281 | 0.08652749 |
| ENSGALG00010017491 | CCDC28B | 419647 | 0.54529946 | 2.3463189 | 0.00167043 | 0.08652749 |
| ENSGALG00010021172 | SLC29A4 | 425593 | 0.305188572 | 6.01915173 | 0.0016717 | 0.08652749 |
| ENSGALG00010009646 | MVD | 425359 | 0.271652442 | 5.97614471 | 0.0016808 | 0.08664629 |
| ENSGALG00010014606 | NA | NA | -0.598282842 | 2.9689042 | 0.00168531 | 0.08664629 |
| ENSGALG00010004416 | STXBP6 | 423299 | -0.328410977 | 5.12355601 | 0.00172097 | 0.08814286 |
| ENSGALG00010014712 | NA | NA | -0.741272319 | 1.05675126 | 0.00173327 | 0.08814286 |
| ENSGALG00010024559 | NA | NA | -0.435483762 | 3.15208568 | 0.00173641 | 0.08814286 |
| ENSGALG00010017368 | ERLIN2 | 426769 | 0.276856768 | 8.83180608 | 0.00173743 | 0.08814286 |
| ENSGALG00010025536 | LEXM | 424660 | -0.834054394 | 1.02228817 | 0.00177827 | 0.08972963 |
| ENSGALG00010000476 | LOC107050559 | 107050559 | 0.237774272 | 5.74319872 | 0.00178042 | 0.08972963 |
| ENSGALG00010008213 | ISCA1 | 427463 | -0.269125628 | 6.38133005 | 0.00179165 | 0.08986637 |
| ENSGALG00010010072 | NA | NA | -1.172953128 | -0.1928433 | 0.00180097 | 0.08986637 |
| ENSGALG00010003902 | LOC100857859 | 100857859 | 0.354268441 | 4.45643234 | 0.00180459 | 0.08986637 |
| ENSGALG00010019140 | NA | NA | -0.68471855 | 1.26931228 | 0.00180804 | 0.08986637 |
| ENSGALG00010024603 | PNPLA1 | 428259 | -1.164524233 | -0.1459138 | 0.00181246 | 0.08986637 |
| ENSGALG00010029170 | DCXR | 374066 | 0.515825879 | 3.034472 | 0.00182794 | 0.09034144 |
| ENSGALG00010007399 | WWC2 | 422555 | 0.258460407 | 5.76629785 | 0.00184747 | 0.09072489 |
| ENSGALG00010013214 | RAI14 | 427431 | -0.268007935 | 5.11822066 | 0.00184754 | 0.09072489 |
| ENSGALG00010022760 | ELAVL4 | 395634 | -0.241665171 | 8.3422338 | 0.00185593 | 0.09084549 |
| ENSGALG00010022481 | CORO1B | 107056441 | 0.276108605 | 6.17106537 | 0.00190309 | 0.09285749 |
| ENSGALG00010018440 | MEA1 | 107052849 | 0.329542173 | 5.04440837 | 0.00192438 | 0.09358113 |
| ENSGALG00010024944 | CNPY2 | 100858933 | 0.284997288 | 5.67927985 | 0.00193014 | 0.09358113 |
| ENSGALG00010012070 | DNAJC28 | 770607 | 0.473582842 | 2.65254781 | 0.00193761 | 0.0936471 |

| | | | | | | |
|---|---|---|---|---|---|---|
| ENSGALG00010023199 | RPS13 | 414782 | 0.239098402 | 8.62654939 | 0.00195426 | 0.09400648 |
| ENSGALG00010014562 | NA | NA | 0.548488873 | 2.38234192 | 0.00195732 | 0.09400648 |
| ENSGALG00010008333 | APC | 415607 | -0.221503707 | 7.77958573 | 0.00198043 | 0.09481905 |
| ENSGALG00010005241 | C4H4orf54 | 112532442 | -0.224916811 | 6.72090398 | 0.0019893 | 0.09494736 |
| ENSGALG00010018323 | GOT1 | 396261 | 0.253674049 | 5.75781187 | 0.00200034 | 0.09501804 |
| ENSGALG00010012301 | NA | NA | -0.5064635 | 3.35626126 | 0.00200803 | 0.09501804 |
| ENSGALG00010023058 | PPARA | 374120 | -0.676835148 | 1.69393967 | 0.00201231 | 0.09501804 |
| ENSGALG00010003358 | DNAJC15 | 418838 | -0.23701671 | 5.79311652 | 0.00201559 | 0.09501804 |
| ENSGALG00010009795 | NDUFA6 | 427897 | 0.277193931 | 4.9869803 | 0.00202605 | 0.0952181 |
| ENSGALG00010019310 | CRELD1 | 100858613 | 0.361940122 | 4.7627654 | 0.00206054 | 0.09639512 |
| ENSGALG00010017517 | LIN9 | 421316 | -0.249585791 | 5.02019788 | 0.00206967 | 0.09639512 |
| ENSGALG00010001285 | NA | NA | 0.362032176 | 3.6781371 | 0.00206997 | 0.09639512 |
| ENSGALG00010022770 | HAT1 | 374037 | -0.2408289 | 6.09490404 | 0.0020946 | 0.09704948 |
| ENSGALG00010010826 | NCOA1 | 422016 | 0.234453895 | 6.7208139 | 0.00209669 | 0.09704948 |
| ENSGALG00010007691 | LOC121108447 | 121108447 | -0.42949733 | 3.08431584 | 0.00213047 | 0.09831601 |
| ENSGALG00010021469 | CSPG4 | 425524 | 0.782356737 | 0.96634539 | 0.00213848 | 0.09838933 |
| ENSGALG00010002824 | NXPH1 | 420582 | 0.946011336 | 0.40275391 | 0.0022056 | 0.1011738 |
| ENSGALG00010016861 | FGR | 777583 | 0.535525053 | 2.86785536 | 0.00223841 | 0.10214074 |
| ENSGALG00010012259 | IL1RAPL2 | 422379 | -0.598983736 | 2.64628939 | 0.00224002 | 0.10214074 |
| ENSGALG00010020236 | SLC35G1 | 423800 | -0.416621379 | 2.98638806 | 0.00226389 | 0.10292293 |
| ENSGALG00010020292 | NPM2 | 107050133 | 0.311322501 | 4.3005352 | 0.00228119 | 0.10340282 |
| ENSGALG00010009809 | SMDT1 | 770755 | 0.306379839 | 4.30679052 | 0.00229189 | 0.10355826 |
| ENSGALG00010019194 | NA | NA | 0.460745169 | 3.06023371 | 0.00229814 | 0.10355826 |
| ENSGALG00010024690 | LOC100857858 | 100857858 | 0.30578407 | 6.42492994 | 0.00233992 | 0.10503612 |
| ENSGALG00010012091 | LOC121108229 | 121108229 | 0.742555329 | 4.09727855 | 0.00234873 | 0.10503612 |
| ENSGALG00010023218 | SSH3 | 101752044 | 0.405910311 | 3.65174805 | 0.0023515 | 0.10503612 |
| ENSGALG00010021362 | ARPC1B | 416490 | 0.553131558 | 2.19926821 | 0.00243186 | 0.10830958 |
| ENSGALG00010012441 | FST | 396119 | -0.361618754 | 3.68860807 | 0.00243908 | 0.10831624 |
| ENSGALG00010008215 | SELENOW | 100310814 | 0.233108443 | 7.11082844 | 0.00249916 | 0.11040945 |
| ENSGALG00010002586 | GPR63 | 100857561 | -0.221108288 | 6.67616629 | 0.00250738 | 0.11040945 |
| ENSGALG00010024907 | PHLDB1 | 419790 | 0.350704469 | 5.02872421 | 0.00250783 | 0.11040945 |
| ENSGALG00010009015 | TMEM231 | 425108 | -0.229015862 | 5.62850072 | 0.00252488 | 0.11052547 |
| ENSGALG00010017563 | ALK | 421297 | -0.434664698 | 3.82628881 | 0.0025249 | 0.11052547 |
| ENSGALG00010021316 | PLA2G12B | 423705 | -1.029574433 | 0.08335823 | 0.00254571 | 0.11096059 |
| ENSGALG00010021735 | PIGBOS1 | 105694155 | 0.323939856 | 4.08687028 | 0.00255627 | 0.11096059 |
| ENSGALG00010002968 | NA | NA | -0.924875032 | 0.403705 | 0.00255656 | 0.11096059 |
| ENSGALG00010003110 | BF1 | 693260 | 0.308544874 | 4.71981703 | 0.0025887 | 0.11203817 |
| ENSGALG00010007171 | NA | NA | -0.747095455 | 4.81654172 | 0.00260731 | 0.11252558 |
| ENSGALG00010027228 | SSPO | 420367 | 0.689571054 | 1.73371098 | 0.00261591 | 0.11257971 |
| ENSGALG00010001024 | NA | NA | -0.518169888 | 3.9194878 | 0.00264054 | 0.11332139 |
| ENSGALG00010018556 | MRPL2 | 421256 | 0.300750288 | 5.49561124 | 0.00269579 | 0.11512007 |
| ENSGALG00010016550 | FBLIM1 | 419458 | 0.323540452 | 4.1655289 | 0.00269748 | 0.11512007 |
| ENSGALG00010017609 | TRIM71 | 428445 | -0.224615975 | 9.75402667 | 0.00272107 | 0.11570245 |
| ENSGALG00010018296 | PIK3CA | 424971 | -0.236594088 | 6.28987484 | 0.00272623 | 0.11570245 |
| ENSGALG00010004701 | NA | NA | -0.572963212 | 2.90441986 | 0.00279262 | 0.11819275 |
| ENSGALG00010021741 | RPL19 | 420003 | 0.23721701 | 9.81405325 | 0.00284088 | 0.11990378 |
| ENSGALG00010008933 | NA | NA | -0.392681271 | 5.27489461 | 0.00288261 | 0.121331 |
| ENSGALG00010020364 | MRPL16 | 428840 | 0.256205245 | 5.01806793 | 0.00290365 | 0.12185201 |
| ENSGALG00010024966 | AGAP3 | 768507 | 0.269672999 | 5.04980625 | 0.0029109 | 0.12185201 |
| ENSGALG00010017063 | TMPPE | 101750979 | 0.2749182 | 4.98171479 | 0.00295319 | 0.12328567 |
| ENSGALG00010014029 | NA | NA | -0.344849312 | 3.89364726 | 0.00297428 | 0.12347513 |
| ENSGALG00010025403 | MTERF4 | 107051861 | 0.362158307 | 3.98062019 | 0.00298083 | 0.12347513 |
| ENSGALG00010029598 | SUPT4H1 | 417470 | 0.228209727 | 5.34476857 | 0.00298191 | 0.12347513 |

| | | | | | | |
|---|---|---|---|---|---|---|
| ENSGALG00010025473 | PARS2 | 424659 | 0.22747735 | 5.93785855 | 0.00299066 | 0.12350376 |
| ENSGALG00010000835 | NA | NA | -0.400436085 | 5.3529253 | 0.00300722 | 0.1238537 |
| ENSGALG00010027423 | PEX11B | 112530296 | 0.363250802 | 3.83929688 | 0.00307066 | 0.12561032 |
| ENSGALG00010013379 | ZNF593 | 419549 | 0.246012984 | 5.89178411 | 0.00307127 | 0.12561032 |
| ENSGALG00010021797 | SLC25A34 | 107054933 | 0.438050227 | 3.14518502 | 0.00307447 | 0.12561032 |
| ENSGALG00010024789 | SUOX | 107055404 | 0.358114461 | 3.80092479 | 0.00310049 | 0.12633656 |
| ENSGALG00010025298 | NACAD | 420409 | 0.35215824 | 5.34845147 | 0.00311938 | 0.12665915 |
| ENSGALG00010027311 | NFASC | 419824 | 0.411640489 | 4.77578078 | 0.00312494 | 0.12665915 |
| ENSGALG00010025286 | ITGA3 | 373946 | 0.258257355 | 5.54841448 | 0.0031485 | 0.12718086 |
| ENSGALG00010014614 | GHR | 408184 | -0.288971448 | 4.92921266 | 0.00315441 | 0.12718086 |
| ENSGALG00010006091 | RTKN | 107056271 | 0.397678223 | 3.97249784 | 0.00317558 | 0.12748747 |
| ENSGALG00010009501 | HGH1 | 107050713 | 0.289359613 | 4.45515036 | 0.00317866 | 0.12748747 |
| ENSGALG00010011247 | CDYL | 420877 | -0.282458301 | 6.18107833 | 0.00320234 | 0.12810192 |
| ENSGALG00010000179 | NA | NA | 0.373826506 | 3.36974447 | 0.00322772 | 0.12878091 |
| ENSGALG00010028129 | CYB561D2 | 100857844 | 0.421477891 | 3.53202322 | 0.003314 | 0.13163893 |
| ENSGALG00010000623 | LOC101748630 | 101748630 | 0.317151365 | 4.40616622 | 0.00331654 | 0.13163893 |
| ENSGALG00010014442 | SHISA3 | 100857319 | 0.657270167 | 1.34232307 | 0.00332769 | 0.13174021 |
| ENSGALG00010018851 | NA | NA | -0.228331984 | 6.42501137 | 0.00335025 | 0.13177443 |
| ENSGALG00010028485 | USF1 | 425126 | 0.236382457 | 6.12129223 | 0.00335636 | 0.13177443 |
| ENSGALG00010029293 | DNAH17 | 768861 | 0.342355074 | 5.5612904 | 0.00335777 | 0.13177443 |
| ENSGALG00010024234 | ARHGEF4 | 424761 | 0.997605439 | 0.23471743 | 0.00337064 | 0.13177443 |
| ENSGALG00010003170 | C4 | 426611 | 0.226679851 | 6.41022413 | 0.00337155 | 0.13177443 |
| ENSGALG00010017778 | PPP3R1 | 378804 | -0.241362842 | 7.41175387 | 0.00338756 | 0.13206295 |
| ENSGALG00010020691 | BOLA3 | 107054980 | 0.216092487 | 6.49936225 | 0.00341406 | 0.13275841 |
| ENSGALG00010001570 | SMPD1 | 419077 | 0.425380161 | 3.44096978 | 0.0034278 | 0.1329551 |
| ENSGALG00010026121 | GSTT1L | 769847 | 0.406846337 | 3.22576952 | 0.00345831 | 0.13377484 |
| ENSGALG00010002061 | NA | NA | -0.374488665 | 4.44728705 | 0.00346787 | 0.13377484 |
| ENSGALG00010000119 | LOC124416997 | 124416997 | 0.699468739 | 2.76512481 | 0.00348287 | 0.13377484 |
| ENSGALG00010015390 | CCNB3 | 396167 | 0.276239044 | 5.53553346 | 0.0034844 | 0.13377484 |
| ENSGALG00010001605 | APBB1 | 100859045 | 0.307724876 | 4.7014562 | 0.00349259 | 0.13377484 |
| ENSGALG00010006580 | ALDH6A1 | 423345 | 0.29613389 | 4.24792417 | 0.003517 | 0.13414081 |
| ENSGALG00010006025 | TNFAIP3 | 421684 | -0.574722454 | 2.32497591 | 0.00352378 | 0.13414081 |
| ENSGALG00010018858 | EPB41L5 | 424235 | 0.199574879 | 7.72374236 | 0.00352841 | 0.13414081 |
| ENSGALG00010020450 | ANKRD39 | 107057371 | 0.433063604 | 3.05459036 | 0.00354818 | 0.13455865 |
| ENSGALG00010014997 | RAE1 | 419324 | -0.232723098 | 7.24035548 | 0.00358919 | 0.13534988 |
| ENSGALG00010025713 | LOC107054297 | 107054297 | -0.591668037 | 5.02073312 | 0.003593 | 0.13534988 |
| ENSGALG00010021659 | SPOCK2 | 100858414 | 0.275637863 | 5.63583688 | 0.00359555 | 0.13534988 |
| ENSGALG00010009470 | ZNF469 | 101748749 | -0.399774058 | 3.70868045 | 0.00360988 | 0.13555632 |
| ENSGALG00010006172 | TTC31 | 100858003 | 0.264324403 | 4.87627651 | 0.00362683 | 0.13581345 |
| ENSGALG00010027329 | FAM188B | 420383 | 0.350382801 | 3.57168647 | 0.00363446 | 0.13581345 |
| ENSGALG00010015239 | NA | NA | 0.239673836 | 5.5712575 | 0.00366988 | 0.1368036 |
| ENSGALG00010023080 | MCRS1 | 426664 | 0.245567564 | 6.85437791 | 0.00371407 | 0.13811477 |
| ENSGALG00010003724 | NA | NA | 0.428310034 | 4.1642739 | 0.00373382 | 0.13851312 |
| ENSGALG00010017200 | KCNQ4 | 419643 | 0.340010565 | 4.163534 | 0.00378834 | 0.13993789 |
| ENSGALG00010011825 | FAM19A2 | 771745 | -0.294375034 | 5.54476977 | 0.00379621 | 0.13993789 |
| ENSGALG00010025316 | SLC25A36 | 424817 | -0.240943195 | 6.91734672 | 0.00380523 | 0.13993789 |
| ENSGALG00010004389 | ZNF71 | 425244 | 0.396996942 | 3.57272598 | 0.00380877 | 0.13993789 |
| ENSGALG00010019082 | POLR3K | 768505 | 0.367668494 | 3.59743789 | 0.00385866 | 0.14143174 |
| ENSGALG00010021010 | LEO1 | 769405 | 0.211214476 | 6.91461552 | 0.00388688 | 0.14212603 |
| ENSGALG00010021837 | NA | NA | -1.327983983 | 0.40009201 | 0.00390275 | 0.14219226 |
| ENSGALG00010028085 | GPATCH4 | 100859444 | 0.223146463 | 6.02924171 | 0.00390725 | 0.14219226 |
| ENSGALG00010024745 | PMEL | 396007 | 0.327965384 | 5.88928018 | 0.00393633 | 0.14291116 |
| ENSGALG00010009205 | SIRT2 | 548628 | 0.276420376 | 5.47476219 | 0.003974 | 0.14337965 |

| | | | | | | |
|---|---|---|---|---|---|---|
| ENSGALG00010023053 | MAN1A2 | 418265 | -0.207388739 | 6.68427963 | 0.00397606 | 0.14337965 |
| ENSGALG00010026488 | NA | NA | 1.684850108 | 3.30473931 | 0.0039911 | 0.14337965 |
| ENSGALG00010027751 | ANGPTL4 | 769087 | 0.31281346 | 4.78920244 | 0.00399505 | 0.14337965 |
| ENSGALG00010020149 | FAHD2A | 426684 | 0.294082495 | 4.91577513 | 0.00401096 | 0.14337965 |
| ENSGALG00010028054 | RPS15 | 396448 | 0.243142041 | 8.96347265 | 0.00401247 | 0.14337965 |
| ENSGALG00010016415 | RASGRF1 | 415578 | 0.708567626 | 0.97971445 | 0.00401474 | 0.14337965 |
| ENSGALG00010024218 | NGFR | 425805 | 0.409062237 | 5.06197457 | 0.00403525 | 0.143777 |
| ENSGALG00010028602 | ZNF687 | 429831 | 0.244248983 | 5.73312336 | 0.00409391 | 0.14552864 |
| ENSGALG00010027814 | DAPK3 | 101748881 | 0.23938133 | 5.70106228 | 0.00412073 | 0.14614268 |
| ENSGALG00010024625 | SPRYD3 | 100858430 | 0.263388179 | 5.46766931 | 0.00414633 | 0.14658478 |
| ENSGALG00010006623 | LOC107052071 | 107052071 | -0.582117288 | 2.05007739 | 0.00415233 | 0.14658478 |
| ENSGALG00010007899 | APRT | 100857155 | 0.295487158 | 4.34827646 | 0.00416943 | 0.14685019 |
| ENSGALG00010001253 | ZBBX | 771469 | 0.304401148 | 3.96422549 | 0.00419865 | 0.14753999 |
| ENSGALG00010022111 | PCGF2 | 425919 | 0.724448924 | 1.13633802 | 0.00421649 | 0.14782795 |
| ENSGALG00010003098 | NA | NA | -0.380855167 | 3.2945162 | 0.00422875 | 0.14791945 |
| ENSGALG00010002036 | NA | NA | -0.321985229 | 4.00289608 | 0.00425661 | 0.14828766 |
| ENSGALG00010029610 | ERN1 | 417414 | -0.238769482 | 5.39545076 | 0.00425864 | 0.14828766 |
| ENSGALG00010025368 | LRP1 | 396170 | 0.236390398 | 9.1956936 | 0.00429096 | 0.14863625 |
| ENSGALG00010029211 | SDF2 | 427832 | 0.30297834 | 5.1139832 | 0.00429291 | 0.14863625 |
| ENSGALG00010021534 | KIFC2 | 415630 | 0.249222416 | 5.15555171 | 0.00429775 | 0.14863625 |
| ENSGALG00010000532 | JPH2 | 770867 | 0.553541272 | 2.30025975 | 0.00431754 | 0.14888053 |
| ENSGALG00010019049 | POLD2 | 107057377 | 0.255317871 | 6.17663704 | 0.00432425 | 0.14888053 |
| ENSGALG00010005830 | USP12 | 428078 | 0.266751772 | 4.89190271 | 0.00433554 | 0.14893461 |
| ENSGALG00010017647 | ITGA9 | 420757 | 0.409035634 | 3.48599228 | 0.00440042 | 0.15069286 |
| ENSGALG00010009160 | NA | NA | -1.094860958 | -0.2407951 | 0.0044064 | 0.15069286 |
| ENSGALG00010003468 | ILVBL | 101747528 | 0.297840891 | 4.92484473 | 0.00442374 | 0.1509491 |
| ENSGALG00010011559 | MTX3 | 427176 | -0.263095363 | 4.99461518 | 0.00443654 | 0.15104923 |
| ENSGALG00010028599 | VPS72 | 425660 | 0.257861978 | 6.28331858 | 0.00455408 | 0.15470752 |
| ENSGALG00010028240 | NA | NA | -0.730032554 | 1.35759468 | 0.00456857 | 0.15485647 |
| ENSGALG00010009493 | MXD3 | 427629 | 0.299679801 | 4.47872271 | 0.0045907 | 0.15491508 |
| ENSGALG00010005995 | NA | NA | -0.61369575 | 1.45053142 | 0.00459893 | 0.15491508 |
| ENSGALG00010017738 | NDUFS5 | 771510 | 0.217370016 | 5.61336751 | 0.00460212 | 0.15491508 |
| ENSGALG00010022565 | ARF3 | 107049507 | 0.211903125 | 5.88945418 | 0.00461075 | 0.15491508 |
| ENSGALG00010000693 | NA | NA | 0.249700258 | 5.72172828 | 0.00463041 | 0.15523523 |
| ENSGALG00010011386 | CTSB | 396329 | 0.233496502 | 7.02586431 | 0.00464156 | 0.15526913 |
| ENSGALG00010026879 | NA | NA | -0.353838867 | 3.55877089 | 0.00467163 | 0.15559622 |
| ENSGALG00010014176 | ABHD17B | 427252 | -0.298213671 | 5.50406578 | 0.00467164 | 0.15559622 |
| ENSGALG00010014170 | CEMIP2 | 427250 | -0.215492549 | 7.38639429 | 0.00469734 | 0.15611282 |
| ENSGALG00010023833 | PIP4K2C | 121108118 | 0.268637171 | 5.28623212 | 0.0047155 | 0.1562807 |
| ENSGALG00010000051 | NA | NA | 0.436094821 | 2.51157711 | 0.0047228 | 0.1562807 |
| ENSGALG00010012015 | LOC424014 | 424014 | 2.090732483 | 2.0051881 | 0.00475946 | 0.15715449 |
| ENSGALG00010003400 | PKIA | 396333 | -0.280845777 | 5.11506599 | 0.00482268 | 0.15889863 |
| ENSGALG00010023179 | KAT2A | 374232 | 0.225573797 | 5.8134217 | 0.00483302 | 0.15889863 |
| ENSGALG00010013470 | RPL31 | 418710 | 0.20192752 | 8.04207479 | 0.00488536 | 0.15979533 |
| ENSGALG00010017702 | ECEL1 | 424937 | 0.406206878 | 3.55662059 | 0.00488781 | 0.15979533 |
| ENSGALG00010023514 | ULK3 | 769780 | 0.256634649 | 4.74907598 | 0.00489793 | 0.15979533 |
| ENSGALG00010013502 | PAFAH2 | 769963 | 0.86846158 | 0.47476162 | 0.00490343 | 0.15979533 |
| ENSGALG00010003271 | AKAP11 | 418835 | -0.223996484 | 5.85899438 | 0.00491354 | 0.15979533 |
| ENSGALG00010014167 | SLC38A9 | 427139 | -0.220619046 | 6.13328936 | 0.00492288 | 0.15979533 |
| ENSGALG00010009489 | LOC101749352 | 101749352 | 0.225497706 | 6.73314599 | 0.0049695 | 0.16073647 |
| ENSGALG00010028032 | CAMK4L | 427267 | 0.343321819 | 3.8072195 | 0.00497285 | 0.16073647 |
| ENSGALG00010009552 | NA | NA | -0.78987691 | 0.78985439 | 0.00501156 | 0.16164639 |
| ENSGALG00010020160 | SYT7 | 101748423 | 0.300973952 | 4.80563499 | 0.00503424 | 0.16184836 |

| | | | | | | |
|---|---|---|---|---|---|---|
| ENSGALG00010009361 | LRRK2 | 427844 | -0.26920896 | 4.451032 | 0.0050475 | 0.16184836 |
| ENSGALG00010011096 | LDB2 | 395631 | 0.334149532 | 3.6825841 | 0.0050556 | 0.16184836 |
| ENSGALG00010002857 | DCAF15 | 112530997 | 0.238421169 | 5.27979609 | 0.00506007 | 0.16184836 |
| ENSGALG00010023874 | HEXIM1 | 100857796 | 0.220875668 | 6.44902041 | 0.00512195 | 0.16348608 |
| ENSGALG00010009170 | RPL26L1 | 396400 | 0.229781891 | 7.83577282 | 0.00517358 | 0.16477939 |
| ENSGALG00010025428 | VANGL1 | 418337 | -0.198789981 | 7.12383918 | 0.00518397 | 0.16477939 |
| ENSGALG00010024389 | AAAS | 100859661 | 0.224269557 | 6.37477436 | 0.00519816 | 0.16488825 |
| ENSGALG00010001878 | XK | 427991 | -0.375777095 | 3.44472986 | 0.00527113 | 0.16661223 |
| ENSGALG00010028194 | LOC107055115 | 107055115 | 0.4340676 | 3.80655001 | 0.00527426 | 0.16661223 |
| ENSGALG00010029207 | SPAG5 | 771702 | 0.219106286 | 7.57484517 | 0.00528835 | 0.16671349 |
| ENSGALG00010027590 | SEMA3F | 374109 | 0.21575119 | 6.82105224 | 0.00530756 | 0.16697572 |
| ENSGALG00010022622 | MPPED2 | 421604 | -0.23592505 | 7.49906526 | 0.00533485 | 0.16701503 |
| ENSGALG00010027717 | ATXN7L2 | 429691 | 0.472311155 | 2.86834793 | 0.00534024 | 0.16701503 |
| ENSGALG00010028482 | CD320 | 420066 | 0.294456285 | 4.42088305 | 0.00534216 | 0.16701503 |
| ENSGALG00010029726 | SEPTIN9 | 417347 | 0.210058152 | 6.39469215 | 0.00535242 | 0.16701503 |
| ENSGALG00010025098 | ATP5B | 426673 | 0.237679986 | 9.56270182 | 0.00540086 | 0.16818417 |
| ENSGALG00010022518 | ECHS1 | 770828 | 0.353532583 | 4.48070709 | 0.00541804 | 0.16837687 |
| ENSGALG00010001209 | CDH7 | 374007 | -0.33866493 | 4.17103173 | 0.00546647 | 0.16953811 |
| ENSGALG00010016706 | RTN4 | 378790 | 0.216106169 | 8.21165904 | 0.00549931 | 0.17010836 |
| ENSGALG00010016791 | LRRTM1 | 771502 | -0.658602239 | 1.98851517 | 0.00550707 | 0.17010836 |
| ENSGALG00010023823 | NA | NA | -0.242090865 | 4.82967366 | 0.00556788 | 0.1708258 |
| ENSGALG00010016490 | NA | NA | 0.259785299 | 4.59528291 | 0.00557216 | 0.1708258 |
| ENSGALG00010004838 | NA | NA | -1.162481813 | -0.0992437 | 0.00557261 | 0.1708258 |
| ENSGALG00010005530 | NA | NA | -0.269536958 | 4.52321487 | 0.00557489 | 0.1708258 |
| ENSGALG00010024822 | RPS26 | 100857770 | 0.242155105 | 8.88818504 | 0.00560223 | 0.1710862 |
| ENSGALG00010015892 | UBE2U | 100859413 | 0.290289704 | 4.21517552 | 0.00561678 | 0.1710862 |
| ENSGALG00010020781 | SNN | 416429 | 0.293743522 | 4.15853436 | 0.00562551 | 0.1710862 |
| ENSGALG00010014797 | ZDHHC9 | 422139 | 0.255487445 | 7.21965298 | 0.00562806 | 0.1710862 |
| ENSGALG00010021688 | ITPR1 | 395412 | -0.234865773 | 5.85562443 | 0.00564011 | 0.17111315 |
| ENSGALG00010017528 | RAI1 | 427664 | 0.271351275 | 5.20364327 | 0.00567431 | 0.17181035 |
| ENSGALG00010026438 | TOMM6 | 419924 | 0.231028297 | 6.34976989 | 0.00571076 | 0.1725731 |
| ENSGALG00010019282 | IGFBP2 | 396315 | 0.225295691 | 5.69869726 | 0.00572274 | 0.17259476 |
| ENSGALG00010014980 | LOC121107687 | 121107687 | 0.248033824 | 5.62298128 | 0.0057957 | 0.17381 |
| ENSGALG00010029697 | GIT1 | 417584 | 0.273661682 | 6.45287645 | 0.00579945 | 0.17381 |
| ENSGALG00010024263 | PHKB | 415741 | 0.235027199 | 5.07336494 | 0.00580562 | 0.17381 |
| ENSGALG00010025348 | HIPK1 | 419880 | -0.199597347 | 6.11888738 | 0.00580842 | 0.17381 |
| ENSGALG00010022354 | FAR2 | 419043 | 0.321422749 | 4.06070655 | 0.00582757 | 0.17404332 |
| ENSGALG00010012426 | MIEF1 | 418012 | 0.220777148 | 5.06802078 | 0.00594438 | 0.17718656 |
| ENSGALG00010018484 | NLGN1 | 424980 | -0.259199589 | 4.8871464 | 0.00597546 | 0.177767 |
| ENSGALG00010021263 | STOML1 | 415303 | 0.489542006 | 2.38402792 | 0.00602008 | 0.17874745 |
| ENSGALG00010003280 | NA | NA | -0.294320772 | 4.4513241 | 0.00604454 | 0.17912647 |
| ENSGALG00010013446 | ATP5I | 769146 | 0.205232045 | 6.24349519 | 0.00607924 | 0.17980689 |
| ENSGALG00010016378 | WDR76 | 426208 | 0.209399203 | 6.22939488 | 0.00612618 | 0.18084631 |
| ENSGALG00010022956 | PBLD | 423685 | -0.341994364 | 3.62705257 | 0.00613936 | 0.18088665 |
| ENSGALG00010024910 | UBTF | 100859792 | 0.210105798 | 7.25343943 | 0.00617991 | 0.18173218 |
| ENSGALG00010024158 | ALDH3B1 | 428812 | 0.305283598 | 4.36048801 | 0.00621403 | 0.18238521 |
| ENSGALG00010015013 | SNX18 | 768969 | -0.233823723 | 5.04522084 | 0.00622622 | 0.18239376 |
| ENSGALG00010024204 | PTBP2 | 424479 | -0.262530619 | 7.04589957 | 0.00626834 | 0.1826578 |
| ENSGALG00010013815 | SRXN1 | 100858692 | 0.601594327 | 1.78339058 | 0.00628617 | 0.1826578 |
| ENSGALG00010020140 | SECISBP2L | 415435 | -0.228248479 | 5.1465461 | 0.00628903 | 0.1826578 |
| ENSGALG00010024121 | NRG4 | 415354 | -0.640985475 | 1.11666108 | 0.00628983 | 0.1826578 |
| ENSGALG00010001025 | NA | NA | -0.476887323 | 4.58972238 | 0.00630438 | 0.1826578 |
| ENSGALG00010021721 | TP53I11 | 426399 | 0.197702472 | 8.11529697 | 0.00631338 | 0.1826578 |

| | | | | | | |
|---|---|---|---|---|---|---|
| ENSGALG00010006944 | MAB21L2 | 374011 | 0.543640251 | 1.85726889 | 0.00632614 | 0.1826578 |
| ENSGALG00010027731 | OLFML2A | 417098 | 0.240030447 | 5.29296379 | 0.0063473 | 0.1826578 |
| ENSGALG00010001835 | MECP2 | 112530992 | 0.231570464 | 6.30795915 | 0.00635913 | 0.1826578 |
| ENSGALG00010029143 | SH2D3C | 417225 | 0.574199737 | 1.48614422 | 0.00636442 | 0.1826578 |
| ENSGALG00010027955 | LOC107050229 | 107050229 | 0.271526936 | 5.24764847 | 0.00636638 | 0.1826578 |
| ENSGALG00010022165 | FAM192A | 415647 | 0.259860845 | 6.37107999 | 0.0063917 | 0.18304141 |
| ENSGALG00010010924 | ADAMTSL2L | 424420 | 0.291491487 | 5.06141386 | 0.00643431 | 0.18315246 |
| ENSGALG00010004412 | NA | NA | -0.399462682 | 2.9430103 | 0.00645088 | 0.18315246 |
| ENSGALG00010007998 | RPL13 | 395849 | 0.25823554 | 8.94493964 | 0.00645527 | 0.18315246 |
| ENSGALG00010010843 | ZNF395 | 422017 | 0.234723726 | 6.16116626 | 0.00647997 | 0.18315246 |
| ENSGALG00010000286 | TCN2 | 429737 | 0.538291865 | 1.86636123 | 0.00648055 | 0.18315246 |
| ENSGALG00010000005 | NA | NA | -0.414482604 | 11.5511753 | 0.00648722 | 0.18315246 |
| ENSGALG00010001612 | HPX | 419076 | 0.744092611 | 0.7334753 | 0.00648879 | 0.18315246 |
| ENSGALG00010029548 | BAIAP2 | 422077 | 0.326008604 | 4.25338554 | 0.00649121 | 0.18315246 |
| ENSGALG00010011597 | TBC1D14 | 422862 | -0.195347133 | 8.93956548 | 0.00651413 | 0.18346125 |
| ENSGALG00010027903 | NDUFA8 | 417112 | 0.214219501 | 5.79154005 | 0.00656076 | 0.18443572 |
| ENSGALG00010017909 | NTHL1 | 416551 | 0.263009277 | 4.78286607 | 0.00659 | 0.18449352 |
| ENSGALG00010013286 | RNF14 | 416340 | 0.195085536 | 6.33154622 | 0.00659139 | 0.18449352 |
| ENSGALG00010008205 | NA | NA | 0.553815097 | 2.0361497 | 0.00659895 | 0.18449352 |
| ENSGALG00010013044 | NOS1AP | 424371 | 0.408643034 | 6.07068852 | 0.00662657 | 0.18492847 |
| ENSGALG00010016411 | CTSH | 770109 | 0.267290253 | 4.45673661 | 0.0066454 | 0.18511672 |
| ENSGALG00010024166 | LOC772005 | 772005 | -0.530283689 | 1.81472727 | 0.00669592 | 0.18602071 |
| ENSGALG00010008897 | NSMCE2 | 420333 | -0.276733927 | 4.65603074 | 0.00670304 | 0.18602071 |
| ENSGALG00010012792 | CBY1 | 771364 | 0.34502133 | 3.60747192 | 0.00671428 | 0.18602071 |
| ENSGALG00010005333 | PLXNA4 | 427941 | 0.406172705 | 2.86717455 | 0.00674267 | 0.18609965 |
| ENSGALG00010022059 | NPTX2 | 416438 | 0.290266636 | 4.36177542 | 0.00675876 | 0.18609965 |
| ENSGALG00010000242 | CHIR2AB2 | 425238 | 0.32112555 | 3.68509983 | 0.00676176 | 0.18609965 |
| ENSGALG00010012749 | RGS4 | 378901 | 0.388618113 | 2.83681014 | 0.00676751 | 0.18609965 |
| ENSGALG00010015033 | BMP7 | 395494 | 0.273095994 | 4.48355376 | 0.00677786 | 0.18609965 |
| ENSGALG00010020653 | NDUFAF3 | 107054424 | 0.367289644 | 3.61614059 | 0.00679252 | 0.18616859 |
| ENSGALG00010024410 | C12orf10 | 426187 | 0.251737476 | 4.96633639 | 0.00683121 | 0.18682937 |
| ENSGALG00010000289 | QPRT | 107051294 | 0.283562145 | 4.85798499 | 0.00684102 | 0.18682937 |
| ENSGALG00010012245 | RNF151 | 420226 | 0.194227461 | 5.89007825 | 0.00686179 | 0.18706321 |
| ENSGALG00010013967 | MLLT3 | 427234 | -0.199393695 | 6.1968134 | 0.00690782 | 0.18798354 |
| ENSGALG00010013250 | E2F5 | 420206 | -0.222435175 | 5.15933846 | 0.0070978 | 0.19266524 |
| ENSGALG00010029495 | RPL38 | 771528 | 0.225474656 | 7.30629625 | 0.00710951 | 0.19266524 |
| ENSGALG00010020126 | NADK | 419403 | -0.246427974 | 5.82044861 | 0.00711759 | 0.19266524 |
| ENSGALG00010026981 | NA | NA | -1.101847783 | -0.3987863 | 0.0071826 | 0.19358164 |
| ENSGALG00010007933 | CEP126 | 100859489 | 1.167799791 | -0.5977835 | 0.0072121 | 0.19358164 |
| ENSGALG00010017456 | NA | NA | -0.448558318 | 2.34402125 | 0.00721393 | 0.19358164 |
| ENSGALG00010009149 | FUT4 | 428092 | 0.730388433 | 1.15426433 | 0.00722527 | 0.19358164 |
| ENSGALG00010029200 | GAS7 | 417305 | -0.219567 | 5.95669178 | 0.00723522 | 0.19358164 |
| ENSGALG00010019934 | TKFC | 426629 | 0.303415449 | 4.33718753 | 0.00723717 | 0.19358164 |
| ENSGALG00010008028 | KCNK7 | 421519 | 0.219115927 | 6.75273995 | 0.00723989 | 0.19358164 |
| ENSGALG00010010662 | MGST3 | 424404 | 0.211152473 | 5.59602523 | 0.00725343 | 0.19360596 |
| ENSGALG00010012125 | RNF152 | 420909 | -0.192250322 | 6.30418848 | 0.00730938 | 0.19476009 |
| ENSGALG00010009520 | SPINK4 | 768734 | 0.777890915 | 0.50467545 | 0.00736445 | 0.19557961 |
| ENSGALG00010013645 | RRAGC | 419605 | -0.198069793 | 6.64622375 | 0.00736986 | 0.19557961 |
| ENSGALG00010018081 | DIO3 | 395939 | 0.720472261 | 0.74118522 | 0.00737844 | 0.19557961 |
| ENSGALG00010021930 | NA | NA | -0.395547159 | 2.84064542 | 0.00741492 | 0.19585347 |
| ENSGALG00010003862 | PCK2 | 396457 | 0.242831002 | 5.87674556 | 0.00741663 | 0.19585347 |
| ENSGALG00010014680 | LOC107053920 | 107053920 | -0.198095205 | 7.44956829 | 0.00742712 | 0.19585347 |
| ENSGALG00010029241 | STXBP4 | 417393 | -0.196579247 | 5.77685682 | 0.00750156 | 0.19747662 |

| | | | | | | |
|---|---|---|---|---|---|---|
| ENSGALG00010029577 | NOG | 373912 | 0.921572433 | 0.32770185 | 0.00754817 | 0.1983629 |
| ENSGALG00010006075 | SOX21 | 107051027 | -0.20828816 | 6.80976564 | 0.00758686 | 0.19840054 |
| ENSGALG00010000449 | PLBD2 | 417031 | 0.303812205 | 4.88825733 | 0.0075903 | 0.19840054 |
| ENSGALG00010018582 | RCC2 | 419361 | 0.198961778 | 7.34590068 | 0.00759151 | 0.19840054 |
| ENSGALG00010005409 | PRKCQ | 776459 | 0.272025905 | 4.37195434 | 0.0076047 | 0.19840054 |
| ENSGALG00010021360 | PTPRN | 424201 | 0.509075679 | 1.85143342 | 0.00761435 | 0.19840054 |
| ENSGALG00010005492 | TNPO3 | 426277 | 0.220535394 | 7.00216805 | 0.00764746 | 0.19892488 |
| ENSGALG00010025645 | NBEAL2 | 425984 | 0.237383093 | 6.44585422 | 0.00767264 | 0.19924155 |
| ENSGALG00010016315 | RPS4Y1 | 396001 | 0.214997233 | 9.52975983 | 0.0076864 | 0.19926121 |
| ENSGALG00010024989 | TIMELESS | 101751192 | 0.211827472 | 6.27682872 | 0.0077151 | 0.19966732 |
| ENSGALG00010008327 | TGFB1 | 100873157 | 0.223103889 | 5.65640024 | 0.00772872 | 0.19968242 |
| ENSGALG00010000546 | FLRT1 | 112530908 | 0.399671824 | 2.78434115 | 0.00778616 | 0.20082788 |
| ENSGALG00010000452 | SLC8B1 | 417032 | 0.51328163 | 2.30380073 | 0.00782998 | 0.20161879 |
| ENSGALG00010013871 | NA | NA | 0.81440574 | 0.475525 | 0.00787697 | 0.20201674 |
| ENSGALG00010009777 | ST3GAL1 | 396140 | -0.313742106 | 3.67048794 | 0.00788678 | 0.20201674 |
| ENSGALG00010027979 | RPL12 | 417264 | 0.195229944 | 8.78011985 | 0.00788989 | 0.20201674 |
| ENSGALG00010027813 | LRRC71 | 100857743 | 0.332282162 | 3.59811376 | 0.00789818 | 0.20201674 |
| ENSGALG00010000619 | NA | NA | 0.30266378 | 4.16768756 | 0.0079571 | 0.20318449 |
| ENSGALG00010003850 | PLA2G12A | 772082 | -0.215538291 | 5.80360256 | 0.00801232 | 0.2035045 |
| ENSGALG00010009338 | KYNU | 424302 | -0.53963637 | 1.75931274 | 0.00801726 | 0.2035045 |
| ENSGALG00010014656 | NA | NA | -0.405785445 | 3.18299201 | 0.00802399 | 0.2035045 |
| ENSGALG00010015391 | SYN3 | 100857420 | 0.975142222 | -0.2418387 | 0.00803829 | 0.2035045 |
| ENSGALG00010027171 | LOC107055991 | 107055991 | 0.325775033 | 3.66822088 | 0.00805705 | 0.2035045 |
| ENSGALG00010019820 | BBOX1 | 426932 | 0.22491836 | 5.72628078 | 0.00805838 | 0.2035045 |
| ENSGALG00010003030 | PPEF1 | 427999 | 0.322043866 | 4.0393134 | 0.0080629 | 0.2035045 |
| ENSGALG00010021813 | WDR72 | 427497 | -0.709329656 | 0.92908445 | 0.00807589 | 0.2035045 |
| ENSGALG00010000063 | SERF2 | 107054222 | 0.214885245 | 6.01338876 | 0.00809247 | 0.20352215 |
| ENSGALG00010013506 | RNF149 | 418712 | -0.238038544 | 5.30802791 | 0.00810316 | 0.20352215 |
| ENSGALG00010023978 | TFDP2 | 424780 | -0.19282645 | 8.38638204 | 0.00812459 | 0.20353887 |
| ENSGALG00010009560 | RAB24 | 427627 | 0.223823241 | 5.12693323 | 0.0081304 | 0.20353887 |
| ENSGALG00010028073 | ST6GALNAC4 | 395182 | 0.366267488 | 3.65229702 | 0.00818707 | 0.20373893 |
| ENSGALG00010027261 | NA | NA | 0.239456237 | 5.07179105 | 0.00819405 | 0.20373893 |
| ENSGALG00010022269 | CNTD1 | 420019 | -0.220529034 | 5.08578815 | 0.00819798 | 0.20373893 |
| ENSGALG00010001694 | NA | NA | 0.673203865 | 3.24224554 | 0.00821283 | 0.20373893 |
| ENSGALG00010003762 | EVI5L | 430998 | 0.326220512 | 4.52743313 | 0.00821425 | 0.20373893 |
| ENSGALG00010025929 | NELFE | 100859071 | 0.235888859 | 5.21956472 | 0.00821818 | 0.20373893 |
| ENSGALG00010021459 | ARHGEF16 | 419386 | 0.278327728 | 4.48374976 | 0.00826415 | 0.20451069 |
| ENSGALG00010008103 | RBPMS | 428738 | 0.206336351 | 6.38035474 | 0.00827927 | 0.20451069 |
| ENSGALG00010023655 | C1R | 418295 | 0.285435442 | 4.28598354 | 0.00828935 | 0.20451069 |
| ENSGALG00010005489 | G2E3 | 423305 | -0.273689772 | 8.81487409 | 0.00831967 | 0.20492881 |
| ENSGALG00010024583 | LOC107049666 | 107049666 | 0.600077622 | 1.40728741 | 0.0085123 | 0.20907469 |
| ENSGALG00010000147 | LOC107050985 | 107050985 | 0.272292945 | 5.0794267 | 0.00851528 | 0.20907469 |
| ENSGALG00010027980 | NUP210 | 415977 | -0.192617278 | 6.41980793 | 0.00852976 | 0.20909503 |
| ENSGALG00010025329 | MYO1A | 396072 | 0.392241243 | 4.01934018 | 0.00858787 | 0.21018321 |
| ENSGALG00010015296 | RPL35A | 424924 | 0.222401649 | 8.80673365 | 0.00861667 | 0.21049157 |
| ENSGALG00010001529 | NA | NA | 0.286182654 | 4.27984815 | 0.00864286 | 0.21049157 |
| ENSGALG00010019521 | NA | NA | -0.350845092 | 4.20191537 | 0.00866801 | 0.21049157 |
| ENSGALG00010015175 | MAPK9 | 395983 | -0.229797453 | 5.84984606 | 0.00866802 | 0.21049157 |
| ENSGALG00010015816 | MITD1 | 418698 | -0.357297441 | 3.26452712 | 0.00870592 | 0.21049157 |
| ENSGALG00010018506 | LOC107051359 | 107051359 | 0.257897091 | 4.59468952 | 0.00872146 | 0.21049157 |
| ENSGALG00010001496 | PTP4A1 | 421877 | -0.226339633 | 7.72043263 | 0.00872173 | 0.21049157 |
| ENSGALG00010022240 | SPINT1 | 423206 | 0.243906213 | 4.59175579 | 0.00872448 | 0.21049157 |
| ENSGALG00010007183 | VOPP1 | 776358 | -0.204223074 | 5.43840367 | 0.0087325 | 0.21049157 |

| | | | | | | |
|---|---|---|---|---|---|---|
| ENSGALG00010020522 | NA | NA | -0.426228591 | 3.1654153 | 0.00873785 | 0.21049157 |
| ENSGALG00010014317 | RAD23B | 431623 | -0.21854183 | 8.53567729 | 0.00877155 | 0.21097168 |
| ENSGALG00010017541 | SREBF1 | 373915 | 0.241697333 | 6.23068357 | 0.00880158 | 0.21097972 |
| ENSGALG00010024238 | ATF7 | 769661 | 0.322286175 | 4.10731676 | 0.00881049 | 0.21097972 |
| ENSGALG00010015306 | BCAT1 | 418193 | 0.24238283 | 5.65719699 | 0.0088132 | 0.21097972 |
| ENSGALG00010021861 | NA | NA | 0.205177859 | 5.54894851 | 0.00885694 | 0.2116961 |
| ENSGALG00010028568 | MLLT1 | 420089 | 0.186635595 | 6.73792779 | 0.00887827 | 0.21187537 |
| ENSGALG00010022546 | MTG1 | 791224 | 0.268869892 | 4.48929099 | 0.00892339 | 0.21262097 |
| ENSGALG00010016127 | ACSL6 | 416324 | 0.194793603 | 7.2162312 | 0.0090464 | 0.21521715 |
| ENSGALG00010029581 | DGKE | 770911 | 0.196908375 | 5.967408 | 0.0091099 | 0.21602705 |
| ENSGALG00010028060 | NA | NA | 0.307635356 | 4.00002415 | 0.00912791 | 0.21602705 |
| ENSGALG00010009559 | NA | NA | -0.300354348 | 4.26692521 | 0.00912823 | 0.21602705 |
| ENSGALG00010024700 | TOLLIP | 423099 | -0.210615201 | 9.16343598 | 0.00913684 | 0.21602705 |
| ENSGALG00010011043 | NA | NA | -0.877371433 | 0.10924681 | 0.00915519 | 0.21612746 |
| ENSGALG00010004072 | SGK3 | 420167 | -0.185490038 | 5.80537431 | 0.00917847 | 0.21628244 |
| ENSGALG00010004429 | LOC112531814 | 112531814 | 0.767369962 | 0.47601989 | 0.00919052 | 0.21628244 |
| ENSGALG00010010430 | ASTN1 | 429070 | 0.290523072 | 4.54274528 | 0.00920411 | 0.21628244 |
| ENSGALG00010007985 | APELA | 770154 | 0.194957872 | 6.51644705 | 0.00925491 | 0.21671009 |
| ENSGALG00010028609 | MRPS21 | 426666 | 0.216646644 | 5.1768353 | 0.0092571 | 0.21671009 |
| ENSGALG00010016303 | ATP5E | 768660 | 0.249633394 | 5.42237291 | 0.00926474 | 0.21671009 |
| ENSGALG00010018266 | GNOT2 | 396117 | 0.235779678 | 5.61999467 | 0.00931654 | 0.21758959 |
| ENSGALG00010009367 | DMXL1 | 427385 | -0.199612036 | 6.21847694 | 0.00934721 | 0.21797361 |
| ENSGALG00010020682 | ZDHHC16 | 423843 | 0.255122179 | 4.79303303 | 0.00942697 | 0.21949944 |
| ENSGALG00010023060 | ALDH1A2 | 395844 | -0.236254735 | 4.53763822 | 0.00947542 | 0.2202928 |
| ENSGALG00010006352 | GEMIN2 | 423336 | 0.221555355 | 5.14796987 | 0.00953481 | 0.22133763 |
| ENSGALG00010000475 | RNFT2 | 427723 | 0.274644139 | 4.64185391 | 0.00957914 | 0.22140159 |
| ENSGALG00010021952 | NA | NA | -0.580622505 | 1.29820059 | 0.00961315 | 0.22140159 |
| ENSGALG00010027264 | LOC420374 | 420374 | 0.219466789 | 6.05034354 | 0.00964698 | 0.22140159 |
| ENSGALG00010000316 | NA | NA | -0.506034748 | 2.41622169 | 0.00965291 | 0.22140159 |
| ENSGALG00010027562 | SLC39A3 | 429050 | 0.356013145 | 3.6034152 | 0.00965549 | 0.22140159 |
| ENSGALG00010025252 | PPP1R19B | 100859037 | 0.247403278 | 4.7115577 | 0.00965978 | 0.22140159 |
| ENSGALG00010002790 | CXorf23 | 100858121 | -0.196452747 | 6.26483579 | 0.00965987 | 0.22140159 |
| ENSGALG00010014272 | PARD6B | 419352 | -0.287741885 | 3.99176532 | 0.00966262 | 0.22140159 |
| ENSGALG00010019334 | ATL1 | 423577 | 0.30094843 | 3.60342653 | 0.00970057 | 0.22140159 |
| ENSGALG00010003248 | MRI1 | 107049831 | 0.20483693 | 6.7605712 | 0.00971354 | 0.22140159 |
| ENSGALG00010017056 | AGTR1 | 396065 | -0.188280742 | 6.64347968 | 0.00971555 | 0.22140159 |
| ENSGALG00010009123 | EMILIN1 | 100858014 | 0.213911649 | 5.11944918 | 0.00972425 | 0.22140159 |
| ENSGALG00010013084 | SGSM3 | 418008 | 0.297801111 | 4.25867651 | 0.00972543 | 0.22140159 |
| ENSGALG00010026297 | NA | NA | -0.918400749 | 0.20966687 | 0.00985407 | 0.22372691 |
| ENSGALG00010019280 | MESDC1 | 415473 | 0.193215479 | 6.30242069 | 0.00989021 | 0.22372691 |
| ENSGALG00010012991 | MFSD9 | 418720 | -0.263881517 | 4.55565406 | 0.00989066 | 0.22372691 |
| ENSGALG00010000548 | NA | NA | 0.296871261 | 4.03356621 | 0.00990906 | 0.22372691 |
| ENSGALG00010023890 | SASS6 | 424471 | -0.207793686 | 5.92615295 | 0.00992314 | 0.22372691 |
| ENSGALG00010011298 | NA | NA | -0.549493446 | 1.94258135 | 0.00993291 | 0.22372691 |
| ENSGALG00010003528 | GDAP1 | 420191 | -0.341497705 | 4.13119225 | 0.00996196 | 0.22372691 |
| ENSGALG00010011694 | NA | NA | -0.493382398 | 2.25363531 | 0.0099635 | 0.22372691 |
| ENSGALG00010012970 | UNC5A | 426419 | 0.382137516 | 4.1788067 | 0.0099746 | 0.22372691 |
| ENSGALG00010014945 | TXN2 | 426978 | 0.229348441 | 5.12642001 | 0.009991 | 0.22372691 |
| ENSGALG00010001681 | NAPG | 421051 | -0.215408565 | 6.11047851 | 0.00999994 | 0.22372691 |
| ENSGALG00010020007 | NA | NA | -0.184914901 | 6.70487957 | 0.0100028 | 0.22372691 |
| ENSGALG00010017226 | PEF1 | 419644 | 0.226482485 | 5.07801371 | 0.01002749 | 0.22395213 |
| ENSGALG00010015526 | NA | NA | -0.297752994 | 3.83196586 | 0.01006332 | 0.22442513 |
| ENSGALG00010022608 | P3H4 | 430099 | 0.298362533 | 3.69103291 | 0.01008592 | 0.22460235 |

| | | | | | | |
|---|---|---|---|---|---|---|
| ENSGALG00010029703 | MIF4GD | 100858848 | 0.230008023 | 5.12537207 | 0.01015942 | 0.22591059 |
| ENSGALG00010009151 | CCNJL | 416155 | -0.212463748 | 6.10792888 | 0.01017741 | 0.22592252 |
| ENSGALG00010027184 | MPND | 100858729 | 0.620789349 | 1.65470839 | 0.01018944 | 0.22592252 |
| ENSGALG00010004765 | DMBT1 | 426819 | 0.27696375 | 5.17568475 | 0.01020913 | 0.22603193 |
| ENSGALG00010017509 | RASD1 | 416507 | 0.425649558 | 3.11353327 | 0.01023671 | 0.22631537 |
| ENSGALG00010023931 | CSRNP2 | 100859835 | 0.266146592 | 5.262288 | 0.01030521 | 0.22750155 |
| ENSGALG00010027543 | TUBB4B | 417255 | 0.21734405 | 8.49715773 | 0.01032616 | 0.22763613 |
| ENSGALG00010020619 | QARS | 416057 | 0.200540845 | 7.68582217 | 0.01035078 | 0.22785109 |
| ENSGALG00010026350 | LOC101749943 | 101749943 | -0.248844596 | 4.9581192 | 0.01037415 | 0.22803772 |
| ENSGALG00010020765 | GLT1D1 | 416807 | -0.202515529 | 6.87248666 | 0.01042614 | 0.22858044 |
| ENSGALG00010019392 | NIN | 428928 | 0.344821305 | 3.77855106 | 0.01044337 | 0.22858044 |
| ENSGALG00010013126 | NA | NA | -0.321169546 | 6.29962024 | 0.0104534 | 0.22858044 |
| ENSGALG00010009065 | EIF2B4 | 425800 | 0.23262202 | 5.96146156 | 0.01045851 | 0.22858044 |
| ENSGALG00010013258 | C8orf59 | 101751649 | 0.794748199 | 0.40290868 | 0.01048821 | 0.22890291 |
| ENSGALG00010017398 | SLC22A5 | 416328 | -0.201047827 | 5.57580939 | 0.01052867 | 0.22945915 |
| ENSGALG00010008096 | CPSF1 | 112531811 | 0.222493836 | 6.52894646 | 0.01056033 | 0.2298222 |
| ENSGALG00010021506 | MAX | 100858676 | 0.313791391 | 4.0787791 | 0.01068728 | 0.23193355 |
| ENSGALG00010017757 | SVBP | 101749360 | 0.234247069 | 5.95542877 | 0.01068762 | 0.23193355 |
| ENSGALG00010003126 | LOC430303 | 430303 | 0.449280734 | 3.14377497 | 0.01072771 | 0.23228293 |
| ENSGALG00010003006 | CYP7B1 | 420164 | -0.18951796 | 5.79789472 | 0.01073405 | 0.23228293 |
| ENSGALG00010028282 | NA | NA | -1.146730879 | -0.5559365 | 0.0107601 | 0.23234062 |
| ENSGALG00010004375 | NA | NA | -0.620241132 | 1.47680926 | 0.01076704 | 0.23234062 |
| ENSGALG00010010885 | SNRPD2 | 107049207 | 0.243770979 | 6.75618123 | 0.01081321 | 0.23255318 |
| ENSGALG00010011283 | LONRF3 | 422375 | -0.261461615 | 4.27672208 | 0.01082323 | 0.23255318 |
| ENSGALG00010028903 | ASTN2 | 417242 | 0.322040449 | 4.0686883 | 0.01084507 | 0.23255318 |
| ENSGALG00010001558 | CCKBR | 414885 | 0.529486804 | 1.74182582 | 0.01085124 | 0.23255318 |
| ENSGALG00010012879 | RCAN1 | 418510 | -0.226635257 | 5.6978038 | 0.01085806 | 0.23255318 |
| ENSGALG00010010720 | NA | NA | -0.499522589 | 2.08360484 | 0.01087817 | 0.23255318 |
| ENSGALG00010012538 | RAP1B | 417840 | 0.201653161 | 8.3830034 | 0.01091791 | 0.23255318 |
| ENSGALG00010006120 | MOGS | 107049491 | 0.288764698 | 4.70625196 | 0.01091804 | 0.23255318 |
| ENSGALG00010000790 | 6-Mar | 420927 | -0.187273579 | 7.81520487 | 0.01092191 | 0.23255318 |
| ENSGALG00010017332 | FGFR1 | 396516 | 0.199422302 | 7.09500288 | 0.01092868 | 0.23255318 |
| ENSGALG00010003292 | GADD45GIP1 | 107055382 | 0.231518431 | 5.33183176 | 0.01095445 | 0.23277815 |
| ENSGALG00010017595 | CCNF | 416749 | 0.200164443 | 6.94388907 | 0.01101116 | 0.23365917 |
| ENSGALG00010022154 | CPN1 | 769143 | 0.300769343 | 4.02649882 | 0.01104863 | 0.23413006 |
| ENSGALG00010007446 | NA | NA | -0.453708366 | 3.43915295 | 0.01110291 | 0.23495531 |
| ENSGALG00010007769 | NA | NA | -0.314388793 | 3.70007292 | 0.01121708 | 0.23704389 |
| ENSGALG00010021244 | PPP3CB | 378890 | -0.199607601 | 5.30014785 | 0.01126663 | 0.23776313 |
| ENSGALG00010004076 | COX7A2 | 772260 | 0.288811037 | 5.15313546 | 0.01128771 | 0.23788038 |
| ENSGALG00010010587 | NA | NA | 0.52948072 | 1.49482865 | 0.01131939 | 0.23808891 |
| ENSGALG00010023770 | SCRN2 | 425759 | 0.380596567 | 2.81694989 | 0.01134285 | 0.23808891 |
| ENSGALG00010011192 | RPL17 | 426845 | 0.208279671 | 9.16978069 | 0.01134971 | 0.23808891 |
| ENSGALG00010025080 | BAZ2A | 101747779 | 0.198633638 | 7.35342114 | 0.01135977 | 0.23808891 |
| ENSGALG00010027996 | GPX4 | 374056 | 0.202146841 | 7.34846357 | 0.01139211 | 0.23829263 |
| ENSGALG00010027605 | LOC101748987 | 101748987 | 0.321763208 | 3.61274642 | 0.01140902 | 0.23829263 |
| ENSGALG00010018876 | GMFB | 423560 | -0.19763321 | 7.34437724 | 0.01141615 | 0.23829263 |
| ENSGALG00010020992 | KCNH7 | 424184 | -0.27284818 | 4.70759167 | 0.01147667 | 0.23894108 |
| ENSGALG00010023638 | DCTN2 | 395587 | 0.197653709 | 6.99094732 | 0.01148982 | 0.23894108 |
| ENSGALG00010019634 | GNG2 | 423581 | 0.285182036 | 4.43745048 | 0.011494 | 0.23894108 |
| ENSGALG00010021599 | ARPC4 | 416051 | 0.231338473 | 5.68681555 | 0.01153745 | 0.23949793 |
| ENSGALG00010029069 | NA | NA | -1.072011752 | -0.4749009 | 0.01155205 | 0.23949793 |
| ENSGALG00010000233 | HORMAD2 | 417012 | 0.704890008 | 0.54428549 | 0.01158133 | 0.23978038 |
| ENSGALG00010016016 | ZDHHC2 | 422734 | -0.211119309 | 5.58423411 | 0.01160895 | 0.24002786 |

| | | | | | | |
|---|---|---|---|---|---|---|
| ENSGALG00010009453 | GSAP | 417724 | -0.230853643 | 4.75125611 | 0.01165118 | 0.24006136 |
| ENSGALG00010001629 | NA | NA | -0.387293135 | 3.34865598 | 0.01168811 | 0.24006136 |
| ENSGALG00010001991 | TOMM7 | 420610 | 0.234683819 | 5.55643752 | 0.01170279 | 0.24006136 |
| ENSGALG00010024882 | MYL6 | 100996929 | 0.221519772 | 7.5916446 | 0.01170295 | 0.24006136 |
| ENSGALG00010012670 | NA | NA | -0.231567762 | 6.15782478 | 0.01171404 | 0.24006136 |
| ENSGALG00010001617 | TRIM3 | 425131 | 0.276360592 | 4.64663433 | 0.01171563 | 0.24006136 |
| ENSGALG00010004928 | HDDC2 | 421719 | 0.224888544 | 5.14867334 | 0.01172025 | 0.24006136 |
| ENSGALG00010005464 | SLC39A8 | 430411 | -0.211482786 | 5.95797007 | 0.01181133 | 0.24160405 |
| ENSGALG00010024245 | TARBP2 | 107055413 | 0.243744656 | 4.5330086 | 0.0118353 | 0.2417715 |
| ENSGALG00010009129 | PRKACA | 100859181 | 0.366107198 | 3.18003903 | 0.01187041 | 0.24181562 |
| ENSGALG00010024477 | NA | NA | -0.872169489 | 0.06993149 | 0.01187207 | 0.24181562 |
| ENSGALG00010009037 | LOC100858919 | 100858919 | 0.870574995 | 0.08347516 | 0.01188481 | 0.24181562 |
| ENSGALG00010003470 | METTL3 | 107050363 | 0.21832791 | 6.22221994 | 0.01195545 | 0.24293027 |
| ENSGALG00010020434 | NA | NA | -0.552614966 | 1.44062822 | 0.01200047 | 0.24327227 |
| ENSGALG00010024082 | NA | NA | -0.638047721 | 1.1309121 | 0.01200404 | 0.24327227 |
| ENSGALG00010000660 | NA | NA | 0.247666091 | 5.15876165 | 0.01203261 | 0.24352921 |
| ENSGALG00010017201 | AP5M1 | 423548 | -0.209145086 | 4.88721604 | 0.01208965 | 0.24436089 |
| ENSGALG00010010366 | CLU | 395722 | 0.230022738 | 5.24411926 | 0.01215742 | 0.24487782 |
| ENSGALG00010016079 | NA | NA | -0.302899395 | 4.15673808 | 0.01217451 | 0.24487782 |
| ENSGALG00010027317 | MPV17L2 | 100857696 | 0.286693456 | 3.97297778 | 0.01219031 | 0.24487782 |
| ENSGALG00010017407 | EPN2 | 416524 | 0.191879056 | 6.14169666 | 0.01220658 | 0.24487782 |
| ENSGALG00010001116 | RNF13 | 396303 | 0.203491751 | 7.76828719 | 0.01223334 | 0.24487782 |
| ENSGALG00010027355 | ADCYAP1R1 | 420386 | -0.177447521 | 7.88463092 | 0.01223526 | 0.24487782 |
| ENSGALG00010017664 | MARK1 | 421342 | -0.225309539 | 5.11722147 | 0.01225343 | 0.24487782 |
| ENSGALG00010011726 | WDR55 | 100858589 | 0.208110432 | 5.95040886 | 0.012282 | 0.24487782 |
| ENSGALG00010024893 | ARL6IP4 | 427695 | 0.204586684 | 5.63211279 | 0.0122826 | 0.24487782 |
| ENSGALG00010023281 | B2M | 414830 | 0.196070591 | 6.31569212 | 0.01228536 | 0.24487782 |
| ENSGALG00010017957 | THUMPD1 | 107049049 | -0.176212695 | 7.56192402 | 0.01229104 | 0.24487782 |
| ENSGALG00010012742 | TUBB2A | 768337 | 0.226111901 | 6.00874844 | 0.01233686 | 0.24547154 |
| ENSGALG00010026874 | TEAD3 | 395810 | 0.215016833 | 5.08723482 | 0.01235861 | 0.2455853 |
| ENSGALG00010005742 | GPR183 | 769212 | -0.71190687 | 0.67368828 | 0.01239863 | 0.24585809 |
| ENSGALG00010011685 | NDUFA2 | 768860 | 0.28388151 | 4.26078835 | 0.01240443 | 0.24585809 |
| ENSGALG00010029329 | LOC101747660 | 101747660 | 0.427387639 | 2.1706074 | 0.01247183 | 0.24687446 |
| ENSGALG00010019257 | SLC1A4 | 769612 | -0.202002525 | 7.93388102 | 0.01250404 | 0.24719271 |
| ENSGALG00010009444 | NFIL3 | 395326 | -0.188766849 | 5.80140358 | 0.01255871 | 0.24795354 |
| ENSGALG00010028311 | CTXN1 | 420051 | 0.59656401 | 1.67035068 | 0.01262916 | 0.24875067 |
| ENSGALG00010002193 | NA | NA | -0.620067542 | 0.93107124 | 0.01263155 | 0.24875067 |
| ENSGALG00010018931 | NA | NA | -0.211592962 | 5.30891407 | 0.01264827 | 0.24876015 |
| ENSGALG00010024073 | RABGGTB | 424723 | -0.206771324 | 7.08260588 | 0.01269541 | 0.24913707 |
| ENSGALG00010028040 | TDRKH | 425667 | 0.350658234 | 4.52404276 | 0.01271934 | 0.24913707 |
| ENSGALG00010022264 | SCN8A | 426869 | 0.541581262 | 1.51484293 | 0.01272435 | 0.24913707 |
| ENSGALG00010003887 | GIPC1 | 112530986 | 0.236349155 | 6.09642551 | 0.01273248 | 0.24913707 |
| ENSGALG00010023955 | NA | NA | 0.366080961 | 3.13114029 | 0.01276827 | 0.24939152 |
| ENSGALG00010021860 | SLA2 | 100858750 | 0.491931894 | 2.29284208 | 0.01277804 | 0.24939152 |
| ENSGALG00010004000 | NA | NA | 0.19402356 | 6.01012139 | 0.01282737 | 0.25003581 |
| ENSGALG00010028497 | ADMP | 395622 | 0.243109524 | 6.86202825 | 0.01285895 | 0.25013839 |
| ENSGALG00010027165 | LOC100857637 | 100857637 | 0.249340492 | 5.25351241 | 0.01286529 | 0.25013839 |
| ENSGALG00010025832 | ENKD1 | 415643 | 0.221877518 | 5.00362682 | 0.01288202 | 0.25014623 |
| ENSGALG00010008424 | PPIP5K2 | 427276 | -0.187046856 | 7.17502599 | 0.01292417 | 0.25064711 |
| ENSGALG00010028738 | MIDN | 107055352 | 0.22778674 | 5.05930666 | 0.0129657 | 0.25095775 |
| ENSGALG00010000028 | ND4L | 63549492 | -0.309017084 | 9.11488681 | 0.01298836 | 0.25095775 |
| ENSGALG00010007867 | ZNF516 | 421009 | 0.177777602 | 6.5851999 | 0.01298933 | 0.25095775 |
| ENSGALG00010014750 | LONRF1 | 422742 | -0.238086495 | 4.73943355 | 0.01301652 | 0.25116636 |

| | | | | | | |
|---|---|---|---|---|---|---|
| ENSGALG00010008235 | MAEL | 418446 | -0.169999378 | 8.01973376 | 0.01305443 | 0.25139323 |
| ENSGALG00010005895 | MLLT10 | 420509 | -0.174141161 | 6.36750079 | 0.01307717 | 0.25139323 |
| ENSGALG00010012229 | FSBP | 770634 | -0.227821997 | 4.59727745 | 0.0130775 | 0.25139323 |
| ENSGALG00010007830 | SMIM19 | 422492 | 0.282003256 | 4.0743986 | 0.01310628 | 0.25163075 |
| ENSGALG00010010232 | LOC107054313 | 107054313 | -0.227907151 | 5.26273936 | 0.01313706 | 0.25180748 |
| ENSGALG00010027986 | CGN | 425665 | 0.364551186 | 3.86727173 | 0.01314836 | 0.25180748 |
| ENSGALG00010000412 | RFC5 | 100859719 | 0.205364108 | 5.96827034 | 0.01320607 | 0.25259696 |
| ENSGALG00010005326 | MASTL | 420487 | -0.201436602 | 7.10169879 | 0.01324046 | 0.25293899 |
| ENSGALG00010000363 | NA | NA | 0.773249168 | 0.83868567 | 0.01325775 | 0.25295392 |
| ENSGALG00010023736 | LOC107050879 | 107050879 | 0.342870856 | 3.80536745 | 0.01331141 | 0.2536619 |
| ENSGALG00010027092 | FAM72A | 419841 | 0.192265251 | 5.8652639 | 0.01333126 | 0.25372451 |
| ENSGALG00010024808 | CELSR2 | 430126 | 0.204840644 | 7.34303709 | 0.01334839 | 0.25373537 |
| ENSGALG00010017119 | BTBD6 | 101752037 | 0.202774124 | 6.03087783 | 0.01343228 | 0.25481376 |
| ENSGALG00010019990 | 8-Mar | 423770 | -0.355447306 | 3.34086292 | 0.01343839 | 0.25481376 |
| ENSGALG00010021828 | NA | NA | -0.914664448 | -0.0686692 | 0.01345761 | 0.25486287 |
| ENSGALG00010024829 | ERBB3 | 693245 | 0.208078399 | 5.38699857 | 0.01361707 | 0.25756431 |
| ENSGALG00010018749 | NA | NA | -0.178591964 | 6.10352055 | 0.01364427 | 0.25774839 |
| ENSGALG00010010287 | STAM2 | 424317 | -0.180153265 | 7.48120725 | 0.01366045 | 0.25774839 |
| ENSGALG00010020316 | PDPK1 | 416588 | -0.183662663 | 5.92212202 | 0.0139001 | 0.26193282 |
| ENSGALG00010014524 | NA | NA | -0.882315488 | -0.0843375 | 0.01391641 | 0.26193282 |
| ENSGALG00010003115 | UBE3D | 421843 | -0.284789443 | 4.13335669 | 0.01396177 | 0.26246408 |
| ENSGALG00010024698 | LOC107055405 | 107055405 | 0.743710344 | 0.22240696 | 0.01404921 | 0.26369656 |
| ENSGALG00010014134 | RPAP3 | 417811 | -0.184351516 | 6.58172903 | 0.01406175 | 0.26369656 |
| ENSGALG00010000003 | NA | NA | -0.351772289 | 10.5604679 | 0.01408946 | 0.26389316 |
| ENSGALG00010023909 | FOXRED1 | 419712 | 0.282835436 | 4.14035865 | 0.01411935 | 0.2641302 |
| ENSGALG00010020298 | PBDC1 | 100858203 | 0.199610932 | 6.47736945 | 0.01415724 | 0.26446606 |
| ENSGALG00010017717 | LLGL1 | 100858027 | 0.20758165 | 5.58979341 | 0.01417183 | 0.26446606 |
| ENSGALG00010000388 | HNRNPC | 121108713 | 0.208574771 | 5.7834617 | 0.01421802 | 0.26456926 |
| ENSGALG00010005288 | CNKSR3 | 421646 | -0.179819865 | 7.53405815 | 0.01421949 | 0.26456926 |
| ENSGALG00010007789 | WWOX | 415801 | 0.235136309 | 4.41558968 | 0.01423349 | 0.26456926 |
| ENSGALG00010027608 | GNAI2 | 396367 | 0.19312125 | 6.77884348 | 0.01427252 | 0.26456926 |
| ENSGALG00010021664 | TYSND1 | 769373 | 0.296448757 | 4.23843794 | 0.01427675 | 0.26456926 |
| ENSGALG00010026293 | NA | NA | -0.231038195 | 4.75447139 | 0.01428097 | 0.26456926 |
| ENSGALG00010010445 | LMAN1L | 426849 | -0.170965437 | 7.87721332 | 0.01431771 | 0.26492943 |
| ENSGALG00010028512 | NDUFA13 | 100859320 | 0.244194645 | 5.24596425 | 0.01451037 | 0.26817051 |
| ENSGALG00010010705 | NA | NA | 0.493621528 | 1.55827189 | 0.01459418 | 0.26913193 |
| ENSGALG00010011776 | IL17RA | 418158 | 0.197245468 | 5.90275327 | 0.01459752 | 0.26913193 |
| ENSGALG00010007102 | CMTM3 | 414829 | 0.247955721 | 4.17481208 | 0.01466843 | 0.26987013 |
| ENSGALG00010017338 | MBNL3 | 422233 | -0.171762895 | 7.25495241 | 0.01467279 | 0.26987013 |
| ENSGALG00010027227 | ZNF467 | 107052491 | 0.233264082 | 4.80542523 | 0.01476886 | 0.27127568 |
| ENSGALG00010012638 | NA | NA | 0.216339715 | 5.0617445 | 0.01478462 | 0.27127568 |
| ENSGALG00010001169 | SLC25A51 | 100859749 | -0.17627045 | 6.55077817 | 0.01483062 | 0.27137151 |
| ENSGALG00010004367 | TSPAN13 | 420595 | -0.190968306 | 7.51695743 | 0.01484166 | 0.27137151 |
| ENSGALG00010008999 | TRAPPC9 | 420312 | 0.27220327 | 3.8998936 | 0.01485817 | 0.27137151 |
| ENSGALG00010004357 | NA | NA | -0.335157515 | 4.80640349 | 0.01486069 | 0.27137151 |
| ENSGALG00010008708 | ZC3H4 | 101748292 | 0.238694796 | 5.82601541 | 0.01489489 | 0.27150992 |
| ENSGALG00010019801 | FIBIN | 426933 | 0.254061219 | 6.68107631 | 0.01491512 | 0.27150992 |
| ENSGALG00010027403 | ACKR2 | 420393 | 0.90156657 | 0.0299495 | 0.01492144 | 0.27150992 |
| ENSGALG00010000402 | PFKFB1 | 415906 | 0.558474438 | 1.77594035 | 0.01502091 | 0.27289021 |
| ENSGALG00010000771 | NA | NA | 0.207982634 | 5.44758934 | 0.01505212 | 0.27289021 |
| ENSGALG00010005836 | NEBL | 395148 | -0.289326309 | 3.99057381 | 0.0150564 | 0.27289021 |
| ENSGALG00010022767 | LOC107050029 | 107050029 | 0.744246464 | 0.2220685 | 0.01507177 | 0.27289021 |
| ENSGALG00010003383 | UBE2J1 | 395528 | -0.197799216 | 5.46842861 | 0.01508635 | 0.27289021 |

| | | | | | | |
|---|---|---|---|---|---|---|
| ENSGALG00010020938 | SMG1 | 416605 | -0.177265385 | 7.47605071 | 0.01512743 | 0.27331046 |
| ENSGALG00010019013 | LRRK1 | 415520 | -0.219862868 | 5.0126913 | 0.01518888 | 0.27392874 |
| ENSGALG00010024866 | TRAPPC4 | 419788 | 0.231810345 | 4.7712 | 0.0151974 | 0.27392874 |
| ENSGALG00010001969 | PLEKHA8 | 420637 | -0.298940392 | 4.89771224 | 0.01523173 | 0.27422488 |
| ENSGALG00010026013 | RABIF | 421170 | 0.231677567 | 4.93186088 | 0.01528453 | 0.27465359 |
| ENSGALG00010000471 | WRAP53 | 121108908 | 0.212439734 | 5.10324759 | 0.0152914 | 0.27465359 |
| ENSGALG00010015804 | CKB | 396248 | 0.192539262 | 7.2984364 | 0.01542603 | 0.27650309 |
| ENSGALG00010006341 | HHIP | 422460 | -0.406933596 | 3.04588599 | 0.01544596 | 0.27650309 |
| ENSGALG00010000205 | RNF215 | 417009 | 0.259057642 | 3.95507662 | 0.01544851 | 0.27650309 |
| ENSGALG00010029928 | USP43 | 101751148 | 0.267960686 | 5.0496275 | 0.01553025 | 0.27718448 |
| ENSGALG00010015756 | PCM1 | 395204 | -0.168373233 | 8.43760985 | 0.01554505 | 0.27718448 |
| ENSGALG00010008219 | NIP7 | 415863 | 0.221010624 | 5.76193681 | 0.01555048 | 0.27718448 |
| ENSGALG00010017033 | TMEM159 | 427005 | -0.17586418 | 6.40381543 | 0.01555895 | 0.27718448 |
| ENSGALG00010026453 | NA | NA | 0.894780651 | -0.1935212 | 0.01560622 | 0.27768208 |
| ENSGALG00010015056 | NA | NA | 0.686637021 | 0.5043809 | 0.01562313 | 0.27768208 |
| ENSGALG00010023231 | ASIC1 | 426883 | 0.371809911 | 3.07671362 | 0.01568454 | 0.27830695 |
| ENSGALG00010028723 | MIR1647 | 100315761 | 0.411844154 | 2.86018813 | 0.01569462 | 0.27830695 |
| ENSGALG00010015912 | NA | NA | -1.070529675 | -0.4757844 | 0.01572439 | 0.27851254 |
| ENSGALG00010027167 | ASIC1L | 107055878 | 0.356599607 | 2.80254282 | 0.01583 | 0.28005934 |
| ENSGALG00010027291 | MST1 | 396135 | 0.298510613 | 4.31649437 | 0.01590108 | 0.28077675 |
| ENSGALG00010000468 | TESC | 771113 | 0.503751736 | 2.08541906 | 0.01595 | 0.28077675 |
| ENSGALG00010019287 | AHSA2 | 421194 | -0.271094306 | 7.49172056 | 0.01596743 | 0.28077675 |
| ENSGALG00010024883 | ING5 | 424852 | 0.171824875 | 6.85491993 | 0.01597499 | 0.28077675 |
| ENSGALG00010009231 | NA | NA | -1.019162292 | -0.1912542 | 0.01598497 | 0.28077675 |
| ENSGALG00010024327 | PPP1R1B | 426639 | 0.502377522 | 1.75101853 | 0.01602035 | 0.28077675 |
| ENSGALG00010028125 | HMG20B | 426515 | 0.242041522 | 5.06282402 | 0.01602047 | 0.28077675 |
| ENSGALG00010006323 | ANAPC10 | 422461 | -0.211017891 | 5.07281603 | 0.0160218 | 0.28077675 |
| ENSGALG00010021766 | AGRN | 396538 | 0.224828781 | 6.8982274 | 0.01603549 | 0.28077675 |
| ENSGALG00010001935 | ABRAXAS2 | 423954 | -0.229647942 | 4.621376 | 0.01610324 | 0.28164133 |
| ENSGALG00010003471 | ADI1 | 421918 | -0.179455463 | 6.11558149 | 0.01615846 | 0.28228482 |
| ENSGALG00010019183 | ANGEL2 | 421363 | -0.17302263 | 5.76610022 | 0.0162399 | 0.28289349 |
| ENSGALG00010006653 | CFL2 | 423320 | -0.202939408 | 7.47553242 | 0.0162465 | 0.28289349 |
| ENSGALG00010003830 | DMB2 | 417051 | 0.328720599 | 3.29906523 | 0.01624991 | 0.28289349 |
| ENSGALG00010000585 | SPTBN2 | 101750255 | 0.20825425 | 7.25568076 | 0.01626716 | 0.28289349 |
| ENSGALG00010013253 | SHB | 427280 | -0.209186705 | 5.41286912 | 0.01630585 | 0.28295123 |
| ENSGALG00010018241 | RALGAPA2 | 421240 | -0.181687782 | 6.05865567 | 0.01631876 | 0.28295123 |
| ENSGALG00010010845 | LOC121108177 | 121108177 | 0.21187176 | 4.94295863 | 0.01632589 | 0.28295123 |
| ENSGALG00010017911 | NA | NA | -0.822283192 | 0.48565219 | 0.01639427 | 0.28361238 |
| ENSGALG00010018696 | NA | NA | -0.229130835 | 4.52102505 | 0.01640106 | 0.28361238 |
| ENSGALG00010027609 | RPL36 | 373936 | 0.256987231 | 7.43267835 | 0.01642708 | 0.2837421 |
| ENSGALG00010013416 | NDUFB2 | 418118 | 0.233106545 | 4.83967021 | 0.0164529 | 0.28379963 |
| ENSGALG00010028059 | GZMM | 100859124 | 0.779001085 | 0.3383657 | 0.01650671 | 0.28379963 |
| ENSGALG00010016754 | NA | NA | -0.300247157 | 3.48939213 | 0.01655915 | 0.28379963 |
| ENSGALG00010024101 | CNEP1R1 | 415736 | -0.228127014 | 4.69144786 | 0.01656037 | 0.28379963 |
| ENSGALG00010015449 | HDX | 422272 | -0.267067751 | 3.8120003 | 0.01656757 | 0.28379963 |
| ENSGALG00010029750 | TEN1 | 100529060 | 0.256586915 | 4.71344205 | 0.01657404 | 0.28379963 |
| ENSGALG00010018340 | DNPEP | 424200 | 0.213055611 | 5.84270795 | 0.0165776 | 0.28379963 |
| ENSGALG00010009351 | GTF3C2 | 430980 | 0.218579673 | 5.53363893 | 0.0165831 | 0.28379963 |
| ENSGALG00010018231 | SMYD5 | 422951 | 0.177845658 | 6.33962131 | 0.01659712 | 0.28379963 |
| ENSGALG00010025232 | PRIM1 | 426646 | 0.233627133 | 4.75864111 | 0.01661826 | 0.28381286 |
| ENSGALG00010011795 | NA | NA | -0.358257696 | 2.95713341 | 0.01665165 | 0.28381286 |
| ENSGALG00010007905 | MRPL32 | 420773 | 0.249165466 | 4.47829554 | 0.016664 | 0.28381286 |
| ENSGALG00010009166 | CLGN | 422447 | -0.227655962 | 6.39120297 | 0.01667199 | 0.28381286 |

| | | | | | | |
|---|---|---|---|---|---|---|
| ENSGALG00010009847 | TMEM14C | 420851 | 0.213753414 | 4.71528274 | 0.01676773 | 0.28509067 |
| ENSGALG00010022244 | ENTPD7 | 430337 | 0.31040538 | 3.2371444 | 0.01678427 | 0.28509067 |
| ENSGALG00010020265 | SPTBN5 | 423225 | -0.931549017 | -0.1448471 | 0.0169651 | 0.28766585 |
| ENSGALG00010022061 | NA | NA | -1.15582051 | -0.4191416 | 0.01697343 | 0.28766585 |
| ENSGALG00010023182 | EPHB6 | 418308 | 0.244549454 | 5.01322822 | 0.01699367 | 0.28769065 |
| ENSGALG00010010216 | ATXN7L1 | 427847 | -0.21161037 | 5.02341873 | 0.0170251 | 0.28790465 |
| ENSGALG00010009050 | PIK3CG | 417706 | -0.602539494 | 1.83473732 | 0.01705779 | 0.28813931 |
| ENSGALG00010023841 | OSBPL7 | 100857443 | 0.381087031 | 3.3539313 | 0.01707921 | 0.28818348 |
| ENSGALG00010015574 | NA | NA | -0.400160579 | 2.80852598 | 0.01715605 | 0.2886584 |
| ENSGALG00010024857 | SLC37A4 | 419789 | 0.189640758 | 8.27164545 | 0.01715918 | 0.2886584 |
| ENSGALG00010029720 | TIMM22 | 417594 | 0.18494579 | 5.84924136 | 0.01716388 | 0.2886584 |
| ENSGALG00010024334 | USP28 | 428246 | 0.18226281 | 6.93554889 | 0.01723673 | 0.28956575 |
| ENSGALG00010001192 | MLF1 | 425019 | -0.215067748 | 5.88812601 | 0.01726937 | 0.28979632 |
| ENSGALG00010016447 | ID3 | 395281 | 0.326307862 | 3.67586791 | 0.01733919 | 0.29059142 |
| ENSGALG00010003107 | GNG11 | 771999 | 0.307864526 | 3.26283278 | 0.01735469 | 0.29059142 |
| ENSGALG00010023397 | LOC107049267 | 107049267 | 0.543276996 | 1.39290646 | 0.01742796 | 0.29149973 |
| ENSGALG00010023696 | LRRC4C | 428862 | -0.212544839 | 5.99787286 | 0.01747067 | 0.29154553 |
| ENSGALG00010021473 | SLC8A3 | 395759 | -0.215431149 | 4.99607108 | 0.01748613 | 0.29154553 |
| ENSGALG00010026344 | TIMM10 | 425202 | 0.243981764 | 4.37919512 | 0.01750084 | 0.29154553 |
| ENSGALG00010017414 | LOC107056278 | 107056278 | 0.364809345 | 3.15158265 | 0.01750681 | 0.29154553 |
| ENSGALG00010020328 | SLC17A6 | 422971 | -0.526609588 | 1.54842744 | 0.0175276 | 0.29157482 |
| ENSGALG00010022593 | MANBAL | 428138 | 0.200446061 | 5.24100937 | 0.01759634 | 0.2924007 |
| ENSGALG00010012956 | FABP3 | 419557 | 0.285206586 | 3.93949501 | 0.01765874 | 0.29311978 |
| ENSGALG00010027540 | PPOX | 107051238 | 0.278515057 | 4.28844948 | 0.01773002 | 0.29398452 |
| ENSGALG00010009707 | NA | NA | -0.629971786 | 0.79952388 | 0.01777739 | 0.29445125 |
| ENSGALG00010017430 | CACNA1H | 416526 | 0.214378135 | 5.52563458 | 0.01790021 | 0.29513684 |
| ENSGALG00010024772 | ELAPOR1 | 429496 | 0.252497645 | 4.30544418 | 0.01791457 | 0.29513684 |
| ENSGALG00010013515 | RBP4 | 396454 | 0.609729421 | 0.80761712 | 0.01791892 | 0.29513684 |
| ENSGALG00010008335 | HNRNPUL1 | 776596 | 0.206381325 | 6.87775355 | 0.01793058 | 0.29513684 |
| ENSGALG00010009157 | ABHD1 | 107053155 | 0.297731781 | 4.37743092 | 0.0179522 | 0.29513684 |
| ENSGALG00010028454 | C8G | 770691 | 0.404523125 | 3.20692566 | 0.01797261 | 0.29513684 |
| ENSGALG00010010317 | LOC121110417 | 121110417 | 0.192644961 | 5.70566808 | 0.01799473 | 0.29513684 |
| ENSGALG00010026023 | THY1 | 378897 | 0.398551434 | 3.38607904 | 0.01800152 | 0.29513684 |
| ENSGALG00010009024 | PMAIP1 | 770077 | -0.550039874 | 1.34958453 | 0.01801096 | 0.29513684 |
| ENSGALG00010009326 | DLD | 417699 | -0.167708803 | 5.99664933 | 0.01801142 | 0.29513684 |
| ENSGALG00010013322 | NA | NA | 0.351289574 | 3.8272364 | 0.01805432 | 0.29539061 |
| ENSGALG00010019390 | TVP23A | 771988 | -0.518550349 | 1.34435184 | 0.01806547 | 0.29539061 |
| ENSGALG00010014785 | NA | NA | -0.746814707 | 1.93423092 | 0.0181364 | 0.2956507 |
| ENSGALG00010008408 | SMARCA4 | 395932 | 0.170364917 | 8.91538636 | 0.01816464 | 0.2956507 |
| ENSGALG00010029502 | BTBD17 | 427813 | 0.253461358 | 4.47847502 | 0.0181856 | 0.2956507 |
| ENSGALG00010017250 | COL16A1 | 430477 | 0.485194672 | 3.25602582 | 0.01819398 | 0.2956507 |
| ENSGALG00010024657 | DPY19L3 | 768782 | 0.228824045 | 5.6753889 | 0.01821098 | 0.2956507 |
| ENSGALG00010027985 | PSMB4 | 429986 | 0.197877939 | 7.55300147 | 0.01821337 | 0.2956507 |
| ENSGALG00010016832 | DYNC1H1 | 423461 | -0.1732029 | 8.64011474 | 0.01822484 | 0.2956507 |
| ENSGALG00010027520 | FANCE | 419891 | 0.31757798 | 3.50440028 | 0.01823701 | 0.2956507 |
| ENSGALG00010014169 | NA | NA | -0.764503463 | 0.8614349 | 0.01825652 | 0.2956507 |
| ENSGALG00010012824 | HIBCH | 423979 | 0.196922778 | 5.67213549 | 0.01827434 | 0.2956507 |
| ENSGALG00010027346 | SIN3B | 420140 | 0.184696607 | 5.61244419 | 0.01829724 | 0.29570892 |
| ENSGALG00010018848 | MEF2A | 395670 | -0.220958596 | 4.52007844 | 0.01835004 | 0.29624964 |
| ENSGALG00010010781 | NAPA | 100858304 | 0.193820643 | 6.11386799 | 0.01837077 | 0.29627222 |
| ENSGALG00010008200 | GCOG8 | 769684 | 0.196083593 | 5.06973323 | 0.01841286 | 0.29639027 |
| ENSGALG00010027567 | PARP3 | 771158 | 0.454497162 | 1.83410514 | 0.01841678 | 0.29639027 |
| ENSGALG00010027917 | PPARG | 373928 | -0.36920829 | 2.98984382 | 0.01848314 | 0.29696837 |

| | | | | | | |
|---|---|---|---|---|---|---|
| ENSGALG00010021947 | CDK2AP2 | 107053360 | 0.296138658 | 3.52977903 | 0.01850384 | 0.29696837 |
| ENSGALG00010013131 | SPATA2 | 419355 | -0.16560078 | 6.13889621 | 0.01851085 | 0.29696837 |
| ENSGALG00010008914 | NA | NA | -0.193297156 | 5.27352199 | 0.01857093 | 0.29728019 |
| ENSGALG00010024878 | RHPN2 | 415771 | -0.220889268 | 5.67145585 | 0.01858642 | 0.29728019 |
| ENSGALG00010028368 | TIMM13 | 100859352 | 0.19972486 | 5.18288625 | 0.0185885 | 0.29728019 |
| ENSGALG00010009089 | AGBL5 | 770678 | 0.21600378 | 5.28428568 | 0.01869446 | 0.29838705 |
| ENSGALG00010000353 | NDUFB11 | 121108932 | 0.225381374 | 5.52740712 | 0.01869666 | 0.29838705 |
| ENSGALG00010026329 | SMTNL1 | 101750657 | 0.271744678 | 3.97677256 | 0.0187311 | 0.29862558 |
| ENSGALG00010012136 | NA | NA | -0.945895084 | -0.2257055 | 0.01876683 | 0.29888414 |
| ENSGALG00010024579 | ZNF740 | 100859343 | 0.184576708 | 5.79652657 | 0.01880381 | 0.29916215 |
| ENSGALG00010005427 | NA | NA | 0.256809627 | 4.60346028 | 0.01888287 | 0.29943051 |
| ENSGALG00010014984 | LPP | 429148 | -0.214544387 | 5.25139845 | 0.01888719 | 0.29943051 |
| ENSGALG00010010685 | STIM2 | 422799 | -0.191420762 | 5.82868255 | 0.01888875 | 0.29943051 |
| ENSGALG00010029192 | DTX2 | 417515 | 0.241125623 | 4.44656768 | 0.01891517 | 0.29943051 |
| ENSGALG00010024695 | TRABD2B | 424623 | -0.180685523 | 5.67504611 | 0.0189184 | 0.29943051 |
| ENSGALG00010029146 | NEK6 | 417101 | 0.194013225 | 5.05078507 | 0.01897779 | 0.29981842 |
| ENSGALG00010000140 | SEPHS3 | 113219448 | 0.268390485 | 6.17868198 | 0.01898204 | 0.29981842 |
| ENSGALG00010003330 | HEY1 | 428365 | 0.252718341 | 4.25499966 | 0.01902083 | 0.29986373 |
| ENSGALG00010000483 | LOC107051159 | 107051159 | 0.392291749 | 3.30743878 | 0.01902406 | 0.29986373 |
| ENSGALG00010014806 | MAP7D1 | 419624 | 0.230925094 | 5.61313344 | 0.01915601 | 0.30104336 |
| ENSGALG00010029761 | MRPL38 | 770172 | 0.199263738 | 5.347883 | 0.01915706 | 0.30104336 |
| ENSGALG00010012196 | CENPK | 427162 | -0.194601791 | 5.1868996 | 0.01915784 | 0.30104336 |
| ENSGALG00010014250 | LOC101750517 | 101750517 | -0.69779261 | 0.49565918 | 0.01919864 | 0.30108279 |
| ENSGALG00010006131 | LOC101749299 | 101749299 | 0.289373672 | 3.76639347 | 0.01920917 | 0.30108279 |
| ENSGALG00010025917 | SCUBE3 | 419896 | 0.412667687 | 2.28662405 | 0.01921931 | 0.30108279 |
| ENSGALG00010019399 | CD248 | 421234 | 0.476483214 | 1.46506706 | 0.01927345 | 0.30136512 |
| ENSGALG00010000279 | NA | NA | -0.293546735 | 3.82076326 | 0.01929246 | 0.30136512 |
| ENSGALG00010008183 | ALG10 | 426734 | -0.214150872 | 5.25895685 | 0.01931386 | 0.30136512 |
| ENSGALG00010003617 | COG3 | 418846 | 0.185487335 | 5.90439326 | 0.01931601 | 0.30136512 |
| ENSGALG00010003372 | ENOX1 | 418839 | -0.418160576 | 2.16212972 | 0.01938698 | 0.30216471 |
| ENSGALG00010015975 | BTK | 374075 | 0.338384983 | 2.8553586 | 0.01943256 | 0.30227493 |
| ENSGALG00010028197 | INTS3 | 771890 | 0.180840009 | 7.54586205 | 0.01943351 | 0.30227493 |
| ENSGALG00010001818 | ADGRB3 | 421875 | 0.485182889 | 1.65495203 | 0.01948868 | 0.30264842 |
| ENSGALG00010020889 | PPRC1 | 423848 | 0.2002759 | 6.66755734 | 0.01949703 | 0.30264842 |
| ENSGALG00010014060 | NA | NA | -0.491779334 | 1.53910795 | 0.01953572 | 0.30294211 |
| ENSGALG00010020206 | MRPS11 | 415498 | 0.271490614 | 3.95050699 | 0.01959449 | 0.30354619 |
| ENSGALG00010021387 | SLC2A14 | 396517 | 0.174999029 | 10.1323199 | 0.01962553 | 0.30362767 |
| ENSGALG00010005364 | MAP7 | 421689 | -0.311798036 | 3.43412891 | 0.01963939 | 0.30362767 |
| ENSGALG00010022032 | KBTBD4 | 423180 | 0.242673607 | 4.30315137 | 0.01971161 | 0.30418792 |
| ENSGALG00010018510 | MRPL43 | 100858689 | 0.279272317 | 4.34797765 | 0.01971533 | 0.30418792 |
| ENSGALG00010002684 | NA | NA | -0.571112197 | 1.02910765 | 0.01975318 | 0.30446522 |
| ENSGALG00010003860 | TMEM70 | 420188 | -0.190445311 | 7.41465642 | 0.01979271 | 0.30476789 |
| ENSGALG00010024821 | NECAP1 | 770083 | -0.190583258 | 5.5487902 | 0.01984518 | 0.30485339 |
| ENSGALG00010014482 | THOC3 | 416232 | 0.189727753 | 5.44589876 | 0.01984598 | 0.30485339 |
| ENSGALG00010027467 | GLYCTK | 427577 | 0.272821041 | 4.23032045 | 0.01985795 | 0.30485339 |
| ENSGALG00010026532 | NA | NA | 0.276805018 | 4.6495109 | 0.01990589 | 0.30510061 |
| ENSGALG00010008106 | NA | NA | 0.434518172 | 2.06219383 | 0.01991388 | 0.30510061 |
| ENSGALG00010021237 | FSCN1 | 416485 | 0.183367575 | 8.01301207 | 0.01995895 | 0.30517985 |
| ENSGALG00010024576 | CSAD | 426184 | 0.260570815 | 4.03039928 | 0.01995913 | 0.30517985 |
| ENSGALG00010020760 | TXNDC11 | 416428 | -0.295663075 | 3.32763496 | 0.01997881 | 0.30517985 |
| ENSGALG00010020497 | OSBPL11 | 424262 | -0.173791532 | 5.80267303 | 0.02001905 | 0.30534143 |
| ENSGALG00010017254 | NA | NA | -0.196940932 | 5.371294 | 0.0200504 | 0.30534143 |
| ENSGALG00010003131 | TAP1 | 427727 | 0.294006936 | 4.297735 | 0.02006278 | 0.30534143 |

| | | | | | | |
|---|---|---|---|---|---|---|
| ENSGALG00010003652 | MYCN | 421948 | -0.158018971 | 8.78033289 | 0.02006911 | 0.30534143 |
| ENSGALG00010006107 | PGF | 771832 | -0.408094793 | 2.33398018 | 0.02010516 | 0.30558642 |
| ENSGALG00010028856 | ETV3 | 426718 | 0.202805992 | 6.67392304 | 0.02017283 | 0.30631106 |
| ENSGALG00010017642 | 1-Mar | 428563 | 0.166580863 | 7.18050219 | 0.02030236 | 0.30672049 |
| ENSGALG00010022734 | ATP5G2 | 100858216 | 0.183649808 | 6.09468285 | 0.02031467 | 0.30672049 |
| ENSGALG00010024751 | MTMR10 | 415382 | -0.174576517 | 5.50206283 | 0.02032326 | 0.30672049 |
| ENSGALG00010029655 | PHKG1 | 417543 | 0.194370052 | 5.49095596 | 0.02032791 | 0.30672049 |
| ENSGALG00010027380 | GPX1 | 100857115 | 0.223480779 | 6.63614002 | 0.0203463 | 0.30672049 |
| ENSGALG00010020648 | TET3 | 425829 | 0.178084459 | 6.08545838 | 0.02036464 | 0.30672049 |
| ENSGALG00010004360 | NA | NA | -0.494161696 | 2.38881295 | 0.02038813 | 0.30672049 |
| ENSGALG00010008561 | EEF1D | 107049004 | 0.162191585 | 8.75768614 | 0.0203947 | 0.30672049 |
| ENSGALG00010007780 | SIRT5 | 420834 | 0.236673178 | 4.7201703 | 0.02040982 | 0.30672049 |
| ENSGALG00010002431 | BCKDHB | 395375 | 0.269694679 | 3.76173179 | 0.02041722 | 0.30672049 |
| ENSGALG00010019149 | COX5B | 107049545 | 0.237139241 | 5.99678741 | 0.02042001 | 0.30672049 |
| ENSGALG00010018520 | LOC421255 | 421255 | 0.262470611 | 4.36534855 | 0.02050199 | 0.30762445 |
| ENSGALG00010000817 | LOC121108888 | 121108888 | 0.194366062 | 7.45496993 | 0.02052034 | 0.30762445 |
| ENSGALG00010020778 | SLIT1 | 395293 | 0.30138547 | 4.92136609 | 0.02055159 | 0.30775294 |
| ENSGALG00010007103 | NA | NA | -0.520748459 | 1.83040006 | 0.02058509 | 0.30775294 |
| ENSGALG00010029926 | KIAA0100 | 417573 | 0.185961404 | 6.31636763 | 0.02058918 | 0.30775294 |
| ENSGALG00010011949 | ABCA12 | 424011 | -0.33566579 | 2.90780202 | 0.02062982 | 0.30804341 |
| ENSGALG00010026262 | NAV1 | 430162 | 0.217885987 | 5.95354872 | 0.02068177 | 0.30804341 |
| ENSGALG00010021519 | FNTB | 100857683 | 0.199158739 | 5.76357359 | 0.02068656 | 0.30804341 |
| ENSGALG00010026422 | P2RX3 | 428856 | 0.472052347 | 2.3769488 | 0.02068903 | 0.30804341 |
| ENSGALG00010024684 | CADNL | 427547 | -0.237744264 | 5.11872409 | 0.02071876 | 0.30818651 |
| ENSGALG00010002550 | NA | NA | -0.75172797 | 0.2094693 | 0.02077699 | 0.30875296 |
| ENSGALG00010013135 | TTC3 | 418518 | -0.165684317 | 6.64450923 | 0.02080504 | 0.30887013 |
| ENSGALG00010029073 | NA | NA | -0.756862359 | 0.11077256 | 0.02088882 | 0.30976199 |
| ENSGALG00010016763 | UBALD1 | 416661 | 0.256717425 | 4.95343867 | 0.02090555 | 0.30976199 |
| ENSGALG00010000432 | LOC100859273 | 100859273 | 0.208460242 | 7.08727681 | 0.02094147 | 0.30999446 |
| ENSGALG00010024709 | TEAD4 | 395542 | 0.259851416 | 4.39194909 | 0.02097775 | 0.31014972 |
| ENSGALG00010016461 | HPCAL4 | 419676 | 0.606574964 | 1.30603606 | 0.02099245 | 0.31014972 |
| ENSGALG00010010214 | FAF2 | 416225 | 0.165661672 | 5.79816685 | 0.02101822 | 0.3102313 |
| ENSGALG00010019882 | NA | NA | -0.740756665 | 1.07551124 | 0.02104431 | 0.31031742 |
| ENSGALG00010000007 | ND1 | 63549479 | -0.274575936 | 10.9039564 | 0.0211048 | 0.31091017 |
| ENSGALG00010015942 | AP3S2 | 415581 | 0.240302246 | 4.51881554 | 0.02116834 | 0.31097263 |
| ENSGALG00010010371 | SOCS1L | 417978 | 0.24902956 | 4.35999629 | 0.02122143 | 0.31097263 |
| ENSGALG00010015951 | NA | NA | 0.578227896 | 1.08320641 | 0.02131305 | 0.31097263 |
| ENSGALG00010005516 | LARP4B | 420457 | -0.184026978 | 8.21415388 | 0.02131739 | 0.31097263 |
| ENSGALG00010013517 | STMN1 | 396057 | 0.159000052 | 8.88948363 | 0.02131754 | 0.31097263 |
| ENSGALG00010005424 | NA | NA | 0.322375375 | 3.49688724 | 0.02132444 | 0.31097263 |
| ENSGALG00010022937 | BLOC1S1 | 100857285 | 0.243657157 | 4.69663209 | 0.02134845 | 0.31097263 |
| ENSGALG00010001222 | ZNF800 | 417747 | -0.262298796 | 3.81621608 | 0.02136003 | 0.31097263 |
| ENSGALG00010029475 | LOC107057183 | 107057183 | 0.821166588 | 1.98906622 | 0.02138687 | 0.31097263 |
| ENSGALG00010025263 | SAMD14 | 100857537 | 0.295029564 | 4.22609758 | 0.02140346 | 0.31097263 |
| ENSGALG00010024058 | SLC44A5 | 429117 | -0.286395344 | 3.85787757 | 0.02140744 | 0.31097263 |
| ENSGALG00010008375 | PLCG1L | 776273 | 0.201376382 | 5.84013081 | 0.02143486 | 0.31097263 |
| ENSGALG00010017380 | MARCKSL1 | 770764 | 0.181377938 | 7.60853507 | 0.02144205 | 0.31097263 |
| ENSGALG00010018247 | NOTO | 396302 | 0.240506408 | 7.78683069 | 0.02145214 | 0.31097263 |
| ENSGALG00010017422 | NA | NA | 0.71041122 | 0.32604937 | 0.02146049 | 0.31097263 |
| ENSGALG00010028086 | MATK | 769076 | 0.579122368 | 0.98104976 | 0.02146212 | 0.31097263 |
| ENSGALG00010023177 | LZTS2 | 428973 | 0.200796141 | 5.30479071 | 0.02146479 | 0.31097263 |
| ENSGALG00010013720 | SLC16A10 | 421753 | -0.173434149 | 6.10670019 | 0.0214861 | 0.31097263 |
| ENSGALG00010012760 | PLEKHF2 | 420234 | -0.21663462 | 4.89664555 | 0.02149953 | 0.31097263 |

| | | | | | | |
|---|---|---|---|---|---|---|
| ENSGALG00010011278 | TDH | 422034 | -0.348601883 | 4.09268467 | 0.02151498 | 0.31097263 |
| ENSGALG00010001982 | DMD | 396236 | -0.159705681 | 7.54906417 | 0.02153823 | 0.31101525 |
| ENSGALG00010020626 | MORN4 | 425290 | 0.3393084 | 3.25809148 | 0.02160842 | 0.31104539 |
| ENSGALG00010021617 | SORBS1 | 423749 | 0.246395983 | 5.08362814 | 0.02163298 | 0.31104539 |
| ENSGALG00010024799 | LOC415641 | 415641 | -0.176395741 | 8.05184703 | 0.02164225 | 0.31104539 |
| ENSGALG00010003928 | NA | NA | 0.290513443 | 4.7327203 | 0.02164934 | 0.31104539 |
| ENSGALG00010024194 | NKAIN4 | 419240 | 0.235943104 | 4.47148993 | 0.02166515 | 0.31104539 |
| ENSGALG00010029140 | NPDC1 | 770620 | 0.180744585 | 6.51091466 | 0.02167183 | 0.31104539 |
| ENSGALG00010026213 | AKR1A1 | 424599 | 0.199157407 | 5.23128852 | 0.02168243 | 0.31104539 |
| ENSGALG00010027692 | PHF2 | 415981 | -0.205863216 | 5.1286526 | 0.02172703 | 0.31139373 |
| ENSGALG00010026977 | ELK4 | 419832 | -0.301289602 | 3.31115684 | 0.02179283 | 0.31185716 |
| ENSGALG00010013329 | LIFR | 395262 | 0.185497371 | 5.49229419 | 0.02180008 | 0.31185716 |
| ENSGALG00010024116 | MCAM | 448832 | 0.240939895 | 5.06980037 | 0.02191524 | 0.31321214 |
| ENSGALG00010025558 | GALNT3 | 424177 | -0.270273101 | 4.18826661 | 0.02195264 | 0.31334373 |
| ENSGALG00010016168 | PLIN2 | 427237 | -0.191772806 | 6.20965377 | 0.02196792 | 0.31334373 |
| ENSGALG00010023063 | KCTD5 | 416589 | 0.172997108 | 6.74904004 | 0.02199259 | 0.31334373 |
| ENSGALG00010021327 | IGSF1 | 419114 | -0.584744082 | 0.81544478 | 0.02200626 | 0.31334373 |
| ENSGALG00010012904 | ERMP1 | 426644 | -0.170396946 | 8.10528092 | 0.02207879 | 0.3140846 |
| ENSGALG00010023447 | C12orf57 | 771099 | 0.251370648 | 4.38336223 | 0.0221475 | 0.31476983 |
| ENSGALG00010018545 | NA | NA | 0.35701669 | 3.16759746 | 0.02228772 | 0.31612607 |
| ENSGALG00010024951 | LOC124417104 | 124417104 | -0.477354817 | 1.99903767 | 0.02229231 | 0.31612607 |
| ENSGALG00010013768 | FAM110A | 100858650 | 0.438609794 | 2.64683919 | 0.02233128 | 0.31612607 |
| ENSGALG00010002855 | RACK1 | 417044 | 0.174669469 | 9.92245182 | 0.0223628 | 0.31612607 |
| ENSGALG00010026555 | NA | NA | -0.499885658 | 1.57797764 | 0.02236794 | 0.31612607 |
| ENSGALG00010018767 | NDUFB6 | 416391 | 0.223230886 | 5.09136395 | 0.02238343 | 0.31612607 |
| ENSGALG00010024610 | PFKL | 769850 | 0.183306431 | 6.58300114 | 0.02238736 | 0.31612607 |
| ENSGALG00010008224 | LOC100859629 | 100859629 | 0.202633565 | 7.27949241 | 0.02247652 | 0.31709278 |
| ENSGALG00010024431 | FEZ1 | 425587 | 0.327944778 | 4.32484608 | 0.02251116 | 0.31728926 |
| ENSGALG00010013561 | AGK | 418121 | -0.221602245 | 8.02767992 | 0.02265771 | 0.31875063 |
| ENSGALG00010028068 | POLE4 | 107055326 | 0.199910139 | 6.51787281 | 0.02266066 | 0.31875063 |
| ENSGALG00010001172 | TMEM200C | 421054 | -0.24188158 | 4.30468875 | 0.02267725 | 0.31875063 |
| ENSGALG00010024656 | NA | NA | 0.273371168 | 3.66957146 | 0.02273899 | 0.31932545 |
| ENSGALG00010017346 | GM2A | 100857973 | 0.198603916 | 5.07273639 | 0.02282143 | 0.3198074 |
| ENSGALG00010027081 | PDE4B | 424700 | 0.276988655 | 4.42631715 | 0.02282626 | 0.3198074 |
| ENSGALG00010015108 | FAAH2 | 100859568 | 0.188896122 | 5.43142772 | 0.02283593 | 0.3198074 |
| ENSGALG00010015364 | CLOCK | 373991 | -0.200744008 | 4.85987272 | 0.02289662 | 0.32009875 |
| ENSGALG00010000232 | TMEM147 | 112531206 | 0.303543744 | 4.37682496 | 0.02289852 | 0.32009875 |
| ENSGALG00010015205 | BPIFB3 | 419289 | 0.779799894 | 0.07008161 | 0.0229865 | 0.32091829 |
| ENSGALG00010003842 | WIZ | 426166 | 0.171079487 | 7.93747857 | 0.0230043 | 0.32091829 |
| ENSGALG00010001499 | TIPARP | 425026 | -0.20904227 | 4.67431765 | 0.02301999 | 0.32091829 |
| ENSGALG00010028478 | LOC107049501 | 107049501 | 0.340185066 | 3.12528581 | 0.02304881 | 0.32102799 |
| ENSGALG00010023354 | SEPTIN2 | 424843 | -0.18206468 | 7.43683501 | 0.02314713 | 0.32201789 |
| ENSGALG00010000126 | LOC121108714 | 121108714 | 0.515138989 | 2.01364322 | 0.02317167 | 0.32201789 |
| ENSGALG00010030046 | NA | NA | 0.450505713 | 1.7001175 | 0.02318293 | 0.32201789 |
| ENSGALG00010006288 | ARF5 | 396265 | 0.187255706 | 5.79640673 | 0.0232457 | 0.32259731 |
| ENSGALG00010021115 | ALPL | 396317 | 0.208392621 | 5.34135154 | 0.02328545 | 0.32285641 |
| ENSGALG00010027214 | PLIN4 | 100857433 | 0.18836463 | 6.9295282 | 0.02330684 | 0.32286086 |
| ENSGALG00010008587 | NA | NA | -0.552873614 | 1.54882268 | 0.02333231 | 0.32292174 |
| ENSGALG00010016760 | DLK1 | 423459 | -0.43319313 | 1.83554378 | 0.02339191 | 0.32345434 |
| ENSGALG00010010813 | LOC107049991 | 107049991 | 0.313238692 | 3.22828926 | 0.0234439 | 0.32388094 |
| ENSGALG00010027582 | LOC101748032 | 101748032 | 0.765031423 | 0.32731447 | 0.02349796 | 0.32433535 |
| ENSGALG00010003097 | TFPI2 | 420561 | -0.219672277 | 4.52024031 | 0.02358625 | 0.32526097 |
| ENSGALG00010021479 | SKAP1 | 101752206 | 0.686338694 | 1.22046436 | 0.02361618 | 0.32538077 |

| | | | | | | |
|---|---|---|---|---|---|---|
| ENSGALG00010020783 | NA | NA | -0.700448115 | 0.39106584 | 0.02368612 | 0.32565936 |
| ENSGALG00010021135 | CLK3 | 415295 | 0.221998503 | 4.62271914 | 0.02369935 | 0.32565936 |
| ENSGALG00010003019 | TAPBP | 417048 | 0.382389063 | 2.52213733 | 0.02370016 | 0.32565936 |
| ENSGALG00010008603 | NA | NA | -0.45534674 | 2.17416914 | 0.02379932 | 0.32672887 |
| ENSGALG00010029364 | NPB | 769277 | 0.281989118 | 3.88666691 | 0.02403503 | 0.32945646 |
| ENSGALG00010026149 | LASP1 | 420002 | 0.191461692 | 5.15339934 | 0.02404973 | 0.32945646 |
| ENSGALG00010005511 | AHCYL2 | 416675 | 0.159356964 | 6.42932576 | 0.02406251 | 0.32945646 |
| ENSGALG00010028445 | SYT11 | 429166 | 0.157226596 | 7.81691352 | 0.02410957 | 0.32980606 |
| ENSGALG00010027901 | MOB3A | 100857182 | 0.241981926 | 5.14358137 | 0.02413961 | 0.32992234 |
| ENSGALG00010007874 | NA | NA | 0.356746661 | 2.75136305 | 0.02416424 | 0.32996469 |
| ENSGALG00010013881 | NA | NA | -0.468427639 | 3.27300427 | 0.02430944 | 0.33165181 |
| ENSGALG00010024831 | SLC7A9 | 415768 | -0.238141844 | 5.23577044 | 0.02445203 | 0.33260823 |
| ENSGALG00010024933 | CS | 100858903 | 0.191934722 | 6.69361169 | 0.02445458 | 0.33260823 |
| ENSGALG00010021315 | COX6A1 | 416978 | 0.184433854 | 7.68611749 | 0.02446125 | 0.33260823 |
| ENSGALG00010028013 | CBARP | 100858130 | 0.377501602 | 2.96607986 | 0.02446778 | 0.33260823 |
| ENSGALG00010029791 | GALK1 | 417373 | 0.267909302 | 3.80283985 | 0.02450653 | 0.33260823 |
| ENSGALG00010029774 | RFLNB | 417623 | -0.48776269 | 1.98898585 | 0.0245098 | 0.33260823 |
| ENSGALG00010026076 | C1orf35 | 769525 | 0.26774669 | 4.09924796 | 0.02457147 | 0.33314994 |
| ENSGALG00010027974 | ABCA7 | 770610 | 0.401233413 | 2.85004561 | 0.02459719 | 0.3332038 |
| ENSGALG00010007203 | SLC30A6 | 421480 | -0.213020201 | 4.63439641 | 0.0246769 | 0.33398827 |
| ENSGALG00010021531 | RAB15 | 429955 | 0.247053148 | 4.39837996 | 0.02484369 | 0.33563602 |
| ENSGALG00010004053 | NA | NA | -0.495373851 | 1.74315776 | 0.02484655 | 0.33563602 |
| ENSGALG00010008602 | NA | NA | -0.348587216 | 5.19069699 | 0.02487271 | 0.33563602 |
| ENSGALG00010002716 | NHLRC3 | 418890 | 0.261598982 | 3.90649745 | 0.02490367 | 0.33563602 |
| ENSGALG00010004433 | AHR1A | 373907 | -0.293190271 | 3.22068386 | 0.02490818 | 0.33563602 |
| ENSGALG00010016319 | RBM41 | 428698 | 0.358835601 | 2.70192341 | 0.02494161 | 0.33579117 |
| ENSGALG00010028182 | SMG5 | 425473 | 0.182498048 | 6.24888399 | 0.02500868 | 0.33639856 |
| ENSGALG00010024806 | ANKRD27 | 415766 | -0.249034884 | 3.81978702 | 0.02516022 | 0.33814015 |
| ENSGALG00010005305 | SCAF8 | 421647 | -0.165598128 | 6.77461753 | 0.02522982 | 0.33877837 |
| ENSGALG00010002958 | DHRS12 | 770438 | 0.554298182 | 0.92991966 | 0.02529242 | 0.33892108 |
| ENSGALG00010006304 | NA | NA | -0.852556119 | -0.0994621 | 0.0253034 | 0.33892108 |
| ENSGALG00010000575 | SEC14L3 | 101751363 | 0.254486794 | 4.57125083 | 0.02530682 | 0.33892108 |
| ENSGALG00010013173 | KCNJ8 | 395973 | 0.427168415 | 1.96919792 | 0.02539429 | 0.33979554 |
| ENSGALG00010025600 | LOC423110 | 423110 | 0.172251773 | 5.58938516 | 0.02570267 | 0.34263071 |
| ENSGALG00010023263 | SRP14 | 423284 | 0.155763567 | 6.16868578 | 0.02571458 | 0.34263071 |
| ENSGALG00010003010 | NA | NA | -0.677857667 | 0.72423575 | 0.02578561 | 0.34263071 |
| ENSGALG00010004734 | NA | NA | -0.450068304 | 2.29589262 | 0.02582429 | 0.34263071 |
| ENSGALG00010028430 | NR2C2AP | 100858401 | 0.268728531 | 3.81834977 | 0.02584305 | 0.34263071 |
| ENSGALG00010027161 | C4BPG | 419853 | 0.35588171 | 2.89874579 | 0.0258478 | 0.34263071 |
| ENSGALG00010010416 | SLC35B3 | 420863 | -0.19929675 | 5.12700243 | 0.02590028 | 0.34263071 |
| ENSGALG00010012584 | NA | NA | -0.802669291 | 0.0289751 | 0.02592158 | 0.34263071 |
| ENSGALG00010023506 | RPE65 | 395700 | 0.606277743 | 0.78200735 | 0.02594047 | 0.34263071 |
| ENSGALG00010019673 | NID2 | 423583 | 0.214323505 | 6.79560901 | 0.02594091 | 0.34263071 |
| ENSGALG00010011250 | RESF1 | 418136 | -0.188329774 | 5.7777565 | 0.02594093 | 0.34263071 |
| ENSGALG00010028677 | LOC107055358 | 107055358 | 0.320741416 | 3.08027134 | 0.02595979 | 0.34263071 |
| ENSGALG00010014687 | CUL4A | 418744 | -0.177923694 | 5.09425369 | 0.02598462 | 0.34263071 |
| ENSGALG00010010469 | NA | NA | -0.721995343 | 0.48507468 | 0.02598695 | 0.34263071 |
| ENSGALG00010009334 | SS18L2 | 769135 | -0.419041989 | 1.93401969 | 0.02598837 | 0.34263071 |
| ENSGALG00010007993 | LRRC14 | 121110037 | 0.27408665 | 4.08442425 | 0.02599816 | 0.34263071 |
| ENSGALG00010016139 | LURAP1L | 768654 | -0.275913039 | 4.13636009 | 0.02600866 | 0.34263071 |
| ENSGALG00010012635 | NA | NA | -0.931277766 | -0.1452476 | 0.02601081 | 0.34263071 |
| ENSGALG00010028510 | CILP2 | 430666 | 0.651939493 | 1.15358243 | 0.02603108 | 0.34263071 |
| ENSGALG00010006531 | FAM161B | 423341 | 0.314524592 | 2.83050514 | 0.02606232 | 0.34274746 |

| | | | | | | |
|---|---|---|---|---|---|---|
| ENSGALG00010023884 | C2orf69 | 424064 | -0.219315912 | 4.94223505 | 0.02612515 | 0.3432791 |
| ENSGALG00010013046 | SLC12A5 | 777252 | 0.54889878 | 1.66812722 | 0.02617832 | 0.34368293 |
| ENSGALG00010010800 | BICRA | 101749950 | 0.2021295 | 5.67853148 | 0.02621417 | 0.34385898 |
| ENSGALG00010021397 | NA | NA | -0.535935584 | 1.15452212 | 0.02624333 | 0.34394704 |
| ENSGALG00010020295 | LGI1 | 423802 | -0.419076255 | 2.11786206 | 0.02630795 | 0.34448566 |
| ENSGALG00010025212 | P2RX4 | 374166 | 0.29161015 | 3.30391753 | 0.0263294 | 0.34448566 |
| ENSGALG00010020833 | RANBP1 | 416787 | 0.185684135 | 9.41170571 | 0.02636002 | 0.34459207 |
| ENSGALG00010000155 | MAZ | 107051057 | 0.205211826 | 6.80244342 | 0.02644537 | 0.345413 |
| ENSGALG00010024022 | NA | NA | -0.403697123 | 2.0320852 | 0.02649187 | 0.34572568 |
| ENSGALG00010014123 | NA | NA | -0.848448709 | 0.05583077 | 0.02652567 | 0.34587216 |
| ENSGALG00010020171 | RRN3 | 416414 | -0.156715893 | 6.88733726 | 0.02655262 | 0.34592919 |
| ENSGALG00010027278 | H1FX | 107054440 | 0.201904737 | 8.59336118 | 0.02662956 | 0.34652602 |
| ENSGALG00010005159 | ARHGAP21 | 420500 | -0.158088776 | 6.49137581 | 0.02664367 | 0.34652602 |
| ENSGALG00010027151 | C19orf70 | 100858091 | 0.259135388 | 3.65986716 | 0.02667292 | 0.34661222 |
| ENSGALG00010023692 | NA | NA | -0.361097383 | 2.59333489 | 0.02682927 | 0.34834851 |
| ENSGALG00010018060 | CASKIN1 | 416556 | 0.295220932 | 4.25489536 | 0.02692479 | 0.34929272 |
| ENSGALG00010023811 | MRPL44 | 424795 | 0.169267305 | 6.80673328 | 0.02705712 | 0.3502834 |
| ENSGALG00010006080 | LOC107056280 | 107056280 | 0.260039327 | 3.71108001 | 0.02707125 | 0.3502834 |
| ENSGALG00010029258 | TMEM100 | 417398 | 0.474003125 | 1.46501368 | 0.0270971 | 0.3502834 |
| ENSGALG00010002220 | NA | NA | -0.402849568 | 2.05405706 | 0.02710453 | 0.3502834 |
| ENSGALG00010018845 | NA | NA | -0.494917607 | 1.72755482 | 0.02713354 | 0.3502834 |
| ENSGALG00010023304 | GHDC | 420028 | 0.362805473 | 3.02697033 | 0.02713833 | 0.3502834 |
| ENSGALG00010024534 | VWF | 419031 | -0.190171683 | 6.337848 | 0.02728219 | 0.35168764 |
| ENSGALG00010023489 | ETV5 | 395749 | 0.198845261 | 7.55016746 | 0.02732893 | 0.35168764 |
| ENSGALG00010022224 | NHEJ1 | 424205 | 0.631698573 | 0.4753522 | 0.0273328 | 0.35168764 |
| ENSGALG00010027173 | AES | 100858542 | 0.17439674 | 5.53543372 | 0.02736569 | 0.35168764 |
| ENSGALG00010008113 | DBNDD1 | 415857 | 0.452705834 | 2.00680064 | 0.02736969 | 0.35168764 |
| ENSGALG00010022422 | RPS6KB2 | 429557 | 0.259545129 | 4.58149695 | 0.02739014 | 0.35168764 |
| ENSGALG00010019042 | PEX6 | 416709 | 0.203158043 | 5.48466424 | 0.02740781 | 0.35168764 |
| ENSGALG00010010271 | HUNK | 418493 | 0.167931409 | 5.82495531 | 0.02744091 | 0.35181778 |
| ENSGALG00010023116 | PROC | 395085 | -0.566803915 | 0.80643165 | 0.02759176 | 0.35340479 |
| ENSGALG00010022978 | ARL14EP | 421605 | -0.165799975 | 6.55687499 | 0.02761083 | 0.35340479 |
| ENSGALG00010016840 | LRRC34 | 424996 | -0.236778442 | 5.98341672 | 0.02764015 | 0.35348481 |
| ENSGALG00010013354 | WWP1 | 420212 | -0.193722234 | 5.15710533 | 0.02768244 | 0.35371483 |
| ENSGALG00010027304 | PTPN23 | 100858513 | 0.184754513 | 6.08906797 | 0.02770431 | 0.35371483 |
| ENSGALG00010000031 | NA | NA | -0.70728075 | 1.06934724 | 0.02773903 | 0.35373557 |
| ENSGALG00010017497 | NA | NA | -0.550758612 | 1.74665908 | 0.02778423 | 0.35373557 |
| ENSGALG00010017706 | FGFRL1 | 395107 | 0.303377439 | 3.03002602 | 0.02779251 | 0.35373557 |
| ENSGALG00010017189 | NA | NA | -0.617787928 | 0.92472789 | 0.02780497 | 0.35373557 |
| ENSGALG00010024178 | N4BP1 | 415739 | -0.164657278 | 5.65875479 | 0.02784378 | 0.35373557 |
| ENSGALG00010008331 | CCDC97 | 101748852 | 0.228848184 | 4.87992404 | 0.02786394 | 0.35373557 |
| ENSGALG00010013911 | TNIK | 424987 | 0.166183167 | 6.96181679 | 0.02788489 | 0.35373557 |
| ENSGALG00010015749 | ATR | 424777 | -0.15681315 | 6.41112851 | 0.02790867 | 0.35373557 |
| ENSGALG00010005984 | IDH3B | 426573 | 0.177734281 | 5.59378004 | 0.02792373 | 0.35373557 |
| ENSGALG00010010708 | NA | NA | -0.614498345 | 0.92431517 | 0.02797705 | 0.35373557 |
| ENSGALG00010017873 | HSPG2 | 429806 | 0.210006788 | 8.12538547 | 0.02799094 | 0.35373557 |
| ENSGALG00010019504 | NA | NA | -0.901598873 | -0.456067 | 0.02800112 | 0.35373557 |
| ENSGALG00010008762 | NA | NA | 0.169236801 | 6.12969609 | 0.02805246 | 0.35373557 |
| ENSGALG00010012599 | DMGDH | 416370 | -0.357349272 | 2.54285813 | 0.02807115 | 0.35373557 |
| ENSGALG00010014162 | COL18A1 | 373978 | 0.189256649 | 6.63714042 | 0.02808798 | 0.35373557 |
| ENSGALG00010007271 | CADM2 | 418463 | 0.477504696 | 1.36513063 | 0.02808893 | 0.35373557 |
| ENSGALG00010000011 | ND2 | 63549482 | -0.293602073 | 10.7312047 | 0.02809844 | 0.35373557 |
| ENSGALG00010002430 | NUMA1 | 768717 | 0.188440807 | 7.55289262 | 0.02814434 | 0.35379713 |

| | | | | | | |
|---|---|---|---|---|---|---|
| ENSGALG00010002443 | TYK2 | 430113 | 0.19799563 | 5.08713026 | 0.02814951 | 0.35379713 |
| ENSGALG00010006068 | PACRG | 421576 | 0.456788043 | 2.14381488 | 0.02827371 | 0.35506682 |
| ENSGALG00010015397 | SRD5A3 | 422750 | -0.238058615 | 3.97053132 | 0.02830504 | 0.35516914 |
| ENSGALG00010022025 | NA | NA | -0.560107571 | 0.7824658 | 0.02835499 | 0.35517627 |
| ENSGALG00010006850 | C5H14orf4 | 429422 | 0.155009822 | 6.73844005 | 0.02838942 | 0.35517627 |
| ENSGALG00010016871 | CNGA1 | 396143 | -0.662559544 | 0.47551612 | 0.02840733 | 0.35517627 |
| ENSGALG00010028198 | UCK1 | 771901 | 0.205324804 | 4.91857168 | 0.02843182 | 0.35517627 |
| ENSGALG00010029411 | SARM1 | 417568 | 0.312413451 | 4.12650373 | 0.02845006 | 0.35517627 |
| ENSGALG00010011621 | WASHC4 | 418077 | 0.15532613 | 6.93640678 | 0.02845584 | 0.35517627 |
| ENSGALG00010023626 | LOXL1 | 426411 | 0.222860028 | 5.80262178 | 0.02846788 | 0.35517627 |
| ENSGALG00010021719 | BHLHE40 | 416108 | 0.367074155 | 2.79125669 | 0.02852972 | 0.35538034 |
| ENSGALG00010006382 | PNN | 423338 | 0.16858232 | 7.84897169 | 0.02853063 | 0.35538034 |
| ENSGALG00010028500 | VPS45 | 430741 | 0.173982835 | 5.85323912 | 0.02860253 | 0.35598649 |
| ENSGALG00010016414 | KLB | 425467 | -0.7117492 | 0.45477923 | 0.02872955 | 0.35688844 |
| ENSGALG00010029506 | OXLD1 | 427815 | 0.318774576 | 2.80112464 | 0.02875769 | 0.35688844 |
| ENSGALG00010016620 | C15ORF40 | 415460 | -0.264006334 | 3.73746438 | 0.0287714 | 0.35688844 |
| ENSGALG00010021213 | FBXO44 | 419494 | 0.65672331 | 1.29810606 | 0.02877972 | 0.35688844 |
| ENSGALG00010004370 | ATP6V1H | 426199 | 0.188816552 | 5.94906755 | 0.02879147 | 0.35688844 |
| ENSGALG00010010983 | MED28 | 425350 | 0.209707659 | 4.79444211 | 0.02883281 | 0.35711194 |
| ENSGALG00010003457 | NA | NA | -0.477204068 | 1.53401234 | 0.0288622 | 0.35718722 |
| ENSGALG00010018533 | PCYOX1 | 426297 | 0.193750525 | 5.02828891 | 0.02892683 | 0.35742124 |
| ENSGALG00010023134 | KNL1 | 423007 | -0.174012113 | 8.2919318 | 0.02892777 | 0.35742124 |
| ENSGALG00010005457 | SEMA3D | 396332 | 0.36122872 | 2.79283147 | 0.02896203 | 0.35755627 |
| ENSGALG00010007980 | 1-Mar | 422420 | 0.182379755 | 5.25940914 | 0.02899606 | 0.35768809 |
| ENSGALG00010026291 | SERPING1 | 423132 | 0.240235099 | 4.9646572 | 0.02918791 | 0.359645 |
| ENSGALG00010025186 | TMEM120B | 416847 | 0.193078721 | 5.03042872 | 0.02920164 | 0.359645 |
| ENSGALG00010028892 | FAM129B | 771307 | 0.358761317 | 2.667174 | 0.0293396 | 0.36096209 |
| ENSGALG00010005695 | KIF5B | 420472 | -0.169168544 | 6.88169471 | 0.0293607 | 0.36096209 |
| ENSGALG00010024607 | LOC107051537 | 107051537 | 0.185444181 | 5.14884864 | 0.02937927 | 0.36096209 |
| ENSGALG00010020594 | PQLC2 | 419472 | 0.27516102 | 3.3111428 | 0.02945011 | 0.36154263 |
| ENSGALG00010011155 | NA | NA | 0.822291392 | -0.1456628 | 0.02951153 | 0.36200648 |
| ENSGALG00010016644 | NMRAL1 | 416672 | 0.227906525 | 4.85317704 | 0.02957216 | 0.3620775 |
| ENSGALG00010017894 | KAT6A | 426789 | -0.151162571 | 7.29952924 | 0.02959996 | 0.3620775 |
| ENSGALG00010015389 | NA | NA | -0.467924785 | 1.64958537 | 0.02963284 | 0.3620775 |
| ENSGALG00010009136 | KHK | 107053156 | 0.421928561 | 3.65466274 | 0.02964647 | 0.3620775 |
| ENSGALG00010003058 | NA | NA | -0.566742055 | 0.78421934 | 0.02966539 | 0.3620775 |
| ENSGALG00010018505 | CHTF18 | 416531 | 0.158358219 | 6.27188056 | 0.02971018 | 0.3620775 |
| ENSGALG00010001553 | CNGA4 | 430630 | 0.845825535 | -0.1303357 | 0.02971491 | 0.3620775 |
| ENSGALG00010022266 | COA3 | 772251 | 0.281520532 | 3.83901421 | 0.02975535 | 0.3620775 |
| ENSGALG00010020686 | C1orf52 | 424529 | 0.194545544 | 5.17616339 | 0.02976733 | 0.3620775 |
| ENSGALG00010017810 | HIF1A | 374177 | -0.175671074 | 7.305109 | 0.0297686 | 0.3620775 |
| ENSGALG00010001798 | PSTK | 771778 | 0.279734631 | 3.76621981 | 0.02977728 | 0.3620775 |
| ENSGALG00010025376 | LOC112529918 | 112529918 | -0.715793501 | 0.31418954 | 0.02982734 | 0.36239868 |
| ENSGALG00010025417 | B3GAT1 | 373949 | 0.466780326 | 1.84642633 | 0.02986454 | 0.36256305 |
| ENSGALG00010024827 | NA | NA | -0.161256266 | 6.12015897 | 0.02991244 | 0.36285703 |
| ENSGALG00010011895 | PHF21B | 427935 | 0.440927921 | 1.65898126 | 0.02994274 | 0.36293724 |
| ENSGALG00010021425 | PTPN9 | 415308 | 0.175069349 | 5.85105069 | 0.03001795 | 0.36356126 |
| ENSGALG00010014570 | KIF27 | 427456 | 0.248789331 | 3.78143544 | 0.03008206 | 0.36404992 |
| ENSGALG00010020129 | MGAT2 | 428925 | 0.183571047 | 5.38495345 | 0.03012547 | 0.36428563 |
| ENSGALG00010017585 | NA | NA | 0.344959892 | 2.78100077 | 0.03014909 | 0.36428563 |
| ENSGALG00010023079 | SLC25A16 | 423689 | -0.250280268 | 5.15253877 | 0.03019902 | 0.36460146 |
| ENSGALG00010020399 | ATP13A2 | 419466 | 0.239074201 | 5.63961139 | 0.030256 | 0.36500171 |
| ENSGALG00010000129 | NA | NA | 0.805579468 | 0.25847659 | 0.03028169 | 0.36502415 |

| | | | | | | |
|---|---|---|---|---|---|---|
| ENSGALG00010028119 | HYAL2 | 415908 | 0.221185902 | 4.86007957 | 0.03033409 | 0.36536838 |
| ENSGALG00010016907 | SQSTM1 | 770805 | -0.2405438 | 4.31137468 | 0.03039553 | 0.36544757 |
| ENSGALG00010009800 | NA | NA | -0.38597514 | 2.88662317 | 0.03041815 | 0.36544757 |
| ENSGALG00010022583 | FKBP11 | 107049506 | 0.381663527 | 2.27541367 | 0.03043076 | 0.36544757 |
| ENSGALG00010015038 | PCSK5 | 395456 | 0.157846611 | 6.40302019 | 0.03043848 | 0.36544757 |
| ENSGALG00010012889 | RIC1 | 427226 | -0.158772673 | 6.38749931 | 0.03045993 | 0.36544757 |
| ENSGALG00010016519 | MEX3B | 107054196 | 0.162094575 | 6.33438373 | 0.03052842 | 0.36598277 |
| ENSGALG00010006273 | ARL13B | 418431 | -0.19468968 | 5.23172429 | 0.03061882 | 0.36648075 |
| ENSGALG00010026117 | LZTR1 | 421166 | 0.195336354 | 5.26940856 | 0.03063891 | 0.36648075 |
| ENSGALG00010010342 | NA | NA | 0.816427783 | -0.1451899 | 0.03064172 | 0.36648075 |
| ENSGALG00010024696 | TRADD | 415700 | 0.327079904 | 3.45461369 | 0.03068412 | 0.36670159 |
| ENSGALG00010001685 | MIR16-2 | 777826 | -0.863128403 | -0.0125239 | 0.03076045 | 0.36718095 |
| ENSGALG00010021354 | ARHGAP39L | 419132 | 0.16905562 | 6.00205691 | 0.03077217 | 0.36718095 |
| ENSGALG00010017343 | DLG1 | 424890 | -0.159313524 | 7.13048492 | 0.03082632 | 0.36754092 |
| ENSGALG00010027872 | KDM5B | 421168 | 0.166474127 | 9.16720357 | 0.03089246 | 0.36804309 |
| ENSGALG00010013281 | BIRC8 | 395280 | -0.188715727 | 5.25361543 | 0.03099028 | 0.36865724 |
| ENSGALG00010009026 | RAPGEF2 | 422418 | -0.17628467 | 5.29710383 | 0.03099611 | 0.36865724 |
| ENSGALG00010002295 | YAF2 | 417790 | -0.20020248 | 6.0585038 | 0.03103735 | 0.36865724 |
| ENSGALG00010015660 | KLHL3 | 416303 | 0.202582471 | 4.60829769 | 0.031067 | 0.36865724 |
| ENSGALG00010005123 | MTIF3 | 418930 | 0.251844349 | 4.01468744 | 0.03107863 | 0.36865724 |
| ENSGALG00010015830 | NA | NA | 0.798059206 | -0.2921661 | 0.03113602 | 0.36865724 |
| ENSGALG00010024004 | EIF4B | 100858349 | 0.159802152 | 8.44879548 | 0.03113795 | 0.36865724 |
| ENSGALG00010026585 | NA | NA | -0.803792019 | -0.1291989 | 0.03114588 | 0.36865724 |
| ENSGALG00010018128 | SEPTIN12 | 416408 | 0.287639322 | 3.80920777 | 0.03122777 | 0.36865724 |
| ENSGALG00010010973 | PGAP1 | 424054 | -0.233025758 | 4.33200947 | 0.03122966 | 0.36865724 |
| ENSGALG00010021679 | MASP2 | 407089 | -0.295618682 | 3.20817595 | 0.03126528 | 0.36865724 |
| ENSGALG00010012719 | COL10A1 | 100858979 | 0.846438541 | -0.3622367 | 0.03129047 | 0.36865724 |
| ENSGALG00010000310 | NA | NA | 0.41446995 | 2.14931085 | 0.03131208 | 0.36865724 |
| ENSGALG00010022852 | NA | NA | -0.315219512 | 3.77268526 | 0.03131807 | 0.36865724 |
| ENSGALG00010029570 | TNFAIP1 | 417672 | -0.185415525 | 7.53779177 | 0.03131932 | 0.36865724 |
| ENSGALG00010024034 | LRP2 | 424168 | -0.187473701 | 9.39951469 | 0.03132901 | 0.36865724 |
| ENSGALG00010010347 | FBXW11 | 416181 | -0.165234288 | 5.87273505 | 0.0313746 | 0.36891044 |
| ENSGALG00010002269 | MCPH1 | 100125976 | -0.179835579 | 5.24670797 | 0.03144341 | 0.36943598 |
| ENSGALG00010020612 | RPL27A | 770018 | 0.223148931 | 8.56776555 | 0.03153926 | 0.37027817 |
| ENSGALG00010018284 | SPEG | 429033 | 0.258957846 | 3.68042633 | 0.03171459 | 0.37205149 |
| ENSGALG00010013878 | ARID2 | 417804 | -0.163745496 | 5.61097124 | 0.03175748 | 0.37226955 |
| ENSGALG00010021256 | PML | 415302 | 0.373024 | 3.43644495 | 0.03190236 | 0.37349793 |
| ENSGALG00010005430 | MCM4 | 426764 | -0.176026469 | 7.68069221 | 0.03193578 | 0.37349793 |
| ENSGALG00010006445 | NA | NA | -0.33203029 | 3.53969362 | 0.03196291 | 0.37349793 |
| ENSGALG00010006903 | B3GLCT | 428075 | 0.167864038 | 5.59233574 | 0.03201987 | 0.37349793 |
| ENSGALG00010029385 | CLIP2 | 417488 | 0.188322683 | 5.78771262 | 0.03202821 | 0.37349793 |
| ENSGALG00010026950 | LHFPL5 | 378916 | 0.383120085 | 2.12914379 | 0.0320626 | 0.37349793 |
| ENSGALG00010007632 | NA | NA | 0.415286315 | 2.52354133 | 0.0320717 | 0.37349793 |
| ENSGALG00010008186 | GLTSCR2 | 107049993 | 0.210030753 | 5.47136963 | 0.03207594 | 0.37349793 |
| ENSGALG00010001178 | ZBTB5 | 431260 | -0.151648686 | 6.21747996 | 0.03208167 | 0.37349793 |
| ENSGALG00010021252 | XPC | 416039 | -0.156977056 | 5.64905686 | 0.03217166 | 0.37381485 |
| ENSGALG00010021493 | UNC93B1 | 395244 | 0.253824332 | 4.18288546 | 0.03220164 | 0.37381485 |
| ENSGALG00010029508 | CCDC137 | 417454 | 0.217519377 | 4.70253194 | 0.03221524 | 0.37381485 |
| ENSGALG00010016868 | NIPAL1 | 428783 | -0.280241214 | 5.66515731 | 0.0322225 | 0.37381485 |
| ENSGALG00010029410 | VTN | 395935 | 0.210472454 | 5.90416316 | 0.03223089 | 0.37381485 |
| ENSGALG00010020383 | BAAT | 769879 | 0.172651882 | 5.89842133 | 0.03236358 | 0.37495513 |
| ENSGALG00010004181 | BBS12 | 422667 | -0.178706235 | 5.5237425 | 0.03240005 | 0.37495513 |
| ENSGALG00010028183 | CRTC1 | 771015 | 0.20132965 | 4.75782041 | 0.03241923 | 0.37495513 |

| | | | | | | |
|---|---|---|---|---|---|---|
| ENSGALG00010006740 | BRCA2 | 374139 | -0.163468932 | 7.00911122 | 0.0324271 | 0.37495513 |
| ENSGALG00010019622 | C2orf42 | 426298 | 0.253895642 | 3.79103066 | 0.03254055 | 0.37592434 |
| ENSGALG00010008099 | PANK3 | 416170 | -0.177594692 | 8.49953989 | 0.03255999 | 0.37592434 |
| ENSGALG00010010821 | RGS8 | 426557 | -0.495398278 | 1.34349343 | 0.03261858 | 0.37611879 |
| ENSGALG00010024479 | ERBB2 | 426130 | 0.235514583 | 4.75443079 | 0.03264091 | 0.37611879 |
| ENSGALG00010016577 | ZBTB17 | 396541 | 0.175483124 | 5.69417948 | 0.03265048 | 0.37611879 |
| ENSGALG00010028070 | MEX3A | 101747646 | 0.187325474 | 5.78364372 | 0.03274294 | 0.37681381 |
| ENSGALG00010000323 | AP1B1 | 417021 | 0.15348061 | 7.11092951 | 0.03276 | 0.37681381 |
| ENSGALG00010011091 | YARS2 | 418131 | -0.14742937 | 6.60714395 | 0.03278935 | 0.37686842 |
| ENSGALG00010023621 | CD79B | 419940 | 0.924086653 | 0.08349136 | 0.03284167 | 0.37718686 |
| ENSGALG00010001581 | NA | NA | -0.523845073 | 1.42306194 | 0.03287735 | 0.37728676 |
| ENSGALG00010010257 | NA | NA | -0.254013329 | 6.18050445 | 0.03289962 | 0.37728676 |
| ENSGALG00010010938 | FZD6 | 378788 | -0.193161495 | 4.94006135 | 0.03303046 | 0.37825355 |
| ENSGALG00010012835 | NA | NA | -0.554539679 | 0.71657438 | 0.03306148 | 0.37825355 |
| ENSGALG00010029521 | ACTG1L | 776816 | 0.166067553 | 11.3879211 | 0.03306674 | 0.37825355 |
| ENSGALG00010002338 | CD99 | 418662 | -0.183917455 | 5.45046595 | 0.03308268 | 0.37825355 |
| ENSGALG00010014469 | TMCO3 | 395675 | -0.219420294 | 4.05490741 | 0.03313273 | 0.37835688 |
| ENSGALG00010027040 | CLIP1 | 395784 | -0.156036911 | 6.38034833 | 0.03315021 | 0.37835688 |
| ENSGALG00010006072 | LOC107056274 | 107056274 | 0.199121415 | 5.6008655 | 0.03316763 | 0.37835688 |
| ENSGALG00010001627 | ARFIP2 | 419075 | 0.186327454 | 5.57194115 | 0.0331905 | 0.37835688 |
| ENSGALG00010015083 | CEP135 | 422746 | -0.190640122 | 4.81696643 | 0.03329099 | 0.37886598 |
| ENSGALG00010022248 | FIGNL2 | 431296 | 0.213295575 | 5.32526396 | 0.03331636 | 0.37886598 |
| ENSGALG00010013187 | KDF1 | 769268 | 0.185827166 | 5.57101669 | 0.0333551 | 0.37886598 |
| ENSGALG00010017928 | NA | NA | 0.820352878 | -0.3266903 | 0.033359 | 0.37886598 |
| ENSGALG00010013431 | TMEM175 | 427293 | -0.14752113 | 6.42803241 | 0.03340789 | 0.37886598 |
| ENSGALG00010001364 | SMCHD1 | 421057 | -0.187940552 | 8.00939767 | 0.03341245 | 0.37886598 |
| ENSGALG00010016802 | TMEM222 | 419582 | 0.208969063 | 4.94850911 | 0.03343097 | 0.37886598 |
| ENSGALG00010029534 | NF1 | 396085 | 0.155309918 | 6.82172148 | 0.03343299 | 0.37886598 |
| ENSGALG00010002411 | XNDC1N | 100857925 | 0.281049128 | 3.48098247 | 0.03345807 | 0.37887001 |
| ENSGALG00010000311 | NA | NA | 0.795084011 | -0.1611927 | 0.03354357 | 0.37955762 |
| ENSGALG00010005534 | IDI2 | 420459 | -0.2394204 | 6.52305557 | 0.03359663 | 0.37986148 |
| ENSGALG00010022683 | NA | NA | -0.352312682 | 2.37774906 | 0.03362235 | 0.37986148 |
| ENSGALG00010025738 | SOX13 | 419820 | 0.205827217 | 5.33703997 | 0.0336448 | 0.37986148 |
| ENSGALG00010029317 | KPNA2 | 417420 | -0.18296184 | 8.15335434 | 0.03367142 | 0.37988201 |
| ENSGALG00010000697 | LOC107050463 | 107050463 | 0.207460801 | 4.82773477 | 0.03372029 | 0.38001706 |
| ENSGALG00010027889 | RBM8A | 107050968 | 0.156501835 | 6.9399083 | 0.03373299 | 0.38001706 |
| ENSGALG00010002908 | S1PR2 | 100857976 | 0.268247795 | 4.24900244 | 0.03380287 | 0.38028999 |
| ENSGALG00010000054 | VPS33B | 425858 | 0.158688024 | 6.89086062 | 0.03380686 | 0.38028999 |
| ENSGALG00010025245 | RAP1A | 419867 | -0.233169905 | 3.81792805 | 0.03383712 | 0.38035108 |
| ENSGALG00010009373 | ARHGAP15 | 424303 | -0.408586487 | 2.53029361 | 0.0338781 | 0.38053255 |
| ENSGALG00010000038 | NA | NA | -0.459119795 | 1.79224548 | 0.03411044 | 0.38278135 |
| ENSGALG00010021330 | COMMD4 | 770048 | 0.271371616 | 3.8040849 | 0.03415076 | 0.38278135 |
| ENSGALG00010014894 | RAB22A | 419318 | -0.208174439 | 4.50019287 | 0.03417771 | 0.38278135 |
| ENSGALG00010018544 | HACD2 | 424251 | -0.154850026 | 7.75315104 | 0.03419107 | 0.38278135 |
| ENSGALG00010028174 | GMPPB | 415924 | 0.265278243 | 3.71712783 | 0.03420323 | 0.38278135 |
| ENSGALG00010024152 | ZNF423 | 415738 | 0.15794447 | 5.97917695 | 0.03431601 | 0.38376318 |
| ENSGALG00010019264 | HK2 | 374044 | 0.217055059 | 5.2364296 | 0.0343587 | 0.38396031 |
| ENSGALG00010027079 | PM20D1 | 419838 | 0.304815542 | 3.44762444 | 0.03438882 | 0.38401679 |
| ENSGALG00010019371 | TMEM87B | 423231 | -0.15526201 | 7.58572987 | 0.03450137 | 0.38499309 |
| ENSGALG00010029487 | CDC42EP4 | 771509 | -0.334023762 | 2.42242507 | 0.03459066 | 0.38534389 |
| ENSGALG00010011374 | RPEL1 | 770045 | 0.214479603 | 5.02973707 | 0.03461906 | 0.38534389 |
| ENSGALG00010021452 | SLC25A1 | 416764 | 0.296701226 | 3.42490944 | 0.03463643 | 0.38534389 |
| ENSGALG00010021466 | HES4 | 395128 | 0.198808113 | 5.31828207 | 0.03465713 | 0.38534389 |

| | | | | | | |
|---|---|---|---|---|---|---|
| ENSGALG00010001698 | RPL15 | 428442 | 0.174563006 | 9.32216363 | 0.03466993 | 0.38534389 |
| ENSGALG00010029689 | NA | NA | 0.190776432 | 5.83052572 | 0.03470583 | 0.38534389 |
| ENSGALG00010007695 | RASGRP3 | 421460 | -0.627733996 | 0.69060343 | 0.03470887 | 0.38534389 |
| ENSGALG00010024797 | IKZF4 | 101747801 | 0.285971635 | 3.5999314 | 0.03478951 | 0.38595947 |
| ENSGALG00010000350 | LOC100857358 | 100857358 | 0.171302068 | 8.32648737 | 0.03503882 | 0.38844412 |
| ENSGALG00010011329 | PCDHGC3 | 408051 | 0.212800667 | 4.79744647 | 0.03509671 | 0.38880453 |
| ENSGALG00010010985 | COL4A3BP | 427209 | -0.158134149 | 7.01437417 | 0.03519107 | 0.38938314 |
| ENSGALG00010021367 | GRIP2 | 416043 | -0.40324712 | 2.75776833 | 0.03519977 | 0.38938314 |
| ENSGALG00010013964 | SLC38A1 | 417806 | -0.146565647 | 7.86353134 | 0.03533236 | 0.39032277 |
| ENSGALG00010017017 | NA | NA | 0.25775096 | 3.69017967 | 0.03533566 | 0.39032277 |
| ENSGALG00010000501 | SCYL1 | 107049850 | 0.273695394 | 4.42063803 | 0.03537856 | 0.39051508 |
| ENSGALG00010028674 | NA | NA | 0.214314682 | 4.16926889 | 0.03541702 | 0.39065815 |
| ENSGALG00010005816 | NA | NA | -0.708153087 | 0.08391307 | 0.03544359 | 0.39066997 |
| ENSGALG00010028124 | SUGP2 | 420113 | -0.174447965 | 5.45004873 | 0.03549195 | 0.39073904 |
| ENSGALG00010010186 | TNPO1 | 427218 | -0.157415448 | 7.33380084 | 0.03550663 | 0.39073904 |
| ENSGALG00010010982 | ZBTB21 | 427986 | -0.155079093 | 6.60076723 | 0.03556124 | 0.39073904 |
| ENSGALG00010004869 | NA | NA | -0.516717451 | 1.37827525 | 0.03556199 | 0.39073904 |
| ENSGALG00010017085 | SRSF7 | 100859609 | 0.195657053 | 4.78085339 | 0.03557737 | 0.39073904 |
| ENSGALG00010020544 | BMERB1 | 416594 | -0.148785124 | 6.62292565 | 0.03572974 | 0.39213137 |
| ENSGALG00010020770 | B4GALNT4 | 770601 | 0.15337178 | 7.59551666 | 0.03583079 | 0.39281616 |
| ENSGALG00010009640 | NPR2 | 100859339 | 0.220102807 | 4.87917153 | 0.03584342 | 0.39281616 |
| ENSGALG00010014908 | PPP4R1L | 419319 | -0.160204632 | 6.33366585 | 0.03589078 | 0.39305411 |
| ENSGALG00010012905 | NKAIN1 | 107049081 | 0.201615251 | 5.50575555 | 0.03602321 | 0.39422252 |
| ENSGALG00010006909 | NA | NA | -0.487154535 | 1.40354828 | 0.03609363 | 0.39446851 |
| ENSGALG00010027870 | HAX1 | 107055138 | 0.271333023 | 4.45709553 | 0.03610197 | 0.39446851 |
| ENSGALG00010001761 | ELOB | 121108722 | 0.163740482 | 5.99553301 | 0.03616194 | 0.39446851 |
| ENSGALG00010003165 | BMP2K | 422578 | -0.147186524 | 6.76529574 | 0.03619904 | 0.39446851 |
| ENSGALG00010005813 | NA | NA | -0.817365098 | -0.0839207 | 0.03620692 | 0.39446851 |
| ENSGALG00010018029 | LOC121107471 | 121107471 | 0.468468407 | 1.50933319 | 0.0362243 | 0.39446851 |
| ENSGALG00010013136 | NA | NA | -0.250641863 | 3.64343646 | 0.03624068 | 0.39446851 |
| ENSGALG00010015317 | NA | NA | -0.367139592 | 2.11886129 | 0.0362927 | 0.39446851 |
| ENSGALG00010004352 | CHMP2A | 776036 | 0.196124226 | 6.16242334 | 0.03630684 | 0.39446851 |
| ENSGALG00010023473 | BOK | 395445 | 0.177100995 | 4.91677792 | 0.0363742 | 0.39446851 |
| ENSGALG00010002063 | NA | NA | 0.285623916 | 3.9634184 | 0.03637505 | 0.39446851 |
| ENSGALG00010028109 | TJP3 | 420070 | 0.321875359 | 3.56067947 | 0.03639395 | 0.39446851 |
| ENSGALG00010021978 | NR2C2 | 395253 | -0.177697677 | 5.28386489 | 0.0364112 | 0.39446851 |
| ENSGALG00010024618 | VPS13C | 415376 | -0.183856845 | 5.74926664 | 0.03644942 | 0.39446851 |
| ENSGALG00010010499 | NR4A2 | 424320 | -0.468393286 | 1.39298312 | 0.03649041 | 0.39446851 |
| ENSGALG00010007887 | CDT1 | 426254 | 0.171319361 | 7.09013368 | 0.03649387 | 0.39446851 |
| ENSGALG00010009480 | NA | NA | -0.648554049 | 0.8873572 | 0.03651467 | 0.39446851 |
| ENSGALG00010021383 | MSI1 | 416979 | 0.14936863 | 6.75363642 | 0.0365192 | 0.39446851 |
| ENSGALG00010022831 | LRRC8C | 424515 | -0.271462926 | 3.6428989 | 0.03653488 | 0.39446851 |
| ENSGALG00010028326 | CRB2 | 417104 | -0.17261364 | 5.57204931 | 0.03660335 | 0.39492947 |
| ENSGALG00010023903 | TBKBP1 | 430679 | 0.288132345 | 3.86015322 | 0.03664436 | 0.39509377 |
| ENSGALG00010015935 | EPHX1 | 421321 | 0.553401515 | 1.55009067 | 0.03671193 | 0.39554396 |
| ENSGALG00010025054 | NA | NA | -0.532133926 | 0.97264914 | 0.03679587 | 0.39589968 |
| ENSGALG00010027120 | UNC13A | 430487 | 0.701562069 | 0.09701192 | 0.03679663 | 0.39589968 |
| ENSGALG00010010346 | TFB2M | 421488 | -0.230490866 | 3.8585226 | 0.03689188 | 0.39664591 |
| ENSGALG00010001406 | EML3 | 107049450 | 0.280591555 | 3.22627686 | 0.03696166 | 0.39711753 |
| ENSGALG00010016308 | RPS6 | 396148 | 0.164491433 | 10.5159435 | 0.03703216 | 0.39736757 |
| ENSGALG00010021528 | BRPF1 | 427590 | 0.189288742 | 5.39777845 | 0.03705819 | 0.39736757 |
| ENSGALG00010001167 | SMC4 | 395189 | -0.180147244 | 5.69233372 | 0.03706274 | 0.39736757 |
| ENSGALG00010002143 | SLC16A13 | 425860 | 0.481021201 | 1.36461458 | 0.03712957 | 0.39780573 |

| | | | | | | |
|---|---|---|---|---|---|---|
| ENSGALG00010025765 | NA | NA | 0.276565821 | 3.18741268 | 0.0371586 | 0.3978385 |
| ENSGALG00010012798 | MFSD6 | 423978 | -0.186779761 | 4.73474637 | 0.03721569 | 0.39804363 |
| ENSGALG00010004046 | IPO4 | 101750157 | 0.183198229 | 5.3610626 | 0.03722972 | 0.39804363 |
| ENSGALG00010003108 | PPP1R3B | 422630 | 0.165625408 | 7.09542017 | 0.03728768 | 0.39838536 |
| ENSGALG00010024864 | MPHOSPH9 | 416829 | -0.161693439 | 5.41117281 | 0.03738635 | 0.39896765 |
| ENSGALG00010014813 | SASH3 | 422138 | 0.276406724 | 3.21204441 | 0.03739427 | 0.39896765 |
| ENSGALG00010010867 | QSOX1 | 373914 | 0.167294627 | 6.25981443 | 0.03753758 | 0.39971989 |
| ENSGALG00010009079 | KCTD15 | 415779 | 0.16462169 | 5.75046897 | 0.03754019 | 0.39971989 |
| ENSGALG00010027303 | APEH | 415926 | 0.194271303 | 5.80933136 | 0.03754304 | 0.39971989 |
| ENSGALG00010006270 | STAM | 420517 | -0.154285719 | 7.25129658 | 0.03760717 | 0.40012465 |
| ENSGALG00010007923 | YAP1 | 396171 | 0.19534273 | 7.24392105 | 0.03764061 | 0.40017836 |
| ENSGALG00010005594 | MRPL50 | 427307 | 0.215302368 | 4.72621786 | 0.03766446 | 0.40017836 |
| ENSGALG00010023482 | SOGA1 | 419159 | 0.237270008 | 5.39087555 | 0.03783993 | 0.40158253 |
| ENSGALG00010009499 | LOC101751218 | 101751218 | 0.312273004 | 3.55871344 | 0.03789172 | 0.40158253 |
| ENSGALG00010009273 | MAL2 | 420280 | -0.209303382 | 5.0006721 | 0.03792551 | 0.40158253 |
| ENSGALG00010015122 | COL23A1 | 425481 | 0.273998602 | 3.17411169 | 0.03792944 | 0.40158253 |
| ENSGALG00010008231 | ZCCHC6 | 100857368 | -0.150903795 | 6.19872981 | 0.03793986 | 0.40158253 |
| ENSGALG00010012671 | NBN | 374246 | -0.224848353 | 5.08274947 | 0.03795389 | 0.40158253 |
| ENSGALG00010004128 | MCMDC2 | 428360 | -0.64474643 | 0.38259589 | 0.0380698 | 0.40253096 |
| ENSGALG00010012378 | RPS19BP1 | 418011 | 0.234686999 | 4.94897687 | 0.0382525 | 0.40418378 |
| ENSGALG00010020900 | PDHB | 416066 | 0.153605907 | 6.39185072 | 0.03829942 | 0.4044007 |
| ENSGALG00010021329 | PSMC3IP | 772119 | 0.168522329 | 5.02359483 | 0.03846781 | 0.40575125 |
| ENSGALG00010006254 | PROS1 | 418430 | -0.1730729 | 5.68396048 | 0.03848029 | 0.40575125 |
| ENSGALG00010015570 | LIMCH1 | 422780 | -0.188904261 | 4.51552031 | 0.03856995 | 0.40605141 |
| ENSGALG00010024682 | LOC100859737 | 100859737 | 0.195164518 | 10.0821181 | 0.03864378 | 0.40605141 |
| ENSGALG00010017594 | CEP68 | 421274 | 0.226556244 | 4.35233972 | 0.03865933 | 0.40605141 |
| ENSGALG00010027270 | COL7A1 | 427584 | 0.233844732 | 4.27718623 | 0.03866606 | 0.40605141 |
| ENSGALG00010004500 | NA | NA | 0.925633351 | -0.7301285 | 0.0386907 | 0.40605141 |
| ENSGALG00010025268 | ZBTB39 | 100858734 | 0.171158676 | 6.25605521 | 0.03873515 | 0.40605141 |
| ENSGALG00010003533 | LOC121107019 | 121107019 | -1.061635036 | 2.47530787 | 0.03873767 | 0.40605141 |
| ENSGALG00010019237 | SORCS3 | 423884 | -0.164720951 | 6.36298861 | 0.03879331 | 0.40605141 |
| ENSGALG00010019620 | NA | NA | -0.276856293 | 3.97276083 | 0.03885619 | 0.40605141 |
| ENSGALG00010003680 | TRAPPC5 | 100858483 | 0.162806523 | 7.2916099 | 0.03887877 | 0.40605141 |
| ENSGALG00010024766 | CLEC18B | 431137 | 0.478939882 | 1.68923392 | 0.03887922 | 0.40605141 |
| ENSGALG00010028573 | KCNN1 | 429723 | 0.221454291 | 4.4503316 | 0.03890316 | 0.40605141 |
| ENSGALG00010028464 | TARS2 | 770407 | 0.214909212 | 5.84128697 | 0.03891055 | 0.40605141 |
| ENSGALG00010028471 | TPGS1 | 100857535 | 0.304530432 | 2.82075356 | 0.03892356 | 0.40605141 |
| ENSGALG00010029394 | CYGB | 427802 | 0.410536146 | 2.99297007 | 0.03892527 | 0.40605141 |
| ENSGALG00010015500 | NA | NA | -0.656988241 | 0.66463657 | 0.03893281 | 0.40605141 |
| ENSGALG00010002803 | NA | NA | -0.244630045 | 4.55180335 | 0.03909407 | 0.40740519 |
| ENSGALG00010013320 | CALCRL | 423990 | -0.174639455 | 5.70398718 | 0.03911979 | 0.40740519 |
| ENSGALG00010018198 | BTBD3 | 427688 | -0.148331038 | 5.89115759 | 0.03914734 | 0.40740519 |
| ENSGALG00010027733 | RPL35 | 374133 | 0.179409314 | 8.31100501 | 0.03916897 | 0.40740519 |
| ENSGALG00010023637 | RHBDD1 | 424800 | -0.19551822 | 4.56741454 | 0.03923459 | 0.40781082 |
| ENSGALG00010027388 | URM1 | 417221 | 0.197878238 | 6.30587745 | 0.03929974 | 0.40813647 |
| ENSGALG00010008437 | NA | NA | 0.213912906 | 5.02248486 | 0.03932241 | 0.40813647 |
| ENSGALG00010019679 | RHOBTB2 | 419536 | 0.159682399 | 5.77721455 | 0.03934584 | 0.40813647 |
| ENSGALG00010009407 | PTPN12 | 417725 | -0.173382736 | 8.89950603 | 0.03952203 | 0.40968679 |
| ENSGALG00010015413 | CARNMT1 | 427256 | -0.187274836 | 7.37652978 | 0.03956006 | 0.40980365 |
| ENSGALG00010018177 | KIF20B | 423793 | -0.14215392 | 7.801495 | 0.039601 | 0.40995058 |
| ENSGALG00010026338 | NA | NA | 0.403062912 | 2.00276214 | 0.03966474 | 0.41033325 |
| ENSGALG00010001307 | LOC107050717 | 107050717 | 0.165896154 | 6.94406673 | 0.03970292 | 0.410451 |
| ENSGALG00010024741 | RILPL2 | 416824 | 0.252032998 | 4.28612098 | 0.03975095 | 0.41067044 |

| | | | | | | |
|---|---|---|---|---|---|---|
| ENSGALG00010029636 | SPATA20 | 422100 | 0.278473905 | 4.84597018 | 0.03984169 | 0.41096151 |
| ENSGALG00010019786 | R3HDM1 | 424292 | -0.145392498 | 9.18129703 | 0.03984857 | 0.41096151 |
| ENSGALG00010012973 | GLDC | 374222 | 0.144471422 | 6.9390335 | 0.03987578 | 0.41096151 |
| ENSGALG00010028167 | NA | NA | -0.200380985 | 6.13213807 | 0.03988642 | 0.41096151 |
| ENSGALG00010020252 | DMTN | 776542 | 0.271287103 | 4.23875671 | 0.04003385 | 0.41153474 |
| ENSGALG00010027168 | SSBP4 | 107055368 | 0.17344637 | 5.90780704 | 0.04005048 | 0.41153474 |
| ENSGALG00010005382 | RAB32 | 421616 | 0.162355672 | 5.50918357 | 0.04005888 | 0.41153474 |
| ENSGALG00010005024 | NA | NA | -0.873425342 | 0.36089166 | 0.04008519 | 0.41153474 |
| ENSGALG00010020009 | MACROD2 | 421259 | 0.264446234 | 3.77198833 | 0.0400907 | 0.41153474 |
| ENSGALG00010006865 | EFCAB2 | 421491 | -0.957458165 | -0.7305142 | 0.0401137 | 0.41153474 |
| ENSGALG00010023836 | NDUFS3 | 423179 | 0.172325919 | 5.55585083 | 0.04013008 | 0.41153474 |
| ENSGALG00010016967 | ERLEC1 | 101751309 | 0.156070062 | 5.63087539 | 0.04016133 | 0.41157975 |
| ENSGALG00010000643 | VKORC1 | 404205 | 0.211615006 | 4.86569613 | 0.04026182 | 0.41195979 |
| ENSGALG00010023547 | NA | NA | -0.18820025 | 5.52282856 | 0.04026802 | 0.41195979 |
| ENSGALG00010014367 | OPHN1 | 422162 | 0.18511371 | 5.25846413 | 0.04027908 | 0.41195979 |
| ENSGALG00010014439 | LOC112532140 | 112532140 | 0.353854373 | 2.61045588 | 0.04034856 | 0.4122581 |
| ENSGALG00010027290 | CSPG5 | 395466 | 0.233094866 | 4.24256283 | 0.04036206 | 0.4122581 |
| ENSGALG00010011358 | POU4F1L | 395521 | 0.558442396 | 0.67297693 | 0.04047685 | 0.41299346 |
| ENSGALG00010021229 | GLO1 | 421428 | -0.206019708 | 6.33525784 | 0.04048797 | 0.41299346 |
| ENSGALG00010023016 | LOC112533478 | 112533478 | -0.548787253 | 0.85516291 | 0.0405718 | 0.41339282 |
| ENSGALG00010026457 | SSRP1 | 396509 | 0.14970497 | 8.7316756 | 0.04058109 | 0.41339282 |
| ENSGALG00010005082 | LOC418927 | 418927 | -0.150346496 | 7.13476146 | 0.04074008 | 0.4142106 |
| ENSGALG00010006800 | NA | NA | -0.723693434 | 0.16017592 | 0.04074872 | 0.4142106 |
| ENSGALG00010011877 | NA | NA | -0.292463407 | 4.36389147 | 0.04076248 | 0.4142106 |
| ENSGALG00010013171 | SLC9A8 | 419356 | -0.164720532 | 5.27205367 | 0.04077729 | 0.4142106 |
| ENSGALG00010029376 | MXRA7 | 769894 | 0.188663693 | 4.71447878 | 0.04079654 | 0.4142106 |
| ENSGALG00010023334 | RPS17 | 374053 | 0.18217717 | 9.06656896 | 0.04100026 | 0.41573405 |
| ENSGALG00010016692 | PRKCI | 424991 | -0.174933759 | 6.90519386 | 0.04100086 | 0.41573405 |
| ENSGALG00010029350 | MRPL12 | 769031 | 0.21451609 | 5.43248009 | 0.04112551 | 0.41672219 |
| ENSGALG00010019123 | PLEKHO2 | 769198 | 0.381618156 | 1.98092952 | 0.04119926 | 0.41712917 |
| ENSGALG00010024507 | NA | NA | -0.469136417 | 1.2056309 | 0.04124104 | 0.41712917 |
| ENSGALG00010010217 | NA | NA | -0.634592985 | 0.48539277 | 0.04124735 | 0.41712917 |
| ENSGALG00010024090 | LOC112532597 | 112532597 | -0.488456583 | 1.51197435 | 0.04128218 | 0.41713829 |
| ENSGALG00010007414 | ROBO2 | 395291 | -0.211505705 | 4.60762316 | 0.04130949 | 0.41713829 |
| ENSGALG00010028595 | SCNM1 | 425744 | 0.250216563 | 4.15039081 | 0.04133642 | 0.41713829 |
| ENSGALG00010004109 | AGMO | 772374 | 0.667123011 | 0.04329505 | 0.04135716 | 0.41713829 |
| ENSGALG00010027964 | RABEPK | 772357 | 0.167695374 | 5.10672634 | 0.0413858 | 0.41715254 |
| ENSGALG00010005274 | LOC772017 | 772017 | 0.462341968 | 1.5627566 | 0.04145576 | 0.41758295 |
| ENSGALG00010005985 | DKK2 | 422531 | -0.589765519 | 0.58251004 | 0.04158976 | 0.41820381 |
| ENSGALG00010014610 | NA | NA | -0.174650146 | 5.55075986 | 0.04159166 | 0.41820381 |
| ENSGALG00010013247 | ATG5 | 421784 | 0.153377723 | 5.60477984 | 0.04159928 | 0.41820381 |
| ENSGALG00010004240 | LOC421792 | 421792 | 0.26481749 | 3.55614126 | 0.04164612 | 0.41840017 |
| ENSGALG00010014180 | SLC19A1 | 424034 | 0.166810113 | 5.89835788 | 0.04173715 | 0.41862887 |
| ENSGALG00010026712 | NA | NA | -0.596362092 | 0.71722077 | 0.04175108 | 0.41862887 |
| ENSGALG00010009991 | LHFPL3 | 100858127 | 0.173117681 | 5.01377412 | 0.04177314 | 0.41862887 |
| ENSGALG00010009738 | LOC101747507 | 101747507 | 0.235741608 | 3.8830178 | 0.04177854 | 0.41862887 |
| ENSGALG00010028692 | RALGPS1 | 417089 | -0.171639559 | 5.04303741 | 0.041808 | 0.41862887 |
| ENSGALG00010027899 | PMVK | 101747962 | 0.211869138 | 4.6669136 | 0.04183283 | 0.41862887 |
| ENSGALG00010017729 | MIEF2 | 416516 | 0.293802003 | 2.96806154 | 0.04189513 | 0.41897867 |
| ENSGALG00010001343 | LOC121108665 | 121108665 | -0.760167811 | -0.1141247 | 0.04194163 | 0.41917009 |
| ENSGALG00010024518 | LOC771136 | 771136 | -0.806937678 | -0.2586748 | 0.04202417 | 0.41972115 |
| ENSGALG00010020697 | NA | NA | -0.675122893 | 0.19756675 | 0.04214368 | 0.42021717 |
| ENSGALG00010019683 | TMEM184A | 769655 | 0.393665104 | 1.81010938 | 0.0421534 | 0.42021717 |

| | | | | | | |
|---|---|---|---|---|---|---|
| ENSGALG00010017733 | TRIM62 | 429807 | 0.224923945 | 4.90955577 | 0.04217915 | 0.42021717 |
| ENSGALG00010002370 | BEND2 | 772331 | -0.216486266 | 4.38865452 | 0.04220507 | 0.42021717 |
| ENSGALG00010002983 | NSMAF | 421137 | -0.214628965 | 4.36972145 | 0.04222 | 0.42021717 |
| ENSGALG00010012587 | MBLAC2 | 431565 | -0.165807206 | 6.37702663 | 0.04223839 | 0.42021717 |
| ENSGALG00010025624 | C1orf228 | 101751863 | -0.56524889 | 0.63745694 | 0.04232067 | 0.42047238 |
| ENSGALG00010017881 | KCNK10 | 428902 | 0.287351847 | 2.981259 | 0.0423576 | 0.42047238 |
| ENSGALG00010027861 | PNPLA7 | 427774 | 0.144427512 | 5.90570735 | 0.0423862 | 0.42047238 |
| ENSGALG00010028547 | ZNF618 | 770984 | -0.150497269 | 6.09860292 | 0.04239807 | 0.42047238 |
| ENSGALG00010005444 | ACTC1 | 423298 | -0.361323069 | 2.39977967 | 0.04240127 | 0.42047238 |
| ENSGALG00010017845 | NA | NA | 0.827686361 | -0.0118867 | 0.04250367 | 0.42120336 |
| ENSGALG00010018611 | NUDCD3 | 770002 | 0.18698868 | 5.41496274 | 0.04252997 | 0.42120336 |
| ENSGALG00010014390 | SPARC | 386571 | 0.182249588 | 7.81037171 | 0.04257172 | 0.42122693 |
| ENSGALG00010024934 | G6PC3 | 100859356 | 0.183272929 | 5.85284595 | 0.04258915 | 0.42122693 |
| ENSGALG00010024276 | NA | NA | -0.344098073 | 2.42673728 | 0.0426206 | 0.42122693 |
| ENSGALG00010011469 | CPS1 | 428994 | -0.223882973 | 3.89388762 | 0.04265701 | 0.42122693 |
| ENSGALG00010000730 | INTS5 | 101750875 | 0.169041949 | 5.57856297 | 0.04269401 | 0.42122693 |
| ENSGALG00010000742 | NA | NA | -0.265572376 | 4.86742627 | 0.04269731 | 0.42122693 |
| ENSGALG00010025243 | PAXIP1 | 420441 | -0.165965176 | 5.69059286 | 0.04276477 | 0.42137046 |
| ENSGALG00010003524 | NA | NA | -0.646966776 | 0.35990465 | 0.04279557 | 0.42137046 |
| ENSGALG00010003480 | NA | NA | 0.289725687 | 3.31520712 | 0.0428311 | 0.42137046 |
| ENSGALG00010021632 | LIMD2 | 419955 | 0.270081793 | 3.23395909 | 0.04284761 | 0.42137046 |
| ENSGALG00010027293 | ELP6 | 420379 | 0.307439998 | 2.9440688 | 0.04284937 | 0.42137046 |
| ENSGALG00010025904 | RPL10A | 419895 | 0.161472244 | 9.06002155 | 0.04288509 | 0.4214512 |
| ENSGALG00010009811 | PJA2 | 415612 | -0.157086149 | 6.1379665 | 0.04295069 | 0.4217259 |
| ENSGALG00010029269 | WSCD1 | 427842 | 0.349952641 | 3.11590473 | 0.04296809 | 0.4217259 |
| ENSGALG00010029029 | NA | NA | -0.27682535 | 3.59132154 | 0.04300429 | 0.42181101 |
| ENSGALG00010009570 | LOC101751687 | 101751687 | 0.15600307 | 5.88712788 | 0.04308581 | 0.42234016 |
| ENSGALG00010029615 | TMEM120A | 417519 | 0.257873793 | 3.94435077 | 0.04313797 | 0.4225471 |
| ENSGALG00010024472 | LDHA | 396221 | 0.285336876 | 8.8848283 | 0.04318634 | 0.4225471 |
| ENSGALG00010026066 | SELENOP2 | 424593 | 0.638481207 | 0.37015082 | 0.04318966 | 0.4225471 |
| ENSGALG00010017726 | TTI2 | 426923 | 0.290946496 | 3.27865984 | 0.04324685 | 0.42283665 |
| ENSGALG00010002912 | RPS3 | 419069 | 0.188814595 | 9.42255397 | 0.04327825 | 0.42287377 |
| ENSGALG00010001745 | CADPS2 | 417756 | -0.241206551 | 3.92756516 | 0.04332253 | 0.42303667 |
| ENSGALG00010005831 | SEC61A2 | 426369 | -0.169300894 | 6.58136728 | 0.04343671 | 0.42374424 |
| ENSGALG00010015422 | DGKK | 422129 | 0.388051322 | 2.63100082 | 0.04345031 | 0.42374424 |
| ENSGALG00010014793 | TLR2A | 374141 | 0.305543466 | 2.72842776 | 0.04352493 | 0.42411924 |
| ENSGALG00010026006 | ITGA2B | 101749783 | 0.331916741 | 4.06621746 | 0.04354413 | 0.42411924 |
| ENSGALG00010028552 | LOC769139 | 769139 | 0.398766428 | 1.81538559 | 0.04362592 | 0.42464595 |
| ENSGALG00010014880 | SFRP2 | 395546 | 0.198553029 | 5.19963572 | 0.04370472 | 0.42514286 |
| ENSGALG00010023496 | PLEKHA3 | 424128 | -0.163934384 | 5.91773802 | 0.04376984 | 0.42550614 |
| ENSGALG00010018448 | KLHDC3 | 421251 | 0.270372675 | 3.48046412 | 0.04387884 | 0.42596711 |
| ENSGALG00010027229 | MDM4 | 419823 | -0.171333384 | 5.61705925 | 0.04389154 | 0.42596711 |
| ENSGALG00010028116 | HYAL1 | 426739 | 0.349125124 | 2.9272325 | 0.04394979 | 0.42596711 |
| ENSGALG00010013433 | NA | NA | -0.416060301 | 2.21589677 | 0.04397873 | 0.42596711 |
| ENSGALG00010021261 | JUP | 429710 | 0.160595339 | 6.92101033 | 0.0439835 | 0.42596711 |
| ENSGALG00010020113 | CIAO1 | 426332 | 0.194052739 | 5.13636479 | 0.04402011 | 0.42596711 |
| ENSGALG00010024644 | EIF6 | 419140 | 0.141776565 | 7.82622246 | 0.04402248 | 0.42596711 |
| ENSGALG00010021972 | PISD | 416960 | -0.157075153 | 7.06173487 | 0.04405047 | 0.42596711 |
| ENSGALG00010022643 | SP3 | 395302 | -0.158869282 | 7.73549744 | 0.04406748 | 0.42596711 |
| ENSGALG00010007671 | FAM206A | 420983 | -0.24007237 | 3.49179232 | 0.04413061 | 0.42610982 |
| ENSGALG00010023382 | NA | NA | -0.864931189 | -0.5337966 | 0.04413787 | 0.42610982 |
| ENSGALG00010022433 | LNPK | 424141 | 0.170348375 | 5.12900411 | 0.04418254 | 0.42627245 |
| ENSGALG00010002589 | SLC4A7 | 420658 | -0.139498231 | 7.46142733 | 0.04421812 | 0.42634728 |

| | | | | | | |
|---|---|---|---|---|---|---|
| ENSGALG00010027524 | TOMM40L | 426278 | 0.183252403 | 6.19019666 | 0.04428793 | 0.42668112 |
| ENSGALG00010028585 | FAM63A | 100859849 | 0.171723797 | 6.4505231 | 0.04430844 | 0.42668112 |
| ENSGALG00010015594 | LOC112530791 | 112530791 | -0.799716749 | -0.3094955 | 0.0443434 | 0.42674947 |
| ENSGALG00010002226 | OXNAD1 | 420644 | 0.214973834 | 4.09155271 | 0.0446993 | 0.4295412 |
| ENSGALG00010006851 | TMEM167A | 770790 | -0.209184738 | 7.20061158 | 0.04470549 | 0.4295412 |
| ENSGALG00010024872 | RUNDC3A | 100858314 | 0.255879004 | 4.47810255 | 0.04471759 | 0.4295412 |
| ENSGALG00010026434 | PRICKLE4 | 100859351 | 0.380963119 | 2.04198908 | 0.04477154 | 0.4297899 |
| ENSGALG00010023791 | VTI1B | 423273 | 0.211036181 | 4.07431435 | 0.04482947 | 0.42987294 |
| ENSGALG00010027402 | AP4B1 | 419879 | 0.161435032 | 5.91909917 | 0.0448363 | 0.42987294 |
| ENSGALG00010022062 | MITF | 395886 | -0.286328339 | 3.35673094 | 0.04490489 | 0.43012771 |
| ENSGALG00010000584 | RBM14 | 426080 | 0.168225337 | 7.1117827 | 0.04491902 | 0.43012771 |
| ENSGALG00010023221 | AMY1A | 414139 | 0.14583556 | 7.53776872 | 0.04504036 | 0.43102022 |
| ENSGALG00010027231 | ATP2B2 | 415958 | -0.248286138 | 4.41360167 | 0.04509287 | 0.43125331 |
| ENSGALG00010021969 | CCDC103 | 419970 | 0.409708206 | 1.78169455 | 0.04520068 | 0.43185966 |
| ENSGALG00010029351 | SLC25A10 | 769048 | 0.212929259 | 5.42217642 | 0.04523585 | 0.43185966 |
| ENSGALG00010002536 | LOC121106913 | 121106913 | 0.26468775 | 3.19527008 | 0.04524083 | 0.43185966 |
| ENSGALG00010022073 | PTPMT1 | 423181 | 0.271159593 | 3.5787452 | 0.04527392 | 0.43190645 |
| ENSGALG00010015738 | CRACDL | 100858050 | -0.426526632 | 2.44975326 | 0.04538199 | 0.43266801 |
| ENSGALG00010002599 | KIFC1 | 417037 | 0.15191354 | 7.6409845 | 0.04541177 | 0.43268263 |
| ENSGALG00010014661 | NA | NA | 0.5054568 | 1.04349574 | 0.04549852 | 0.43274541 |
| ENSGALG00010007548 | HSBP1 | 415813 | 0.162117757 | 6.74577613 | 0.04550566 | 0.43274541 |
| ENSGALG00010029777 | RPH3AL | 772026 | 0.228146561 | 3.82525698 | 0.04552884 | 0.43274541 |
| ENSGALG00010003352 | FARSA | 100859604 | 0.163496067 | 6.10042985 | 0.04553134 | 0.43274541 |
| ENSGALG00010022038 | GAPDH | 374193 | 0.18861224 | 11.2061544 | 0.04563803 | 0.43319191 |
| ENSGALG00010001330 | NA | NA | 0.199317298 | 5.69174729 | 0.04568169 | 0.43319191 |
| ENSGALG00010029550 | CHMP6 | 428673 | 0.156206127 | 5.69562453 | 0.04569474 | 0.43319191 |
| ENSGALG00010014971 | SMC5 | 431608 | -0.180719153 | 7.36747336 | 0.04575347 | 0.43319191 |
| ENSGALG00010001268 | CBLN2 | 421019 | -0.189647609 | 4.55918507 | 0.0457573 | 0.43319191 |
| ENSGALG00010004798 | PTPRK | 421711 | -0.150599225 | 6.49437982 | 0.04575733 | 0.43319191 |
| ENSGALG00010026693 | NA | NA | -0.367072346 | 2.26607844 | 0.04577624 | 0.43319191 |
| ENSGALG00010019705 | NA | NA | -0.261280484 | 4.11778609 | 0.04581717 | 0.43331161 |
| ENSGALG00010022783 | STAG1 | 424735 | 0.163647207 | 5.06027792 | 0.04591115 | 0.4339326 |
| ENSGALG00010005079 | NA | NA | -0.48211914 | 0.99508846 | 0.04596248 | 0.43407604 |
| ENSGALG00010017327 | CLN6 | 415558 | 0.245415815 | 4.22508675 | 0.04598299 | 0.43407604 |
| ENSGALG00010013686 | UQCRQ | 416336 | 0.148382883 | 6.16414704 | 0.04603187 | 0.43426991 |
| ENSGALG00010022055 | E2F1 | 396142 | -0.201095534 | 4.88456105 | 0.04606627 | 0.43432693 |
| ENSGALG00010017358 | NA | NA | -0.812932122 | -0.39902 | 0.04610227 | 0.43434767 |
| ENSGALG00010000044 | MAP1A | 415588 | 0.149735996 | 6.65996558 | 0.04612517 | 0.43434767 |
| ENSGALG00010028425 | DAP3 | 425065 | 0.158916539 | 6.11966289 | 0.04636045 | 0.43614045 |
| ENSGALG00010021118 | USMG5 | 423873 | 0.179382597 | 5.14900441 | 0.04640139 | 0.43614045 |
| ENSGALG00010016800 | RPL14 | 374134 | 0.143287002 | 8.10842444 | 0.04640391 | 0.43614045 |
| ENSGALG00010023589 | NA | NA | -0.384423841 | 2.04318235 | 0.04646302 | 0.43614045 |
| ENSGALG00010009101 | HIKESHI | 427034 | 0.157913599 | 5.36382866 | 0.04648366 | 0.43614045 |
| ENSGALG00010022223 | DOK4 | 430610 | 0.227544222 | 3.75623642 | 0.04651324 | 0.43614045 |
| ENSGALG00010021789 | CHD9 | 415722 | -0.151935694 | 6.73247228 | 0.04651482 | 0.43614045 |
| ENSGALG00010014601 | NA | NA | -0.446333989 | 2.39361821 | 0.04668451 | 0.43697778 |
| ENSGALG00010004131 | NA | NA | -0.631844677 | 0.23531459 | 0.04669276 | 0.43697778 |
| ENSGALG00010004034 | NEDD8 | 100858776 | 0.181653508 | 5.74775588 | 0.04670709 | 0.43697778 |
| ENSGALG00010004477 | TULP4 | 421658 | -0.149349671 | 7.74057563 | 0.04675943 | 0.43697778 |
| ENSGALG00010028890 | LOC101747382 | 101747382 | 0.414212387 | 3.26614035 | 0.0468038 | 0.43697778 |
| ENSGALG00010004096 | COL12A1 | 395875 | -0.527886063 | 1.2104361 | 0.04681121 | 0.43697778 |
| ENSGALG00010001270 | LOC107050177 | 107050177 | 0.150636973 | 6.85591558 | 0.04682064 | 0.43697778 |
| ENSGALG00010007513 | GALNT12 | 420978 | -0.20279911 | 4.55209431 | 0.04684168 | 0.43697778 |

| | | | | | | |
|---|---|---|---|---|---|---|
| ENSGALG00010020912 | PDE8A | 415455 | -0.364634863 | 1.90404311 | 0.04686793 | 0.43697778 |
| ENSGALG00010000098 | NA | NA | 0.203477709 | 4.35111738 | 0.04689916 | 0.43697778 |
| ENSGALG00010021572 | MYL9 | 396215 | 0.20656486 | 5.19543698 | 0.04691786 | 0.43697778 |
| ENSGALG00010028763 | CTSK | 395818 | 0.570114336 | 1.05069315 | 0.0469508 | 0.43701892 |
| ENSGALG00010010164 | KIF28P | 101748750 | -0.300351822 | 3.13332169 | 0.04700998 | 0.4373041 |
| ENSGALG00010026081 | KPNA1 | 418271 | -0.138131393 | 6.31710814 | 0.04707076 | 0.43760384 |
| ENSGALG00010002423 | LOC121106436 | 121106436 | 0.841208619 | -0.4761118 | 0.04711289 | 0.43772989 |
| ENSGALG00010021238 | NDUFS8 | 769492 | 0.188021368 | 5.43019257 | 0.04716547 | 0.43795284 |
| ENSGALG00010014344 | LSG1 | 424897 | 0.138567519 | 6.65960022 | 0.04724945 | 0.43813728 |
| ENSGALG00010012047 | LMAN2 | 100859676 | 0.151167465 | 6.88366574 | 0.04725764 | 0.43813728 |
| ENSGALG00010005292 | TAF3 | 419107 | -0.153503318 | 6.98022332 | 0.04727577 | 0.43813728 |
| ENSGALG00010028131 | TMEM161A | 100857264 | 0.201287333 | 4.91346483 | 0.0473106 | 0.43813728 |
| ENSGALG00010005558 | TAGAP | 421662 | -0.959608016 | -0.4176038 | 0.04733205 | 0.43813728 |
| ENSGALG00010029127 | LOC112530003 | 112530003 | 0.517531998 | 0.77417989 | 0.04737635 | 0.43813728 |
| ENSGALG00010028103 | APBA3 | 420069 | 0.187750457 | 4.91700003 | 0.04738551 | 0.43813728 |
| ENSGALG00010001653 | NA | NA | -0.56584173 | 0.54507127 | 0.04742762 | 0.43826208 |
| ENSGALG00010013435 | SDC3 | 396343 | 0.157349783 | 6.2970622 | 0.04753165 | 0.43895869 |
| ENSGALG00010019223 | AAMP | 769880 | 0.151928405 | 5.64789085 | 0.04767728 | 0.4400383 |
| ENSGALG00010025389 | SNED1 | 424846 | 0.182520931 | 4.65070904 | 0.04770884 | 0.44006447 |
| ENSGALG00010000055 | PRC1 | 770336 | 0.15746891 | 7.01765426 | 0.04783633 | 0.44067183 |
| ENSGALG00010020420 | NA | NA | 0.886263774 | -0.2928995 | 0.04785232 | 0.44067183 |
| ENSGALG00010029338 | INPP5K | 771549 | -0.145373522 | 6.45888649 | 0.04786097 | 0.44067183 |
| ENSGALG00010019578 | NA | NA | -0.818047675 | -0.2087849 | 0.04793252 | 0.44106553 |
| ENSGALG00010015095 | RBFOX2 | 418055 | 0.139305455 | 6.97236256 | 0.04800011 | 0.44142238 |
| ENSGALG00010017798 | BAG4 | 426776 | 0.249049754 | 3.30993669 | 0.04805369 | 0.44165004 |
| ENSGALG00010017627 | NA | NA | -0.796021408 | -0.3096456 | 0.0481916 | 0.44253651 |
| ENSGALG00010020786 | DDX19 | 426504 | 0.179734542 | 5.58599282 | 0.04821843 | 0.44253651 |
| ENSGALG00010014091 | CHIC2 | 422751 | -0.17357796 | 5.73216401 | 0.0482368 | 0.44253651 |
| ENSGALG00010010136 | EPHX2 | 421999 | 0.248156493 | 4.19115168 | 0.04832197 | 0.44274957 |
| ENSGALG00010010859 | LOC426385 | 426385 | 0.166740838 | 5.48376581 | 0.04833451 | 0.44274957 |
| ENSGALG00010027951 | FAM32A | 420150 | 0.158616021 | 5.56691425 | 0.04834672 | 0.44274957 |
| ENSGALG00010027107 | KIF21B | 421178 | 0.154035044 | 5.7843886 | 0.04840066 | 0.44297883 |
| ENSGALG00010003081 | DOPEY2 | 421842 | -0.146841058 | 5.88186009 | 0.04860081 | 0.44389095 |
| ENSGALG00010019785 | ACSL5 | 423896 | -0.787074024 | -0.3628646 | 0.04862635 | 0.44389095 |
| ENSGALG00010013263 | ATP11A | 418749 | -0.24497248 | 3.69992491 | 0.04864852 | 0.44389095 |
| ENSGALG00010027215 | NA | NA | 0.273060931 | 3.70049843 | 0.04866098 | 0.44389095 |
| ENSGALG00010007134 | NA | NA | -0.334950017 | 2.29937579 | 0.04868624 | 0.44389095 |
| ENSGALG00010025072 | VAMP2 | 424827 | 0.695246659 | -0.1301711 | 0.04871424 | 0.44389095 |
| ENSGALG00010017093 | NA | NA | 0.733338333 | -0.0692268 | 0.04874884 | 0.44389095 |
| ENSGALG00010024679 | MSL1 | 777194 | 0.15691643 | 7.18646604 | 0.0487532 | 0.44389095 |
| ENSGALG00010019398 | SLC35F5 | 424281 | -0.191461207 | 5.16390587 | 0.04876108 | 0.44389095 |
| ENSGALG00010012421 | LOC768614 | 768614 | -0.803796125 | -0.4556837 | 0.0488181 | 0.44414613 |
| ENSGALG00010000811 | LOC770405 | 770405 | 0.511392278 | 0.87035807 | 0.04890093 | 0.44443719 |
| ENSGALG00010027118 | ABHD6 | 416009 | 0.368606144 | 2.55982031 | 0.04890811 | 0.44443719 |
| ENSGALG00010015569 | NA | NA | -0.601459995 | 0.46554746 | 0.04895613 | 0.44460989 |
| ENSGALG00010026242 | CCDC141 | 770766 | 0.241932989 | 3.5147182 | 0.0490332 | 0.44478543 |
| ENSGALG00010021426 | SLC38A7 | 107052162 | 0.192889776 | 5.46861869 | 0.04908671 | 0.44478543 |
| ENSGALG00010019271 | CDKL1 | 423575 | 0.416484743 | 1.53956876 | 0.04910883 | 0.44478543 |
| ENSGALG00010009330 | EIF3K | 101750374 | 0.15644887 | 8.1521973 | 0.04911698 | 0.44478543 |
| ENSGALG00010021492 | PARP6 | 769472 | 0.235446566 | 4.41773077 | 0.04912062 | 0.44478543 |
| ENSGALG00010005441 | UBE2V2 | 421109 | -0.166134793 | 7.01405552 | 0.04920842 | 0.44510205 |
| ENSGALG00010027233 | NA | NA | 0.237317851 | 3.50539997 | 0.04926756 | 0.44510205 |
| ENSGALG00010026333 | NA | NA | -0.681515835 | 0.07031569 | 0.04931539 | 0.44510205 |

| | | | | | | |
|---|---|---|---|---|---|---|
| ENSGALG00010019822 | NA | NA | -0.73538416 | -0.2086363 | 0.04932025 | 0.44510205 |
| ENSGALG00010004385 | ADAMTS3 | 428760 | -0.290089113 | 3.42764743 | 0.04932745 | 0.44510205 |
| ENSGALG00010022673 | KLHL42 | 419046 | 0.217125239 | 3.77996128 | 0.04937119 | 0.44510205 |
| ENSGALG00010028939 | NA | NA | 0.169860853 | 5.66035242 | 0.04940458 | 0.44510205 |
| ENSGALG00010016825 | CD164L2 | 101748332 | 0.346635807 | 2.19374216 | 0.04941686 | 0.44510205 |
| ENSGALG00010027427 | UBAC1 | 427754 | 0.168581924 | 7.04206439 | 0.04941705 | 0.44510205 |
| ENSGALG00010016603 | LOC422901 | 422901 | -0.165817739 | 6.19293594 | 0.04962143 | 0.44577465 |
| ENSGALG00010002728 | RP2 | 418675 | -0.200013478 | 4.39008134 | 0.0496503 | 0.44577465 |
| ENSGALG00010022450 | NA | NA | -0.544290538 | 0.69118623 | 0.04968996 | 0.44577465 |
| ENSGALG00010012653 | CBX7 | 101752246 | -0.219002837 | 3.90774108 | 0.04976567 | 0.44577465 |
| ENSGALG00010011199 | LRRIQ1 | 417882 | 0.374335004 | 2.35923788 | 0.04977513 | 0.44577465 |
| ENSGALG00010001386 | FKBP14 | 420636 | -0.151198553 | 7.70895338 | 0.04977711 | 0.44577465 |
| ENSGALG00010001725 | MTA2 | 101747888 | 0.153753302 | 7.96800017 | 0.04978503 | 0.44577465 |
| ENSGALG00010008123 | LOC121112980 | 121112980 | 0.168975923 | 5.65090894 | 0.04979244 | 0.44577465 |
| ENSGALG00010024222 | KCNK15 | 419191 | 0.417614443 | 1.86229178 | 0.04984458 | 0.44577465 |
| ENSGALG00010017859 | EXOC6 | 423807 | -0.171422175 | 4.85165026 | 0.04984871 | 0.44577465 |
| ENSGALG00010013285 | ATP2B1 | 374244 | -0.158478066 | 9.73411675 | 0.04985495 | 0.44577465 |
| ENSGALG00010020023 | FOSL2 | 421416 | -0.205944804 | 4.59728561 | 0.04986584 | 0.44577465 |
| ENSGALG00010016773 | RPS27A | 395796 | 0.154134925 | 8.79285804 | 0.04990133 | 0.44577465 |
| ENSGALG00010025727 | GLI1 | 396045 | 0.297586651 | 3.44434248 | 0.04991055 | 0.44577465 |
| ENSGALG00010015806 | NA | NA | -0.804301632 | -0.4554987 | 0.04997215 | 0.44577465 |
| ENSGALG00010019789 | ARNT2 | 415481 | 0.154882522 | 6.93288281 | 0.04997236 | 0.44577465 |
| ENSGALG00010023086 | PCSK7 | 395455 | 0.200772292 | 4.20959295 | 0.04998635 | 0.44577465 |

Supplementally table S8. DEGs comparing sgSoprano#2 and sgOVA in CRISPRi PGCs (p-adj < 0.05)

| Ensembl_ID | Symbol | entrez_ID | logFC | logCPM | p-adj_value | FDR |
|---|---|---|---|---|---|---|
| ENSGALG00010018908 | NA | NA | -1.950726546 | 6.90855291 | 1.12E-23 | 1.72E-19 |
| ENSGALG00010013595 | WEE2 | 427918 | -1.219627419 | 5.922567298 | 4.92E-20 | 3.78E-16 |
| ENSGALG00010007287 | LOC124418405 | 124418405 | -0.965771627 | 9.395789125 | 6.17E-18 | 3.16E-14 |
| ENSGALG00010004371 | NA | NA | -0.65115379 | 9.111981987 | 2.31E-17 | 7.59E-14 |
| ENSGALG00010011693 | NA | NA | -0.884315124 | 5.343643777 | 2.47E-17 | 7.59E-14 |
| ENSGALG00010006797 | NA | NA | -1.290810178 | 5.54818629 | 8.41E-17 | 2.15E-13 |
| ENSGALG00010001989 | NA | NA | -0.962510422 | 7.632941115 | 9.03E-16 | 1.98E-12 |
| ENSGALG00010007105 | NA | NA | -1.355698457 | 5.154967767 | 3.52E-15 | 6.76E-12 |
| ENSGALG00010002503 | NA | NA | -0.949517329 | 8.608837271 | 3.58E-14 | 6.11E-11 |
| ENSGALG00010006859 | NA | NA | -1.008206784 | 3.464165979 | 4.71E-14 | 7.24E-11 |
| ENSGALG00010018244 | CHM | 422266 | 0.528374505 | 5.832871247 | 1.33E-13 | 1.85E-10 |
| ENSGALG00010009553 | NA | NA | -1.066681327 | 7.139067283 | 3.91E-13 | 5.00E-10 |
| ENSGALG00010014283 | NA | NA | -1.34728179 | 2.880648249 | 1.02E-12 | 1.20E-09 |
| ENSGALG00010013729 | NA | NA | -1.087127043 | 3.43166642 | 5.98E-11 | 6.56E-08 |
| ENSGALG00010002195 | NA | NA | -1.018078628 | 4.484459324 | 9.34E-11 | 9.57E-08 |
| ENSGALG00010017060 | CLSTN1 | 395372 | 0.478741278 | 8.266644051 | 2.61E-10 | 2.51E-07 |
| ENSGALG00010004208 | NA | NA | -1.349785698 | 1.882672316 | 3.77E-10 | 3.41E-07 |
| ENSGALG00010010905 | ENC1 | 427203 | 0.465769799 | 7.259087732 | 5.63E-10 | 4.81E-07 |
| ENSGALG00010005001 | GUCA1C | 427960 | -0.864383911 | 5.524030724 | 9.93E-10 | 8.03E-07 |
| ENSGALG00010006798 | NA | NA | -1.057039394 | 4.196701607 | 1.54E-09 | 1.18E-06 |
| ENSGALG00010013561 | AGK | 418121 | -0.560033034 | 7.876752316 | 7.06E-09 | 5.17E-06 |
| ENSGALG00010019204 | ARF6 | 428927 | 0.460281362 | 7.220676099 | 7.49E-09 | 5.23E-06 |
| ENSGALG00010026334 | CALCOCO2 | 419993 | 0.454449072 | 5.316978639 | 8.59E-09 | 5.74E-06 |
| ENSGALG00010006916 | NA | NA | -1.101450468 | 3.50098676 | 1.88E-08 | 1.20E-05 |
| ENSGALG00010013572 | DENND11 | 427917 | -0.722693267 | 6.674033998 | 2.13E-08 | 1.31E-05 |
| ENSGALG00010011694 | NA | NA | -1.065136923 | 1.997509239 | 2.68E-08 | 1.59E-05 |
| ENSGALG00010007167 | NA | NA | -0.991687735 | 4.550722695 | 3.80E-08 | 2.16E-05 |
| ENSGALG00010015173 | ENS-1 | 769415 | -1.169384506 | 10.72187043 | 3.98E-08 | 2.18E-05 |
| ENSGALG00010008262 | CYB5B | 415865 | 0.453146039 | 6.124599807 | 4.42E-08 | 2.34E-05 |
| ENSGALG00010002022 | ERNI | 769673 | -1.000576995 | 10.04906694 | 4.73E-08 | 2.42E-05 |
| ENSGALG00010023092 | NA | NA | -1.348115111 | 3.095050112 | 5.10E-08 | 2.53E-05 |
| ENSGALG00010012578 | NDUFS4 | 374122 | 0.451864529 | 6.523686416 | 7.30E-08 | 3.51E-05 |
| ENSGALG00010008642 | NA | NA | -3.048508934 | -1.029650195 | 8.02E-08 | 3.73E-05 |
| ENSGALG00010016645 | NA | NA | -1.168202947 | 7.580318808 | 8.70E-08 | 3.78E-05 |
| ENSGALG00010006179 | NA | NA | -1.188798634 | 6.041121544 | 8.76E-08 | 3.78E-05 |
| ENSGALG00010025365 | SIKE1 | 419887 | 0.432099543 | 6.682549524 | 8.85E-08 | 3.78E-05 |
| ENSGALG00010004527 | NA | NA | -0.929508835 | 4.596873544 | 9.92E-08 | 4.12E-05 |
| ENSGALG00010003982 | GLCCI1 | 420580 | 0.392440839 | 7.680872443 | 1.14E-07 | 4.61E-05 |
| ENSGALG00010004570 | LOC107051826 | 107051826 | -1.010786766 | 10.59129364 | 1.77E-07 | 6.97E-05 |
| ENSGALG00010003570 | LOC121107016 | 121107016 | -1.485188234 | 2.046756243 | 1.98E-07 | 7.62E-05 |
| ENSGALG00010024824 | DNAJC5 | 419256 | 0.508092463 | 4.505466611 | 2.05E-07 | 7.68E-05 |
| ENSGALG00010000554 | LOC121108774 | 121108774 | -0.882615999 | 2.897383278 | 2.64E-07 | 9.67E-05 |
| ENSGALG00010007161 | NA | NA | -0.885095572 | 4.233662423 | 6.07E-07 | 0.000215056 |
| ENSGALG00010007049 | NA | NA | -1.076086148 | 9.214219818 | 6.16E-07 | 0.000215056 |
| ENSGALG00010011997 | DENND5B | 418139 | 0.469810486 | 4.928261748 | 7.26E-07 | 0.000248001 |
| ENSGALG00010009771 | NA | NA | -0.865097219 | 3.189767041 | 9.44E-07 | 0.000310862 |
| ENSGALG00010006214 | NA | NA | -1.065664756 | 7.719161657 | 9.51E-07 | 0.000310862 |
| ENSGALG00010014226 | SLC16A7 | 417815 | -0.675701806 | 3.165806096 | 1.02E-06 | 0.000325023 |
| ENSGALG00010023095 | NA | NA | -1.174405329 | 2.794806079 | 1.11E-06 | 0.000348568 |
| ENSGALG00010028355 | RBM18 | 417111 | 0.379265542 | 5.773305757 | 1.33E-06 | 0.000407419 |
| ENSGALG00010005396 | MICU2 | 418946 | 0.388575994 | 5.039457514 | 2.13E-06 | 0.000643013 |
| ENSGALG00010008982 | NA | NA | -1.126731665 | 1.53094995 | 2.25E-06 | 0.000666262 |
| ENSGALG00010009327 | NID1 | 395531 | 0.375814386 | 7.339167555 | 2.37E-06 | 0.000688607 |
| ENSGALG00010024154 | VPS29L | 416931 | 0.517213463 | 4.429961153 | 2.98E-06 | 0.000847054 |
| ENSGALG00010007504 | TMPRSS15 | 427967 | -1.405791839 | 1.065613345 | 3.03E-06 | 0.000847054 |
| ENSGALG00010021861 | NA | NA | 0.340844928 | 5.622145574 | 3.60E-06 | 0.000987503 |

| | | | | | | |
|---|---|---|---|---|---|---|
| ENSGALG00010015392 | TMEM165 | 428780 | 0.337188796 | 6.218588009 | 4.50E-06 | 0.001212467 |
| ENSGALG00010009863 | NA | NA | -1.505955429 | 0.341568083 | 5.79E-06 | 0.001535557 |
| ENSGALG00010001828 | LOC100858869 | 100858869 | -1.273375101 | 7.647833236 | 6.17E-06 | 0.001606634 |
| ENSGALG00010006059 | BIVM | 374099 | 0.330761284 | 6.843365881 | 6.39E-06 | 0.001635772 |
| ENSGALG00010009859 | NA | NA | -1.384143632 | 2.408831141 | 6.74E-06 | 0.001698359 |
| ENSGALG00010005493 | KLF6 | 420463 | -0.351237571 | 5.338073749 | 7.46E-06 | 0.001846602 |
| ENSGALG00010006086 | LOC100857117 | 100857117 | -1.036014077 | 9.193474462 | 7.57E-06 | 0.001846602 |
| ENSGALG00010001138 | SPTSSB | 425008 | 0.364965625 | 5.313343389 | 9.95E-06 | 0.002389459 |
| ENSGALG00010002078 | NA | NA | -1.750989633 | -0.331695443 | 1.15E-05 | 0.002730799 |
| ENSGALG00010007115 | CD69L | 417816 | -0.409496454 | 5.215136262 | 1.21E-05 | 0.002816564 |
| ENSGALG00010013772 | KIAA0319L | 419634 | 0.318283057 | 5.983502624 | 1.23E-05 | 0.002820268 |
| ENSGALG00010006902 | NA | NA | -1.136892207 | 2.400418847 | 1.29E-05 | 0.002924728 |
| ENSGALG00010018537 | LOC112530649 | 112530649 | -1.166823063 | 8.829559765 | 1.40E-05 | 0.00311968 |
| ENSGALG00010014282 | NA | NA | -1.488935318 | 1.392139221 | 1.58E-05 | 0.003464271 |
| ENSGALG00010003639 | NA | NA | -1.203661518 | 9.091325378 | 1.66E-05 | 0.003594691 |
| ENSGALG00010017976 | GPR137C | 428932 | -0.440865033 | 4.881259351 | 1.69E-05 | 0.00361589 |
| ENSGALG00010006266 | CENPW | 421716 | 0.38527331 | 6.721747564 | 2.02E-05 | 0.004247083 |
| ENSGALG00010019767 | SUCLG2 | 416087 | 0.331558925 | 6.950964553 | 2.31E-05 | 0.004803511 |
| ENSGALG00010002060 | NA | NA | -0.609831065 | 2.909385295 | 2.35E-05 | 0.004814326 |
| ENSGALG00010001418 | RNMT | 421046 | 0.350272456 | 7.413409346 | 2.66E-05 | 0.005374436 |
| ENSGALG00010012538 | RAP1B | 417840 | 0.310391988 | 8.439240638 | 2.88E-05 | 0.005758567 |
| ENSGALG00010013385 | PGRMC1 | 772196 | 0.345506255 | 5.882351372 | 3.31E-05 | 0.006514329 |
| ENSGALG00010019752 | GPAM | 423895 | 0.321510525 | 7.236046117 | 3.49E-05 | 0.006787026 |
| ENSGALG00010010296 | NA | NA | -0.481472393 | 4.030964862 | 3.84E-05 | 0.007362504 |
| ENSGALG00010012294 | LOC100857299 | 100857299 | -0.896491528 | 7.790026268 | 3.88E-05 | 0.007362504 |
| ENSGALG00010014169 | NA | NA | -1.368868327 | 0.584726916 | 4.58E-05 | 0.00857746 |
| ENSGALG00010015483 | SGCD | 416251 | -0.320851974 | 5.390488954 | 4.90E-05 | 0.009067781 |
| ENSGALG00010006623 | LOC107052071 | 107052071 | -0.934120382 | 1.872381522 | 5.19E-05 | 0.009499735 |
| ENSGALG00010007171 | NA | NA | -1.075061526 | 4.677266366 | 5.33E-05 | 0.009632069 |
| ENSGALG00010005008 | LOC418414 | 418414 | -1.640495327 | 0.657356754 | 6.15E-05 | 0.010989401 |
| ENSGALG00010021837 | NA | NA | -1.639287321 | 0.236802548 | 6.94E-05 | 0.012264731 |
| ENSGALG00010006910 | NA | NA | -1.167736364 | 0.886709555 | 7.69E-05 | 0.013381047 |
| ENSGALG00010001483 | LOC121110182 | 121110182 | -1.00101135 | 5.248739686 | 7.75E-05 | 0.013381047 |
| ENSGALG00010003307 | COL1A2 | 396243 | 0.429802663 | 7.559544626 | 8.64E-05 | 0.014644984 |
| ENSGALG00010017063 | TMPPE | 101750979 | 0.355793022 | 5.025903829 | 8.67E-05 | 0.014644984 |
| ENSGALG00010025449 | LOC121108098 | 121108098 | 0.742602364 | 1.917416714 | 9.19E-05 | 0.015357659 |
| ENSGALG00010008968 | NA | NA | -1.067203549 | 7.553615223 | 9.80E-05 | 0.016188706 |
| ENSGALG00010001116 | RNF13 | 396303 | 0.289566165 | 7.812148658 | 0.000100541 | 0.016439444 |
| ENSGALG00010009928 | NA | NA | -1.16634301 | 5.617755432 | 0.00010965 | 0.017740221 |
| ENSGALG00010004159 | NA | NA | -1.255112364 | 2.465477459 | 0.000116177 | 0.018600429 |
| ENSGALG00010006087 | LOC107051465 | 107051465 | -0.909121627 | 9.778210997 | 0.000119018 | 0.018858856 |
| ENSGALG00010019724 | HES5 | 419392 | -1.171572918 | 0.35653571 | 0.000141931 | 0.022259965 |
| ENSGALG00010012301 | NA | NA | -1.51532493 | 2.952449137 | 0.000143708 | 0.02231104 |
| ENSGALG00010007963 | LDAH | 421967 | 0.293906828 | 5.795841294 | 0.000191733 | 0.029469312 |
| ENSGALG00010023231 | ASIC1 | 426883 | 0.530633084 | 3.177182396 | 0.000195385 | 0.029733325 |
| ENSGALG00010015239 | NA | NA | 0.302353637 | 5.603731664 | 0.000200415 | 0.030199845 |
| ENSGALG00010014931 | RNF145 | 416239 | -0.256717041 | 6.081021972 | 0.00020856 | 0.03112206 |
| ENSGALG00010028040 | TDRKH | 425667 | 0.417589474 | 4.563314077 | 0.000217775 | 0.032184635 |
| ENSGALG00010028240 | NA | NA | -0.901041491 | 1.248414021 | 0.000220902 | 0.032335808 |
| ENSGALG00010027092 | FAM72A | 419841 | 0.292090551 | 5.918117252 | 0.000240001 | 0.034800109 |
| ENSGALG00010022478 | ATP6V1A | 395821 | 0.276182153 | 8.60053235 | 0.000243653 | 0.034999474 |
| ENSGALG00010007985 | APELA | 770154 | 0.275393045 | 6.557458233 | 0.000263238 | 0.037154346 |
| ENSGALG00010029610 | ERN1 | 417414 | -0.294146685 | 5.365074814 | 0.000263489 | 0.037154346 |
| ENSGALG00010021387 | SLC2A14 | 396517 | 0.309460405 | 10.20197436 | 0.000266143 | 0.037187388 |
| ENSGALG00010008650 | NA | NA | -0.991090703 | 4.155281726 | 0.000287481 | 0.039807031 |
| ENSGALG00010012753 | GTPBP1 | 418022 | -0.272944965 | 6.701594477 | 0.000319456 | 0.043839658 |
| ENSGALG00010022059 | NPTX2 | 416438 | 0.37972728 | 4.412714492 | 0.000333394 | 0.045347465 |

| | | | | | | |
|---|---|---|---|---|---|---|
| ENSGALG00010021070 | RNF141 | 423039 | 0.318931378 | 5.016339926 | 0.000342687 | 0.045876007 |
| ENSGALG00010017667 | SLC2A1 | 396130 | 0.312725929 | 5.20901383 | 0.000343504 | 0.045876007 |
| ENSGALG00010023499 | PEBP1 | 416990 | -0.331768694 | 7.732600935 | 0.000346234 | 0.045876007 |
| ENSGALG00010029216 | RAB34 | 100858849 | 0.36915611 | 4.201040898 | 0.000361884 | 0.047539857 |
| ENSGALG00010001209 | CDH7 | 374007 | -0.416206261 | 4.126060015 | 0.000365787 | 0.047645254 |
| ENSGALG00010017647 | ITGA9 | 420757 | 0.491901762 | 3.536138183 | 0.000376224 | 0.048593002 |
| ENSGALG00010018834 | SAMD4A | 423559 | 0.334364227 | 5.024080475 | 0.000417574 | 0.053484314 |
| ENSGALG00010023546 | SLC11A2 | 751817 | 0.316822167 | 7.030642833 | 0.000430525 | 0.054687311 |
| ENSGALG00010028013 | CBARP | 100858130 | 0.521855667 | 3.054098787 | 0.000442248 | 0.055716032 |
| ENSGALG00010021233 | NA | NA | 0.263617759 | 5.950315656 | 0.000448201 | 0.056006921 |
| ENSGALG00010017368 | ERLIN2 | 426769 | 0.301929387 | 8.842381583 | 0.000459245 | 0.056924149 |
| ENSGALG00010015806 | NA | NA | -1.612927967 | -0.781019876 | 0.000482333 | 0.059307604 |
| ENSGALG00010001416 | AP4M1 | 396046 | 0.34665316 | 4.275951119 | 0.000489326 | 0.059690049 |
| ENSGALG00010010720 | NA | NA | -0.670306989 | 1.983565791 | 0.00051459 | 0.062277496 |
| ENSGALG00010013168 | DSCR3 | 418519 | 0.307505081 | 5.114837867 | 0.000525516 | 0.063102951 |
| ENSGALG00010019480 | NA | NA | -0.271670633 | 5.46263191 | 0.000538547 | 0.064166368 |
| ENSGALG00010023063 | KCTD5 | 416589 | 0.262012987 | 6.794332178 | 0.000546934 | 0.064664452 |
| ENSGALG00010029424 | MAP2K6 | 417445 | 0.524198735 | 3.496255451 | 0.000604043 | 0.070871306 |
| ENSGALG00010002837 | DNMT1 | 396011 | 0.256695936 | 7.649106365 | 0.00061328 | 0.071409904 |
| ENSGALG00010013214 | RAI14 | 427431 | -0.287479172 | 5.101914152 | 0.000772905 | 0.089319881 |
| ENSGALG00010005830 | USP12 | 428078 | 0.325629608 | 4.923704471 | 0.00080636 | 0.09249071 |
| ENSGALG00010008028 | KCNK7 | 421519 | 0.302255221 | 6.795772822 | 0.000831454 | 0.094662587 |
| ENSGALG00010011371 | LOC424430 | 424430 | 0.385030781 | 3.786086471 | 0.000843909 | 0.095374149 |
| ENSGALG00010002947 | LOC121107416 | 121107416 | -0.690883671 | 10.64644311 | 0.000866336 | 0.096596155 |
| ENSGALG00010014122 | ABCA1 | 373945 | 0.240319668 | 6.108296677 | 0.000867291 | 0.096596155 |
| ENSGALG00010010189 | NA | NA | -0.829810996 | 3.718481044 | 0.000942761 | 0.104246301 |
| ENSGALG00010023317 | RAPGEF3 | 426888 | 0.786135764 | 3.58786559 | 0.001007404 | 0.110598572 |
| ENSGALG00010027034 | NA | NA | -1.002197671 | 0.417995834 | 0.001015288 | 0.110673592 |
| ENSGALG00010009801 | LOC101750251 | 101750251 | -0.65634203 | 9.191495025 | 0.001053559 | 0.114036622 |
| ENSGALG00010006800 | NA | NA | -1.143388572 | -0.055819016 | 0.001062102 | 0.114157446 |
| ENSGALG00010004370 | ATP6V1H | 426199 | 0.246579705 | 5.978409498 | 0.00108888 | 0.116222845 |
| ENSGALG00010006382 | PNN | 423338 | 0.256069102 | 7.892919949 | 0.00110243 | 0.116626533 |
| ENSGALG00010002795 | NA | NA | -0.435789242 | 3.19168025 | 0.001124187 | 0.116626533 |
| ENSGALG00010006079 | LOC107055200 | 107055200 | -1.081889689 | 7.555895136 | 0.001124889 | 0.116626533 |
| ENSGALG00010029073 | NA | NA | -1.074125978 | -0.069583088 | 0.001126853 | 0.116626533 |
| ENSGALG00010011825 | FAM19A2 | 771745 | -0.288033707 | 5.541940804 | 0.001130602 | 0.116626533 |
| ENSGALG00010013164 | FAM83F | 770747 | 0.227278706 | 6.013549376 | 0.001158671 | 0.118725138 |
| ENSGALG00010027718 | LOC100857333 | 100857333 | -0.79018935 | 0.875859653 | 0.001186717 | 0.120742335 |
| ENSGALG00010000170 | LOC107051372 | 107051372 | -0.358318684 | 4.810911539 | 0.001194069 | 0.120742335 |
| ENSGALG00010020126 | NADK | 419403 | -0.249839937 | 5.814061661 | 0.001214051 | 0.12196059 |
| ENSGALG00010009470 | ZNF469 | 101748749 | -0.4191194 | 3.687170175 | 0.001228887 | 0.122148693 |
| ENSGALG00010015503 | NA | NA | -1.161846114 | -0.279259065 | 0.001231818 | 0.122148693 |
| ENSGALG00010012245 | RNF151 | 420226 | 0.228105482 | 5.906073336 | 0.001250909 | 0.12324666 |
| ENSGALG00010024657 | DPY19L3 | 768782 | 0.300831215 | 5.712729856 | 0.001317223 | 0.128953628 |
| ENSGALG00010016361 | NELFCD | 419312 | 0.220972122 | 6.072655214 | 0.001367368 | 0.133015436 |
| ENSGALG00010013243 | NA | NA | -0.786320599 | 2.722323652 | 0.001376906 | 0.133100871 |
| ENSGALG00010003358 | DNAJC15 | 418838 | -0.253517629 | 5.781348973 | 0.001401061 | 0.134589386 |
| ENSGALG00010029556 | NPTX1 | 422078 | 0.429222541 | 3.546207881 | 0.00147201 | 0.139864857 |
| ENSGALG00010009039 | SLC36A4 | 419007 | 0.348987566 | 4.15083592 | 0.001474177 | 0.139864857 |
| ENSGALG00010008093 | EFR3A | 420327 | 0.251531018 | 5.579263309 | 0.001528932 | 0.144169867 |
| ENSGALG00010007548 | HSBP1 | 415813 | 0.268491676 | 6.800447883 | 0.001550048 | 0.145269736 |
| ENSGALG00010021804 | NA | NA | -0.48676274 | 4.810157541 | 0.001631044 | 0.151798915 |
| ENSGALG00010017563 | ALK | 421297 | -0.378651752 | 3.838285353 | 0.001644062 | 0.151798915 |
| ENSGALG00010020691 | BOLA3 | 107054980 | 0.264271478 | 6.523204447 | 0.001649344 | 0.151798915 |
| ENSGALG00010026881 | LOC121111826 | 121111826 | -0.238907651 | 6.083844115 | 0.001689675 | 0.153792187 |
| ENSGALG00010004838 | NA | NA | -1.3865527 | -0.268117144 | 0.001691014 | 0.153792187 |
| ENSGALG00010002034 | WWC3 | 418649 | 0.233248651 | 6.545515995 | 0.001724669 | 0.155057046 |

| | | | | | | |
|---|---|---|---|---|---|---|
| ENSGALG00010001581 | NA | NA | -0.672395982 | 1.32303563 | 0.001725098 | 0.155057046 |
| ENSGALG00010024263 | PHKB | 415741 | 0.257796265 | 5.083946898 | 0.00174711 | 0.156122587 |
| ENSGALG00010020009 | MACROD2 | 421259 | 0.408235272 | 3.856054797 | 0.001790872 | 0.159108123 |
| ENSGALG00010011298 | NA | NA | -0.695512736 | 1.855277927 | 0.001883381 | 0.166365331 |
| ENSGALG00010018502 | ARHGEF10L | 419360 | 0.382051654 | 4.03512059 | 0.002015796 | 0.175800517 |
| ENSGALG00010019282 | IGFBP2 | 396315 | 0.299403396 | 5.737149001 | 0.002016286 | 0.175800517 |
| ENSGALG00010017866 | RALA | 420765 | 0.220953766 | 6.502621123 | 0.002024508 | 0.175800517 |
| ENSGALG00010013727 | NA | NA | -1.287843047 | -0.34587786 | 0.002051031 | 0.177103039 |
| ENSGALG00010019073 | LOC101750242 | 101750242 | 1.193544059 | -0.247322402 | 0.002080859 | 0.178674844 |
| ENSGALG00010027169 | ABCB8 | 107055879 | 0.316815792 | 4.913530345 | 0.002100559 | 0.179364377 |
| ENSGALG00010018386 | NA | NA | -0.869114123 | 0.665260553 | 0.002160858 | 0.183493823 |
| ENSGALG00010016706 | RTN4 | 378790 | 0.238571104 | 8.220752405 | 0.002232772 | 0.188558816 |
| ENSGALG00010022165 | FAM192A | 415647 | 0.274001893 | 6.376280614 | 0.002245094 | 0.188563354 |
| ENSGALG00010001629 | NA | NA | -0.538558452 | 3.268492931 | 0.002304972 | 0.191625716 |
| ENSGALG00010014656 | NA | NA | -0.471230604 | 3.138147982 | 0.002312949 | 0.191625716 |
| ENSGALG00010005382 | RAB32 | 421616 | 0.227117529 | 5.542104094 | 0.002318958 | 0.191625716 |
| ENSGALG00010002378 | NA | NA | -0.749943716 | 3.882579524 | 0.002349962 | 0.19183067 |
| ENSGALG00010014157 | NA | NA | -1.252429877 | 3.057030683 | 0.002357285 | 0.19183067 |
| ENSGALG00010003115 | UBE3D | 421843 | -0.336783047 | 4.100799635 | 0.002359766 | 0.19183067 |
| ENSGALG00010006220 | SGCG | 418945 | 0.374529753 | 3.599730943 | 0.002371362 | 0.19183067 |
| ENSGALG00010001162 | CD226 | 772097 | 0.431773219 | 3.040202682 | 0.002395182 | 0.19274314 |
| ENSGALG00010000831 | NA | NA | -0.901986749 | 7.410598458 | 0.002412046 | 0.193089346 |
| ENSGALG00010016852 | AHDC1 | 112530201 | 0.329161629 | 4.315658803 | 0.002443239 | 0.194572972 |
| ENSGALG00010006088 | NA | NA | -0.532131185 | 5.001918837 | 0.00249037 | 0.19730404 |
| ENSGALG00010003533 | LOC121107019 | 121107019 | -2.046423728 | 2.206400386 | 0.002600655 | 0.203285175 |
| ENSGALG00010021802 | NA | NA | -0.373095173 | 4.991519279 | 0.002605342 | 0.203285175 |
| ENSGALG00010027213 | KCNC4 | 419806 | 0.354940707 | 3.999618722 | 0.002605542 | 0.203285175 |
| ENSGALG00010005985 | DKK2 | 422531 | -0.914817005 | 0.401183321 | 0.002644798 | 0.204348883 |
| ENSGALG00010003617 | COG3 | 418846 | 0.228555678 | 5.925294759 | 0.002645766 | 0.204348883 |
| ENSGALG00010026003 | NECTIN1 | 100857875 | 0.217996119 | 6.687915153 | 0.002687502 | 0.206534491 |
| ENSGALG00010020399 | ATP13A2 | 419466 | 0.256518979 | 5.646715619 | 0.002780044 | 0.21200952 |
| ENSGALG00010019132 | H2AZ2 | 426617 | 0.27149844 | 6.873846504 | 0.002788805 | 0.21200952 |
| ENSGALG00010024534 | VWF | 419031 | -0.243467227 | 6.309141651 | 0.002800126 | 0.21200952 |
| ENSGALG00010029524 | BAHCC1 | 776014 | 0.316569987 | 4.31373203 | 0.002819168 | 0.21240493 |
| ENSGALG00010023525 | NA | NA | 1.980101284 | 0.435137094 | 0.002845014 | 0.21330666 |
| ENSGALG00010018296 | PIK3CA | 424971 | -0.235827101 | 6.28566714 | 0.002869521 | 0.213650008 |
| ENSGALG00010015218 | PLCL1 | 424060 | 0.245005417 | 6.808099688 | 0.002877394 | 0.213650008 |
| ENSGALG00010021964 | NA | NA | -0.347163799 | 5.597698838 | 0.002942623 | 0.217442864 |
| ENSGALG00010022546 | MTG1 | 791224 | 0.28427657 | 4.497170522 | 0.003019102 | 0.222026805 |
| ENSGALG00010029492 | SDK2 | 395215 | 0.307654236 | 4.382672812 | 0.00305415 | 0.223534658 |
| ENSGALG00010017642 | 1-Mar | 428563 | 0.22196945 | 7.207244876 | 0.003087483 | 0.224903389 |
| ENSGALG00010020833 | RANBP1 | 416787 | 0.251368783 | 9.443776965 | 0.003133951 | 0.226425825 |
| ENSGALG00010021010 | LEO1 | 769405 | 0.224675955 | 6.919000827 | 0.003137847 | 0.226425825 |
| ENSGALG00010008024 | MRS2 | 420820 | 0.253858066 | 5.044971891 | 0.003192494 | 0.229292683 |
| ENSGALG00010026213 | AKR1A1 | 424599 | 0.241104654 | 5.251969405 | 0.003249642 | 0.231666282 |
| ENSGALG00010004648 | LOC421806 | 421806 | -0.393525988 | 8.121851327 | 0.00327446 | 0.231666282 |
| ENSGALG00010001570 | SMPD1 | 419077 | 0.361642104 | 3.407957461 | 0.003281997 | 0.231666282 |
| ENSGALG00010003010 | NA | NA | -0.829059019 | 0.611499418 | 0.003285833 | 0.231666282 |
| ENSGALG00010006635 | NA | NA | -0.88831392 | 0.781240015 | 0.00331799 | 0.232865305 |
| ENSGALG00010029581 | DGKE | 770911 | 0.209080373 | 5.971209221 | 0.003366531 | 0.235198069 |
| ENSGALG00010022807 | LOC107050028 | 107050028 | 0.282622066 | 4.742913574 | 0.003463468 | 0.24087555 |
| ENSGALG00010008227 | DUSP1 | 374192 | -0.471533953 | 2.680926891 | 0.003492263 | 0.241784164 |
| ENSGALG00010016490 | NA | NA | 0.270193597 | 4.59996819 | 0.00361423 | 0.249106369 |
| ENSGALG00010024218 | NGFR | 425805 | 0.393771974 | 5.056246444 | 0.003642093 | 0.249906124 |
| ENSGALG00010000468 | TESC | 771113 | 0.558155635 | 2.13351077 | 0.003680734 | 0.250809096 |
| ENSGALG00010023287 | RGS19 | 374070 | 0.558849834 | 1.80075744 | 0.003687889 | 0.250809096 |
| ENSGALG00010018441 | CRIPT | 421412 | 0.235926314 | 5.29229218 | 0.003750087 | 0.253915584 |

| | | | | | | |
|---|---|---|---|---|---|---|
| ENSGALG00010028194 | LOC107055115 | 107055115 | 0.429539717 | 3.813143815 | 0.003775696 | 0.254528258 |
| ENSGALG00010015502 | LOC107049084 | 107049084 | -0.91005903 | 0.184724014 | 0.003865247 | 0.25873919 |
| ENSGALG00010021397 | NA | NA | -0.658524848 | 1.06352612 | 0.003871829 | 0.25873919 |
| ENSGALG00010000449 | PLBD2 | 417031 | 0.278699405 | 4.873142765 | 0.003989967 | 0.26547961 |
| ENSGALG00010011206 | LOC112532760 | 112532760 | -0.375494274 | 3.268369655 | 0.004031179 | 0.266111281 |
| ENSGALG00010012441 | FST | 396119 | -0.333438097 | 3.68879224 | 0.004045293 | 0.266111281 |
| ENSGALG00010024507 | NA | NA | -0.764532496 | 1.037312163 | 0.004051401 | 0.266111281 |
| ENSGALG00010027908 | PBXIP1 | 100857958 | 0.372554101 | 3.596367754 | 0.004094656 | 0.267342676 |
| ENSGALG00010024856 | SORT1 | 100858592 | 0.26277844 | 4.916819308 | 0.00410719 | 0.267342676 |
| ENSGALG00010023034 | NA | NA | -1.258897586 | -0.683180523 | 0.00412233 | 0.267342676 |
| ENSGALG00010002968 | NA | NA | -1.040651813 | 0.286525149 | 0.004197202 | 0.270875096 |
| ENSGALG00010024583 | LOC107049666 | 107049666 | 0.619723832 | 1.435260397 | 0.004218081 | 0.270875096 |
| ENSGALG00010010169 | NA | NA | -1.183209882 | 3.177557979 | 0.004234681 | 0.270875096 |
| ENSGALG00010004281 | VIM | 420519 | 0.259050823 | 10.09300802 | 0.004247293 | 0.270875096 |
| ENSGALG00010021155 | RADIL | 416479 | 0.974316589 | 0.134325264 | 0.004329353 | 0.274644738 |
| ENSGALG00010026242 | CCDC141 | 770766 | 0.353660316 | 3.578312862 | 0.004342139 | 0.274644738 |
| ENSGALG00010001479 | TMEM168 | 417777 | -0.221510809 | 5.379339837 | 0.004388907 | 0.275472141 |
| ENSGALG00010025355 | STAT6 | 100859196 | 0.373282537 | 3.279822031 | 0.004413384 | 0.275472141 |
| ENSGALG00010003549 | LOC121107023 | 121107023 | 2.261949414 | 2.330551009 | 0.004419476 | 0.275472141 |
| ENSGALG00010026297 | NA | NA | -0.989157378 | 0.110977648 | 0.004430593 | 0.275472141 |
| ENSGALG00010011096 | LDB2 | 395631 | 0.380993618 | 3.713459306 | 0.004444834 | 0.275472141 |
| ENSGALG00010022770 | HAT1 | 374037 | -0.219226596 | 6.1010763 | 0.004571528 | 0.281877842 |
| ENSGALG00010015390 | CCNB3 | 396167 | 0.240230641 | 5.514467937 | 0.004584871 | 0.281877842 |
| ENSGALG00010022216 | ROMO1 | 768833 | 0.311838558 | 5.273489675 | 0.004621764 | 0.282680211 |
| ENSGALG00010016896 | SPTBN1 | 421216 | 0.226303928 | 8.196624888 | 0.004634705 | 0.282680211 |
| ENSGALG00010025992 | MMP11 | 101751203 | 0.502918929 | 2.304315277 | 0.004733848 | 0.287585929 |
| ENSGALG00010020476 | TMEM132A | 107053338 | 0.211737475 | 5.903945205 | 0.004790889 | 0.289905364 |
| ENSGALG00010018749 | NA | NA | -0.198331663 | 6.090153722 | 0.004812834 | 0.290091236 |
| ENSGALG00010006408 | CCDC169 | 101751930 | -1.303562106 | -0.639944854 | 0.004864796 | 0.292077794 |
| ENSGALG00010019064 | DPY19L1 | 420743 | 0.246363407 | 7.007162487 | 0.004955122 | 0.295365175 |
| ENSGALG00010029577 | NOG | 373912 | 0.875180553 | 0.314545689 | 0.004957984 | 0.295365175 |
| ENSGALG00010009552 | NA | NA | -0.862458018 | 0.696244976 | 0.004990578 | 0.296159035 |
| ENSGALG00010024389 | AAAS | 100859661 | 0.196744588 | 6.35769263 | 0.005094217 | 0.301146585 |
| ENSGALG00010028912 | EXFABP | 396393 | 0.344991641 | 4.317478924 | 0.005138821 | 0.302152365 |
| ENSGALG00010012798 | MFSD6 | 423978 | -0.263934792 | 4.691332775 | 0.005150548 | 0.302152365 |
| ENSGALG00010020571 | MRTO4 | 428192 | 0.30761542 | 4.478187812 | 0.005191881 | 0.303017137 |
| ENSGALG00010009024 | PMAIP1 | 770077 | -0.652024786 | 1.265298424 | 0.005204719 | 0.303017137 |
| ENSGALG00010011757 | NA | NA | 0.367169718 | 3.706808056 | 0.005270688 | 0.303384437 |
| ENSGALG00010022409 | CARNS1 | 100359387 | 0.567454025 | 1.855189768 | 0.005276187 | 0.303384437 |
| ENSGALG00010018542 | LOC107054982 | 107054982 | 0.20245694 | 6.215922011 | 0.005288891 | 0.303384437 |
| ENSGALG00010003817 | BF2 | 425389 | 0.197167398 | 5.866950369 | 0.005289982 | 0.303384437 |
| ENSGALG00010002036 | NA | NA | -0.311220739 | 3.997886217 | 0.005334337 | 0.304770925 |
| ENSGALG00010017346 | GM2A | 100857973 | 0.28986753 | 5.120257251 | 0.005353816 | 0.304770925 |
| ENSGALG00010000217 | NA | NA | -0.441352058 | 3.816603246 | 0.005480515 | 0.309896192 |
| ENSGALG00010016907 | SQSTM1 | 770805 | -0.500954977 | 4.179652903 | 0.005484175 | 0.309896192 |
| ENSGALG00010012125 | RNF152 | 420909 | -0.19704257 | 6.297691679 | 0.00568787 | 0.320229156 |
| ENSGALG00010026532 | NA | NA | 0.277942067 | 4.649411538 | 0.005738054 | 0.321623267 |
| ENSGALG00010029351 | SLC25A10 | 769048 | 0.222243923 | 5.425324968 | 0.005754483 | 0.321623267 |
| ENSGALG00010000824 | NA | NA | -0.576935587 | 1.715597396 | 0.00578716 | 0.322277699 |
| ENSGALG00010017119 | BTBD6 | 101752037 | 0.215165718 | 6.035353756 | 0.005910661 | 0.327594071 |
| ENSGALG00010018340 | DNPEP | 424200 | 0.21572948 | 5.842297074 | 0.005936448 | 0.327594071 |
| ENSGALG00010001423 | FAM210A | 421045 | -0.221003917 | 5.965289067 | 0.005946568 | 0.327594071 |
| ENSGALG00010017250 | COL16A1 | 430477 | 0.467404679 | 3.258335404 | 0.005978748 | 0.328190575 |
| ENSGALG00010010214 | FAF2 | 416225 | 0.195062824 | 5.811354577 | 0.006067558 | 0.331880294 |
| ENSGALG00010013056 | SPATA46 | 424370 | 0.350572228 | 3.50809584 | 0.006113343 | 0.332373807 |
| ENSGALG00010021679 | MASP2 | 407089 | -0.400117593 | 3.147595987 | 0.00611983 | 0.332373807 |
| ENSGALG00010012835 | NA | NA | -0.750357098 | 0.588897671 | 0.00615819 | 0.332985223 |

| | | | | | | |
|---|---|---|---|---|---|---|
| ENSGALG00010000003 | NA | NA | -0.433591918 | 10.52110054 | 0.006174417 | 0.332985223 |
| ENSGALG00010014123 | NA | NA | -1.04534414 | -0.091405916 | 0.00622189 | 0.333414988 |
| ENSGALG00010000452 | SLC8B1 | 417032 | 0.482166109 | 2.296341023 | 0.006245362 | 0.333414988 |
| ENSGALG00010023311 | ETS1 | 396235 | 0.21913033 | 5.363388554 | 0.006254299 | 0.333414988 |
| ENSGALG00010027156 | ISYNA1 | 107055367 | 0.208471817 | 6.016034056 | 0.006269156 | 0.333414988 |
| ENSGALG00010029425 | SPNS2 | 417492 | -0.678145326 | 0.704217493 | 0.006335583 | 0.335443939 |
| ENSGALG00010010469 | NA | NA | -0.86935524 | 0.364663491 | 0.006365183 | 0.335443939 |
| ENSGALG00010002716 | NHLRC3 | 418890 | 0.31827033 | 3.938776151 | 0.00639543 | 0.335443939 |
| ENSGALG00010028686 | TAGLN2 | 121106445 | 0.213309051 | 5.989539394 | 0.006438282 | 0.335443939 |
| ENSGALG00010005326 | MASTL | 420487 | -0.20568614 | 7.095911991 | 0.006441355 | 0.335443939 |
| ENSGALG00010029277 | WFIKKN2 | 422097 | -0.603485207 | 2.234559867 | 0.006462536 | 0.335443939 |
| ENSGALG00010003814 | TPP1 | 425576 | 0.220752493 | 6.717351789 | 0.006467274 | 0.335443939 |
| ENSGALG00010001253 | ZBBX | 771469 | 0.301573334 | 3.963158712 | 0.006481903 | 0.335443939 |
| ENSGALG00010014250 | LOC101750517 | 101750517 | -0.802452865 | 0.39725693 | 0.006510665 | 0.335531277 |
| ENSGALG00010018379 | SRBD1 | 421408 | 0.207117153 | 6.860864777 | 0.006527251 | 0.335531277 |
| ENSGALG00010029932 | RASA4B | 417513 | 0.480579344 | 3.18792713 | 0.006557247 | 0.335949608 |
| ENSGALG00010007789 | WWOX | 415801 | 0.267380097 | 4.43201164 | 0.006787034 | 0.346298514 |
| ENSGALG00010022433 | LNPK | 424141 | 0.232809289 | 5.160614379 | 0.006809544 | 0.346298514 |
| ENSGALG00010004137 | CROT | 420533 | 0.214915721 | 5.685819616 | 0.006826835 | 0.346298514 |
| ENSGALG00010002220 | NA | NA | -0.509014029 | 1.980534488 | 0.006905049 | 0.346338121 |
| ENSGALG00010028341 | NA | NA | -0.972968786 | -0.2634017 | 0.006921974 | 0.346338121 |
| ENSGALG00010010705 | NA | NA | 0.553105513 | 1.606085808 | 0.0069398 | 0.346338121 |
| ENSGALG00010026422 | P2RX3 | 428856 | 0.523482005 | 2.420923225 | 0.006966222 | 0.346338121 |
| ENSGALG00010018029 | LOC121107471 | 121107471 | 0.773707839 | 1.752417248 | 0.007009499 | 0.346338121 |
| ENSGALG00010004128 | MCMDC2 | 428360 | -0.817663736 | 0.25906148 | 0.00702005 | 0.346338121 |
| ENSGALG00010026149 | LASP1 | 420002 | 0.226958563 | 5.171408135 | 0.007027145 | 0.346338121 |
| ENSGALG00010015631 | GSKIP | 423442 | 0.362344076 | 3.235112124 | 0.007051647 | 0.346338121 |
| ENSGALG00010001991 | TOMM7 | 420610 | 0.252219174 | 5.564053071 | 0.00705915 | 0.346338121 |
| ENSGALG00010027899 | PMVK | 101747962 | 0.246853839 | 4.685145304 | 0.007082885 | 0.346338121 |
| ENSGALG00010009493 | MXD3 | 427629 | 0.293151413 | 4.474988881 | 0.007086064 | 0.346338121 |
| ENSGALG00010012070 | DNAJC28 | 770607 | 0.410575497 | 2.621336037 | 0.007098016 | 0.346338121 |
| ENSGALG00010028568 | MLLT1 | 420089 | 0.196974342 | 6.740888748 | 0.007121706 | 0.346394352 |
| ENSGALG00010021095 | NT5C2 | 423871 | 0.206704249 | 6.341536223 | 0.00716581 | 0.347440074 |
| ENSGALG00010011679 | OVAL | 396058 | 0.34272264 | 4.979341534 | 0.007297365 | 0.351642882 |
| ENSGALG00010013502 | PAFAH2 | 769963 | 0.840188275 | 0.485259054 | 0.007298248 | 0.351642882 |
| ENSGALG00010003928 | NA | NA | 0.268537393 | 4.719993759 | 0.007410033 | 0.355913157 |
| ENSGALG00010027228 | SSPO | 420367 | 0.581490077 | 1.687093738 | 0.007459717 | 0.356876139 |
| ENSGALG00010028301 | FBN3 | 395079 | 0.275242585 | 5.972885578 | 0.007479411 | 0.356876139 |
| ENSGALG00010020328 | SLC17A6 | 422971 | -0.59016067 | 1.485359841 | 0.007509294 | 0.356876139 |
| ENSGALG00010008587 | NA | NA | -0.610180989 | 1.485269417 | 0.007541601 | 0.356876139 |
| ENSGALG00010027706 | GSN | 395774 | 0.201741802 | 5.736181548 | 0.007565651 | 0.356876139 |
| ENSGALG00010009160 | NA | NA | -1.125851688 | -0.346345152 | 0.007569396 | 0.356876139 |
| ENSGALG00010006025 | TNFAIP3 | 421684 | -0.523218426 | 2.315076299 | 0.007621482 | 0.358232956 |
| ENSGALG00010001172 | TMEM200C | 421054 | -0.282713111 | 4.277356257 | 0.007649031 | 0.358431744 |
| ENSGALG00010000060 | HYPK | 770307 | 0.200628371 | 6.198336611 | 0.007756564 | 0.361730507 |
| ENSGALG00010008272 | DCP2 | 769972 | -0.189654721 | 6.200313678 | 0.007766498 | 0.361730507 |
| ENSGALG00010010174 | NA | NA | -1.31714746 | 2.452313153 | 0.007819486 | 0.363098175 |
| ENSGALG00010005241 | C4H4orf54 | 112532442 | -0.199946537 | 6.728747331 | 0.007864857 | 0.364104995 |
| ENSGALG00010025255 | NA | NA | -0.766406799 | 1.649536151 | 0.007898742 | 0.364575571 |
| ENSGALG00010008914 | NA | NA | -0.268161915 | 5.233470893 | 0.007984249 | 0.366880052 |
| ENSGALG00010021947 | CDK2AP2 | 107053360 | 0.328793182 | 3.549166071 | 0.00799641 | 0.366880052 |
| ENSGALG00010017493 | YPEL5 | 421299 | 0.22546628 | 7.327217844 | 0.008125154 | 0.371677439 |
| ENSGALG00010012259 | IL1RAPL2 | 422379 | -0.553921309 | 2.650799762 | 0.008249584 | 0.376249574 |
| ENSGALG00010023549 | NA | NA | 0.456153164 | 5.819527697 | 0.008296984 | 0.376348895 |
| ENSGALG00010013435 | SDC3 | 396343 | 0.199790652 | 6.316904848 | 0.008300734 | 0.376348895 |
| ENSGALG00010003850 | PLA2G12A | 772082 | -0.202186369 | 5.805658755 | 0.008417398 | 0.379825258 |
| ENSGALG00010018533 | PCYOX1 | 426297 | 0.228721238 | 5.045518872 | 0.008426832 | 0.379825258 |

| | | | | | | |
|---|---|---|---|---|---|---|
| ENSGALG00010029647 | RSAD1 | 422101 | 0.360241826 | 3.188212958 | 0.008473441 | 0.380809309 |
| ENSGALG00010015292 | NA | NA | 0.261189174 | 6.404341106 | 0.008527844 | 0.381824568 |
| ENSGALG00010003090 | DMA | 417050 | 0.46250016 | 2.031922229 | 0.008562533 | 0.381824568 |
| ENSGALG00010001343 | LOC121108665 | 121108665 | -0.978012537 | -0.262793193 | 0.008605697 | 0.381824568 |
| ENSGALG00010007399 | WWC2 | 422555 | 0.236312988 | 5.752152779 | 0.008611349 | 0.381824568 |
| ENSGALG00010018899 | ACSS1A | 416714 | 0.322220598 | 4.337351581 | 0.00862912 | 0.381824568 |
| ENSGALG00010023655 | C1R | 418295 | 0.274351272 | 4.279842344 | 0.008656362 | 0.381824568 |
| ENSGALG00010002618 | RGNL | 428008 | -1.045685553 | -0.648492047 | 0.008670873 | 0.381824568 |
| ENSGALG00010027692 | PHF2 | 415981 | -0.282674382 | 5.08696447 | 0.008703384 | 0.381824568 |
| ENSGALG00010015669 | INPP4A | 418690 | 0.209500905 | 5.828749594 | 0.008719611 | 0.381824568 |
| ENSGALG00010006091 | RTKN | 107056271 | 0.310982241 | 3.925343569 | 0.00875844 | 0.382435295 |
| ENSGALG00010028615 | NA | NA | -0.371760623 | 3.239898853 | 0.0088097 | 0.383583806 |
| ENSGALG00010010826 | NCOA1 | 422016 | 0.192040935 | 6.695418393 | 0.008848062 | 0.383599407 |
| ENSGALG00010008867 | NA | NA | -0.675786073 | 0.666280268 | 0.008859973 | 0.383599407 |
| ENSGALG00010008718 | NA | NA | -1.01651242 | -0.518100958 | 0.008924336 | 0.385300691 |
| ENSGALG00010007102 | CMTM3 | 414829 | 0.291806284 | 4.199019002 | 0.008958192 | 0.385679006 |
| ENSGALG00010025185 | NA | NA | -0.344792351 | 3.27915037 | 0.009024021 | 0.386907711 |
| ENSGALG00010000982 | NA | NA | 0.315685066 | 3.477139144 | 0.009037077 | 0.386907711 |
| ENSGALG00010001588 | ARHGAP33L1 | 100858315 | 1.094924216 | -0.563029597 | 0.00910386 | 0.388684239 |
| ENSGALG00010007923 | YAP1 | 396171 | 0.241926444 | 7.265934822 | 0.009145741 | 0.389390694 |
| ENSGALG00010029341 | SLC43A2 | 417557 | 0.259891436 | 4.50016567 | 0.009186072 | 0.390027401 |
| ENSGALG00010005765 | NA | NA | -0.476857441 | 2.349676178 | 0.009216442 | 0.390238867 |
| ENSGALG00010020097 | RETSAT | 426549 | 0.320235817 | 3.850977582 | 0.009339764 | 0.394374098 |
| ENSGALG00010012843 | NA | NA | -1.006958506 | -0.004036724 | 0.009408109 | 0.396171614 |
| ENSGALG00010000005 | NA | NA | -0.416557873 | 11.54630766 | 0.009492619 | 0.398638137 |
| ENSGALG00010021723 | COPS3 | 416504 | 0.214276597 | 7.284148276 | 0.009547174 | 0.399836703 |
| ENSGALG00010015709 | RBMXL3 | 416302 | 0.24247576 | 8.707349706 | 0.009646089 | 0.402881489 |
| ENSGALG00010018374 | HPS1 | 429879 | 0.362715567 | 3.629473349 | 0.009681792 | 0.403276803 |
| ENSGALG00010000242 | CHIR2AB2 | 425238 | 0.324499048 | 3.687262488 | 0.009795131 | 0.406895017 |
| ENSGALG00010023930 | SRPRA | 419713 | 0.214517959 | 6.698414959 | 0.009900074 | 0.407623812 |
| ENSGALG00010004076 | COX7A2 | 772260 | 0.404642008 | 5.213175261 | 0.009903647 | 0.407623812 |
| ENSGALG00010026291 | SERPING1 | 423132 | 0.242819662 | 4.964013939 | 0.009925087 | 0.407623812 |
| ENSGALG00010007695 | RASGRP3 | 421460 | -0.697018531 | 0.611134456 | 0.009926732 | 0.407623812 |
| ENSGALG00010003524 | NA | NA | -0.942416332 | 0.175566751 | 0.009967437 | 0.407623812 |
| ENSGALG00010006926 | ID4 | 374150 | 0.266999514 | 4.890427092 | 0.009971799 | 0.407623812 |
| ENSGALG00010011247 | CDYL | 420877 | -0.244042651 | 6.194019296 | 0.010067359 | 0.410438471 |
| ENSGALG00010003678 | ZNFY1 | 120623203 | 0.388768507 | 3.125075267 | 0.010236915 | 0.41484079 |
| ENSGALG00010021172 | SLC29A4 | 425593 | 0.227429413 | 5.97431471 | 0.010252788 | 0.41484079 |
| ENSGALG00010012905 | NKAIN1 | 107049081 | 0.22194967 | 5.514420981 | 0.010256311 | 0.41484079 |
| ENSGALG00010014750 | LONRF1 | 422742 | -0.24575747 | 4.728815501 | 0.010342642 | 0.417234665 |
| ENSGALG00010000015 | NA | NA | -1.603375962 | -0.612761262 | 0.01037339 | 0.417379606 |
| ENSGALG00010005730 | NA | NA | -0.806308433 | 0.289479297 | 0.010447954 | 0.419282141 |
| ENSGALG00010000315 | NA | NA | 0.721328935 | 2.013867141 | 0.010527231 | 0.421349138 |
| ENSGALG00010003616 | LOC121107011 | 121107011 | -1.060389029 | 2.824138446 | 0.010554289 | 0.421349138 |
| ENSGALG00010023841 | OSBPL7 | 100857443 | 0.356095612 | 3.343779964 | 0.010600527 | 0.422098717 |
| ENSGALG00010000412 | RFC5 | 100859719 | 0.212512809 | 5.969670106 | 0.01066464 | 0.423315648 |
| ENSGALG00010009147 | HNMT | 424298 | -0.845906204 | 0.214637106 | 0.010687097 | 0.423315648 |
| ENSGALG00010024121 | NRG4 | 415354 | -0.603930112 | 1.087121308 | 0.010713714 | 0.423315648 |
| ENSGALG00010024695 | TRABD2B | 424623 | -0.210458701 | 5.656865572 | 0.010763348 | 0.423794853 |
| ENSGALG00010015816 | MITD1 | 418698 | -0.336448788 | 3.259853061 | 0.010780988 | 0.423794853 |
| ENSGALG00010018080 | BRICD5 | 107054618 | -0.917654683 | -0.082694061 | 0.010914931 | 0.42539342 |
| ENSGALG00010019634 | GNG2 | 423581 | 0.264051051 | 4.425026217 | 0.010948749 | 0.42539342 |
| ENSGALG00010012749 | RGS4 | 378901 | 0.386105067 | 2.838523223 | 0.010972871 | 0.42539342 |
| ENSGALG00010012426 | MIEF1 | 418012 | 0.210634217 | 5.061364963 | 0.011002279 | 0.42539342 |
| ENSGALG00010025044 | TMEM9 | 421157 | 0.530004705 | 1.582266261 | 0.011006093 | 0.42539342 |
| ENSGALG00010011895 | PHF21B | 427935 | 0.531588657 | 1.72731899 | 0.011013441 | 0.42539342 |
| ENSGALG00010027231 | ATP2B2 | 415958 | -0.287511399 | 4.387180222 | 0.011034935 | 0.42539342 |

| | | | | | | |
|---|---|---|---|---|---|---|
| ENSGALG00010022942 | NEU2 | 107049056 | -0.711568594 | 0.589470467 | 0.011043069 | 0.42539342 |
| ENSGALG00010005531 | SUSD1 | 427334 | 0.232503672 | 4.98920796 | 0.01109874 | 0.426469077 |
| ENSGALG00010000811 | LOC770405 | 770405 | 0.695330848 | 1.006775423 | 0.011365817 | 0.43386798 |
| ENSGALG00010001810 | OPN2SW | 396525 | 0.211437106 | 5.434904451 | 0.011373509 | 0.43386798 |
| ENSGALG00010016447 | ID3 | 395281 | 0.345079757 | 3.686974218 | 0.011425335 | 0.43386798 |
| ENSGALG00010006304 | NA | NA | -0.92754723 | -0.202044583 | 0.011443981 | 0.43386798 |
| ENSGALG00010009991 | LHFPL3 | 100858127 | 0.225467633 | 5.039201419 | 0.011455489 | 0.43386798 |
| ENSGALG00010017307 | TMEM186 | 416641 | -0.200730312 | 5.40194081 | 0.011460664 | 0.43386798 |
| ENSGALG00010015680 | FRG1BP | 422726 | -0.179310735 | 5.980255615 | 0.011546625 | 0.434896941 |
| ENSGALG00010016411 | CTSH | 770109 | 0.260598091 | 4.452405605 | 0.011563514 | 0.434896941 |
| ENSGALG00010008081 | TOP1MT | 408025 | 0.781422048 | 0.414865687 | 0.011572729 | 0.434896941 |
| ENSGALG00010014894 | RAB22A | 419318 | -0.255659452 | 4.4707736 | 0.011704582 | 0.437840399 |
| ENSGALG00010012062 | HMGA2 | 395851 | 0.230111589 | 4.738542879 | 0.011708029 | 0.437840399 |
| ENSGALG00010003155 | NA | NA | -0.481276149 | 2.627489724 | 0.011776932 | 0.439348149 |
| ENSGALG00010011340 | PKNOX1 | 418542 | -0.203306269 | 5.394905681 | 0.011862754 | 0.441478287 |
| ENSGALG00010000729 | EDNRB | 408082 | 0.232232833 | 7.53284179 | 0.011907589 | 0.442076447 |
| ENSGALG00010028244 | NA | NA | -1.049676065 | -0.588283117 | 0.011955732 | 0.442794223 |
| ENSGALG00010019280 | MESDC1 | 415473 | 0.18401542 | 6.294819306 | 0.011999844 | 0.443359638 |
| ENSGALG00010021477 | LOC100858626 | 100858626 | 0.353258156 | 3.392707083 | 0.012035114 | 0.443596405 |
| ENSGALG00010008103 | RBPMS | 428738 | 0.194894574 | 6.372009169 | 0.012093359 | 0.444676876 |
| ENSGALG00010003231 | NA | NA | -0.615439133 | 0.91026203 | 0.012124344 | 0.444752205 |
| ENSGALG00010018177 | KIF20B | 423793 | -0.192294888 | 7.774259229 | 0.012184786 | 0.44590515 |
| ENSGALG00010028445 | SYT11 | 429166 | 0.18502561 | 7.828592559 | 0.01226868 | 0.445997158 |
| ENSGALG00010023179 | KAT2A | 374232 | 0.183424369 | 5.788076742 | 0.012274907 | 0.445997158 |
| ENSGALG00010015370 | ARHGAP20B | 422171 | -0.223440429 | 7.177156343 | 0.012299507 | 0.445997158 |
| ENSGALG00010012230 | PTPRQ | 772163 | -1.167973875 | -0.547417474 | 0.012320304 | 0.445997158 |
| ENSGALG00010022518 | ECHS1 | 770828 | 0.324412026 | 4.467303451 | 0.012353986 | 0.445997158 |
| ENSGALG00010029269 | WSCD1 | 427842 | 0.364469512 | 3.130180102 | 0.012361405 | 0.445997158 |
| ENSGALG00010025850 | NA | NA | 0.990311474 | -0.393430398 | 0.012416155 | 0.446923429 |
| ENSGALG00010015574 | NA | NA | -0.493895007 | 2.744230271 | 0.0124728 | 0.447913419 |
| ENSGALG00010011043 | NA | NA | -0.856798 | 0.042918687 | 0.012592056 | 0.451141947 |
| ENSGALG00010000096 | NA | NA | -1.196088195 | -0.222899531 | 0.012693431 | 0.452431935 |
| ENSGALG00010023199 | RPS13 | 414782 | 0.20224707 | 8.603565017 | 0.012706005 | 0.452431935 |
| ENSGALG00010026874 | TEAD3 | 395810 | 0.207560265 | 5.08176883 | 0.012728526 | 0.452431935 |
| ENSGALG00010015391 | SYN3 | 100857420 | 0.896702246 | -0.276246998 | 0.012765965 | 0.452431935 |
| ENSGALG00010002958 | DHRS12 | 770438 | 0.62072524 | 0.984851408 | 0.012809101 | 0.452431935 |
| ENSGALG00010021955 | NA | NA | -0.728671969 | 0.461106362 | 0.012828983 | 0.452431935 |
| ENSGALG00010000483 | LOC107051159 | 107051159 | 0.374067255 | 3.304808604 | 0.01284581 | 0.452431935 |
| ENSGALG00010018693 | NA | NA | 0.35122054 | 3.271728834 | 0.012870119 | 0.452431935 |
| ENSGALG00010013336 | MTNR1A | 396319 | -0.552727645 | 1.627220794 | 0.012892986 | 0.452431935 |
| ENSGALG00010017139 | COL9A2 | 396524 | 0.235754658 | 5.023533551 | 0.012973774 | 0.45419209 |
| ENSGALG00010015093 | NA | NA | -1.004708766 | -0.573726246 | 0.013002246 | 0.45419209 |
| ENSGALG00010024789 | SUOX | 107055404 | 0.298350446 | 3.768306142 | 0.013096024 | 0.455720757 |
| ENSGALG00010016069 | GALK2 | 415596 | 0.398764423 | 3.120033496 | 0.013105307 | 0.455720757 |
| ENSGALG00010007691 | LOC121108447 | 121108447 | -0.504780437 | 3.022044293 | 0.013196524 | 0.457001805 |
| ENSGALG00010021688 | ITPR1 | 395412 | -0.232881618 | 5.851446847 | 0.013201614 | 0.457001805 |
| ENSGALG00010019691 | NA | NA | -0.400637065 | 3.641364611 | 0.01323824 | 0.457239869 |
| ENSGALG00010027173 | AES | 100858542 | 0.214367675 | 5.554723843 | 0.013308694 | 0.458642655 |
| ENSGALG00010019801 | FIBIN | 426933 | 0.269245904 | 6.687548843 | 0.013347622 | 0.458955155 |
| ENSGALG00010011597 | TBC1D14 | 422862 | -0.181979457 | 8.942428596 | 0.013416517 | 0.460294359 |
| ENSGALG00010001553 | CNGA4 | 430630 | 0.937982015 | -0.053699587 | 0.013760958 | 0.470534928 |
| ENSGALG00010011949 | ABCA12 | 424011 | -0.367792025 | 2.87687704 | 0.013776234 | 0.470534928 |
| ENSGALG00010025536 | LEXM | 424660 | -0.670320311 | 1.026640424 | 0.013862923 | 0.471522137 |
| ENSGALG00010025428 | VANGL1 | 418337 | -0.194089855 | 7.122397863 | 0.013866494 | 0.471522137 |
| ENSGALG00010023218 | SSH3 | 101752044 | 0.31414692 | 3.603036415 | 0.013923134 | 0.472403034 |
| ENSGALG00010006341 | HHIP | 422460 | -0.400071543 | 3.030132162 | 0.013978822 | 0.472658114 |
| ENSGALG00010021509 | DDIT4 | 107053650 | 0.383024258 | 5.911120383 | 0.01402201 | 0.472658114 |

| | | | | | | |
|---|---|---|---|---|---|---|
| ENSGALG00010024831 | SLC7A9 | 415768 | -0.286279246 | 5.20728094 | 0.014022908 | 0.472658114 |
| ENSGALG00010022343 | GATA5 | 396391 | -0.605332927 | 1.423993932 | 0.014110063 | 0.474555062 |
| ENSGALG00010025327 | TMEM59 | 424656 | 0.196088226 | 7.379888666 | 0.014248635 | 0.478169255 |
| ENSGALG00010008708 | ZC3H4 | 101748292 | 0.183631178 | 5.793369508 | 0.014373444 | 0.481306825 |
| ENSGALG00010015038 | PCSK5 | 395456 | 0.177298963 | 6.410379881 | 0.014407783 | 0.481407879 |
| ENSGALG00010027562 | SLC39A3 | 429050 | 0.336454024 | 3.592518553 | 0.014465915 | 0.482301754 |
| ENSGALG00010022034 | BRI3 | 427655 | 0.270553593 | 4.314032347 | 0.01467301 | 0.488147548 |
| ENSGALG00010004928 | HDDC2 | 421719 | 0.212955886 | 5.140171924 | 0.014711578 | 0.488373555 |
| ENSGALG00010010578 | GLUD2 | 423612 | 0.189043375 | 5.647643978 | 0.015030768 | 0.497033663 |
| ENSGALG00010025316 | SLC25A36 | 424817 | -0.200532867 | 6.932130047 | 0.015077405 | 0.497033663 |
| ENSGALG00010021473 | SLC8A3 | 395759 | -0.20261415 | 4.995512799 | 0.015082385 | 0.497033663 |
| ENSGALG00010016861 | FGR | 777583 | 0.391401555 | 2.793358682 | 0.015101804 | 0.497033663 |
| ENSGALG00010003437 | PNRC1 | 421817 | 0.198923529 | 7.610748607 | 0.015203538 | 0.499263602 |
| ENSGALG00010028326 | CRB2 | 417104 | -0.18773518 | 5.560636944 | 0.015243423 | 0.499263602 |
| ENSGALG00010026474 | LOC121107673 | 121107673 | 1.751382606 | 0.582574335 | 0.015267007 | 0.499263602 |
| ENSGALG00010027311 | NFASC | 419824 | 0.362196016 | 4.753220228 | 0.015321709 | 0.499988694 |
| ENSGALG00010027171 | LOC107055991 | 107055991 | 0.282358349 | 3.645323415 | 0.015385522 | 0.501007354 |
| ENSGALG00010024684 | CADNL | 427547 | -0.253649293 | 5.105269297 | 0.01550936 | 0.501320793 |
| ENSGALG00010018851 | NA | NA | -0.190011641 | 6.438471273 | 0.015527846 | 0.501320793 |
| ENSGALG00010011621 | WASHC4 | 418077 | 0.182107797 | 6.947535142 | 0.015537161 | 0.501320793 |
| ENSGALG00010026266 | NA | NA | -0.849393929 | 0.234853508 | 0.015545233 | 0.501320793 |
| ENSGALG00010019600 | MAP3K19 | 429048 | -0.548636349 | 1.197418164 | 0.015589525 | 0.501320793 |
| ENSGALG00010005409 | PRKCQ | 776459 | 0.277239803 | 4.375288691 | 0.015600298 | 0.501320793 |
| ENSGALG00010017226 | PEF1 | 419644 | 0.203259621 | 5.063918509 | 0.015623465 | 0.501320793 |
| ENSGALG00010027717 | ATXN7L2 | 429691 | 0.376476304 | 2.819559861 | 0.016015461 | 0.512828414 |
| ENSGALG00010021806 | NA | NA | -0.336646862 | 3.364691737 | 0.016112564 | 0.514865107 |
| ENSGALG00010022732 | NA | NA | -0.696119461 | 0.37442914 | 0.016168646 | 0.515147056 |
| ENSGALG00010011776 | IL17RA | 418158 | 0.170458335 | 5.885770151 | 0.016219883 | 0.515147056 |
| ENSGALG00010005351 | DTNA | 421104 | 0.185710482 | 5.793477375 | 0.016230502 | 0.515147056 |
| ENSGALG00010029487 | CDC42EP4 | 771509 | -0.396926107 | 2.373950467 | 0.016255454 | 0.515147056 |
| ENSGALG00010003063 | GGH | 421144 | 0.18480713 | 5.324605977 | 0.016299858 | 0.515491409 |
| ENSGALG00010015500 | NA | NA | -0.774621563 | 0.558367478 | 0.016399141 | 0.517566305 |
| ENSGALG00010021781 | RAPGEF4 | 429022 | -0.236791543 | 4.384352418 | 0.016499204 | 0.518305896 |
| ENSGALG00010025329 | MYO1A | 396072 | 0.338789876 | 3.995801483 | 0.016529221 | 0.518305896 |
| ENSGALG00010003618 | LOC112533598 | 112533598 | -0.922809729 | 2.282558237 | 0.016549191 | 0.518305896 |
| ENSGALG00010001956 | NA | NA | 0.919145255 | -0.185495926 | 0.016620447 | 0.518305896 |
| ENSGALG00010028070 | MEX3A | 101747646 | 0.204800519 | 5.790356377 | 0.016643029 | 0.518305896 |
| ENSGALG00010011968 | LOC427400 | 427400 | 0.90522367 | -0.370174629 | 0.016644458 | 0.518305896 |
| ENSGALG00010028573 | KCNN1 | 429723 | 0.250985144 | 4.465519757 | 0.016658628 | 0.518305896 |
| ENSGALG00010021617 | SORBS1 | 423749 | 0.270161593 | 5.095799754 | 0.016787079 | 0.521247296 |
| ENSGALG00010012759 | ATP1B4 | 422365 | -0.604167098 | 0.770288024 | 0.016939172 | 0.524814795 |
| ENSGALG00010028363 | NA | NA | -0.523247758 | 3.132058524 | 0.017002415 | 0.524814795 |
| ENSGALG00010007545 | NA | NA | -0.829785915 | -0.208853213 | 0.017004409 | 0.524814795 |
| ENSGALG00010024907 | PHLDB1 | 419790 | 0.257326158 | 4.976798764 | 0.017044638 | 0.524838651 |
| ENSGALG00010008213 | ISCA1 | 427463 | -0.187239736 | 6.415295031 | 0.017073476 | 0.524838651 |
| ENSGALG00010025645 | NBEAL2 | 425984 | 0.19061614 | 6.417628722 | 0.017195637 | 0.526907561 |
| ENSGALG00010027955 | LOC107050229 | 107050229 | 0.192706165 | 5.202313855 | 0.017295523 | 0.526907561 |
| ENSGALG00010006610 | NA | NA | -0.618193645 | 0.923942862 | 0.017295845 | 0.526907561 |
| ENSGALG00010024374 | CLDN15 | 424768 | -0.909152507 | -0.368138497 | 0.017381533 | 0.526907561 |
| ENSGALG00010027219 | LOC101749138 | 101749138 | 0.176261883 | 5.882827212 | 0.017412307 | 0.526907561 |
| ENSGALG00010014797 | ZDHHC9 | 422139 | 0.215835088 | 7.19566738 | 0.017520295 | 0.526907561 |
| ENSGALG00010008384 | NA | NA | -0.568286041 | 2.328710855 | 0.017560389 | 0.526907561 |
| ENSGALG00010027923 | AMPD2 | 112530327 | 0.188241538 | 5.41845769 | 0.017567994 | 0.526907561 |
| ENSGALG00010018858 | EPB41L5 | 424235 | 0.186225407 | 7.713554805 | 0.017574531 | 0.526907561 |
| ENSGALG00010000321 | NA | NA | 1.224892145 | -0.187577911 | 0.017582452 | 0.526907561 |
| ENSGALG00010021828 | NA | NA | -0.878214119 | -0.125458095 | 0.017605609 | 0.526907561 |
| ENSGALG00010009166 | CLGN | 422447 | -0.201268852 | 6.399581445 | 0.017620987 | 0.526907561 |

| | | | | | | |
|---|---|---|---|---|---|---|
| ENSGALG00010025072 | VAMP2 | 424827 | 0.806553423 | -0.059602008 | 0.017634087 | 0.526907561 |
| ENSGALG00010014572 | IL1RAP | 424908 | 0.276898642 | 4.382979021 | 0.017663883 | 0.526907561 |
| ENSGALG00010020579 | NA | NA | 0.749305566 | 0.137061442 | 0.017688212 | 0.526907561 |
| ENSGALG00010027329 | FAM188B | 420383 | 0.334097258 | 3.566933814 | 0.017714563 | 0.526907561 |
| ENSGALG00010027883 | ATP8B2 | 770035 | 0.187889224 | 6.264124264 | 0.017730742 | 0.526907561 |
| ENSGALG00010015802 | TRPC7 | 427641 | -0.52315943 | 1.373044142 | 0.017757848 | 0.526907561 |
| ENSGALG00010013329 | LIFR | 395262 | 0.212886541 | 5.504767468 | 0.017809154 | 0.527411752 |
| ENSGALG00010029703 | MIF4GD | 100858848 | 0.218334071 | 5.117188778 | 0.017924595 | 0.529450377 |
| ENSGALG00010012831 | NA | NA | -0.649824939 | 0.684334588 | 0.017992302 | 0.529450377 |
| ENSGALG00010019050 | TNS1 | 396439 | 0.308549136 | 3.888479461 | 0.018054301 | 0.529450377 |
| ENSGALG00010020103 | PSAP | 395602 | 0.222406925 | 6.937992881 | 0.018060186 | 0.529450377 |
| ENSGALG00010027120 | UNC13A | 430487 | 0.740292875 | 0.128336664 | 0.018076232 | 0.529450377 |
| ENSGALG00010007899 | APRT | 100857155 | 0.280374058 | 4.338301407 | 0.01809021 | 0.529450377 |
| ENSGALG00010017422 | NA | NA | 0.707530728 | 0.337950468 | 0.018119122 | 0.529450377 |
| ENSGALG00010019416 | NA | NA | -0.886891289 | -0.015410959 | 0.018155214 | 0.529498378 |
| ENSGALG00010029074 | NA | NA | -0.847180501 | -0.455430848 | 0.018322139 | 0.53335468 |
| ENSGALG00010013247 | ATG5 | 421784 | 0.182387569 | 5.616788539 | 0.018452616 | 0.535318333 |
| ENSGALG00010021263 | STOML1 | 415303 | 0.426348301 | 2.360320407 | 0.018459253 | 0.535318333 |
| ENSGALG00010003400 | PKIA | 396333 | -0.225484612 | 5.134557608 | 0.018629461 | 0.537560361 |
| ENSGALG00010008999 | TRAPPC9 | 420312 | 0.265057963 | 3.896174378 | 0.018646286 | 0.537560361 |
| ENSGALG00010022783 | STAG1 | 424735 | 0.190597802 | 5.07249318 | 0.018677659 | 0.537560361 |
| ENSGALG00010020695 | LOC395991 | 395991 | -0.802277788 | -0.036500289 | 0.018689766 | 0.537560361 |
| ENSGALG00010009891 | NA | NA | -0.414680996 | 3.497375322 | 0.018739154 | 0.537560361 |
| ENSGALG00010010228 | NA | NA | -0.671053485 | 0.578551196 | 0.018765155 | 0.537560361 |
| ENSGALG00010020763 | NA | NA | -0.67056073 | 0.589921581 | 0.018786589 | 0.537560361 |
| ENSGALG00010027605 | LOC101748987 | 101748987 | 0.2960094 | 3.59941654 | 0.018816361 | 0.537560361 |
| ENSGALG00010019001 | ALDH1A3 | 395389 | 0.485676478 | 1.698766113 | 0.018891495 | 0.538705524 |
| ENSGALG00010011302 | NA | NA | -0.642690226 | 0.660515433 | 0.01902469 | 0.540508981 |
| ENSGALG00010016813 | LOC422929 | 422929 | -0.809613025 | -0.329646923 | 0.019025072 | 0.540508981 |
| ENSGALG00010005457 | SEMA3D | 396332 | 0.409282551 | 2.830814991 | 0.019187379 | 0.543372024 |
| ENSGALG00010018453 | TMX4 | 416729 | 0.191508472 | 6.816965792 | 0.019196552 | 0.543372024 |
| ENSGALG00010004975 | SNW1 | 423381 | -0.205417691 | 7.312733819 | 0.019232127 | 0.543378283 |
| ENSGALG00010024477 | NA | NA | -0.773611735 | 0.037059236 | 0.019325646 | 0.544096359 |
| ENSGALG00010013018 | ID1 | 395282 | 0.416840869 | 2.433234144 | 0.019345552 | 0.544096359 |
| ENSGALG00010001653 | NA | NA | -0.653894728 | 0.463023897 | 0.019363742 | 0.544096359 |
| ENSGALG00010008284 | BCKDHA | 374210 | 0.216889612 | 4.83574461 | 0.019455079 | 0.545535419 |
| ENSGALG00010001492 | NA | NA | -0.784090836 | -0.168451301 | 0.019485943 | 0.545535419 |
| ENSGALG00010011662 | RIDA | 426536 | 0.265575834 | 4.540908574 | 0.019599714 | 0.547017691 |
| ENSGALG00010006155 | TNFRSF19 | 418943 | 0.342878906 | 3.239077529 | 0.019612121 | 0.547017691 |
| ENSGALG00010009198 | KCNN2 | 395579 | 0.472920839 | 1.909488801 | 0.019645658 | 0.547017691 |
| ENSGALG00010009781 | NA | NA | -0.637926501 | 1.155288248 | 0.019732437 | 0.547962674 |
| ENSGALG00010022977 | NA | NA | -0.434415803 | 3.159848708 | 0.019818327 | 0.547962674 |
| ENSGALG00010017609 | TRIM71 | 428445 | -0.208330026 | 9.758094248 | 0.01982254 | 0.547962674 |
| ENSGALG00010016970 | SYNE3 | 423439 | -0.981763377 | -0.723740816 | 0.019835769 | 0.547962674 |
| ENSGALG00010014680 | LOC107053920 | 107053920 | -0.194300418 | 7.447637798 | 0.01990448 | 0.547962674 |
| ENSGALG00010025077 | NA | NA | -0.510547398 | 1.854589724 | 0.019917373 | 0.547962674 |
| ENSGALG00010002454 | NEK5 | 418881 | 0.446558745 | 2.043531372 | 0.019929156 | 0.547962674 |
| ENSGALG00010029394 | CYGB | 427802 | 0.413867131 | 3.007324406 | 0.019973481 | 0.548200724 |
| ENSGALG00010027046 | HESX1 | 395864 | -0.433349936 | 3.665384125 | 0.020028292 | 0.548419892 |
| ENSGALG00010009751 | NA | NA | -0.740186286 | 0.154107339 | 0.020052829 | 0.548419892 |
| ENSGALG00010018590 | TGFA | 414743 | 0.221378804 | 4.975584658 | 0.020114101 | 0.549118519 |
| ENSGALG00010029534 | NF1 | 396085 | 0.187087327 | 6.835632711 | 0.02024949 | 0.551834507 |
| ENSGALG00010023182 | EPHB6 | 418308 | 0.243242468 | 5.011503924 | 0.020337312 | 0.553246885 |
| ENSGALG00010023692 | NA | NA | -0.44651823 | 2.535086457 | 0.020482579 | 0.556214213 |
| ENSGALG00010004307 | NPC2 | 423353 | 0.229037354 | 4.492203751 | 0.020644823 | 0.559631256 |
| ENSGALG00010017407 | EPN2 | 416524 | 0.179666934 | 6.132843688 | 0.020944811 | 0.566763639 |
| ENSGALG00010008765 | NA | NA | -0.650689981 | 0.696267355 | 0.021019065 | 0.56756748 |

| | | | | | | |
|---|---|---|---|---|---|---|
| ENSGALG00010022115 | E2F4 | 769430 | 0.201174972 | 5.327981963 | 0.021048371 | 0.56756748 |
| ENSGALG00010009809 | SMDT1 | 770755 | 0.233168301 | 4.266193111 | 0.021122517 | 0.567775121 |
| ENSGALG00010016225 | P4HA2 | 416326 | -0.210203789 | 4.996599703 | 0.021129952 | 0.567775121 |
| ENSGALG00010014270 | NA | NA | -0.20674859 | 6.806647308 | 0.021181466 | 0.568166018 |
| ENSGALG00010018568 | NA | NA | 0.196464532 | 5.54243537 | 0.021275633 | 0.569431299 |
| ENSGALG00010020457 | IL5RA | 771801 | -0.33235123 | 3.53084871 | 0.021330756 | 0.569431299 |
| ENSGALG00010023263 | SRP14 | 423284 | 0.166125787 | 6.171952625 | 0.021339781 | 0.569431299 |
| ENSGALG00010011148 | LOC107050579 | 107050579 | -0.608626059 | 3.928350139 | 0.021421535 | 0.569856703 |
| ENSGALG00010003372 | ENOX1 | 418839 | -0.397815509 | 2.150011551 | 0.021463369 | 0.569856703 |
| ENSGALG00010003110 | BF1 | 693260 | 0.207351117 | 4.662783376 | 0.021498525 | 0.569856703 |
| ENSGALG00010014827 | LOC112532963 | 112532963 | -0.876793278 | -0.353309394 | 0.021504027 | 0.569856703 |
| ENSGALG00010028068 | POLE4 | 107055326 | 0.203124495 | 6.517020803 | 0.02155179 | 0.570139432 |
| ENSGALG00010002937 | GALNT7 | 422561 | 0.15973387 | 5.978782919 | 0.021681354 | 0.572581464 |
| ENSGALG00010026981 | NA | NA | -0.86087866 | -0.404443712 | 0.021745992 | 0.573303427 |
| ENSGALG00010021314 | ZP3 | 378906 | 0.559815224 | 1.183990264 | 0.021899319 | 0.575435401 |
| ENSGALG00010029636 | SPATA20 | 422100 | 0.241976723 | 4.825671552 | 0.021901738 | 0.575435401 |
| ENSGALG00010025658 | PDE1B | 777528 | 0.23877192 | 4.059671041 | 0.022053457 | 0.576974378 |
| ENSGALG00010025575 | TRPM2 | 424835 | -0.425586925 | 2.872900921 | 0.02209723 | 0.576974378 |
| ENSGALG00010025868 | NA | NA | -0.472422887 | 2.41710139 | 0.022118778 | 0.576974378 |
| ENSGALG00010015141 | PFDN4 | 419337 | -0.176073017 | 7.239890296 | 0.022123443 | 0.576974378 |
| ENSGALG00010003248 | MRI1 | 107049831 | 0.24412121 | 6.778441078 | 0.022164642 | 0.576974378 |
| ENSGALG00010010172 | NA | NA | -0.968602614 | -0.201877765 | 0.022245149 | 0.576974378 |
| ENSGALG00010010255 | CACNB4 | 374259 | -0.200705469 | 4.723859473 | 0.022288973 | 0.576974378 |
| ENSGALG00010020696 | MOB1A | 429657 | -0.448700424 | 2.975886326 | 0.022292919 | 0.576974378 |
| ENSGALG00010018169 | NA | NA | -0.3727604 | 2.930403475 | 0.022300402 | 0.576974378 |
| ENSGALG00010012306 | LOC121108207 | 121108207 | -0.750891316 | 4.103604891 | 0.022420414 | 0.576974378 |
| ENSGALG00010004892 | HEY2 | 421718 | 0.263223817 | 3.716719547 | 0.022432572 | 0.576974378 |
| ENSGALG00010029356 | LOC124417312 | 124417312 | 0.334615813 | 2.930356257 | 0.022439618 | 0.576974378 |
| ENSGALG00010019115 | DBNL | 425994 | 0.174922548 | 6.206853132 | 0.02244832 | 0.576974378 |
| ENSGALG00010019072 | AEBP1 | 100859431 | 0.262058863 | 5.443279119 | 0.02249844 | 0.577297187 |
| ENSGALG00010022849 | NA | NA | 0.51340761 | 3.615162311 | 0.022553406 | 0.577743086 |
| ENSGALG00010003845 | CFI | 428773 | -0.312895397 | 3.696989341 | 0.02260912 | 0.57806344 |
| ENSGALG00010005068 | PLCXD2 | 418417 | -0.367464377 | 2.5155652 | 0.022641132 | 0.57806344 |
| ENSGALG00010027427 | UBAC1 | 427754 | 0.176486729 | 7.043457922 | 0.022758188 | 0.578702918 |
| ENSGALG00010017595 | CCNF | 416749 | 0.176084418 | 6.928170221 | 0.022770184 | 0.578702918 |
| ENSGALG00010016496 | PLP1 | 396214 | 0.357397309 | 2.594737634 | 0.022838501 | 0.578702918 |
| ENSGALG00010029547 | HIP1 | 770683 | 0.199591246 | 6.111147623 | 0.02284861 | 0.578702918 |
| ENSGALG00010020277 | IDH2 | 431056 | 0.172173254 | 6.316289571 | 0.02289051 | 0.578702918 |
| ENSGALG00010029726 | SEPTIN9 | 417347 | 0.163199799 | 6.367124099 | 0.022892087 | 0.578702918 |
| ENSGALG00010015059 | ALG3 | 424952 | 0.241176671 | 4.318860769 | 0.023220708 | 0.585514778 |
| ENSGALG00010025186 | TMEM120B | 416847 | 0.20026727 | 5.032481939 | 0.023316497 | 0.585514778 |
| ENSGALG00010019882 | NA | NA | -0.698991492 | 1.045117684 | 0.023316968 | 0.585514778 |
| ENSGALG00010028642 | APH1A | 431243 | 0.199058779 | 5.154698316 | 0.023343163 | 0.585514778 |
| ENSGALG00010028088 | APOA1BP | 426569 | 0.201383295 | 5.798102403 | 0.023395733 | 0.585514778 |
| ENSGALG00010009286 | MPV17 | 100857465 | 0.260731091 | 4.01218431 | 0.023416526 | 0.585514778 |
| ENSGALG00010020664 | ESYT3 | 424271 | -0.322685718 | 2.990352979 | 0.023481138 | 0.585514778 |
| ENSGALG00010028129 | CYB561D2 | 100857844 | 0.288106871 | 3.459642715 | 0.023488268 | 0.585514778 |
| ENSGALG00010001631 | NA | NA | -0.223582495 | 4.430667702 | 0.023504399 | 0.585514778 |
| ENSGALG00010016289 | IRF1 | 396384 | -0.478573577 | 1.372472322 | 0.023547798 | 0.585646704 |
| ENSGALG00010016805 | FOXI3 | 428801 | -0.163009486 | 5.667315239 | 0.023613176 | 0.585846798 |
| ENSGALG00010005913 | NFKBIA | 396093 | -0.300708876 | 3.690942414 | 0.023657988 | 0.585846798 |
| ENSGALG00010000163 | NA | NA | -0.918886939 | 0.753012585 | 0.023670193 | 0.585846798 |
| ENSGALG00010023118 | LRRC27 | 426464 | 0.515558885 | 1.443722161 | 0.023786837 | 0.587673638 |
| ENSGALG00010026917 | AMT | 395566 | 0.210399285 | 4.69605648 | 0.023820473 | 0.587673638 |
| ENSGALG00010003692 | ZNFY2 | 107057099 | 0.799486298 | 0.264260977 | 0.023862632 | 0.587770268 |
| ENSGALG00010000228 | NA | NA | 0.753922839 | 0.207724776 | 0.024034726 | 0.588740128 |
| ENSGALG00010015804 | CKB | 396248 | 0.20165699 | 7.300301934 | 0.024091935 | 0.588740128 |

| | | | | | | |
|---|---|---|---|---|---|---|
| ENSGALG00010024696 | TRADD | 415700 | 0.304366017 | 3.444462109 | 0.024108906 | 0.588740128 |
| ENSGALG00010025212 | P2RX4 | 374166 | 0.290961019 | 3.30535463 | 0.024110416 | 0.588740128 |
| ENSGALG00010014214 | REEP2 | 771891 | -0.190531047 | 4.883891352 | 0.024120623 | 0.588740128 |
| ENSGALG00010015033 | BMP7 | 395494 | 0.230759854 | 4.459554148 | 0.024131833 | 0.588740128 |
| ENSGALG00010012667 | SCAMP1 | 416368 | 0.168098393 | 6.110928486 | 0.024259808 | 0.59043588 |
| ENSGALG00010002703 | PIN1 | 100859153 | -0.864948848 | 1.175804185 | 0.02427817 | 0.59043588 |
| ENSGALG00010017702 | ECEL1 | 424937 | 0.321741854 | 3.513862309 | 0.02446732 | 0.593492235 |
| ENSGALG00010013808 | HRASLS | 424903 | 0.404592235 | 2.888786904 | 0.024481072 | 0.593492235 |
| ENSGALG00010024878 | RHPN2 | 415771 | -0.19606199 | 5.678513714 | 0.024529996 | 0.593741788 |
| ENSGALG00010005789 | RNASET2 | 421569 | 0.168540182 | 5.634113292 | 0.024675917 | 0.595722802 |
| ENSGALG00010017497 | NA | NA | -0.688548259 | 1.634803179 | 0.024689358 | 0.595722802 |
| ENSGALG00010014170 | CEMIP2 | 427250 | -0.167883479 | 7.405045276 | 0.024789575 | 0.596291539 |
| ENSGALG00010020007 | NA | NA | -0.166382298 | 6.709887729 | 0.02479052 | 0.596291539 |
| ENSGALG00010019822 | NA | NA | -0.785064056 | -0.295664658 | 0.024878007 | 0.597460896 |
| ENSGALG00010023116 | PROC | 395085 | -0.579940456 | 0.756708979 | 0.024976114 | 0.598881242 |
| ENSGALG00010019422 | IMMP1L | 771365 | -0.24979754 | 4.06577826 | 0.02502281 | 0.599066337 |
| ENSGALG00010027745 | BICD2 | 415990 | 0.176194043 | 6.114133914 | 0.025106076 | 0.600125017 |
| ENSGALG00010027980 | NUP210 | 415977 | -0.156578844 | 6.43252376 | 0.025180934 | 0.600979745 |
| ENSGALG00010014029 | NA | NA | -0.253765495 | 3.925024935 | 0.025268078 | 0.601283992 |
| ENSGALG00010013126 | NA | NA | -0.862909457 | 6.067772101 | 0.025271923 | 0.601283992 |
| ENSGALG00010028339 | BCAN | 425976 | 0.631182479 | 0.700813922 | 0.025432237 | 0.602442243 |
| ENSGALG00010028867 | REXO4 | 417153 | 0.197787611 | 4.963412556 | 0.025455089 | 0.602442243 |
| ENSGALG00010015422 | DGKK | 422129 | 0.409547619 | 2.656713341 | 0.025461059 | 0.602442243 |
| ENSGALG00010023685 | NA | NA | -0.78486562 | -0.286578802 | 0.025518997 | 0.602442243 |
| ENSGALG00010017421 | NA | NA | 0.309417528 | 3.723012336 | 0.025548199 | 0.602442243 |
| ENSGALG00010007446 | NA | NA | -0.451647149 | 3.414554388 | 0.02555578 | 0.602442243 |
| ENSGALG00010018219 | SLC24A5 | 415431 | -0.729382729 | 0.014723038 | 0.025689204 | 0.604660134 |
| ENSGALG00010029146 | NEK6 | 417101 | 0.187382959 | 5.045469875 | 0.025829695 | 0.60703733 |
| ENSGALG00010011954 | LOC770141 | 770141 | -0.596570139 | 0.771190432 | 0.0259279 | 0.608415002 |
| ENSGALG00010014608 | NA | NA | -0.744769517 | 0.111746294 | 0.026138522 | 0.612422391 |
| ENSGALG00010010688 | IQGAP2 | 427211 | 0.166318902 | 6.179079595 | 0.026292441 | 0.615091053 |
| ENSGALG00010026711 | NA | NA | -0.454911484 | 2.025045105 | 0.026332815 | 0.615099346 |
| ENSGALG00010018931 | NA | NA | -0.179297758 | 5.318832645 | 0.026422911 | 0.616267286 |
| ENSGALG00010024644 | EIF6 | 419140 | 0.1675849 | 7.836765598 | 0.026521412 | 0.616443783 |
| ENSGALG00010014908 | PPP4R1L | 419319 | -0.154906294 | 6.332035621 | 0.026568601 | 0.616443783 |
| ENSGALG00010029186 | ZNF302 | 417303 | -0.273130522 | 3.555727727 | 0.026577472 | 0.616443783 |
| ENSGALG00010013286 | RNF14 | 416340 | 0.179618384 | 6.32063467 | 0.026590906 | 0.616443783 |
| ENSGALG00010018582 | RCC2 | 419361 | 0.163131906 | 7.323982763 | 0.026992169 | 0.624803671 |
| ENSGALG00010002324 | SNTG2 | 421913 | -0.862147248 | -0.354797578 | 0.027138854 | 0.627254422 |
| ENSGALG00010014705 | ARAP2 | 428790 | -0.272133183 | 3.488132089 | 0.027267997 | 0.629292974 |
| ENSGALG00010011450 | PTDSS1 | 428374 | -0.196222324 | 5.329323358 | 0.027343392 | 0.630086854 |
| ENSGALG00010022760 | ELAVL4 | 395634 | -0.165689411 | 8.374201841 | 0.027500493 | 0.632472694 |
| ENSGALG00010018231 | SMYD5 | 422951 | 0.165445962 | 6.330661274 | 0.027529228 | 0.632472694 |
| ENSGALG00010014243 | HIGD2A | 416227 | 0.399536261 | 3.360950296 | 0.027725743 | 0.635299521 |
| ENSGALG00010010384 | NA | NA | 0.187908049 | 5.275937237 | 0.027734937 | 0.635299521 |
| ENSGALG00010000402 | PFKFB1 | 415906 | 0.541169087 | 1.754401737 | 0.027852477 | 0.636379554 |
| ENSGALG00010011019 | NECAB1 | 420219 | 0.693071333 | 0.843228871 | 0.027864895 | 0.636379554 |
| ENSGALG00010023487 | GRN | 101748909 | 0.184570091 | 5.65704559 | 0.0280375 | 0.639371482 |
| ENSGALG00010000031 | NA | NA | -1.398533731 | 0.70054945 | 0.02813176 | 0.640570585 |
| ENSGALG00010029558 | NA | NA | -0.839348677 | -0.574846798 | 0.028414433 | 0.644343411 |
| ENSGALG00010009554 | ABCB10 | 421537 | -0.185297556 | 5.142596328 | 0.028420281 | 0.644343411 |
| ENSGALG00010014985 | LOC100857197 | 100857197 | 0.270021275 | 3.812282374 | 0.028458392 | 0.644343411 |
| ENSGALG00010018958 | PELI3 | 421279 | -0.172988471 | 5.134135629 | 0.028476833 | 0.644343411 |
| ENSGALG00010003468 | ILVBL | 101747528 | 0.235779123 | 4.889401477 | 0.028513264 | 0.644343411 |
| ENSGALG00010010651 | NUAK1 | 418074 | 0.187924262 | 5.267199462 | 0.028548983 | 0.644343411 |
| ENSGALG00010021367 | GRIP2 | 416043 | -0.392472065 | 2.747784837 | 0.028603283 | 0.64462238 |
| ENSGALG00010022264 | SCN8A | 426869 | 0.479509359 | 1.490628313 | 0.028683262 | 0.645032734 |

| | | | | | | |
|---|---|---|---|---|---|---|
| ENSGALG00010005814 | UPF2 | 419100 | 0.159253913 | 8.170306567 | 0.028705425 | 0.645032734 |
| ENSGALG00010016832 | DYNC1H1 | 423461 | -0.195299901 | 8.626238352 | 0.028779493 | 0.645753011 |
| ENSGALG00010023522 | EMG1 | 418292 | -0.511588189 | 1.146451256 | 0.028906088 | 0.647185624 |
| ENSGALG00010009795 | NDUFA6 | 427897 | 0.213130315 | 4.950593751 | 0.028927555 | 0.647185624 |
| ENSGALG00010021479 | SKAP1 | 101752206 | 0.675465038 | 1.267158979 | 0.0290595 | 0.649192607 |
| ENSGALG00010010843 | ZNF395 | 422017 | 0.189515049 | 6.134417546 | 0.029200706 | 0.651400362 |
| ENSGALG00010016647 | EGR1 | 373931 | -0.483688146 | 2.852169038 | 0.0292976 | 0.651659969 |
| ENSGALG00010012066 | STXBP3 | 424341 | -0.183943864 | 5.606161434 | 0.029379498 | 0.651659969 |
| ENSGALG00010006580 | ALDH6A1 | 423345 | 0.223357432 | 4.20720156 | 0.029437939 | 0.651659969 |
| ENSGALG00010000898 | NA | NA | -0.50617657 | 2.012336631 | 0.029503095 | 0.651659969 |
| ENSGALG00010027738 | NA | NA | 0.717822501 | -0.034905387 | 0.029504156 | 0.651659969 |
| ENSGALG00010014487 | OSTM1 | 421773 | 0.195430238 | 5.146792368 | 0.029504529 | 0.651659969 |
| ENSGALG00010001878 | XK | 427991 | -0.307658547 | 3.461484143 | 0.029546003 | 0.651659969 |
| ENSGALG00010002987 | PCDH8 | 428065 | 0.351559655 | 2.717642532 | 0.029551529 | 0.651659969 |
| ENSGALG00010008101 | CENPQ | 422048 | 0.205964501 | 4.969912992 | 0.029596001 | 0.651705632 |
| ENSGALG00010007921 | FAM120B | 421552 | 0.250339838 | 4.015369171 | 0.029705856 | 0.653188845 |
| ENSGALG00010011457 | SYCP3 | 418097 | -0.378498036 | 2.036501057 | 0.029757637 | 0.653392686 |
| ENSGALG00010027312 | LOC120017336 | 120017336 | -0.755655103 | -0.311517265 | 0.029895365 | 0.655480389 |
| ENSGALG00010008333 | APC | 415607 | -0.168824799 | 7.800593784 | 0.030006878 | 0.656307071 |
| ENSGALG00010007723 | LOC112530788 | 112530788 | -0.589772352 | 0.645864034 | 0.030047483 | 0.656307071 |
| ENSGALG00010003762 | EVI5L | 430998 | 0.263329847 | 4.492884881 | 0.03006117 | 0.656307071 |
| ENSGALG00010024204 | PTBP2 | 424479 | -0.189036756 | 7.076003265 | 0.030362347 | 0.661771323 |
| ENSGALG00010026878 | LOC112532592 | 112532592 | -0.74027601 | 0.087291689 | 0.030398526 | 0.661771323 |
| ENSGALG00010023890 | SASS6 | 424471 | -0.156050833 | 5.946891113 | 0.030480346 | 0.661771323 |
| ENSGALG00010006903 | B3GLCT | 428075 | 0.18366125 | 5.597742438 | 0.030483676 | 0.661771323 |
| ENSGALG00010004701 | NA | NA | -0.470554789 | 2.913890326 | 0.030568636 | 0.662679743 |
| ENSGALG00010004389 | ZNF71 | 425244 | 0.294642795 | 3.518848622 | 0.030696276 | 0.663420564 |
| ENSGALG00010003473 | LOC112531056 | 112531056 | 0.149941783 | 5.950915417 | 0.0307164 | 0.663420564 |
| ENSGALG00010000232 | TMEM147 | 112531206 | 0.305858744 | 4.375807256 | 0.030732299 | 0.663420564 |
| ENSGALG00010006120 | MOGS | 107049491 | 0.197492494 | 4.656232126 | 0.030860306 | 0.663764629 |
| ENSGALG00010010072 | NA | NA | -0.732251474 | -0.125538594 | 0.030927943 | 0.663764629 |
| ENSGALG00010014220 | C9orf85 | 770483 | -0.164715581 | 5.835782212 | 0.030961256 | 0.663764629 |
| ENSGALG00010020617 | DGUOK | 107054975 | 0.247537023 | 3.910128831 | 0.031068818 | 0.663764629 |
| ENSGALG00010000357 | NA | NA | -0.608935224 | 2.302811053 | 0.031091468 | 0.663764629 |
| ENSGALG00010005829 | NPAS3 | 423315 | -0.339198304 | 2.659534532 | 0.031138768 | 0.663764629 |
| ENSGALG00010020843 | HMG20A | 415346 | 0.156008027 | 7.220861651 | 0.031144551 | 0.663764629 |
| ENSGALG00010025298 | NACAD | 420409 | 0.238787759 | 5.285474215 | 0.031144583 | 0.663764629 |
| ENSGALG00010023625 | NA | NA | -0.285699317 | 3.529773429 | 0.031223513 | 0.663764629 |
| ENSGALG00010029111 | NA | NA | 0.922205882 | -0.175764953 | 0.031241204 | 0.663764629 |
| ENSGALG00010015027 | CLSPN | 428223 | -0.155632941 | 6.015116289 | 0.031287645 | 0.663764629 |
| ENSGALG00010025583 | NA | NA | -0.425644403 | 1.755153275 | 0.031316259 | 0.663764629 |
| ENSGALG00010012130 | TMEM67 | 420224 | 0.267509963 | 3.67396573 | 0.031342797 | 0.663764629 |
| ENSGALG00010020293 | SEMA4B | 415582 | 0.265056593 | 3.619024706 | 0.031393911 | 0.663764629 |
| ENSGALG00010020626 | MORN4 | 425290 | 0.306852755 | 3.24332219 | 0.031396024 | 0.663764629 |
| ENSGALG00010013343 | KIAA1644 | 100857276 | 0.236893975 | 4.03454874 | 0.031490657 | 0.664128767 |
| ENSGALG00010017604 | GATM | 395504 | 0.362820253 | 2.46683874 | 0.031499666 | 0.664128767 |
| ENSGALG00010005926 | NUDT5 | 426368 | 0.311253873 | 3.199731871 | 0.031743757 | 0.668358282 |
| ENSGALG00010018668 | OVOL3 | 416720 | 0.284370975 | 3.480597176 | 0.031823247 | 0.669115336 |
| ENSGALG00010024571 | KCNA3 | 404303 | 0.204342376 | 4.383720093 | 0.031894145 | 0.669532463 |
| ENSGALG00010001969 | PLEKHA8 | 420637 | -0.247612653 | 4.913719753 | 0.031949574 | 0.669532463 |
| ENSGALG00010006463 | SCO2 | 429671 | 0.241875691 | 3.92947938 | 0.032089996 | 0.669532463 |
| ENSGALG00010005477 | TMEM161B | 426919 | -0.169145095 | 5.938239014 | 0.032099428 | 0.669532463 |
| ENSGALG00010024788 | SIK3 | 419764 | -0.186440282 | 5.309902217 | 0.032102601 | 0.669532463 |
| ENSGALG00010021452 | SLC25A1 | 416764 | 0.284086316 | 3.418953812 | 0.032104452 | 0.669532463 |
| ENSGALG00010022815 | LRRC8D | 424513 | -0.179268429 | 5.878907015 | 0.03246536 | 0.675618682 |
| ENSGALG00010019432 | NA | NA | -0.90713998 | -0.664907623 | 0.032484203 | 0.675618682 |
| ENSGALG00010001318 | NA | NA | -0.768525353 | 0.264122857 | 0.032537327 | 0.675809078 |

| | | | | | | |
|---|---|---|---|---|---|---|
| ENSGALG00010017528 | RAI1 | 427664 | 0.190197856 | 5.157011346 | 0.032692505 | 0.678103627 |
| ENSGALG00010019088 | NA | NA | -0.663677952 | 0.133072111 | 0.032736037 | 0.678103627 |
| ENSGALG00010010983 | MED28 | 425350 | 0.227220152 | 4.802261784 | 0.032787875 | 0.678263302 |
| ENSGALG00010011437 | DMRT1 | 769693 | 0.166675167 | 7.295427894 | 0.032940909 | 0.680513135 |
| ENSGALG00010025204 | TRAF6 | 423163 | -0.175264227 | 5.23715759 | 0.033085397 | 0.682580598 |
| ENSGALG00010012988 | RPL18A | 417823 | 0.204419381 | 8.952068499 | 0.033189989 | 0.682969008 |
| ENSGALG00010029066 | LOC121107075 | 121107075 | -0.162701568 | 6.660384584 | 0.033198298 | 0.682969008 |
| ENSGALG00010026229 | CRHR2 | 395105 | 0.353187273 | 2.440099312 | 0.033237529 | 0.682969008 |
| ENSGALG00010022382 | CALB2 | 396255 | -0.442206415 | 1.734976156 | 0.033291821 | 0.683171276 |
| ENSGALG00010024866 | TRAPPC4 | 419788 | 0.19684853 | 4.751229934 | 0.033380454 | 0.683856017 |
| ENSGALG00010002211 | ACER3 | 419087 | 0.158224715 | 6.703032825 | 0.033414175 | 0.683856017 |
| ENSGALG00010002910 | NA | NA | -0.443437126 | 1.954304385 | 0.033473189 | 0.684152807 |
| ENSGALG00010028174 | GMPPB | 415924 | 0.261245596 | 3.714610856 | 0.033632515 | 0.686496362 |
| ENSGALG00010022194 | LOC776275 | 776275 | -0.514659843 | 1.303048985 | 0.033825643 | 0.688468206 |
| ENSGALG00010022248 | FIGNL2 | 431296 | 0.191590034 | 5.311171023 | 0.033858734 | 0.688468206 |
| ENSGALG00010008368 | SNCA | 395393 | 0.343922141 | 3.327612488 | 0.033904705 | 0.688468206 |
| ENSGALG00010013206 | AFF2 | 422394 | 0.22075467 | 4.380437512 | 0.033908291 | 0.688468206 |
| ENSGALG00010028856 | ETV3 | 426718 | 0.18093669 | 6.659335183 | 0.033953928 | 0.688485328 |
| ENSGALG00010017358 | NA | NA | -0.794827187 | -0.468670917 | 0.034040527 | 0.689331894 |
| ENSGALG00010012824 | HIBCH | 423979 | 0.179787324 | 5.659793234 | 0.034118709 | 0.689810226 |
| ENSGALG00010020236 | SLC35G1 | 423800 | -0.29264583 | 3.0263897 | 0.034182241 | 0.689810226 |
| ENSGALG00010013084 | SGSM3 | 418008 | 0.217197899 | 4.213737216 | 0.03420749 | 0.689810226 |
| ENSGALG00010023572 | MYBL2 | 396258 | -0.1489329 | 6.426776485 | 0.03424367 | 0.689810226 |
| ENSGALG00010009560 | RAB24 | 427627 | 0.188260009 | 5.105535584 | 0.034500364 | 0.693455393 |
| ENSGALG00010004477 | TULP4 | 421658 | -0.17124299 | 7.726580987 | 0.034545008 | 0.693455393 |
| ENSGALG00010019310 | CRELD1 | 100858613 | 0.212897691 | 4.679835087 | 0.034559976 | 0.693455393 |
| ENSGALG00010018606 | DZANK1 | 416722 | -0.363069802 | 2.451178998 | 0.034640266 | 0.694160217 |
| ENSGALG00010022707 | SALL1 | 395446 | 0.212010569 | 7.215389604 | 0.034755871 | 0.694818245 |
| ENSGALG00010006107 | PGF | 771832 | -0.41103713 | 2.306316045 | 0.034796623 | 0.694818245 |
| ENSGALG00010019893 | LOC112532451 | 112532451 | -0.744403078 | -0.062191327 | 0.034808721 | 0.694818245 |
| ENSGALG00010025933 | SKIV2L | 421171 | 0.180498613 | 6.080756767 | 0.034943275 | 0.695985861 |
| ENSGALG00010003723 | NA | NA | -0.283584355 | 3.431322434 | 0.03495778 | 0.695985861 |
| ENSGALG00010024816 | NFIA | 396210 | 0.343315553 | 2.889447296 | 0.035097446 | 0.697862543 |
| ENSGALG00010005463 | SKA3 | 418948 | -0.148757931 | 6.457885472 | 0.035334315 | 0.700949118 |
| ENSGALG00010028188 | SMAD4 | 107055136 | 0.176826385 | 5.736492021 | 0.035343889 | 0.700949118 |
| ENSGALG00010003862 | PCK2 | 396457 | 0.179122949 | 5.83986169 | 0.03542842 | 0.701720126 |
| ENSGALG00010014148 | KCNG1 | 419347 | 0.173454153 | 6.250319809 | 0.035475871 | 0.701755645 |
| ENSGALG00010015013 | SNX18 | 768969 | -0.177479918 | 5.065364336 | 0.035629149 | 0.703881775 |
| ENSGALG00010021301 | SMURF1 | 416487 | -0.155094234 | 5.74742147 | 0.035756105 | 0.705483108 |
| ENSGALG00010012638 | NA | NA | 0.199968035 | 5.050770199 | 0.035848805 | 0.706405307 |
| ENSGALG00010003131 | TAP1 | 427727 | 0.231507101 | 4.263466757 | 0.036199209 | 0.711610344 |
| ENSGALG00010001697 | TAF10 | 107052460 | 0.23462113 | 4.205309212 | 0.036205549 | 0.711610344 |
| ENSGALG00010011469 | CPS1 | 428994 | -0.233205515 | 3.880129795 | 0.03634861 | 0.713272289 |
| ENSGALG00010005407 | DPH6 | 423294 | -0.218112825 | 4.270076251 | 0.036412381 | 0.713272289 |
| ENSGALG00010025917 | SCUBE3 | 419896 | 0.365700798 | 2.263985281 | 0.036429326 | 0.713272289 |
| ENSGALG00010028442 | CHTOP | 426357 | 0.153576256 | 7.430787032 | 0.036595448 | 0.715613287 |
| ENSGALG00010017467 | SFRP1 | 395237 | 0.156287954 | 6.372762562 | 0.036660436 | 0.71567716 |
| ENSGALG00010005875 | CDK8 | 428080 | 0.170513034 | 6.381370947 | 0.03669769 | 0.71567716 |
| ENSGALG00010006575 | SSBP2 | 770958 | -0.167781813 | 7.476991276 | 0.036779576 | 0.71567716 |
| ENSGALG00010022451 | CHP1 | 423211 | 0.149178098 | 5.806333709 | 0.036796698 | 0.71567716 |
| ENSGALG00010024194 | NKAIN4 | 419240 | 0.234737005 | 4.469861542 | 0.036831531 | 0.71567716 |
| ENSGALG00010021256 | PML | 415302 | 0.296775071 | 3.398409434 | 0.036881916 | 0.715751319 |
| ENSGALG00010008311 | SRP19 | 770061 | -0.179968339 | 5.485987912 | 0.037097991 | 0.718336507 |
| ENSGALG00010024205 | ANKRD13D | 107049270 | 0.373452407 | 2.678570745 | 0.037319219 | 0.718336507 |
| ENSGALG00010006045 | NA | NA | -0.483349283 | 1.543515592 | 0.037332026 | 0.718336507 |
| ENSGALG00010021470 | NA | NA | 0.352775932 | 2.866745183 | 0.037348365 | 0.718336507 |
| ENSGALG00010017616 | PDCD7 | 415528 | 0.200397296 | 4.681381645 | 0.03741683 | 0.718336507 |

| | | | | | | |
|---|---|---|---|---|---|---|
| ENSGALG00010009002 | RNF19AL | 421988 | 0.226026079 | 4.01785907 | 0.037448744 | 0.718336507 |
| ENSGALG00010014427 | TMEM267 | 427193 | 0.193096653 | 4.707235056 | 0.037451885 | 0.718336507 |
| ENSGALG00010020140 | SECISBP2L | 415435 | -0.178955864 | 5.163967933 | 0.037462624 | 0.718336507 |
| ENSGALG00010028485 | USF1 | 425126 | 0.152851721 | 6.074068365 | 0.037462652 | 0.718336507 |
| ENSGALG00010017388 | PROSC | 426770 | 0.169653969 | 5.034533941 | 0.03751807 | 0.718336507 |
| ENSGALG00010020013 | ADAM8 | 423672 | 0.775133043 | -0.081892497 | 0.037548648 | 0.718336507 |
| ENSGALG00010027132 | NA | NA | -0.225912082 | 4.194289543 | 0.037612329 | 0.718336507 |
| ENSGALG00010027543 | TUBB4B | 417255 | 0.181125857 | 8.474515783 | 0.037786054 | 0.718336507 |
| ENSGALG00010001995 | LRRC1 | 421891 | 0.218951662 | 4.341797333 | 0.037825207 | 0.718336507 |
| ENSGALG00010006850 | C5H14orf4 | 429422 | 0.165389131 | 6.741428674 | 0.037856479 | 0.718336507 |
| ENSGALG00010002550 | NA | NA | -0.662321328 | 0.182679315 | 0.037856928 | 0.718336507 |
| ENSGALG00010008256 | NA | NA | -0.720183113 | -0.071853656 | 0.037866146 | 0.718336507 |
| ENSGALG00010010810 | GALT | 431509 | 0.303312846 | 3.741914143 | 0.037936149 | 0.718336507 |
| ENSGALG00010024224 | NA | NA | -0.598714353 | 0.510605979 | 0.037954481 | 0.718336507 |
| ENSGALG00010013173 | KCNJ8 | 395973 | 0.423848419 | 1.982395603 | 0.03798903 | 0.718336507 |
| ENSGALG00010021115 | ALPL | 396317 | 0.173294218 | 5.319999949 | 0.038082636 | 0.718336507 |
| ENSGALG00010018009 | LOC107057413 | 107057413 | 1.02579514 | 0.420467072 | 0.038110478 | 0.718336507 |
| ENSGALG00010005587 | LEF1 | 395865 | 0.179696318 | 4.865033864 | 0.038169011 | 0.718336507 |
| ENSGALG00010007069 | NA | NA | -0.988676243 | -0.763845421 | 0.038169266 | 0.718336507 |
| ENSGALG00010004869 | NA | NA | -0.499890906 | 1.352561715 | 0.038214933 | 0.718336507 |
| ENSGALG00010028003 | NA | NA | -0.162473569 | 7.204730088 | 0.038352754 | 0.718336507 |
| ENSGALG00010020265 | SPTBN5 | 423225 | -0.745819969 | -0.152401164 | 0.038421991 | 0.718336507 |
| ENSGALG00010026950 | LHFPL5 | 378916 | 0.387770939 | 2.137311271 | 0.038422649 | 0.718336507 |
| ENSGALG00010023173 | LOC107049856 | 107049856 | -0.808645349 | 0.916341517 | 0.038438451 | 0.718336507 |
| ENSGALG00010018105 | NA | NA | 0.345801695 | 2.385516512 | 0.038455781 | 0.718336507 |
| ENSGALG00010022269 | CNTD1 | 420019 | -0.188027604 | 5.096397611 | 0.038522977 | 0.718336507 |
| ENSGALG00010028073 | ST6GALNAC4 | 395182 | 0.277962472 | 3.602147629 | 0.038532697 | 0.718336507 |
| ENSGALG00010029553 | MYH1C | 417310 | -0.570829502 | 0.608932838 | 0.038569517 | 0.718336507 |
| ENSGALG00010021413 | NA | NA | 0.256856577 | 4.016888358 | 0.038649912 | 0.718336507 |
| ENSGALG00010020548 | CMTR1 | 421434 | -0.15418397 | 6.492877058 | 0.038650897 | 0.718336507 |
| ENSGALG00010012326 | SGTB | 427166 | 0.196595862 | 4.467996304 | 0.038817595 | 0.718788341 |
| ENSGALG00010004262 | NA | NA | -0.293518605 | 3.789338417 | 0.038848206 | 0.718788341 |
| ENSGALG00010004375 | NA | NA | -0.537149921 | 1.466203543 | 0.038891431 | 0.718788341 |
| ENSGALG00010002827 | SBSPON | 420181 | 0.813632151 | -0.188760668 | 0.038914893 | 0.718788341 |
| ENSGALG00010022222 | MZT2B | 768838 | 0.322194198 | 2.593512166 | 0.038981673 | 0.718788341 |
| ENSGALG00010019049 | POLD2 | 107057377 | 0.153620399 | 6.119472472 | 0.039030023 | 0.718788341 |
| ENSGALG00010028109 | TJP3 | 420070 | 0.289598286 | 3.547215513 | 0.039048544 | 0.718788341 |
| ENSGALG00010020676 | CYP2C23a | 414746 | -0.720648534 | -0.16713202 | 0.039049334 | 0.718788341 |
| ENSGALG00010019140 | NA | NA | -0.44909195 | 1.327874718 | 0.039101953 | 0.718895951 |
| ENSGALG00010000078 | LOC112531364 | 112531364 | -0.307143936 | 4.343170669 | 0.039170088 | 0.719288242 |
| ENSGALG00010011681 | MAP1B | 396174 | 0.156108804 | 6.691654549 | 0.039411504 | 0.722280971 |
| ENSGALG00010007026 | NA | NA | 0.400569581 | 1.852884643 | 0.039427048 | 0.722280971 |
| ENSGALG00010016016 | ZDHHC2 | 422734 | -0.152947328 | 5.607076908 | 0.039519358 | 0.723110163 |
| ENSGALG00010000896 | NA | NA | -0.378752853 | 2.006461842 | 0.039571091 | 0.723169737 |
| ENSGALG00010002603 | FAM196A | 101748342 | -0.266005851 | 3.673801975 | 0.039656194 | 0.723169737 |
| ENSGALG00010027229 | MDM4 | 419823 | -0.213144837 | 5.592704201 | 0.039663766 | 0.723169737 |
| ENSGALG00010002186 | ICK | 421896 | -0.143012333 | 5.991076943 | 0.039786499 | 0.723912946 |
| ENSGALG00010027751 | ANGPTL4 | 769087 | 0.215049407 | 4.734151335 | 0.03982989 | 0.723912946 |
| ENSGALG00010019390 | TVP23A | 771988 | -0.44338937 | 1.341689385 | 0.03989512 | 0.723912946 |
| ENSGALG00010016791 | LRRTM1 | 771502 | -0.606072757 | 1.982791234 | 0.039952002 | 0.723912946 |
| ENSGALG00010008190 | TSNARE1 | 420304 | 0.208041252 | 4.837517924 | 0.039965673 | 0.723912946 |
| ENSGALG00010022202 | CX3CL1 | 415651 | -0.687210523 | -0.145799388 | 0.040045432 | 0.723912946 |
| ENSGALG00010003740 | PITX2 | 395862 | 0.413673251 | 2.232063726 | 0.040053505 | 0.723912946 |
| ENSGALG00010000099 | NA | NA | -0.993487583 | 0.692750174 | 0.040120065 | 0.723912946 |
| ENSGALG00010001351 | CAPN1 | 693249 | 0.167394753 | 5.21114641 | 0.040184874 | 0.723912946 |
| ENSGALG00010017688 | SPDYA | 101748777 | -0.250530115 | 4.224561769 | 0.040253053 | 0.723912946 |
| ENSGALG00010025589 | LOC121111680 | 121111680 | -0.263881682 | 3.534436038 | 0.040263494 | 0.723912946 |

| | | | | | | |
|---|---|---|---|---|---|---|
| ENSGALG00010005419 | NA | NA | 0.388713754 | 2.405826139 | 0.040276215 | 0.723912946 |
| ENSGALG00010021329 | PSMC3IP | 772119 | 0.181964708 | 5.029071914 | 0.040316817 | 0.723912946 |
| ENSGALG00010029069 | NA | NA | -0.868399116 | -0.498081451 | 0.040490513 | 0.726183417 |
| ENSGALG00010019820 | BBOX1 | 426932 | 0.20351681 | 5.712784536 | 0.040734048 | 0.728454843 |
| ENSGALG00010000870 | DACH1 | 373935 | 0.153677266 | 5.43685434 | 0.040783948 | 0.728454843 |
| ENSGALG00010008862 | ZNF367 | 100858851 | -0.164630422 | 5.351080024 | 0.040822835 | 0.728454843 |
| ENSGALG00010004131 | NA | NA | -0.658567803 | 0.166142553 | 0.040852931 | 0.728454843 |
| ENSGALG00010025640 | SLC38A11 | 424181 | 0.150849091 | 5.887783987 | 0.040854136 | 0.728454843 |
| ENSGALG00010005213 | NA | NA | 0.704231547 | -0.038531901 | 0.040913755 | 0.728672548 |
| ENSGALG00010023514 | ULK3 | 769780 | 0.190004573 | 4.711056428 | 0.041086235 | 0.729875124 |
| ENSGALG00010018696 | NA | NA | -0.236526016 | 4.51008536 | 0.041135139 | 0.729875124 |
| ENSGALG00010009326 | DLD | 417699 | -0.151431431 | 5.999540526 | 0.041230362 | 0.729875124 |
| ENSGALG00010029774 | RFLNB | 417623 | -0.394549658 | 2.008119904 | 0.041322697 | 0.729875124 |
| ENSGALG00010008161 | TMEFF1 | 420985 | 0.241590736 | 4.127099809 | 0.041348389 | 0.729875124 |
| ENSGALG00010023397 | LOC107049267 | 107049267 | 0.510559047 | 1.367744393 | 0.041355702 | 0.729875124 |
| ENSGALG00010002824 | NXPH1 | 420582 | 0.666474062 | 0.251090867 | 0.041358864 | 0.729875124 |
| ENSGALG00010022229 | RAPSN | 395986 | -0.605972328 | 0.7692269 | 0.041361173 | 0.729875124 |
| ENSGALG00010021287 | COL17A1 | 396503 | -0.319996554 | 2.745152215 | 0.041514595 | 0.730953677 |
| ENSGALG00010016754 | NA | NA | -0.303613479 | 3.474259914 | 0.041517408 | 0.730953677 |
| ENSGALG00010020324 | PLEKHH2 | 421399 | 0.260199389 | 4.233432697 | 0.041568645 | 0.731018383 |
| ENSGALG00010003271 | AKAP11 | 418835 | -0.148903455 | 5.889607986 | 0.041959428 | 0.734845332 |
| ENSGALG00010016812 | LRRC31 | 424994 | -0.161985254 | 6.114438051 | 0.041965324 | 0.734845332 |
| ENSGALG00010016992 | NA | NA | -0.450772583 | 1.372634309 | 0.041984446 | 0.734845332 |
| ENSGALG00010027271 | FAM212B | 419868 | -0.520736675 | 0.630477687 | 0.04200401 | 0.734845332 |
| ENSGALG00010000753 | TRIO | 420919 | -0.16190586 | 6.136951272 | 0.042025312 | 0.734845332 |
| ENSGALG00010011386 | CTSB | 396329 | 0.178542494 | 6.993655031 | 0.042204663 | 0.737142808 |
| ENSGALG00010019347 | NA | NA | 0.184515715 | 5.498173275 | 0.042335248 | 0.738584299 |
| ENSGALG00010020474 | NA | NA | -0.806835658 | -0.240919875 | 0.042476201 | 0.740203187 |
| ENSGALG00010007704 | NA | NA | -0.720100316 | -0.122413954 | 0.042606362 | 0.741262261 |
| ENSGALG00010022354 | FAR2 | 419043 | 0.227395012 | 4.00885569 | 0.042660432 | 0.741262261 |
| ENSGALG00010024611 | NA | NA | 0.23989135 | 4.345721966 | 0.042796238 | 0.741262261 |
| ENSGALG00010028322 | STRBP | 417105 | -0.167967945 | 7.172205213 | 0.042856504 | 0.741262261 |
| ENSGALG00010008381 | C18orf21 | 420954 | 0.16387672 | 5.058837601 | 0.042863206 | 0.741262261 |
| ENSGALG00010000822 | BASP1 | 373905 | 0.1527102 | 6.731626514 | 0.042868097 | 0.741262261 |
| ENSGALG00010016866 | ZNRF2 | 420678 | -0.180838542 | 4.741587532 | 0.042907914 | 0.741262261 |
| ENSGALG00010019334 | ATL1 | 423577 | 0.25688561 | 3.579221363 | 0.042922798 | 0.741262261 |
| ENSGALG00010012760 | PLEKHF2 | 420234 | -0.171543049 | 4.911624735 | 0.043080062 | 0.742354489 |
| ENSGALG00010007596 | LOC121109412 | 121109412 | -0.699193892 | 0.575096522 | 0.043087134 | 0.742354489 |
| ENSGALG00010018829 | PTPN4 | 424234 | 0.144940206 | 6.25897178 | 0.043131203 | 0.742354489 |
| ENSGALG00010024090 | LOC112532597 | 112532597 | -0.587688311 | 1.413919383 | 0.043194972 | 0.742354489 |
| ENSGALG00010025767 | NA | NA | -0.364369799 | 2.56607551 | 0.043253385 | 0.742354489 |
| ENSGALG00010027382 | PALM | 101750199 | 0.32169453 | 2.571760092 | 0.043275837 | 0.742354489 |
| ENSGALG00010026320 | GGT5 | 416943 | -0.38476397 | 1.791916066 | 0.043492812 | 0.745244722 |
| ENSGALG00010021492 | PARP6 | 769472 | 0.210376739 | 4.402952588 | 0.043577784 | 0.745869201 |
| ENSGALG00010027538 | NA | NA | 0.269095291 | 3.35566436 | 0.043662291 | 0.746102276 |
| ENSGALG00010020540 | PCGF5 | 423796 | -0.211028521 | 4.284013531 | 0.043729735 | 0.746102276 |
| ENSGALG00010024008 | CCNE1 | 426117 | -0.163414474 | 5.104922529 | 0.043799746 | 0.746102276 |
| ENSGALG00010001402 | NUP42 | 420614 | -0.182180824 | 5.054947526 | 0.043827601 | 0.746102276 |
| ENSGALG00010022956 | PBLD | 423685 | -0.237281367 | 3.664251812 | 0.043834907 | 0.746102276 |
| ENSGALG00010007980 | 1-Mar | 422420 | 0.183103382 | 5.25706435 | 0.043882658 | 0.746102276 |
| ENSGALG00010029416 | UNC119 | 771690 | 0.253741936 | 3.585021538 | 0.044045397 | 0.748041718 |
| ENSGALG00010004170 | NA | NA | -0.639078291 | -0.105730955 | 0.04417787 | 0.749463424 |
| ENSGALG00010020149 | FAHD2A | 426684 | 0.180623444 | 4.851797109 | 0.044281447 | 0.750392321 |
| ENSGALG00010005141 | CCDC170 | 421639 | -0.763984534 | -0.543169039 | 0.044432012 | 0.751533366 |
| ENSGALG00010023759 | CALCA | 396256 | -0.268660089 | 3.32073347 | 0.044506578 | 0.751533366 |
| ENSGALG00010012584 | NA | NA | -0.651691545 | 0.021897686 | 0.044560866 | 0.751533366 |
| ENSGALG00010003536 | NA | NA | -0.350081857 | 5.599885428 | 0.044602404 | 0.751533366 |

| | | | | | | |
|---|---|---|---|---|---|---|
| ENSGALG00010021721 | TP53I11 | 426399 | 0.153403743 | 8.088693532 | 0.04463412 | 0.751533366 |
| ENSGALG00010005695 | KIF5B | 420472 | -0.158136153 | 6.88317254 | 0.044654923 | 0.751533366 |
| ENSGALG00010003191 | NA | NA | 0.18612083 | 4.9132759 | 0.044728039 | 0.751533366 |
| ENSGALG00010027299 | PGPEP1 | 420127 | -0.176344512 | 6.403211112 | 0.04473995 | 0.751533366 |
| ENSGALG00010027594 | ITPR3 | 419910 | 0.375984572 | 2.420550066 | 0.044797442 | 0.751677608 |
| ENSGALG00010028877 | FAM163B | 100858326 | 0.424502287 | 1.506127368 | 0.044878943 | 0.752189216 |
| ENSGALG00010021425 | PTPN9 | 415308 | 0.15258788 | 5.836463181 | 0.044972214 | 0.752189216 |
| ENSGALG00010000050 | CIB1 | 770162 | 0.189042798 | 4.51938163 | 0.044985593 | 0.752189216 |
| ENSGALG00010028386 | NCAN | 395493 | 0.234933353 | 3.950925874 | 0.045041623 | 0.752189216 |
| ENSGALG00010026421 | NA | NA | -0.543083422 | 1.276997664 | 0.045101714 | 0.752189216 |
| ENSGALG00010004110 | NA | NA | -0.333015546 | 2.353248464 | 0.045121565 | 0.752189216 |
| ENSGALG00010029591 | SUZ12 | 417406 | -0.147920156 | 6.330518308 | 0.045525946 | 0.756780588 |
| ENSGALG00010008022 | SLC35F3 | 421518 | 0.18774085 | 4.977798414 | 0.045541019 | 0.756780588 |
| ENSGALG00010003479 | SYDE1 | 429892 | 0.309357407 | 2.629216851 | 0.0455447 | 0.756780588 |
| ENSGALG00010024110 | CPNE1 | 419134 | -0.161669894 | 5.607366999 | 0.045675777 | 0.757398159 |
| ENSGALG00010007762 | SH2D4A | 422487 | -0.568357214 | 0.509700363 | 0.045800173 | 0.757398159 |
| ENSGALG00010010232 | LOC107054313 | 107054313 | -0.181044503 | 5.279367571 | 0.045835603 | 0.757398159 |
| ENSGALG00010011235 | RPP40 | 420876 | 0.224256879 | 4.04208001 | 0.045868313 | 0.757398159 |
| ENSGALG00010016599 | LOC416263 | 416263 | 0.312144271 | 3.424047635 | 0.045953306 | 0.757398159 |
| ENSGALG00010013881 | NA | NA | -0.393790681 | 3.284500552 | 0.045955938 | 0.757398159 |
| ENSGALG00010009079 | KCTD15 | 415779 | 0.176099455 | 5.753680256 | 0.045973099 | 0.757398159 |
| ENSGALG00010020522 | NA | NA | -0.302413376 | 3.205400631 | 0.045976089 | 0.757398159 |
| ENSGALG00010002969 | NA | NA | -0.526966787 | 0.797667837 | 0.046074822 | 0.757556003 |
| ENSGALG00010008752 | GOLM1 | 427462 | 0.184313417 | 5.398968258 | 0.046084246 | 0.757556003 |
| ENSGALG00010021678 | DNAAF4 | 415409 | 0.322084133 | 2.904983621 | 0.046159447 | 0.757981511 |
| ENSGALG00010027397 | RFX5 | 100857912 | 0.278986193 | 3.195761874 | 0.046461463 | 0.761087681 |
| ENSGALG00010008474 | NA | NA | 0.204131005 | 5.282040963 | 0.046488991 | 0.761087681 |
| ENSGALG00010027362 | RPS27 | 107055140 | 0.227965415 | 7.638991678 | 0.046497159 | 0.761087681 |
| ENSGALG00010021972 | PISD | 416960 | -0.158437033 | 7.057518239 | 0.046579382 | 0.76128217 |
| ENSGALG00010005742 | GPR183 | 769212 | -0.523557479 | 0.703534028 | 0.046649673 | 0.76128217 |
| ENSGALG00010010769 | HEXB | 427204 | 0.164471654 | 5.023007301 | 0.046657632 | 0.76128217 |
| ENSGALG00010020155 | MFGE8 | 415494 | 0.280035949 | 4.615480522 | 0.046727494 | 0.761613561 |
| ENSGALG00010013645 | RRAGC | 419605 | -0.147446851 | 6.66653922 | 0.046817426 | 0.762271011 |
| ENSGALG00010024212 | RIMS4 | 428142 | -0.192889158 | 4.784996915 | 0.046895859 | 0.762740055 |
| ENSGALG00010024965 | STAT2 | 100858732 | 0.250100271 | 3.686747288 | 0.046952974 | 0.762861741 |
| ENSGALG00010003292 | GADD45GIP1 | 107055382 | 0.174440618 | 5.299314715 | 0.047251762 | 0.766905577 |
| ENSGALG00010022921 | INPP1 | 107049248 | 0.430587579 | 1.527696227 | 0.047382523 | 0.768216639 |
| ENSGALG00010024625 | SPRYD3 | 100858430 | 0.17752895 | 5.419498885 | 0.047523505 | 0.768887607 |
| ENSGALG00010011329 | PCDHGC3 | 408051 | 0.186651424 | 4.780811693 | 0.047528552 | 0.768887607 |
| ENSGALG00010006455 | NA | NA | -0.395798731 | 1.534653765 | 0.047573983 | 0.768887607 |
| ENSGALG00010011183 | NA | NA | -0.414031024 | 1.569067409 | 0.047650211 | 0.768981854 |
| ENSGALG00010022325 | MIGA1 | 771457 | 0.221841704 | 4.101635452 | 0.047710678 | 0.768981854 |
| ENSGALG00010015562 | IRF9 | 395243 | 0.435457443 | 1.282579182 | 0.047743297 | 0.768981854 |
| ENSGALG00010029127 | LOC112530003 | 112530003 | 0.515470395 | 0.782259167 | 0.047809019 | 0.768981854 |
| ENSGALG00010029293 | DNAH17 | 768861 | 0.242611559 | 5.506828163 | 0.047829971 | 0.768981854 |
| ENSGALG00010009847 | TMEM14C | 420851 | 0.208424375 | 4.710898419 | 0.04791797 | 0.769591639 |
| ENSGALG00010010371 | SOCS1L | 417978 | 0.192140542 | 4.327833349 | 0.04812378 | 0.772090294 |
| ENSGALG00010028310 | NA | NA | -0.577139583 | 1.85362604 | 0.048431403 | 0.775926216 |
| ENSGALG00010011717 | NA | NA | -0.39886667 | 2.962303374 | 0.048463837 | 0.775926216 |
| ENSGALG00010017538 | H3F3C | 427887 | 0.167308128 | 8.898527314 | 0.048612552 | 0.777290185 |
| ENSGALG00010010966 | LOC768760 | 768760 | -0.698684512 | -0.007330875 | 0.048650173 | 0.777290185 |
| ENSGALG00010020984 | PSEN1 | 373977 | 0.15912019 | 5.881698578 | 0.048791303 | 0.778221525 |
| ENSGALG00010027954 | KIAA1257 | 100857401 | -0.345012052 | 2.384376798 | 0.048826865 | 0.778221525 |
| ENSGALG00010028311 | CTXN1 | 420051 | 0.470738764 | 1.612006902 | 0.048860363 | 0.778221525 |
| ENSGALG00010018530 | FAM161A | 769188 | 0.42398974 | 1.564368968 | 0.049055635 | 0.77997227 |
| ENSGALG00010009285 | NA | NA | -0.753829173 | 0.05590363 | 0.049108637 | 0.77997227 |
| ENSGALG00010009367 | DMXL1 | 427385 | -0.157451583 | 6.233815898 | 0.049166066 | 0.77997227 |

| | | | | | | | |
|---|---|---|---|---|---|---|---|
| ENSGALG00010028507 | ANP32E | | 426109 | 0.17388359 | 9.93862065 | 0.049230356 | 0.77997227 |
| ENSGALG00010026315 | SYT2 | | 396327 | 0.525145199 | 0.611513892 | 0.04924301 | 0.77997227 |
| ENSGALG00010001688 | NA | NA | | -0.391796744 | 3.341799249 | 0.049274761 | 0.77997227 |
| ENSGALG00010027356 | NA | NA | | 0.176296172 | 5.211831305 | 0.049486587 | 0.781215662 |
| ENSGALG00010012478 | NA | NA | | -0.552697371 | 0.681017956 | 0.049492019 | 0.781215662 |
| ENSGALG00010028213 | NA | NA | | -0.297308097 | 3.162725712 | 0.049569552 | 0.781215662 |
| ENSGALG00010024192 | ALG6 | | 395078 | 0.158074363 | 5.687641756 | 0.049643627 | 0.781215662 |
| ENSGALG00010005310 | LAMTOR3 | | 425210 | 0.175444525 | 5.640326661 | 0.049723214 | 0.781215662 |
| ENSGALG00010025829 | PTPRVP | | 421174 | 0.508221059 | 1.331196651 | 0.049884265 | 0.781215662 |
| ENSGALG00010016393 | BLM | | 415577 | -0.151551619 | 6.765764079 | 0.049941641 | 0.781215662 |
| ENSGALG00010019912 | ARG2 | | 423272 | 0.254439074 | 3.524569579 | 0.049985857 | 0.781215662 |

Supplementally table S9. DEGs comparing 3x sgSoprano and 3x sgOVA in CRISPRi PGCs (p-adj < 0.05)

| Ensembl_ID | Symbol | entrez_ID | logFC | logCPM | p-adj_value | FDR |
|---|---|---|---|---|---|---|
| ENSGALG00010006179 | NA | NA | -5.1451683 | 5.26852166 | 1.25E-262 | 1.98E-258 |
| ENSGALG00010013561 | AGK | 418121 | -3.1186859 | 7.65006793 | 8.71E-237 | 6.91E-233 |
| ENSGALG00010007161 | NA | NA | -4.9134288 | 4.39989465 | 3.92E-206 | 1.95E-202 |
| ENSGALG00010008650 | NA | NA | -5.2667915 | 4.3295373 | 4.92E-206 | 1.95E-202 |
| ENSGALG00010006088 | NA | NA | -4.9136474 | 4.60836089 | 1.58E-191 | 5.00E-188 |
| ENSGALG00010007171 | NA | NA | -5.3472482 | 4.24547801 | 9.54E-178 | 2.52E-174 |
| ENSGALG00010009928 | NA | NA | -4.2759581 | 5.57383218 | 2.87E-174 | 6.49E-171 |
| ENSGALG00010012301 | NA | NA | -5.7823439 | 4.07150108 | 1.12E-158 | 2.23E-155 |
| ENSGALG00010002195 | NA | NA | -4.6943396 | 4.07331042 | 8.05E-157 | 1.42E-153 |
| ENSGALG00010013572 | DENND11 | 427917 | -2.1056004 | 6.78138913 | 1.19E-142 | 1.88E-139 |
| ENSGALG00010013126 | NA | NA | -4.7721593 | 6.21483883 | 2.81E-128 | 4.06E-125 |
| ENSGALG00010010174 | NA | NA | -5.6016726 | 3.20687563 | 1.30E-124 | 1.72E-121 |
| ENSGALG00010007287 | LOC124418405 | 124418405 | -2.5503477 | 9.45215404 | 1.16E-120 | 1.42E-117 |
| ENSGALG00010007167 | NA | NA | -4.6702163 | 3.6857215 | 7.90E-114 | 8.95E-111 |
| ENSGALG00010000706 | NA | NA | 7.12039851 | 7.46667038 | 2.52E-113 | 2.66E-110 |
| ENSGALG00010004343 | NA | NA | -2.3731139 | 6.6055838 | 2.17E-111 | 2.15E-108 |
| ENSGALG00010006214 | NA | NA | -4.4238255 | 7.86952777 | 2.40E-111 | 2.24E-108 |
| ENSGALG00010005803 | NA | NA | -1.8551348 | 6.28200845 | 3.05E-109 | 2.69E-106 |
| ENSGALG00010009891 | NA | NA | -3.8980733 | 3.46393356 | 6.99E-107 | 5.83E-104 |
| ENSGALG00010012306 | LOC121108207 | 121108207 | -5.6390964 | 4.99561711 | 1.36E-100 | 1.08E-97 |
| ENSGALG00010000831 | NA | NA | -4.0699985 | 6.99628392 | 5.64E-95 | 4.26E-92 |
| ENSGALG00010006087 | LOC107051465 | 107051465 | -3.890844 | 9.62488925 | 1.09E-93 | 7.88E-91 |
| ENSGALG00010014283 | NA | NA | -5.3681976 | 2.58316668 | 3.07E-93 | 2.12E-90 |
| ENSGALG00010009559 | NA | NA | -3.1524539 | 4.57850895 | 1.53E-92 | 1.01E-89 |
| ENSGALG00010003098 | NA | NA | -4.8394583 | 2.89728644 | 3.92E-91 | 2.49E-88 |
| ENSGALG00010006086 | LOC100857117 | 100857117 | -4.0529589 | 9.20502562 | 6.00E-91 | 3.66E-88 |
| ENSGALG00010009771 | NA | NA | -4.5027329 | 3.2542177 | 1.10E-87 | 6.44E-85 |
| ENSGALG00010010169 | NA | NA | -5.3382947 | 2.92891654 | 3.20E-84 | 1.81E-81 |
| ENSGALG00010016645 | NA | NA | -5.6234397 | 6.82432822 | 8.15E-84 | 4.46E-81 |
| ENSGALG00010008384 | NA | NA | -4.063161 | 3.17479998 | 7.96E-82 | 4.21E-79 |
| ENSGALG00010010193 | NA | NA | -4.7924412 | 4.86279748 | 1.75E-81 | 8.94E-79 |
| ENSGALG00010009800 | NA | NA | -4.7718714 | 2.63352284 | 3.58E-81 | 1.77E-78 |
| ENSGALG00010006902 | NA | NA | -5.1786069 | 2.56521121 | 9.49E-81 | 4.56E-78 |
| ENSGALG00010002947 | LOC121107416 | 121107416 | -4.9425344 | 10.3554524 | 2.16E-80 | 1.01E-77 |
| ENSGALG00010016320 | NA | NA | -4.7056619 | 2.4632954 | 5.24E-80 | 2.38E-77 |
| ENSGALG00010005001 | GUCA1C | 427960 | -5.0736145 | 5.70730247 | 2.40E-79 | 1.06E-76 |
| ENSGALG00010013729 | NA | NA | -4.5156536 | 2.79227289 | 2.97E-78 | 1.27E-75 |
| ENSGALG00010013432 | NA | NA | -3.2943004 | 4.17159884 | 1.55E-76 | 6.48E-74 |
| ENSGALG00010015173 | ENS-1 | 769415 | -3.6310882 | 10.9717104 | 7.97E-75 | 3.24E-72 |
| ENSGALG00010009553 | NA | NA | -4.4533976 | 7.36519704 | 3.47E-74 | 1.37E-71 |
| ENSGALG00010003639 | NA | NA | -4.8780062 | 9.50006124 | 1.01E-73 | 3.89E-71 |
| ENSGALG00010012294 | LOC100857299 | 100857299 | -4.1537095 | 7.52125876 | 2.60E-73 | 9.81E-71 |
| ENSGALG00010006916 | NA | NA | -5.925496 | 2.16246518 | 7.76E-72 | 2.83E-69 |
| ENSGALG00010006079 | LOC107055200 | 107055200 | -4.4119861 | 7.45283641 | 7.85E-72 | 2.83E-69 |
| ENSGALG00010013595 | WEE2 | 427918 | -4.5164448 | 5.3877298 | 1.67E-70 | 5.87E-68 |
| ENSGALG00010002503 | NA | NA | -4.8917996 | 8.08112611 | 1.70E-70 | 5.87E-68 |
| ENSGALG00010009878 | NA | NA | -4.683191 | 2.31554372 | 2.73E-70 | 9.20E-68 |
| ENSGALG00010018537 | LOC112530649 | 112530649 | -4.3531604 | 8.63131252 | 2.58E-69 | 8.54E-67 |
| ENSGALG00010008602 | NA | NA | -3.8346349 | 5.03522123 | 7.89E-69 | 2.55E-66 |
| ENSGALG00010002022 | ERNI | 769673 | -4.3793441 | 9.83914161 | 9.62E-69 | 3.05E-66 |
| ENSGALG00010007049 | NA | NA | -4.9744867 | 9.54666382 | 1.97E-68 | 6.12E-66 |
| ENSGALG00010014157 | NA | NA | -4.8017145 | 3.63758908 | 1.62E-66 | 4.95E-64 |
| ENSGALG00010004159 | NA | NA | -5.9910387 | 2.38911656 | 2.62E-66 | 7.84E-64 |
| ENSGALG00010001828 | LOC100858869 | 100858869 | -4.7875665 | 7.20937223 | 7.97E-65 | 2.34E-62 |
| ENSGALG00010006859 | NA | NA | -4.3768506 | 2.92403387 | 1.51E-64 | 4.35E-62 |
| ENSGALG00010001989 | NA | NA | -5.1515252 | 7.39051323 | 4.80E-64 | 1.36E-61 |
| ENSGALG00010003323 | STMN2 | 396095 | -2.3565621 | 4.19145601 | 6.88E-63 | 1.91E-60 |
| ENSGALG00010009801 | LOC101750251 | 101750251 | -4.7132114 | 8.90505761 | 1.69E-59 | 4.61E-57 |
| ENSGALG00010004527 | NA | NA | -4.3493272 | 4.04877594 | 1.21E-58 | 3.25E-56 |

| | | | | | | |
|---|---|---|---|---|---|---|
| ENSGALG00010000768 | NA | NA | -2.9551981 | 6.08972753 | 1.66E-58 | 4.40E-56 |
| ENSGALG00010001026 | NA | NA | -5.0791368 | 4.57764628 | 1.49E-57 | 3.88E-55 |
| ENSGALG00010018908 | NA | NA | -5.8379386 | 7.40491476 | 3.81E-57 | 9.74E-55 |
| ENSGALG00010001024 | NA | NA | -4.6575868 | 3.27242234 | 1.57E-56 | 3.96E-54 |
| ENSGALG00010000830 | NA | NA | -2.5808638 | 5.14203335 | 5.85E-56 | 1.45E-53 |
| ENSGALG00010001103 | NA | NA | -2.4780945 | 6.30588572 | 7.76E-56 | 1.89E-53 |
| ENSGALG00010000825 | NA | NA | -2.3963216 | 3.73370628 | 9.71E-56 | 2.33E-53 |
| ENSGALG00010004570 | LOC107051826 | 107051826 | -4.7005305 | 10.4099972 | 1.21E-55 | 2.86E-53 |
| ENSGALG00010007105 | NA | NA | -1.292598 | 5.94268585 | 3.29E-55 | 7.67E-53 |
| ENSGALG00010003536 | NA | NA | -2.8815569 | 5.16277682 | 3.66E-53 | 8.41E-51 |
| ENSGALG00010005791 | NA | NA | -5.42458 | 1.84169948 | 5.71E-53 | 1.29E-50 |
| ENSGALG00010010189 | NA | NA | -2.8801974 | 3.96221062 | 1.51E-50 | 3.37E-48 |
| ENSGALG00010008968 | NA | NA | -5.2543791 | 7.76240764 | 2.16E-50 | 4.75E-48 |
| ENSGALG00010011694 | NA | NA | -3.8398279 | 2.02249052 | 3.27E-50 | 7.10E-48 |
| ENSGALG00010001025 | NA | NA | -4.7527299 | 4.08279583 | 8.91E-50 | 1.91E-47 |
| ENSGALG00010000835 | NA | NA | -2.5406945 | 5.44989889 | 2.27E-49 | 4.80E-47 |
| ENSGALG00010025713 | LOC107054297 | 107054297 | -2.2262124 | 5.12897201 | 3.99E-48 | 8.33E-46 |
| ENSGALG00010023095 | NA | NA | -4.3759877 | 2.61189151 | 1.07E-46 | 2.20E-44 |
| ENSGALG00010019013 | LRRK1 | 415520 | -1.4367851 | 4.9787885 | 1.12E-46 | 2.28E-44 |
| ENSGALG00010014282 | NA | NA | -5.9609393 | 1.28195184 | 2.80E-46 | 5.63E-44 |
| ENSGALG00010023092 | NA | NA | -4.0675768 | 2.73502406 | 3.57E-46 | 7.08E-44 |
| ENSGALG00010001483 | LOC121110182 | 121110182 | -4.1326264 | 5.29419594 | 3.85E-46 | 7.53E-44 |
| ENSGALG00010011693 | NA | NA | -3.6986695 | 4.92078198 | 2.60E-45 | 5.03E-43 |
| ENSGALG00010004357 | NA | NA | -2.7209747 | 4.71543029 | 6.49E-45 | 1.24E-42 |
| ENSGALG00010004371 | NA | NA | -4.081432 | 8.00880947 | 4.09E-44 | 7.72E-42 |
| ENSGALG00010013433 | NA | NA | -4.745358 | 1.47012223 | 7.53E-44 | 1.41E-41 |
| ENSGALG00010004208 | NA | NA | -5.5416309 | 1.41744017 | 1.45E-43 | 2.68E-41 |
| ENSGALG00010006306 | NA | NA | -2.5136223 | 2.66846879 | 4.07E-42 | 7.43E-40 |
| ENSGALG00010008603 | NA | NA | -3.2631099 | 2.16693424 | 1.90E-39 | 3.43E-37 |
| ENSGALG00010011849 | NA | NA | -1.7733916 | 7.27863391 | 4.80E-39 | 8.56E-37 |
| ENSGALG00010014785 | NA | NA | -3.214027 | 1.82447556 | 5.31E-39 | 9.35E-37 |
| ENSGALG00010006798 | NA | NA | -1.376995 | 5.07218993 | 7.73E-39 | 1.35E-36 |
| ENSGALG00010001031 | NA | NA | -2.5370103 | 3.60821299 | 1.31E-37 | 2.26E-35 |
| ENSGALG00010008982 | NA | NA | -5.3580645 | 1.22419773 | 5.61E-37 | 9.57E-35 |
| ENSGALG00010008933 | NA | NA | -3.6867821 | 5.04938249 | 5.98E-36 | 1.01E-33 |
| ENSGALG00010007115 | CD69L | 417816 | -0.959298 | 5.66815366 | 1.97E-34 | 3.28E-32 |
| ENSGALG00010003524 | NA | NA | -8.927294 | 0.82782252 | 8.07E-34 | 1.33E-31 |
| ENSGALG00010010172 | NA | NA | -6.1678031 | 0.65057625 | 2.00E-32 | 3.27E-30 |
| ENSGALG00010006797 | NA | NA | -1.3200494 | 6.4383579 | 3.31E-32 | 5.36E-30 |
| ENSGALG00010001209 | CDH7 | 374007 | -1.4392055 | 4.13944843 | 1.87E-31 | 3.00E-29 |
| ENSGALG00010005090 | NA | NA | -2.0036152 | 6.95362121 | 7.18E-31 | 1.14E-28 |
| ENSGALG00010005493 | KLF6 | 420463 | -1.402775 | 5.29566331 | 1.86E-29 | 2.93E-27 |
| ENSGALG00010017006 | DNAH3 | 427004 | -1.3446981 | 5.24188211 | 2.57E-29 | 4.00E-27 |
| ENSGALG00010008989 | NA | NA | -4.2441452 | 0.99550286 | 9.20E-29 | 1.42E-26 |
| ENSGALG00010014606 | NA | NA | -2.0120719 | 2.58726968 | 2.12E-27 | 3.24E-25 |
| ENSGALG00010007504 | TMPRSS15 | 427967 | -5.0062883 | 0.66661384 | 2.00E-26 | 3.02E-24 |
| ENSGALG00010009787 | LOC112532404 | 112532404 | -2.2030687 | 2.864074 | 3.23E-26 | 4.83E-24 |
| ENSGALG00010013128 | NA | NA | -3.9318728 | 0.72893184 | 4.11E-26 | 6.09E-24 |
| ENSGALG00010007068 | NCAM2 | 427968 | -1.884319 | 3.51422348 | 2.61E-25 | 3.83E-23 |
| ENSGALG00010009552 | NA | NA | -4.283 | 0.44599068 | 2.61E-24 | 3.79E-22 |
| ENSGALG00010001477 | NLGN4 | 428006 | -1.4440579 | 3.39023948 | 3.66E-24 | 5.27E-22 |
| ENSGALG00010013113 | NA | NA | -2.0862773 | 2.7827625 | 4.76E-24 | 6.79E-22 |
| ENSGALG00010008454 | NA | NA | -3.1502353 | 1.79857038 | 4.79E-24 | 6.79E-22 |
| ENSGALG00010005024 | NA | NA | -4.5436588 | 0.47825087 | 1.41E-23 | 1.98E-21 |
| ENSGALG00010003261 | PEX2 | 420192 | -0.9704661 | 7.21745398 | 4.30E-23 | 5.99E-21 |
| ENSGALG00010006910 | NA | NA | -4.6552228 | 0.35873603 | 1.00E-22 | 1.39E-20 |
| ENSGALG00010018386 | NA | NA | -4.8805488 | 0.27590662 | 1.95E-22 | 2.67E-20 |
| ENSGALG00010002390 | MYT1L | 421915 | -2.2472514 | 2.02274382 | 2.13E-22 | 2.88E-20 |
| ENSGALG00010001745 | CADPS2 | 417756 | -1.7118328 | 3.72342617 | 1.86E-21 | 2.50E-19 |
| ENSGALG00010014169 | NA | NA | -5.5273789 | 0.5217508 | 6.38E-21 | 8.51E-19 |

| | | | | | | |
|---|---|---|---|---|---|---|
| ENSGALG00010017820 | FLRT2 | 428901 | -1.4391206 | 7.19548353 | 2.43E-20 | 3.22E-18 |
| ENSGALG00010009859 | NA | NA | -2.9278271 | 1.62671958 | 5.54E-20 | 7.27E-18 |
| ENSGALG00010005008 | LOC418414 | 418414 | -3.846973 | 0.5296823 | 8.94E-20 | 1.16E-17 |
| ENSGALG00010013270 | NA | NA | -1.4747806 | 3.20270093 | 1.03E-19 | 1.33E-17 |
| ENSGALG00010023591 | NA | NA | -1.6860853 | 2.27496325 | 7.18E-19 | 9.18E-17 |
| ENSGALG00010004360 | NA | NA | -2.8658399 | 1.23419691 | 7.42E-19 | 9.41E-17 |
| ENSGALG00010009781 | NA | NA | -2.7612446 | 0.97515024 | 2.71E-18 | 3.41E-16 |
| ENSGALG00010014226 | SLC16A7 | 417815 | -1.5665524 | 2.5503349 | 1.16E-17 | 1.45E-15 |
| ENSGALG00010005072 | NA | NA | -2.0394732 | 1.74269509 | 1.35E-17 | 1.67E-15 |
| ENSGALG00010007183 | VOPP1 | 776358 | -0.8896031 | 5.56906159 | 2.47E-17 | 3.03E-15 |
| ENSGALG00010021622 | PTHLH | 396281 | -2.043095 | 1.66603305 | 9.57E-17 | 1.17E-14 |
| ENSGALG00010007691 | LOC121108447 | 121108447 | -1.2921579 | 3.04518851 | 1.13E-16 | 1.37E-14 |
| ENSGALG00010015806 | NA | NA | -5.6598764 | -0.4997522 | 4.10E-16 | 4.93E-14 |
| ENSGALG00010003372 | ENOX1 | 418839 | -1.9690172 | 1.95425813 | 1.13E-15 | 1.35E-13 |
| ENSGALG00010002061 | NA | NA | -0.8287715 | 4.89913469 | 2.55E-15 | 3.02E-13 |
| ENSGALG00010022969 | TTN | 424126 | 0.74228629 | 7.04159001 | 4.02E-15 | 4.72E-13 |
| ENSGALG00010007446 | NA | NA | -1.6518804 | 2.88947131 | 4.39E-15 | 5.11E-13 |
| ENSGALG00010002078 | NA | NA | -7.4073022 | -0.5406237 | 9.85E-15 | 1.14E-12 |
| ENSGALG00010002570 | ADAM12 | 425843 | 0.75455813 | 6.14478228 | 2.18E-14 | 2.51E-12 |
| ENSGALG00010004631 | NA | NA | 1.18174519 | 4.23228489 | 2.54E-14 | 2.90E-12 |
| ENSGALG00010010166 | NA | NA | -1.2452195 | 4.50889671 | 8.57E-14 | 9.71E-12 |
| ENSGALG00010014782 | NA | NA | -1.2271547 | 2.8897675 | 1.49E-13 | 1.68E-11 |
| ENSGALG00010008645 | NA | NA | -7.1907831 | -0.7198614 | 6.84E-13 | 7.64E-11 |
| ENSGALG00010028374 | FDPS | 425061 | -0.6065988 | 5.53118791 | 8.14E-13 | 9.03E-11 |
| ENSGALG00010001945 | NA | NA | -5.4394854 | -0.6729037 | 1.00E-12 | 1.10E-10 |
| ENSGALG00010008427 | NA | NA | -5.2967485 | -0.7939661 | 1.13E-12 | 1.23E-10 |
| ENSGALG00010012921 | EBF1 | 395512 | 0.85339961 | 5.0600475 | 1.13E-12 | 1.23E-10 |
| ENSGALG00010009480 | NA | NA | -1.9515235 | 1.21506745 | 1.31E-12 | 1.41E-10 |
| ENSGALG00010010061 | CYP2D6 | 417981 | 0.82728846 | 4.38937398 | 1.54E-12 | 1.65E-10 |
| ENSGALG00010005079 | NA | NA | -1.9268435 | 1.38432834 | 1.23E-11 | 1.30E-09 |
| ENSGALG00010025490 | NA | NA | 6.33681403 | 2.88266527 | 1.23E-11 | 1.30E-09 |
| ENSGALG00010014656 | NA | NA | -1.7758936 | 1.97999138 | 1.62E-11 | 1.70E-09 |
| ENSGALG00010027117 | RASSF5 | 419844 | 0.56623955 | 5.97901629 | 3.08E-11 | 3.22E-09 |
| ENSGALG00010011001 | ANGPT1 | 395129 | -3.3819315 | 0.04284901 | 3.29E-11 | 3.41E-09 |
| ENSGALG00010000697 | LOC107050463 | 107050463 | 0.72951454 | 4.77161586 | 4.50E-11 | 4.63E-09 |
| ENSGALG00010001292 | LOX | 396474 | 0.66737667 | 5.80804333 | 7.17E-11 | 7.34E-09 |
| ENSGALG00010000847 | NA | NA | -2.8997977 | -0.2413056 | 1.53E-10 | 1.56E-08 |
| ENSGALG00010004335 | NA | NA | -4.3593766 | -0.8985826 | 1.99E-10 | 2.02E-08 |
| ENSGALG00010014601 | NA | NA | -1.3902964 | 2.48733489 | 2.58E-10 | 2.59E-08 |
| ENSGALG00010007103 | NA | NA | -1.595245 | 1.60049332 | 2.75E-10 | 2.75E-08 |
| ENSGALG00010000381 | LOC107051255 | 107051255 | 0.81186219 | 3.81278329 | 3.33E-10 | 3.30E-08 |
| ENSGALG00010010072 | NA | NA | -6.6679695 | -1.1298887 | 5.11E-10 | 5.03E-08 |
| ENSGALG00010009556 | NA | NA | -2.3498799 | 0.58060764 | 9.73E-10 | 9.52E-08 |
| ENSGALG00010029333 | NA | NA | -0.4221693 | 7.2603609 | 1.52E-09 | 1.48E-07 |
| ENSGALG00010000826 | NA | NA | -2.4285212 | 0.16214978 | 3.06E-09 | 2.96E-07 |
| ENSGALG00010009720 | RELN | 427850 | 0.59935276 | 4.93477551 | 4.44E-09 | 4.27E-07 |
| ENSGALG00010000017 | COX1 | 63549487 | -0.5498613 | 13.6624804 | 6.77E-09 | 6.47E-07 |
| ENSGALG00010013245 | NA | NA | -2.4784174 | 1.03084246 | 7.32E-09 | 6.96E-07 |
| ENSGALG00010015500 | NA | NA | -2.3081682 | 0.11873418 | 7.84E-09 | 7.40E-07 |
| ENSGALG00010011206 | LOC112532760 | 112532760 | 1.00018373 | 3.53619055 | 7.92E-09 | 7.43E-07 |
| ENSGALG00010028386 | NCAN | 395493 | 0.60155603 | 4.66774863 | 1.49E-08 | 1.39E-06 |
| ENSGALG00010001300 | MRPL49 | 100858532 | 0.7433464 | 3.64762631 | 1.57E-08 | 1.46E-06 |
| ENSGALG00010014042 | SGCZ | 422739 | -0.5709055 | 5.06396008 | 2.05E-08 | 1.89E-06 |
| ENSGALG00010001817 | SLC25A6 | 374072 | -0.4269458 | 8.88634209 | 2.08E-08 | 1.91E-06 |
| ENSGALG00010015596 | UCHL1 | 770302 | -0.3908701 | 8.6232442 | 2.20E-08 | 2.01E-06 |
| ENSGALG00010020570 | NA | NA | -2.0468825 | 0.59571967 | 2.22E-08 | 2.01E-06 |
| ENSGALG00010011353 | LRP1B | 424301 | -0.4738912 | 7.39390491 | 2.78E-08 | 2.51E-06 |
| ENSGALG00010000023 | ATP6 | 63549466 | -0.6600451 | 12.8735458 | 3.03E-08 | 2.72E-06 |
| ENSGALG00010007107 | NA | NA | 0.97154593 | 3.69852341 | 3.72E-08 | 3.31E-06 |
| ENSGALG00010001380 | SCRN1 | 420635 | -0.6302054 | 6.2999454 | 3.75E-08 | 3.32E-06 |

| | | | | | | |
|---|---|---|---|---|---|---|
| ENSGALG00010017621 | SYNE2 | 423509 | 0.3806537 | 7.3038452 | 5.58E-08 | 4.92E-06 |
| ENSGALG00010009744 | DAPK1 | 431670 | -0.5676139 | 5.64578989 | 6.58E-08 | 5.77E-06 |
| ENSGALG00010000018 | NA | NA | -2.0009408 | 0.75908231 | 1.14E-07 | 9.97E-06 |
| ENSGALG00010028301 | FBN3 | 395079 | 0.50590927 | 6.26225972 | 1.15E-07 | 1.00E-05 |
| ENSGALG00010000034 | CYTB | 63549472 | -0.6227377 | 13.0676779 | 1.19E-07 | 1.02E-05 |
| ENSGALG00010029736 | LOC101747587 | 101747587 | 0.43678097 | 9.08540927 | 1.35E-07 | 1.16E-05 |
| ENSGALG00010011825 | FAM19A2 | 771745 | -0.5106496 | 6.00383206 | 1.49E-07 | 1.27E-05 |
| ENSGALG00010020685 | RPL6 | 373957 | -0.4035481 | 9.4265242 | 1.74E-07 | 1.47E-05 |
| ENSGALG00010004838 | NA | NA | -2.9976264 | -0.4007129 | 1.90E-07 | 1.60E-05 |
| ENSGALG00010025629 | ST6GAL1 | 396169 | 0.70721893 | 3.60813291 | 1.96E-07 | 1.64E-05 |
| ENSGALG00010029411 | SARM1 | 417568 | -0.5177933 | 5.08179116 | 2.04E-07 | 1.70E-05 |
| ENSGALG00010000090 | NA | NA | 1.44149028 | 1.96678073 | 2.09E-07 | 1.74E-05 |
| ENSGALG00010026590 | NA | NA | -0.8532349 | 4.20557583 | 2.32E-07 | 1.91E-05 |
| ENSGALG00010027046 | HESX1 | 395864 | -0.7350358 | 3.49047906 | 2.41E-07 | 1.98E-05 |
| ENSGALG00010000824 | NA | NA | -1.3977773 | 1.05565374 | 2.64E-07 | 2.16E-05 |
| ENSGALG00010000033 | ND5 | 63549495 | -0.6249694 | 12.9877944 | 2.70E-07 | 2.19E-05 |
| ENSGALG00010000328 | NA | NA | -0.9485384 | 2.3929929 | 2.83E-07 | 2.29E-05 |
| ENSGALG00010030101 | MIR3064 | 102465325 | -0.7156867 | 3.46617543 | 2.89E-07 | 2.33E-05 |
| ENSGALG00010021615 | TATDN3 | 421366 | -0.5639458 | 7.47165856 | 2.92E-07 | 2.34E-05 |
| ENSGALG00010002060 | NA | NA | -0.7514383 | 3.79152473 | 2.97E-07 | 2.37E-05 |
| ENSGALG00010009441 | ZNF518B | 422841 | 0.56203877 | 5.2390016 | 4.10E-07 | 3.25E-05 |
| ENSGALG00010018484 | NLGN1 | 424980 | 0.60268417 | 5.06764065 | 5.64E-07 | 4.45E-05 |
| ENSGALG00010029270 | TIMP2 | 374178 | -0.5614627 | 5.26602156 | 6.56E-07 | 5.15E-05 |
| ENSGALG00010019282 | IGFBP2 | 396315 | 0.46096814 | 5.57980431 | 7.16E-07 | 5.60E-05 |
| ENSGALG00010029191 | GSG1L2 | 768455 | 1.35920647 | 1.94905708 | 7.31E-07 | 5.68E-05 |
| ENSGALG00010000386 | NA | NA | 0.5129706 | 5.0186356 | 7.47E-07 | 5.77E-05 |
| ENSGALG00010028684 | NA | NA | -1.8884929 | 0.38717362 | 7.49E-07 | 5.77E-05 |
| ENSGALG00010026418 | NA | NA | 1.14739079 | 1.55767607 | 7.58E-07 | 5.81E-05 |
| ENSGALG00010006623 | LOC107052071 | 107052071 | -1.3707388 | 1.86737519 | 7.63E-07 | 5.82E-05 |
| ENSGALG00010027165 | LOC100857637 | 100857637 | -0.5545777 | 5.58246973 | 8.60E-07 | 6.53E-05 |
| ENSGALG00010000029 | ND4 | 63549470 | -0.6236921 | 12.6368602 | 9.55E-07 | 7.22E-05 |
| ENSGALG00010006909 | NA | NA | -1.4511633 | 1.39240202 | 9.80E-07 | 7.37E-05 |
| ENSGALG00010005534 | IDI2 | 420459 | -0.4419338 | 6.00577906 | 1.02E-06 | 7.61E-05 |
| ENSGALG00010014334 | HMGCS1 | 396379 | -0.4345933 | 7.71452012 | 1.03E-06 | 7.70E-05 |
| ENSGALG00010017272 | NRXN1 | 395398 | 0.42583035 | 6.84887179 | 1.06E-06 | 7.83E-05 |
| ENSGALG00010013715 | IL1R1 | 396481 | 0.73488456 | 3.65211404 | 1.11E-06 | 8.16E-05 |
| ENSGALG00010030080 | LOC101751181 | 101751181 | 0.34316402 | 7.38176224 | 1.13E-06 | 8.28E-05 |
| ENSGALG00010014124 | AK9 | 771077 | 0.71351188 | 4.43642243 | 1.22E-06 | 8.89E-05 |
| ENSGALG00010005319 | NOL4L | 769466 | 1.05735181 | 1.95761219 | 1.22E-06 | 8.89E-05 |
| ENSGALG00010028153 | MANF | 107048996 | -0.7284689 | 6.46363422 | 1.35E-06 | 9.77E-05 |
| ENSGALG00010008343 | TNFRSF21 | 422062 | -0.44264 | 6.2967635 | 1.51E-06 | 0.0001087 |
| ENSGALG00010027704 | NA | NA | 0.76768854 | 3.18828143 | 1.71E-06 | 0.00012244 |
| ENSGALG00010016022 | EPHB3 | 396179 | 0.45968133 | 5.85882389 | 1.94E-06 | 0.00013857 |
| ENSGALG00010027785 | NA | NA | 0.45076461 | 5.36890323 | 1.96E-06 | 0.00013908 |
| ENSGALG00010028025 | TNNC1 | 396032 | -0.6852324 | 3.56514551 | 2.13E-06 | 0.00015058 |
| ENSGALG00010023662 | LOC121106465 | 121106465 | 2.05875819 | -0.0413957 | 2.14E-06 | 0.00015058 |
| ENSGALG00010015908 | CHRD | 395828 | 0.76217067 | 3.61360583 | 2.70E-06 | 0.00018949 |
| ENSGALG00010000484 | NA | NA | 1.83553029 | 0.23781981 | 2.76E-06 | 0.00019316 |
| ENSGALG00010006076 | PODXL | 395755 | 0.73217656 | 3.78694518 | 2.89E-06 | 0.00020132 |
| ENSGALG00010011046 | NA | NA | 1.24312133 | 1.09811787 | 3.13E-06 | 0.00021705 |
| ENSGALG00010020760 | TXNDC11 | 416428 | -0.6183928 | 4.23915652 | 3.48E-06 | 0.00024033 |
| ENSGALG00010008301 | FAM91A1 | 770621 | -0.4161099 | 5.60229795 | 3.78E-06 | 0.00025967 |
| ENSGALG00010030076 | NA | NA | 0.82918511 | 2.58301425 | 3.89E-06 | 0.00026608 |
| ENSGALG00010011810 | FGFR4 | 395603 | -0.4686644 | 5.09261647 | 4.12E-06 | 0.00027965 |
| ENSGALG00010027969 | HSPA5 | 396487 | -0.6003365 | 9.42998922 | 4.13E-06 | 0.00027965 |
| ENSGALG00010027865 | SELENOK | 415995 | -0.5655297 | 4.6605316 | 4.15E-06 | 0.00028009 |
| ENSGALG00010024204 | PTBP2 | 424479 | -0.3737301 | 6.91564465 | 4.23E-06 | 0.000284 |
| ENSGALG00010007920 | FAT3 | 395167 | 0.46155783 | 5.98618066 | 4.49E-06 | 0.00030076 |
| ENSGALG00010001138 | SPTSSB | 425008 | 0.50971441 | 5.20078461 | 4.63E-06 | 0.00030777 |
| ENSGALG00010011321 | MAP2 | 424001 | 0.40019903 | 6.20763963 | 4.64E-06 | 0.00030777 |

| | | | | | | |
|---|---|---|---|---|---|---|
| ENSGALG00010020640 | PTPN14 | 421360 | 0.3950771 | 5.31405849 | 4.70E-06 | 0.00031078 |
| ENSGALG00010010003 | NA | NA | 0.66141947 | 5.0478394 | 4.72E-06 | 0.00031086 |
| ENSGALG00010013243 | NA | NA | -1.0200919 | 1.93708752 | 4.76E-06 | 0.00031217 |
| ENSGALG00010013175 | HIPK2 | 374138 | 0.35726547 | 6.00976785 | 4.96E-06 | 0.00032348 |
| ENSGALG00010019618 | NPHP4 | 419377 | 0.46591286 | 4.78968509 | 5.20E-06 | 0.00033834 |
| ENSGALG00010010885 | SNRPD2 | 107049207 | 0.39444847 | 6.67352179 | 5.65E-06 | 0.0003656 |
| ENSGALG00010022682 | EIF1 | 420037 | -0.3372303 | 7.69424352 | 5.74E-06 | 0.00037025 |
| ENSGALG00010005856 | NA | NA | 1.22949811 | 1.84091171 | 6.18E-06 | 0.00039718 |
| ENSGALG00010002367 | STK31 | 420621 | 0.38147537 | 8.02665195 | 6.21E-06 | 0.00039728 |
| ENSGALG00010015084 | FOXO6 | 100857767 | 0.39731018 | 5.42789277 | 6.31E-06 | 0.00040192 |
| ENSGALG00010003099 | NA | NA | -1.6612802 | 0.51890024 | 6.71E-06 | 0.00042582 |
| ENSGALG00010018284 | SPEG | 429033 | -0.5370721 | 4.20335473 | 7.56E-06 | 0.00047781 |
| ENSGALG00010009841 | GABRA1 | 374214 | 0.82528253 | 2.22342497 | 7.60E-06 | 0.00047829 |
| ENSGALG00010007400 | LOC121109235 | 121109235 | -0.8860843 | 2.13846591 | 7.68E-06 | 0.00048176 |
| ENSGALG00010006341 | HHIP | 422460 | -0.7711968 | 3.62920206 | 8.32E-06 | 0.00051943 |
| ENSGALG00010017497 | NA | NA | -1.4016698 | 0.52847468 | 8.91E-06 | 0.00055413 |
| ENSGALG00010016431 | B3GNT5 | 424960 | 0.43444527 | 6.66685776 | 9.13E-06 | 0.00056573 |
| ENSGALG00010019281 | LOC421195 | 421195 | 0.4826269 | 5.57631956 | 9.21E-06 | 0.00056847 |
| ENSGALG00010021788 | RPL5 | 395269 | -0.3834216 | 9.49758286 | 9.43E-06 | 0.00057953 |
| ENSGALG00010014390 | SPARC | 386571 | -0.5109932 | 7.65540262 | 1.03E-05 | 0.00062914 |
| ENSGALG00010022831 | LRRC8C | 424515 | -0.6981997 | 3.35750465 | 1.15E-05 | 0.00070238 |
| ENSGALG00010023974 | PREX1 | 419210 | 0.35496168 | 5.61127023 | 1.18E-05 | 0.00071668 |
| ENSGALG00010018724 | RRBP1 | 396414 | -0.4233396 | 6.38495464 | 1.23E-05 | 0.00074684 |
| ENSGALG00010014608 | NA | NA | -1.5364425 | 0.1816398 | 1.25E-05 | 0.0007538 |
| ENSGALG00010001092 | NA | NA | -1.5043626 | 0.39338389 | 1.27E-05 | 0.00076444 |
| ENSGALG00010001419 | KCND2 | 374143 | -0.7560111 | 2.85553193 | 1.30E-05 | 0.00077819 |
| ENSGALG00010001220 | PLXNB2 | 425938 | 0.3374411 | 7.0621563 | 1.37E-05 | 0.00081571 |
| ENSGALG00010000037 | ND6 | 63549497 | -0.6214133 | 8.25199331 | 1.40E-05 | 0.00083223 |
| ENSGALG00010015759 | EIF5 | 423479 | -0.2957021 | 9.33693668 | 1.48E-05 | 0.00087483 |
| ENSGALG00010006492 | GAP43 | 427955 | 0.57006019 | 3.59375331 | 1.58E-05 | 0.0009327 |
| ENSGALG00010001666 | PIEZO2 | 421052 | 0.63142193 | 3.99270924 | 1.63E-05 | 0.00095849 |
| ENSGALG00010023311 | ETS1 | 396235 | 0.62941712 | 4.91843364 | 1.68E-05 | 0.00098053 |
| ENSGALG00010029263 | TOB1 | 395225 | -0.4081693 | 6.51954597 | 1.71E-05 | 0.00099975 |
| ENSGALG00010017943 | NA | NA | -0.5892719 | 4.82004184 | 1.79E-05 | 0.00103788 |
| ENSGALG00010023309 | PIK3R3 | 429096 | 0.37416113 | 5.07599811 | 1.80E-05 | 0.00104409 |
| ENSGALG00010001224 | DNAJB9 | 417783 | -0.46688 | 5.95811248 | 1.92E-05 | 0.00110517 |
| ENSGALG00010026252 | ENO2 | 395689 | -0.7531597 | 3.6973848 | 1.93E-05 | 0.00110921 |
| ENSGALG00010015643 | LOC423474 | 423474 | 1.18578303 | 1.5315893 | 1.94E-05 | 0.00110981 |
| ENSGALG00010017650 | NA | NA | -1.986772 | -0.3853159 | 1.95E-05 | 0.00111031 |
| ENSGALG00010016535 | BNC2 | 431604 | 1.67877792 | 0.06025788 | 2.04E-05 | 0.00116005 |
| ENSGALG00010021867 | CXCL12 | 395180 | 0.52077093 | 6.80043949 | 2.05E-05 | 0.00116356 |
| ENSGALG00010002358 | FREM2 | 418893 | -0.6675217 | 3.95028024 | 2.09E-05 | 0.00117627 |
| ENSGALG00010009382 | NA | NA | 0.76784536 | 2.50111291 | 2.09E-05 | 0.00117627 |
| ENSGALG00010019657 | PPP1R17 | 420753 | -0.6858532 | 4.82154613 | 2.13E-05 | 0.00119204 |
| ENSGALG00010015218 | PLCL1 | 424060 | 0.34854204 | 6.54578287 | 2.16E-05 | 0.00120657 |
| ENSGALG00010007165 | LOC101747821 | 101747821 | -0.7163282 | 4.78292975 | 2.24E-05 | 0.00123933 |
| ENSGALG00010015163 | PLK2 | 427150 | -0.680816 | 4.54243129 | 2.24E-05 | 0.00123933 |
| ENSGALG00010025219 | NACA | 396544 | -0.3154863 | 8.37242308 | 2.24E-05 | 0.00123933 |
| ENSGALG00010009327 | NID1 | 395531 | 0.6894583 | 7.7004735 | 2.28E-05 | 0.00125805 |
| ENSGALG00010028653 | NA | NA | -0.5559198 | 3.75430534 | 2.34E-05 | 0.00128529 |
| ENSGALG00010000011 | ND2 | 63549482 | -0.5485227 | 11.7913525 | 2.41E-05 | 0.00131749 |
| ENSGALG00010024883 | ING5 | 424852 | -0.45575 | 6.31627398 | 2.45E-05 | 0.00133368 |
| ENSGALG00010000020 | COX2 | 63549489 | -0.5025305 | 12.7811775 | 2.59E-05 | 0.00140393 |
| ENSGALG00010009210 | PCDH15 | 423644 | 0.41960483 | 5.31306737 | 2.59E-05 | 0.00140393 |
| ENSGALG00010006304 | NA | NA | -1.8512459 | -0.2606433 | 2.63E-05 | 0.00141877 |
| ENSGALG00010013237 | SVOPL | 418105 | 0.71819455 | 2.67216242 | 2.68E-05 | 0.00144226 |
| ENSGALG00010017534 | ATP11C | 422254 | 0.30731022 | 7.24491167 | 2.70E-05 | 0.00144843 |
| ENSGALG00010024693 | ANXA2 | 396297 | -0.3025215 | 6.54003002 | 2.73E-05 | 0.00145578 |
| ENSGALG00010007302 | CDH11 | 415797 | -0.4269779 | 5.25682408 | 2.75E-05 | 0.00146617 |
| ENSGALG00010010909 | SLIT3 | 374173 | 0.37195303 | 4.96114834 | 2.82E-05 | 0.00149792 |

| | | | | | | |
|---|---|---|---|---|---|---|
| ENSGALG00010007467 | CHODL | 418474 | -1.0390479 | 2.69852365 | 2.87E-05 | 0.00151773 |
| ENSGALG00010024974 | CD44 | 395666 | 0.78704015 | 3.98609943 | 2.91E-05 | 0.00153557 |
| ENSGALG00010022784 | DMRTA2 | 107054041 | -0.4840266 | 4.49838901 | 3.10E-05 | 0.00162764 |
| ENSGALG00010000646 | NA | NA | 0.54987659 | 6.04961259 | 3.13E-05 | 0.00163858 |
| ENSGALG00010016983 | ZFAND2A | 425104 | -0.6882171 | 3.77473988 | 3.21E-05 | 0.00167412 |
| ENSGALG00010029200 | GAS7 | 417305 | 0.29971706 | 6.37415651 | 3.39E-05 | 0.00176292 |
| ENSGALG00010010779 | DOCK2 | 427612 | 0.54246033 | 4.00653596 | 3.40E-05 | 0.00176292 |
| ENSGALG00010005267 | CYP51A1 | 420548 | -0.3922041 | 6.63689935 | 3.42E-05 | 0.00176488 |
| ENSGALG00010023237 | GAD1 | 395743 | 0.61901533 | 3.53132533 | 3.54E-05 | 0.00182192 |
| ENSGALG00010029778 | DOC2B | 417624 | -0.6640764 | 4.86724657 | 3.63E-05 | 0.00186329 |
| ENSGALG00010010370 | NA | NA | -1.4685483 | 0.3848228 | 3.76E-05 | 0.00192437 |
| ENSGALG00010020396 | CHL1 | 416102 | -0.4450325 | 5.23433375 | 3.86E-05 | 0.00196919 |
| ENSGALG00010000430 | NA | NA | -1.1140536 | 1.82519387 | 3.94E-05 | 0.00200146 |
| ENSGALG00010024946 | KIF1A | 424849 | -0.4383146 | 5.43229887 | 3.95E-05 | 0.00200299 |
| ENSGALG00010007414 | ROBO2 | 395291 | -0.4765374 | 4.11227736 | 3.99E-05 | 0.00201294 |
| ENSGALG00010020532 | SIX3 | 378786 | -0.9411089 | 1.55643334 | 4.00E-05 | 0.00201294 |
| ENSGALG00010028348 | NES | 395890 | -0.6170041 | 6.87291139 | 4.01E-05 | 0.00201372 |
| ENSGALG00010015766 | TLE1 | 427450 | -0.3681895 | 5.17453386 | 4.17E-05 | 0.00208485 |
| ENSGALG00010014630 | NA | NA | 0.54037358 | 4.18578881 | 4.26E-05 | 0.00212572 |
| ENSGALG00010027097 | SRGAP2 | 419843 | 0.35128194 | 6.71686702 | 4.49E-05 | 0.00223375 |
| ENSGALG00010008561 | EEF1D | 107049004 | -0.3644701 | 8.72821141 | 4.60E-05 | 0.00228093 |
| ENSGALG00010007855 | RNF182 | 771221 | 0.34086667 | 6.37150185 | 4.75E-05 | 0.00234413 |
| ENSGALG00010012165 | NA | NA | -1.0552198 | 1.43849658 | 4.76E-05 | 0.00234413 |
| ENSGALG00010026633 | SLC5A1 | 395496 | 0.73956631 | 3.27342813 | 4.96E-05 | 0.00243797 |
| ENSGALG00010020143 | KCNK5 | 421424 | 0.41104656 | 5.02589385 | 5.01E-05 | 0.00245305 |
| ENSGALG00010028024 | RBM3 | 425789 | -0.3100523 | 8.31489747 | 5.11E-05 | 0.0024933 |
| ENSGALG00010028174 | GMPPB | 415924 | -0.6105561 | 4.0856993 | 5.15E-05 | 0.0025036 |
| ENSGALG00010021427 | RPLP0 | 395835 | -0.3278811 | 9.64857346 | 5.26E-05 | 0.00255392 |
| ENSGALG00010007877 | MSMO1 | 422423 | -0.473562 | 7.34579599 | 5.33E-05 | 0.00257195 |
| ENSGALG00010015569 | NA | NA | 1.20215943 | 0.84809936 | 5.33E-05 | 0.00257195 |
| ENSGALG00010027220 | ANKRD24 | 428331 | 0.56094772 | 3.6325469 | 5.36E-05 | 0.00257694 |
| ENSGALG00010003412 | ASB5 | 422572 | -0.639742 | 3.85612073 | 5.62E-05 | 0.00269544 |
| ENSGALG00010018627 | MYLK | 396445 | -0.593943 | 4.54316192 | 5.67E-05 | 0.00270992 |
| ENSGALG00010028434 | COL5A1 | 395568 | 0.54837681 | 4.13742043 | 5.78E-05 | 0.0027516 |
| ENSGALG00010017244 | NA | NA | 0.69793083 | 2.98271427 | 5.93E-05 | 0.00281668 |
| ENSGALG00010024660 | KMT2D | 425846 | 0.29094769 | 8.11326821 | 6.09E-05 | 0.00288509 |
| ENSGALG00010015793 | RSRP1 | 419592 | -0.4693137 | 5.16713126 | 6.19E-05 | 0.00292009 |
| ENSGALG00010006704 | NA | NA | 1.10893546 | 1.14108705 | 6.22E-05 | 0.00292777 |
| ENSGALG00010005260 | ANXA5 | 428767 | -0.5261628 | 4.2167645 | 6.39E-05 | 0.00299677 |
| ENSGALG00010003270 | LOC429798 | 429798 | 1.62879979 | 0.34940614 | 6.68E-05 | 0.00312383 |
| ENSGALG00010008056 | GTF2E2 | 771227 | -0.4195814 | 5.57179704 | 6.79E-05 | 0.00316571 |
| ENSGALG00010029850 | MYO15L | 417377 | 0.80304159 | 1.94942971 | 6.81E-05 | 0.00316571 |
| ENSGALG00010010280 | HSP90B1 | 374163 | -0.5779367 | 9.97879344 | 7.03E-05 | 0.00326169 |
| ENSGALG00010014989 | NA | NA | 1.2891633 | 0.8930372 | 7.13E-05 | 0.00329637 |
| ENSGALG00010011096 | LDB2 | 395631 | 0.81772818 | 3.77909163 | 7.16E-05 | 0.00330253 |
| ENSGALG00010009135 | TENM4 | 395406 | 0.49533867 | 3.83547664 | 7.33E-05 | 0.00336789 |
| ENSGALG00010004877 | NA | NA | 1.00805289 | 1.34159477 | 7.44E-05 | 0.00341073 |
| ENSGALG00010024778 | RAB5B | 100529061 | -0.3464979 | 5.84623923 | 7.60E-05 | 0.00346843 |
| ENSGALG00010025904 | RPL10A | 419895 | -0.3503866 | 8.606138 | 7.61E-05 | 0.00346843 |
| ENSGALG00010015110 | LOC422177 | 422177 | -1.4476336 | 0.43814531 | 7.75E-05 | 0.00351861 |
| ENSGALG00010005178 | EPHA5 | 395997 | -0.4521518 | 6.29071469 | 7.76E-05 | 0.00351861 |
| ENSGALG00010016487 | NA | NA | 0.62854199 | 2.79275392 | 7.83E-05 | 0.00352889 |
| ENSGALG00010008926 | NA | NA | 0.84111836 | 1.77753017 | 7.83E-05 | 0.00352889 |
| ENSGALG00010029582 | SCPEP1 | 417403 | 0.31576633 | 5.68770502 | 7.96E-05 | 0.00357502 |
| ENSGALG00010001115 | PFN2 | 771904 | -0.3020977 | 8.09195064 | 8.02E-05 | 0.00359406 |
| ENSGALG00010016418 | NFIB | 396209 | 0.7594507 | 2.9164088 | 8.09E-05 | 0.00361039 |
| ENSGALG00010025954 | TUBA1A | 429035 | -0.3971611 | 8.09329174 | 8.10E-05 | 0.00361039 |
| ENSGALG00010002433 | GRHL1 | 428651 | 0.39746001 | 5.12081029 | 8.17E-05 | 0.00362984 |
| ENSGALG00010025645 | NBEAL2 | 425984 | 0.30557698 | 6.50692143 | 8.62E-05 | 0.003818 |
| ENSGALG00010002873 | MRAP2 | 421836 | 0.54994521 | 3.67353758 | 8.91E-05 | 0.00393724 |

| | | | | | | |
|---|---|---|---|---|---|---|
| ENSGALG00010011043 | NA | NA | 1.08435919 | 1.08302657 | 8.94E-05 | 0.00394013 |
| ENSGALG00010005785 | RPL21 | 418933 | -0.3040675 | 8.9037975 | 8.98E-05 | 0.00394569 |
| ENSGALG00010003980 | CHCHD7 | 770709 | -0.4270977 | 5.60302167 | 9.08E-05 | 0.00397881 |
| ENSGALG00010007908 | IRX1 | 374185 | -1.1077324 | 1.3106939 | 9.12E-05 | 0.00398525 |
| ENSGALG00010003214 | PRDX4 | 418601 | -0.3780807 | 7.10801934 | 9.20E-05 | 0.00401125 |
| ENSGALG00010015870 | EIF5B | 395514 | -0.4435025 | 8.27158454 | 9.26E-05 | 0.00402293 |
| ENSGALG00010025862 | PTCH2 | 374187 | -0.543804 | 3.66799702 | 9.30E-05 | 0.00403229 |
| ENSGALG00010014839 | TM7SF3 | 418215 | 0.35327169 | 5.41443373 | 9.39E-05 | 0.00405798 |
| ENSGALG00010019399 | CD248 | 421234 | 0.97753734 | 1.93800802 | 9.46E-05 | 0.00406701 |
| ENSGALG00010010019 | NA | NA | -0.7602543 | 2.48064203 | 9.51E-05 | 0.00406701 |
| ENSGALG00010024370 | HERPUD1 | 415644 | -0.6467761 | 5.96677858 | 9.51E-05 | 0.00406701 |
| ENSGALG00010027702 | FOXP4 | 100858652 | 0.5919899 | 3.31317293 | 9.51E-05 | 0.00406701 |
| ENSGALG00010019709 | LOC124417823 | 124417823 | 0.57264372 | 3.43673609 | 9.78E-05 | 0.00416986 |
| ENSGALG00010016718 | LIN28A | 428206 | -0.3906901 | 8.41464096 | 9.81E-05 | 0.00417211 |
| ENSGALG00010000024 | COX3 | 63549467 | -0.4605637 | 12.4615936 | 9.91E-05 | 0.00420413 |
| ENSGALG00010012932 | ATF6B | 424365 | -0.4209497 | 4.9288252 | 9.99E-05 | 0.00422602 |
| ENSGALG00010002896 | OCA2 | 428009 | -1.4209297 | 0.3894849 | 0.00010042 | 0.00423287 |
| ENSGALG00010018030 | POLB | 426794 | -0.3667241 | 6.13698208 | 0.00010061 | 0.00423287 |
| ENSGALG00010021237 | FSCN1 | 416485 | -0.418656 | 7.64688492 | 0.00010174 | 0.00426932 |
| ENSGALG00010016426 | HELZ2 | 419265 | -1.0534927 | 3.62272781 | 0.00010295 | 0.00430858 |
| ENSGALG00010014836 | LOC112532960 | 112532960 | -1.4217117 | 0.67922338 | 0.00010547 | 0.00440243 |
| ENSGALG00010016551 | NA | NA | 0.45751136 | 5.06698525 | 0.00010721 | 0.00446356 |
| ENSGALG00010025092 | COL1A1 | 395532 | 0.28929462 | 6.05546153 | 0.00010781 | 0.00447658 |
| ENSGALG00010026821 | NA | NA | 1.37237483 | 0.40237147 | 0.00010998 | 0.00455494 |
| ENSGALG00010002335 | TMEM47 | 418585 | -0.4477666 | 5.14970223 | 0.00011565 | 0.00477708 |
| ENSGALG00010024688 | CADM1 | 419762 | 0.28127373 | 8.04970668 | 0.00011808 | 0.00486477 |
| ENSGALG00010000588 | NA | NA | -0.5607349 | 11.8406794 | 0.00012072 | 0.00496096 |
| ENSGALG00010019880 | LOC112532332 | 112532332 | 0.62166154 | 5.0860017 | 0.00012208 | 0.00500371 |
| ENSGALG00010028445 | SYT11 | 429166 | -0.2827436 | 7.89416402 | 0.00012431 | 0.0050818 |
| ENSGALG00010015716 | MGAT4A | 418692 | 0.45806259 | 4.56240033 | 0.00012684 | 0.00517218 |
| ENSGALG00010021861 | NA | NA | 0.31477788 | 5.65070608 | 0.00012725 | 0.00517527 |
| ENSGALG00010001773 | NRCAM | 396202 | 0.98569406 | 1.1815826 | 0.00012839 | 0.00520425 |
| ENSGALG00010025481 | TMEM39A | 418340 | -0.395341 | 5.92744672 | 0.00012861 | 0.00520425 |
| ENSGALG00010027375 | FUT7 | 427782 | 1.34079162 | 0.75042308 | 0.000131 | 0.00528718 |
| ENSGALG00010021825 | ONECUT1 | 107054176 | 0.61105065 | 2.86797935 | 0.00013202 | 0.00531512 |
| ENSGALG00010026636 | NA | NA | 1.81444326 | -0.1554493 | 0.0001344 | 0.00539721 |
| ENSGALG00010006735 | NR3C2 | 374131 | 0.38959618 | 4.73454635 | 0.00013986 | 0.0055945 |
| ENSGALG00010014338 | BHLHE41 | 101750689 | 1.32996833 | 0.5432672 | 0.00014002 | 0.0055945 |
| ENSGALG00010003557 | LOC121107015 | 121107015 | 0.35887623 | 5.37598169 | 0.00014069 | 0.0056072 |
| ENSGALG00010019774 | NA | NA | 0.55926135 | 3.78116016 | 0.00014467 | 0.00575107 |
| ENSGALG00010016967 | ERLEC1 | 101751309 | -0.5723961 | 6.08185238 | 0.0001461 | 0.00579342 |
| ENSGALG00010016797 | SEC62 | 424993 | -0.3158447 | 7.58903148 | 0.00015194 | 0.00601017 |
| ENSGALG00010015047 | NPY2R | 422405 | 0.4097671 | 4.54752505 | 0.00015349 | 0.00605627 |
| ENSGALG00010004446 | ACAT2 | 421587 | -0.2855169 | 6.05372513 | 0.00015517 | 0.00610741 |
| ENSGALG00010019461 | MGAT5 | 424286 | 0.27622913 | 6.12216887 | 0.0001574 | 0.00617996 |
| ENSGALG00010008103 | RBPMS | 428738 | -0.2767327 | 6.56980755 | 0.00015819 | 0.00619552 |
| ENSGALG00010010832 | NA | NA | 0.69678802 | 2.28634071 | 0.00015931 | 0.00622408 |
| ENSGALG00010022048 | NA | NA | 0.84722959 | 2.11355468 | 0.0001667 | 0.00649686 |
| ENSGALG00010016433 | TMED3 | 415580 | -0.7422559 | 3.69235861 | 0.00016828 | 0.00654236 |
| ENSGALG00010003610 | NA | NA | 0.99534629 | 1.73540589 | 0.00016894 | 0.00655186 |
| ENSGALG00010010406 | PTCH1 | 395806 | -0.4941938 | 3.67440466 | 0.00017119 | 0.00662299 |
| ENSGALG00010011097 | NA | NA | 1.67879389 | 0.02700007 | 0.00017327 | 0.00667228 |
| ENSGALG00010023807 | MRPL10 | 426521 | -0.327725 | 5.0515657 | 0.00017331 | 0.00667228 |
| ENSGALG00010001877 | OAT | 426430 | -0.538195 | 6.27497813 | 0.00017534 | 0.00673435 |
| ENSGALG00010020723 | RPS24 | 423726 | -0.3200994 | 8.9210273 | 0.00017614 | 0.00674844 |
| ENSGALG00010009863 | NA | NA | -1.5859431 | 0.09163718 | 0.00018236 | 0.00697009 |
| ENSGALG00010002544 | THSD1 | 418884 | 0.54461339 | 3.18114426 | 0.00018432 | 0.00702793 |
| ENSGALG00010018099 | SMAD3 | 395132 | 0.30432371 | 7.40531416 | 0.00018712 | 0.00710973 |
| ENSGALG00010027060 | LOC112532950 | 112532950 | -1.8040912 | -0.6777255 | 0.00018736 | 0.00710973 |
| ENSGALG00010022656 | FKBP10 | 427013 | -0.4799218 | 4.93389398 | 0.00019129 | 0.00722908 |

| | | | | | | |
|---|---|---|---|---|---|---|
| ENSGALG00010007717 | GALR1 | 428512 | 1.86760497 | -0.475799 | 0.00019141 | 0.00722908 |
| ENSGALG00010014439 | LOC112532140 | 112532140 | -0.9975228 | 3.93183918 | 0.00019259 | 0.00725621 |
| ENSGALG00010023913 | ZMYND8 | 419207 | 0.26662329 | 7.15919435 | 0.00019553 | 0.00734957 |
| ENSGALG00010026199 | TBX6 | 395792 | 0.27152655 | 8.77933077 | 0.00019908 | 0.00746528 |
| ENSGALG00010008148 | ABCD2 | 417694 | 0.48709282 | 3.57604112 | 0.00020501 | 0.00766953 |
| ENSGALG00010017195 | NA | NA | 0.76933644 | 2.68248986 | 0.00020688 | 0.00772139 |
| ENSGALG00010016377 | FRMPD3 | 428699 | 0.82152281 | 2.32920602 | 0.00020849 | 0.00775812 |
| ENSGALG00010003558 | TPT1 | 396363 | -0.332182 | 9.43168539 | 0.00020885 | 0.00775812 |
| ENSGALG00010029328 | PITPNC1 | 100859655 | 0.32235099 | 6.66602957 | 0.00020971 | 0.00777207 |
| ENSGALG00010016447 | ID3 | 395281 | -0.4640869 | 4.52521422 | 0.00021285 | 0.00786981 |
| ENSGALG00010004852 | LOC426820 | 426820 | 0.76288593 | 1.84136877 | 0.0002153 | 0.00794215 |
| ENSGALG00010003305 | CALR | 100859104 | -0.4761592 | 9.22834298 | 0.00021675 | 0.00797712 |
| ENSGALG00010009076 | TMEM214 | 428658 | -0.4000524 | 6.12301028 | 0.00022064 | 0.00810136 |
| ENSGALG00010021516 | SRSF5 | 423258 | -0.3441271 | 6.1875357 | 0.00022289 | 0.00816522 |
| ENSGALG00010016127 | ACSL6 | 416324 | -0.3155769 | 7.06320775 | 0.00022467 | 0.00821147 |
| ENSGALG00010025565 | LOC424577 | 424577 | 0.90145156 | 1.90707614 | 0.000226 | 0.00824084 |
| ENSGALG00010005991 | FARP1 | 418781 | 0.36900522 | 5.35231463 | 0.00022665 | 0.00824285 |
| ENSGALG00010010900 | SLC34A2 | 395131 | 0.2679785 | 8.34725275 | 0.00022709 | 0.00824285 |
| ENSGALG00010027548 | PCBP4 | 100859674 | -0.3097445 | 6.27660094 | 0.00023242 | 0.00841716 |
| ENSGALG00010023739 | NA | NA | 0.54596735 | 3.27393517 | 0.00023488 | 0.00848659 |
| ENSGALG00010023642 | PAX6 | 395943 | -0.6765186 | 2.68723648 | 0.00023678 | 0.00853595 |
| ENSGALG00010022240 | SPINT1 | 423206 | 0.36862257 | 4.72648962 | 0.00023948 | 0.00861378 |
| ENSGALG00010010504 | NA | NA | 0.88214095 | 1.91632155 | 0.00024657 | 0.00884861 |
| ENSGALG00010002918 | FAM84A | 421946 | -0.3856017 | 4.42358819 | 0.000249 | 0.00891564 |
| ENSGALG00010010420 | NPM1 | 396203 | -0.2843413 | 10.7029299 | 0.00025597 | 0.00914444 |
| ENSGALG00010019068 | SNRPA1 | 415523 | -0.31855 | 6.55056615 | 0.00025722 | 0.00916863 |
| ENSGALG00010012682 | SLC5A3 | 112530700 | 0.28303554 | 5.84980523 | 0.0002586 | 0.00919725 |
| ENSGALG00010021158 | ACTG1 | 415296 | -0.268793 | 8.74812632 | 0.00025923 | 0.00919876 |
| ENSGALG00010019726 | PEBP4 | 101747939 | 1.19436268 | 0.58187149 | 0.00025994 | 0.00920339 |
| ENSGALG00010026014 | SOX14 | 395526 | 0.77026436 | 1.99995993 | 0.00026551 | 0.00937969 |
| ENSGALG00010028760 | SSR2 | 425656 | -0.3797661 | 7.03871251 | 0.00026682 | 0.00940525 |
| ENSGALG00010023863 | EYA2 | 395745 | 0.92052673 | 1.61400987 | 0.00027247 | 0.00958307 |
| ENSGALG00010028055 | LOC121107542 | 121107542 | -0.5225682 | 4.91307461 | 0.00027567 | 0.00967416 |
| ENSGALG00010012610 | RPL3 | 418016 | -0.2627445 | 9.58893683 | 0.00027724 | 0.00970783 |
| ENSGALG00010016086 | TDRD9 | 423488 | 0.27446776 | 7.86019884 | 0.00027919 | 0.00972604 |
| ENSGALG00010014213 | ETFRF1 | 776112 | 0.64904466 | 2.74536778 | 0.00027982 | 0.00972604 |
| ENSGALG00010028497 | ADMP | 395622 | 0.38417102 | 6.69182649 | 0.00027993 | 0.00972604 |
| ENSGALG00010020078 | LOC112531986 | 112531986 | -1.4712656 | 0.08811746 | 0.00028022 | 0.00972604 |
| ENSGALG00010005229 | CIPC | 771996 | 0.25191858 | 7.68628168 | 0.00028263 | 0.00978821 |
| ENSGALG00010022900 | MMP17 | 416801 | 0.38372551 | 5.64861697 | 0.00028402 | 0.00981518 |
| ENSGALG00010008321 | ZFHX3 | 395682 | 0.24583925 | 8.179897 | 0.00029402 | 0.0101386 |
| ENSGALG00010000324 | LOC100857850 | 100857850 | 0.31708242 | 5.88872548 | 0.00029634 | 0.01019654 |
| ENSGALG00010029218 | RPL23A | 417574 | -0.3623853 | 8.77427428 | 0.00029852 | 0.01024919 |
| ENSGALG00010005309 | ASXL3 | 421102 | 0.39982622 | 4.32765395 | 0.00030511 | 0.01045266 |
| ENSGALG00010000486 | NA | NA | -1.4458356 | 2.91119866 | 0.00030946 | 0.01057896 |
| ENSGALG00010013452 | LSS | 424037 | -0.2804378 | 6.72087841 | 0.0003142 | 0.01071805 |
| ENSGALG00010029330 | AFMID | 769005 | -0.3462405 | 4.95636599 | 0.00032096 | 0.01092504 |
| ENSGALG00010007369 | NA | NA | -0.3650837 | 5.03549094 | 0.000322 | 0.01093693 |
| ENSGALG00010014867 | TP63 | 374269 | 0.41586555 | 4.32237332 | 0.00032511 | 0.01101357 |
| ENSGALG00010000559 | NA | NA | -0.716411 | 3.20828538 | 0.00032564 | 0.01101357 |
| ENSGALG00010013226 | DUSP6 | 374272 | -0.273449 | 7.07691111 | 0.00032683 | 0.01103024 |
| ENSGALG00010017532 | MIXL1 | 395838 | 1.10417054 | 3.08449608 | 0.00032964 | 0.01110145 |
| ENSGALG00010023104 | EFCAB4B | 423106 | 0.27319182 | 6.2283389 | 0.00033164 | 0.01114112 |
| ENSGALG00010024728 | SLC5A9 | 424625 | 0.87147231 | 1.57932644 | 0.00033223 | 0.01114112 |
| ENSGALG00010016490 | NA | NA | 0.44484511 | 4.11822168 | 0.00033394 | 0.01117512 |
| ENSGALG00010017751 | ST3GAL5 | 407775 | -0.311476 | 6.10002961 | 0.00033515 | 0.01119177 |
| ENSGALG00010012654 | FAM46B | 430080 | 0.63808233 | 4.7037214 | 0.00033666 | 0.01121874 |
| ENSGALG00010010094 | SLC2A9 | 422843 | 0.2940687 | 6.41515359 | 0.00033914 | 0.01127778 |
| ENSGALG00010010115 | NEB | 101748402 | 0.30277405 | 7.48700388 | 0.00033994 | 0.0112805 |
| ENSGALG00010010824 | PKP4 | 424328 | -0.2651173 | 6.45127726 | 0.00034764 | 0.01151191 |

| | | | | | | |
|---|---|---|---|---|---|---|
| ENSGALG00010016773 | RPS27A | 395796 | -0.3161973 | 8.65936304 | 0.00034897 | 0.01152832 |
| ENSGALG00010019752 | GPAM | 423895 | 0.28831076 | 7.22066842 | 0.00034959 | 0.01152832 |
| ENSGALG00010017205 | NA | NA | 0.59761072 | 3.06683733 | 0.00035182 | 0.01157806 |
| ENSGALG00010018573 | SEC23B | 416724 | -0.3030114 | 6.40864678 | 0.00035776 | 0.01174917 |
| ENSGALG00010006806 | NA | NA | 1.85224187 | -0.5110549 | 0.00035859 | 0.01175193 |
| ENSGALG00010019800 | EXT2 | 425859 | -0.303098 | 8.59352069 | 0.00036004 | 0.01177527 |
| ENSGALG00010007237 | NA | NA | 1.45414675 | 0.2204463 | 0.0003639 | 0.01187692 |
| ENSGALG00010022441 | FICD | 416889 | -0.640586 | 3.38728596 | 0.00037124 | 0.01209157 |
| ENSGALG00010005236 | TMTC4 | 418772 | 0.26791126 | 7.27515822 | 0.00037323 | 0.01213148 |
| ENSGALG00010020973 | GNB5 | 415417 | -0.4912507 | 3.65420581 | 0.0003749 | 0.01216073 |
| ENSGALG00010026753 | NA | NA | 0.8583295 | 1.6309487 | 0.00037798 | 0.01223576 |
| ENSGALG00010022246 | RHOV | 428868 | 0.37900316 | 5.10411761 | 0.00037989 | 0.01227253 |
| ENSGALG00010004477 | TULP4 | 421658 | 0.30758201 | 7.26245802 | 0.00038343 | 0.01236184 |
| ENSGALG00010005265 | DYNC2H1 | 418979 | 0.38172939 | 4.73535448 | 0.00038469 | 0.01237714 |
| ENSGALG00010016800 | RPL14 | 374134 | -0.301723 | 8.16396602 | 0.00038592 | 0.01239158 |
| ENSGALG00010021617 | SORBS1 | 423749 | 0.32358035 | 5.52525632 | 0.00039165 | 0.01255005 |
| ENSGALG00010001132 | SELENOT | 425041 | -0.3371177 | 7.54179427 | 0.00039261 | 0.01255547 |
| ENSGALG00010025376 | LOC112529918 | 112529918 | -1.5141988 | 0.19788418 | 0.0003934 | 0.0125555 |
| ENSGALG00010019820 | BBOX1 | 426932 | 0.38138363 | 5.7757128 | 0.00039492 | 0.01257875 |
| ENSGALG00010001267 | BCHE | 395358 | 0.36377224 | 4.79742685 | 0.0003994 | 0.01268715 |
| ENSGALG00010016868 | NIPAL1 | 428783 | -0.5619009 | 5.3505887 | 0.00039992 | 0.01268715 |
| ENSGALG00010014291 | OSTN | 424907 | -1.6310689 | 1.27477372 | 0.00040213 | 0.01273182 |
| ENSGALG00010014778 | TMPRSS6 | 418046 | 1.74419263 | 0.08865298 | 0.00040341 | 0.01274691 |
| ENSGALG00010006410 | ADM2 | 112532887 | -0.8297799 | 3.08771544 | 0.00040482 | 0.01276605 |
| ENSGALG00010013720 | SLC16A10 | 421753 | -0.3177621 | 6.19997643 | 0.00040664 | 0.01279786 |
| ENSGALG00010014603 | TSC22D4 | 768091 | -1.0872575 | 3.3813545 | 0.00040873 | 0.01283826 |
| ENSGALG00010019243 | SERTAD2 | 421276 | 0.33825433 | 4.67353052 | 0.0004101 | 0.01285572 |
| ENSGALG00010001972 | FAM204A | 423925 | -0.2915307 | 5.68117818 | 0.00041504 | 0.0129848 |
| ENSGALG00010010986 | NARS | 426856 | -0.2582457 | 7.43860785 | 0.00041991 | 0.01308645 |
| ENSGALG00010012638 | NA | NA | -0.2996803 | 5.48330088 | 0.00042013 | 0.01308645 |
| ENSGALG00010002441 | USO1 | 422575 | -0.3216605 | 7.86727615 | 0.00042076 | 0.01308645 |
| ENSGALG00010018936 | SYT14 | 428566 | 0.63226283 | 3.03618814 | 0.00042258 | 0.01310932 |
| ENSGALG00010001268 | CBLN2 | 421019 | -0.3898748 | 4.2459944 | 0.00042315 | 0.01310932 |
| ENSGALG00010007529 | CDH13 | 414849 | -0.6068751 | 6.31151119 | 0.00042456 | 0.01311952 |
| ENSGALG00010024396 | NA | NA | -0.8225754 | 2.32891492 | 0.00042513 | 0.01311952 |
| ENSGALG00010015829 | SMAD5 | 395679 | 0.249645 | 8.15145898 | 0.00043196 | 0.01329425 |
| ENSGALG00010000039 | PDIA3 | 373899 | -0.5296425 | 9.54432525 | 0.00043247 | 0.01329425 |
| ENSGALG00010011019 | NECAB1 | 420219 | 0.95215331 | 1.76487665 | 0.00043684 | 0.01340042 |
| ENSGALG00010004429 | LOC112531814 | 112531814 | -1.5750023 | 1.56226626 | 0.00043761 | 0.01340042 |
| ENSGALG00010019989 | SLC6A14 | 100857521 | 1.05795681 | 0.85991804 | 0.00044139 | 0.01349015 |
| ENSGALG00010022184 | PRKAR1B | 416446 | 0.52852242 | 3.70336959 | 0.00044615 | 0.0136093 |
| ENSGALG00010022583 | FKBP11 | 107049506 | -1.1281478 | 4.02001337 | 0.00044774 | 0.0136316 |
| ENSGALG00010015325 | NA | NA | -0.327982 | 5.79864927 | 0.00045499 | 0.01382506 |
| ENSGALG00010009191 | NA | NA | 0.66441312 | 2.89633764 | 0.00045632 | 0.01382506 |
| ENSGALG00010008211 | NA | NA | 0.29134676 | 5.50882166 | 0.00045723 | 0.01382506 |
| ENSGALG00010013325 | CA2 | 396257 | -1.2867767 | 2.82933436 | 0.00045758 | 0.01382506 |
| ENSGALG00010007935 | LARP1B | 422500 | -0.2897117 | 5.57275775 | 0.00045875 | 0.01383402 |
| ENSGALG00010027114 | FLNB | 378913 | 0.24973312 | 7.74285281 | 0.00046311 | 0.01393884 |
| ENSGALG00010024561 | ROBO3 | 100858578 | -0.6062334 | 4.70880995 | 0.00046626 | 0.01400725 |
| ENSGALG00010016677 | MEAF6 | 419617 | -0.2559251 | 6.88409996 | 0.00047318 | 0.01418819 |
| ENSGALG00010017558 | ANP32A | 415562 | -0.2493165 | 7.18693193 | 0.0004767 | 0.01426673 |
| ENSGALG00010019801 | FIBIN | 426933 | 0.34611698 | 6.79012126 | 0.00048148 | 0.0143737 |
| ENSGALG00010017918 | SIX5 | 395601 | -0.2952566 | 5.5233763 | 0.00048208 | 0.0143737 |
| ENSGALG00010003474 | RPS7 | 421919 | -0.297012 | 8.55329015 | 0.00048355 | 0.01439031 |
| ENSGALG00010009172 | SPOPL | 424299 | 0.27436364 | 7.82418483 | 0.00048995 | 0.0145534 |
| ENSGALG00010018983 | MDGA2 | 423563 | 0.25395609 | 6.60138749 | 0.00049337 | 0.01462779 |
| ENSGALG00010013389 | NAP1L1 | 417864 | -0.2388806 | 9.55769298 | 0.00049822 | 0.01474406 |
| ENSGALG00010024664 | DNAH10 | 416818 | 0.25379429 | 6.58921049 | 0.00049969 | 0.0147598 |
| ENSGALG00010018474 | LOC112532599 | 112532599 | -1.33595 | -0.1794686 | 0.00050327 | 0.01483796 |
| ENSGALG00010000022 | ATP8 | 63549490 | -0.5480984 | 7.82862148 | 0.00050548 | 0.01487568 |

| | | | | | | |
|---|---|---|---|---|---|---|
| ENSGALG00010014014 | PARP8 | 427198 | -0.3778441 | 4.41717513 | 0.00050999 | 0.01495963 |
| ENSGALG00010005442 | SASH1 | 421621 | 0.30211967 | 5.35971551 | 0.00051022 | 0.01495963 |
| ENSGALG00010028547 | ZNF618 | 770984 | 0.28644444 | 6.25228729 | 0.0005148 | 0.01506606 |
| ENSGALG00010028992 | NA | NA | 0.91376775 | 1.42190096 | 0.0005178 | 0.01512579 |
| ENSGALG00010020282 | MAGI1 | 416083 | 0.27147607 | 7.75121881 | 0.00052339 | 0.01526108 |
| ENSGALG00010020928 | AACS | 416811 | -0.3377244 | 4.78995431 | 0.00052486 | 0.01527593 |
| ENSGALG00010024330 | SDF2L1 | 416770 | -0.4186703 | 5.57751724 | 0.00052773 | 0.01533136 |
| ENSGALG00010005914 | NA | NA | 0.42535056 | 4.01691525 | 0.00052915 | 0.01534429 |
| ENSGALG00010017380 | MARCKSL1 | 770764 | -0.3522092 | 8.02742537 | 0.00053103 | 0.01537061 |
| ENSGALG00010029528 | TBL2 | 101751077 | -0.3039163 | 5.12372792 | 0.00053283 | 0.01537061 |
| ENSGALG00010005245 | KIAA1217 | 420501 | 0.35807149 | 5.11410059 | 0.00053296 | 0.01537061 |
| ENSGALG00010004701 | NA | NA | 0.66462108 | 2.92683377 | 0.000536 | 0.01543013 |
| ENSGALG00010014729 | MRPS15 | 770261 | -0.2914212 | 6.83320702 | 0.00054052 | 0.01553198 |
| ENSGALG00010000553 | NA | NA | -0.5466902 | 3.93404051 | 0.00054203 | 0.01554742 |
| ENSGALG00010012149 | SFXN1 | 416221 | -0.5519994 | 6.29214735 | 0.00054563 | 0.01562239 |
| ENSGALG00010028806 | SPACA9 | 772069 | 0.76914475 | 1.53640985 | 0.0005576 | 0.01593632 |
| ENSGALG00010024726 | TMED2 | 416821 | -0.4430063 | 8.64491439 | 0.00056673 | 0.01616821 |
| ENSGALG00010029190 | DHRS7C | 417304 | 1.29587082 | 0.54189909 | 0.00056806 | 0.01617707 |
| ENSGALG00010020074 | LOC421419 | 421419 | 0.25742861 | 6.35618826 | 0.0005755 | 0.01634231 |
| ENSGALG00010011029 | HMGCR | 395145 | -0.3948904 | 8.25636373 | 0.00057593 | 0.01634231 |
| ENSGALG00010022482 | PRTG | 415407 | 0.30922462 | 7.03639436 | 0.00058708 | 0.01661984 |
| ENSGALG00010010396 | NA | NA | -0.3891658 | 4.54122793 | 0.0005878 | 0.01661984 |
| ENSGALG00010011363 | FDFT1 | 422038 | -0.3336399 | 10.2079814 | 0.00058903 | 0.01662481 |
| ENSGALG00010000026 | ND3 | 63549491 | -0.3868698 | 8.76629046 | 0.00059217 | 0.01665835 |
| ENSGALG00010028854 | RPL7A | 417158 | -0.2941189 | 9.06145368 | 0.00059232 | 0.01665835 |
| ENSGALG00010028309 | NA | NA | 0.53603753 | 3.04073243 | 0.00059386 | 0.01667222 |
| ENSGALG00010001744 | NA | NA | 1.0709454 | 0.67193763 | 0.00059786 | 0.01675473 |
| ENSGALG00010023429 | BMF | 769140 | 0.48043958 | 5.46409942 | 0.00060182 | 0.0168362 |
| ENSGALG00010027349 | NA | NA | 1.24260017 | 0.35962452 | 0.00060493 | 0.01689318 |
| ENSGALG00010013372 | SPRY4 | 427643 | -0.5483443 | 2.81898146 | 0.00060736 | 0.01693136 |
| ENSGALG00010002492 | RPL8 | 418568 | -0.272558 | 9.38640614 | 0.00062377 | 0.01735824 |
| ENSGALG00010011059 | ANKRD31 | 427208 | 0.2652949 | 6.84753536 | 0.00062873 | 0.01746555 |
| ENSGALG00010004835 | HHLA2 | 107054979 | -0.5743683 | 4.42811609 | 0.00063697 | 0.01766362 |
| ENSGALG00010015733 | COL4A6 | 422350 | 0.25825478 | 7.59705833 | 0.00063901 | 0.01768927 |
| ENSGALG00010008932 | VASP | 107057642 | 0.4275172 | 3.90436826 | 0.00064138 | 0.01772411 |
| ENSGALG00010011875 | PSD2 | 427605 | 0.39416009 | 4.10522397 | 0.00064307 | 0.01773965 |
| ENSGALG00010017832 | POU3F1 | 430270 | -0.6655685 | 2.71634061 | 0.00065831 | 0.01809764 |
| ENSGALG00010000943 | NA | NA | 1.51556213 | -0.0638564 | 0.00065832 | 0.01809764 |
| ENSGALG00010002880 | RASGEF1B | 100859744 | 0.32094288 | 6.72135707 | 0.00066192 | 0.01816493 |
| ENSGALG00010007751 | HSPA13 | 418469 | -0.448187 | 4.14222025 | 0.0006638 | 0.01818129 |
| ENSGALG00010008298 | LOC420992 | 420992 | 0.28667034 | 5.50085611 | 0.00066749 | 0.01818129 |
| ENSGALG00010010185 | PHYHIPL | 423603 | 0.73214818 | 1.74938411 | 0.00066766 | 0.01818129 |
| ENSGALG00010029229 | CBX4 | 395354 | -0.9056918 | 2.65400836 | 0.00066809 | 0.01818129 |
| ENSGALG00010028368 | TIMM13 | 100859352 | -0.3485197 | 4.74405538 | 0.00066829 | 0.01818129 |
| ENSGALG00010009656 | NA | NA | 0.629337 | 4.13294172 | 0.00066939 | 0.01818129 |
| ENSGALG00010002843 | NA | NA | -0.8192644 | 2.39978484 | 0.00067415 | 0.01825844 |
| ENSGALG00010013151 | LOC107055505 | 107055505 | 0.93453161 | 3.97847513 | 0.00067453 | 0.01825844 |
| ENSGALG00010024872 | RUNDC3A | 100858314 | -0.4338688 | 4.36457456 | 0.00068135 | 0.01838081 |
| ENSGALG00010027267 | MYO9B | 420135 | -0.2851286 | 7.55851147 | 0.00068137 | 0.01838081 |
| ENSGALG00010018867 | RPS2 | 416544 | -0.2466752 | 9.78592274 | 0.00069 | 0.01858202 |
| ENSGALG00010023533 | FKBP7 | 424129 | -0.5860605 | 2.63493801 | 0.00069522 | 0.0186908 |
| ENSGALG00010028054 | RPS15 | 396448 | -0.2991216 | 8.61929988 | 0.00070104 | 0.01880611 |
| ENSGALG00010026005 | DERL3 | 416934 | -0.8338207 | 5.74493896 | 0.00070188 | 0.01880611 |
| ENSGALG00010017421 | NA | NA | 0.36446936 | 4.49431487 | 0.00070615 | 0.01887949 |
| ENSGALG00010023620 | LOC107051846 | 107051846 | 0.28398681 | 5.70115626 | 0.000707 | 0.01887949 |
| ENSGALG00010019093 | RPL32 | 416122 | -0.2922207 | 8.1885406 | 0.00071242 | 0.01899223 |
| ENSGALG00010028070 | MEX3A | 101747646 | 0.31682124 | 5.30694186 | 0.00072979 | 0.01942274 |
| ENSGALG00010008444 | SLC7A11 | 428731 | -1.1552624 | 3.15779701 | 0.00073433 | 0.0195107 |
| ENSGALG00010004925 | NA | NA | 1.34701362 | 0.0843956 | 0.00074085 | 0.0196512 |
| ENSGALG00010026562 | NA | NA | 0.94274499 | 1.29869613 | 0.00074524 | 0.01973464 |

| | | | | | | |
|---|---|---|---|---|---|---|
| ENSGALG00010013170 | COL3A1 | 396340 | 0.25223399 | 7.40065971 | 0.00075273 | 0.01989968 |
| ENSGALG00010001712 | ATP6AP1 | 427854 | 0.43451335 | 4.37389651 | 0.00075469 | 0.01991823 |
| ENSGALG00010028603 | SH3GLB2 | 417197 | -0.4843863 | 3.16897742 | 0.00076754 | 0.02015871 |
| ENSGALG00010015881 | GAB3 | 428717 | 0.50688403 | 3.14004728 | 0.00076768 | 0.02015871 |
| ENSGALG00010016845 | MIA3 | 421337 | -0.3035899 | 6.31433885 | 0.00076809 | 0.02015871 |
| ENSGALG00010028525 | TM6SF2 | 425719 | 1.54214779 | -0.2283205 | 0.00076888 | 0.02015871 |
| ENSGALG00010015088 | LOC107050476 | 107050476 | -1.2033024 | 1.80931709 | 0.0007755 | 0.02029853 |
| ENSGALG00010023386 | NEUROD4 | 395959 | -1.2708586 | 0.34691141 | 0.00078105 | 0.02041033 |
| ENSGALG00010023238 | NUCB2 | 423071 | -0.3331502 | 7.31061271 | 0.00078618 | 0.02048133 |
| ENSGALG00010029652 | LRRC59 | 422103 | -0.3517644 | 4.83008333 | 0.00078764 | 0.02048133 |
| ENSGALG00010010753 | LOC107051636 | 107051636 | -1.1975799 | 2.12800744 | 0.00078764 | 0.02048133 |
| ENSGALG00010010547 | BST1 | 422828 | 0.8907017 | 1.32400211 | 0.00079847 | 0.02069193 |
| ENSGALG00010015852 | ZFAND5 | 427254 | -0.2611282 | 7.88743574 | 0.00079876 | 0.02069193 |
| ENSGALG00010005444 | ACTC1 | 423298 | -0.7365596 | 2.18864567 | 0.00079966 | 0.02069193 |
| ENSGALG00010017895 | MNAT1 | 423521 | -0.2814375 | 6.24528937 | 0.00080512 | 0.0207994 |
| ENSGALG00010008142 | CD247 | 395911 | 1.00275379 | 0.86638692 | 0.00080802 | 0.02084024 |
| ENSGALG00010010858 | LHX4 | 776024 | 0.97606766 | 2.38216725 | 0.00081047 | 0.02086957 |
| ENSGALG00010029868 | NA | NA | 0.59873023 | 2.74534156 | 0.00081651 | 0.02099098 |
| ENSGALG00010026809 | NA | NA | 0.83916892 | 1.55766698 | 0.00081955 | 0.02101008 |
| ENSGALG00010017791 | CCNL2 | 395124 | -0.2737432 | 6.84732801 | 0.0008199 | 0.02101008 |
| ENSGALG00010004890 | ACAT1 | 418968 | -0.2334348 | 6.55294432 | 0.00082471 | 0.02109919 |
| ENSGALG00010012978 | HSD17B7 | 424367 | -0.5028643 | 4.53268437 | 0.00084423 | 0.02153891 |
| ENSGALG00010002533 | LPAR2 | 426086 | -0.350957 | 4.93768837 | 0.00084461 | 0.02153891 |
| ENSGALG00010025381 | MRPL37 | 772313 | -0.2765731 | 5.78466181 | 0.00085251 | 0.0217054 |
| ENSGALG00010009346 | STC2 | 416208 | -0.4364078 | 8.10447237 | 0.00085551 | 0.02174699 |
| ENSGALG00010023823 | NA | NA | 0.28618825 | 5.5670131 | 0.00085802 | 0.02174757 |
| ENSGALG00010012001 | FAM102B | 768922 | 0.5261104 | 3.60284335 | 0.00085943 | 0.02174757 |
| ENSGALG00010011357 | UNC80 | 424003 | 0.32745137 | 5.03816834 | 0.00086101 | 0.02174757 |
| ENSGALG00010021328 | NA | NA | -0.986748 | 1.07859153 | 0.00086102 | 0.02174757 |
| ENSGALG00010013866 | KCTD8 | 770600 | -0.3467618 | 4.81546267 | 0.00086432 | 0.02179612 |
| ENSGALG00010009357 | LYST | 421514 | 0.3032019 | 5.32584222 | 0.00087793 | 0.02210436 |
| ENSGALG00010012780 | FABP7 | 396246 | -0.4161135 | 5.47726843 | 0.00088106 | 0.02212652 |
| ENSGALG00010027940 | ZBTB7B | 121107583 | -0.8334041 | 2.24236917 | 0.0008816 | 0.02212652 |
| ENSGALG00010023822 | CHEK2 | 768913 | -0.3329863 | 5.3383942 | 0.00089585 | 0.02244865 |
| ENSGALG00010024776 | MPHOSPH10 | 415384 | -0.2601611 | 6.20935185 | 0.00090213 | 0.02254365 |
| ENSGALG00010016947 | CANX | 416288 | -0.2696928 | 9.57989428 | 0.00090306 | 0.02254365 |
| ENSGALG00010005524 | STRIP2 | 416676 | 0.64764307 | 3.32533257 | 0.00090391 | 0.02254365 |
| ENSGALG00010012938 | UPF3B | 100858231 | -0.2546118 | 6.76698073 | 0.00091152 | 0.02269779 |
| ENSGALG00010010826 | NCOA1 | 422016 | 0.26729753 | 6.65005356 | 0.00092218 | 0.0229273 |
| ENSGALG00010000450 | NA | NA | 0.42824259 | 5.20153604 | 0.00092516 | 0.02293199 |
| ENSGALG00010020554 | ZFP36L1 | 101751828 | -0.2457957 | 7.62268433 | 0.00092526 | 0.02293199 |
| ENSGALG00010029588 | RAB11FIP4 | 427808 | 0.24097881 | 6.45191709 | 0.00093255 | 0.02306915 |
| ENSGALG00010006748 | FAM177A1 | 423323 | -0.3489638 | 5.68206662 | 0.0009337 | 0.02306915 |
| ENSGALG00010028019 | RPL29 | 770249 | -0.3130112 | 6.79338227 | 0.00093971 | 0.02318147 |
| ENSGALG00010005629 | NRP1 | 395560 | 0.28146557 | 5.34148363 | 0.00094291 | 0.02320248 |
| ENSGALG00010006608 | KL | 418909 | -0.8693957 | 1.00948478 | 0.00094773 | 0.02320248 |
| ENSGALG00010025407 | MLXIP | 416857 | 0.24585819 | 6.35143603 | 0.00094843 | 0.02320248 |
| ENSGALG00010024181 | FOXD3 | 395794 | -0.4695942 | 4.06172844 | 0.00094856 | 0.02320248 |
| ENSGALG00010020978 | BCL2L10 | 395193 | -0.2584613 | 6.35725921 | 0.00094897 | 0.02320248 |
| ENSGALG00010000019 | NA | NA | -1.811829 | 0.60078431 | 0.00095018 | 0.02320248 |
| ENSGALG00010026166 | WNT9A | 395829 | 0.56646581 | 2.57423186 | 0.0009508 | 0.02320248 |
| ENSGALG00010011873 | RFX3 | 427351 | 0.46450293 | 3.62354078 | 0.0009551 | 0.02327146 |
| ENSGALG00010015472 | LGMN | 423418 | 0.32688603 | 6.65229097 | 0.00096363 | 0.02344338 |
| ENSGALG00010002102 | PAK1 | 419083 | 0.29578598 | 6.26334641 | 0.00096694 | 0.02348788 |
| ENSGALG00010017109 | ITGB1BP3 | 415448 | -0.2626613 | 8.64189981 | 0.00097054 | 0.02353924 |
| ENSGALG00010028884 | VAV2 | 373938 | 0.27752091 | 5.86816152 | 0.00097434 | 0.02359548 |
| ENSGALG00010008062 | VPS9D1 | 415839 | -0.3057749 | 4.85847212 | 0.00098829 | 0.02386142 |
| ENSGALG00010002956 | SCRG1 | 422563 | 0.94272041 | 1.29138659 | 0.00098833 | 0.02386142 |
| ENSGALG00010027789 | UAP1L1 | 417295 | -0.3225496 | 4.86194259 | 0.00099446 | 0.02397292 |
| ENSGALG00010008705 | NA | NA | 0.37781518 | 4.45607694 | 0.00100024 | 0.02404397 |

| | | | | | | |
|---|---|---|---|---|---|---|
| ENSGALG00010027172 | GDF15 | 101748931 | -0.7220865 | 4.14896983 | 0.00100044 | 0.02404397 |
| ENSGALG00010027073 | DNAH12 | 416004 | -0.3994226 | 6.01869612 | 0.00100814 | 0.02417576 |
| ENSGALG00010003792 | NA | NA | 0.89279391 | 1.40330619 | 0.00101014 | 0.02417576 |
| ENSGALG00010011316 | PARPBP | 418092 | 0.40340099 | 3.89994709 | 0.0010105 | 0.02417576 |
| ENSGALG00010009668 | TFAP2A | 395982 | 0.74344094 | 1.60930135 | 0.00101595 | 0.02426947 |
| ENSGALG00010010298 | KCNIP4 | 395239 | -0.4761119 | 3.05007353 | 0.00101998 | 0.02432914 |
| ENSGALG00010008227 | DUSP1 | 374192 | -0.6426992 | 2.51891325 | 0.00102161 | 0.02433144 |
| ENSGALG00010018885 | EFEMP1 | 428543 | 0.96611663 | 1.38379212 | 0.00102366 | 0.0243437 |
| ENSGALG00010023061 | NA | NA | -0.4917029 | 3.36579817 | 0.00102913 | 0.02443711 |
| ENSGALG00010008784 | NA | NA | -0.9794581 | 0.6028343 | 0.00103372 | 0.02450957 |
| ENSGALG00010000472 | NA | NA | -1.029468 | 1.1593706 | 0.00103867 | 0.02459002 |
| ENSGALG00010024058 | SLC44A5 | 429117 | 0.39048196 | 4.14510756 | 0.00104133 | 0.02459671 |
| ENSGALG00010017860 | CYSLTR1 | 428691 | 0.6208503 | 3.52436636 | 0.00104205 | 0.02459671 |
| ENSGALG00010019955 | MIR363 | 114482517 | -1.4121099 | -0.3521788 | 0.00105168 | 0.02474998 |
| ENSGALG00010024662 | MOB2 | 423097 | -0.2625722 | 6.97655237 | 0.00105212 | 0.02474998 |
| ENSGALG00010020562 | LRRN1 | 771599 | -0.3567051 | 7.86149831 | 0.00105401 | 0.02474998 |
| ENSGALG00010002585 | NA | NA | 0.55731453 | 2.54090484 | 0.00105478 | 0.02474998 |
| ENSGALG00010024504 | CASC3 | 426135 | -0.3006329 | 7.51191981 | 0.00105787 | 0.02478561 |
| ENSGALG00010029376 | MXRA7 | 769894 | -0.4353066 | 4.64682709 | 0.00106351 | 0.02488102 |
| ENSGALG00010020692 | TUB | 423061 | -0.2873155 | 5.92701033 | 0.00107964 | 0.02522125 |
| ENSGALG00010024781 | CEP19 | 424858 | -0.4797707 | 3.28455695 | 0.00108702 | 0.02533757 |
| ENSGALG00010022412 | CHAC1 | 423209 | -0.5830976 | 5.50313893 | 0.00108781 | 0.02533757 |
| ENSGALG00010013301 | PARVB | 418235 | 0.30922335 | 6.39278816 | 0.00110365 | 0.02566864 |
| ENSGALG00010010214 | FAF2 | 416225 | -0.2474776 | 5.99643208 | 0.00110734 | 0.02571679 |
| ENSGALG00010007228 | NA | NA | 1.52117441 | -0.342513 | 0.00113233 | 0.02625868 |
| ENSGALG00010019264 | HK2 | 374044 | 0.3053321 | 5.41204717 | 0.00115401 | 0.02669543 |
| ENSGALG00010024873 | HYOU1 | 428251 | -0.4686267 | 9.35181298 | 0.00115452 | 0.02669543 |
| ENSGALG00010009993 | NA | NA | -0.4646861 | 3.3896604 | 0.00116401 | 0.02687568 |
| ENSGALG00010008974 | SMCO4 | 772206 | -0.4732383 | 3.55673622 | 0.00116935 | 0.0269369 |
| ENSGALG00010028529 | GRIN1 | 404296 | 0.98742312 | 0.65206123 | 0.00117372 | 0.0269369 |
| ENSGALG00010023612 | GPT2 | 415746 | -0.728413 | 6.34875678 | 0.00117483 | 0.0269369 |
| ENSGALG00010015115 | CLDN2 | 422292 | -0.4033154 | 4.53061118 | 0.00117522 | 0.0269369 |
| ENSGALG00010017702 | ECEL1 | 424937 | -0.6097589 | 3.5246017 | 0.00117614 | 0.0269369 |
| ENSGALG00010001127 | SIAH2 | 429164 | -0.5849741 | 3.88756525 | 0.00117686 | 0.0269369 |
| ENSGALG00010007100 | NA | NA | -1.303363 | -0.1951699 | 0.00118975 | 0.02719289 |
| ENSGALG00010006009 | MALRD1 | 420512 | 1.19406213 | 0.36286634 | 0.00119587 | 0.02727195 |
| ENSGALG00010000044 | MAP1A | 415588 | 0.49029138 | 6.53420504 | 0.00119837 | 0.02727195 |
| ENSGALG00010011202 | NA | NA | 1.0713043 | 0.65324041 | 0.00119837 | 0.02727195 |
| ENSGALG00010004416 | STXBP6 | 423299 | -0.3880387 | 5.18295402 | 0.00121701 | 0.02763875 |
| ENSGALG00010010461 | SLC7A5 | 415832 | -0.6717826 | 6.81613798 | 0.00121797 | 0.02763875 |
| ENSGALG00010004765 | DMBT1 | 426819 | 0.28529199 | 5.54483328 | 0.00122756 | 0.02781641 |
| ENSGALG00010030104 | P4HB | 374091 | -0.4190521 | 9.58074267 | 0.00123218 | 0.02788145 |
| ENSGALG00010000636 | ZNF646 | 121108148 | 0.28884602 | 5.84151795 | 0.00124245 | 0.02807371 |
| ENSGALG00010019024 | CNPY3 | 100858990 | -0.3297087 | 5.23013632 | 0.00124989 | 0.02818328 |
| ENSGALG00010018858 | EPB41L5 | 424235 | 0.23997396 | 7.6863837 | 0.00125085 | 0.02818328 |
| ENSGALG00010022940 | NA | NA | 1.3913168 | 0.06822034 | 0.00125838 | 0.02831267 |
| ENSGALG00010011502 | NA | NA | -1.5199755 | -0.2587374 | 0.00126519 | 0.02841089 |
| ENSGALG00010002211 | ACER3 | 419087 | 0.25444677 | 6.86541434 | 0.00126633 | 0.02841089 |
| ENSGALG00010004044 | RNF138 | 421096 | 0.26874531 | 6.80731887 | 0.00127434 | 0.0285245 |
| ENSGALG00010005792 | ADGRG6 | 421673 | -0.624355 | 2.83906021 | 0.00127499 | 0.0285245 |
| ENSGALG00010005852 | NA | NA | -1.3372507 | -0.3331821 | 0.0013023 | 0.02909438 |
| ENSGALG00010016578 | GRIK3 | 419619 | 0.55612565 | 3.71494237 | 0.00130552 | 0.02912547 |
| ENSGALG00010029618 | TEX2 | 417415 | 0.28773338 | 6.61662551 | 0.00131195 | 0.02919978 |
| ENSGALG00010001698 | RPL15 | 428442 | -0.2655538 | 9.34358431 | 0.00131382 | 0.02919978 |
| ENSGALG00010001171 | POLR1E | 426805 | -0.2403701 | 6.27383113 | 0.00131438 | 0.02919978 |
| ENSGALG00010002474 | HPCAL1 | 396161 | 0.85175607 | 1.72326324 | 0.00131879 | 0.02922342 |
| ENSGALG00010016644 | NMRAL1 | 416672 | -0.4663618 | 5.28040506 | 0.00131913 | 0.02922342 |
| ENSGALG00010017037 | NA | NA | 1.63252369 | -0.3014722 | 0.00132349 | 0.02927928 |
| ENSGALG00010015737 | NA | NA | 1.11770159 | 0.25573419 | 0.00132837 | 0.0293462 |
| ENSGALG00010012104 | SMC1B | 418243 | 0.52600339 | 3.3909673 | 0.00133606 | 0.02945962 |

| | | | | | | |
|---|---|---|---|---|---|---|
| ENSGALG00010017163 | CDV3 | 420693 | -0.2420957 | 8.32604321 | 0.00133722 | 0.02945962 |
| ENSGALG00010005477 | TMEM161B | 426919 | -0.2687462 | 6.34872954 | 0.00134092 | 0.02948623 |
| ENSGALG00010008098 | TMEM126A | 426392 | -0.2762334 | 5.17368653 | 0.00134214 | 0.02948623 |
| ENSGALG00010007998 | RPL13 | 395849 | -0.290204 | 8.95208158 | 0.00134697 | 0.02955134 |
| ENSGALG00010009170 | RPL26L1 | 396400 | -0.3283583 | 8.54307613 | 0.00135883 | 0.02974858 |
| ENSGALG00010007539 | RPS3A | 422477 | -0.2632788 | 9.90958059 | 0.00135971 | 0.02974858 |
| ENSGALG00010011118 | GMPR2 | 420842 | 0.39942982 | 4.35011917 | 0.00136551 | 0.02983435 |
| ENSGALG00010026205 | A2ML4 | 100858010 | 0.36183044 | 4.75433487 | 0.00137133 | 0.02992029 |
| ENSGALG00010011984 | FN1 | 396133 | 0.26603902 | 7.08463344 | 0.00137564 | 0.02997302 |
| ENSGALG00010025445 | B3GNT4 | 100858043 | -0.2677621 | 8.06307034 | 0.00138163 | 0.03006232 |
| ENSGALG00010024509 | SEPTIN5 | 416778 | -0.3700058 | 8.36972187 | 0.00138507 | 0.0300959 |
| ENSGALG00010012188 | ADAMTS6 | 427160 | 0.2976084 | 4.83525242 | 0.00139064 | 0.03017564 |
| ENSGALG00010014470 | RPS6KA1 | 386579 | 0.28190641 | 5.68879697 | 0.00139299 | 0.03018519 |
| ENSGALG00010027527 | MKRN3 | 419892 | -0.5591681 | 2.28470144 | 0.00139954 | 0.03024888 |
| ENSGALG00010008340 | PDIA4 | 420785 | -0.4454964 | 8.87006397 | 0.00139974 | 0.03024888 |
| ENSGALG00010012631 | GLRX2 | 424357 | -0.2469242 | 6.78915635 | 0.00140899 | 0.03040724 |
| ENSGALG00010009253 | SLC45A4 | 420308 | 0.28095332 | 5.18976716 | 0.00142207 | 0.03064781 |
| ENSGALG00010017944 | NA | NA | 1.00520764 | 0.72836561 | 0.00142492 | 0.03066759 |
| ENSGALG00010003560 | ADGRL3 | 422616 | 0.38440054 | 4.26102344 | 0.00143794 | 0.03090601 |
| ENSGALG00010005250 | SLC35A5 | 418368 | 0.3171671 | 4.84120088 | 0.00144344 | 0.03098214 |
| ENSGALG00010014038 | NA | NA | 0.32512461 | 4.77683747 | 0.0014532 | 0.03114959 |
| ENSGALG00010022972 | A2M | 418251 | -1.0816797 | 0.38411915 | 0.00146781 | 0.03142013 |
| ENSGALG00010012998 | MAFF | 395519 | -0.5567972 | 3.46799355 | 0.00147346 | 0.03149876 |
| ENSGALG00010004243 | KIAA1109 | 422670 | 0.23156145 | 7.60874505 | 0.00147689 | 0.03150764 |
| ENSGALG00010027077 | ARF4 | 769725 | -0.2754381 | 8.14706364 | 0.0014779 | 0.03150764 |
| ENSGALG00010010080 | CDC42EP3 | 428577 | 0.58132632 | 3.03099987 | 0.00147984 | 0.03150764 |
| ENSGALG00010017212 | TINAGL1 | 430648 | -0.6618938 | 3.10969557 | 0.00148714 | 0.03158005 |
| ENSGALG00010006359 | SGK1 | 395133 | 0.36056964 | 5.14317702 | 0.00148722 | 0.03158005 |
| ENSGALG00010021741 | RPL19 | 420003 | -0.2527922 | 9.05583259 | 0.0014935 | 0.03167088 |
| ENSGALG00010020383 | BAAT | 769879 | -0.2414418 | 6.28634897 | 0.00150029 | 0.03177245 |
| ENSGALG00010027110 | IKBKE | 430480 | 0.41418404 | 4.03348372 | 0.00150475 | 0.03182436 |
| ENSGALG00010012955 | RPL39L | 374132 | -0.3335528 | 7.85186354 | 0.00151818 | 0.03206566 |
| ENSGALG00010021419 | MANSC1 | 426428 | 0.44199742 | 3.96379564 | 0.00152567 | 0.03218105 |
| ENSGALG00010027569 | ATP5PB | 419866 | -0.2327939 | 7.26275809 | 0.00157926 | 0.03326726 |
| ENSGALG00010028649 | LOC101751615 | 101751615 | 0.73489229 | 3.545271 | 0.00158226 | 0.03328228 |
| ENSGALG00010027186 | PLXNA2 | 419857 | 0.53332015 | 3.30703623 | 0.00158417 | 0.03328228 |
| ENSGALG00010018054 | SMAD6 | 374096 | -0.42698 | 3.83504155 | 0.00158874 | 0.03333406 |
| ENSGALG00010010399 | FGF18 | 395453 | 0.69238428 | 1.61707878 | 0.00159865 | 0.03349772 |
| ENSGALG00010012876 | KDELR3 | 418026 | -0.5982633 | 4.27426129 | 0.00160562 | 0.03355181 |
| ENSGALG00010028514 | FNBP1 | 417187 | -0.2201579 | 7.71648968 | 0.00160718 | 0.03355181 |
| ENSGALG00010024714 | SRPRB | 101750392 | -0.4560936 | 10.0598634 | 0.00160758 | 0.03355181 |
| ENSGALG00010016875 | SESN2 | 100858796 | -0.7187581 | 4.95599728 | 0.00161167 | 0.03359118 |
| ENSGALG00010026320 | GGT5 | 416943 | 0.59236293 | 2.32865955 | 0.0016137 | 0.03359118 |
| ENSGALG00010012830 | ARRDC3 | 427107 | -0.2437811 | 6.13502399 | 0.00161695 | 0.03361468 |
| ENSGALG00010008652 | NA | NA | -0.5246618 | 3.50210254 | 0.00162591 | 0.03375673 |
| ENSGALG00010027996 | GPX4 | 374056 | -0.3937319 | 7.55429445 | 0.00162954 | 0.03378783 |
| ENSGALG00010020747 | DBI | 395263 | -0.2410062 | 6.17029598 | 0.0016324 | 0.03380297 |
| ENSGALG00010015736 | NA | NA | 0.94113726 | 1.03109365 | 0.00163461 | 0.03380474 |
| ENSGALG00010013380 | PHGDH | 424381 | -0.4712598 | 7.43756541 | 0.00164223 | 0.03391794 |
| ENSGALG00010016593 | OTX1 | 100858585 | -0.504611 | 3.44180934 | 0.00164812 | 0.03399548 |
| ENSGALG00010004306 | PNISR | 421796 | -0.2339443 | 7.00454132 | 0.00166616 | 0.03432297 |
| ENSGALG00010027427 | UBAC1 | 427754 | -0.2419701 | 6.90409926 | 0.00168034 | 0.03457008 |
| ENSGALG00010008161 | TMEFF1 | 420985 | 0.50326371 | 3.93189026 | 0.00169476 | 0.03479318 |
| ENSGALG00010006522 | NA | NA | -0.6606623 | 3.81205227 | 0.00169635 | 0.03479318 |
| ENSGALG00010029331 | TK1 | 395719 | -0.2320298 | 6.20571229 | 0.00169776 | 0.03479318 |
| ENSGALG00010016338 | PTPRU | 395153 | 0.37080783 | 5.07917246 | 0.00170724 | 0.03488172 |
| ENSGALG00010013171 | SLC9A8 | 419356 | -0.3540984 | 5.79062115 | 0.00170822 | 0.03488172 |
| ENSGALG00010015533 | PSAT1 | 427263 | -0.5308394 | 6.82493836 | 0.00170868 | 0.03488172 |
| ENSGALG00010011757 | NA | NA | 0.45069082 | 3.5657336 | 0.00171443 | 0.03495412 |
| ENSGALG00010020365 | UNC5B | 423716 | 0.37233953 | 3.97515962 | 0.00172255 | 0.03507465 |

| | | | | | | |
|---|---|---|---|---|---|---|
| ENSGALG00010008180 | POU2F1 | 396463 | 0.31000076 | 5.11829035 | 0.00172599 | 0.03509952 |
| ENSGALG00010001525 | EEF1A1 | 373963 | -0.2364887 | 8.22161747 | 0.00173742 | 0.0352484 |
| ENSGALG00010015401 | NA | NA | 1.04367532 | 0.37744838 | 0.00173775 | 0.0352484 |
| ENSGALG00010025867 | NA | NA | 0.99323223 | 0.45346292 | 0.00174188 | 0.0352568 |
| ENSGALG00010027895 | LOC107051565 | 107051565 | 0.31836768 | 5.4892541 | 0.00174261 | 0.0352568 |
| ENSGALG00010017381 | CA6 | 428187 | 0.65810278 | 2.16548491 | 0.00175035 | 0.03536826 |
| ENSGALG00010001627 | ARFIP2 | 419075 | -0.2433596 | 5.75501554 | 0.00175511 | 0.03541917 |
| ENSGALG00010029492 | SDK2 | 395215 | -0.3574316 | 4.97720413 | 0.00175941 | 0.03546097 |
| ENSGALG00010021156 | DDOST | 425542 | -0.3034776 | 8.0528504 | 0.00176351 | 0.03549849 |
| ENSGALG00010029418 | ABCA5 | 417444 | 0.24479079 | 6.29748458 | 0.00176688 | 0.0355212 |
| ENSGALG00010027307 | JUND | 107055370 | -0.3187091 | 6.1759164 | 0.00177164 | 0.03557182 |
| ENSGALG00010015729 | RHCE | 395121 | -0.43933 | 3.41407558 | 0.00177419 | 0.03557791 |
| ENSGALG00010024481 | LOC112532944 | 112532944 | -1.210505 | 0.06027538 | 0.00178849 | 0.03581858 |
| ENSGALG00010008999 | TRAPPC9 | 420312 | 0.34149781 | 4.40029005 | 0.0017907 | 0.03581858 |
| ENSGALG00010003571 | RPL7 | 420182 | -0.2232568 | 9.38640919 | 0.00180006 | 0.03596047 |
| ENSGALG00010010713 | NSA2 | 427206 | -0.2403111 | 7.24265108 | 0.00181461 | 0.03620549 |
| ENSGALG00010000619 | NA | NA | 0.38741358 | 4.06390052 | 0.00182426 | 0.0363417 |
| ENSGALG00010010512 | PLXNC1 | 417901 | 0.44821246 | 5.4360501 | 0.00182602 | 0.0363417 |
| ENSGALG00010028562 | ACSBG2 | 420090 | -0.5423849 | 5.08886557 | 0.00184078 | 0.03658962 |
| ENSGALG00010014324 | COL4A2 | 418752 | 0.3609471 | 5.55888851 | 0.0018713 | 0.03708542 |
| ENSGALG00010004975 | SNW1 | 423381 | -0.2465968 | 7.14765923 | 0.00187185 | 0.03708542 |
| ENSGALG00010000279 | NA | NA | -0.3783089 | 3.9173891 | 0.00187274 | 0.03708542 |
| ENSGALG00010020708 | FAM214A | 415416 | 0.23710485 | 7.19376075 | 0.00188367 | 0.03725534 |
| ENSGALG00010013004 | UAP1 | 424368 | -0.2746642 | 5.89439909 | 0.00189291 | 0.03739147 |
| ENSGALG00010012982 | DDR2 | 396404 | -0.6943364 | 3.58923173 | 0.00189697 | 0.03742513 |
| ENSGALG00010029943 | HS3ST3A1 | 769490 | 0.68292883 | 1.71951323 | 0.00194046 | 0.03823558 |
| ENSGALG00010001732 | NA | NA | 0.29659344 | 5.40158204 | 0.00195845 | 0.03854207 |
| ENSGALG00010001506 | DPYSL4 | 374022 | -0.3323932 | 5.67963319 | 0.00197323 | 0.03878475 |
| ENSGALG00010015695 | ASAH1 | 422727 | -0.3109322 | 5.68477629 | 0.00197929 | 0.03885584 |
| ENSGALG00010024425 | MRPL40 | 100858795 | -0.3259696 | 4.68066529 | 0.00198728 | 0.03892458 |
| ENSGALG00010004035 | NA | NA | 0.71665434 | 1.52760623 | 0.0019877 | 0.03892458 |
| ENSGALG00010026148 | RPS10 | 419904 | -0.2674831 | 8.93986735 | 0.00200496 | 0.0391558 |
| ENSGALG00010012135 | CAMSAP2 | 424345 | -0.2862243 | 6.35438411 | 0.00200592 | 0.0391558 |
| ENSGALG00010017605 | GFPT1 | 419514 | -0.23116 | 8.20494784 | 0.00200691 | 0.0391558 |
| ENSGALG00010006061 | RPL24 | 418401 | -0.2287164 | 7.77745926 | 0.00201612 | 0.03928707 |
| ENSGALG00010012670 | NA | NA | 0.28864399 | 5.46954915 | 0.00202596 | 0.03939773 |
| ENSGALG00010011607 | TSEN15 | 424448 | -0.5619894 | 2.26243322 | 0.00202945 | 0.03939773 |
| ENSGALG00010000616 | PSENEN | 121108876 | 0.7396773 | 2.60167557 | 0.00203069 | 0.03939773 |
| ENSGALG00010000579 | NA | NA | 0.47751647 | 2.93433534 | 0.00203173 | 0.03939773 |
| ENSGALG00010007673 | FHDC1 | 422484 | 0.24341444 | 5.60104093 | 0.00204575 | 0.03962102 |
| ENSGALG00010025745 | COX14 | 426880 | -0.5033 | 3.57428972 | 0.00204961 | 0.03964739 |
| ENSGALG00010021625 | HCN4 | 427478 | 0.50314517 | 3.37758286 | 0.0020579 | 0.03975927 |
| ENSGALG00010024554 | ST6GALNAC3 | 404746 | 0.43072907 | 3.56275969 | 0.00206168 | 0.03978387 |
| ENSGALG00010003680 | TRAPPC5 | 100858483 | 0.26239182 | 7.05949847 | 0.00206746 | 0.03984689 |
| ENSGALG00010015573 | MAN1C1 | 419588 | 0.55876921 | 2.83516152 | 0.00207026 | 0.03985253 |
| ENSGALG00010018973 | NEFM | 396206 | -0.397245 | 3.71926185 | 0.00208446 | 0.04007723 |
| ENSGALG00010001414 | CCDC126 | 420619 | -0.2921952 | 5.35143552 | 0.00210623 | 0.04042287 |
| ENSGALG00010003198 | PRKG2 | 428753 | 0.30870356 | 5.9867823 | 0.00210754 | 0.04042287 |
| ENSGALG00010011636 | GRIK1 | 418489 | 0.60623343 | 2.50944658 | 0.00211703 | 0.04055586 |
| ENSGALG00010014878 | TMCO1 | 427100 | -0.2684712 | 5.29366642 | 0.00212292 | 0.04061965 |
| ENSGALG00010023141 | LOC121108102 | 121108102 | 2.43759751 | 0.66872558 | 0.00212837 | 0.04064897 |
| ENSGALG00010025640 | SLC38A11 | 424181 | 0.30217745 | 6.10740822 | 0.00212957 | 0.04064897 |
| ENSGALG00010025621 | PPP1CC | 416872 | -0.2861335 | 7.88407941 | 0.00213763 | 0.04075375 |
| ENSGALG00010013604 | FYN | 396294 | 0.28262134 | 5.29317352 | 0.00215443 | 0.04098903 |
| ENSGALG00010027385 | CNTN2 | 419825 | -0.8088685 | 1.84757175 | 0.00215514 | 0.04098903 |
| ENSGALG00010028114 | PFKFB4 | 100859653 | 0.26390553 | 6.06457387 | 0.0021734 | 0.0412249 |
| ENSGALG00010024870 | TPD52L2 | 419257 | -0.2192404 | 7.01628583 | 0.00217475 | 0.0412249 |
| ENSGALG00010009194 | NA | NA | 0.86682752 | 0.94735758 | 0.00217647 | 0.0412249 |
| ENSGALG00010029451 | LOC107057175 | 107057175 | -1.2438862 | 2.54274112 | 0.00217794 | 0.0412249 |
| ENSGALG00010025685 | NCKAP1L | 425499 | 0.28587332 | 5.16050222 | 0.00218107 | 0.04123492 |

| | | | | | | |
|---|---|---|---|---|---|---|
| ENSGALG00010016647 | EGR1 | 373931 | 0.52274704 | 2.72960678 | 0.0021898 | 0.04133743 |
| ENSGALG00010011157 | NA | NA | 1.17648635 | 0.22375352 | 0.0021917 | 0.04133743 |
| ENSGALG00010005360 | CAMK1D | 425522 | 0.6274562 | 3.3621243 | 0.00222636 | 0.0418778 |
| ENSGALG00010004245 | ARFGEF3 | 421681 | -0.3847974 | 5.66075128 | 0.00222878 | 0.0418778 |
| ENSGALG00010022416 | CCDC186 | 423905 | 0.23210401 | 7.99197686 | 0.00223007 | 0.0418778 |
| ENSGALG00010020371 | NA | NA | 1.5251214 | -0.3570943 | 0.00223348 | 0.0418778 |
| ENSGALG00010016591 | HOMER2 | 415458 | -0.2211562 | 6.62815346 | 0.00223355 | 0.0418778 |
| ENSGALG00010004222 | SPAG6 | 420505 | 0.49230414 | 3.52885801 | 0.00223638 | 0.04188131 |
| ENSGALG00010019310 | CRELD1 | 100858613 | -0.2992459 | 5.11274234 | 0.00224296 | 0.04195501 |
| ENSGALG00010023259 | TSPAN15 | 423699 | 0.45487269 | 3.44206225 | 0.00226147 | 0.0422514 |
| ENSGALG00010028877 | FAM163B | 100858326 | 0.66448066 | 1.90039824 | 0.00226989 | 0.04235873 |
| ENSGALG00010020996 | NA | NA | 0.70940547 | 1.71523748 | 0.00228015 | 0.04250021 |
| ENSGALG00010014906 | IFT27 | 418053 | 0.31620022 | 5.1143103 | 0.0022862 | 0.0425631 |
| ENSGALG00010021693 | PPA1 | 769424 | -0.2489751 | 7.1022344 | 0.00229728 | 0.04271926 |
| ENSGALG00010009489 | LOC101749352 | 101749352 | 0.23320068 | 6.92846771 | 0.00230876 | 0.04287463 |
| ENSGALG00010017852 | LETM2 | 426779 | -0.3970129 | 3.91447886 | 0.00231105 | 0.04287463 |
| ENSGALG00010003767 | LOC121106918 | 121106918 | 1.79840794 | -0.4821344 | 0.00231839 | 0.04290773 |
| ENSGALG00010011192 | RPL17 | 426845 | -0.290275 | 9.07245098 | 0.00231865 | 0.04290773 |
| ENSGALG00010008350 | LRRC6 | 420324 | 0.87336631 | 1.1069866 | 0.00232095 | 0.04290773 |
| ENSGALG00010022169 | TOX2 | 419156 | -0.7009731 | 3.53317572 | 0.00233957 | 0.04320161 |
| ENSGALG00010020636 | NBL1 | 373952 | -0.3558017 | 4.65312762 | 0.00234898 | 0.04332504 |
| ENSGALG00010005911 | TBXT | 395782 | 0.53524952 | 4.15585987 | 0.00236219 | 0.04347588 |
| ENSGALG00010003978 | ANK2 | 422689 | 0.23679603 | 7.62314428 | 0.00236264 | 0.04347588 |
| ENSGALG00010005472 | PFKP | 428411 | -0.218374 | 8.17216918 | 0.00237712 | 0.04361973 |
| ENSGALG00010017509 | RASD1 | 416507 | -0.5491913 | 2.90744764 | 0.00237849 | 0.04361973 |
| ENSGALG00010016394 | PDIK1L | 419604 | -0.294552 | 4.83443067 | 0.00237871 | 0.04361973 |
| ENSGALG00010013214 | RAI14 | 427431 | 0.32045745 | 5.10339792 | 0.00238719 | 0.04372469 |
| ENSGALG00010024078 | KIRREL3 | 419719 | 0.52957082 | 3.80581717 | 0.00239634 | 0.04381499 |
| ENSGALG00010015889 | SH3BP5 | 420682 | 0.30718588 | 4.90470319 | 0.00239764 | 0.04381499 |
| ENSGALG00010012322 | MYO3AL | 427419 | -0.2569933 | 6.36781291 | 0.00240726 | 0.04394015 |
| ENSGALG00010010577 | PBX1 | 395505 | 0.21048411 | 8.38936946 | 0.00241214 | 0.04397859 |
| ENSGALG00010011449 | DUSP22 | 420887 | 0.22080484 | 7.85245696 | 0.00241806 | 0.04403585 |
| ENSGALG00010024742 | PAK2 | 424860 | -0.3111936 | 6.8089261 | 0.00242678 | 0.04414393 |
| ENSGALG00010012489 | GBX2 | 395950 | 0.60755322 | 2.31327754 | 0.00243999 | 0.04433354 |
| ENSGALG00010024582 | TLN2 | 415374 | 0.217867 | 7.5946852 | 0.00244591 | 0.04439016 |
| ENSGALG00010029584 | TRIM25 | 417401 | -0.6219745 | 2.40704095 | 0.00245125 | 0.04443633 |
| ENSGALG00010028048 | S100A10 | 396506 | -0.604567 | 2.96081158 | 0.00245419 | 0.04443885 |
| ENSGALG00010021721 | TP53I11 | 426399 | 0.22354269 | 7.66198247 | 0.00245703 | 0.04443946 |
| ENSGALG00010023668 | RCN1 | 428602 | -0.3658429 | 5.63302852 | 0.00247431 | 0.04470099 |
| ENSGALG00010027134 | DYRK3 | 419846 | 0.24598701 | 6.78769957 | 0.00248769 | 0.04489169 |
| ENSGALG00010024964 | ITM2C | 424831 | 0.42116826 | 3.76357384 | 0.00249537 | 0.04497896 |
| ENSGALG00010029546 | WSB1 | 395685 | -0.2511336 | 6.57547762 | 0.00249831 | 0.04498095 |
| ENSGALG00010006283 | DNAJC25 | 431639 | -0.2878838 | 5.26256998 | 0.00250293 | 0.045013 |
| ENSGALG00010001272 | NA | NA | 0.56593264 | 2.6044027 | 0.00250583 | 0.04501414 |
| ENSGALG00010007859 | CENPN | 693246 | -0.285816 | 4.96886035 | 0.0025134 | 0.04509898 |
| ENSGALG00010025318 | NUAK2 | 419828 | 0.52212556 | 2.90733095 | 0.00252429 | 0.04524333 |
| ENSGALG00010028120 | TBXA2R | 100857635 | -0.3100215 | 4.81195462 | 0.00253524 | 0.0453882 |
| ENSGALG00010019268 | NSL1 | 421367 | -0.2825575 | 4.9459821 | 0.00254051 | 0.04543135 |
| ENSGALG00010021683 | HFM1 | 424509 | 0.78087518 | 1.32514357 | 0.00255011 | 0.04555162 |
| ENSGALG00010024351 | SP1 | 395303 | 0.23356106 | 6.0035925 | 0.00255427 | 0.04557465 |
| ENSGALG00010019679 | RHOBTB2 | 419536 | 0.26392853 | 5.82397198 | 0.00257604 | 0.04591137 |
| ENSGALG00010001541 | NA | NA | -0.5683529 | 2.50932389 | 0.00260465 | 0.04632338 |
| ENSGALG00010009013 | LIPG | 426846 | -0.7170261 | 3.05412059 | 0.002605 | 0.04632338 |
| ENSGALG00010022808 | PSMA7 | 395318 | -0.2191945 | 7.48448425 | 0.00260992 | 0.04632887 |
| ENSGALG00010024340 | NETO2 | 415743 | 0.33200731 | 4.58583031 | 0.00261115 | 0.04632887 |
| ENSGALG00010028017 | LOC121107546 | 121107546 | -0.7710983 | 1.32405007 | 0.0026252 | 0.04652608 |
| ENSGALG00010013822 | NA | NA | -0.7327704 | 1.87627743 | 0.00263367 | 0.04662417 |
| ENSGALG00010016040 | RPSAP58 | 395181 | -0.2380554 | 10.5620309 | 0.00264719 | 0.04681124 |
| ENSGALG00010018574 | NA | NA | -0.8088622 | 1.80575211 | 0.00267304 | 0.04721569 |
| ENSGALG00010011681 | MAP1B | 396174 | -0.4540406 | 6.2940503 | 0.00269138 | 0.04748675 |

| | | | | | | |
|---|---|---|---|---|---|---|
| ENSGALG00010021199 | TMEM258 | 422964 | -0.2755587 | 5.91837536 | 0.00270069 | 0.04759811 |
| ENSGALG00010006572 | POU4F2 | 428735 | 1.14893817 | 0.20995231 | 0.00270449 | 0.04761224 |
| ENSGALG00010029903 | MIR22 | 100315903 | -1.3392554 | -0.5297824 | 0.00272557 | 0.04789873 |
| ENSGALG00010027059 | ASB14 | 416003 | -0.348829 | 4.34925601 | 0.0027268 | 0.04789873 |
| ENSGALG00010024649 | NCOR2 | 416815 | 0.23822824 | 7.0774241 | 0.00273592 | 0.04800566 |
| ENSGALG00010012242 | CCDC77 | 418149 | 0.33596409 | 4.57193853 | 0.00275182 | 0.04819835 |
| ENSGALG00010005545 | RPL34 | 422526 | -0.2775483 | 8.60148687 | 0.00275773 | 0.04819835 |
| ENSGALG00010020457 | IL5RA | 771801 | -0.5050641 | 3.57124158 | 0.00275811 | 0.04819835 |
| ENSGALG00010002912 | RPS3 | 419069 | -0.2734012 | 9.14759423 | 0.00275905 | 0.04819835 |
| ENSGALG00010029761 | MRPL38 | 770172 | -0.2866513 | 5.23915406 | 0.00276661 | 0.04827714 |
| ENSGALG00010023441 | C15orf52 | 423013 | 0.78495328 | 1.55655395 | 0.00277157 | 0.04830055 |
| ENSGALG00010021083 | SCD | 395706 | -0.4117771 | 8.01971191 | 0.00277404 | 0.04830055 |
| ENSGALG00010011790 | AMY2A | 414140 | 0.29033819 | 5.28589654 | 0.00277888 | 0.04833174 |
| ENSGALG00010020337 | NA | NA | -0.667826 | 2.26846279 | 0.00280008 | 0.0486119 |
| ENSGALG00010024102 | MSH4 | 424725 | -0.229416 | 7.22318243 | 0.00280111 | 0.0486119 |
| ENSGALG00010001532 | NA | NA | -0.7300808 | 2.34610126 | 0.00282137 | 0.04890985 |
| ENSGALG00010024386 | TPPP3 | 415669 | -0.889963 | 1.5275703 | 0.00283115 | 0.04902594 |
| ENSGALG00010015152 | BOLL | 424061 | 0.33134738 | 5.51678919 | 0.00285764 | 0.04943058 |
| ENSGALG00010002399 | CSMD1 | 421899 | -0.3844487 | 4.81246822 | 0.00287047 | 0.04957012 |
| ENSGALG00010007181 | GJB2 | 428083 | 0.51548852 | 2.51036055 | 0.00287409 | 0.04957012 |
| ENSGALG00010020410 | NA | NA | 1.25943083 | 0.22592633 | 0.00287508 | 0.04957012 |
| ENSGALG00010004648 | LOC421806 | 421806 | 0.27133073 | 9.41332821 | 0.00288301 | 0.0496528 |
| ENSGALG00010015443 | PAK3 | 422342 | 0.24476299 | 6.60013004 | 0.00288707 | 0.04966891 |
| ENSGALG00010016407 | RPL9 | 425468 | -0.2379198 | 8.92696913 | 0.00289106 | 0.04968365 |
| ENSGALG00010013355 | FAM171B | 423991 | -0.2278065 | 7.11503425 | 0.00289949 | 0.04975326 |
| ENSGALG00010015376 | NA | NA | 0.4981752 | 3.79914927 | 0.00290138 | 0.04975326 |
| ENSGALG00010001298 | DOCK4 | 417779 | 0.28347332 | 5.4480542 | 0.00291624 | 0.04995398 |
| ENSGALG00010012801 | DMC1 | 427903 | 0.45486215 | 3.19612569 | 0.00292201 | 0.04999881 |
| ENSGALG00010008209 | PPARGC1B | 416148 | 0.48143627 | 3.01285658 | 0.00293225 | 0.05009534 |
| ENSGALG00010024377 | CIB2 | 415363 | -0.4912471 | 2.5758207 | 0.00293397 | 0.05009534 |
| ENSGALG00010021348 | SLC4A3 | 424194 | 0.37804279 | 4.24321072 | 0.00293918 | 0.05013036 |
| ENSGALG00010010743 | KLHL20 | 425091 | 0.27348851 | 5.07284447 | 0.00295022 | 0.0502647 |
| ENSGALG00010016502 | YIPF5 | 416344 | -0.2673204 | 6.33144726 | 0.00295555 | 0.0503014 |
| ENSGALG00010025130 | ATP2A2 | 396446 | -0.2613797 | 8.84354965 | 0.00296472 | 0.05040337 |
| ENSGALG00010024512 | CCNDBP1 | 419217 | -0.2648625 | 5.50862463 | 0.00300148 | 0.05097379 |
| ENSGALG00010018557 | TBCC | 428567 | -0.3104819 | 7.57674 | 0.00300915 | 0.05104941 |
| ENSGALG00010000383 | NA | NA | -0.6506597 | 4.92998971 | 0.00304688 | 0.05163411 |
| ENSGALG00010029475 | LOC107057183 | 107057183 | -0.799436 | 2.51918504 | 0.00306895 | 0.05195271 |
| ENSGALG00010020866 | LCOR | 423847 | 0.25452344 | 5.24764891 | 0.00308781 | 0.05221627 |
| ENSGALG00010018310 | APOOL | 422271 | -0.2256555 | 6.31160195 | 0.00309451 | 0.05226808 |
| ENSGALG00010005264 | ITIH2 | 419110 | 0.51533047 | 2.54153853 | 0.00309747 | 0.05226808 |
| ENSGALG00010000007 | ND1 | 63549479 | -0.3692288 | 12.0776809 | 0.00311294 | 0.05247337 |
| ENSGALG00010010190 | RNF44 | 416224 | 0.28299183 | 6.26154328 | 0.00315066 | 0.0530529 |
| ENSGALG00010027214 | PLIN4 | 100857433 | -0.2257161 | 6.59139885 | 0.00316962 | 0.05326047 |
| ENSGALG00010008737 | NA | NA | 1.4943444 | -0.5177497 | 0.00316971 | 0.05326047 |
| ENSGALG00010011936 | NA | NA | -0.2936746 | 7.91216262 | 0.0031733 | 0.05326437 |
| ENSGALG00010026568 | BYSL | 419927 | -0.2472446 | 6.42970544 | 0.00319385 | 0.0535527 |
| ENSGALG00010000065 | NA | NA | -0.7193494 | 1.68336071 | 0.00320717 | 0.05366766 |
| ENSGALG00010026350 | LOC101749943 | 101749943 | -0.2947975 | 5.46030869 | 0.00320747 | 0.05366766 |
| ENSGALG00010004224 | NT5E | 421833 | 0.53053018 | 3.00534168 | 0.00321225 | 0.05369097 |
| ENSGALG00010007417 | NA | NA | 1.20075762 | 0.54496832 | 0.00322653 | 0.05387281 |
| ENSGALG00010026899 | HOOK1 | 424675 | -0.3480116 | 5.30427358 | 0.00323418 | 0.05394372 |
| ENSGALG00010012407 | ATF4 | 395693 | -0.2807592 | 8.70388188 | 0.00324789 | 0.05411554 |
| ENSGALG00010014128 | COL4A1 | 395530 | 0.29640711 | 6.72596345 | 0.00326346 | 0.05431793 |
| ENSGALG00010010505 | CD38 | 422827 | 0.47465815 | 4.20446181 | 0.00327941 | 0.05447377 |
| ENSGALG00010003724 | NA | NA | 0.40400683 | 3.91048844 | 0.00327969 | 0.05447377 |
| ENSGALG00010023244 | PPIB | 396447 | -0.4076199 | 7.87265277 | 0.00328594 | 0.05447667 |
| ENSGALG00010004068 | NA | NA | 1.01918238 | 0.76981155 | 0.00328673 | 0.05447667 |
| ENSGALG00010007981 | TDRD15 | 101751663 | 0.29507935 | 5.69273447 | 0.0032935 | 0.0545318 |
| ENSGALG00010024822 | RPS26 | 100857770 | -0.2383025 | 8.37934069 | 0.00330376 | 0.054551 |

| | | | | | | |
|---|---|---|---|---|---|---|
| ENSGALG00010021848 | LARGE2 | 423200 | -0.291654 | 5.97427333 | 0.00330467 | 0.054551 |
| ENSGALG00010017495 | NT5M | 416505 | 0.33821218 | 4.41196401 | 0.00330498 | 0.054551 |
| ENSGALG00010014493 | NA | NA | 0.69006256 | 1.56015988 | 0.00331318 | 0.0546003 |
| ENSGALG00010009147 | HNMT | 424298 | 0.86631696 | 0.97822676 | 0.00331485 | 0.0546003 |
| ENSGALG00010013405 | RPS14 | 416275 | -0.2123099 | 8.46365273 | 0.00332459 | 0.05470397 |
| ENSGALG00010025370 | HPD | 416852 | -0.440182 | 3.68203518 | 0.00333308 | 0.05478686 |
| ENSGALG00010010146 | PHACTR1 | 395093 | 0.63131973 | 2.21039816 | 0.00336184 | 0.05520231 |
| ENSGALG00010021313 | KPNA7 | 416488 | -0.9486627 | 0.97802972 | 0.00336908 | 0.05526408 |
| ENSGALG00010001113 | IGSF10 | 425035 | -0.5934435 | 1.90808956 | 0.00337302 | 0.05527145 |
| ENSGALG00010020649 | MYH11 | 396211 | 0.29885611 | 5.52573625 | 0.00337983 | 0.05532594 |
| ENSGALG00010020382 | MFAP2 | 100858555 | -0.8571956 | 0.93538045 | 0.00339092 | 0.05543485 |
| ENSGALG00010016761 | GPR160 | 424992 | -0.255835 | 6.50805842 | 0.00339347 | 0.05543485 |
| ENSGALG00010007088 | NA | NA | -0.770957 | 1.65942473 | 0.0033984 | 0.05545817 |
| ENSGALG00010013056 | SPATA46 | 424370 | -0.4417228 | 3.3272942 | 0.00343426 | 0.05598582 |
| ENSGALG00010004361 | NPFFR2 | 428759 | 0.64240056 | 1.65346841 | 0.00343815 | 0.05599168 |
| ENSGALG00010013213 | SH2D1A | 100857869 | 0.71210039 | 2.13359806 | 0.00345015 | 0.05612951 |
| ENSGALG00010023921 | GRK7 | 429125 | 0.45981543 | 3.17271529 | 0.00346504 | 0.05625974 |
| ENSGALG00010007984 | EIF3H | 420288 | -0.2290612 | 7.59896271 | 0.00346525 | 0.05625974 |
| ENSGALG00010020614 | FAIM | 424270 | 0.36420785 | 4.11163861 | 0.00347379 | 0.05630642 |
| ENSGALG00010018383 | METRN | 427650 | 0.28667777 | 5.32722263 | 0.00347522 | 0.05630642 |
| ENSGALG00010012330 | RFX6 | 421737 | 1.01055976 | 0.39028179 | 0.00349494 | 0.05656812 |
| ENSGALG00010021121 | PLA2R1 | 404304 | 0.35720924 | 4.26050408 | 0.00350073 | 0.05660404 |
| ENSGALG00010019016 | PLEKHG5 | 426013 | -0.6549906 | 2.7046238 | 0.00354119 | 0.05714028 |
| ENSGALG00010005114 | PHLDB2 | 418418 | 0.21597091 | 6.31337042 | 0.00354431 | 0.05714028 |
| ENSGALG00010026002 | FTH1 | 395970 | -0.2051576 | 8.66979711 | 0.0035447 | 0.05714028 |
| ENSGALG00010010453 | DRC1 | 422009 | 0.48305705 | 3.01551126 | 0.00355606 | 0.05726513 |
| ENSGALG00010022850 | ULK1 | 416800 | 0.27944119 | 5.10855624 | 0.0035742 | 0.0574989 |
| ENSGALG00010018036 | RPL4 | 415551 | -0.2159417 | 10.3794558 | 0.00358405 | 0.05759899 |
| ENSGALG00010025212 | P2RX4 | 374166 | -0.4142239 | 3.31548543 | 0.00360703 | 0.05790967 |
| ENSGALG00010013429 | PDE4DIP | 425735 | -0.2479175 | 6.15790246 | 0.00364155 | 0.0584047 |
| ENSGALG00010021490 | ACACB | 430557 | 0.6231899 | 1.9649752 | 0.0036518 | 0.05850998 |
| ENSGALG00010002143 | SLC16A13 | 425860 | 0.57192995 | 2.25543348 | 0.0036687 | 0.05868056 |
| ENSGALG00010004146 | NAAA | 422641 | 0.7403628 | 1.62732015 | 0.00366985 | 0.05868056 |
| ENSGALG00010024849 | CCDC84 | 419787 | -0.3905759 | 3.41496858 | 0.00368028 | 0.05876483 |
| ENSGALG00010009620 | TMOD1 | 395883 | 0.41564446 | 4.25676939 | 0.00368253 | 0.05876483 |
| ENSGALG00010027718 | LOC100857333 | 100857333 | 0.77502912 | 1.09079746 | 0.0036882 | 0.05879615 |
| ENSGALG00010015482 | NA | NA | 0.73159961 | 1.19963093 | 0.00369706 | 0.05887331 |
| ENSGALG00010016139 | LURAP1L | 768654 | -0.3217072 | 4.63116643 | 0.00370046 | 0.05887331 |
| ENSGALG00010011929 | ELK3 | 417919 | 0.31330531 | 5.07579485 | 0.00370559 | 0.0588958 |
| ENSGALG00010028211 | GATAD2B | 426098 | 0.26177208 | 6.45671855 | 0.00372763 | 0.05918679 |
| ENSGALG00010022879 | LOC415616 | 415616 | 0.64568636 | 2.0469292 | 0.00375631 | 0.05958258 |
| ENSGALG00010000431 | AIFM3 | 416999 | 0.55784817 | 2.59014635 | 0.00376075 | 0.05959341 |
| ENSGALG00010016812 | LRRC31 | 424994 | -0.2141368 | 6.31087901 | 0.00377769 | 0.05980208 |
| ENSGALG00010006075 | SOX21 | 107051027 | -0.2208656 | 7.09410691 | 0.00380491 | 0.06014697 |
| ENSGALG00010019315 | PRRT3 | 416053 | -0.7315532 | 1.65085024 | 0.00380706 | 0.06014697 |
| ENSGALG00010029235 | TRIM37 | 417628 | 0.21814394 | 6.76373539 | 0.00382673 | 0.06039762 |
| ENSGALG00010018561 | ACTR1A | 423862 | -0.2398305 | 7.95825619 | 0.00383127 | 0.0604092 |
| ENSGALG00010019898 | NUDT1 | 416467 | -0.2613395 | 4.97487881 | 0.00383684 | 0.06043694 |
| ENSGALG00010005227 | EXOSC9 | 619530 | -0.2275496 | 6.25335009 | 0.00388226 | 0.06109165 |
| ENSGALG00010018397 | FAM173A | 416404 | 0.5888708 | 1.9862125 | 0.00391052 | 0.06145048 |
| ENSGALG00010007867 | ZNF516 | 421009 | 0.21064476 | 6.38168663 | 0.00391281 | 0.06145048 |
| ENSGALG00010016803 | NA | NA | 0.52751622 | 2.56768757 | 0.00396401 | 0.06215636 |
| ENSGALG00010022173 | RPL23 | 420001 | -0.2764969 | 8.16591498 | 0.00396559 | 0.06215636 |
| ENSGALG00010009972 | PTK2B | 421998 | 1.11979243 | 0.11909676 | 0.00397868 | 0.06226762 |
| ENSGALG00010026614 | NA | NA | 0.69851415 | 1.42549193 | 0.00398054 | 0.06226762 |
| ENSGALG00010014876 | PCNT | 424039 | 0.20336604 | 8.10750112 | 0.00398978 | 0.06235068 |
| ENSGALG00010005364 | MAP7 | 421689 | -0.4505743 | 3.30997606 | 0.00399921 | 0.06243651 |
| ENSGALG00010008235 | MAEL | 418446 | -0.2393508 | 8.05182304 | 0.00400614 | 0.06248322 |
| ENSGALG00010029632 | TMEM132E | 771389 | 0.40332849 | 4.40099976 | 0.00402036 | 0.06250157 |
| ENSGALG00010022597 | SLCO4A1 | 419231 | -0.4976215 | 4.09161506 | 0.00402202 | 0.06250157 |

| | | | | | | |
|---|---|---|---|---|---|---|
| ENSGALG00010023281 | B2M | 414830 | -0.2932439 | 6.18198884 | 0.00402333 | 0.06250157 |
| ENSGALG00010001119 | COMMD2 | 425044 | -0.2261942 | 7.08937855 | 0.00402543 | 0.06250157 |
| ENSGALG00010028855 | SURF6 | 417160 | -0.2813061 | 5.51372749 | 0.00402793 | 0.06250157 |
| ENSGALG00010026960 | SEC61A1 | 416023 | -0.449018 | 9.20471453 | 0.00403096 | 0.06250157 |
| ENSGALG00010016543 | TCEA3 | 100858897 | -0.9461998 | 1.26220002 | 0.00405211 | 0.06276807 |
| ENSGALG00010013402 | MARCKS | 396473 | -0.2231685 | 6.4614331 | 0.0040597 | 0.0628244 |
| ENSGALG00010024595 | MYBPHL | 421154 | 0.48149187 | 2.70537641 | 0.0040654 | 0.06285123 |
| ENSGALG00010014980 | LOC121107687 | 121107687 | -0.4265358 | 5.10880876 | 0.00408627 | 0.06311233 |
| ENSGALG00010013183 | CLTA | 427284 | -0.2217118 | 7.81110079 | 0.00409744 | 0.06316688 |
| ENSGALG00010016225 | P4HA2 | 416326 | -0.2834445 | 5.38286029 | 0.00409776 | 0.06316688 |
| ENSGALG00010003375 | YWHAQ | 421932 | -0.1991142 | 9.21293796 | 0.0041101 | 0.06329561 |
| ENSGALG00010022449 | COPZ1 | 107055416 | 0.721842 | 1.31263421 | 0.0041253 | 0.06337106 |
| ENSGALG00010000939 | NA | NA | 0.31773065 | 4.78730036 | 0.00412783 | 0.06337106 |
| ENSGALG00010009099 | CXCR4 | 395324 | -0.3272254 | 4.62661063 | 0.00412989 | 0.06337106 |
| ENSGALG00010011523 | PIGO | 101748924 | -0.2190684 | 6.26150164 | 0.00413464 | 0.06337106 |
| ENSGALG00010024758 | SPATA6 | 424626 | 0.32219511 | 4.93651893 | 0.00413498 | 0.06337106 |
| ENSGALG00010030016 | NA | NA | 0.22002664 | 6.22552126 | 0.00416597 | 0.06373219 |
| ENSGALG00010017188 | ITGAD | 101748640 | 0.35544096 | 5.10687547 | 0.00416842 | 0.06373219 |
| ENSGALG00010025642 | RPS8 | 424584 | -0.2652029 | 9.23374646 | 0.0041728 | 0.06373219 |
| ENSGALG00010028462 | EDF1 | 417286 | -0.2828926 | 7.44246679 | 0.00417462 | 0.06373219 |
| ENSGALG00010005719 | NA | NA | 0.98865938 | 1.00239637 | 0.00419485 | 0.06396321 |
| ENSGALG00010011625 | ERBB4 | 395671 | 0.32659408 | 4.7769185 | 0.00419781 | 0.06396321 |
| ENSGALG00010019832 | MAD2L1BP | 421448 | -0.2126845 | 6.24342614 | 0.00420494 | 0.06401037 |
| ENSGALG00010004801 | NA | NA | -0.5554441 | 2.26431582 | 0.00420974 | 0.06402194 |
| ENSGALG00010001229 | ZNF407 | 421011 | 0.24609751 | 6.22913997 | 0.00421527 | 0.0640447 |
| ENSGALG00010009250 | ERO1B | 421509 | -0.4200202 | 6.12972966 | 0.00423409 | 0.064269 |
| ENSGALG00010025458 | BCAS2 | 419884 | -0.239253 | 5.62480497 | 0.00423816 | 0.06426932 |
| ENSGALG00010023263 | SRP14 | 423284 | -0.2362166 | 6.59189266 | 0.00424944 | 0.06437884 |
| ENSGALG00010024169 | FAM168B | 424759 | 0.24282482 | 6.95990302 | 0.00427564 | 0.06471395 |
| ENSGALG00010004003 | HNRNPDL | 422601 | -0.2564524 | 8.06795244 | 0.00432373 | 0.06537941 |
| ENSGALG00010000028 | ND4L | 63549492 | -0.3891395 | 9.75366221 | 0.00434823 | 0.06568727 |
| ENSGALG00010015848 | UBE2B | 100857678 | -0.3625454 | 4.03671075 | 0.00435879 | 0.06578419 |
| ENSGALG00010006847 | SEL1L | 423391 | -0.4026425 | 7.69469934 | 0.00437171 | 0.06591641 |
| ENSGALG00010020880 | PXK | 416065 | -0.3396515 | 3.98400846 | 0.00438367 | 0.06603396 |
| ENSGALG00010006845 | CALML3 | 416692 | 1.05654932 | 0.38005282 | 0.00442745 | 0.06663022 |
| ENSGALG00010002706 | ROCK2 | 374182 | -0.2189671 | 5.81724752 | 0.00445383 | 0.0669637 |
| ENSGALG00010005343 | RNF17 | 418961 | -0.4404453 | 8.99652433 | 0.00446003 | 0.06699334 |
| ENSGALG00010018834 | SAMD4A | 423559 | 0.26372566 | 5.445387 | 0.00448462 | 0.06729895 |
| ENSGALG00010011335 | VCP | 427410 | -0.2004911 | 9.02405134 | 0.0045072 | 0.06750966 |
| ENSGALG00010001066 | SLC33A1 | 425029 | -0.3259585 | 5.04161771 | 0.00451047 | 0.06750966 |
| ENSGALG00010021600 | FMN1 | 386747 | 0.32731592 | 5.09402292 | 0.00451143 | 0.06750966 |
| ENSGALG00010004081 | RPS20 | 430990 | -0.2779277 | 9.11005546 | 0.00452626 | 0.06766774 |
| ENSGALG00010015661 | NA | NA | -0.3358947 | 4.23029026 | 0.00454106 | 0.06770515 |
| ENSGALG00010022315 | NA | NA | -0.314727 | 5.54405825 | 0.00454134 | 0.06770515 |
| ENSGALG00010013166 | LOC121108557 | 121108557 | 1.2909257 | -0.25833 | 0.00454156 | 0.06770515 |
| ENSGALG00010025315 | LRRC42 | 424655 | -0.2143854 | 6.20859787 | 0.00455112 | 0.06778385 |
| ENSGALG00010017980 | NA | NA | -0.3895171 | 3.39433475 | 0.00455839 | 0.06782851 |
| ENSGALG00010017131 | TLR3 | 422720 | 0.55893463 | 2.9328494 | 0.00462604 | 0.06877054 |
| ENSGALG00010013897 | AMOT | 422336 | 0.28581191 | 5.38345277 | 0.00464775 | 0.06899361 |
| ENSGALG00010020618 | NA | NA | 0.84892778 | 1.16416173 | 0.00465236 | 0.06899361 |
| ENSGALG00010016785 | PDGFRB | 770488 | 0.40309919 | 3.74512391 | 0.00465409 | 0.06899361 |
| ENSGALG00010022932 | OPCML | 395422 | 0.27312151 | 5.99278994 | 0.0046651 | 0.06909227 |
| ENSGALG00010028444 | MEF2B | 426373 | -1.0550841 | 2.531713 | 0.00467655 | 0.06919729 |
| ENSGALG00010014917 | EIF3D | 771356 | -0.2024937 | 7.58957158 | 0.00469048 | 0.06933871 |
| ENSGALG00010001136 | SERP1 | 107054146 | -0.3688726 | 7.3473773 | 0.00470006 | 0.06941552 |
| ENSGALG00010010991 | C2CD2 | 418532 | 0.25823266 | 5.40556009 | 0.00470639 | 0.06944449 |
| ENSGALG00010012903 | NA | NA | 0.23147235 | 6.11186958 | 0.00471471 | 0.06950247 |
| ENSGALG00010024690 | LOC100857858 | 100857858 | -0.2096706 | 6.07057498 | 0.0047407 | 0.06982079 |
| ENSGALG00010023199 | RPS13 | 414782 | -0.229323 | 7.86870463 | 0.00474706 | 0.06984954 |
| ENSGALG00010020700 | MTHFD2 | 426126 | -0.2660294 | 6.72984572 | 0.00476866 | 0.07010237 |

| | | | | | | | |
|---|---|---|---|---|---|---|---|
| ENSGALG00010005253 | POMT2 | | 423373 | 0.21843784 | 5.91201705 | 0.0047819 | 0.07023187 |
| ENSGALG00010022044 | PXMP4 | | 419153 | -0.2531772 | 5.13634252 | 0.00478938 | 0.0702767 |
| ENSGALG00010015869 | SLC7A2 | | 422730 | 0.4548596 | 3.15073597 | 0.00482475 | 0.07073034 |
| ENSGALG00010015637 | NA | NA | | 0.66848129 | 1.61168683 | 0.00483454 | 0.07077256 |
| ENSGALG00010004255 | RPS12 | | 421698 | -0.2653653 | 8.40634173 | 0.00483656 | 0.07077256 |
| ENSGALG00010025067 | SLC1A2 | | 423156 | 0.71645834 | 1.38812048 | 0.00484453 | 0.07082394 |
| ENSGALG00010016308 | RPS6 | | 396148 | -0.2513462 | 10.0477099 | 0.00485143 | 0.07085947 |
| ENSGALG00010007695 | RASGRP3 | | 421460 | -0.7138028 | 1.4133672 | 0.00487218 | 0.07109692 |
| ENSGALG00010012786 | TEF | | 396342 | 0.24609067 | 6.23642807 | 0.00487665 | 0.07109692 |
| ENSGALG00010011002 | RPRM | | 419946 | -0.6928009 | 1.28564514 | 0.00489775 | 0.07133552 |
| ENSGALG00010024803 | CLK1 | | 100858576 | -0.2505763 | 6.19747234 | 0.00490201 | 0.07133552 |
| ENSGALG00010006285 | FZD8 | | 374059 | -0.2367094 | 6.03915389 | 0.00492183 | 0.07155827 |
| ENSGALG00010022154 | CPN1 | | 769143 | -0.3178223 | 4.22664899 | 0.00493028 | 0.07161548 |
| ENSGALG00010010845 | LOC121108177 | | 121108177 | 0.30176865 | 4.92980867 | 0.00499835 | 0.07253779 |
| ENSGALG00010021229 | GLO1 | | 421428 | -0.2378443 | 6.27505258 | 0.00500395 | 0.07255268 |
| ENSGALG00010019570 | NA | NA | | 0.57742525 | 2.08211625 | 0.00502113 | 0.07273527 |
| ENSGALG00010024842 | ZFYVE9 | | 424639 | 0.23500381 | 6.18133322 | 0.00506072 | 0.07324189 |
| ENSGALG00010005555 | PRSS12 | | 422685 | -0.3194571 | 4.55691295 | 0.0050826 | 0.07349146 |
| ENSGALG00010025968 | FADS2 | | 423122 | -0.28046 | 6.22347048 | 0.00511674 | 0.07391773 |
| ENSGALG00010027286 | RPN1 | | 416017 | -0.259952 | 8.75497143 | 0.00517524 | 0.07462759 |
| ENSGALG00010013802 | PYROXD1 | | 418191 | 0.42846737 | 2.96382916 | 0.00517528 | 0.07462759 |
| ENSGALG00010029384 | WDR81 | | 417560 | -0.2397836 | 6.06590944 | 0.0051861 | 0.07471563 |
| ENSGALG00010012291 | NR5A2 | | 395961 | -0.7517268 | 1.49106901 | 0.00519617 | 0.07479276 |
| ENSGALG00010008236 | NA | NA | | 0.6198448 | 2.48767715 | 0.00520792 | 0.0748565 |
| ENSGALG00010011362 | NA | NA | | 0.4295635 | 3.64490988 | 0.00521447 | 0.0748565 |
| ENSGALG00010027725 | BTN1A1 | | 420061 | 0.70964966 | 1.14302923 | 0.00521732 | 0.0748565 |
| ENSGALG00010013006 | GRIK2 | | 428628 | 1.03494784 | 0.16665308 | 0.00521947 | 0.0748565 |
| ENSGALG00010017583 | MTHFD1 | | 423508 | 0.21490904 | 7.2843877 | 0.00524149 | 0.07510429 |
| ENSGALG00010005356 | DNAJC3 | | 418787 | -0.3687584 | 7.54138779 | 0.00524626 | 0.07510487 |
| ENSGALG00010029416 | UNC119 | | 771690 | -0.3918586 | 3.81743707 | 0.00527718 | 0.07539824 |
| ENSGALG00010028000 | LOC121107538 | | 121107538 | -0.3771662 | 4.80721545 | 0.00527769 | 0.07539824 |
| ENSGALG00010016789 | DISP3 | | 419497 | 0.4501395 | 2.81366398 | 0.00528101 | 0.07539824 |
| ENSGALG00010007570 | NA | NA | | 0.78575339 | 1.07732797 | 0.00530521 | 0.07567563 |
| ENSGALG00010022090 | TRRAP | | 416440 | 0.20516273 | 8.60669415 | 0.00531541 | 0.07575289 |
| ENSGALG00010005871 | EGLN3 | | 423316 | 0.20524003 | 6.97163739 | 0.00532559 | 0.07582996 |
| ENSGALG00010011360 | LOC121108224 | | 121108224 | 0.60849938 | 3.80157913 | 0.00533358 | 0.07587554 |
| ENSGALG00010008220 | NA | NA | | 0.78436085 | 0.86776764 | 0.00537834 | 0.07644381 |
| ENSGALG00010005241 | C4H4orf54 | | 112532442 | 0.26151434 | 6.95554606 | 0.00540823 | 0.07675326 |
| ENSGALG00010012207 | GRIA3 | | 396138 | 0.74884901 | 1.13236178 | 0.00540979 | 0.07675326 |
| ENSGALG00010020890 | RPS15A | | 427675 | -0.2672719 | 8.20688296 | 0.00541537 | 0.07676376 |
| ENSGALG00010018889 | NA | NA | | 1.34044959 | -0.4708373 | 0.00542781 | 0.07687135 |
| ENSGALG00010003795 | LCP1 | | 418852 | 0.64504994 | 3.14655494 | 0.00543874 | 0.07690399 |
| ENSGALG00010004887 | MYH15 | | 395534 | -0.659043 | 3.16545287 | 0.00543981 | 0.07690399 |
| ENSGALG00010010588 | SLC25A3 | | 417924 | -0.1971029 | 8.56292443 | 0.00546015 | 0.07708817 |
| ENSGALG00010022070 | NA | NA | | 1.42964376 | -0.414076 | 0.00546256 | 0.07708817 |
| ENSGALG00010009514 | PRELID1 | | 395819 | -0.2117331 | 6.55924324 | 0.00547948 | 0.07721167 |
| ENSGALG00010025832 | ENKD1 | | 415643 | -0.2543031 | 5.39460759 | 0.00548105 | 0.07721167 |
| ENSGALG00010005842 | NA | NA | | 0.88831854 | 0.8040035 | 0.00549517 | 0.07728874 |
| ENSGALG00010010604 | HIVEP1 | | 420856 | 0.20206804 | 6.29944009 | 0.00549626 | 0.07728874 |
| ENSGALG00010028046 | SMIM4 | | 415896 | -0.3425974 | 4.28098279 | 0.005531 | 0.07766864 |
| ENSGALG00010005228 | ZNF521 | | 421083 | 0.22582033 | 7.31543564 | 0.00553307 | 0.07766864 |
| ENSGALG00010027685 | LOC100857714 | | 100857714 | -1.1321633 | -0.2991546 | 0.00554967 | 0.07776537 |
| ENSGALG00010011904 | KIF24 | | 427398 | 0.25738421 | 5.72657254 | 0.00554977 | 0.07776537 |
| ENSGALG00010018051 | GNPNAT1 | | 423588 | -0.282702 | 4.46799338 | 0.00555992 | 0.07782144 |
| ENSGALG00010003061 | ABCB5 | | 420606 | 0.36732191 | 3.89492358 | 0.00556358 | 0.07782144 |
| ENSGALG00010025019 | MIP | | 374124 | -0.768204 | 0.98388335 | 0.0055741 | 0.07789985 |
| ENSGALG00010013296 | AGXT2 | | 431666 | 0.36304806 | 4.12412552 | 0.00558329 | 0.07793738 |
| ENSGALG00010026621 | NA | NA | | 0.58311842 | 2.42559002 | 0.00558661 | 0.07793738 |
| ENSGALG00010029184 | DYNLL2 | | 417663 | -0.2177356 | 6.29189332 | 0.00559678 | 0.07801072 |
| ENSGALG00010018819 | LRRC28 | | 415512 | 0.2252439 | 5.73056221 | 0.00561946 | 0.07821975 |

| | | | | | | |
|---|---|---|---|---|---|---|
| ENSGALG00010023285 | SLC48A1 | 426887 | 0.34931427 | 3.84634088 | 0.00562164 | 0.07821975 |
| ENSGALG00010024684 | CADNL | 427547 | 0.24128503 | 5.42742854 | 0.0056321 | 0.07823572 |
| ENSGALG00010027622 | FCN3 | 427762 | 0.49936818 | 3.7101582 | 0.00563567 | 0.07823572 |
| ENSGALG00010017635 | NPPC | 101748596 | -0.7628639 | 2.62289657 | 0.00563759 | 0.07823572 |
| ENSGALG00010019021 | LRR1 | 423565 | -0.2706993 | 4.93343363 | 0.0056456 | 0.07827846 |
| ENSGALG00010017348 | NA | NA | -1.1297195 | 0.15864045 | 0.00565107 | 0.0782858 |
| ENSGALG00010000310 | NA | NA | 0.51819272 | 3.22656912 | 0.00566165 | 0.07836391 |
| ENSGALG00010009269 | NA | NA | -0.4489744 | 3.59719716 | 0.00567803 | 0.07838407 |
| ENSGALG00010021113 | SYP | 396359 | 0.45167925 | 3.35368689 | 0.00567836 | 0.07838407 |
| ENSGALG00010029646 | MRPS17 | 417538 | -0.1977981 | 6.3141898 | 0.00568628 | 0.07838407 |
| ENSGALG00010015177 | TC2N | 423412 | 0.67475512 | 1.77156618 | 0.00568686 | 0.07838407 |
| ENSGALG00010012857 | NSDHL | 422302 | -0.3597743 | 4.66364709 | 0.00568781 | 0.07838407 |
| ENSGALG00010005509 | COCH | 395779 | -0.7515711 | 4.39531096 | 0.00570997 | 0.07862109 |
| ENSGALG00010027672 | CNTRL | 417121 | 0.21304396 | 6.83775617 | 0.00572151 | 0.07871169 |
| ENSGALG00010007698 | ADAMTS18 | 100858277 | -0.6494555 | 3.45846404 | 0.00574534 | 0.07897104 |
| ENSGALG00010028747 | NA | NA | 0.48271534 | 3.20545823 | 0.00575749 | 0.07899766 |
| ENSGALG00010015407 | TIMP3 | 396483 | 0.4175885 | 3.01366005 | 0.00576005 | 0.07899766 |
| ENSGALG00010023273 | HDAC7 | 426885 | 0.24120516 | 5.21159926 | 0.00576222 | 0.07899766 |
| ENSGALG00010008860 | SQLE | 420335 | -0.2220731 | 7.56019776 | 0.00577569 | 0.07911396 |
| ENSGALG00010020811 | MCOLN2 | 424531 | 0.64157049 | 1.72164009 | 0.00579826 | 0.07933953 |
| ENSGALG00010005547 | LRP11 | 421629 | 0.32112834 | 4.95718964 | 0.00580666 | 0.07933953 |
| ENSGALG00010022427 | RPL27 | 396280 | -0.1973782 | 8.34504712 | 0.00580727 | 0.07933953 |
| ENSGALG00010016384 | GNAS | 419313 | 0.22376503 | 5.63340748 | 0.00581216 | 0.07933953 |
| ENSGALG00010029935 | SYNRG | 417658 | -0.3120626 | 5.7320661 | 0.00581794 | 0.07935011 |
| ENSGALG00010024437 | HERC1 | 415367 | 0.20403845 | 8.68189616 | 0.00587153 | 0.08001227 |
| ENSGALG00010003701 | SARAF | 422624 | -0.2667392 | 6.23316645 | 0.00587666 | 0.08001339 |
| ENSGALG00010004888 | FOXA1 | 101752179 | -0.7403209 | 1.74059783 | 0.00588935 | 0.08011743 |
| ENSGALG00010025569 | PPTC7 | 416869 | -0.229493 | 5.43414114 | 0.00589748 | 0.08012141 |
| ENSGALG00010008302 | NA | NA | 0.28091836 | 4.84562949 | 0.00589975 | 0.08012141 |
| ENSGALG00010009601 | ZHX2 | 420351 | 0.20983106 | 8.07267447 | 0.00590483 | 0.08012181 |
| ENSGALG00010000078 | LOC112531364 | 112531364 | -0.3542215 | 4.47991512 | 0.00592885 | 0.08037894 |
| ENSGALG00010020990 | UBB | 396190 | 0.27948596 | 8.71025343 | 0.0059391 | 0.08044913 |
| ENSGALG00010023098 | NA | NA | 0.87211337 | 0.69158071 | 0.00594915 | 0.08051655 |
| ENSGALG00010028242 | NA | NA | 0.2363598 | 5.60016596 | 0.00596941 | 0.08072185 |
| ENSGALG00010023835 | INAFM2 | 107053356 | -0.2870455 | 4.87874876 | 0.00598158 | 0.08081758 |
| ENSGALG00010017155 | BFSP2 | 396141 | -0.403142 | 4.00449458 | 0.00600951 | 0.08112578 |
| ENSGALG00010010445 | LMAN1L | 426849 | -0.2690441 | 7.8287197 | 0.00604827 | 0.08153828 |
| ENSGALG00010018007 | DNAAF9 | 422945 | -0.5172272 | 6.40766635 | 0.00605034 | 0.08153828 |
| ENSGALG00010010754 | DNAI1 | 431654 | 0.38601947 | 3.86897997 | 0.006085 | 0.0819357 |
| ENSGALG00010024914 | NA | NA | 1.31427523 | 0.14435209 | 0.00609585 | 0.0819678 |
| ENSGALG00010029524 | BAHCC1 | 776014 | 0.32294166 | 4.96309115 | 0.00609772 | 0.0819678 |
| ENSGALG00010014984 | LPP | 429148 | 0.26079091 | 5.2664979 | 0.00614545 | 0.08247096 |
| ENSGALG00010001886 | SPCS2 | 419056 | -0.2478775 | 6.37301079 | 0.00614555 | 0.08247096 |
| ENSGALG00010010997 | LAP3 | 425306 | -0.2532976 | 6.04055443 | 0.00616219 | 0.08262434 |
| ENSGALG00010013340 | LIN28B | 421786 | 0.20265858 | 7.75358069 | 0.00620005 | 0.08306182 |
| ENSGALG00010010715 | LOC124417187 | 124417187 | 0.8562365 | 0.68767967 | 0.00620844 | 0.08310403 |
| ENSGALG00010027405 | DCLRE1B | 428269 | 0.21531847 | 6.40161394 | 0.00621464 | 0.08311681 |
| ENSGALG00010024526 | TGM6 | 771144 | 0.57309916 | 2.21780202 | 0.0062265 | 0.08320536 |
| ENSGALG00010004092 | NA | NA | 0.43302936 | 5.95309671 | 0.00623195 | 0.08320808 |
| ENSGALG00010024676 | NA | NA | 0.29665882 | 4.25997395 | 0.00627294 | 0.08358755 |
| ENSGALG00010013378 | SH3RF3 | 418731 | 0.45674189 | 2.78429922 | 0.00627585 | 0.08358755 |
| ENSGALG00010025741 | NA | NA | 0.9911738 | 0.18039093 | 0.00627618 | 0.08358755 |
| ENSGALG00010003206 | NA | NA | 1.38484123 | -0.597869 | 0.0062957 | 0.08377722 |
| ENSGALG00010014687 | CUL4A | 418744 | -0.2976161 | 4.79888425 | 0.0063092 | 0.08388647 |
| ENSGALG00010023937 | XBP1 | 416918 | -0.2757427 | 6.6545064 | 0.00632145 | 0.08397895 |
| ENSGALG00010015321 | NA | NA | -0.8326961 | 0.62772823 | 0.00632891 | 0.08399391 |
| ENSGALG00010023625 | NA | NA | -0.3317942 | 4.22626273 | 0.00633317 | 0.08399391 |
| ENSGALG00010030066 | NA | NA | -0.6527915 | 2.3725923 | 0.00635148 | 0.08416634 |
| ENSGALG00010020994 | DHCR7 | 422982 | -0.2729352 | 6.98779564 | 0.00636684 | 0.08429945 |
| ENSGALG00010029142 | RGS3 | 395110 | 1.22041775 | -0.2278017 | 0.00638441 | 0.08431332 |

| | | | | | | |
|---|---|---|---|---|---|---|
| ENSGALG00010023377 | OSER1 | 419203 | -0.3510943 | 4.1265927 | 0.00638664 | 0.08431332 |
| ENSGALG00010008429 | NA | NA | 0.45326946 | 3.04860801 | 0.00639025 | 0.08431332 |
| ENSGALG00010004948 | NA | NA | 0.82829661 | 0.92890087 | 0.00639977 | 0.08431332 |
| ENSGALG00010021792 | KLHDC8B | 416054 | -0.5194571 | 3.77400367 | 0.00640069 | 0.08431332 |
| ENSGALG00010008580 | PANX1 | 772164 | 0.24129576 | 5.69609858 | 0.00640303 | 0.08431332 |
| ENSGALG00010001133 | EIF2A | 425042 | -0.1944254 | 6.49390371 | 0.00640509 | 0.08431332 |
| ENSGALG00010011378 | TUBB2B | 420883 | -0.220568 | 6.33355065 | 0.00641904 | 0.08442693 |
| ENSGALG00010012147 | ARFGAP3 | 418224 | -0.3743454 | 6.97432857 | 0.00643901 | 0.0845456 |
| ENSGALG00010010382 | SEC11C | 426850 | -0.4479666 | 4.59345417 | 0.00644396 | 0.0845456 |
| ENSGALG00010009113 | RPLP1 | 396262 | -0.2341967 | 8.85863182 | 0.00644576 | 0.0845456 |
| ENSGALG00010012637 | NA | NA | -0.5919927 | 1.90167694 | 0.00645414 | 0.0845456 |
| ENSGALG00010000043 | PPIP5K1 | 415587 | 0.51692799 | 2.23544536 | 0.00645472 | 0.0845456 |
| ENSGALG00010003858 | SUCLA2 | 418857 | -0.2340702 | 6.29520495 | 0.00646713 | 0.08457267 |
| ENSGALG00010007593 | SEC61B | 420997 | -0.3887134 | 6.39290658 | 0.00646745 | 0.08457267 |
| ENSGALG00010023837 | STT3A | 100857165 | -0.3191652 | 8.1121038 | 0.00650671 | 0.08497086 |
| ENSGALG00010013192 | B4GALT5 | 107054826 | -0.2446703 | 6.19692448 | 0.00650861 | 0.08497086 |
| ENSGALG00010024195 | PSMA4 | 415357 | -0.1996391 | 7.11584536 | 0.00651584 | 0.0849953 |
| ENSGALG00010022622 | MPPED2 | 421604 | -0.2088415 | 6.85045618 | 0.00652752 | 0.08507765 |
| ENSGALG00010005670 | SGMS2 | 422529 | -0.3396114 | 5.0875314 | 0.00654456 | 0.08522967 |
| ENSGALG00010011541 | NA | NA | -0.9672945 | 1.44855011 | 0.00655845 | 0.08534055 |
| ENSGALG00010014018 | DPYSL3 | 395154 | -0.2123153 | 8.75851465 | 0.00656574 | 0.0853654 |
| ENSGALG00010027258 | SHC2 | 770341 | -0.4950049 | 2.7278619 | 0.00658589 | 0.08555718 |
| ENSGALG00010011345 | CRMP1 | 395156 | 0.19115723 | 6.58847781 | 0.00663399 | 0.08611163 |
| ENSGALG00010011699 | TIAM1 | 418491 | 0.25250873 | 5.22813666 | 0.00663961 | 0.086114 |
| ENSGALG00010016575 | CWH43 | 422763 | -0.40449 | 3.39290369 | 0.00665368 | 0.08618732 |
| ENSGALG00010025936 | GOSR2 | 419973 | -0.2685902 | 6.51819906 | 0.00665613 | 0.08618732 |
| ENSGALG00010014576 | HRH2 | 427635 | 1.08623207 | 0.07128794 | 0.00666574 | 0.08624137 |
| ENSGALG00010001697 | TAF10 | 107052460 | -0.3716163 | 4.82837418 | 0.00679001 | 0.08777763 |
| ENSGALG00010016919 | ARHGDIG | 100858938 | -0.4280855 | 3.22313554 | 0.00680181 | 0.0878585 |
| ENSGALG00010019520 | NA | NA | -0.4084169 | 3.4734196 | 0.00682756 | 0.08799327 |
| ENSGALG00010018302 | RAB11FIP5 | 422952 | 0.19913501 | 7.9114872 | 0.00683742 | 0.08799327 |
| ENSGALG00010024920 | VIPR2 | 420453 | 0.45383677 | 3.52377235 | 0.00684048 | 0.08799327 |
| ENSGALG00010007915 | HECW1 | 420774 | 0.29531231 | 5.09351722 | 0.00684265 | 0.08799327 |
| ENSGALG00010024834 | INSIG1 | 420442 | -0.3760561 | 4.93637489 | 0.00684516 | 0.08799327 |
| ENSGALG00010010645 | FAM219A | 768856 | -0.3213154 | 4.51119687 | 0.00684552 | 0.08799327 |
| ENSGALG00010019322 | EGR3 | 102725543 | 0.30772463 | 4.37934693 | 0.00685278 | 0.08801517 |
| ENSGALG00010009415 | HMP19 | 416212 | 0.89484923 | 0.64708115 | 0.00686799 | 0.08813921 |
| ENSGALG00010013086 | CXCL14 | 395451 | 0.66357612 | 2.71587532 | 0.00687856 | 0.08820347 |
| ENSGALG00010003087 | TWISTNB | 420603 | -0.3374212 | 4.09854997 | 0.00692783 | 0.08876355 |
| ENSGALG00010000059 | MFAP1 | 415586 | -0.2182025 | 6.57392243 | 0.00693529 | 0.08878738 |
| ENSGALG00010003817 | BF2 | 425389 | -0.2810163 | 5.71992087 | 0.00695058 | 0.08891136 |
| ENSGALG00010017678 | YARS | 419666 | -0.2360513 | 6.71351534 | 0.00696367 | 0.08900709 |
| ENSGALG00010016500 | ATPIF1 | 100857945 | -0.2394618 | 6.19673951 | 0.0069978 | 0.08937119 |
| ENSGALG00010013920 | NA | NA | 1.16884095 | 0.26094907 | 0.00700385 | 0.08937653 |
| ENSGALG00010027164 | complement C3b/C4b receptor 1 | 419854 | 1.28720464 | -0.493216 | 0.00701153 | 0.08940259 |
| ENSGALG00010022233 | FAM46C | 426544 | 0.38923266 | 3.93434151 | 0.00703127 | 0.0895823 |
| ENSGALG00010027606 | NA | NA | -0.6956142 | 1.19072289 | 0.00710763 | 0.0904825 |
| ENSGALG00010007182 | NA | NA | -0.4767445 | 3.37761799 | 0.00714849 | 0.09092976 |
| ENSGALG00010007190 | NA | NA | 0.57716239 | 2.05373443 | 0.00715769 | 0.09093213 |
| ENSGALG00010012969 | SLC12A2 | 427123 | -0.2214697 | 6.48803529 | 0.00716015 | 0.09093213 |
| ENSGALG00010003987 | NA | NA | 0.77195589 | 1.91979246 | 0.00717051 | 0.09099086 |
| ENSGALG00010019104 | POLL | 423853 | 0.33383878 | 4.2416403 | 0.00719622 | 0.09124416 |
| ENSGALG00010013438 | RPL37 | 427186 | -0.2784227 | 8.43834544 | 0.00720816 | 0.09132256 |
| ENSGALG00010019208 | PLXND1 | 416124 | 0.45481233 | 2.73317058 | 0.00724084 | 0.09166337 |
| ENSGALG00010005283 | ATP5C1 | 419108 | -0.1952901 | 6.94923919 | 0.00726319 | 0.09183609 |
| ENSGALG00010028138 | LOC101747452 | 101747452 | 0.74856752 | 0.9145106 | 0.00726606 | 0.09183609 |
| ENSGALG00010009204 | NA | NA | 0.57987076 | 2.42886785 | 0.00728986 | 0.09206349 |
| ENSGALG00010027543 | TUBB4B | 417255 | -0.2083824 | 8.69620341 | 0.00731401 | 0.09223102 |
| ENSGALG00010010291 | TCF20 | 417980 | 0.19161319 | 7.82670945 | 0.00731475 | 0.09223102 |
| ENSGALG00010002293 | GYG2 | 418660 | 0.26500792 | 5.10925615 | 0.00732883 | 0.09230441 |

| | | | | | | |
|---|---|---|---|---|---|---|
| ENSGALG00010019029 | CENPF | 395357 | 0.2008569 | 7.10330664 | 0.00733647 | 0.09230441 |
| ENSGALG00010017226 | PEF1 | 419644 | -0.2352591 | 5.17515873 | 0.00733803 | 0.09230441 |
| ENSGALG00010016209 | ADAMTS7 | 415570 | 0.25438307 | 5.71198898 | 0.00736355 | 0.092552 |
| ENSGALG00010018560 | KLC2 | 769965 | -0.2016539 | 6.13063177 | 0.00737272 | 0.09259391 |
| ENSGALG00010024953 | NCAPG2 | 100859557 | 0.20543964 | 6.1181998 | 0.00745196 | 0.09351504 |
| ENSGALG00010000569 | SIPA1 | 101749913 | 0.21233968 | 5.92943092 | 0.00746609 | 0.09361825 |
| ENSGALG00010006681 | CRISPLD2 | 415818 | 0.65070603 | 1.74950449 | 0.00748823 | 0.09375101 |
| ENSGALG00010007811 | PRCP | 428096 | 0.31921324 | 3.97445735 | 0.0074885 | 0.09375101 |
| ENSGALG00010002725 | TSPAN7 | 418577 | 0.34646292 | 3.57791856 | 0.00752951 | 0.09412968 |
| ENSGALG00010022847 | NA | NA | 0.81947764 | 0.90783797 | 0.00753061 | 0.09412968 |
| ENSGALG00010022882 | NA | NA | 0.71406616 | 1.79270955 | 0.00754533 | 0.09423938 |
| ENSGALG00010029168 | RFNG | 395789 | -0.2221429 | 5.46488516 | 0.00755236 | 0.09425293 |
| ENSGALG00010022506 | EBNA1BP2 | 424558 | -0.1996353 | 7.0591426 | 0.00759127 | 0.09465601 |
| ENSGALG00010000071 | LOC112532885 | 112532885 | 1.10400261 | 0.3509924 | 0.00759685 | 0.09465601 |
| ENSGALG00010000121 | LOC121108659 | 121108659 | 0.22192199 | 6.46355966 | 0.00760387 | 0.09465601 |
| ENSGALG00010003438 | KCNS3 | 428653 | 0.50809571 | 2.38996558 | 0.00761002 | 0.09465601 |
| ENSGALG00010027942 | KHDC4 | 425070 | -0.2023031 | 6.51902825 | 0.00761449 | 0.09465601 |
| ENSGALG00010023180 | EEF1A2 | 419244 | -0.5930309 | 2.32372808 | 0.0076404 | 0.09490365 |
| ENSGALG00010011683 | NIPAL2 | 771428 | 1.05324664 | -0.0184146 | 0.00765169 | 0.09496957 |
| ENSGALG00010006575 | SSBP2 | 770958 | 0.23220563 | 7.47097551 | 0.0077033 | 0.09553536 |
| ENSGALG00010025597 | SCN2A | 395945 | 1.4104966 | -0.5113341 | 0.00771192 | 0.09556756 |
| ENSGALG00010007588 | TTC27 | 421462 | 0.21286995 | 6.02965582 | 0.00772296 | 0.09558655 |
| ENSGALG00010028901 | PAPPA | 417245 | 0.6705937 | 1.41688495 | 0.00772551 | 0.09558655 |
| ENSGALG00010013316 | KDM7A | 418110 | 0.20551863 | 5.97600539 | 0.00773377 | 0.09561423 |
| ENSGALG00010001126 | FAM188B2 | 771731 | -0.6184212 | 1.42753463 | 0.00774476 | 0.09567556 |
| ENSGALG00010017215 | FRMD7 | 428710 | 0.94478031 | 1.0042763 | 0.00776077 | 0.09579874 |
| ENSGALG00010021874 | TMCC1 | 416125 | 0.22439806 | 5.92831582 | 0.00777955 | 0.09595584 |
| ENSGALG00010001554 | ARHGAP33L4 | 100858531 | 0.34331935 | 3.53657506 | 0.0078554 | 0.09681616 |
| ENSGALG00010027271 | FAM212B | 419868 | 0.68532331 | 1.13229982 | 0.00787382 | 0.09696781 |
| ENSGALG00010021386 | NA | NA | 0.70919166 | 2.46209164 | 0.00788578 | 0.09703976 |
| ENSGALG00010021346 | RPS21 | 419228 | -0.2594068 | 6.51726707 | 0.00789548 | 0.09708377 |
| ENSGALG00010012428 | CCNE2 | 420231 | 0.31222 | 4.37522562 | 0.00790213 | 0.09709036 |
| ENSGALG00010027966 | CLK2 | 425060 | -0.3161741 | 4.84797435 | 0.00790965 | 0.09710748 |
| ENSGALG00010018005 | NA | NA | 0.62514999 | 2.33516399 | 0.00791659 | 0.09711757 |
| ENSGALG00010029367 | DUSP14 | 417657 | -0.2432998 | 6.79933661 | 0.00793606 | 0.09728115 |
| ENSGALG00010017033 | TMEM159 | 427005 | -0.2733883 | 6.52508486 | 0.00794309 | 0.09729216 |
| ENSGALG00010006955 | HELT | 101750337 | -1.0835992 | -0.1019763 | 0.00795708 | 0.09738831 |
| ENSGALG00010024016 | CRYZ | 772289 | -0.2705315 | 4.65803611 | 0.00797854 | 0.09753763 |
| ENSGALG00010022170 | RSPRY1 | 415648 | 0.20187926 | 7.12524017 | 0.00798158 | 0.09753763 |
| ENSGALG00010008197 | STARD4 | 769322 | -0.2465679 | 5.16001617 | 0.0080005 | 0.09769023 |
| ENSGALG00010009396 | AUH | 427269 | 0.38712271 | 3.95186108 | 0.00801119 | 0.09769023 |
| ENSGALG00010022438 | RPN2 | 419177 | -0.2782846 | 8.62973182 | 0.00801255 | 0.09769023 |
| ENSGALG00010004734 | NA | NA | -0.3790797 | 3.12248452 | 0.00801879 | 0.09769126 |
| ENSGALG00010023240 | PIWIL1 | 416804 | 0.20334058 | 8.01650577 | 0.00803783 | 0.09784803 |
| ENSGALG00010016655 | TRANK1 | 420728 | 0.20777362 | 6.03904706 | 0.00804835 | 0.097901 |
| ENSGALG00010000063 | SERF2 | 107054222 | -0.4415044 | 6.26888132 | 0.00808743 | 0.09830094 |
| ENSGALG00010012376 | VGLL2 | 421735 | 0.81611409 | 0.66193447 | 0.00811826 | 0.09849346 |
| ENSGALG00010027778 | NA | NA | 1.25802565 | -0.3037861 | 0.00811913 | 0.09849346 |
| ENSGALG00010028783 | DDX31 | 427759 | 0.23047809 | 5.59497519 | 0.00812189 | 0.09849346 |
| ENSGALG00010011549 | KIAA0232 | 422860 | 0.22385763 | 6.72866672 | 0.00813617 | 0.09849891 |
| ENSGALG00010009490 | SNTB1 | 404774 | -0.462716 | 2.43651604 | 0.00813984 | 0.09849891 |
| ENSGALG00010023547 | NA | NA | 0.24659664 | 6.01886113 | 0.00814225 | 0.09849891 |
| ENSGALG00010028425 | DAP3 | 425065 | -0.2272471 | 6.07054766 | 0.00814718 | 0.09849891 |
| ENSGALG00010029473 | COG1 | 417449 | -0.2703077 | 5.15998474 | 0.00818775 | 0.09891398 |
| ENSGALG00010024711 | DDX55 | 416820 | -0.2612254 | 4.60642563 | 0.00821443 | 0.0991295 |
| ENSGALG00010015143 | EXOC1 | 422747 | -0.2274684 | 5.59843109 | 0.00822019 | 0.0991295 |
| ENSGALG00010004893 | NA | NA | 0.57139345 | 2.82369289 | 0.00822434 | 0.0991295 |
| ENSGALG00010010301 | CDHR3 | 427846 | 0.45808037 | 3.20234859 | 0.00823487 | 0.09918115 |
| ENSGALG00010008159 | SUPT3H | 422053 | -0.2666789 | 4.6370347 | 0.00824485 | 0.09922591 |
| ENSGALG00010021143 | ABCA4 | 424490 | 0.36765468 | 5.3602707 | 0.0082532 | 0.0992378 |

| | | | | | | |
|---|---|---|---|---|---|---|
| ENSGALG00010000125 | NA | NA | -0.797289 | 0.62126606 | 0.00825835 | 0.0992378 |
| ENSGALG00010002609 | PDIA6 | 421940 | -0.3196724 | 8.75229544 | 0.00826604 | 0.09925502 |
| ENSGALG00010019878 | MCFD2 | 421413 | -0.3076845 | 7.5285173 | 0.00827331 | 0.09926719 |
| ENSGALG00010018255 | MFN1 | 424973 | -0.2144785 | 6.27643054 | 0.00831901 | 0.09974009 |
| ENSGALG00010026159 | NA | NA | -1.1641262 | -0.1306367 | 0.00832906 | 0.09978518 |
| ENSGALG00010027173 | AES | 100858542 | -0.2057436 | 5.65825961 | 0.00838419 | 0.10036978 |
| ENSGALG00010019471 | NUBP1 | 416634 | -0.2396397 | 5.14429501 | 0.00841308 | 0.10061225 |
| ENSGALG00010011253 | KDM4C | 427231 | -0.5374039 | 4.97479535 | 0.00842241 | 0.10061225 |
| ENSGALG00010011997 | DENND5B | 418139 | 0.30896469 | 4.98932874 | 0.00842347 | 0.10061225 |
| ENSGALG00010016315 | RPS4Y1 | 396001 | -0.2181295 | 9.35321996 | 0.00844431 | 0.10073661 |
| ENSGALG00010001367 | IMMP2L | 417780 | 0.7263145 | 1.05123196 | 0.00844658 | 0.10073661 |
| ENSGALG00010012011 | HOOK3 | 427297 | -0.2146266 | 7.15897291 | 0.00848883 | 0.10116436 |
| ENSGALG00010029091 | NA | NA | 0.99442456 | 0.35284079 | 0.00850783 | 0.10127867 |
| ENSGALG00010002220 | NA | NA | -0.467656 | 3.07520246 | 0.00851393 | 0.10127867 |
| ENSGALG00010026195 | NA | NA | 0.93206912 | 0.58968533 | 0.00851757 | 0.10127867 |
| ENSGALG00010023212 | RIMBP2 | 374105 | 0.25311146 | 5.6522699 | 0.00856798 | 0.10180168 |
| ENSGALG00010017610 | PSMC1 | 395804 | -0.1986024 | 7.46062348 | 0.00857763 | 0.10184015 |
| ENSGALG00010007787 | C1orf198 | 768787 | -0.2080473 | 5.7390923 | 0.00861279 | 0.10218101 |
| ENSGALG00010028881 | SARDH | 417146 | 0.56889186 | 2.29507391 | 0.00863682 | 0.10238956 |
| ENSGALG00010027979 | RPL12 | 417264 | -0.2026007 | 8.3977525 | 0.00865374 | 0.10251353 |
| ENSGALG00010010807 | LOC112530796 | 112530796 | -1.035521 | 0.21697091 | 0.00866377 | 0.10255576 |
| ENSGALG00010027972 | TMEM259 | 420101 | -0.1881346 | 6.43776663 | 0.00867576 | 0.10262115 |
| ENSGALG00010029513 | WBSCR22 | 431053 | -0.2507799 | 4.90149556 | 0.00868882 | 0.10269903 |
| ENSGALG00010021184 | DRAXIN | 419492 | 0.94178949 | 1.12175585 | 0.00870091 | 0.10276527 |
| ENSGALG00010006752 | ZAR1L | 771155 | -0.2686224 | 5.38347076 | 0.00871751 | 0.10288482 |
| ENSGALG00010006139 | GAREM1 | 421098 | -0.3987233 | 3.00693219 | 0.00873758 | 0.10300938 |
| ENSGALG00010003442 | SPCS3 | 396234 | -0.3037923 | 6.57096656 | 0.00874176 | 0.10300938 |
| ENSGALG00010016795 | GUCA2B | 419498 | 0.65995522 | 1.25900904 | 0.00874755 | 0.10300938 |
| ENSGALG00010021780 | LOC101750658 | 101750658 | 0.89852334 | 0.35743266 | 0.00876241 | 0.10310777 |
| ENSGALG00010009560 | RAB24 | 427627 | -0.260087 | 5.17403217 | 0.00878984 | 0.10335386 |
| ENSGALG00010005281 | TMED8 | 423375 | 0.21651905 | 6.07875197 | 0.0088108 | 0.10352359 |
| ENSGALG00010014111 | GORAB | 770613 | -0.2379546 | 5.2032343 | 0.0088259 | 0.10362428 |
| ENSGALG00010004560 | NA | NA | 1.03020053 | 0.12580524 | 0.00885697 | 0.10385435 |
| ENSGALG00010004728 | TSC22D1 | 396005 | 0.20907664 | 5.84196402 | 0.00885859 | 0.10385435 |
| ENSGALG00010007337 | PRORP | 423325 | -0.1935887 | 6.07676566 | 0.00888416 | 0.1040063 |
| ENSGALG00010021711 | NA | NA | 0.41907567 | 2.93785082 | 0.00888466 | 0.1040063 |
| ENSGALG00010018854 | LYSMD4 | 427510 | -0.2216497 | 5.96896199 | 0.00889216 | 0.1040173 |
| ENSGALG00010026565 | RSPH14 | 416954 | 0.52347767 | 2.57647701 | 0.00891006 | 0.10414986 |
| ENSGALG00010027269 | CDC25A | 420375 | 0.23127885 | 5.41020158 | 0.0089343 | 0.10432575 |
| ENSGALG00010003642 | NA | NA | -0.7027507 | 0.8924967 | 0.00894469 | 0.10432575 |
| ENSGALG00010019704 | NA | NA | 1.18091512 | -0.285775 | 0.00894484 | 0.10432575 |
| ENSGALG00010019630 | AJAP1 | 770310 | 0.32207235 | 4.41112974 | 0.00896646 | 0.10450104 |
| ENSGALG00010020612 | RPL27A | 770018 | -0.2252724 | 8.33876938 | 0.00897921 | 0.10457281 |
| ENSGALG00010024559 | NA | NA | 0.32368906 | 3.89564916 | 0.00899583 | 0.10468956 |
| ENSGALG00010003382 | ASNS | 420574 | -0.3690046 | 7.54224835 | 0.00905037 | 0.10524702 |
| ENSGALG00010024218 | NGFR | 425805 | 0.26831479 | 4.9192263 | 0.00906606 | 0.10535223 |
| ENSGALG00010008009 | GPLD1 | 420819 | 0.94231679 | 0.79957371 | 0.00907493 | 0.1053781 |
| ENSGALG00010006220 | SGCG | 418945 | 0.34870497 | 3.73042375 | 0.00909676 | 0.10555438 |
| ENSGALG00010019665 | NA | NA | 1.3001684 | -0.3261496 | 0.0091137 | 0.10561133 |
| ENSGALG00010004985 | LOC769841 | 769841 | 1.04298824 | 0.52215951 | 0.00911499 | 0.10561133 |
| ENSGALG00010027668 | RANBP3 | 420058 | -0.1950292 | 7.64550981 | 0.00913004 | 0.10570855 |
| ENSGALG00010024731 | AOX1 | 424071 | -0.3577672 | 3.55978395 | 0.00915331 | 0.10584691 |
| ENSGALG00010016287 | BCL11B | 423444 | 0.93081134 | 0.19031006 | 0.00915989 | 0.10584691 |
| ENSGALG00010005346 | FBXO30 | 421612 | 0.20808448 | 6.38306884 | 0.00916375 | 0.10584691 |
| ENSGALG00010026609 | NA | NA | -0.3752083 | 3.41394413 | 0.0091767 | 0.10584691 |
| ENSGALG00010011328 | NA | NA | 0.66043204 | 1.81748859 | 0.00918258 | 0.10584691 |
| ENSGALG00010030070 | NA | NA | 0.98881161 | 0.31755683 | 0.00918928 | 0.10584691 |
| ENSGALG00010024611 | NA | NA | 0.2532819 | 5.07001978 | 0.00918969 | 0.10584691 |
| ENSGALG00010013021 | SLC25A5 | 772225 | -0.2074725 | 7.39406967 | 0.00919538 | 0.10584691 |
| ENSGALG00010012677 | NA | NA | 0.90290238 | 0.53933125 | 0.00924151 | 0.10630078 |

| | | | | | | |
|---|---|---|---|---|---|---|
| ENSGALG00010022508 | USF3 | 418320 | 0.20012153 | 6.19344886 | 0.00926407 | 0.10648313 |
| ENSGALG00010004283 | DCK | 422648 | 0.21526404 | 6.50264254 | 0.00930254 | 0.10680651 |
| ENSGALG00010025225 | DMRTB1 | 429107 | 0.31795339 | 6.71471463 | 0.00930567 | 0.10680651 |
| ENSGALG00010008022 | SLC35F3 | 421518 | 0.27743493 | 4.52987723 | 0.00934074 | 0.10713147 |
| ENSGALG00010005724 | PROSER2 | 419101 | 0.2621931 | 4.62271365 | 0.00935232 | 0.10713281 |
| ENSGALG00010009727 | CDH20 | 373927 | 0.25945905 | 5.79504157 | 0.00935437 | 0.10713281 |
| ENSGALG00010020309 | GXYLT2 | 416097 | -0.2181366 | 5.37745516 | 0.00936593 | 0.10715734 |
| ENSGALG00010009431 | SMN | 374025 | -0.1902373 | 7.71312084 | 0.00937002 | 0.10715734 |
| ENSGALG00010016049 | ZFYVE21 | 423486 | -0.2920131 | 4.76381671 | 0.00939423 | 0.1073357 |
| ENSGALG00010024251 | TIGAR | 419040 | -0.4952313 | 2.65240138 | 0.00940122 | 0.1073357 |
| ENSGALG00010003632 | TECRL | 422618 | 1.00968314 | 0.80202297 | 0.00940592 | 0.1073357 |
| ENSGALG00010011966 | SLC25A24 | 424337 | 0.25976472 | 4.89351647 | 0.00943627 | 0.1076047 |
| ENSGALG00010029685 | ATP5H | 422115 | -0.2156759 | 5.54107692 | 0.00948474 | 0.10807972 |
| ENSGALG00010027951 | FAM32A | 420150 | -0.2642056 | 5.50075049 | 0.00950965 | 0.10828575 |
| ENSGALG00010023740 | PPP1R16B | 419170 | 0.37921048 | 3.39989087 | 0.00951808 | 0.10830402 |
| ENSGALG00010016856 | NOD1 | 420677 | 0.28011591 | 4.28174877 | 0.00952895 | 0.10834998 |
| ENSGALG00010013648 | NAV3 | 417869 | 0.24784147 | 5.30755161 | 0.00954392 | 0.10844009 |
| ENSGALG00010007297 | NA | NA | -1.0184631 | -0.0570906 | 0.00955055 | 0.10844009 |
| ENSGALG00010021228 | ESRP2 | 415710 | -0.1939274 | 7.96553205 | 0.00955957 | 0.10846486 |
| ENSGALG00010000232 | TMEM147 | 112531206 | 0.25498274 | 4.58244783 | 0.00957555 | 0.10851913 |
| ENSGALG00010027085 | PIM1 | 100858838 | -0.2882805 | 5.49919894 | 0.00957839 | 0.10851913 |
| ENSGALG00010026949 | NA | NA | 0.25935691 | 5.3062894 | 0.00958488 | 0.10851913 |
| ENSGALG00010007686 | NA | NA | 0.51340344 | 2.67697097 | 0.0095991 | 0.10860267 |
| ENSGALG00010009638 | RND3 | 424312 | 0.19921698 | 6.98892852 | 0.00962886 | 0.10886167 |
| ENSGALG00010020514 | CAMKMT | 421406 | -0.3997741 | 3.01323899 | 0.00966199 | 0.10915845 |
| ENSGALG00010028467 | MADCAM1 | 100858236 | 0.58322979 | 1.72523105 | 0.00975237 | 0.10993598 |
| ENSGALG00010010388 | NA | NA | 0.4831026 | 2.74206734 | 0.00975436 | 0.10993598 |
| ENSGALG00010026062 | ITGA4 | 424121 | -0.2981326 | 5.55995014 | 0.00976602 | 0.10993598 |
| ENSGALG00010009330 | EIF3K | 101750374 | -0.2360666 | 7.2458335 | 0.00977176 | 0.10993598 |
| ENSGALG00010012358 | STK10 | 416204 | -0.5806546 | 2.751377 | 0.00977238 | 0.10993598 |
| ENSGALG00010020015 | ATP6V1D | 423280 | -0.1849438 | 6.62431023 | 0.00977426 | 0.10993598 |
| ENSGALG00010003189 | ANTXR2 | 422580 | 0.25046167 | 4.84712901 | 0.00977933 | 0.10993598 |
| ENSGALG00010004375 | NA | NA | 0.4963528 | 2.5232811 | 0.00979612 | 0.11004678 |
| ENSGALG00010029542 | AATK | 422076 | 0.30730825 | 3.80733144 | 0.00983732 | 0.11039542 |
| ENSGALG00010025233 | WDR12 | 424089 | -0.187315 | 6.29919067 | 0.00984722 | 0.11039542 |
| ENSGALG00010017140 | UQCRC2 | 427009 | -0.1937867 | 6.99710109 | 0.00984804 | 0.11039542 |
| ENSGALG00010013000 | NA | NA | 1.12740945 | 0.03502771 | 0.009858 | 0.11041261 |
| ENSGALG00010023119 | NDRG4 | 768618 | -0.3097065 | 4.01554716 | 0.00986349 | 0.11041261 |
| ENSGALG00010011386 | CTSB | 396329 | -0.2265257 | 6.88884498 | 0.00987538 | 0.11046775 |
| ENSGALG00010021771 | ANKRD61 | 101752127 | -0.1971122 | 6.19963498 | 0.00989359 | 0.11059342 |
| ENSGALG00010026095 | MIF | 100857237 | -0.2421789 | 7.71252868 | 0.00991858 | 0.11079476 |
| ENSGALG00010027319 | MST1R | 427573 | -0.3640351 | 3.85931249 | 0.00998128 | 0.11141659 |
| ENSGALG00010003737 | SRP72 | 422626 | -0.2110764 | 7.27376371 | 0.00999741 | 0.11151819 |
| ENSGALG00010000786 | ANKRD33B | 420924 | 1.17519876 | -0.3422493 | 0.01002558 | 0.11175389 |
| ENSGALG00010000711 | SPRY2 | 395584 | 0.23145167 | 6.12542071 | 0.01004122 | 0.11184963 |
| ENSGALG00010019261 | NA | NA | -1.1040973 | 0.3414291 | 0.01009781 | 0.11238466 |
| ENSGALG00010001771 | WDR73 | 425698 | 0.27282432 | 5.26810873 | 0.01010343 | 0.11238466 |
| ENSGALG00010006944 | MAB21L2 | 374011 | -0.5708196 | 1.7813042 | 0.01014229 | 0.1127379 |
| ENSGALG00010021243 | DYL1 | 427708 | -0.1868002 | 7.08198091 | 0.0101995 | 0.11329449 |
| ENSGALG00010002853 | CEP162 | 421835 | 0.24479138 | 5.49347708 | 0.0102728 | 0.11402772 |
| ENSGALG00010024212 | RIMS4 | 428142 | -0.2527151 | 5.1881674 | 0.01028101 | 0.11402772 |
| ENSGALG00010021893 | FAM43B | 100858896 | -0.41505 | 3.33378807 | 0.01028708 | 0.11402772 |
| ENSGALG00010016903 | PPM1A | 423525 | -0.1902062 | 7.02521754 | 0.01031341 | 0.11415098 |
| ENSGALG00010016658 | ZAR1 | 100857604 | -0.2306492 | 6.91945254 | 0.01032126 | 0.11415098 |
| ENSGALG00010025905 | CPT1A | 423118 | 0.27013427 | 4.83954025 | 0.0103227 | 0.11415098 |
| ENSGALG00010017419 | B9D1 | 416525 | 0.20341677 | 5.59349146 | 0.01032699 | 0.11415098 |
| ENSGALG00010017141 | AFTPH | 421277 | -0.2225453 | 5.53362804 | 0.01033604 | 0.11417147 |
| ENSGALG00010001273 | SI | 425007 | 0.40447265 | 3.38550222 | 0.01035333 | 0.11428287 |
| ENSGALG00010024463 | FBXL22 | 101749169 | 0.34250967 | 4.04038438 | 0.01037911 | 0.11448783 |
| ENSGALG00010002522 | HPSE | 373981 | 0.44586018 | 2.40703078 | 0.01039825 | 0.1146192 |

| | | | | | | |
|---|---|---|---|---|---|---|
| ENSGALG00010010320 | SYPL1 | 417708 | 0.64337576 | 1.52483784 | 0.01043367 | 0.11486444 |
| ENSGALG00010000554 | LOC121108774 | 121108774 | -0.6037184 | 2.70840896 | 0.01044087 | 0.11486444 |
| ENSGALG00010002186 | ICK | 421896 | 0.24157972 | 6.22414889 | 0.01044308 | 0.11486444 |
| ENSGALG00010026690 | KRT14 | 408039 | 1.06970441 | -0.2205519 | 0.01044946 | 0.11486444 |
| ENSGALG00010024363 | EDEM2 | 770034 | -0.2934366 | 4.49413433 | 0.01049115 | 0.11524277 |
| ENSGALG00010019276 | NA | NA | 0.61873342 | 1.45741162 | 0.01050258 | 0.11524943 |
| ENSGALG00010014068 | NA | NA | 0.90386918 | 0.49508212 | 0.01050628 | 0.11524943 |
| ENSGALG00010017135 | LOC100859848 | 100859848 | -0.342542 | 3.72267541 | 0.01051863 | 0.11530509 |
| ENSGALG00010022772 | NA | NA | 0.43552544 | 3.68515246 | 0.01053132 | 0.11536449 |
| ENSGALG00010012320 | DPY19L4 | 420229 | 0.18230179 | 7.12130451 | 0.01054739 | 0.11546079 |
| ENSGALG00010029288 | PYCR1 | 769203 | -0.2496831 | 7.69404237 | 0.01056165 | 0.11553721 |
| ENSGALG00010029617 | POR | 417520 | 0.23837103 | 5.90236303 | 0.01058499 | 0.11571264 |
| ENSGALG00010009991 | LHFPL3 | 100858127 | 0.26612661 | 5.1319014 | 0.01059255 | 0.11571561 |
| ENSGALG00010018822 | TRAF5 | 374055 | 0.24872673 | 4.78398563 | 0.01060102 | 0.11572843 |
| ENSGALG00010013881 | NA | NA | -0.399578 | 3.44188567 | 0.01060968 | 0.1157433 |
| ENSGALG00010020568 | PLA2G4F | 428871 | 0.71457723 | 0.92370111 | 0.01062391 | 0.11581882 |
| ENSGALG00010002520 | TTC32 | 769943 | 0.31909251 | 3.82380778 | 0.01066056 | 0.11613861 |
| ENSGALG00010026723 | NA | NA | 1.25640422 | -0.5959977 | 0.01068611 | 0.11626362 |
| ENSGALG00010021725 | SH2D6 | 395733 | -0.2432608 | 4.8781611 | 0.0106867 | 0.11626362 |
| ENSGALG00010009470 | ZNF469 | 101748749 | 0.58059059 | 3.41024654 | 0.01073945 | 0.11671271 |
| ENSGALG00010023422 | WLS | 424707 | 0.20945656 | 7.0673302 | 0.01074269 | 0.11671271 |
| ENSGALG00010028463 | RPS28 | 768930 | -0.2215554 | 7.37221451 | 0.01075103 | 0.11672337 |
| ENSGALG00010020340 | DCAF5 | 428876 | -0.2291541 | 5.67260377 | 0.01077348 | 0.11685043 |
| ENSGALG00010008892 | LOC421982 | 421982 | -0.3333246 | 4.15045971 | 0.01077747 | 0.11685043 |
| ENSGALG00010016126 | NA | NA | 1.11463353 | 0.36890845 | 0.01080128 | 0.11702863 |
| ENSGALG00010011434 | NPL | 429074 | -0.2456177 | 6.0927497 | 0.01081279 | 0.1170734 |
| ENSGALG00010005445 | SFT2D2 | 418373 | 0.19332307 | 7.58867273 | 0.0108473 | 0.11736692 |
| ENSGALG00010007324 | GAN | 430553 | 0.26705806 | 4.63616363 | 0.01087551 | 0.1175277 |
| ENSGALG00010008311 | SRP19 | 770061 | -0.2347451 | 5.08093553 | 0.01087698 | 0.1175277 |
| ENSGALG00010025404 | CERS6 | 771503 | -0.2494534 | 6.15613701 | 0.01089019 | 0.11759038 |
| ENSGALG00010024525 | GP1BB | 107051576 | 0.56740553 | 1.63276256 | 0.01090517 | 0.11767192 |
| ENSGALG00010013147 | LOC107051951 | 107051951 | 0.38252123 | 3.28095888 | 0.01092641 | 0.11782105 |
| ENSGALG00010029728 | TNRC6C | 417348 | 0.19302731 | 7.05409945 | 0.0109594 | 0.11806906 |
| ENSGALG00010002419 | CTBP2 | 423958 | -0.3197802 | 5.0905432 | 0.0109643 | 0.11806906 |
| ENSGALG00010023168 | SPHKAP | 424789 | 0.32266766 | 3.9117891 | 0.01098734 | 0.11823684 |
| ENSGALG00010012898 | FAM114A2 | 416262 | -0.3007817 | 5.58832298 | 0.01103902 | 0.11867237 |
| ENSGALG00010010022 | KMT2E | 417712 | -0.1976725 | 6.41284166 | 0.01104277 | 0.11867237 |
| ENSGALG00010001222 | ZNF800 | 417747 | 0.35497109 | 3.77699916 | 0.01110968 | 0.11931061 |
| ENSGALG00010011333 | KCTD12B | 425504 | -0.600022 | 1.40432719 | 0.01112092 | 0.11934917 |
| ENSGALG00010012749 | RGS4 | 378901 | 0.38128299 | 3.34639746 | 0.011131 | 0.11934917 |
| ENSGALG00010008123 | LOC121112980 | 121112980 | 0.20363446 | 6.0620712 | 0.01114022 | 0.11934917 |
| ENSGALG00010029676 | FADS6 | 428683 | -0.6423976 | 1.64989709 | 0.01114337 | 0.11934917 |
| ENSGALG00010008421 | OPN5 | 428670 | 1.10368505 | -0.2685244 | 0.0111647 | 0.11949694 |
| ENSGALG00010013470 | RPL31 | 418710 | -0.2385386 | 7.98798452 | 0.01118717 | 0.11965675 |
| ENSGALG00010025738 | SOX13 | 419820 | -0.2386707 | 5.19732268 | 0.01121886 | 0.11988493 |
| ENSGALG00010020336 | OSBP | 101747893 | -0.1930955 | 6.62696224 | 0.01122362 | 0.11988493 |
| ENSGALG00010022954 | ADRM1 | 395365 | -0.1832191 | 6.26018958 | 0.01127567 | 0.12035977 |
| ENSGALG00010016286 | ULK4 | 420713 | 0.4134461 | 3.5430823 | 0.01131569 | 0.12070527 |
| ENSGALG00010025231 | NA | NA | 0.77378972 | 1.03226079 | 0.01132325 | 0.12070527 |
| ENSGALG00010029353 | GCGR | 425670 | -0.4663032 | 3.03031594 | 0.01139934 | 0.12143469 |
| ENSGALG00010020981 | CIT | 416984 | 0.19286205 | 6.80738373 | 0.01140892 | 0.12145527 |
| ENSGALG00010009388 | TMED7 | 769360 | -0.2149915 | 8.67220827 | 0.0115265 | 0.12258114 |
| ENSGALG00010022138 | SPAG17 | 418315 | 0.52737624 | 2.30828144 | 0.01153014 | 0.12258114 |
| ENSGALG00010025974 | EEF1B2 | 395723 | 0.90161046 | 0.45654796 | 0.0115729 | 0.12294799 |
| ENSGALG00010012880 | CLK4 | 416295 | -0.1939245 | 5.85389054 | 0.01158015 | 0.12294799 |
| ENSGALG00010003473 | LOC112531056 | 112531056 | 0.20507839 | 7.00595694 | 0.01160664 | 0.1231468 |
| ENSGALG00010018815 | NA | NA | -0.2275419 | 5.32275708 | 0.01162295 | 0.1232375 |
| ENSGALG00010026591 | NA | NA | 0.76396087 | 0.82387146 | 0.01163489 | 0.12328163 |
| ENSGALG00010012755 | CLVS2 | 428616 | -0.4273092 | 4.01465293 | 0.01167065 | 0.12357803 |
| ENSGALG00010003853 | NA | NA | 0.44831955 | 2.54713427 | 0.01169985 | 0.1238045 |

| | | | | | | |
|---|---|---|---|---|---|---|
| ENSGALG00010003528 | GDAP1 | 420191 | -0.3430816 | 4.14241332 | 0.01172949 | 0.12403544 |
| ENSGALG00010007878 | PSMB1 | 421551 | -0.1813417 | 6.54947097 | 0.01175843 | 0.12421844 |
| ENSGALG00010020955 | FHAD1 | 771735 | 0.91848867 | 0.21791554 | 0.01176246 | 0.12421844 |
| ENSGALG00010016894 | BTBD7 | 423424 | 0.18672236 | 6.85061699 | 0.01180639 | 0.12454338 |
| ENSGALG00010018212 | LOC112530238 | 112530238 | 0.93950526 | 0.22103653 | 0.01180893 | 0.12454338 |
| ENSGALG00010005089 | NA | NA | 0.88053268 | 0.19393025 | 0.01181872 | 0.1245597 |
| ENSGALG00010023502 | NA | NA | 0.64813787 | 1.68062631 | 0.01183206 | 0.1245597 |
| ENSGALG00010014148 | KCNG1 | 419347 | 0.23979977 | 6.05256773 | 0.01183404 | 0.1245597 |
| ENSGALG00010025246 | NA | NA | 0.60665636 | 3.13360593 | 0.01187117 | 0.12486767 |
| ENSGALG00010016523 | LOC121107350 | 121107350 | 0.75664902 | 1.13792896 | 0.01188584 | 0.12488929 |
| ENSGALG00010025073 | IL18 | 395312 | -0.6156693 | 2.01740645 | 0.01188897 | 0.12488929 |
| ENSGALG00010021239 | USP54 | 423746 | 0.17969157 | 6.91704622 | 0.01189798 | 0.12490117 |
| ENSGALG00010028807 | NA | NA | 0.3475234 | 4.74375401 | 0.01191335 | 0.12497985 |
| ENSGALG00010001461 | ASB9 | 418625 | -0.9650741 | 0.04811748 | 0.01198594 | 0.12565826 |
| ENSGALG00010006510 | LOC121110508 | 121110508 | -0.3274641 | 4.08350832 | 0.0120272 | 0.12600755 |
| ENSGALG00010020728 | TMEM37 | 770956 | 0.33918426 | 3.63988127 | 0.01204174 | 0.12607662 |
| ENSGALG00010028762 | CACNA1B | 374169 | 0.53030665 | 1.93245662 | 0.01205785 | 0.12616197 |
| ENSGALG00010024528 | TBX1 | 771328 | 0.40908718 | 2.93880498 | 0.0120842 | 0.12635437 |
| ENSGALG00010025383 | SNORD12C | 100858469 | -0.8118023 | 1.21957402 | 0.01211005 | 0.12654122 |
| ENSGALG00010005223 | GRIA4 | 396137 | 0.61297 | 1.45328167 | 0.0121722 | 0.12704349 |
| ENSGALG00010013181 | DYRK1A | 395233 | 0.18265575 | 6.8060447 | 0.01217933 | 0.12704349 |
| ENSGALG00010023886 | CYP2C18 | 414833 | 0.98016208 | 0.07911219 | 0.01218214 | 0.12704349 |
| ENSGALG00010017622 | RAB1B | 421273 | -0.1913315 | 8.2431814 | 0.01220079 | 0.12711115 |
| ENSGALG00010011774 | ANKRA2 | 426153 | -0.2196093 | 6.66594441 | 0.01220466 | 0.12711115 |
| ENSGALG00010022190 | ACVR1B | 768878 | 0.21752742 | 5.53688346 | 0.01221917 | 0.12717877 |
| ENSGALG00010003674 | AKIRIN2 | 421821 | -0.1769838 | 7.99169373 | 0.01223637 | 0.12721565 |
| ENSGALG00010007357 | NOX4 | 419012 | 0.37239405 | 3.62520158 | 0.01223875 | 0.12721565 |
| ENSGALG00010023696 | LRRC4C | 428862 | 0.22725252 | 5.46452515 | 0.01229558 | 0.12770822 |
| ENSGALG00010000230 | LOC107051217 | 107051217 | 0.23137879 | 7.51308602 | 0.01230224 | 0.12770822 |
| ENSGALG00010017644 | RBBP6 | 416569 | -0.1772664 | 7.8966047 | 0.01232127 | 0.12775183 |
| ENSGALG00010015674 | RFKL | 431449 | -0.2111688 | 6.22672162 | 0.01232801 | 0.12775183 |
| ENSGALG00010027918 | CSNK1G2 | 100857862 | -0.2100531 | 5.98738753 | 0.01233061 | 0.12775183 |
| ENSGALG00010021799 | LOC416055 | 416055 | -0.3254076 | 4.48366881 | 0.01239202 | 0.1283043 |
| ENSGALG00010004442 | NA | NA | 0.55029802 | 2.14723674 | 0.01241443 | 0.12845253 |
| ENSGALG00010027156 | ISYNA1 | 107055367 | 0.20418309 | 5.96658082 | 0.01243206 | 0.12848693 |
| ENSGALG00010020992 | KCNH7 | 424184 | 0.33669794 | 4.18323377 | 0.01243396 | 0.12848693 |
| ENSGALG00010018061 | NDUFAF5 | 416742 | -0.3080264 | 3.75559456 | 0.01245948 | 0.12856199 |
| ENSGALG00010024101 | CNEP1R1 | 415736 | -0.2839828 | 4.83093417 | 0.01246853 | 0.12856199 |
| ENSGALG00010013157 | NA | NA | 0.38290196 | 3.46536701 | 0.01247125 | 0.12856199 |
| ENSGALG00010029285 | CUX1 | 101749861 | 0.23163672 | 5.18317085 | 0.01247834 | 0.12856199 |
| ENSGALG00010029812 | SAP30BP | 417375 | -0.1952932 | 5.88823276 | 0.01248175 | 0.12856199 |
| ENSGALG00010025874 | MDH1B | 424097 | 0.56824168 | 2.92552629 | 0.01249248 | 0.12857611 |
| ENSGALG00010026973 | MGLL | 416024 | 0.59768506 | 2.2215474 | 0.01249933 | 0.12857611 |
| ENSGALG00010029221 | AXIN2 | 395152 | -0.3809424 | 3.69889247 | 0.01253987 | 0.1289095 |
| ENSGALG00010022857 | NA | NA | 0.44454034 | 2.38970177 | 0.01255937 | 0.12902639 |
| ENSGALG00010017599 | KCNK13 | 772210 | -0.6456404 | 2.25775292 | 0.01259228 | 0.12928077 |
| ENSGALG00010015127 | RPL11 | 419682 | -0.2134913 | 8.03706511 | 0.01270992 | 0.13040408 |
| ENSGALG00010022675 | DMBX1 | 395271 | 0.30645713 | 4.89488527 | 0.01276041 | 0.13083753 |
| ENSGALG00010001696 | FAM135A | 421871 | 0.22692352 | 6.01381138 | 0.01278549 | 0.13089446 |
| ENSGALG00010018912 | NA | NA | 0.38196122 | 3.55610756 | 0.01279254 | 0.13089446 |
| ENSGALG00010024040 | FER1L4 | 419135 | -0.590927 | 1.5741461 | 0.01279335 | 0.13089446 |
| ENSGALG00010018959 | UBA5 | 414879 | -0.2578991 | 5.77374208 | 0.01279897 | 0.13089446 |
| ENSGALG00010003077 | EIF2S3 | 418597 | -0.2165073 | 8.88276027 | 0.01281538 | 0.13093001 |
| ENSGALG00010002385 | MPP6 | 420622 | 0.19096511 | 6.10818973 | 0.01281896 | 0.13093001 |
| ENSGALG00010024276 | NA | NA | -0.5130119 | 2.31295362 | 0.01284437 | 0.13110515 |
| ENSGALG00010009362 | IL15 | 395258 | 0.56255872 | 2.16514832 | 0.01285912 | 0.13117133 |
| ENSGALG00010006103 | NXPE3 | 418403 | 0.24209235 | 4.87352518 | 0.01288133 | 0.13126361 |
| ENSGALG00010012768 | EPS8 | 418174 | 0.24879341 | 4.50924896 | 0.01288864 | 0.13126361 |
| ENSGALG00010028473 | CDC34 | 425483 | -0.2059472 | 5.80890341 | 0.01289489 | 0.13126361 |
| ENSGALG00010008255 | C2H9ORF30 | 420991 | 0.35131477 | 4.29323061 | 0.01290813 | 0.13126361 |

| | | | | | | |
|---|---|---|---|---|---|---|
| ENSGALG00010028037 | LOC427783 | 427783 | 0.37536565 | 3.29332249 | 0.0129156 | 0.13126361 |
| ENSGALG00010015624 | ITGA1 | 395951 | 0.81618056 | 0.41182597 | 0.01291782 | 0.13126361 |
| ENSGALG00010019542 | NA | NA | 0.364924 | 3.15618259 | 0.01295819 | 0.13156579 |
| ENSGALG00010000334 | NA | NA | 0.7306766 | 1.08828474 | 0.01296415 | 0.13156579 |
| ENSGALG00010006446 | LSAMP | 395687 | 0.31595983 | 3.92588524 | 0.01297912 | 0.13161221 |
| ENSGALG00010016571 | NA | NA | 1.04334104 | 0.03435592 | 0.01298532 | 0.13161221 |
| ENSGALG00010002353 | CMTM8 | 420665 | 0.40968321 | 3.08699153 | 0.01301047 | 0.13165785 |
| ENSGALG00010014819 | LOC112532945 | 112532945 | -0.8124237 | 2.03872694 | 0.01301136 | 0.13165785 |
| ENSGALG00010027414 | PBX3 | 417093 | -0.4781571 | 2.91543094 | 0.01301472 | 0.13165785 |
| ENSGALG00010014834 | LOC112532956 | 112532956 | -0.8746804 | 0.90379138 | 0.01304022 | 0.13183172 |
| ENSGALG00010001831 | CSF2RA | 418666 | 0.9684253 | 0.41712529 | 0.01306358 | 0.13196566 |
| ENSGALG00010016879 | RNF130 | 416281 | 0.21266144 | 5.78817028 | 0.01307011 | 0.13196566 |
| ENSGALG00010028588 | ABHD8 | 100857580 | -0.5177923 | 2.77893767 | 0.01314375 | 0.13253364 |
| ENSGALG00010023155 | CCDC113 | 768642 | -0.2928025 | 4.26980738 | 0.01314391 | 0.13253364 |
| ENSGALG00010008695 | NA | NA | 0.38987178 | 3.01484957 | 0.01315143 | 0.13253364 |
| ENSGALG00010019743 | KCNU1 | 100857499 | 0.35601695 | 3.45385437 | 0.01320686 | 0.13291815 |
| ENSGALG00010005170 | GPATCH2L | 423368 | 0.19086416 | 5.86008772 | 0.01321623 | 0.13291815 |
| ENSGALG00010015628 | UBQLN1 | 427454 | -0.1873779 | 6.88610696 | 0.01321987 | 0.13291815 |
| ENSGALG00010025563 | TSPEAR | 424836 | -0.3332884 | 3.60607706 | 0.0132345 | 0.13291815 |
| ENSGALG00010010393 | FANCC | 427468 | 0.20167008 | 7.3896582 | 0.01323592 | 0.13291815 |
| ENSGALG00010017283 | INF2 | 423492 | -0.3131528 | 4.7037658 | 0.01323986 | 0.13291815 |
| ENSGALG00010026472 | NA | NA | -0.4410065 | 2.55735609 | 0.01325441 | 0.13294069 |
| ENSGALG00010009451 | ROR2 | 769516 | -0.206421 | 6.61491132 | 0.01325887 | 0.13294069 |
| ENSGALG00010022642 | SEC13 | 416119 | -0.1930477 | 6.8338219 | 0.01327925 | 0.13306096 |
| ENSGALG00010023861 | SLC2A11L5 | 416916 | -0.2479387 | 4.69040959 | 0.01332871 | 0.13343319 |
| ENSGALG00010028340 | NA | NA | 0.34854011 | 3.52225617 | 0.0133386 | 0.13343319 |
| ENSGALG00010008739 | VSTM5 | 100859821 | 0.42066978 | 2.68062285 | 0.01334164 | 0.13343319 |
| ENSGALG00010028090 | NA | NA | 0.70930367 | 1.95429064 | 0.01340913 | 0.13395011 |
| ENSGALG00010028661 | NA | NA | -0.5186354 | 1.90244483 | 0.01341021 | 0.13395011 |
| ENSGALG00010029765 | KCNJ16 | 427812 | 1.01477317 | -0.1526057 | 0.01342759 | 0.13403933 |
| ENSGALG00010013538 | E2F7 | 417868 | 0.2868899 | 4.33311818 | 0.013453 | 0.13420849 |
| ENSGALG00010008399 | INO80C | 420956 | 0.29587532 | 4.52822852 | 0.0134755 | 0.13434849 |
| ENSGALG00010025159 | NA | NA | 0.84910041 | 0.60669353 | 0.01348633 | 0.13437194 |
| ENSGALG00010018171 | LOC101748601 | 101748601 | 0.98614824 | -0.2185209 | 0.01350049 | 0.13442858 |
| ENSGALG00010008295 | ATP5SL | 107050327 | 0.66497361 | 1.14709646 | 0.01350947 | 0.13443364 |
| ENSGALG00010001692 | ALDH18A1 | 423976 | -0.3816831 | 8.69322693 | 0.01353299 | 0.13458328 |
| ENSGALG00010001928 | KCTD21 | 428098 | 0.34059039 | 4.69588892 | 0.0135515 | 0.13463311 |
| ENSGALG00010024530 | RDH8L | 429086 | 0.61673572 | 1.79563735 | 0.01355498 | 0.13463311 |
| ENSGALG00010024857 | SLC37A4 | 419789 | -0.1819174 | 8.25724746 | 0.01356777 | 0.13464203 |
| ENSGALG00010005033 | USP14 | 421063 | -0.1815995 | 7.08170876 | 0.01357285 | 0.13464203 |
| ENSGALG00010017553 | EDA | 769069 | -0.339974 | 4.86450104 | 0.01359095 | 0.13473723 |
| ENSGALG00010019532 | NA | NA | 0.59967671 | 1.27513766 | 0.01361204 | 0.13486204 |
| ENSGALG00010011166 | SOX7 | 771337 | 0.52845331 | 2.7716961 | 0.01364249 | 0.13507941 |
| ENSGALG00010014077 | LOC121106448 | 121106448 | 0.21687565 | 5.38544208 | 0.01365313 | 0.13510036 |
| ENSGALG00010004384 | LOC101751416 | 101751416 | -1.1547496 | -0.5471336 | 0.01371006 | 0.13557917 |
| ENSGALG00010011285 | NA | NA | -0.5216792 | 1.68189881 | 0.01372103 | 0.1356031 |
| ENSGALG00010018704 | PGPEP1L | 100859846 | 0.42567203 | 2.47142324 | 0.01375612 | 0.13583836 |
| ENSGALG00010021381 | DNAJB2 | 427022 | -0.2371865 | 4.9844636 | 0.01376196 | 0.13583836 |
| ENSGALG00010003289 | NA | NA | 0.59301685 | 1.33461053 | 0.01378365 | 0.1359678 |
| ENSGALG00010007897 | EFHC1 | 422041 | 0.44539024 | 3.05941351 | 0.013809 | 0.13605217 |
| ENSGALG00010023843 | NA | NA | 0.5632592 | 1.57763438 | 0.01381818 | 0.13605217 |
| ENSGALG00010019624 | SRGAP3 | 416115 | -0.2120175 | 5.91569514 | 0.01382014 | 0.13605217 |
| ENSGALG00010022165 | FAM192A | 415647 | -0.200013 | 6.44650566 | 0.01382651 | 0.13605217 |
| ENSGALG00010029604 | CASP1 | 395764 | -0.361557 | 3.85769258 | 0.01384628 | 0.13614803 |
| ENSGALG00010016486 | NA | NA | 0.69856197 | 1.01546494 | 0.01385342 | 0.13614803 |
| ENSGALG00010011465 | PPP2R2C | 422858 | 0.26073748 | 4.75890871 | 0.01390202 | 0.13654111 |
| ENSGALG00010021931 | ITGA6 | 396226 | -0.1977165 | 7.95762254 | 0.01395125 | 0.13693978 |
| ENSGALG00010006178 | RPS23 | 427323 | -0.1902577 | 8.34012014 | 0.0139621 | 0.13696156 |
| ENSGALG00010018492 | TMEM121L | 421253 | -0.4377175 | 2.88467052 | 0.01399603 | 0.13720954 |
| ENSGALG00010022633 | NA | NA | -0.9347076 | -0.0283075 | 0.0140331 | 0.13748797 |

| | | | | | | |
|---|---|---|---|---|---|---|
| ENSGALG00010019689 | NA | NA | 0.52316769 | 2.19572217 | 0.01406703 | 0.13773532 |
| ENSGALG00010021574 | TTLL4 | 424209 | 0.20495972 | 6.20594039 | 0.01408397 | 0.13781565 |
| ENSGALG00010013005 | SLC40A1 | 423984 | 0.35636058 | 3.36936683 | 0.01409261 | 0.13781565 |
| ENSGALG00010001381 | MDFIC | 417774 | 0.27938573 | 4.77973359 | 0.0141098 | 0.13789869 |
| ENSGALG00010029206 | CACNG4 | 373924 | -0.4723194 | 2.6559176 | 0.01412762 | 0.13798785 |
| ENSGALG00010012477 | LOC430496 | 430496 | -0.7445548 | 0.88332924 | 0.01414535 | 0.13805322 |
| ENSGALG00010023070 | SIDT2 | 770233 | -0.295865 | 4.21990016 | 0.01415172 | 0.13805322 |
| ENSGALG00010028357 | SPPL2C | 420056 | 0.20464573 | 7.49150643 | 0.01418236 | 0.13818839 |
| ENSGALG00010000925 | NA | NA | -0.3299104 | 3.58644267 | 0.014183 | 0.13818839 |
| ENSGALG00010023254 | MYO3B | 424157 | 0.70969036 | 1.05899132 | 0.01423508 | 0.13861067 |
| ENSGALG00010014171 | DLC1 | 422740 | 0.42451122 | 2.75146426 | 0.01426696 | 0.13881331 |
| ENSGALG00010015410 | HTR2B | 395581 | 0.85608974 | 0.36061076 | 0.01427339 | 0.13881331 |
| ENSGALG00010010626 | ADTRP | 771464 | 0.92745123 | 0.73937837 | 0.01437338 | 0.13962177 |
| ENSGALG00010014769 | CCDC69 | 416268 | -1.122871 | 0.1127672 | 0.01437412 | 0.13962177 |
| ENSGALG00010027708 | LOC101750889 | 101750889 | 0.65806466 | 1.49422848 | 0.01438398 | 0.13963203 |
| ENSGALG00010012722 | ZSWIM6 | 770670 | 0.21753157 | 5.05993437 | 0.0143993 | 0.13969518 |
| ENSGALG00010028440 | PSMB7 | 378915 | -0.1769779 | 7.55842833 | 0.01449527 | 0.14034956 |
| ENSGALG00010018665 | IL13RA2 | 422219 | 0.71829267 | 1.16959968 | 0.01449963 | 0.14034956 |
| ENSGALG00010015549 | LOC101750367 | 101750367 | 0.82725762 | 0.61786379 | 0.01450672 | 0.14034956 |
| ENSGALG00010021703 | CD82 | 423172 | -0.3065372 | 5.58974733 | 0.0145273 | 0.14034956 |
| ENSGALG00010018458 | SEMA5B | 424248 | 0.19049089 | 5.83818793 | 0.01452752 | 0.14034956 |
| ENSGALG00010021489 | CNPPD1 | 424204 | -0.2237097 | 5.74776633 | 0.01454217 | 0.14034956 |
| ENSGALG00010015837 | GDA | 427253 | 0.35129301 | 3.6333592 | 0.01454698 | 0.14034956 |
| ENSGALG00010003783 | NA | NA | 0.94287638 | 0.9457418 | 0.01454701 | 0.14034956 |
| ENSGALG00010011388 | GPR50 | 396318 | 0.4997198 | 2.06695861 | 0.01455314 | 0.14034956 |
| ENSGALG00010001906 | SKAP2 | 420629 | -0.2238336 | 5.56989992 | 0.01455523 | 0.14034956 |
| ENSGALG00010017783 | STAR | 395421 | -0.7133429 | 1.05078541 | 0.01456437 | 0.14035236 |
| ENSGALG00010024930 | TMEM237 | 424084 | 0.23614366 | 6.62738619 | 0.01464166 | 0.14101158 |
| ENSGALG00010023195 | COL24A1 | 424526 | -0.527441 | 4.46825183 | 0.0146599 | 0.14110158 |
| ENSGALG00010024467 | RPESL | 771089 | -0.5561844 | 1.52746757 | 0.01468283 | 0.14123655 |
| ENSGALG00010021773 | LOC423822 | 423822 | 0.51038269 | 3.23398647 | 0.01471606 | 0.14140652 |
| ENSGALG00010001654 | RPS11 | 419049 | -0.2133521 | 8.81219532 | 0.01471833 | 0.14140652 |
| ENSGALG00010019391 | GANC | 423232 | -0.322326 | 4.03594919 | 0.01473538 | 0.14148457 |
| ENSGALG00010007230 | NA | NA | -0.5320149 | 1.57600997 | 0.01479079 | 0.14193078 |
| ENSGALG00010026945 | NA | NA | 0.70331767 | 0.92564571 | 0.01482964 | 0.14221751 |
| ENSGALG00010015318 | SALL4 | 769286 | 0.19440897 | 9.36939168 | 0.01489564 | 0.14266367 |
| ENSGALG00010013209 | KIAA1549 | 100857885 | 0.18645469 | 6.72096123 | 0.01489994 | 0.14266367 |
| ENSGALG00010016411 | CTSH | 770109 | -0.3103009 | 4.32155331 | 0.01490315 | 0.14266367 |
| ENSGALG00010003719 | RPGR | 418578 | 0.22966416 | 5.95832834 | 0.0149234 | 0.14276005 |
| ENSGALG00010015039 | FAM114A1 | 426196 | -0.6453004 | 3.26996947 | 0.01493121 | 0.14276005 |
| ENSGALG00010016008 | TAF7 | 422192 | -0.2204638 | 5.36690731 | 0.01494759 | 0.14283049 |
| ENSGALG00010019307 | ATF3 | 421369 | -0.5478693 | 4.6322264 | 0.01498046 | 0.142997 |
| ENSGALG00010018615 | HTR2C | 428707 | 0.40467107 | 2.95642539 | 0.01498304 | 0.142997 |
| ENSGALG00010016927 | MGAT4B | 416285 | 0.21882499 | 5.75153973 | 0.0150316 | 0.14337416 |
| ENSGALG00010022250 | SLC25A28 | 423832 | -0.2387732 | 5.4136413 | 0.01504251 | 0.14339201 |
| ENSGALG00010002462 | RRM2 | 421936 | 0.24400477 | 7.07360516 | 0.01510138 | 0.14386669 |
| ENSGALG00010026988 | NA | NA | 0.54545935 | 1.98467232 | 0.01513516 | 0.14410199 |
| ENSGALG00010025938 | ERP29 | 416882 | -0.2290236 | 7.10896562 | 0.01521248 | 0.14474468 |
| ENSGALG00010020346 | PCBD1 | 395729 | -0.2670535 | 4.87270348 | 0.01522564 | 0.14474468 |
| ENSGALG00010016549 | PTPRA | 396060 | 0.17644019 | 7.03279791 | 0.01523004 | 0.14474468 |
| ENSGALG00010018317 | ZMAT3 | 425428 | 0.31175718 | 3.74261574 | 0.01528266 | 0.14515781 |
| ENSGALG00010017049 | TRAF3 | 423471 | 0.36296085 | 3.94329738 | 0.01531466 | 0.14537468 |
| ENSGALG00010027034 | NA | NA | 0.70250808 | 1.0117842 | 0.01532593 | 0.14537573 |
| ENSGALG00010000038 | NA | NA | -0.9899095 | 0.96286268 | 0.0153331 | 0.14537573 |
| ENSGALG00010017765 | SYT16 | 423516 | 0.28618919 | 4.15088767 | 0.01539274 | 0.14572408 |
| ENSGALG00010004671 | NA | NA | 0.29749519 | 4.14665609 | 0.0153941 | 0.14572408 |
| ENSGALG00010010475 | CCDC71L | 101750427 | 0.32228451 | 3.54348776 | 0.0153974 | 0.14572408 |
| ENSGALG00010028008 | NTMT1 | 425539 | -0.2059206 | 5.18468844 | 0.01547804 | 0.14638965 |
| ENSGALG00010008006 | CPNE7 | 415851 | -0.7462047 | 0.59256295 | 0.01548618 | 0.14638965 |
| ENSGALG00010021769 | TCHP | 425209 | -0.3248395 | 3.91349423 | 0.01551728 | 0.14659624 |

| | | | | | | |
|---|---|---|---|---|---|---|
| ENSGALG00010017378 | LOC422926 | 422926 | 0.58837102 | 2.19436918 | 0.01553416 | 0.14666832 |
| ENSGALG00010016491 | DNAJC8 | 419569 | -0.1720422 | 7.07343375 | 0.01554726 | 0.14670474 |
| ENSGALG00010003753 | SLCO5A1 | 420174 | 0.26346386 | 4.69200625 | 0.01556439 | 0.14673317 |
| ENSGALG00010013359 | BTG1 | 396295 | -0.3321918 | 6.79544774 | 0.01557428 | 0.14673317 |
| ENSGALG00010021817 | EVI5 | 424500 | -0.2409007 | 5.30303415 | 0.01557803 | 0.14673317 |
| ENSGALG00010023247 | LOC121106503 | 121106503 | 0.22716113 | 5.67243315 | 0.01559969 | 0.14684999 |
| ENSGALG00010001901 | PRPS2 | 418639 | 0.75866187 | 0.96643019 | 0.01569583 | 0.14766073 |
| ENSGALG00010027380 | GPX1 | 100857115 | -0.2281776 | 6.53677409 | 0.01571227 | 0.14766073 |
| ENSGALG00010027292 | KDM4B | 428354 | -0.1695509 | 6.52812759 | 0.01571374 | 0.14766073 |
| ENSGALG00010013681 | NA | NA | -0.5764327 | 1.90396359 | 0.01573362 | 0.14776001 |
| ENSGALG00010028548 | MYO1F | 396187 | 0.96315944 | 0.31431594 | 0.01583775 | 0.14857228 |
| ENSGALG00010022157 | EPS8L2 | 770081 | 0.39239934 | 3.15738374 | 0.01584267 | 0.14857228 |
| ENSGALG00010018773 | NA | NA | 0.79851618 | 0.53217725 | 0.01584821 | 0.14857228 |
| ENSGALG00010006803 | COX4I1 | 415826 | -0.1978152 | 7.18622776 | 0.01588551 | 0.14883397 |
| ENSGALG00010006277 | GNG10 | 769603 | -0.2224757 | 8.2425616 | 0.01599088 | 0.1497328 |
| ENSGALG00010029594 | CRLF3 | 417407 | 0.18261663 | 6.84893098 | 0.01600138 | 0.14974272 |
| ENSGALG00010006101 | INO80B | 107049493 | -0.2542512 | 4.65369038 | 0.01608166 | 0.15038433 |
| ENSGALG00010026099 | NA | NA | 0.93633905 | 0.25706846 | 0.01609325 | 0.15038433 |
| ENSGALG00010020179 | PSMD6 | 416079 | -0.1814142 | 6.75546626 | 0.01609839 | 0.15038433 |
| ENSGALG00010008453 | AMOTL1 | 428091 | 0.2762665 | 4.10907399 | 0.01615962 | 0.15076768 |
| ENSGALG00010021092 | VPS13D | 419481 | 0.16899314 | 7.06536665 | 0.01615986 | 0.15076768 |
| ENSGALG00010010908 | STK17B | 424049 | 0.21015358 | 5.14946131 | 0.01616794 | 0.15076768 |
| ENSGALG00010002792 | LOC395611 | 395611 | 0.68417157 | 1.09687248 | 0.01617854 | 0.15077786 |
| ENSGALG00010009328 | SLC22A15L | 423653 | 0.25783396 | 4.53682446 | 0.01622641 | 0.15113522 |
| ENSGALG00010027537 | SGTA | 426396 | -0.177008 | 7.29138184 | 0.01627198 | 0.15147072 |
| ENSGALG00010027402 | AP4B1 | 419879 | -0.205242 | 5.99727391 | 0.01631315 | 0.15176491 |
| ENSGALG00010021583 | NFE2L1 | 417987 | -0.2214719 | 7.8408062 | 0.0163245 | 0.15178144 |
| ENSGALG00010010770 | SLC8A2 | 107057643 | 0.86346564 | 0.22814728 | 0.01634788 | 0.15190978 |
| ENSGALG00010009313 | UNC13B | 100858799 | -0.1829464 | 8.63692427 | 0.01639634 | 0.15227089 |
| ENSGALG00010012449 | SUB1 | 427425 | -0.1916134 | 7.99114231 | 0.01641754 | 0.15237059 |
| ENSGALG00010020806 | RPP14 | 416064 | -0.2088938 | 5.30606695 | 0.01642628 | 0.15237059 |
| ENSGALG00010002868 | PPP1R9A | 420568 | 0.39045778 | 3.37028737 | 0.01645398 | 0.15240852 |
| ENSGALG00010014149 | LGALS2 | 425107 | -0.8735459 | 0.04209668 | 0.01645698 | 0.15240852 |
| ENSGALG00010027131 | UBA52 | 395958 | -0.2476902 | 7.63537289 | 0.0164592 | 0.15240852 |
| ENSGALG00010018893 | LOC107054097 | 107054097 | 0.92458832 | 0.05008187 | 0.01653293 | 0.15291432 |
| ENSGALG00010025849 | OBSCN | 420395 | 0.25458587 | 6.54145938 | 0.0165331 | 0.15291432 |
| ENSGALG00010012094 | GART-B | 427977 | 0.42811588 | 2.48997684 | 0.01657692 | 0.15323026 |
| ENSGALG00010027877 | OAZ1 | 395746 | -0.1925506 | 8.22568015 | 0.01661435 | 0.15348676 |
| ENSGALG00010010416 | SLC35B3 | 420863 | -0.2121739 | 5.11771203 | 0.01663838 | 0.15361932 |
| ENSGALG00010025026 | DENR | 771794 | -0.1853064 | 6.39568385 | 0.01664864 | 0.15362461 |
| ENSGALG00010002781 | EIF3G | 101747413 | -0.2142974 | 7.25992778 | 0.01672436 | 0.15406337 |
| ENSGALG00010013382 | ASB12 | 422313 | 0.45615619 | 2.29011741 | 0.01673097 | 0.15406337 |
| ENSGALG00010012831 | NA | NA | -0.741326 | 1.11495395 | 0.0167426 | 0.15406337 |
| ENSGALG00010016955 | LOC424998 | 424998 | 0.32529471 | 6.86992307 | 0.01674605 | 0.15406337 |
| ENSGALG00010017253 | NA | NA | 0.22293105 | 5.51542024 | 0.01674735 | 0.15406337 |
| ENSGALG00010027103 | ACVR2B | 374213 | 0.22794986 | 5.48150092 | 0.01675446 | 0.15406337 |
| ENSGALG00010013619 | IL1R2 | 418715 | 0.70969722 | 0.71577489 | 0.01679819 | 0.1543286 |
| ENSGALG00010029730 | TTC19 | 417597 | -0.3517453 | 3.13191593 | 0.01680277 | 0.1543286 |
| ENSGALG00010008264 | CNTNAP2 | 768456 | 0.35836498 | 4.75131791 | 0.01681888 | 0.15438721 |
| ENSGALG00010011116 | TAPT1 | 422824 | -0.1910413 | 5.81910318 | 0.01688287 | 0.15485437 |
| ENSGALG00010028254 | NA | NA | -0.361267 | 3.51504109 | 0.0168893 | 0.15485437 |
| ENSGALG00010006720 | SRP54 | 423322 | -0.233757 | 6.57133996 | 0.01694103 | 0.15520886 |
| ENSGALG00010012378 | RPS19BP1 | 418011 | -0.232868 | 4.85798754 | 0.01695145 | 0.15520886 |
| ENSGALG00010011917 | HSF2BP | 418548 | 0.25247624 | 5.15726822 | 0.01695732 | 0.15520886 |
| ENSGALG00010020867 | LOC112533229 | 112533229 | -1.211862 | -0.6326889 | 0.01708823 | 0.15617982 |
| ENSGALG00010016901 | MECOM | 424997 | 0.28351376 | 4.25218813 | 0.01709006 | 0.15617982 |
| ENSGALG00010001835 | MECP2 | 112530992 | 0.2291715 | 6.16674157 | 0.01709294 | 0.15617982 |
| ENSGALG00010022236 | SHF | 415339 | 0.169899 | 6.53769587 | 0.01713687 | 0.15636452 |
| ENSGALG00010000075 | NA | NA | 0.58569816 | 1.73020684 | 0.01715023 | 0.15636452 |
| ENSGALG00010008717 | NA | NA | 1.00610519 | -0.3220112 | 0.01715257 | 0.15636452 |

| | | | | | | |
|---|---|---|---|---|---|---|
| ENSGALG00010021651 | RNF113B | 419956 | -0.2618624 | 4.3817582 | 0.01715258 | 0.15636452 |
| ENSGALG00010019212 | RPL22 | 373937 | -0.1971736 | 7.0736017 | 0.01721105 | 0.15680737 |
| ENSGALG00010010511 | GABRP | 416174 | -0.2575826 | 4.48130828 | 0.01726609 | 0.15715391 |
| ENSGALG00010005913 | NFKBIA | 396093 | -0.404402 | 2.60215911 | 0.0172689 | 0.15715391 |
| ENSGALG00010009059 | MED29 | 107055391 | -0.2259516 | 4.95887562 | 0.01733714 | 0.15768451 |
| ENSGALG00010028113 | PIP5K1C | 420072 | -0.2005439 | 5.88784219 | 0.01738584 | 0.15800371 |
| ENSGALG00010003245 | NANOS3 | 112530983 | 0.26893349 | 5.7701193 | 0.01739216 | 0.15800371 |
| ENSGALG00010000284 | NA | NA | -0.3423642 | 3.59319488 | 0.0174165 | 0.15813426 |
| ENSGALG00010018226 | JAG1 | 396043 | 0.25204436 | 4.86684698 | 0.0174287 | 0.1581545 |
| ENSGALG00010015855 | NIPAL3 | 419697 | -0.1701336 | 6.80118473 | 0.0174549 | 0.15830164 |
| ENSGALG00010018137 | NDUFB5 | 424978 | -0.1994456 | 6.09048363 | 0.01749724 | 0.15859494 |
| ENSGALG00010000552 | NA | NA | 0.40548494 | 2.9746241 | 0.01751133 | 0.15863203 |
| ENSGALG00010018224 | LOC112530237 | 112530237 | -0.8432264 | 0.98744681 | 0.01758492 | 0.15918104 |
| ENSGALG00010029380 | RGS9 | 417433 | 0.29398753 | 5.16815016 | 0.01759934 | 0.15918104 |
| ENSGALG00010002796 | PRDM8 | 777386 | 0.64121344 | 1.04648414 | 0.01760204 | 0.15918104 |
| ENSGALG00010022340 | GFOD2 | 415642 | -0.2100292 | 6.91432636 | 0.01766641 | 0.15958241 |
| ENSGALG00010020602 | DENND2B | 423055 | 0.28470827 | 4.54284165 | 0.01768212 | 0.15958241 |
| ENSGALG00010002330 | KHDRBS2 | 428643 | -0.4393591 | 4.69749087 | 0.01768644 | 0.15958241 |
| ENSGALG00010024344 | YEATS2 | 424764 | 0.16907491 | 7.29059685 | 0.01769244 | 0.15958241 |
| ENSGALG00010001865 | CYBB | 418581 | 0.2410509 | 4.92573933 | 0.01770068 | 0.15958241 |
| ENSGALG00010019982 | PLCE1 | 425224 | 0.17814346 | 6.77502642 | 0.01770679 | 0.15958241 |
| ENSGALG00010000548 | NA | NA | -0.5151981 | 3.86124212 | 0.01776943 | 0.160056 |
| ENSGALG00010028104 | GOLGA1 | 417097 | -0.2192213 | 5.30800982 | 0.01785376 | 0.16072438 |
| ENSGALG00010022182 | CNTNAP1 | 428309 | 0.4509944 | 2.28462178 | 0.01787434 | 0.16081838 |
| ENSGALG00010000031 | NA | NA | -0.5632815 | 1.39948123 | 0.01791024 | 0.16105002 |
| ENSGALG00010004350 | MOXD1 | 395333 | 0.24288228 | 5.07772134 | 0.01796496 | 0.16144269 |
| ENSGALG00010021775 | CCDC36 | 101750778 | -0.5002078 | 1.73172372 | 0.01798476 | 0.16144269 |
| ENSGALG00010029404 | SRR | 771635 | -0.3553914 | 3.38555743 | 0.01798902 | 0.16144269 |
| ENSGALG00010010920 | UBN2 | 417970 | 0.18770916 | 5.60026527 | 0.01799462 | 0.16144269 |
| ENSGALG00010025750 | NA | NA | -1.1628819 | -0.5959251 | 0.01801413 | 0.16152635 |
| ENSGALG00010026553 | PLPP3 | 424666 | 0.29147514 | 4.4254152 | 0.01805738 | 0.16182268 |
| ENSGALG00010026046 | NA | NA | 0.37876333 | 3.08788209 | 0.01810569 | 0.162164 |
| ENSGALG00010001501 | CAPZA2 | 417771 | -0.1905093 | 7.00574807 | 0.01815174 | 0.16248474 |
| ENSGALG00010024804 | WDR53 | 770128 | -0.1909815 | 5.7771073 | 0.01816408 | 0.16250345 |
| ENSGALG00010024191 | DPYD | 429083 | 0.17480829 | 6.62716417 | 0.01824582 | 0.16314268 |
| ENSGALG00010013379 | ZNF593 | 419549 | -0.2818322 | 5.57423445 | 0.01834936 | 0.16397604 |
| ENSGALG00010005820 | NA | NA | -0.2603955 | 5.52485085 | 0.01836957 | 0.16406429 |
| ENSGALG00010009279 | ELMOD2 | 422449 | 0.21796172 | 5.05685579 | 0.01841238 | 0.16435405 |
| ENSGALG00010012384 | NRK | 422380 | 0.27493443 | 4.45173284 | 0.0184349 | 0.16446257 |
| ENSGALG00010022358 | NA | NA | -0.4727279 | 1.87495531 | 0.01847113 | 0.16469315 |
| ENSGALG00010017434 | FGF13 | 414831 | 0.19051131 | 6.78253342 | 0.01849353 | 0.1648002 |
| ENSGALG00010015519 | LOC101747532 | 101747532 | 0.29315558 | 3.77635529 | 0.01850397 | 0.16480062 |
| ENSGALG00010022735 | NA | NA | 0.38855509 | 2.96444535 | 0.01853886 | 0.16501872 |
| ENSGALG00010001758 | COL19A1 | 421873 | 0.29871099 | 4.05383045 | 0.01859655 | 0.16543938 |
| ENSGALG00010000005 | NA | NA | -0.3171635 | 10.3257146 | 0.01862955 | 0.16563723 |
| ENSGALG00010013549 | PCP4 | 771220 | 0.25335011 | 5.30899134 | 0.01865101 | 0.16563723 |
| ENSGALG00010025276 | LOC112533226 | 112533226 | -0.7801637 | 1.30040857 | 0.01865258 | 0.16563723 |
| ENSGALG00010009364 | NA | NA | -0.3039324 | 5.10723595 | 0.01866987 | 0.16563723 |
| ENSGALG00010023231 | ASIC1 | 426883 | -0.4607058 | 3.10691662 | 0.018671 | 0.16563723 |
| ENSGALG00010029203 | ALDOC | 395492 | -0.1952864 | 8.59443579 | 0.01872566 | 0.16602289 |
| ENSGALG00010004233 | NHSL1 | 421680 | 0.17213327 | 8.35749785 | 0.01874255 | 0.16602289 |
| ENSGALG00010016808 | ENTPD3 | 428447 | 0.79267712 | 0.53759928 | 0.01874587 | 0.16602289 |
| ENSGALG00010025274 | LOC107050147 | 107050147 | 0.37713652 | 3.69675259 | 0.01876358 | 0.166087 |
| ENSGALG00010023652 | SLC30A7 | 424464 | -0.2198387 | 5.52275847 | 0.01883613 | 0.16660416 |
| ENSGALG00010009371 | PHTF2 | 417728 | 0.28208921 | 5.5496871 | 0.01884301 | 0.16660416 |
| ENSGALG00010016200 | MICU3 | 422736 | -0.2503778 | 4.51939806 | 0.01886583 | 0.16671294 |
| ENSGALG00010027899 | PMVK | 101747962 | -0.2558558 | 5.02098887 | 0.01888302 | 0.16677198 |
| ENSGALG00010000311 | NA | NA | 0.94933126 | 0.13532492 | 0.01890861 | 0.16690502 |
| ENSGALG00010017576 | AKAP5 | 423506 | 0.27130459 | 5.66469237 | 0.01894528 | 0.16711275 |
| ENSGALG00010004929 | ACCSL | 420553 | 0.47770884 | 2.47907642 | 0.01896247 | 0.16711275 |

| | | | | | | |
|---|---|---|---|---|---|---|
| ENSGALG00010016336 | SLC24A2 | 414891 | -0.6325699 | 1.17735405 | 0.01896375 | 0.16711275 |
| ENSGALG00010021666 | CCDC47 | 419958 | -0.1985663 | 5.36745977 | 0.01899564 | 0.16729566 |
| ENSGALG00010005359 | MEIS2 | 395588 | -0.5525705 | 1.6464809 | 0.0190056 | 0.16729566 |
| ENSGALG00010018495 | NA | NA | 0.32949524 | 4.01579227 | 0.019031 | 0.16742636 |
| ENSGALG00010029223 | NME2 | 395916 | -0.1860293 | 8.74855553 | 0.01905383 | 0.1675343 |
| ENSGALG00010018360 | NA | NA | 0.38565012 | 2.91603249 | 0.01907355 | 0.16758945 |
| ENSGALG00010003945 | NA | NA | 0.97191715 | -0.1108077 | 0.01908419 | 0.16758945 |
| ENSGALG00010000653 | NSUN2 | 420938 | -0.211004 | 5.85500153 | 0.01909949 | 0.16758945 |
| ENSGALG00010018043 | PSMC6 | 423586 | -0.1746475 | 7.13252346 | 0.01910237 | 0.16758945 |
| ENSGALG00010020681 | BCL10 | 424528 | -0.1716434 | 6.26884184 | 0.01913397 | 0.16777392 |
| ENSGALG00010008119 | CCDC83 | 425232 | 0.37964347 | 2.92097116 | 0.01915709 | 0.16788388 |
| ENSGALG00010022546 | MTG1 | 791224 | -0.24596 | 4.68353833 | 0.01916989 | 0.16790326 |
| ENSGALG00010014484 | NA | NA | 0.23363653 | 5.32744106 | 0.01918689 | 0.16795941 |
| ENSGALG00010021366 | NA | NA | 0.28571473 | 4.00036807 | 0.01922128 | 0.16809774 |
| ENSGALG00010027732 | EEF2 | 396325 | -0.2165735 | 10.7105603 | 0.01922389 | 0.16809774 |
| ENSGALG00010026536 | BCR | 416952 | 0.20860477 | 5.80587676 | 0.01926697 | 0.16836694 |
| ENSGALG00010023714 | EIF3M | 421602 | -0.1688927 | 7.7005906 | 0.01927843 | 0.16836694 |
| ENSGALG00010009065 | EIF2B4 | 425800 | -0.177403 | 5.99866743 | 0.0192937 | 0.16836694 |
| ENSGALG00010005504 | LATS1 | 421627 | -0.1936416 | 6.09220716 | 0.01929713 | 0.16836694 |
| ENSGALG00010007023 | NA | NA | 0.4224427 | 3.1440564 | 0.01932704 | 0.16853523 |
| ENSGALG00010003444 | NA | NA | 0.81740442 | 0.39665226 | 0.01935811 | 0.16871339 |
| ENSGALG00010024035 | OSBPL5 | 770816 | 0.44180832 | 2.78216229 | 0.01937401 | 0.16875926 |
| ENSGALG00010001404 | NUP42L4 | 420615 | -0.6829628 | 1.89673792 | 0.01943308 | 0.16918083 |
| ENSGALG00010015551 | VWA3B | 418688 | 0.43094117 | 2.68413824 | 0.01946125 | 0.16933313 |
| ENSGALG00010018923 | VPS54 | 421280 | -0.2063611 | 6.15742599 | 0.01947207 | 0.16933439 |
| ENSGALG00010029966 | RABGEF1 | 417548 | 0.17665877 | 6.20619665 | 0.01959724 | 0.17027756 |
| ENSGALG00010010075 | NA | NA | 0.34048293 | 3.22170097 | 0.01961012 | 0.17027756 |
| ENSGALG00010028001 | SBNO2 | 420104 | -0.2548258 | 4.86346229 | 0.01961273 | 0.17027756 |
| ENSGALG00010004610 | TGS1 | 421126 | 0.2291118 | 6.88362953 | 0.0196391 | 0.17041324 |
| ENSGALG00010008313 | PSMD7 | 415869 | -0.1750445 | 6.87216641 | 0.0196685 | 0.17057499 |
| ENSGALG00010010868 | SLC39A10 | 429005 | 0.18759884 | 6.13979389 | 0.01981478 | 0.17174972 |
| ENSGALG00010018967 | LOC112533095 | 112533095 | -0.6420252 | 1.11707911 | 0.01986878 | 0.17212374 |
| ENSGALG00010012778 | TBC1D15 | 417857 | -0.2336071 | 6.13526096 | 0.01991753 | 0.17245188 |
| ENSGALG00010002605 | EOMES | 428443 | 0.20008375 | 7.68187327 | 0.01996079 | 0.17273215 |
| ENSGALG00010013305 | THOC2 | 422361 | -0.1688156 | 7.79282984 | 0.01998338 | 0.17275146 |
| ENSGALG00010015752 | NA | NA | -0.2904276 | 6.38265642 | 0.0199848 | 0.17275146 |
| ENSGALG00010024802 | CD59 | 423148 | -0.349628 | 3.90487139 | 0.02000685 | 0.17282443 |
| ENSGALG00010028167 | NA | NA | -0.2215643 | 6.67072191 | 0.02001589 | 0.17282443 |
| ENSGALG00010028170 | COPE | 420117 | -0.1847155 | 7.56428199 | 0.02002593 | 0.17282443 |
| ENSGALG00010015029 | TFRC | 396191 | 0.17375566 | 7.76900996 | 0.02013376 | 0.17366051 |
| ENSGALG00010002331 | C1HXORF59 | 418583 | 0.2992693 | 4.34235609 | 0.0201767 | 0.17387795 |
| ENSGALG00010024800 | KMT5A | 416825 | -0.2190539 | 4.90583786 | 0.02018089 | 0.17387795 |
| ENSGALG00010014477 | ITPR2 | 418212 | -0.2682978 | 6.63171406 | 0.02021487 | 0.17407336 |
| ENSGALG00010022290 | NEXN | 424552 | 0.35873499 | 3.29896533 | 0.02022552 | 0.17407336 |
| ENSGALG00010021055 | IRAG1 | 423037 | 0.98869695 | -0.0967824 | 0.02028497 | 0.17449033 |
| ENSGALG00010012709 | PPP1R8 | 419564 | -0.209227 | 6.19638621 | 0.02030873 | 0.17460002 |
| ENSGALG00010021645 | ASCC1 | 423709 | -0.2416401 | 4.78357168 | 0.02038896 | 0.17513395 |
| ENSGALG00010005157 | IMPA1 | 420199 | -0.2016254 | 5.52030995 | 0.02039292 | 0.17513395 |
| ENSGALG00010017101 | CSMD2 | 419640 | 0.53239643 | 1.99888659 | 0.02043856 | 0.17543096 |
| ENSGALG00010011247 | CDYL | 420877 | -0.2230841 | 6.04645653 | 0.02045322 | 0.17546188 |
| ENSGALG00010007119 | NA | NA | 0.23177008 | 8.06758903 | 0.02049475 | 0.1757231 |
| ENSGALG00010027425 | CFD | 100857593 | -0.3771768 | 3.62908875 | 0.02053762 | 0.17590796 |
| ENSGALG00010024416 | VDR | 395988 | 0.46294408 | 2.2034424 | 0.02053849 | 0.17590796 |
| ENSGALG00010014275 | NA | NA | 0.7333515 | 0.68505128 | 0.02057691 | 0.17614193 |
| ENSGALG00010000350 | LOC100857358 | 100857358 | 0.18369692 | 8.58265238 | 0.02060247 | 0.17626554 |
| ENSGALG00010017803 | LOC422151 | 422151 | -0.2209268 | 6.83706449 | 0.02069667 | 0.17697605 |
| ENSGALG00010027395 | BCL2L15 | 772088 | 0.88990389 | 0.07618999 | 0.02071646 | 0.17700932 |
| ENSGALG00010027787 | DCP1A | 415992 | 0.23994748 | 5.09750752 | 0.02072288 | 0.17700932 |
| ENSGALG00010027391 | TPM3 | 770103 | -0.166954 | 8.52636476 | 0.02074143 | 0.17707238 |
| ENSGALG00010018946 | NA | NA | 0.48103599 | 2.18712007 | 0.02075685 | 0.17710876 |

| | | | | | | |
|---|---|---|---|---|---|---|
| ENSGALG00010007133 | BLVRA | 420776 | 0.32367885 | 3.79489234 | 0.02081978 | 0.17755019 |
| ENSGALG00010020697 | NA | NA | 0.61822214 | 0.90550812 | 0.02085376 | 0.17774441 |
| ENSGALG00010028600 | PSMD4 | 100216364 | -0.1886276 | 6.52213143 | 0.02087781 | 0.17785385 |
| ENSGALG00010019097 | STC1 | 428201 | -0.8138677 | 0.15059421 | 0.02094503 | 0.17833072 |
| ENSGALG00010025900 | FAM237A | 424098 | -0.8221306 | 1.72778701 | 0.02095707 | 0.17833747 |
| ENSGALG00010024597 | DNAJB11 | 424868 | -0.2727353 | 6.77266751 | 0.02110197 | 0.17947423 |
| ENSGALG00010028298 | NA | NA | -0.6602904 | 0.99937272 | 0.02113928 | 0.17954442 |
| ENSGALG00010020246 | ADAMTS9 | 416080 | 0.17789673 | 7.68633051 | 0.02113957 | 0.17954442 |
| ENSGALG00010028861 | SURF4 | 374273 | -0.2011019 | 8.44987571 | 0.02114418 | 0.17954442 |
| ENSGALG00010029181 | MYO19 | 417649 | 0.18420314 | 5.73084927 | 0.02119455 | 0.17983558 |
| ENSGALG00010004141 | KPNA3 | 418870 | 0.17077234 | 6.24176122 | 0.02120114 | 0.17983558 |
| ENSGALG00010028377 | PDCL | 772371 | -0.1905809 | 5.49765831 | 0.02121323 | 0.17984192 |
| ENSGALG00010023414 | SUDS3 | 427713 | -0.1727265 | 6.73969415 | 0.02123607 | 0.17988857 |
| ENSGALG00010014400 | FAM65C | 428169 | 0.31373159 | 3.7903699 | 0.02124141 | 0.17988857 |
| ENSGALG00010022930 | GTF2B | 424518 | -0.2147981 | 5.05582221 | 0.02125412 | 0.17990009 |
| ENSGALG00010016608 | NA | NA | 0.70183163 | 0.61801367 | 0.02128381 | 0.17992154 |
| ENSGALG00010027567 | PARP3 | 771158 | -0.4588128 | 2.04915459 | 0.02128541 | 0.17992154 |
| ENSGALG00010024680 | BRSK1 | 423098 | 0.88310359 | 0.25854037 | 0.02129068 | 0.17992154 |
| ENSGALG00010028727 | POLR3GL | 121107555 | 0.25235904 | 4.4569441 | 0.02132775 | 0.18002726 |
| ENSGALG00010015959 | TRNAU1AP | 419597 | -0.3771391 | 3.35405739 | 0.0213465 | 0.18002726 |
| ENSGALG00010003220 | SPP1 | 395210 | 0.63065903 | 1.06017812 | 0.0213485 | 0.18002726 |
| ENSGALG00010025291 | DUS2 | 415712 | 0.23461471 | 4.66937104 | 0.02135305 | 0.18002726 |
| ENSGALG00010004386 | CFAP44 | 418359 | 0.38494649 | 2.53787962 | 0.02135994 | 0.18002726 |
| ENSGALG00010023854 | SHCBP1 | 415751 | 0.2212706 | 4.85572793 | 0.02139301 | 0.18013352 |
| ENSGALG00010025255 | NA | NA | 0.69673904 | 1.15058925 | 0.02139526 | 0.18013352 |
| ENSGALG00010021604 | SPTB | 769410 | -0.2161979 | 5.04386085 | 0.02142048 | 0.18025019 |
| ENSGALG00010021425 | PTPN9 | 415308 | -0.2072734 | 5.21660402 | 0.02145363 | 0.18043348 |
| ENSGALG00010011178 | RSPO2 | 420271 | 0.2738721 | 4.03725026 | 0.021545 | 0.18110585 |
| ENSGALG00010016981 | CCDC110 | 101750861 | 0.57489803 | 1.35581581 | 0.02155997 | 0.18113568 |
| ENSGALG00010009884 | GFOD1 | 770283 | -0.3421402 | 3.00507823 | 0.02157289 | 0.18114828 |
| ENSGALG00010012442 | CPLX1 | 427286 | 0.609884 | 1.43388312 | 0.02159046 | 0.1811999 |
| ENSGALG00010019877 | NA | NA | 0.20231679 | 5.39790915 | 0.02163023 | 0.18143769 |
| ENSGALG00010026493 | NA | NA | -0.7763092 | 0.62464827 | 0.02165002 | 0.18149268 |
| ENSGALG00010002621 | SLC25A15 | 418886 | -0.1874168 | 5.78879317 | 0.02165967 | 0.18149268 |
| ENSGALG00010018902 | NOXO1 | 416546 | 0.63694459 | 1.12070545 | 0.02167738 | 0.18154523 |
| ENSGALG00010023423 | NA | NA | 0.85096467 | 0.68191377 | 0.02170843 | 0.18165471 |
| ENSGALG00010013828 | PTN | 418125 | -0.4165106 | 3.29181809 | 0.02171336 | 0.18165471 |
| ENSGALG00010016820 | EIF2AK3 | 422930 | -0.4352427 | 5.23599748 | 0.0217519 | 0.18181111 |
| ENSGALG00010000810 | UCHL3 | 395626 | -0.1737497 | 5.94067962 | 0.02175661 | 0.18181111 |
| ENSGALG00010014145 | METTL11B | 429066 | 0.39786671 | 3.00473674 | 0.02176644 | 0.18181111 |
| ENSGALG00010023918 | MFSD14A | 429081 | -0.1623957 | 7.27147621 | 0.02183139 | 0.18223084 |
| ENSGALG00010008293 | PLA2G7 | 395816 | -0.2095413 | 5.00636558 | 0.02187493 | 0.18223084 |
| ENSGALG00010020126 | NADK | 419403 | -0.2504207 | 6.13624872 | 0.02188362 | 0.18223084 |
| ENSGALG00010007102 | CMTM3 | 414829 | 0.22807686 | 4.60484012 | 0.02188686 | 0.18223084 |
| ENSGALG00010000789 | ROPN1L | 420926 | 0.58094814 | 1.16010259 | 0.02189326 | 0.18223084 |
| ENSGALG00010005674 | NA | NA | -0.4483412 | 2.27090486 | 0.02189943 | 0.18223084 |
| ENSGALG00010019081 | PCSK6 | 395454 | -0.2552308 | 6.42530467 | 0.02194304 | 0.18223084 |
| ENSGALG00010015709 | RBMXL3 | 416302 | 0.18589688 | 8.33989521 | 0.02194783 | 0.18223084 |
| ENSGALG00010022097 | INPP5J | 416996 | -0.2521357 | 4.69151066 | 0.02195458 | 0.18223084 |
| ENSGALG00010001743 | LOC107049019 | 107049019 | 0.42689492 | 3.71701204 | 0.02195731 | 0.18223084 |
| ENSGALG00010009003 | ALG12 | 417733 | -0.1876479 | 6.3686382 | 0.02195752 | 0.18223084 |
| ENSGALG00010008867 | NA | NA | 0.63031265 | 1.1151648 | 0.02196573 | 0.18223084 |
| ENSGALG00010028460 | CRLF1 | 100858730 | 0.34500645 | 5.11748702 | 0.02196928 | 0.18223084 |
| ENSGALG00010027571 | RBM10 | 415947 | -0.1613474 | 8.09573167 | 0.02197753 | 0.18223084 |
| ENSGALG00010001215 | PRDM6 | 426800 | 0.95528498 | -0.2239968 | 0.02199219 | 0.1822571 |
| ENSGALG00010020648 | TET3 | 425829 | 0.18417601 | 5.74593499 | 0.02202231 | 0.18241142 |
| ENSGALG00010023043 | CNP | 395921 | 0.45143015 | 2.30891136 | 0.02203782 | 0.18244465 |
| ENSGALG00010012823 | NA | NA | -0.2749978 | 3.99398725 | 0.02208258 | 0.18263282 |
| ENSGALG00010013777 | CPLANE1 | 427440 | 0.16230037 | 6.66848277 | 0.02208358 | 0.18263282 |
| ENSGALG00010004902 | ANKRD29 | 428523 | 0.86266275 | 0.28323541 | 0.02211254 | 0.18277698 |

| | | | | | | |
|---|---|---|---|---|---|---|
| ENSGALG00010016504 | SNRK | 420675 | 0.22563792 | 4.74235262 | 0.02215223 | 0.18294385 |
| ENSGALG00010019641 | CAPN11 | 396240 | -0.2498215 | 7.1949977 | 0.02215579 | 0.18294385 |
| ENSGALG00010014047 | HM13 | 100859573 | -0.3135787 | 6.67774582 | 0.02219003 | 0.18313123 |
| ENSGALG00010004147 | SPRYD7 | 418871 | -0.1867013 | 5.95592606 | 0.02220405 | 0.18315167 |
| ENSGALG00010019524 | HHIPL2 | 421339 | 1.05332628 | 0.12722778 | 0.02222432 | 0.18322355 |
| ENSGALG00010017467 | SFRP1 | 395237 | -0.1649363 | 6.37565013 | 0.02224437 | 0.18329361 |
| ENSGALG00010021707 | DLX2 | 107053847 | -0.4200773 | 2.29038503 | 0.02226717 | 0.18338625 |
| ENSGALG00010001647 | ARL8BL | 769134 | -0.1941087 | 5.60163657 | 0.02230653 | 0.18361506 |
| ENSGALG00010024844 | TM4SF19 | 424855 | -0.5300671 | 1.65087962 | 0.02234181 | 0.18378526 |
| ENSGALG00010020624 | MINOS1 | 419474 | -0.1837523 | 5.82538636 | 0.02235038 | 0.18378526 |
| ENSGALG00010029217 | CEP112 | 101749632 | 0.31772969 | 4.61228729 | 0.02237264 | 0.18387293 |
| ENSGALG00010011351 | KANK1 | 395475 | 0.17154797 | 8.47853412 | 0.02238935 | 0.18391496 |
| ENSGALG00010000720 | RBM26 | 418804 | -0.1963223 | 7.15161849 | 0.02240398 | 0.18393991 |
| ENSGALG00010021535 | SUSD6 | 423259 | 0.16853006 | 7.10676065 | 0.02249115 | 0.1845601 |
| ENSGALG00010000272 | LOC112531323 | 112531323 | 0.40743148 | 2.45005396 | 0.02250991 | 0.18461853 |
| ENSGALG00010020258 | NA | NA | 0.73701691 | 0.55291889 | 0.02257535 | 0.18505951 |
| ENSGALG00010021803 | TLL2 | 107053676 | 0.35730418 | 3.97816029 | 0.02260318 | 0.18519196 |
| ENSGALG00010029766 | MRM3 | 417620 | -0.3094114 | 3.6678761 | 0.02263226 | 0.18533446 |
| ENSGALG00010007027 | NA | NA | 0.98930128 | -0.3249871 | 0.02265252 | 0.1854047 |
| ENSGALG00010013119 | RNF114 | 414881 | -0.1882453 | 6.81934063 | 0.02267297 | 0.18547635 |
| ENSGALG00010015299 | GALE | 419686 | -0.2384585 | 4.98987842 | 0.02270419 | 0.18554331 |
| ENSGALG00010009484 | RTKN2 | 423660 | -0.1644069 | 7.25714838 | 0.02270827 | 0.18554331 |
| ENSGALG00010002459 | G3BP2 | 422576 | -0.1746634 | 8.63118189 | 0.02272724 | 0.18554331 |
| ENSGALG00010025550 | NA | NA | 0.53808524 | 2.27267689 | 0.02272794 | 0.18554331 |
| ENSGALG00010021811 | NA | NA | -1.0739565 | -0.5897973 | 0.02278572 | 0.18572422 |
| ENSGALG00010023939 | NA | NA | 0.99790984 | -0.1069438 | 0.02279144 | 0.18572422 |
| ENSGALG00010014716 | NA | NA | -0.3376689 | 3.50137332 | 0.02279615 | 0.18572422 |
| ENSGALG00010019836 | MINAR1 | 415485 | 1.10667092 | -0.4541124 | 0.02279694 | 0.18572422 |
| ENSGALG00010003315 | EPGN | 497270 | 0.58503491 | 1.87882931 | 0.02283135 | 0.18590904 |
| ENSGALG00010017826 | PABPC4 | 419674 | -0.2390905 | 5.42341672 | 0.02285913 | 0.18603978 |
| ENSGALG00010003918 | HTR1E | 771973 | 0.50135629 | 1.76915259 | 0.02287302 | 0.18605737 |
| ENSGALG00010011250 | RESF1 | 418136 | -0.1905288 | 5.69389401 | 0.02288891 | 0.18609115 |
| ENSGALG00010009749 | NDRG1 | 420321 | -0.3795669 | 5.20661786 | 0.02291796 | 0.18616562 |
| ENSGALG00010023186 | MIER1 | 424702 | 0.16252734 | 6.76828066 | 0.02292154 | 0.18616562 |
| ENSGALG00010010122 | THBS4 | 396306 | 0.18530305 | 5.89233034 | 0.02294292 | 0.18624387 |
| ENSGALG00010025674 | GRB14 | 424183 | -0.5536166 | 1.51680403 | 0.02297921 | 0.18644306 |
| ENSGALG00010025426 | SMARCD3 | 430336 | 0.28823819 | 4.34364543 | 0.02300307 | 0.18648672 |
| ENSGALG00010022373 | AK5 | 424555 | 0.30295171 | 4.06263465 | 0.0230081 | 0.18648672 |
| ENSGALG00010009493 | MXD3 | 427629 | -0.2289169 | 4.72504421 | 0.02302134 | 0.18649869 |
| ENSGALG00010018085 | PGP | 416559 | -0.1847388 | 5.5801482 | 0.02305306 | 0.18666035 |
| ENSGALG00010027621 | TSPO2 | 419914 | 0.41815034 | 2.35378463 | 0.02322836 | 0.18775636 |
| ENSGALG00010027649 | FBLN2 | 427583 | 0.19123397 | 7.84286054 | 0.02323087 | 0.18775636 |
| ENSGALG00010016980 | NPHP1 | 421223 | 0.22840167 | 5.1933852 | 0.02323415 | 0.18775636 |
| ENSGALG00010029141 | C5 | 417122 | 0.31598753 | 3.93435966 | 0.02323577 | 0.18775636 |
| ENSGALG00010012976 | SLC9A2 | 418719 | 0.80541271 | 0.32752641 | 0.02326906 | 0.18784011 |
| ENSGALG00010001505 | SSR3 | 425027 | -0.2249279 | 7.08904375 | 0.02326982 | 0.18784011 |
| ENSGALG00010024721 | MUC5L | 107053414 | 0.48478365 | 2.16462584 | 0.02332796 | 0.18821372 |
| ENSGALG00010014425 | NA | NA | 0.54926039 | 1.94207741 | 0.02337084 | 0.18846376 |
| ENSGALG00010023124 | NSUN4 | 424613 | -0.2437423 | 4.33506558 | 0.02338648 | 0.1884941 |
| ENSGALG00010010934 | HARS2 | 768802 | -0.3013344 | 3.71105392 | 0.0234462 | 0.18885175 |
| ENSGALG00010006287 | ECPAS | 427336 | -0.1589556 | 7.5472559 | 0.02345467 | 0.18885175 |
| ENSGALG00010006958 | KIF26B | 100859813 | -0.2405593 | 5.6408316 | 0.02347302 | 0.18887016 |
| ENSGALG00010010970 | TPM2 | 396430 | -0.2806007 | 4.7632822 | 0.02348641 | 0.18887016 |
| ENSGALG00010017229 | NA | NA | -0.4415382 | 2.01589135 | 0.02349268 | 0.18887016 |
| ENSGALG00010017332 | FGFR1 | 396516 | -0.1985124 | 6.97510591 | 0.023511 | 0.18887041 |
| ENSGALG00010010821 | RGS8 | 426557 | 0.54172688 | 1.76672175 | 0.02351652 | 0.18887041 |
| ENSGALG00010011657 | ALDH1L2 | 418078 | -0.226488 | 6.61963365 | 0.02354147 | 0.1889751 |
| ENSGALG00010015936 | RPL36A | 100859914 | -0.2122291 | 6.81773817 | 0.02355993 | 0.18902765 |
| ENSGALG00010023631 | TMPRSS4 | 770454 | 0.73774141 | 0.39305388 | 0.02357262 | 0.18903381 |
| ENSGALG00010022256 | PSMC3 | 423182 | -0.1674942 | 7.298115 | 0.02362049 | 0.18923242 |

| | | | | | | |
|---|---|---|---|---|---|---|
| ENSGALG00010024162 | SRCIN1 | 426281 | -0.2461252 | 5.84326736 | 0.02362125 | 0.18923242 |
| ENSGALG00010027109 | C19orf44 | 420145 | -0.2239463 | 4.97930867 | 0.02368926 | 0.18968153 |
| ENSGALG00010006585 | TTC29 | 422468 | 0.62793359 | 1.19309575 | 0.02372165 | 0.18978602 |
| ENSGALG00010010349 | RREB1 | 395920 | 0.26914486 | 4.73061045 | 0.02372624 | 0.18978602 |
| ENSGALG00010000405 | NA | NA | 0.91125309 | 0.19745933 | 0.02375295 | 0.18990391 |
| ENSGALG00010003895 | THSD7A | 420585 | 0.61548625 | 1.50737469 | 0.0237951 | 0.19014403 |
| ENSGALG00010028676 | LOC112530443 | 112530443 | -0.4410938 | 4.49582746 | 0.02380696 | 0.19014403 |
| ENSGALG00010003818 | GAB2 | 419027 | 0.36316292 | 3.56798474 | 0.02384738 | 0.19037097 |
| ENSGALG00010017443 | NA | NA | 0.30059163 | 3.88113239 | 0.02386412 | 0.19040875 |
| ENSGALG00010006240 | NA | NA | -0.8427529 | 0.57970142 | 0.02390604 | 0.19064738 |
| ENSGALG00010019305 | SNRNP27 | 771295 | -0.1654069 | 6.31724536 | 0.02392777 | 0.19072475 |
| ENSGALG00010018154 | SLC16A12 | 423791 | -0.4984105 | 1.77956745 | 0.0239652 | 0.1909272 |
| ENSGALG00010001313 | EMILIN2 | 421058 | -0.2492514 | 6.19832712 | 0.02398787 | 0.19101184 |
| ENSGALG00010007224 | DHFR | 427317 | 0.18608639 | 5.84201367 | 0.02400629 | 0.19106257 |
| ENSGALG00010029718 | ABR | 771897 | -0.3057416 | 6.43250744 | 0.02405455 | 0.19135068 |
| ENSGALG00010024659 | BEND5 | 424628 | 0.19581017 | 6.73112573 | 0.02408323 | 0.19148283 |
| ENSGALG00010019256 | BBS9 | 420745 | 0.22992569 | 5.00672409 | 0.02409682 | 0.19149483 |
| ENSGALG00010028157 | PRRC2B | 417177 | -0.1624019 | 9.53995266 | 0.02412786 | 0.19159062 |
| ENSGALG00010028068 | POLE4 | 107055326 | 0.17300339 | 6.47520575 | 0.02413303 | 0.19159062 |
| ENSGALG00010029405 | POLDIP2 | 417565 | -0.162254 | 6.83455993 | 0.0241726 | 0.19180879 |
| ENSGALG00010024767 | CTTN | 396455 | -0.1606553 | 7.75042757 | 0.02420184 | 0.19191678 |
| ENSGALG00010017016 | ZIC1 | 374103 | -0.8742997 | -0.0852876 | 0.02421041 | 0.19191678 |
| ENSGALG00010016702 | WDR25 | 428908 | 0.48291691 | 1.9384998 | 0.02428984 | 0.1924503 |
| ENSGALG00010009323 | RNF150 | 100858072 | 0.22494272 | 4.89326133 | 0.02432026 | 0.19259508 |
| ENSGALG00010003744 | NA | NA | 0.44277523 | 2.80974998 | 0.02433633 | 0.19261195 |
| ENSGALG00010019183 | ANGEL2 | 421363 | -0.1913359 | 5.76498181 | 0.02434668 | 0.19261195 |
| ENSGALG00010021204 | NAGLU | 107055308 | -0.336378 | 3.76275804 | 0.0243619 | 0.19263631 |
| ENSGALG00010016240 | EIF4A2 | 395232 | -0.1633974 | 9.48337764 | 0.02439957 | 0.19283808 |
| ENSGALG00010002270 | NA | NA | 0.50144107 | 1.78988957 | 0.02445458 | 0.19312158 |
| ENSGALG00010017597 | CNOT10 | 420669 | 0.20919796 | 5.43218629 | 0.02445979 | 0.19312158 |
| ENSGALG00010029624 | RASL10B | 771592 | 0.59880749 | 1.69776723 | 0.02452498 | 0.1935334 |
| ENSGALG00010019797 | NA | NA | 1.16594088 | -0.1939741 | 0.02453636 | 0.1935334 |
| ENSGALG00010012970 | UNC5A | 426419 | 0.3293996 | 4.05604433 | 0.02460297 | 0.193926 |
| ENSGALG00010016674 | SLC10A4 | 770263 | -0.2712743 | 4.10740688 | 0.02461058 | 0.193926 |
| ENSGALG00010014328 | RASSF8 | 418210 | -0.3009401 | 4.20125499 | 0.0246708 | 0.19422566 |
| ENSGALG00010017706 | FGFRL1 | 395107 | 0.45508939 | 2.58826988 | 0.02468208 | 0.19422566 |
| ENSGALG00010001759 | C10orf88 | 427084 | -0.1706681 | 5.82608853 | 0.02468534 | 0.19422566 |
| ENSGALG00010015810 | LIPT1 | 418697 | 0.28473034 | 3.79977744 | 0.02471396 | 0.19435438 |
| ENSGALG00010026585 | NA | NA | 0.92662145 | 0.34392216 | 0.02473668 | 0.19443671 |
| ENSGALG00010002848 | TRIB2 | 378919 | -0.2283185 | 7.19703088 | 0.02479426 | 0.19478123 |
| ENSGALG00010010430 | ASTN1 | 429070 | -0.4248838 | 4.23745843 | 0.02480507 | 0.19478123 |
| ENSGALG00010011696 | ZFC3H1 | 417855 | -0.1600308 | 7.34211619 | 0.02482258 | 0.19481049 |
| ENSGALG00010021461 | SNX33 | 415309 | -0.2958635 | 4.09667389 | 0.02483913 | 0.19481049 |
| ENSGALG00010013358 | RPS16 | 417871 | -0.2363978 | 8.76679795 | 0.02485524 | 0.19481049 |
| ENSGALG00010015677 | GRHL3 | 419695 | 0.46443215 | 2.88388034 | 0.02487367 | 0.19481049 |
| ENSGALG00010001161 | B3GALNT1 | 425011 | 0.16605969 | 6.72610581 | 0.02490413 | 0.19481049 |
| ENSGALG00010017090 | NA | NA | -0.9259217 | -0.2633214 | 0.02490507 | 0.19481049 |
| ENSGALG00010017859 | EXOC6 | 423807 | -0.2223211 | 4.88181436 | 0.02490747 | 0.19481049 |
| ENSGALG00010008051 | ZNF830 | 420966 | -0.19742 | 5.46600879 | 0.0249088 | 0.19481049 |
| ENSGALG00010012704 | SMIM15 | 770642 | -0.3297251 | 3.39255845 | 0.02493511 | 0.19481049 |
| ENSGALG00010028201 | LOC112533358 | 112533358 | -0.4040665 | 3.41631732 | 0.02494033 | 0.19481049 |
| ENSGALG00010004297 | SULF1 | 426312 | -0.4644126 | 2.70558665 | 0.0249439 | 0.19481049 |
| ENSGALG00010022365 | PLCG1 | 419175 | 0.20113714 | 6.3160694 | 0.02496767 | 0.19490016 |
| ENSGALG00010025796 | NA | NA | 0.89528697 | -0.2184032 | 0.02502047 | 0.19521629 |
| ENSGALG00010029527 | NDUFAF8 | 422071 | -0.3603279 | 3.071132 | 0.02513683 | 0.19602775 |
| ENSGALG00010002899 | LOC107053745 | 107053745 | -0.987804 | -0.4293618 | 0.02518621 | 0.19631628 |
| ENSGALG00010022727 | ACP2 | 428864 | -0.1717846 | 6.02710044 | 0.02522445 | 0.1965178 |
| ENSGALG00010021391 | TUBA4AL | 100857214 | 0.67585516 | 0.83188488 | 0.02524857 | 0.19660913 |
| ENSGALG00010003986 | FNDC3A | 418863 | 0.18844038 | 5.51997141 | 0.02526998 | 0.19666628 |
| ENSGALG00010019047 | GHRL | 408185 | 0.43392298 | 2.40291697 | 0.02528071 | 0.19666628 |

| | | | | | | |
|---|---|---|---|---|---|---|
| ENSGALG00010021847 | P3H1 | 414142 | -0.2210443 | 4.95769621 | 0.02535101 | 0.19711652 |
| ENSGALG00010024190 | LOC101750690 | 101750690 | 0.24779947 | 4.78592613 | 0.02536671 | 0.19714193 |
| ENSGALG00010026898 | PNPLA2 | 431066 | 0.17627463 | 5.59221415 | 0.0254096 | 0.19737862 |
| ENSGALG00010022691 | ASIC2 | 419945 | 0.66129599 | 0.91111357 | 0.02545877 | 0.19752998 |
| ENSGALG00010020389 | ACTN1 | 373918 | 0.19386888 | 7.12471938 | 0.02546025 | 0.19752998 |
| ENSGALG00010001586 | ZBTB14 | 396070 | 0.22489116 | 5.09647107 | 0.02546645 | 0.19752998 |
| ENSGALG00010024566 | MTSS1L | 415679 | -0.202249 | 5.3774562 | 0.02553526 | 0.1979669 |
| ENSGALG00010016564 | SPEN | 419460 | 0.15543438 | 7.81977714 | 0.02564862 | 0.19874861 |
| ENSGALG00010014806 | MAP7D1 | 419624 | -0.1835466 | 6.17597915 | 0.02568542 | 0.19893655 |
| ENSGALG00010017565 | ANKFN1L | 416752 | 0.32564936 | 3.11641203 | 0.02571652 | 0.19906888 |
| ENSGALG00010026090 | USP2 | 395766 | -0.4336118 | 2.06639651 | 0.0257276 | 0.19906888 |
| ENSGALG00010017314 | CALML4 | 427515 | 0.37997363 | 2.80208096 | 0.02580981 | 0.1996076 |
| ENSGALG00010009097 | C1QTNF2 | 416156 | 0.98364393 | -0.2197131 | 0.02585898 | 0.19973831 |
| ENSGALG00010025239 | RAG2 | 423165 | 0.31036907 | 4.46011446 | 0.02585913 | 0.19973831 |
| ENSGALG00010005895 | MLLT10 | 420509 | -0.173714 | 6.54282684 | 0.02586449 | 0.19973831 |
| ENSGALG00010012900 | RBX1 | 418001 | -0.2116112 | 5.39544212 | 0.02587985 | 0.19975966 |
| ENSGALG00010011040 | CDH6 | 414842 | -0.8580169 | 0.12916364 | 0.02591571 | 0.19993917 |
| ENSGALG00010017963 | LPAR4 | 422149 | 0.40165576 | 3.38128807 | 0.02594538 | 0.20007079 |
| ENSGALG00010011653 | NR2C1 | 373913 | -0.1722594 | 6.53101191 | 0.02597325 | 0.20018839 |
| ENSGALG00010015960 | CHRNA9 | 395522 | 0.38633498 | 3.46435542 | 0.026082 | 0.20092893 |
| ENSGALG00010006627 | SF3B5 | 769794 | -0.1821245 | 6.29368375 | 0.02610294 | 0.20099261 |
| ENSGALG00010010706 | SREBF2 | 395304 | -0.1953488 | 8.26280186 | 0.02612999 | 0.20110327 |
| ENSGALG00010029332 | YWHAE | 417554 | -0.1761362 | 9.26143015 | 0.02617555 | 0.20135627 |
| ENSGALG00010003255 | CALCR | 428425 | 0.39645078 | 2.71532687 | 0.02623215 | 0.20162622 |
| ENSGALG00010009068 | AGO2 | 100858102 | 0.22028066 | 5.37988138 | 0.02623955 | 0.20162622 |
| ENSGALG00010012975 | KRR1 | 417863 | -0.1805058 | 6.14075783 | 0.02624878 | 0.20162622 |
| ENSGALG00010007090 | CEBPZ | 421474 | -0.1668636 | 6.36346277 | 0.0263936 | 0.20264051 |
| ENSGALG00010016131 | SMYD1 | 373960 | 0.98688891 | -0.3252097 | 0.02642837 | 0.20277499 |
| ENSGALG00010000048 | MAN2A2 | 415585 | -0.2139113 | 5.40910108 | 0.02643668 | 0.20277499 |
| ENSGALG00010025010 | SLC7A10 | 415774 | -0.7167674 | 0.8910137 | 0.02644979 | 0.20277748 |
| ENSGALG00010010939 | ZCCHC4 | 422806 | -0.264022 | 4.12395317 | 0.02649403 | 0.20301519 |
| ENSGALG00010017386 | NA | NA | 0.95754875 | 0.01158773 | 0.0265064 | 0.20301519 |
| ENSGALG00010021154 | CHST13 | 416032 | 0.29274821 | 3.68234252 | 0.02654068 | 0.20317964 |
| ENSGALG00010028198 | UCK1 | 771901 | -0.2022034 | 5.07168746 | 0.02657285 | 0.20327127 |
| ENSGALG00010015893 | TCF7 | 395229 | 0.23180587 | 5.20103211 | 0.02657828 | 0.20327127 |
| ENSGALG00010004154 | COPS5 | 426579 | -0.1675901 | 6.06627312 | 0.02660724 | 0.20339469 |
| ENSGALG00010025833 | NA | NA | 0.49471264 | 2.97270525 | 0.0267001 | 0.20397088 |
| ENSGALG00010013686 | UQCRQ | 416336 | -0.1684008 | 5.81204956 | 0.02671726 | 0.20397088 |
| ENSGALG00010007054 | NA | NA | 0.16953903 | 7.54405618 | 0.02672119 | 0.20397088 |
| ENSGALG00010002855 | RACK1 | 417044 | -0.2045234 | 9.79295298 | 0.0267875 | 0.20437867 |
| ENSGALG00010002801 | NA | NA | 0.69267388 | 0.73739726 | 0.0268177 | 0.20451078 |
| ENSGALG00010023966 | PEAK1 | 100857260 | 0.15941437 | 6.78162034 | 0.02684044 | 0.2045858 |
| ENSGALG00010014897 | COL8A2 | 428221 | 0.29300828 | 3.99271317 | 0.02685652 | 0.20461002 |
| ENSGALG00010023703 | PITPNB | 416912 | -0.1702821 | 6.06417323 | 0.0268931 | 0.20479041 |
| ENSGALG00010023693 | SRSF11 | 424714 | -0.1864006 | 8.01786027 | 0.02695531 | 0.20516561 |
| ENSGALG00010003160 | KIDINS220 | 421924 | 0.17221599 | 7.49169342 | 0.02711562 | 0.20628676 |
| ENSGALG00010021332 | OIT3 | 423706 | 0.30409149 | 3.730033 | 0.02718968 | 0.20675109 |
| ENSGALG00010020116 | GNA12 | 769924 | 0.1716857 | 6.7808453 | 0.02722122 | 0.20689171 |
| ENSGALG00010005355 | FDX1 | 373947 | 0.18266793 | 5.67955918 | 0.0272429 | 0.20695733 |
| ENSGALG00010020651 | LMO1 | 100858792 | 0.48260398 | 2.08732516 | 0.02726894 | 0.20705599 |
| ENSGALG00010023087 | SH3GLB1 | 424522 | -0.208769 | 7.0219069 | 0.02730614 | 0.20723926 |
| ENSGALG00010013348 | MKRN4P | 395500 | -0.1621833 | 8.57051659 | 0.02737834 | 0.20768782 |
| ENSGALG00010015101 | CAP1 | 419680 | -0.1574494 | 7.49381328 | 0.02741519 | 0.20771205 |
| ENSGALG00010015050 | HIVEP3 | 428208 | 0.26160102 | 6.70280372 | 0.02742012 | 0.20771205 |
| ENSGALG00010029523 | FSCN2 | 429198 | 0.98669551 | -0.3252085 | 0.02742774 | 0.20771205 |
| ENSGALG00010009538 | CRIM1 | 395067 | -0.3096366 | 3.90060716 | 0.02743391 | 0.20771205 |
| ENSGALG00010008214 | CCNG2 | 422512 | -0.182779 | 6.39417327 | 0.02747059 | 0.20789047 |
| ENSGALG00010025753 | SORL1 | 395998 | -0.2292919 | 6.48518129 | 0.02749598 | 0.20798343 |
| ENSGALG00010011422 | KATNAL2 | 416384 | 0.42000119 | 2.19126594 | 0.02754862 | 0.20815435 |
| ENSGALG00010018308 | ACAP3 | 419420 | 0.45698029 | 2.46660124 | 0.02754947 | 0.20815435 |

| | | | | | | |
|---|---|---|---|---|---|---|
| ENSGALG00010003476 | NA | NA | 0.84879666 | 0.3330183 | 0.02756185 | 0.20815435 |
| ENSGALG00010013400 | NA | NA | -0.3133488 | 3.89426331 | 0.02757107 | 0.20815435 |
| ENSGALG00010014767 | LRRC15 | 768899 | 0.59004729 | 1.14403185 | 0.02765551 | 0.2086315 |
| ENSGALG00010010633 | C21orf62 | 418501 | 0.94457398 | -0.2377873 | 0.02766448 | 0.2086315 |
| ENSGALG00010012718 | NA | NA | 0.56369571 | 1.37271398 | 0.02767373 | 0.2086315 |
| ENSGALG00010013163 | NA | NA | -0.4756192 | 1.65955394 | 0.02771714 | 0.20882412 |
| ENSGALG00010006998 | POLH | 395102 | -0.2323214 | 4.8206697 | 0.02772918 | 0.20882412 |
| ENSGALG00010022269 | CNTD1 | 420019 | 0.20501775 | 4.96301331 | 0.02773877 | 0.20882412 |
| ENSGALG00010006355 | PTGR1 | 427337 | 0.16901256 | 7.33612508 | 0.02775427 | 0.20884163 |
| ENSGALG00010014089 | FIG4 | 421762 | 0.19368251 | 5.28621666 | 0.02778434 | 0.20896878 |
| ENSGALG00010003090 | DMA | 417050 | -0.5122189 | 2.71953639 | 0.0278165 | 0.20902652 |
| ENSGALG00010006243 | ARHGEF38 | 428744 | 0.44408699 | 2.36065082 | 0.02781837 | 0.20902652 |
| ENSGALG00010024369 | MYOD1 | 374048 | 0.75019876 | 0.83629365 | 0.02784913 | 0.20915855 |
| ENSGALG00010026754 | NA | NA | 0.71918649 | 0.6875235 | 0.02789601 | 0.20941151 |
| ENSGALG00010022310 | DCLRE1A | 423902 | -0.1903351 | 5.31953083 | 0.02791572 | 0.20946035 |
| ENSGALG00010020603 | VPS39 | 423230 | -0.1608524 | 6.54026599 | 0.02804967 | 0.2102201 |
| ENSGALG00010021960 | CDKN1B | 374106 | -0.2161564 | 5.94622651 | 0.02805277 | 0.2102201 |
| ENSGALG00010019130 | RPS6KC1 | 421362 | 0.17345827 | 5.86622275 | 0.02805674 | 0.2102201 |
| ENSGALG00010025204 | TRAF6 | 423163 | -0.1866946 | 5.5052467 | 0.02821811 | 0.21132938 |
| ENSGALG00010024572 | CLDN18 | 429135 | 1.27708959 | -0.385125 | 0.02823537 | 0.21134171 |
| ENSGALG00010012988 | RPL18A | 417823 | -0.1671438 | 8.65849791 | 0.0282464 | 0.21134171 |
| ENSGALG00010016059 | EPDR1 | 420731 | 0.32690948 | 3.30601727 | 0.02826891 | 0.21138842 |
| ENSGALG00010029410 | VTN | 395935 | -0.2001794 | 6.01468272 | 0.0282793 | 0.21138842 |
| ENSGALG00010020023 | FOSL2 | 421416 | -0.2582004 | 4.57647578 | 0.02829312 | 0.21139211 |
| ENSGALG00010008356 | NA | NA | 0.44381934 | 2.48288052 | 0.02834046 | 0.21164613 |
| ENSGALG00010019073 | LOC101750242 | 101750242 | 0.75539162 | 0.41050432 | 0.02838214 | 0.21185762 |
| ENSGALG00010014802 | LOC121111544 | 121111544 | 0.86500302 | -0.0787263 | 0.02840746 | 0.21193171 |
| ENSGALG00010001262 | NA | NA | 0.19603521 | 5.70274937 | 0.02841878 | 0.21193171 |
| ENSGALG00010008807 | BRD1 | 417731 | 0.18195363 | 6.43010955 | 0.02843979 | 0.21198868 |
| ENSGALG00010027707 | HIGD1C | 768335 | -0.1909719 | 5.6538412 | 0.02852393 | 0.21251602 |
| ENSGALG00010019004 | ECE2 | 100857430 | -0.3288861 | 3.1686607 | 0.02856727 | 0.21273901 |
| ENSGALG00010002422 | TRAPPC12 | 421917 | -0.182546 | 5.61739568 | 0.02863471 | 0.21314114 |
| ENSGALG00010013095 | ADSL | 396540 | -0.1637668 | 6.02318017 | 0.02867078 | 0.21326244 |
| ENSGALG00010015686 | CNOT8 | 416255 | -0.158562 | 6.31872709 | 0.0286779 | 0.21326244 |
| ENSGALG00010018156 | NA | NA | -0.7476498 | 0.34255616 | 0.02870328 | 0.21335117 |
| ENSGALG00010020677 | CYR61 | 429089 | 0.61594188 | 1.4291263 | 0.02874485 | 0.21356011 |
| ENSGALG00010009029 | IGSF3 | 418451 | 0.18035063 | 6.83013696 | 0.02876502 | 0.2136035 |
| ENSGALG00010009195 | CLEC5A | 100859051 | -0.7287248 | 0.3085874 | 0.02878881 | 0.2136035 |
| ENSGALG00010004869 | NA | NA | -0.5006865 | 1.68798517 | 0.02879109 | 0.2136035 |
| ENSGALG00010003459 | NA | NA | -0.872953 | -0.0571697 | 0.02881333 | 0.21366856 |
| ENSGALG00010023698 | NA | NA | 0.27491619 | 3.84391078 | 0.02888271 | 0.21404198 |
| ENSGALG00010020063 | VSIR | 769500 | -0.4982885 | 1.6412513 | 0.02889067 | 0.21404198 |
| ENSGALG00010021513 | SDF4 | 419423 | -0.1738778 | 7.72566871 | 0.02892566 | 0.21420114 |
| ENSGALG00010019906 | ZBTB42 | 428913 | 0.2184459 | 4.81588355 | 0.02894324 | 0.21423132 |
| ENSGALG00010009439 | B3GALNT2 | 430770 | 0.21075124 | 4.92604917 | 0.02896105 | 0.21426311 |
| ENSGALG00010024187 | LOC101751887 | 101751887 | -0.7356531 | 0.52638088 | 0.02897856 | 0.2142927 |
| ENSGALG00010010599 | F2R | 427212 | 0.20918112 | 6.14157363 | 0.02899817 | 0.21429407 |
| ENSGALG00010025549 | DHCR24 | 424661 | -0.2058666 | 6.95814783 | 0.02901771 | 0.21429407 |
| ENSGALG00010023334 | RPS17 | 374053 | -0.2198217 | 8.7612197 | 0.02901993 | 0.21429407 |
| ENSGALG00010024361 | H2AFX | 100857391 | -0.6582162 | 0.69209099 | 0.02903278 | 0.21429407 |
| ENSGALG00010020544 | BMERB1 | 416594 | 0.19104938 | 6.48691278 | 0.02904644 | 0.21429516 |
| ENSGALG00010017790 | R3HCC1L | 423835 | 0.293061 | 4.01474649 | 0.02907891 | 0.214435 |
| ENSGALG00010003495 | NA | NA | 0.70141846 | 1.5693359 | 0.029152 | 0.21487405 |
| ENSGALG00010005176 | BAMBI | 428413 | -0.40893 | 2.44094716 | 0.02917738 | 0.21496124 |
| ENSGALG00010018826 | NA | NA | 0.9357574 | -0.1280129 | 0.02923791 | 0.21530718 |
| ENSGALG00010022725 | KRT222 | 420046 | 0.62988751 | 0.80166589 | 0.02928304 | 0.2155395 |
| ENSGALG00010014974 | LIG4 | 418764 | -0.6012205 | 1.63837465 | 0.02930201 | 0.21557905 |
| ENSGALG00010000060 | HYPK | 770307 | -0.1997194 | 6.14182912 | 0.02933415 | 0.21564273 |
| ENSGALG00010018901 | TCF7L2 | 395508 | -0.273444 | 5.91364477 | 0.02935047 | 0.21564273 |
| ENSGALG00010027820 | NA | NA | 0.45297487 | 2.29752289 | 0.02935145 | 0.21564273 |

| | | | | | | |
|---|---|---|---|---|---|---|
| ENSGALG00010011486 | AQP7 | 426892 | 1.08442966 | -0.6178327 | 0.0294165 | 0.21596044 |
| ENSGALG00010008111 | SYTL2 | 426394 | 0.34226702 | 3.32223552 | 0.02942192 | 0.21596044 |
| ENSGALG00010027699 | VANGL2 | 426903 | -0.1869035 | 6.7027557 | 0.0295362 | 0.21649891 |
| ENSGALG00010008688 | HBEGF | 395654 | -0.7796951 | 0.32437991 | 0.02954335 | 0.21649891 |
| ENSGALG00010023788 | RTCA | 424468 | -0.2102448 | 4.9017519 | 0.02954549 | 0.21649891 |
| ENSGALG00010017813 | DDHD2 | 426777 | -0.2334987 | 6.12926177 | 0.02955097 | 0.21649891 |
| ENSGALG00010017503 | MED9 | 416506 | -0.264438 | 5.04933344 | 0.02956353 | 0.21649891 |
| ENSGALG00010021382 | TPRG1L | 419384 | -0.1684518 | 6.40294865 | 0.02970208 | 0.21741318 |
| ENSGALG00010029227 | UTP18 | 417388 | -0.1690604 | 6.49634351 | 0.02972194 | 0.21745821 |
| ENSGALG00010004434 | TCP1 | 421586 | -0.1519722 | 7.9172744 | 0.02977003 | 0.21767588 |
| ENSGALG00010020861 | CDA | 771849 | -0.3054886 | 3.94219728 | 0.02978268 | 0.21767588 |
| ENSGALG00010000610 | MED10 | 420940 | -0.1570123 | 6.27130241 | 0.02979605 | 0.21767588 |
| ENSGALG00010023408 | ZBTB44 | 101751641 | 0.16446616 | 7.39718878 | 0.02980658 | 0.21767588 |
| ENSGALG00010008150 | TERT | 420972 | 0.26759347 | 4.15372017 | 0.02986494 | 0.21795548 |
| ENSGALG00010004591 | RRS1 | 107052739 | -0.2222204 | 6.24412099 | 0.02989187 | 0.21795548 |
| ENSGALG00010002087 | POLA1 | 418593 | 0.16011434 | 7.60112024 | 0.02989558 | 0.21795548 |
| ENSGALG00010001252 | SNCAIP | 426796 | 0.18311847 | 5.74721575 | 0.02989983 | 0.21795548 |
| ENSGALG00010023060 | ALDH1A2 | 395844 | -0.2977031 | 5.00325768 | 0.02996432 | 0.21832527 |
| ENSGALG00010029988 | NA | NA | 0.74971421 | 0.46535756 | 0.03012685 | 0.21931139 |
| ENSGALG00010021727 | MVK | 768555 | -0.1720211 | 6.14342721 | 0.03012732 | 0.21931139 |
| ENSGALG00010028252 | NA | NA | -0.2319439 | 4.34611864 | 0.0301778 | 0.21943777 |
| ENSGALG00010019277 | IGFBP5 | 424220 | -0.3606366 | 3.79409352 | 0.03021976 | 0.21943777 |
| ENSGALG00010000070 | NA | NA | 0.40829988 | 2.98679067 | 0.03023848 | 0.21943777 |
| ENSGALG00010011238 | NA | NA | -0.6472814 | 0.66982955 | 0.03024146 | 0.21943777 |
| ENSGALG00010021688 | ITPR1 | 395412 | -0.21415 | 5.8980224 | 0.0302425 | 0.21943777 |
| ENSGALG00010021892 | NDUFV1 | 430210 | -0.2555681 | 4.23231901 | 0.03024255 | 0.21943777 |
| ENSGALG00010001156 | ALDH7A1 | 426812 | -0.2444486 | 7.35303986 | 0.03024764 | 0.21943777 |
| ENSGALG00010022252 | COQ9 | 415636 | -0.2701783 | 4.28187591 | 0.03025535 | 0.21943777 |
| ENSGALG00010005635 | MOS | 428537 | 0.60375096 | 0.95868534 | 0.03032289 | 0.21973243 |
| ENSGALG00010024431 | FEZ1 | 425587 | -0.2430968 | 4.57229257 | 0.03033394 | 0.21973243 |
| ENSGALG00010026827 | FAM117A | 112530395 | 0.21274255 | 6.06652406 | 0.03033754 | 0.21973243 |
| ENSGALG00010028003 | NA | NA | -0.1612003 | 7.35450469 | 0.030377 | 0.21991783 |
| ENSGALG00010021602 | PLEKHG4 | 100859373 | 0.15254557 | 7.62519085 | 0.03051014 | 0.22078097 |
| ENSGALG00010026600 | LOC107053412 | 107053412 | -0.413963 | 2.08738303 | 0.03056807 | 0.22095198 |
| ENSGALG00010007184 | VGLL3 | 418462 | 0.41468178 | 2.55254831 | 0.03057667 | 0.22095198 |
| ENSGALG00010003821 | RUBCNL | 418854 | 0.31249318 | 3.74835536 | 0.03057855 | 0.22095198 |
| ENSGALG00010021485 | SMOC1 | 768688 | 0.23377925 | 4.7168279 | 0.03058949 | 0.22095198 |
| ENSGALG00010020574 | DTL | 421373 | 0.26219237 | 5.4005721 | 0.03067135 | 0.2214424 |
| ENSGALG00010017667 | SLC2A1 | 396130 | 0.21207376 | 5.79895769 | 0.03068712 | 0.22145544 |
| ENSGALG00010027264 | LOC420374 | 420374 | -0.1648082 | 6.11639226 | 0.03070673 | 0.22149622 |
| ENSGALG00010021414 | BUD31 | 416492 | -0.1848849 | 5.3234052 | 0.03084087 | 0.22236267 |
| ENSGALG00010008513 | NA | NA | 1.1084895 | -0.5326573 | 0.03086606 | 0.2223897 |
| ENSGALG00010002946 | GDPD5 | 419070 | 0.45845611 | 1.80260089 | 0.03087266 | 0.2223897 |
| ENSGALG00010002894 | FCHSD2 | 419068 | -0.1992329 | 6.29136458 | 0.03089168 | 0.22242574 |
| ENSGALG00010028399 | SLC35E1 | 420143 | -0.1919017 | 5.99716189 | 0.0309669 | 0.22277861 |
| ENSGALG00010000612 | NA | NA | 0.93751035 | -0.2009064 | 0.03096878 | 0.22277861 |
| ENSGALG00010012640 | PRRC1 | 427127 | -0.1929461 | 6.42701374 | 0.03105281 | 0.22327385 |
| ENSGALG00010020083 | CFAP74 | 419400 | 0.30950554 | 3.71443325 | 0.03106578 | 0.22327385 |
| ENSGALG00010002871 | LOC423967 | 423967 | -0.2521221 | 5.12830454 | 0.03114547 | 0.22374519 |
| ENSGALG00010008198 | GLRB | 422411 | 0.365317 | 2.54499748 | 0.03117941 | 0.22388764 |
| ENSGALG00010016609 | NA | NA | -0.8095728 | -0.0302036 | 0.03122702 | 0.22412805 |
| ENSGALG00010012212 | RAD54B | 395449 | 0.21685989 | 5.37630982 | 0.03127818 | 0.22431351 |
| ENSGALG00010009683 | ADAMTS19 | 415600 | -0.401991 | 3.00715309 | 0.03128114 | 0.22431351 |
| ENSGALG00010029512 | TSPAN10 | 417455 | 0.87457526 | 0.2926156 | 0.03137658 | 0.22489619 |
| ENSGALG00010009020 | ARHGAP35 | 425730 | 0.15821255 | 6.70766343 | 0.031415 | 0.22498608 |
| ENSGALG00010005899 | RASA1 | 427327 | -0.1540028 | 7.42208751 | 0.03141749 | 0.22498608 |
| ENSGALG00010006219 | CLEC14A | 121110894 | -0.5031374 | 1.67721087 | 0.03143668 | 0.22501337 |
| ENSGALG00010012890 | MFAP3 | 416261 | -0.1625276 | 6.61259317 | 0.03144967 | 0.22501337 |
| ENSGALG00010018367 | PAK5 | 416734 | -0.2548876 | 3.80113024 | 0.03148839 | 0.22518886 |
| ENSGALG00010006393 | NA | NA | -0.7260424 | 0.58355969 | 0.03151647 | 0.22519926 |

| | | | | | | | |
|---|---|---|---|---|---|---|---|
| ENSGALG00010007348 | NA | | NA | 0.70388115 | 0.50388741 | 0.03152691 | 0.22519926 |
| ENSGALG00010028620 | NA | | NA | 0.63107356 | 0.83972529 | 0.0315333 | 0.22519926 |
| ENSGALG00010029361 | ALYREF | | 769169 | -0.1618122 | 7.64824389 | 0.03154664 | 0.22519926 |
| ENSGALG00010024734 | ZPR1 | | 428256 | -0.2082829 | 5.64672232 | 0.03161733 | 0.22560235 |
| ENSGALG00010018396 | TBL1XR1 | | 426284 | 0.16143977 | 7.64139298 | 0.03167881 | 0.22583839 |
| ENSGALG00010028596 | DDA1 | | 426942 | -0.1515471 | 7.12626353 | 0.03167888 | 0.22583839 |
| ENSGALG00010018013 | LOC121107473 | | 121107473 | 0.62058164 | 1.2909827 | 0.03170734 | 0.22586828 |
| ENSGALG00010003027 | SCIN | | 420588 | 0.43097757 | 2.26398993 | 0.03171155 | 0.22586828 |
| ENSGALG00010005029 | FAT4 | | 395166 | 0.24381195 | 4.13510322 | 0.0317496 | 0.22603779 |
| ENSGALG00010021469 | CSPG4 | | 425524 | 0.64757189 | 1.23308497 | 0.03180532 | 0.22604484 |
| ENSGALG00010000718 | EEF1G | | 100526660 | -0.1685201 | 8.83981361 | 0.03181276 | 0.22604484 |
| ENSGALG00010009798 | YIPF4 | | 421464 | -0.2980212 | 5.585367 | 0.03181367 | 0.22604484 |
| ENSGALG00010008164 | TG | | 100857838 | 0.36950525 | 2.72395454 | 0.03181939 | 0.22604484 |
| ENSGALG00010017688 | SPDYA | | 101748777 | -0.2517504 | 4.16453021 | 0.03182185 | 0.22604484 |
| ENSGALG00010002673 | PQLC3 | | 421941 | 0.19222816 | 5.18075096 | 0.0318652 | 0.22625151 |
| ENSGALG00010012939 | GJA1 | | 395278 | 0.14821189 | 7.49485801 | 0.0320127 | 0.22719706 |
| ENSGALG00010024024 | AGL | | 424474 | -0.1833888 | 5.92457172 | 0.03203571 | 0.22725869 |
| ENSGALG00010017630 | NA | | NA | 0.81168537 | 0.6628812 | 0.03207126 | 0.22740915 |
| ENSGALG00010017707 | AK2 | | 428227 | -0.1558407 | 6.89305528 | 0.03209578 | 0.22748135 |
| ENSGALG00010004356 | TRAPPC8 | | 421095 | -0.1568549 | 6.57030161 | 0.03212209 | 0.22756613 |
| ENSGALG00010019323 | LRIG1 | | 427592 | 0.19409752 | 5.24676576 | 0.03221829 | 0.22814574 |
| ENSGALG00010027991 | LOC107049643 | | 107049643 | -0.2577892 | 4.28017914 | 0.03225796 | 0.22832476 |
| ENSGALG00010016852 | AHDC1 | | 112530201 | -0.2263603 | 4.64943842 | 0.03232142 | 0.22867186 |
| ENSGALG00010021686 | CCPG1 | | 427496 | -0.2637888 | 7.46788886 | 0.03234148 | 0.22871183 |
| ENSGALG00010015680 | FRG1BP | | 422726 | -0.1716168 | 6.04066617 | 0.03240736 | 0.22902041 |
| ENSGALG00010004697 | NA | | NA | 0.79778895 | 0.57639347 | 0.03242683 | 0.22902041 |
| ENSGALG00010004055 | FILIP1 | | 421860 | 0.3339401 | 2.93062705 | 0.03242844 | 0.22902041 |
| ENSGALG00010017185 | NAA30 | | 423547 | 0.1562495 | 6.33856785 | 0.03247302 | 0.2292332 |
| ENSGALG00010028078 | LAMTOR2 | | 100859842 | -0.2978794 | 4.78455353 | 0.03252524 | 0.22949968 |
| ENSGALG00010001606 | PIGA | | 418624 | 0.22177344 | 4.60879884 | 0.03258642 | 0.22973066 |
| ENSGALG00010013758 | FAM3B | | 418527 | 0.83856177 | 0.00540734 | 0.03258694 | 0.22973066 |
| ENSGALG00010004851 | SOGA3 | | 421713 | -0.2267035 | 4.48906257 | 0.03263498 | 0.22996714 |
| ENSGALG00010010270 | SSR1 | | 420871 | -0.2347582 | 9.44753348 | 0.03266251 | 0.23005892 |
| ENSGALG00010016342 | NA | | NA | 0.98431757 | -0.3959403 | 0.03269984 | 0.23021961 |
| ENSGALG00010023317 | RAPGEF3 | | 426888 | 0.34798362 | 3.79968919 | 0.03272181 | 0.23027214 |
| ENSGALG00010011740 | NA | | NA | 0.18955547 | 6.24712099 | 0.03276747 | 0.23049119 |
| ENSGALG00010012353 | PIGG | | 427290 | 0.16918558 | 6.12462238 | 0.03278953 | 0.23050415 |
| ENSGALG00010013343 | KIAA1644 | | 100857276 | 0.28991506 | 3.52858003 | 0.0328124 | 0.23050415 |
| ENSGALG00010026474 | LOC121107673 | | 121107673 | -0.8716921 | 0.18263029 | 0.03283605 | 0.23050415 |
| ENSGALG00010028512 | NDUFA13 | | 100859320 | -0.24166 | 5.3033928 | 0.03283901 | 0.23050415 |
| ENSGALG00010012571 | AGAP1 | | 424024 | 0.17257363 | 5.99599998 | 0.03285225 | 0.23050415 |
| ENSGALG00010027929 | CDK5RAP2 | | 771445 | 0.1696799 | 6.66791611 | 0.03285651 | 0.23050415 |
| ENSGALG00010017541 | SREBF1 | | 373915 | -0.2435691 | 6.34340115 | 0.0329927 | 0.23135731 |
| ENSGALG00010009618 | NA | | NA | -0.9532673 | -0.1940886 | 0.03302883 | 0.23150828 |
| ENSGALG00010026649 | LOC112530406 | | 112530406 | -0.6192168 | 0.69158993 | 0.03305701 | 0.23156676 |
| ENSGALG00010018344 | SFXN5 | | 422954 | 0.21221539 | 4.69180274 | 0.03306637 | 0.23156676 |
| ENSGALG00010006078 | SPATA13 | | 418940 | 0.2043685 | 5.4182063 | 0.03309449 | 0.23166143 |
| ENSGALG00010022599 | RNLS | | 423674 | -0.3676261 | 2.54829733 | 0.0331285 | 0.23179719 |
| ENSGALG00010019579 | NA | | NA | 0.72040909 | 0.51374659 | 0.03315935 | 0.23184492 |
| ENSGALG00010013266 | KIAA1755 | | 101750666 | 0.59763201 | 1.61807912 | 0.03317777 | 0.23184492 |
| ENSGALG00010026569 | GNAZ | | 770226 | -0.383147 | 3.07711793 | 0.03317917 | 0.23184492 |
| ENSGALG00010027731 | OLFML2A | | 417098 | -0.1984599 | 5.39367006 | 0.0332127 | 0.23197706 |
| ENSGALG00010018528 | NA | | NA | 0.42356551 | 3.10089146 | 0.03331437 | 0.23258473 |
| ENSGALG00010016378 | WDR76 | | 426208 | 0.16718872 | 6.08729626 | 0.03336274 | 0.232642 |
| ENSGALG00010003522 | THRSPB | | 404756 | -0.6848354 | 0.42731176 | 0.03336568 | 0.232642 |
| ENSGALG00010012745 | | 3-Mar | 768965 | -0.247113 | 4.33770289 | 0.03336657 | 0.232642 |
| ENSGALG00010029371 | AUTS2 | | 417481 | 0.19609071 | 6.1876632 | 0.03342281 | 0.23293173 |
| ENSGALG00010001442 | LDLRAD4 | | 421044 | 0.21443597 | 5.81603209 | 0.03347718 | 0.2332082 |
| ENSGALG00010005335 | AHSA1 | | 423378 | -0.1639991 | 6.60491805 | 0.03351033 | 0.23333666 |
| ENSGALG00010029780 | TEX14 | | 417625 | 0.16003873 | 6.5604843 | 0.03353861 | 0.23343106 |

| | | | | | | |
|---|---|---|---|---|---|---|
| ENSGALG00010025658 | PDE1B | 777528 | -0.3097928 | 4.84275686 | 0.03357484 | 0.23356962 |
| ENSGALG00010017449 | TRMT6 | 421313 | -0.1755332 | 5.46212298 | 0.03358797 | 0.23356962 |
| ENSGALG00010011366 | NA | NA | 0.30342397 | 4.00741593 | 0.03361556 | 0.23364985 |
| ENSGALG00010009511 | ZFPM1 | 415835 | 0.28115508 | 3.91679634 | 0.03362896 | 0.23364985 |
| ENSGALG00010016322 | LOC107050199 | 107050199 | 0.19467814 | 6.02423382 | 0.03367101 | 0.23383958 |
| ENSGALG00010012071 | NA | NA | -0.2070153 | 5.42151388 | 0.03370038 | 0.23394111 |
| ENSGALG00010015617 | SRSF10 | 419689 | 0.35264555 | 3.2167698 | 0.03375098 | 0.23418986 |
| ENSGALG00010015175 | MAPK9 | 395983 | 0.18988001 | 5.64009434 | 0.03378834 | 0.23423704 |
| ENSGALG00010013539 | NSFL1C | 419268 | -0.1557575 | 6.3057702 | 0.03380448 | 0.23423704 |
| ENSGALG00010026268 | GARS | 428403 | -0.2159688 | 8.16207133 | 0.03381233 | 0.23423704 |
| ENSGALG00010006135 | ZC2HC1C | 428890 | 0.36546068 | 2.42131086 | 0.03381685 | 0.23423704 |
| ENSGALG00010010308 | PACRGL | 422818 | 0.34805447 | 2.68265007 | 0.03385447 | 0.23439526 |
| ENSGALG00010029327 | CRK | 107054794 | -0.1645402 | 7.57887857 | 0.0338893 | 0.23445273 |
| ENSGALG00010027925 | MKRN2 | 374162 | -0.2011493 | 5.14887826 | 0.03389317 | 0.23445273 |
| ENSGALG00010025429 | SCN3B | 419744 | -0.4132614 | 3.75317545 | 0.03391758 | 0.23445273 |
| ENSGALG00010003812 | BG8 | 417035 | 0.29798006 | 3.86184729 | 0.03392189 | 0.23445273 |
| ENSGALG00010001561 | NA | NA | 0.72666653 | 0.5859241 | 0.03401944 | 0.23502452 |
| ENSGALG00010010499 | NR4A2 | 424320 | 0.6329344 | 1.48806329 | 0.03406126 | 0.23517349 |
| ENSGALG00010011085 | UGT1A1 | 424028 | -0.7226192 | 0.0956508 | 0.03407065 | 0.23517349 |
| ENSGALG00010027982 | ENTPD2 | 395797 | -0.3690852 | 2.55697634 | 0.03410508 | 0.23530874 |
| ENSGALG00010021086 | PSMD14 | 424189 | -0.1669042 | 7.33478676 | 0.03417191 | 0.23558018 |
| ENSGALG00010016907 | SQSTM1 | 770805 | -0.2705418 | 3.63510267 | 0.03417413 | 0.23558018 |
| ENSGALG00010029603 | RHBDL3 | 417411 | 0.27643323 | 3.90421321 | 0.03420252 | 0.23567346 |
| ENSGALG00010028388 | NA | NA | 0.8435432 | -0.0930088 | 0.03427779 | 0.23602723 |
| ENSGALG00010028455 | ENSA | 425073 | -0.1600901 | 5.92221331 | 0.03428362 | 0.23602723 |
| ENSGALG00010019659 | MRPL14 | 421443 | -0.2003492 | 4.93275907 | 0.03431829 | 0.23613632 |
| ENSGALG00010023801 | LOC101751605 | 101751605 | 0.8956945 | -0.2063798 | 0.03433892 | 0.23613632 |
| ENSGALG00010024986 | PTS | 374144 | -0.5048926 | 1.36608211 | 0.03435303 | 0.23613632 |
| ENSGALG00010030103 | C1QTNF1 | 768784 | -0.5253516 | 1.5947569 | 0.03436429 | 0.23613632 |
| ENSGALG00010022833 | HRH3 | 428147 | 0.49998492 | 1.4138976 | 0.03439069 | 0.23613632 |
| ENSGALG00010004669 | ELMOD1 | 418971 | 0.23576456 | 4.28378756 | 0.0344003 | 0.23613632 |
| ENSGALG00010021136 | WEE1 | 423045 | 0.15032808 | 8.5118488 | 0.03440367 | 0.23613632 |
| ENSGALG00010019344 | DNAJC24 | 421603 | 0.18444843 | 5.378868 | 0.03445559 | 0.23624076 |
| ENSGALG00010029866 | NA | NA | -0.8305209 | 0.20436092 | 0.03446342 | 0.23624076 |
| ENSGALG00010020388 | MYOF | 423805 | 0.24765982 | 4.19102005 | 0.03446357 | 0.23624076 |
| ENSGALG00010027861 | PNPLA7 | 427774 | 0.15735198 | 6.0013049 | 0.03447878 | 0.23624297 |
| ENSGALG00010029290 | ABCC3 | 422099 | 0.21241612 | 5.09164386 | 0.03450626 | 0.23632913 |
| ENSGALG00010029238 | CAMKK1 | 771003 | 0.34141905 | 3.66947441 | 0.03454919 | 0.2364334 |
| ENSGALG00010029684 | HTR3AL | 100859449 | -0.8084552 | -0.1655363 | 0.03455129 | 0.2364334 |
| ENSGALG00010016763 | UBALD1 | 416661 | -0.2272054 | 5.37018283 | 0.03458959 | 0.23658106 |
| ENSGALG00010029626 | AP2B1 | 417525 | 0.15270115 | 7.66403123 | 0.03461407 | 0.23658106 |
| ENSGALG00010012439 | PAPD4 | 427178 | -0.1846791 | 5.66457159 | 0.03461762 | 0.23658106 |
| ENSGALG00010019812 | NA | NA | 0.69201552 | 1.10801868 | 0.03467227 | 0.2368525 |
| ENSGALG00010014123 | NA | NA | -0.9618522 | -0.4869532 | 0.0347416 | 0.23708796 |
| ENSGALG00010017504 | NA | NA | -0.9654711 | -0.390952 | 0.03474672 | 0.23708796 |
| ENSGALG00010004520 | NA | NA | -0.6927613 | 1.35550733 | 0.03475496 | 0.23708796 |
| ENSGALG00010030046 | NA | NA | -0.4876557 | 1.53794726 | 0.03477377 | 0.23708796 |
| ENSGALG00010021178 | NDST2 | 423738 | -0.1648147 | 5.83002863 | 0.03478147 | 0.23708796 |
| ENSGALG00010014201 | NA | NA | -0.6976684 | 0.68744404 | 0.03482292 | 0.23717502 |
| ENSGALG00010006621 | DACT2 | 421561 | 0.26977941 | 4.01989788 | 0.03482415 | 0.23717502 |
| ENSGALG00010016428 | ADRA1D | 422933 | 0.25740747 | 4.00861635 | 0.03483948 | 0.23717758 |
| ENSGALG00010018266 | GNOT2 | 396117 | -0.2021482 | 5.89844506 | 0.03492077 | 0.23756455 |
| ENSGALG00010028106 | ARPC5L | 101752218 | -0.1892514 | 5.0418476 | 0.03492627 | 0.23756455 |
| ENSGALG00010021679 | MASP2 | 407089 | 0.31808838 | 3.82320915 | 0.03495677 | 0.23767009 |
| ENSGALG00010024348 | LOC101750667 | 101750667 | 0.81373947 | 0.29347674 | 0.03500522 | 0.23789752 |
| ENSGALG00010007643 | COTL1 | 768420 | -0.2366232 | 7.43012775 | 0.03508393 | 0.23833029 |
| ENSGALG00010027358 | RNF126 | 420085 | -0.1650721 | 5.61618185 | 0.03516439 | 0.23860311 |
| ENSGALG00010026088 | SNRPC | 419901 | -0.1856523 | 7.05254374 | 0.03516778 | 0.23860311 |
| ENSGALG00010022382 | CALB2 | 396255 | -0.4536413 | 1.67768503 | 0.03516922 | 0.23860311 |
| ENSGALG00010014473 | NA | NA | 0.79071888 | 0.05257024 | 0.03518518 | 0.23860936 |

| | | | | | | |
|---|---|---|---|---|---|---|
| ENSGALG00010000743 | NA | NA | -0.8074965 | -0.011534 | 0.03520304 | 0.23862845 |
| ENSGALG00010007374 | CMIP | 425774 | 0.22449498 | 5.4280523 | 0.03522649 | 0.23868543 |
| ENSGALG00010029642 | NAT9 | 422109 | -0.1917376 | 5.43898771 | 0.03529696 | 0.23906077 |
| ENSGALG00010029369 | MFSD11 | 417352 | 0.17569096 | 6.09155371 | 0.03534438 | 0.23926954 |
| ENSGALG00010011996 | LRRC70 | 107049022 | 0.41845112 | 2.15375723 | 0.03535795 | 0.23926954 |
| ENSGALG00010017289 | ADGRB2 | 426163 | 0.40431404 | 2.64539789 | 0.03540065 | 0.23945637 |
| ENSGALG00010017050 | L3HYPDH | 423531 | -0.1964165 | 5.50650292 | 0.03546428 | 0.23974582 |
| ENSGALG00010018926 | SLC22A7 | 416712 | 0.85214059 | -0.250061 | 0.03547368 | 0.23974582 |
| ENSGALG00010015092 | HSPE1 | 395948 | -0.1471061 | 7.7018895 | 0.0355604 | 0.24022959 |
| ENSGALG00010007658 | BIRC2 | 374012 | 0.15610662 | 6.75314825 | 0.03558325 | 0.24028161 |
| ENSGALG00010019172 | NEMF | 428926 | -0.1652634 | 5.62627247 | 0.03561175 | 0.24037173 |
| ENSGALG00010019046 | SELENOS | 415522 | -0.2455244 | 5.48629838 | 0.03565784 | 0.24048496 |
| ENSGALG00010001966 | ARHGAP6 | 418642 | 0.28933705 | 3.58170255 | 0.03567786 | 0.24048496 |
| ENSGALG00010029709 | SLC6A4 | 404747 | 0.67464818 | 0.47794059 | 0.03568203 | 0.24048496 |
| ENSGALG00010026917 | AMT | 395566 | -0.2051527 | 4.87260253 | 0.03569811 | 0.24048496 |
| ENSGALG00010009677 | SLC26A5 | 417715 | 0.18190767 | 6.0471019 | 0.03570433 | 0.24048496 |
| ENSGALG00010023054 | AQP9 | 415402 | -1.487369 | -0.2163302 | 0.03572726 | 0.24053726 |
| ENSGALG00010029873 | NA | NA | 0.29144507 | 3.69756299 | 0.03574791 | 0.24055675 |
| ENSGALG00010007923 | YAP1 | 396171 | 0.16180774 | 7.30416008 | 0.03576269 | 0.24055675 |
| ENSGALG00010024213 | SGO2 | 424070 | 0.39443166 | 3.40499243 | 0.03581186 | 0.24055675 |
| ENSGALG00010027118 | ABHD6 | 416009 | 0.38839313 | 2.60626034 | 0.03581359 | 0.24055675 |
| ENSGALG00010022579 | TNFAIP8L3 | 427499 | 0.41053359 | 2.07674978 | 0.03583169 | 0.24055675 |
| ENSGALG00010029388 | ST6GALNAC2 | 396159 | -0.20308 | 4.94518118 | 0.03583631 | 0.24055675 |
| ENSGALG00010022585 | CIR1 | 426529 | -0.1633586 | 7.00154239 | 0.03583631 | 0.24055675 |
| ENSGALG00010024532 | TMEM86A | 428847 | 0.36845177 | 2.68617053 | 0.03591108 | 0.24077576 |
| ENSGALG00010024885 | SLC4A1 | 396532 | 0.62803062 | 0.77689203 | 0.03591249 | 0.24077576 |
| ENSGALG00010027711 | WNK2 | 415984 | -0.1554021 | 8.1664952 | 0.03592172 | 0.24077576 |
| ENSGALG00010016340 | RGS12 | 395619 | 0.22682146 | 4.37657516 | 0.03592966 | 0.24077576 |
| ENSGALG00010016304 | EIF3J | 415573 | -0.1667623 | 6.8930634 | 0.035954 | 0.24083715 |
| ENSGALG00010006008 | NA | NA | 0.97442982 | -0.3269348 | 0.03600566 | 0.24105414 |
| ENSGALG00010011190 | LOC107050560 | 107050560 | -0.7899211 | 0.4275719 | 0.03601679 | 0.24105414 |
| ENSGALG00010029543 | NSUN5 | 417462 | -0.2039254 | 5.27388319 | 0.03606441 | 0.24127104 |
| ENSGALG00010026174 | NA | NA | 0.60196084 | 1.40301175 | 0.03611789 | 0.2415168 |
| ENSGALG00010018767 | NDUFB6 | 416391 | -0.2155717 | 4.67264355 | 0.0361316 | 0.2415168 |
| ENSGALG00010001716 | TRIL | 428436 | 0.16280997 | 6.23710637 | 0.03619019 | 0.24180655 |
| ENSGALG00010004409 | NA | NA | 0.34093324 | 3.21316491 | 0.03628487 | 0.2423371 |
| ENSGALG00010020118 | GNB1 | 419402 | -0.1433827 | 8.83457155 | 0.0363514 | 0.24267922 |
| ENSGALG00010008106 | NA | NA | 0.75763232 | 0.41859799 | 0.03640372 | 0.2428976 |
| ENSGALG00010025908 | NA | NA | 0.58792284 | 0.88465466 | 0.03641473 | 0.2428976 |
| ENSGALG00010029211 | SDF2 | 427832 | -0.2076561 | 5.0127622 | 0.03649198 | 0.24331052 |
| ENSGALG00010021831 | TSNAXIP1 | 769262 | 0.41367372 | 2.43120443 | 0.03652035 | 0.24339737 |
| ENSGALG00010001689 | ILK | 374018 | -0.1742764 | 5.51174648 | 0.03661039 | 0.24389498 |
| ENSGALG00010023330 | TSPAN1 | 424607 | -0.5076517 | 4.96309309 | 0.03665487 | 0.24400454 |
| ENSGALG00010020131 | SNRNP200 | 503574 | 0.15907395 | 8.97018148 | 0.0366576 | 0.24400454 |
| ENSGALG00010008433 | EBAG9 | 420276 | -0.169036 | 5.6751476 | 0.03670699 | 0.24423083 |
| ENSGALG00010016667 | WARS | 423458 | -0.1749938 | 6.12955471 | 0.03675967 | 0.24447874 |
| ENSGALG00010011203 | C18orf32 | 426844 | -0.202168 | 5.78280387 | 0.03678818 | 0.24453072 |
| ENSGALG00010012839 | TGFBI | 395897 | -0.2351469 | 7.27783162 | 0.03679863 | 0.24453072 |
| ENSGALG00010022897 | NA | NA | -0.1525623 | 7.09439881 | 0.03682758 | 0.24453072 |
| ENSGALG00010025216 | DPP6 | 420440 | -0.2496425 | 3.91598389 | 0.03684796 | 0.24453072 |
| ENSGALG00010024866 | TRAPPC4 | 419788 | -0.221058 | 4.89010235 | 0.03685998 | 0.24453072 |
| ENSGALG00010001521 | TBC1D5 | 420646 | 0.15706692 | 6.79820105 | 0.03686851 | 0.24453072 |
| ENSGALG00010022293 | DNAJC10 | 424115 | -0.2290928 | 7.41040069 | 0.03690406 | 0.24453072 |
| ENSGALG00010029056 | NA | NA | 0.52049417 | 1.80829461 | 0.03690505 | 0.24453072 |
| ENSGALG00010019621 | LOC421441 | 421441 | -0.2483211 | 7.55884249 | 0.03690623 | 0.24453072 |
| ENSGALG00010003921 | COLEC12 | 421061 | 0.2533527 | 4.58631675 | 0.03694186 | 0.24458287 |
| ENSGALG00010009340 | SYK | 427272 | -0.5238108 | 1.83885463 | 0.03695971 | 0.24458287 |
| ENSGALG00010014766 | F10 | 395876 | -0.2794161 | 3.42095264 | 0.03696036 | 0.24458287 |
| ENSGALG00010012097 | GART | 395315 | 0.15768245 | 7.14873349 | 0.03698484 | 0.24460403 |
| ENSGALG00010026396 | NA | NA | 0.69051134 | 0.58180981 | 0.03699439 | 0.24460403 |

| | | | | | | |
|---|---|---|---|---|---|---|
| ENSGALG00010001617 | TRIM3 | 425131 | -0.1915985 | 5.23254369 | 0.03705854 | 0.24484637 |
| ENSGALG00010015158 | PIN4 | 422134 | -0.2234435 | 4.61062886 | 0.03706192 | 0.24484637 |
| ENSGALG00010023288 | NA | NA | 0.82815013 | -0.1536417 | 0.03711832 | 0.24501151 |
| ENSGALG00010011866 | KLF8 | 101749773 | 0.24856145 | 4.23503735 | 0.0371352 | 0.24501151 |
| ENSGALG00010002763 | SEPTIN11 | 422635 | 0.21151877 | 5.49596998 | 0.03716632 | 0.24501151 |
| ENSGALG00010001402 | NUP42 | 420614 | -0.1969761 | 4.76297401 | 0.03717727 | 0.24501151 |
| ENSGALG00010025025 | NA | NA | 0.71633559 | 0.38096668 | 0.03721457 | 0.24501151 |
| ENSGALG00010016696 | DOK7 | 422875 | 0.43573117 | 2.6060756 | 0.03721762 | 0.24501151 |
| ENSGALG00010026296 | NA | NA | 0.20764565 | 4.71816106 | 0.03722809 | 0.24501151 |
| ENSGALG00010018466 | DYSF | 425353 | -0.1658241 | 7.16621387 | 0.037234 | 0.24501151 |
| ENSGALG00010008594 | NA | NA | 0.8326933 | 0.08841513 | 0.03723878 | 0.24501151 |
| ENSGALG00010024082 | NA | NA | -0.5811767 | 1.04942836 | 0.03724138 | 0.24501151 |
| ENSGALG00010023702 | COL4A4 | 424799 | -0.2582265 | 4.47159561 | 0.03730829 | 0.24531823 |
| ENSGALG00010008708 | ZC3H4 | 101748292 | 0.16441013 | 6.33641106 | 0.03733146 | 0.24531823 |
| ENSGALG00010026010 | ZSWIM5 | 424591 | -0.1846932 | 5.44398334 | 0.0373344 | 0.24531823 |
| ENSGALG00010019209 | VCPKMT | 423571 | -0.3612938 | 2.71627612 | 0.03745992 | 0.24587987 |
| ENSGALG00010010653 | NA | NA | -0.9990368 | -0.2493524 | 0.03746687 | 0.24587987 |
| ENSGALG00010013282 | SEC22B | 424377 | -0.1693266 | 7.83068422 | 0.0374745 | 0.24587987 |
| ENSGALG00010021560 | NA | NA | 0.80978509 | 0.02353008 | 0.03748188 | 0.24587987 |
| ENSGALG00010004060 | ETV1 | 395748 | 0.19317754 | 4.95192689 | 0.03751127 | 0.24588468 |
| ENSGALG00010010682 | SMIM13 | 119694942 | 0.17869574 | 5.21991308 | 0.03751361 | 0.24588468 |
| ENSGALG00010021043 | PTPRG | 374208 | 0.17478707 | 5.64406274 | 0.03756662 | 0.24613044 |
| ENSGALG00010005484 | GINM1 | 395778 | -0.4134883 | 2.71306311 | 0.03761585 | 0.24623988 |
| ENSGALG00010023599 | NA | NA | -0.5519708 | 1.43444589 | 0.03762354 | 0.24623988 |
| ENSGALG00010018841 | NA | NA | 0.82048945 | -0.0619311 | 0.0376299 | 0.24623988 |
| ENSGALG00010021352 | CFAP43 | 423879 | 0.34058042 | 3.44381958 | 0.03764805 | 0.24625706 |
| ENSGALG00010011895 | PHF21B | 427935 | 0.42776988 | 1.96034424 | 0.03766915 | 0.2462935 |
| ENSGALG00010015641 | NA | NA | 0.83133162 | -0.095371 | 0.03783142 | 0.24725259 |
| ENSGALG00010012081 | CHIC1 | 422318 | 0.21493804 | 5.17757397 | 0.03796467 | 0.24802126 |
| ENSGALG00010006413 | NA | NA | 0.95354554 | -0.357543 | 0.03802867 | 0.24833707 |
| ENSGALG00010023805 | GALNT6 | 429501 | 0.30761913 | 3.47294161 | 0.03806128 | 0.24844491 |
| ENSGALG00010015785 | FGL1A | 422728 | 0.44887082 | 1.65019726 | 0.03807651 | 0.24844491 |
| ENSGALG00010002954 | CHST7 | 107051963 | 0.86482668 | -0.1851951 | 0.03818243 | 0.24903357 |
| ENSGALG00010005468 | PPIL4 | 421625 | -0.1800044 | 5.79758495 | 0.03827995 | 0.24946973 |
| ENSGALG00010016762 | EML6 | 421215 | 0.41281603 | 2.6915636 | 0.03829436 | 0.24946973 |
| ENSGALG00010022518 | ECHS1 | 770828 | -0.2196397 | 4.53743438 | 0.03829648 | 0.24946973 |
| ENSGALG00010012702 | PTPRO | 373911 | 0.54908489 | 2.82120622 | 0.03836818 | 0.24957919 |
| ENSGALG00010027900 | REN | 431167 | 0.33415997 | 2.83564612 | 0.0383699 | 0.24957919 |
| ENSGALG00010017645 | SYNC | 419664 | -0.371925 | 2.33977966 | 0.03837369 | 0.24957919 |
| ENSGALG00010026291 | SERPING1 | 423132 | -0.208895 | 5.48881933 | 0.03837738 | 0.24957919 |
| ENSGALG00010029815 | NA | NA | 0.33503103 | 2.71387495 | 0.03839196 | 0.24957919 |
| ENSGALG00010014645 | SLC36A1 | 770250 | 0.19747108 | 5.17444276 | 0.03843919 | 0.24978386 |
| ENSGALG00010012303 | CA9 | 770004 | 0.26186628 | 4.61658846 | 0.03855101 | 0.25030812 |
| ENSGALG00010009608 | AFAP1 | 374034 | -0.3181874 | 5.75190982 | 0.03857352 | 0.25030812 |
| ENSGALG00010003852 | TCEB1 | 420187 | 0.1695485 | 6.51730986 | 0.03858558 | 0.25030812 |
| ENSGALG00010018209 | GMPPA | 769292 | -0.285156 | 5.22563895 | 0.03859869 | 0.25030812 |
| ENSGALG00010025644 | PI16 | 419818 | -0.7215689 | 0.65573407 | 0.03859877 | 0.25030812 |
| ENSGALG00010011916 | MCAT | 427934 | -0.2078408 | 5.3314287 | 0.03864963 | 0.25053552 |
| ENSGALG00010027775 | MVB12B | 417092 | 0.17705954 | 7.48744203 | 0.03869952 | 0.25073401 |
| ENSGALG00010021796 | NA | NA | 0.48020369 | 2.43811887 | 0.03871187 | 0.25073401 |
| ENSGALG00010029933 | FAM18B1 | 417324 | -0.1665465 | 8.02524913 | 0.03876102 | 0.25089241 |
| ENSGALG00010017428 | PSMA3 | 423542 | -0.1517121 | 7.23894044 | 0.03876795 | 0.25089241 |
| ENSGALG00010004085 | RB1CC1 | 421116 | 0.17188105 | 5.81464825 | 0.03879524 | 0.2509604 |
| ENSGALG00010013423 | ZNF804A | 112532789 | 0.29350417 | 4.4846692 | 0.0388101 | 0.2509604 |
| ENSGALG00010011729 | FAM129A | 424451 | 0.33960672 | 3.0753533 | 0.03886816 | 0.25112602 |
| ENSGALG00010017054 | GALM | 426268 | -0.504319 | 2.09178168 | 0.03887042 | 0.25112602 |
| ENSGALG00010007243 | BEAN1 | 415796 | -0.3982744 | 2.23606767 | 0.0388989 | 0.25112602 |
| ENSGALG00010024755 | PIGX | 424859 | -0.2793556 | 3.62206826 | 0.03889904 | 0.25112602 |
| ENSGALG00010024482 | USP3 | 415369 | 0.19537306 | 5.96726426 | 0.03891704 | 0.25113996 |
| ENSGALG00010026878 | LOC112532592 | 112532592 | -0.7505927 | 0.12662656 | 0.0390804 | 0.25189428 |

| | | | | | | |
|---|---|---|---|---|---|---|
| ENSGALG00010007017 | SRD5A2 | 772291 | 0.77495589 | 0.16894587 | 0.03909249 | 0.25189428 |
| ENSGALG00010013749 | NELL2 | 417799 | 0.16436519 | 6.65702349 | 0.03909408 | 0.25189428 |
| ENSGALG00010001140 | NMD3 | 425009 | 0.16586549 | 6.10014356 | 0.03909745 | 0.25189428 |
| ENSGALG00010027600 | KIRREL1 | 425302 | -0.3123518 | 4.61183644 | 0.03911362 | 0.25189617 |
| ENSGALG00010015803 | SYF2 | 419593 | -0.1645103 | 5.9382507 | 0.03923233 | 0.2525581 |
| ENSGALG00010005537 | GRIN3A | 769001 | -0.195763 | 4.97606557 | 0.03930472 | 0.25285509 |
| ENSGALG00010013051 | XPOT | 417824 | -0.157014 | 7.33198521 | 0.03935201 | 0.25285509 |
| ENSGALG00010026968 | NA | NA | 0.92693631 | -0.2530782 | 0.03935768 | 0.25285509 |
| ENSGALG00010025098 | ATP5B | 426673 | -0.1551813 | 9.39501975 | 0.03936092 | 0.25285509 |
| ENSGALG00010026740 | DAB1 | 374083 | 0.16224701 | 6.88630211 | 0.03936709 | 0.25285509 |
| ENSGALG00010011531 | MTDH | 101751823 | -0.2138253 | 8.18131016 | 0.03937411 | 0.25285509 |
| ENSGALG00010003862 | PCK2 | 396457 | 0.1591816 | 6.20036574 | 0.03943948 | 0.25312582 |
| ENSGALG00010000105 | LOC101752139 | 101752139 | 0.17415309 | 6.05524308 | 0.03944818 | 0.25312582 |
| ENSGALG00010027043 | IL17RD | 395186 | -0.1528119 | 8.05184043 | 0.03948813 | 0.25321826 |
| ENSGALG00010022843 | NA | NA | 0.66751015 | 1.24568839 | 0.0395136 | 0.25321826 |
| ENSGALG00010019267 | STARD5 | 770108 | -0.2271649 | 4.48909758 | 0.03953774 | 0.25321826 |
| ENSGALG00010002355 | UFM1 | 418894 | -0.2129041 | 6.45035303 | 0.03954304 | 0.25321826 |
| ENSGALG00010012102 | SON | 770637 | 0.15521585 | 8.97699479 | 0.03955237 | 0.25321826 |
| ENSGALG00010005237 | RAP2A | 769797 | -0.20742 | 7.147857 | 0.03955837 | 0.25321826 |
| ENSGALG00010021041 | GLTP | 429534 | -0.2378031 | 4.19024668 | 0.03957786 | 0.25324083 |
| ENSGALG00010014609 | NA | NA | -1.3315138 | -0.5104159 | 0.03966203 | 0.25362054 |
| ENSGALG00010028057 | APC2 | 429363 | -0.1998571 | 5.35435911 | 0.03969469 | 0.25362054 |
| ENSGALG00010010932 | GHITM | 423621 | -0.1473018 | 7.65210529 | 0.03969823 | 0.25362054 |
| ENSGALG00010019510 | NA | NA | 0.75086459 | 0.03580636 | 0.03970116 | 0.25362054 |
| ENSGALG00010005341 | UGGT2 | 418785 | 0.18774128 | 5.66852667 | 0.0397447 | 0.25379644 |
| ENSGALG00010008469 | NOCT | 404779 | -0.2002808 | 5.1183846 | 0.03979284 | 0.2539843 |
| ENSGALG00010003129 | NA | NA | -0.9116726 | -0.1482702 | 0.0398178 | 0.2539843 |
| ENSGALG00010015792 | MARK3 | 423480 | 0.18831248 | 7.86832396 | 0.03982215 | 0.2539843 |
| ENSGALG00010010698 | GPRIN2 | 395648 | 0.5608631 | 1.03890835 | 0.0399078 | 0.25442828 |
| ENSGALG00010024877 | PRPF38A | 771795 | -0.2410849 | 5.42907763 | 0.03996459 | 0.25468797 |
| ENSGALG00010011588 | COLGALT1 | 424447 | -0.2381782 | 4.30401599 | 0.0400543 | 0.25515713 |
| ENSGALG00010000204 | LOC107049147 | 107049147 | -0.1749848 | 6.81483888 | 0.04010994 | 0.25522602 |
| ENSGALG00010013635 | BCL6 | 424912 | 0.22987841 | 4.133554 | 0.04013415 | 0.25522602 |
| ENSGALG00010009583 | LYPD6B | 424310 | -0.2857615 | 4.28396334 | 0.04015975 | 0.25522602 |
| ENSGALG00010002647 | FAM168A | 419066 | 0.16403803 | 6.04595444 | 0.04016404 | 0.25522602 |
| ENSGALG00010025538 | ANO1 | 423144 | 0.56466103 | 0.91011696 | 0.0401655 | 0.25522602 |
| ENSGALG00010003187 | RHOG | 100859599 | 0.23334501 | 4.13527416 | 0.04017719 | 0.25522602 |
| ENSGALG00010002356 | WNT11 | 395562 | 0.34061432 | 2.88081607 | 0.04017774 | 0.25522602 |
| ENSGALG00010023624 | PSMA1 | 395874 | -0.1483669 | 6.31628554 | 0.04030153 | 0.25583769 |
| ENSGALG00010023758 | LOC107053670 | 107053670 | 0.51535527 | 1.66516634 | 0.04031159 | 0.25583769 |
| ENSGALG00010020709 | NA | NA | 0.71987161 | 0.49725315 | 0.04032984 | 0.25583769 |
| ENSGALG00010019572 | NFKBIE | 421438 | -0.4284568 | 1.87707452 | 0.04033855 | 0.25583769 |
| ENSGALG00010024462 | ZBTB32 | 419759 | 0.35260638 | 3.50967052 | 0.04041056 | 0.25619196 |
| ENSGALG00010027609 | RPL36 | 373936 | -0.1857278 | 6.88594475 | 0.04042858 | 0.25619979 |
| ENSGALG00010016774 | PEX14 | 771421 | -0.1587903 | 5.70194246 | 0.0404477 | 0.25619979 |
| ENSGALG00010002348 | THAP12 | 428122 | 0.19407303 | 5.36238737 | 0.04046917 | 0.25619979 |
| ENSGALG00010028577 | CERS2 | 100859441 | -0.1693071 | 6.96977737 | 0.04049534 | 0.25619979 |
| ENSGALG00010019598 | KCNAB2 | 396068 | 0.30618192 | 3.4657678 | 0.04050589 | 0.25619979 |
| ENSGALG00010003406 | C1GALT1 | 420575 | -0.1977589 | 7.93143528 | 0.04051403 | 0.25619979 |
| ENSGALG00010020155 | MFGE8 | 415494 | -0.3231362 | 4.79595396 | 0.04052486 | 0.25619979 |
| ENSGALG00010000120 | NA | NA | 0.33481571 | 2.88191171 | 0.04057001 | 0.25635268 |
| ENSGALG00010011720 | IKZF2 | 395309 | 0.41473712 | 2.7094461 | 0.04058136 | 0.25635268 |
| ENSGALG00010021468 | LOC107052934 | 107052934 | 1.03343531 | -0.5107192 | 0.04063782 | 0.25658211 |
| ENSGALG00010007802 | LOC770839 | 770839 | 0.62429561 | 1.7981292 | 0.04065003 | 0.25658211 |
| ENSGALG00010011689 | NA | NA | 0.72706058 | 0.74720729 | 0.04069383 | 0.25672921 |
| ENSGALG00010028574 | ENDOG | 417204 | -0.3352092 | 3.14138121 | 0.040709 | 0.25672921 |
| ENSGALG00010000717 | NDFIP2 | 418803 | 0.15131699 | 7.73563934 | 0.04072189 | 0.25672921 |
| ENSGALG00010006301 | NA | NA | -0.421743 | 1.75044425 | 0.04074508 | 0.25677335 |
| ENSGALG00010001180 | SPIRE1 | 421033 | 0.24406652 | 4.40552327 | 0.04076529 | 0.25679868 |
| ENSGALG00010004039 | ALPK1 | 100858173 | 0.77626683 | 0.20701625 | 0.04081545 | 0.25701254 |

| | | | | | | |
|---|---|---|---|---|---|---|
| ENSGALG00010007056 | NA | NA | 0.77647548 | 0.21560813 | 0.04087146 | 0.25726311 |
| ENSGALG00010028175 | POMT1 | 417175 | 0.22903187 | 4.98030107 | 0.04090386 | 0.2572815 |
| ENSGALG00010015913 | CHST10 | 395206 | 0.21027115 | 4.50025417 | 0.04090974 | 0.2572815 |
| ENSGALG00010008330 | EZH2 | 420784 | 0.18005119 | 6.89606262 | 0.04092304 | 0.2572815 |
| ENSGALG00010020053 | LFNG | 395790 | -0.3584415 | 2.87662119 | 0.04105336 | 0.25799858 |
| ENSGALG00010026264 | FZD7 | 374060 | 0.22420478 | 4.77329898 | 0.04109667 | 0.25816844 |
| ENSGALG00010016676 | NLRC3 | 416673 | 0.5213308 | 1.98918398 | 0.04112931 | 0.25827121 |
| ENSGALG00010029925 | GGNBP2 | 425053 | -0.1808799 | 5.22825848 | 0.04119754 | 0.25859733 |
| ENSGALG00010023712 | ARHGAP40 | 419168 | 0.45448874 | 1.69824135 | 0.04126223 | 0.25890092 |
| ENSGALG00010019830 | JAG2 | 776138 | 0.24954458 | 4.17842981 | 0.04128634 | 0.25894975 |
| ENSGALG00010018032 | ERO1A | 423585 | -0.2076642 | 4.85678636 | 0.04143591 | 0.25969752 |
| ENSGALG00010021054 | FEZF2 | 416074 | -0.3891545 | 3.35524967 | 0.04143831 | 0.25969752 |
| ENSGALG00010017498 | NA | NA | -0.4609271 | 2.01928773 | 0.04146033 | 0.25973294 |
| ENSGALG00010028517 | RNF208 | 417279 | -0.2621358 | 5.62371395 | 0.04152691 | 0.25989744 |
| ENSGALG00010000440 | C14ORF166B | 427063 | 0.45972013 | 2.06264136 | 0.04153515 | 0.25989744 |
| ENSGALG00010011390 | LOC100857497 | 100857497 | 0.31904772 | 3.30569931 | 0.04154899 | 0.25989744 |
| ENSGALG00010029673 | TMEM248 | 417549 | -0.1825642 | 8.42898555 | 0.0415541 | 0.25989744 |
| ENSGALG00010011972 | IPO11 | 769679 | 0.14493354 | 7.35177649 | 0.04158193 | 0.25989744 |
| ENSGALG00010000255 | NA | NA | -0.1556161 | 7.2283917 | 0.0415849 | 0.25989744 |
| ENSGALG00010011617 | MAMLD1 | 422389 | 0.22398141 | 4.70485547 | 0.04163134 | 0.26008519 |
| ENSGALG00010005904 | NA | NA | 0.50029626 | 1.41081945 | 0.04167764 | 0.26023706 |
| ENSGALG00010021999 | ANKRD22 | 771419 | 0.64236419 | 0.51852961 | 0.04168846 | 0.26023706 |
| ENSGALG00010018721 | LRCH2 | 422220 | 0.20859937 | 5.64717266 | 0.04171634 | 0.26030867 |
| ENSGALG00010029738 | TRPV2 | 417603 | 0.45401132 | 2.16053972 | 0.04174845 | 0.26040657 |
| ENSGALG00010001508 | CD109 | 421862 | 0.44855804 | 1.6464918 | 0.04183189 | 0.26062699 |
| ENSGALG00010000382 | NA | NA | 0.2375291 | 4.88097738 | 0.04184138 | 0.26062699 |
| ENSGALG00010016374 | FRMD5 | 415576 | 0.31345788 | 3.19070129 | 0.04184795 | 0.26062699 |
| ENSGALG00010015673 | LOC112532082 | 112532082 | 0.3213936 | 2.93934059 | 0.04184951 | 0.26062699 |
| ENSGALG00010016519 | MEX3B | 107054196 | 0.16335292 | 6.37670019 | 0.04190174 | 0.26084982 |
| ENSGALG00010017390 | DDX24 | 423429 | -0.1422457 | 7.39931878 | 0.04194571 | 0.26094297 |
| ENSGALG00010014613 | NA | NA | 0.72785611 | 0.60950868 | 0.0419496 | 0.26094297 |
| ENSGALG00010000581 | LOC425562 | 425562 | -0.1788925 | 6.36789888 | 0.04200277 | 0.26117128 |
| ENSGALG00010005217 | CCNA2 | 396172 | 0.14805339 | 8.13454156 | 0.04204529 | 0.26133321 |
| ENSGALG00010015274 | NA | NA | -0.5058149 | 1.52165909 | 0.04208197 | 0.26145873 |
| ENSGALG00010027265 | IP6K3 | 419908 | -0.6816647 | 0.34478821 | 0.04210102 | 0.26145873 |
| ENSGALG00010018373 | NA | NA | 0.70147378 | 0.61833643 | 0.04213266 | 0.26145873 |
| ENSGALG00010021714 | TSPAN18 | 395586 | 0.35962681 | 3.12269022 | 0.04214695 | 0.26145873 |
| ENSGALG00010012662 | SLC35E3 | 417842 | -0.1470615 | 7.70125307 | 0.0421479 | 0.26145873 |
| ENSGALG00010021203 | FUT11 | 395071 | 0.25662908 | 4.01690308 | 0.04218163 | 0.26156569 |
| ENSGALG00010027597 | EXOSC2 | 417182 | -0.1565467 | 6.27253554 | 0.04219831 | 0.26156688 |
| ENSGALG00010017927 | PCSK2 | 395136 | -0.2531601 | 4.32517714 | 0.04226819 | 0.26180951 |
| ENSGALG00010022347 | SEC11A | 415334 | -0.1450245 | 6.99499187 | 0.04227194 | 0.26180951 |
| ENSGALG00010007781 | MBP | 396217 | -0.2777629 | 4.40918374 | 0.04228697 | 0.26180951 |
| ENSGALG00010008448 | TNS3 | 420792 | 0.19470714 | 4.93137239 | 0.04231419 | 0.26187579 |
| ENSGALG00010020005 | EOGT | 426961 | 0.18039827 | 6.01200535 | 0.04235888 | 0.26205013 |
| ENSGALG00010004365 | TPD52 | 770339 | -0.1462601 | 8.40628653 | 0.04240708 | 0.26215983 |
| ENSGALG00010018340 | DNPEP | 424200 | -0.1679987 | 5.7743163 | 0.04241051 | 0.26215983 |
| ENSGALG00010023083 | HARBI1 | 423193 | -0.3163519 | 3.92913682 | 0.04245097 | 0.26215983 |
| ENSGALG00010001482 | TRA2A | 420618 | -0.1484988 | 7.88408047 | 0.04246351 | 0.26215983 |
| ENSGALG00010007653 | NA | NA | 0.25814315 | 3.8758773 | 0.04247106 | 0.26215983 |
| ENSGALG00010004689 | BBX | 418407 | 0.19398876 | 5.20667292 | 0.04247578 | 0.26215983 |
| ENSGALG00010008258 | CCDC82 | 426136 | 0.4521216 | 1.74550339 | 0.04256957 | 0.26262002 |
| ENSGALG00010029245 | SKA2P1 | 417629 | -0.2354399 | 4.14231353 | 0.04259839 | 0.26262002 |
| ENSGALG00010029286 | MAFG | 769355 | -0.2202422 | 5.78005303 | 0.0426083 | 0.26262002 |
| ENSGALG00010010841 | PMM1 | 417989 | -0.1683647 | 5.53623357 | 0.04262216 | 0.26262002 |
| ENSGALG00010009256 | CREBRF | 416206 | 0.22604396 | 5.53415901 | 0.04264054 | 0.26262002 |
| ENSGALG00010029639 | CCLI7 | 417535 | 0.948164 | -0.3443054 | 0.04265817 | 0.26262002 |
| ENSGALG00010002917 | PGLS | 430442 | -0.1596832 | 6.82369237 | 0.04266623 | 0.26262002 |
| ENSGALG00010003466 | TSC22D1 | 396005 | 0.23029177 | 7.42219932 | 0.04277293 | 0.26317464 |
| ENSGALG00010008780 | KARS | 415885 | -0.1399973 | 8.19672418 | 0.04281028 | 0.26328221 |

| | | | | | | |
|---|---|---|---|---|---|---|
| ENSGALG00010000119 | LOC124416997 | 124416997 | -0.4646476 | 2.67961849 | 0.04282361 | 0.26328221 |
| ENSGALG00010015540 | ACSL4 | 422345 | -0.1724131 | 6.53250756 | 0.04286625 | 0.26339643 |
| ENSGALG00010023362 | GADD45A | 693255 | -0.3266183 | 3.07004202 | 0.0428754 | 0.26339643 |
| ENSGALG00010025340 | METTL21A | 424094 | -0.2281404 | 4.15106736 | 0.04289957 | 0.2634429 |
| ENSGALG00010000808 | ADCY2 | 420935 | 0.39577001 | 2.17254113 | 0.04295701 | 0.26355272 |
| ENSGALG00010004116 | SPRY1 | 395583 | 0.20488745 | 5.43044167 | 0.04296025 | 0.26355272 |
| ENSGALG00010006271 | PAX4 | 107054978 | 0.25067154 | 4.14668528 | 0.04296957 | 0.26355272 |
| ENSGALG00010024401 | KCNC1 | 423076 | 0.63862391 | 0.92393566 | 0.04298392 | 0.26355272 |
| ENSGALG00010002297 | NA | NA | 0.49559874 | 1.65604234 | 0.04302019 | 0.26367318 |
| ENSGALG00010008623 | TRPS1 | 420289 | 0.21900983 | 4.84870191 | 0.04304345 | 0.26371385 |
| ENSGALG00010020404 | NA | NA | 0.33189836 | 2.84827323 | 0.0430868 | 0.26387751 |
| ENSGALG00010011374 | RPEL1 | 770045 | -0.1883137 | 5.24875716 | 0.04312187 | 0.26399037 |
| ENSGALG00010001166 | TRIM59 | 425013 | 0.1523092 | 7.65900595 | 0.04331636 | 0.26505033 |
| ENSGALG00010017266 | PMM2 | 427679 | -0.1942581 | 5.30273848 | 0.04333162 | 0.26505033 |
| ENSGALG00010027510 | RHOC | 395869 | -0.4591869 | 2.14335909 | 0.04334514 | 0.26505033 |
| ENSGALG00010009612 | DERL1 | 420350 | -0.2287956 | 8.47400585 | 0.04337888 | 0.26515447 |
| ENSGALG00010021210 | SEC24C | 423742 | -0.1719105 | 7.44666298 | 0.04342498 | 0.26533401 |
| ENSGALG00010025214 | LRP8 | 396102 | 0.15361479 | 6.51576798 | 0.04348537 | 0.2656007 |
| ENSGALG00010028917 | NR6A1 | 772325 | -0.1809161 | 8.65754798 | 0.04355113 | 0.26585701 |
| ENSGALG00010016639 | NA | NA | 0.35884118 | 2.71711517 | 0.04356086 | 0.26585701 |
| ENSGALG00010024472 | LDHA | 396221 | 0.15067274 | 9.44977245 | 0.0435885 | 0.26590222 |
| ENSGALG00010019140 | NA | NA | 0.55498831 | 1.12379746 | 0.0436018 | 0.26590222 |
| ENSGALG00010017647 | ITGA9 | 420757 | 0.28680093 | 3.50839152 | 0.04362678 | 0.26595234 |
| ENSGALG00010004362 | FBXL4 | 421798 | 0.25941281 | 3.75947747 | 0.04368679 | 0.26621586 |
| ENSGALG00010027154 | PLPP2 | 100858471 | -0.2307603 | 4.56963572 | 0.04389205 | 0.26736392 |
| ENSGALG00010027357 | GHRHR-LR | 428397 | -0.4194396 | 4.03222489 | 0.04398866 | 0.26783042 |
| ENSGALG00010011880 | U2AF1 | 395370 | 0.14400184 | 7.54352722 | 0.0440024 | 0.26783042 |
| ENSGALG00010005366 | AUP1 | 769206 | -0.1691173 | 5.77763478 | 0.04403043 | 0.26786845 |
| ENSGALG00010014909 | FBXL14 | 427926 | -0.1963226 | 4.68860717 | 0.04404242 | 0.26786845 |
| ENSGALG00010019744 | KBP | 396300 | 0.57980818 | 0.8322638 | 0.0440943 | 0.26808118 |
| ENSGALG00010021256 | PML | 415302 | 0.26417439 | 3.74221491 | 0.04412941 | 0.26819187 |
| ENSGALG00010000340 | LOC107049486 | 107049486 | -0.193797 | 5.74135932 | 0.04416735 | 0.26831961 |
| ENSGALG00010025119 | NA | NA | 0.7862346 | 0.04102935 | 0.04421951 | 0.26853366 |
| ENSGALG00010013019 | DPYS | 420266 | 0.44809264 | 2.28152231 | 0.04423949 | 0.26855213 |
| ENSGALG00010023264 | RACGAP1 | 426884 | -0.1440444 | 6.25312167 | 0.04426517 | 0.26860522 |
| ENSGALG00010022958 | SFSWAP | 416802 | -0.1460068 | 6.63300893 | 0.04434366 | 0.26889392 |
| ENSGALG00010021076 | TBR1 | 429029 | -0.3949216 | 1.95330964 | 0.04434665 | 0.26889392 |
| ENSGALG00010010225 | NA | NA | 0.40685379 | 3.04151056 | 0.04445662 | 0.26945771 |
| ENSGALG00010027468 | BSG | 770363 | -0.1599202 | 8.6102303 | 0.04452002 | 0.26968092 |
| ENSGALG00010001191 | TOMM5 | 426803 | -0.1595187 | 6.78012262 | 0.04452749 | 0.26968092 |
| ENSGALG00010026242 | CCDC141 | 770766 | 0.25335864 | 4.02247332 | 0.04457016 | 0.26968092 |
| ENSGALG00010018955 | ZBTB48 | 419370 | -0.2055188 | 4.90808318 | 0.04458068 | 0.26968092 |
| ENSGALG00010021105 | LOC112532477 | 112532477 | 0.44821284 | 1.95653401 | 0.0445889 | 0.26968092 |
| ENSGALG00010005748 | AIG1 | 421671 | -0.2337435 | 5.58143591 | 0.04460951 | 0.26968092 |
| ENSGALG00010014214 | REEP2 | 771891 | -0.1783103 | 5.42488731 | 0.04462757 | 0.26968092 |
| ENSGALG00010003384 | WDR17 | 422570 | 1.07970664 | -0.2205724 | 0.04462945 | 0.26968092 |
| ENSGALG00010019848 | MAD1L1 | 427658 | 0.19742477 | 5.62697151 | 0.04466944 | 0.26974615 |
| ENSGALG00010015946 | TLE4 | 374080 | -0.174854 | 5.812705 | 0.04467426 | 0.26974615 |
| ENSGALG00010014125 | PLPP1 | 427138 | 0.16398735 | 7.65733406 | 0.04472784 | 0.26996686 |
| ENSGALG00010016891 | LOC107052288 | 107052288 | -0.594182 | 3.05813563 | 0.0447956 | 0.27027302 |
| ENSGALG00010008375 | PLCG1L | 776273 | -0.232591 | 6.40724746 | 0.04482756 | 0.27036304 |
| ENSGALG00010020862 | NA | NA | 0.48704797 | 1.95465663 | 0.04487355 | 0.27049914 |
| ENSGALG00010008383 | PLD5 | 421501 | 0.33876124 | 3.20005534 | 0.04488423 | 0.27049914 |
| ENSGALG00010001228 | THAP5 | 417784 | -0.1837562 | 5.94642935 | 0.0449142 | 0.27057694 |
| ENSGALG00010005150 | ALCAM | 396092 | -0.215327 | 4.93307151 | 0.04494177 | 0.27064025 |
| ENSGALG00010008012 | CCDC90B | 419023 | -0.1480157 | 5.97673266 | 0.04496759 | 0.27069294 |
| ENSGALG00010017771 | PNO1 | 769547 | -0.1821094 | 5.11347928 | 0.04500089 | 0.27079064 |
| ENSGALG00010029757 | ULK2 | 417614 | 0.16064774 | 6.04975429 | 0.04502706 | 0.27084538 |
| ENSGALG00010005495 | TXN | 396437 | -0.140941 | 8.04198517 | 0.04509689 | 0.27116255 |
| ENSGALG00010021735 | PIGBOS1 | 105694155 | -0.2141679 | 4.69549303 | 0.04512778 | 0.2712455 |

| | | | | | | |
|---|---|---|---|---|---|---|
| ENSGALG00010019113 | KLHDC1 | 423568 | -0.1665513 | 5.71103831 | 0.04515231 | 0.2712901 |
| ENSGALG00010000100 | NA | NA | -0.491473 | 2.28531732 | 0.04527372 | 0.27191659 |
| ENSGALG00010024608 | DENND2D | 421153 | 0.22181037 | 4.69840271 | 0.04532309 | 0.27196956 |
| ENSGALG00010018545 | NA | NA | 0.2335678 | 4.05255931 | 0.04533815 | 0.27196956 |
| ENSGALG00010010263 | RAB4A | 421532 | 0.19980457 | 5.15927482 | 0.04533977 | 0.27196956 |
| ENSGALG00010016653 | EPHA2 | 771550 | 0.18159487 | 5.74643537 | 0.04535112 | 0.27196956 |
| ENSGALG00010028010 | ASB6 | 417194 | -0.1766268 | 5.07917066 | 0.04541197 | 0.27223157 |
| ENSGALG00010024423 | SERGEF | 423077 | 0.163711 | 6.13149422 | 0.04549845 | 0.27264691 |
| ENSGALG00010002619 | RAB6A | 419063 | -0.2090269 | 7.52475081 | 0.04557057 | 0.27284275 |
| ENSGALG00010011650 | RPL30 | 425416 | -0.1821764 | 7.61822547 | 0.04557453 | 0.27284275 |
| ENSGALG00010010099 | TNR | 396213 | 0.53757297 | 1.27374098 | 0.04560313 | 0.27284275 |
| ENSGALG00010003001 | MAP6 | 395753 | 0.17095351 | 5.64801108 | 0.04560425 | 0.27284275 |
| ENSGALG00010014644 | NA | NA | -0.5688971 | 0.93158893 | 0.04561713 | 0.27284275 |
| ENSGALG00010009676 | ACO1 | 373916 | -0.1446758 | 7.26702623 | 0.0457049 | 0.27326463 |
| ENSGALG00010005428 | ZNF770 | 423295 | -0.2055785 | 4.37859373 | 0.04582524 | 0.27382172 |
| ENSGALG00010023214 | SUPV3L1 | 423697 | -0.1830052 | 5.74375203 | 0.0458326 | 0.27382172 |
| ENSGALG00010007960 | TSNAX | 421526 | -0.1678863 | 6.71579205 | 0.04585855 | 0.27387364 |
| ENSGALG00010013369 | MTMR8 | 422312 | 0.14500572 | 7.36888301 | 0.04587626 | 0.27387627 |
| ENSGALG00010018922 | EEF2K | 416619 | -0.2198293 | 4.98688756 | 0.04590372 | 0.27389433 |
| ENSGALG00010025268 | ZBTB39 | 100858734 | -0.1623672 | 6.07374975 | 0.04591382 | 0.27389433 |
| ENSGALG00010013323 | NOTCH2 | 374031 | 0.14163751 | 7.24483924 | 0.04600761 | 0.27435065 |
| ENSGALG00010003478 | COLEC11 | 428649 | -0.3674739 | 2.37973423 | 0.04611367 | 0.27487198 |
| ENSGALG00010017201 | AP5M1 | 423548 | -0.2029828 | 4.90557287 | 0.04613183 | 0.27487198 |
| ENSGALG00010024748 | TRPM1 | 427494 | 0.79489928 | 0.89174944 | 0.04616029 | 0.27487198 |
| ENSGALG00010006177 | NEK9 | 428891 | 0.13757964 | 8.70833851 | 0.04616435 | 0.27487198 |
| ENSGALG00010010371 | SOCS1L | 417978 | 0.21388681 | 4.50083568 | 0.04626939 | 0.27539404 |
| ENSGALG00010011371 | LOC424430 | 424430 | 0.24993763 | 3.8750151 | 0.04630987 | 0.27553157 |
| ENSGALG00010023449 | NA | NA | -0.382384 | 2.57016753 | 0.04636116 | 0.27556195 |
| ENSGALG00010011638 | GRPEL1 | 422865 | -0.2661 | 4.2269893 | 0.04637019 | 0.27556195 |
| ENSGALG00010006977 | NA | NA | 0.69142661 | 0.86508271 | 0.04638416 | 0.27556195 |
| ENSGALG00010022805 | A2ML1 | 418254 | -0.271394 | 3.73826962 | 0.04639236 | 0.27556195 |
| ENSGALG00010000585 | SPTBN2 | 101750255 | 0.15482862 | 7.22976842 | 0.04640184 | 0.27556195 |
| ENSGALG00010015151 | LOC771085 | 771085 | -0.5242432 | 1.47104202 | 0.04647472 | 0.27589146 |
| ENSGALG00010025238 | IGSF9B | 428237 | 0.34539641 | 4.61679794 | 0.04649717 | 0.2759011 |
| ENSGALG00010003596 | CRISPLD1 | 395369 | 0.18342441 | 5.36971607 | 0.04654432 | 0.2759011 |
| ENSGALG00010006002 | NA | NA | -0.5812843 | 0.98574982 | 0.04655268 | 0.2759011 |
| ENSGALG00010016937 | DHRS7 | 423527 | -0.1758548 | 5.13261998 | 0.04656887 | 0.2759011 |
| ENSGALG00010014919 | NA | NA | 0.14566652 | 7.17446602 | 0.04657502 | 0.2759011 |
| ENSGALG00010022929 | LRP4 | 423189 | 0.16284895 | 5.88637679 | 0.04658855 | 0.2759011 |
| ENSGALG00010026106 | TESK2 | 100858275 | 0.22870012 | 4.11497927 | 0.0465981 | 0.2759011 |
| ENSGALG00010020635 | NDE1 | 416597 | 0.17410187 | 5.56232025 | 0.04664228 | 0.27605963 |
| ENSGALG00010008735 | LOC112530752 | 112530752 | -0.8082901 | -0.0391475 | 0.04666025 | 0.276063 |
| ENSGALG00010013921 | PPP2R2B | 416350 | -0.2279407 | 4.26695063 | 0.04670722 | 0.27622487 |
| ENSGALG00010003455 | NA | NA | 0.30965336 | 3.28016038 | 0.04673728 | 0.27622487 |
| ENSGALG00010003407 | CHMP4C | 420201 | 0.45442001 | 1.89877823 | 0.04673985 | 0.27622487 |
| ENSGALG00010009200 | ADGRA3 | 422817 | -0.1563462 | 7.00059761 | 0.04678182 | 0.27633509 |
| ENSGALG00010021812 | TEAD1 | 403089 | 0.20924119 | 6.38182307 | 0.04679335 | 0.27633509 |
| ENSGALG00010018057 | SSTR4 | 428547 | 0.79861275 | 0.28146493 | 0.04687943 | 0.27666409 |
| ENSGALG00010002129 | NA | NA | 0.95038424 | -0.4746702 | 0.04688394 | 0.27666409 |
| ENSGALG00010025035 | NA | NA | -0.6352519 | 0.46148051 | 0.04694281 | 0.27690843 |
| ENSGALG00010007325 | NA | NA | -0.4840409 | 1.36092891 | 0.04701217 | 0.2770431 |
| ENSGALG00010016517 | NA | NA | 0.33733783 | 2.83898479 | 0.04701832 | 0.2770431 |
| ENSGALG00010017657 | DIS3L2 | 424934 | 0.15342626 | 6.30461125 | 0.04702275 | 0.2770431 |
| ENSGALG00010019649 | TMEM63B | 421442 | -0.1675676 | 6.49783434 | 0.04704782 | 0.2770431 |
| ENSGALG00010018264 | CFAP61 | 421242 | 0.53423826 | 1.19640817 | 0.04705296 | 0.2770431 |
| ENSGALG00010019415 | NA | NA | 0.42892556 | 2.35305588 | 0.04709106 | 0.27716449 |
| ENSGALG00010011979 | NEDD1 | 417922 | 0.16263111 | 6.69160896 | 0.04714724 | 0.27730838 |
| ENSGALG00010028503 | NDOR1 | 417280 | -0.2722626 | 4.34604522 | 0.04715984 | 0.27730838 |
| ENSGALG00010021252 | XPC | 416039 | -0.1552014 | 5.78193292 | 0.04716795 | 0.27730838 |
| ENSGALG00010008186 | GLTSCR2 | 107049993 | -0.1898664 | 5.34422138 | 0.04722175 | 0.27751067 |

| | | | | | | |
|---|---|---|---|---|---|---|
| ENSGALG00010010966 | LOC768760 | 768760 | 0.95267102 | -0.0010955 | 0.04725437 | 0.27751067 |
| ENSGALG00010016932 | OGT | 422203 | -0.1380779 | 8.91161537 | 0.04727036 | 0.27751067 |
| ENSGALG00010010227 | OLIG2 | 428612 | 0.94901775 | -0.4748219 | 0.04727234 | 0.27751067 |
| ENSGALG00010009054 | RYR2 | 421508 | -0.2639963 | 4.83090725 | 0.04733645 | 0.27768498 |
| ENSGALG00010005641 | NA | NA | -0.5828724 | 1.39633668 | 0.04734923 | 0.27768498 |
| ENSGALG00010029634 | DDX5 | 395629 | -0.133018 | 9.75574931 | 0.04735455 | 0.27768498 |
| ENSGALG00010025541 | TTC21B | 424176 | 0.24178616 | 5.26076846 | 0.04740978 | 0.27790611 |
| ENSGALG00010019026 | CHSY1 | 415521 | -0.1526442 | 6.33683971 | 0.04744648 | 0.27798004 |
| ENSGALG00010022946 | KRT17 | 100858439 | 0.8867811 | -0.2848801 | 0.04745785 | 0.27798004 |
| ENSGALG00010003251 | ELOF1 | 107057620 | 0.34370652 | 3.97152661 | 0.04748586 | 0.27798004 |
| ENSGALG00010012896 | STK38L | 418217 | -0.2379199 | 7.55263186 | 0.04749249 | 0.27798004 |
| ENSGALG00010024966 | AGAP3 | 768507 | -0.2114188 | 5.26927715 | 0.04753006 | 0.27804782 |
| ENSGALG00010019697 | EIF4EBP1 | 426773 | -0.1419061 | 6.52391425 | 0.04753913 | 0.27804782 |
| ENSGALG00010002235 | MXRA5 | 418657 | -0.4378607 | 1.97229566 | 0.04761468 | 0.27838705 |
| ENSGALG00010027050 | GSTT1 | 396322 | -0.2928806 | 3.22519223 | 0.04763656 | 0.27841233 |
| ENSGALG00010000465 | CUNH17orf49 | 112531156 | 0.19025976 | 6.13855054 | 0.04779824 | 0.2792544 |
| ENSGALG00010016949 | LUC7L | 416654 | -0.1704091 | 5.58289153 | 0.04783759 | 0.27938138 |
| ENSGALG00010018289 | HEYL | 771101 | 0.45343921 | 1.90576479 | 0.04788562 | 0.27955898 |
| ENSGALG00010007391 | NANP | 421467 | -0.2569894 | 5.18328922 | 0.04793373 | 0.27973689 |
| ENSGALG00010015362 | ATP9A | 419345 | -0.1696262 | 6.93713195 | 0.04810965 | 0.28066024 |
| ENSGALG00010001958 | RAB11FIP2 | 423924 | 0.1894865 | 4.8289598 | 0.04818401 | 0.28099072 |
| ENSGALG00010024446 | GRM4 | 419906 | 0.30225498 | 2.97399852 | 0.04823529 | 0.28118638 |
| ENSGALG00010020130 | PDXDC1 | 416413 | -0.16595 | 6.1378897 | 0.04828871 | 0.28132367 |
| ENSGALG00010025362 | G6PC2 | 424171 | 0.30531012 | 3.27781109 | 0.04829431 | 0.28132367 |
| ENSGALG00010004944 | ALKBH1 | 428898 | -0.1951423 | 4.57164384 | 0.04832946 | 0.28142505 |
| ENSGALG00010004949 | RAB39A | 101748614 | 0.52584877 | 1.10891505 | 0.04839181 | 0.2815885 |
| ENSGALG00010023493 | LOC121106536 | 121106536 | 0.32388023 | 3.21999509 | 0.04839303 | 0.2815885 |
| ENSGALG00010024937 | ILKAP | 424850 | -0.1624083 | 6.35599043 | 0.04842697 | 0.28168264 |
| ENSGALG00010018539 | CHD2 | 415507 | -0.1714783 | 6.78590769 | 0.04846042 | 0.28175831 |
| ENSGALG00010001335 | NA | NA | 0.39355241 | 2.18653597 | 0.04849407 | 0.28175831 |
| ENSGALG00010029326 | NCBP3 | 417552 | -0.1598349 | 5.89137072 | 0.0484956 | 0.28175831 |
| ENSGALG00010005738 | ELAVL2 | 770158 | 0.17757344 | 7.23727452 | 0.04851103 | 0.28175831 |
| ENSGALG00010005836 | NEBL | 395148 | -0.2048754 | 4.4157669 | 0.04857056 | 0.28191533 |
| ENSGALG00010007381 | EGFR | 396494 | 0.1658582 | 5.52985772 | 0.04857361 | 0.28191533 |
| ENSGALG00010028896 | ATP6V1G1 | 417249 | -0.1478645 | 6.552608 | 0.04859486 | 0.28193548 |
| ENSGALG00010014987 | KLF9 | 770238 | 0.55548362 | 0.92969063 | 0.04862058 | 0.28198161 |
| ENSGALG00010029704 | SLC25A19 | 422123 | -0.2577646 | 3.79543389 | 0.04867231 | 0.28211095 |
| ENSGALG00010000094 | NA | NA | 0.62281416 | 0.7706871 | 0.04868454 | 0.28211095 |
| ENSGALG00010001285 | NA | NA | 0.27225955 | 3.59614495 | 0.04869917 | 0.28211095 |
| ENSGALG00010017886 | DDRGK1 | 422941 | -0.16567 | 5.91520313 | 0.04871403 | 0.28211095 |
| ENSGALG00010015508 | FUCA1 | 419687 | -0.2933136 | 3.79297883 | 0.04878552 | 0.28233391 |
| ENSGALG00010012899 | CCDC107 | 417860 | -0.2320841 | 4.32327443 | 0.04879285 | 0.28233391 |
| ENSGALG00010027884 | SF3A2 | 420077 | -0.1599383 | 6.50840526 | 0.04881835 | 0.28233391 |
| ENSGALG00010015025 | TRAF3IP1 | 424029 | -0.1688235 | 5.8607438 | 0.04882372 | 0.28233391 |
| ENSGALG00010006953 | MAP1LC3B | 427559 | -0.1643088 | 6.92069366 | 0.04887226 | 0.28250389 |
| ENSGALG00010023193 | PPDPF | 419245 | -0.3213336 | 4.19486849 | 0.04888874 | 0.28250389 |
| ENSGALG00010023338 | HDLBP | 395984 | -0.1491022 | 9.99656038 | 0.04897154 | 0.28281617 |
| ENSGALG00010007758 | DEK | 420828 | -0.1411902 | 7.49557614 | 0.04897844 | 0.28281617 |
| ENSGALG00010006382 | PNN | 423338 | -0.1538904 | 7.90285873 | 0.04904377 | 0.28308076 |
| ENSGALG00010029476 | NA | NA | 0.92148304 | 1.62232014 | 0.04907108 | 0.28308076 |
| ENSGALG00010029204 | HELZ | 417427 | 0.13736977 | 6.58190767 | 0.04909381 | 0.28308076 |
| ENSGALG00010028772 | TTF1 | 417169 | -0.190545 | 4.9691272 | 0.04909565 | 0.28308076 |
| ENSGALG00010000281 | OSBP2 | 417002 | 0.27248614 | 3.77779709 | 0.04918097 | 0.28346969 |
| ENSGALG00010012940 | FBN2 | 374203 | 0.21017634 | 4.62747329 | 0.04920902 | 0.2835283 |
| ENSGALG00010016215 | SPATA17 | 421354 | 0.45348842 | 1.93887079 | 0.04924655 | 0.28364151 |
| ENSGALG00010013938 | NA | NA | -0.4390025 | 1.64260191 | 0.04929065 | 0.2837925 |
| ENSGALG00010021296 | GATC | 416976 | -0.2826556 | 3.49591778 | 0.0493599 | 0.28408806 |
| ENSGALG00010021934 | EFTUD2 | 428281 | 0.1506174 | 7.89888194 | 0.04943266 | 0.28440363 |
| ENSGALG00010015884 | MIR7450 | 102465726 | 0.75252338 | 0.37799062 | 0.04946515 | 0.28448739 |
| ENSGALG00010027128 | EIF2D | 419845 | -0.1763475 | 5.34070609 | 0.04951266 | 0.2846574 |

| | | | | | | |
|---|---|---|---|---|---|---|
| ENSGALG00010013994 | CHRM2 | 418126 | 0.69570335 | 1.07114058 | 0.04953297 | 0.28467101 |
| ENSGALG00010027096 | YIPF1 | 424653 | -0.2441383 | 5.41020469 | 0.04959338 | 0.28491493 |
| ENSGALG00010003125 | APOO | 418599 | -0.1735258 | 5.33260177 | 0.04964546 | 0.28503341 |
| ENSGALG00010008821 | TAF1D | 419002 | -0.2965213 | 4.0333512 | 0.04964994 | 0.28503341 |
| ENSGALG00010022800 | KLHL10 | 420033 | -0.5784628 | 0.73794393 | 0.04978288 | 0.28569322 |
| ENSGALG00010005560 | RSPH3 | 421663 | 0.27049202 | 3.72029792 | 0.04981571 | 0.28577823 |
| ENSGALG00010000356 | LOC112533553 | 112533553 | -0.3637144 | 2.59988497 | 0.04987416 | 0.28601009 |
| ENSGALG00010023607 | NA | NA | 0.76153458 | 0.28559259 | 0.04993583 | 0.2861799 |
| ENSGALG00010012812 | DECR1 | 420218 | -0.2764835 | 5.02177837 | 0.04993986 | 0.2861799 |

### Supplementally table S10. GO analysis of down-regulated genes comparing 3x sgSoprano and 3x sgOVA in CRISPRi PGCs

| source | term_name | term_id | adjusted_p_value | negative_log10_of_adjusted_p_value | term_size | query_size | intersection_size |
|---|---|---|---|---|---|---|---|
| GO:MF | structural constituent of ribosome | GO:0003735 | 4.11E-26 | 25.38658745 | 202 | 491 | 52 |
| GO:MF | structural molecule activity | GO:0005198 | 9.99E-12 | 11.00049568 | 582 | 491 | 64 |
| GO:MF | oxidoreduction-driven active transmembrane transporter activity | GO:0015453 | 1.90E-07 | 6.721829008 | 24 | 491 | 11 |
| GO:MF | electron transfer activity | GO:0009055 | 1.52974E-05 | 4.815382671 | 34 | 491 | 11 |
| GO:MF | proton transmembrane transporter activity | GO:0015078 | 1.74786E-05 | 4.757494259 | 110 | 491 | 19 |
| GO:MF | RNA binding | GO:0003723 | 0.000122044 | 3.913483561 | 803 | 491 | 61 |
| GO:MF | NADH dehydrogenase (ubiquinone) activity | GO:0008137 | 0.000282049 | 3.549675292 | 15 | 491 | 7 |
| GO:MF | mRNA binding | GO:0003729 | 0.000415352 | 3.381584074 | 187 | 491 | 23 |
| GO:MF | oxidoreductase activity, acting on NAD(P)H, quinone or similar compound as acceptor | GO:0016655 | 0.0004335 | 3.363011092 | 22 | 491 | 8 |
| GO:MF | ubiquitin ligase inhibitor activity | GO:1990948 | 0.004960436 | 2.304480106 | 5 | 491 | 4 |
| GO:MF | ribosome binding | GO:0043022 | 0.013902374 | 1.856911021 | 64 | 491 | 11 |
| GO:MF | oxidoreductase activity, acting on NAD(P)H | GO:0016651 | 0.01450916 | 1.83835772 | 43 | 491 | 9 |
| GO:MF | intramolecular oxidoreductase activity | GO:0016860 | 0.019896663 | 1.701219745 | 26 | 491 | 7 |
| GO:MF | intramolecular oxidoreductase activity, transposing S-S bonds | GO:0016864 | 0.024801794 | 1.605516911 | 12 | 491 | 5 |
| GO:MF | protein disulfide isomerase activity | GO:0003756 | 0.024801794 | 1.605516911 | 12 | 491 | 5 |
| GO:MF | smoothened binding | GO:0005119 | 0.026621625 | 1.574765433 | 3 | 491 | 3 |
| GO:MF | translation factor activity, RNA binding | GO:0008135 | 0.03609661 | 1.44253358 | 59 | 491 | 10 |
| GO:MF | mRNA 5'-UTR binding | GO:0048027 | 0.036641698 | 1.436024411 | 20 | 491 | 6 |
| GO:BP | translation | GO:0006412 | 1.44E-22 | 21.84126534 | 439 | 491 | 72 |
| GO:BP | response to endoplasmic reticulum stress | GO:0034976 | 1.09E-09 | 8.961995644 | 149 | 491 | 29 |
| GO:BP | ribosome biogenesis | GO:0042254 | 1.47E-07 | 6.832765395 | 192 | 491 | 30 |
| GO:BP | protein metabolic process | GO:0019538 | 6.43E-07 | 6.191597615 | 2987 | 491 | 178 |
| GO:BP | cytoplasmic translation | GO:0002181 | 1.85275E-05 | 4.732182184 | 71 | 491 | 16 |
| GO:BP | ribonucleoprotein complex biogenesis | GO:0022613 | 0.000665456 | 3.176880563 | 335 | 491 | 34 |
| GO:BP | endoplasmic reticulum unfolded protein response | GO:0030968 | 0.002447876 | 2.611210569 | 47 | 491 | 11 |
| GO:BP | oxidative phosphorylation | GO:0006119 | 0.002496163 | 2.602727084 | 77 | 491 | 14 |
| GO:BP | retrograde protein transport, ER to cytosol | GO:0030970 | 0.003680658 | 2.434074541 | 24 | 491 | 8 |
| GO:BP | endoplasmic reticulum to cytosol transport | GO:1903513 | 0.003680658 | 2.434074541 | 24 | 491 | 8 |
| GO:BP | ribosomal small subunit biogenesis | GO:0042274 | 0.005421664 | 2.265867368 | 71 | 491 | 13 |
| GO:BP | ER-nucleus signaling pathway | GO:0006984 | 0.005621494 | 2.250148272 | 33 | 491 | 9 |
| GO:BP | response to unfolded protein | GO:0006986 | 0.008386035 | 2.076443314 | 63 | 491 | 12 |
| GO:BP | protein targeting to ER | GO:0045047 | 0.010004881 | 1.999788057 | 27 | 491 | 8 |
| GO:BP | rRNA processing | GO:0006364 | 0.012348663 | 1.908380054 | 112 | 491 | 16 |
| GO:BP | protein localization to endoplasmic reticulum | GO:0070972 | 0.014932572 | 1.82586537 | 46 | 491 | 10 |
| GO:BP | cellular response to unfolded protein | GO:0034620 | 0.015167641 | 1.819081957 | 56 | 491 | 11 |
| GO:BP | rRNA metabolic process | GO:0016072 | 0.019465216 | 1.710740771 | 129 | 491 | 17 |
| GO:BP | protein exit from endoplasmic reticulum | GO:0032527 | 0.019920826 | 1.700692659 | 38 | 491 | 9 |
| GO:BP | ERAD pathway | GO:0036503 | 0.022318097 | 1.651342848 | 48 | 491 | 10 |
| GO:BP | establishment of protein localization to endoplasmic reticulum | GO:0072599 | 0.023755347 | 1.624238628 | 30 | 491 | 8 |
| GO:BP | biosynthetic process | GO:0009058 | 0.032309287 | 1.490672625 | 5073 | 491 | 245 |
| GO:BP | response to topologically incorrect protein | GO:0035966 | 0.040354727 | 1.394105584 | 73 | 491 | 12 |
| GO:CC | cytosolic ribosome | GO:0022626 | 6.64E-37 | 36.17752215 | 79 | 496 | 43 |
| GO:CC | ribosomal subunit | GO:0044391 | 5.41E-36 | 35.26656785 | 125 | 496 | 51 |
| GO:CC | ribosome | GO:0005840 | 6.79E-26 | 25.16844398 | 250 | 496 | 58 |
| GO:CC | cytosolic large ribosomal subunit | GO:0022625 | 4.44E-25 | 24.35268957 | 39 | 496 | 26 |
| GO:CC | large ribosomal subunit | GO:0015934 | 6.31E-21 | 20.19996475 | 84 | 496 | 32 |
| GO:CC | cytoplasm | GO:0005737 | 4.55E-17 | 16.34239884 | 6577 | 496 | 363 |
| GO:CC | cytosolic small ribosomal subunit | GO:0022627 | 1.58E-14 | 13.80131286 | 25 | 496 | 16 |
| GO:CC | intracellular anatomical structure | GO:0005622 | 4.14E-14 | 13.38290898 | 9235 | 496 | 446 |
| GO:CC | endoplasmic reticulum | GO:0005783 | 5.65E-14 | 13.24780393 | 921 | 496 | 92 |
| GO:CC | small ribosomal subunit | GO:0015935 | 2.13E-13 | 12.67180349 | 43 | 496 | 19 |
| GO:CC | ribonucleoprotein complex | GO:1990904 | 5.88E-11 | 10.23070987 | 659 | 496 | 69 |
| GO:CC | endoplasmic reticulum protein-containing complex | GO:0140534 | 2.58E-09 | 8.588079583 | 75 | 496 | 20 |
| GO:CC | nuclear outer membrane-endoplasmic reticulum membrane network | GO:0042175 | 1.72E-07 | 6.765554704 | 419 | 496 | 46 |
| GO:CC | intracellular organelle | GO:0043229 | 1.91E-07 | 6.718143585 | 8187 | 496 | 393 |
| GO:CC | endoplasmic reticulum membrane | GO:0005789 | 2.12E-07 | 6.674212441 | 407 | 496 | 45 |
| GO:CC | endoplasmic reticulum subcompartment | GO:0098827 | 3.66E-07 | 6.436418854 | 414 | 496 | 45 |
| GO:CC | organelle | GO:0043226 | 1.75459E-06 | 5.755825096 | 8395 | 496 | 397 |
| GO:CC | organelle subcompartment | GO:0031984 | 0.000141336 | 3.849747669 | 589 | 496 | 50 |
| GO:CC | endomembrane system | GO:0012505 | 0.000567097 | 3.246342808 | 2282 | 496 | 134 |
| GO:CC | mitochondrial inner membrane | GO:0005743 | 0.000966962 | 3.014590807 | 208 | 496 | 24 |
| GO:CC | respiratory chain complex | GO:0098803 | 0.001019846 | 2.99146542 | 25 | 496 | 8 |
| GO:CC | endoplasmic reticulum lumen | GO:0005788 | 0.0019809 | 2.703137367 | 44 | 496 | 10 |
| GO:CC | intracellular membrane-bounded organelle | GO:0043231 | 0.002176791 | 2.662183209 | 7131 | 496 | 336 |
| GO:CC | organelle membrane | GO:0031090 | 0.002227906 | 2.652103158 | 1540 | 496 | 96 |
| GO:CC | cytosol | GO:0005829 | 0.004882495 | 2.311358161 | 2355 | 496 | 133 |
| GO:CC | preribosome | GO:0030684 | 0.006319881 | 2.199291093 | 71 | 496 | 12 |
| GO:CC | organelle inner membrane | GO:0019866 | 0.006666744 | 2.176086245 | 233 | 496 | 24 |
| GO:CC | mitochondrial envelope | GO:0005740 | 0.007089302 | 2.149396536 | 361 | 496 | 32 |
| GO:CC | mitochondrial membrane | GO:0031966 | 0.00879428 | 2.055799691 | 332 | 496 | 30 |
| GO:CC | endoplasmic reticulum chaperone complex | GO:0034663 | 0.00934685 | 2.029334732 | 6 | 496 | 4 |
| GO:CC | protein-containing complex | GO:0032991 | 0.021811717 | 1.661310138 | 3764 | 496 | 192 |
| GO:CC | membrane-bounded organelle | GO:0043227 | 0.02435219 | 1.613461983 | 7412 | 496 | 341 |
| GO:CC | synapse | GO:0045202 | 0.027459824 | 1.561302249 | 733 | 496 | 51 |
| GO:CC | small-subunit processome | GO:0032040 | 0.029014412 | 1.537386221 | 59 | 496 | 10 |

### Supplementally table S11. GO analysis of up-regulated genes comparing 3x sgSoprano and 3x sgOVA in CRISPRi PGCs

| source | term_name | term_id | adjusted_p_value | negative_log10_of_adjusted_p_value | term_size | query_size | intersection_size |
|---|---|---|---|---|---|---|---|
| GO:MF | semaphorin receptor activity | GO:0017154 | 0.002733338 | 2.563306684 | 10 | 399 | 5 |
| GO:MF | platelet-derived growth factor binding | GO:0048407 | 0.044136414 | 1.355202953 | 9 | 399 | 4 |
| GO:BP | system development | GO:0048731 | 5.52E-09 | 8.257885062 | 2305 | 363 | 122 |
| GO:BP | cell motility | GO:0048870 | 1.46E-08 | 7.837036097 | 994 | 363 | 69 |
| GO:BP | cell migration | GO:0016477 | 8.33E-08 | 7.079478674 | 896 | 363 | 63 |
| GO:BP | multicellular organismal process | GO:0032501 | 7.75E-07 | 6.110905947 | 3848 | 363 | 168 |
| GO:BP | multicellular organism development | GO:0007275 | 1.01766E-06 | 5.992395666 | 2718 | 363 | 130 |
| GO:BP | locomotion | GO:0040011 | 2.58523E-06 | 5.587500346 | 724 | 363 | 52 |
| GO:BP | regulation of cell migration | GO:0030334 | 4.84443E-06 | 5.314757039 | 565 | 363 | 44 |
| GO:BP | nervous system development | GO:0007399 | 1.02956E-05 | 4.98734921 | 1380 | 363 | 78 |
| GO:BP | regulation of cell motility | GO:2000145 | 2.00945E-05 | 4.696923353 | 592 | 363 | 44 |
| GO:BP | regulation of locomotion | GO:0040012 | 2.39163E-05 | 4.621306859 | 617 | 363 | 45 |
| GO:BP | anatomical structure development | GO:0048856 | 3.05029E-05 | 4.51565887 | 3372 | 363 | 147 |
| GO:BP | neuron projection development | GO:0031175 | 3.8376E-05 | 4.415940596 | 562 | 363 | 42 |
| GO:BP | developmental process | GO:0032502 | 5.26564E-05 | 4.278548573 | 3583 | 363 | 153 |
| GO:BP | neuron differentiation | GO:0030182 | 5.59973E-05 | 4.251832536 | 839 | 363 | 54 |
| GO:BP | plasma membrane bounded cell projection organization | GO:0120036 | 5.92025E-05 | 4.227660135 | 935 | 363 | 58 |
| GO:BP | neuron development | GO:0048666 | 7.13116E-05 | 4.146839814 | 662 | 363 | 46 |
| GO:BP | generation of neurons | GO:0048699 | 7.34838E-05 | 4.133808534 | 893 | 363 | 56 |
| GO:BP | anatomical structure morphogenesis | GO:0009653 | 7.90186E-05 | 4.102270553 | 1657 | 363 | 86 |
| GO:BP | cell projection organization | GO:0030030 | 9.28291E-05 | 4.03231568 | 947 | 363 | 58 |
| GO:BP | neuron projection morphogenesis | GO:0048812 | 0.00017846 | 3.748459956 | 364 | 363 | 31 |
| GO:BP | plasma membrane bounded cell projection morphogenesis | GO:0120039 | 0.000328333 | 3.483685155 | 374 | 363 | 31 |
| GO:BP | cell projection morphogenesis | GO:0048858 | 0.000348502 | 3.457794947 | 375 | 363 | 31 |
| GO:BP | cell adhesion | GO:0007155 | 0.000371233 | 3.430353128 | 793 | 363 | 50 |
| GO:BP | neurogenesis | GO:0022008 | 0.000380811 | 3.419290021 | 1036 | 363 | 60 |
| GO:BP | circulatory system development | GO:0072359 | 0.000538936 | 3.268462575 | 709 | 363 | 46 |
| GO:BP | anatomical structure formation involved in morphogenesis | GO:0048646 | 0.000659402 | 3.180849748 | 714 | 363 | 46 |
| GO:BP | regulation of neuron projection development | GO:0010975 | 0.000962936 | 3.016402544 | 220 | 363 | 22 |
| GO:BP | cell junction organization | GO:0034330 | 0.001054798 | 2.976830782 | 415 | 363 | 32 |
| GO:BP | cell morphogenesis involved in neuron differentiation | GO:0048667 | 0.001348324 | 2.870205805 | 338 | 363 | 28 |
| GO:BP | cell morphogenesis | GO:0000902 | 0.002329282 | 2.632777851 | 584 | 363 | 39 |
| GO:BP | cell differentiation | GO:0030154 | 0.00265098 | 2.576593615 | 2365 | 363 | 106 |
| GO:BP | cellular developmental process | GO:0048869 | 0.00265098 | 2.576593615 | 2365 | 363 | 106 |
| GO:BP | axon guidance | GO:0007411 | 0.003932143 | 2.405370663 | 135 | 363 | 16 |
| GO:BP | neuron projection guidance | GO:0097485 | 0.004339699 | 2.362540391 | 136 | 363 | 16 |
| GO:BP | cell junction assembly | GO:0034329 | 0.006756228 | 2.170295673 | 247 | 363 | 22 |
| GO:BP | regulation of plasma membrane bounded cell projection organization | GO:0120035 | 0.007108928 | 2.148195876 | 347 | 363 | 27 |
| GO:BP | regulation of cell projection organization | GO:0031344 | 0.009796259 | 2.008939732 | 353 | 363 | 27 |
| GO:BP | tube morphogenesis | GO:0035239 | 0.010560051 | 1.976333968 | 552 | 363 | 36 |
| GO:BP | regulation of developmental process | GO:0050793 | 0.012192201 | 1.91391787 | 1445 | 363 | 71 |
| GO:BP | regulation of biological process | GO:0050789 | 0.014009544 | 1.853576009 | 7293 | 363 | 253 |
| GO:BP | blood vessel development | GO:0001568 | 0.016664184 | 1.778215955 | 450 | 363 | 31 |
| GO:BP | axonogenesis | GO:0007409 | 0.018625888 | 1.729883003 | 263 | 363 | 22 |
| GO:BP | vasculature development | GO:0001944 | 0.020429237 | 1.689747851 | 477 | 363 | 32 |
| GO:BP | positive regulation of biological process | GO:0048518 | 0.02377034 | 1.623964602 | 3676 | 363 | 145 |
| GO:BP | cell surface receptor signaling pathway | GO:0007166 | 0.024863442 | 1.604438751 | 1558 | 363 | 74 |
| GO:BP | regulation of cellular process | GO:0050794 | 0.029229368 | 1.534180578 | 7022 | 363 | 244 |
| GO:BP | animal organ development | GO:0048513 | 0.033548779 | 1.474323287 | 1740 | 363 | 80 |
| GO:BP | cell development | GO:0048468 | 0.042294259 | 1.373718584 | 1610 | 363 | 75 |
| GO:BP | angiogenesis | GO:0001525 | 0.043630641 | 1.360208402 | 319 | 363 | 24 |
| GO:BP | positive regulation of cellular process | GO:0048522 | 0.049230491 | 1.307765835 | 3471 | 363 | 137 |
| GO:CC | cell periphery | GO:0071944 | 0.000211026 | 3.675664827 | 2860 | 355 | 123 |
| GO:CC | plasma membrane | GO:0005886 | 0.004557926 | 2.34123273 | 2604 | 355 | 109 |
| GO:CC | focal adhesion | GO:0005925 | 0.004980675 | 2.302711807 | 111 | 355 | 13 |
| GO:CC | collagen trimer | GO:0005581 | 0.005508632 | 2.258956201 | 21 | 355 | 6 |
| GO:CC | cell-substrate junction | GO:0030055 | 0.009606045 | 2.017455397 | 118 | 355 | 13 |
| GO:CC | neuron to neuron synapse | GO:0098984 | 0.027143684 | 1.566331215 | 167 | 355 | 15 |
| GO:CC | asymmetric synapse | GO:0032279 | 0.028372643 | 1.547100205 | 149 | 355 | 14 |
| GO:CC | complex of collagen trimers | GO:0098644 | 0.03254091 | 1.487570301 | 10 | 355 | 4 |

Supplementally table S12. Intersection of DEGs comparing sgSoprano#1, sgSoprano#2 and 3x sgSoprano in CRISPRi PGCs (p-adj 0.05 and lFCl > 2 in 3x sgSoprano)

| Ensembl_ID | Symbol | entrez_ID | logFC | p-adj_value | FDR | Discriptions |
|---|---|---|---|---|---|---|
| ENSGALG00010007287 | LOC124418405 | 124418405 | -2.5503477 | 1.16E-120 | 1.42E-117 | ENS-1/ERNI-like |
| ENSGALG00010006087 | LOC107051465 | 107051465 | -3.890844 | 1.09E-93 | 7.88E-91 | ENS-1/ERNI-like |
| ENSGALG00010006086 | LOC100857117 | 100857117 | -4.0529589 | 6.00E-91 | 3.66E-88 | ENS-1/ERNI-like |
| ENSGALG00010011206 | LOC112532760 | 112532760 | 1.00018373 | 7.92E-09 | 7.43E-07 | ncRNA |
| ENSGALG00010007691 | LOC121108447 | 121108447 | -1.2921579 | 1.13E-16 | 1.37E-14 | ncRNA |
| ENSGALG00010004570 | LOC107051826 | 107051826 | -4.7005305 | 1.21E-55 | 2.86E-53 | ENS-1/ERNI-like |
| ENSGALG00010001209 | CDH7 | 374007 | -1.4392055 | 1.87E-31 | 3.00E-29 | Cadherin 7 |
| ENSGALG00010013572 | DENND11 | 427917 | -2.1056004 | 1.19E-142 | 1.88E-139 | DENN domain containing 11 |
| ENSGALG00010005001 | GUCA1C | 427960 | -5.0736145 | 2.40E-79 | 1.06E-76 | guanylate cyclase activator 1C |
| ENSGALG00010013595 | WEE2 | 427918 | -4.5164448 | 1.67E-70 | 5.87E-68 | WEE2 Oocyte Meiosis Inhibiting Kinase |
| ENSGALG00010012294 | LOC100857299 | 100857299 | -4.1537095 | 2.60E-73 | 9.81E-71 | ENS-1/ERNI-like |
| ENSGALG00010002947 | LOC121107416 | 121107416 | -4.9425344 | 2.16E-80 | 1.01E-77 | ENS-1/ERNI-like |
| ENSGALG00010006623 | LOC107052071 | 107052071 | -1.3707388 | 7.63E-07 | 5.82E-05 | ncRNA |
| ENSGALG00010013561 | AGK | 418121 | -3.1186859 | 8.71E-237 | 6.91E-233 | mitochondrial acylglycerol kinase |
| ENSGALG00010018537 | LOC112530649 | 112530649 | -4.3531604 | 2.58E-69 | 8.54E-67 | ENS-1/ERNI-like |
| ENSGALG00010009801 | LOC101750251 | 101750251 | -4.7132114 | 1.69E-59 | 4.61E-57 | ENS-1/ERNI-like |
| ENSGALG00010014226 | SLC16A7 | 417815 | -1.5665524 | 1.16E-17 | 1.45E-15 | Monocarboxylate transporter 2 |
| ENSGALG00010005008 | LOC418414 | 418414 | -3.846973 | 8.94E-20 | 1.16E-17 | protein coding |
| ENSGALG00010007504 | TMPRSS15 | 427967 | -5.0062883 | 2.00E-26 | 3.02E-24 | transmembrane serine protease 15 |
| ENSGALG00010015173 | ENS-1 | 769415 | -3.6310882 | 7.97E-75 | 3.24E-72 | ENS-1 |
| ENSGALG00010003372 | ENOX1 | 418839 | -1.9690172 | 1.13E-15 | 1.35E-13 | Ecto-NOX disulfide-thiol exchanger 1 |
| ENSGALG00010001828 | LOC100858869 | 100858869 | -4.7875665 | 7.97E-65 | 2.34E-62 | ENS-1/ERNI-like |
| ENSGALG00010006079 | LOC107055200 | 107055200 | -4.4119861 | 7.85E-72 | 2.83E-69 | ENS-1/ERNI-like |
| ENSGALG00010012306 | LOC121108207 | 121108207 | -5.6390964 | 1.36E-100 | 1.08E-97 | ENS-1/ERNI-like |
| ENSGALG00010002022 | ERNI | 769673 | -4.3793441 | 9.62E-69 | 3.05E-66 | ERNI |

### Supplementally table S13. Sequences of Soprano clone

| Clone | Sequence |
|---|---|
| #1 | TTTTATATCATTGACTCAAAGTTTGCTGAGGAACAAGTCCAGGCAAGTCCTGGGCAAAGGCAGAGAAATCTTTTGTCTTGAGGACACTGATGGACAGGTCCTGGCTAAGGATTGTGAAATCCTCTAAGGAGCACAGATGGAC<br>AAGGCCAGGGGCATCGAGAGAGAGATAAGCTGCTGATAATGGCCGGGAAACGGTCTTTTTGTGTGGACTTATCTCGAGGAAGATGGCCATCTCAGGACGTATGCACAGGACTCTTGCTCAAGCACCCAGGAATGTCACGT<br>AGGCAGGAGAAAATGGAGGATAAAAGAGGGTCCAACAACCACGGCAAGGTAGAGGCTTATCACTCACCACGACAGACTTGAGGGGTTCTCTGCCATCTGATCCTTCACTGCAATGTGGAGACTGATCTCTCATCGCAACAG<br>TGGAGACTGATCTCTACGTGGAGACTGATCTCTCACCGCAACGTGGAGGCTGATCTCTCACCGCAACACAGAGGCTGATCTCTCGATCTCTCGCCACGACACGAGCTTCCTGCCTTCAGATCCTTCTCTACGGACTGTTTG<br>CTGACGGACTTCCCTGGGCCTGCTACCTGAGACCTGCTGCTTCCTCCCCGACCTGCACCCTCTTGTTGCCCAAGACCGGCCTCGCTGCTCCTGCCCTTTGGCCTCGGACCGTCGGAACGTCGTGCAACGGGACTGCTG<br>CCGGATCCTGGTGGTGACTATCCCCGCTTTACGCAATTCTTGCCTCTTTCTATCTTTTCTATCGCTCGCCTTCCCTTCCCCATCACCCCAATTCGTAATAGTGTCCGTTCTCCCCTTTCCCCATCTCCCTTATTAACATTTGTA<br>ATAAACTGGTTGGACCAACATTTGAACCGTTGTTTCTTAATCTCACGCCGGGCATACATATTTCAAAGAACCACCTCTACCTCCTATAAATTGGAGCGAGACA |
| #2 | TTTTATATCATTGACTCAAAGTTTGCTGAGGAACAAGTCCAGGCAAGTCCTGGGCAAAGGCAGAGAAATCTTTTGTCCTTTGATGGACAGGTCCTGGCTAAGGATTGTGAAATCCTCTAAGGAGCACAGATGGACAAGGCCA<br>GGGGCAACGAGAGAGAAGATAAGCTGCTGCTAATGGCTGGGAAACGGTCTTTTGTGTGGACTTATCTTGAGGAAGATGGCCATCTCAGGAGGTATGCCCAGGACTCTTGCTCAAGCACCCAGGAATGTCATGTAGGCAG<br>GAGAAAATGGAGGATAAAAGAGGGTCCAACAACCACGGCGAGGTAGAGGCTTATCTCTCACCACGACAGACTTGAGGGGTTCTCTGCCATCTGATCCTTCACCACAACGTGGAGACTGATCTCTCACCACGACACGAGCTT<br>CCTGCCTTCAGATTCTTCTCTACAGACCGTTTGCTGACGGACTTCCCTGGGCCTGCTACCTGAGACCTGCTGCTTCCTCCCTGACCTGCACCCTCTCGTTGCCCAAGACCGGCCCCGCTGCTCCTGCCCTTCGGCCTCAG<br>ACCGTCGGAACATCGTGTAACGGGACTGCTGCTGGATCCTGGTGGTGACTATCCCTGCTTTACGCAATTCTTGCCTCTTTCTATCTTTTCTATCGCTCGCCTTCCCTTCCCCATCACCCCAATCCTTAATAGTGTCCGTCCT<br>CCCCTTTCCCCATCTCCCTTATTAACATTTGTAATAAACTGGTCGGACCAACATTTGAACCGTTGTTTCTTAATCTCACGCTGGGCATACATATTTCAAAGAACCTCCTCTCCCTCCTATAAATTGGAGCGAGACA |
| #3 | TTTTATATCATTGACTCAAAGTTTGCTGAGGAACAAGTCCAGGGCAAAGGTAGAGAAATCTTCTGTCTGGAGGACACTGATGGACGGGTCCTGGCTAAGGGTTGTGAAATCCTTTTAGGAGCACAGATGGACAAGGCCAGG<br>GGCAAGGAGAGAGGGATAAGCTGCTGATAATGGCCGGGAAACGGTCTTTTGTGTGGACTTATCTCGAGGAAGATGGCCATCCCAGGAGGTATGCACATGTCCATCCCATCACCCCAATCTGTAATAGTGTCCATCCTCCC<br>CTTTCTTCATCTCCATTATTAACATTTGTAATAAACTGGTCAGACCAACATTTGAACCGTTGTTTCTTAATCTCACACCGAGCATACATATTTCTAAGAACCTCCTCTCCCTCCTATAAATTGGAGCGAGACA |
| #4 | TTTTATATCATTGACTCAAAGTTTGCTGAGGAACAAGTCCAGGCAAGTCCTGGGCAAAGGCAGAGAAATCTTCTGTCTGGAGGACACTGAGGGACAGGTCCCAGCTAAGGTTGTGAAATCCTTTAAGGAGCACAGATGTACA<br>AGGTCGGGGCAATGAGAGACGGATAAGCTGCTGATAATGGCCGGGATTGGACTTTTGTGTGGACTTATCTCGAGGAAGATGGCCATCTCAGGAGGTACGCACATGACTCTTGCTCAAGCACCCAGGGATGTCGTGTAGG<br>CAGGAAAAAAAGGAGGATAAAAGAGGGTCCAACAACCATGAGGAGAGAGGGTTTCTGCCATCAGACCCCTCACCACGACGGACTAAGGGGTTTTCTGCCATCTGATCTCTCACCATGACGGACTCGAGGTTTCTGCCATCA<br>GATCCTTCACTACGGACCGTATGCTGACAGAGTTCCCTGGTCATGCTACCTGAGACCTGCTGCTTCCTCCCAGACTTACTCTGTGGCTTCTTCCGGGACCCACCCAGTACCTGCACCCTCTCGTTGCCCAAGACCGGCCT<br>CGCTGCTCCTGCCCTTCGGCTTCGGACCATCAGAACGTCGTGTAACGGGACTACTGTCGCATCTTGGTGGTGACTATCCCCGCTTTACACAATTCTTGCTTCTTTCTATCTTTTCTATTGCTCGCCTTCCCTTCCCCATCAC<br>CCCAATCCGTAATAGTGTCCATCCTCCCCTTTCTTCATCTCCCTTGTTAACATTTGTAATAAACTGGTTGGACCAACATTTGAACTGTTGTTTCTTAATCTCACTCCGGGCATACATACTTCAAAGAACCTCCTCTCCCTCCTAT<br>AAATTGGAGCGAGACA |
| #5 | TTTTATATCATTGACTCAAAGTTTGCTGAGGAACAAGTCCAGGCAAGTCCTGGGCAAAGGTAGAGAAATCTTCTGTCTGGAAGACAGTGATGGACAGGTCCTGGCTAAGGGTTGTGAAATCCTTTAAGGAGCACAGATGGA<br>CAAGGCCAGGGGCAACAAGAGAGGGATAAGCTGCTGATAATGGCCGGGAAATGGTCTTTTGTGTGGACTTAGCTCGAGGAAGATGGCCATCTCAGGAGGTACGCACATGACTCTTGCTCAAGCACCCAGGGATGTCATG<br>TAGGCAGGAAAAAAGGAGGATAAAAGAGGGTCCAACAACCATGAGGAGAGGGTTTCTCCATCAGATCCCTCACCACAATGGACTCGAGGTGTCTGCCATCAGATCCCTCACTACAGGCTGCATGCTGACGGACTTCCCTG<br>GGCCTGCTGCTTCCTCCCAGACTTACTCTGCGGATTCTTCCTGGACCCACCCGGTACCTGCACCCTCTCATTGCCCAAGACTGGCCTCGCTGCTCCTGCCCTTCAGCCTTGGACCATCGGAACGTTGTGCAACGGGACTA<br>CTGCCGGATCCTGGTGGTGACTATCCCCGCTTTACGCAATTCTTGCTTCTTTCTATCTTTTCTATTGCTCTCCTTCCCTTCCCCATCACCCCAGTCCGTAATAGTGTCCATCCTCCACCTTCTTCATCTCCCTTATTAACATTT<br>GTAATAAACTGGTCGGACCAACATTTGAACCGTCGTTTCTTAATCTCACGCTGGGCATACATATTTCAAAGAACCTCCTCTCCCTCCTATAAATTGGAGCGAGACA |
| #6 | TTTTATATCATTGACTCAAAGTTTGCTGAGGAACAAGTCCAGGCAAGTCCTGGGCAAAGGCAGAGAAATCTCTTGTCTTGAGGACACTGATGGACAGGTCCTGGCTAAGGATTGTGATATCCTCTAAGGAGCACAGATGGAC<br>AAGGCCAGGGGCATCGAGAGAGAGATAAGCTGCTGCTAATGGCCGGGAAGCAGTCTTTTTGTGTGGACTTATCTCAAGGAAAATGGCCATCTCAGGAGGTATGCACAGGACTCTTGCTCAAGCACCCAGGAATGTCACGT<br>AGGCAGCAGAAAATGGAGGATAAAAGAGGGTCCAACAACCACGGCATCGTAGAGGCTTATCTCTCACCACGACAGTCTTGAGGGACCGTCGGAACGTCGTGCAACGGGACTGCTGCCGGATCCTGGTGGTGACTAACCC<br>CGCTTTACGCAATTCTTGCCTCTTCCTATCTTTTCTATCTCCCACCTTCCCTTTCCCATCACCCCAATCCTTAACAGTGTCCATCCTCCCCTTTCCCCATCTCCCTTATTAACATTTGTAATAAACTGGTCAGACCAACATTTGAA<br>CCGTTGTTTCTTAATCTCACGCCGGGCATACATATTTCAAAGAACCTCCTCTCCCTCCTATAAATTGGAGCGAGACA |
| #7 | TTTTATATCATTGACTCAAAGTTTGCTGAGGAACAAGTCCAGGCAAGTCCTGGGCAAAGGCAGAGGAATCTTTTGTCTTGAGGACACTGATGGACAGGTCCTGGCTAAGGATTGTGAAATCCTTTAAGGAGCACAGATGGAC<br>AAGGCCAGGGGCATCGAGAGAGAGATAAGCTGCCGCTAATGGCCGGGAAACGGTCTTTTTGTGTGGACTTATCTCAAGGAAAATGGCCATCTCAGGAGGTATGCACAGGACTCTTGCTCAAGCCCCCAGGAATGTCACGT<br>AGGCAGCAGAAAATGGAGGATAAAAGAGGGTCCAATAACCACAACGGTGGAAGCTGATCCTTCACCACAACCACGGCAACGGGGGAGGCTTATCTCTCACCACGACAGACTTGAGGGGTTCTCTGCCATCTGATCCTTCAC<br>CGCAACAGTGGAGACTGATCTCTACGTGTAGACGGATCGCTACGTGGAGACTGATCTCTCACCACGACACGAGCTTCCTGCCTTCCGATCCTCCTCTACGGACCGTTTGCTGACGGACTTCCCTGGGCCTGCTACCTGAG<br>ACCTGCTGCTTCCTCCCTGGCCTGCATCCTCTCGCTGCCCCAGACCGGCCTCACTGCTCCTGCCCTTCGGCCTCGGACCGTCGGAACGTCGTGCAACGGGACTGCTGCCGGATCCTGGTGGTGACTATCCCCGCTTTA<br>CGCAATTCTTGCCTCTTTCTATCTTTTCTATCGCTCGCCTTCCCTTCCCCATCACCCCAATCCTTAATAGCGTCCGTCCTCCCCTTTCCCCATCTCTCTTATTAACATTTATAATAAACTGGTCGGACCAACATTTGAACCGCTG<br>TTTCTTAATCTCACGCCGGGCATACATATTTCTAAGAACCTCCTCTCCCTCCTATAAATTGGAGCGAGACA |
| #8 | TTTTATATCATTGACTCAAAGTTTGCTGAGGAACAAGTCCAGGCAAGTCCTGGGCAAAGGCAGAGAAATCTTTTGTCTTGAGGACACTGATGGACAGGTCCTGGCTAAGGATTGTGAAATCCTCTAAGGAGCACAGATGGAC<br>AAGGCCAGGGGCATCGAGAGAGAGATAAGCTGCCGCTAATGGCCGGTAAACGGTCTTTTTGTGTGGACTTATCTCAAGGAAAATGGCCATCTCAGGAGGTATGCACAGGACTCTTGCTCAAGCACCCAGGAATGTCACGT<br>AGGCAGCAGAAAATGGAGGATAAAAGAGGGTCCAATAACCACAACGGTGGAAGCTGATCCTTCACCACAACCACGGCAACGGGAGAGGCTTATCTCTCACCACGACAGACTTGAGGGGTTCTCTGCCAGCTGATCCTTCAC<br>TGCAACGTGGAGACGGATCTCTACGTGGAGACTGATCTCTCACCACGACACGAGCTTCCTGCCTTCCAATCCTCCTCTACGGACCGTTTGCTGACGGACTTCCCTGGGCCTGCTACCTGAGACCTGCTGCTTCCTCCCCG<br>ACCTGCACCCTCTCGTTGCCCAAGACCGGCCTCGCTGCTCCTGCCCTTCGGCCTCGGACTGTCGGAACATCGTGCAACAGGACTGCTGCCGGATCCTGGTGGTGACTATCCCCGCTTTACGCAATTCTTGCCTCTTTCTA<br>TCTTTTCTATCGCTCGCCTCCCCTTCCCCATCACCCCAATCCTTAATAGCGTCCATCCTCCCCTTTCCCCATCTCCCTTATTAACATTTGTAATAAACTGGTCGGACCAACATTTGAACCGTTGCTTCTTAATCTCACGCCGGG<br>CATACATATTTCAAAGAACCTCTTCTCCCTCCTATAAATTGGAGCGAGACA |
| #9 | TTTTATATCATTGACTCAAAGTTTGCTGAGGAACAAGTCCAGGCAAGTCCTGGGCAAAGGCAGAGAAATCTTTTGTCTTGAGGACACTGATGGACAGGTCCTGGCTAAGGATTGTGATATCCTCTAAGGAGCACAGGTGGA<br>CAAGGCCAGGGACAACGAGAGAGAGATAAGCTGCTGCTAATGGCCGGGAAACGGTCTTTTTGTGTGGACTTATCTCGAGGAAGATGGCCATCTCAGGAGGTATGCACAGGACTCTTGCTTAAGCACCCAGGAATGTCACG<br>TAGGTAGGAGAAAATGCAGGATAAAAGAGGGTCCAACAACCACGGCTATGTGGAGACTGATCTCTCACTGCAACGTGGAGGCTGATCTCTCACCGCAACACGGGGGCTGATTGATCTCTCACCACGACACGAGCTTTCTG<br>CCTTCAGATCCTTCTCTACGGACCGTTTGCTGACGGACTTCCCTGGGCCTGCTACCTGAGACCTGCTGCTTCCTCCCTGACCTGCACCCTCTCGCTGCCCAAGACTGGCCTCGCTGCTCCTGACTGTCGGAACATCGTGC<br>AACGGGACTGCTGCCGGATCCTGGTGGTGACTATCCCCGCTTTACGCAATTCTTGCCTCTTTCTATCTTTTCTATCGCTCACCTTCCCTTCCCCATCACCCCAATCCTTAATAGTGTCCGTCCTCCCCTTTCCCCATCTCCTT<br>TATTAACATTTGTAATAAACTGGTCGGACCAACATTTGAACCGTTGTTTCTTAATCTCACGCCGGGCATACATATTTCAAAGACCCTCCTCTCCCTCCTATAAATTGGAGCGAGACA |
| #10 | TTTTATATCATTGACTCAAAGTTTGCTGAGGAACAAGTCCAGGCAAGTCCTTGGCAAAGGTAGAGAAATCTGCTGTCTGGAGGACACTGATGGACAGGTCCTGGCTAAGGGTTGTGAAATCCTTTAAGGAGCACAGATGGA<br>CAAGGCCAGGGGCAACAAGAGAGGGATAAGCTGCTGATAATGGCCGGGAAATGGTCTTTTGTGTGGACTTGTCTCGAGGAAGATGGCCATCTCAGGAGGTACGCACATGACTCTTGCTCAAGCACCCAGGGATGTCATG<br>TAGACAGGAAAAAAGGAGGATAAAAGAGGGTCCAACAACCATGAGGATAGGTTTCTGCCATCAGATCCCTCCCCACAACAGACGCAAGGTTTTTGCCATCTGATCCTTCACCTCAAAAGGCCTGCATGCTGACAGACTTCCC<br>TGGGCCTGCTACCTGAGACCTGCTGTTTCCTCCCAGGCTTACTCTGCGGCTTCTTCCTGGACCCACCCGGTACCTGCACCCTCTCGTTGCCCAAGGCCAGCCTCGCTGCTCCTGCCCTTCAGCCTCGGACCATTGGAAC<br>ATCGTGCAACAGGACTACTGCCGGATCCTGGTGGTGACTATCCTCACTTTACGCAGTTCTTGCTTCTTTCTAGCTTTTCTATCACTCGCCTTCCCTTCCCCATCACCCCAATCCATAATAGTGTCCATCCTCCCCTTTCTTCAT<br>CTCCCTTATTAACATTTGTAATAAACTGGTCGGACCAACATTTGAACCGTTGTTTCTTAATCTCACGCCGGGCATACATATTTCAAAGAACCTCCTCTCCCTCCTATAAATTGGAGCGAGACA |
| #11 | TTTTATATCATTGACTCAAAGTTTGCTGAGGAACAAGTCCAGGCAAGTCCTGGGCAAAGGCAGAGAAATCTTTTGTCTTGAGGACACTGATGGACAGGTCCTGGCTAAGGATTGTGAAATCCTTTAAGGAGCACAGATGGAC<br>AAGGCCAGGGGCATCGAGAGAGAGATAAGCTGCCGCTAATGGCCGGGAAACGGTCTTTTTGTGTGGACTTATCTCAAGGAAAATGGCCATCTCAGGAGGTATGCACAGGACTCTTGCTCAAGCCCCCAGGAATGTCACGT<br>AGGCAGCAGAAAATGGAGGATAAAAGAGGGTCCAATAACCACAACGGTGGAAGCTGATCCTTCACCACAACCACGGCAACGGGAGAGGCTTATCTCTCACCACGACAGACTTGAGGGGTTCTCTCTGCCAACTGATCTCTC<br>ACCGCAACGTGTAGACGGATCTCTACGTGGAGACTGATCTCTCACCACGACACGAGCTTCCTGCCTTCCAACCCTCCTCTACGGACCGTTTGCTGACGGACTTCCCTGGGCCTGCTACCTGAGACCTGCTGCTTCCTCCC<br>CGACCTGCACCCTCTCGTTGCCCAAGACCGGCCTCGCTGCTCCTGCCCTTCGGCCTCGGACCGTCGGAACATCGTGCAACGGGACTGCTGCCGGATCCTGGTGGTGACTATCCCCGCTTTACGCAATTCTTGCCTCTTT<br>CTATCTTTTCTATCGCTCGCCTTCCCTTCCCCATCACCCCAATCCTTAATAGCGTCCGTCCTCCCCTTTCCCCATCTCCCTTATTAACATTTGTAATAAACTGGTCGGACCAACATTTGAACCGCTGTTTCTTAATCTCACGCC<br>GGGCATACATATTTCAAAGAACCTCCTCTCCCTCCTATAAATTGGAGCGAGACA |
| #12 | TTTTATATCATTGACTCAAAGTTTGCTGAGGAACAAGTCCAGGCAAGTCCTGGGCAAAGGTAGAGAAATCTTCTGTCTGGAGGACACTGATGGAGAGGTCCTGGCTAAGGGTTGTGAAATCCTTTTAGGAGCACAGATGGA<br>CAAGACCAGGGGTAACGAGAGAGGGATAAGCTGCTGATAATGGCCGGGAAATGGTCTTTTGTGTGGACTTATCTCGAGGAAGATGGCCATCTCAGGAGGTACGCACATGACTCTTGCTCAAGCACCCAGGGATGTCACGT<br>AGGCAGGAAAAAAGGAGGACAAAAGAGAGTCCAACAACCATGAGGATAGGTTTCTGCCATCAGATCCCTCACCACGACGGACTCGAGGTTTCTGCCTTCAGATCCCTCACTACGGACCGTATGCTGACGGACTTCCCTGGT<br>CCTGCTACCTGAGGCCTGCTGCTTCCTCCGGGCTTACTCTGAGGCTTCTTCCTGGACCCACCCGGTACCTGCACCCTCTCATTGCCCAAGACCAGCCTCGCTGCTCCTGCCCTTCGGCCTCAGACCATCGGAACGGCAT<br>GCAACGGGACTACTGCCAGATCCTGGTGGTGACTATCCCCGCTTTACGCAATTCTTGCTTCTTTCTATCTTTTCAATCACTCACCTTCCCTTCCCCATCACTCCAATCCGTAATAGTGTCCGTCCTCCCTTTCTTCATCTCCCT<br>TATTAACATTTGTAATAAACTGGTTGGACCAACATTTGAATCATTGTTTCTTAATCTCATGCGAGGCATGCATATTTCAAAGAACCTCCTCTCCCTCCTATAAATTGGAGCGAGACA |
| #13 | TTTTATATCATTGACTCAAAGTTTGCTGAGGAACAAGTCCAGGCAAGTCCTGGACAAAGGTAGAGAAATCTTCTGTCTGGAGGACACTGATGGACAGGTCCTGGCTAAGCGTAGTGAAATCCTTTTAGGAGCACAGATGGAC<br>AAGGCCAGGGGCAACGAGAGAGGGATAAGCTGCTGATAATGGCCGGGAAACGGTCTTTTGTGTGGACTTATCTCAAGGAAGATGGCCATCTCAGGAGGTACACGCATGACACTTGCTCAAGCACCCAGGGATGTCACAT<br>GGGCAGGAAAAAGGAGGATACAATAGGGTCCAACAACAATGAGGAAAGCTTTCTGCCAGCAGATCCTTCACCACGAAAGGCTTGCATGCTGATGGACTCGAGGTTTCTGCCATCAGATCCTTCACCACGAAAGTCCTGCA<br>TGCTGACAGACTCAAGGTTTCTGCTATCAGATCCCTCACCACGAAAGGCCTGCATGCTGACGGACTCGAGGTGTCTACCATCAGATCCCTCACAACGGACCGTATGCTGACGGACTTCCCTGGGCCTGCTACCTGAGACC<br>TGCTGCTTCCTCCCAGACTTACTCTGCGACTTCTTCCTGGACCCACCCGGTACCTGCACCCTCTCGTTGCCCAAGACCGGCCTCACTGCTCCTGCCCTTCGACCATCAGAACATCGTGCAACGGGACTACTGCCGGATCC<br>TGGTGGTGACTATCCCCGCTTTATGCAATTCTTGCTTCTTTCTATCTTTTCTATCGGTCACCTTCCCTTCCCCATCACCCCAACCCATAATAGTGTCCGTCCTCCCCTTTCTTCATCTCCCTTATTAACATTTGTAATAAACTGG<br>TCGGACCAAAATTTGAACTGTTGTTTCTTAATCTCACTCTGGGCATACATATTTCAAAGAACCTCCTCTCCCTCCTATAAATTGGAGCGAGACA |
| #14 | TTTTATATCATTGACTCAAAGTTTGCTGAGGAACAAGTCCAGGCAAGTCCTGGGCAAAGGTAGAGAAATCTTTTGTCTTGAGGACACCGATGGACAGGTCCTGGCTAAGGATTGTGAAATCCTCTAAGGAGCACAGATGGAC<br>AAGGCCAGGGGCAACGAGAGAGAGATAAGCTGCTGCTAATGGCCGGGAAACGGTCTTTTTGTGTGGACTTATCTCGAGGAAGATGGCCATCTCAGGAGGTATGCACATGACTCTTGCTCAAGCACTCAGGGATGTCACGT<br>AGGCAGGAGAAAATGGAGGATAAAAGAGGGTCCAACAACCACGGCGAGGTAGAGGCTGATCTCTCACCGCAACCGAGAGGCTGATCTCTCATGATCTCTCACCACGATTCGAGCTTTCTGTCTTCAGATCCTTCTCTACGG<br>ACCGTTTGCTGACGGACTTCCCTGGGCCTGCTACCTGAGACCTGCTGCTTCCTCCCTGACTTACTCGGCGGCTTCTTCCTGGACCCACCCGGTACCTGCACTCTCTCTCGTTGCCCAAGACCGGCCTCGCTGCTCCTGC<br>CCTTCGGCCTCGGACCGTCGGAACGTCGTGCAACGGGACTGCTGCCGGATCCTGGTGGTGACTATCCCTGCTTTACGCAATTCTTGCCTCTTTCTATCTTTTCTATCGCTCGCCTTCCCTTCCCCATCACCCCAATCCTTA<br>ATAGTGTCCGTCCTCCCCTTTCTTCATCTCCCTTATTAACATTTGTAATAAACTGGTCGGACCAACATTTGAACCGTTGTTTCTTAATCTCACGCCGGGCATACATATTTCAAAGAACCTCCTCTCCCTCCTATAAATTGGAGCG<br>AGATA |

#15

TTTTATATCATTGACTCAAAGTTTGCTGAGGAACAAGTCCAGGCAAGTCCTGGGCAAACGTAGAGAAATCTTCTGTCTGGAGGACACTGATGGACAGGTCCTGGCTAAGGGTTGTGAAATCCTTTAAGGAGCACAGATGGA
CAAGGCCAGGGGCAAGAAGAGAGTGATAAGCTGCTGATAATGGCTGGGAAACGGTCTTTTGTGTGGACTTATCTTGAGGAAGATGGCCATCTCAGGAGGTATGCACATGACTCTTGCTCAAGCACCCAGGGATGTCAGGT
AGGCAAGAAAAAAAGGAGGATAAAAGAGGGTCCAACAATCATGAGGAGGGGTTTCTGCCATCAGTTCCCTCTCCATGACGGACTCGAGGTTACTGCCTTCAGATCCCTCACTACGGACTGTATGCTGACAGACTTCCCTGG
GCCTGCTACCTGAGACCTGCTGCTTCCTCCCGGGCTTACTCTGCGGCTTCTTCCTGAACCCACCCAGTAACTGCACCCTCTTGTTGCCCAAGACCGGCCTCGCTGCTCCTGCCCTTCGGCCTCGAACCATCAGAACGTCA
TGCAATGGGACTACTGCCGGATCCTGGTGGTGACTATCCCTACTTTATGCATTTCTTGCTTCTTTCTATCTTTTCTATCGCTCGCCTTCCCTTCCCCATCACCCCAATCCAAAATAGTGTCCGTCTTCCCCTTTCTTCATTTCC
CTTATTAACATTTGTAATAAACTGGTAGAACCAACATTTGAACCGTTGTTTCTTAATCTCATGCCGGGCATACATATTTCAAAGAACCTCCTCTCCCTCCTATAAATTGGAGCGAGACA

#16

TTTTATATCATTGACTCAAAGTTGCTGAGAACAAGTCCAGGCAAGTCCTGGGCAAAGGCAGAGAAATCTTTTGTCTTGAGGACACTGATGGACAGGTCCTGGCTAAGGATTGTGAAATCCTTCAAGGAGCACAGATGGACAA
GGCCAGAGGCATCGAGAGAGAGAGATAAGCTGCCGCTAATGGCCGGGAAACGGTCTTTTTGTGTGGACTTATCTCAAGGAAAATGGCCATCTCAGGAGGTATGCACAGGACTCTTGCTCAAGCACCCAGGAATGTCACGT
AGGCAGCAGAAGATGGAGGATAAAAGAGGGTCCAATAACCACAACGGTGAAAGCTGATCCTTCACCACAACCACGGCAACGGTAGAGGCTTATCTCTCACCACGACAGACTTGAGGGGTTCTCTGCCAACTGATCTCTCAC
CGCAACGTGTAGACGGATCTCTACGTGGAGACTGATCTCTCACCACGACACGAGCTTCCTGCCTTCCGATCCTCCTCTACGGACCGTTTGCTGACGGACTTCCCTGGGCCTGCTACCTGAGACCTGCTGCTTCCTCCCTG
ACCTGCATCCTCTCGCTGCCCCAGACCGGCCTCGCTGCTCCTGCCCTTCGGCCTCGGACCGTCGGAACGTCGTGCAACGGGACTGCTGCCGGATCCTGGTGGTGACTATCCCCGCTTTACGCAATTCTTGCCTCTTTCT
ATCTTTTCTATCGCTCGCCTTCCCTTCCCCATCACCCCAATCCTTAATAGCGTCCGTCCTCCCCTTTCCCCATCTCCCTTATTAACATTTGTAATAAACTGGTCGGACCAACATTTGAACCGCTGTTTCTTAATCTCACGCCGG
GCATACATATTTCTAAGAACCTCCTCTCCCTCCTATAAATTGGAGCGAGACA

#17

TTTTATATCATTGACTCAAAGTTTGCTGAGGAACAAGTCCAGGCAAGTCCGGGGCAAAGGCAGAGAAATCTTTTGTCTTGAGGACACTGATGGACAGGTCCTGGCTAAGGATTGTGAAATCCTCTAAGGAGCACAGATGGA
CAAGGCCAGGGGCATCGAGAGAGAGATAAGCTGCTGCTAATGGCCGGGAAGCGGTCTTTTTGTGTGGACTTATCTCAAGGAAAATGGCCATCTCAGGAGGTATGCACAGGACTCTTGCTCAAGCACCCAGGAATGTCACG
TAGACAGCAGAAAATGGAGAATAAAAGAGGGTCCAATAACCACAACGCTGGAAGCTGATCCTTCACCACAACCACGGCAACGGGAGAGGCTTATCTCTCACCACGACAGACTTGAGGGGTTCTCTGCCAACTGATCTCTCAT
CGCAACAGTGGAGACTGATCTCTCACCGCAACGTGTAGACGGATCTCTACGTGGAGACTGATCTCTCACCACGGCACGAGCTTCCTGCCTTCCGATCCTCCTCTACGGACCGTTTGCTGACGGACTTCCCTGGGCCTGCT
ACCTGAGACCTGCTGCTTCCTCCCTGACCTGCACCCTCTCGCTGCCCCAGACCGGCCTCGCTGCTCCTGCCCTTCGGCCTCGGACCGTCGGAACATCGTGCAACGGGACTGCTGCCGGATCCTGGTGGTGACTATCCC
CGCTTTACGCAATTCTTGCCTCTTTCTATCTTTTCTATCGCTCGCCTTCCCTTCCCCATCACCCCAATCCTTAATAGCGTCCGTCCTCCCCTTTCCCCATCTCCCTTATTAACATTTGTAATAAACTGGTCGGACCAACATTTGA
ACCGCTGTTTCTTAATCTCACGCCGGGCATACATATTTCAAAGAACCTCATCTCCCTCCTATAAATTGGAGCGAGACA

#18

TTTTATATCATTGACTCAAAGTTTGCTGAGGACAAGTCCAGGCAAGTCCTGGCAAAGGCAGAGAAATCTTTTGTCTTGAGGACACTGATGGACAGGTCCTGGCTAAGGATTGTGAAATCCTTTAAGGAGCACAGATGGACAA
GGCCAGGGGCATCGAGAGAGAGATAAGCTGCCGCTAATGGCCGGGAAACGGTCTTTTTGTGTGGACTTATCTCAAGGAAAATGGCCATCTCAGGAGGTATGCACAGGACTCTTGCTCAAGCCCCCAGGAATGTCACGTA
GGCAGCAGAAAATGGAGGATAAAAGAGGGTCCAATAACCACAACGGTGGAAGCTGATCCTTCACCACAACCACGGCAACGGGAGAGGCTTATCTCTCACCACGACAGACTTGAGGGGTTCTCTCTGCCAACTGATCTCTCA
CCGCAACGTGGAGACGGATCTCTACGTGGAGACTGATCTCTCACCACGACACGAGCTTCCTGCCTTCCGATCCTCCTCTACGGACCGTTTGCCGACGGACTTCCCTGGGCCTGCTACCTGAGACCTGCTGCTTCCTCCC
CGACCTGCATCCTCTCGCTGCCCAAGACCAGCCTCGCTGCTCCTGCCCTTCGGCCTCGGACCGTCGGAACGTCGTGCAACGGGACGTCCCTGCTGCCGGATCCTGGTGGTGACTATCCCCGCTTTACGCAATTCTTGC
CTCTTTCTATCTTTTCTATCGCTCGCCTTCCCTTTCCCATCACCCCAATCCTTAATAGCGTCCGTCCTCCCCTTTCCCCATCTCCCTTATTAACATTTGTAATAAACTGGTCGGACCAACATTTGAACCGTTGTTTCTTAATCTC
ACGCCGGGCATACATATTTCAAAGAACCTCCTCTCCCTCCTATAAATTGGAGCGAGACA

#19

TTTTATATCATTGACTCAAAGTTTGCTGAGGAACAAGTCCAGGCAAGTCCTGGGCAAAGGCAGAGAAATCTTTTGTCTTGAGGACACTGATGGACAGGTCCTGGCTAAGGATTGTGAAATCCTCTAAGATGGACAAGGCCAG
GGGCAACGAGAGAGAGAGATAAGCTGCTGATAATGGCCGGGAAACGGTCTTTTTGTGTGGACTTATCTCGAGGAAGATGGCCATCTCAGGAGGTATGCACATGACTCTTGCTCAAGCACCCAGGGATGTCACGTAGGCAGG
AAAAAAAGGAGGATAAAAGAGGGTCCAACAACCATGAGGAGAGGGTTTCTGCCATCAGATCCCTCACTACAGACCGTATGCTGACGGACTTCCCTGGGCCTGCTGCCTGAGACCTGCTGCTTCCTCCCAGACTTACTCTGC
GGCTTCTTCCTGGACCCTCCTGGTACCTGCACCCTCTCGTTGCCCAAGACCGGCCTCGCTGCTCCTGCCCTTCGGCCTCGGACCATCGGAACGTTGTGCAACAGGACTACTGCCGGATCCTGGTGGTGACTATCCCCGC
TTTACGCAATTCTTGCTTCTTTCTATCTTTTCTATCGCTCGCCTTCCCTTCCCCATCACCCCAATTTATAATAGTGTCCATCCTCCCCTTTCTTCATCTCCCTTATTAACATTTGTAATAAATTGGTCAGACCAACATTTGAACTG
TTGTTTCTTAATCTCACGCCGGGCATACATATTTCAAAGAACCTCCTCTCCCTCCTATAAATTGGAGCGAGACA

#20

TTTTATATCATTGACTCAAAGTTTGCTGAGGAACAAGTCCAGGCAAGTCCTGGGCAAAGGCAGAGAAATCTTTTGTCTTGAGGACACTGATGGACAGGTCCTGGCTAAGGATTGTGAAATCCTTTAAGGAGCACAGATGGAC
AAGGCCAGGGGCATCGAGAGAGAGATAAGCTGCCGCTAATGGCCGGGAAACGGTCTTTTTGTGTGGACTTATCTCAAGGAAAATGGCCATCTCAGGAGGTATGCACAGGACTCTTGCTCAAGCCCCCAGGAATGTCACGT
AGGCAGCAGAAAATGGAGGATAAAAGAGGGTCCAATAACCACAACGGTGGAAGCTGATCCTTCACCACAACCACGGCAACGGGAGAGGCTTATCTCTCACCACGACAGACTTGAGGGGTTCTCTGCCAACTGATCTCTCAC
CGCAATGGGTAGACGGATCTCTACGTGGAGACTGATCTCTCACCACGACACGAGCTTCCTGCCTTCCGATCCTCCTCTACGGACCGTTTGCTGACGGACTTCCCTGGGCCTGCTACCTGAGACCTGCTGCTTCCTCCCTG
ACCTGCACCCTCTCGTTGCCCAAGACCGGCCTCGCTGCTCCTGCCCTTCGGCCTCGGACCGTCGGAACATCGTGCAACGGGACTGCTGCCGGATCCTGGTGGTGACTATCCCCGCTTTACGCAATTCTTGCCTCTTTCT
ATCTTTTCTATCGCTCGCCTTCCCTTCCCCATCACCCCAATCCTTAATAGCGTCCGTCCTCCCCTTTCCCCATCTCCCTTATTAACATTTGTAATAAACTGGTCGGACCAACATTTGAACCGCTGTTTCTTAATCTCACACCGG
GCATACATATTTCTAAGAACCTCCTCTCCCTCCTATAAATTGGAGCGAGACA

#21

TTTTATATCATTGACTCAAAGTTTGCTGAGGAACAAGTCCAGGCAAGTCCTGGGCAAAGGTAGAGAAATCTTCTGTCTTGAGGACACTGATGGACAGGTCATGGCTAAGGGTTGTGAAATCACTTAAGGAGCACGGATGGA
CAAGGCCAGGGGCAATGAGAGAGGGATAAGCTGCTGATCATGGCCGGGAAATGGTCTTTTGTGTGGACTTATCTCAAAGAAGATGGCCATCTCAGGAGGTACGCATGTGACTCTTGCTCAAGCACGCAGGGATGTCACG
TAGGCAGGAAAAAAGGAGGATAAAAGAGGGTCCAACAACCATGAGGATAGGATTTCTGCCATCAGATCCCTCACCACGACAGACTCAAGGTTTCTGCCATCAGATCCCTCACCACGACAGACTTGAGGTGTCTGCCTTCAGA
TCCCTCACTACGGACCGTATGTTGATGGACTTCCCTGGGCCTGCTACCTGAGACCTGCTGCTTCCTCCTGGGCTTACTCTGCGGCTTCTTCCTGGACCCACCCAGTACCTGTACCCTCTCATTGCCCAAGACTGGCCTCG
CTGCTCCTGCCCTTAGGCCTCGGACCATCGGAACGGCGTGCAACAGGACTACTGCCGGATCCTGGTGGTGACTATCCCTGCTTTATGCAATTCGTGCTTCTTTTTATCTTTTCTATCACTCACCTTCCCTTCCCCATCACCC
CAATCCTTAATAGTGTCCGTTCTCCCCTTTCTTCATCTCCCTTATTAACATTTGTAATAAACTGGTCGGACCAACATTTGAATCGTTGTTTCTTAATCTCATGCCAGGCATACATATTTCAAAGAACCTCCTCTCCCTCCTATAAA
TTGGAGCGAGAC

#22

TTTTATATCATTGACTCAAAGTTTGCTGAGGAACAAGTCCAGGCAAGTCCTGGGCAAAGGCAGAGAAATCTTTTGTCTTGAGGACACTGATGGACAGGTCCTGGCTAAGGATTGTGATATCCTCTAAGGAGCACAGATGGAC
AAGGCCAGGGGCAACGAGAGAGAGATAAGCTGCTGCTAATGGCCGGGAAACGGTCTTTTTGTGTGGACTTATCTCGAGGAAGATGGCCATCTCAGGAGGTATGCACAGGACTCTTGCTCAAGCACCCAGGAATGTCATGT
AGGCAGGAGAAAATGGAGGATAAAAGAGGGTCCAACAACCACGGCAACAGTAGAGGCTTATCTCTCACCACGACAGACTTGAGGGGTTCTCTGCCATCTGATCCTTCACCACAACCGAGAGGCTGATCTCTCATGATCTCT
CACCACGACACGAGCTTTCTGCCTTCAGATCCTTCTCTACGGACCGTTTGCTGACGGACTTCCCTGGGCCTGCTACCTGAGACCTGCTGCTTCCTCCCTGACCTGCACCCTCTCGTTGCCCAAGACCAGCCTCGCTGCTC
CTGCCCTTCGGCCTCGGACCATCGGAACGTCGTGCAACGGGACCACTGCCGGATCCTGGTGGTGACTATCCCCGCTTTACGCAATTCTTGCCTCTTTCTATCTTTTCTATCGCTCGCCTTCCCTTCCCCATCACCCCAATC
CTTAATAGTGTCCGTCCTCCCCTTTCCCCATCTCCCTTATTAACATTTGTAATAAACTGGTCGGACCACCATTTGAACCATTGTTTCTTAATCTCACGCCGGGCATACATATTTCAAAGAACCTCCTCTCCCTCCTATAAATTGG
AGCGAGACA

#23

TTTTATATCATTGACTCAAAGTTTGCTGAGGAACAAGTCCAGGCAAGTCCTGGGCAAAGGCAGAGAAATCTTCTGTCTGGAGGACACTGATGGACATGTCCTGGCTAAGGGTTGTGAAATCCTTTAGAAGCACAGATGGACA
AGGCCAGGGGCAACGAGAGAGGGATAAGCTGCTGATAATGGCTGGGAAACGGTCTTTTGTGTGGACTTATCTCGAGGAAGATGGCCTTCTCAGGAAGTACGCACATGACTCTTGCTCAAGCACCCAGGGATGTCACGTA
GGCAGGAAAAAAAGGAGGATAAAAGAGGGTCCAACAACCATGAAGATAGGTGTCTGCCATCAGATCTCTCACTACGGACCGCATGCTGACGGACTTCCCTGGGCCTGCTACCTGAGACCTCCTGCTTCCTCTCGGACTCA
CTCTGCGGCTTCTTCCTGGACCCACCTGGTACCTGCACCCTCTCGTTGCCCAAGACCATCCTCACTGCTCCTGCCCTTTGGCTGTGGACCATTGAAACGGCGTGCAACAGACTCCTGCTGGATCTTGGTGGTGACTTATC
CCTGCTTTAAACAATTCTTGCTTCTTTGCTATTGCCTTCCTTCCCTCCCCCTTTACCCTAACTCATAATAGCGTCCATCCTCCCCTTTCTTCATCTCCCTTATTAACATTTGTAATAGACTGGTCAGACCAACATTTGAACCGTT
GTTTCTTAATCTCAGGCCTGGTATACATACATCAAAGAACCTCTTCTCCCTCCTATAAATTGGAGCGAGACA

#24

TTTTATATCATTGACTCAAAGTTTGCTGAGGAACAAGTCCAGGCAAGTCCTGGGCAAAGGTAGAGAAATCTTCTGTCTGGAGGACACTGATGGACAGGTCCTGGCTAAGGGTTGTGAAATCCTTTAAGGAGCACAGATGGA
CAAGGCCAGGGGCAATGAGAGAGAGATAAGCTGCTGCTAATGGCCGGGAAATGGTCTTTTGTGTGGACTTGAGGAAGATGGCCATCTCAGGAGGCACGCACATGACTCTTGCTCAAGCACCCAGGGATGTCACGTAGGC
AGGAAAAAAAAGAGGATAAAAGAGGGTCCAACAACCAAGAGAGTAGGGGGGTTTCTGCCATCAGATCCCTCACCATGACAGACTCGAGGTTTCTGCCATCAGGTCGTTCACTACGGACCATATACTGACGGACTTCCCTAG
GCCTGCTACCTGAGACCTGCTGCTTCCTCCCGGGCTTACTCTGCGGCCTCTTCCTGGACCGACCTGGTACCTGCACCCTCTCATTGCCCAAGACCGGCCTCGTTGCTCCTGCCCTTTGGCCTTGGACCATCGGAACGTC
GTGCAACGGGACTACTGCTGGATCCTGGTGGTGATTATCCCCGCTTTACGCAATTCTTGCTTCTTTCTATCTTTTCAATCGCTCACCTTCCCTTCCCCATCACCCCAATCTGTAATAGTGTCTGTCCTCTCCTTTCTTCATCT
CCCTTATTAACATTTGTAATAAACTGGTCGGACCAACATTTTAACCGTTGTTTCTTAATCTCACGCCGGGCATACATATTTCAAAGAACCTCCTCTCCCTCCTATAAATTGGGAGCGAGACA

#25

TTTTATATCATTGACTCAAAGTTTGCTGAGGAACAAGTCCTGGCAAGTCCTGGGCAAAGGCAGAGAAATCTTTTGTCTTGAGGACACTGATGGACAGGTCCTAGCTAAGGATTGTGAAATCCTCTAAGGAGCACCGATGGAA
AAGGCCAGGGACAACGAAAGAAACGGTCTTTTTGTGTGGACTTATCTCGAGGAAGATGGCCATCTCAGGAGGTATGCACAGGACTCTTGCTCAAGCACCCAGGAATGTCACGTAGGCAGGAGAAAATGGAGGATAAAAGA
GGGTCCAACAACCACGGCTAAGTAGAGGCTTATCTCTCACCATGACAGACTTGAGGGGTTCTCTCACCGAAACACAGAGGCTGATCTCTCATGATCTCTCACCACGACACGAGCTTCCTGCCTTCAGATCCTTCTCTACGGA
CCGTTTGCTGACGGACTTCCCTGGGCCTGCTACCTGAGACATGCTGCTTCCTCCCTGACCTGCACCCTTTCGTTGCCCAAGACCGGCCTCGCTGCTCCTGCCCTTCGGCCTCGGACCGTCGGAACGTCGTGCAACGGG
ACTGCTGCCGGATCCTGGTGGTGACTATCCCTGCTTTATGCAATTCTTGCTTCTTTCTACCTTTTCTATCGCTCGCCTTCCCTTCCCCAGCACCCCAATCCGTAATAGTGTCCGTCCTCCCCTTTCTTCATCTCCCTTATTAA
CATTTGTAATAAAATGGTCGGACCAACATTTGAACGGCTGTTTCTTAATCTCACGCCGGGCATACATATTTCAAAGAACCTCTCCCTCCTATAAATTGGAGCGAGACA

#26

TTTTATATCATTGACTCAAAGTTTGCTGAGGAACAAGTCCAGGCAAGTCCTGGGCAAAGGCAGAGAAATCTTTTGTCTTGAGGACACTGATGGACAGGTCCTGACTAAGGATTGTGAAATCCTTTAAGGAGCGCAGATGGAC
AAGGCCAGGGGCATCGAGAGAGAGATAAGCTGCCGCTAATGGCCGGGAAACGGTCTTTTTGTGTGGACTTATCTCAAGGAAAATGGCCATCTCAGGAGGTATGCACAGGACTCTTGCTCAAGCCCCCAGGAATGTCACGT
AGGCAGCAGAAAATGGAGGATAAAAGAGGGTCCAATAACCACAACGGTGGAAGCTGATCCTTCACCACAACCACGGCAACGGTGGAGGCTTATCTCTCACCACGACAGACTTGAGGGGTTCTCTGCCATCTGATCCTTCAC
CACAATGGTGGAGGCTGATCTCTCACCGCAACACAGAGGCTGATCTCTCATGATCTCTCACCACGACACGAGCTTCCTGCCTTCCGATCCTTCTCTACGGACCGTTTGCTGACGGACTTCCCTGGGCCTGCTACCTGAGA
CCTGCTGCTTCCTCCCCGACCTGCACCCTCTCGTTGCCCAAGACCGGCCTCGCTGCTTCTGCCCTTCGGCCTTGGACCGTCGGAACATCGTGCAACGGGACTGCTGCCAGATCCTGGTGGTGACTATCCCCGCTTTACG
CGATTCTTGCCTCTTTCTATCTTTTCTATCGCTCGCCTTCCCTTCCCCATCACCCCAATCCTTAATAGTGTCCGTCCTCCCCTTTCCCCATCTCCCTTATTAACATTTGTAATAAACTGGTCGGACCAACATTTGAACCGCTGTT
TCTTAATCTCACGCCGGGCATACATATTTCAAAGAACCTCCTCTCCCTCCTATAAATTGGAGCGAGACA

#27

TTTTATATCATTGACTCAAAGTTTGCTGAGGAACAAGTCCAGGCAAGTCCTGGGCAAAGGCAGAGAAATCTTTTGTCTTGAGGACACTGATGGACAGGTCCTGGCTAAGGATTGTGAAATCCTCTAAGGAGCACAGATGGAC
AAGGCCAGGGGCATCGAGAGAGAGATAAGCTGCTGCTAATGGCCGGGAAGCGGTCTTTTTGTGTGGACTTATCTCAAGGAAAATGGCCATCTCAGGAGGTATGCACAGGATTCTTGCTCAAGCACCCAGGAATGTCACGT
AGGCAGCAGAAAATGGAGAATAAAAGAGGGTCCAATAACCACGGCAACGGGAGAGGCTTATCTCTCACCACGACAGACTTGAGGGGTTCTCTGCCAACTGATCTCTCACCGCAACGGGGAGACGGATCTCTACGTGGAGA
CTGATCTCTCACCACGACACGAGCTTCCTGCCTTCAGATCCTCCTCTACGGACCGTTTGCTGACGGACTTCCCTGGGCCTGCTACCTGAGACCTGCTGCTTCCTCCCCGACCTGCACCCTCTCGTTGCCCAAGACCGGCC
TCGCTGCTCCTGCCCTTCGGCCTCGGACCGTCGGAACATCGTGCAACGGGACTGCTGCCGGATCCCGGTGGTGACTATCCCCGCTTTACGCAATTCTTGCCTCTTTCTATCATTTCTATCGCTCGCCTTCCCTTCCCCAT
CACCCCAATCCTTAATAGCGTCCGTCCTCCCCTTTCCCCATCTCCCTTATTAACATTTGTAATAAACTGGTCGGACCAACATTTGAACCGTTGTTTCTTAATCTCACGCCGGGCATACATATTTCAAAGAACCTCCTCTCCCTC
CTATAAATTGGAGCGAGACA

| | |
|---|---|
| #28 | TTTTATATCATTGACTCAAAGTTTGCTGAGGAACAAGTCCAGGCAAGTCCTGGGCAAAGGCAGAGAAATCTTTTGTCTTGAGGACACTGATGGACAGGTCCTGGCTGACAAGGCCAGGGGCATCGAGAGAGAGATAAGCTG<br>CTGCTAATGGCCGGGAAGCGGTCTTTTTGTGTGGACTTATCTCAAGGAAAATGGCCATCTCAGGAGGTATGCACAGGACTCTTGCTCAAGCACCCAGGAATGTCACGTAGGCAGCAGAAGATGGAGGATAAAAGAGGGTC<br>CAATAACCACAACGGTGGAAGCTGATCCTTCACCACAACCACGGCAACGAGAGACGCTTATCTCTCACCACGACAGACTTGAGGGGTTCTCTGCCANCTGATCTCTCACCGCAACGTGTAGACGGATCTCTACGTGGAGAC<br>TGATCTCTCACCACGACACGAGCTTCCTGCCTTCCGATCCTCCTCTACGGATCGTTTGCTGACGGACTTCCCTGGGCCTGCTACCTGAGACCTGCTGCTTCCTCCCTGGCCTGCATCCTCTCGCTGCCCCAGACCGGCCT<br>CGCTGCTCCTGCCCTTCGGCCTCGGACCGTCGGAACGTCGTGCAACGGGACTGCTGCCGGATCCTGGTGGTGACTATCCCCGCTTTAGGCAATTCTTGCCTCTTTCTATCTTTTCTATCGCTCGCCTTCCCTTCCCCATC<br>ACCCCAATCCTTAATAGCGTCCGTCCTCCCCTTTCCCCATCTCCCTTATTAACATTTATAATAAACTGGTCGGACCAACATTTGAACCGCTGTTTCTTAATCTCACGCCGGGCATACATATTTCAAAGAACCTCCTCTCCCTCCT<br>ATAAATTGGAGCGAGACA |
| #29 | TTTTATATCATTGACTCAAAGTTTGCTGAGGAACAAGTCCAGGCAAGTCCTGGGCAAAGGCAGAGAAATCTTTTGTCTTGAGGACACTGATGGACAGGTCCTGGCTAAGGACTGTGAAATCATCTAAGGACCACAGATGGAC<br>AAGGCCAGGGGCATCGAGAGAGAGATAAGCTGCTGCTAATGGCCGGGAAGCGGTCTTTTTGTGTGGACTTATCTCAAGGAAAATGGCCATCTCAGGAGGTATGCACAGGACTCTTGCTCAAGCACCCAGGAATGTCACGT<br>AGGCAGCAGAAAATGGAGGATAAAAGAGGGTCCAATAACCACAACGGTGGAAGCTGATCCTTCACCACAACCACGGCAACGGGAGAGGCTTATCTCTCACCACGACAGACTTGAGGGGTTCTCTGCCAACTGATCTCTCAC<br>CGCAACGTGTAGACGGATCTCTACGTGGAGACTGATCTCTCACCACGACACGAGCTTCCTGCCTTCCGATCCTCCTCTACGGACCGTTTGCTGACGGACTTCCCTGGGCCTGCTACCTGAGACCTGCTGCTTCCTCCCTG<br>ACCTGCATCCTCTCGCTGCCCCAGACCGGCCTCGCTGCTCCTGCCCTTCGGCCTCGGACCGTCGGAACGTCGTGCAACGGGACTGCTGCCGGATCCTGGTGGTGACTATCCCCGCTTTACGCAATTCTTGCCTCTTTCT<br>ATCTTTTCTATCGCTCGCCTTCCCTTCCCCATCACCCCAATCCTTAATAGCGTCCGTCCTCCCCTTTCCCCATCTCTCTTATTAACATTTGTAATAAACTGGTCGGACCAACATTTGAACCGTTGTTTCTTAATCTCACGCCGG<br>GCATACATATTTCAAAGAACCTCCTCTCCCTCCTATAAATTGGAGCGAGACA |
| #30 | TTTTATATCATGACTCAAAGTTTGCTGAGGAACAAGTCCAGGCAAGTCCTGGGCAAAGGCAGAGAAATCTTTTGTCTTGAGGACACTGATGGACAGGTCCTGGCGGACAAGACCAGGGGTATCGAGAGAGAGATAAGCTGC<br>TGCTAATGGCCGGGAAGCGGTCTTTTTGTGTGGACTTATCTCAAGGAAAATGGCCATCTCAGGAGGTATGCACAGGACTCTTGCTCAAGCCCCCAGGAATGTCACTTAGGCAGCAGAAAATGGAGAATAAAAGAGGGTCCA<br>ATAACCACAACGGTGGAAGCTGATCCTTCACCACAACCACGGCAACGGGAGAGGTTTATCTCTCACCACGACAGACTTGAGGGGCTCTCTGCCAACTGATCTCTCACCGCAACGTGTAGACGGATCTCTACGTGGAGACTG<br>ATCTCTCACCACGACACGAGCTTCCTGCCTTCCGATCCTCCTCTACGGACCGTTTGCTGACGGACTTCCCTGGGCCTGCTACCTGAGACCTGCTGCTTCCTCCCTGGCCTGCATCCTCTCGCTGCCCCAGACCGGCCTC<br>GCTGCTCCTGCCCTTCGGCCTCGGACCGTCGGAACGTCGTGCAACGCGACTGCTGCCGGATCCTGGTGGTGACTATCCCCGCTTTACGCAATTCTTGCCTCTTTCTATCTTTTCTATCGCTCGCCTTCCCTTCCCCATCA<br>CCCCAATCCTTAATAGCGTCCGTCCTCCCCTTTCCCCATCTCTCTTATTAACATTTATAATAAACTGGTCGGACCAACATTTGAACCGCTGTTTCTTAATCTCACGCCGGGCATACATATTTCAAAGAACCTCTTCTCCCCCTCC<br>TATAAATTGGAGCGAGACA |
| #31 | TTTTATATCATTGACTCAAAGTTTGCTGAGGAACAAGTCCAGGCAAGTCCTGGGCAAAGGTAGAAAAATCTTTTGTCTTGAGGACACTGATGGACAGGTCCTGGCTAAGAACTGTGAAATCCTCTAAGGAGCACAGATGGAC<br>AAGGCCAGGGGCAACGAGAGAGAGATAAGCTGCTGCTAATGGCCGAGAAACGATCTTTTGGTGTGGACTTATCTCGAGGAAGATGGCCATCTCAGGAGGTATGCACATGACTCTTGCTCAAGCAGCCAGGGATGTCACGT<br>AGACAGGAGAAAATGGAGGATAAAAGAGGGTCCAACAACCCCGACAGATCAGAAGGGTTTCGACCATCAGATCCCTCACCACGACGGACTCGAGGTTTCTGCCATCAGATACGGACCGTATGCTGACGGACTTCCCTGGG<br>CCTGCTACCTGAGATCTGCTGCTTCCTCTCAGACTTACTCTGCGGCTTCTTCCTGGACCCACCCGGTACCTGCGCCTTCTCGTTGCTCAAGACCGGCCCCGTTGCTCCTGCCCTTCGGCCTCGGACCATCGGAACGTCG<br>TGCAACGGGACTACTGCCGGATCCTGGTGGTGACTATCCCCACTTTACGCAATTCTTGTTTCTTTCTATCTTTTCTATTGCTCGCCTTCCCTTCCCCATCACCCCAATCCGTAATAGTGTCCGTCCTCCCCTTTCTTCATCTC<br>CCTTATTAACATTTGTAATAAACTGGTCGGACCAACATTTGAACCGTTGTTTCTTAATCTCACGCCGGGCATACATATTTCAAAGAACCTCCTCTCCCTCCTATAAATTGGAGCGAGACA |
| #32 | TTTTATATCATGACTCAAAGTTTGCTGAGGAACAAGTCCAGGCAAGTCCTGGGCAAAGGTAGAGAAATCTTTTGTCTTGAGGACACTGATGGACAGGTCCTGGCTACGGATTGTGATATCCTCTAAGGAGCACAGATGGACA<br>AGGCCAGGGGCATCGAGAGAGAGATAAGCTGCTGCTAATGGCCAGGAAACGGTCTTCAAGGAAAATGGCCATCTCAGGAGGTATGCACAGGACTCTTGCTCAAGCCCCCAGGAATGTCACGTAGGCAGCAGAAAATGGA<br>GGATAAAAGAGGGTCCAATAACCACAACGGTAAAAGCTGATCCTTCACCACAACCACGGCAACGGGAGAGGCTTATCTCTCACCACGACAGACTTGAGGGGTTCTCTGCCAACTGATCTCTCACCGCAACGGGGAGACGGA<br>TCTCTACGTGGAGACTGATCTCTCACCACGACACGAGCTTCCTGCCTTCCGATCCTCCTCTACGGACCGTTTGCTGACGGACTTCCCTGGGCCTGCTACCTGAGACCTGCTGCTTCCTCCCTGGCCTGCATCCTCTCGCT<br>GCCCCAGACCGGCCTCGCTGCTCCTGCCCTTCGGCCTCGGACCGTCGGAACGTCGTGCAACGGGACTGCTGCCGGATCCTGGTGGTGACTATCCCCGCTTTACGCAATTCTTGCCTCTTTCTATCTTTTCTATCGCTCG<br>CCTTCCCTTCCCCATTACCCCAATCCTTAATAGCGTCCGTCCTCCCCTTTCCCCATCTCCCTTATTAACATTTGTAATAAACTGGTCGGACCAACATTTGAACCGCTGTTTCTTAATCTCACGCCGGGCATACATATTTCAAAGA<br>ACCTCCTCTCCCTCCTATAAATTGGAGCGAGACA |
| #33 | TTTTATATCATTGACTCAAAGTTTGCTGAGGAACAAGTCCAGGCAAGTCCTGGGCAAAGGTAGAGAAATCTTCTGTCTGGAGGACACTGATGGACAGGTCCTGGCTAAGGGTTGTGAAATCCTTTAAGGAGCACAGATGGA<br>CAAGGCCGGGGGCAACGAGAGAGGGATAAGCTGCTGATAATGGCCGGGAAACGGTCTTTTGTGTGGACTTATCTCGAGGAAGATGGCCATCTCAGGAGGTATGCACATGACTCTTGCTCAAGCAGCCAGGGATGTCACG<br>TAGGCAGGAAAAAAAGGAGGATTAAAGAGGGTCCAACAACCGACAGACTAGAGGGGTTTTCTGCCATCTGATCTCTCACCACGACAGACTAGAGGGGTTTTCTGCCATCTAATCTCTCACCACGACAGGCTGGAGAAGGGT<br>TTTCTGCCATCAGATCCCTCACCACGACGGACTCGAGGTTTCTGCCTTCAGATCATTCTCTACGGACTGTATGCTGACAGGCTTCCCTGGGTCTGCTACCTGAGACCTGCTGCTTCCTCCCGGGCTTACTCTGCGGCTTCT<br>TCCTGGACCCACCCGGTACCTGCACCTTCTCGTTGCCCAAGACCGGCCTCGCTGCTCCTGCCCTTCAGCCTCGGACCATCGGAACGTCGTGCAACAGGACTACTGCCGGATCCTGGTGGTGACTGTCCCCGCTTTACGC<br>AATTCTTGCTTCTTTCTATCTTTTCTATCGCTCGCCTTCCCTTCCCCATCACCCCAATCCGTAATAGTGTCTGTCCTCTCCTTTCCTCATCTCCCTTATTAACATTTGTAATAAACTGGTCGGACCAACATTTGAACCGTTGTTT<br>CTTAATCTCACGCCGGGCATACATATTTCAGAGAACCTCCTCTCCCTCCTATAAATTGGAGCGAGACA |
| #34 | TTTTATATCATTGACTCAAAGTTTGCTGAGGAACAAGTCCAGGCAAGTCCTGGGCAAAGGCAGAGAAATCTTTTGTCTTGAGGACACTGATGGACAGGTCCTGGCTAAGGATTGTGAAATCCTCTAAGGAGCACAGATGGAC<br>AAGGCCAGGGGCATCAAGAGAGAGATAAGCTGCTGCTAATGGCCGGGAAATGGTCTTTTTTGTGTGGACTTATCTCGAGGAAGATGGCCATCTCAGGAGGTATGCACAGGACTCTTCCTCAGGCACCCAGGAATGCCACG<br>TAGGCAGGAGAAAATGGAGGATAAAAGAGGGTCCAACAACCATGGCGAGGTAGAGGCTTATCTCTCACCACGACAGACTTGAGGGGTTCTCTGCCATCTGATCCTTCACCACAACAGTGGAGGCTGATCTCTCACCGCAAC<br>GTGAAGGCTGATCTCTCACCGCAATGTGGAGGCTGATCTCTCACCGCAACACAGAGGCTGATCTCTCATGATCTCTCACCACAACACGAGCTTCCTGCCTTCAGATCCTTCTCTACGGCCCGTCTGCTGATGGACTTCCCT<br>GGGCCTGCTACCTGAGACCTGCTGCTTCCTCCCTGACCTGCACCCTCTCGTTGCCCAAGACCGCTGCTCCTCGCTGCTCCTGCCCTTCGGCCTCGGACCGTCGGAACATCATGCAGCGGGACTGCTGCCGGATCCTGG<br>TGGTGACTATCCCCGCTTTATGCAATTCTTGCCTCCTTCTATCTTTTCTATCACTCGCCTTCCCTTCCCCATCACCCCAATCCTTAATAGTGTCCGTCCTCCCCTTTCCCCATCTCCCTTATTAACATTTGTAATAAACTGGTCG<br>GACCAACATTTGAACCGTTGTTTCTTAATCTCACGTCGGGCATACATATTTCAAAGAACCTCCTCTCCCTCCTATAAATTGGAGCGAGACA |
| #35 | TTTTATATCATTGACTCAAAGTTTGCTGAGGAACAAGTCCAGGCAAGTCCTGGGCAAAGGTGGAGAAATCTTCTGTCTGGAGGACACTGATGGACAGGTCCTGGCTAAGGGTTGTGAAATCCTTTAAGGAGCACAGATGGA<br>CAAGGCCATGGGCAATGAGAGAGGGATAAGCTGCTGATAATGGCCAGGAAACGGTCTTTTGTGTGGACTTATCTCGAGGAAGATGGCCATCTCAGGAGGTACGCACATGACTCTTGCTCAAGCACCCAGGGATGTCACGT<br>AGGCAGGAAAAAAGAAGGATAAAAGAGGGTCCAACAACCTTGAGGAGAGGGTTTTCTGCCATCAGATCCCTCACCATGATGGACTCAAGGTTTCTGCCATCAGATCCCTCACTATGGACCATATGCTGGCAGACTTCCCTG<br>GGCCTGCTACCTGAAACCTGCTGCTTCCTCCCGGGCTTACTCTGTGGCTTCTTCCTGAACCCACCCGGTACCTGCACCCTCTCGTTGCCCAAGACCGTTCTCACTGCTCCTGCCCTTCGGCCTCGGACCATCAGAATAGC<br>ATGCAACGGGACTGCTGCTGGATCCTGGTGGTGACTATCCCCGCTTTACGCAATTCTTGCCTCTTTCTATCTTTTCTATCGCTCGCCTTCCCTTCCCCATCACCCCAATCCTTAATAGTGTCCGTCCTCCCCTTTCTTCATCT<br>CCCTTATTAACATTTGTAATAAACTGGTCAGACCAACATTTGAACCGTTGTTTCTTAATCTCACGCCGGGCATACATATTTCAAAGAACCTCCTCTCCCTCCTATAAATTGGAGCGAGACA |
| #36 | TTTTATATCATGACTCAAAGTTTGCTGAGGAACAAGTCCAGGCAAGTCCTGGGCAAAGGCAGAGAAATCTTTTGTCTTGAGGACACTGATGGACAGGTCCTGGCTAAGGATTGTGAAATCCTTTAAGAAGCACAGATGGACA<br>AGGCCAGGGGCAACGAGAGAGAGATAAGCTGCTGATAATGGCCGGGAAACGGTCTTATCTCGAGGAAGATGGCCATCTCAGGAGGTATGCACAGGACTCTTGCTCAAGCACCCAGGAATGTCACGTAGGCAGCAGAAAA<br>TGGAGGATAAAAGAGGGTCCAATAACCACAACGGTGGAAGCTGATCCTTCACCACAACCACGGCAACGGGAGAGGCTTATCTCTCACCACGACAGACTTGAGGGGTTCTCTGCCAACTGATCTCTCATCGCAACAGTGGAG<br>ACTGATTTCTCACCACGGCACGAGCTTCCTGCCTTCCGATCCTCCTCTACGGACCGTTTGCTGACGGACTTCCCTGGGCCTGCTACCTGAGACCTGCTGCTTCCTCCCCGACCTGCACCCTCTCGTTGCCCAAGACCGGC<br>CTCGCTGCTCCTGCCCTTCGGCCTCGGACCGTCGGAACATCGTGCAACGGGACTGCTGCCGGATCCTGGTGGTGACTATCCCCGCTTTACGCAATTCTTGCCTCTTTCTATCATTTCTATCGCTCGCCTTCCCTTCCCCA<br>TCACCCCAATCCTTAATAGCGTCCGTCCTCTCCTTTCCCCATCTCCCTTATTAACATTTGTAATAAACTGGTCGGACCAACATTTGAACCGCTGTTTCTTAATCTCACGCCGGGCATACATATTTCAAAGAACCTCCTCTCCCTC<br>CTATAAATTGGAGCGAGACA |
| #37 | TTTTATATCATTGACTCAAAGTTTGCTGAGGAACAAGTCCAGGGCAAAGGTAGAGAAATCTTCTGTCTGGAGGACACTGATGGACGGGTCCTGGCTAAGGGTTGTGAAATCCTTTAAGGAGCACAGATGGACAAGGCCAGG<br>GGCAACAAGAGAGAGAAAAGCTGCTGATAATGGCCGGGAAACGATCTTTTGTGTGGACTTATCTCGAGGAAGATGGCCATCTCAGGAGGCACGCACATGACTTTTGCTCAAGCAACCAGGGATGTCACGTAGGCAGGAAA<br>AAAAGGAGGATAAAAGAGGGTCCAACAACCAAGAGGCTAGAGGTTTCTGCCATCAGATCCCTCATCACTATGGACCGTATGCTGATGGACTTCCCTGGGCCTGCTACCTGAGACCTGCTGCTTCCTCCCAGACTTACTCTG<br>CGGCTTCTTCCTGGACCCACCCGGTACCTGCACCCTCTCGTTGCCCAAGACCAGCCTCGCTGCTCCTGCCCTTCGGCTTCGGACCATCAGAACGTTGTGCAACGGGACTACTGTCAAATCCTGGTGGTGACTATCCCCG<br>CTTTACGCAATTCTTGCTTCTTTCTATCTTTTCTCTCGCTCTCCTTCCCTTCCCCATCACCCCAATCCGTAATAGTGTCCGTCCTACCCTTTCTTCATCTCCCTTATTAACATTTGTAATAAACTGGTCGGACCAACATTTGAAC<br>CGTTGTTTCTTAATCTCACTCCGGGCATACATATTTCAAAGAACCTCCTCTCCCTCCTATAAATTGGAGCGAGACA |
| #38 | TTTTATATCATTGACTCAAAGTTTGCTGAGGAACAAGTCCAGGGCAAAGGTAGAGAAATCTTCTGTCTGGAGGACACTGATGGACGGGTCCTGGCTAAGGGTTGTGAAATCCTTTTAGGAGCACAGATGGACAAGGCCAGG<br>GGCAAGGAGAGAGGGATAAGCTGCTGATAATGGCCGGGAAACGGTCTTTTGTGTGGACTTATCTCGAGGAAGATGGCCATCCCAGGAGGTATGCACATGTCCATCCCATCACCCCAATCTGTAATAGTGTCCATCCTCCC<br>CTTTCTTCATCTCCATTATTAACATTTGTAATAAACTGGTTGGACCAACATTTGAACTGTTGTTTCTTAATCTCACTCCGGGCATACATATTTCAAAGAACCTCCTCTCCCTCCTATAAATTGGAGCGAGACA |
| #39 | TTTTATATCATTGACTCAAAGTTTGCTGAGGAACAAGTCCAGGCAAGTCCTGGGCAAAGGCAGAGAAATCTTTTGTCTTGAGGACACTGATGGACAGGTCCTGGCTAAGGATTGTGAAATCCTTTAAGGAGCACAGATGGAC<br>AAGGCCAGGGGCATCGAGAGAGAGATAAGCTGCCGCTAATGGCCGGGAAACGGTCTTTTTGTGTGGACTTATCTCAAGGAAAATGGCCATCTCAGGAGGTATGCACATGACTCTTGCTCAAGCACCCAGGAATGTCACGT<br>AGGCAGCAGAAGATGGAGGATAAAAGAGGGTCCAATAACCACAACGGTGGAAGCTGATCCTTCACCACAACCACGGCAACGGGAGAGGCTTATCTCTCACCACGACAGACTTGAGGGGTTCTCTGCCAACTGATCTCTCAC<br>CGCAACGTGTAGACGGATCTCTACGTGGAGACTGATCTCTCACCACGACACGAGCTTCCTGCCTTCCGATCCTCCTCTACGGACCGTTTGCTGACGGACTTCCCTGGGCCTGCTACCTGAGACCTGCTGCTTCCTCCCTG<br>ACCTGCATCCTCTCGCTGCCCCAGACCGGCCTTGCTGCTCCTGCCCTTCGGCCCTGGACCGTCGGAACATCGTGCAACGGGACTGCTGCCGGATCCTGGTGGTGACTATCCCCGCTTTACGCAATTCTTGCCTCTTTCT<br>ATCTTTTCTATCGCTCGCCTTCCCTTCCCCATCACCCCAATCCTTAATAGTGTCCGTCCTCCCCTTTCCCCATCTCCCTTATTAACATTTGTAATAAACTGGTCGGACCAACATTTGAACCGCTGTTTCTTAATCTCACGCCGG<br>GCATACATATTTCAAAGAACCTCTTCTCCCTCCTATAAATTGGAGCGAGACA |
| #40 | TTTTATATCATTGACTCAAAGTTTGCTGAGGAACAAGTCCAGGCAAGTCCTGGGCAAAGGCAGAGAAATCTTTTGTCTTGAGGACACTGATGGACAGGTCCTGGCTAAGGATTGTGAAATCCTTTAAGGAGCACAGATGGAC<br>AAGGCCAGGGGCATCGAGAGAGAGATAAGCTGCCGCTAATGGCCGGTAAACGGTCTTTTTGTGTGGACTTATCTCAAGGAAAATGGCCATCTCAGGAGGTATGCACCGGACTCTTGCTCAAGCCCCCAGGAATGTCACGT<br>AGGCAGCAGAAAATGGAGGATAAAAGAGGGTCCAATAACCACAACGGTGGAAGCTGATCCTTCACCACAACCACGGCAACGGGAGAGGCTTATCTCTCGNCACGACAGACTTGAGGGGTTCTCTCTGCCAACTGATCTCTC<br>ACCGCAACGTGGAGACGGATCTCTACGTGGAGACTGATCTCTCACCACGGCACGAGCTTCCTGCCTTCCGATCCTCCTCTACGGACCGTTTGCTGACGGACTTCCCTGGGCCTGCTACCTGAGACCTGCTGCTTCCTCC<br>CTGACCTGCATCCTCTCATTGCCCAAGACCGGCCCCGCTGCTCCTGCCCTTCGGCCTCGGACCGTCGGAACATCGTGCAACGGGACTGCTGCCGGATCCTGGTGGTGACTATCCCCGCTTTACGCAATTCTTGCCTCTT<br>TCTATCTTTTCTATCGCTCGCCTTCCCTTCCCCATCACCCCAATCCTTAATAGTGTCCGTCCTCCCCTTTCCCCATCTCCCTTATTAACATTTGTAATAAACTGGTCGGACCAACATTTGAACCGTTGTTTCTTAATCTCACGCC<br>AGGCATACATATTTCAAAGAACCTCCTCTCCCTCCTATAAATTGGAGCGAGACA |
| #41 | TTTTATATCATTGACTCAAAGTTTGCTGAGGAACAAGTCCAGGCAAGTCCTGGGCAAAGGCAGAGAAATCTTTTGTCTTGAGGACACTGATGGACAGGTCCTGGCTAAGGATTGTGAAATCCTCTAAGGAGCACAGATGGAC<br>AAGGCCAGGGGCAACGAGAGAGAGATAAGCTGCTGCTAATGGCTGGGAAACGGTCTTTTTGTGTGGACTTATCTCGAGGAAGATGGCCATCTCAGGAGGTATGCACAGGACTCTTGCTCAAGCACCCAGGAATGTCACGT<br>AGGCAGGAGAAAATGGAGGATAAAAGAGGGTCCAACAACCACGGCGGTGTGGAGGTTGATCCTTCACCACAACTGAGAGGCTCATCTCTCACCACGACAGACTTGAGGGGTTCTCTGCCATCTGATCCTTCACCACAACA<br>GTGGAGGCTGATCTCTCACCGCAACACAGAGGCTGATCTCTCATGATCTCTCACCACGACACGAGCTTCCTGCCTTCAGATCCTTCTCTATGGACCGTTTGCTGACGGACTTCCCTGGGCCTGCTACCTGAGACCTGCTG<br>CTTCCTCCCTGACCTGCACCCTCTCATTGCCAAGACCGGCCTCGCTGCTCCTGCCCTTCGGCCTCGGACCGTCAGAACGTCGTGCAACGGGACTACTGCCGGATCCTGGTGGTGACTATCCCCACTTTACGCAATTCTTG<br>CTTCTTTCTATCTTTTCTATCGCTCGCTTTCCCTTCCCCAACACCCCAATCCGTAATAGTGTCTGTCCTCCCCTTTCTTCTTCTCCCTTATTAACATTTGTAATAAACTGGTTGGACCAACATTTGAACCATTGTTTCTTAATCTC<br>ACGCCGGGCATACATATTTCAAAGAACCTCCTCTCCCTCCTATAAATTGGAGCGAGACA |
| 3xmotif7_core | ACAAGGCCAGGGGCATCGAGAGAGAGATAAGCTGCCGCTAATGGCCGGGATCGAGAGAGAGATAAGCTGCCGCTAATGGCCGGGATCGAGAGAGAGATAAGCTGCCGCTAATGGCCGGGAAACGGTCTTTTTGTG |

Supplementally table S14. Transcription factor binding sites prediction of motif 7 sequence

| | |
|---|---|
| Input sequence (motif7) | ACAAGGCCAGGGGCATCGAGAGAGAGATAAGCTGCCGCTAATGGCCGGGA |

| ID | NAME | Score | Relative Score | start | end | strand | sequence |
|---|---|---|---|---|---|---|---|
| MA1111.1 | MA1111.1.NR2F2 | 6.6912417 | 0.81822634 | 1 | 11 | + | ACAAGGCCAGG |
| MA0442.1 | MA0442.1.SOX10 | 7.093567 | 0.9193686 | 1 | 6 | - | CCTTGT |
| MA0442.3 | MA0442.3.SOX10 | 5.210161 | 0.8400845 | 1 | 6 | + | ACAAGG |
| MA0592.4 | MA0592.4.ESRRA | 7.8106885 | 0.83449465 | 1 | 9 | + | ACAAGGCCA |
| MA0592.1 | MA0592.1.ESRRA | 7.0254955 | 0.8028078 | 1 | 11 | + | ACAAGGCCAGG |
| MA1981.2 | MA1981.2.ZNF530 | 15.069937 | 0.90924543 | 1 | 14 | + | ACAAGGCCAGGGGC |
| MA1981.1 | MA1981.1.ZNF530 | 14.031356 | 0.86623096 | 1 | 18 | + | ACAAGGCCAGGGGCATCG |
| MA1710.1 | MA1710.1.ZNF257 | 4.8707857 | 0.8075619 | 1 | 12 | + | ACAAGGCCAGGG |
| MA0442.1 | MA0442.1.SOX10 | 7.093567 | 0.9193686 | 1 | 6 | - | CCTTGT |
| MA0505.3 | MA0505.3.Nr5A2 | 10.102723 | 0.8832274 | 1 | 9 | - | TGGCCTTGT |
| MA0512.1 | MA0512.1.Rxra | 6.31979 | 0.85536015 | 1 | 11 | + | ACAAGGCCAGG |
| MA0643.2 | MA0643.2.Esrrg | 6.091519 | 0.8512928 | 1 | 9 | + | ACAAGGCCA |
| MA0643.1 | MA0643.1.Esrrg | 4.456385 | 0.8217325 | 1 | 10 | + | ACAAGGCCAG |
| MA1996.1 | MA1996.1.Nr1H2 | 4.8434944 | 0.80952555 | 1 | 11 | + | ACAAGGCCAGG |
| MA1112.3 | MA1112.3.NR4A1 | 5.117354 | 0.81147635 | 2 | 9 | + | CAAGGCCA |
| MA1541.2 | MA1541.2.NR6A1 | 12.675077 | 0.867303 | 2 | 15 | + | CAAGGCCAGGGGCA |
| MA1535.1 | MA1535.1.NR2C1 | 2.381361 | 0.8172728 | 2 | 10 | + | CAAGGCCAG |
| MA0484.1 | MA0484.1.HNF4G | 4.280618 | 0.8044322 | 2 | 16 | + | CAAGGCCAGGGGCAT |
| MA0017.3 | MA0017.3.NR2F1 | 4.6194096 | 0.8109275 | 2 | 13 | + | CAAGGCCAGGGG |
| MA0160.3 | MA0160.3.NR4A2 | 5.0754676 | 0.80202174 | 2 | 9 | + | CAAGGCCA |
| MA1541.2 | MA1541.2.NR6A1 | 12.675077 | 0.867303 | 2 | 15 | + | CAAGGCCAGGGGCA |
| MA1535.1 | MA1535.1.NR2C1 | 2.381361 | 0.8172728 | 2 | 10 | + | CAAGGCCAG |
| MA0039.1 | MA0039.1.Klf4 | 5.5982046 | 0.8211851 | 2 | 11 | + | CAAGGCCAGG |
| MA1110.2 | MA1110.2.Nr1H4 | 3.8622274 | 0.81830287 | 2 | 11 | + | CAAGGCCAGG |
| MA0671.1 | MA0671.1.NFIX | 6.3595567 | 0.895226 | 3 | 11 | + | AAGGCCAGG |
| MA1111.2 | MA1111.2.NR2F2 | 7.0944977 | 0.85886484 | 3 | 9 | + | AAGGCCA |
| MA1536.1 | MA1536.1.NR2C2 | 0.716456 | 0.82629126 | 3 | 10 | + | AAGGCCAG |
| MA0160.1 | MA0160.1.NR4A2 | 8.129536 | 0.87670153 | 3 | 10 | + | AAGGCCAG |
| MA1536.1 | MA1536.1.NR2C2 | 0.716456 | 0.82629126 | 3 | 10 | + | AAGGCCAG |
| MA1710.2 | MA1710.2.ZNF257 | 6.0860734 | 0.8366518 | 3 | 12 | + | AAGGCCAGGG |
| MA0160.1 | MA0160.1.NR4A2 | 8.129536 | 0.87670153 | 3 | 10 | + | AAGGCCAG |
| MA0065.3 | MA0065.3.Pparg::Rxra | 7.8783884 | 0.81435686 | 3 | 15 | + | AAGGCCAGGGGCA |
| MA0505.1 | MA0505.1.Nr5a2 | 7.9381347 | 0.808703 | 3 | 17 | + | AAGGCCAGGGGCATC |
| MA0146.1 | MA0146.1.Zfx | 6.628974 | 0.8083232 | 3 | 16 | - | ATGCCCCTGGCCTT |
| MA0146.2 | MA0146.2.Zfx | 6.622901 | 0.80815655 | 3 | 16 | - | ATGCCCCTGGCCTT |
| MA0039.1 | MA0039.1.Klf4 | 5.0854774 | 0.80613446 | 3 | 12 | + | AAGGCCAGGG |
| MA2338.1 | MA2338.1.Ppara | 5.662796 | 0.8290682 | 3 | 9 | + | AAGGCCA |
| MA0810.1 | MA0810.1.TFAP2A | 9.27623 | 0.8969335 | 4 | 15 | - | TGCCCCTGGCCT |
| MA0811.1 | MA0811.1.TFAP2B | 8.814459 | 0.88014466 | 4 | 15 | - | TGCCCCTGGCCT |
| MA0811.1 | MA0811.1.TFAP2B | 8.788611 | 0.8797347 | 4 | 15 | + | AGGCCAGGGGCA |
| MA0811.2 | MA0811.2.TFAP2B | 7.891571 | 0.8741564 | 4 | 14 | - | GCCCCTGGCCT |
| MA0810.1 | MA0810.1.TFAP2A | 6.956968 | 0.8609372 | 4 | 15 | + | AGGCCAGGGGCA |
| MA0810.2 | MA0810.2.TFAP2A | 6.0281863 | 0.85618174 | 4 | 14 | + | AGGCCAGGGGC |
| MA0830.2 | MA0830.2.TCF4 | 7.5247426 | 0.823848 | 4 | 16 | - | ATGCCCCTGGCCT |
| MA1536.2 | MA1536.2.NR2C2 | 0.661936 | 0.8306202 | 4 | 9 | + | AGGCCA |
| MA1535.2 | MA1535.2.NR2C1 | 1.9623798 | 0.82428306 | 4 | 9 | + | AGGCCA |
| MA0524.1 | MA0524.1.TFAP2C | 10.028958 | 0.8902031 | 4 | 18 | - | CGATGCCCCTGGCCT |
| MA0524.2 | MA0524.2.TFAP2C | 9.560668 | 0.8895406 | 4 | 15 | + | AGGCCAGGGGCA |
| MA0524.2 | MA0524.2.TFAP2C | 8.495872 | 0.87223715 | 4 | 15 | - | TGCCCCTGGCCT |
| MA0524.3 | MA0524.3.TFAP2C | 7.559289 | 0.8654068 | 4 | 14 | - | GCCCCTGGCCT |
| MA0498.2 | MA0498.2.MEIS1 | 1.0118972 | 0.8021741 | 4 | 10 | + | AGGCCAG |
| MA0499.2 | MA0499.2.MYOD1 | 6.014278 | 0.8000991 | 4 | 16 | - | ATGCCCCTGGCCT |
| MA0003.2 | MA0003.2.TFAP2A | 9.272595 | 0.84573364 | 4 | 18 | - | CGATGCCCCTGGCCT |
| MA0003.4 | MA0003.4.TFAP2A | 7.297412 | 0.8003344 | 4 | 17 | - | GATGCCCCTGGCCT |
| MA1536.2 | MA1536.2.NR2C2 | 0.661936 | 0.8306202 | 4 | 9 | + | AGGCCA |
| MA1535.2 | MA1535.2.NR2C1 | 1.9623798 | 0.82428306 | 4 | 9 | + | AGGCCA |
| MA1631.1 | MA1631.1.ASCL1 | 6.202894 | 0.8063441 | 4 | 16 | - | ATGCCCCTGGCCT |
| MA0146.3 | MA0146.3.Zfx | 8.742937 | 0.8747475 | 4 | 13 | - | CCCCTGGCCT |
| MA0141.2 | MA0141.2.Esrrb | 7.243574 | 0.8368991 | 4 | 15 | + | AGGCCAGGGGCA |
| MA0141.1 | MA0141.1.Esrrb | 7.2546544 | 0.83600986 | 4 | 15 | + | AGGCCAGGGGCA |
| MA2337.1 | MA2337.1.Nr1h3 | 4.2773867 | 0.8531281 | 4 | 9 | + | AGGCCA |
| MA1110.3 | MA1110.3.Nr1H4 | 3.5065694 | 0.82409453 | 4 | 9 | + | AGGCCA |
| MA1996.2 | MA1996.2.Nr1H2 | 4.3566766 | 0.82107645 | 4 | 9 | + | AGGCCA |
| MA0671.2 | MA0671.2.NFIX | 6.32249 | 0.90594786 | 5 | 10 | + | GGCCAG |
| MA0810.2 | MA0810.2.TFAP2A | 9.842387 | 0.9174111 | 5 | 15 | - | TGCCCCTGGCC |
| MA0811.2 | MA0811.2.TFAP2B | 8.936252 | 0.89155364 | 5 | 15 | + | GGCCAGGGGCA |
| MA0814.1 | MA0814.1.TFAP2C | 6.3595576 | 0.8498663 | 5 | 15 | + | GGCCAGGGGCA |
| MA0812.1 | MA0812.1.TFAP2B | 5.5890255 | 0.83491045 | 5 | 15 | - | TGCCCCTGGCC |
| MA0814.1 | MA0814.1.TFAP2C | 5.234773 | 0.831827 | 5 | 15 | - | TGCCCCTGGCC |
| MA0812.1 | MA0812.1.TFAP2B | 3.8323958 | 0.80827916 | 5 | 15 | + | GGCCAGGGGCA |
| MA1569.1 | MA1569.1.TFAP2E | 4.247269 | 0.8582885 | 5 | 15 | - | TGCCCCTGGCC |
| MA1569.1 | MA1569.1.TFAP2E | 4.028171 | 0.85536283 | 5 | 15 | + | GGCCAGGGGCA |

| | | | | | | | |
|---|---|---|---|---|---|---|---|
| MA1569.2 | MA1569.2.TFAP2E | 0.26566607 | 0.81091505 | 5 | 13 | + | GGCCAGGGG |
| MA1569.2 | MA1569.2.TFAP2E | 0.1608316 | 0.80944 | 5 | 13 | - | CCCCTGGCC |
| MA0524.3 | MA0524.3.TFAP2C | 9.73621 | 0.9026767 | 5 | 15 | + | GGCCAGGGGCA |
| MA0161.1 | MA0161.1.NFIC | 6.6678586 | 0.8985289 | 5 | 10 | - | CTGGCC |
| MA0003.3 | MA0003.3.TFAP2A | 7.4503756 | 0.8719181 | 5 | 15 | - | TGCCCCTGGCC |
| MA0003.3 | MA0003.3.TFAP2A | 3.7610755 | 0.81392246 | 5 | 15 | + | GGCCAGGGGCA |
| MA0154.2 | MA0154.2.EBF1 | 0.34467325 | 0.81170136 | 5 | 15 | - | TGCCCCTGGCC |
| MA0155.1 | MA0155.1.INSM1 | 7.915667 | 0.81009734 | 5 | 16 | + | GGCCAGGGGCAT |
| MA1569.1 | MA1569.1.TFAP2E | 4.247269 | 0.8582885 | 5 | 15 | - | TGCCCCTGGCC |
| MA1569.1 | MA1569.1.TFAP2E | 4.028171 | 0.85536283 | 5 | 15 | + | GGCCAGGGGCA |
| MA1569.2 | MA1569.2.TFAP2E | 0.26566607 | 0.81091505 | 5 | 13 | + | GGCCAGGGG |
| MA1569.2 | MA1569.2.TFAP2E | 0.1608316 | 0.80944 | 5 | 13 | - | CCCCTGGCC |
| MA0812.2 | MA0812.2.TFAP2B | 7.183233 | 0.86883986 | 6 | 14 | - | GCCCCTGGC |
| MA0830.1 | MA0830.1.TCF4 | 3.7106338 | 0.8441797 | 6 | 15 | - | TGCCCCTGGC |
| MA0812.2 | MA0812.2.TFAP2B | 5.3741527 | 0.8391104 | 6 | 14 | + | GCCAGGGGC |
| MA0814.3 | MA0814.3.TFAP2C | 6.9789968 | 0.8205242 | 6 | 14 | + | GCCAGGGGC |
| MA1100.2 | MA1100.2.ASCL1 | 4.4624534 | 0.8107925 | 6 | 15 | - | TGCCCCTGGC |
| MA0820.1 | MA0820.1.FIGLA | 4.349767 | 0.80282074 | 6 | 15 | - | TGCCCCTGGC |
| MA1569.2 | MA1569.2.TFAP2E | 5.051339 | 0.87825114 | 6 | 14 | - | GCCCCTGGC |
| MA1569.2 | MA1569.2.TFAP2E | 4.9168587 | 0.876359 | 6 | 14 | + | GCCAGGGGC |
| MA1540.1 | MA1540.1.NR5A1 | 6.0451827 | 0.8209527 | 6 | 16 | + | GCCAGGGGCAT |
| MA0522.2 | MA0522.2.TCF3 | 3.3915277 | 0.8265048 | 6 | 15 | - | TGCCCCTGGC |
| MA0499.3 | MA0499.3.MYOD1 | 5.7407036 | 0.80857533 | 6 | 14 | - | GCCCCTGGC |
| MA0003.1 | MA0003.1.TFAP2A | 8.823754 | 0.9488304 | 6 | 14 | + | GCCAGGGGC |
| MA0003.1 | MA0003.1.TFAP2A | 8.6296425 | 0.9424961 | 6 | 14 | - | GCCCCTGGC |
| MA0003.5 | MA0003.5.TFAP2A | 7.197854 | 0.8213118 | 6 | 14 | - | GCCCCTGGC |
| MA0162.2 | MA0162.2.EGR1 | 2.3430543 | 0.8052722 | 6 | 19 | - | TCGATGCCCCTGGC |
| MA1569.2 | MA1569.2.TFAP2E | 5.051339 | 0.87825114 | 6 | 14 | - | GCCCCTGGC |
| MA1569.2 | MA1569.2.TFAP2E | 4.9168587 | 0.876359 | 6 | 14 | + | GCCAGGGGC |
| MA1540.1 | MA1540.1.NR5A1 | 6.0451827 | 0.8209527 | 6 | 16 | + | GCCAGGGGCAT |
| MA1631.2 | MA1631.2.ASCL1 | 6.2955065 | 0.82368463 | 6 | 14 | - | GCCCCTGGC |
| MA0830.3 | MA0830.3.TCF4 | 6.944005 | 0.8429033 | 7 | 14 | - | GCCCCTGG |
| MA1100.3 | MA1100.3.ASCL1 | 4.6310854 | 0.82482314 | 7 | 14 | - | GCCCCTGG |
| MA1100.3 | MA1100.3.ASCL1 | 3.701287 | 0.80523014 | 7 | 14 | + | CCAGGGGC |
| MA0592.1 | MA0592.1.ESRRA | 8.548125 | 0.8321985 | 7 | 17 | + | CCAGGGGCATC |
| MA0512.1 | MA0512.1.Rxra | 4.471038 | 0.82873076 | 7 | 17 | + | CCAGGGGCATC |
| MA0643.1 | MA0643.1.Esrrg | 3.3550744 | 0.8055314 | 7 | 16 | + | CCAGGGGCAT |
| MA0738.1 | MA0738.1.HIC2 | 7.9631376 | 0.90590775 | 8 | 16 | - | ATGCCCCTG |
| MA0820.2 | MA0820.2.FIGLA | 6.0417624 | 0.8816823 | 8 | 13 | - | CCCCTG |
| MA1117.1 | MA1117.1.RELB | 6.6722445 | 0.801272 | 8 | 18 | - | CGATGCCCCTG |
| MA1535.1 | MA1535.1.NR2C1 | 6.189681 | 0.8906118 | 8 | 16 | + | CAGGGGCAT |
| MA0597.3 | MA0597.3.THAP1 | 5.4647593 | 0.8112446 | 8 | 15 | + | CAGGGGCA |
| MA0103.4 | MA0103.4.ZEB1 | 2.5357227 | 0.811137 | 8 | 13 | - | CCCCTG |
| MA1535.1 | MA1535.1.NR2C1 | 6.189681 | 0.8906118 | 8 | 16 | + | CAGGGGCAT |
| MA0739.1 | MA0739.1.Hic1 | 5.806898 | 0.8347071 | 8 | 16 | - | ATGCCCCTG |
| MA1536.1 | MA1536.1.NR2C2 | 6.380095 | 0.9297215 | 9 | 16 | + | AGGGGCAT |
| MA1536.1 | MA1536.1.NR2C2 | 6.380095 | 0.9297215 | 9 | 16 | + | AGGGGCAT |
| MA0739.2 | MA0739.2.Hic1 | 6.3469577 | 0.8578637 | 9 | 16 | - | ATGCCCCT |
| MA1117.2 | MA1117.2.RELB | 7.3175187 | 0.8508877 | 10 | 16 | - | ATGCCCC |
| MA1536.2 | MA1536.2.NR2C2 | 5.825279 | 0.92812663 | 10 | 15 | + | GGGGCA |
| MA1535.2 | MA1535.2.NR2C1 | 5.488384 | 0.8964561 | 10 | 15 | + | GGGGCA |
| MA1517.2 | MA1517.2.KLF6 | 5.3420486 | 0.8226365 | 10 | 18 | - | CGATGCCCC |
| MA0527.1 | MA0527.1.ZBTB33 | 9.933018 | 0.8388569 | 10 | 24 | - | CTCTCTCGATGCCCC |
| MA0528.1 | MA0528.1.ZNF263 | 8.833209 | 0.8290015 | 10 | 30 | + | GGGGCATCGAGAGAGAGATAA |
| MA1536.2 | MA1536.2.NR2C2 | 5.825279 | 0.92812663 | 10 | 15 | + | GGGGCA |
| MA1535.2 | MA1535.2.NR2C1 | 5.488384 | 0.8964561 | 10 | 15 | + | GGGGCA |
| MA1517.2 | MA1517.2.KLF6 | 5.3420486 | 0.8226365 | 10 | 18 | - | CGATGCCCC |
| MA2337.1 | MA2337.1.Nr1h3 | 4.3381734 | 0.8544442 | 10 | 15 | + | GGGGCA |
| MA0738.2 | MA0738.2.HIC2 | 9.837441 | 1 | 11 | 16 | - | ATGCCC |
| MA1563.1 | MA1563.1.SOX18 | 6.5160456 | 0.8991971 | 13 | 20 | - | CTCGATGC |
| MA1563.1 | MA1563.1.SOX18 | 6.5160456 | 0.8991971 | 13 | 20 | - | CTCGATGC |
| MA0035.1 | MA0035.1.Gata1 | 6.597768 | 0.97241 | 13 | 18 | - | CGATGC |
| MA0527.2 | MA0527.2.ZBTB33 | 8.0358305 | 0.8378458 | 14 | 23 | - | TCTCTCGATG |
| MA0036.1 | MA0036.1.GATA2 | 5.55889 | 0.93657154 | 14 | 18 | - | CGATG |
| MA0140.2 | MA0140.2.GATA1::TAL1 | 7.5852513 | 0.8210826 | 14 | 31 | - | CTTATCTCTCTCTCGATG |
| MA0140.3 | MA0140.3.GATA1::TAL1 | 6.8749294 | 0.8178819 | 14 | 30 | - | TTATCTCTCTCTCGATG |
| MA0140.1 | MA0140.1.Gata1::Tal1 | 12.563282 | 0.8967722 | 14 | 31 | + | CATCGAGAGAGAGATAAG |
| MA0673.1 | MA0673.1.NKX2-8 | 1.6418386 | 0.80022925 | 15 | 23 | - | TCTCTCGAT |
| MA0673.2 | MA0673.2.NKX2-8 | 3.0161836 | 0.8292545 | 16 | 23 | - | TCTCTCGA |
| MA0508.3 | MA0508.3.PRDM1 | 6.6321983 | 0.8232267 | 16 | 26 | - | CTCTCTCTCGA |
| MA2003.2 | MA2003.2.NKX2-4 | 4.3385158 | 0.8117606 | 16 | 23 | - | TCTCTCGA |
| MA0037.1 | MA0037.1.GATA3 | 4.5302625 | 0.83815706 | 17 | 22 | + | CGAGAG |
| MA0039.1 | MA0039.1.Klf4 | 5.692202 | 0.82394433 | 17 | 26 | + | CGAGAGAGAG |
| MA0508.4 | MA0508.4.PRDM1 | 8.155882 | 0.8855621 | 18 | 24 | - | CTCTCTC |
| MA0508.3 | MA0508.3.PRDM1 | 9.121172 | 0.87104386 | 18 | 28 | - | ATCTCTCTCTC |
| MA0471.1 | MA0471.1.E2F6 | 2.7671793 | 0.80184823 | 18 | 28 | + | GAGAGAGAGAT |

| | | | | | | | |
|---|---|---|---|---|---|---|---|
| MA1731.2 | MA1731.2.ZNF768 | 8.027947 | 0.8550021 | 18 | 26 | - | CTCTCTCTC |
| MA0037.1 | MA0037.1.GATA3 | 5.2857265 | 0.8698499 | 19 | 24 | + | AGAGAG |
| MA0508.4 | MA0508.4.PRDM1 | 8.155882 | 0.8855621 | 20 | 26 | - | CTCTCTC |
| MA1731.2 | MA1731.2.ZNF768 | 6.877582 | 0.83208734 | 20 | 28 | - | ATCTCTCTC |
| MA0037.1 | MA0037.1.GATA3 | 5.2857265 | 0.8698499 | 21 | 26 | + | AGAGAG |
| MA1104.1 | MA1104.1.GATA6 | 12.036442 | 0.9204205 | 22 | 34 | + | GAGAGATAAGCTG |
| MA0033.1 | MA0033.1.FOXL1 | 5.3230734 | 0.8462748 | 22 | 29 | + | GAGAGATA |
| MA0482.1 | MA0482.1.Gata4 | 14.057045 | 0.9738922 | 22 | 32 | - | GCTTATCTCTC |
| MA1104.2 | MA1104.2.GATA6 | 11.594254 | 0.8943952 | 23 | 35 | - | GCAGCTTATCTCT |
| MA0891.1 | MA0891.1.GSC2 | 5.9211655 | 0.8343217 | 23 | 32 | - | GCTTATCTCT |
| MA0482.2 | MA0482.2.GATA4 | 11.620038 | 0.90677756 | 23 | 34 | - | CAGCTTATCTCT |
| MA0635.1 | MA0635.1.BARHL2 | 3.802522 | 0.837378 | 23 | 32 | - | GCTTATCTCT |
| MA0648.1 | MA0648.1.GSC | 5.2353644 | 0.8186342 | 23 | 32 | - | GCTTATCTCT |
| MA0036.3 | MA0036.3.GATA2 | 13.068148 | 0.9714925 | 23 | 33 | - | AGCTTATCTCT |
| MA0036.2 | MA0036.2.GATA2 | 13.081162 | 0.9299424 | 23 | 36 | - | GGCAGCTTATCTCT |
| MA0035.4 | MA0035.4.GATA1 | 10.762284 | 0.91447955 | 23 | 33 | - | AGCTTATCTCT |
| MA1970.1 | MA1970.1.TRPS1 | 11.659926 | 0.9078272 | 23 | 34 | - | CAGCTTATCTCT |
| MA0035.2 | MA0035.2.Gata1 | 12.180768 | 0.95631444 | 23 | 33 | + | AGAGATAAGCT |
| MA0035.3 | MA0035.3.Gata1 | 13.138827 | 0.94785124 | 23 | 33 | - | AGCTTATCTCT |
| MA0037.4 | MA0037.4.Gata3 | 11.844524 | 0.9196418 | 23 | 34 | - | CAGCTTATCTCT |
| MA0766.2 | MA0766.2.GATA5 | 10.435503 | 0.94904095 | 24 | 33 | + | GAGATAAGCT |
| MA0719.1 | MA0719.1.RHOXF1 | 5.623186 | 0.91266996 | 24 | 31 | - | CTTATCTC |
| MA0712.1 | MA0712.1.OTX2 | 5.0443745 | 0.8511165 | 24 | 31 | - | CTTATCTC |
| MA0711.1 | MA0711.1.OTX1 | 3.0480316 | 0.8187575 | 24 | 31 | - | CTTATCTC |
| MA0714.1 | MA0714.1.PITX3 | 4.5250573 | 0.8052406 | 24 | 32 | - | GCTTATCTC |
| MA0877.2 | MA0877.2.BARHL1 | 3.2366128 | 0.8243014 | 24 | 31 | - | CTTATCTC |
| MA1480.1 | MA1480.1.DPRX | 9.67678 | 0.8934484 | 24 | 33 | - | AGCTTATCTC |
| MA1480.1 | MA1480.1.DPRX | 9.67678 | 0.8934484 | 24 | 33 | - | AGCTTATCTC |
| MA0766.3 | MA0766.3.GATA5 | 11.098037 | 0.97757965 | 25 | 32 | + | AGATAAGC |
| MA0766.1 | MA0766.1.GATA5 | 10.12808 | 0.97034866 | 25 | 32 | + | AGATAAGC |
| MA0719.2 | MA0719.2.RHOXF1 | 5.331678 | 0.91555077 | 25 | 30 | - | TTATCT |
| MA0711.2 | MA0711.2.OTX1 | 2.8238444 | 0.82846344 | 25 | 30 | - | TTATCT |
| MA1104.3 | MA1104.3.GATA6 | 11.393329 | 0.93583244 | 25 | 32 | - | GCTTATCT |
| MA0891.2 | MA0891.2.GSC2 | 4.9261847 | 0.8395931 | 25 | 30 | - | TTATCT |
| MA1480.2 | MA1480.2.DPRX | 9.208557 | 0.8913453 | 25 | 33 | - | AGCTTATCT |
| MA1480.2 | MA1480.2.DPRX | 7.9756775 | 0.8666903 | 25 | 33 | + | AGATAAGCT |
| MA1480.1 | MA1480.1.DPRX | 8.058306 | 0.8617105 | 25 | 34 | + | AGATAAGCTG |
| MA0482.3 | MA0482.3.GATA4 | 11.500963 | 0.93496716 | 25 | 32 | - | GCTTATCT |
| MA0635.2 | MA0635.2.BARHL2 | 2.5784996 | 0.8401126 | 25 | 30 | - | TTATCT |
| MA0036.4 | MA0036.4.GATA2 | 12.923377 | 1 | 25 | 31 | - | CTTATCT |
| MA0036.1 | MA0036.1.GATA2 | 6.2320204 | 0.9756508 | 25 | 29 | + | AGATA |
| MA0037.3 | MA0037.3.GATA3 | 10.987059 | 0.9738492 | 25 | 32 | + | AGATAAGC |
| MA0035.5 | MA0035.5.GATA1 | 11.376707 | 0.9693649 | 25 | 31 | - | CTTATCT |
| MA0037.2 | MA0037.2.GATA3 | 12.732875 | 0.96934724 | 25 | 32 | + | AGATAAGC |
| MA0037.1 | MA0037.1.GATA3 | 7.1894307 | 0.9497132 | 25 | 30 | + | AGATAA |
| MA1480.2 | MA1480.2.DPRX | 9.208557 | 0.8913453 | 25 | 33 | - | AGCTTATCT |
| MA1480.2 | MA1480.2.DPRX | 7.9756775 | 0.8666903 | 25 | 33 | + | AGATAAGCT |
| MA1480.1 | MA1480.1.DPRX | 8.058306 | 0.8617105 | 25 | 34 | + | AGATAAGCTG |
| MA1970.2 | MA1970.2.TRPS1 | 11.337377 | 0.92992425 | 25 | 32 | - | GCTTATCT |
| MA0037.5 | MA0037.5.Gata3 | 11.528673 | 0.9351444 | 25 | 32 | - | GCTTATCT |
| MA0035.1 | MA0035.1.Gata1 | 4.4595475 | 0.8560758 | 25 | 30 | + | AGATAA |
| MA0714.1 | MA0714.1.PITX3 | 4.741932 | 0.8103673 | 26 | 34 | + | GATAAGCTG |
| MA0842.1 | MA0842.1.NRL | 5.721393 | 0.8246058 | 26 | 36 | + | GATAAGCTGCC |
| MA0891.1 | MA0891.1.GSC2 | 4.9547377 | 0.81051326 | 26 | 35 | + | GATAAGCTGC |
| MA0635.1 | MA0635.1.BARHL2 | 4.391176 | 0.8483359 | 26 | 35 | + | GATAAGCTGC |
| MA0500.1 | MA0500.1.Myog | 3.316181 | 0.8384537 | 26 | 36 | - | GGCAGCTTATC |
| MA0521.1 | MA0521.1.Tcf12 | 3.1155736 | 0.8243787 | 26 | 36 | - | GGCAGCTTATC |
| MA0719.1 | MA0719.1.RHOXF1 | 6.152899 | 0.9228801 | 27 | 34 | + | ATAAGCTG |
| MA0711.1 | MA0711.1.OTX1 | 5.932823 | 0.8749971 | 27 | 34 | + | ATAAGCTG |
| MA0712.1 | MA0712.1.OTX2 | 5.7966957 | 0.86642426 | 27 | 34 | + | ATAAGCTG |
| MA0682.2 | MA0682.2.PITX1 | 3.2264004 | 0.814453 | 27 | 34 | + | ATAAGCTG |
| MA0877.2 | MA0877.2.BARHL1 | 2.9380724 | 0.8175352 | 27 | 34 | + | ATAAGCTG |
| MA1472.1 | MA1472.1.BHLHA15 | 4.520576 | 0.8114588 | 27 | 36 | + | ATAAGCTGCC |
| MA1472.1 | MA1472.1.BHLHA15 | 4.520576 | 0.8114588 | 27 | 36 | + | ATAAGCTGCC |
| MA0499.1 | MA0499.1.Myod1 | 4.0075607 | 0.84173006 | 27 | 39 | + | ATAAGCTGCCGCT |
| MA0063.1 | MA0063.1.Nkx2-5 | 4.9718757 | 0.8250539 | 27 | 33 | + | ATAAGCT |
| MA0633.2 | MA0633.2.Twist2 | 1.7549466 | 0.8184241 | 27 | 36 | + | ATAAGCTGCC |
| MA0633.2 | MA0633.2.Twist2 | 1.7549466 | 0.8184241 | 27 | 36 | - | GGCAGCTTAT |
| MA0467.1 | MA0467.1.Crx | 5.248869 | 0.81226134 | 27 | 37 | - | CGGCAGCTTAT |
| MA0522.1 | MA0522.1.Tcf3 | 1.7334361 | 0.8083049 | 27 | 37 | + | ATAAGCTGCCG |
| MA0682.1 | MA0682.1.Pitx1 | 4.563924 | 0.8082841 | 27 | 34 | + | ATAAGCTG |
| MA1997.1 | MA1997.1.Olig2 | 1.6292045 | 0.8175178 | 27 | 36 | + | ATAAGCTGCC |
| MA1997.1 | MA1997.1.Olig2 | 1.6292045 | 0.8175178 | 27 | 36 | - | GGCAGCTTAT |
| MA1993.1 | MA1993.1.Neurod2 | 1.4584854 | 0.81585103 | 27 | 36 | + | ATAAGCTGCC |
| MA1993.1 | MA1993.1.Neurod2 | 1.4584854 | 0.81585103 | 27 | 36 | - | GGCAGCTTAT |
| MA1684.1 | MA1684.1.Foxn1 | 0.7151981 | 0.8022139 | 27 | 32 | + | ATAAGC |

| | | | | | | | |
|---|---|---|---|---|---|---|---|
| MA0719.2 | MA0719.2.RHOXF1 | 6.2764587 | 0.9343377 | 28 | 33 | + | TAAGCT |
| MA0711.2 | MA0711.2.OTX1 | 5.2761655 | 0.88030976 | 28 | 33 | + | TAAGCT |
| MA0714.2 | MA0714.2.PITX3 | 5.7855716 | 0.863991 | 28 | 33 | + | TAAGCT |
| MA0682.3 | MA0682.3.PITX1 | 4.152888 | 0.85010177 | 28 | 33 | + | TAAGCT |
| MA0891.2 | MA0891.2.GSC2 | 4.4868627 | 0.82658464 | 28 | 33 | + | TAAGCT |
| MA0877.4 | MA0877.4.BARHL1 | 2.9996986 | 0.8088255 | 28 | 33 | - | AGCTTA |
| MA0635.2 | MA0635.2.BARHL2 | 3.4035308 | 0.85749346 | 28 | 33 | + | TAAGCT |
| MA0648.2 | MA0648.2.GSC | 3.335787 | 0.8019492 | 28 | 33 | + | TAAGCT |
| MA0100.2 | MA0100.2.Myb | 4.0202394 | 0.8139977 | 28 | 37 | + | TAAGCTGCCG |
| MA0164.1 | MA0164.1.Nr2e3 | 6.2397103 | 0.8061964 | 28 | 34 | + | TAAGCTG |
| MA0820.2 | MA0820.2.FIGLA | 4.7212787 | 0.85025436 | 29 | 34 | - | CAGCTT |
| MA1583.1 | MA1583.1.ZFP57 | 6.966908 | 0.8451742 | 29 | 41 | + | AAGCTGCCGCTAA |
| MA1583.1 | MA1583.1.ZFP57 | 6.966908 | 0.8451742 | 29 | 41 | + | AAGCTGCCGCTAA |
| MA1635.2 | MA1635.2.BHLHE22 | 2.8410106 | 0.8330293 | 29 | 34 | - | CAGCTT |
| MA1635.2 | MA1635.2.BHLHE22 | 2.8406963 | 0.8330236 | 29 | 34 | + | AAGCTG |
| MA0633.3 | MA0633.3.Twist2 | 2.465053 | 0.8359876 | 29 | 34 | + | AAGCTG |
| MA0633.3 | MA0633.3.Twist2 | 2.465053 | 0.8359876 | 29 | 34 | - | CAGCTT |
| MA0164.2 | MA0164.2.Nr2e3 | 6.4044814 | 0.83294225 | 29 | 34 | + | AAGCTG |
| MA0164.2 | MA0164.2.Nr2e3 | 6.4044814 | 0.83294225 | 29 | 34 | - | CAGCTT |
| MA0521.3 | MA0521.3.Tcf12 | 4.325149 | 0.8256369 | 29 | 34 | - | CAGCTT |
| MA0521.3 | MA0521.3.Tcf12 | 4.302039 | 0.82508963 | 29 | 34 | + | AAGCTG |
| MA1993.2 | MA1993.2.Neurod2 | 2.3762918 | 0.8357511 | 29 | 34 | + | AAGCTG |
| MA1993.2 | MA1993.2.Neurod2 | 2.3762918 | 0.8357511 | 29 | 34 | - | CAGCTT |
| MA1997.2 | MA1997.2.Olig2 | 2.1639824 | 0.83396477 | 29 | 34 | + | AAGCTG |
| MA1997.2 | MA1997.2.Olig2 | 2.1639824 | 0.83396477 | 29 | 34 | - | CAGCTT |
| MA1542.1 | MA1542.1.OSR1 | 4.7005806 | 0.82084703 | 30 | 39 | + | AGCTGCCGCT |
| MA0470.1 | MA0470.1.E2F4 | 4.232801 | 0.8118414 | 30 | 40 | - | TAGCGGCAGCT |
| MA0471.1 | MA0471.1.E2F6 | 3.3093023 | 0.80984664 | 30 | 40 | - | TAGCGGCAGCT |
| MA1542.1 | MA1542.1.OSR1 | 4.7005806 | 0.82084703 | 30 | 39 | + | AGCTGCCGCT |
| MA1728.2 | MA1728.2.ZNF549 | 6.207406 | 0.8080607 | 30 | 37 | + | AGCTGCCG |
| MA0816.1 | MA0816.1.Ascl2 | 6.004277 | 0.8005746 | 30 | 39 | + | AGCTGCCGCT |
| MA0671.1 | MA0671.1.NFIX | 4.2065425 | 0.84453857 | 31 | 39 | - | AGCGGCAGC |
| MA0764.4 | MA0764.4.ETV4 | 4.3732667 | 0.8092915 | 31 | 39 | - | AGCGGCAGC |
| MA0759.3 | MA0759.3.ELK3 | 3.7454429 | 0.8062963 | 31 | 39 | - | AGCGGCAGC |
| MA0765.4 | MA0765.4.ETV5 | 3.296537 | 0.8006485 | 31 | 39 | - | AGCGGCAGC |
| MA1484.2 | MA1484.2.ETS2 | 3.3178673 | 0.83800584 | 31 | 39 | - | AGCGGCAGC |
| MA1484.1 | MA1484.1.ETS2 | 2.8134139 | 0.8277422 | 31 | 40 | - | TAGCGGCAGC |
| MA1542.2 | MA1542.2.OSR1 | 4.4538064 | 0.8260085 | 31 | 38 | + | GCTGCCGC |
| MA0024.2 | MA0024.2.E2F1 | 8.938323 | 0.9024963 | 31 | 41 | - | TTAGCGGCAGC |
| MA1484.2 | MA1484.2.ETS2 | 3.3178673 | 0.83800584 | 31 | 39 | - | AGCGGCAGC |
| MA1484.1 | MA1484.1.ETS2 | 2.8134139 | 0.8277422 | 31 | 40 | - | TAGCGGCAGC |
| MA1542.2 | MA1542.2.OSR1 | 4.4538064 | 0.8260085 | 31 | 38 | + | GCTGCCGC |
| MA1684.1 | MA1684.1.Foxn1 | 1.5127487 | 0.81909716 | 31 | 36 | - | GGCAGC |
| MA0671.2 | MA0671.2.NFIX | 3.716774 | 0.8420332 | 32 | 37 | - | CGGCAG |
| MA0865.3 | MA0865.3.E2F8 | 7.7640343 | 0.85034144 | 32 | 40 | + | CTGCCGCTA |
| MA0865.2 | MA0865.2.E2F8 | 7.0703626 | 0.8050668 | 32 | 43 | + | CTGCCGCTAATG |
| MA1536.1 | MA1536.1.NR2C2 | -0.18933833 | 0.8097495 | 32 | 39 | - | AGCGGCAG |
| MA0498.2 | MA0498.2.MEIS1 | 1.1931098 | 0.8060117 | 32 | 38 | + | CTGCCGC |
| MA0080.1 | MA0080.1.SPI1 | 4.3853064 | 0.81848 | 32 | 37 | - | CGGCAG |
| MA1536.1 | MA1536.1.NR2C2 | -0.18933833 | 0.8097495 | 32 | 39 | - | AGCGGCAG |
| MA1721.2 | MA1721.2.ZNF93 | 8.587485 | 0.8059193 | 32 | 45 | - | GCCATTAGCGGCAG |
| MA1583.2 | MA1583.2.ZFP57 | 8.307053 | 0.90036076 | 33 | 39 | + | TGCCGCT |
| MA1554.2 | MA1554.2.RFX7 | 4.3269086 | 0.81658685 | 33 | 40 | + | TGCCGCTA |
| MA1536.2 | MA1536.2.NR2C2 | -0.26876622 | 0.8130445 | 33 | 38 | - | GCGGCA |
| MA0498.3 | MA0498.3.MEIS1 | 1.6050415 | 0.82704395 | 33 | 37 | + | TGCCG |
| MA0081.1 | MA0081.1.SPIB | 5.103546 | 0.81667465 | 33 | 39 | - | AGCGGCA |
| MA1583.2 | MA1583.2.ZFP57 | 8.307053 | 0.90036076 | 33 | 39 | + | TGCCGCT |
| MA1554.2 | MA1554.2.RFX7 | 4.3269086 | 0.81658685 | 33 | 40 | + | TGCCGCTA |
| MA1536.2 | MA1536.2.NR2C2 | -0.26876622 | 0.8130445 | 33 | 38 | - | GCGGCA |
| MA1684.1 | MA1684.1.Foxn1 | 0.81124043 | 0.804247 | 33 | 38 | + | TGCCGC |
| MA0776.1 | MA0776.1.MYBL1 | 5.8928413 | 0.8107634 | 34 | 45 | - | GCCATTAGCGGC |
| MA0893.3 | MA0893.3.GSX2 | 1.4522122 | 0.8111782 | 34 | 40 | + | GCCGCTA |
| MA0893.2 | MA0893.2.GSX2 | 1.8419366 | 0.8109837 | 34 | 41 | + | GCCGCTAA |
| MA1545.2 | MA1545.2.OVOL2 | 5.694909 | 0.870813 | 34 | 40 | + | GCCGCTA |
| MA0635.1 | MA0635.1.BARHL2 | 4.601733 | 0.8522556 | 34 | 43 | - | CATTAGCGGC |
| MA0594.2 | MA0594.2.HOXA9 | 4.811472 | 0.835043 | 34 | 43 | + | GCCGCTAATG |
| MA0594.3 | MA0594.3.HOXA9 | 3.5463839 | 0.820725 | 34 | 40 | + | GCCGCTA |
| MA1545.2 | MA1545.2.OVOL2 | 5.694909 | 0.870813 | 34 | 40 | + | GCCGCTA |
| MA0877.2 | MA0877.2.BARHL1 | 5.1086016 | 0.8667282 | 35 | 42 | - | ATTAGCGG |
| MA0145.1 | MA0145.1.Tfcp2l1 | 6.395767 | 0.824612 | 35 | 48 | - | CCGGCCATTAGCGG |
| MA0145.2 | MA0145.2.Tfcp2l1 | 6.3816233 | 0.8240674 | 35 | 48 | - | CCGGCCATTAGCGG |
| MA0635.2 | MA0635.2.BARHL2 | 3.9533534 | 0.86907655 | 36 | 41 | - | TTAGCG |
| MA0594.2 | MA0594.2.HOXA9 | 5.480538 | 0.8510605 | 36 | 45 | - | GCCATTAGCG |
| MA0158.1 | MA0158.1.HOXA5 | 8.92601 | 0.96724313 | 36 | 43 | + | CGCTAATG |
| MA0158.1 | MA0158.1.HOXA5 | 8.92601 | 0.96724313 | 36 | 43 | + | CGCTAATG |
| MA1608.1 | MA1608.1.Isl1 | 12.999704 | 0.98265874 | 36 | 46 | - | GGCCATTAGCG |

| | | | | | | | |
|---|---|---|---|---|---|---|---|
| MA1476.2 | MA1476.2.Dlx5 | 8.996206 | 0.8622528 | 36 | 47 | - | CGGCCATTAGCG |
| MA0885.2 | MA0885.2.Dlx2 | 9.04979 | 0.86183226 | 36 | 47 | - | CGGCCATTAGCG |
| MA0706.1 | MA0706.1.MEOX2 | 7.232636 | 0.8721193 | 37 | 46 | - | GGCCATTAGC |
| MA0699.1 | MA0699.1.LBX2 | 7.249927 | 0.8719977 | 37 | 46 | - | GGCCATTAGC |
| MA0707.1 | MA0707.1.MNX1 | 7.057372 | 0.870884 | 37 | 46 | - | GGCCATTAGC |
| MA0700.1 | MA0700.1.LHX2 | 6.2097645 | 0.8672761 | 37 | 46 | - | GGCCATTAGC |
| MA0700.1 | MA0700.1.LHX2 | 5.909695 | 0.8622129 | 37 | 46 | + | GCTAATGGCC |
| MA0710.1 | MA0710.1.NOTO | 5.052614 | 0.85707986 | 37 | 46 | - | GGCCATTAGC |
| MA0710.1 | MA0710.1.NOTO | 3.2498431 | 0.8251998 | 37 | 46 | + | GCTAATGGCC |
| MA0718.1 | MA0718.1.RAX | 4.895288 | 0.82104796 | 37 | 46 | - | GGCCATTAGC |
| MA0706.1 | MA0706.1.MEOX2 | 5.0719204 | 0.81809783 | 37 | 46 | + | GCTAATGGCC |
| MA0903.1 | MA0903.1.HOXB3 | 9.5972395 | 0.95731235 | 37 | 46 | - | GGCCATTAGC |
| MA0900.1 | MA0900.1.HOXA2 | 9.365544 | 0.942174 | 37 | 46 | - | GGCCATTAGC |
| MA0902.1 | MA0902.1.HOXB2 | 8.875398 | 0.94153196 | 37 | 46 | - | GGCCATTAGC |
| MA0888.1 | MA0888.1.EVX2 | 9.458034 | 0.93860126 | 37 | 46 | - | GGCCATTAGC |
| MA0887.1 | MA0887.1.EVX1 | 9.304435 | 0.9343212 | 37 | 46 | - | GGCCATTAGC |
| MA0886.1 | MA0886.1.EMX2 | 8.429402 | 0.9128133 | 37 | 46 | - | GGCCATTAGC |
| MA0893.1 | MA0893.1.GSX2 | 7.497275 | 0.88630205 | 37 | 46 | - | GGCCATTAGC |
| MA0892.1 | MA0892.1.GSX1 | 7.047585 | 0.88495576 | 37 | 46 | - | GGCCATTAGC |
| MA0890.1 | MA0890.1.GBX2 | 6.975975 | 0.880332 | 37 | 46 | - | GGCCATTAGC |
| MA0889.1 | MA0889.1.GBX1 | 7.1567607 | 0.87960553 | 37 | 46 | - | GGCCATTAGC |
| MA0891.1 | MA0891.1.GSC2 | 6.3296757 | 0.84438556 | 37 | 46 | + | GCTAATGGCC |
| MA0894.1 | MA0894.1.HESX1 | 4.349231 | 0.8251787 | 37 | 46 | + | GCTAATGGCC |
| MA0878.1 | MA0878.1.CDX1 | 3.881263 | 0.8067364 | 37 | 45 | - | GCCATTAGC |
| MA1497.1 | MA1497.1.HOXA6 | 9.991935 | 0.94672376 | 37 | 46 | - | GGCCATTAGC |
| MA1501.1 | MA1501.1.HOXB7 | 11.142902 | 0.9374041 | 37 | 46 | - | GGCCATTAGC |
| MA1530.1 | MA1530.1.NKX6-3 | 7.5420613 | 0.89850914 | 37 | 45 | - | GCCATTAGC |
| MA1542.2 | MA1542.2.OSR1 | 3.589477 | 0.81068575 | 37 | 44 | + | GCTAATGG |
| MA0642.1 | MA0642.1.EN2 | 9.23017 | 0.925407 | 37 | 46 | - | GGCCATTAGC |
| MA0612.1 | MA0612.1.EMX1 | 9.162516 | 0.92056274 | 37 | 46 | + | GCTAATGGCC |
| MA0661.1 | MA0661.1.MEOX1 | 7.341605 | 0.8768736 | 37 | 46 | - | GGCCATTAGC |
| MA0661.1 | MA0661.1.MEOX1 | 7.0138206 | 0.87012017 | 37 | 46 | + | GCTAATGGCC |
| MA0662.1 | MA0662.1.MIXL1 | 6.037306 | 0.86234075 | 37 | 46 | - | GGCCATTAGC |
| MA0644.1 | MA0644.1.ESX1 | 6.5525856 | 0.8600237 | 37 | 46 | - | GGCCATTAGC |
| MA0648.1 | MA0648.1.GSC | 6.9668217 | 0.85660744 | 37 | 46 | + | GCTAATGGCC |
| MA0635.1 | MA0635.1.BARHL2 | 2.284319 | 0.8091163 | 37 | 46 | + | GCTAATGGCC |
| MA0658.1 | MA0658.1.LHX6 | 3.1053588 | 0.8019141 | 37 | 46 | - | GGCCATTAGC |
| MA0658.1 | MA0658.1.LHX6 | 3.0916517 | 0.80167973 | 37 | 46 | + | GCTAATGGCC |
| MA0642.1 | MA0642.1.EN2 | 3.456605 | 0.80029017 | 37 | 46 | + | GCTAATGGCC |
| MA0003.1 | MA0003.1.TFAP2A | 6.1033154 | 0.86005676 | 37 | 45 | - | GCCATTAGC |
| MA1497.1 | MA1497.1.HOXA6 | 9.991935 | 0.94672376 | 37 | 46 | - | GGCCATTAGC |
| MA1501.1 | MA1501.1.HOXB7 | 11.142902 | 0.9374041 | 37 | 46 | - | GGCCATTAGC |
| MA1530.1 | MA1530.1.NKX6-3 | 7.5420613 | 0.89850914 | 37 | 45 | - | GCCATTAGC |
| MA1542.2 | MA1542.2.OSR1 | 3.589477 | 0.81068575 | 37 | 44 | + | GCTAATGG |
| MA0877.1 | MA0877.1.Barhl1 | 9.096096 | 0.9451992 | 37 | 46 | + | GCTAATGGCC |
| MA0879.1 | MA0879.1.Dlx1 | 5.7487025 | 0.8494294 | 37 | 46 | - | GGCCATTAGC |
| MA0723.2 | MA0723.2.VAX2 | 8.624679 | 0.9616891 | 38 | 45 | - | GCCATTAG |
| MA0707.2 | MA0707.2.MNX1 | 7.44623 | 0.93537104 | 38 | 45 | - | GCCATTAG |
| MA0723.1 | MA0723.1.VAX2 | 8.348263 | 0.9147653 | 38 | 45 | + | CTAATGGC |
| MA0701.1 | MA0701.1.LHX9 | 7.5998907 | 0.9094126 | 38 | 45 | - | GCCATTAG |
| MA0724.1 | MA0724.1.VENTX | 8.437543 | 0.9062262 | 38 | 46 | - | GGCCATTAG |
| MA0722.1 | MA0722.1.VAX1 | 7.885804 | 0.90396327 | 38 | 45 | + | CTAATGGC |
| MA0722.2 | MA0722.2.VAX1 | 7.3509865 | 0.90105736 | 38 | 44 | + | CTAATGG |
| MA0675.1 | MA0675.1.NKX6-2 | 7.017501 | 0.89382416 | 38 | 45 | - | GCCATTAG |
| MA0708.2 | MA0708.2.MSX2 | 6.2681227 | 0.8831076 | 38 | 45 | - | GCCATTAG |
| MA0674.1 | MA0674.1.NKX6-1 | 6.568898 | 0.87961 | 38 | 45 | - | GCCATTAG |
| MA0674.2 | MA0674.2.NKX6-1 | 6.168579 | 0.87709564 | 38 | 44 | - | CCATTAG |
| MA0708.1 | MA0708.1.MSX2 | 6.9147024 | 0.86876386 | 38 | 45 | - | GCCATTAG |
| MA0701.2 | MA0701.2.LHX9 | 5.940269 | 0.86735207 | 38 | 45 | - | GCCATTAG |
| MA0717.1 | MA0717.1.RAX2 | 6.014216 | 0.85673773 | 38 | 45 | - | GCCATTAG |
| MA0711.1 | MA0711.1.OTX1 | 4.661749 | 0.8502173 | 38 | 45 | + | CTAATGGC |
| MA0706.2 | MA0706.2.MEOX2 | 5.43227 | 0.8500467 | 38 | 44 | + | CTAATGG |
| MA0716.1 | MA0716.1.PRRX1 | 5.2605414 | 0.83667934 | 38 | 45 | - | GCCATTAG |
| MA0725.1 | MA0725.1.VSX1 | 5.9511294 | 0.8362451 | 38 | 45 | + | CTAATGGC |
| MA0725.2 | MA0725.2.VSX1 | 5.8843107 | 0.835849 | 38 | 44 | + | CTAATGG |
| MA0710.2 | MA0710.2.NOTO | 2.9112473 | 0.8329023 | 38 | 44 | + | CTAATGG |
| MA0721.1 | MA0721.1.UNCX | 5.2314086 | 0.83192974 | 38 | 45 | - | GCCATTAG |
| MA0701.3 | MA0701.3.LHX9 | 3.3176491 | 0.82475287 | 38 | 44 | + | CTAATGG |
| MA0703.1 | MA0703.1.LMX1B | 4.722879 | 0.81042475 | 38 | 45 | - | GCCATTAG |
| MA0701.2 | MA0701.2.LHX9 | 2.7418666 | 0.8082178 | 38 | 45 | + | CTAATGGC |
| MA0726.2 | MA0726.2.VSX2 | 4.723573 | 0.80592513 | 38 | 44 | + | CTAATGG |
| MA0701.1 | MA0701.1.LHX9 | 3.6034558 | 0.8000963 | 38 | 45 | + | CTAATGGC |
| MA0875.1 | MA0875.1.BARX1 | 10.402158 | 0.9836232 | 38 | 45 | - | GCCATTAG |
| MA0893.2 | MA0893.2.GSX2 | 10.181031 | 0.9753747 | 38 | 45 | - | GCCATTAG |
| MA0902.2 | MA0902.2.HOXB2 | 10.201051 | 0.9709236 | 38 | 45 | - | GCCATTAG |
| MA0876.1 | MA0876.1.BSX | 9.292447 | 0.96336514 | 38 | 45 | - | GCCATTAG |

| | | | | | | | |
|---|---|---|---|---|---|---|---|
| MA0910.2 | MA0910.2.HOXD8 | 10.36608 | 0.96334547 | 38 | 45 | - | GCCATTAG |
| MA0879.2 | MA0879.2.DLX1 | 8.432682 | 0.94431704 | 38 | 45 | - | GCCATTAG |
| MA0912.2 | MA0912.2.HOXD3 | 9.725193 | 0.92887 | 38 | 45 | + | CTAATGGC |
| MA0904.2 | MA0904.2.HOXB5 | 7.8296103 | 0.917769 | 38 | 45 | + | CTAATGGC |
| MA0904.2 | MA0904.2.HOXB5 | 7.814363 | 0.91746426 | 38 | 45 | - | GCCATTAG |
| MA0900.2 | MA0900.2.HOXA2 | 7.4725647 | 0.91739 | 38 | 45 | - | GCCATTAG |
| MA0882.1 | MA0882.1.DLX6 | 6.6399493 | 0.88533807 | 38 | 45 | - | GCCATTAG |
| MA0893.3 | MA0893.3.GSX2 | 2.3971798 | 0.8305796 | 38 | 44 | + | CTAATGG |
| MA0914.1 | MA0914.1.ISL2 | 5.4609494 | 0.82956725 | 38 | 45 | - | GCCATTAG |
| MA1500.1 | MA1500.1.HOXB6 | 10.047016 | 0.9749131 | 38 | 45 | + | CTAATGGC |
| MA1498.1 | MA1498.1.HOXA7 | 10.364698 | 0.9746022 | 38 | 45 | - | GCCATTAG |
| MA1496.1 | MA1496.1.HOXA4 | 9.8640785 | 0.9730046 | 38 | 45 | - | GCCATTAG |
| MA1502.1 | MA1502.1.HOXB8 | 10.510669 | 0.9718083 | 38 | 45 | - | GCCATTAG |
| MA1504.1 | MA1504.1.HOXC4 | 9.6640215 | 0.9708456 | 38 | 45 | - | GCCATTAG |
| MA1507.1 | MA1507.1.HOXD4 | 9.792811 | 0.9683188 | 38 | 45 | - | GCCATTAG |
| MA1498.2 | MA1498.2.HOXA7 | 9.368576 | 0.96629936 | 38 | 45 | - | GCCATTAG |
| MA1499.1 | MA1499.1.HOXB4 | 9.975273 | 0.95963824 | 38 | 45 | - | GCCATTAG |
| MA1518.1 | MA1518.1.LHX1 | 8.183356 | 0.94109136 | 38 | 45 | - | GCCATTAG |
| MA1495.1 | MA1495.1.HOXA1 | 7.2372394 | 0.9204784 | 38 | 45 | + | CTAATGGC |
| MA1505.1 | MA1505.1.HOXC8 | 7.07423 | 0.9177366 | 38 | 45 | + | CTAATGGC |
| MA1505.1 | MA1505.1.HOXC8 | 7.06026 | 0.91745883 | 38 | 45 | - | GCCATTAG |
| MA1495.1 | MA1495.1.HOXA1 | 7.0106645 | 0.9161489 | 38 | 45 | - | GCCATTAG |
| MA1530.2 | MA1530.2.NKX6-3 | 7.8708324 | 0.9156398 | 38 | 45 | - | GCCATTAG |
| MA1519.1 | MA1519.1.LHX5 | 5.4537063 | 0.86577976 | 38 | 45 | - | GCCATTAG |
| MA1476.1 | MA1476.1.DLX5 | 4.7902765 | 0.86487544 | 38 | 45 | - | GCCATTAG |
| MA1481.1 | MA1481.1.DRGX | 4.524179 | 0.8566995 | 38 | 45 | - | GCCATTAG |
| MA1500.2 | MA1500.2.HOXB6 | 2.1012437 | 0.81547 | 38 | 44 | - | CCATTAG |
| MA1577.1 | MA1577.1.TLX2 | 3.0535464 | 0.8146359 | 38 | 45 | - | GCCATTAG |
| MA1500.1 | MA1500.1.HOXB6 | 2.048446 | 0.8073234 | 38 | 45 | - | GCCATTAG |
| MA1496.2 | MA1496.2.HOXA4 | 1.2899168 | 0.803513 | 38 | 44 | + | CTAATGG |
| MA1481.1 | MA1481.1.DRGX | 1.8444849 | 0.8031982 | 38 | 45 | + | CTAATGGC |
| MA0612.2 | MA0612.2.EMX1 | 7.9054494 | 0.91290957 | 38 | 45 | - | GCCATTAG |
| MA0666.1 | MA0666.1.MSX1 | 7.8345776 | 0.9035045 | 38 | 45 | - | GCCATTAG |
| MA0666.2 | MA0666.2.MSX1 | 6.174872 | 0.8770587 | 38 | 45 | - | GCCATTAG |
| MA0630.1 | MA0630.1.SHOX | 6.4822965 | 0.86120456 | 38 | 45 | + | CTAATGGC |
| MA0661.2 | MA0661.2.MEOX1 | 5.5582066 | 0.86074674 | 38 | 44 | + | CTAATGG |
| MA0654.1 | MA0654.1.ISX | 5.5217304 | 0.858669 | 38 | 45 | - | GCCATTAG |
| MA0644.2 | MA0644.2.ESX1 | 4.2961063 | 0.85249496 | 38 | 45 | - | GCCATTAG |
| MA0618.1 | MA0618.1.LBX1 | 4.4627457 | 0.81829864 | 38 | 45 | + | CTAATGGC |
| MA0502.3 | MA0502.3.NFYB | 5.680759 | 0.8060634 | 38 | 46 | + | CTAATGGCC |
| MA0644.3 | MA0644.3.ESX1 | 1.3601639 | 0.8019592 | 38 | 44 | + | CTAATGG |
| MA0132.2 | MA0132.2.PDX1 | 9.789959 | 0.96033883 | 38 | 45 | - | GCCATTAG |
| MA0158.2 | MA0158.2.HOXA5 | 10.092203 | 0.957795 | 38 | 45 | - | GCCATTAG |
| MA0027.2 | MA0027.2.EN1 | 6.3650007 | 0.8798867 | 38 | 45 | - | GCCATTAG |
| MA0158.2 | MA0158.2.HOXA5 | 1.7485955 | 0.80267864 | 38 | 45 | + | CTAATGGC |
| MA2119.1 | MA2119.1.HOXA3 | 6.473275 | 0.89610463 | 38 | 44 | + | CTAATGG |
| MA1500.1 | MA1500.1.HOXB6 | 10.047016 | 0.9749131 | 38 | 45 | + | CTAATGGC |
| MA1498.1 | MA1498.1.HOXA7 | 10.364698 | 0.9746022 | 38 | 45 | - | GCCATTAG |
| MA1496.1 | MA1496.1.HOXA4 | 9.8640785 | 0.9730046 | 38 | 45 | - | GCCATTAG |
| MA1502.1 | MA1502.1.HOXB8 | 10.510669 | 0.9718083 | 38 | 45 | - | GCCATTAG |
| MA1504.1 | MA1504.1.HOXC4 | 9.6640215 | 0.9708456 | 38 | 45 | - | GCCATTAG |
| MA1507.1 | MA1507.1.HOXD4 | 9.792811 | 0.9683188 | 38 | 45 | - | GCCATTAG |
| MA1498.2 | MA1498.2.HOXA7 | 9.368576 | 0.96629936 | 38 | 45 | - | GCCATTAG |
| MA1499.1 | MA1499.1.HOXB4 | 9.975273 | 0.95963824 | 38 | 45 | - | GCCATTAG |
| MA1518.1 | MA1518.1.LHX1 | 8.183356 | 0.94109136 | 38 | 45 | - | GCCATTAG |
| MA1495.1 | MA1495.1.HOXA1 | 7.2372394 | 0.9204784 | 38 | 45 | + | CTAATGGC |
| MA1505.1 | MA1505.1.HOXC8 | 7.07423 | 0.9177366 | 38 | 45 | + | CTAATGGC |
| MA1505.1 | MA1505.1.HOXC8 | 7.06026 | 0.91745883 | 38 | 45 | - | GCCATTAG |
| MA1495.1 | MA1495.1.HOXA1 | 7.0106645 | 0.9161489 | 38 | 45 | - | GCCATTAG |
| MA1530.2 | MA1530.2.NKX6-3 | 7.8708324 | 0.9156398 | 38 | 45 | - | GCCATTAG |
| MA1519.1 | MA1519.1.LHX5 | 5.4537063 | 0.86577976 | 38 | 45 | - | GCCATTAG |
| MA1476.1 | MA1476.1.DLX5 | 4.7902765 | 0.86487544 | 38 | 45 | - | GCCATTAG |
| MA1481.1 | MA1481.1.DRGX | 4.524179 | 0.8566995 | 38 | 45 | - | GCCATTAG |
| MA1500.2 | MA1500.2.HOXB6 | 2.1012437 | 0.81547 | 38 | 44 | - | CCATTAG |
| MA1577.1 | MA1577.1.TLX2 | 3.0535464 | 0.8146359 | 38 | 45 | - | GCCATTAG |
| MA1500.1 | MA1500.1.HOXB6 | 2.048446 | 0.8073234 | 38 | 45 | - | GCCATTAG |
| MA1496.2 | MA1496.2.HOXA4 | 1.2899168 | 0.803513 | 38 | 44 | + | CTAATGG |
| MA1481.1 | MA1481.1.DRGX | 1.8444849 | 0.8031982 | 38 | 45 | + | CTAATGGC |
| MA2093.1 | MA2093.1.HOXB1 | 6.3045144 | 0.8943404 | 38 | 44 | + | CTAATGG |
| MA1963.2 | MA1963.2.SATB1 | 3.635183 | 0.80131817 | 38 | 44 | + | CTAATGG |
| MA0132.1 | MA0132.1.Pdx1 | 8.729877 | 0.9654329 | 38 | 43 | + | CTAATG |
| MA0709.1 | MA0709.1.Msx3 | 6.141421 | 0.8670296 | 38 | 45 | - | GCCATTAG |
| MA0720.1 | MA0720.1.Shox2 | 5.6885877 | 0.8638926 | 38 | 45 | - | GCCATTAG |
| MA0704.1 | MA0704.1.Lhx4 | 5.8006396 | 0.8577672 | 38 | 45 | - | GCCATTAG |
| MA0705.1 | MA0705.1.Lhx8 | 4.154091 | 0.85569835 | 38 | 45 | - | GCCATTAG |
| MA0705.1 | MA0705.1.Lhx8 | 3.9426327 | 0.8517041 | 38 | 45 | + | CTAATGGC |

| | | | | | | | |
|---|---|---|---|---|---|---|---|
| MA0075.2 | MA0075.2.Prrx2 | 5.9703703 | 0.850297 | 38 | 45 | - | GCCATTAG |
| MA1608.2 | MA1608.2.Isl1 | 12.6508465 | 1 | 38 | 44 | - | CCATTAG |
| MA0881.1 | MA0881.1.Dlx4 | 7.0524826 | 0.89727163 | 38 | 45 | - | GCCATTAG |
| MA0885.1 | MA0885.1.Dlx2 | 6.604795 | 0.89112103 | 38 | 45 | - | GCCATTAG |
| MA0880.1 | MA0880.1.Dlx3 | 6.8357754 | 0.8875854 | 38 | 45 | - | GCCATTAG |
| MA1476.3 | MA1476.3.Dlx5 | 9.176074 | 0.88447505 | 38 | 45 | - | GCCATTAG |
| MA0885.3 | MA0885.3.Dlx2 | 9.201587 | 0.8833738 | 38 | 45 | - | GCCATTAG |
| MA0675.2 | MA0675.2.NKX6-2 | 8.366475 | 0.96726143 | 39 | 44 | - | CCATTA |
| MA0723.3 | MA0723.3.VAX2 | 7.812914 | 0.95923865 | 39 | 44 | - | CCATTA |
| MA0707.3 | MA0707.3.MNX1 | 6.93994 | 0.9436445 | 39 | 44 | - | CCATTA |
| MA0699.2 | MA0699.2.LBX2 | 6.76798 | 0.90276426 | 39 | 44 | - | CCATTA |
| MA0706.2 | MA0706.2.MEOX2 | 7.32203 | 0.9012769 | 39 | 45 | - | GCCATTA |
| MA0708.3 | MA0708.3.MSX2 | 6.0327024 | 0.89843357 | 39 | 44 | - | CCATTA |
| MA0717.2 | MA0717.2.RAX2 | 6.12805 | 0.88441014 | 39 | 44 | - | CCATTA |
| MA0716.2 | MA0716.2.PRRX1 | 5.6275578 | 0.871014 | 39 | 44 | - | CCATTA |
| MA0721.2 | MA0721.2.UNCX | 5.737259 | 0.8680766 | 39 | 44 | - | CCATTA |
| MA0701.3 | MA0701.3.LHX9 | 5.586999 | 0.8678444 | 39 | 45 | - | GCCATTA |
| MA0711.2 | MA0711.2.OTX1 | 4.437145 | 0.8625714 | 39 | 44 | + | TAATGG |
| MA0700.3 | MA0700.3.LHX2 | 5.456798 | 0.85414034 | 39 | 44 | - | CCATTA |
| MA0748.2 | MA0748.2.YY2 | 8.607961 | 0.8494159 | 39 | 49 | + | TAATGGCCGGG |
| MA0710.2 | MA0710.2.NOTO | 3.726513 | 0.84821653 | 39 | 45 | - | GCCATTA |
| MA0748.1 | MA0748.1.YY2 | 8.935658 | 0.8463984 | 39 | 49 | - | CCCGGCCATTA |
| MA0718.2 | MA0718.2.RAX | 4.132467 | 0.83900243 | 39 | 44 | - | CCATTA |
| MA0910.3 | MA0910.3.HOXD8 | 11.068592 | 0.99444085 | 39 | 45 | - | GCCATTA |
| MA0876.2 | MA0876.2.BSX | 9.110977 | 0.98364615 | 39 | 44 | - | CCATTA |
| MA0875.2 | MA0875.2.BARX1 | 9.50878 | 0.9826615 | 39 | 44 | - | CCATTA |
| MA0893.3 | MA0893.3.GSX2 | 9.399768 | 0.9743518 | 39 | 45 | - | GCCATTA |
| MA0902.3 | MA0902.3.HOXB2 | 8.656557 | 0.96841115 | 39 | 44 | - | CCATTA |
| MA0903.2 | MA0903.2.HOXB3 | 8.463723 | 0.96729696 | 39 | 44 | - | CCATTA |
| MA0892.2 | MA0892.2.GSX1 | 7.5967455 | 0.94914377 | 39 | 44 | - | CCATTA |
| MA0879.3 | MA0879.3.DLX1 | 7.9927025 | 0.9467285 | 39 | 44 | - | CCATTA |
| MA0888.2 | MA0888.2.EVX2 | 7.187567 | 0.9305606 | 39 | 44 | - | CCATTA |
| MA0887.2 | MA0887.2.EVX1 | 7.051072 | 0.92398274 | 39 | 44 | - | CCATTA |
| MA0904.3 | MA0904.3.HOXB5 | 6.632622 | 0.91381854 | 39 | 44 | + | TAATGG |
| MA0900.3 | MA0900.3.HOXA2 | 6.160971 | 0.91252905 | 39 | 44 | - | CCATTA |
| MA0882.2 | MA0882.2.DLX6 | 7.047838 | 0.912509 | 39 | 44 | - | CCATTA |
| MA0904.3 | MA0904.3.HOXB5 | 6.522933 | 0.91141343 | 39 | 44 | - | CCATTA |
| MA0886.2 | MA0886.2.EMX2 | 6.384897 | 0.90922934 | 39 | 44 | - | CCATTA |
| MA0890.2 | MA0890.2.GBX2 | 5.374399 | 0.88136 | 39 | 44 | - | CCATTA |
| MA0889.2 | MA0889.2.GBX1 | 5.3540573 | 0.8734543 | 39 | 45 | - | GCCATTA |
| MA0891.2 | MA0891.2.GSC2 | 4.713759 | 0.8333031 | 39 | 44 | + | TAATGG |
| MA0914.2 | MA0914.2.ISL2 | 4.254517 | 0.8304505 | 39 | 44 | - | CCATTA |
| MA0894.2 | MA0894.2.HESX1 | 2.8480856 | 0.8235998 | 39 | 44 | + | TAATGG |
| MA0865.2 | MA0865.2.E2F8 | 6.938762 | 0.8026159 | 39 | 50 | - | TCCCGGCCATTA |
| MA1502.2 | MA1502.2.HOXB8 | 10.93254 | 0.9918023 | 39 | 45 | - | GCCATTA |
| MA1498.3 | MA1498.3.HOXA7 | 8.968124 | 0.99136084 | 39 | 44 | - | CCATTA |
| MA1500.2 | MA1500.2.HOXB6 | 10.058779 | 0.9868973 | 39 | 45 | + | TAATGGC |
| MA1496.2 | MA1496.2.HOXA4 | 9.418008 | 0.97234315 | 39 | 45 | - | GCCATTA |
| MA1504.2 | MA1504.2.HOXC4 | 9.001886 | 0.9703839 | 39 | 44 | - | CCATTA |
| MA1507.2 | MA1507.2.HOXD4 | 8.934631 | 0.9671691 | 39 | 44 | - | CCATTA |
| MA1497.2 | MA1497.2.HOXA6 | 9.092035 | 0.96146935 | 39 | 45 | - | GCCATTA |
| MA1499.2 | MA1499.2.HOXB4 | 8.712382 | 0.9584892 | 39 | 44 | - | CCATTA |
| MA1505.2 | MA1505.2.HOXC8 | 7.0299196 | 0.9415059 | 39 | 44 | - | CCATTA |
| MA1505.2 | MA1505.2.HOXC8 | 6.917952 | 0.9391473 | 39 | 44 | + | TAATGG |
| MA1501.2 | MA1501.2.HOXB7 | 9.172599 | 0.9353736 | 39 | 45 | - | GCCATTA |
| MA1495.2 | MA1495.2.HOXA1 | 6.3656745 | 0.92126167 | 39 | 44 | + | TAATGG |
| MA1495.2 | MA1495.2.HOXA1 | 6.218827 | 0.91830766 | 39 | 44 | - | CCATTA |
| MA1481.2 | MA1481.2.DRGX | 5.2122383 | 0.88524586 | 39 | 44 | - | CCATTA |
| MA1519.2 | MA1519.2.LHX5 | 5.718004 | 0.8797561 | 39 | 45 | - | GCCATTA |
| MA1577.2 | MA1577.2.TLX2 | 2.3491423 | 0.81136566 | 39 | 44 | - | CCATTA |
| MA0642.3 | MA0642.3.EN2 | 8.2832775 | 0.9242366 | 39 | 45 | - | GCCATTA |
| MA0642.2 | MA0642.2.EN2 | 8.220523 | 0.9135616 | 39 | 46 | - | GGCCATTA |
| MA0594.3 | MA0594.3.HOXA9 | 6.9888473 | 0.91153294 | 39 | 45 | - | GCCATTA |
| MA0612.3 | MA0612.3.EMX1 | 6.6334095 | 0.9100351 | 39 | 44 | - | CCATTA |
| MA0666.3 | MA0666.3.MSX1 | 5.955944 | 0.8939224 | 39 | 44 | - | CCATTA |
| MA0630.2 | MA0630.2.SHOX | 6.624642 | 0.89190876 | 39 | 44 | + | TAATGG |
| MA0661.2 | MA0661.2.MEOX1 | 6.8831544 | 0.8903459 | 39 | 45 | - | GCCATTA |
| MA0654.2 | MA0654.2.ISX | 5.5732846 | 0.88042647 | 39 | 44 | - | CCATTA |
| MA0662.2 | MA0662.2.MIXL1 | 5.712163 | 0.8788821 | 39 | 44 | - | CCATTA |
| MA0634.2 | MA0634.2.ALX3 | 5.612903 | 0.86559427 | 39 | 44 | - | CCATTA |
| MA0618.2 | MA0618.2.LBX1 | 5.3554945 | 0.8511317 | 39 | 45 | + | TAATGGC |
| MA0644.3 | MA0644.3.ESX1 | 3.9262066 | 0.8499202 | 39 | 45 | - | GCCATTA |
| MA0597.1 | MA0597.1.THAP1 | 5.940984 | 0.8494567 | 39 | 47 | - | CGGCCATTA |
| MA0635.2 | MA0635.2.BARHL2 | 2.1591628 | 0.83127856 | 39 | 44 | + | TAATGG |
| MA0648.2 | MA0648.2.GSC | 4.457343 | 0.8309553 | 39 | 44 | + | TAATGG |
| MA0132.3 | MA0132.3.PDX1 | 8.654493 | 0.9577921 | 39 | 44 | - | CCATTA |

| | | | | | | | |
|---|---|---|---|---|---|---|---|
| MA0027.3 | MA0027.3.EN1 | 5.5449796 | 0.88449544 | 39 | 44 | - | CCATTA |
| MA2119.1 | MA2119.1.HOXA3 | 7.953136 | 0.9263085 | 39 | 45 | - | GCCATTA |
| MA1502.2 | MA1502.2.HOXB8 | 10.93254 | 0.9918023 | 39 | 45 | - | GCCATTA |
| MA1498.3 | MA1498.3.HOXA7 | 8.968124 | 0.99136084 | 39 | 44 | - | CCATTA |
| MA1500.2 | MA1500.2.HOXB6 | 10.058779 | 0.9868973 | 39 | 45 | + | TAATGGC |
| MA1496.2 | MA1496.2.HOXA4 | 9.418008 | 0.97234315 | 39 | 45 | - | GCCATTA |
| MA1504.2 | MA1504.2.HOXC4 | 9.001886 | 0.9703839 | 39 | 44 | - | CCATTA |
| MA1507.2 | MA1507.2.HOXD4 | 8.934631 | 0.9671691 | 39 | 44 | - | CCATTA |
| MA1497.2 | MA1497.2.HOXA6 | 9.092035 | 0.96146935 | 39 | 45 | - | GCCATTA |
| MA1499.2 | MA1499.2.HOXB4 | 8.712382 | 0.9584892 | 39 | 44 | - | CCATTA |
| MA1505.2 | MA1505.2.HOXC8 | 7.0299196 | 0.9415059 | 39 | 44 | - | CCATTA |
| MA1505.2 | MA1505.2.HOXC8 | 6.917952 | 0.9391473 | 39 | 44 | + | TAATGG |
| MA1501.2 | MA1501.2.HOXB7 | 9.172599 | 0.9353736 | 39 | 45 | - | GCCATTA |
| MA1495.2 | MA1495.2.HOXA1 | 6.3656745 | 0.92126167 | 39 | 44 | + | TAATGG |
| MA1495.2 | MA1495.2.HOXA1 | 6.218827 | 0.91830766 | 39 | 44 | - | CCATTA |
| MA1481.2 | MA1481.2.DRGX | 5.2122383 | 0.88524586 | 39 | 44 | - | CCATTA |
| MA1519.2 | MA1519.2.LHX5 | 5.718004 | 0.8797561 | 39 | 45 | - | GCCATTA |
| MA1577.2 | MA1577.2.TLX2 | 2.3491423 | 0.81136566 | 39 | 44 | - | CCATTA |
| MA2093.1 | MA2093.1.HOXB1 | 7.3222885 | 0.9142894 | 39 | 45 | - | GCCATTA |
| MA0125.2 | MA0125.2.Nobox | 7.1938314 | 0.90052944 | 39 | 44 | + | TAATGG |
| MA0704.2 | MA0704.2.Lhx4 | 6.095402 | 0.89003617 | 39 | 44 | - | CCATTA |
| MA0125.1 | MA0125.1.Nobox | 7.885545 | 0.8853156 | 39 | 46 | + | TAATGGCC |
| MA0720.2 | MA0720.2.Shox2 | 5.673306 | 0.8838594 | 39 | 44 | - | CCATTA |
| MA0705.2 | MA0705.2.Lhx8 | 3.858031 | 0.87139773 | 39 | 44 | - | CCATTA |
| MA0709.2 | MA0709.2.Msx3 | 5.5862036 | 0.8697775 | 39 | 44 | - | CCATTA |
| MA0705.2 | MA0705.2.Lhx8 | 3.072413 | 0.8550231 | 39 | 44 | + | TAATGG |
| MA0075.1 | MA0075.1.Prrx2 | 5.342571 | 0.84313756 | 39 | 43 | - | CATTA |
| MA0881.2 | MA0881.2.Dlx4 | 7.1609955 | 0.9143908 | 39 | 44 | - | CCATTA |
| MA0880.2 | MA0880.2.Dlx3 | 7.0805902 | 0.9079615 | 39 | 44 | - | CCATTA |
| MA2339.1 | MA2339.1.Nanog | 5.369332 | 0.80714804 | 39 | 45 | - | GCCATTA |
| MA0671.1 | MA0671.1.NFIX | 3.2760122 | 0.8226315 | 40 | 48 | - | CCGGCCATT |
| MA0095.1 | MA0095.1.YY1 | 7.219338 | 0.94056094 | 40 | 45 | - | GCCATT |
| MA0748.3 | MA0748.3.YY2 | 9.06184 | 0.90647644 | 41 | 47 | + | ATGGCCG |
| MA0671.2 | MA0671.2.NFIX | 3.9440665 | 0.8476084 | 41 | 46 | - | GGCCAT |
| MA0738.2 | MA0738.2.HIC2 | 2.5845835 | 0.81601536 | 41 | 46 | + | ATGGCC |
| MA0161.1 | MA0161.1.NFIC | 4.377613 | 0.8218204 | 41 | 46 | + | ATGGCC |
| MA0740.2 | MA0740.2.KLF14 | 6.0892816 | 0.84053344 | 42 | 50 | + | TGGCCGGGA |
| MA0865.3 | MA0865.3.E2F8 | 5.7615986 | 0.80881864 | 42 | 50 | - | TCCCGGCCA |
| MA0516.3 | MA0516.3.SP2 | 4.4059386 | 0.80261517 | 42 | 50 | + | TGGCCGGGA |
| MA2341.1 | MA2341.1.FEZF2 | 6.4996877 | 0.8117953 | 42 | 49 | - | CCCGGCCA |
| MA1513.2 | MA1513.2.KLF15 | 4.508929 | 0.83367336 | 43 | 50 | - | TCCCGGCC |
| MA0471.3 | MA0471.3.E2F6 | 5.2871466 | 0.80621666 | 43 | 50 | + | GGCCGGGA |
| MA1513.2 | MA1513.2.KLF15 | 4.508929 | 0.83367336 | 43 | 50 | - | TCCCGGCC |

### Supplementally table S15. Sequences list (primers, DsiRNAs, sgRNA spacer sequences, and ERNI cDNA)

primers for qRT-PCR

| Gene Nane | Direction | Sequence |
|---|---|---|
| GAPDH | Fwd | GGGCACGCCATCACTATC |
| | Rev | GTGAAGACACCAGTGGACTCC |
| POU5F3 | Fwd | AACGCTGGAGAGCTTCTTCCG |
| | Rev | GAACCAGACCCGGACAACGTC |
| NANOG | Fwd | AGCCGGTTTCAGAGCCAGAAG |
| | Rev | AAACCACGTCTTCACCTGCT |
| DDX4 | Fwd | TGACTTATGTCCCCCCTCCT |
| | Rev | GTAATGGTGCTGGAGGGTCA |
| DAZL | Fwd | AGAATGTGCTGTCCCAGAGC |
| | Rev | CAAAGGTGTTCCTCAGACGG |
| PRDM14 | Fwd | TGCAATAAGGCGTTCACAGC |
| | Rev | TCCTCCAGCTCTGGGAAGTG |
| Soprano | Fwd | CTTATCTCTCACCACGACAGACTTG |
| | Rev | GAGGAGGATCGGAAGGCAGG |
| ENS-1/ERNI_CDS | Fwd | GGGGACTTCCTGAATCGTTAAAACC |
| | Rev | TACAGATCCCTCCAACGCTGC |
| AGK | Fwd | GGTCACCTCCACGGAAAGAA |
| | Rev | AGTCCTCAGTGCGTGTCAGA |
| CDH7 | Fwd | AGAGTCTGTGACTGCGATGC |
| | Rev | TCAGCAAGACAAGAACTAGCAA |
| DENND11 | Fwd | GGAGAACCAGGTCAGACACC |
| | Rev | AGGTGGTGGTGCCCTTACTA |
| ENOX1 | Fwd | GGGCCCTGAGTTCGATGAC |
| | Rev | TTCCCAAACCATCAGCAGCA |
| GUCA1C | Fwd | TGCTGTGGGAACAGCACTTA |
| | Rev | GTTTCCAGCATCCTCCTCCA |
| LOC418414 | Fwd | ACGGTGGCTGAGCAGATTAG |
| | Rev | GTGTGTCCCAGACTGCTGTAG |
| SLC16A7 | Fwd | AAATTCATAGCCCCGCTCAC |
| | Rev | TCTTCTCCAATTAGGCTGGTCTC |
| TMPRSS15 | Fwd | ACAGCATACAGATCACAGTGGTT |
| | Rev | TGGAGTTGATGCCAGAGAATTTG |
| WEE1 | Fwd | GAATACTGCAATGGCGGCAG |
| | Rev | GAGCCACCTGAAGCAGTAGG |
| WEE2 | Fwd | TCAAGCGCCTTGATGGATGT |
| | Rev | GGTGTCCAAGGACTGCATGA |

Primers for sex identification

| | | |
|---|---|---|
| CHD1 | Fwd | TATCGTCAGTTTCCTTTTCAGGT |
| | Rev | CCTTTTATTGATCCATCAAGCCT |

DsiRNAs

| | | |
|---|---|---|
| siENS-1/ERNI#1 | passenger | CAAGGAAAGAUACAAGCCAUGUCTC |
| | guide | GAGACAUGGCUUGUAUCUUUCCUUGGA |
| siENS-1/ERNI#2 | passenger | GAUGAAUGGAUUACCCACAGUCAGA |
| | guide | UCUGACUGUGGGUAAUCCAUUCAUCAA |

blue:DNA red:RNA

sgRNA spacer sequences

| Name | Sequence | discription |
|---|---|---|
| sgSoprano#1 | GATAAGCTGCCGCTAATGGC | for CRISPRi with synthesized gRNA |
| sgSoprano#2 | GACCGTTTACCGGCCATTAG | for CRISPRi with synthesized gRNA |
| sgSoprano#3 | AGCCCCCAGGAATGTCACGT | for CRISPRi with plasmid transfection |
| sgSoprano#4 | CGTTGCCGTGGTTGTGGTGA | for CRISPRi with plasmid transfection |
| sgSoprano#5 | AGAGGGTCCAATAACCACAA | for CRISPRi with plasmid transfection |
| sgENS-1/ERNI#1 | CCTGAATCGTTAAAACCTAT | for knockout |
| sgENS-1/ERNI#2 | CCTATAGGTATACAACTCCA | for knockout |
| sgENS-1/ERNI#3 | ACATGGCTTGTATCTTTCCT | for knockout |
| sgOVA#1 | TGCTGTACATAGTACCATGC | for knockout & CRISPRi synthesized gRNA and plasmid transfection control |
| sgOVA#2 | GCTCTAGCCATGGTATACCT | for CRISPRi with plasmid transfection control |
| sgOVA#3 | GATTATACTCTCAAGCTGCA | for CRISPRi with plasmid transfection control |

ERNI_cDNA

ATGAGCAACAGTATGGCCAGTATGAAAAGTGAAGATGTATTATTTGATCTTTTAGAAAAGCATGGTGCTCGGCCTTCTGTATCAGGGGTGGATTGGGCACGACAGAACTGGTATAATTTGCAAAGTGTTTCAGAC
CGTATTCGTGTTTTACAAAATGAGGCTCGTACTCGGGCCGGAAAAGGGAAATCTTTTATTTGTGCAGTACTCGGTGCTGCTTTAAAAGCAGCTGTGGAGTTCCGAGAGGAAAAGAACTCTACGGAAACCCAGACT
ATTCAAGCATTACAGGAATCGGTTAAAGTGACGCAAGAATTGGTAAAATCTCTGAGAAGCCAAATAATGAGTCTTGAGGATCAATTAGAAAGAGAAAAACATAATTCGGTTTTGTTGCAAACAGCTTTTAAGGAGCT
GATAACGTGTAAGGACACCGGTGACGCTGTTATCCACAGTGCACCTCAAGAAAAAGTTTATCCTCAAGGGAAATTACAAGAGGTGAAGGAAAGGCTAGATAAATTAGAGGCCTCTCCAGCCCACATTCGTCCTTT
GATAAAAACTGAATATACTTTCGATAACAGTGAGAATCTAGATCCTCAAATGAATGTTAAGGAAATTCCCTTTTCGGCCACTGAACTGGCCAAACTGAAAAAGGATTTCAGTCGCTCCCCAAAGGAGTCTGAAACAG
AGTACGTCTGGAGAGTTAGTCTCACTGGCGGAGACCAGATCCTACTAACAGAGAAAGAAGCTGAAGGTTACTGGGGACCAGGAGTATTTTTAACCACTGGCAATAATCGTGCTCCCTGGTCCTTAACACAGAGG
GCTGCCTATTGGGCAGGGGGTCTCAACCCTTTAGAAAGGGGGGACCCTCTTGCTATTACTGGAACTATCGACCAGTTAGTGGAGAATGTTCAGAAAACTGCTTGTCTCCAAATGATGTATGATAGAAAGTTGCAA
CCACATCATGAATCACCCATGATGTTACCTGTTAATCCGGAGAGATTGACACCTCTAATTAGGGGACTTCCTGAATCGTTAAAACCTATAGGTATACAACTCCAAGGAAAGATACAAGCCATGTCTCAGGGAGAGA
GAACCTGGGCAGCGTTGGAGGGATCTGTAGCCCCTAACCACCAGTCAGGACCCAAAGTGTGGACTTGGGGAGAGGTTGCCCAAGAATTAATTAACTATGGAAGAAAATATGGGCCGGTGGTTTCTACCTCCAG
TAAATTTGAGCCAAGAGGAGTAAGGCTTGCAGTAGCCAGCCTTGCCTCCAGGCCTCCTAGCCCAAGACTTATTGGAACCAAAAAGGTTTCATCCCCAGTAAAAACGGGGACACGATGCATTGATCATAAACGCAA
TGGACTTTGGACTCTGGGCTGGACAAAGGGTATTCCACGAGATTTGATGAATGGATTACCCACAGTCAGATTAGAGAAATTAGTTAACTGCTGGCCAGAACAAAAGCTCAAGGGGAGCTGA
